# Core Activation Program and Selective Regional Responsiveness of Microglia in Aging and Parabiosis

**DOI:** 10.1101/2025.10.29.685417

**Authors:** Huma Naz, Robert Pálovics, Shinnosuke Yamada, Nannan Lu, Tony Wyss-Coray, Qingyun Li, Guoyan Zhao

## Abstract

Aging is associated with immune dysregulation in brain and is the biggest risk factor for many neurodegenerative diseases whereas rejuvenation interventions can mediate beneficial effects. Microglia are considered as a major player in the development of neurodegenerative disease yet, the molecular changes underlying brain aging and rejuvenation remain poorly understood at the single cell level. We identified and benchmarked several reproducible microglial states and a core set of genes leading to microglia activation in the mice brain. We investigated microglial heterogeneity and studied the impact of aging and parabiosis-mediated exposure of young and old blood on microglia subpopulations across four different brain regions including cerebellum, cortex, hippocampus, and striatum. We revealed region-specific differences in microglia subpopulation composition and age-related changes, with cerebellum and striatum displaying the most distinctive profiles and dynamic shifts compared to other brain regions. We consistently observed cerebellum as the most responsive, while striatum appeared distinctive by its minimal responsiveness to these interventions. Our findings highlighted the role of microglia in brain regional vulnerability and provided a foundation for microglia-targeted treatment for modulating brain aging.

**Highlights:**

- Defined the composition of different microglial populations reproducible in aging and parabiosis, benchmarking a reference for the field.
- Uncovered an under-appreciated core activation gene signature of microglia shared in all reactive states and regions during normal aging and old blood-induced aging.
- Identified region-specific gene expression changes and associated biological processes in microglia during aging and parabiosis
- Discovered microglial regional selectivity in response to aging and parabiosis, showing cerebellum as the most sensitive region and the striatum as the least affected.

## Introduction

Aging is the primary risk factor for most neurodegenerative diseases, including Alzheimer’s disease (AD), the most common form of dementia, affecting ∼55 million people worldwide^1,2^. The central nervous system (CNS) undergoes numerous changes during normal aging, including cortical shrinkage, limited neurogenesis, reduced synaptic density, decreased glucose metabolism, elevated levels of proinflammatory cytokines, decreased levels of anti-inflammatory cytokines, and increased blood-brain barrier (BBB) permeability^3^. Recent studies using bulk RNA sequencing from 15 brain regions have highlighted the dynamic and region-specific nature of molecular changes in the aging brain, revealing widespread shifts in gene expression patterns, particularly in microglial cells^4^. Microglia, the brain resident macrophages, play important roles in maintaining brain homeostasis by modulating neuronal activity, clearing debris, and responding to foreign stimuli^5^. Microglial dysfunction is increasingly recognized as a key contributor to aging and neurodegenerative disease pathogenesis^6–8^. Regional differences in microglial responses to injury and neurodegenerative conditions have been reported, suggesting that transcriptomic variations may underlie the selective vulnerability of specific brain regions^6,9–11^. However, regional differences of microglia during aging and rejuvenation interventions have not been well characterized at the single-cell level. Elucidating the key pathways that regulate microglial regional heterogeneity and transcriptomic changes during aging and rejuvenation may provide crucial insights into regional susceptibility to degeneration and aid in the development of novel therapeutic strategies.

Extensive studies using *in vitro* systems and mouse models have significantly advanced our understanding of microglial heterogeneity during development and across various neurodegenerative diseases^12–14^. Under physiological conditions, microglia typically maintain a homeostatic state with a ramified morphology, playing an important role in sustaining brain homeostasis^15^. As the brain’s resident immune cells, microglia express a wide array of receptors that act as molecular sensors, enabling them to recognize exogenous or endogenous insults and initiate immune responses. These responses include changes in cellular morphology, secretion of chemokines and cytotoxic factors, and proliferation, collectively referred to as “reactive” or “activated” microglia^16,17^. Microglia exhibit significant heterogeneity, adopting diverse activation states depending on the stimuli, and performing both detrimental and beneficial functions based on their activation status^12,15^. For example, studies have identified distinct transcriptional and functional signatures associated with specific microglial states, such as homeostatic microglia (HS) ^18–20^, disease-associated microglia (DAM) ^8,21,22^, interferon-response microglia (IRM)^23^. However, how microglial gene expression changes during the aging process and the effects of rejuvenation on different microglial subpopulations remain poorly understood.

Pioneering research using techniques such as heterochronic parabiosis and plasmapheresis has provided compelling evidence of a strong connection between the systemic environment and brain aging^24–26^. Heterochronic parabiosis involves surgically connecting the circulatory systems of two mice, allowing shared blood flow^27^. In this context, heterochronic parabionts refer to a young mouse paired with an old mouse (treatment groups: heterochronic young and heterochronic aged), while isochronic parabionts refer to mice of the same age paired together (control groups: Isochronic young and Isochronic aged). Heterochronic parabiosis has demonstrated significant health benefits for aged parabionts connected to young partners, including enhanced bone regrowth, neurogenesis, and cognitive function, but has also revealed detrimental effects on young parabionts connected to aged ones^28,29^. Our recent single-cell transcriptomic studies of 20 major organs and various cell types during aging and heterochronic parabiosis in mice enabled comparisons of normal aging, accelerated aging induced by old blood, and rejuvenation driven by young blood across tissues. These studies uncovered both global and tissue- or cell-type-specific aging signatures, as well as cell-type-specific responses to young and old blood ^30,31^. However, due to the scale of the work, detailed investigation into changes of microglia, particularly at the level of specific cell subpopulations have not been performed.

The *Tabula Muris Senis* and the parabiosis datasets included microglia isolated from four different brains regions–cerebellum, cortex, hippocampus, and striatum–of young, old and parabiotic mice^31,32^. In this study, we leveraged these high-quality datasets to investigate microglial heterogeneity and their transcriptomic changes during aging and rejuvenation across the four brain regions. We identified eight distinct microglial subpopulations and seven of which were validated in a separate aging dataset. We nominated combinatorial signaling codes governing different reactive microglial states and defined the core microglial activation gene program. Additionally, we uncovered shared and region-specific aging signatures across the four brain regions and identified region-specific differential gene expression associated with cellular senescence, aging, amyloid metabolism and response to unfolded proteins. Our analysis revealed that parabiosis-mediated accelerated aging and rejuvenation had distinct effects on gene expression in specific brain regions, with the cerebellum consistently emerging as the most sensitive region and the striatum as the least affected. Pathways involved in apoptotic signaling, inflammatory responses, glial cell activation, synaptic plasticity, and phagocytic activity were differentially impacted across brain regions in response to aged and young blood exposure. This work highlights the regional variation in microglial subpopulation transcriptomic dynamics during aging and under rejuvenation conditions, providing insights into differential brain regional vulnerability and offering potential avenues for microglia-targeted modulation of brain aging.

## Results

### Distinct microglial subpopulations shared across brain regions and lifespan

To investigate microglial subpopulations and their transcriptomic changes during aging and rejuvenation, we took advantage of our published scRNA-seq data from the aging study, *Tabula Muris Senis* (referred to as the “AGE” data herein)^32^ and the parabiosis study (referred to as the “PB” data herein)^31^. The AGE dataset included microglia collected from three young mice of age 3-4 months and six old mice of age 18-24 months, which corresponded to human ages of 18-20 and 65-75 years, respectively. The PB dataset included three isochronic parabiosis animal pairs and two heterochronic parabiosis animal pairs^31^. Heterochronic parabionts were referred to as heterochronic young (HY) and heterochronic aged (HA), respectively. Isochronic parabionts served as the controls for the parabiosis surgery effects, and depending on the age of the animals, they were referred to as isochronic young (IY) or isochronic aged (IA). These datasets enabled the comparison of parabiosis-mediated rejuvenation (REJ, heterochronic aged vs. isochronic aged) and accelerated ageing (ACC, heterochronic young vs. isochronic young) to normal ageing (AGE, aged vs young) (Fig. 1a).

**Fig. 1:**
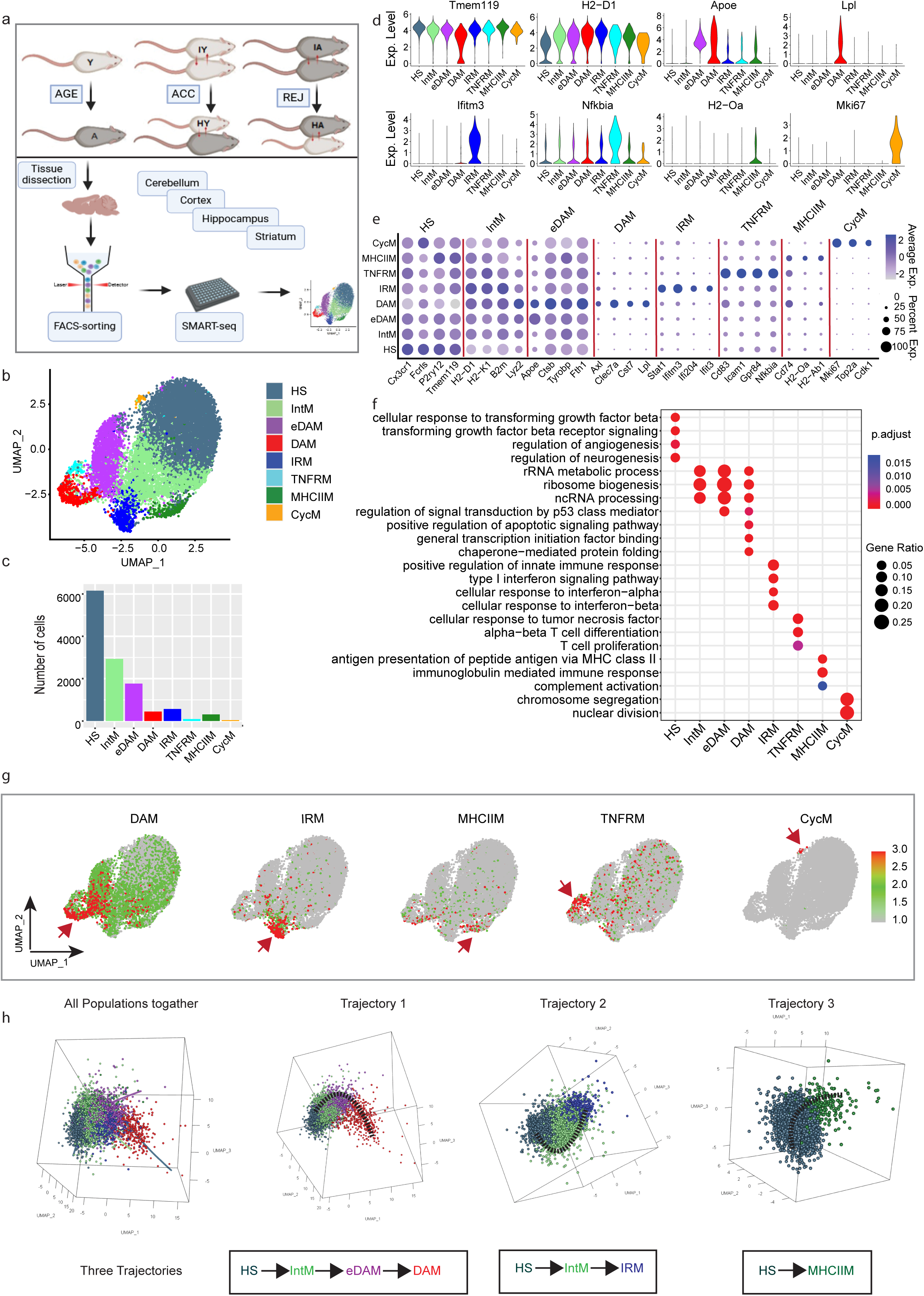
Identification and characterization of microglia subpopulations. **a**) Experimental workflow. **b**) Unsupervised clustering of scRNA seq data from AGE+Parabiosis study and UMAP (Uniform Manifold Approximation and Projection) representation of different microglia subpopulation (total cells= 12,310) colored by cluster identity. UMAP plots were generated using default parameters except reduction = ‘pca’, dims = 1:30, res=0.35. **c**) Number of cells in each microglia subpopulation. **d**) Violin plot of known marker gene expression. **e**) Dot plot showing the expression of marker genes of each cluster. **f**) Gene ontology showing enrichment of unique biological process of identified microglia subpopulations. **g**) Projection of module score activity calculated using reported subpopulation-specific feature genes. **h**) Pseudotime analysis showing potential microglia activation trajectories for different microglia subpopulations. For all data, the experiment was performed once. FindMarkers using Wilcoxon rank sum test and Seurat R package, adjusted P value < 0.05. HS; homeostatic microglia, IntM; intermediate activated microglia, eDAM; early disease associated microglia, DAM; disease associated microglia, IRM; interferon-response microglia, TNFRM; tumor necrosis factor response microglia, MHCIIM; major histocompatibility complex class II-positive microglia and CycM; cycling microglia.

First, we performed integrated analysis of AGE and PB combined data (referred to as “AGE + PB” data) using single-cell genomics tool Seurat^33^. After removing low quality cells and doublets, a total of 12,310 cells were obtained. We performed unsupervised cell clustering and identified eight microglial subpopulations based on their distinct expression profiles (Fig. 1b, c, Fig S1a-c). We identified marker genes highly expressed in each microglial subpopulation (Supplementary Table 1) and annotated cell clusters by comparing their marker genes with the feature genes of known microglial states. Homeostatic microglia (HS) were annotated based on the high expression of multiple conventional homeostatic microglia markers such as *Cx3cr1, Fcrls, P2ry12* and *Tmem119* (Fig. 1d, e). Gene ontology (GO) enrichment analysis on the cluster marker genes revealed unique enrichment of the response to transforming growth factor beta (TGFβ) signaling pathway (Fig. 1f), consistent with the known function of TGFβ in regulating microglial homeostatic state under basal nonpathological conditions^34^. Disease-associated microglia (DAM) was identified by the unique expression of multiple DAM signature^8^ genes such as *Axl, Clec7a, Csf1, Cst7,* and *Lpl* (Fig. 1d, e). Early disease-associated microglia (eDAM) was named because of the strongest enrichment of *Apoe* expression and the shared markers with the reported stage 1 DAM^8^ such as *Ctsb, Lyz2, Tyrobp, Fth1*, and *B2m* (Fig. 1d, e). DAM signature gene activity was most strongly enriched in eDAM and DAM (Fig. 1g)^8^. Type I interferon-response microglia (IRM) had unique expression of many interferon-response genes such as *Ifitm3, Ifit3*, and *Ifi204* (Fig. 1d, e) and were uniquely enriched for type 1 interferon signaling pathway (IFNα and IFNβ) and positive regulation of innate immune response pathways (Fig. 1f)^35^. IRM have been identified from scRNA-seq studies on murine AD models and their signature features were most strongly enriched in IRM cluster (Fig. 1g)^23,36^. The major histocompatibility complex class II microglia (MHCIIM) were named for the expression of multiple MHC class II genes such as *H2-Oa, Cd74*, and *H2-Ab1* as well as the unique enrichment of antigen presentation via MHC class II pathway in the maker genes of this population (Fig. 1e. f). Reported MHCIIM feature gene activities were also mapped to this population (Fig. 1g)^23,36,37^. Tumor necrosis factor response microglia (TNFRM) was named for the unique enrichment of cellular response to tumor necrosis factor pathway (Fig. 1f). Multiple previously reported disease inflammatory macrophages (DIM) features such as *Cd83, Icam1,* and *Nfkbia* were highly expressed in this population (Fig. 1d, e) and DIM feature activities were mapped to TNFRM (Fig. 1g)^38^. Cell proliferation marker genes, *Mki67*, *Top2a,* and *Cdk1*, distinguished proliferating microglia (CycM) from all other subpopulations with multiple cell-cycle related pathways uniquely enriched in this population (Fig. 1d-g)^36,39^. Intermediate activated microglia (IntM) were characterized by the increased expression of *H2-D1, H2-K1, B2m, Lyz2.* This population shared many genes that were broadly upregulated in different types of activated microglia without any gene uniquely enriched in this population. It was therefore named intermediate activated microglia, which may represent parainflammation state, an intermediate immune state where immune responses are directed toward reestablishing homeostasis that is distinctive from pathological inflammation^40,41^. To investigate the relationships among different microglial subpopulations, we performed pseudotime analysis using Slingshot^42^. Three pseudotime trajectories were identified which revealed that the IntM population served as an intermediate state linking other activated microglia, such as eDAM, DAM, and IRM (Fig. 1h).

To validate the identified microglial subpopulations, we independently analyzed another scRNA-seq aging dataset published by Hammond *et al.* (referred to as the “TRH” data) ^43^ which compared microglia from whole brain tissues in six young (3 month) and six old (21 month) mice. Using the same analysis algorithm and parameters, we identified 9 microglial subpopulations based on their distinct expression profiles (Fig S1d-g). Seven out of nine (HS, IntM, eDAM, DAM, IRM, TNFRM and CycM) had corresponding populations in our data with highly correlated cluster marker gene expression (Fig. S1h). Pseudotime analysis identified four trajectories consistent with our data that the IntM population served as an intermediate state linking other activated microglia states (Fig. S1i). Two microglial states with unique expression of *Notch2* and *Kcnq1ot1* were observed only in TRH data, which may be due to difference in brain regions being sampled--whole brain for TRH data versus four brain regions in our study. The lack of MHCIIM in the whole-brain data could be due to the low abundance of this population in brain regions other than the four included in our data. Overall, this analysis defined distinct microglial subpopulations characterized by specific transcriptional profiles in young, aged and parabiosis mice.

### Microglia activation state dynamics during aging and rejuvenation

Given that many of the activated microglia subpopulations, such as DAM and IRM were initially identified in disease models or aged animals^23,43,44^, we first analyzed population dynamic change during aging. All microglia subpopulations exist in both young and aged animals in both datasets (Fig. S2a) suggesting that these distinct microglia subpopulations exist in young mice under physiological conditions. Next, we tested cell proportion changes between young and old mice using Propeller test which is robust to the existence of outliers and performs multiple comparison correction^45^. For the AGE data, we analyzed cells from all brain regions together and found a significant decrease of eDAM but an increase of TNFRM and MHCIIM populations during aging (Fig. S2a). We observed the same population shift in TRH data for eDAM and TNFRM. In addition, HS and proliferating microglia populations decreased whereas, DAM and IRM increased significantly in aged mice in TRH whole brain data. The regional information in the AGE data enabled us to examine microglia population dynamics in each brain region separately. We found that cerebellum had a distinct cellular composition than other brain regions and the TRH whole brain data (Fig. S2b) with much lower proportions of HS microglia but higher proportions of IntM and eDAM. No significant differences of microglia composition between young and aged mice were detected in the cerebellum or hippocampus. Striatum had the most dramatic population shift with decreased HS population and increased proportions of eDAM, IRM, TNFRM, and MHCIIM microglia. TRH whole brain data recapitulated these findings except the MHCIIM microglia, which was absent in the TRH data. However, MHCIIM microglia were consistently increased in the cortex and striatum as well as in the combined AGE data. IRM has been shown to accrual over time in an age-dependent manner accompanying progressive β-amyloidosis ^35^. Interestingly, Chen et al. found that the percentages of the IRM and the number of microglia positive for MHC class II proteins was significantly elevated in brain regions with tau pathology in TE4 mice which is associated with T cell infiltration and neurodegeneration^46^. In summary, our results revealed region-specific differences in microglia subpopulation composition and age-related changes, with cerebellum and striatum displaying the most distinctive profiles and dynamic shifts compared to other brain regions. These regional differences may contribute to the varying vulnerability of brain regions to neurodegenerative disease processes.

Next, we examined microglia subpopulation proportion changes during aging and parabiosis using Propeller test considering hetero-young (HY) and hetero aged (HA) mice as intermediate aged mice between young and old mice. The striatum remains the most dramatically affected brain region with significant changes in HS, IntM, eDAM, IRM, TNFRM, and MHICIIM (Fig. S2c). Exposure to old blood significantly reduced the HS population but increased eDAM proportions whereas exposure to young blood brought back the HS population but significantly reduced the IntM proportions. Furthermore, we used the Jonckheere-Terpstra test to determine whether the shifts in microglia subpopulation proportions exhibited a statistically significant trend for each population independently. We found a consistent increases in the MHCIIM population in the cortex, hippocampus and striatum (Fig. S2d). Similarly, the IRM population consistently increased in the hippocampus and striatum with the largest changes in aged mice. These results are consistent with the above observation, suggesting the exposure of young blood significantly impacted the microglia subpopulation structure. We also observed a significant decrease in the DAM population in the cortex and hippocampus (Fig. S2d). Given the observed protective role of DAM the decrease in DAM populations during aging may underlie the increased vulnerability of these brain regions. No population structure changes were observed for the cerebellum.

### Combinatorial signaling code underlying the activation states of microglia

To better understand properties of different microglia subpopulations, we compared gene expression of activated microglia populations and proliferating microglia to that of the homeostatic microglia. We identified 100-1221 genes differential expressed in various microglia subpopulations (referred to as subpopulation DEGs herein) (Fig. 2a, Fig. S3a, Supplementary Table 2). Most of the DEGs were unique to certain subpopulations suggesting distinct gene expression programs associated with these microglia activation states (Fig. 2b). This is consistent with the current view that microglia exist in diverse, dynamic, and multidimensional states depending on the context, including local environment^47^. We also observed subpopulation DEGs shared across multiple subpopulations and there were few overlaps between up- and downregulated DEGs across different subpopulations suggesting that microglia may employ a conserved gene program when became activated (Fig. 2b). To identify the core microglia activation program shared by all activated microglia subpopulations, we identified subpopulation DEGs in at least 5 out of 6 activated microglia subpopulations in our dataset. We obtained 75 genes including upregulation of *H2-D1, H2-K1, Cd74, Cd52, Cd9, Ctsz, Fth1, Fcer1g, Selplg* and downregulation of *P2ry12, P2ry13, Cx3cr1, Fcrls* in activated microglial states (Fig. 2c, d, Supplementary Table 3). We performed the same analysis on TRH data and identified 32-218 subpopulation DEGs compared to the HS microglia (Fig. S3a). Many more subpopulation DEGs were identified in our data compared to the TRH data for each corresponding subpopulation likely due to deeper sequencing depth of our data (Fig. S3a). The gene expression fold changes showed the highest correlation between equivalent subpopulations of the two datasets (Fig. S3b) demonstrating reproducibility of characteristic gene expression programs associated with each microglial activation state. Many shared subpopulation DEGs identified in our dataset were also recapitulated in the TRH dataset consistent with the presence of a core microglial activation program (Fig. S3c, d)^48,49^.

**Fig. 2:**
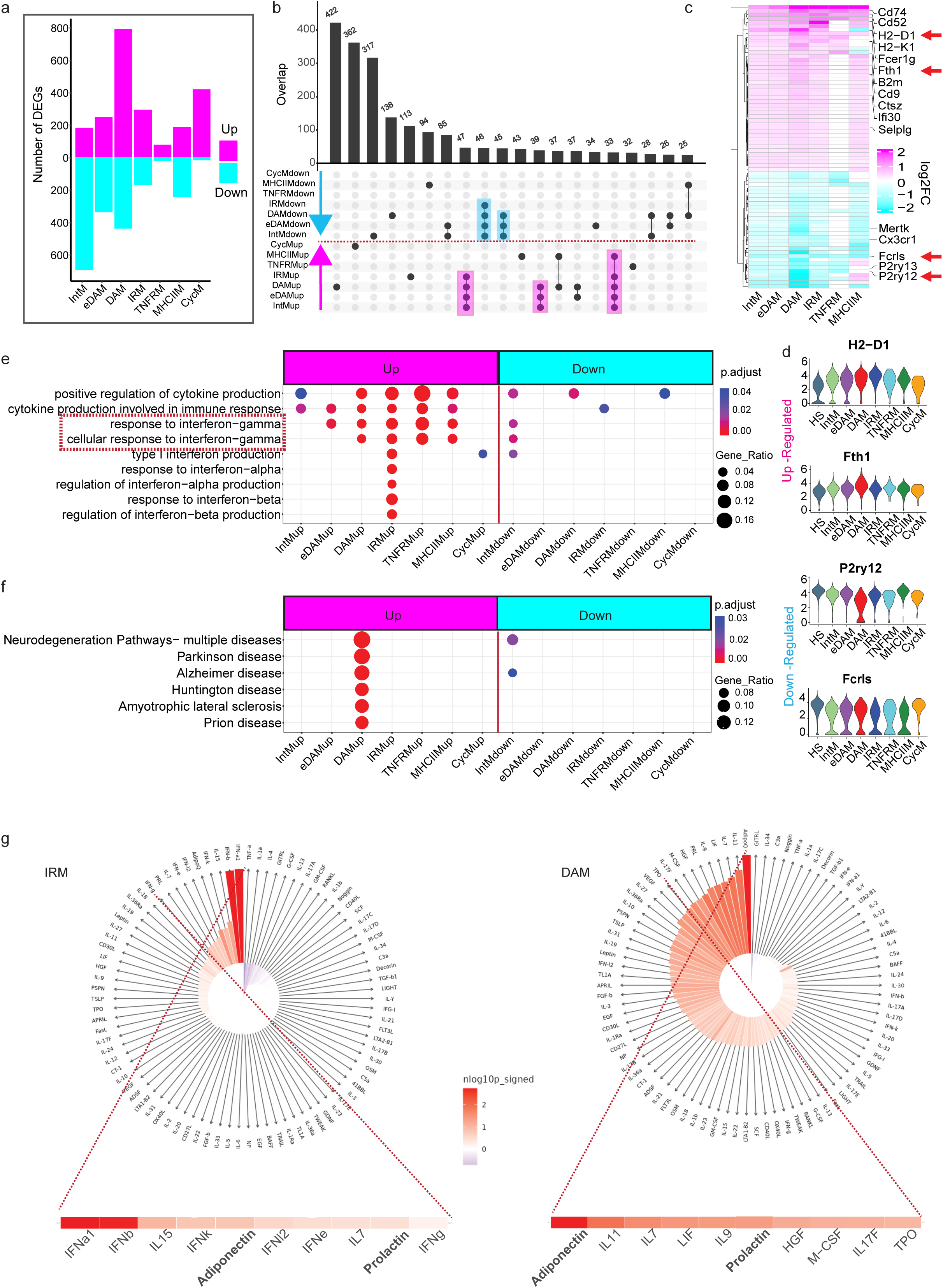
Comparison of genes and pathways associated with microglia activation. **a**) Bar plot showing the number of up and down regulated DEGs (differential expressed genes) of all activated microglia and CycM compared to HS microglia (Wilcoxon rank sum test, FDR-adjusted *P* value < 0.05 and absolute log2FC > 0.25). **b**) Upset plot showing overlap of up and downregulated DEGs. **c**) Heatmap of DEGs log2FC showing core set of DEGs shared by at least 5/6 activated microglia subpopulations. **d**) Violin plot showing examples of up and down regulated core activated microglia genes. **e**) GO (Gene Ontology) enrichment analysis on the subpopulation DEGs showing some shared and unique biological pathways. **f**) KEGG (Kyoto Encyclopedia of Genes and Genomes) pathways showing unique enrichment of neurodegenerative disease related pathways. g) IREA (Immune response enrichment analysis) showing top enrichment cytokine response genes in the DEGs of IRM and DAM compared to HS microglia. DEGs were calculated using Wilcoxon rank sum test and Seurat R package, adjusted P value < 0.05 and absolute Log2FC>0.25. Pathways with FDR-adjusted P value < 0.05 (hypergeometric test) and at least five query genes were considered statistically significant. Up: upregulated; down: downregulated.

Next, to understand the functional implication of the gene programs engaged during microglial activation, we performed GO enrichment analysis on the subpopulation DEGs. Interestingly, cytokines production and response to interferon-gamma (IFN-γ) pathways were upregulated in every activated microglia population in our AGE+PB data (Fig. 2e) and multiple activated microglia populations in the TRH data (Fig. S3f). We also observed various type I interferon pathways uniquely enriched in IRM upregulated genes (Fig. 2e) whereas, several neurodegenerative disease pathways were uniquely upregulated in DAM, including Alzheimer’s disease (AD), Parkinson’s disease (PD), Huntington disease (HD) and amyloid lateral sclerosis (ALS) (Fig. 2f). IFN-γ has been shown to play a critical role in regulating microglial activity and inflammatory response^50,51^. Induction of IFN-γ leads to the activation of Type I interferon, TGFβ, and other cytokines which can either synergize or antagonize IFN-γ function depending on the context^52–54^. It has been shown that certain combinations of cytokines induce a synergistic immune state which is distinct from that induced by either one alone and can be preferentially used in specific biological pathways^55^. For example, IFN-γ synergizes with Toll-like receptor (TLR) signaling and drives TLR-activated microglia into neurotoxic phenotypes that can induce severe neural dysfunction and cell death^51^. IFN-γ can lead to induction of many cell surface molecules and chemokines, such as MHC class II^56^, *Icam-I*^57^, *Bst2*^58^, *Cxcl16*^59^, and *Ccl6*^60^ depending on the cell type and the presence of additional signals. Interestingly, these molecules and *Tlr2* were uniquely enriched in different microglial subpopulations in both datasets (Fig. S3e).

To better understand the signaling pathways that control microglial activation, we leveraged the recently published Immune Dictionary^61^. Cui *et al.* systematically profiled single-cell transcriptomes of more than 17 immune cell types in response to each of 86 cytokines (>1,400 cytokine–cell type combinations) in mouse lymph nodes *in vivo* and created the Immune Dictionary^61^. The accompanying software, Immune response enrichment analysis (IREA) enables the inference of immune cell polarization and cytokine responses from transcriptomic data^61^. We queried the Immune Dictionary using the subpopulation DEGs to infer the potential cytokines which elicited the transcriptomic responses for each microglial subpopulation. As expected, IRM showed highest enrichment of type 1 interferon (IFN-a1 and IFN-b) response (Fig. 2g). Interestingly, adiponectin (AdipoQ) was the top enriched cytokine response in all activated microglia populations (Fig. 2g, S4a). Adiponectin is a circulating hormone predominantly expressed in adipocytes, which can enter the brain by crossing the blood–brain barrier (BBB) and modulate microglia-mediated neuroinflammation with both neuroprotective and anti-inflammatory effects^62–65^. Elevated circulating adiponectin levels lead to reduce age-related tissue inflammation and a prolonged healthspan and median lifespan whereas adiponectin deficiency accelerates brain aging with significantly shortened lifespan in mice^66,67^. Prolactin (PRL) response was also highly ranked in almost all microglial subpopulations. It has been studied extensively and has been shown to mediate both proinflammatory and anti-inflammatory effects in a variety of cell types including microglia^68^. These dual effects are highly influenced by the type of target cells and the molecular composition of the milieu. Prolactin can mitigate deficiencies of retinal function associated with aging^69^. Response to pro-inflammatory cytokine IL-11 was also enrichment in multiple activated microglial subpopulations including IntM, eDAM, DAM, and MHCIIM. In a recent study using a range of genetic and pharmacological approaches, IL-11 signaling has been shown to have a negative effect on healthspan and lifespan in mice^70^. Consistent with different combinations of cytokine responses in different microglial subpopulations, distinct combinations of transcription factors, transcription coactivator activity, and NF-KappaB activity regulatory pathways were enriched in different microglial subpopulations which could be responsible for their distinct cytokine and chemokine expression profiles and responses (Fig. S4b). In summary, this analysis suggests a combinatorial signaling code associated with the different activation states of microglia.

### Regional commonalities and differences in age-related transcriptomic changes

Next, we investigated age-related transcriptomic changes in microglia. Because TRH data were obtained from the whole brain, we first analyzed our AGE data combining all four brain regions to identify genes differentially expressed between microglia from aged mice and the young mice (aging DEGs) and then compared with the TRH data. Although there are fewer cells in our AGE data (5579 cells) compared to the TRH data (22411 cells), we identified more aging DEGs (2181 DEGs) compared to the TRH data (63 DEGs) likely due to the deeper sequencing depth of our data (Fig. S3a, S5a, b). Over 95% of the genes were detected in larger proportions of cells in our AGE data than in the TRH data (Fig. S5c), further demonstrating the higher sequencing depth of the AGE data which likely underlies the higher number of detected aging DEGs. Over 80% of the aging DEGs from the TRH data (51 out of 63 genes, *p* value < 0.01, hypergeometric test) overlapped with those in our AGE data. We also retrieved microglia aging DEGs from Ximerakis *et al.* study which is another single-cell study of brain aging using whole brain tissues (referred to as XIM data)^71^. We found over 75% of the XIM DEGs (37 out of 49 genes, *p* value < 0.01, hypergeometric test) overlapped with those in our AGE DEGs. The XIM data had an average number of UMI (nUMI) and nonzero genes (nFeature) of 2,877 and 1,113, respectively, which represented similar sequencing depth as the TRH data. Our AGE DEGs shared with other datasets in the upregulation of genes such as *Lyz2, Cd52, Cd63, Cd9,* and *Csf1r*^43,71^ during aging (Fig. S5d). Nonetheless, the vast majority of AGE DEGs were only identified in this current study, highlighting the power of deep scRNA-seq technologies.

The high quality of the AGE data and the regional information of the sampled microglia enabled us to investigate regional variation of age-related transcriptomic changes in microglia. We performed DEG analysis comparing microglia of aged mice with those of the young mice from the cerebellum (Cb), cortex (Ctx), hippocampus (Hipc), and striatum (Str) separately and identified 560-1274 genes differentially expressed during aging in different brain regions (referred to as regional AGE DEGs herein) (Fig. S5e, f, Supplementary Table 4). We found that most DEGs were in the upregulation direction during aging and many of the upregulated genes were shared across multiple brain regions (Fig. 3a, b, c, magenta shade). Pearson correlation of gene expression log fold changes showed that the four brain regions had similar transcriptomic dynamics during ageing (Fig. 3d). Consistently, most of the DEGs that were shared across brain regions changed in the same direction (Fig. 3c, magenta shade) including 254 genes that were upregulated in all four brain regions and an additional 237 (128+109) genes shared by at least 3 brain regions (Fig. 3c, e, Supplementary Table 5). Interestingly, 53% of core microglia activation genes (40/75) were overlapped with these shared up regulated aging DEGs (including *Cd9, B2m, Fth1, Selplg, Fcer1g* etc.) across all brain regions (Fig.3e, f), suggesting a concerted increase of the core microglial activation program throughout the brain during aging. A recent study demonstrated that *CD9* induces synapse engulfment by microglia in the thalamus and causes synaptic loss and recognition memory deficits^72^. In addition, AD risk genes (*Abi3, Bin1*) and other critical regulators of microglial gene expression and function (*Irf8* and *C1qa*)^73^ were upregulated in almost all brain regions during aging. On the other hand, we found a set of genes that were downregulated in at least 3 brain regions including multiple insulin pathway related genes such as insulin I and II (*Ins1, Ins2), Egr1, Rgs1* ^74^ (Fig. 3g, Supplementary Table 6). This is consistent with decreased insulin secretion in older compared with younger individuals and the concept that insulin pathway serves as an evolutionarily conserved mechanism controlling aging and longevity ^75,76^. Proper functioning of these genes is required to maintain brain homeostasis and their downregulation may underlie age-related neurodegeneration^77^. We also observed significant downregulation of Neuroplastin (*Nptn*) in three brain regions and trending downregulation in the cortex (Fig. 3f, g). This gene is important for learning and memory and has been shown to be involved in multiple neurological disorders including dementia^78^. Moreover, age-dependent dysregulation of FKBP Prolyl Isomerase 5 (*Fkbp5*) and ATP Binding Cassette Subfamily A Member 1 (*Abca1)* may underlie their roles during aging and cellular senescence^79,80^. In addition to these shared changes, we also observed region-specific gene expression changes, such as upregulation of *Picalm*, *Golm1*, *Nf1* and *Braf* in Cb, Ctx, Hipc and Str respectively, indicating potential roles of microglia in driving differential vulnerability of brain regions in aging and age-related disease^81–84^ (Fig. 3h, Supplementary Table 7). Cb had the highest number of unique upregulated genes followed by Hipc and Str (Fig. 3c, red shade). Some genes upregulated in the Cb were found to be downregulated in the Hipc and/or Str (Fig. 3c, gray shade) consistent with Cb transcriptomic changes being the least correlated with other brain regions (Fig. 3d), and that the Cb ages differentially than other brain regions^44,85,86^.

**Fig. 3:**
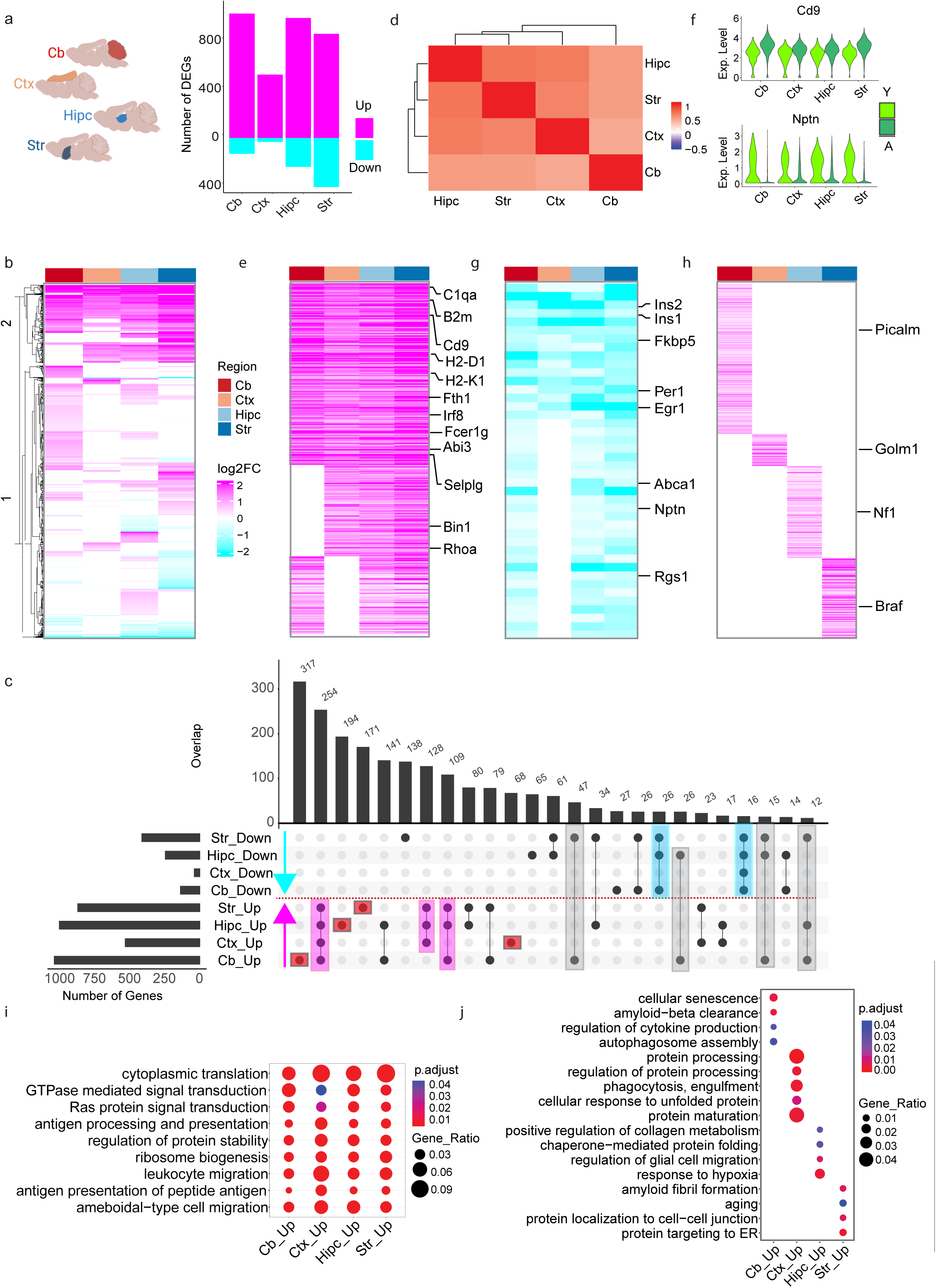
Age-related transcriptomic changes of microglia in different brain regions. **a)** Number of aging DEGs (comparing aged vs young) in the cerebellum (Cb), cortex (Ctx), hippocampus (Hipc), and striatum (Str). Up: upregulated; down: downregulated. DEGs were calculated using Wilcoxon rank sum test and Seurat R package, adjusted P value < 0.05 and absolute Log2FC>0.25. **b)** Heatmap showing the log2FC of aging DEGs. **c**) Upset plot showing overlap of up- and downregulated DEGs during aging. **d**) Heatmap of Pearson’s correlation coefficient of genome-wide gene expression log2FC of different brain regions. The color represents the correlation’s directionality, and the shade of color represents the significant levels. (FDR-correlated P value < 0.05). **e**) Heatmap of up regulated genes shared by at least three brain regions. **f**) Violin plot showing example of up-(*Cd9*) and down-(*Nptn*) regulated genes during aging. **g**) Heatmap of downregulated genes shared by at least three brain regions. **h**) Heatmap of region specific DEGs during aging. **i**) Top three GO terms in the Biological Process category enriched in the aging DEGs. **j**) Region-specific GO terms in the Biological Process (BP) and Molecular function (MF) category enriched in the aging DEGs.

GO enrichment analysis of regional aging DEGs revealed that many top enriched pathways were shared across all brain regions including cytoplasmic translation, antigen processing and presentation, regulation of protein stability (Fig. 3i). Dysregulation of cytoplasmic translation, impaired protein homeostasis and ribosome biogenesis genes have been identified as hallmarks of aged cells^87^. We also observed region-specific enrichment of pathways related to cellular senescence, amyloid metabolism, phagocytosis, hypoxia, and protein trafficking, respectively (Fig. 3j) suggesting divergent roles of microglia in different brain regions during aging. For example, the Cb upregulated genes were uniquely enriched for cytokines production, amyloid beta clearance, and cellular senescence. This is consistent with a previous study showing that microglia in the cerebellum differ from microglia in other brain regions such as cortex, hippocampus and striatum in their transcriptomes and immune function^44,88^. The Ctx upregulated genes showed enrichment of phagocytosis and multiple protein processing and unfolded response pathways. In summary, different brain regions shared a common aging signature but also have their unique transcriptomic changes which may underlie brain selective regional vulnerability.

### Regional variation of parabiosis-mediated transcriptomic changes

To investigate the effect of old blood-mediated transcriptomic change in microglia, we compared gene expression of the young mice of heterochronic parabionts to that of isochronic young parabionts for each brain region separately, referred to as parabiosis-mediated DEGs by old blood (PB-Old DEGs, HY vs. IY). We identified 101-512 PB-Old DEGs, with Cb containing the most and Str displaying the fewest DEGs (Fig. 4a, b, Supplementary Table 8). In contrast to AGE effects, where shared DEGs across regions were prevalent (Fig. 3c), most of the PB-Old DEGs were unique to each brain region pointing to largely region-specific transcriptomic changes when exposed to old blood (Fig. 4d). Interestingly, gene expression fold changes in Str microglia were the least correlated with other brain regions (even a negative correlation with Hipc) (Fig. 4c) suggesting differential and limited effect of old blood on striatum. While a few DEGs in striatum were shared with those found in the Cb and Ctx (Fig. 4d, magenta and blue shades), roughly equal numbers of DEGs in striatum demonstrated changes in the opposite direction compared to DEGs in other brain regions (Fig. 4d, grey shade). These results suggest that Cb, Ctx, and Hipc microglia preserve a certain level of similarity in their response to old blood with Cb being the most sensitive brain region, whereas Str appears to be the least sensitive and display distinct responses to old blood.

**Fig. 4:**
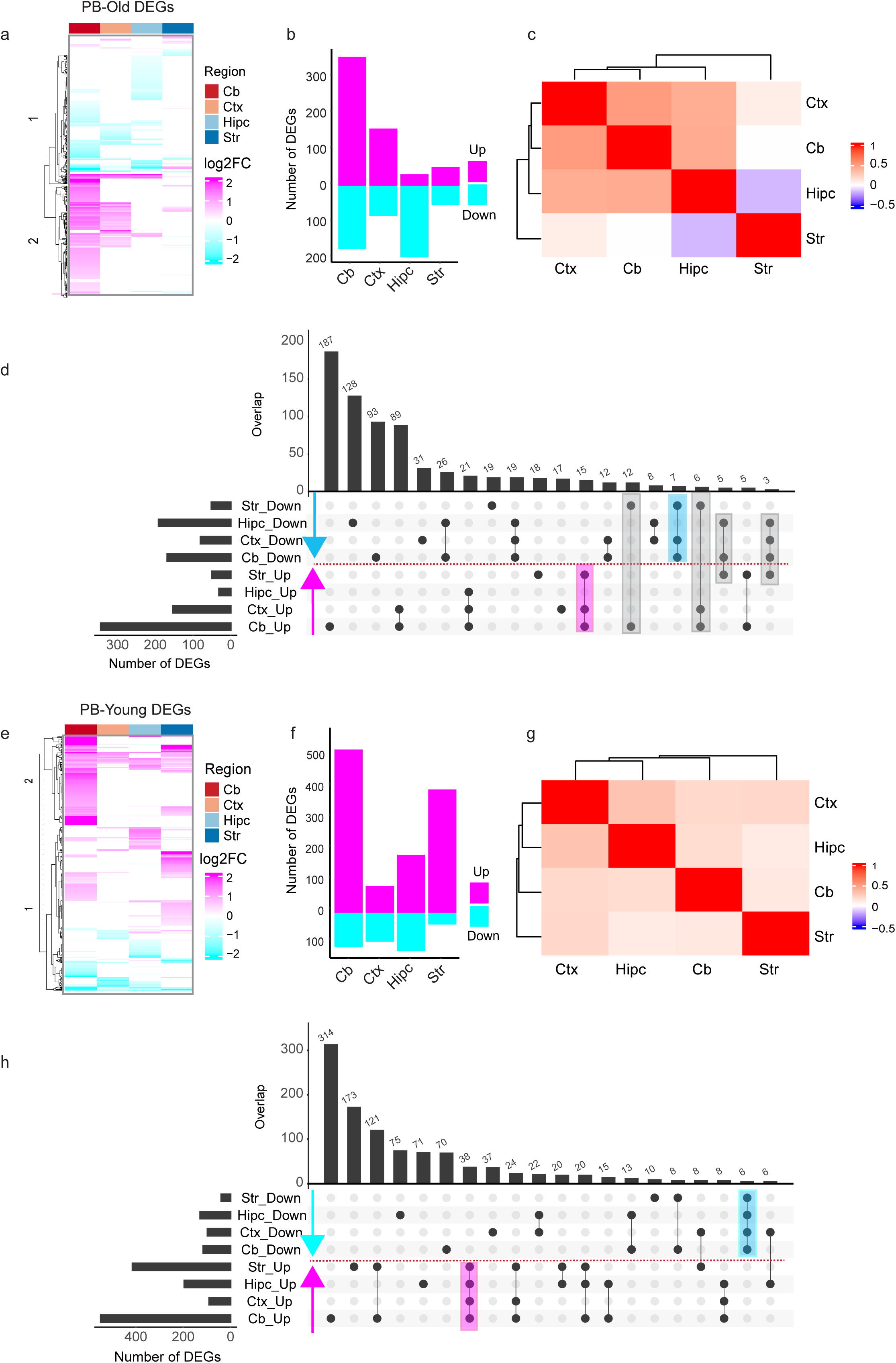
Regional variation of parabiosis-mediated transcriptomic changes. **a)** Heatmap of PB-Old DEGs (DEGs mediated by the exposure of old blood, HY vs IY). DEGs were calculated using Wilcoxon rank sum test and Seurat R package, adjusted P value < 0.05 and absolute Log2FC>0.25. **b**) Bar plot showing number PB-Old DEGs. Up: upregulated; down: downregulated. **c**) Heatmap of Pearson’s correlation coefficient of genome-wide gene expression log2FC of PB-Old DEGs across different brain regions. The color represents the correlation’s directionality, and the shade of color represents the significant levels. FDR-correlated P value < 0.05. **d**) Upset plot showing overlap of up- and downregulated PB-Old DEGs. **e**) Heatmap of PB-Young DEGs (DEGs mediated by the exposure of young blood, HA vs IA). **f**) Bar plot showing number of up- and downregulated PB-Young DEGs. **g**) Heatmap of Pearson’s correlation coefficient of genome-wide gene expression log2FC of PB-Young DEGs across different brain regions. The color represents the correlation’s directionality, and the shade of color represents the significant levels. FDR-correlated P value < 0.05. **h**) Upset plot showing overlap of up- and downregulated PB-Young DEGs.

Likewise, we investigated the effect of young blood in old mice by comparing microglia in the old mice of heterochronic parabionts to microglia in isochronic old parabionts for each brain region separately, referred to as parabiosis-mediated DEGs by young blood (PB-Young DEGs, HA vs. IA). We identified 187-663 differentially expressed genes that were largely distinct to each brain region (Fig. 4e, f, Supplementary Table 9). Again, Cb had the most DEGs suggesting that Cb is the most sensitive brain region to young blood (Fig. 4f). Interestingly, Str remained to be the brain region in which gene expression fold changes were the least correlated with other brain regions (Fig. 4g), even though the striatum had a similar number of DEGs compared to the Hipc and more DEGs than the Ctx. The high number of unique DEGs and the low correlation of gene expression changes with other brain regions suggested that Str also had distinct gene expression changes when exposed to young blood compared to other brain regions. Despite these differences, we found that 38 genes (Fig. 4h, magenta shade), including *Pfn1, Oaz1, Ctsz,* and *Tmsb4x*, were upregulated in all brain regions, when exposed to the young blood. On the other hand, 6 genes (Fig. 4h, blue shade), including *Rn7sk, Lars2 and* Sun2, were downregulated in all brain regions. Previous studies have showed that knockdown of *Rn7SK* in mesenchymal stem cells leads to delayed senescence, while its overexpression shows the opposite effects^89^. Taken together, our analysis demonstrated regional heterogeneity of microglia in response to young or old blood exposure, where cerebellum is the most sensitive brain region by showing the biggest changes and the striatum exhibits the most distinctive response compared to other regions.

### ACC: Recapitulating age-related changes with reduced intensity

To explore how exposure to old blood may lead to accelerated aging in microglia, we extracted the genes shared by PB-Old DEGs and AGE DEGs (referred to as parabiosis-mediated accelerated aging or ACC DEGs) (Fig. 5a, purple dots, and Fig. S6a, b). We identified 80 to 297 ACC DEGs in each brain region (Fig. 5b, red and black circles). Both Cb and Ctx had much more ACC DEGs compared to Hipc and Str (Fig. 5b). Additionally, Cb and Ctx contained higher percentages of ACC DEGs that showed changes in the same direction with AGE DEGs (81% and 93%, respectively), compared to Hipc and Str (42% and 41% respectively) (Fig. 5c). Since only the genes that change in the same direction during young cell exposure to old blood (PB-Old) and chronological aging (AGE) align with accelerate aging effect of plasma factors (Fig. S7a), this data suggests that Cb and Ctx are the two brain regions more affected in young mice by old blood exposure (bigger ACC effect). Next, we were curious about the strength of this pro-aging effect by old blood compared to true biological aging. We focused on ACC DEGs that showed changes in the same direction in Cb and Ctx (Fig. 5d, S7a). Fold changes of these genes were highly correlated between PB-Old and AGE conditions for both Cb (Pearson’s correlation coefficient r = 0.81) and Ctx (Pearson’s correlation coefficient r = 0.68), suggesting overall concerted changes among these genes. Interestingly, we found that multiple DAM signature genes (*Cd9, Fth1, Ftl1, B2m, Tyrobp, Ctss*) were upregulated in both AGE and PB-Old (Fig. 5d, Fig. S7b). Absolute gene expression fold changes were significantly higher for AGE than PB-Old (Fig. 5e, *t* test, *p* value < 2.2e-16), suggesting that old blood-mediated aging was less pronounced than true biological aging. When comparing the effect of ACC between Cb and Ctx, Cb had a larger slope (0.74 in Cb vs. 0.61 in Ctx) (Fig. 5d). Consistently, using the PB-Old over AGE ratio of gene expression log fold changes to measure the strength of ACC effect relative to biological aging, we found that Cb was prone to a significantly larger ACC effect than Ctx (Fig. 5f, *t* test, *p* value< 0.05), suggesting that microglia from Cb were more preferentially affected by old blood in young mice. GO enrichment analysis demonstrated that multiple neuroinflammation and microglia activation-related pathways were enriched mostly in the Cb and less so in the Ctx (Fig. 5g). Taken together, old blood-mediated accelerated aging leads to similar, but to a lesser extent, microglial gene expression changes as in biological aging, and there is a significant regional difference in ACC effect with Cb being the most sensitive brain region.

**Fig. 5:**
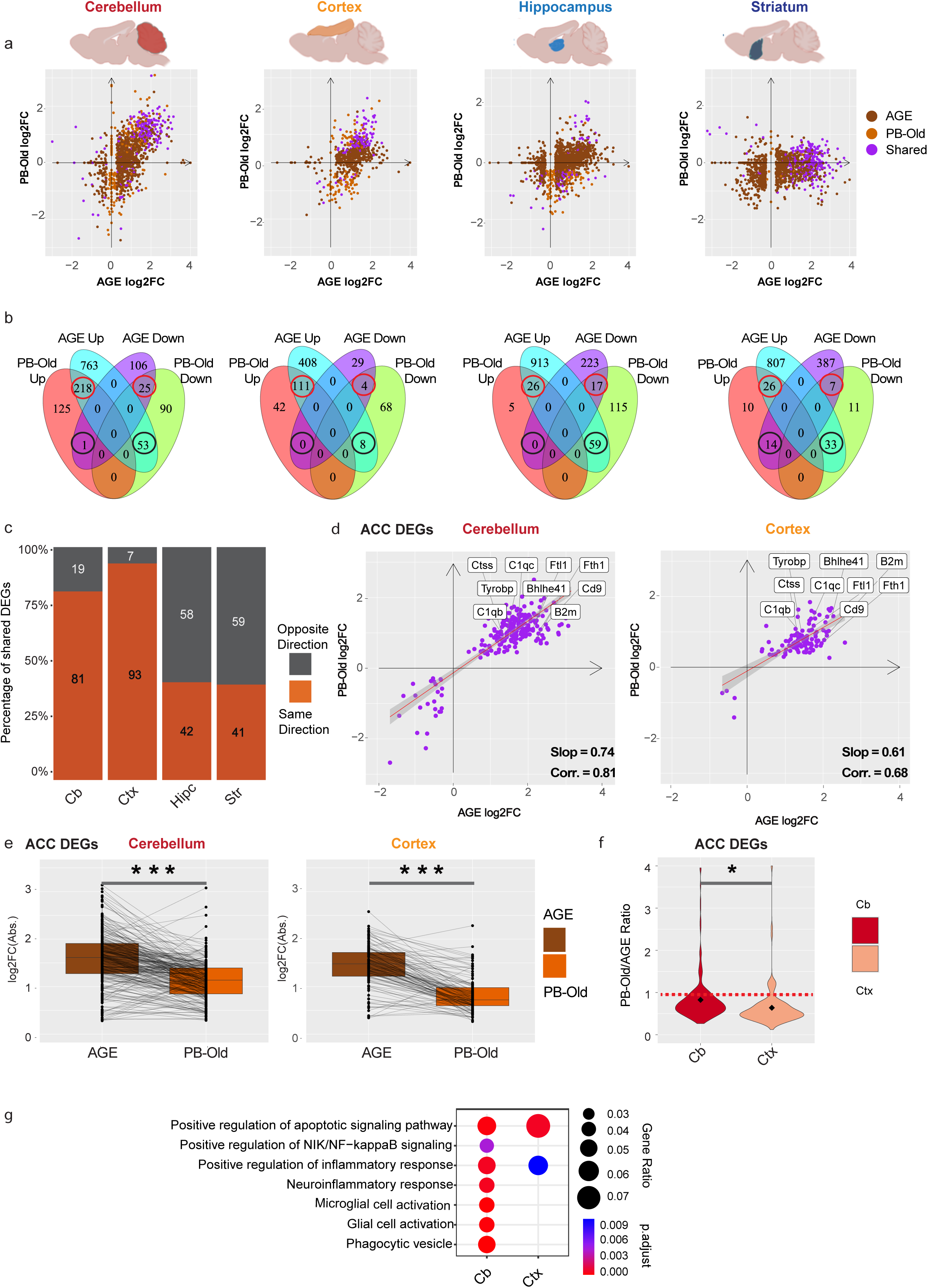
Evaluation of accelerated aging (ACC) effects by comparing transcriptomic changes in aging and the exposure to old blood. **a)** Scatter plot comparing log2FC of PB-Old DEGs vs AGE DEGs (dark brown = AGE DEGs only, light brown = PB-Old DEGs only, purple = shared DEGs between AGE and PB-Old). **b**) Venn diagram showing the number of DEGs unique and shared between PB-Old and AGE DEGs (red circles; shared DEGs with the same direction of change, black circles; shared DEGs in opposite direction of change). **c**) Percentage of shared DEGs with the same (brown shade) and opposite (grey shade) direction of change in each brain region. **d**) Scatter plot showing log2FC of shared DEGs between PB-Old and AGE in the cerebellum and cortex. **e**) Box plot showing Absolute (Abs.) Log2FC of shared DEGs between AGE and PB-Old (paired *t test,* \*\*\**P* < 0.001, \*\**P* < 0.01 \**P* < 0.05). **f**) Ratio of Abs. log2FC of PB-Old to AGE DEGs. **g**) GO analysis showing biological process (BP) terms enriched for shared DEGs between AGE and PB-Old in the cerebellum and cortex.

### REJ: Combatting the effects of aging by young blood

To investigate how exposure to young blood leads to reversal of aging effects in microglia, we identified the genes shared by PB-Young DEGs and AGE DEGs that exhibited opposing directionality (referred to as parabiosis-mediated rejuvenation or REJ DEGs) (Fig. S8a). We identified 3-41 REJ DEGs for the four brain regions (Fig. 6a, S8b). Cb contained the highest number of REJ DEGs (41 genes) followed by Ctx and Hipc (25 DEGs in each), and Str had the least REJ DEGs (3 genes). Cb was also the only region showing higher absolute log fold changes of gene expression in PB-Young compared to AGE (Fig. 6b, Fig. S8c). Using the PB-Young over AGE ratios of gene expression log fold changes to measure the effect of rejuvenation relative to biological aging, we found that, again, Cb had the biggest effect of rejuvenation compared to Ctx and Hipc with the PB-Young vs. AGE ratio >1 (Fig. 6c). These results suggest that Cb is the most sensitive brain region in response to young blood, and blood-mediated rejuvenation can reverse some aspects of biological aging, particularly in Cb (even overshooting in reversal of aging). We found very few REJ DEGs shared between different brain regions (Fig. 6d), with only one gene, encoding a lysosomal gene *Prosaposin* (*Psap*), shared across three brain regions. We observed upregulation of *Psap* during aging but downregulation during blood-mediated rejuvenation in Cb, Ctx and Hipc (Fig. 6e). *Psap* is a multifunctional neuroprotective secreted protein and its levels increase in neurons in experimental models of neuronal injury^90^. Knockdown of *Psap* causes the formation of lipofuscin, a hallmark of aging^91^. It plays a role in maintaining lipid homeostasis in dopamine neurons and counteracting experimental parkinsonism in rodents^92^. The function of *Psap* in microglia in the context of aging remains unclear. Furthermore, we found 5 REJ DEGs shared between Cb and Hipc including *Sipa1l2*, *Sall3* (AGE-up, PB-Young-down) as well as *Fos*, *Jun*, *Dusp1* (AGE-down, PB-Young-Up). *Sipa1l2* has been reported to be a risk gene for age-related neurodegeneration including AD and PD^93,94^. On the other hand, we identified many region-specific REJ DEGs showing upregulation in AGE and downregulation in PB-Young, which have known functions in aging. For example, in Cb, dedicator of cytokinesis (*Dock2* and *Dock10*) have been reported to regulate microglia-mediated inflammatory processes during aging^95,96^. In Ctx, telomere repeat-binding factor 1 (*Trf1*) plays a role in promoting telomere shortening, a hallmark of cellular senescence^97^. Interferon gamma receptor (*Ifngr1)* mediates signals causing neuroinflammation^98^. In Hipc, multiple microglia activation related genes (*H2-D1, H2-K1, Wnk1*)^41,99^ as well as phospholipase C γ (*Plcg2*), were observed as REJ DEGs^100^. Upregulation of these genes during aging may be associated with brain dysfunction and reversal of these changes by young-blood exposure may generate beneficial outcomes in that particular brain region. GO enrichment analysis demonstrated that young blood-mediated rejuvenation modified distinct pathways including synaptic plasticity and phagocytic activity across different brain regions (Fig. 6f). Overall, these results suggested that young blood-mediated rejuvenation have distinct region-specific effects on microglial gene expression with the most pronounced effect seen in the Cb, moderate effects in the Ctx and Hipc, and the least in the Str.

**Fig. 6:**
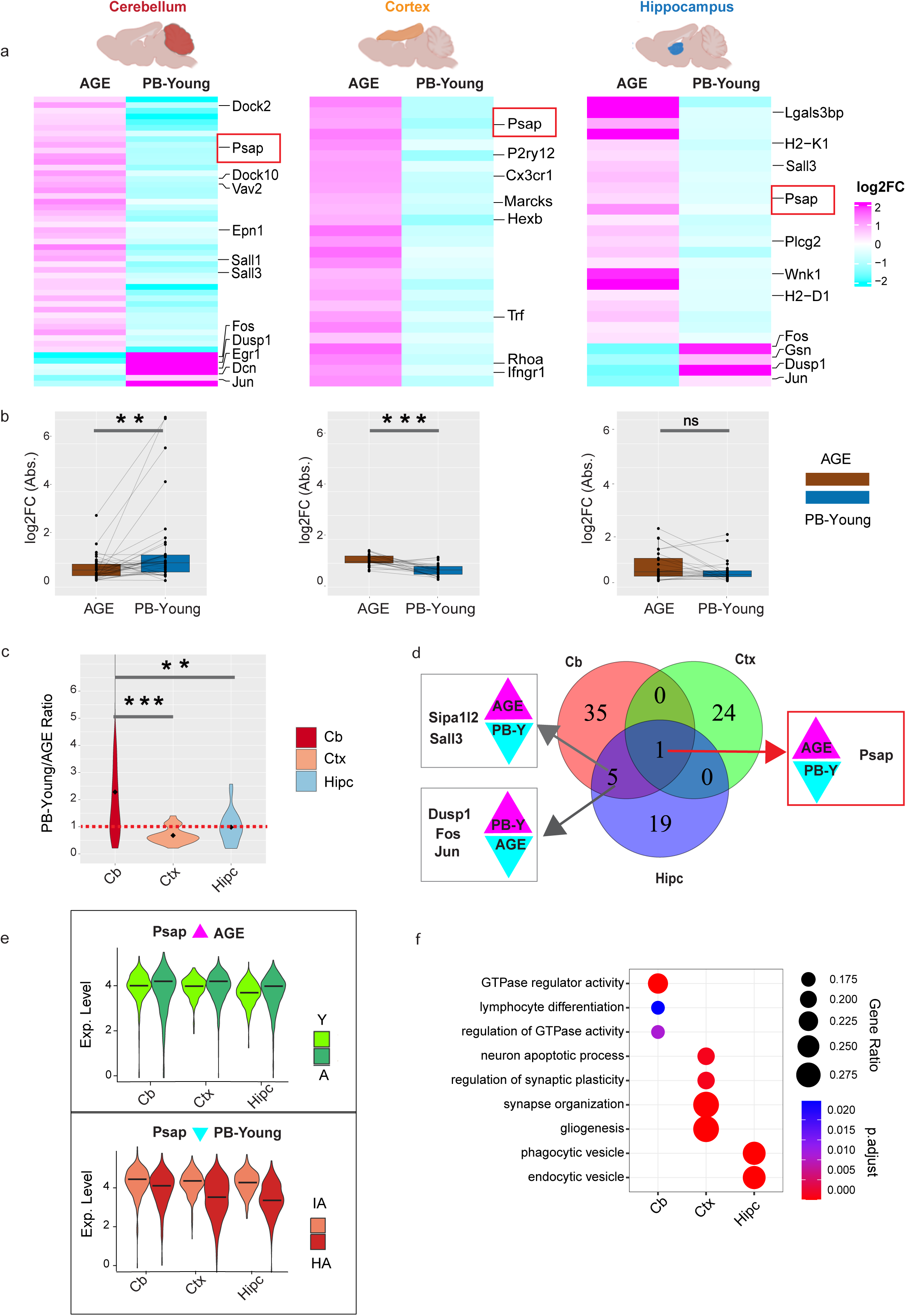
Evaluation of rejuvenation (REJ) effects by comparing transcriptomic changes in aging and the exposure to young blood. **a)** Heatmap showing log2FC of AGE DEGs that were reversed by PB-Young (referred to as REJ DEGs). **b**) Box plot showing absolute (Abs.) Log2FC of REJ DEGs (paired *t test,* \*\*\**P* < 0.001, \*\**P* < 0.01 \**P* < 0.05). **c)** Ratio of log2FC PB-Young to AGE DEGs. **d**) Venn diagram demonstrating overlap of REJ DEGs among the three brain regions (Cb, Ctx and Hipc). **e**) Violin plot showing the expression of *Psap* gene in AGE and PB-Young condition. **f**) GO terms in the Biological Process (BP) and Molecular function (MF) category enriched for REJ DEGs.

Surprisingly, we also observed upregulation of several microglial homeostatic genes (*P2ry12, Cx3cr1, Sall1, Sall3, Hexb*) during aging and downregulation of them during young-blood mediated rejuvenation in Cb, Ctx, and Hipc (Fig. 6a). To validate this observation, we reanalyzed a recently published transcriptomics profiling dataset ^101^ covering 15 brain regions across the mouse lifespan (from 3 to 28 months old). Consistent with our current study, all these homeostatic genes in this dataset exhibited varying degrees of upregulation at least in some brain regions examined during aging (Fig. S8d).

### Effects of AGE, ACC, and REJ at the microglial subpopulation level

As we observed regional differences of microglial transcriptomic changes in aging and parabiosis mediated exposure of old and young blood, we further sought to examine these differences by microglial subpopulations and asked which subpopulations were the driving factors. We focused on five microglial subpopulations: HS, IntM, eDAM, DAM, and IRM, each containing a sufficient number of cells. We first calculated DEGs in each condition (i.e. AGE, PB-Old, and PB-Young) as described above for each subpopulation in each brain region separately (Fig. S9). Gene expression fold changes of AGE DEGs were highly correlated across all subpopulations and brain regions suggesting similar effects of aging contributed by all microglia subpopulations in different brain regions (Fig. 7a, AGE). Subpopulation level changes were consistent with what we observed in the whole population level analysis (Fig. 3), suggesting concerted transcriptomic changes across microglia subpopulations and brain regions during aging. However, in parabiosis-mediated transcriptomic changes by old blood, Str had the most distinct transcriptomic changes compared to other brain regions (Fig 7a, PB-old). Most of the subpopulations had negative correlations of gene expression fold changes with microglia subpopulations in Hipc and Cb, and they had positive correlations with only two subpopulations (IntM and IRM) in Ctx. These data were reminiscent of our observations in the global level analysis (Fig. 4a-c), suggesting that striatal microglia, regardless of their states, respond to old blood differently compared to microglia from other brain regions. In contrast, parabiosis-mediated transcriptomic changes by young blood were highly correlated across all subpopulations and brain regions, suggesting similar effects of young blood on microglia across all microglia subpopulations and brain regions (Fig. 7a, PB-Young).

**Fig. 7:**
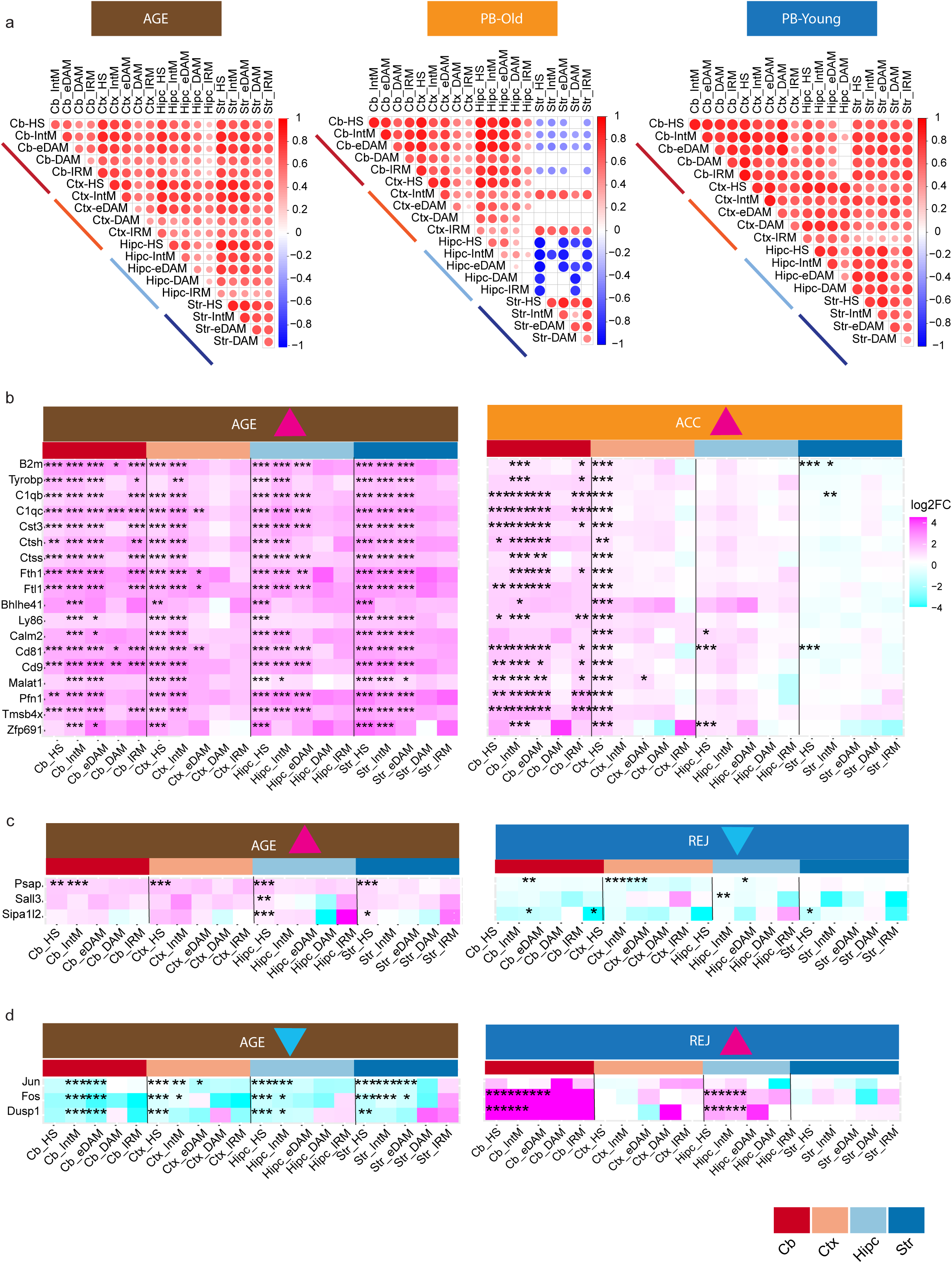
Effects of AGE, ACC, and REJ at the microglial subpopulation level. **a)** Heatmap of Pearson’s correlation coefficient of genome-wide gene expression log2FC of AGE, PB-Old and PB-Young DEGs at subpopulation level in each brain regions. The color represents the correlation’s directionality, and the shade of color represents the significant levels. FDR-correlated P value < 0.05. **b**) Heatmap showing log2FC of some important shared aging genes between AGE and PB-Old condition in each brain region at subpopulations level. (\*\*\**P* < 0.001, \*\**P* < 0.01 \**P* < 0.05). **c**) Heatmap showing log2FC of REJ DEGs shared between AGE and PB-Young condition in each brain region at subpopulation level.

Next, we zoomed in on individual DEGs to examine contributions of different microglial subpopulations from each brain region to the ACC and REJ effects. For ACC, we focused on genes shared by Cb and Ctx (Fig. S7a), excluding ribosomal and mitochondrial genes. Interestingly, many of these genes were DAM signature genes which have established roles in aging and age-related disease. Again, we found upregulation of these genes during AGE in nearly all microglial subpopulations across regions (Fig. 7b, left panel). During ACC, Cb microglial subpopulations showed the strongest upregulation, followed by Ctx microglia (mainly in HS). Hipc microglial subpopulations mainly showed trending significance in the upregulation of these genes, whereas Str microglial subpopulations displayed the opposite trends (Fig. 7b, right panel). These data support our conclusions that Cb microglia are the most sensitive and Str microglia are the least sensitive to old blood-mediated ACC (Fig. 5), and biological aging had stronger effects on microglial gene expression than the exposure to old blood. Meanwhile, these data also suggest that such effects are driven by changes across multiple microglial subpopulations rather than a particular state.

Regarding REJ, we focused on shared DEGs (in at least two brain regions) that showed the opposite directionality in AGE vs. PB-Young (Fig. 6d). Only 6 genes fell into this category (Fig. 7c, d). Nonetheless, changes on these genes appeared to be driven by various microglial subpopulations together across brain regions. Altogether, our analyses demonstrated varying sensitivity of microglia from different brain regions to aging and parabiosis which may underlie distinct regional vulnerabilities to age-related diseases.

## Discussion

Through comprehensive analyses of our deep scRNA-seq datasets and cross-validation with another widely used dataset containing much more microglial cells (albeit at shallower sequencing depth)^43^, we benchmarked various microglial states that were reproducibly present in the adult brain. We found all identified microglia states in both AGE and PB data, which suggest that they represent basal-level heterogeneity of microglia (Fig. S1a-c). Microglia play diverse and crucial roles in the healthy brain, encompassing functions such as neuronal activity modulation, synapse elimination, and debris clearance^102^. However, the growing list of microglial states from scRNA-seq experiments and inconsistent nomenclature may hinder research progress without cross-validation^47^. Our analysis provides the field with a robust reference for future investigations requiring microglial state annotations. Many of these activated microglial subpopulations, such as DAM, IRM, and MHCIIM, were initially identified in various neurodegenerative disease models and aging^8,23^. Recent studies demonstrated that IRM in the developing somatosensory cortex actively engulf whole neurons^103^, defining an important role for IRM during critical windows of brain development. This supports the notion that each microglial state may serve unique roles in the brain under normal conditions, contributing to specific cellular functions and exhibiting differential responses to various stimuli and pathological conditions.

Reactive microglial states are defined by unique combinations of differentially expressed genes and often show overlaps in their gene expression across populations. However, to what extent do they overlap remain poorly understood. Here, we identify a core microglia activation signature (Fig. 2c), including 13 genes (*Fth1, B2m, Cd9*, *Fcer1g, Selplg* etc.) that are upregulated in aging, old blood exposure, and across all reactive microglial populations within all brain regions tested (Fig. S7b). This points to a context-independent, conserved mechanism regulating microglial reactivity. *B2M*, *FTH1*, and *CD74* are involved in microglia transition from a homeostatic to an activated stage associated with neurodegeneration in humans and mice^8,21,104^, whereas *CD9*-dependent microglial state plays crucial role in thalamic synapse loss and recognition memory deficits^72^. Given the widespread induction of IFN-γ pathways across most activated microglia population in our AGE+PB data (Fig. 2e) and the TRH data (Fig. S3f) as well as the critical role of IFN-γ in regulating microglial activity and inflammatory response^50,51^, IFN-γ might be a key regulator that drives this core gene program for microglial reactivity^50,51^. This deserves future investigation.

Moreover, *in vivo* tracking of reactive microglial states in disease have demonstrated their extraordinary plasticity^105^. This raises a question about dynamic changes of microglial states in a healthy brain. Here, through 3-D trajectory inference, we uncovered three trajectories of state progression (Fig.1h), again largely reproduced in another independent dataset (Fig. S1g), in which IntM microglia appeared to serve as an intermediate state that transits towards either DAM or IRM. The IntM may act as a transitional immune state with a tug-of-war dynamic between different states. Signals from disease or injury may tip the balance to generate reactive states suitable for the contexts leading to development of DAM in helping injury repair ^105^ or to IRM which exacerbates damages ^40^. Future studies can leverage the plasticity of microglia to enhance their beneficial reactive outcomes.

One interesting observation in our study is selective regional sensitivity of microglia during aging and parabiosis. Cerebellum is consistently the most responsive brain region to aging, young or old blood exposure. Previous studies indicated that the cerebellum ages at a slower rate compared to other brain regions^86^. A growing body of evidence highlights differences between various brain regions, particularly the cerebral cortex and the cerebellum, in structural and functional changes associated with age-related diseases affecting cognition^106^, such as Alzheimer’s disease^107^, frontotemporal dementia^108^, and Parkinson’s disease^109^. The consistently high number of DEGs observed in cerebellum as well as large fraction of these DEGs as DAM signature genes (if DAM are being protective)^8,105^, it might represents a feedback mechanism for the cerebellum to adapt to the challenges. On the other hand, striatum was found the least responsive to parabiosis. Different studies have shown the role of striatum leading to motor function disabilities (hallmarks of PD and HD)^110,111^. Our findings of regional differences of microglial response to aging and perturbations offer new insights into the selective regional vulnerability of brain aging and neurodegenerative disease pathogenesis.

Lastly, young blood seems to have health benefits to rejuvenate the brain, which is a tantalizing agent for potentially slowing down age-related deleterious effects. However, blood is complex, and transferring young blood may also lead to unpredictable outcomes. In our analysis, we focused on genes that were consistent with a presumable rejuvenation effect (changing in the opposite direction between AGE and PB-Young). Despite the short gene list, it does point to potential targets which young blood may act on to induce the beneficial effects in the brain innate immune system. For example, in cortex, we found downregulation of Interferon gamma receptor (*Ifngr1)* (Fig. 6a), that is reported to mediate neuroinflammation signals^98^. Similarly, we found *Psap* gene to be rejuvenating gene shared by cerebellum, cortex and hippocampus. Future studies are required to dissect the underlying role of microglial *Psap* in regulating age-related phenotypes. Our study may help advance initiatives to comprehend and modulate the aging process and in exploring molecular and cellular therapeutic targets for neurodegenerative diseases associated with aging.

### Limitations of the study

Given that region-specific and age-related changes in gene expression may vary among species, the conclusions drawn from mouse data may not be fully translatable to humans. The analysis of our scRNA seq data is all from male mice, potentially lacking subtle sex-specific gene expression differences. We suggest further studies including both sexes to see the effect of systemic factors during aging on murine microglia transcriptomics. Microglial changes observed may also be due to the indirect effects from parabiosis impacting other cells in the brain.

## Acknowledgements

This work was supported by the NIH R03AG070474, R21AG077643, and R01NS123571 in part support of G.Z.. The experimental work was funded by N.L. AHA-Allen Brain Health and Cognitive Impairment Cross-Network Collaborative Grants (23BHCICG1188316). N.L. is a MAC3 Dementia and Ageing Fellow supported by MAC3 Impact Philanthropies. This publication is solely the responsibility of the authors and does not necessarily represent the official view of the National Institute of Health. The funders had no role in study design, data collection and analysis, decisions to publish or preparation of the manuscript.

## Author contributions

G.Z. and Q.L. contributed to study concept, design and supervision. H.N., G.Z., and S.Y. contributed to bioinformatics analysis. H.N., G.Z. and Q.L. contributed to qualitative analysis of data. Q.L., N.L., and R.P., prepared tissue and snRNA-seq library construction. H.N., G.Z., Q.L., N.L., and T.W.C. contributed to critical manuscript revision for important intellectual content. H.N., Q.L., and G.Z., wrote the manuscript, with feedback from all authors. All authors read and approved the final manuscript.

## Methods

### Quality control and cell clustering

Microglia cells were extracted from our published scRNA-seq data from the aging study, *Tabula Muris Senis* (referred to as the “AGE” data herein)^1^ and the parabiosis study (referred to as the “PB” data herein)^2^. We performed further quality control by filtered out cells that were doublets or low quality after unsupervised clustering. Cell clusters with a mixed expression of markers from microglia with markers of other cell types and clusters with low quality were removed, such as clusters with a high percentage of mitochondrial genes. After these quality control procedures, we obtained 12,310 high-quality single cells gene expression profiles and detected a median of 2259 genes and 824,948 transcripts per cell. We used the same parameters for Hammond *et al*^3^ data and obtained 22,411 high-quality single cells with a median 1126 genes and 2157 transcripts per cell.

For integrative analysis, we followed the workflow described in the Seurat V4 guided analysis for performing integration on datasets normalized with SCTransform^4^. We use SCTransform to normalize gene expression levels and to regress out variations from mitochondrial gene expressions. To integrate the single-cell data from individual samples, we used function SelectIntegrationFeatures (nfeatures = 3,000) to identify highly variable genes. Functions PrepSCTIntegration (anchor.features = selected.features), FindIntegrationAnchors (anchor.features = selected.features, normalization.method = ‘SCT’, dims = 1:30) and IntegrateData (anchorset = selected.anchors, normalization.method = ‘SCT’) were implemented. The top 3,000 most variable genes were selected as integration features and used for integration anchor selection. Principal component analysis was performed using the top 30 PCAs. UMAP analysis was performed with the top 30 dimensions. Clusters were identified with the functions FindNeighbors (dims = 1:30) and FindClusters (resolution = 0.35). A resolution of 0.35 was selected for the downstream analysis because clusters were clearly separated and matched visual inspection. Default parameters were used unless noted.

### Microglia subpopulation identification

FindMarkers function (assay = ‘SCT’, slot = ‘data’, test.use = ‘wilcox’, min.pct = 0.2) was used to determine genes differentially expressed in each microglia subpopulation. A gene with a Benjamini– Hochberg (FDR) adjusted *P* value < 0.05 and a log 2 fold change > 0.25 or < −0.25 was determined to be statistically significant. Cluster marker analysis of the microglia subpopulation confirmed they were transcriptionally distinct, with their unique marker genes used to annotate cell clusters by comparing identified marker genes with the expression patterns of known microglia-subpopulation-specific markers. TNFRM clusters were visually separate from the other clusters but were clustered together with DAM. It was manually selected using the CellSelector function of the *Seurat* R package. The following markers were used for microglia annotation: *Cx3cr1, Fcrls, P2ry12, Tmem119 for* homeostatic microglia (HS), *H2-D1, H2-K1, B2m,* and *Lyz2* were used for intermediate activated microglia (IntM). The downregulation of *Cx3cr1, P2ry12, Tmem119,* and upregulation of *Tyrobp, Ctsb, Apoe, B2m, Fth1, Lyz2* were used for early disease associated microglia (eDAM) or stage I DAM whereas *Axl, Cst7,Ctsl, Lpl, Cd9, Csf1, Ccl6, Itgax, Clec7a, Lilrb4a, Timp2,* and *Trem2* were used for disease associated microglia (DAM) as reported ^5^. *Bst2, Ccl12, Ifi204, Ifi27l2a, Ifit3, Ifitm3, Irf7, Isg15, Lgals3bp, Oasl2, Rtp4, Slfn2, Usp18* were used for interferon-response microglia (IRM) ^6,7^ *Vcam1, Cd83, Marcksl1, Tnf, Gpr84, Nfkbia, Icam1, Slamf8, Tlr2* were used for tumor necrosis factor response microglia (TNFRM)^8^. *H2-Oa, H2-Ab1, Cd74, Lyz2, C3ar1, Mafb, Notch1* were used for major histocompatibility complex class II-positive microglia (MHCIIM)^7^. *Mki67, Top2a, Cdk1, Ccnb2, Ccna2, Cdca8, Cdc20, Cdkn1a, Cdca2* were used for cycling (proliferating) microglia (CycM) ^9^. Signature gene enrichment was evaluated using the hypergeometric test implemented in the *phyper* function in the R *Hypergeometric* package with lower.tail= FALSE.

### Trajectory inference

Slingshot^10^ was used to perform pseudotime analysis for trajectory inference. Pseudotime analysis was performed using different numbers of dimensionality from uniform manifold approximation and projection (UMAP) analysis to test the stability of trajectory ranging from the first two dimensions up to ten dimensions with HS microglia as the starting cluster. In its first step, Slingshot identifies trajectories by treating clusters of cells as nodes in a graph and drawing a minimum spanning tree (MST) between the nodes. Trajectories are then defined as ordered sets of clusters created by tracing paths through the MST, starting from a given root node (HS microglia). The final stable trajectory using dimension 1:10 was reported.

### Differential gene expression analysis during AGE, PB-Old and PB-Young

FindMarkers function (assay = ‘SCT’, slot = ‘data’, test.use = ‘wilcox’, min.pct = 0.2) was used to determine differentially expressed genes (DEGs) during AGE (A vs Y), PB-Old (HY vs IY) and PB-Young (HA vs IA) in all brain region combined and in each brain region separately. We used logarithms base = 2 (the default parameter for calculating fold change in the FindMarkers function implemented in Seurat v4.0) for all the comparisons to keep the analysis consistent within the manuscript. A gene with a Benjamini–Hochberg (FDR) adjusted *P* value < 0.05 and a log 2 fold change > 0.25 or < −0.25 was determined to be statistically significant.

### Gene set enrichment analysis

Gene ontology (GO)^11^ enrichment analyses were performed using the R package clusterProfiler v3.16.1 ^12^. Results with FDR-corrected *P* value < 0.05 and at least five query genes were reported as significantly enriched pathways. We performed GO term enrichment analysis under the following three sub-ontologies: biological process (BP), molecular function (MF) and cellular component (CC). The total number of features that were detected at least once in the cell population being analyzed was used as the background gene set in GO enrichment analysis.

### Immune Response Enrichment Analysis (IREA)

The Immune response enrichment analysis (IREA) was used to query the Immune Dictionary^13^ to infer immune cell polarization and cytokine responses from transcriptomic data. Subpopulation DEGs comparing each microglia cluster to the homeostatic microglia (HS) was used as inputs to infer the potential cytokines which elicited the transcriptomic responses for each microglial subpopulation.

### Statistics and reproducibility

R version 4.2.2, R package Seurat version 4.4.0, was used for statistical analysis and plotting (R Core Team (2013). R: A language and environment for statistical computing. R Foundation for Statistical Computing, Vienna, Austria; http://www.R-project.org/). Correlation analysis was performed using the “corr” function in R and “corrplot” package was used for plotting correlation results. Paired *t* test in R was used for two group comparisions. Wilcoxon rank sum test is a nonparametric test, which does not require normal distribution of the data. Our findings were replicated in an independent dataset for microglia subpopulations, suggesting that our findings are representative.

### Data availability

Aging data from The *Tabula Muris Senis* is from NCBI under accession number GSE109774 (https://www.ncbi.nlm.nih.gov/geo/query/acc.cgi?acc=GSE109774). The parabiosis raw sequencing data is available in the Gene Expression Omnibus under accession code <u>GSE132042</u> (https://www.ncbi.nlm.nih.gov/geo/query/acc.cgi?acc=GSE132042). Processed data is available from figshare (https://figshare.com/projects/Tabula_Muris_Senis/64982). Hammonds *et al.,* Aging data for comparison was downloaded from NCBI under accession number <u>GSE121654</u> (https://www.ncbi.nlm.nih.gov/geo/query/acc.cgi?acc=GSE121654). We also retrieved microglia aging DEGs from Ximerakis *et al.*^14^ supplementary table 6 (41593_2019_491_MOESM8_ESM.xlsx).

---

**Fig. S1:**
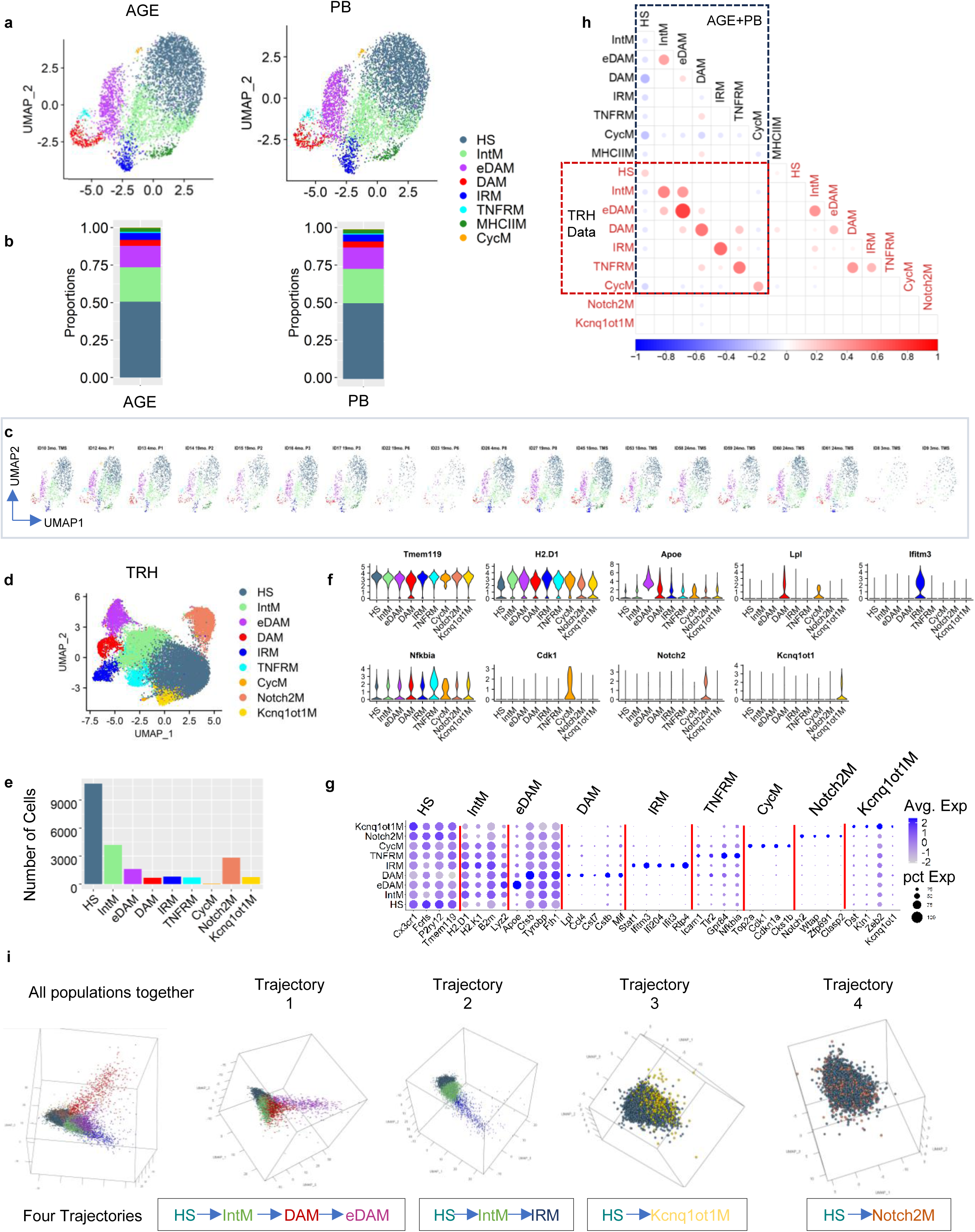
Cross comparison of microglia subpopulations in independent datasets. **a)** UMAP (Uniform Manifold Approximation and Projection) representation of different microglia subpopulation in the AGE+PB data split by datasets (aging dataset=AGE, parabiosis dataset= PB). **b**) Proportion of microglia subpopulations in AGE and PB dataset. **c**) UMAP of microglia subpopulation in the AGE+PB data split by individual samples. **d**) Unsupervised clustering of scRNA-seq aging dataset published by Hammond *et al.,* (referred to as the “TRH” data) and UMAP (Uniform Manifold Approximation and Projection) representation of different microglia subpopulation (total cells= 22,411) colored by cluster identity. UMAP plots were generated using default parameters except reduction = ‘pca’, dims = 1:30, res=0.35. **e**) Number of cells in each of the microglia subpopulation in the TRH data. **f**) Violin plot of known marker gene expression. **g**) Dot plot showing expression of unique marker geness. **h**) Heatmap of Pearson’s correlation coefficient of cluster marker gene expression between the AGE+PB data and TRH data. i) Pseudotime analysis showing four potential microglia activation trajectories for different microglia subpopulations in the TRH data.

**Fig. S2:**
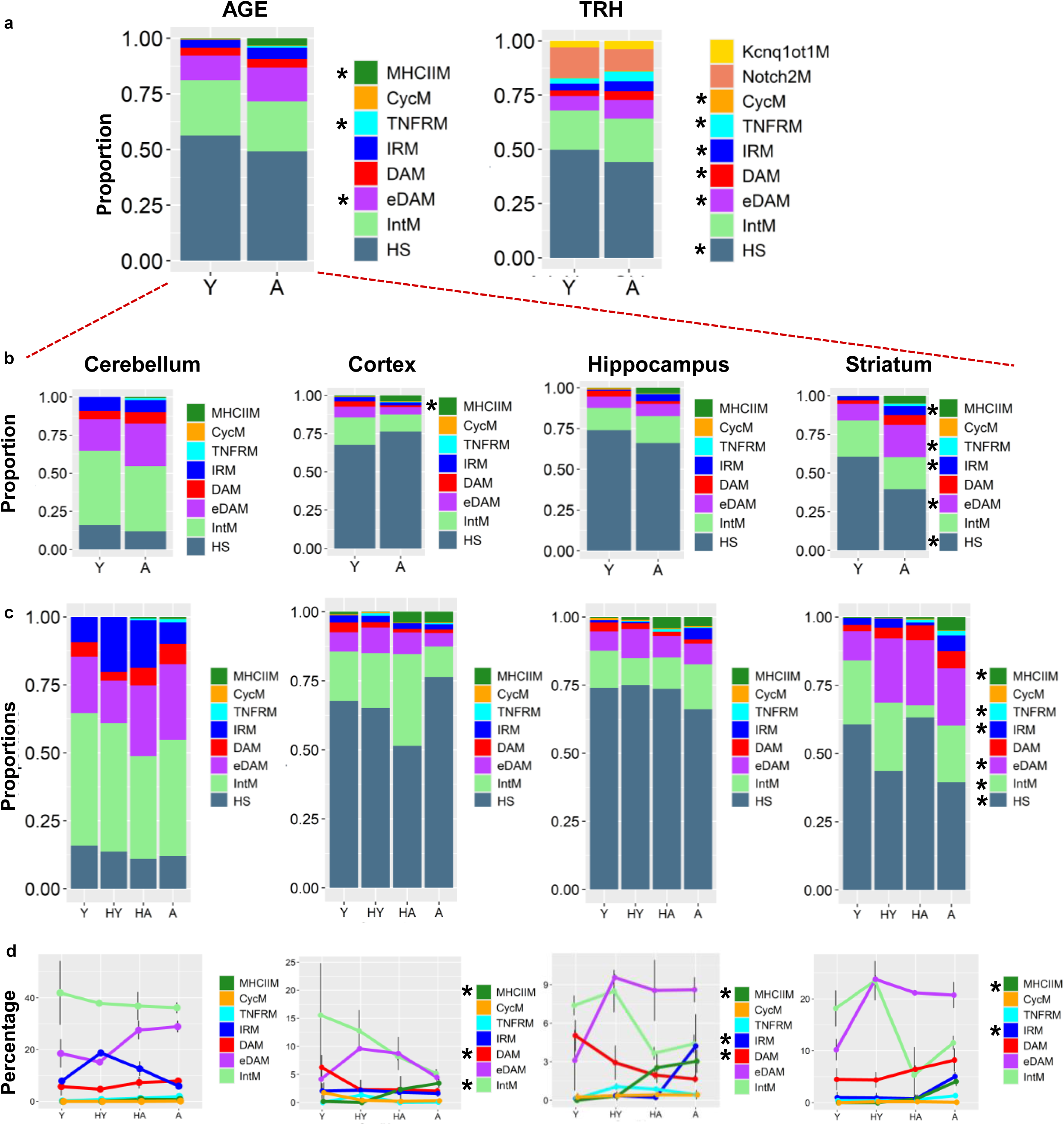
Microglia subpopulations dynamics during aging and rejuvenation. **a)** Stack Bar plot showing the proportion of different microglia subpopulations during aging in AGE and TRH data. **b**) Proportion of different microglia subpopulations during aging in AGE data in each brain region. **c**) Proportion of different microglia subpopulations in young (Y), young blood mediated parabiosis (HY), old blood mediated parabiosis (HA) as well as in old (A) in each brain region. **d**) Line plot showing percentage of changes in microglia subpopulation during Y, HY, HA and A in each brain region. Propeller test and Jonckheere-Terpstra test were used to determine statistical significance.

**Fig. S3:**
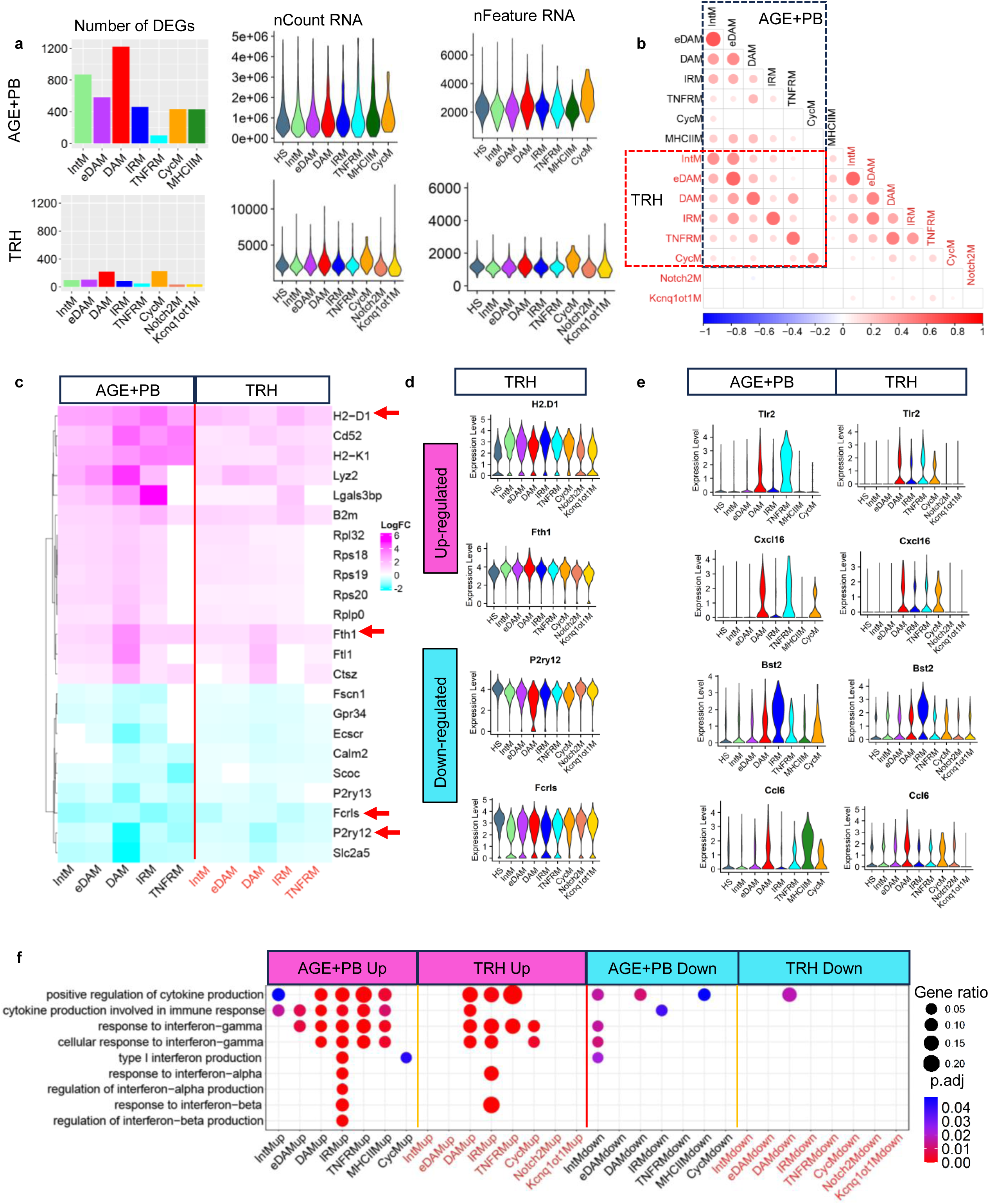
Comparison of genes and pathways associated with microglia activation between the AGE+PB and TRH data. **a)** Bar plot showing the number of DEGs in each microglia subpopulation compared to HS microglia. Violin plots showing nCount and nFeature of the AGE+PB data (upper panel) and TRH data (lower panel). **b**) Heatmap of Pearson’s correlation coefficient of log2FC of DEGs of all microglia subpopulations compared to HS microglia of the AGE+PB and TRH data respectively. **c**) Heatmap of log2FC showing core set of DEGs shared by at least 8/10 activated microglia subpopulations in the AGE+PB and TRH data. **d**) Violin plot showing examples of up- and downregulated genes in activated microglia in TRH data. **e**) Violin plot showing examples of interferon-gamma (IFN-γ) pathways related genes expression in the AGE+PB and TRH data. **f**) GO (Gene Ontology) enrichment analysis showing shared biological pathways across subpopulations of both datasets.

**Fig. S4:**
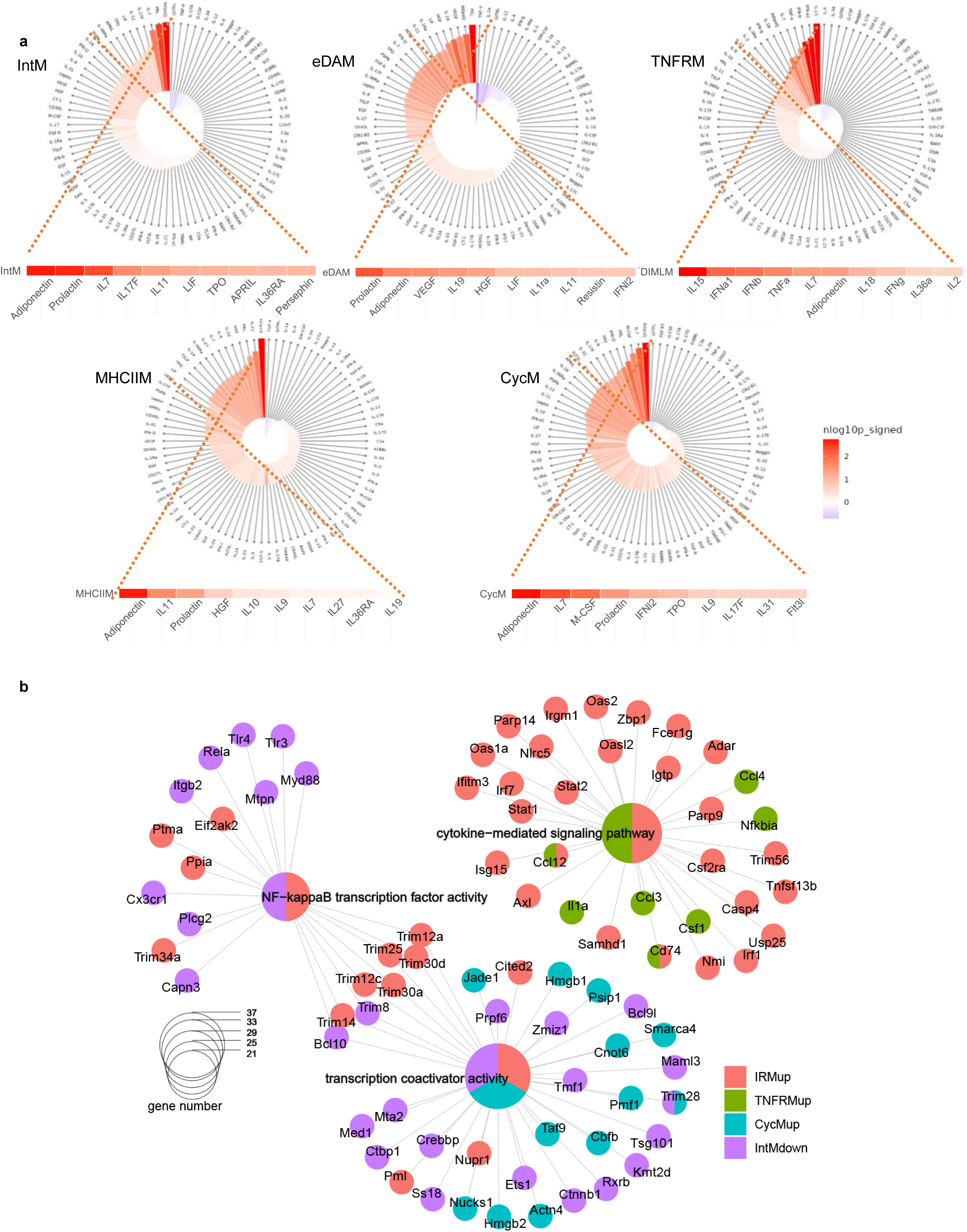
Unique combinations of different cytokines and transcription factors in different microglia subpopulations. **a)** IREA (Immune response enrichment analysis) showing enrichment of top cytokine responses by the DEGs of different microglia subpopulations compared to HS microglia. **b**) Gene regulatory network analysis using DEGs compared to HS microglia. Central nodes showing term enriched (Node sizes reflect the number of genes present in that term), while outer nodes are showing component genes included in the term.

**Fig. S5:**
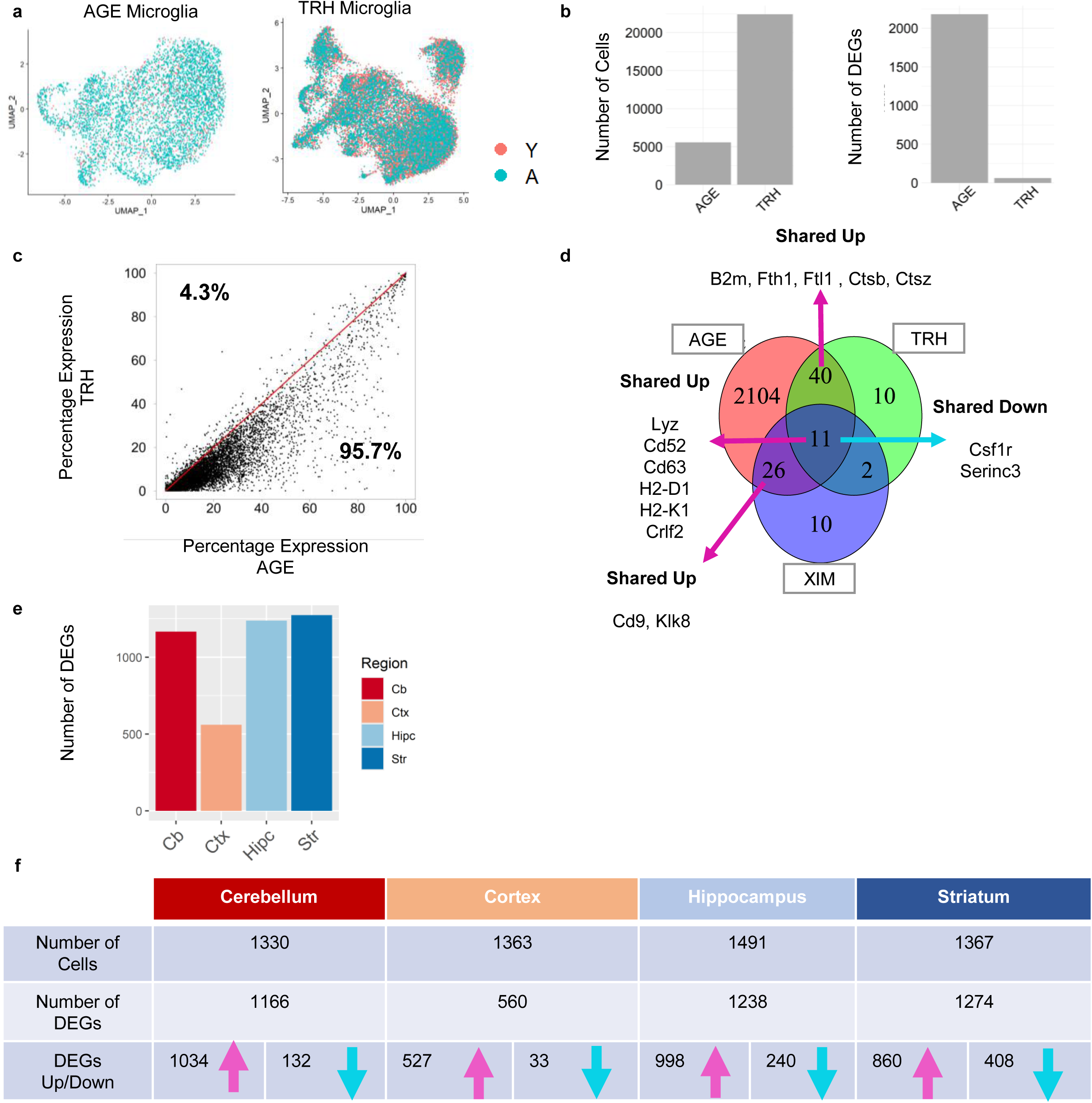
Comparison of age associated transcriptomic changes. **a)** UMAP showing distribution of young and aged cells in the AGE and TRH datasets. **b**) Bar plot showing the number of cells and Aging DEGs (Aged vs young) in the AGE and TRH dataset. **c**) Scatter plot showing percentage of cells with detected gene expression in our AGE data and TRH data. **d**) Venn diagram demonstrating the overlap of aging DEGs across our AGE data, TRH data and XIM (Ximerakis *et al*., 2019) data. **e**) Bar plot showing number of aging DEGs in each brain region in our AGE data. f) Table showing number of cells, up- and downregulated DEGs in each brain region. Cb=Cerebellum, Ctx=Cortex, Hipc=Hippocampus, Str=Striatum

**Fig. S6:**
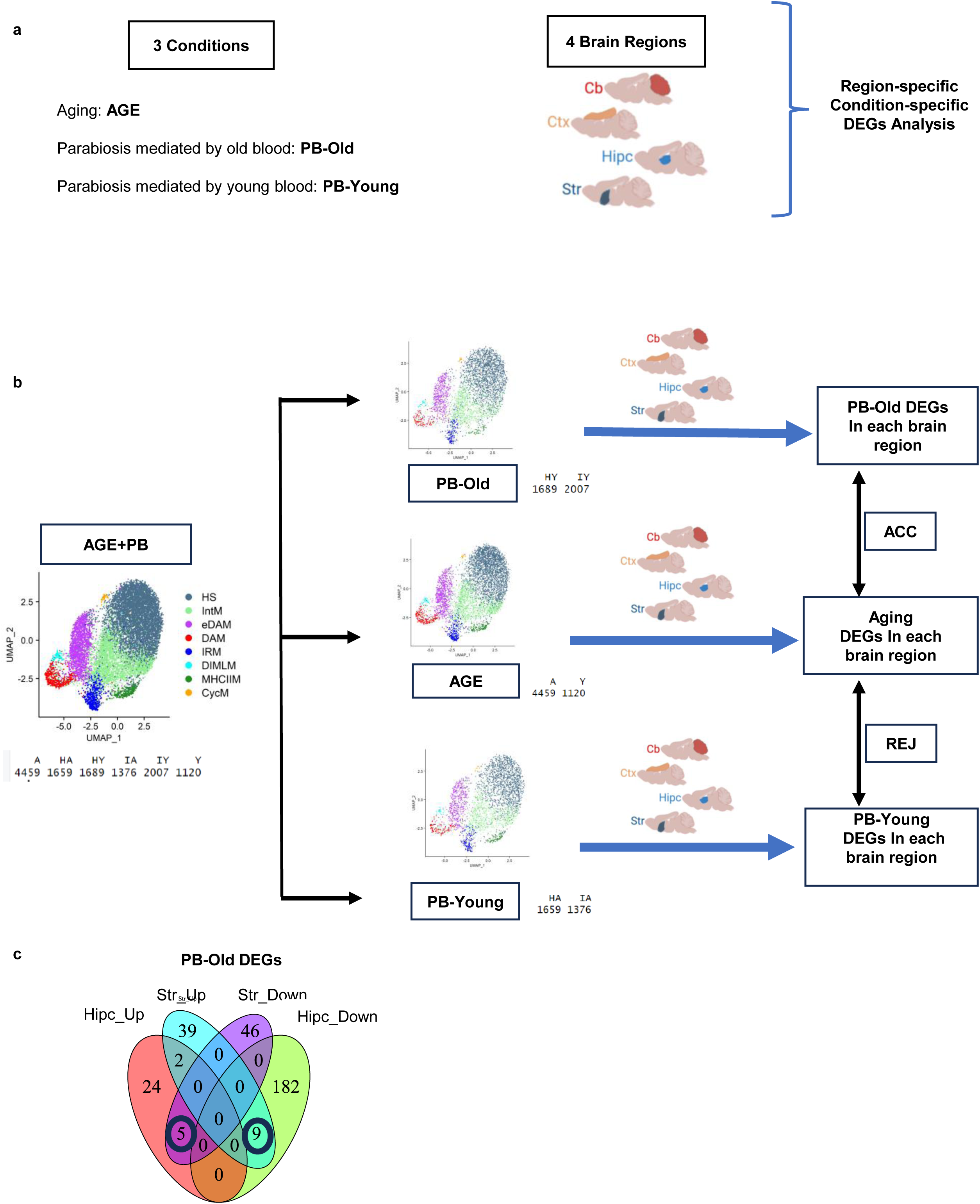
Overview of comparative evaluations across conditions and brain regions in the AGE+PB data. **a)** Three conditions including AGE (aged vs young; A vs Y), PB-Old (heterochronic young vs isochronic young; HY vs IY), and PB-Young (heterochronic aged vs isochronic aged; HA vs IA) and four brain regions under study. **b**) Overview of comparison strategies between different conditions in each brain region. **c**) Venn diagram demonstrating the overlap of PB-Old DEGs between hippocampus and striatum.

**Fig. S7:**
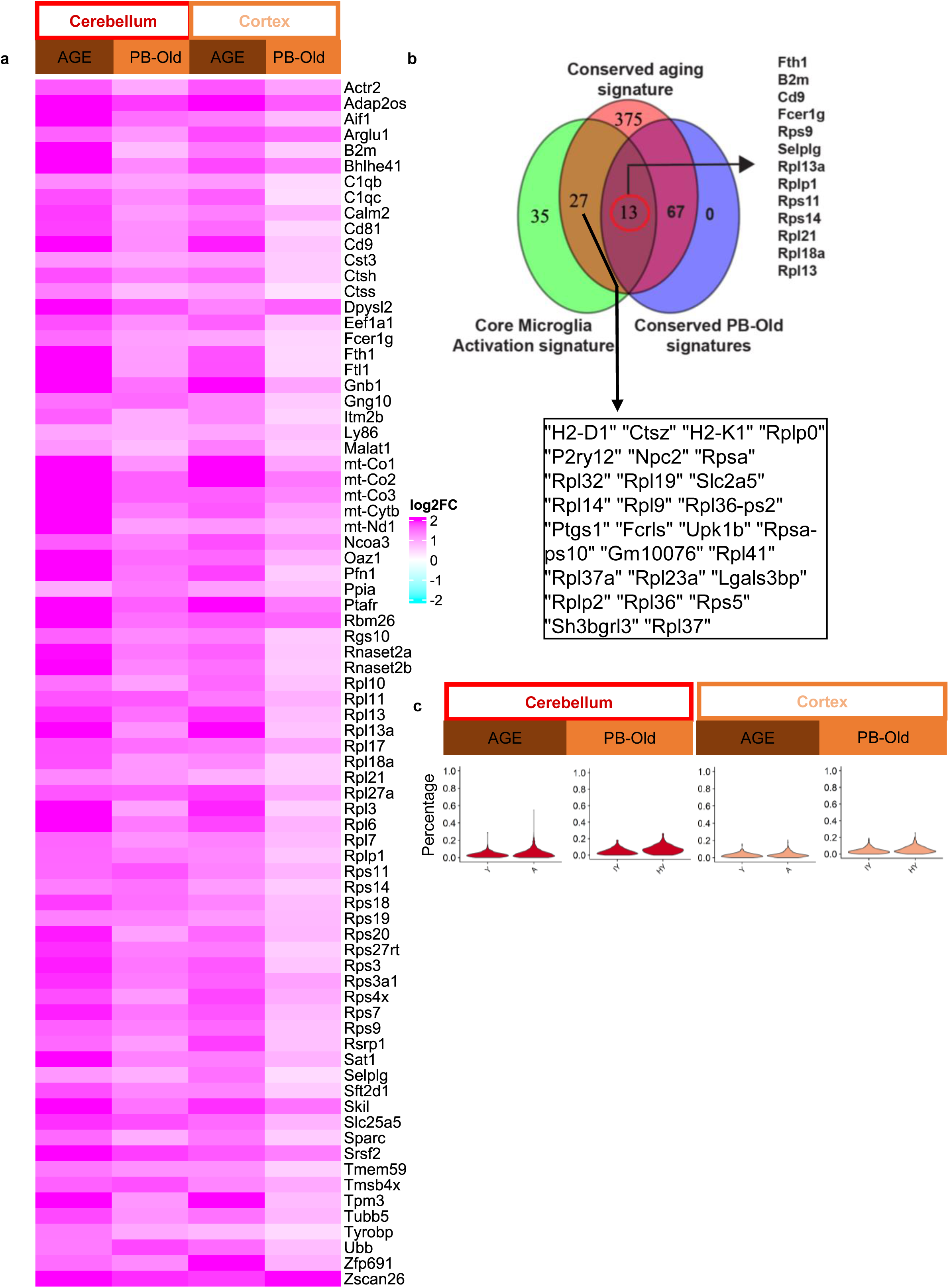
Commonalities between cerebellum and cortex AGE and PB-Old DEGs. **a**) Heatmap showing log2FC of shared upregulated DEGs between cerebellum and cortex in AGE and PB-Old condition. a) Venn diagram demonstrating the overlap of core active microglial activation genes across all microglia subpopulations, conserved upregulated DEGs during aging in all brain region (Cb, Ctx, Hipc, Str), and conserved upregulated genes mediated by old blood exposure between Cb and Ctx. **c**) Violin plot showing percentage of ribosomal genes in cerebellum and cortex in AGE and PB-Old condition.

**Fig. S8:**
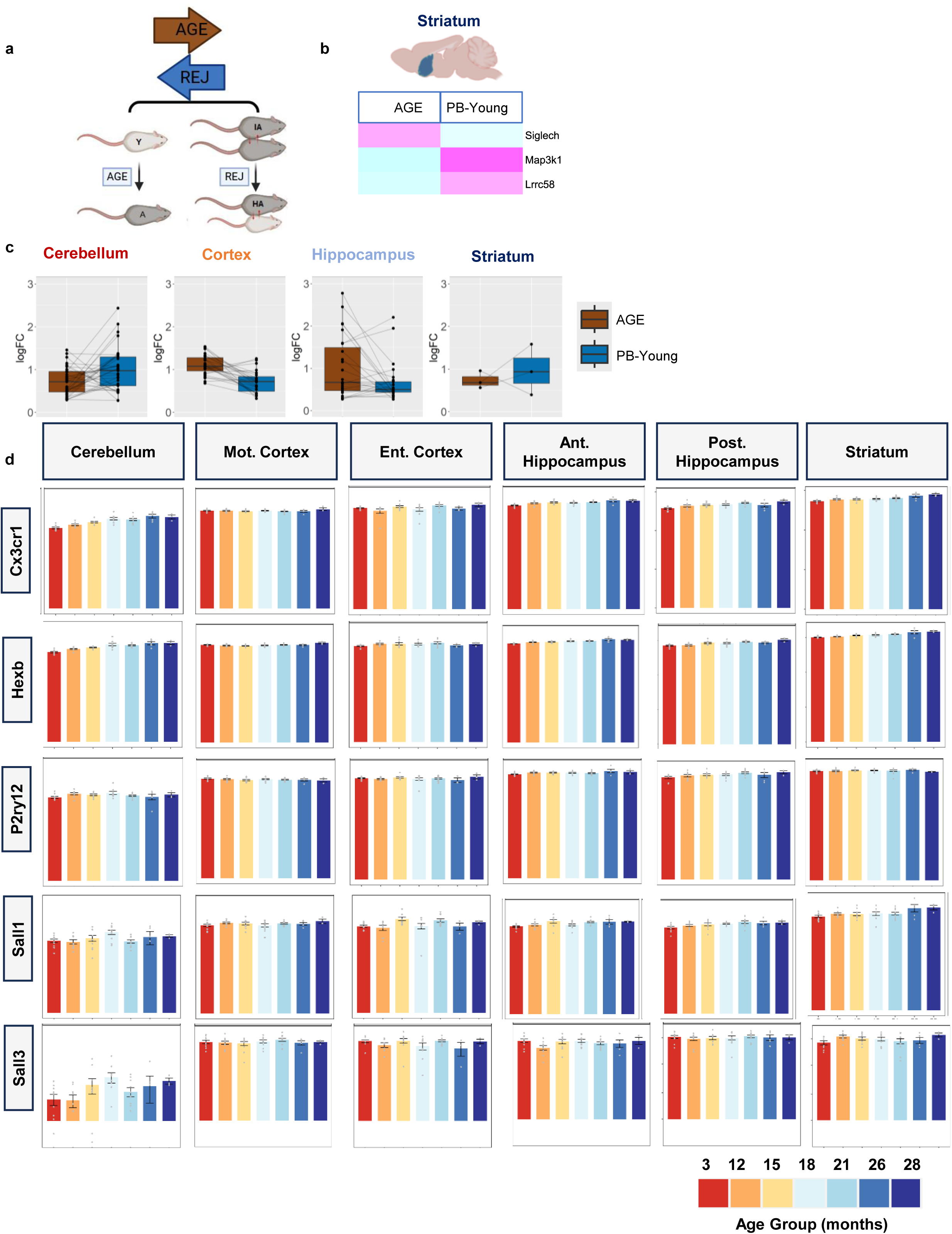
Rejuvenation effects mediated by the exposure to young blood. **a)** Strategy to find genes mediating rejuvenating effects by comparing DEGs with opposite trends in AGE and PB-Young conditions. **b**) Heatmap showing log2FC of AGE DEGs that were reversed by PB-Young (REJ DEGs) in Striatum. **c**) Box plot showing absolute (Abs.) log2FC of REJ DEGs after removal of outliers in the cerebellum as shown in Fig. 6b. **d**) Comparing expression of microglial homeostatic genes across different brain regions and age groups published by Oliver *et al*., 2023 using interactive shiny app website (https://twc-stanford.shinyapps.io/spatiotemporal_brain_map/). Mot. Cortex: Motor cortex; Ent. Cortex: Entorhinal cortex; Ant. Hippocampus: Anterior hippocampus; Post Hippocampus: Posterior hippocampus.

**Fig. S9:**
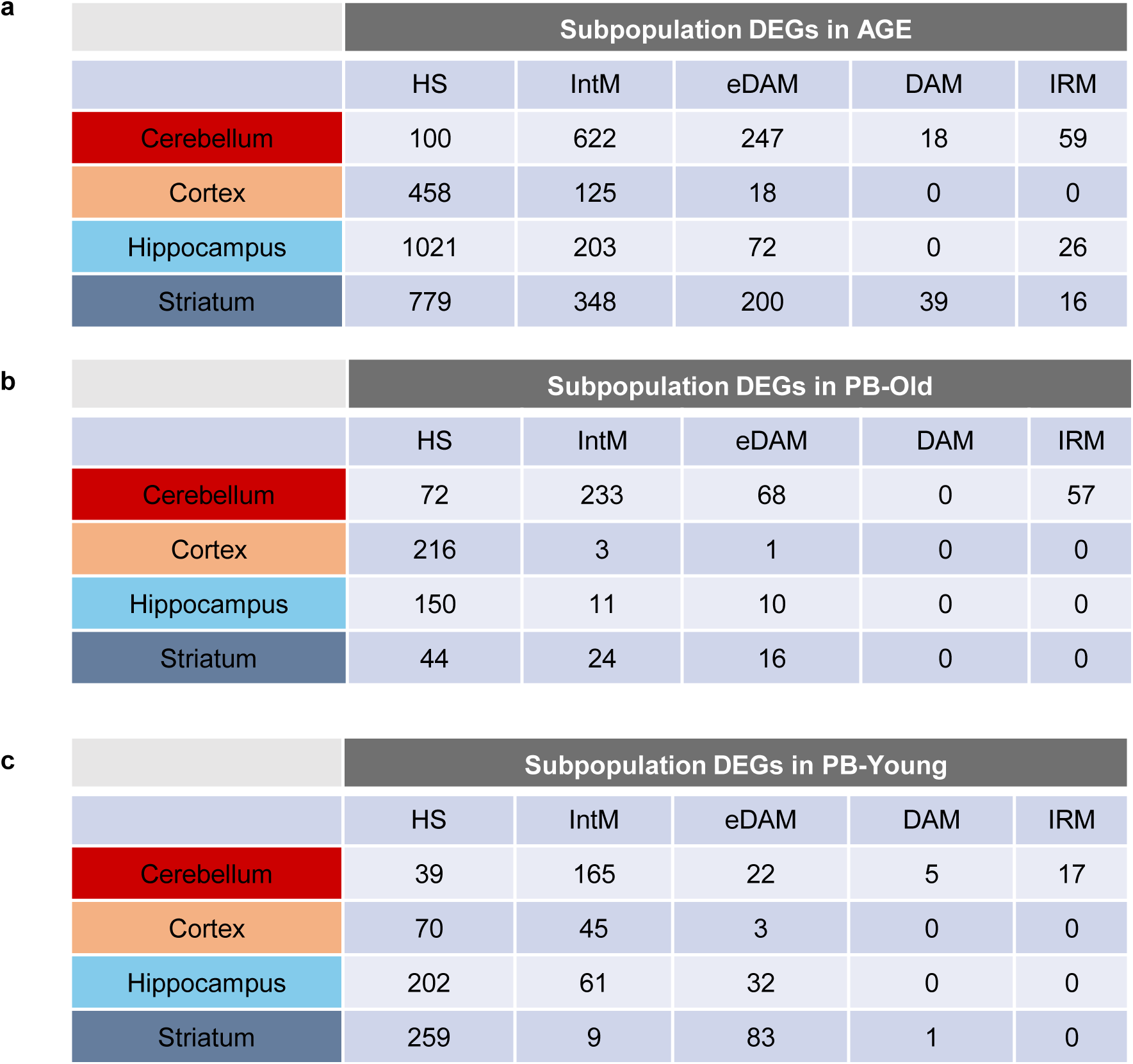
Effects of aging and parabiosis mediated transcriptomics changes at subpopulations level. **a**) Number of DEGs identifies in each brain region for HS, IntM, eDAM, DAM and IRM in AGE (A vs Y), **b**) in PB-Old (HY vs IY) and **c**) in PB-Young (HA vs IA).

