## Supplementary _Tables for "Core Activation Program and Selective Regional Responsiveness of Microglia in Aging and Parabiosis"

#### Supplementry Table 1. Marker genes for HS

| Genenames | p_val | avg_log2FC | pct.1 | pct.2 | p_val_adj |
|---|---|---|---|---|---|
| Fcrls | 0 | 1.164151 | 0.975 | 0.835 | 0 |
| P2ry12 | 0 | 0.683877 | 1 | 0.992 | 0 |
| Slc2a5 | 1.1E-265 | 0.805048 | 0.944 | 0.778 | 2.6E-261 |
| Cd164 | 4.2E-198 | 0.637272 | 0.946 | 0.832 | 9.9E-194 |
| P2ry13 | 1.6E-187 | 0.672599 | 0.977 | 0.912 | 3.8E-183 |
| Gpr34 | 4.8E-156 | 0.525926 | 0.992 | 0.971 | 1.1E-151 |
| Tmem119 | 3.3E-146 | 0.436684 | 1 | 0.989 | 7.8E-142 |
| Cx3cr1 | 3.2E-138 | 0.368682 | 1 | 0.999 | 7.5E-134 |
| Ptgs1 | 1.5E-132 | 0.617568 | 0.946 | 0.859 | 3.5E-128 |
| Upk1b | 2E-119 | 1.108185 | 0.473 | 0.285 | 4.7E-115 |
| Selplg | 2.2E-118 | 0.346517 | 1 | 0.997 | 5.3E-114 |
| Rhob | 1.1E-111 | 0.442272 | 0.928 | 0.837 | 2.5E-107 |
| Adgrg1 | 2.1E-104 | 0.452595 | 0.905 | 0.787 | 4.9E-100 |
| Stab1 | 4.4E-103 | 0.5169 | 0.822 | 0.651 | 1.04E-98 |
| Dusp6 | 4.2E-100 | 0.72768 | 0.731 | 0.592 | 9.89E-96 |
| Selenop | 7.63E-96 | 0.394549 | 0.991 | 0.977 | 1.8E-91 |
| Cmtm6 | 2.28E-93 | 0.504687 | 0.854 | 0.719 | 5.37E-89 |
| Ecscr | 6.61E-91 | 0.437621 | 0.706 | 0.521 | 1.56E-86 |
| Csf1r | 2.36E-86 | 0.319901 | 1 | 0.998 | 5.58E-82 |
| Lrba | 3.81E-83 | 0.792829 | 0.547 | 0.386 | 8.99E-79 |
| Olfml3 | 5.36E-81 | 0.354676 | 0.996 | 0.984 | 1.26E-76 |
| Fscn1 | 4.1E-76 | 0.386426 | 0.859 | 0.726 | 9.68E-72 |
| Slc40a1 | 1.78E-75 | 0.637155 | 0.522 | 0.363 | 4.2E-71 |
| Plxdc2 | 4.39E-75 | 0.409537 | 0.931 | 0.862 | 1.04E-70 |
| Txnip | 8.81E-75 | 0.632475 | 0.699 | 0.574 | 2.08E-70 |
| Scoc | 2.13E-74 | 0.445565 | 0.831 | 0.705 | 5.03E-70 |
| Slc7a8 | 4.93E-74 | 0.461301 | 0.723 | 0.573 | 1.16E-69 |
| Fam102b | 1.36E-73 | 0.667319 | 0.665 | 0.527 | 3.2E-69 |
| Fam110a | 7.62E-71 | 0.818828 | 0.364 | 0.221 | 1.8E-66 |
| Tmem204 | 1.7E-70 | 1.283354 | 0.225 | 0.108 | 4.01E-66 |
| Gpr165 | 2.39E-69 | 0.856521 | 0.267 | 0.139 | 5.64E-65 |
| Ubc | 1.69E-66 | 0.326818 | 0.974 | 0.941 | 3.99E-62 |
| Entpd1 | 2.05E-66 | 0.370484 | 0.94 | 0.873 | 4.83E-62 |
| Maf | 2.53E-66 | 0.51273 | 0.897 | 0.82 | 5.97E-62 |
| Crybb1 | 1.19E-65 | 0.276509 | 0.794 | 0.639 | 2.8E-61 |
| Vsir | 4.14E-65 | 0.308311 | 0.985 | 0.958 | 9.77E-61 |
| Golm1 | 6.34E-65 | 0.370533 | 0.877 | 0.769 | 1.49E-60 |
| Glul | 1.25E-61 | 0.26772 | 0.938 | 0.864 | 2.94E-57 |
| Gal3st4 | 1.11E-60 | 0.401926 | 0.58 | 0.432 | 2.62E-56 |
| Rtn4rl1 | 1.54E-60 | 0.468931 | 0.684 | 0.55 | 3.64E-56 |
| Qk | 1.87E-59 | 0.388145 | 0.936 | 0.875 | 4.4E-55 |
| Scamp5 | 6.34E-59 | 0.488286 | 0.578 | 0.435 | 1.5E-54 |
| Sall1 | 1.97E-58 | 0.363611 | 0.856 | 0.756 | 4.65E-54 |

| | | | | | |
|---|---|---|---|---|---|
| Sirpa | 9.66E-58 | 0.346027 | 0.958 | 0.912 | 2.28E-53 |
| Pla2g15 | 4.94E-57 | 0.451404 | 0.803 | 0.72 | 1.17E-52 |
| Rps6ka1 | 5.26E-57 | 0.403746 | 0.813 | 0.717 | 1.24E-52 |
| Il16 | 7.75E-57 | 0.631431 | 0.574 | 0.45 | 1.83E-52 |
| Zfhx3 | 3.3E-56 | 0.360951 | 0.948 | 0.889 | 7.78E-52 |
| Serinc3 | 3.98E-56 | 0.308423 | 0.996 | 0.973 | 9.39E-52 |
| Hpgd | 4.98E-56 | 0.478191 | 0.746 | 0.643 | 1.18E-51 |
| Ivns1abp | 7.53E-56 | 0.353558 | 0.852 | 0.764 | 1.78E-51 |
| Rnf13 | 4.5E-55 | 0.430901 | 0.776 | 0.691 | 1.06E-50 |
| Lpcat2 | 4.82E-55 | 0.262118 | 0.984 | 0.96 | 1.14E-50 |
| 1700017B0 | 5.61E-55 | 0.406011 | 0.694 | 0.566 | 1.32E-50 |
| Elmo1 | 1.18E-54 | 0.409739 | 0.724 | 0.604 | 2.79E-50 |
| Ifngr1 | 2.12E-54 | 0.267656 | 0.972 | 0.945 | 4.99E-50 |
| Bin2 | 7.37E-54 | 0.352016 | 0.909 | 0.839 | 1.74E-49 |
| Cyth4 | 8.19E-54 | 0.395225 | 0.894 | 0.839 | 1.93E-49 |
| Slco2b1 | 1.03E-52 | 0.329592 | 0.937 | 0.869 | 2.44E-48 |
| Bank1 | 2.55E-52 | 1.205105 | 0.2 | 0.103 | 6.03E-48 |
| Hfe | 4.49E-52 | 0.561348 | 0.452 | 0.322 | 1.06E-47 |
| Rab3il1 | 2.57E-51 | 0.398494 | 0.828 | 0.747 | 6.06E-47 |
| Pmepa1 | 2.82E-51 | 0.277433 | 0.845 | 0.735 | 6.64E-47 |
| Dusp7 | 4.88E-51 | 0.417557 | 0.588 | 0.458 | 1.15E-46 |
| Ddit4 | 1.07E-50 | 0.749708 | 0.25 | 0.143 | 2.53E-46 |
| Sema4b | 2.19E-50 | 0.785352 | 0.213 | 0.113 | 5.16E-46 |
| Ccr5 | 2.26E-50 | 0.419122 | 0.717 | 0.607 | 5.33E-46 |
| Hspa5 | 2.72E-50 | 0.298546 | 0.866 | 0.796 | 6.42E-46 |
| Sgk1 | 1.46E-49 | 0.608645 | 0.584 | 0.482 | 3.44E-45 |
| Sox4 | 2.47E-49 | 0.684201 | 0.422 | 0.301 | 5.84E-45 |
| Chsy1 | 1.46E-48 | 0.58925 | 0.496 | 0.37 | 3.45E-44 |
| Mertk | 1.73E-47 | 0.318743 | 0.872 | 0.795 | 4.09E-43 |
| Fermt3 | 1.29E-46 | 0.460391 | 0.773 | 0.687 | 3.03E-42 |
| Tm9sf2 | 1.14E-45 | 0.30651 | 0.769 | 0.672 | 2.69E-41 |
| Ubl3 | 1.31E-45 | 0.510589 | 0.691 | 0.601 | 3.1E-41 |
| Adrb2 | 1.71E-45 | 0.396873 | 0.652 | 0.541 | 4.02E-41 |
| Tsc22d4 | 1.92E-45 | 0.331944 | 0.667 | 0.553 | 4.54E-41 |
| Prpsap2 | 3.39E-45 | 0.652506 | 0.359 | 0.246 | 8E-41 |
| Pdia3 | 5.99E-45 | 0.268321 | 0.961 | 0.925 | 1.41E-40 |
| Eng | 1.33E-43 | 0.557093 | 0.519 | 0.409 | 3.14E-39 |
| Plod1 | 9.34E-43 | 0.444195 | 0.628 | 0.526 | 2.2E-38 |
| Eef2k | 1.3E-42 | 0.642957 | 0.448 | 0.336 | 3.07E-38 |
| St3gal5 | 1.37E-42 | 0.491178 | 0.611 | 0.508 | 3.24E-38 |
| Arhgap5 | 1.69E-42 | 0.323528 | 0.761 | 0.662 | 3.98E-38 |
| Zfp710 | 2.34E-42 | 0.435203 | 0.727 | 0.623 | 5.52E-38 |
| Itpripl1 | 4.09E-42 | 0.3881 | 0.729 | 0.611 | 9.66E-38 |
| Mylip | 2.48E-41 | 0.420518 | 0.699 | 0.601 | 5.85E-37 |
| Abcc3 | 7.11E-41 | 0.509275 | 0.427 | 0.311 | 1.68E-36 |
| Lyn | 2.76E-40 | 0.335306 | 0.871 | 0.811 | 6.51E-36 |
| Prkab1 | 1.06E-39 | 0.696638 | 0.381 | 0.277 | 2.5E-35 |

| | | | | | |
|---|---|---|---|---|---|
| Cttnbp2nl | 2.89E-39 | 0.309375 | 0.755 | 0.65 | 6.82E-35 |
| Rap1b | 3.81E-39 | 0.3998 | 0.773 | 0.706 | 8.98E-35 |
| Fcrl1 | 5.09E-39 | 0.721482 | 0.287 | 0.189 | 1.2E-34 |
| Bin1 | 5.4E-39 | 0.285104 | 0.913 | 0.862 | 1.27E-34 |
| St3gal6 | 1.73E-38 | 0.314481 | 0.804 | 0.72 | 4.08E-34 |
| Tuba1a | 2.38E-38 | 0.465076 | 0.61 | 0.509 | 5.62E-34 |
| Nrip1 | 2.8E-38 | 0.25787 | 0.654 | 0.539 | 6.61E-34 |
| Ccng2 | 4.58E-38 | 0.576445 | 0.365 | 0.261 | 1.08E-33 |
| 0610040J0 | 8.37E-38 | 0.462621 | 0.543 | 0.445 | 1.98E-33 |
| Sipa1 | 9.18E-38 | 0.34507 | 0.635 | 0.529 | 2.17E-33 |
| Mtss1 | 1.47E-37 | 0.45904 | 0.37 | 0.266 | 3.47E-33 |
| Gas6 | 3.47E-37 | 0.579763 | 0.482 | 0.387 | 8.18E-33 |
| Sh2b3 | 7.17E-37 | 0.495781 | 0.449 | 0.341 | 1.69E-32 |
| Vav1 | 1.91E-36 | 0.459322 | 0.661 | 0.578 | 4.51E-32 |
| Csnk1e | 2.41E-36 | 0.348621 | 0.622 | 0.517 | 5.68E-32 |
| Numb | 3.9E-36 | 0.275439 | 0.592 | 0.479 | 9.2E-32 |
| Slc35c2 | 4.17E-36 | 0.582575 | 0.464 | 0.367 | 9.84E-32 |
| Tanc2 | 1.12E-35 | 0.280638 | 0.831 | 0.751 | 2.63E-31 |
| Zfp706 | 2.86E-35 | 0.310843 | 0.661 | 0.56 | 6.75E-31 |
| Cdk5r1 | 3.12E-35 | 0.320043 | 0.283 | 0.187 | 7.37E-31 |
| Idh2 | 3.15E-35 | 0.523171 | 0.444 | 0.342 | 7.44E-31 |
| Hnrnpk | 4.1E-35 | 0.316522 | 0.829 | 0.761 | 9.66E-31 |
| Gna15 | 5.09E-35 | 0.344084 | 0.689 | 0.593 | 1.2E-30 |
| Ncf1 | 9.82E-35 | 0.348283 | 0.735 | 0.658 | 2.32E-30 |
| Sesn1 | 6.2E-34 | 0.632588 | 0.439 | 0.342 | 1.46E-29 |
| Capza2 | 8.29E-34 | 0.283427 | 0.89 | 0.853 | 1.95E-29 |
| Gm7964 | 1.03E-33 | 0.324737 | 0.644 | 0.538 | 2.43E-29 |
| Btg2 | 1.43E-33 | 0.432699 | 0.584 | 0.497 | 3.38E-29 |
| Siglece | 1.97E-33 | 0.334585 | 0.372 | 0.271 | 4.65E-29 |
| Tram1 | 2.44E-33 | 0.318631 | 0.673 | 0.575 | 5.76E-29 |
| Snx18 | 4.53E-33 | 0.390938 | 0.648 | 0.554 | 1.07E-28 |
| Fli1 | 4.64E-33 | 0.413362 | 0.645 | 0.552 | 1.09E-28 |
| Tnfrsf21 | 7.44E-33 | 0.432048 | 0.513 | 0.416 | 1.76E-28 |
| C5ar2 | 7.47E-33 | 0.353663 | 0.658 | 0.553 | 1.76E-28 |
| Rnf167 | 9.29E-33 | 0.443715 | 0.524 | 0.433 | 2.19E-28 |
| Snn | 1.03E-32 | 0.267437 | 0.382 | 0.279 | 2.43E-28 |
| Nrros | 1.24E-32 | 0.399065 | 0.64 | 0.557 | 2.93E-28 |
| Daglb | 1.87E-32 | 0.325049 | 0.753 | 0.688 | 4.42E-28 |
| Sla | 1.93E-32 | 0.390499 | 0.545 | 0.45 | 4.54E-28 |
| Tmem63a | 2.32E-32 | 0.3987 | 0.469 | 0.369 | 5.48E-28 |
| Slc44a2 | 3.7E-32 | 0.516288 | 0.443 | 0.344 | 8.72E-28 |
| Nav3 | 4.19E-32 | 0.275558 | 0.656 | 0.558 | 9.88E-28 |
| Gp9 | 5.38E-32 | 0.48953 | 0.228 | 0.144 | 1.27E-27 |
| Epb41l2 | 9.38E-32 | 0.255783 | 0.89 | 0.835 | 2.21E-27 |
| Man1a | 2.27E-31 | 0.375456 | 0.401 | 0.302 | 5.35E-27 |
| Ggta1 | 3.61E-31 | 0.848348 | 0.206 | 0.129 | 8.52E-27 |
| Mef2a | 3.74E-31 | 0.300211 | 0.81 | 0.744 | 8.82E-27 |

| | | | | | |
|---|---|---|---|---|---|
| Abl1 | 9.05E-31 | 0.53173 | 0.327 | 0.235 | 2.13E-26 |
| Ddx5 | 3.5E-30 | 0.323182 | 0.789 | 0.723 | 8.25E-26 |
| Serinc1 | 4.1E-30 | 0.346479 | 0.657 | 0.576 | 9.67E-26 |
| Tspan14 | 4.97E-30 | 0.405659 | 0.571 | 0.482 | 1.17E-25 |
| Gcn1l1 | 5.36E-30 | 0.485596 | 0.464 | 0.368 | 1.26E-25 |
| Chst7 | 5.59E-30 | 0.636523 | 0.286 | 0.201 | 1.32E-25 |
| Gcnt1 | 7.36E-30 | 0.252243 | 0.524 | 0.422 | 1.74E-25 |
| Rtn1 | 8.25E-30 | 0.5493 | 0.391 | 0.301 | 1.95E-25 |
| Cables1 | 8.97E-30 | 0.596228 | 0.29 | 0.206 | 2.12E-25 |
| Tnfrsf1b | 1.11E-29 | 0.347102 | 0.559 | 0.461 | 2.62E-25 |
| Sqor | 1.37E-29 | 0.475578 | 0.319 | 0.231 | 3.22E-25 |
| Arhgef40 | 1.51E-29 | 0.312972 | 0.469 | 0.368 | 3.57E-25 |
| Rilpl1 | 2.65E-29 | 0.305933 | 0.328 | 0.236 | 6.25E-25 |
| Snx5 | 3.87E-29 | 0.296503 | 0.81 | 0.758 | 9.13E-25 |
| Tifab | 4.82E-29 | 0.336659 | 0.577 | 0.485 | 1.14E-24 |
| Tpst2 | 4.94E-29 | 0.279891 | 0.701 | 0.619 | 1.16E-24 |
| Ms4a6b | 8.15E-29 | 0.326202 | 0.372 | 0.276 | 1.92E-24 |
| Cdh23 | 8.28E-29 | 0.633794 | 0.26 | 0.178 | 1.95E-24 |
| Snx17 | 8.53E-29 | 0.426827 | 0.501 | 0.414 | 2.01E-24 |
| Twf2 | 1.1E-28 | 0.277916 | 0.619 | 0.521 | 2.6E-24 |
| Etv5 | 1.1E-28 | 0.48683 | 0.418 | 0.324 | 2.6E-24 |
| Pald1 | 1.17E-28 | 0.407546 | 0.473 | 0.377 | 2.76E-24 |
| Zfp652 | 1.19E-28 | 0.456481 | 0.694 | 0.618 | 2.82E-24 |
| Lpcat3 | 1.36E-28 | 0.31286 | 0.547 | 0.452 | 3.2E-24 |
| Lpar6 | 1.9E-28 | 0.27646 | 0.632 | 0.543 | 4.49E-24 |
| Fchsd2 | 3.17E-28 | 0.491544 | 0.429 | 0.335 | 7.47E-24 |
| Foxn3 | 3.87E-28 | 0.263809 | 0.872 | 0.813 | 9.14E-24 |
| Dnm2 | 3.94E-28 | 0.339471 | 0.687 | 0.608 | 9.29E-24 |
| Gm6277 | 4.04E-28 | 0.606108 | 0.219 | 0.143 | 9.52E-24 |
| Edem2 | 5.92E-28 | 0.333807 | 0.586 | 0.494 | 1.4E-23 |
| Sash3 | 6.52E-28 | 0.424992 | 0.412 | 0.321 | 1.54E-23 |
| St6galnac4 | 7.45E-28 | 0.310921 | 0.467 | 0.371 | 1.76E-23 |
| Parvg | 1.17E-27 | 0.477001 | 0.457 | 0.375 | 2.77E-23 |
| Trim8 | 1.43E-27 | 0.420743 | 0.493 | 0.406 | 3.37E-23 |
| Irf2bpl | 1.6E-27 | 0.271666 | 0.378 | 0.288 | 3.78E-23 |
| Cask | 2.07E-27 | 0.342939 | 0.405 | 0.315 | 4.88E-23 |
| Sort1 | 4.66E-27 | 0.34995 | 0.52 | 0.424 | 1.1E-22 |
| Mlph | 5.17E-27 | 0.593429 | 0.411 | 0.326 | 1.22E-22 |
| Arhgap27 | 6.11E-27 | 0.510043 | 0.342 | 0.256 | 1.44E-22 |
| Sft2d2 | 6.67E-27 | 0.328738 | 0.427 | 0.332 | 1.57E-22 |
| Cd33 | 6.85E-27 | 0.261227 | 0.775 | 0.707 | 1.62E-22 |
| Ggt5 | 8.2E-27 | 0.630771 | 0.263 | 0.185 | 1.93E-22 |
| Mafg | 8.76E-27 | 0.491158 | 0.525 | 0.444 | 2.07E-22 |
| Man1c1 | 9.27E-27 | 0.347955 | 0.521 | 0.43 | 2.19E-22 |
| Tnfrsf13b | 1.84E-26 | 0.615662 | 0.331 | 0.25 | 4.33E-22 |
| Ykt6 | 2.09E-26 | 0.250243 | 0.405 | 0.314 | 4.94E-22 |
| Surf4 | 2.36E-26 | 0.286592 | 0.618 | 0.531 | 5.56E-22 |

| | | | | | |
|---|---|---|---|---|---|
| Fgd2 | 2.39E-26 | 0.323803 | 0.747 | 0.681 | 5.65E-22 |
| Rcsd1 | 2.98E-26 | 0.414197 | 0.46 | 0.372 | 7.03E-22 |
| Abcg1 | 3.64E-26 | 0.286793 | 0.657 | 0.571 | 8.6E-22 |
| Ctsf | 4.88E-26 | 0.311546 | 0.666 | 0.592 | 1.15E-21 |
| Inpp4b | 5.15E-26 | 0.712379 | 0.252 | 0.176 | 1.22E-21 |
| Tcf7l2 | 1.31E-25 | 0.323139 | 0.452 | 0.362 | 3.1E-21 |
| Ctc1 | 1.47E-25 | 0.263921 | 0.63 | 0.545 | 3.48E-21 |
| Ubash3b | 1.75E-25 | 0.434285 | 0.608 | 0.535 | 4.13E-21 |
| Nptn | 1.92E-25 | 0.290143 | 0.619 | 0.545 | 4.53E-21 |
| Clstn1 | 2.24E-25 | 0.429958 | 0.432 | 0.348 | 5.28E-21 |
| Gm10790 | 3.01E-25 | 0.318034 | 0.605 | 0.518 | 7.11E-21 |
| Klf3 | 3.08E-25 | 0.318585 | 0.588 | 0.506 | 7.27E-21 |
| Tgm2 | 3.22E-25 | 0.652485 | 0.27 | 0.196 | 7.59E-21 |
| Gm10134 | 5.6E-25 | 0.555531 | 0.388 | 0.305 | 1.32E-20 |
| Ncoa4 | 6.31E-25 | 0.398116 | 0.254 | 0.179 | 1.49E-20 |
| Cib1 | 7.44E-25 | 0.320602 | 0.292 | 0.211 | 1.75E-20 |
| Zfp36l2 | 8.7E-25 | 0.327942 | 0.504 | 0.413 | 2.05E-20 |
| Tm6sf1 | 9.35E-25 | 0.429437 | 0.459 | 0.377 | 2.21E-20 |
| Vapa | 1.02E-24 | 0.278431 | 0.648 | 0.568 | 2.41E-20 |
| Casp8 | 1.03E-24 | 0.349359 | 0.474 | 0.391 | 2.44E-20 |
| Soga1 | 1.19E-24 | 0.264686 | 0.586 | 0.495 | 2.8E-20 |
| Tle3 | 1.23E-24 | 0.383028 | 0.317 | 0.234 | 2.9E-20 |
| Itga9 | 1.3E-24 | 0.519464 | 0.291 | 0.21 | 3.08E-20 |
| Mgat1 | 1.61E-24 | 0.334236 | 0.574 | 0.497 | 3.8E-20 |
| Slc16a6 | 3.02E-24 | 0.35322 | 0.548 | 0.464 | 7.12E-20 |
| Dtx4 | 3.28E-24 | 0.418167 | 0.458 | 0.371 | 7.74E-20 |
| Asb2 | 4.45E-24 | 0.421393 | 0.375 | 0.289 | 1.05E-19 |
| G530011O( | 4.77E-24 | 0.26517 | 0.385 | 0.298 | 1.13E-19 |
| Rgmb | 5.19E-24 | 0.32951 | 0.461 | 0.372 | 1.22E-19 |
| Mgll | 5.42E-24 | 0.311592 | 0.361 | 0.278 | 1.28E-19 |
| Nckap5l | 7.15E-24 | 0.561017 | 0.24 | 0.168 | 1.69E-19 |
| Gsk3b | 9.14E-24 | 0.355106 | 0.606 | 0.527 | 2.16E-19 |
| Fez2 | 1.17E-23 | 0.330629 | 0.501 | 0.417 | 2.77E-19 |
| Rin2 | 1.37E-23 | 0.435989 | 0.55 | 0.477 | 3.23E-19 |
| Irf5 | 1.62E-23 | 0.275102 | 0.686 | 0.617 | 3.83E-19 |
| Ets1 | 1.67E-23 | 0.322281 | 0.351 | 0.269 | 3.95E-19 |
| Mtus1 | 1.95E-23 | 0.278985 | 0.592 | 0.503 | 4.59E-19 |
| Dpysl2 | 2.15E-23 | 0.302657 | 0.878 | 0.832 | 5.08E-19 |
| Rapgef5 | 2.19E-23 | 0.353141 | 0.374 | 0.289 | 5.17E-19 |
| Akirin1 | 3.77E-23 | 0.416068 | 0.375 | 0.296 | 8.88E-19 |
| Irf2bp2 | 5.13E-23 | 0.38806 | 0.381 | 0.299 | 1.21E-18 |
| Rhoh | 7.66E-23 | 0.288771 | 0.48 | 0.391 | 1.81E-18 |
| Atmin | 7.95E-23 | 0.317217 | 0.206 | 0.138 | 1.87E-18 |
| Mef2c | 1.27E-22 | 0.260219 | 0.846 | 0.802 | 2.99E-18 |
| Rpn2 | 1.68E-22 | 0.252745 | 0.735 | 0.679 | 3.97E-18 |
| Zmiz1 | 2.56E-22 | 0.360651 | 0.642 | 0.568 | 6.03E-18 |
| Lcp2 | 3.79E-22 | 0.270361 | 0.595 | 0.515 | 8.94E-18 |

| | | | | | |
|---|---|---|---|---|---|
| Mfap3 | 4.01E-22 | 0.256484 | 0.615 | 0.537 | 9.46E-18 |
| Tjp1 | 4.59E-22 | 0.305824 | 0.342 | 0.262 | 1.08E-17 |
| Hs6st1 | 5.73E-22 | 0.302239 | 0.317 | 0.239 | 1.35E-17 |
| Adgre1 | 8.97E-22 | 0.253242 | 0.629 | 0.552 | 2.12E-17 |
| Cnot8 | 1.15E-21 | 0.401594 | 0.496 | 0.421 | 2.7E-17 |
| Vcp | 1.2E-21 | 0.299582 | 0.576 | 0.503 | 2.83E-17 |
| Plod3 | 1.26E-21 | 0.313732 | 0.575 | 0.499 | 2.98E-17 |
| Rab1a | 1.36E-21 | 0.259081 | 0.659 | 0.586 | 3.21E-17 |
| Tagap | 1.38E-21 | 0.373942 | 0.376 | 0.299 | 3.25E-17 |
| Slc25a45 | 1.57E-21 | 0.491042 | 0.338 | 0.265 | 3.71E-17 |
| Tmem206 | 1.69E-21 | 0.44843 | 0.353 | 0.28 | 3.99E-17 |
| Snx6 | 2.37E-21 | 0.30121 | 0.478 | 0.4 | 5.6E-17 |
| Ak1 | 2.39E-21 | 0.628379 | 0.253 | 0.185 | 5.64E-17 |
| Cep68 | 3.26E-21 | 0.432365 | 0.32 | 0.242 | 7.69E-17 |
| Map2k1 | 3.38E-21 | 0.497584 | 0.459 | 0.393 | 7.97E-17 |
| Cd300a | 3.52E-21 | 0.296575 | 0.579 | 0.505 | 8.3E-17 |
| Nr3c1 | 4.15E-21 | 0.268107 | 0.528 | 0.447 | 9.8E-17 |
| Ddx6 | 4.16E-21 | 0.433351 | 0.381 | 0.309 | 9.82E-17 |
| Ap1b1 | 4.43E-21 | 0.308731 | 0.459 | 0.378 | 1.04E-16 |
| Mfng | 5.32E-21 | 0.368409 | 0.422 | 0.346 | 1.25E-16 |
| Scn1b | 6.64E-21 | 0.440344 | 0.316 | 0.244 | 1.57E-16 |
| Snta1 | 7.4E-21 | 0.446682 | 0.319 | 0.244 | 1.75E-16 |
| Amfr | 7.6E-21 | 0.368858 | 0.444 | 0.371 | 1.79E-16 |
| Pacsin2 | 9.04E-21 | 0.417652 | 0.456 | 0.386 | 2.13E-16 |
| Gpr155 | 1.03E-20 | 0.505124 | 0.295 | 0.224 | 2.43E-16 |
| Dip2b | 1.37E-20 | 0.341166 | 0.603 | 0.525 | 3.23E-16 |
| Tlr12 | 1.48E-20 | 0.516028 | 0.279 | 0.211 | 3.49E-16 |
| Nfam1 | 1.88E-20 | 0.255608 | 0.632 | 0.566 | 4.43E-16 |
| Xbp1 | 3.51E-20 | 0.262194 | 0.63 | 0.557 | 8.28E-16 |
| Ptpn1 | 3.68E-20 | 0.267307 | 0.63 | 0.563 | 8.69E-16 |
| Itpr2 | 3.82E-20 | 0.420454 | 0.517 | 0.44 | 9.02E-16 |
| Ptp4a3 | 4.21E-20 | 0.323268 | 0.531 | 0.459 | 9.93E-16 |
| Abcd3 | 6.21E-20 | 0.587483 | 0.225 | 0.16 | 1.47E-15 |
| Stard3 | 1.08E-19 | 0.297485 | 0.472 | 0.397 | 2.54E-15 |
| Csnk1a1 | 1.08E-19 | 0.25938 | 0.519 | 0.443 | 2.55E-15 |
| I830077J02 | 1.21E-19 | 0.366956 | 0.41 | 0.335 | 2.85E-15 |
| Cep152 | 1.47E-19 | 0.667505 | 0.221 | 0.161 | 3.48E-15 |
| Bcl9l | 2.42E-19 | 0.316169 | 0.243 | 0.177 | 5.71E-15 |
| Ppp3ca | 2.96E-19 | 0.274941 | 0.533 | 0.463 | 6.98E-15 |
| Gm33699 | 3.22E-19 | 0.265437 | 0.258 | 0.19 | 7.6E-15 |
| Gtf2a1 | 3.45E-19 | 0.478216 | 0.389 | 0.32 | 8.13E-15 |
| Sppl3 | 5.07E-19 | 0.402464 | 0.469 | 0.397 | 1.2E-14 |
| Pdlim4 | 5.42E-19 | 0.250514 | 0.356 | 0.281 | 1.28E-14 |
| Atp6ap2 | 6.9E-19 | 0.252493 | 0.639 | 0.579 | 1.63E-14 |
| Kdm6b | 7.62E-19 | 0.38724 | 0.292 | 0.223 | 1.8E-14 |
| Klhl9 | 8.5E-19 | 0.330661 | 0.488 | 0.413 | 2.01E-14 |
| Bid | 9.48E-19 | 0.463883 | 0.259 | 0.194 | 2.24E-14 |

| | | | | | |
|---|---|---|---|---|---|
| Ralb | 1.6E-18 | 0.430634 | 0.445 | 0.377 | 3.78E-14 |
| Rin3 | 2.5E-18 | 0.257827 | 0.34 | 0.267 | 5.89E-14 |
| Map7d1 | 3.06E-18 | 0.489586 | 0.34 | 0.275 | 7.22E-14 |
| Plcg1 | 3.49E-18 | 0.600873 | 0.204 | 0.146 | 8.24E-14 |
| Sypl | 3.89E-18 | 0.355945 | 0.463 | 0.397 | 9.18E-14 |
| Btg1 | 3.93E-18 | 0.28589 | 0.689 | 0.632 | 9.27E-14 |
| Aftph | 4.13E-18 | 0.345723 | 0.37 | 0.299 | 9.73E-14 |
| Arl8b | 4.46E-18 | 0.32226 | 0.494 | 0.426 | 1.05E-13 |
| Nars | 5.23E-18 | 0.34236 | 0.507 | 0.437 | 1.23E-13 |
| Tor4a | 5.6E-18 | 0.515538 | 0.262 | 0.199 | 1.32E-13 |
| Pld1 | 5.64E-18 | 0.259461 | 0.461 | 0.38 | 1.33E-13 |
| Slc46a1 | 1.12E-17 | 0.345195 | 0.455 | 0.389 | 2.65E-13 |
| Numa1 | 1.13E-17 | 0.364841 | 0.401 | 0.33 | 2.67E-13 |
| Ski | 1.25E-17 | 0.267996 | 0.53 | 0.462 | 2.95E-13 |
| Tmem100 | 1.65E-17 | 0.348726 | 0.362 | 0.294 | 3.89E-13 |
| Tmem64 | 1.78E-17 | 0.354955 | 0.218 | 0.158 | 4.2E-13 |
| Stt3a | 1.87E-17 | 0.270135 | 0.576 | 0.517 | 4.41E-13 |
| Ssr1 | 1.88E-17 | 0.364509 | 0.652 | 0.594 | 4.43E-13 |
| Hhex | 2.86E-17 | 0.252461 | 0.394 | 0.318 | 6.74E-13 |
| Csnk2a1 | 3.8E-17 | 0.295982 | 0.494 | 0.429 | 8.97E-13 |
| Zdhhc14 | 3.98E-17 | 0.511763 | 0.201 | 0.145 | 9.38E-13 |
| Glg1 | 5.66E-17 | 0.288853 | 0.393 | 0.323 | 1.34E-12 |
| Mxi1 | 5.67E-17 | 0.536824 | 0.203 | 0.148 | 1.34E-12 |
| Abhd17c | 8.8E-17 | 0.280048 | 0.474 | 0.406 | 2.08E-12 |
| Acss1 | 9E-17 | 0.387022 | 0.247 | 0.186 | 2.12E-12 |
| Orai2 | 1E-16 | 0.466496 | 0.233 | 0.174 | 2.37E-12 |
| Usp22 | 1.02E-16 | 0.251189 | 0.364 | 0.295 | 2.4E-12 |
| Wdr1 | 1.02E-16 | 0.321044 | 0.529 | 0.474 | 2.41E-12 |
| Sh3bp1 | 1.23E-16 | 0.358366 | 0.318 | 0.253 | 2.89E-12 |
| Atp6v0a2 | 1.36E-16 | 0.319246 | 0.432 | 0.363 | 3.21E-12 |
| Poldip3 | 1.37E-16 | 0.293652 | 0.389 | 0.322 | 3.24E-12 |
| Slc25a37 | 1.4E-16 | 0.352009 | 0.236 | 0.175 | 3.31E-12 |
| Spsb1 | 2.07E-16 | 0.459728 | 0.325 | 0.262 | 4.89E-12 |
| Pxk | 2.12E-16 | 0.574728 | 0.234 | 0.179 | 5E-12 |
| Sf1 | 2.31E-16 | 0.354377 | 0.337 | 0.271 | 5.45E-12 |
| Prrc2a | 2.55E-16 | 0.253836 | 0.371 | 0.306 | 6.01E-12 |
| Golt1b | 2.64E-16 | 0.480972 | 0.323 | 0.262 | 6.24E-12 |
| Slc9a9 | 2.68E-16 | 0.298431 | 0.373 | 0.307 | 6.33E-12 |
| Yipf4 | 2.81E-16 | 0.296661 | 0.348 | 0.284 | 6.62E-12 |
| Dnajb9 | 2.89E-16 | 0.349183 | 0.333 | 0.27 | 6.81E-12 |
| Fry | 2.99E-16 | 0.326982 | 0.269 | 0.207 | 7.04E-12 |
| Samsn1 | 3.78E-16 | 0.348095 | 0.464 | 0.391 | 8.91E-12 |
| Mpi | 4.04E-16 | 0.346188 | 0.298 | 0.233 | 9.53E-12 |
| Sbf2 | 4.19E-16 | 0.296645 | 0.357 | 0.29 | 9.88E-12 |
| Cryl1 | 5E-16 | 0.298114 | 0.333 | 0.269 | 1.18E-11 |
| Nek6 | 5.36E-16 | 0.259918 | 0.432 | 0.361 | 1.26E-11 |
| Rab35 | 6.41E-16 | 0.315617 | 0.278 | 0.213 | 1.51E-11 |

| | | | | | |
|---|---|---|---|---|---|
| Tmem144 | 6.7E-16 | 0.642079 | 0.203 | 0.149 | 1.58E-11 |
| Tfe3 | 6.89E-16 | 0.345448 | 0.477 | 0.414 | 1.63E-11 |
| Retreg1 | 7.79E-16 | 0.268946 | 0.501 | 0.433 | 1.84E-11 |
| Cul1 | 8.17E-16 | 0.263887 | 0.412 | 0.347 | 1.93E-11 |
| Cln8 | 9.97E-16 | 0.418211 | 0.337 | 0.275 | 2.35E-11 |
| P3h2 | 1.04E-15 | 0.430873 | 0.239 | 0.183 | 2.45E-11 |
| Rassf5 | 1.12E-15 | 0.372324 | 0.262 | 0.205 | 2.65E-11 |
| Pdia4 | 1.39E-15 | 0.377217 | 0.392 | 0.329 | 3.27E-11 |
| Cerk | 1.51E-15 | 0.35101 | 0.417 | 0.352 | 3.57E-11 |
| Lpxn | 1.7E-15 | 0.280256 | 0.39 | 0.325 | 4.01E-11 |
| Mapre1 | 1.89E-15 | 0.308514 | 0.467 | 0.405 | 4.47E-11 |
| Il21r | 2.37E-15 | 0.345305 | 0.473 | 0.41 | 5.59E-11 |
| Mapk14 | 3.05E-15 | 0.291203 | 0.382 | 0.319 | 7.18E-11 |
| Hjurp | 3.38E-15 | 0.392627 | 0.222 | 0.167 | 7.97E-11 |
| Filip1l | 4E-15 | 0.386947 | 0.295 | 0.235 | 9.45E-11 |
| Gbp7 | 4.47E-15 | 0.313331 | 0.275 | 0.218 | 1.05E-10 |
| March1 | 4.71E-15 | 0.262974 | 0.334 | 0.27 | 1.11E-10 |
| Arl1 | 5E-15 | 0.41414 | 0.347 | 0.286 | 1.18E-10 |
| Crebbp | 1.04E-14 | 0.315936 | 0.569 | 0.511 | 2.45E-10 |
| Tmem135 | 1.21E-14 | 0.274004 | 0.371 | 0.31 | 2.86E-10 |
| Gmpr | 1.28E-14 | 0.328439 | 0.216 | 0.161 | 3.01E-10 |
| Msrb2 | 1.28E-14 | 0.309811 | 0.255 | 0.197 | 3.02E-10 |
| Acads | 1.46E-14 | 0.334394 | 0.283 | 0.223 | 3.44E-10 |
| Plxna4 | 1.51E-14 | 0.339624 | 0.227 | 0.173 | 3.56E-10 |
| Gosr2 | 1.62E-14 | 0.384987 | 0.206 | 0.155 | 3.83E-10 |
| Mta2 | 1.8E-14 | 0.3677 | 0.287 | 0.231 | 4.24E-10 |
| Extl3 | 1.93E-14 | 0.313796 | 0.437 | 0.371 | 4.54E-10 |
| Npl | 2.51E-14 | 0.332198 | 0.224 | 0.17 | 5.91E-10 |
| Ncstn | 2.79E-14 | 0.263849 | 0.521 | 0.464 | 6.57E-10 |
| Capn3 | 3.03E-14 | 0.308686 | 0.422 | 0.36 | 7.15E-10 |
| Ccdc93 | 3.88E-14 | 0.418962 | 0.284 | 0.226 | 9.15E-10 |
| Arhgap12 | 4.45E-14 | 0.439636 | 0.265 | 0.207 | 1.05E-09 |
| Ppp1r9a | 5.03E-14 | 0.274698 | 0.382 | 0.319 | 1.19E-09 |
| Mbnl2 | 5.53E-14 | 0.383817 | 0.44 | 0.381 | 1.3E-09 |
| Wac | 5.68E-14 | 0.294202 | 0.36 | 0.302 | 1.34E-09 |
| Ddah2 | 6.21E-14 | 0.356675 | 0.209 | 0.157 | 1.46E-09 |
| Lrp10 | 6.59E-14 | 0.281908 | 0.472 | 0.415 | 1.55E-09 |
| Zfp703 | 6.73E-14 | 0.416614 | 0.233 | 0.181 | 1.59E-09 |
| Lpcat1 | 6.95E-14 | 0.260019 | 0.519 | 0.458 | 1.64E-09 |
| Wdr45b | 8.32E-14 | 0.759462 | 0.246 | 0.196 | 1.96E-09 |
| Bicd2 | 9.46E-14 | 0.375389 | 0.268 | 0.212 | 2.23E-09 |
| Ube2h | 9.52E-14 | 0.301208 | 0.353 | 0.293 | 2.25E-09 |
| Cltc | 9.62E-14 | 0.297066 | 0.644 | 0.594 | 2.27E-09 |
| Zbtb38 | 1.36E-13 | 0.387585 | 0.265 | 0.21 | 3.21E-09 |
| Rnf145 | 1.63E-13 | 0.506099 | 0.218 | 0.168 | 3.86E-09 |
| Atxn1 | 1.76E-13 | 0.492303 | 0.222 | 0.171 | 4.15E-09 |
| Dbnl | 2.07E-13 | 0.277079 | 0.434 | 0.375 | 4.88E-09 |

| | | | | | |
|---|---|---|---|---|---|
| Tgfa | 2.12E-13 | 0.259049 | 0.28 | 0.222 | 5E-09 |
| Myd88 | 2.65E-13 | 0.34324 | 0.265 | 0.211 | 6.24E-09 |
| Elk3 | 2.66E-13 | 0.27597 | 0.401 | 0.342 | 6.27E-09 |
| Prag1 | 2.8E-13 | 0.459376 | 0.25 | 0.197 | 6.6E-09 |
| Ppp3r1 | 2.88E-13 | 0.475695 | 0.238 | 0.188 | 6.8E-09 |
| Trim28 | 3.57E-13 | 0.27212 | 0.321 | 0.264 | 8.43E-09 |
| Dok3 | 4.04E-13 | 0.287165 | 0.286 | 0.231 | 9.52E-09 |
| Crot | 4.76E-13 | 0.600525 | 0.217 | 0.167 | 1.12E-08 |
| Irgm2 | 5.14E-13 | 0.329606 | 0.383 | 0.322 | 1.21E-08 |
| Bcl2l1 | 5.16E-13 | 0.336655 | 0.415 | 0.362 | 1.22E-08 |
| Il1a | 5.59E-13 | 0.544656 | 0.21 | 0.162 | 1.32E-08 |
| Ptbp1 | 6.33E-13 | 0.539486 | 0.303 | 0.254 | 1.49E-08 |
| Lifr | 8.35E-13 | 0.355694 | 0.326 | 0.268 | 1.97E-08 |
| Rab18 | 9.01E-13 | 0.413222 | 0.271 | 0.219 | 2.12E-08 |
| Rela | 9.65E-13 | 0.25914 | 0.397 | 0.339 | 2.28E-08 |
| Wdfy2 | 9.7E-13 | 0.289748 | 0.288 | 0.235 | 2.29E-08 |
| Maea | 1.01E-12 | 0.306779 | 0.286 | 0.232 | 2.39E-08 |
| Pnrc2 | 1.18E-12 | 0.509329 | 0.323 | 0.273 | 2.78E-08 |
| Zfp90 | 1.23E-12 | 0.36091 | 0.239 | 0.188 | 2.89E-08 |
| Tsg101 | 1.31E-12 | 0.271721 | 0.333 | 0.277 | 3.09E-08 |
| Kat6a | 1.34E-12 | 0.484375 | 0.334 | 0.278 | 3.15E-08 |
| Frmd8 | 1.37E-12 | 0.586308 | 0.227 | 0.18 | 3.23E-08 |
| Kcnk13 | 1.38E-12 | 0.404814 | 0.332 | 0.279 | 3.26E-08 |
| Tspyl1 | 1.74E-12 | 0.390056 | 0.284 | 0.232 | 4.11E-08 |
| Spred1 | 1.77E-12 | 0.566224 | 0.23 | 0.182 | 4.17E-08 |
| 4833439L1 | 1.84E-12 | 0.475735 | 0.336 | 0.285 | 4.35E-08 |
| Ncbp3 | 1.95E-12 | 0.402315 | 0.239 | 0.189 | 4.6E-08 |
| Agmo | 2.16E-12 | 0.455362 | 0.245 | 0.195 | 5.1E-08 |
| Ctso | 2.37E-12 | 0.287367 | 0.393 | 0.339 | 5.6E-08 |
| Gm38158 | 2.38E-12 | 0.266099 | 0.315 | 0.259 | 5.62E-08 |
| Tspan18 | 2.67E-12 | 0.415367 | 0.207 | 0.161 | 6.31E-08 |
| Zbtb18 | 2.83E-12 | 0.364318 | 0.231 | 0.181 | 6.68E-08 |
| Crat | 3.02E-12 | 0.442362 | 0.262 | 0.211 | 7.11E-08 |
| Cnot2 | 3.27E-12 | 0.371064 | 0.326 | 0.276 | 7.71E-08 |
| Pdcd4 | 3.82E-12 | 0.37016 | 0.424 | 0.37 | 9.01E-08 |
| Slc35a5 | 4.19E-12 | 0.389303 | 0.219 | 0.171 | 9.88E-08 |
| Fbrsl1 | 4.55E-12 | 0.355161 | 0.414 | 0.361 | 1.07E-07 |
| Sidt2 | 4.68E-12 | 0.270502 | 0.308 | 0.255 | 1.1E-07 |
| Mfsd14a | 5.07E-12 | 0.406778 | 0.235 | 0.187 | 1.2E-07 |
| Maml3 | 5.13E-12 | 0.275495 | 0.277 | 0.223 | 1.21E-07 |
| Bckdha | 6.12E-12 | 0.357234 | 0.217 | 0.169 | 1.44E-07 |
| Tlr4 | 7.02E-12 | 0.300664 | 0.213 | 0.166 | 1.66E-07 |
| Il6st | 7.7E-12 | 0.290025 | 0.507 | 0.453 | 1.82E-07 |
| BC035044 | 9.08E-12 | 0.314423 | 0.258 | 0.207 | 2.14E-07 |
| Galnt12 | 1.57E-11 | 0.491229 | 0.204 | 0.159 | 3.71E-07 |
| Brox | 1.66E-11 | 0.338637 | 0.215 | 0.168 | 3.92E-07 |
| Zfp266 | 1.83E-11 | 0.336838 | 0.225 | 0.178 | 4.31E-07 |

| | | | | | |
|---|---|---|---|---|---|
| Ubqln1 | 2.57E-11 | 0.301028 | 0.299 | 0.248 | 6.07E-07 |
| Tbl1xr1 | 2.98E-11 | 0.471505 | 0.248 | 0.2 | 7.02E-07 |
| Klhdc3 | 3.33E-11 | 0.389951 | 0.219 | 0.174 | 7.87E-07 |
| Mxd4 | 3.37E-11 | 0.277729 | 0.33 | 0.278 | 7.96E-07 |
| Atp2a2 | 3.88E-11 | 0.296496 | 0.316 | 0.263 | 9.16E-07 |
| Gorasp2 | 3.99E-11 | 0.355906 | 0.346 | 0.296 | 9.41E-07 |
| B4galt4 | 4.78E-11 | 0.268298 | 0.307 | 0.255 | 1.13E-06 |
| Usf2 | 5.05E-11 | 0.266539 | 0.285 | 0.234 | 1.19E-06 |
| Aars | 6.6E-11 | 0.289629 | 0.355 | 0.304 | 1.56E-06 |
| Pttg1ip | 7.38E-11 | 0.259294 | 0.47 | 0.425 | 1.74E-06 |
| Zmynd11 | 7.81E-11 | 0.424985 | 0.274 | 0.227 | 1.84E-06 |
| Sipa1l1 | 7.9E-11 | 0.537666 | 0.218 | 0.173 | 1.86E-06 |
| Ppp2r1a | 8.46E-11 | 0.303288 | 0.448 | 0.4 | 2E-06 |
| Rhoq | 9.02E-11 | 0.32134 | 0.244 | 0.196 | 2.13E-06 |
| Flcn | 9.1E-11 | 0.286887 | 0.353 | 0.303 | 2.15E-06 |
| Actr1a | 1.1E-10 | 0.386386 | 0.331 | 0.284 | 2.6E-06 |
| Ppp2ca | 1.31E-10 | 0.271575 | 0.517 | 0.474 | 3.09E-06 |
| Ndfip2 | 1.44E-10 | 0.357555 | 0.333 | 0.287 | 3.41E-06 |
| Dnajb11 | 1.59E-10 | 0.269069 | 0.387 | 0.336 | 3.74E-06 |
| Plekha2 | 1.66E-10 | 0.372195 | 0.242 | 0.195 | 3.92E-06 |
| Psen1 | 1.78E-10 | 0.269214 | 0.391 | 0.339 | 4.19E-06 |
| Wsb1 | 1.79E-10 | 0.303249 | 0.407 | 0.362 | 4.22E-06 |
| Uba2 | 1.86E-10 | 0.334711 | 0.284 | 0.236 | 4.4E-06 |
| Bnip2 | 1.91E-10 | 0.290431 | 0.307 | 0.258 | 4.52E-06 |
| Mtmr3 | 1.92E-10 | 0.464282 | 0.217 | 0.173 | 4.54E-06 |
| Spcs3 | 2.03E-10 | 0.284659 | 0.284 | 0.236 | 4.79E-06 |
| Clcn4 | 2.04E-10 | 0.274432 | 0.406 | 0.358 | 4.81E-06 |
| Csnk1d | 2.25E-10 | 0.259288 | 0.351 | 0.3 | 5.31E-06 |
| D6Wsu163 | 2.32E-10 | 0.426738 | 0.2 | 0.157 | 5.48E-06 |
| Mknk1 | 2.54E-10 | 0.327608 | 0.493 | 0.452 | 6E-06 |
| Rab5a | 2.57E-10 | 0.514316 | 0.249 | 0.206 | 6.06E-06 |
| Cdc42se1 | 3E-10 | 0.264936 | 0.379 | 0.329 | 7.07E-06 |
| Ext2 | 3.21E-10 | 0.438387 | 0.265 | 0.222 | 7.58E-06 |
| Stau1 | 3.3E-10 | 0.395549 | 0.318 | 0.272 | 7.79E-06 |
| Ss18 | 3.33E-10 | 0.525744 | 0.237 | 0.196 | 7.86E-06 |
| Yod1 | 3.36E-10 | 0.524532 | 0.203 | 0.161 | 7.93E-06 |
| Dync1i2 | 3.44E-10 | 0.26171 | 0.432 | 0.377 | 8.11E-06 |
| Rnf139 | 3.44E-10 | 0.38904 | 0.253 | 0.211 | 8.12E-06 |
| Jun | 3.6E-10 | 0.402399 | 0.39 | 0.347 | 8.48E-06 |
| Ddb1 | 4.03E-10 | 0.30799 | 0.224 | 0.179 | 9.5E-06 |
| Kat7 | 4.15E-10 | 0.340223 | 0.236 | 0.193 | 9.78E-06 |
| Vhl | 4.29E-10 | 0.330992 | 0.237 | 0.193 | 1.01E-05 |
| Exosc10 | 5.17E-10 | 0.358343 | 0.216 | 0.173 | 1.22E-05 |
| Vbp1 | 5.28E-10 | 0.410105 | 0.225 | 0.183 | 1.25E-05 |
| Rnf2 | 5.5E-10 | 0.339607 | 0.211 | 0.168 | 1.3E-05 |
| G3bp1 | 6.96E-10 | 0.314487 | 0.431 | 0.385 | 1.64E-05 |
| Vps26a | 7.71E-10 | 0.263197 | 0.26 | 0.216 | 1.82E-05 |

| | | | | | |
|---|---|---|---|---|---|
| Galnt10 | 8.75E-10 | 0.25015 | 0.221 | 0.177 | 2.06E-05 |
| Ceacam1 | 9.48E-10 | 0.276656 | 0.234 | 0.191 | 2.24E-05 |
| Wapl | 9.76E-10 | 0.326227 | 0.402 | 0.359 | 2.3E-05 |
| Med13l | 1.01E-09 | 0.310788 | 0.297 | 0.251 | 2.39E-05 |
| Usf1 | 1.16E-09 | 0.309871 | 0.247 | 0.202 | 2.74E-05 |
| Pdk1 | 1.18E-09 | 0.28699 | 0.3 | 0.252 | 2.79E-05 |
| Yme1l1 | 1.27E-09 | 0.294704 | 0.409 | 0.362 | 3E-05 |
| Tk2 | 1.37E-09 | 0.453145 | 0.251 | 0.208 | 3.24E-05 |
| Jade2 | 1.42E-09 | 0.336413 | 0.244 | 0.2 | 3.35E-05 |
| Sh3gl1 | 1.5E-09 | 0.301868 | 0.243 | 0.2 | 3.54E-05 |
| Ppp1r21 | 1.97E-09 | 0.426421 | 0.288 | 0.245 | 4.66E-05 |
| Fbxw11 | 2.44E-09 | 0.341349 | 0.314 | 0.267 | 5.75E-05 |
| Ugt1a7c | 2.53E-09 | 0.321763 | 0.317 | 0.273 | 5.96E-05 |
| B3galnt1 | 2.53E-09 | 0.404092 | 0.209 | 0.17 | 5.97E-05 |
| Slc37a2 | 2.58E-09 | 0.299067 | 0.443 | 0.398 | 6.08E-05 |
| Srpr | 2.75E-09 | 0.300801 | 0.362 | 0.314 | 6.48E-05 |
| Npc1 | 2.93E-09 | 0.306906 | 0.38 | 0.33 | 6.92E-05 |
| Nfkb1 | 3.12E-09 | 0.281135 | 0.483 | 0.436 | 7.37E-05 |
| Slc4a7 | 3.18E-09 | 0.486907 | 0.236 | 0.195 | 7.5E-05 |
| Pla2g4a | 3.61E-09 | 0.506678 | 0.205 | 0.167 | 8.52E-05 |
| Hdac1 | 3.64E-09 | 0.336475 | 0.289 | 0.245 | 8.6E-05 |
| Ccdc71 | 3.75E-09 | 0.316404 | 0.2 | 0.161 | 8.84E-05 |
| Ddx3x | 4.19E-09 | 0.294637 | 0.397 | 0.352 | 9.87E-05 |
| Rxrb | 4.45E-09 | 0.317596 | 0.257 | 0.214 | 0.000105 |
| Tnfaip1 | 4.82E-09 | 0.470068 | 0.228 | 0.188 | 0.000114 |
| Tmem250- | 6.34E-09 | 0.267051 | 0.541 | 0.495 | 0.00015 |
| Smox | 6.4E-09 | 0.305763 | 0.319 | 0.276 | 0.000151 |
| Foxj2 | 6.67E-09 | 0.312552 | 0.224 | 0.182 | 0.000157 |
| Kdm5b | 7.22E-09 | 0.268605 | 0.216 | 0.175 | 0.00017 |
| Hsd17b4 | 7.22E-09 | 0.299893 | 0.348 | 0.307 | 0.00017 |
| 2900097C1 | 7.95E-09 | 0.356287 | 0.217 | 0.178 | 0.000188 |
| Vps4b | 8.26E-09 | 0.299415 | 0.335 | 0.29 | 0.000195 |
| Ctns | 8.66E-09 | 0.402252 | 0.22 | 0.181 | 0.000204 |
| Ufsp2 | 8.72E-09 | 0.33448 | 0.345 | 0.302 | 0.000206 |
| Snx2 | 9.48E-09 | 0.264247 | 0.406 | 0.362 | 0.000224 |
| Tbl1x | 9.64E-09 | 0.519576 | 0.235 | 0.197 | 0.000227 |
| Med1 | 1.02E-08 | 0.38194 | 0.404 | 0.367 | 0.000242 |
| Vwa5a | 1.13E-08 | 0.251653 | 0.223 | 0.181 | 0.000265 |
| Rnf216 | 1.13E-08 | 0.31272 | 0.307 | 0.264 | 0.000267 |
| Ssfa2 | 1.15E-08 | 0.274985 | 0.234 | 0.193 | 0.000271 |
| Zfp120 | 1.28E-08 | 0.526562 | 0.201 | 0.163 | 0.000303 |
| Letm1 | 1.36E-08 | 0.443472 | 0.217 | 0.179 | 0.000322 |
| Rraga | 1.59E-08 | 0.257209 | 0.259 | 0.217 | 0.000374 |
| Hnrnpu | 1.69E-08 | 0.274395 | 0.447 | 0.403 | 0.000398 |
| Smpd1 | 2.13E-08 | 0.411117 | 0.238 | 0.2 | 0.000502 |
| Ankrd44 | 2.31E-08 | 0.258022 | 0.471 | 0.426 | 0.000546 |
| Zfp532 | 2.44E-08 | 0.319968 | 0.346 | 0.299 | 0.000577 |

| | | | | | |
|---|---|---|---|---|---|
| Itfg1 | 2.7E-08 | 0.30629 | 0.34 | 0.297 | 0.000636 |
| Nrbp1 | 2.77E-08 | 0.385993 | 0.23 | 0.193 | 0.000654 |
| Papola | 2.92E-08 | 0.27674 | 0.445 | 0.404 | 0.000689 |
| Gna13 | 3.2E-08 | 0.374003 | 0.326 | 0.287 | 0.000756 |
| Hk3 | 3.89E-08 | 0.389636 | 0.251 | 0.212 | 0.000917 |
| Ppm1g | 3.9E-08 | 0.330119 | 0.306 | 0.264 | 0.000921 |
| Cds1 | 4.59E-08 | 0.272876 | 0.213 | 0.174 | 0.001083 |
| Mettl7a1 | 4.95E-08 | 0.357845 | 0.213 | 0.176 | 0.001167 |
| Fundc1 | 5.04E-08 | 0.386617 | 0.205 | 0.168 | 0.001188 |
| Kpnb1 | 5.25E-08 | 0.303308 | 0.272 | 0.23 | 0.001238 |
| Mib2 | 5.27E-08 | 0.335108 | 0.21 | 0.172 | 0.001244 |
| Srp68 | 5.32E-08 | 0.338698 | 0.204 | 0.167 | 0.001254 |
| Tpbgl | 5.37E-08 | 0.25244 | 0.393 | 0.351 | 0.001266 |
| Qrich1 | 5.94E-08 | 0.367159 | 0.277 | 0.238 | 0.0014 |
| 2510039O1 | 6.8E-08 | 0.345924 | 0.299 | 0.261 | 0.001605 |
| Dirc2 | 7.14E-08 | 0.263158 | 0.287 | 0.246 | 0.001685 |
| Slc2a8 | 7.94E-08 | 0.301731 | 0.205 | 0.169 | 0.001874 |
| Nelfb | 7.95E-08 | 0.514612 | 0.204 | 0.169 | 0.001875 |
| Nipa2 | 7.96E-08 | 0.416093 | 0.225 | 0.189 | 0.001877 |
| Serp1 | 8.03E-08 | 0.255499 | 0.426 | 0.39 | 0.001894 |
| Tbc1d14 | 8.09E-08 | 0.268432 | 0.296 | 0.253 | 0.001908 |
| Sar1a | 8.6E-08 | 0.342715 | 0.314 | 0.276 | 0.002028 |
| Chmp1b | 8.75E-08 | 0.345905 | 0.259 | 0.22 | 0.002065 |
| Clcn3 | 8.78E-08 | 0.317934 | 0.255 | 0.216 | 0.002071 |
| Tmem106b | 1.06E-07 | 0.351463 | 0.22 | 0.182 | 0.002512 |
| Map3k11 | 1.43E-07 | 0.273116 | 0.261 | 0.221 | 0.003385 |
| Mlxipl | 1.46E-07 | 0.266648 | 0.401 | 0.36 | 0.003449 |
| Pum1 | 1.68E-07 | 0.309389 | 0.218 | 0.181 | 0.003955 |
| Aagab | 1.9E-07 | 0.565466 | 0.233 | 0.2 | 0.004486 |
| Rp2 | 1.99E-07 | 0.324473 | 0.246 | 0.207 | 0.004702 |
| Mgrn1 | 2.16E-07 | 0.291435 | 0.35 | 0.31 | 0.005102 |
| Ccz1 | 2.23E-07 | 0.353609 | 0.289 | 0.249 | 0.00527 |
| Usp24 | 2.34E-07 | 0.264121 | 0.263 | 0.225 | 0.005514 |
| Mknk2 | 2.37E-07 | 0.283933 | 0.233 | 0.197 | 0.005586 |
| Txndc5 | 2.43E-07 | 0.400656 | 0.216 | 0.18 | 0.005733 |
| Gnaq | 2.84E-07 | 0.342725 | 0.446 | 0.416 | 0.006706 |
| Prpf6 | 3.02E-07 | 0.427322 | 0.215 | 0.18 | 0.007112 |
| Zfp148 | 3.41E-07 | 0.273367 | 0.245 | 0.208 | 0.008041 |
| Golga7 | 3.78E-07 | 0.35273 | 0.231 | 0.197 | 0.008918 |
| Etv1 | 4.11E-07 | 0.295896 | 0.221 | 0.185 | 0.009689 |
| Rnf180 | 4.38E-07 | 0.400879 | 0.221 | 0.188 | 0.010325 |
| Hnrnph2 | 4.54E-07 | 0.508829 | 0.259 | 0.227 | 0.010716 |
| Cat | 5.44E-07 | 0.334379 | 0.231 | 0.197 | 0.012834 |
| Stxbp3 | 5.67E-07 | 0.441042 | 0.206 | 0.175 | 0.013376 |
| Ggps1 | 5.99E-07 | 0.349811 | 0.256 | 0.219 | 0.014127 |
| Dennd1c | 6.43E-07 | 0.256356 | 0.289 | 0.25 | 0.01516 |
| BC037034 | 6.82E-07 | 0.256572 | 0.333 | 0.295 | 0.016089 |

| | | | | | |
|---|---|---|---|---|---|
| Thumpd3 | 6.98E-07 | 0.25359 | 0.26 | 0.225 | 0.016466 |
| Slmap | 6.99E-07 | 0.372228 | 0.251 | 0.216 | 0.016479 |
| Scaf11 | 7.28E-07 | 0.255057 | 0.4 | 0.364 | 0.017172 |
| Vps18 | 8.31E-07 | 0.264478 | 0.271 | 0.233 | 0.019609 |
| Mfsd14b | 8.59E-07 | 0.284794 | 0.206 | 0.174 | 0.020257 |
| Wbp1l | 8.64E-07 | 0.426075 | 0.222 | 0.188 | 0.020385 |
| Tmx2 | 8.65E-07 | 0.296124 | 0.273 | 0.239 | 0.020412 |
| Vamp3 | 8.88E-07 | 0.256675 | 0.347 | 0.308 | 0.020948 |
| Arnt | 9.25E-07 | 0.343047 | 0.205 | 0.171 | 0.021818 |
| Slc46a3 | 1.01E-06 | 0.277747 | 0.324 | 0.286 | 0.02389 |
| Psmd1 | 1.14E-06 | 0.307565 | 0.307 | 0.272 | 0.026823 |
| Dpf2 | 1.19E-06 | 0.251483 | 0.227 | 0.194 | 0.028159 |
| Phactr2 | 1.23E-06 | 0.346781 | 0.268 | 0.234 | 0.029014 |
| Rab6a | 1.43E-06 | 0.386899 | 0.205 | 0.173 | 0.033844 |
| Fnbp1 | 1.54E-06 | 0.29803 | 0.501 | 0.463 | 0.036363 |
| Zbtb1 | 1.64E-06 | 0.367647 | 0.234 | 0.2 | 0.03863 |
| Slc35a4 | 1.79E-06 | 0.372434 | 0.212 | 0.18 | 0.042283 |
| Dars | 1.81E-06 | 0.40243 | 0.22 | 0.188 | 0.042711 |
| Sumf1 | 1.91E-06 | 0.336188 | 0.237 | 0.205 | 0.045083 |
| Gramd1a | 2.07E-06 | 0.408473 | 0.294 | 0.259 | 0.048797 |

### Supplementry Table 1. Marker genes for IntM

| Genenames | p_val | avg_log2FC | pct.1 | pct.2 | p_val_adj |
|---|---|---|---|---|---|
| H2-D1 | 1.22E-87 | 0.647789 | 0.947 | 0.9 | 2.87E-83 |
| Lyz2 | 1.17E-46 | 0.473459 | 0.755 | 0.663 | 2.77E-42 |
| Malat1 | 8.89E-45 | 0.36874 | 0.984 | 0.989 | 2.1E-40 |
| H2-K1 | 2.14E-44 | 0.537281 | 0.874 | 0.811 | 5.05E-40 |
| B2m | 2.47E-42 | 0.378746 | 0.999 | 0.998 | 5.82E-38 |
| Ctss | 1.33E-33 | 0.261354 | 1 | 1 | 3.13E-29 |
| Fth1 | 1.48E-32 | 0.252863 | 0.996 | 0.994 | 3.49E-28 |
| Gm10800 | 5.47E-30 | 0.482009 | 0.753 | 0.711 | 1.29E-25 |
| Rpl18a | 1.18E-29 | 0.369343 | 0.989 | 0.986 | 2.79E-25 |
| Rplp1 | 1.42E-24 | 0.374559 | 0.956 | 0.957 | 3.36E-20 |
| Rps11 | 1.71E-21 | 0.369985 | 0.915 | 0.921 | 4.03E-17 |
| Rpl37a | 4.63E-19 | 0.325911 | 0.945 | 0.939 | 1.09E-14 |
| Gm24270 | 6.76E-19 | 0.318493 | 0.779 | 0.749 | 1.59E-14 |
| Gm26740 | 6.59E-18 | 0.731211 | 0.207 | 0.142 | 1.55E-13 |
| Gm26917 | 8.84E-18 | 0.448333 | 0.943 | 0.947 | 2.09E-13 |
| Rps27 | 7.1E-17 | 0.365709 | 0.957 | 0.957 | 1.67E-12 |
| Rps28 | 8.9E-17 | 0.368065 | 0.921 | 0.908 | 2.1E-12 |
| Gm23935 | 1.33E-16 | 0.294449 | 0.962 | 0.965 | 3.14E-12 |
| Gm10801 | 5.19E-16 | 0.286428 | 0.328 | 0.258 | 1.22E-11 |
| Rpl13 | 1.33E-15 | 0.348819 | 0.943 | 0.945 | 3.14E-11 |
| Rpl32 | 1.54E-15 | 0.357151 | 0.895 | 0.88 | 3.63E-11 |
| AU020206 | 2.2E-15 | 0.478241 | 0.584 | 0.523 | 5.19E-11 |
| Lsp1 | 9.66E-15 | 0.589646 | 0.404 | 0.332 | 2.28E-10 |
| Rpl21 | 1.19E-14 | 0.343641 | 0.918 | 0.921 | 2.8E-10 |
| Lars2 | 2.58E-14 | 0.33544 | 0.933 | 0.924 | 6.08E-10 |
| Rps29 | 3.08E-14 | 0.304746 | 0.989 | 0.977 | 7.26E-10 |
| Emp3 | 4.94E-14 | 0.699011 | 0.218 | 0.161 | 1.16E-09 |
| Rps9 | 7.61E-14 | 0.284444 | 0.962 | 0.958 | 1.79E-09 |
| Rps12 | 9.72E-14 | 0.356111 | 0.838 | 0.833 | 2.29E-09 |
| Rps18 | 9.97E-14 | 0.357744 | 0.854 | 0.854 | 2.35E-09 |
| Rplp2 | 2.58E-13 | 0.322778 | 0.897 | 0.907 | 6.09E-09 |
| Rps14 | 2.59E-13 | 0.330277 | 0.919 | 0.926 | 6.12E-09 |
| Rpl41 | 2.86E-13 | 0.289415 | 0.936 | 0.947 | 6.74E-09 |
| Rpl37 | 4.1E-13 | 0.343696 | 0.899 | 0.9 | 9.67E-09 |
| Ctsh | 1.13E-11 | 0.259041 | 0.954 | 0.95 | 2.66E-07 |
| Rpl36 | 8.19E-11 | 0.366526 | 0.823 | 0.823 | 1.93E-06 |
| Gm10076 | 1.06E-10 | 0.252702 | 0.902 | 0.909 | 2.5E-06 |
| Gm12435 | 1.26E-10 | 0.344037 | 0.157 | 0.213 | 2.97E-06 |
| Rps23 | 2.69E-10 | 0.284532 | 0.895 | 0.901 | 6.33E-06 |
| Gm15564 | 3.4E-10 | 0.287843 | 0.601 | 0.545 | 8.03E-06 |
| Rps13 | 5.25E-10 | 0.435671 | 0.778 | 0.785 | 1.24E-05 |
| Rps20 | 6.48E-10 | 0.364429 | 0.83 | 0.83 | 1.53E-05 |
| Tpt1 | 7.72E-10 | 0.347088 | 0.87 | 0.885 | 1.82E-05 |

| | | | | | |
|---|---|---|---|---|---|
| Tmsb4x | 9.13E-10 | 0.257716 | 1 | 1 | 2.15E-05 |
| H2-M3 | 1.77E-09 | 0.603922 | 0.259 | 0.209 | 4.18E-05 |
| Rpl30 | 2.09E-09 | 0.366033 | 0.862 | 0.871 | 4.93E-05 |
| Neat1 | 3.71E-09 | 0.393897 | 0.258 | 0.208 | 8.76E-05 |
| Rpl23a | 4.63E-09 | 0.313535 | 0.773 | 0.77 | 0.000109 |
| Ifi30 | 2.15E-08 | 0.414669 | 0.529 | 0.491 | 0.000508 |
| Rps21 | 2.6E-08 | 0.326974 | 0.781 | 0.776 | 0.000613 |
| Rps19 | 3.75E-08 | 0.289313 | 0.864 | 0.88 | 0.000884 |
| Rpl35 | 4.21E-08 | 0.364522 | 0.816 | 0.825 | 0.000993 |
| Rps24 | 4.87E-08 | 0.344586 | 0.843 | 0.839 | 0.00115 |
| Fau | 8E-08 | 0.283317 | 0.896 | 0.91 | 0.001888 |
| Rpl23 | 8.32E-08 | 0.346344 | 0.826 | 0.839 | 0.001963 |
| Rplp0 | 2.75E-07 | 0.287367 | 0.891 | 0.9 | 0.006492 |
| Rn7sk | 8.38E-07 | 0.343878 | 0.587 | 0.561 | 0.019772 |
| Gm15536 | 9.49E-07 | 0.28685 | 0.27 | 0.233 | 0.02239 |
| Rpl34 | 9.57E-07 | 0.316595 | 0.785 | 0.797 | 0.022575 |
| Gm28437 | 1.3E-06 | 0.273542 | 0.82 | 0.832 | 0.030654 |

### Supplementry Table 1. Marker genes for eDAM

| Genenames | p_val | avg_log2FC | pct.1 | pct.2 | p_val_adj |
|---|---|---|---|---|---|
| Apoe | 0 | 3.028703 | 0.998 | 0.411 | 0 |
| Lyz2 | 8.6E-105 | 1.005077 | 0.831 | 0.661 | 2E-100 |
| Cd63 | 2.48E-48 | 0.544221 | 0.837 | 0.718 | 5.85E-44 |
| H2-D1 | 7.83E-46 | 0.622658 | 0.946 | 0.905 | 1.85E-41 |
| Cd63-ps | 2E-38 | 0.431696 | 0.661 | 0.514 | 4.71E-34 |
| Rpl18a | 3.39E-35 | 0.366637 | 0.988 | 0.987 | 8.01E-31 |
| Ctss | 9.65E-35 | 0.298607 | 1 | 1 | 2.28E-30 |
| Fth1 | 6.56E-32 | 0.326757 | 0.995 | 0.995 | 1.55E-27 |
| Rps28 | 2.99E-31 | 0.393874 | 0.943 | 0.905 | 7.06E-27 |
| Rps9 | 5.93E-31 | 0.396612 | 0.977 | 0.956 | 1.4E-26 |
| B2m | 1.46E-29 | 0.348715 | 0.998 | 0.998 | 3.44E-25 |
| Rps27 | 3.2E-29 | 0.38439 | 0.975 | 0.954 | 7.56E-25 |
| Rpl32 | 5.76E-28 | 0.412582 | 0.918 | 0.878 | 1.36E-23 |
| Rpl41 | 3.29E-25 | 0.348018 | 0.97 | 0.94 | 7.75E-21 |
| Gm10076 | 2.42E-24 | 0.404364 | 0.941 | 0.902 | 5.72E-20 |
| Rps12 | 3.1E-24 | 0.421412 | 0.878 | 0.827 | 7.32E-20 |
| Rps18 | 4.42E-24 | 0.399186 | 0.891 | 0.848 | 1.04E-19 |
| Rps19 | 5.87E-24 | 0.397288 | 0.91 | 0.871 | 1.38E-19 |
| Rps23 | 6.43E-24 | 0.356239 | 0.927 | 0.895 | 1.52E-19 |
| Rps29 | 7.42E-24 | 0.287992 | 0.986 | 0.979 | 1.75E-19 |
| Rpl37a | 1.81E-23 | 0.367474 | 0.96 | 0.937 | 4.28E-19 |
| Ctsl | 1.6E-22 | 0.305581 | 0.965 | 0.948 | 3.78E-18 |
| Rpl35 | 3.31E-22 | 0.384847 | 0.87 | 0.815 | 7.8E-18 |
| Gm4149 | 4.25E-22 | 0.405139 | 0.876 | 0.829 | 1E-17 |
| Rplp2 | 3.5E-21 | 0.329552 | 0.929 | 0.9 | 8.25E-17 |
| Rps14 | 5.24E-21 | 0.329938 | 0.952 | 0.919 | 1.24E-16 |
| Rplp1 | 6.62E-20 | 0.292956 | 0.964 | 0.956 | 1.56E-15 |
| Fau | 2.65E-19 | 0.295973 | 0.934 | 0.902 | 6.26E-15 |
| H2-K1 | 6.76E-19 | 0.470708 | 0.859 | 0.821 | 1.59E-14 |
| Rpl37 | 1.35E-18 | 0.328887 | 0.926 | 0.895 | 3.18E-14 |
| Gm15427 | 2.28E-18 | 0.440115 | 0.964 | 0.946 | 5.37E-14 |
| Gm3511 | 2.59E-18 | 0.373901 | 0.654 | 0.562 | 6.1E-14 |
| Rps24 | 5.68E-18 | 0.347369 | 0.873 | 0.834 | 1.34E-13 |
| Rps20 | 6.72E-18 | 0.365256 | 0.874 | 0.823 | 1.58E-13 |
| Rpl30 | 7.64E-18 | 0.359753 | 0.901 | 0.864 | 1.8E-13 |
| Gm14303 | 9.83E-18 | 0.269602 | 0.833 | 0.771 | 2.32E-13 |
| Gm9843 | 1.91E-17 | 0.401072 | 0.88 | 0.844 | 4.51E-13 |
| Rpl23 | 2.72E-17 | 0.405444 | 0.874 | 0.83 | 6.41E-13 |
| Rps11 | 3.36E-17 | 0.292804 | 0.939 | 0.917 | 7.91E-13 |
| Npc2 | 9.28E-17 | 0.373548 | 0.879 | 0.834 | 2.19E-12 |
| Ctsh | 1.19E-16 | 0.329812 | 0.958 | 0.949 | 2.8E-12 |
| Rpl21 | 1.99E-16 | 0.308456 | 0.936 | 0.918 | 4.7E-12 |
| Rpl17 | 5.19E-16 | 0.398885 | 0.834 | 0.789 | 1.23E-11 |

| | | | | | |
|---|---|---|---|---|---|
| Rpl13 | 1.01E-15 | 0.292544 | 0.956 | 0.942 | 2.38E-11 |
| Rps25 | 1.65E-15 | 0.341402 | 0.888 | 0.857 | 3.89E-11 |
| Ftl1-ps1 | 3.92E-15 | 0.270175 | 0.918 | 0.887 | 9.24E-11 |
| Rps27rt | 5.01E-15 | 0.344799 | 0.783 | 0.704 | 1.18E-10 |
| Rpl36 | 5.7E-15 | 0.349596 | 0.865 | 0.816 | 1.34E-10 |
| Rps21 | 2.4E-14 | 0.374848 | 0.821 | 0.77 | 5.66E-10 |
| Rpl10a | 3.17E-14 | 0.462519 | 0.718 | 0.666 | 7.47E-10 |
| Gm29216 | 3.71E-14 | 0.285938 | 0.806 | 0.766 | 8.75E-10 |
| Rpl23a | 6.45E-14 | 0.327426 | 0.808 | 0.764 | 1.52E-09 |
| Gm11361 | 7.52E-14 | 0.35918 | 0.542 | 0.463 | 1.77E-09 |
| Rps16 | 9.71E-14 | 0.279101 | 0.815 | 0.773 | 2.29E-09 |
| Gm10443 | 1.17E-13 | 0.314123 | 0.479 | 0.402 | 2.76E-09 |
| Rps17 | 1.38E-13 | 0.308622 | 0.78 | 0.731 | 3.25E-09 |
| Cd9 | 2.16E-13 | 0.271557 | 0.978 | 0.967 | 5.08E-09 |
| Gm5905 | 2.78E-13 | 0.321284 | 0.773 | 0.708 | 6.56E-09 |
| Eef1a1 | 3.81E-13 | 0.251479 | 0.993 | 0.994 | 8.99E-09 |
| Rpl34 | 5.06E-13 | 0.310841 | 0.828 | 0.789 | 1.19E-08 |
| 2410006H1 | 9.62E-13 | 0.849156 | 0.282 | 0.216 | 2.27E-08 |
| Rps4x | 1.97E-12 | 0.372113 | 0.843 | 0.802 | 4.65E-08 |
| Rpl13a | 4.56E-12 | 0.315672 | 0.866 | 0.829 | 1.08E-07 |
| Gm2000 | 7.2E-12 | 0.351404 | 0.655 | 0.588 | 1.7E-07 |
| Rps16-ps2 | 1.04E-11 | 0.305831 | 0.65 | 0.579 | 2.45E-07 |
| Gas5 | 1.13E-11 | 0.411243 | 0.432 | 0.355 | 2.68E-07 |
| Gm10132 | 1.14E-11 | 0.417636 | 0.654 | 0.576 | 2.7E-07 |
| Gm28437 | 1.18E-11 | 0.355641 | 0.855 | 0.825 | 2.78E-07 |
| Rps3a1 | 1.33E-11 | 0.33516 | 0.866 | 0.839 | 3.13E-07 |
| Rpl12 | 1.59E-11 | 0.367205 | 0.731 | 0.675 | 3.74E-07 |
| Rpl19 | 2.35E-11 | 0.277027 | 0.842 | 0.806 | 5.53E-07 |
| Rps26 | 2.49E-11 | 0.37174 | 0.729 | 0.692 | 5.87E-07 |
| Rpl10a-ps1 | 2.62E-11 | 0.407736 | 0.618 | 0.55 | 6.19E-07 |
| Rps5 | 3.19E-11 | 0.378279 | 0.767 | 0.736 | 7.53E-07 |
| Lag3 | 3.41E-11 | 0.295218 | 0.802 | 0.766 | 8.05E-07 |
| Gm10736 | 3.78E-11 | 0.256152 | 0.594 | 0.518 | 8.91E-07 |
| Rps7 | 4.3E-11 | 0.319512 | 0.822 | 0.787 | 1.01E-06 |
| Rpl31 | 4.88E-11 | 0.289813 | 0.823 | 0.791 | 1.15E-06 |
| Rps27a | 7.23E-11 | 0.302001 | 0.826 | 0.794 | 1.71E-06 |
| mt-Co3 | 1.49E-10 | 0.392843 | 0.925 | 0.901 | 3.5E-06 |
| Rps18-ps3 | 2.19E-10 | 0.434792 | 0.642 | 0.577 | 5.17E-06 |
| Uba52 | 2.76E-10 | 0.29606 | 0.802 | 0.742 | 6.51E-06 |
| Lsp1 | 2.99E-10 | 0.522421 | 0.407 | 0.339 | 7.06E-06 |
| Cd52 | 4.13E-10 | 0.39627 | 0.545 | 0.476 | 9.75E-06 |
| Gm9794 | 6.95E-10 | 0.265054 | 0.684 | 0.628 | 1.64E-05 |
| Rpl14 | 8.29E-10 | 0.262897 | 0.813 | 0.772 | 1.96E-05 |
| mt-Co1 | 9.73E-10 | 0.266906 | 0.989 | 0.985 | 2.3E-05 |
| Rps10 | 9.94E-10 | 0.333161 | 0.836 | 0.798 | 2.34E-05 |
| Gm7331 | 1.11E-09 | 0.339663 | 0.624 | 0.556 | 2.63E-05 |
| Lgals3bp | 2.1E-09 | 0.258679 | 0.644 | 0.582 | 4.96E-05 |

| | | | | | |
|---|---|---|---|---|---|
| Zfas1 | 2.8E-09 | 0.607461 | 0.203 | 0.15 | 6.59E-05 |
| Rps13 | 2.98E-09 | 0.29696 | 0.807 | 0.779 | 7.02E-05 |
| Tmsb10 | 3.09E-09 | 0.338332 | 0.492 | 0.428 | 7.28E-05 |
| Rpl27a | 3.19E-09 | 0.274959 | 0.782 | 0.744 | 7.52E-05 |
| Rpl19-ps11 | 3.32E-09 | 0.344481 | 0.67 | 0.611 | 7.84E-05 |
| Gm15710 | 4.13E-09 | 0.287168 | 0.789 | 0.74 | 9.75E-05 |
| Naalad2 | 4.73E-09 | 0.743007 | 0.22 | 0.166 | 0.000112 |
| Rpl39 | 4.79E-09 | 0.315277 | 0.768 | 0.713 | 0.000113 |
| Gm11478 | 5.27E-09 | 0.288306 | 0.729 | 0.674 | 0.000124 |
| Tpt1 | 5.58E-09 | 0.269294 | 0.895 | 0.879 | 0.000132 |
| Timp2 | 5.91E-09 | 0.266752 | 0.854 | 0.829 | 0.000139 |
| Rpl11 | 6.87E-09 | 0.300755 | 0.76 | 0.725 | 0.000162 |
| mt-Atp6 | 7.83E-09 | 0.336701 | 0.857 | 0.826 | 0.000185 |
| Rpl26 | 1.1E-08 | 0.311081 | 0.797 | 0.765 | 0.000259 |
| Rpl38 | 1.17E-08 | 0.254545 | 0.691 | 0.64 | 0.000276 |
| mt-Cytb | 1.65E-08 | 0.279459 | 0.972 | 0.962 | 0.00039 |
| Rpsa | 1.74E-08 | 0.290931 | 0.82 | 0.797 | 0.000411 |
| Hebp1 | 1.81E-08 | 0.588667 | 0.227 | 0.174 | 0.000426 |
| Rps2 | 2.11E-08 | 0.282895 | 0.737 | 0.689 | 0.000499 |
| Gm7536 | 2.41E-08 | 0.377723 | 0.641 | 0.581 | 0.000569 |
| Rps10-ps2 | 2.77E-08 | 0.350442 | 0.646 | 0.59 | 0.000654 |
| Rpl31-ps8 | 2.95E-08 | 0.286555 | 0.64 | 0.591 | 0.000695 |
| Rpl15 | 3.67E-08 | 0.272624 | 0.73 | 0.682 | 0.000865 |
| Gm6977 | 3.95E-08 | 0.357787 | 0.579 | 0.519 | 0.000931 |
| Gm5805 | 4.24E-08 | 0.300681 | 0.556 | 0.501 | 0.000999 |
| Gm8730 | 5.29E-08 | 0.275518 | 0.776 | 0.736 | 0.001248 |
| Gm10073 | 6.78E-08 | 0.305061 | 0.651 | 0.597 | 0.001599 |
| Gm10275 | 8.72E-08 | 0.325557 | 0.465 | 0.404 | 0.002058 |
| Gm10925 | 9.66E-08 | 0.361372 | 0.854 | 0.822 | 0.00228 |
| Rpl36a | 1.23E-07 | 0.356574 | 0.578 | 0.528 | 0.00291 |
| Rpl17-ps3 | 1.24E-07 | 0.33101 | 0.449 | 0.397 | 0.002936 |
| Gm8430 | 1.45E-07 | 0.376399 | 0.573 | 0.517 | 0.00343 |
| Rpl6 | 1.48E-07 | 0.28194 | 0.79 | 0.761 | 0.003494 |
| Rpl22 | 1.79E-07 | 0.303883 | 0.594 | 0.542 | 0.004221 |
| Rpl7 | 1.81E-07 | 0.25987 | 0.832 | 0.798 | 0.004271 |
| Gm11808 | 2.92E-07 | 0.300055 | 0.551 | 0.497 | 0.00688 |
| Gm9616 | 4.23E-07 | 0.266857 | 0.525 | 0.47 | 0.009967 |
| Rpl3 | 4.31E-07 | 0.280533 | 0.836 | 0.832 | 0.010158 |
| Rpl22l1 | 4.49E-07 | 0.30179 | 0.307 | 0.251 | 0.010588 |
| Rps15a | 5.04E-07 | 0.352135 | 0.648 | 0.612 | 0.01188 |
| Rpsa-ps10 | 5.25E-07 | 0.322658 | 0.611 | 0.563 | 0.012387 |
| Rps15 | 7.81E-07 | 0.250441 | 0.776 | 0.754 | 0.018433 |
| Gm6136 | 1.11E-06 | 0.271686 | 0.581 | 0.532 | 0.026089 |
| Gm28661 | 1.14E-06 | 0.279719 | 0.751 | 0.735 | 0.026934 |
| mt-Co2 | 1.36E-06 | 0.322305 | 0.863 | 0.834 | 0.032046 |
| Rpl19-ps1 | 1.57E-06 | 0.279412 | 0.275 | 0.228 | 0.037018 |
| Gm10095 | 1.6E-06 | 0.278807 | 0.487 | 0.434 | 0.037767 |

| | | | | | |
|---|---|---|---|---|---|
| AU020206 | 2.12E-06 | 0.362464 | 0.581 | 0.53 | 0.04999 |

### Supplementry Table 1. Marker genes for DAM

| Genenames | p_val | avg_log2FC | pct.1 | pct.2 | p_val_adj |
|---|---|---|---|---|---|
| Clec7a | 0 | 4.28328 | 0.7 | 0.072 | 0 |
| Lpl | 0 | 6.013262 | 0.565 | 0.072 | 0 |
| Gas2l3 | 1E-227 | 4.210664 | 0.24 | 0.014 | 2.5E-223 |
| Itgax | 1.2E-227 | 3.732557 | 0.289 | 0.021 | 2.9E-223 |
| Cst7 | 1.5E-209 | 3.780326 | 0.473 | 0.07 | 3.5E-205 |
| Lilrb4a | 3.2E-203 | 3.525115 | 0.283 | 0.023 | 7.5E-199 |
| Lgals3 | 7.5E-180 | 6.592606 | 0.424 | 0.069 | 1.8E-175 |
| Csf1 | 2.6E-162 | 3.432179 | 0.377 | 0.055 | 6.2E-158 |
| Cd63 | 2.7E-149 | 2.171881 | 0.989 | 0.725 | 6.5E-145 |
| Fam20c | 1.4E-147 | 3.479756 | 0.316 | 0.042 | 3.4E-143 |
| Axl | 2E-146 | 2.987874 | 0.408 | 0.072 | 4.7E-142 |
| Ccl4 | 3.7E-140 | 5.570311 | 0.251 | 0.028 | 8.7E-136 |
| Cd63-ps | 6E-136 | 1.95588 | 0.935 | 0.52 | 1.4E-131 |
| Cd74 | 3.9E-127 | 3.206625 | 0.614 | 0.195 | 9.2E-123 |
| Fgr | 6E-123 | 3.357827 | 0.24 | 0.029 | 1.4E-118 |
| Ctsb | 6.5E-117 | 1.33963 | 1 | 0.998 | 1.5E-112 |
| Olfr111 | 5.4E-115 | 2.346728 | 0.28 | 0.042 | 1.3E-110 |
| Ccl3 | 6E-102 | 5.342624 | 0.307 | 0.057 | 1.42E-97 |
| Ftl1 | 1.1E-100 | 1.651806 | 1 | 0.993 | 2.69E-96 |
| Ank | 1.26E-96 | 2.48576 | 0.401 | 0.096 | 2.97E-92 |
| Cstb | 1.45E-96 | 2.202868 | 0.686 | 0.268 | 3.41E-92 |
| Capg | 3.29E-96 | 2.657219 | 0.554 | 0.188 | 7.75E-92 |
| Chst2 | 7.77E-96 | 2.900751 | 0.251 | 0.04 | 1.83E-91 |
| Plaur | 1.16E-94 | 3.045302 | 0.33 | 0.069 | 2.73E-90 |
| Cd9 | 2.03E-88 | 1.37603 | 1 | 0.967 | 4.79E-84 |
| Ftl2-ps | 5.66E-88 | 1.600443 | 0.998 | 0.926 | 1.33E-83 |
| Cox6a2 | 7.24E-88 | 2.610622 | 0.209 | 0.03 | 1.71E-83 |
| Ftl1-ps1 | 2.15E-84 | 1.564791 | 0.991 | 0.888 | 5.07E-80 |
| Cxcl16 | 2.77E-82 | 2.265556 | 0.632 | 0.26 | 6.53E-78 |
| Cd83 | 1.21E-80 | 2.001037 | 0.684 | 0.301 | 2.85E-76 |
| Pkm | 6.61E-79 | 1.921542 | 0.872 | 0.557 | 1.56E-74 |
| Pld3 | 1.22E-75 | 1.764691 | 0.879 | 0.569 | 2.88E-71 |
| Mif | 1.95E-71 | 2.057516 | 0.693 | 0.349 | 4.61E-67 |
| Ctsz | 6.6E-71 | 1.063891 | 1 | 0.993 | 1.56E-66 |
| Fxyd5 | 2.66E-69 | 3.299336 | 0.312 | 0.082 | 6.27E-65 |
| Il2rg | 2.97E-68 | 2.734948 | 0.253 | 0.055 | 7.01E-64 |
| Mt1 | 2.99E-68 | 1.573815 | 0.664 | 0.302 | 7.06E-64 |
| Gm6560 | 2.7E-65 | 1.297371 | 0.628 | 0.291 | 6.37E-61 |
| Mfsd12 | 6.41E-63 | 2.093964 | 0.433 | 0.149 | 1.51E-58 |
| Fth1 | 2.42E-62 | 1.469027 | 1 | 0.995 | 5.72E-58 |
| Aldoa | 4.35E-62 | 1.483335 | 0.917 | 0.69 | 1.03E-57 |
| Aplp2 | 4.5E-62 | 2.032 | 0.578 | 0.261 | 1.06E-57 |
| Pgk1 | 1.47E-61 | 1.67698 | 0.48 | 0.175 | 3.46E-57 |

| | | | | | |
|---|---|---|---|---|---|
| Tpi1 | 1.76E-61 | 1.66514 | 0.626 | 0.287 | 4.16E-57 |
| Gm15590 | 1.66E-60 | 1.07384 | 0.776 | 0.487 | 3.93E-56 |
| Mir692-2 | 3.59E-59 | 0.926781 | 0.657 | 0.337 | 8.47E-55 |
| Gapdh | 9.65E-58 | 1.253292 | 0.969 | 0.888 | 2.28E-53 |
| C3ar1 | 9.68E-58 | 1.561206 | 0.825 | 0.557 | 2.28E-53 |
| Apoe | 1.04E-57 | 3.563038 | 0.744 | 0.486 | 2.46E-53 |
| Ctsd | 4.29E-57 | 0.870972 | 1 | 1 | 1.01E-52 |
| Rhoc | 2.16E-56 | 2.146177 | 0.321 | 0.094 | 5.1E-52 |
| Clic4 | 4.08E-56 | 3.323458 | 0.231 | 0.055 | 9.62E-52 |
| Cd52 | 5.81E-56 | 1.667604 | 0.778 | 0.475 | 1.37E-51 |
| C230062I1( | 2.68E-54 | 0.30587 | 0.466 | 0.179 | 6.33E-50 |
| Atp6v0c-ps | 3.69E-54 | 1.120067 | 0.883 | 0.637 | 8.7E-50 |
| Prdx1 | 1.69E-52 | 1.512396 | 0.791 | 0.499 | 3.98E-48 |
| Cd68 | 5.44E-51 | 1.024354 | 0.991 | 0.948 | 1.28E-46 |
| Cd72 | 2.49E-50 | 2.984443 | 0.204 | 0.047 | 5.88E-46 |
| Atp6v0c | 1.56E-49 | 1.122184 | 0.948 | 0.819 | 3.69E-45 |
| Ftl1-ps2 | 4.08E-49 | 0.864773 | 0.583 | 0.305 | 9.63E-45 |
| C530043K1 | 5.97E-49 | 2.303621 | 0.265 | 0.073 | 1.41E-44 |
| Ldha | 3.1E-48 | 1.639972 | 0.783 | 0.531 | 7.31E-44 |
| Apbb2 | 5.53E-48 | 1.604343 | 0.397 | 0.147 | 1.31E-43 |
| Anxa5 | 7.53E-47 | 2.083092 | 0.43 | 0.18 | 1.78E-42 |
| Adssl1 | 1.15E-45 | 2.12853 | 0.249 | 0.07 | 2.71E-41 |
| Tubb6 | 6.13E-45 | 2.755581 | 0.229 | 0.062 | 1.45E-40 |
| Hmox1 | 2.32E-44 | 1.99231 | 0.363 | 0.134 | 5.46E-40 |
| Lyz2 | 1E-43 | 1.974834 | 0.89 | 0.677 | 2.36E-39 |
| Pea15a | 5.17E-43 | 1.251592 | 0.693 | 0.401 | 1.22E-38 |
| Hif1a | 6.94E-42 | 1.615637 | 0.603 | 0.322 | 1.64E-37 |
| Myo1e | 8.65E-42 | 1.831366 | 0.262 | 0.08 | 2.04E-37 |
| Cpd | 2.65E-41 | 1.853981 | 0.274 | 0.087 | 6.25E-37 |
| Gm2574 | 3.96E-41 | 1.056326 | 0.691 | 0.421 | 9.34E-37 |
| Glipr1 | 1.1E-40 | 2.009726 | 0.224 | 0.063 | 2.61E-36 |
| Gm6977 | 3.4E-40 | 1.165279 | 0.756 | 0.519 | 8.02E-36 |
| Rpsa | 4.2E-40 | 1.048754 | 0.928 | 0.796 | 9.91E-36 |
| Slc16a3 | 6.05E-40 | 1.746696 | 0.422 | 0.181 | 1.43E-35 |
| Sulf2 | 7.89E-40 | 1.970995 | 0.224 | 0.064 | 1.86E-35 |
| Id2 | 9.41E-39 | 2.454527 | 0.359 | 0.147 | 2.22E-34 |
| Gm5559 | 6.86E-38 | 1.066533 | 0.897 | 0.7 | 1.62E-33 |
| Cadm1 | 7.55E-38 | 1.189754 | 0.661 | 0.393 | 1.78E-33 |
| Nrp2 | 4.1E-37 | 1.854931 | 0.33 | 0.127 | 9.67E-33 |
| Hsp90ab1 | 1.37E-36 | 0.938506 | 0.944 | 0.838 | 3.24E-32 |
| Atp1a3 | 4.74E-36 | 1.619679 | 0.211 | 0.061 | 1.12E-31 |
| Tlr2 | 8.64E-36 | 1.853061 | 0.608 | 0.357 | 2.04E-31 |
| Rps2 | 6.34E-35 | 0.999458 | 0.877 | 0.689 | 1.5E-30 |
| Nme1 | 8.15E-35 | 1.262861 | 0.437 | 0.201 | 1.92E-30 |
| H2-K1 | 1.58E-34 | 0.899597 | 0.948 | 0.822 | 3.73E-30 |
| Gnas | 1.63E-34 | 1.11575 | 0.859 | 0.705 | 3.84E-30 |
| Bola2 | 1.91E-34 | 0.482864 | 0.41 | 0.174 | 4.5E-30 |

| | | | | | |
|---|---|---|---|---|---|
| Bcl2a1d | 2.15E-34 | 2.1473 | 0.352 | 0.151 | 5.06E-30 |
| Gm7336 | 2.3E-34 | 0.984857 | 0.574 | 0.324 | 5.43E-30 |
| Bcl2a1b | 7.35E-34 | 0.889603 | 0.738 | 0.462 | 1.73E-29 |
| Nceh1 | 9.72E-34 | 1.71825 | 0.238 | 0.079 | 2.29E-29 |
| Selenow | 1.16E-33 | 1.193417 | 0.415 | 0.187 | 2.73E-29 |
| Slamf8 | 6.54E-33 | 1.75311 | 0.294 | 0.111 | 1.54E-28 |
| H2-D1 | 8.63E-33 | 0.844049 | 0.975 | 0.909 | 2.03E-28 |
| Pcna | 1.88E-32 | 1.060404 | 0.446 | 0.211 | 4.44E-28 |
| Eno1 | 3.53E-32 | 0.952042 | 0.872 | 0.692 | 8.32E-28 |
| Slc11a2 | 4.99E-32 | 1.805853 | 0.24 | 0.081 | 1.18E-27 |
| Rap2b | 1.47E-31 | 1.413887 | 0.278 | 0.103 | 3.46E-27 |
| Lgals1 | 4.86E-31 | 2.770859 | 0.211 | 0.07 | 1.15E-26 |
| Rpsa-ps10 | 1.3E-30 | 0.865511 | 0.796 | 0.561 | 3.08E-26 |
| Hnrnpa1 | 4.86E-30 | 1.377057 | 0.507 | 0.281 | 1.15E-25 |
| Atp5g1 | 5.11E-29 | 1.067528 | 0.583 | 0.34 | 1.21E-24 |
| Slc15a3 | 1.07E-28 | 1.563687 | 0.592 | 0.367 | 2.52E-24 |
| Gm22774 | 1.99E-28 | 0.43816 | 0.285 | 0.114 | 4.7E-24 |
| Ncl | 2.75E-28 | 1.070962 | 0.776 | 0.557 | 6.48E-24 |
| Tpm4 | 6.13E-28 | 1.897947 | 0.312 | 0.137 | 1.45E-23 |
| Gm10359 | 9.48E-28 | 0.692689 | 0.52 | 0.302 | 2.24E-23 |
| Mir692-1 | 1.07E-27 | 0.340245 | 0.249 | 0.093 | 2.52E-23 |
| Dpp7 | 1.43E-27 | 1.097484 | 0.336 | 0.148 | 3.37E-23 |
| Ccl9 | 1.65E-27 | 1.915833 | 0.556 | 0.355 | 3.9E-23 |
| Ly9 | 2.98E-27 | 1.434667 | 0.289 | 0.119 | 7.02E-23 |
| Pdgfa | 3.42E-27 | 1.123482 | 0.307 | 0.129 | 8.06E-23 |
| Gm47428 | 4.26E-27 | 0.402341 | 0.226 | 0.082 | 1E-22 |
| Atox1 | 4.59E-27 | 0.829694 | 0.731 | 0.484 | 1.08E-22 |
| Nme2 | 7.34E-27 | 0.799625 | 0.59 | 0.342 | 1.73E-22 |
| Ttyh2 | 7.89E-27 | 1.80992 | 0.209 | 0.073 | 1.86E-22 |
| Hsp90aa1 | 1.05E-26 | 1.063777 | 0.724 | 0.517 | 2.47E-22 |
| Dtnbp1 | 1.66E-26 | 1.030912 | 0.413 | 0.205 | 3.91E-22 |
| Pgam1 | 1.74E-26 | 1.05634 | 0.525 | 0.297 | 4.1E-22 |
| Eif4a1 | 2.16E-26 | 1.194787 | 0.776 | 0.594 | 5.1E-22 |
| Npm1 | 2.17E-26 | 1.005185 | 0.744 | 0.527 | 5.12E-22 |
| Gpi1 | 4.39E-26 | 1.044082 | 0.561 | 0.337 | 1.04E-21 |
| Csf2ra | 4.93E-26 | 1.203937 | 0.502 | 0.295 | 1.16E-21 |
| Rps2-ps10 | 6.35E-26 | 0.711572 | 0.478 | 0.262 | 1.5E-21 |
| Cd9-ps | 1.26E-25 | 0.357635 | 0.26 | 0.103 | 2.97E-21 |
| Kcnj2 | 1.49E-25 | 1.440154 | 0.22 | 0.08 | 3.52E-21 |
| Gm12164 | 1.63E-25 | 0.345614 | 0.316 | 0.143 | 3.86E-21 |
| Iqgap1 | 1.77E-25 | 2.286258 | 0.242 | 0.097 | 4.17E-21 |
| Gm12346 | 2.67E-25 | 0.708968 | 0.478 | 0.268 | 6.3E-21 |
| Myl6 | 3.19E-25 | 0.789868 | 0.803 | 0.6 | 7.54E-21 |
| Renbp | 3.21E-25 | 0.981974 | 0.536 | 0.312 | 7.57E-21 |
| Gm12537 | 3.53E-25 | 0.767036 | 0.5 | 0.292 | 8.32E-21 |
| Eno1b | 3.94E-25 | 0.878045 | 0.742 | 0.522 | 9.28E-21 |
| Rplp0 | 4.5E-25 | 0.814142 | 0.962 | 0.895 | 1.06E-20 |

| | | | | | |
|---|---|---|---|---|---|
| Plekhm2 | 4.96E-25 | 1.789517 | 0.258 | 0.106 | 1.17E-20 |
| Atf3 | 5.82E-25 | 3.230463 | 0.283 | 0.128 | 1.37E-20 |
| Cyba | 7.01E-25 | 0.628535 | 0.991 | 0.97 | 1.65E-20 |
| Gm3699 | 8.74E-25 | 0.740533 | 0.572 | 0.355 | 2.06E-20 |
| Sdc4 | 8.89E-25 | 1.77502 | 0.359 | 0.182 | 2.1E-20 |
| Gm49369 | 1.18E-24 | 0.888949 | 0.85 | 0.769 | 2.77E-20 |
| Scpep1 | 1.5E-24 | 0.851466 | 0.605 | 0.377 | 3.54E-20 |
| Dgkz | 1.61E-24 | 1.273277 | 0.475 | 0.262 | 3.79E-20 |
| Itga5 | 1.79E-24 | 2.041061 | 0.222 | 0.085 | 4.23E-20 |
| Rpl41 | 7.85E-24 | 0.683526 | 0.978 | 0.943 | 1.85E-19 |
| Uap1l1 | 2.27E-23 | 1.243167 | 0.357 | 0.176 | 5.35E-19 |
| Lgals3bp | 2.74E-23 | 0.972737 | 0.771 | 0.584 | 6.46E-19 |
| Rack1 | 6.18E-23 | 0.866671 | 0.854 | 0.704 | 1.46E-18 |
| S1pr1 | 8.74E-23 | 0.75219 | 0.496 | 0.289 | 2.06E-18 |
| Ctsa | 8.89E-23 | 0.542662 | 0.989 | 0.963 | 2.1E-18 |
| B2m | 9.58E-23 | 0.54089 | 1 | 0.998 | 2.26E-18 |
| Plek | 1E-22 | 1.223217 | 0.702 | 0.508 | 2.36E-18 |
| Gm4609 | 1.44E-22 | 0.753566 | 0.504 | 0.301 | 3.4E-18 |
| Crip1 | 2.8E-22 | 2.162364 | 0.233 | 0.099 | 6.6E-18 |
| Got1 | 2.84E-22 | 1.268629 | 0.24 | 0.098 | 6.71E-18 |
| Anp32b | 3.1E-22 | 1.143726 | 0.435 | 0.237 | 7.32E-18 |
| Gm13456 | 3.66E-22 | 0.673876 | 0.978 | 0.946 | 8.63E-18 |
| Ddx39 | 4.18E-22 | 1.534213 | 0.222 | 0.088 | 9.85E-18 |
| Gm8730 | 4.96E-22 | 0.753804 | 0.87 | 0.737 | 1.17E-17 |
| Gm3839 | 4.99E-22 | 0.351181 | 0.325 | 0.157 | 1.18E-17 |
| Eef1a1 | 5.05E-22 | 0.666923 | 0.998 | 0.993 | 1.19E-17 |
| Hspe1 | 6.31E-22 | 0.860363 | 0.605 | 0.388 | 1.49E-17 |
| Plekho2 | 7.17E-22 | 1.613992 | 0.296 | 0.138 | 1.69E-17 |
| Eif3a | 1.15E-21 | 1.362163 | 0.478 | 0.289 | 2.71E-17 |
| Cotl1 | 1.44E-21 | 0.771893 | 0.895 | 0.774 | 3.39E-17 |
| Fblim1 | 1.55E-21 | 1.369923 | 0.249 | 0.105 | 3.65E-17 |
| Slc2a1 | 1.84E-21 | 1.417601 | 0.381 | 0.205 | 4.35E-17 |
| Ndufa1 | 2.42E-21 | 0.779039 | 0.48 | 0.276 | 5.72E-17 |
| Cdkn1a | 3.24E-21 | 2.006339 | 0.262 | 0.122 | 7.64E-17 |
| Tyrobp | 4.07E-21 | 0.546053 | 0.998 | 0.995 | 9.59E-17 |
| Gm5138 | 4.1E-21 | 0.962424 | 0.626 | 0.442 | 9.68E-17 |
| Gm21399 | 5.56E-21 | 0.367871 | 0.343 | 0.173 | 1.31E-16 |
| Gm4735 | 7.89E-21 | 0.817791 | 0.563 | 0.362 | 1.86E-16 |
| Gm11808 | 9.68E-21 | 0.670324 | 0.691 | 0.498 | 2.28E-16 |
| Eif4a-ps4 | 1.2E-20 | 0.913997 | 0.574 | 0.381 | 2.83E-16 |
| Eef1g | 1.6E-20 | 0.928613 | 0.592 | 0.393 | 3.78E-16 |
| Rpl10a | 1.71E-20 | 0.564034 | 0.85 | 0.667 | 4.03E-16 |
| Anxa4 | 2.8E-20 | 1.270081 | 0.267 | 0.12 | 6.59E-16 |
| Pebp1 | 4.07E-20 | 0.88079 | 0.489 | 0.297 | 9.6E-16 |
| AI506816 | 4.14E-20 | 1.423031 | 0.419 | 0.256 | 9.76E-16 |
| Rps20 | 4.15E-20 | 0.639974 | 0.93 | 0.826 | 9.79E-16 |
| Ptma | 4.54E-20 | 0.682911 | 0.791 | 0.627 | 1.07E-15 |

| | | | | | |
|---|---|---|---|---|---|
| Uba52 | 6.19E-20 | 0.613004 | 0.886 | 0.746 | 1.46E-15 |
| Gm9118 | 6.83E-20 | 0.652804 | 0.35 | 0.179 | 1.61E-15 |
| Gm4691 | 7.58E-20 | 0.355676 | 0.222 | 0.093 | 1.79E-15 |
| Rps5 | 7.65E-20 | 0.779704 | 0.865 | 0.736 | 1.81E-15 |
| Scd2 | 9.63E-20 | 0.8577 | 0.28 | 0.13 | 2.27E-15 |
| Pcna-ps2 | 1.08E-19 | 0.567643 | 0.26 | 0.117 | 2.54E-15 |
| Sec61g | 1.28E-19 | 0.53652 | 0.596 | 0.375 | 3.02E-15 |
| Atp6v1a | 1.41E-19 | 1.360414 | 0.372 | 0.198 | 3.34E-15 |
| Gm10076 | 1.46E-19 | 0.704227 | 0.96 | 0.905 | 3.45E-15 |
| Gusb | 1.54E-19 | 0.755367 | 0.771 | 0.604 | 3.63E-15 |
| Gm29667 | 1.57E-19 | 0.627618 | 0.489 | 0.3 | 3.71E-15 |
| Sdc3 | 2.02E-19 | 0.938827 | 0.318 | 0.158 | 4.77E-15 |
| Rpl3 | 2.41E-19 | 0.588084 | 0.942 | 0.828 | 5.7E-15 |
| Emp3 | 2.54E-19 | 1.059684 | 0.33 | 0.169 | 6E-15 |
| Tpd52 | 2.57E-19 | 0.796724 | 0.561 | 0.364 | 6.07E-15 |
| Gaa | 4.31E-19 | 0.892662 | 0.534 | 0.347 | 1.02E-14 |
| Rps27l | 4.34E-19 | 0.711321 | 0.543 | 0.34 | 1.02E-14 |
| Uqcr11 | 5.69E-19 | 0.732555 | 0.594 | 0.392 | 1.34E-14 |
| Gm2000 | 6.37E-19 | 0.708314 | 0.765 | 0.591 | 1.5E-14 |
| Siglecf | 6.73E-19 | 1.476675 | 0.217 | 0.092 | 1.59E-14 |
| Ccl6 | 1.17E-18 | 1.364902 | 0.599 | 0.42 | 2.75E-14 |
| Eprs | 1.19E-18 | 0.819276 | 0.419 | 0.24 | 2.8E-14 |
| Rpl38 | 1.33E-18 | 0.69037 | 0.809 | 0.641 | 3.15E-14 |
| Commd4 | 1.71E-18 | 0.78556 | 0.361 | 0.192 | 4.02E-14 |
| Rpl3-ps1 | 1.83E-18 | 0.688846 | 0.857 | 0.701 | 4.32E-14 |
| Fam162a | 2.49E-18 | 1.138039 | 0.242 | 0.109 | 5.88E-14 |
| P2rx4 | 2.51E-18 | 0.814038 | 0.587 | 0.403 | 5.93E-14 |
| Bax | 3.02E-18 | 0.864413 | 0.439 | 0.26 | 7.13E-14 |
| Myo9b | 4.28E-18 | 0.69811 | 0.392 | 0.217 | 1.01E-13 |
| Gm4366 | 4.84E-18 | 0.680266 | 0.43 | 0.251 | 1.14E-13 |
| Rps8 | 4.98E-18 | 0.601363 | 0.922 | 0.825 | 1.18E-13 |
| Rpl5 | 5.23E-18 | 0.664488 | 0.877 | 0.78 | 1.23E-13 |
| Ranbp1 | 5.37E-18 | 0.835471 | 0.446 | 0.263 | 1.27E-13 |
| Gnl3 | 6.86E-18 | 1.187157 | 0.28 | 0.137 | 1.62E-13 |
| Gm9625 | 7.43E-18 | 0.75818 | 0.684 | 0.501 | 1.75E-13 |
| Cndp2 | 7.53E-18 | 0.755446 | 0.652 | 0.455 | 1.78E-13 |
| Bcl2a1a | 8.49E-18 | 0.830346 | 0.325 | 0.168 | 2E-13 |
| Pgam1-ps2 | 9.32E-18 | 0.3836 | 0.309 | 0.158 | 2.2E-13 |
| H2-Q7 | 1.07E-17 | 1.386992 | 0.247 | 0.116 | 2.52E-13 |
| Atp5k | 1.11E-17 | 0.436562 | 0.466 | 0.273 | 2.62E-13 |
| Hmgn2 | 1.19E-17 | 0.591014 | 0.556 | 0.35 | 2.81E-13 |
| AU020206 | 1.75E-17 | 0.698617 | 0.715 | 0.531 | 4.13E-13 |
| Rpl35 | 2.45E-17 | 0.531862 | 0.93 | 0.819 | 5.78E-13 |
| Rpl32 | 2.56E-17 | 0.513926 | 0.948 | 0.881 | 6.04E-13 |
| Atp6v1b2 | 2.74E-17 | 0.850859 | 0.498 | 0.31 | 6.46E-13 |
| Ybx1 | 3E-17 | 0.648723 | 0.76 | 0.579 | 7.07E-13 |
| Slc25a5 | 3.13E-17 | 0.608438 | 0.89 | 0.787 | 7.37E-13 |

| | | | | | |
|---|---|---|---|---|---|
| Rpl12 | 3.4E-17 | 0.687342 | 0.814 | 0.678 | 8.03E-13 |
| Npc2 | 3.87E-17 | 0.612932 | 0.944 | 0.837 | 9.13E-13 |
| Bst2 | 4.04E-17 | 0.67853 | 0.612 | 0.414 | 9.53E-13 |
| Mien1 | 4.51E-17 | 0.75756 | 0.332 | 0.174 | 1.06E-12 |
| Lipa | 5.64E-17 | 1.057635 | 0.576 | 0.409 | 1.33E-12 |
| Hsd17b12 | 5.89E-17 | 0.621651 | 0.482 | 0.301 | 1.39E-12 |
| C5ar1 | 7.43E-17 | 1.218711 | 0.576 | 0.408 | 1.75E-12 |
| Gm5526 | 1.01E-16 | 0.439958 | 0.513 | 0.329 | 2.39E-12 |
| Srm | 1.09E-16 | 0.986987 | 0.365 | 0.207 | 2.57E-12 |
| Cfl1 | 1.2E-16 | 0.624211 | 0.973 | 0.96 | 2.84E-12 |
| Rps26 | 1.23E-16 | 0.555128 | 0.859 | 0.691 | 2.9E-12 |
| Rps19 | 1.36E-16 | 0.562922 | 0.937 | 0.874 | 3.22E-12 |
| Gm5844 | 1.85E-16 | 0.80765 | 0.379 | 0.218 | 4.37E-12 |
| Gm10039 | 1.91E-16 | 0.484927 | 0.386 | 0.214 | 4.5E-12 |
| Gm6477 | 2.26E-16 | 0.668985 | 0.26 | 0.127 | 5.33E-12 |
| Hint1 | 3.6E-16 | 0.696163 | 0.574 | 0.386 | 8.5E-12 |
| H2-Q6 | 4.22E-16 | 1.367917 | 0.22 | 0.102 | 9.96E-12 |
| Cd84 | 4.36E-16 | 0.751967 | 0.818 | 0.696 | 1.03E-11 |
| Aprt | 4.77E-16 | 1.056091 | 0.305 | 0.163 | 1.12E-11 |
| Cox8a | 5.11E-16 | 0.590355 | 0.776 | 0.609 | 1.21E-11 |
| Alas1 | 5.58E-16 | 0.968164 | 0.386 | 0.227 | 1.32E-11 |
| Tagln2 | 5.75E-16 | 0.885621 | 0.399 | 0.24 | 1.36E-11 |
| Slc11a1 | 6.32E-16 | 0.727179 | 0.765 | 0.6 | 1.49E-11 |
| Ran | 7.25E-16 | 0.83993 | 0.543 | 0.368 | 1.71E-11 |
| Akr1a1 | 7.32E-16 | 0.666293 | 0.868 | 0.73 | 1.73E-11 |
| Pabpc1 | 7.34E-16 | 0.59406 | 0.816 | 0.659 | 1.73E-11 |
| Hspa9 | 9.82E-16 | 1.105037 | 0.388 | 0.23 | 2.32E-11 |
| Cycs | 1.07E-15 | 0.772679 | 0.464 | 0.283 | 2.52E-11 |
| Pfkl | 1.17E-15 | 0.798311 | 0.401 | 0.239 | 2.76E-11 |
| Rpl18 | 1.2E-15 | 0.547024 | 0.908 | 0.801 | 2.83E-11 |
| Slc6a6 | 1.22E-15 | 0.663213 | 0.554 | 0.371 | 2.89E-11 |
| Rps18 | 1.25E-15 | 0.557757 | 0.942 | 0.851 | 2.94E-11 |
| Ifi27l2a | 1.27E-15 | 0.508949 | 0.305 | 0.159 | 2.99E-11 |
| Txn1 | 1.32E-15 | 0.718102 | 0.408 | 0.243 | 3.12E-11 |
| Atp6v1c1 | 1.32E-15 | 0.760754 | 0.525 | 0.342 | 3.12E-11 |
| Flna | 1.37E-15 | 0.501766 | 0.249 | 0.12 | 3.24E-11 |
| Gm10180 | 1.38E-15 | 0.374077 | 0.583 | 0.419 | 3.26E-11 |
| Rps28 | 1.42E-15 | 0.498439 | 0.978 | 0.908 | 3.36E-11 |
| Mybbp1a | 1.64E-15 | 0.782949 | 0.321 | 0.175 | 3.88E-11 |
| Calm1 | 1.8E-15 | 0.616214 | 0.863 | 0.733 | 4.25E-11 |
| Grn | 2.06E-15 | 0.418351 | 1 | 0.993 | 4.87E-11 |
| Amdhd2 | 2.2E-15 | 1.373648 | 0.238 | 0.117 | 5.2E-11 |
| Elob | 2.48E-15 | 0.581725 | 0.652 | 0.466 | 5.84E-11 |
| Gng12 | 3.51E-15 | 1.281035 | 0.274 | 0.144 | 8.28E-11 |
| Clcn7 | 3.68E-15 | 0.654851 | 0.502 | 0.322 | 8.68E-11 |
| Tceal9 | 5.35E-15 | 1.251094 | 0.22 | 0.105 | 1.26E-10 |
| Gm15710 | 5.42E-15 | 0.4645 | 0.863 | 0.743 | 1.28E-10 |

| | | | | | |
|---|---|---|---|---|---|
| Sars | 5.42E-15 | 0.790795 | 0.417 | 0.256 | 1.28E-10 |
| Psen2 | 6.2E-15 | 1.29276 | 0.238 | 0.117 | 1.46E-10 |
| Rpl14 | 6.41E-15 | 0.541671 | 0.895 | 0.774 | 1.51E-10 |
| Smpdl3a | 6.66E-15 | 1.259043 | 0.224 | 0.107 | 1.57E-10 |
| Gm5915 | 7.13E-15 | 0.403645 | 0.354 | 0.204 | 1.68E-10 |
| Creld2 | 7.58E-15 | 0.923439 | 0.41 | 0.249 | 1.79E-10 |
| Soat1 | 7.66E-15 | 0.91101 | 0.48 | 0.312 | 1.81E-10 |
| Dnase2a | 8.98E-15 | 0.540295 | 0.632 | 0.453 | 2.12E-10 |
| Rpl23 | 9.12E-15 | 0.557113 | 0.933 | 0.832 | 2.15E-10 |
| Napa | 9.91E-15 | 0.89996 | 0.453 | 0.292 | 2.34E-10 |
| Rpl10a-ps1 | 9.92E-15 | 0.468467 | 0.72 | 0.554 | 2.34E-10 |
| Gm10224 | 1.38E-14 | 0.383471 | 0.318 | 0.176 | 3.26E-10 |
| Lamp1 | 1.57E-14 | 0.4539 | 0.98 | 0.955 | 3.71E-10 |
| Ctsl | 1.7E-14 | 0.613832 | 0.982 | 0.949 | 4.01E-10 |
| Colgalt1 | 1.87E-14 | 0.609624 | 0.451 | 0.284 | 4.42E-10 |
| Gba | 2.24E-14 | 0.684585 | 0.572 | 0.407 | 5.3E-10 |
| Grina | 2.37E-14 | 0.522701 | 0.722 | 0.559 | 5.59E-10 |
| Anxa2 | 2.42E-14 | 2.782998 | 0.222 | 0.113 | 5.7E-10 |
| Gm1840 | 2.43E-14 | 0.395684 | 0.287 | 0.155 | 5.73E-10 |
| Actr3 | 2.6E-14 | 0.725637 | 0.661 | 0.512 | 6.12E-10 |
| Naa50 | 2.64E-14 | 1.319128 | 0.233 | 0.116 | 6.24E-10 |
| Mitf | 2.97E-14 | 0.672624 | 0.413 | 0.254 | 7.01E-10 |
| Rps27a | 3.16E-14 | 0.583613 | 0.906 | 0.794 | 7.45E-10 |
| Creg1 | 3.88E-14 | 0.518846 | 0.834 | 0.683 | 9.14E-10 |
| mt-Tp | 3.94E-14 | 0.955918 | 0.307 | 0.172 | 9.3E-10 |
| Atp1a1 | 4.12E-14 | 0.565339 | 0.623 | 0.452 | 9.71E-10 |
| Tmem256 | 4.27E-14 | 0.429413 | 0.444 | 0.273 | 1.01E-09 |
| Tomm20 | 4.98E-14 | 0.717528 | 0.498 | 0.326 | 1.18E-09 |
| Dusp3 | 6.35E-14 | 0.887133 | 0.518 | 0.358 | 1.5E-09 |
| Gm10275 | 7.02E-14 | 0.621991 | 0.572 | 0.407 | 1.66E-09 |
| Rplp2 | 7.21E-14 | 0.401662 | 0.955 | 0.902 | 1.7E-09 |
| Hspd1 | 7.38E-14 | 0.817085 | 0.37 | 0.223 | 1.74E-09 |
| Gm9844 | 7.4E-14 | 0.303841 | 0.231 | 0.113 | 1.75E-09 |
| Eif5a | 7.42E-14 | 0.674784 | 0.742 | 0.582 | 1.75E-09 |
| Banf1 | 7.85E-14 | 0.627015 | 0.401 | 0.242 | 1.85E-09 |
| Txndc17 | 7.91E-14 | 0.577405 | 0.397 | 0.239 | 1.87E-09 |
| Gm2606 | 7.95E-14 | 0.251698 | 0.28 | 0.15 | 1.87E-09 |
| Gm11539 | 8.23E-14 | 0.447613 | 0.46 | 0.301 | 1.94E-09 |
| C1qbp | 8.44E-14 | 0.636065 | 0.361 | 0.211 | 1.99E-09 |
| Rbbp7 | 9.75E-14 | 0.724817 | 0.417 | 0.262 | 2.3E-09 |
| Rps13 | 1.07E-13 | 0.434056 | 0.899 | 0.779 | 2.52E-09 |
| Eif5 | 1.38E-13 | 0.61593 | 0.574 | 0.402 | 3.25E-09 |
| Lrpap1 | 1.39E-13 | 0.86622 | 0.581 | 0.424 | 3.29E-09 |
| Rpl14-ps1 | 1.62E-13 | 0.507646 | 0.744 | 0.584 | 3.81E-09 |
| Ssr4 | 1.73E-13 | 0.554918 | 0.776 | 0.623 | 4.08E-09 |
| Rpl19 | 1.82E-13 | 0.49022 | 0.913 | 0.808 | 4.28E-09 |
| Rps25 | 1.86E-13 | 0.531724 | 0.955 | 0.858 | 4.38E-09 |

| | | | | | |
|---|---|---|---|---|---|
| Hspa8 | 1.93E-13 | 0.519342 | 0.948 | 0.927 | 4.54E-09 |
| Rps15 | 1.97E-13 | 0.478768 | 0.877 | 0.752 | 4.66E-09 |
| Glmp | 2.09E-13 | 0.484311 | 0.702 | 0.535 | 4.93E-09 |
| Polr2l | 2.43E-13 | 0.463649 | 0.291 | 0.157 | 5.74E-09 |
| Rpl13 | 2.66E-13 | 0.451271 | 0.975 | 0.943 | 6.27E-09 |
| Cox5a | 2.66E-13 | 0.584093 | 0.453 | 0.285 | 6.28E-09 |
| Txn2 | 3.09E-13 | 0.694429 | 0.372 | 0.226 | 7.28E-09 |
| Cln5 | 3.17E-13 | 0.636842 | 0.406 | 0.247 | 7.49E-09 |
| Cox6c | 3.21E-13 | 0.620287 | 0.621 | 0.446 | 7.58E-09 |
| Rps16-ps2 | 3.24E-13 | 0.630648 | 0.715 | 0.584 | 7.63E-09 |
| Olfr77 | 3.55E-13 | 0.425853 | 0.489 | 0.326 | 8.39E-09 |
| Ppia | 3.96E-13 | 0.428027 | 0.982 | 0.966 | 9.35E-09 |
| Timm50 | 4.01E-13 | 0.754565 | 0.303 | 0.17 | 9.47E-09 |
| Rpl17 | 4.28E-13 | 0.501257 | 0.872 | 0.793 | 1.01E-08 |
| Atp6v1g1 | 4.3E-13 | 0.484622 | 0.776 | 0.589 | 1.01E-08 |
| Atp5e | 4.96E-13 | 0.647793 | 0.657 | 0.479 | 1.17E-08 |
| Prune2 | 5.12E-13 | 0.546676 | 0.413 | 0.256 | 1.21E-08 |
| Vim | 5.28E-13 | 2.764471 | 0.209 | 0.107 | 1.25E-08 |
| Nop56 | 5.43E-13 | 0.796006 | 0.345 | 0.205 | 1.28E-08 |
| Mettl1 | 5.44E-13 | 0.966864 | 0.211 | 0.103 | 1.28E-08 |
| Myl12a | 5.52E-13 | 0.64632 | 0.648 | 0.474 | 1.3E-08 |
| Sdf2l1 | 5.79E-13 | 0.704106 | 0.599 | 0.436 | 1.37E-08 |
| Gm7331 | 5.99E-13 | 0.340574 | 0.704 | 0.56 | 1.41E-08 |
| Morf4l2 | 6.28E-13 | 0.80429 | 0.43 | 0.277 | 1.48E-08 |
| Rplp1 | 6.36E-13 | 0.385 | 0.982 | 0.956 | 1.5E-08 |
| Ifrd1 | 6.89E-13 | 1.393445 | 0.231 | 0.12 | 1.62E-08 |
| Uqcrq | 7.54E-13 | 0.565369 | 0.518 | 0.364 | 1.78E-08 |
| Oxct1 | 7.82E-13 | 0.640063 | 0.63 | 0.477 | 1.84E-08 |
| Gm11836 | 7.94E-13 | 0.258741 | 0.211 | 0.104 | 1.87E-08 |
| Gpx1 | 8.21E-13 | 0.477105 | 0.85 | 0.704 | 1.94E-08 |
| Rpl15 | 8.51E-13 | 0.605035 | 0.807 | 0.684 | 2.01E-08 |
| Tmsb10 | 9.39E-13 | 0.60802 | 0.592 | 0.431 | 2.21E-08 |
| Gm5529 | 1.02E-12 | 0.421896 | 0.471 | 0.328 | 2.41E-08 |
| Eif2s1 | 1.02E-12 | 0.858641 | 0.368 | 0.226 | 2.41E-08 |
| Ctdnep1 | 1.08E-12 | 1.219432 | 0.296 | 0.174 | 2.55E-08 |
| Nhp2 | 1.28E-12 | 0.967081 | 0.231 | 0.12 | 3.03E-08 |
| Tmem160 | 1.33E-12 | 0.298352 | 0.496 | 0.321 | 3.13E-08 |
| Cops9 | 1.38E-12 | 0.37006 | 0.567 | 0.396 | 3.25E-08 |
| Nap1l1 | 1.42E-12 | 0.74691 | 0.274 | 0.152 | 3.35E-08 |
| Degs1 | 1.48E-12 | 0.735429 | 0.439 | 0.288 | 3.5E-08 |
| Timp2 | 1.84E-12 | 0.504308 | 0.913 | 0.83 | 4.33E-08 |
| Rps16 | 1.84E-12 | 0.482605 | 0.872 | 0.776 | 4.34E-08 |
| Tomm70a | 1.85E-12 | 0.646038 | 0.307 | 0.176 | 4.38E-08 |
| Eef1b2 | 1.95E-12 | 0.562468 | 0.61 | 0.457 | 4.61E-08 |
| Higd1a | 2.2E-12 | 0.646305 | 0.291 | 0.163 | 5.2E-08 |
| Ufm1 | 2.34E-12 | 0.9202 | 0.229 | 0.12 | 5.51E-08 |
| Rpl9 | 2.35E-12 | 0.484122 | 0.874 | 0.761 | 5.54E-08 |

| | | | | | |
|---|---|---|---|---|---|
| Gm4149 | 2.42E-12 | 0.335063 | 0.917 | 0.833 | 5.7E-08 |
| Asph | 2.62E-12 | 0.57569 | 0.767 | 0.621 | 6.18E-08 |
| Spg21 | 2.63E-12 | 0.814361 | 0.37 | 0.228 | 6.2E-08 |
| Prmt1 | 2.67E-12 | 0.648535 | 0.318 | 0.189 | 6.29E-08 |
| Galnt7 | 2.69E-12 | 1.173082 | 0.217 | 0.11 | 6.34E-08 |
| Gm15500 | 2.69E-12 | 0.808286 | 0.74 | 0.605 | 6.35E-08 |
| P4hb | 2.86E-12 | 0.525045 | 0.901 | 0.803 | 6.76E-08 |
| Bri3bp | 3.01E-12 | 1.194767 | 0.202 | 0.101 | 7.1E-08 |
| Lat2 | 3.14E-12 | 0.589367 | 0.536 | 0.383 | 7.4E-08 |
| Slc25a3 | 3.85E-12 | 0.531335 | 0.697 | 0.548 | 9.08E-08 |
| Hspd1-ps3 | 4.17E-12 | 0.504565 | 0.213 | 0.109 | 9.83E-08 |
| Eif1a | 4.56E-12 | 0.96265 | 0.296 | 0.171 | 1.07E-07 |
| Plin2 | 4.77E-12 | 1.088918 | 0.22 | 0.115 | 1.13E-07 |
| Ctsc | 4.81E-12 | 0.48951 | 0.91 | 0.847 | 1.13E-07 |
| Tcp1 | 4.88E-12 | 0.466647 | 0.408 | 0.256 | 1.15E-07 |
| Atp5b | 4.92E-12 | 0.517576 | 0.836 | 0.739 | 1.16E-07 |
| Rpl4 | 4.95E-12 | 0.446248 | 0.915 | 0.832 | 1.17E-07 |
| Hspa5 | 6.04E-12 | 0.739596 | 0.888 | 0.829 | 1.43E-07 |
| Hnrnpab | 6.24E-12 | 0.57319 | 0.679 | 0.526 | 1.47E-07 |
| Cox5b | 6.5E-12 | 0.499709 | 0.543 | 0.391 | 1.53E-07 |
| Fam221b | 6.8E-12 | 0.515599 | 0.583 | 0.425 | 1.6E-07 |
| Tent5c | 6.97E-12 | 0.728398 | 0.511 | 0.37 | 1.64E-07 |
| Tgif1 | 6.99E-12 | 0.790041 | 0.321 | 0.192 | 1.65E-07 |
| Atp6v1f | 7.79E-12 | 0.456793 | 0.693 | 0.514 | 1.84E-07 |
| Eif4ebp1 | 7.85E-12 | 0.822204 | 0.287 | 0.163 | 1.85E-07 |
| Uqcc2 | 8.11E-12 | 0.642561 | 0.368 | 0.226 | 1.91E-07 |
| Rps10 | 8.3E-12 | 0.439035 | 0.922 | 0.799 | 1.96E-07 |
| Gtf2f1 | 9.53E-12 | 0.604008 | 0.24 | 0.128 | 2.25E-07 |
| Txndc5 | 1.04E-11 | 1.07775 | 0.314 | 0.193 | 2.45E-07 |
| Sqstm1 | 1.11E-11 | 0.810534 | 0.711 | 0.58 | 2.61E-07 |
| Syngr1 | 1.11E-11 | 0.540746 | 0.901 | 0.815 | 2.62E-07 |
| Ifitm3 | 1.14E-11 | 0.582101 | 0.318 | 0.194 | 2.69E-07 |
| Phgdh | 1.16E-11 | 0.762358 | 0.507 | 0.36 | 2.73E-07 |
| Psmd1 | 1.18E-11 | 0.854504 | 0.428 | 0.284 | 2.77E-07 |
| Ube2l3 | 1.25E-11 | 0.450503 | 0.561 | 0.39 | 2.95E-07 |
| Tmbim1 | 1.27E-11 | 0.834338 | 0.318 | 0.191 | 3E-07 |
| Il11ra1 | 1.31E-11 | 0.88995 | 0.296 | 0.173 | 3.09E-07 |
| Nutf2 | 1.35E-11 | 0.551226 | 0.372 | 0.227 | 3.17E-07 |
| Atp5mpl | 1.42E-11 | 0.52908 | 0.419 | 0.272 | 3.35E-07 |
| Vegfb | 1.42E-11 | 0.641373 | 0.404 | 0.264 | 3.35E-07 |
| Rps15a | 1.47E-11 | 0.47243 | 0.738 | 0.613 | 3.46E-07 |
| St13 | 1.51E-11 | 0.382851 | 0.78 | 0.616 | 3.56E-07 |
| Arpc1b | 1.51E-11 | 0.500856 | 0.87 | 0.765 | 3.56E-07 |
| Gpr84 | 1.55E-11 | 1.055284 | 0.567 | 0.438 | 3.65E-07 |
| Cox6a1 | 2.04E-11 | 0.487548 | 0.697 | 0.551 | 4.82E-07 |
| Atp5l | 2.05E-11 | 0.464502 | 0.637 | 0.471 | 4.83E-07 |
| Tlr1 | 2.05E-11 | 0.62603 | 0.291 | 0.167 | 4.84E-07 |

| | | | | | |
|---|---|---|---|---|---|
| Cox7b | 2.06E-11 | 0.610231 | 0.442 | 0.293 | 4.85E-07 |
| Elp5 | 2.13E-11 | 0.774465 | 0.22 | 0.114 | 5.03E-07 |
| Edf1 | 2.55E-11 | 0.571425 | 0.578 | 0.421 | 6.03E-07 |
| Wdr43 | 2.63E-11 | 0.642203 | 0.291 | 0.17 | 6.2E-07 |
| Elovl1 | 2.64E-11 | 0.49726 | 0.511 | 0.355 | 6.22E-07 |
| Psmb6 | 2.77E-11 | 0.60399 | 0.587 | 0.438 | 6.53E-07 |
| Rpl6 | 2.79E-11 | 0.475556 | 0.861 | 0.761 | 6.58E-07 |
| Arl5c | 3.22E-11 | 1.643663 | 0.211 | 0.111 | 7.6E-07 |
| 1810022K0 | 3.55E-11 | 0.430091 | 0.379 | 0.232 | 8.37E-07 |
| Sdcbp | 3.85E-11 | 0.575162 | 0.814 | 0.69 | 9.09E-07 |
| Rps17 | 3.86E-11 | 0.401215 | 0.861 | 0.733 | 9.11E-07 |
| Rpl28 | 4.04E-11 | 0.454599 | 0.845 | 0.733 | 9.53E-07 |
| Rps7 | 4.4E-11 | 0.511872 | 0.863 | 0.79 | 1.04E-06 |
| Gm26825 | 4.46E-11 | 0.466267 | 0.807 | 0.7 | 1.05E-06 |
| Rps15a-ps6 | 4.66E-11 | 0.576248 | 0.386 | 0.249 | 1.1E-06 |
| Rpl11 | 5.07E-11 | 0.370515 | 0.877 | 0.724 | 1.2E-06 |
| Gng5 | 5.36E-11 | 0.431248 | 0.796 | 0.658 | 1.26E-06 |
| Usmg5 | 5.43E-11 | 0.468254 | 0.491 | 0.334 | 1.28E-06 |
| Gm5879 | 6.25E-11 | 0.464021 | 0.484 | 0.34 | 1.48E-06 |
| Asna1 | 6.26E-11 | 0.695481 | 0.249 | 0.138 | 1.48E-06 |
| Mfsd1 | 6.56E-11 | 0.597313 | 0.563 | 0.412 | 1.55E-06 |
| Hpf1 | 6.72E-11 | 0.610441 | 0.303 | 0.18 | 1.58E-06 |
| Glb1 | 6.78E-11 | 0.872508 | 0.379 | 0.249 | 1.6E-06 |
| Slirp | 7.13E-11 | 0.323648 | 0.271 | 0.152 | 1.68E-06 |
| Mrpl33 | 7.54E-11 | 0.681911 | 0.305 | 0.185 | 1.78E-06 |
| Sh3bgrl3 | 8.49E-11 | 0.499934 | 0.906 | 0.814 | 2E-06 |
| Gm9794 | 9.83E-11 | 0.576133 | 0.765 | 0.631 | 2.32E-06 |
| Pdia4 | 9.92E-11 | 0.601129 | 0.504 | 0.355 | 2.34E-06 |
| Polr2f | 1E-10 | 0.480505 | 0.283 | 0.164 | 2.36E-06 |
| Sgpl1 | 1.02E-10 | 0.61807 | 0.583 | 0.43 | 2.41E-06 |
| Cox7a2 | 1.04E-10 | 0.404165 | 0.487 | 0.336 | 2.45E-06 |
| Dnajb14 | 1.06E-10 | 0.497565 | 0.341 | 0.212 | 2.5E-06 |
| Gsto1 | 1.21E-10 | 0.749189 | 0.238 | 0.131 | 2.84E-06 |
| Gm8822 | 1.26E-10 | 0.445803 | 0.352 | 0.226 | 2.97E-06 |
| Mapkapk2 | 1.27E-10 | 0.583984 | 0.357 | 0.229 | 2.99E-06 |
| Rpl9-ps6 | 1.28E-10 | 0.469775 | 0.76 | 0.597 | 3.02E-06 |
| Hmgn1 | 1.31E-10 | 0.477779 | 0.466 | 0.324 | 3.09E-06 |
| Atp5j | 1.33E-10 | 0.349749 | 0.668 | 0.498 | 3.14E-06 |
| Coro1c | 1.34E-10 | 0.363602 | 0.363 | 0.225 | 3.16E-06 |
| Srgn | 1.35E-10 | 0.491128 | 0.599 | 0.459 | 3.19E-06 |
| Tbca | 1.42E-10 | 0.651702 | 0.325 | 0.203 | 3.34E-06 |
| Pitpna | 1.43E-10 | 0.737279 | 0.54 | 0.409 | 3.37E-06 |
| Gm11560 | 1.45E-10 | 0.270239 | 0.415 | 0.273 | 3.42E-06 |
| Hacd2 | 1.48E-10 | 0.694876 | 0.354 | 0.226 | 3.48E-06 |
| Agpat5 | 1.53E-10 | 1.361331 | 0.226 | 0.126 | 3.6E-06 |
| Usp50 | 1.53E-10 | 0.401238 | 0.267 | 0.15 | 3.61E-06 |
| Plbd2 | 1.56E-10 | 0.45452 | 0.504 | 0.352 | 3.69E-06 |

| | | | | | |
|---|---|---|---|---|---|
| Ptms | 1.58E-10 | 0.61696 | 0.83 | 0.689 | 3.73E-06 |
| Gm4617 | 1.6E-10 | 0.285217 | 0.41 | 0.274 | 3.78E-06 |
| Arpc2 | 1.68E-10 | 0.505154 | 0.91 | 0.821 | 3.96E-06 |
| Rnf128 | 1.8E-10 | 1.271102 | 0.226 | 0.127 | 4.24E-06 |
| Rpl7a | 1.86E-10 | 0.550478 | 0.8 | 0.688 | 4.4E-06 |
| Rpl31 | 2.01E-10 | 0.383822 | 0.886 | 0.792 | 4.74E-06 |
| Gm14494 | 2.07E-10 | 0.48835 | 0.296 | 0.176 | 4.89E-06 |
| Krtcap2 | 2.08E-10 | 0.442066 | 0.585 | 0.43 | 4.92E-06 |
| Tkt | 2.16E-10 | 0.369109 | 0.496 | 0.345 | 5.09E-06 |
| Cct3 | 2.2E-10 | 0.445972 | 0.43 | 0.289 | 5.18E-06 |
| Rrp1 | 2.32E-10 | 0.473262 | 0.493 | 0.356 | 5.48E-06 |
| Mydgf | 2.63E-10 | 0.31644 | 0.504 | 0.352 | 6.21E-06 |
| Psap | 2.78E-10 | 0.370152 | 1 | 0.998 | 6.57E-06 |
| Psmb2 | 2.88E-10 | 0.433087 | 0.489 | 0.337 | 6.79E-06 |
| Snrpa1 | 3.02E-10 | 0.792092 | 0.222 | 0.122 | 7.11E-06 |
| Etf1 | 3.03E-10 | 0.85164 | 0.276 | 0.164 | 7.15E-06 |
| Fnip2 | 3.09E-10 | 1.040378 | 0.249 | 0.144 | 7.29E-06 |
| Gm12070 | 3.14E-10 | 0.518081 | 0.217 | 0.121 | 7.4E-06 |
| Gm7901 | 3.23E-10 | 0.556148 | 0.379 | 0.247 | 7.62E-06 |
| Eif3b | 3.3E-10 | 0.546068 | 0.377 | 0.246 | 7.78E-06 |
| Sod2 | 3.44E-10 | 0.618822 | 0.3 | 0.182 | 8.11E-06 |
| Snrpf | 3.61E-10 | 0.352548 | 0.352 | 0.221 | 8.52E-06 |
| Rpl39 | 3.69E-10 | 0.314283 | 0.832 | 0.717 | 8.7E-06 |
| Rpl36a | 3.81E-10 | 0.47484 | 0.688 | 0.529 | 8.98E-06 |
| Pa2g4 | 3.91E-10 | 0.623402 | 0.327 | 0.205 | 9.21E-06 |
| Gm14760 | 4.03E-10 | 0.252527 | 0.233 | 0.131 | 9.5E-06 |
| Cox4i1 | 4.82E-10 | 0.407695 | 0.863 | 0.773 | 1.14E-05 |
| Rasgef1b | 4.82E-10 | 0.569701 | 0.2 | 0.105 | 1.14E-05 |
| Vps29 | 5.27E-10 | 0.549958 | 0.469 | 0.328 | 1.24E-05 |
| Bag1 | 5.29E-10 | 0.53391 | 0.446 | 0.311 | 1.25E-05 |
| Rps19-ps6 | 5.44E-10 | 0.370582 | 0.78 | 0.646 | 1.28E-05 |
| Mrpl12 | 5.53E-10 | 0.693384 | 0.28 | 0.166 | 1.3E-05 |
| Gabarap | 5.79E-10 | 0.430319 | 0.841 | 0.718 | 1.37E-05 |
| Eif1ax | 6.03E-10 | 0.719075 | 0.202 | 0.108 | 1.42E-05 |
| Eif3i | 6.11E-10 | 0.79386 | 0.363 | 0.24 | 1.44E-05 |
| Eif3c | 6.12E-10 | 0.377005 | 0.572 | 0.426 | 1.44E-05 |
| H2-Ab1 | 6.24E-10 | 2.570214 | 0.26 | 0.164 | 1.47E-05 |
| Ndufa4 | 6.55E-10 | 0.41709 | 0.561 | 0.402 | 1.54E-05 |
| Rps19-ps4 | 6.81E-10 | 0.479332 | 0.217 | 0.121 | 1.61E-05 |
| Rars | 7.16E-10 | 0.636899 | 0.213 | 0.116 | 1.69E-05 |
| Rpl19-ps11 | 7.59E-10 | 0.479103 | 0.751 | 0.615 | 1.79E-05 |
| Syncrip | 7.62E-10 | 0.901758 | 0.296 | 0.183 | 1.8E-05 |
| Vmp1 | 7.73E-10 | 0.41582 | 0.383 | 0.248 | 1.82E-05 |
| Gm10080 | 7.86E-10 | 0.272141 | 0.334 | 0.212 | 1.85E-05 |
| Cd48 | 7.94E-10 | 0.494604 | 0.549 | 0.402 | 1.87E-05 |
| Ndufa12 | 9.08E-10 | 0.590598 | 0.341 | 0.216 | 2.14E-05 |
| Rnf149 | 9.23E-10 | 0.563101 | 0.312 | 0.192 | 2.18E-05 |

| | | | | | |
|---|---|---|---|---|---|
| Atp1b3 | 9.53E-10 | 0.446963 | 0.774 | 0.65 | 2.25E-05 |
| Bcl2l1 | 1.01E-09 | 0.626479 | 0.522 | 0.384 | 2.37E-05 |
| Gm13436 | 1.03E-09 | 0.470307 | 0.617 | 0.474 | 2.42E-05 |
| Retreg2 | 1.08E-09 | 0.625155 | 0.386 | 0.255 | 2.54E-05 |
| Cd180 | 1.09E-09 | 0.547275 | 0.552 | 0.426 | 2.58E-05 |
| Psme1 | 1.1E-09 | 0.454708 | 0.646 | 0.498 | 2.59E-05 |
| Rpl37a | 1.18E-09 | 0.292099 | 0.964 | 0.94 | 2.78E-05 |
| Nop10 | 1.19E-09 | 0.448857 | 0.339 | 0.211 | 2.81E-05 |
| Gm10123 | 1.2E-09 | 0.49532 | 0.948 | 0.925 | 2.84E-05 |
| Rpsa-ps2 | 1.21E-09 | 0.31817 | 0.327 | 0.212 | 2.84E-05 |
| Prelid1 | 1.25E-09 | 0.727097 | 0.549 | 0.441 | 2.95E-05 |
| Eif5b | 1.27E-09 | 0.523033 | 0.381 | 0.255 | 2.99E-05 |
| Ddx21 | 1.33E-09 | 1.045314 | 0.269 | 0.161 | 3.15E-05 |
| Srp14 | 1.35E-09 | 0.42624 | 0.457 | 0.316 | 3.18E-05 |
| Snu13 | 1.35E-09 | 0.566805 | 0.345 | 0.224 | 3.19E-05 |
| Hypk | 1.36E-09 | 0.584018 | 0.265 | 0.156 | 3.21E-05 |
| Mrpl39 | 1.37E-09 | 0.844392 | 0.222 | 0.124 | 3.22E-05 |
| Taf10 | 1.41E-09 | 0.330705 | 0.516 | 0.382 | 3.32E-05 |
| Rbm3 | 1.46E-09 | 0.505075 | 0.426 | 0.291 | 3.45E-05 |
| Rpl23a-ps3 | 1.71E-09 | 0.419542 | 0.628 | 0.492 | 4.03E-05 |
| Slc35b1 | 1.71E-09 | 0.61917 | 0.374 | 0.25 | 4.04E-05 |
| mt-Nd1 | 1.78E-09 | 0.508158 | 0.955 | 0.927 | 4.19E-05 |
| Hnrnpm | 1.78E-09 | 0.450781 | 0.531 | 0.395 | 4.2E-05 |
| Ybx3 | 1.78E-09 | 1.062557 | 0.202 | 0.113 | 4.21E-05 |
| Gm9347 | 1.8E-09 | 0.257211 | 0.3 | 0.186 | 4.25E-05 |
| Rpl38-ps2 | 1.84E-09 | 0.33633 | 0.343 | 0.224 | 4.34E-05 |
| Erh | 1.84E-09 | 0.426811 | 0.473 | 0.323 | 4.35E-05 |
| Rps3 | 1.85E-09 | 0.410367 | 0.951 | 0.901 | 4.35E-05 |
| Phb2 | 1.87E-09 | 0.545311 | 0.401 | 0.272 | 4.42E-05 |
| Minos1 | 1.91E-09 | 0.441448 | 0.516 | 0.377 | 4.5E-05 |
| Gm4294 | 1.91E-09 | 0.38668 | 0.487 | 0.347 | 4.51E-05 |
| Ndufb7 | 1.92E-09 | 0.317547 | 0.543 | 0.385 | 4.52E-05 |
| Cct8 | 2.04E-09 | 0.50002 | 0.507 | 0.373 | 4.82E-05 |
| Eif3l | 2.16E-09 | 0.635887 | 0.401 | 0.275 | 5.09E-05 |
| H2afz | 2.26E-09 | 0.473213 | 0.614 | 0.484 | 5.32E-05 |
| Gm14586 | 2.3E-09 | 0.429057 | 0.59 | 0.453 | 5.42E-05 |
| Noc2l | 2.35E-09 | 0.752851 | 0.291 | 0.18 | 5.54E-05 |
| Gm8129 | 2.37E-09 | 0.458883 | 0.413 | 0.288 | 5.58E-05 |
| mt-Rnr2 | 2.43E-09 | 0.361644 | 0.987 | 0.99 | 5.73E-05 |
| Tpt1 | 2.52E-09 | 0.445136 | 0.93 | 0.879 | 5.95E-05 |
| Gm15501 | 2.72E-09 | 0.530703 | 0.621 | 0.487 | 6.42E-05 |
| Nop58 | 2.73E-09 | 0.85736 | 0.267 | 0.163 | 6.43E-05 |
| Anapc15 | 2.77E-09 | 1.124943 | 0.202 | 0.112 | 6.53E-05 |
| Rps18-ps3 | 2.77E-09 | 0.5122 | 0.686 | 0.583 | 6.53E-05 |
| Cct2 | 2.84E-09 | 0.42374 | 0.41 | 0.281 | 6.7E-05 |
| Neat1 | 3.19E-09 | 0.802048 | 0.325 | 0.216 | 7.52E-05 |
| Clip1 | 3.24E-09 | 0.551196 | 0.475 | 0.343 | 7.64E-05 |

| | | | | | |
|---|---|---|---|---|---|
| Pdcd5 | 3.28E-09 | 0.508452 | 0.37 | 0.244 | 7.73E-05 |
| Rpl27 | 3.31E-09 | 0.421519 | 0.807 | 0.693 | 7.8E-05 |
| Bri3 | 3.31E-09 | 0.442922 | 0.726 | 0.577 | 7.82E-05 |
| Rpl23a | 3.35E-09 | 0.443809 | 0.886 | 0.766 | 7.89E-05 |
| Sirt2 | 3.37E-09 | 0.67064 | 0.444 | 0.323 | 7.94E-05 |
| Echs1 | 3.47E-09 | 0.522201 | 0.35 | 0.229 | 8.19E-05 |
| Rpl31-ps8 | 3.76E-09 | 0.490781 | 0.72 | 0.593 | 8.86E-05 |
| Set | 3.79E-09 | 0.372317 | 0.713 | 0.589 | 8.93E-05 |
| Gm9800 | 3.82E-09 | 0.363768 | 0.348 | 0.227 | 9.01E-05 |
| Phb | 3.87E-09 | 0.657783 | 0.377 | 0.257 | 9.13E-05 |
| H13 | 4.08E-09 | 0.482874 | 0.527 | 0.377 | 9.63E-05 |
| Gm10053 | 4.16E-09 | 0.297859 | 0.262 | 0.158 | 9.8E-05 |
| Calm3 | 4.46E-09 | 0.404615 | 0.359 | 0.236 | 0.000105 |
| Cct5 | 4.65E-09 | 0.571555 | 0.491 | 0.364 | 0.00011 |
| Plekha1 | 4.84E-09 | 0.639162 | 0.298 | 0.191 | 0.000114 |
| B4galnt1 | 5.04E-09 | 0.804369 | 0.213 | 0.121 | 0.000119 |
| Ndufa7 | 5.06E-09 | 0.462114 | 0.531 | 0.392 | 0.000119 |
| Nedd8 | 5.17E-09 | 0.552593 | 0.525 | 0.389 | 0.000122 |
| Slamf9 | 5.17E-09 | 0.550214 | 0.473 | 0.344 | 0.000122 |
| Prkcsh | 5.22E-09 | 0.836677 | 0.374 | 0.26 | 0.000123 |
| Gm10800 | 5.45E-09 | 0.270987 | 0.617 | 0.725 | 0.000128 |
| Snx3 | 5.67E-09 | 0.423164 | 0.578 | 0.431 | 0.000134 |
| Actg1 | 5.69E-09 | 0.618012 | 0.879 | 0.832 | 0.000134 |
| Abracl | 6.33E-09 | 0.322615 | 0.305 | 0.187 | 0.000149 |
| Atp6v0e | 6.45E-09 | 0.458579 | 0.612 | 0.465 | 0.000152 |
| Nop16 | 6.49E-09 | 0.780572 | 0.226 | 0.13 | 0.000153 |
| Gm6170 | 6.54E-09 | 0.310039 | 0.484 | 0.36 | 0.000154 |
| Nutf2-ps1 | 7.34E-09 | 0.425721 | 0.253 | 0.15 | 0.000173 |
| Cmtm3 | 7.37E-09 | 0.449764 | 0.502 | 0.371 | 0.000174 |
| Gm11361 | 8.69E-09 | 0.339745 | 0.594 | 0.47 | 0.000205 |
| Mlf2 | 8.72E-09 | 0.398284 | 0.5 | 0.358 | 0.000206 |
| Hspa4 | 8.79E-09 | 0.470692 | 0.484 | 0.351 | 0.000207 |
| Rpl24 | 8.8E-09 | 0.422339 | 0.731 | 0.616 | 0.000208 |
| Iah1 | 8.81E-09 | 0.875221 | 0.224 | 0.13 | 0.000208 |
| Rbx1 | 9.54E-09 | 0.303629 | 0.576 | 0.436 | 0.000225 |
| Ubl5 | 1.04E-08 | 0.370956 | 0.54 | 0.403 | 0.000246 |
| Rpl36-ps2 | 1.05E-08 | 0.370521 | 0.637 | 0.514 | 0.000248 |
| Galc | 1.06E-08 | 0.556939 | 0.202 | 0.115 | 0.00025 |
| Crlf2 | 1.07E-08 | 0.57757 | 0.525 | 0.397 | 0.000252 |
| Btf3 | 1.07E-08 | 0.518442 | 0.601 | 0.464 | 0.000252 |
| Actn4 | 1.09E-08 | 0.593204 | 0.354 | 0.24 | 0.000257 |
| Calr | 1.09E-08 | 0.411829 | 0.939 | 0.898 | 0.000258 |
| Lrrc59 | 1.09E-08 | 0.531165 | 0.253 | 0.154 | 0.000258 |
| Gm8430 | 1.13E-08 | 0.372683 | 0.648 | 0.521 | 0.000266 |
| Gm12715 | 1.14E-08 | 0.71924 | 0.751 | 0.683 | 0.00027 |
| Snrpg | 1.22E-08 | 0.338522 | 0.5 | 0.356 | 0.000288 |
| 1500011K1 | 1.22E-08 | 0.388922 | 0.249 | 0.148 | 0.000288 |

| | | | | | |
|---|---|---|---|---|---|
| Mvp | 1.23E-08 | 0.53414 | 0.274 | 0.169 | 0.000291 |
| Gm3788 | 1.25E-08 | 0.348541 | 0.224 | 0.132 | 0.000296 |
| Sod1 | 1.26E-08 | 0.526226 | 0.731 | 0.602 | 0.000297 |
| M6pr | 1.3E-08 | 0.483108 | 0.682 | 0.57 | 0.000307 |
| Eif3e | 1.33E-08 | 0.378016 | 0.397 | 0.273 | 0.000313 |
| Atp5d | 1.52E-08 | 0.333935 | 0.567 | 0.423 | 0.000358 |
| Rpl26 | 1.61E-08 | 0.35643 | 0.848 | 0.767 | 0.00038 |
| Mrpl54 | 1.63E-08 | 0.444969 | 0.231 | 0.136 | 0.000384 |
| Por | 1.66E-08 | 0.780828 | 0.352 | 0.24 | 0.000391 |
| Ucp2 | 1.66E-08 | 0.467657 | 0.717 | 0.624 | 0.000393 |
| Rpl28-ps1 | 1.67E-08 | 0.498331 | 0.578 | 0.438 | 0.000393 |
| Hnrnpu | 1.7E-08 | 0.565992 | 0.545 | 0.421 | 0.0004 |
| Impdh2 | 1.76E-08 | 0.854251 | 0.231 | 0.138 | 0.000415 |
| Tomm40 | 1.84E-08 | 0.720058 | 0.274 | 0.172 | 0.000434 |
| Gm8186 | 1.92E-08 | 0.383943 | 0.419 | 0.287 | 0.000452 |
| Rps6-ps4 | 1.93E-08 | 0.414211 | 0.614 | 0.505 | 0.000456 |
| Rpl27a | 1.95E-08 | 0.390811 | 0.848 | 0.746 | 0.000459 |
| Uqcr10 | 1.96E-08 | 0.268604 | 0.442 | 0.307 | 0.000461 |
| Kars | 1.99E-08 | 0.492597 | 0.314 | 0.199 | 0.00047 |
| Dnmt3a | 2E-08 | 0.429356 | 0.552 | 0.408 | 0.000472 |
| Mfsd11 | 2.01E-08 | 0.407323 | 0.502 | 0.373 | 0.000475 |
| Oaz1 | 2.03E-08 | 0.287393 | 0.886 | 0.768 | 0.000478 |
| Ndufa11 | 2.04E-08 | 0.281029 | 0.374 | 0.245 | 0.000482 |
| Got2 | 2.07E-08 | 0.439455 | 0.262 | 0.162 | 0.000489 |
| Pfkfb3 | 2.17E-08 | 0.630909 | 0.357 | 0.239 | 0.000512 |
| Gm10443 | 2.18E-08 | 0.250111 | 0.538 | 0.409 | 0.000514 |
| App | 2.19E-08 | 0.521241 | 0.552 | 0.439 | 0.000516 |
| Rpl18-ps2 | 2.28E-08 | 0.393231 | 0.484 | 0.364 | 0.000538 |
| Fau | 2.29E-08 | 0.35666 | 0.955 | 0.905 | 0.000541 |
| Cdv3 | 2.32E-08 | 0.290381 | 0.469 | 0.332 | 0.000548 |
| Rps10-ps2 | 2.39E-08 | 0.432713 | 0.726 | 0.593 | 0.000565 |
| Commd9 | 2.46E-08 | 0.474238 | 0.247 | 0.148 | 0.000579 |
| Chd4 | 2.58E-08 | 0.355094 | 0.439 | 0.319 | 0.000609 |
| D8Ertd738 | 2.82E-08 | 0.356603 | 0.498 | 0.358 | 0.000664 |
| Gtf2h5 | 2.9E-08 | 0.47978 | 0.217 | 0.125 | 0.000683 |
| Dohh | 2.97E-08 | 0.786856 | 0.213 | 0.123 | 0.000701 |
| Rpl34 | 3.09E-08 | 0.324335 | 0.899 | 0.791 | 0.00073 |
| Gpr137b-p | 3.13E-08 | 0.500045 | 0.457 | 0.336 | 0.000739 |
| Gm3531 | 3.48E-08 | 0.296023 | 0.39 | 0.268 | 0.000821 |
| Litaf | 3.87E-08 | 0.49379 | 0.578 | 0.454 | 0.000914 |
| Emc1 | 3.97E-08 | 0.738433 | 0.204 | 0.118 | 0.000937 |
| Cyc1 | 4.12E-08 | 0.628517 | 0.404 | 0.287 | 0.000972 |
| Sigmar1 | 4.22E-08 | 0.589708 | 0.298 | 0.194 | 0.000996 |
| Cox6b1 | 4.39E-08 | 0.337782 | 0.641 | 0.5 | 0.001035 |
| Wdr83os | 4.39E-08 | 0.327849 | 0.357 | 0.239 | 0.001035 |
| Denr | 4.52E-08 | 0.340453 | 0.39 | 0.267 | 0.001066 |
| Dbi | 4.75E-08 | 0.677706 | 0.341 | 0.232 | 0.00112 |

| | | | | | |
|---|---|---|---|---|---|
| Gm6180 | 4.86E-08 | 0.467449 | 0.507 | 0.398 | 0.001146 |
| Tspo | 4.92E-08 | 0.435503 | 0.446 | 0.324 | 0.001161 |
| Edem1 | 5.36E-08 | 0.57389 | 0.648 | 0.563 | 0.001265 |
| Gm12918 | 5.48E-08 | 0.378211 | 0.632 | 0.521 | 0.001292 |
| Ndufa13 | 5.52E-08 | 0.313826 | 0.614 | 0.47 | 0.001302 |
| Rps25-ps1 | 5.55E-08 | 0.256041 | 0.406 | 0.289 | 0.001309 |
| Ugp2 | 5.56E-08 | 0.818221 | 0.247 | 0.154 | 0.001311 |
| Tcof1 | 5.71E-08 | 0.50664 | 0.278 | 0.177 | 0.001347 |
| Hsp90b1 | 6.1E-08 | 0.331465 | 0.942 | 0.918 | 0.001439 |
| Rps14 | 6.39E-08 | 0.423779 | 0.964 | 0.922 | 0.001507 |
| Fbl | 6.53E-08 | 0.652945 | 0.3 | 0.196 | 0.001541 |
| Ptp4a1 | 6.59E-08 | 0.727653 | 0.217 | 0.131 | 0.001555 |
| Hnrnpf | 6.64E-08 | 0.358419 | 0.697 | 0.564 | 0.001567 |
| Rps23 | 6.65E-08 | 0.307879 | 0.955 | 0.897 | 0.001569 |
| Fis1 | 6.69E-08 | 0.524269 | 0.484 | 0.357 | 0.001579 |
| Ndufb6 | 6.94E-08 | 0.524331 | 0.291 | 0.185 | 0.001636 |
| Gm10146 | 6.94E-08 | 0.365953 | 0.2 | 0.114 | 0.001638 |
| Rps12 | 7.08E-08 | 0.409561 | 0.901 | 0.832 | 0.001671 |
| Rps6 | 7.36E-08 | 0.512468 | 0.841 | 0.755 | 0.001735 |
| Llph | 7.96E-08 | 0.509541 | 0.296 | 0.192 | 0.001878 |
| Gm10132 | 8.13E-08 | 0.430089 | 0.704 | 0.583 | 0.001919 |
| Drap1 | 8.36E-08 | 0.416576 | 0.433 | 0.308 | 0.001972 |
| Oaz1-ps | 8.39E-08 | 0.277019 | 0.756 | 0.626 | 0.001979 |
| Gng5-ps | 8.61E-08 | 0.459843 | 0.374 | 0.262 | 0.002032 |
| Rab7 | 8.75E-08 | 0.327752 | 0.682 | 0.55 | 0.002064 |
| Tomm7 | 9.01E-08 | 0.389869 | 0.43 | 0.304 | 0.002126 |
| Capns1 | 9.03E-08 | 0.634136 | 0.419 | 0.307 | 0.00213 |
| Atxn10 | 9.2E-08 | 0.521147 | 0.466 | 0.341 | 0.002169 |
| Gm7536 | 9.26E-08 | 0.276828 | 0.704 | 0.585 | 0.002184 |
| Gm9531 | 1.01E-07 | 0.398489 | 0.204 | 0.119 | 0.002374 |
| Tuba1c | 1.02E-07 | 1.36656 | 0.457 | 0.383 | 0.002401 |
| Smdt1 | 1.03E-07 | 0.275365 | 0.415 | 0.287 | 0.002432 |
| Rpl22 | 1.05E-07 | 0.415367 | 0.684 | 0.545 | 0.002485 |
| G3bp1 | 1.06E-07 | 0.54551 | 0.516 | 0.404 | 0.002504 |
| Psmd7 | 1.1E-07 | 0.314554 | 0.437 | 0.314 | 0.00259 |
| Tmed3 | 1.15E-07 | 0.483582 | 0.592 | 0.474 | 0.002715 |
| Rsl1d1 | 1.3E-07 | 0.549935 | 0.359 | 0.247 | 0.003066 |
| Ggh | 1.46E-07 | 0.526691 | 0.309 | 0.207 | 0.003453 |
| Naa20 | 1.5E-07 | 0.429021 | 0.417 | 0.294 | 0.003539 |
| Tns3 | 1.5E-07 | 0.696789 | 0.435 | 0.322 | 0.003549 |
| Ubxn1 | 1.57E-07 | 0.590834 | 0.372 | 0.267 | 0.003705 |
| Psme2 | 1.64E-07 | 0.574297 | 0.585 | 0.494 | 0.003878 |
| Ssr2 | 1.66E-07 | 0.48143 | 0.502 | 0.387 | 0.003906 |
| Arpc3 | 1.67E-07 | 0.322967 | 0.722 | 0.603 | 0.003929 |
| Slc23a2 | 1.69E-07 | 0.707223 | 0.3 | 0.199 | 0.003985 |
| Sdf4 | 1.79E-07 | 0.367251 | 0.7 | 0.579 | 0.004225 |
| Sec61b | 1.82E-07 | 0.259092 | 0.424 | 0.297 | 0.004291 |

| | | | | | |
|---|---|---|---|---|---|
| Tpt1-ps3 | 1.9E-07 | 0.366615 | 0.735 | 0.636 | 0.004481 |
| Tma7 | 1.96E-07 | 0.25002 | 0.529 | 0.395 | 0.004632 |
| Gm3511 | 1.97E-07 | 0.355418 | 0.682 | 0.572 | 0.004637 |
| Rpl29 | 2.03E-07 | 0.535794 | 0.76 | 0.652 | 0.004784 |
| Cd34 | 2.03E-07 | 0.41262 | 0.628 | 0.512 | 0.004795 |
| Rpl8 | 2.05E-07 | 0.484315 | 0.8 | 0.712 | 0.004827 |
| Ptges3-ps | 2.21E-07 | 0.47822 | 0.274 | 0.177 | 0.00521 |
| Naca | 2.25E-07 | 0.352617 | 0.731 | 0.597 | 0.005299 |
| mt-Co1 | 2.25E-07 | 0.472454 | 0.987 | 0.986 | 0.005304 |
| Rpl30 | 2.26E-07 | 0.310638 | 0.933 | 0.867 | 0.005327 |
| Hebp1 | 2.34E-07 | 0.337446 | 0.278 | 0.178 | 0.005519 |
| Igbp1 | 2.43E-07 | 0.485551 | 0.247 | 0.155 | 0.005728 |
| 2410015M | 2.68E-07 | 0.266514 | 0.39 | 0.273 | 0.006316 |
| Dad1 | 2.8E-07 | 0.358074 | 0.695 | 0.578 | 0.006604 |
| Copg1 | 2.83E-07 | 0.86942 | 0.26 | 0.172 | 0.006677 |
| Gltp | 2.99E-07 | 0.412472 | 0.498 | 0.38 | 0.007064 |
| Arhgdia | 3.07E-07 | 0.351274 | 0.881 | 0.799 | 0.007243 |
| Gm13509 | 3.12E-07 | 0.257163 | 0.424 | 0.303 | 0.007352 |
| Pgd | 3.24E-07 | 0.526553 | 0.46 | 0.344 | 0.007644 |
| H2afj | 3.26E-07 | 0.36303 | 0.426 | 0.303 | 0.007693 |
| Copa | 3.44E-07 | 0.590864 | 0.444 | 0.33 | 0.008119 |
| Mtch2 | 3.47E-07 | 0.386796 | 0.307 | 0.206 | 0.008189 |
| Ndufb9 | 3.56E-07 | 0.444955 | 0.491 | 0.375 | 0.008387 |
| Rpl22l1 | 3.63E-07 | 0.314667 | 0.361 | 0.255 | 0.008551 |
| Atp6v0d1 | 3.63E-07 | 0.98501 | 0.498 | 0.392 | 0.008567 |
| Gm5560 | 3.64E-07 | 0.309324 | 0.453 | 0.336 | 0.008595 |
| Impact | 3.7E-07 | 0.696936 | 0.343 | 0.242 | 0.00874 |
| Eef2 | 3.74E-07 | 0.291473 | 0.839 | 0.764 | 0.00882 |
| 1810058I2 | 3.8E-07 | 0.584336 | 0.296 | 0.194 | 0.00896 |
| Nup62 | 3.82E-07 | 0.435484 | 0.209 | 0.125 | 0.00902 |
| Tns1 | 3.85E-07 | 0.608233 | 0.341 | 0.237 | 0.009071 |
| Rpl35a | 4.15E-07 | 0.362835 | 0.814 | 0.74 | 0.009799 |
| Snrpd2 | 4.2E-07 | 0.643382 | 0.294 | 0.192 | 0.009898 |
| Atp6ap2 | 4.28E-07 | 0.40132 | 0.715 | 0.605 | 0.010101 |
| Frrs1 | 4.36E-07 | 0.492648 | 0.48 | 0.365 | 0.010294 |
| Neu1 | 4.39E-07 | 0.532756 | 0.318 | 0.216 | 0.010359 |
| Mat2a | 4.48E-07 | 1.034861 | 0.334 | 0.238 | 0.010556 |
| Ndufs6 | 4.53E-07 | 0.337308 | 0.37 | 0.259 | 0.01068 |
| Tmem167 | 4.53E-07 | 0.29613 | 0.318 | 0.218 | 0.010692 |
| Gm28437 | 4.61E-07 | 0.45699 | 0.881 | 0.827 | 0.010879 |
| Rps24 | 4.81E-07 | 0.382636 | 0.91 | 0.837 | 0.011345 |
| Kras | 4.96E-07 | 0.56738 | 0.334 | 0.233 | 0.011703 |
| Ddx1 | 5.03E-07 | 0.562843 | 0.253 | 0.163 | 0.011865 |
| Mrps24 | 5.3E-07 | 0.656397 | 0.274 | 0.184 | 0.012492 |
| Ssr3 | 5.33E-07 | 0.469914 | 0.401 | 0.301 | 0.012568 |
| Tars | 5.47E-07 | 0.659827 | 0.217 | 0.135 | 0.012901 |
| tpr | 5.47E-07 | 0.343552 | 0.399 | 0.289 | 0.012908 |

| | | | | | |
|---|---|---|---|---|---|
| Rpl36 | 5.68E-07 | 0.293958 | 0.899 | 0.82 | 0.013395 |
| Nupr1 | 5.69E-07 | 0.994394 | 0.235 | 0.153 | 0.013424 |
| Rpl27-ps3 | 5.79E-07 | 0.29283 | 0.478 | 0.364 | 0.013662 |
| Cisd2 | 5.9E-07 | 0.447078 | 0.39 | 0.285 | 0.013918 |
| Tec | 5.92E-07 | 0.35424 | 0.204 | 0.124 | 0.013973 |
| Snrpd3 | 6.14E-07 | 0.458121 | 0.332 | 0.228 | 0.014489 |
| Gars | 6.3E-07 | 0.630608 | 0.276 | 0.185 | 0.014857 |
| Dnajb11 | 6.55E-07 | 0.447333 | 0.475 | 0.357 | 0.015445 |
| Cmas | 6.86E-07 | 0.319797 | 0.289 | 0.192 | 0.016185 |
| Rpl7 | 7.35E-07 | 0.301211 | 0.879 | 0.8 | 0.017331 |
| Cux1 | 7.38E-07 | 0.30583 | 0.599 | 0.475 | 0.017416 |
| Pi4ka | 7.44E-07 | 0.655 | 0.231 | 0.146 | 0.017546 |
| Rpl10 | 7.44E-07 | 0.349165 | 0.948 | 0.902 | 0.017556 |
| Gm10736 | 7.49E-07 | 0.394418 | 0.626 | 0.525 | 0.017675 |
| Igsf8 | 7.86E-07 | 0.408428 | 0.323 | 0.222 | 0.018533 |
| Pole4 | 7.86E-07 | 0.483657 | 0.235 | 0.148 | 0.018537 |
| Rps13-ps2 | 8.01E-07 | 0.450338 | 0.641 | 0.526 | 0.018891 |
| Ccdc86 | 8.01E-07 | 0.435016 | 0.213 | 0.131 | 0.018904 |
| Eif3j1 | 8.17E-07 | 0.466828 | 0.43 | 0.32 | 0.019261 |
| Gm11942 | 8.48E-07 | 0.250086 | 0.545 | 0.44 | 0.020004 |
| Mroh1 | 8.54E-07 | 0.631437 | 0.226 | 0.144 | 0.020133 |
| Psmb4 | 8.63E-07 | 0.615841 | 0.462 | 0.353 | 0.020352 |
| Gpr137b | 8.72E-07 | 0.387095 | 0.339 | 0.238 | 0.020572 |
| Gm11518 | 9.18E-07 | 0.259596 | 0.623 | 0.486 | 0.021649 |
| Cltb | 9.31E-07 | 0.66566 | 0.271 | 0.181 | 0.021961 |
| Ssrp1 | 9.32E-07 | 0.792625 | 0.305 | 0.205 | 0.021993 |
| Uqcrb | 1.02E-06 | 0.504649 | 0.473 | 0.356 | 0.024173 |
| Zcrb1 | 1.05E-06 | 0.68923 | 0.253 | 0.166 | 0.024824 |
| Clptm1l | 1.07E-06 | 0.485697 | 0.442 | 0.335 | 0.025214 |
| Gm26917 | 1.07E-06 | 0.864902 | 0.89 | 0.948 | 0.025358 |
| Dpagt1 | 1.08E-06 | 0.344217 | 0.296 | 0.2 | 0.025424 |
| Gm7931 | 1.1E-06 | 0.323267 | 0.229 | 0.147 | 0.025927 |
| Aurkaip1 | 1.1E-06 | 0.492147 | 0.28 | 0.188 | 0.025937 |
| Psmc1 | 1.18E-06 | 0.386565 | 0.352 | 0.25 | 0.027823 |
| Gm16286 | 1.19E-06 | 0.254876 | 0.422 | 0.31 | 0.02802 |
| Atp2b1 | 1.25E-06 | 0.457017 | 0.585 | 0.483 | 0.029458 |
| Rpl36al | 1.25E-06 | 0.366255 | 0.617 | 0.502 | 0.02952 |
| Serbp1 | 1.25E-06 | 0.268723 | 0.639 | 0.52 | 0.029601 |
| Ak6 | 1.31E-06 | 0.4134 | 0.318 | 0.22 | 0.030987 |
| Polr2g | 1.32E-06 | 0.440117 | 0.312 | 0.214 | 0.031073 |
| Polr2i | 1.35E-06 | 0.667976 | 0.244 | 0.156 | 0.031904 |
| Slc7a6 | 1.4E-06 | 0.419724 | 0.22 | 0.139 | 0.033119 |
| Eif1 | 1.44E-06 | 0.382159 | 0.935 | 0.871 | 0.033904 |
| Myo1f | 1.46E-06 | 0.435861 | 0.643 | 0.542 | 0.034396 |
| Tubb5 | 1.49E-06 | 0.300047 | 0.872 | 0.785 | 0.035172 |
| Mrpl23 | 1.49E-06 | 0.315032 | 0.361 | 0.256 | 0.035213 |
| Lsm4 | 1.53E-06 | 0.539557 | 0.265 | 0.174 | 0.036093 |

| | | | | | |
|---|---|---|---|---|---|
| Pgls | 1.54E-06 | 0.347889 | 0.401 | 0.288 | 0.036257 |
| Ormdl2 | 1.59E-06 | 0.399876 | 0.327 | 0.227 | 0.037495 |
| Gm8995 | 1.64E-06 | 0.512314 | 0.312 | 0.214 | 0.038684 |
| Snd1 | 1.78E-06 | 0.411937 | 0.339 | 0.242 | 0.042066 |
| Atp5g3 | 1.87E-06 | 0.329169 | 0.473 | 0.369 | 0.044189 |
| Txnl1 | 1.91E-06 | 0.529259 | 0.334 | 0.24 | 0.045125 |
| Gpx4 | 1.95E-06 | 0.347886 | 0.608 | 0.499 | 0.045899 |
| Sdhc | 1.99E-06 | 0.423777 | 0.419 | 0.312 | 0.046873 |
| Atp5h | 2.04E-06 | 0.397569 | 0.563 | 0.44 | 0.048162 |
| Meaf6 | 2.07E-06 | 0.467967 | 0.217 | 0.137 | 0.048739 |

### Supplementry Table 1. Marker genes for IRM

| Genenames | p_val | avg_log2FC | pct.1 | pct.2 | p_val_adj |
|---|---|---|---|---|---|
| Ifit3 | 0 | 4.895551 | 0.487 | 0.047 | 0 |
| Ifit3b | 0 | 5.0447 | 0.349 | 0.021 | 0 |
| Ifitm3 | 0 | 4.338424 | 0.781 | 0.17 | 0 |
| Irf7 | 0 | 4.346364 | 0.422 | 0.041 | 0 |
| Isg15 | 0 | 3.794183 | 0.446 | 0.043 | 0 |
| Oasl2 | 0 | 4.293938 | 0.517 | 0.051 | 0 |
| Rtp4 | 0 | 3.396895 | 0.716 | 0.12 | 0 |
| Zbp1 | 0 | 4.789052 | 0.339 | 0.02 | 0 |
| Ifi204 | 3.7E-301 | 3.670117 | 0.494 | 0.063 | 8.8E-297 |
| Ifit2 | 5E-296 | 4.789524 | 0.321 | 0.024 | 1.2E-291 |
| Usp18 | 9.7E-284 | 4.217784 | 0.54 | 0.085 | 2.3E-279 |
| Ifi27l2a | 8.9E-262 | 3.410583 | 0.658 | 0.141 | 2.1E-257 |
| Oas1a | 2E-260 | 3.382052 | 0.457 | 0.062 | 4.8E-256 |
| Ifi206 | 2E-256 | 4.018006 | 0.22 | 0.011 | 4.8E-252 |
| Ifi213 | 2.3E-254 | 4.653445 | 0.247 | 0.015 | 5.4E-250 |
| Oas2 | 5.1E-237 | 3.782403 | 0.233 | 0.015 | 1.2E-232 |
| Lgals3bp | 1.7E-235 | 2.542407 | 0.984 | 0.572 | 4E-231 |
| Sdc3 | 2.7E-191 | 2.886647 | 0.591 | 0.143 | 6.4E-187 |
| Ifit1 | 9.6E-191 | 4.794673 | 0.215 | 0.016 | 2.3E-186 |
| Tgtp2 | 2.5E-190 | 2.676698 | 0.21 | 0.015 | 5.8E-186 |
| Mx1 | 3E-190 | 4.237498 | 0.243 | 0.022 | 7.2E-186 |
| Nlrc5 | 5.6E-183 | 3.032914 | 0.316 | 0.04 | 1.3E-178 |
| Slfn5 | 4.2E-182 | 3.895581 | 0.39 | 0.066 | 1E-177 |
| Ifi209 | 6.7E-182 | 3.517153 | 0.344 | 0.049 | 1.6E-177 |
| Phf11b | 9.6E-180 | 3.25062 | 0.471 | 0.096 | 2.3E-175 |
| Sp100 | 8.1E-172 | 2.913023 | 0.519 | 0.119 | 1.9E-167 |
| Gm4951 | 2.1E-154 | 3.150663 | 0.34 | 0.056 | 4.8E-150 |
| Stat1 | 2.7E-152 | 2.727359 | 0.661 | 0.219 | 6.3E-148 |
| Bst2 | 4E-150 | 2.542321 | 0.817 | 0.402 | 9.4E-146 |
| Ifi207 | 1.4E-147 | 3.926342 | 0.247 | 0.031 | 3.2E-143 |
| Slfn2 | 1.7E-140 | 2.104587 | 0.564 | 0.159 | 3.9E-136 |
| Xaf1 | 1.6E-136 | 2.318882 | 0.533 | 0.148 | 3.9E-132 |
| Ccl12 | 3.1E-135 | 3.951927 | 0.57 | 0.191 | 7.2E-131 |
| Isg20 | 3.6E-129 | 4.020138 | 0.215 | 0.026 | 8.4E-125 |
| Phf11d | 5.6E-121 | 3.193813 | 0.384 | 0.09 | 1.3E-116 |
| Ube2l6 | 5.9E-116 | 2.663796 | 0.363 | 0.081 | 1.4E-111 |
| Dhx58 | 7.7E-109 | 2.747853 | 0.342 | 0.076 | 1.8E-104 |
| Rnf213 | 8.8E-107 | 2.198063 | 0.473 | 0.141 | 2.1E-102 |
| Parp14 | 4E-103 | 2.621492 | 0.515 | 0.175 | 9.5E-99 |
| Stat2 | 6.8E-99 | 2.793213 | 0.404 | 0.114 | 1.59E-94 |
| Fgl2 | 1.31E-97 | 3.150725 | 0.282 | 0.058 | 3.09E-93 |
| Ly6a | 4.11E-94 | 3.541378 | 0.42 | 0.134 | 9.69E-90 |
| H2-D1 | 5.63E-92 | 1.310796 | 0.993 | 0.907 | 1.33E-87 |

| | | | | | |
|---|---|---|---|---|---|
| Trim30a | 6.85E-92 | 2.181983 | 0.679 | 0.32 | 1.62E-87 |
| Gm12250 | 3.72E-89 | 3.037779 | 0.219 | 0.038 | 8.77E-85 |
| H2-K1 | 7.94E-85 | 1.308292 | 0.975 | 0.819 | 1.87E-80 |
| H2-Q7 | 3.95E-84 | 2.371466 | 0.372 | 0.109 | 9.32E-80 |
| H2-Q6 | 1.33E-82 | 2.301329 | 0.344 | 0.095 | 3.13E-78 |
| B2m | 2.69E-81 | 0.87779 | 1 | 0.998 | 6.34E-77 |
| Eif2ak2 | 8.42E-79 | 2.192796 | 0.416 | 0.138 | 1.99E-74 |
| Herc6 | 6.99E-78 | 2.599548 | 0.397 | 0.128 | 1.65E-73 |
| Ly6e | 1.62E-73 | 1.064615 | 0.986 | 0.951 | 3.81E-69 |
| Irgm1 | 2.06E-72 | 1.982119 | 0.434 | 0.156 | 4.87E-68 |
| Tor3a | 2.19E-71 | 2.283914 | 0.545 | 0.251 | 5.17E-67 |
| Axl | 8.27E-68 | 1.908933 | 0.279 | 0.074 | 1.95E-63 |
| H2-T23 | 4.43E-61 | 1.417627 | 0.859 | 0.662 | 1.05E-56 |
| Trim30d | 1.54E-59 | 1.804448 | 0.534 | 0.248 | 3.64E-55 |
| Prr5l | 5.89E-56 | 2.919328 | 0.206 | 0.051 | 1.39E-51 |
| Ifi35 | 1.77E-55 | 1.916987 | 0.48 | 0.222 | 4.18E-51 |
| Ddx60 | 2.46E-54 | 1.932424 | 0.231 | 0.062 | 5.81E-50 |
| Tap1 | 1.19E-53 | 1.82146 | 0.462 | 0.202 | 2.81E-49 |
| Atrip | 5.32E-53 | 0.719905 | 0.481 | 0.211 | 1.26E-48 |
| Tspo | 1.52E-52 | 1.662275 | 0.577 | 0.316 | 3.58E-48 |
| Ddx58 | 2.07E-52 | 1.948064 | 0.342 | 0.125 | 4.87E-48 |
| Shisa5 | 4.46E-52 | 1.19908 | 0.843 | 0.624 | 1.05E-47 |
| Fcgr1 | 1.66E-48 | 1.251475 | 0.861 | 0.723 | 3.92E-44 |
| Pml | 8.07E-48 | 2.530919 | 0.257 | 0.083 | 1.9E-43 |
| Igtp | 8.56E-47 | 1.792919 | 0.519 | 0.266 | 2.02E-42 |
| Clec2d | 1.39E-44 | 1.840932 | 0.242 | 0.076 | 3.27E-40 |
| Epsti1 | 1.63E-43 | 1.487752 | 0.386 | 0.165 | 3.84E-39 |
| Gm8995 | 2.72E-42 | 1.728926 | 0.432 | 0.207 | 6.41E-38 |
| Adar | 3.67E-42 | 1.432997 | 0.425 | 0.197 | 8.66E-38 |
| Parp12 | 3.79E-41 | 1.540607 | 0.42 | 0.195 | 8.93E-37 |
| Slfn8 | 1.11E-40 | 1.712859 | 0.21 | 0.064 | 2.62E-36 |
| Ifih1 | 4.14E-38 | 2.161194 | 0.266 | 0.1 | 9.76E-34 |
| Znfx1 | 5.28E-38 | 1.739284 | 0.459 | 0.237 | 1.25E-33 |
| Lgals9 | 1.43E-36 | 1.079719 | 0.824 | 0.676 | 3.38E-32 |
| Gm15753 | 2.67E-36 | 1.753074 | 0.302 | 0.125 | 6.29E-32 |
| Dtx3l | 1.2E-35 | 1.288855 | 0.42 | 0.205 | 2.84E-31 |
| Parp9 | 1.55E-35 | 1.433984 | 0.347 | 0.154 | 3.67E-31 |
| Samhd1 | 6.18E-35 | 1.564276 | 0.481 | 0.261 | 1.46E-30 |
| Sp110 | 2.33E-33 | 1.602936 | 0.494 | 0.283 | 5.49E-29 |
| H2-T24 | 3.31E-32 | 1.282075 | 0.46 | 0.245 | 7.8E-28 |
| Ogfr | 1.24E-31 | 1.399247 | 0.36 | 0.172 | 2.93E-27 |
| Grn | 2.49E-31 | 0.518801 | 0.996 | 0.993 | 5.87E-27 |
| Trim34a | 2.02E-30 | 1.783387 | 0.261 | 0.107 | 4.76E-26 |
| Nampt | 1.52E-28 | 1.769722 | 0.245 | 0.101 | 3.6E-24 |
| Zc3hav1 | 3.35E-28 | 1.257348 | 0.526 | 0.323 | 7.89E-24 |
| Ctsc | 5.91E-27 | 0.811977 | 0.922 | 0.846 | 1.39E-22 |
| Cd52 | 6.69E-27 | 1.007979 | 0.661 | 0.478 | 1.58E-22 |

| | | | | | |
|---|---|---|---|---|---|
| Trim25 | 1.83E-26 | 1.485658 | 0.224 | 0.091 | 4.32E-22 |
| AU020206 | 7.64E-25 | 0.781622 | 0.723 | 0.528 | 1.8E-20 |
| Irf1 | 2.68E-24 | 1.656154 | 0.323 | 0.166 | 6.31E-20 |
| Trim14 | 5.76E-24 | 1.173856 | 0.309 | 0.152 | 1.36E-19 |
| Trim12c | 7.25E-24 | 1.295009 | 0.337 | 0.177 | 1.71E-19 |
| Zup1 | 6.55E-23 | 1.270044 | 0.527 | 0.348 | 1.54E-18 |
| Psmb8 | 1.83E-22 | 0.736788 | 0.741 | 0.568 | 4.31E-18 |
| Uba7 | 2.34E-22 | 1.089005 | 0.367 | 0.201 | 5.51E-18 |
| Psme1 | 4.07E-22 | 0.9562 | 0.674 | 0.496 | 9.6E-18 |
| 1600014C1 | 5.54E-22 | 1.216851 | 0.374 | 0.21 | 1.31E-17 |
| 4930599N2 | 1.24E-21 | 1.417439 | 0.238 | 0.111 | 2.92E-17 |
| Psme2 | 7.54E-21 | 0.921298 | 0.653 | 0.49 | 1.78E-16 |
| Hck | 2.64E-20 | 0.8391 | 0.728 | 0.594 | 6.22E-16 |
| Tmem106a | 4.89E-20 | 1.328192 | 0.265 | 0.132 | 1.15E-15 |
| Tor1aip1 | 1.94E-19 | 0.980979 | 0.684 | 0.547 | 4.58E-15 |
| Selenow | 2.21E-19 | 1.280439 | 0.335 | 0.188 | 5.22E-15 |
| Trim12a | 1.71E-18 | 1.136403 | 0.561 | 0.409 | 4.04E-14 |
| Cd47 | 1.47E-17 | 0.865074 | 0.744 | 0.624 | 3.47E-13 |
| Ctss | 2.22E-17 | 0.308881 | 0.996 | 1 | 5.24E-13 |
| Psmb9 | 2.71E-17 | 0.841316 | 0.473 | 0.313 | 6.4E-13 |
| Calm1 | 2.51E-16 | 0.680412 | 0.808 | 0.734 | 5.91E-12 |
| Tmem184b | 5.38E-16 | 0.762799 | 0.469 | 0.315 | 1.27E-11 |
| Dopey2 | 9.77E-16 | 1.341326 | 0.272 | 0.15 | 2.3E-11 |
| Tent5a | 2.36E-15 | 0.874382 | 0.543 | 0.402 | 5.56E-11 |
| Trafd1 | 2.65E-15 | 0.922545 | 0.476 | 0.326 | 6.26E-11 |
| Ccdc86 | 1.8E-14 | 1.152674 | 0.24 | 0.129 | 4.24E-10 |
| Irf9 | 3.04E-14 | 1.018523 | 0.326 | 0.202 | 7.17E-10 |
| Fth1 | 7.62E-14 | 0.347424 | 0.996 | 0.995 | 1.8E-09 |
| Tor1aip2 | 8.4E-14 | 0.872146 | 0.432 | 0.296 | 1.98E-09 |
| Gm46224 | 1.04E-13 | 0.859759 | 0.261 | 0.146 | 2.45E-09 |
| Casp4 | 1.28E-13 | 1.35492 | 0.212 | 0.112 | 3.01E-09 |
| Hsp90ab1 | 1.49E-13 | 0.561675 | 0.901 | 0.839 | 3.52E-09 |
| Etnk1 | 1.76E-13 | 0.938215 | 0.466 | 0.328 | 4.14E-09 |
| Csf2ra | 1.81E-13 | 0.732794 | 0.436 | 0.296 | 4.27E-09 |
| Tmem219 | 5.4E-13 | 0.721823 | 0.499 | 0.364 | 1.27E-08 |
| Sppl2a | 5.43E-13 | 0.677808 | 0.684 | 0.555 | 1.28E-08 |
| Luc7l3 | 5.49E-13 | 0.867387 | 0.515 | 0.385 | 1.3E-08 |
| Cebpb | 6.85E-13 | 0.663671 | 0.647 | 0.529 | 1.62E-08 |
| Rnf114 | 8.96E-13 | 0.842476 | 0.464 | 0.331 | 2.11E-08 |
| Unc93b1 | 1.5E-12 | 0.349109 | 0.991 | 0.987 | 3.53E-08 |
| 9930111J2 | 1.53E-12 | 1.152045 | 0.441 | 0.316 | 3.6E-08 |
| Apobec3 | 1.68E-12 | 0.77946 | 0.624 | 0.488 | 3.97E-08 |
| Nmi | 1.85E-12 | 1.151979 | 0.245 | 0.142 | 4.37E-08 |
| Ly86 | 3.37E-12 | 0.307963 | 0.993 | 0.989 | 7.95E-08 |
| Naaa | 1.27E-11 | 0.935063 | 0.367 | 0.249 | 3E-07 |
| Anxa6 | 2.14E-11 | 0.696916 | 0.3 | 0.187 | 5.04E-07 |
| Rnase6 | 2.51E-11 | 1.100442 | 0.217 | 0.124 | 5.92E-07 |

| | | | | | |
|---|---|---|---|---|---|
| Gm26917 | 4.81E-11 | 0.60963 | 0.97 | 0.945 | 1.13E-06 |
| Irgm2 | 6.66E-11 | 0.778665 | 0.471 | 0.347 | 1.57E-06 |
| Pik3ap1 | 2.84E-10 | 0.850414 | 0.354 | 0.244 | 6.71E-06 |
| Apobec1 | 4.21E-10 | 0.560702 | 0.374 | 0.255 | 9.92E-06 |
| Usp25 | 4.46E-10 | 0.568491 | 0.295 | 0.187 | 1.05E-05 |
| Ms4a6c | 5.62E-10 | 1.271012 | 0.206 | 0.123 | 1.32E-05 |
| Hmox2 | 8.11E-10 | 0.689504 | 0.531 | 0.427 | 1.91E-05 |
| Cited2 | 1E-09 | 0.860186 | 0.295 | 0.194 | 2.36E-05 |
| Ms4a6b | 1.45E-09 | 0.724011 | 0.427 | 0.319 | 3.43E-05 |
| Tdrd7 | 1.54E-09 | 0.949458 | 0.222 | 0.133 | 3.63E-05 |
| Lrch3 | 1.58E-09 | 1.067029 | 0.21 | 0.124 | 3.72E-05 |
| Tnfsf13b | 1.89E-09 | 0.620118 | 0.3 | 0.196 | 4.47E-05 |
| Naalad2 | 2.94E-09 | 0.969221 | 0.263 | 0.169 | 6.93E-05 |
| Tmpo | 3.12E-09 | 0.845921 | 0.287 | 0.192 | 7.36E-05 |
| March5 | 3.54E-09 | 0.904252 | 0.231 | 0.144 | 8.35E-05 |
| Ankle2 | 4.86E-09 | 0.822697 | 0.464 | 0.357 | 0.000115 |
| Tapbp | 5.26E-09 | 0.52508 | 0.808 | 0.733 | 0.000124 |
| Tlr3 | 5.48E-09 | 0.959139 | 0.323 | 0.225 | 0.000129 |
| Mb21d1 | 6.4E-09 | 0.619602 | 0.277 | 0.181 | 0.000151 |
| Trim56 | 8.35E-09 | 1.035554 | 0.236 | 0.149 | 0.000197 |
| Ly9 | 8.61E-09 | 0.734642 | 0.203 | 0.122 | 0.000203 |
| Acer3 | 1.19E-08 | 0.626312 | 0.545 | 0.435 | 0.000282 |
| Rplp1 | 1.35E-08 | 0.352315 | 0.97 | 0.956 | 0.000317 |
| Ctsz | 1.41E-08 | 0.260497 | 0.995 | 0.993 | 0.000333 |
| Sat1 | 2.05E-08 | 0.326277 | 0.892 | 0.85 | 0.000483 |
| Gatm | 2.54E-08 | 0.616363 | 0.358 | 0.255 | 0.000599 |
| Tap2 | 2.62E-08 | 0.612024 | 0.437 | 0.333 | 0.000618 |
| Gm13456 | 2.67E-08 | 0.254465 | 0.972 | 0.946 | 0.00063 |
| Npc2 | 2.74E-08 | 0.352755 | 0.873 | 0.839 | 0.000646 |
| Gm26825 | 3.19E-08 | 0.545788 | 0.751 | 0.701 | 0.000752 |
| Rpsa | 4.34E-08 | 0.38276 | 0.857 | 0.798 | 0.001024 |
| H2-M3 | 5.18E-08 | 0.606899 | 0.31 | 0.217 | 0.001222 |
| Zfas1 | 5.52E-08 | 0.490939 | 0.238 | 0.154 | 0.001301 |
| Fam111a | 1.28E-07 | 0.858798 | 0.333 | 0.245 | 0.003019 |
| Synj1 | 1.47E-07 | 0.777465 | 0.372 | 0.278 | 0.003472 |
| Ptma | 1.5E-07 | 0.493644 | 0.698 | 0.629 | 0.003549 |
| Atp6v0a2 | 1.51E-07 | 0.675391 | 0.49 | 0.393 | 0.00356 |
| Rnpep | 1.51E-07 | 0.402211 | 0.561 | 0.45 | 0.003564 |
| Tmsb10 | 1.66E-07 | 0.490812 | 0.522 | 0.433 | 0.003909 |
| Nsd3 | 1.75E-07 | 0.683107 | 0.524 | 0.423 | 0.004128 |
| Rpl41 | 1.8E-07 | 0.328538 | 0.972 | 0.943 | 0.004251 |
| Rpl24 | 2.03E-07 | 0.392382 | 0.695 | 0.616 | 0.004777 |
| Hspa8 | 2.18E-07 | 0.422429 | 0.937 | 0.928 | 0.005136 |
| Rrbp1 | 2.96E-07 | 0.393544 | 0.938 | 0.912 | 0.006972 |
| Icosl | 3.16E-07 | 0.857459 | 0.399 | 0.308 | 0.007464 |
| Sgk3 | 3.53E-07 | 0.705725 | 0.296 | 0.209 | 0.008328 |
| Dpysl2 | 3.96E-07 | 0.414242 | 0.907 | 0.852 | 0.009336 |

| | | | | | |
|---|---|---|---|---|---|
| Gm11518 | 5.36E-07 | 0.300148 | 0.578 | 0.487 | 0.012644 |
| Morc3 | 5.63E-07 | 0.675433 | 0.409 | 0.311 | 0.013271 |
| Aif1 | 5.65E-07 | 0.310581 | 0.921 | 0.914 | 0.013335 |
| Gm4617 | 5.67E-07 | 0.335176 | 0.365 | 0.275 | 0.013383 |
| Ctsh | 7.08E-07 | 0.272284 | 0.958 | 0.95 | 0.016691 |
| Ppfia4 | 8.33E-07 | 0.453846 | 0.727 | 0.652 | 0.019652 |
| Cd74 | 8.88E-07 | 0.876076 | 0.282 | 0.207 | 0.020953 |
| Gbp7 | 1.08E-06 | 1.581929 | 0.317 | 0.243 | 0.025536 |
| Chmp4b | 1.2E-06 | 0.501179 | 0.233 | 0.156 | 0.028368 |
| Gm47283 | 1.24E-06 | 0.680264 | 0.381 | 0.292 | 0.029216 |
| Stard3 | 1.28E-06 | 0.529971 | 0.524 | 0.43 | 0.030161 |
| Dnaja1 | 1.49E-06 | 0.609159 | 0.434 | 0.349 | 0.035152 |
| Laptm4a | 1.51E-06 | 0.25578 | 0.91 | 0.891 | 0.0356 |
| Rpl14 | 1.89E-06 | 0.347289 | 0.822 | 0.776 | 0.044466 |
| Emp3 | 1.94E-06 | 0.775936 | 0.245 | 0.171 | 0.045815 |

#### Supplementry Table 1. Marker genes for MHCIIM

| Genenames | p_val | avg_log2FC | pct.1 | pct.2 | p_val_adj |
|---|---|---|---|---|---|
| Siglech | 6.13E-71 | 1.220993 | 0.994 | 0.994 | 1.45E-66 |
| Ccl6 | 6.04E-69 | 2.235079 | 0.792 | 0.417 | 1.42E-64 |
| Chst2 | 1.08E-67 | 2.487171 | 0.253 | 0.042 | 2.54E-63 |
| Cox6a2 | 1.7E-64 | 3.101151 | 0.212 | 0.032 | 4.01E-60 |
| Capg | 9.73E-54 | 2.227875 | 0.516 | 0.193 | 2.3E-49 |
| Haao | 1.42E-53 | 2.853306 | 0.24 | 0.047 | 3.34E-49 |
| C1qb | 2.28E-53 | 0.792697 | 1 | 1 | 5.38E-49 |
| Slc2a5 | 6.48E-52 | 1.197582 | 0.971 | 0.858 | 1.53E-47 |
| C1qa | 3.53E-49 | 0.759647 | 1 | 1 | 8.33E-45 |
| H2-Ab1 | 3.22E-46 | 3.357348 | 0.426 | 0.161 | 7.6E-42 |
| Rad51b | 4.37E-45 | 2.680016 | 0.212 | 0.043 | 1.03E-40 |
| Crlf2 | 1.23E-42 | 1.438956 | 0.718 | 0.393 | 2.91E-38 |
| Ctsd | 1.24E-41 | 0.706068 | 1 | 1 | 2.93E-37 |
| Rnase6 | 3.15E-40 | 1.876122 | 0.369 | 0.122 | 7.43E-36 |
| Gapdh | 3.13E-38 | 1.072482 | 0.965 | 0.889 | 7.38E-34 |
| Ly6e | 4.53E-38 | 1.092048 | 0.99 | 0.952 | 1.07E-33 |
| Cd74 | 2.97E-37 | 3.503403 | 0.458 | 0.204 | 7.01E-33 |
| Lsp1 | 1.43E-32 | 1.387377 | 0.635 | 0.342 | 3.37E-28 |
| Laptm5 | 2.86E-29 | 0.594583 | 1 | 1 | 6.76E-25 |
| Sparc | 2.63E-28 | 0.583826 | 0.997 | 0.998 | 6.2E-24 |
| Tspan7 | 9.61E-26 | 1.24753 | 0.712 | 0.503 | 2.27E-21 |
| Selplg | 2.9E-23 | 0.521993 | 1 | 0.998 | 6.84E-19 |
| H2-D1 | 3.48E-23 | 0.578987 | 0.981 | 0.909 | 8.21E-19 |
| Srgn | 3.72E-23 | 1.096046 | 0.696 | 0.458 | 8.77E-19 |
| Bsg | 5.76E-23 | 0.770061 | 0.939 | 0.851 | 1.36E-18 |
| Cst3 | 7.42E-23 | 0.413822 | 1 | 1 | 1.75E-18 |
| Cd164 | 2.96E-22 | 0.892092 | 0.946 | 0.888 | 6.98E-18 |
| Ighm | 6.12E-22 | 0.944935 | 0.817 | 0.645 | 1.44E-17 |
| Mpc1 | 9.09E-22 | 0.935852 | 0.708 | 0.471 | 2.15E-17 |
| Nupr1 | 1.11E-21 | 1.797325 | 0.337 | 0.151 | 2.63E-17 |
| Syngr1 | 2.07E-21 | 0.867793 | 0.901 | 0.816 | 4.88E-17 |
| Syngr2 | 2.76E-20 | 0.815573 | 0.901 | 0.795 | 6.52E-16 |
| Rnaset2a | 1.04E-19 | 0.644513 | 0.968 | 0.934 | 2.46E-15 |
| Fcer1g | 1.19E-19 | 0.499143 | 1 | 0.991 | 2.81E-15 |
| Upk1b | 1.28E-19 | 0.896469 | 0.609 | 0.373 | 3.01E-15 |
| Ly86 | 3.56E-19 | 0.500731 | 0.99 | 0.989 | 8.4E-15 |
| Cyb561a3 | 2.42E-18 | 0.921603 | 0.663 | 0.453 | 5.72E-14 |
| Mpc1-ps | 2.71E-18 | 0.712342 | 0.628 | 0.402 | 6.39E-14 |
| H2-Oa | 7.92E-18 | 1.574545 | 0.353 | 0.175 | 1.87E-13 |
| Pkm | 8.54E-18 | 1.069913 | 0.724 | 0.565 | 2.01E-13 |
| Tgfbi | 8.83E-18 | 1.523732 | 0.401 | 0.216 | 2.08E-13 |
| Ank | 9.13E-18 | 1.29178 | 0.253 | 0.103 | 2.15E-13 |
| C1qc | 2.07E-17 | 0.432359 | 1 | 1 | 4.87E-13 |

| | | | | | |
|---|---|---|---|---|---|
| Ctss | 3.37E-17 | 0.408272 | 1 | 1 | 7.96E-13 |
| Tmsb4x | 3.55E-17 | 0.439711 | 1 | 1 | 8.37E-13 |
| Ccl9 | 9.59E-17 | 1.071717 | 0.545 | 0.358 | 2.26E-12 |
| Rnase4 | 1.32E-16 | 0.61979 | 0.962 | 0.933 | 3.11E-12 |
| Gm5559 | 3.16E-16 | 0.583848 | 0.885 | 0.703 | 7.46E-12 |
| Gm49369 | 4.48E-16 | 0.633126 | 0.84 | 0.77 | 1.06E-11 |
| Ppia | 8.99E-16 | 0.464908 | 0.987 | 0.966 | 2.12E-11 |
| Cd52 | 2.89E-15 | 0.905658 | 0.66 | 0.482 | 6.83E-11 |
| Cd81 | 3.15E-15 | 0.438264 | 1 | 0.999 | 7.43E-11 |
| Sdf4 | 4.27E-15 | 0.894591 | 0.728 | 0.579 | 1.01E-10 |
| Pafah1b3 | 5.03E-15 | 1.511214 | 0.266 | 0.123 | 1.19E-10 |
| Gp9 | 1.03E-14 | 1.31001 | 0.346 | 0.182 | 2.42E-10 |
| Itm2b | 1.86E-14 | 0.341624 | 1 | 1 | 4.4E-10 |
| Tspan3 | 2.32E-14 | 0.936665 | 0.628 | 0.453 | 5.46E-10 |
| Gm10359 | 3.8E-14 | 0.397167 | 0.49 | 0.306 | 8.96E-10 |
| Tmem176b | 5.63E-14 | 0.682617 | 0.862 | 0.783 | 1.33E-09 |
| Gm7336 | 1.94E-13 | 0.418253 | 0.532 | 0.328 | 4.58E-09 |
| Trem2 | 2.81E-13 | 0.485086 | 0.994 | 0.985 | 6.63E-09 |
| Cbfa2t3 | 3.24E-13 | 1.044534 | 0.519 | 0.335 | 7.63E-09 |
| H2-K1 | 3.41E-13 | 0.588311 | 0.901 | 0.824 | 8.05E-09 |
| Mgat1 | 3.5E-13 | 1.049866 | 0.67 | 0.532 | 8.26E-09 |
| Serpine2 | 4.21E-13 | 0.554881 | 0.929 | 0.895 | 9.92E-09 |
| Gm2574 | 4.43E-13 | 0.475042 | 0.593 | 0.427 | 1.05E-08 |
| Sept11 | 4.51E-13 | 1.73052 | 0.221 | 0.1 | 1.06E-08 |
| P2ry12 | 5.45E-13 | 0.403703 | 1 | 0.996 | 1.28E-08 |
| Lat2 | 8.27E-13 | 0.851035 | 0.548 | 0.384 | 1.95E-08 |
| Klhdc8b | 1.3E-12 | 1.374606 | 0.301 | 0.161 | 3.07E-08 |
| Gm12537 | 1.58E-12 | 0.375002 | 0.478 | 0.295 | 3.72E-08 |
| Fkbp2 | 1.6E-12 | 0.820329 | 0.587 | 0.407 | 3.77E-08 |
| Hexa | 1.72E-12 | 0.470142 | 0.974 | 0.966 | 4.07E-08 |
| Clic1 | 3.21E-12 | 0.50551 | 0.891 | 0.838 | 7.56E-08 |
| Hnrnpc | 3.78E-12 | 0.869677 | 0.638 | 0.484 | 8.91E-08 |
| AW112010 | 8.91E-12 | 1.500849 | 0.228 | 0.108 | 2.1E-07 |
| Laptm4b | 1.28E-11 | 1.385859 | 0.327 | 0.192 | 3.02E-07 |
| Arhgdib | 2.02E-11 | 0.720602 | 0.817 | 0.734 | 4.76E-07 |
| Tmed3 | 2.54E-11 | 0.754147 | 0.619 | 0.475 | 5.99E-07 |
| Sdf2l1 | 2.59E-11 | 0.994904 | 0.567 | 0.438 | 6.1E-07 |
| Actb | 4.4E-11 | 0.381578 | 1 | 1 | 1.04E-06 |
| Tmbim6 | 6.67E-11 | 0.48072 | 0.974 | 0.952 | 1.57E-06 |
| Tspan13 | 7.53E-11 | 0.826946 | 0.625 | 0.497 | 1.78E-06 |
| Tmem119 | 2.4E-10 | 0.339707 | 1 | 0.994 | 5.67E-06 |
| Ltc4s | 4.78E-10 | 0.580647 | 0.894 | 0.808 | 1.13E-05 |
| Tyrobp | 6.61E-10 | 0.376929 | 0.994 | 0.995 | 1.56E-05 |
| Pecam1 | 6.7E-10 | 1.941509 | 0.208 | 0.104 | 1.58E-05 |
| Grap | 1.12E-09 | 0.932256 | 0.41 | 0.273 | 2.64E-05 |
| Thrsp | 1.31E-09 | 0.590126 | 0.433 | 0.28 | 3.1E-05 |
| Rps14 | 1.38E-09 | 0.357444 | 0.939 | 0.924 | 3.26E-05 |

| | | | | | |
|---|---|---|---|---|---|
| Gm6560 | 1.49E-09 | 0.408468 | 0.458 | 0.299 | 3.52E-05 |
| Lrba | 1.82E-09 | 0.653605 | 0.609 | 0.462 | 4.3E-05 |
| Tspan9 | 2.48E-09 | 1.42666 | 0.244 | 0.132 | 5.84E-05 |
| Slc16a3 | 3.76E-09 | 1.183859 | 0.308 | 0.186 | 8.87E-05 |
| Pnp | 4.35E-09 | 0.626587 | 0.654 | 0.536 | 0.000103 |
| Gm10076 | 5.46E-09 | 0.418087 | 0.949 | 0.906 | 0.000129 |
| Gm2606 | 7.04E-09 | 0.414971 | 0.269 | 0.152 | 0.000166 |
| Gngt2 | 7.52E-09 | 0.576549 | 0.699 | 0.58 | 0.000177 |
| Ubb | 7.96E-09 | 0.387961 | 0.984 | 0.971 | 0.000188 |
| Rps27rt | 7.96E-09 | 0.3906 | 0.817 | 0.713 | 0.000188 |
| Edem1 | 8.16E-09 | 1.011816 | 0.663 | 0.563 | 0.000193 |
| Atp5a1 | 9.47E-09 | 0.614071 | 0.702 | 0.591 | 0.000223 |
| Nucb1 | 1.04E-08 | 0.697296 | 0.683 | 0.57 | 0.000245 |
| Selenof | 1.36E-08 | 0.524116 | 0.737 | 0.634 | 0.000321 |
| Maged1 | 1.38E-08 | 1.1846 | 0.276 | 0.162 | 0.000325 |
| Arf1 | 1.46E-08 | 0.490465 | 0.788 | 0.73 | 0.000345 |
| Ly6a | 1.56E-08 | 1.71159 | 0.247 | 0.144 | 0.000368 |
| Gnai2 | 1.57E-08 | 0.412324 | 0.965 | 0.944 | 0.00037 |
| Fscn1 | 1.9E-08 | 0.52929 | 0.869 | 0.791 | 0.000448 |
| Ramp1 | 1.96E-08 | 1.251442 | 0.308 | 0.191 | 0.000463 |
| Ssr4 | 1.99E-08 | 0.659048 | 0.715 | 0.626 | 0.000469 |
| Selenow | 2.31E-08 | 1.070453 | 0.304 | 0.192 | 0.000544 |
| Tmem50a | 2.63E-08 | 0.39119 | 0.917 | 0.853 | 0.000621 |
| Cd37 | 2.75E-08 | 0.463675 | 0.897 | 0.862 | 0.000649 |
| Dad1 | 3.04E-08 | 0.579094 | 0.683 | 0.58 | 0.000717 |
| Ndufa1 | 3.07E-08 | 0.772445 | 0.404 | 0.28 | 0.000724 |
| Rhoa | 3.38E-08 | 0.486217 | 0.897 | 0.831 | 0.000798 |
| Ighj1 | 3.84E-08 | 0.479673 | 0.288 | 0.169 | 0.000907 |
| Tpi1 | 3.96E-08 | 1.003728 | 0.413 | 0.296 | 0.000934 |
| Bri3 | 4.34E-08 | 0.598093 | 0.673 | 0.58 | 0.001023 |
| Krtcap2 | 4.58E-08 | 0.745568 | 0.554 | 0.433 | 0.00108 |
| Oaz1 | 4.92E-08 | 0.482667 | 0.821 | 0.771 | 0.00116 |
| Pfn1 | 5.08E-08 | 0.305034 | 0.974 | 0.952 | 0.001199 |
| Trf | 5.48E-08 | 0.34368 | 0.984 | 0.963 | 0.001293 |
| Gm5138 | 6.11E-08 | 0.352444 | 0.599 | 0.445 | 0.001441 |
| Myl6 | 8.27E-08 | 0.565641 | 0.705 | 0.605 | 0.001951 |
| Csmd3 | 8.33E-08 | 0.586514 | 0.51 | 0.374 | 0.001966 |
| Tpbgl | 8.81E-08 | 0.778483 | 0.5 | 0.369 | 0.002078 |
| Psmb1 | 9.15E-08 | 0.642548 | 0.587 | 0.453 | 0.002159 |
| Rpl41 | 1.13E-07 | 0.300875 | 0.968 | 0.944 | 0.002666 |
| Jup | 1.37E-07 | 0.74382 | 0.433 | 0.306 | 0.003233 |
| Cyb5r3 | 1.39E-07 | 1.126771 | 0.279 | 0.17 | 0.003271 |
| H2afz | 1.42E-07 | 0.59632 | 0.615 | 0.485 | 0.003344 |
| Vars | 1.48E-07 | 1.137395 | 0.292 | 0.18 | 0.003496 |
| Bhlhe41 | 1.55E-07 | 0.874583 | 0.837 | 0.782 | 0.003649 |
| Spcs1 | 2.03E-07 | 0.440756 | 0.635 | 0.502 | 0.004791 |
| Os9 | 2.15E-07 | 0.743367 | 0.574 | 0.465 | 0.00507 |

| | | | | | |
|---|---|---|---|---|---|
| Tmem160 | 2.36E-07 | 0.765475 | 0.446 | 0.324 | 0.005577 |
| Rnaset2b | 2.41E-07 | 0.34022 | 0.981 | 0.939 | 0.005676 |
| Rpl3 | 2.7E-07 | 0.375555 | 0.897 | 0.831 | 0.006362 |
| Vps28 | 3.13E-07 | 0.680729 | 0.506 | 0.387 | 0.007382 |
| Tnfsf13b | 3.23E-07 | 0.831097 | 0.311 | 0.198 | 0.007625 |
| Gpx1 | 3.58E-07 | 0.555705 | 0.785 | 0.707 | 0.008439 |
| Ywhah | 3.82E-07 | 0.333739 | 0.917 | 0.899 | 0.009018 |
| Pdia6 | 3.88E-07 | 0.480117 | 0.756 | 0.703 | 0.009153 |
| Tbcb | 4.87E-07 | 0.763914 | 0.401 | 0.285 | 0.011499 |
| Npc2 | 5.01E-07 | 0.361808 | 0.885 | 0.84 | 0.011822 |
| Padi2 | 5.02E-07 | 0.600089 | 0.349 | 0.226 | 0.011838 |
| S100a13 | 5.04E-07 | 0.661601 | 0.253 | 0.152 | 0.011898 |
| Eno1 | 5.07E-07 | 0.497049 | 0.772 | 0.697 | 0.011968 |
| Tubgcp5 | 5.44E-07 | 0.776703 | 0.561 | 0.43 | 0.012829 |
| Naxe | 5.49E-07 | 0.677483 | 0.429 | 0.318 | 0.01295 |
| Csf3r | 5.81E-07 | 0.461073 | 0.881 | 0.836 | 0.013705 |
| Commd8 | 6.1E-07 | 0.459662 | 0.763 | 0.644 | 0.01439 |
| Hnrnpa1 | 6.2E-07 | 0.613792 | 0.413 | 0.286 | 0.014614 |
| Elob | 6.64E-07 | 0.604767 | 0.59 | 0.469 | 0.015668 |
| Atp5h | 6.65E-07 | 0.459567 | 0.571 | 0.441 | 0.015694 |
| Eef1a1 | 7.38E-07 | 0.329245 | 0.994 | 0.994 | 0.017406 |
| Tspo | 1.08E-06 | 0.452581 | 0.449 | 0.325 | 0.025429 |
| Dnajc3 | 1.72E-06 | 0.628159 | 0.564 | 0.461 | 0.040491 |
| Gm10123 | 1.8E-06 | 0.300626 | 0.946 | 0.925 | 0.042552 |
| Cd53 | 1.91E-06 | 0.344337 | 0.962 | 0.953 | 0.045117 |
| Vkorc1 | 1.96E-06 | 0.625952 | 0.513 | 0.403 | 0.046246 |
| Tfg | 1.97E-06 | 0.582547 | 0.375 | 0.257 | 0.046377 |
| Rpsa | 2.04E-06 | 0.299737 | 0.869 | 0.799 | 0.048117 |
| Rpl13a | 2.07E-06 | 0.398589 | 0.881 | 0.833 | 0.048748 |

### Supplementry Table 1. Marker genes for CycM

| Genenames | p_val | avg_log2FC | pct.1 | pct.2 | p_val_adj |
|---|---|---|---|---|---|
| Mki67 | 0 | 7.657051 | 0.625 | 0.009 | 0 |
| Nuf2 | 0 | 7.306449 | 0.475 | 0.003 | 0 |
| Top2a | 4.8E-241 | 7.970524 | 0.575 | 0.01 | 1.1E-236 |
| Knstrn | 4.9E-234 | 6.163943 | 0.425 | 0.005 | 1.2E-229 |
| Cdca8 | 1.4E-226 | 6.466144 | 0.475 | 0.007 | 3.3E-222 |
| Kifc1 | 2.3E-207 | 6.793222 | 0.4 | 0.005 | 5.4E-203 |
| Racgap1 | 1.2E-205 | 7.454993 | 0.525 | 0.01 | 2.8E-201 |
| Ckap2l | 2.1E-204 | 6.027543 | 0.575 | 0.012 | 4.8E-200 |
| Sgo2a | 3.4E-196 | 4.81287 | 0.25 | 0.002 | 8E-192 |
| Ccna2 | 3.5E-194 | 7.42107 | 0.575 | 0.013 | 8.2E-190 |
| Nusap1 | 2.6E-193 | 5.737882 | 0.45 | 0.008 | 6.1E-189 |
| Foxm1 | 3E-184 | 4.908547 | 0.4 | 0.006 | 7.2E-180 |
| Birc5 | 3.7E-181 | 6.637671 | 0.4 | 0.006 | 8.7E-177 |
| Melk | 3.2E-171 | 5.579418 | 0.2 | 0.001 | 7.6E-167 |
| Prr11 | 6E-168 | 4.591871 | 0.3 | 0.004 | 1.4E-163 |
| Arhgef39 | 2.4E-166 | 5.578434 | 0.45 | 0.009 | 5.6E-162 |
| Prc1 | 8.1E-160 | 6.33607 | 0.425 | 0.008 | 1.9E-155 |
| Cenpe | 9.2E-160 | 6.136417 | 0.425 | 0.008 | 2.2E-155 |
| Fignl1 | 4.4E-156 | 4.709061 | 0.35 | 0.006 | 1E-151 |
| Ndc80 | 5.3E-153 | 6.518448 | 0.4 | 0.008 | 1.3E-148 |
| Lockd | 3E-149 | 3.677916 | 0.275 | 0.004 | 7E-145 |
| Ncapg | 3.3E-149 | 5.675667 | 0.35 | 0.006 | 7.8E-145 |
| Anln | 9.8E-148 | 4.948492 | 0.375 | 0.007 | 2.3E-143 |
| Cip2a | 7.3E-146 | 3.772424 | 0.275 | 0.004 | 1.7E-141 |
| Ccnb2 | 1.4E-143 | 7.110082 | 0.45 | 0.011 | 3.4E-139 |
| Cenpa | 2.5E-143 | 6.082595 | 0.575 | 0.018 | 6E-139 |
| Bub1b | 4.7E-142 | 5.625259 | 0.325 | 0.005 | 1.1E-137 |
| Ncapd2 | 5.3E-141 | 5.613296 | 0.525 | 0.015 | 1.2E-136 |
| Kif23 | 1.4E-140 | 6.212152 | 0.325 | 0.005 | 3.3E-136 |
| 4930579G2 | 1E-139 | 4.826071 | 0.375 | 0.007 | 2.4E-135 |
| Kif20a | 4.2E-132 | 6.196681 | 0.325 | 0.006 | 1E-127 |
| Dlgap5 | 4.9E-128 | 5.844146 | 0.25 | 0.003 | 1.2E-123 |
| Knl1 | 1.4E-125 | 5.866465 | 0.275 | 0.004 | 3.3E-121 |
| Cenpf | 3.1E-125 | 6.107599 | 0.55 | 0.019 | 7.3E-121 |
| Plk1 | 2.6E-123 | 7.233016 | 0.25 | 0.004 | 6.1E-119 |
| Esco2 | 3.6E-122 | 5.076316 | 0.2 | 0.002 | 8.6E-118 |
| Aspm | 8E-121 | 4.457927 | 0.25 | 0.004 | 1.9E-116 |
| Cdca2 | 2.5E-116 | 5.485154 | 0.3 | 0.006 | 5.9E-112 |
| Cenpk | 1.9E-112 | 4.359666 | 0.25 | 0.004 | 4.5E-108 |
| Tpx2 | 3.9E-110 | 5.167709 | 0.225 | 0.003 | 9.2E-106 |
| Cdc20 | 4.3E-109 | 7.276828 | 0.35 | 0.009 | 1E-104 |
| Gm11223 | 5E-109 | 4.926839 | 0.875 | 0.061 | 1.2E-104 |
| Ccnf | 8.5E-108 | 6.471246 | 0.225 | 0.003 | 2E-103 |

| | | | | | |
|---|---|---|---|---|---|
| Pbk | 1E-106 | 6.78304 | 0.2 | 0.003 | 2.3E-102 |
| Clspn | 1.4E-105 | 5.717431 | 0.25 | 0.004 | 3.2E-101 |
| Kif22 | 4.5E-105 | 6.052334 | 0.375 | 0.01 | 1.1E-100 |
| Rad51ap1 | 1.7E-103 | 4.685063 | 0.325 | 0.008 | 4.1E-99 |
| Cdca3 | 1.2E-101 | 5.760851 | 0.275 | 0.006 | 2.83E-97 |
| Kif2c | 6.5E-100 | 4.3512 | 0.25 | 0.005 | 1.53E-95 |
| Tacc3 | 1.52E-98 | 5.076864 | 0.5 | 0.02 | 3.58E-94 |
| Hmmr | 8.92E-98 | 5.070143 | 0.225 | 0.004 | 2.1E-93 |
| Cdk1 | 3.03E-96 | 6.005713 | 0.475 | 0.019 | 7.16E-92 |
| Gen1 | 8.7E-95 | 4.653229 | 0.35 | 0.01 | 2.05E-90 |
| Trim59 | 1.08E-90 | 4.515692 | 0.25 | 0.005 | 2.54E-86 |
| Cit | 1.85E-90 | 5.252338 | 0.375 | 0.012 | 4.38E-86 |
| Rad51 | 1.37E-88 | 5.275682 | 0.3 | 0.008 | 3.23E-84 |
| Mis18bp1 | 1.92E-88 | 5.733982 | 0.25 | 0.005 | 4.54E-84 |
| Lmnb1 | 8.91E-84 | 5.313049 | 0.525 | 0.026 | 2.1E-79 |
| Spc25 | 6.61E-83 | 4.686505 | 0.3 | 0.008 | 1.56E-78 |
| Aurkb | 1.96E-81 | 5.237103 | 0.25 | 0.006 | 4.63E-77 |
| Aurka | 1.67E-80 | 5.232009 | 0.275 | 0.007 | 3.93E-76 |
| Mybl1 | 3.72E-80 | 1.785653 | 0.2 | 0.004 | 8.78E-76 |
| Kif20b | 4.65E-79 | 4.776262 | 0.275 | 0.007 | 1.1E-74 |
| Spc24 | 1.3E-78 | 6.363706 | 0.375 | 0.014 | 3.06E-74 |
| Tcf19 | 1.05E-77 | 6.525636 | 0.3 | 0.009 | 2.48E-73 |
| Mad2l1 | 2.14E-74 | 4.358068 | 0.45 | 0.021 | 5.05E-70 |
| Stmn1 | 5.94E-72 | 5.564543 | 0.9 | 0.102 | 1.4E-67 |
| Kifc5b | 2.47E-67 | 3.471311 | 0.35 | 0.014 | 5.82E-63 |
| Asf1b | 9.5E-66 | 5.292908 | 0.375 | 0.017 | 2.24E-61 |
| Gm4739 | 1.11E-65 | 2.281156 | 0.775 | 0.074 | 2.61E-61 |
| Cks1b | 2.67E-65 | 4.433289 | 0.5 | 0.03 | 6.3E-61 |
| Cdkn2c | 3.31E-65 | 5.088412 | 0.5 | 0.031 | 7.81E-61 |
| Kif15 | 2.22E-64 | 5.236206 | 0.375 | 0.017 | 5.24E-60 |
| Mastl | 3.07E-61 | 3.964582 | 0.275 | 0.01 | 7.24E-57 |
| H2afx | 6.56E-60 | 5.079853 | 0.675 | 0.063 | 1.55E-55 |
| Gm4316 | 3.46E-59 | 3.244736 | 0.425 | 0.025 | 8.16E-55 |
| Cks2 | 3.12E-55 | 4.231665 | 0.6 | 0.053 | 7.36E-51 |
| Zgrf1 | 7.83E-54 | 3.133941 | 0.275 | 0.011 | 1.85E-49 |
| Pclaf | 9.68E-54 | 4.715168 | 0.375 | 0.021 | 2.28E-49 |
| Tk1 | 3.1E-53 | 5.398524 | 0.55 | 0.046 | 7.32E-49 |
| Dtl | 2.15E-51 | 2.777256 | 0.2 | 0.006 | 5.06E-47 |
| Ccne2 | 6.54E-51 | 4.941077 | 0.325 | 0.016 | 1.54E-46 |
| Gm42047 | 2.12E-49 | 4.528543 | 0.45 | 0.033 | 5.01E-45 |
| Gm15452 | 2.62E-49 | 0.434371 | 0.25 | 0.01 | 6.18E-45 |
| Cdkn1a | 3.18E-49 | 5.425426 | 0.825 | 0.125 | 7.51E-45 |
| Smc2 | 4.35E-49 | 4.320552 | 0.5 | 0.04 | 1.03E-44 |
| Ncapg2 | 1.96E-46 | 3.586534 | 0.275 | 0.013 | 4.63E-42 |
| Nsl1 | 1.51E-44 | 3.230953 | 0.375 | 0.024 | 3.56E-40 |
| Uhrf1 | 1.67E-44 | 4.782091 | 0.2 | 0.007 | 3.93E-40 |
| Incenp | 5.19E-44 | 4.99992 | 0.4 | 0.029 | 1.23E-39 |

| | | | | | |
|---|---|---|---|---|---|
| Hist1h2ab | 9.93E-44 | 3.759696 | 0.325 | 0.019 | 2.34E-39 |
| Lmnb2 | 3.73E-43 | 4.25324 | 0.375 | 0.026 | 8.81E-39 |
| Tubb4b-ps: | 2.01E-42 | 1.653998 | 0.675 | 0.084 | 4.74E-38 |
| Smc4 | 4.29E-41 | 4.94446 | 0.55 | 0.06 | 1.01E-36 |
| Atad2 | 6.42E-41 | 4.279569 | 0.275 | 0.015 | 1.51E-36 |
| Bora | 1.05E-39 | 4.013883 | 0.3 | 0.018 | 2.49E-35 |
| Slc43a3 | 3.83E-39 | 5.172658 | 0.45 | 0.041 | 9.04E-35 |
| Gm19945 | 5.16E-39 | 0.434718 | 0.2 | 0.008 | 1.22E-34 |
| Hmgb2 | 5.47E-38 | 4.490704 | 0.875 | 0.189 | 1.29E-33 |
| Ezh2 | 4.35E-37 | 4.240688 | 0.375 | 0.03 | 1.03E-32 |
| Hist2h2ac | 4.96E-37 | 3.985813 | 0.375 | 0.03 | 1.17E-32 |
| Cchcr1 | 2.7E-36 | 3.532809 | 0.225 | 0.011 | 6.38E-32 |
| Prim1 | 5.39E-36 | 3.906911 | 0.275 | 0.017 | 1.27E-31 |
| Fbxo5 | 2.73E-34 | 5.141373 | 0.225 | 0.012 | 6.45E-30 |
| Gas2l3 | 1.43E-33 | 4.216682 | 0.3 | 0.021 | 3.37E-29 |
| Gm42427 | 1.48E-33 | 0.693419 | 0.4 | 0.037 | 3.48E-29 |
| Hspb6 | 2.42E-33 | 3.973755 | 0.25 | 0.015 | 5.7E-29 |
| Gm6750 | 2.91E-33 | 1.220944 | 0.525 | 0.064 | 6.87E-29 |
| Gm12919 | 2.12E-31 | 0.520939 | 0.275 | 0.019 | 5.01E-27 |
| Nt5dc2 | 3.34E-31 | 3.572682 | 0.5 | 0.062 | 7.87E-27 |
| Wdhd1 | 6.46E-31 | 3.382436 | 0.3 | 0.023 | 1.52E-26 |
| Pmf1 | 8.25E-31 | 3.731867 | 0.425 | 0.045 | 1.95E-26 |
| Hist1h2ae | 1.6E-30 | 3.955603 | 0.425 | 0.047 | 3.78E-26 |
| Ac125028.: | 2.57E-30 | 0.768502 | 0.225 | 0.013 | 6.07E-26 |
| Cbx5 | 2.54E-28 | 2.663599 | 0.6 | 0.098 | 6E-24 |
| Enkd1 | 6.56E-28 | 3.520978 | 0.2 | 0.011 | 1.55E-23 |
| Smtn | 4.71E-27 | 5.067293 | 0.225 | 0.015 | 1.11E-22 |
| Hist1h1d | 1.69E-26 | 3.090178 | 0.3 | 0.026 | 3.98E-22 |
| Ddx39 | 1.69E-26 | 2.843529 | 0.575 | 0.092 | 3.99E-22 |
| Fancd2 | 2.12E-26 | 2.988448 | 0.2 | 0.012 | 5.01E-22 |
| Tubb4b | 4.81E-26 | 3.839738 | 0.775 | 0.204 | 1.14E-21 |
| Kcna3 | 6.44E-26 | 2.687152 | 0.275 | 0.023 | 1.52E-21 |
| Tyms | 7.07E-26 | 3.150477 | 0.525 | 0.08 | 1.67E-21 |
| Ncapd3 | 6.67E-25 | 1.921685 | 0.4 | 0.048 | 1.57E-20 |
| Polh | 1E-24 | 2.089035 | 0.425 | 0.054 | 2.36E-20 |
| Dbf4 | 3.3E-24 | 3.807476 | 0.275 | 0.025 | 7.8E-20 |
| Gm20628 | 3.55E-24 | 1.579533 | 0.2 | 0.013 | 8.37E-20 |
| H2afv | 3.77E-24 | 2.860899 | 0.75 | 0.18 | 8.9E-20 |
| Hmgn2-ps1 | 4.87E-24 | 1.326486 | 0.625 | 0.12 | 1.15E-19 |
| Rnf26 | 7.57E-24 | 2.005278 | 0.55 | 0.09 | 1.78E-19 |
| Cdkn2d | 1.27E-23 | 3.549774 | 0.375 | 0.046 | 2.99E-19 |
| Prdx4 | 1.49E-23 | 2.382573 | 0.8 | 0.195 | 3.52E-19 |
| Gm6682 | 1.84E-23 | 1.629609 | 0.6 | 0.119 | 4.35E-19 |
| Mcm7 | 5.3E-23 | 3.147045 | 0.425 | 0.059 | 1.25E-18 |
| Haspin | 7.83E-23 | 3.087642 | 0.4 | 0.052 | 1.85E-18 |
| Lin9 | 8.69E-23 | 2.311559 | 0.425 | 0.06 | 2.05E-18 |
| Rrm1 | 9.11E-23 | 3.52657 | 0.425 | 0.061 | 2.15E-18 |

| | | | | | |
|---|---|---|---|---|---|
| Haus5 | 1.3E-22 | 3.32177 | 0.225 | 0.018 | 3.08E-18 |
| Gm10282 | 1.51E-22 | 2.477907 | 0.65 | 0.142 | 3.56E-18 |
| Atad5 | 1.81E-22 | 2.987534 | 0.25 | 0.022 | 4.26E-18 |
| Hist1h2ap | 3.22E-22 | 2.066427 | 0.425 | 0.064 | 7.59E-18 |
| Mcm2 | 4.06E-22 | 2.879374 | 0.275 | 0.026 | 9.58E-18 |
| Kntc1 | 4.52E-22 | 2.052335 | 0.275 | 0.026 | 1.07E-17 |
| Gm10182 | 8.66E-22 | 0.329386 | 0.25 | 0.022 | 2.04E-17 |
| Hmgn2 | 8.99E-22 | 3.029036 | 0.925 | 0.356 | 2.12E-17 |
| Suv39h1 | 1.75E-21 | 3.233091 | 0.35 | 0.043 | 4.12E-17 |
| Rdm1 | 2.1E-21 | 3.03344 | 0.475 | 0.079 | 4.96E-17 |
| Rfwd3 | 3.35E-21 | 3.451552 | 0.5 | 0.089 | 7.91E-17 |
| Tmpo | 4.99E-21 | 3.102685 | 0.725 | 0.195 | 1.18E-16 |
| Ska2 | 5.4E-21 | 2.960103 | 0.25 | 0.023 | 1.27E-16 |
| Kpna2 | 6.27E-21 | 3.510787 | 0.55 | 0.107 | 1.48E-16 |
| Fen1 | 6.39E-21 | 2.862331 | 0.375 | 0.05 | 1.51E-16 |
| Ptgr1 | 6.9E-21 | 2.242095 | 0.425 | 0.062 | 1.63E-16 |
| Tubb4b-ps: | 1.03E-20 | 0.429871 | 0.2 | 0.015 | 2.42E-16 |
| Cenpl | 5.06E-20 | 2.932315 | 0.275 | 0.029 | 1.19E-15 |
| Rgs12 | 5.22E-20 | 2.636002 | 0.25 | 0.024 | 1.23E-15 |
| Cenpt | 1.01E-19 | 2.379583 | 0.4 | 0.058 | 2.38E-15 |
| Cntrob | 1.2E-19 | 1.897048 | 0.25 | 0.024 | 2.82E-15 |
| Cmc2 | 1.23E-19 | 2.945701 | 0.325 | 0.04 | 2.9E-15 |
| Tex30 | 1.38E-19 | 2.620549 | 0.35 | 0.046 | 3.26E-15 |
| Gm7931 | 1.45E-19 | 1.571545 | 0.625 | 0.148 | 3.42E-15 |
| Mcm5 | 1.96E-19 | 3.024326 | 0.275 | 0.03 | 4.63E-15 |
| Gm10184 | 2.3E-19 | 1.330371 | 0.375 | 0.055 | 5.43E-15 |
| Selenoh | 4.22E-19 | 2.360787 | 0.425 | 0.068 | 9.94E-15 |
| Nemp1 | 6.13E-19 | 2.199106 | 0.3 | 0.036 | 1.45E-14 |
| Ramp1 | 7.99E-19 | 2.887369 | 0.725 | 0.193 | 1.88E-14 |
| Hist1h2ao | 8.89E-19 | 3.518566 | 0.525 | 0.115 | 2.1E-14 |
| Rangap1 | 1.11E-18 | 2.778816 | 0.5 | 0.097 | 2.63E-14 |
| Ube2s | 1.3E-18 | 2.920744 | 0.65 | 0.167 | 3.07E-14 |
| Cep57l1 | 1.98E-18 | 2.05203 | 0.4 | 0.062 | 4.68E-14 |
| Nde1 | 2.03E-18 | 2.273307 | 0.575 | 0.124 | 4.8E-14 |
| Nptxr | 5.72E-18 | 2.288821 | 0.225 | 0.022 | 1.35E-13 |
| Dnajc9 | 6.47E-18 | 3.713121 | 0.4 | 0.066 | 1.53E-13 |
| Tagln2 | 9.2E-18 | 1.992075 | 0.775 | 0.244 | 2.17E-13 |
| Sdk1 | 1.15E-17 | 2.047952 | 0.35 | 0.051 | 2.72E-13 |
| Slc20a1 | 1.42E-17 | 2.610661 | 0.3 | 0.039 | 3.35E-13 |
| Gm3226 | 2.5E-17 | 1.138842 | 0.675 | 0.186 | 5.91E-13 |
| Zfp367 | 2.75E-17 | 3.206583 | 0.35 | 0.053 | 6.48E-13 |
| Rbl1 | 3.04E-17 | 2.830963 | 0.35 | 0.053 | 7.17E-13 |
| Haus3 | 3.21E-17 | 2.404166 | 0.35 | 0.052 | 7.58E-13 |
| Sae1 | 3.27E-17 | 2.459189 | 0.525 | 0.113 | 7.71E-13 |
| AI506816 | 4.27E-17 | 2.409093 | 0.8 | 0.26 | 1.01E-12 |
| Wdr76 | 8.68E-17 | 1.948366 | 0.35 | 0.054 | 2.05E-12 |
| E2f6 | 1.38E-16 | 2.353171 | 0.25 | 0.029 | 3.25E-12 |

| | | | | | |
|---|---|---|---|---|---|
| Reep4 | 3.34E-16 | 2.564207 | 0.45 | 0.088 | 7.88E-12 |
| Nup62 | 4.47E-16 | 2.387693 | 0.55 | 0.126 | 1.05E-11 |
| Cdk5rap2 | 9.99E-16 | 2.245949 | 0.35 | 0.056 | 2.36E-11 |
| Gm10143 | 1.26E-15 | 2.253758 | 0.2 | 0.02 | 2.98E-11 |
| Hdgf | 1.3E-15 | 1.736381 | 0.7 | 0.2 | 3.06E-11 |
| Alg8 | 2.07E-15 | 1.554696 | 0.625 | 0.157 | 4.89E-11 |
| H2afz | 2.17E-15 | 2.628635 | 0.9 | 0.487 | 5.11E-11 |
| Lamb2 | 2.19E-15 | 2.719723 | 0.325 | 0.052 | 5.16E-11 |
| Rfc5 | 2.71E-15 | 2.60765 | 0.275 | 0.037 | 6.39E-11 |
| Cep89 | 3.11E-15 | 3.283615 | 0.225 | 0.026 | 7.33E-11 |
| Gm14150 | 3.62E-15 | 1.095976 | 0.775 | 0.282 | 8.54E-11 |
| 1700025G( | 4.15E-15 | 2.181416 | 0.225 | 0.026 | 9.8E-11 |
| Aaas | 7.99E-15 | 2.160923 | 0.35 | 0.059 | 1.88E-10 |
| Arhgap19 | 1.01E-14 | 1.827267 | 0.25 | 0.032 | 2.39E-10 |
| Nup210 | 1.16E-14 | 4.018247 | 0.225 | 0.027 | 2.73E-10 |
| Poc1a | 1.8E-14 | 3.627352 | 0.225 | 0.027 | 4.24E-10 |
| Jpt2 | 2.88E-14 | 1.959405 | 0.425 | 0.087 | 6.8E-10 |
| Brip1os | 3.22E-14 | 2.248141 | 0.6 | 0.169 | 7.59E-10 |
| Dynlt1a | 3.53E-14 | 3.713324 | 0.25 | 0.034 | 8.34E-10 |
| Tuba1c | 3.55E-14 | 3.632813 | 0.8 | 0.384 | 8.38E-10 |
| Hyls1 | 4.72E-14 | 3.407345 | 0.225 | 0.028 | 1.11E-09 |
| Pomt1 | 4.82E-14 | 2.128906 | 0.425 | 0.088 | 1.14E-09 |
| 4930503L1 | 5.9E-14 | 2.921242 | 0.2 | 0.022 | 1.39E-09 |
| Gm10357 | 5.93E-14 | 0.447249 | 0.225 | 0.028 | 1.4E-09 |
| Ckap5 | 7.49E-14 | 2.339201 | 0.5 | 0.122 | 1.77E-09 |
| Usp1 | 1.1E-13 | 2.76611 | 0.375 | 0.073 | 2.59E-09 |
| Gm13577 | 1.24E-13 | 1.113894 | 0.575 | 0.152 | 2.93E-09 |
| Tuba1b | 1.45E-13 | 2.049399 | 0.975 | 0.752 | 3.43E-09 |
| Pidd1 | 2E-13 | 3.127994 | 0.2 | 0.023 | 4.71E-09 |
| Dna2 | 2.15E-13 | 2.669943 | 0.2 | 0.023 | 5.07E-09 |
| Ncaph | 2.55E-13 | 1.676353 | 0.425 | 0.089 | 6.01E-09 |
| Gm17383 | 3.07E-13 | 2.261995 | 0.925 | 0.53 | 7.25E-09 |
| Jade1 | 5.29E-13 | 2.23053 | 0.375 | 0.074 | 1.25E-08 |
| Tfdp1 | 5.59E-13 | 2.865644 | 0.35 | 0.068 | 1.32E-08 |
| Anp32b | 6.36E-13 | 1.925307 | 0.7 | 0.243 | 1.5E-08 |
| Hjurp | 6.78E-13 | 2.952567 | 0.6 | 0.193 | 1.6E-08 |
| Nucks1 | 6.99E-13 | 1.745795 | 0.75 | 0.278 | 1.65E-08 |
| Rcc1 | 7.64E-13 | 2.957965 | 0.25 | 0.037 | 1.8E-08 |
| Gm13461 | 8.49E-13 | 0.872863 | 0.575 | 0.168 | 2E-08 |
| Lmf2 | 1.21E-12 | 2.500566 | 0.6 | 0.192 | 2.85E-08 |
| Nup107 | 1.36E-12 | 2.026139 | 0.275 | 0.044 | 3.22E-08 |
| Rbmx2 | 1.39E-12 | 1.432583 | 0.2 | 0.024 | 3.28E-08 |
| Arl6ip1 | 1.44E-12 | 2.228523 | 0.95 | 0.736 | 3.4E-08 |
| Calm3 | 2.11E-12 | 1.585732 | 0.725 | 0.239 | 4.97E-08 |
| Orc6 | 2.29E-12 | 3.076625 | 0.275 | 0.045 | 5.4E-08 |
| Lbr | 2.36E-12 | 1.905002 | 0.575 | 0.169 | 5.56E-08 |
| Hirip3 | 2.68E-12 | 2.956131 | 0.2 | 0.025 | 6.31E-08 |

| | | | | | |
|---|---|---|---|---|---|
| Arhgap11a | 2.93E-12 | 0.614338 | 0.275 | 0.044 | 6.9E-08 |
| Anp32e | 3.68E-12 | 1.501623 | 0.6 | 0.175 | 8.69E-08 |
| Cln6 | 3.9E-12 | 2.23299 | 0.35 | 0.07 | 9.21E-08 |
| Gm5620 | 4.51E-12 | 1.185908 | 0.625 | 0.214 | 1.06E-07 |
| Rad54l | 4.66E-12 | 3.315271 | 0.275 | 0.048 | 1.1E-07 |
| Hmgb1-ps7 | 8.32E-12 | 1.140014 | 0.475 | 0.124 | 1.96E-07 |
| Mis12 | 8.63E-12 | 1.492187 | 0.4 | 0.089 | 2.04E-07 |
| Ncaph2 | 1.06E-11 | 1.894209 | 0.65 | 0.242 | 2.5E-07 |
| Dnmt1 | 1.06E-11 | 2.56222 | 0.325 | 0.065 | 2.51E-07 |
| Rad21 | 1.36E-11 | 1.133688 | 0.5 | 0.137 | 3.2E-07 |
| Ran | 1.77E-11 | 1.734376 | 0.825 | 0.373 | 4.18E-07 |
| G2e3 | 1.87E-11 | 2.160822 | 0.3 | 0.056 | 4.4E-07 |
| Ndc1 | 2.21E-11 | 2.394183 | 0.2 | 0.027 | 5.21E-07 |
| Hmgb1-ps2 | 2.84E-11 | 0.79912 | 0.575 | 0.179 | 6.7E-07 |
| Kmt5a | 3.24E-11 | 1.810996 | 0.475 | 0.123 | 7.65E-07 |
| Pgp | 3.32E-11 | 2.263373 | 0.55 | 0.179 | 7.82E-07 |
| Gm3756 | 3.56E-11 | 0.632764 | 0.575 | 0.187 | 8.4E-07 |
| Gm15387 | 3.61E-11 | 0.523701 | 0.45 | 0.115 | 8.51E-07 |
| Ccng1 | 3.73E-11 | 1.446617 | 0.45 | 0.115 | 8.8E-07 |
| Fbxo34 | 3.81E-11 | 2.137471 | 0.35 | 0.077 | 8.99E-07 |
| Pold1 | 5.19E-11 | 3.73579 | 0.225 | 0.035 | 1.22E-06 |
| Dck | 6.45E-11 | 1.887951 | 0.35 | 0.076 | 1.52E-06 |
| Mast2 | 7.44E-11 | 2.549721 | 0.325 | 0.068 | 1.76E-06 |
| Dbr1 | 7.94E-11 | 1.956463 | 0.275 | 0.049 | 1.87E-06 |
| Ccdc34 | 9.08E-11 | 1.66785 | 0.3 | 0.058 | 2.14E-06 |
| Hmgb1-ps1 | 9.13E-11 | 0.315909 | 0.275 | 0.05 | 2.15E-06 |
| Htra3 | 1.04E-10 | 2.430424 | 0.35 | 0.08 | 2.45E-06 |
| Chac2 | 1.13E-10 | 1.40238 | 0.2 | 0.028 | 2.68E-06 |
| Gm21596 | 1.22E-10 | 1.036273 | 0.7 | 0.259 | 2.89E-06 |
| Nudc-ps1 | 1.31E-10 | 0.469214 | 0.425 | 0.104 | 3.09E-06 |
| Trim28 | 1.54E-10 | 1.581597 | 0.725 | 0.291 | 3.64E-06 |
| Glipr1 | 1.58E-10 | 2.285247 | 0.325 | 0.068 | 3.73E-06 |
| Snrpd1 | 1.81E-10 | 1.455572 | 0.525 | 0.154 | 4.26E-06 |
| Cenpq | 2.08E-10 | 2.730498 | 0.225 | 0.036 | 4.92E-06 |
| Cfap43 | 2.16E-10 | 2.676559 | 0.2 | 0.029 | 5.11E-06 |
| Nup188 | 2.48E-10 | 3.039224 | 0.275 | 0.052 | 5.86E-06 |
| Mum1 | 2.58E-10 | 2.246965 | 0.35 | 0.08 | 6.09E-06 |
| Sesn2 | 3.17E-10 | 2.450773 | 0.25 | 0.044 | 7.48E-06 |
| Cdk2 | 3.27E-10 | 2.52165 | 0.35 | 0.083 | 7.72E-06 |
| Smc1a | 3.29E-10 | 1.562783 | 0.625 | 0.217 | 7.77E-06 |
| Raly | 5.09E-10 | 1.43833 | 0.825 | 0.388 | 1.2E-05 |
| Nudc | 5.2E-10 | 1.323377 | 0.725 | 0.296 | 1.23E-05 |
| Sugp2 | 5.45E-10 | 1.70196 | 0.35 | 0.082 | 1.29E-05 |
| Zhx3 | 6.87E-10 | 0.902265 | 0.4 | 0.103 | 1.62E-05 |
| Rbbp8 | 8.01E-10 | 3.372438 | 0.225 | 0.038 | 1.89E-05 |
| Rbm3 | 8.71E-10 | 1.424238 | 0.725 | 0.294 | 2.06E-05 |
| Jpt1 | 9.28E-10 | 1.46321 | 0.825 | 0.403 | 2.19E-05 |

| | | | | | |
|---|---|---|---|---|---|
| Maz | 9.37E-10 | 1.90551 | 0.525 | 0.175 | 2.21E-05 |
| Ppil1 | 9.41E-10 | 2.191394 | 0.25 | 0.046 | 2.22E-05 |
| Rbbp7 | 9.94E-10 | 1.589251 | 0.675 | 0.266 | 2.34E-05 |
| Xpo1 | 1.01E-09 | 1.261209 | 0.4 | 0.105 | 2.37E-05 |
| Pcgf5 | 1.17E-09 | 1.975103 | 0.275 | 0.056 | 2.76E-05 |
| Cbx3 | 1.51E-09 | 1.529908 | 0.8 | 0.392 | 3.56E-05 |
| Dek | 1.57E-09 | 2.236362 | 0.65 | 0.276 | 3.69E-05 |
| Taf9 | 1.87E-09 | 1.229977 | 0.225 | 0.039 | 4.41E-05 |
| Tipin | 2.07E-09 | 2.624708 | 0.275 | 0.056 | 4.89E-05 |
| Tubg1 | 2.29E-09 | 1.909026 | 0.325 | 0.075 | 5.39E-05 |
| Rfc3 | 2.58E-09 | 1.275873 | 0.275 | 0.056 | 6.08E-05 |
| Stom | 2.95E-09 | 3.418643 | 0.2 | 0.034 | 6.96E-05 |
| Calu | 3.06E-09 | 1.452734 | 0.675 | 0.258 | 7.23E-05 |
| Lig1 | 3.15E-09 | 3.380885 | 0.225 | 0.041 | 7.43E-05 |
| Tubb5 | 3.19E-09 | 2.506623 | 0.95 | 0.788 | 7.51E-05 |
| Adgrl1 | 3.19E-09 | 2.621242 | 0.225 | 0.039 | 7.52E-05 |
| Tpm4 | 3.61E-09 | 2.189609 | 0.45 | 0.142 | 8.51E-05 |
| Psat1 | 3.62E-09 | 1.878192 | 0.35 | 0.086 | 8.55E-05 |
| Gm7870 | 3.75E-09 | 0.42038 | 0.325 | 0.074 | 8.84E-05 |
| Nsmce1 | 4.1E-09 | 0.95587 | 0.525 | 0.168 | 9.66E-05 |
| Nemp2 | 4.19E-09 | 0.538005 | 0.225 | 0.039 | 9.88E-05 |
| Cnnm4 | 4.41E-09 | 1.45846 | 0.575 | 0.198 | 0.000104 |
| Gm12346 | 4.54E-09 | 0.927248 | 0.625 | 0.274 | 0.000107 |
| Cnot6 | 4.75E-09 | 2.425803 | 0.35 | 0.088 | 0.000112 |
| Stt3b | 4.85E-09 | 1.871005 | 0.5 | 0.17 | 0.000114 |
| Haus4 | 5.05E-09 | 1.67217 | 0.3 | 0.066 | 0.000119 |
| Gm5844 | 5.43E-09 | 0.625679 | 0.6 | 0.223 | 0.000128 |
| Psip1 | 5.61E-09 | 1.180817 | 0.5 | 0.161 | 0.000132 |
| Myef2 | 5.77E-09 | 1.719053 | 0.425 | 0.126 | 0.000136 |
| Haus1 | 5.8E-09 | 1.983849 | 0.225 | 0.041 | 0.000137 |
| Pcyox1l | 5.85E-09 | 1.628665 | 0.425 | 0.12 | 0.000138 |
| Pla2g4a | 6.46E-09 | 1.069874 | 0.55 | 0.185 | 0.000152 |
| Nsd2 | 6.57E-09 | 2.698265 | 0.375 | 0.105 | 0.000155 |
| Nasp | 6.91E-09 | 0.414476 | 0.3 | 0.067 | 0.000163 |
| Ipo9 | 7.12E-09 | 1.010297 | 0.45 | 0.134 | 0.000168 |
| Eri1 | 7.53E-09 | 2.191235 | 0.35 | 0.091 | 0.000178 |
| Rhno1 | 7.7E-09 | 1.220731 | 0.525 | 0.173 | 0.000182 |
| Hat1 | 7.73E-09 | 1.925568 | 0.375 | 0.102 | 0.000182 |
| Cep135 | 8.28E-09 | 1.442925 | 0.25 | 0.05 | 0.000195 |
| Actn4 | 1.01E-08 | 1.133698 | 0.625 | 0.243 | 0.000239 |
| Ankrd13a | 1.08E-08 | 1.239961 | 0.65 | 0.255 | 0.000256 |
| Pak4 | 1.16E-08 | 1.543882 | 0.25 | 0.05 | 0.000273 |
| Recql | 1.18E-08 | 2.589952 | 0.275 | 0.062 | 0.000279 |
| Stard7 | 1.2E-08 | 1.269752 | 0.4 | 0.111 | 0.000283 |
| Nrm | 1.49E-08 | 1.710245 | 0.575 | 0.219 | 0.000351 |
| Cdc25a | 1.85E-08 | 1.336739 | 0.2 | 0.035 | 0.000437 |
| Hmgn1 | 1.9E-08 | 1.544707 | 0.7 | 0.328 | 0.000448 |

| | | | | | |
|---|---|---|---|---|---|
| Galns | 1.96E-08 | 1.924516 | 0.25 | 0.052 | 0.000463 |
| Nxt2 | 1.98E-08 | 1.397896 | 0.4 | 0.115 | 0.000468 |
| Ssna1 | 2.12E-08 | 1.259111 | 0.525 | 0.18 | 0.0005 |
| Gm9844 | 2.21E-08 | 0.44827 | 0.4 | 0.117 | 0.00052 |
| Uevld | 2.36E-08 | 1.46543 | 0.25 | 0.052 | 0.000556 |
| Fbxo45 | 2.65E-08 | 2.369963 | 0.2 | 0.037 | 0.000625 |
| Tmem129 | 2.82E-08 | 1.071166 | 0.35 | 0.092 | 0.000665 |
| Prpf3 | 2.97E-08 | 2.546303 | 0.3 | 0.073 | 0.000701 |
| Apba1 | 3.09E-08 | 0.70283 | 0.525 | 0.176 | 0.00073 |
| Pcyox1 | 3.22E-08 | 1.159663 | 0.675 | 0.285 | 0.000759 |
| Slc25a25 | 3.47E-08 | 2.690431 | 0.25 | 0.054 | 0.000817 |
| Lsm2 | 4.14E-08 | 1.644784 | 0.4 | 0.121 | 0.000976 |
| Efemp2 | 4.56E-08 | 0.969368 | 0.225 | 0.044 | 0.001076 |
| Gm5518 | 4.95E-08 | 0.708425 | 0.55 | 0.222 | 0.001168 |
| Chuk | 4.99E-08 | 1.133767 | 0.4 | 0.119 | 0.001178 |
| Mcm6 | 5.35E-08 | 1.862533 | 0.25 | 0.054 | 0.001262 |
| Arl4c | 5.35E-08 | 0.943114 | 0.875 | 0.446 | 0.001263 |
| Anapc5 | 5.48E-08 | 1.752723 | 0.6 | 0.256 | 0.001292 |
| Sass6 | 5.89E-08 | 2.271072 | 0.275 | 0.065 | 0.00139 |
| Ranbp3 | 6.12E-08 | 1.308029 | 0.475 | 0.162 | 0.001444 |
| Ifi27l2a | 6.21E-08 | 2.611404 | 0.45 | 0.164 | 0.001464 |
| Gm17201 | 6.46E-08 | 0.563439 | 0.25 | 0.054 | 0.001524 |
| Hsp90aa1 | 7E-08 | 1.734576 | 0.85 | 0.524 | 0.001651 |
| Dtymk | 7.02E-08 | 1.927137 | 0.375 | 0.113 | 0.001655 |
| Hist1h1e | 7.25E-08 | 2.753664 | 0.45 | 0.167 | 0.00171 |
| Alyref | 7.35E-08 | 1.658019 | 0.5 | 0.184 | 0.001733 |
| Ppp2r5d | 7.39E-08 | 1.847123 | 0.4 | 0.121 | 0.001742 |
| Mphosph9 | 7.73E-08 | 1.573225 | 0.225 | 0.045 | 0.001823 |
| Grk6 | 7.82E-08 | 1.800303 | 0.35 | 0.098 | 0.001844 |
| Evi2 | 7.93E-08 | 0.47473 | 0.325 | 0.084 | 0.001871 |
| Pom121 | 8.47E-08 | 0.990359 | 0.6 | 0.239 | 0.001999 |
| Cbfb | 9.02E-08 | 1.012301 | 0.5 | 0.179 | 0.002129 |
| Nap1l1 | 9.58E-08 | 1.658513 | 0.45 | 0.156 | 0.002259 |
| Timm50 | 9.61E-08 | 1.296073 | 0.5 | 0.174 | 0.002268 |
| Cdkn2aipnl | 1.09E-07 | 1.501197 | 0.525 | 0.198 | 0.002562 |
| Miip | 1.09E-07 | 1.89013 | 0.3 | 0.078 | 0.002581 |
| Fkbp2 | 1.1E-07 | 1.194517 | 0.775 | 0.41 | 0.002593 |
| Milr1 | 1.15E-07 | 1.531925 | 0.35 | 0.1 | 0.002723 |
| Cenpb | 1.21E-07 | 1.589461 | 0.4 | 0.126 | 0.002854 |
| Haus8 | 1.26E-07 | 2.025979 | 0.25 | 0.056 | 0.002974 |
| Cenpc1 | 1.32E-07 | 1.70029 | 0.225 | 0.048 | 0.003108 |
| Sfpq | 1.32E-07 | 1.217626 | 0.675 | 0.326 | 0.003119 |
| Tmem134 | 1.34E-07 | 1.167766 | 0.475 | 0.163 | 0.003154 |
| Olfml2b | 1.74E-07 | 1.637708 | 0.4 | 0.124 | 0.004108 |
| Gm16374 | 1.78E-07 | 0.365161 | 0.3 | 0.078 | 0.004202 |
| Tmsb10 | 1.78E-07 | 1.523151 | 0.8 | 0.436 | 0.004208 |
| Clpb | 1.87E-07 | 0.831646 | 0.35 | 0.099 | 0.004421 |

| | | | | | |
|---|---|---|---|---|---|
| Hmgb1 | 1.91E-07 | 1.104665 | 0.95 | 0.739 | 0.004515 |
| Rnaseh2b | 2E-07 | 1.764199 | 0.275 | 0.067 | 0.004721 |
| Pcna-ps2 | 2.02E-07 | 1.156751 | 0.375 | 0.121 | 0.004774 |
| Tmem109 | 2.13E-07 | 0.841984 | 0.85 | 0.425 | 0.005033 |
| Chek2 | 2.15E-07 | 1.437623 | 0.225 | 0.048 | 0.005063 |
| Cdca7 | 2.2E-07 | 1.848699 | 0.3 | 0.081 | 0.005188 |
| Gm4617 | 2.9E-07 | 0.568958 | 0.65 | 0.278 | 0.006831 |
| Cpd | 2.97E-07 | 1.880043 | 0.325 | 0.093 | 0.007015 |
| Gm9800 | 3E-07 | 0.367133 | 0.575 | 0.23 | 0.007078 |
| Ing2 | 3.02E-07 | 1.766426 | 0.3 | 0.079 | 0.007126 |
| Coq2 | 3.12E-07 | 0.905989 | 0.5 | 0.184 | 0.007348 |
| Fam111a | 3.21E-07 | 1.272447 | 0.575 | 0.248 | 0.007573 |
| Ac122335.: | 3.22E-07 | 1.512127 | 0.2 | 0.041 | 0.007591 |
| Zfp131 | 3.85E-07 | 1.059144 | 0.375 | 0.12 | 0.009082 |
| Larp7 | 3.89E-07 | 1.501432 | 0.3 | 0.081 | 0.009185 |
| Fam92a | 3.94E-07 | 2.664948 | 0.2 | 0.041 | 0.009294 |
| H1f0 | 4.02E-07 | 1.541568 | 0.525 | 0.207 | 0.009484 |
| Tbl2 | 4.47E-07 | 1.055961 | 0.35 | 0.105 | 0.010551 |
| Tdp2 | 4.81E-07 | 1.627372 | 0.25 | 0.059 | 0.011342 |
| Eif1ad | 5.01E-07 | 1.081323 | 0.4 | 0.126 | 0.011809 |
| Ruvbl2 | 5.3E-07 | 1.520075 | 0.4 | 0.132 | 0.012507 |
| Abhd2 | 5.84E-07 | 1.392501 | 0.425 | 0.15 | 0.013777 |
| Hist1h2bg | 5.85E-07 | 1.747295 | 0.2 | 0.041 | 0.013802 |
| Dennd2c | 5.89E-07 | 0.532591 | 0.325 | 0.091 | 0.013903 |
| Hmgn3 | 6.2E-07 | 2.42213 | 0.2 | 0.042 | 0.014628 |
| Hist1h4d | 6.41E-07 | 2.723009 | 0.55 | 0.288 | 0.015131 |
| Smarcc1 | 6.54E-07 | 0.992732 | 0.475 | 0.173 | 0.015434 |
| Slc29a1 | 6.98E-07 | 1.689039 | 0.725 | 0.381 | 0.01646 |
| Ipo5 | 7.16E-07 | 0.465886 | 0.375 | 0.117 | 0.01688 |
| Ccdc77 | 7.25E-07 | 2.073091 | 0.2 | 0.042 | 0.017097 |
| Smarca4 | 7.63E-07 | 1.709088 | 0.525 | 0.216 | 0.018003 |
| Ptma | 7.75E-07 | 1.170858 | 0.9 | 0.632 | 0.018271 |
| Men1 | 7.75E-07 | 2.045016 | 0.25 | 0.061 | 0.018283 |
| Kat7 | 7.76E-07 | 0.754714 | 0.55 | 0.213 | 0.018309 |
| Cbx6 | 8.33E-07 | 2.324282 | 0.275 | 0.076 | 0.019645 |
| Hist1h1c | 9.83E-07 | 1.578896 | 0.525 | 0.216 | 0.023183 |
| H2afy | 1.01E-06 | 1.048033 | 0.85 | 0.505 | 0.023911 |
| Srsf7 | 1.03E-06 | 1.161691 | 0.575 | 0.251 | 0.024181 |
| Sf3a2 | 1.03E-06 | 0.324183 | 0.5 | 0.18 | 0.024237 |
| Anapc7 | 1.08E-06 | 1.665975 | 0.4 | 0.133 | 0.025368 |
| Pbdc1 | 1.11E-06 | 1.169125 | 0.3 | 0.083 | 0.026291 |
| tpr | 1.13E-06 | 0.47229 | 0.65 | 0.292 | 0.026602 |
| Hp1bp3 | 1.23E-06 | 0.520031 | 0.85 | 0.431 | 0.028976 |
| Smchd1 | 1.34E-06 | 1.563035 | 0.525 | 0.219 | 0.031565 |
| Nup160 | 1.34E-06 | 1.396257 | 0.225 | 0.054 | 0.03162 |
| Hnrnpab | 1.57E-06 | 1.242194 | 0.875 | 0.531 | 0.036976 |
| Rmi1 | 1.61E-06 | 1.671147 | 0.4 | 0.145 | 0.037972 |

| | | | | | |
|---|---|---|---|---|---|
| Ak6 | 1.64E-06 | 0.903333 | 0.55 | 0.222 | 0.038648 |
| Ccdc88a | 1.66E-06 | 1.255505 | 0.45 | 0.165 | 0.039065 |
| Bex3 | 1.69E-06 | 1.186069 | 0.225 | 0.053 | 0.039942 |
| Kcnk12 | 1.72E-06 | 1.263416 | 0.425 | 0.152 | 0.040582 |
| Gm21399 | 1.75E-06 | 0.650939 | 0.475 | 0.178 | 0.041271 |
| Sgpp1 | 1.77E-06 | 0.917041 | 0.45 | 0.167 | 0.041705 |
| Cnep1r1 | 1.78E-06 | 1.183571 | 0.4 | 0.141 | 0.042093 |
| Slbp | 1.78E-06 | 1.773875 | 0.525 | 0.231 | 0.0421 |
| Tcerg1 | 1.91E-06 | 1.515163 | 0.375 | 0.126 | 0.045104 |
| Sec11c | 2E-06 | 0.703981 | 0.9 | 0.551 | 0.047067 |
| Gapdh | 2.04E-06 | 0.864534 | 1 | 0.891 | 0.048044 |
| Dhx8 | 2.1E-06 | 1.599489 | 0.35 | 0.111 | 0.049508 |

#### Supplementry Table 1. Marker genes for TNFRM

| Genenames | p_val | avg_log2FC | pct.1 | pct.2 | p_val_adj |
|---|---|---|---|---|---|
| Marcksl1 | 1.33E-61 | 4.39226 | 0.554 | 0.086 | 3.14E-57 |
| Cd83 | 2.36E-49 | 3.889189 | 0.87 | 0.311 | 5.56E-45 |
| Vcam1 | 4.7E-40 | 3.937796 | 0.326 | 0.043 | 1.11E-35 |
| Gpr84 | 1.11E-34 | 3.592861 | 0.859 | 0.44 | 2.61E-30 |
| Icam1 | 1.1E-29 | 2.740372 | 0.815 | 0.364 | 2.61E-25 |
| Tlr2 | 2.54E-27 | 2.995078 | 0.783 | 0.362 | 5.98E-23 |
| Relb | 1.67E-26 | 2.293949 | 0.446 | 0.105 | 3.93E-22 |
| Bcl3 | 4.4E-25 | 4.330512 | 0.239 | 0.037 | 1.04E-20 |
| Ccl12 | 8E-25 | 4.071109 | 0.598 | 0.206 | 1.89E-20 |
| Slamf8 | 8.01E-25 | 3.029011 | 0.446 | 0.115 | 1.89E-20 |
| Nfkbia | 4.71E-24 | 2.608087 | 0.848 | 0.474 | 1.11E-19 |
| Nfkbie | 8.42E-24 | 2.643182 | 0.413 | 0.102 | 1.99E-19 |
| Herpud1 | 1.22E-19 | 2.209644 | 0.837 | 0.562 | 2.89E-15 |
| Ccl4 | 1.99E-19 | 4.005062 | 0.207 | 0.035 | 4.69E-15 |
| Cd74 | 8.59E-19 | 2.439102 | 0.554 | 0.208 | 2.03E-14 |
| Tnfaip2 | 2.21E-18 | 3.831356 | 0.337 | 0.087 | 5.21E-14 |
| H2-Q6 | 2.26E-18 | 1.983151 | 0.38 | 0.104 | 5.32E-14 |
| Cxcl16 | 9.8E-18 | 2.592607 | 0.63 | 0.271 | 2.31E-13 |
| Nfkb2 | 1.59E-17 | 3.267085 | 0.391 | 0.114 | 3.74E-13 |
| Hpn | 1.55E-16 | 2.494716 | 0.261 | 0.059 | 3.66E-12 |
| Birc3 | 1.65E-16 | 2.431159 | 0.391 | 0.12 | 3.9E-12 |
| Rab20 | 1.53E-15 | 2.444729 | 0.435 | 0.156 | 3.61E-11 |
| Tnfaip3 | 5.66E-14 | 2.232264 | 0.293 | 0.078 | 1.33E-09 |
| C5ar1 | 1.08E-13 | 2.04583 | 0.663 | 0.412 | 2.54E-09 |
| Rnf19b | 3.71E-13 | 1.579867 | 0.522 | 0.223 | 8.75E-09 |
| Nfkbiz | 4.47E-13 | 3.033928 | 0.315 | 0.096 | 1.06E-08 |
| Bcl2a1a | 5.42E-13 | 1.627158 | 0.446 | 0.171 | 1.28E-08 |
| Arl5c | 6.45E-13 | 3.164676 | 0.337 | 0.113 | 1.52E-08 |
| Sdc4 | 7.7E-13 | 2.64613 | 0.457 | 0.187 | 1.82E-08 |
| H2-Q4 | 1.88E-12 | 3.403031 | 0.315 | 0.101 | 4.44E-08 |
| Ccrl2 | 1E-11 | 2.353545 | 0.217 | 0.055 | 2.36E-07 |
| Slc15a3 | 2.62E-11 | 1.585672 | 0.652 | 0.373 | 6.17E-07 |
| Rrad | 6.56E-11 | 2.460825 | 0.25 | 0.076 | 1.55E-06 |
| Cd52 | 7.84E-11 | 1.300924 | 0.761 | 0.484 | 1.85E-06 |
| Hivep3 | 2.97E-10 | 1.534108 | 0.696 | 0.407 | 7E-06 |
| Bcl6 | 3.8E-10 | 1.407627 | 0.326 | 0.118 | 8.97E-06 |
| Vasp | 4.68E-10 | 1.565322 | 0.63 | 0.36 | 1.1E-05 |
| Mat2a | 5.13E-10 | 1.463992 | 0.511 | 0.239 | 1.21E-05 |
| Plekho2 | 5.71E-10 | 1.810342 | 0.359 | 0.142 | 1.35E-05 |
| H2-K1 | 1.38E-09 | 1.133398 | 0.957 | 0.825 | 3.27E-05 |
| Ier3 | 2.15E-09 | 2.879203 | 0.217 | 0.066 | 5.07E-05 |
| Hcar2 | 2.19E-09 | 2.98893 | 0.217 | 0.066 | 5.17E-05 |
| Csf1 | 2.27E-09 | 2.482801 | 0.217 | 0.066 | 5.37E-05 |

| | | | | | |
|---|---|---|---|---|---|
| Cebpb | 5.48E-09 | 1.289564 | 0.772 | 0.533 | 0.000129 |
| Nfkbib | 1.07E-08 | 1.702927 | 0.402 | 0.182 | 0.000253 |
| Il1a | 1.08E-08 | 1.945324 | 0.402 | 0.184 | 0.000255 |
| Mcl1 | 1.18E-08 | 1.603903 | 0.707 | 0.496 | 0.000278 |
| H2-Q7 | 1.51E-08 | 1.142038 | 0.315 | 0.119 | 0.000355 |
| Ccl3 | 1.9E-08 | 2.114099 | 0.207 | 0.065 | 0.000448 |
| Itga5 | 2.45E-08 | 2.309674 | 0.25 | 0.089 | 0.000577 |
| Stk38l | 4.94E-08 | 2.283015 | 0.217 | 0.073 | 0.001165 |
| Plaur | 5.71E-08 | 1.739749 | 0.228 | 0.077 | 0.001348 |
| Apobec3 | 1.08E-07 | 1.40138 | 0.707 | 0.492 | 0.002544 |
| Fhl3 | 1.94E-07 | 1.23264 | 0.25 | 0.093 | 0.004583 |
| Ifi30 | 2.16E-07 | 1.255206 | 0.739 | 0.498 | 0.005086 |
| Cep85 | 2.98E-07 | 1.476163 | 0.717 | 0.503 | 0.007027 |
| Gm43305 | 3.05E-07 | 2.062425 | 0.75 | 0.581 | 0.007191 |
| Mob3c | 3.19E-07 | 1.578947 | 0.391 | 0.187 | 0.007523 |
| Sqstm1 | 3.33E-07 | 1.042824 | 0.739 | 0.583 | 0.00786 |
| Mvp | 4.78E-07 | 1.474368 | 0.37 | 0.172 | 0.011284 |
| Mthfd2 | 5.12E-07 | 1.521647 | 0.261 | 0.104 | 0.012073 |
| Gm26917 | 5.92E-07 | 0.908026 | 1 | 0.946 | 0.013959 |
| Tnip1 | 6.89E-07 | 1.577847 | 0.207 | 0.072 | 0.016252 |
| Mapkapk2 | 7.05E-07 | 1.645376 | 0.435 | 0.232 | 0.016628 |
| Junb | 7.24E-07 | 1.882321 | 0.652 | 0.498 | 0.017071 |
| Tank | 7.62E-07 | 1.182085 | 0.261 | 0.102 | 0.017964 |
| Hspa5 | 9.92E-07 | 0.969534 | 0.946 | 0.83 | 0.023389 |
| Nadk | 1.16E-06 | 1.85646 | 0.402 | 0.209 | 0.02728 |
| Cacul1 | 1.19E-06 | 1.228158 | 0.25 | 0.097 | 0.028034 |
| Rassf4 | 1.58E-06 | 1.353819 | 0.543 | 0.346 | 0.037335 |

### Supplementry Table 2. IntM DEGs compared to HS

| GeneName | p_val | avg_log2FC | pct.1 | pct.2 | p_val_adj |
|---|---|---|---|---|---|
| Fcrls | 0 | -1.23854 | 0.842 | 0.975 | 0 |
| P2ry12 | 8.2E-235 | -0.69244 | 0.998 | 1 | 1.9E-230 |
| H2-D1 | 1.9E-223 | 1.486782 | 0.947 | 0.867 | 4.4E-219 |
| Slc2a5 | 2.2E-187 | -0.91933 | 0.801 | 0.944 | 5.2E-183 |
| P2ry13 | 4.9E-146 | -0.75731 | 0.915 | 0.977 | 1.2E-141 |
| Cd164 | 9.6E-137 | -0.68232 | 0.847 | 0.946 | 2.3E-132 |
| H2-K1 | 5.4E-126 | 1.274552 | 0.874 | 0.766 | 1.3E-121 |
| Lyz2 | 9.7E-115 | 1.340087 | 0.755 | 0.608 | 2.3E-110 |
| Gpr34 | 2.9E-112 | -0.55565 | 0.978 | 0.992 | 6.9E-108 |
| B2m | 4.4E-109 | 0.719262 | 0.999 | 0.998 | 1E-104 |
| Tmem119 | 2.98E-98 | -0.44204 | 0.997 | 1 | 7.03E-94 |
| Fth1 | 5.94E-95 | 0.656733 | 0.996 | 0.993 | 1.4E-90 |
| Ptgs1 | 7.5E-88 | -0.6365 | 0.886 | 0.946 | 1.77E-83 |
| Upk1b | 2.17E-84 | -1.25455 | 0.278 | 0.473 | 5.12E-80 |
| Hexb | 1.88E-83 | -0.32128 | 1 | 1 | 4.43E-79 |
| Cx3cr1 | 1.19E-82 | -0.34961 | 0.999 | 1 | 2.81E-78 |
| Dusp6 | 6.98E-82 | -0.8844 | 0.586 | 0.731 | 1.65E-77 |
| Cmtm6 | 3.51E-76 | -0.54509 | 0.703 | 0.854 | 8.27E-72 |
| Ctss | 2.74E-75 | 0.43557 | 1 | 1 | 6.47E-71 |
| Selenop | 8.03E-75 | -0.43411 | 0.98 | 0.991 | 1.89E-70 |
| Selplg | 3.06E-72 | -0.34153 | 1 | 1 | 7.22E-68 |
| Adgrg1 | 4.72E-66 | -0.42152 | 0.798 | 0.905 | 1.11E-61 |
| Hspa5 | 3.44E-63 | -0.55537 | 0.778 | 0.866 | 8.11E-59 |
| Rhob | 4.97E-63 | -0.3777 | 0.851 | 0.928 | 1.17E-58 |
| Plxdc2 | 7.54E-63 | -0.47225 | 0.863 | 0.931 | 1.78E-58 |
| Rpl18a | 1.1E-62 | 0.582444 | 0.989 | 0.986 | 2.59E-58 |
| Fscn1 | 1.55E-61 | -0.44398 | 0.711 | 0.859 | 3.65E-57 |
| Ubc | 1.12E-60 | -0.40084 | 0.931 | 0.974 | 2.65E-56 |
| Ftl1 | 1.31E-60 | 0.558514 | 0.996 | 0.991 | 3.08E-56 |
| Sgk1 | 3.59E-59 | -0.95608 | 0.445 | 0.584 | 8.46E-55 |
| Csf1r | 7.94E-59 | -0.32552 | 1 | 1 | 1.87E-54 |
| Hspa8 | 1.98E-58 | -0.42923 | 0.901 | 0.945 | 4.66E-54 |
| Slc7a8 | 1.56E-56 | -0.53414 | 0.565 | 0.723 | 3.68E-52 |
| Rplp1 | 7.69E-56 | 0.604784 | 0.956 | 0.952 | 1.81E-51 |
| Itgb5 | 3.08E-55 | -0.32259 | 0.959 | 0.969 | 7.26E-51 |
| Lrba | 1.15E-54 | -0.82668 | 0.39 | 0.547 | 2.7E-50 |
| Pla2g15 | 1.11E-51 | -0.54088 | 0.708 | 0.803 | 2.63E-47 |
| Rnf13 | 1.67E-50 | -0.53014 | 0.676 | 0.776 | 3.95E-46 |
| Txnip | 4.45E-50 | -0.65537 | 0.583 | 0.699 | 1.05E-45 |
| Hnrnpk | 1.69E-48 | -0.49908 | 0.735 | 0.829 | 3.98E-44 |
| Scoc | 3.58E-48 | -0.40846 | 0.702 | 0.831 | 8.45E-44 |
| Pdia3 | 2.02E-47 | -0.37416 | 0.924 | 0.961 | 4.77E-43 |
| Stab1 | 6.14E-47 | -0.37231 | 0.687 | 0.822 | 1.45E-42 |

| | | | | | |
|---|---|---|---|---|---|
| Malat1 | 1.59E-46 | 0.430693 | 0.984 | 0.992 | 3.74E-42 |
| Gm7964 | 2.85E-46 | -0.51891 | 0.491 | 0.644 | 6.72E-42 |
| Fam102b | 8.95E-46 | -0.6913 | 0.533 | 0.665 | 2.11E-41 |
| Ftl1-ps1 | 8.62E-45 | 0.531199 | 0.897 | 0.874 | 2.03E-40 |
| Rps11 | 2.24E-44 | 0.579932 | 0.915 | 0.911 | 5.28E-40 |
| Lyn | 6.34E-44 | -0.46482 | 0.799 | 0.871 | 1.5E-39 |
| Qk | 7.94E-44 | -0.40945 | 0.872 | 0.936 | 1.87E-39 |
| Gal3st4 | 9.61E-44 | -0.40692 | 0.416 | 0.58 | 2.27E-39 |
| Ubl3 | 1.29E-43 | -0.64838 | 0.585 | 0.691 | 3.05E-39 |
| Rpl37a | 1.95E-43 | 0.52625 | 0.945 | 0.929 | 4.59E-39 |
| Sirpa | 6.21E-43 | -0.35212 | 0.914 | 0.958 | 1.47E-38 |
| Olfml3 | 1.03E-42 | -0.30919 | 0.99 | 0.996 | 2.43E-38 |
| Lpcat2 | 1.19E-42 | -0.27272 | 0.965 | 0.984 | 2.82E-38 |
| Rpl41 | 2.11E-42 | 0.562055 | 0.936 | 0.934 | 4.98E-38 |
| Rpl32 | 2.96E-42 | 0.632672 | 0.895 | 0.863 | 6.98E-38 |
| Gpr165 | 3.96E-42 | -0.93032 | 0.139 | 0.267 | 9.34E-38 |
| Abcg1 | 6.27E-42 | -0.55912 | 0.524 | 0.657 | 1.48E-37 |
| Cyth4 | 1.05E-41 | -0.44298 | 0.837 | 0.894 | 2.48E-37 |
| Rps28 | 1.82E-41 | 0.621193 | 0.921 | 0.892 | 4.3E-37 |
| Marcks | 4.08E-41 | -0.31653 | 0.987 | 0.996 | 9.64E-37 |
| Tmbim6 | 1.59E-40 | -0.31284 | 0.934 | 0.959 | 3.76E-36 |
| Lgals3bp | 2.78E-40 | 0.908913 | 0.636 | 0.503 | 6.57E-36 |
| Capza2 | 1.3E-39 | -0.40256 | 0.838 | 0.89 | 3.08E-35 |
| Ifitm3 | 5.37E-39 | 2.123076 | 0.22 | 0.121 | 1.27E-34 |
| Daglb | 9.83E-39 | -0.48952 | 0.665 | 0.753 | 2.32E-34 |
| Slc40a1 | 1.6E-38 | -0.50711 | 0.38 | 0.522 | 3.76E-34 |
| Rps6ka1 | 2.01E-38 | -0.38615 | 0.724 | 0.813 | 4.73E-34 |
| Chsy1 | 2.25E-38 | -0.62078 | 0.358 | 0.496 | 5.3E-34 |
| Fam110a | 3.26E-38 | -0.69763 | 0.232 | 0.364 | 7.7E-34 |
| Tmem204 | 5.64E-38 | -1.18505 | 0.114 | 0.225 | 1.33E-33 |
| Rps27 | 6.32E-38 | 0.571028 | 0.957 | 0.95 | 1.49E-33 |
| Tm9sf2 | 6.94E-38 | -0.33495 | 0.659 | 0.769 | 1.64E-33 |
| Maf | 6.97E-38 | -0.52264 | 0.839 | 0.897 | 1.64E-33 |
| Tram1 | 9.05E-38 | -0.4136 | 0.541 | 0.673 | 2.14E-33 |
| Golm1 | 9.5E-38 | -0.28009 | 0.766 | 0.877 | 2.24E-33 |
| Rtn4rl1 | 1.35E-37 | -0.56934 | 0.563 | 0.684 | 3.18E-33 |
| Rpl13 | 6.23E-37 | 0.564876 | 0.943 | 0.938 | 1.47E-32 |
| Cdc42 | 1.18E-36 | -0.3343 | 0.847 | 0.899 | 2.79E-32 |
| Bin2 | 1.78E-36 | -0.36265 | 0.843 | 0.909 | 4.21E-32 |
| AU020206 | 1.98E-36 | 0.838021 | 0.584 | 0.473 | 4.66E-32 |
| Rps18 | 3.99E-36 | 0.633077 | 0.854 | 0.833 | 9.41E-32 |
| Lsp1 | 4.5E-36 | 1.103284 | 0.404 | 0.284 | 1.06E-31 |
| Mylip | 5.26E-36 | -0.4489 | 0.582 | 0.699 | 1.24E-31 |
| Vav1 | 6.7E-36 | -0.56406 | 0.557 | 0.661 | 1.58E-31 |
| Rap1b | 2.32E-35 | -0.45174 | 0.694 | 0.773 | 5.47E-31 |
| Gm10076 | 1.6E-34 | 0.530763 | 0.902 | 0.893 | 3.77E-30 |
| Rplp2 | 2.13E-34 | 0.544784 | 0.897 | 0.895 | 5.02E-30 |

| | | | | | |
|---|---|---|---|---|---|
| Sall1 | 2.73E-34 | -0.34455 | 0.777 | 0.856 | 6.44E-30 |
| P4ha1 | 5.65E-34 | -0.39299 | 0.54 | 0.656 | 1.33E-29 |
| Rps9 | 8.42E-34 | 0.472374 | 0.962 | 0.95 | 1.99E-29 |
| Rps14 | 9.04E-34 | 0.554939 | 0.919 | 0.914 | 2.13E-29 |
| Entpd1 | 9.93E-34 | -0.3086 | 0.892 | 0.94 | 2.34E-29 |
| Gm10800 | 1.2E-33 | 0.578165 | 0.753 | 0.711 | 2.84E-29 |
| Fermt3 | 1.91E-33 | -0.46637 | 0.69 | 0.773 | 4.5E-29 |
| Gm26825 | 3.01E-33 | -0.30562 | 0.616 | 0.743 | 7.11E-29 |
| Plod1 | 3.27E-33 | -0.44336 | 0.519 | 0.628 | 7.72E-29 |
| Cd9 | 3.44E-33 | 0.531919 | 0.965 | 0.965 | 8.11E-29 |
| Sox4 | 4.29E-33 | -0.64584 | 0.297 | 0.422 | 1.01E-28 |
| Rps12 | 4.38E-33 | 0.606649 | 0.838 | 0.812 | 1.03E-28 |
| Slc44a2 | 5.01E-33 | -0.64121 | 0.316 | 0.443 | 1.18E-28 |
| Tyrobp | 8.09E-33 | 0.400426 | 0.994 | 0.996 | 1.91E-28 |
| Ywhaz | 8.76E-33 | -0.39087 | 0.811 | 0.88 | 2.07E-28 |
| Scamp5 | 9.43E-33 | -0.39338 | 0.444 | 0.578 | 2.22E-28 |
| Ivns1abp | 1.77E-32 | -0.3105 | 0.773 | 0.852 | 4.17E-28 |
| Bin1 | 2.16E-32 | -0.30674 | 0.859 | 0.913 | 5.1E-28 |
| Rpl21 | 4.74E-32 | 0.540041 | 0.918 | 0.917 | 1.12E-27 |
| Il16 | 1.14E-31 | -0.60814 | 0.467 | 0.574 | 2.7E-27 |
| Gm26917 | 1.19E-31 | 0.733967 | 0.943 | 0.951 | 2.8E-27 |
| Rab3il1 | 1.31E-31 | -0.36071 | 0.759 | 0.828 | 3.09E-27 |
| Hpgd | 1.41E-31 | -0.45599 | 0.661 | 0.746 | 3.34E-27 |
| Rps29 | 2.67E-31 | 0.456568 | 0.989 | 0.972 | 6.3E-27 |
| Dusp7 | 3.63E-31 | -0.32167 | 0.458 | 0.588 | 8.56E-27 |
| Rpl37 | 5.13E-31 | 0.539868 | 0.899 | 0.886 | 1.21E-26 |
| C1qb | 5.15E-31 | 0.293248 | 1 | 1 | 1.22E-26 |
| Canx | 6.62E-31 | -0.31416 | 0.876 | 0.934 | 1.56E-26 |
| Emp3 | 7.71E-31 | 1.302949 | 0.218 | 0.128 | 1.82E-26 |
| Ftl2-ps | 1.36E-30 | 0.470801 | 0.936 | 0.914 | 3.21E-26 |
| Slco2b1 | 6.51E-30 | -0.27652 | 0.878 | 0.937 | 1.54E-25 |
| Ddit4 | 1.27E-29 | -0.61949 | 0.146 | 0.25 | 3E-25 |
| 1700017B0 | 1.29E-29 | -0.26849 | 0.574 | 0.694 | 3.05E-25 |
| Gm15427 | 1.6E-29 | 0.438221 | 0.952 | 0.942 | 3.77E-25 |
| Serinc3 | 1.81E-29 | -0.27436 | 0.979 | 0.996 | 4.28E-25 |
| Trf | 4.74E-29 | -0.26777 | 0.945 | 0.971 | 1.12E-24 |
| Ncf1 | 5.02E-29 | -0.42375 | 0.66 | 0.735 | 1.19E-24 |
| St3gal5 | 5.33E-29 | -0.49039 | 0.513 | 0.611 | 1.26E-24 |
| Lcp1 | 6.39E-29 | -0.31825 | 0.701 | 0.785 | 1.51E-24 |
| Zfp706 | 7.77E-29 | -0.34983 | 0.55 | 0.661 | 1.83E-24 |
| Cd63 | 1.02E-28 | 0.759086 | 0.739 | 0.684 | 2.4E-24 |
| Atp6ap2 | 1.57E-28 | -0.45801 | 0.548 | 0.639 | 3.71E-24 |
| Zfp710 | 1.99E-28 | -0.49748 | 0.627 | 0.727 | 4.69E-24 |
| Elmo1 | 2.06E-28 | -0.32035 | 0.62 | 0.724 | 4.87E-24 |
| Abhd12 | 2.61E-28 | -0.26912 | 0.952 | 0.977 | 6.15E-24 |
| Rps20 | 2.63E-28 | 0.632091 | 0.83 | 0.807 | 6.21E-24 |
| Abcc3 | 2.95E-28 | -0.45771 | 0.305 | 0.427 | 6.95E-24 |

| | | | | | |
|---|---|---|---|---|---|
| Nrros | 4.86E-28 | -0.4472 | 0.546 | 0.64 | 1.15E-23 |
| Zfhx3 | 6.29E-28 | -0.30736 | 0.899 | 0.948 | 1.48E-23 |
| Vapa | 1.03E-27 | -0.3923 | 0.543 | 0.648 | 2.44E-23 |
| Sema4b | 1.06E-27 | -0.57217 | 0.117 | 0.213 | 2.51E-23 |
| Cd52 | 1.08E-27 | 1.003163 | 0.503 | 0.411 | 2.54E-23 |
| Hfe | 1.22E-27 | -0.49762 | 0.335 | 0.452 | 2.88E-23 |
| Etv5 | 1.5E-27 | -0.65335 | 0.303 | 0.418 | 3.54E-23 |
| Irf5 | 1.73E-27 | -0.43847 | 0.605 | 0.686 | 4.09E-23 |
| Man1a | 1.85E-27 | -0.48136 | 0.286 | 0.401 | 4.36E-23 |
| Tspan14 | 1.85E-27 | -0.4771 | 0.469 | 0.571 | 4.36E-23 |
| Rps19 | 2.18E-27 | 0.558618 | 0.864 | 0.863 | 5.14E-23 |
| Rpl36 | 2.42E-27 | 0.593094 | 0.823 | 0.799 | 5.7E-23 |
| Parvg | 2.65E-27 | -0.70949 | 0.361 | 0.457 | 6.24E-23 |
| Bank1 | 2.83E-27 | -1.31576 | 0.111 | 0.2 | 6.69E-23 |
| Dpysl2 | 2.91E-27 | -0.43 | 0.819 | 0.878 | 6.87E-23 |
| Gm26740 | 3.74E-27 | 1.157925 | 0.207 | 0.123 | 8.83E-23 |
| Rps23 | 5.68E-27 | 0.475849 | 0.895 | 0.887 | 1.34E-22 |
| St3gal6 | 7.74E-27 | -0.31136 | 0.728 | 0.804 | 1.82E-22 |
| Asah1 | 7.86E-27 | -0.26659 | 0.809 | 0.845 | 1.85E-22 |
| Sdcbp | 8.17E-27 | -0.40679 | 0.65 | 0.71 | 1.93E-22 |
| Nptn | 9.85E-27 | -0.31239 | 0.517 | 0.619 | 2.32E-22 |
| Ssr1 | 1.35E-26 | -0.51568 | 0.56 | 0.652 | 3.2E-22 |
| Vcp | 1.74E-26 | -0.43626 | 0.471 | 0.576 | 4.11E-22 |
| Ctsh | 1.83E-26 | 0.43584 | 0.954 | 0.947 | 4.32E-22 |
| Saraf | 1.95E-26 | -0.29215 | 0.677 | 0.768 | 4.61E-22 |
| Eng | 2.87E-26 | -0.61909 | 0.423 | 0.519 | 6.77E-22 |
| Gas6 | 3.02E-26 | -0.6296 | 0.387 | 0.482 | 7.13E-22 |
| Rpn2 | 3.26E-26 | -0.37196 | 0.661 | 0.735 | 7.7E-22 |
| Snx18 | 4.61E-26 | -0.42892 | 0.544 | 0.648 | 1.09E-21 |
| Gm12715 | 7.52E-26 | -0.33389 | 0.615 | 0.724 | 1.77E-21 |
| Gna15 | 1.05E-25 | -0.35603 | 0.593 | 0.689 | 2.48E-21 |
| St6galnac4 | 1.29E-25 | -0.43801 | 0.358 | 0.467 | 3.04E-21 |
| Wdr1 | 1.57E-25 | -0.57807 | 0.438 | 0.529 | 3.7E-21 |
| Gm24270 | 1.9E-25 | 0.461459 | 0.779 | 0.749 | 4.48E-21 |
| Idh2 | 2.01E-25 | -0.49116 | 0.336 | 0.444 | 4.75E-21 |
| Prpsap2 | 2.2E-25 | -0.63819 | 0.256 | 0.359 | 5.18E-21 |
| Rac1 | 3.26E-25 | -0.33476 | 0.79 | 0.845 | 7.7E-21 |
| Mgat1 | 3.27E-25 | -0.4947 | 0.48 | 0.574 | 7.71E-21 |
| Actg1 | 3.45E-25 | -0.27875 | 0.798 | 0.854 | 8.14E-21 |
| Pdia6 | 4.5E-25 | -0.26995 | 0.638 | 0.736 | 1.06E-20 |
| Eef2k | 5.56E-25 | -0.55561 | 0.342 | 0.448 | 1.31E-20 |
| Sipa1 | 6.55E-25 | -0.32313 | 0.533 | 0.635 | 1.55E-20 |
| Adam10 | 6.95E-25 | -0.25075 | 0.709 | 0.796 | 1.64E-20 |
| Akirin1 | 2.63E-24 | -0.60094 | 0.274 | 0.375 | 6.21E-20 |
| Gm17383 | 3.05E-24 | -0.41503 | 0.46 | 0.572 | 7.19E-20 |
| Tuba1a | 3.36E-24 | -0.44025 | 0.512 | 0.61 | 7.93E-20 |
| Snx17 | 3.7E-24 | -0.43793 | 0.398 | 0.501 | 8.72E-20 |

| | | | | | |
|---|---|---|---|---|---|
| Rilpl1 | 4.3E-24 | -0.48881 | 0.224 | 0.328 | 1.01E-19 |
| Ccr5 | 5.2E-24 | -0.39848 | 0.647 | 0.717 | 1.23E-19 |
| Pald1 | 5.39E-24 | -0.43239 | 0.364 | 0.473 | 1.27E-19 |
| Rtn3 | 6.09E-24 | -0.28995 | 0.603 | 0.688 | 1.44E-19 |
| Trim8 | 6.57E-24 | -0.49112 | 0.397 | 0.493 | 1.55E-19 |
| Tnfrsf21 | 6.79E-24 | -0.35742 | 0.409 | 0.513 | 1.6E-19 |
| Cask | 7.22E-24 | -0.44249 | 0.3 | 0.405 | 1.7E-19 |
| Hnrnpf | 8.14E-24 | -0.39523 | 0.511 | 0.596 | 1.92E-19 |
| Fli1 | 1.13E-23 | -0.4021 | 0.55 | 0.645 | 2.65E-19 |
| Mrfap1 | 1.19E-23 | -0.28456 | 0.599 | 0.697 | 2.82E-19 |
| Lag3 | 1.24E-23 | -0.35918 | 0.713 | 0.788 | 2.92E-19 |
| Tuba1b | 1.29E-23 | -0.27006 | 0.695 | 0.779 | 3.03E-19 |
| Ctsz | 1.3E-23 | 0.34984 | 0.995 | 0.993 | 3.06E-19 |
| Gm8822 | 1.32E-23 | -0.31302 | 0.166 | 0.261 | 3.11E-19 |
| Irf2bpl | 1.75E-23 | -0.33009 | 0.274 | 0.378 | 4.13E-19 |
| Stt3a | 1.94E-23 | -0.46117 | 0.487 | 0.576 | 4.57E-19 |
| Arl8b | 1.96E-23 | -0.52632 | 0.397 | 0.494 | 4.63E-19 |
| Sec61a1 | 2.66E-23 | -0.33563 | 0.55 | 0.638 | 6.27E-19 |
| Lcp2 | 4.09E-23 | -0.40195 | 0.498 | 0.595 | 9.66E-19 |
| Cd63-ps | 4.26E-23 | 0.558892 | 0.541 | 0.465 | 1E-18 |
| Cttnbp2nl | 4.31E-23 | -0.27436 | 0.661 | 0.755 | 1.02E-18 |
| Gm4149 | 4.54E-23 | 0.386726 | 0.837 | 0.813 | 1.07E-18 |
| Sept2 | 4.82E-23 | -0.42032 | 0.681 | 0.734 | 1.14E-18 |
| Rpl30 | 5.62E-23 | 0.575519 | 0.862 | 0.858 | 1.33E-18 |
| Ccng2 | 5.88E-23 | -0.53551 | 0.266 | 0.365 | 1.39E-18 |
| Serinc1 | 6.34E-23 | -0.38224 | 0.571 | 0.657 | 1.5E-18 |
| Kmt2d | 1.02E-22 | -0.25038 | 0.604 | 0.703 | 2.4E-18 |
| Slc35c2 | 1.03E-22 | -0.55555 | 0.37 | 0.464 | 2.44E-18 |
| Rps13 | 1.4E-22 | 0.656458 | 0.778 | 0.766 | 3.3E-18 |
| Surf4 | 1.42E-22 | -0.30791 | 0.518 | 0.618 | 3.35E-18 |
| Rpl23a | 1.64E-22 | 0.524462 | 0.773 | 0.745 | 3.87E-18 |
| Adrb2 | 2.1E-22 | -0.29357 | 0.561 | 0.652 | 4.96E-18 |
| Gm4202 | 2.44E-22 | -0.37389 | 0.454 | 0.566 | 5.77E-18 |
| Tm9sf3 | 4.68E-22 | -0.33369 | 0.39 | 0.493 | 1.1E-17 |
| Rpl23 | 5.88E-22 | 0.603883 | 0.826 | 0.82 | 1.39E-17 |
| Tnfrsf1b | 5.88E-22 | -0.37135 | 0.458 | 0.559 | 1.39E-17 |
| Rps24 | 5.98E-22 | 0.565428 | 0.843 | 0.818 | 1.41E-17 |
| Fau | 1.06E-21 | 0.469367 | 0.896 | 0.899 | 2.49E-17 |
| Gsk3b | 1.18E-21 | -0.42401 | 0.514 | 0.606 | 2.78E-17 |
| Snx5 | 1.37E-21 | -0.26679 | 0.753 | 0.81 | 3.23E-17 |
| Gp9 | 1.37E-21 | -0.36454 | 0.14 | 0.228 | 3.23E-17 |
| Tlr12 | 1.55E-21 | -0.6366 | 0.19 | 0.279 | 3.66E-17 |
| Sqstm1 | 1.94E-21 | -0.46921 | 0.535 | 0.606 | 4.58E-17 |
| Ppp1cc | 2.01E-21 | -0.39612 | 0.253 | 0.351 | 4.74E-17 |
| Rpl35 | 2.67E-21 | 0.583361 | 0.816 | 0.807 | 6.29E-17 |
| Adgre1 | 2.93E-21 | -0.28991 | 0.536 | 0.629 | 6.92E-17 |
| Tpt1 | 2.96E-21 | 0.534926 | 0.87 | 0.878 | 6.97E-17 |

| | | | | | |
|---|---|---|---|---|---|
| Foxn3 | 3.85E-21 | -0.30051 | 0.811 | 0.872 | 9.08E-17 |
| Golt1b | 3.93E-21 | -0.69111 | 0.233 | 0.323 | 9.28E-17 |
| Ywhae | 4.33E-21 | -0.33526 | 0.717 | 0.777 | 1.02E-16 |
| Snx6 | 4.46E-21 | -0.38918 | 0.38 | 0.478 | 1.05E-16 |
| Arhgap5 | 5.1E-21 | -0.25274 | 0.679 | 0.761 | 1.2E-16 |
| Tmem63a | 7.1E-21 | -0.39102 | 0.371 | 0.469 | 1.67E-16 |
| Itpripl1 | 7.52E-21 | -0.31812 | 0.631 | 0.729 | 1.77E-16 |
| Gm23935 | 7.99E-21 | 0.396945 | 0.962 | 0.965 | 1.89E-16 |
| Prkab1 | 8.57E-21 | -0.61928 | 0.288 | 0.381 | 2.02E-16 |
| Btg2 | 8.71E-21 | -0.42868 | 0.507 | 0.584 | 2.05E-16 |
| Ctsl | 8.73E-21 | 0.381775 | 0.947 | 0.946 | 2.06E-16 |
| Cables1 | 1.04E-20 | -0.73928 | 0.204 | 0.29 | 2.45E-16 |
| Tcn2 | 1.22E-20 | -0.26287 | 0.764 | 0.836 | 2.87E-16 |
| Fcrl1 | 1.31E-20 | -0.70706 | 0.199 | 0.287 | 3.09E-16 |
| Zfp652 | 1.32E-20 | -0.54361 | 0.617 | 0.694 | 3.11E-16 |
| Pdia4 | 1.45E-20 | -0.54106 | 0.298 | 0.392 | 3.41E-16 |
| Itgb2 | 1.88E-20 | -0.30506 | 0.749 | 0.789 | 4.43E-16 |
| Rplp0 | 2.06E-20 | 0.516894 | 0.891 | 0.89 | 4.87E-16 |
| Gm10134 | 2.11E-20 | -0.61365 | 0.294 | 0.388 | 4.99E-16 |
| Tpp1 | 2.46E-20 | -0.29544 | 0.704 | 0.771 | 5.81E-16 |
| Sla | 2.78E-20 | -0.3387 | 0.458 | 0.545 | 6.55E-16 |
| Ncoa4 | 3.89E-20 | -0.40416 | 0.169 | 0.254 | 9.18E-16 |
| Plod3 | 4.21E-20 | -0.35843 | 0.486 | 0.575 | 9.93E-16 |
| Cdh23 | 4.28E-20 | -0.64839 | 0.174 | 0.26 | 1.01E-15 |
| C5ar2 | 4.78E-20 | -0.32209 | 0.556 | 0.658 | 1.13E-15 |
| Ak1 | 5.09E-20 | -0.70903 | 0.169 | 0.253 | 1.2E-15 |
| Rtn1 | 6E-20 | -0.50127 | 0.301 | 0.391 | 1.42E-15 |
| Gcn1l1 | 7.02E-20 | -0.44126 | 0.363 | 0.464 | 1.66E-15 |
| Ddx5 | 7.06E-20 | -0.28815 | 0.721 | 0.789 | 1.67E-15 |
| Cyba | 8.64E-20 | 0.394418 | 0.961 | 0.973 | 2.04E-15 |
| M6pr | 8.74E-20 | -0.412 | 0.522 | 0.59 | 2.06E-15 |
| Tmed2 | 1.02E-19 | -0.27492 | 0.574 | 0.653 | 2.41E-15 |
| Lpcat3 | 1.08E-19 | -0.26846 | 0.447 | 0.547 | 2.54E-15 |
| Eif4g2 | 1.13E-19 | -0.31574 | 0.462 | 0.565 | 2.67E-15 |
| Tifab | 1.3E-19 | -0.28757 | 0.487 | 0.577 | 3.07E-15 |
| Sh2b3 | 1.31E-19 | -0.49459 | 0.352 | 0.449 | 3.1E-15 |
| Stard3 | 1.79E-19 | -0.42203 | 0.384 | 0.472 | 4.23E-15 |
| Rnf167 | 1.81E-19 | -0.4527 | 0.445 | 0.524 | 4.26E-15 |
| Tmsb4x | 1.92E-19 | 0.358849 | 1 | 1 | 4.52E-15 |
| Ppp1ccb | 2E-19 | -0.25016 | 0.187 | 0.276 | 4.71E-15 |
| Mef2c | 2.95E-19 | -0.2849 | 0.802 | 0.846 | 6.95E-15 |
| Nars | 3.4E-19 | -0.47965 | 0.416 | 0.507 | 8.01E-15 |
| Sash3 | 3.53E-19 | -0.47336 | 0.32 | 0.412 | 8.33E-15 |
| Map2k1 | 3.54E-19 | -0.55575 | 0.38 | 0.459 | 8.34E-15 |
| Sf1 | 4.41E-19 | -0.62331 | 0.249 | 0.337 | 1.04E-14 |
| Rpsa | 4.47E-19 | 0.60582 | 0.804 | 0.774 | 1.05E-14 |
| Ctsf | 4.53E-19 | -0.31832 | 0.586 | 0.666 | 1.07E-14 |

| | | | | | |
|---|---|---|---|---|---|
| Chst7 | 4.8E-19 | -0.65157 | 0.203 | 0.286 | 1.13E-14 |
| Tapbp | 5.07E-19 | -0.31669 | 0.692 | 0.753 | 1.2E-14 |
| Xbp1 | 5.69E-19 | -0.34011 | 0.546 | 0.63 | 1.34E-14 |
| Tmem206 | 5.76E-19 | -0.48111 | 0.266 | 0.353 | 1.36E-14 |
| Atp5a1 | 7.43E-19 | -0.35976 | 0.54 | 0.611 | 1.75E-14 |
| Csnk1a1 | 7.53E-19 | -0.29428 | 0.423 | 0.519 | 1.78E-14 |
| Edem2 | 8.03E-19 | -0.3308 | 0.497 | 0.586 | 1.89E-14 |
| Ppp3ca | 8.3E-19 | -0.36775 | 0.445 | 0.533 | 1.96E-14 |
| Ms4a6b | 8.37E-19 | -0.31805 | 0.277 | 0.372 | 1.97E-14 |
| Casp8 | 8.81E-19 | -0.34282 | 0.382 | 0.474 | 2.08E-14 |
| Dtx4 | 9.36E-19 | -0.49274 | 0.365 | 0.458 | 2.21E-14 |
| Dnm2 | 1.01E-18 | -0.36266 | 0.617 | 0.687 | 2.38E-14 |
| Mir692-2 | 1.02E-18 | 0.275946 | 0.38 | 0.301 | 2.41E-14 |
| Rpl19 | 1.39E-18 | 0.515159 | 0.803 | 0.794 | 3.29E-14 |
| Cd47 | 1.42E-18 | -0.37817 | 0.586 | 0.655 | 3.36E-14 |
| Rcsd1 | 1.46E-18 | -0.41129 | 0.371 | 0.46 | 3.44E-14 |
| 4833439L1 | 1.53E-18 | -0.82842 | 0.257 | 0.336 | 3.6E-14 |
| Dnajb11 | 1.87E-18 | -0.54459 | 0.301 | 0.387 | 4.41E-14 |
| Fez2 | 2.01E-18 | -0.41327 | 0.414 | 0.501 | 4.73E-14 |
| Ubash3b | 2.02E-18 | -0.48201 | 0.535 | 0.608 | 4.76E-14 |
| Gm10801 | 2.55E-18 | 0.333634 | 0.328 | 0.248 | 6.01E-14 |
| 0610040J0 | 2.74E-18 | -0.37751 | 0.466 | 0.543 | 6.46E-14 |
| Edem1 | 2.82E-18 | -0.40294 | 0.504 | 0.591 | 6.66E-14 |
| Mta2 | 3E-18 | -0.5631 | 0.204 | 0.287 | 7.07E-14 |
| Irf2bp2 | 3.26E-18 | -0.38196 | 0.289 | 0.381 | 7.7E-14 |
| Copb1 | 3.35E-18 | -0.28659 | 0.256 | 0.347 | 7.91E-14 |
| Mtss1 | 3.47E-18 | -0.29171 | 0.28 | 0.37 | 8.18E-14 |
| Pacsin2 | 3.58E-18 | -0.47718 | 0.376 | 0.456 | 8.44E-14 |
| Fcer1g | 4.11E-18 | 0.331415 | 0.988 | 0.993 | 9.69E-14 |
| Vps35 | 4.29E-18 | -0.36381 | 0.272 | 0.362 | 1.01E-13 |
| Man1c1 | 4.33E-18 | -0.33195 | 0.431 | 0.521 | 1.02E-13 |
| Map7d1 | 4.47E-18 | -0.49649 | 0.254 | 0.34 | 1.05E-13 |
| H2-M3 | 4.83E-18 | 0.893606 | 0.259 | 0.184 | 1.14E-13 |
| Asb2 | 4.87E-18 | -0.46481 | 0.285 | 0.375 | 1.15E-13 |
| Nckap5l | 5.02E-18 | -0.69669 | 0.163 | 0.24 | 1.18E-13 |
| Scn1b | 6.78E-18 | -0.44694 | 0.232 | 0.316 | 1.6E-13 |
| Man2b2 | 7.75E-18 | -0.27705 | 0.387 | 0.47 | 1.83E-13 |
| Amfr | 7.83E-18 | -0.36502 | 0.359 | 0.444 | 1.85E-13 |
| Gm15590 | 8.97E-18 | 0.267115 | 0.529 | 0.453 | 2.12E-13 |
| Sypl | 9.31E-18 | -0.4498 | 0.385 | 0.463 | 2.2E-13 |
| Hnrnpul1 | 1.08E-17 | -0.25223 | 0.414 | 0.507 | 2.55E-13 |
| Mgll | 1.2E-17 | -0.29773 | 0.272 | 0.361 | 2.84E-13 |
| Neat1 | 1.25E-17 | 0.701242 | 0.258 | 0.183 | 2.95E-13 |
| Actr3 | 1.35E-17 | -0.45861 | 0.466 | 0.536 | 3.18E-13 |
| Abcd3 | 2.16E-17 | -0.62334 | 0.149 | 0.225 | 5.11E-13 |
| Samsn1 | 2.78E-17 | -0.46449 | 0.375 | 0.464 | 6.55E-13 |
| Adam15 | 3.51E-17 | -0.32991 | 0.583 | 0.648 | 8.28E-13 |

| | | | | | |
|---|---|---|---|---|---|
| Atp1b3 | 4.05E-17 | -0.25589 | 0.6 | 0.679 | 9.55E-13 |
| Lars2 | 4.07E-17 | 0.459669 | 0.933 | 0.923 | 9.61E-13 |
| Gtf2a1 | 4.13E-17 | -0.65108 | 0.311 | 0.389 | 9.73E-13 |
| Klf3 | 5E-17 | -0.28905 | 0.507 | 0.588 | 1.18E-12 |
| Itpr2 | 5.4E-17 | -0.48369 | 0.432 | 0.517 | 1.27E-12 |
| Sppl2a | 5.4E-17 | -0.33255 | 0.511 | 0.589 | 1.27E-12 |
| Ly6e | 6.34E-17 | 0.500739 | 0.952 | 0.948 | 1.5E-12 |
| Inpp4b | 6.95E-17 | -0.67634 | 0.177 | 0.252 | 1.64E-12 |
| Sort1 | 7.18E-17 | -0.29335 | 0.424 | 0.52 | 1.69E-12 |
| Tspan7 | 7.24E-17 | -0.36905 | 0.451 | 0.532 | 1.71E-12 |
| Ggta1 | 7.75E-17 | -0.61952 | 0.134 | 0.206 | 1.83E-12 |
| Abhd6 | 7.82E-17 | -0.28217 | 0.254 | 0.343 | 1.84E-12 |
| Rps27a | 8.88E-17 | 0.561297 | 0.794 | 0.78 | 2.09E-12 |
| Rpl34 | 8.91E-17 | 0.50101 | 0.785 | 0.778 | 2.1E-12 |
| Spcs3 | 8.96E-17 | -0.57475 | 0.204 | 0.284 | 2.11E-12 |
| Arl1 | 9.87E-17 | -0.54087 | 0.266 | 0.347 | 2.33E-12 |
| Tnfrsf13b | 1.15E-16 | -0.6755 | 0.254 | 0.331 | 2.71E-12 |
| Bcl2l1 | 1.26E-16 | -0.63341 | 0.341 | 0.415 | 2.96E-12 |
| Mef2a | 1.31E-16 | -0.25365 | 0.757 | 0.81 | 3.08E-12 |
| Abca1 | 1.49E-16 | -0.28397 | 0.464 | 0.556 | 3.51E-12 |
| Gm14303 | 1.6E-16 | 0.413291 | 0.788 | 0.753 | 3.76E-12 |
| Bmp2k | 1.67E-16 | -0.32184 | 0.894 | 0.933 | 3.93E-12 |
| G530011O( | 1.68E-16 | -0.25987 | 0.297 | 0.385 | 3.95E-12 |
| Zmiz1 | 1.89E-16 | -0.39516 | 0.563 | 0.642 | 4.45E-12 |
| Gm8399 | 2.02E-16 | -0.26022 | 0.411 | 0.509 | 4.77E-12 |
| Trim28 | 2.06E-16 | -0.44286 | 0.24 | 0.321 | 4.86E-12 |
| Mafg | 2.18E-16 | -0.5466 | 0.454 | 0.525 | 5.15E-12 |
| Rin2 | 2.27E-16 | -0.44712 | 0.478 | 0.55 | 5.37E-12 |
| Cnot8 | 2.56E-16 | -0.43611 | 0.421 | 0.496 | 6.04E-12 |
| Bid | 2.63E-16 | -0.63261 | 0.185 | 0.259 | 6.2E-12 |
| Ier5 | 2.81E-16 | -0.35288 | 0.445 | 0.525 | 6.63E-12 |
| Sesn1 | 3.71E-16 | -0.64634 | 0.363 | 0.439 | 8.74E-12 |
| Abhd17c | 4.13E-16 | -0.30132 | 0.39 | 0.474 | 9.74E-12 |
| Cyb561a3 | 4.13E-16 | -0.32443 | 0.41 | 0.487 | 9.75E-12 |
| Arf1 | 5.1E-16 | -0.2567 | 0.698 | 0.751 | 1.2E-11 |
| Wdr45b | 5.29E-16 | -0.98774 | 0.177 | 0.246 | 1.25E-11 |
| Rps16 | 5.39E-16 | 0.55355 | 0.774 | 0.76 | 1.27E-11 |
| Rps26 | 5.42E-16 | 0.518344 | 0.707 | 0.664 | 1.28E-11 |
| Clptm1 | 5.56E-16 | -0.33097 | 0.467 | 0.541 | 1.31E-11 |
| Rpl17 | 5.56E-16 | 0.564623 | 0.792 | 0.778 | 1.31E-11 |
| Rps8 | 5.94E-16 | 0.520423 | 0.825 | 0.813 | 1.4E-11 |
| Chtf8 | 6.11E-16 | -0.34769 | 0.138 | 0.206 | 1.44E-11 |
| Retreg1 | 6.59E-16 | -0.37263 | 0.413 | 0.501 | 1.55E-11 |
| Tmf1 | 7.15E-16 | -0.3873 | 0.437 | 0.515 | 1.69E-11 |
| Ube2h | 9.16E-16 | -0.33757 | 0.272 | 0.353 | 2.16E-11 |
| Ptp4a2 | 1E-15 | -0.2728 | 0.674 | 0.746 | 2.36E-11 |
| Ssr3 | 1.06E-15 | -0.5763 | 0.25 | 0.326 | 2.51E-11 |

| | | | | | |
|---|---|---|---|---|---|
| Ralb | 1.08E-15 | -0.50978 | 0.369 | 0.445 | 2.55E-11 |
| Klhl9 | 1.22E-15 | -0.37965 | 0.403 | 0.488 | 2.88E-11 |
| Hvcn1 | 1.26E-15 | -0.37305 | 0.462 | 0.531 | 2.98E-11 |
| Tmem30a | 1.32E-15 | -0.35328 | 0.408 | 0.491 | 3.12E-11 |
| Cln8 | 1.59E-15 | -0.51057 | 0.26 | 0.337 | 3.76E-11 |
| G3bp1 | 1.86E-15 | -0.5692 | 0.357 | 0.431 | 4.39E-11 |
| Rps21 | 2.22E-15 | 0.487306 | 0.781 | 0.762 | 5.23E-11 |
| Elk3 | 2.89E-15 | -0.28402 | 0.315 | 0.401 | 6.81E-11 |
| Prkcd | 3.07E-15 | -0.29221 | 0.676 | 0.725 | 7.24E-11 |
| Rpn1 | 3.08E-15 | -0.28247 | 0.534 | 0.617 | 7.27E-11 |
| Glg1 | 3.36E-15 | -0.33139 | 0.311 | 0.393 | 7.93E-11 |
| Gm10073 | 3.44E-15 | 0.346431 | 0.616 | 0.568 | 8.11E-11 |
| Gm6277 | 3.72E-15 | -0.48697 | 0.149 | 0.219 | 8.77E-11 |
| Tagap | 3.79E-15 | -0.3387 | 0.297 | 0.376 | 8.94E-11 |
| Ppia | 3.83E-15 | 0.317982 | 0.956 | 0.969 | 9.03E-11 |
| Sft2d2 | 3.9E-15 | -0.33731 | 0.342 | 0.427 | 9.19E-11 |
| Hs6st1 | 4.32E-15 | -0.27404 | 0.237 | 0.317 | 1.02E-10 |
| Ifi30 | 4.82E-15 | 0.616825 | 0.529 | 0.468 | 1.14E-10 |
| Mfsd14a | 4.96E-15 | -0.52578 | 0.167 | 0.235 | 1.17E-10 |
| Calu | 5.33E-15 | -0.45378 | 0.207 | 0.281 | 1.26E-10 |
| Mtpn | 5.34E-15 | -0.38243 | 0.318 | 0.395 | 1.26E-10 |
| Sqor | 5.98E-15 | -0.31439 | 0.242 | 0.319 | 1.41E-10 |
| Tgm2 | 6.23E-15 | -0.75614 | 0.202 | 0.27 | 1.47E-10 |
| Rps7 | 6.76E-15 | 0.582488 | 0.787 | 0.776 | 1.6E-10 |
| Adam17 | 7.03E-15 | -0.28685 | 0.351 | 0.433 | 1.66E-10 |
| Mlph | 8.29E-15 | -0.56204 | 0.335 | 0.411 | 1.96E-10 |
| Cd300a | 8.79E-15 | -0.26019 | 0.503 | 0.579 | 2.07E-10 |
| Ptges3 | 9.25E-15 | -0.3147 | 0.437 | 0.516 | 2.18E-10 |
| Fchsd2 | 9.27E-15 | -0.38602 | 0.345 | 0.429 | 2.19E-10 |
| Nek6 | 1E-14 | -0.45631 | 0.352 | 0.432 | 2.37E-10 |
| Psmd1 | 1.05E-14 | -0.68326 | 0.237 | 0.307 | 2.47E-10 |
| Bcl9l | 1.12E-14 | -0.26186 | 0.171 | 0.243 | 2.65E-10 |
| Numa1 | 1.26E-14 | -0.4105 | 0.32 | 0.401 | 2.98E-10 |
| Csnk2a1 | 1.38E-14 | -0.30389 | 0.417 | 0.494 | 3.24E-10 |
| Mcfd2 | 1.43E-14 | -0.29405 | 0.375 | 0.453 | 3.37E-10 |
| Prrc2a | 1.51E-14 | -0.27206 | 0.293 | 0.371 | 3.57E-10 |
| Gm5805 | 1.54E-14 | 0.354664 | 0.536 | 0.474 | 3.63E-10 |
| Srsf3 | 1.63E-14 | -0.31689 | 0.409 | 0.487 | 3.85E-10 |
| Zfp36l2 | 1.66E-14 | -0.29341 | 0.418 | 0.504 | 3.91E-10 |
| Hnrnpu | 1.78E-14 | -0.42455 | 0.367 | 0.447 | 4.2E-10 |
| Gm28437 | 1.84E-14 | 0.448509 | 0.82 | 0.825 | 4.34E-10 |
| Abl1 | 2.07E-14 | -0.34612 | 0.25 | 0.327 | 4.89E-10 |
| Pnrc2 | 2.53E-14 | -0.6589 | 0.255 | 0.323 | 5.97E-10 |
| Ptbp1 | 2.88E-14 | -0.77309 | 0.239 | 0.303 | 6.8E-10 |
| Oxct1 | 3.11E-14 | -0.2915 | 0.424 | 0.502 | 7.33E-10 |
| Rassf5 | 3.36E-14 | -0.50735 | 0.195 | 0.262 | 7.92E-10 |
| Gm11361 | 3.52E-14 | 0.318274 | 0.494 | 0.434 | 8.29E-10 |

| | | | | | |
|---|---|---|---|---|---|
| Uba52 | 3.91E-14 | 0.592881 | 0.752 | 0.721 | 9.21E-10 |
| Ddx6 | 4.18E-14 | -0.39409 | 0.307 | 0.381 | 9.87E-10 |
| Ggt5 | 4.49E-14 | -0.56487 | 0.193 | 0.263 | 1.06E-09 |
| Mxi1 | 4.67E-14 | -0.52989 | 0.139 | 0.203 | 1.1E-09 |
| Il21r | 5.01E-14 | -0.44923 | 0.402 | 0.473 | 1.18E-09 |
| Smad3 | 5.28E-14 | -0.26682 | 0.28 | 0.361 | 1.24E-09 |
| Mapre1 | 5.41E-14 | -0.39938 | 0.398 | 0.467 | 1.28E-09 |
| Rps10 | 5.44E-14 | 0.501843 | 0.8 | 0.786 | 1.28E-09 |
| Rpl36-ps2 | 7.51E-14 | 0.455611 | 0.535 | 0.478 | 1.77E-09 |
| Ncstn | 7.58E-14 | -0.30993 | 0.453 | 0.521 | 1.79E-09 |
| Ets1 | 8.15E-14 | -0.26697 | 0.274 | 0.351 | 1.92E-09 |
| Gm3511 | 8.64E-14 | 0.311051 | 0.596 | 0.534 | 2.04E-09 |
| Tle3 | 8.75E-14 | -0.31951 | 0.239 | 0.317 | 2.06E-09 |
| Rela | 8.95E-14 | -0.36654 | 0.322 | 0.397 | 2.11E-09 |
| Atp6v0a2 | 9.21E-14 | -0.40291 | 0.356 | 0.432 | 2.17E-09 |
| Ly86 | 1.08E-13 | 0.274801 | 0.984 | 0.991 | 2.55E-09 |
| Gm6977 | 1.17E-13 | 0.404513 | 0.544 | 0.479 | 2.75E-09 |
| Nrp1 | 1.17E-13 | -0.30454 | 0.36 | 0.437 | 2.76E-09 |
| Tpt1-ps3 | 1.19E-13 | 0.373001 | 0.642 | 0.62 | 2.8E-09 |
| Lrp10 | 1.27E-13 | -0.29651 | 0.399 | 0.472 | 3E-09 |
| Arhgap27 | 1.59E-13 | -0.34999 | 0.267 | 0.342 | 3.76E-09 |
| Med1 | 1.75E-13 | -0.57316 | 0.34 | 0.404 | 4.13E-09 |
| Npc2 | 1.83E-13 | 0.507641 | 0.834 | 0.82 | 4.32E-09 |
| Dbnl | 1.85E-13 | -0.30422 | 0.36 | 0.434 | 4.38E-09 |
| Slc25a45 | 2.18E-13 | -0.46392 | 0.268 | 0.338 | 5.15E-09 |
| Clstn1 | 2.25E-13 | -0.31756 | 0.358 | 0.432 | 5.32E-09 |
| Pfkl | 2.27E-13 | -0.63997 | 0.198 | 0.261 | 5.35E-09 |
| Gm12435 | 2.39E-13 | 0.330986 | 0.157 | 0.225 | 5.64E-09 |
| Gm11942 | 2.5E-13 | 0.360677 | 0.464 | 0.408 | 5.9E-09 |
| Twf1 | 2.7E-13 | -0.3817 | 0.283 | 0.356 | 6.37E-09 |
| Eif4a-ps4 | 2.75E-13 | -0.28315 | 0.331 | 0.411 | 6.49E-09 |
| Sppl3 | 2.78E-13 | -0.4339 | 0.4 | 0.469 | 6.55E-09 |
| Tmem64 | 2.81E-13 | -0.37192 | 0.153 | 0.218 | 6.62E-09 |
| Txndc5 | 2.82E-13 | -0.77313 | 0.154 | 0.216 | 6.65E-09 |
| Tspo | 2.87E-13 | 0.858365 | 0.34 | 0.274 | 6.78E-09 |
| Sh3bgrl3 | 3.49E-13 | 0.458092 | 0.805 | 0.812 | 8.23E-09 |
| Litaf | 3.5E-13 | -0.31055 | 0.413 | 0.484 | 8.26E-09 |
| Dnajb9 | 3.51E-13 | -0.40461 | 0.263 | 0.333 | 8.28E-09 |
| Ddb1 | 3.7E-13 | -0.44003 | 0.158 | 0.224 | 8.74E-09 |
| Yme1l1 | 4.11E-13 | -0.51817 | 0.336 | 0.409 | 9.7E-09 |
| Cryl1 | 4.21E-13 | -0.34174 | 0.262 | 0.333 | 9.94E-09 |
| Btg1 | 4.35E-13 | -0.31659 | 0.638 | 0.689 | 1.03E-08 |
| Tor4a | 4.64E-13 | -0.52668 | 0.196 | 0.262 | 1.09E-08 |
| 2900097C1 | 4.64E-13 | -0.66738 | 0.154 | 0.217 | 1.1E-08 |
| Irgm2 | 4.74E-13 | -0.58792 | 0.309 | 0.383 | 1.12E-08 |
| Cltc | 4.92E-13 | -0.4025 | 0.582 | 0.644 | 1.16E-08 |
| Cd83 | 5.79E-13 | 0.605289 | 0.33 | 0.26 | 1.36E-08 |

| | | | | | |
|---|---|---|---|---|---|
| Ubqln1 | 5.96E-13 | -0.49292 | 0.232 | 0.299 | 1.41E-08 |
| Gm10443 | 7.23E-13 | 0.272436 | 0.433 | 0.37 | 1.71E-08 |
| Acads | 7.43E-13 | -0.27404 | 0.211 | 0.283 | 1.75E-08 |
| Ptp4a3 | 7.46E-13 | -0.33505 | 0.462 | 0.531 | 1.76E-08 |
| Lpcat1 | 7.84E-13 | -0.32621 | 0.448 | 0.519 | 1.85E-08 |
| Rgmb | 8.14E-13 | -0.26991 | 0.381 | 0.461 | 1.92E-08 |
| Pdcd6ip | 8.94E-13 | -0.28177 | 0.26 | 0.334 | 2.11E-08 |
| Il1a | 1.03E-12 | -0.76567 | 0.15 | 0.21 | 2.42E-08 |
| Bicd2 | 1.08E-12 | -0.45942 | 0.201 | 0.268 | 2.54E-08 |
| Eif4h | 1.23E-12 | -0.32859 | 0.423 | 0.491 | 2.9E-08 |
| Tfe3 | 1.24E-12 | -0.34707 | 0.404 | 0.477 | 2.93E-08 |
| Ctnnb1 | 1.29E-12 | -0.42518 | 0.25 | 0.318 | 3.04E-08 |
| Kdm6b | 1.36E-12 | -0.32816 | 0.222 | 0.292 | 3.22E-08 |
| Slc16a6 | 1.38E-12 | -0.25526 | 0.477 | 0.548 | 3.25E-08 |
| Hcls1 | 1.41E-12 | -0.32833 | 0.507 | 0.566 | 3.32E-08 |
| Gm10736 | 1.41E-12 | 0.390064 | 0.542 | 0.489 | 3.33E-08 |
| Fbrsl1 | 1.58E-12 | -0.57086 | 0.351 | 0.414 | 3.73E-08 |
| Serp1 | 1.59E-12 | -0.42057 | 0.365 | 0.426 | 3.74E-08 |
| Zfp90 | 1.65E-12 | -0.45317 | 0.175 | 0.239 | 3.9E-08 |
| Atmin | 1.66E-12 | -0.25476 | 0.143 | 0.206 | 3.92E-08 |
| Lpxn | 1.82E-12 | -0.26331 | 0.316 | 0.39 | 4.28E-08 |
| Gm15564 | 1.83E-12 | 0.410547 | 0.601 | 0.537 | 4.33E-08 |
| Tnfrsf1a | 1.91E-12 | -0.30019 | 0.57 | 0.625 | 4.51E-08 |
| Aftph | 2.23E-12 | -0.29704 | 0.297 | 0.37 | 5.25E-08 |
| Psen1 | 2.8E-12 | -0.39678 | 0.321 | 0.391 | 6.61E-08 |
| Sh3bp1 | 3.02E-12 | -0.45438 | 0.252 | 0.318 | 7.12E-08 |
| Cul1 | 3.2E-12 | -0.25343 | 0.339 | 0.412 | 7.55E-08 |
| Gpr155 | 3.26E-12 | -0.48517 | 0.229 | 0.295 | 7.69E-08 |
| Rpl10 | 3.68E-12 | 0.360847 | 0.894 | 0.901 | 8.69E-08 |
| Pxk | 3.72E-12 | -0.62412 | 0.175 | 0.234 | 8.77E-08 |
| Hdac1 | 3.99E-12 | -0.40494 | 0.224 | 0.289 | 9.41E-08 |
| Ppp2r1a | 4.13E-12 | -0.43331 | 0.385 | 0.448 | 9.74E-08 |
| Rab5a | 4.18E-12 | -0.65196 | 0.189 | 0.249 | 9.86E-08 |
| Gm7331 | 4.38E-12 | 0.435063 | 0.574 | 0.528 | 1.03E-07 |
| Tmem144 | 4.59E-12 | -0.71608 | 0.145 | 0.203 | 1.08E-07 |
| Slc39a1 | 5.52E-12 | -0.31958 | 0.481 | 0.555 | 1.3E-07 |
| mt-Atp6 | 5.58E-12 | 0.415316 | 0.816 | 0.825 | 1.32E-07 |
| Tsg101 | 6.68E-12 | -0.37015 | 0.266 | 0.333 | 1.58E-07 |
| Srpr | 6.92E-12 | -0.39725 | 0.291 | 0.362 | 1.63E-07 |
| Letm1 | 7.47E-12 | -0.7364 | 0.16 | 0.217 | 1.76E-07 |
| Rpl13a | 8.48E-12 | 0.479053 | 0.821 | 0.824 | 2E-07 |
| Elavl1 | 8.87E-12 | -0.28257 | 0.285 | 0.354 | 2.09E-07 |
| Mat2a | 9.04E-12 | -0.29021 | 0.191 | 0.255 | 2.13E-07 |
| Lasp1 | 9.32E-12 | -0.28952 | 0.37 | 0.446 | 2.2E-07 |
| Tmem100 | 1.06E-11 | -0.28601 | 0.293 | 0.362 | 2.49E-07 |
| Orai2 | 1.09E-11 | -0.5742 | 0.172 | 0.233 | 2.57E-07 |
| Rbm4b | 1.21E-11 | -0.40125 | 0.334 | 0.402 | 2.85E-07 |

| | | | | | |
|---|---|---|---|---|---|
| Pgam1 | 1.3E-11 | -0.34085 | 0.249 | 0.314 | 3.07E-07 |
| Fbxw11 | 1.35E-11 | -0.43639 | 0.247 | 0.314 | 3.18E-07 |
| Tor1aip2 | 1.36E-11 | -0.32202 | 0.255 | 0.319 | 3.2E-07 |
| Rpl14 | 1.41E-11 | 0.570669 | 0.773 | 0.756 | 3.32E-07 |
| Dctn2 | 1.47E-11 | -0.28758 | 0.34 | 0.406 | 3.47E-07 |
| Pttg1ip | 1.48E-11 | -0.36608 | 0.414 | 0.47 | 3.48E-07 |
| Gm8730 | 1.59E-11 | 0.385427 | 0.733 | 0.723 | 3.76E-07 |
| Rab7 | 1.61E-11 | -0.29827 | 0.517 | 0.569 | 3.8E-07 |
| Abi1 | 1.63E-11 | -0.28265 | 0.314 | 0.383 | 3.84E-07 |
| Tlr4 | 1.76E-11 | -0.36769 | 0.154 | 0.213 | 4.15E-07 |
| Rab11b | 1.81E-11 | -0.30304 | 0.373 | 0.438 | 4.27E-07 |
| Hnrnph2 | 2.09E-11 | -0.79265 | 0.203 | 0.259 | 4.92E-07 |
| Rps19-ps6 | 2.11E-11 | 0.415353 | 0.647 | 0.618 | 4.99E-07 |
| Ythdf2 | 2.24E-11 | -0.27027 | 0.499 | 0.563 | 5.28E-07 |
| Crebbp | 2.24E-11 | -0.35034 | 0.506 | 0.569 | 5.29E-07 |
| Snx13 | 2.27E-11 | -0.27883 | 0.288 | 0.355 | 5.36E-07 |
| Rps27rt | 2.29E-11 | 0.373101 | 0.711 | 0.687 | 5.4E-07 |
| Frmd8 | 2.3E-11 | -0.79847 | 0.171 | 0.227 | 5.42E-07 |
| Slc9a9 | 2.57E-11 | -0.3027 | 0.305 | 0.373 | 6.06E-07 |
| Cerk | 2.62E-11 | -0.37396 | 0.35 | 0.417 | 6.17E-07 |
| Lrrc8a | 2.66E-11 | -0.30528 | 0.422 | 0.489 | 6.27E-07 |
| Ppp3r1 | 2.72E-11 | -0.54004 | 0.18 | 0.238 | 6.42E-07 |
| Maea | 2.79E-11 | -0.45671 | 0.225 | 0.286 | 6.59E-07 |
| Atp2a2 | 2.94E-11 | -0.3109 | 0.249 | 0.316 | 6.94E-07 |
| Scarb2 | 3.1E-11 | -0.27846 | 0.469 | 0.535 | 7.31E-07 |
| Arf4 | 3.33E-11 | -0.34783 | 0.268 | 0.334 | 7.85E-07 |
| Gm4332 | 3.44E-11 | 0.253787 | 0.266 | 0.212 | 8.11E-07 |
| Tm6sf1 | 3.46E-11 | -0.33307 | 0.398 | 0.459 | 8.16E-07 |
| Cep152 | 3.94E-11 | -0.57228 | 0.165 | 0.221 | 9.28E-07 |
| Extl3 | 4.05E-11 | -0.34978 | 0.368 | 0.437 | 9.56E-07 |
| Cd74 | 4.25E-11 | 1.591875 | 0.2 | 0.153 | 1E-06 |
| Gosr2 | 4.28E-11 | -0.32829 | 0.15 | 0.206 | 1.01E-06 |
| Wdfy3 | 4.36E-11 | -0.31843 | 0.299 | 0.365 | 1.03E-06 |
| Sf3b4 | 4.38E-11 | -0.27018 | 0.245 | 0.312 | 1.03E-06 |
| Gm15536 | 5.13E-11 | 0.413089 | 0.27 | 0.216 | 1.21E-06 |
| Spint1 | 5.38E-11 | -0.26522 | 0.421 | 0.487 | 1.27E-06 |
| Gorasp2 | 5.89E-11 | -0.43879 | 0.284 | 0.346 | 1.39E-06 |
| Tgfa | 5.97E-11 | -0.28843 | 0.217 | 0.28 | 1.41E-06 |
| Mgat2 | 6E-11 | -0.29978 | 0.305 | 0.368 | 1.41E-06 |
| Gm10132 | 6.8E-11 | 0.567813 | 0.591 | 0.555 | 1.6E-06 |
| Hsd17b4 | 6.99E-11 | -0.48764 | 0.289 | 0.348 | 1.65E-06 |
| Srp68 | 7.26E-11 | -0.58724 | 0.149 | 0.204 | 1.71E-06 |
| Phgdh | 7.35E-11 | -0.33907 | 0.314 | 0.38 | 1.73E-06 |
| Itfg1 | 8.34E-11 | -0.48605 | 0.278 | 0.34 | 1.97E-06 |
| Plekha2 | 8.61E-11 | -0.39813 | 0.183 | 0.242 | 2.03E-06 |
| Atp6v1h | 9.14E-11 | -0.52708 | 0.162 | 0.217 | 2.16E-06 |
| Ube2d3 | 9.75E-11 | -0.28099 | 0.579 | 0.616 | 2.3E-06 |

| | | | | | |
|---|---|---|---|---|---|
| Gm9625 | 1.04E-10 | 0.369518 | 0.518 | 0.477 | 2.46E-06 |
| Gm12918 | 1.05E-10 | 0.32851 | 0.543 | 0.496 | 2.49E-06 |
| Chn2 | 1.06E-10 | -0.3599 | 0.258 | 0.321 | 2.49E-06 |
| Spsb1 | 1.1E-10 | -0.46605 | 0.265 | 0.325 | 2.6E-06 |
| Aars | 1.16E-10 | -0.30177 | 0.291 | 0.355 | 2.73E-06 |
| Gna12 | 1.16E-10 | -0.2976 | 0.512 | 0.562 | 2.74E-06 |
| mt-Co3 | 1.17E-10 | 0.459496 | 0.893 | 0.902 | 2.76E-06 |
| Tnfsf13b | 1.21E-10 | 0.566494 | 0.217 | 0.162 | 2.85E-06 |
| Tspyl1 | 1.21E-10 | -0.35766 | 0.223 | 0.284 | 2.85E-06 |
| Sar1a | 1.34E-10 | -0.56705 | 0.255 | 0.314 | 3.17E-06 |
| Mid1ip1 | 1.38E-10 | -0.31039 | 0.35 | 0.413 | 3.26E-06 |
| Tmem135 | 1.4E-10 | -0.29774 | 0.308 | 0.371 | 3.3E-06 |
| Ctso | 1.51E-10 | -0.36713 | 0.335 | 0.393 | 3.56E-06 |
| St13 | 1.54E-10 | -0.25592 | 0.58 | 0.635 | 3.63E-06 |
| Ap1b1 | 1.56E-10 | -0.25049 | 0.39 | 0.459 | 3.68E-06 |
| Rab18 | 1.64E-10 | -0.44822 | 0.214 | 0.271 | 3.86E-06 |
| Gm15710 | 1.65E-10 | 0.317172 | 0.748 | 0.723 | 3.88E-06 |
| Itga9 | 1.67E-10 | -0.33025 | 0.224 | 0.291 | 3.94E-06 |
| Tmem248 | 1.67E-10 | -0.32063 | 0.149 | 0.204 | 3.95E-06 |
| Pdzd8 | 1.88E-10 | -0.27268 | 0.209 | 0.269 | 4.43E-06 |
| Sbf2 | 1.93E-10 | -0.27379 | 0.29 | 0.357 | 4.55E-06 |
| Il6st | 2.11E-10 | -0.25132 | 0.442 | 0.507 | 4.98E-06 |
| Npl | 2.22E-10 | -0.27324 | 0.166 | 0.224 | 5.24E-06 |
| Rbmxl1 | 2.45E-10 | -0.33461 | 0.211 | 0.272 | 5.79E-06 |
| Wsb1 | 2.6E-10 | -0.36881 | 0.35 | 0.407 | 6.14E-06 |
| Ext2 | 2.62E-10 | -0.43323 | 0.208 | 0.265 | 6.17E-06 |
| Bhlhe41 | 2.63E-10 | -0.28529 | 0.751 | 0.795 | 6.2E-06 |
| Ddah2 | 3.08E-10 | -0.27974 | 0.152 | 0.209 | 7.26E-06 |
| Kcnk13 | 3.12E-10 | -0.46836 | 0.275 | 0.332 | 7.35E-06 |
| Rpl38 | 3.16E-10 | 0.455906 | 0.647 | 0.616 | 7.46E-06 |
| Pkm | 3.52E-10 | -0.36646 | 0.517 | 0.559 | 8.3E-06 |
| Rps13-ps2 | 3.63E-10 | 0.320088 | 0.541 | 0.503 | 8.57E-06 |
| Rpl3 | 3.75E-10 | 0.431543 | 0.827 | 0.821 | 8.85E-06 |
| Rab35 | 3.76E-10 | -0.36046 | 0.213 | 0.278 | 8.87E-06 |
| BC035044 | 3.78E-10 | -0.28602 | 0.199 | 0.258 | 8.92E-06 |
| Gm29216 | 3.82E-10 | 0.250987 | 0.76 | 0.759 | 9E-06 |
| Rps15 | 3.83E-10 | 0.49031 | 0.759 | 0.737 | 9.03E-06 |
| Mettl7a1 | 3.87E-10 | -0.47224 | 0.159 | 0.213 | 9.12E-06 |
| Pigt | 3.87E-10 | -0.26736 | 0.261 | 0.318 | 9.13E-06 |
| Wac | 4E-10 | -0.30694 | 0.301 | 0.36 | 9.42E-06 |
| Rbm47 | 4.06E-10 | -0.36514 | 0.306 | 0.371 | 9.58E-06 |
| Gm9843 | 4.21E-10 | 0.377257 | 0.845 | 0.834 | 9.93E-06 |
| Gm38158 | 4.59E-10 | -0.28469 | 0.253 | 0.315 | 1.08E-05 |
| Csf2ra | 4.9E-10 | 0.805947 | 0.32 | 0.266 | 1.16E-05 |
| Fam3a | 5E-10 | -0.34434 | 0.238 | 0.297 | 1.18E-05 |
| Rpl10a | 5.01E-10 | 0.548287 | 0.673 | 0.643 | 1.18E-05 |
| Cldnd1 | 5.32E-10 | -0.27203 | 0.261 | 0.321 | 1.25E-05 |

| | | | | | |
|---|---|---|---|---|---|
| Maml3 | 5.47E-10 | -0.36667 | 0.216 | 0.277 | 1.29E-05 |
| Rps5 | 5.59E-10 | 0.507655 | 0.736 | 0.718 | 1.32E-05 |
| Vps26a | 5.67E-10 | -0.52712 | 0.206 | 0.26 | 1.34E-05 |
| Atp6v1c1 | 6.23E-10 | -0.32001 | 0.301 | 0.359 | 1.47E-05 |
| Rhoq | 6.47E-10 | -0.25709 | 0.185 | 0.244 | 1.53E-05 |
| Rnf145 | 6.49E-10 | -0.61805 | 0.166 | 0.218 | 1.53E-05 |
| Tpbgl | 6.78E-10 | -0.42534 | 0.335 | 0.393 | 1.6E-05 |
| Rnf2 | 6.82E-10 | -0.36648 | 0.156 | 0.211 | 1.61E-05 |
| Crot | 6.96E-10 | -0.70838 | 0.164 | 0.217 | 1.64E-05 |
| Ctbp1 | 7.09E-10 | -0.51129 | 0.262 | 0.316 | 1.67E-05 |
| Gna13 | 7.29E-10 | -0.38435 | 0.267 | 0.326 | 1.72E-05 |
| Papola | 7.4E-10 | -0.34216 | 0.386 | 0.445 | 1.75E-05 |
| Wapl | 7.8E-10 | -0.43241 | 0.348 | 0.402 | 1.84E-05 |
| Clcn4 | 8.2E-10 | -0.34739 | 0.35 | 0.406 | 1.94E-05 |
| Fundc1 | 8.48E-10 | -0.54345 | 0.153 | 0.205 | 2E-05 |
| Rpl12 | 8.8E-10 | 0.518188 | 0.684 | 0.659 | 2.08E-05 |
| Wls | 9.99E-10 | -0.27852 | 0.382 | 0.441 | 2.36E-05 |
| Ufsp2 | 1E-09 | -0.40439 | 0.285 | 0.345 | 2.37E-05 |
| Slc35a5 | 1.01E-09 | -0.43751 | 0.166 | 0.219 | 2.38E-05 |
| Gm9794 | 1.01E-09 | 0.338278 | 0.639 | 0.606 | 2.39E-05 |
| Ncln | 1.09E-09 | -0.55124 | 0.273 | 0.326 | 2.57E-05 |
| Vamp3 | 1.14E-09 | -0.47163 | 0.288 | 0.347 | 2.68E-05 |
| Brox | 1.17E-09 | -0.2864 | 0.161 | 0.215 | 2.75E-05 |
| Mapk14 | 1.17E-09 | -0.31874 | 0.322 | 0.382 | 2.75E-05 |
| Vbp1 | 1.21E-09 | -0.42705 | 0.173 | 0.225 | 2.85E-05 |
| Snd1 | 1.29E-09 | -0.3042 | 0.205 | 0.261 | 3.04E-05 |
| Vhl | 1.35E-09 | -0.45063 | 0.183 | 0.237 | 3.18E-05 |
| Smpd1 | 1.36E-09 | -0.75563 | 0.189 | 0.238 | 3.2E-05 |
| Rnf5 | 1.39E-09 | -0.28985 | 0.199 | 0.259 | 3.28E-05 |
| Dip2b | 1.43E-09 | -0.27922 | 0.54 | 0.603 | 3.37E-05 |
| Nmt1 | 1.52E-09 | -0.27073 | 0.533 | 0.587 | 3.6E-05 |
| Rpl9 | 1.59E-09 | 0.529351 | 0.757 | 0.75 | 3.75E-05 |
| Cap1 | 1.64E-09 | -0.30273 | 0.408 | 0.47 | 3.87E-05 |
| Olfr77 | 1.69E-09 | 0.288002 | 0.346 | 0.294 | 3.98E-05 |
| Plxna4 | 1.71E-09 | -0.31348 | 0.173 | 0.227 | 4.03E-05 |
| Cab39 | 1.76E-09 | -0.3584 | 0.258 | 0.317 | 4.15E-05 |
| Zfp207 | 1.77E-09 | -0.27693 | 0.544 | 0.598 | 4.18E-05 |
| Prag1 | 1.85E-09 | -0.60962 | 0.195 | 0.25 | 4.35E-05 |
| Ddi2 | 1.85E-09 | -0.25005 | 0.327 | 0.385 | 4.37E-05 |
| Foxj2 | 1.88E-09 | -0.34028 | 0.171 | 0.224 | 4.44E-05 |
| Rps18-ps3 | 1.92E-09 | 0.276763 | 0.59 | 0.556 | 4.54E-05 |
| Rpl15 | 1.95E-09 | 0.517509 | 0.678 | 0.672 | 4.59E-05 |
| Dhx15 | 2E-09 | -0.30127 | 0.207 | 0.266 | 4.73E-05 |
| Copb2 | 2.18E-09 | -0.37781 | 0.175 | 0.227 | 5.13E-05 |
| Crat | 2.18E-09 | -0.44918 | 0.207 | 0.262 | 5.15E-05 |
| Tnfaip1 | 2.24E-09 | -0.6737 | 0.177 | 0.228 | 5.29E-05 |
| Tm9sf1 | 2.29E-09 | -0.69444 | 0.178 | 0.227 | 5.39E-05 |

| | | | | | |
|---|---|---|---|---|---|
| Rpl39 | 2.3E-09 | 0.552915 | 0.711 | 0.699 | 5.41E-05 |
| Pla2g4a | 2.35E-09 | -0.70437 | 0.157 | 0.205 | 5.54E-05 |
| Cep68 | 2.36E-09 | -0.36037 | 0.257 | 0.32 | 5.56E-05 |
| Gm2000 | 2.54E-09 | 0.407768 | 0.594 | 0.571 | 5.99E-05 |
| Vwa5a | 2.55E-09 | -0.41012 | 0.17 | 0.223 | 6.01E-05 |
| Med22 | 2.59E-09 | -0.26989 | 0.175 | 0.229 | 6.12E-05 |
| Rpl37rt | 2.72E-09 | 0.566981 | 0.669 | 0.635 | 6.42E-05 |
| Naglu | 3.23E-09 | -0.28472 | 0.415 | 0.467 | 7.62E-05 |
| Actr1a | 3.29E-09 | -0.44816 | 0.277 | 0.331 | 7.76E-05 |
| Cdc40 | 3.31E-09 | -0.33598 | 0.169 | 0.224 | 7.8E-05 |
| Myd88 | 3.42E-09 | -0.30562 | 0.21 | 0.265 | 8.07E-05 |
| Ywhab | 3.47E-09 | -0.25978 | 0.498 | 0.554 | 8.18E-05 |
| Ccdc93 | 3.6E-09 | -0.40254 | 0.227 | 0.284 | 8.5E-05 |
| Acss1 | 3.74E-09 | -0.31962 | 0.192 | 0.247 | 8.81E-05 |
| Ppm1g | 4.27E-09 | -0.45991 | 0.251 | 0.306 | 0.000101 |
| Ppp2ca | 4.95E-09 | -0.27541 | 0.47 | 0.517 | 0.000117 |
| Rpl17-ps3 | 5.08E-09 | 0.354924 | 0.42 | 0.375 | 0.00012 |
| Rpl18 | 5.2E-09 | 0.46665 | 0.785 | 0.799 | 0.000123 |
| B3galnt1 | 5.48E-09 | -0.51038 | 0.161 | 0.209 | 0.000129 |
| Rps3a1 | 5.7E-09 | 0.4005 | 0.823 | 0.837 | 0.000135 |
| Mpi | 5.78E-09 | -0.32456 | 0.238 | 0.298 | 0.000136 |
| Zfp266 | 5.98E-09 | -0.2809 | 0.174 | 0.225 | 0.000141 |
| Yod1 | 6.57E-09 | -0.62242 | 0.155 | 0.203 | 0.000155 |
| Nfkb1 | 6.63E-09 | -0.31865 | 0.428 | 0.483 | 0.000156 |
| Golga7 | 6.63E-09 | -0.44298 | 0.181 | 0.231 | 0.000156 |
| Plcg2 | 6.81E-09 | -0.30104 | 0.322 | 0.379 | 0.000161 |
| Mknk1 | 6.82E-09 | -0.38352 | 0.449 | 0.493 | 0.000161 |
| Trafd1 | 6.83E-09 | -0.36434 | 0.293 | 0.347 | 0.000161 |
| Cct6a | 6.86E-09 | -0.26709 | 0.308 | 0.363 | 0.000162 |
| Naa60 | 7.14E-09 | -0.48074 | 0.204 | 0.255 | 0.000168 |
| Rpl31 | 7.86E-09 | 0.410098 | 0.78 | 0.785 | 0.000185 |
| Csnk1d | 8.35E-09 | -0.32854 | 0.295 | 0.351 | 0.000197 |
| Znrf1 | 8.57E-09 | -0.36114 | 0.177 | 0.228 | 0.000202 |
| Gm24187 | 8.6E-09 | 0.307684 | 0.837 | 0.832 | 0.000203 |
| Clcn3 | 8.89E-09 | -0.46915 | 0.204 | 0.255 | 0.00021 |
| Mkln1 | 9.05E-09 | -0.33941 | 0.362 | 0.413 | 0.000214 |
| Gm10925 | 1.04E-08 | 0.412057 | 0.806 | 0.823 | 0.000246 |
| Filip1l | 1.07E-08 | -0.29945 | 0.24 | 0.295 | 0.000254 |
| Plcg1 | 1.13E-08 | -0.42748 | 0.155 | 0.204 | 0.000266 |
| Eloc | 1.13E-08 | -0.35771 | 0.239 | 0.294 | 0.000267 |
| Tmx2 | 1.15E-08 | -0.36153 | 0.22 | 0.273 | 0.000272 |
| Rnf4 | 1.18E-08 | -0.28795 | 0.183 | 0.235 | 0.000278 |
| Snx2 | 1.2E-08 | -0.29052 | 0.349 | 0.406 | 0.000284 |
| Zdhhc14 | 1.22E-08 | -0.47141 | 0.153 | 0.201 | 0.000287 |
| Snta1 | 1.23E-08 | -0.28108 | 0.262 | 0.319 | 0.000291 |
| Smox | 1.27E-08 | -0.3684 | 0.266 | 0.319 | 0.0003 |
| Slc39a7 | 1.28E-08 | -0.36267 | 0.236 | 0.287 | 0.000302 |

| | | | | | |
|---|---|---|---|---|---|
| Cybc1 | 1.31E-08 | -0.35295 | 0.352 | 0.4 | 0.00031 |
| Mtmr2 | 1.46E-08 | -0.32837 | 0.181 | 0.233 | 0.000344 |
| Slc2a1 | 1.47E-08 | -0.44474 | 0.168 | 0.217 | 0.000346 |
| Ip6k1 | 1.58E-08 | -0.25646 | 0.28 | 0.338 | 0.000372 |
| 2410006H1 | 1.62E-08 | 0.860822 | 0.242 | 0.198 | 0.000383 |
| Metap2 | 1.7E-08 | -0.34212 | 0.314 | 0.372 | 0.000401 |
| Ccdc50 | 1.76E-08 | -0.34969 | 0.367 | 0.422 | 0.000415 |
| Cnot2 | 1.87E-08 | -0.34993 | 0.274 | 0.326 | 0.000441 |
| Slc4a7 | 1.9E-08 | -0.70673 | 0.189 | 0.236 | 0.000448 |
| Rbm7 | 1.94E-08 | -0.34786 | 0.228 | 0.279 | 0.000457 |
| Kat7 | 1.97E-08 | -0.39246 | 0.187 | 0.236 | 0.000465 |
| Gm12184 | 1.98E-08 | -0.33623 | 0.2 | 0.256 | 0.000468 |
| Nrbp1 | 2.02E-08 | -0.50182 | 0.183 | 0.23 | 0.000476 |
| Nono | 2.03E-08 | -0.33418 | 0.292 | 0.344 | 0.000479 |
| D1Ertd622 | 2.07E-08 | -0.29215 | 0.204 | 0.255 | 0.000489 |
| Fam32a | 2.09E-08 | -0.2973 | 0.203 | 0.254 | 0.000494 |
| Aco2 | 2.12E-08 | -0.49512 | 0.226 | 0.277 | 0.0005 |
| Rpl19-ps11 | 2.14E-08 | 0.306757 | 0.618 | 0.589 | 0.000506 |
| Fuca2 | 2.22E-08 | -0.25139 | 0.337 | 0.388 | 0.000524 |
| Cdc42se1 | 2.24E-08 | -0.26205 | 0.324 | 0.379 | 0.00053 |
| Gm8995 | 2.25E-08 | 0.551657 | 0.229 | 0.182 | 0.000531 |
| Rad23a | 2.25E-08 | -0.37774 | 0.194 | 0.243 | 0.000531 |
| Strap | 2.56E-08 | -0.45045 | 0.162 | 0.208 | 0.000604 |
| Flcn | 2.63E-08 | -0.31162 | 0.299 | 0.353 | 0.00062 |
| Mbnl2 | 2.75E-08 | -0.35849 | 0.387 | 0.44 | 0.000648 |
| Slc35f6 | 2.77E-08 | -0.28719 | 0.212 | 0.264 | 0.000653 |
| P3h2 | 2.89E-08 | -0.34134 | 0.191 | 0.239 | 0.000681 |
| Tmx1 | 2.92E-08 | -0.30159 | 0.251 | 0.302 | 0.000689 |
| Mknk2 | 2.94E-08 | -0.39333 | 0.184 | 0.233 | 0.000692 |
| Mgrn1 | 2.95E-08 | -0.41811 | 0.296 | 0.35 | 0.000697 |
| Pik3ap1 | 3.01E-08 | -0.31154 | 0.209 | 0.262 | 0.000711 |
| Rpl7 | 3.12E-08 | 0.405621 | 0.788 | 0.792 | 0.000737 |
| Rsl24d1 | 3.12E-08 | -0.44758 | 0.165 | 0.214 | 0.000737 |
| mt-Cytb | 3.21E-08 | 0.384532 | 0.951 | 0.967 | 0.000756 |
| Psmd2 | 3.26E-08 | -0.54631 | 0.251 | 0.297 | 0.000769 |
| Ncbp3 | 3.33E-08 | -0.34341 | 0.19 | 0.239 | 0.000786 |
| Ss18 | 3.44E-08 | -0.51221 | 0.192 | 0.237 | 0.000811 |
| Bst2 | 3.6E-08 | 0.701481 | 0.412 | 0.369 | 0.00085 |
| Stau1 | 3.67E-08 | -0.48085 | 0.269 | 0.318 | 0.000865 |
| Npc1 | 3.98E-08 | -0.33263 | 0.321 | 0.38 | 0.000938 |
| Rpl11 | 4.15E-08 | 0.467441 | 0.719 | 0.713 | 0.00098 |
| Srp72 | 4.44E-08 | -0.35258 | 0.259 | 0.312 | 0.001047 |
| Slc12a2 | 4.58E-08 | -0.25353 | 0.229 | 0.281 | 0.001081 |
| Tra2b | 4.8E-08 | -0.26916 | 0.541 | 0.594 | 0.001132 |
| Mesd | 5.04E-08 | -0.33783 | 0.245 | 0.297 | 0.001189 |
| Gaa | 5.13E-08 | -0.32226 | 0.314 | 0.365 | 0.001211 |
| Gatad1 | 5.21E-08 | -0.30629 | 0.199 | 0.249 | 0.001228 |

| | | | | | |
|---|---|---|---|---|---|
| Abhd16a | 5.45E-08 | -0.44651 | 0.207 | 0.254 | 0.001285 |
| Pak1 | 5.48E-08 | -0.29514 | 0.187 | 0.238 | 0.001294 |
| Tfg | 5.9E-08 | -0.45063 | 0.225 | 0.272 | 0.001391 |
| Hjurp | 6E-08 | -0.4664 | 0.175 | 0.222 | 0.001416 |
| Slc46a1 | 6.45E-08 | -0.25148 | 0.408 | 0.455 | 0.001521 |
| 9530068E0 | 6.52E-08 | -0.37227 | 0.349 | 0.391 | 0.001539 |
| Glyr1 | 6.72E-08 | -0.40509 | 0.211 | 0.261 | 0.001586 |
| Tlr3 | 6.78E-08 | -0.25849 | 0.198 | 0.25 | 0.001599 |
| Eif3b | 7.27E-08 | -0.47491 | 0.219 | 0.266 | 0.001715 |
| Rpl28 | 7.76E-08 | 0.477668 | 0.723 | 0.723 | 0.00183 |
| Ugt1a7c | 7.77E-08 | -0.35076 | 0.267 | 0.317 | 0.001832 |
| Rpsa-ps10 | 8.02E-08 | 0.352871 | 0.563 | 0.532 | 0.001892 |
| Exosc10 | 8.54E-08 | -0.31495 | 0.169 | 0.216 | 0.002015 |
| Gbp7 | 8.79E-08 | -0.56104 | 0.228 | 0.275 | 0.002073 |
| Rpl35a | 9.12E-08 | 0.462951 | 0.729 | 0.732 | 0.002151 |
| Spred1 | 9.29E-08 | -0.45165 | 0.184 | 0.23 | 0.002192 |
| Gas5 | 9.3E-08 | 0.665367 | 0.378 | 0.333 | 0.002194 |
| Rps17 | 9.34E-08 | 0.440007 | 0.722 | 0.719 | 0.002203 |
| mt-Rnr1 | 1.01E-07 | 0.396671 | 0.872 | 0.888 | 0.002378 |
| Rtf2 | 1.03E-07 | -0.27885 | 0.253 | 0.305 | 0.00244 |
| D6Wsu163 | 1.05E-07 | -0.53641 | 0.155 | 0.2 | 0.002471 |
| Rps3 | 1.05E-07 | 0.317565 | 0.894 | 0.897 | 0.002479 |
| Tcp1 | 1.07E-07 | -0.30186 | 0.221 | 0.27 | 0.002526 |
| Atxn1 | 1.16E-07 | -0.44682 | 0.176 | 0.222 | 0.002736 |
| Rpl10a-ps1 | 1.24E-07 | 0.425481 | 0.557 | 0.528 | 0.002921 |
| Copz1 | 1.25E-07 | -0.36434 | 0.236 | 0.284 | 0.002937 |
| Tmem230 | 1.3E-07 | -0.32064 | 0.156 | 0.2 | 0.00307 |
| Gm3362 | 1.35E-07 | 0.488994 | 0.397 | 0.356 | 0.003184 |
| Rps25 | 1.41E-07 | 0.333881 | 0.845 | 0.852 | 0.003317 |
| Sdha | 1.41E-07 | -0.42765 | 0.372 | 0.409 | 0.003329 |
| Peli1 | 1.48E-07 | -0.29885 | 0.395 | 0.439 | 0.003479 |
| Tbl1xr1 | 1.51E-07 | -0.4895 | 0.201 | 0.248 | 0.003573 |
| Arf3 | 1.56E-07 | -0.44836 | 0.185 | 0.231 | 0.003676 |
| Dars | 1.56E-07 | -0.52494 | 0.177 | 0.22 | 0.00369 |
| Gm6204 | 1.64E-07 | 0.260001 | 0.609 | 0.582 | 0.003879 |
| Kpnb1 | 1.66E-07 | -0.4255 | 0.224 | 0.272 | 0.003914 |
| Klhdc3 | 1.76E-07 | -0.27583 | 0.174 | 0.219 | 0.00414 |
| Rpl22 | 1.84E-07 | 0.460664 | 0.553 | 0.517 | 0.004333 |
| Sumf1 | 1.95E-07 | -0.54205 | 0.194 | 0.237 | 0.004603 |
| Pdk1 | 1.96E-07 | -0.32708 | 0.248 | 0.3 | 0.00462 |
| Papd4 | 1.96E-07 | -0.33346 | 0.158 | 0.203 | 0.004629 |
| Rpl26 | 1.97E-07 | 0.468881 | 0.752 | 0.763 | 0.004642 |
| Lifr | 1.99E-07 | -0.34355 | 0.274 | 0.326 | 0.004683 |
| Hacd3 | 2.02E-07 | -0.31214 | 0.26 | 0.308 | 0.004762 |
| Rpl23a-ps3 | 2.2E-07 | 0.284964 | 0.499 | 0.467 | 0.005187 |
| Rragc | 2.25E-07 | -0.39992 | 0.229 | 0.275 | 0.005306 |
| Ggps1 | 2.25E-07 | -0.32116 | 0.207 | 0.256 | 0.005309 |

| | | | | | |
|---|---|---|---|---|---|
| Xpo6 | 2.34E-07 | -0.35725 | 0.174 | 0.221 | 0.005511 |
| Rsl1d1 | 2.34E-07 | -0.30403 | 0.215 | 0.265 | 0.005512 |
| Zfp703 | 2.43E-07 | -0.37321 | 0.187 | 0.233 | 0.005738 |
| Uba2 | 2.46E-07 | -0.31667 | 0.236 | 0.284 | 0.0058 |
| Usp8 | 2.59E-07 | -0.27802 | 0.266 | 0.318 | 0.006107 |
| Pdcd10 | 2.62E-07 | -0.32059 | 0.206 | 0.252 | 0.00618 |
| Gm28661 | 2.68E-07 | 0.338394 | 0.726 | 0.731 | 0.006324 |
| Tiprl | 2.69E-07 | -0.33153 | 0.17 | 0.216 | 0.00634 |
| Mfsd14b | 2.77E-07 | -0.35774 | 0.164 | 0.206 | 0.006545 |
| Galnt12 | 2.96E-07 | -0.54077 | 0.162 | 0.204 | 0.006987 |
| Rps4x | 2.99E-07 | 0.45957 | 0.794 | 0.802 | 0.007052 |
| Ppp1r9a | 3.3E-07 | -0.30412 | 0.33 | 0.382 | 0.007775 |
| Fndc3a | 3.69E-07 | -0.50155 | 0.193 | 0.237 | 0.008693 |
| Osbpl9 | 4.02E-07 | -0.33336 | 0.228 | 0.273 | 0.009474 |
| Qrich1 | 4.05E-07 | -0.43856 | 0.232 | 0.277 | 0.009565 |
| Gm10095 | 4.95E-07 | 0.440377 | 0.456 | 0.413 | 0.011678 |
| Cep170 | 5.01E-07 | -0.32193 | 0.259 | 0.304 | 0.011808 |
| Zmynd11 | 5.02E-07 | -0.42786 | 0.229 | 0.274 | 0.011835 |
| Clip1 | 5.12E-07 | -0.35309 | 0.316 | 0.362 | 0.012089 |
| Lmbrd1 | 5.27E-07 | -0.27724 | 0.268 | 0.317 | 0.012437 |
| Gnaq | 5.37E-07 | -0.35078 | 0.407 | 0.446 | 0.012669 |
| Pfas | 5.5E-07 | -0.65698 | 0.179 | 0.22 | 0.012972 |
| Capn3 | 5.51E-07 | -0.26403 | 0.375 | 0.422 | 0.013004 |
| Sipa1l1 | 5.85E-07 | -0.55521 | 0.175 | 0.218 | 0.013806 |
| Gm8430 | 5.94E-07 | 0.360906 | 0.526 | 0.496 | 0.014017 |
| Slc35a4 | 6.11E-07 | -0.5276 | 0.17 | 0.212 | 0.014418 |
| Ptpa | 6.18E-07 | -0.25383 | 0.3 | 0.348 | 0.014582 |
| Zfp36 | 6.56E-07 | -0.36094 | 0.431 | 0.476 | 0.015475 |
| Prpf6 | 6.62E-07 | -0.46354 | 0.171 | 0.215 | 0.015612 |
| Ndfip2 | 6.72E-07 | -0.30591 | 0.289 | 0.333 | 0.015852 |
| Ergic1 | 7.02E-07 | -0.37453 | 0.263 | 0.31 | 0.016565 |
| Nelfb | 7.06E-07 | -0.52932 | 0.163 | 0.204 | 0.016656 |
| Ddx3x | 7.07E-07 | -0.3603 | 0.351 | 0.397 | 0.016684 |
| Rnf115 | 7.23E-07 | -0.60973 | 0.166 | 0.207 | 0.017052 |
| Ctns | 7.27E-07 | -0.43447 | 0.178 | 0.22 | 0.017148 |
| Sh3gl1 | 7.42E-07 | -0.29455 | 0.199 | 0.243 | 0.017502 |
| Cd48 | 7.63E-07 | 0.588354 | 0.415 | 0.379 | 0.018002 |
| Tuba1c | 7.91E-07 | 0.32239 | 0.333 | 0.4 | 0.01865 |
| Ppp4r3b | 8.03E-07 | -0.44331 | 0.203 | 0.247 | 0.018941 |
| Chmp1b | 8.34E-07 | -0.36622 | 0.214 | 0.259 | 0.019683 |
| Cdkn1b | 8.64E-07 | -0.26434 | 0.238 | 0.285 | 0.020374 |
| Orai3 | 8.66E-07 | -0.25727 | 0.234 | 0.278 | 0.020438 |
| Cmas | 9.11E-07 | -0.45057 | 0.166 | 0.207 | 0.02148 |
| Tbl1x | 9.11E-07 | -0.4602 | 0.194 | 0.235 | 0.021485 |
| Rab6a | 9.44E-07 | -0.52348 | 0.165 | 0.205 | 0.02226 |
| Cers2 | 9.85E-07 | -0.30699 | 0.416 | 0.454 | 0.023247 |
| Man2a2 | 1.03E-06 | -0.25458 | 0.218 | 0.261 | 0.024273 |

| | | | | | |
|---|---|---|---|---|---|
| Sbno1 | 1.06E-06 | -0.3449 | 0.35 | 0.396 | 0.024907 |
| Paqr7 | 1.09E-06 | -0.25643 | 0.276 | 0.323 | 0.025598 |
| Ganab | 1.2E-06 | -0.27469 | 0.162 | 0.204 | 0.028383 |
| Gm11808 | 1.22E-06 | 0.384032 | 0.504 | 0.473 | 0.028671 |
| Stxbp3 | 1.23E-06 | -0.5292 | 0.168 | 0.206 | 0.029032 |
| Rn7sk | 1.24E-06 | 0.381842 | 0.587 | 0.563 | 0.029173 |
| Bcl10 | 1.28E-06 | -0.28967 | 0.242 | 0.288 | 0.030166 |
| Rpl36-ps12 | 1.28E-06 | 0.390783 | 0.246 | 0.206 | 0.03028 |
| 1600012H( | 1.29E-06 | -0.3678 | 0.168 | 0.21 | 0.030338 |
| Prmt1 | 1.33E-06 | 0.256244 | 0.155 | 0.201 | 0.03127 |
| Tk2 | 1.34E-06 | -0.49866 | 0.209 | 0.251 | 0.031662 |
| Ewsr1 | 1.35E-06 | -0.42212 | 0.355 | 0.395 | 0.031827 |
| Rpl27a | 1.4E-06 | 0.346822 | 0.745 | 0.733 | 0.033037 |
| Hk3 | 1.42E-06 | -0.55109 | 0.209 | 0.251 | 0.033572 |
| Rxrb | 1.6E-06 | -0.27733 | 0.212 | 0.257 | 0.037791 |
| Phc2 | 1.61E-06 | -0.25463 | 0.218 | 0.261 | 0.037893 |
| Gm33858 | 1.61E-06 | -0.26915 | 0.273 | 0.32 | 0.037943 |
| Rps15a | 1.66E-06 | 0.522829 | 0.61 | 0.596 | 0.039263 |
| Slamf9 | 1.67E-06 | 0.596522 | 0.366 | 0.321 | 0.03937 |
| 2510039O1 | 1.69E-06 | -0.42426 | 0.259 | 0.299 | 0.039944 |
| Cs | 1.74E-06 | -0.42477 | 0.194 | 0.235 | 0.041017 |
| Smarca5 | 1.82E-06 | -0.27394 | 0.242 | 0.291 | 0.042824 |
| Usp4 | 1.92E-06 | -0.37458 | 0.284 | 0.329 | 0.045255 |
| Rps16-ps2 | 2.01E-06 | 0.302247 | 0.581 | 0.559 | 0.047461 |
| Morf4l2 | 2.05E-06 | -0.32411 | 0.248 | 0.291 | 0.048243 |
| Mars | 2.12E-06 | -0.38664 | 0.167 | 0.206 | 0.049902 |

#### Supplementry Table 2. eDAM DEGs compared to HS

| GeneName | p_val | avg_log2FC | pct.1 | pct.2 | p_val_adj |
|---|---|---|---|---|---|
| Apoe | 0 | 5.807816 | 0.998 | 0.376 | 0 |
| Fcrls | 2.3E-185 | -1.0043 | 0.853 | 0.975 | 5.4E-181 |
| Lyz2 | 5.6E-174 | 1.804824 | 0.831 | 0.608 | 1.3E-169 |
| H2-D1 | 6.9E-149 | 1.536861 | 0.946 | 0.867 | 1.6E-144 |
| P2ry12 | 5.8E-117 | -0.57154 | 0.997 | 1 | 1.4E-112 |
| Slc2a5 | 4.6E-111 | -0.8212 | 0.789 | 0.944 | 1.1E-106 |
| Cd164 | 3.14E-96 | -0.69844 | 0.811 | 0.946 | 7.4E-92 |
| Cd63 | 2.4E-88 | 1.080009 | 0.837 | 0.684 | 5.65E-84 |
| Fth1 | 3.89E-86 | 0.743374 | 0.995 | 0.993 | 9.18E-82 |
| B2m | 4.67E-86 | 0.733424 | 0.998 | 0.998 | 1.1E-81 |
| H2-K1 | 7.37E-77 | 1.277628 | 0.859 | 0.766 | 1.74E-72 |
| Ctss | 3.55E-73 | 0.492725 | 1 | 1 | 8.38E-69 |
| Ifi27l2a | 5.8E-73 | 2.11211 | 0.221 | 0.074 | 1.37E-68 |
| Cd63-ps | 1.34E-70 | 0.860977 | 0.661 | 0.465 | 3.16E-66 |
| Rpl18a | 5.81E-69 | 0.618053 | 0.988 | 0.986 | 1.37E-64 |
| Tmem119 | 2.3E-59 | -0.42764 | 0.995 | 1 | 5.43E-55 |
| P2ry13 | 6.27E-58 | -0.51959 | 0.922 | 0.977 | 1.48E-53 |
| Rps28 | 3.42E-57 | 0.680078 | 0.943 | 0.892 | 8.07E-53 |
| Rpl32 | 1.41E-55 | 0.714658 | 0.918 | 0.863 | 3.33E-51 |
| Rpl41 | 7.46E-55 | 0.639709 | 0.97 | 0.934 | 1.76E-50 |
| Gpr34 | 1.47E-54 | -0.46419 | 0.977 | 0.992 | 3.47E-50 |
| Ftl1 | 4.58E-54 | 0.587792 | 0.995 | 0.991 | 1.08E-49 |
| Rps9 | 2.39E-52 | 0.593567 | 0.977 | 0.95 | 5.64E-48 |
| Rps27 | 2.89E-51 | 0.623812 | 0.975 | 0.95 | 6.83E-47 |
| Gm10076 | 1.85E-49 | 0.681361 | 0.941 | 0.893 | 4.36E-45 |
| Rplp1 | 2.39E-48 | 0.575806 | 0.964 | 0.952 | 5.63E-44 |
| Rps18 | 2.75E-48 | 0.703674 | 0.891 | 0.833 | 6.49E-44 |
| Upk1b | 2.54E-47 | -1.09015 | 0.29 | 0.473 | 5.98E-43 |
| Rpl37a | 4.4E-47 | 0.593614 | 0.96 | 0.929 | 1.04E-42 |
| Cx3cr1 | 4.62E-47 | -0.30427 | 1 | 1 | 1.09E-42 |
| Selplg | 9.1E-47 | -0.32684 | 0.998 | 1 | 2.15E-42 |
| Rps19 | 3.16E-46 | 0.676841 | 0.91 | 0.863 | 7.46E-42 |
| Rps12 | 5.74E-45 | 0.696595 | 0.878 | 0.812 | 1.35E-40 |
| Stab1 | 2.36E-43 | -0.54638 | 0.646 | 0.822 | 5.56E-39 |
| Rhob | 4.31E-43 | -0.37109 | 0.833 | 0.928 | 1.02E-38 |
| Rps14 | 5.92E-43 | 0.588228 | 0.952 | 0.914 | 1.4E-38 |
| Rplp2 | 6.47E-43 | 0.58314 | 0.929 | 0.895 | 1.53E-38 |
| Lgals3bp | 6.54E-43 | 1.117367 | 0.644 | 0.503 | 1.54E-38 |
| Olfml3 | 1.65E-42 | -0.396 | 0.984 | 0.996 | 3.89E-38 |
| Rps23 | 4.92E-42 | 0.563811 | 0.927 | 0.887 | 1.16E-37 |
| Ftl1-ps1 | 8.11E-41 | 0.607057 | 0.918 | 0.874 | 1.91E-36 |
| Rps29 | 1.45E-40 | 0.473485 | 0.986 | 0.972 | 3.43E-36 |
| Rps11 | 2.66E-39 | 0.554083 | 0.939 | 0.911 | 6.28E-35 |

| | | | | | |
|---|---|---|---|---|---|
| Ctsl | 9.09E-39 | 0.488987 | 0.965 | 0.946 | 2.14E-34 |
| Gm4149 | 1.25E-38 | 0.576993 | 0.876 | 0.813 | 2.94E-34 |
| Lrba | 3.12E-38 | -0.81838 | 0.38 | 0.547 | 7.36E-34 |
| Rpl35 | 1.42E-37 | 0.637366 | 0.87 | 0.807 | 3.35E-33 |
| Rps20 | 2.01E-37 | 0.670066 | 0.874 | 0.807 | 4.74E-33 |
| Cd9 | 8.5E-37 | 0.592123 | 0.978 | 0.965 | 2.01E-32 |
| Rpl13 | 1.77E-36 | 0.553917 | 0.956 | 0.938 | 4.17E-32 |
| Rpl37 | 2.11E-36 | 0.562669 | 0.926 | 0.886 | 4.98E-32 |
| Cd52 | 5.91E-36 | 1.146513 | 0.545 | 0.411 | 1.39E-31 |
| Tyrobp | 6.43E-36 | 0.372225 | 0.994 | 0.996 | 1.52E-31 |
| Fau | 9.27E-35 | 0.508396 | 0.934 | 0.899 | 2.19E-30 |
| Rpl21 | 2.99E-34 | 0.545979 | 0.936 | 0.917 | 7.05E-30 |
| Gm15427 | 3.58E-34 | 0.638829 | 0.964 | 0.942 | 8.43E-30 |
| Hspa5 | 1.38E-33 | -0.46831 | 0.784 | 0.866 | 3.25E-29 |
| Ctsb | 1.41E-33 | 0.398705 | 0.999 | 0.999 | 3.33E-29 |
| Dusp6 | 1.93E-33 | -0.54537 | 0.602 | 0.731 | 4.56E-29 |
| Rps24 | 2.98E-33 | 0.602847 | 0.873 | 0.818 | 7.02E-29 |
| Rpl30 | 3.18E-33 | 0.607822 | 0.901 | 0.858 | 7.51E-29 |
| Txnip | 5.38E-33 | -0.57297 | 0.566 | 0.699 | 1.27E-28 |
| Adgrg1 | 6.42E-33 | -0.33994 | 0.795 | 0.905 | 1.52E-28 |
| Rpl23 | 8.84E-33 | 0.687752 | 0.874 | 0.82 | 2.09E-28 |
| Npc2 | 1.63E-32 | 0.645434 | 0.879 | 0.82 | 3.84E-28 |
| Rpl36 | 1.66E-31 | 0.616679 | 0.865 | 0.799 | 3.92E-27 |
| Ctsh | 2.16E-31 | 0.520507 | 0.958 | 0.947 | 5.1E-27 |
| Ifitm3 | 4E-30 | 2.415055 | 0.218 | 0.121 | 9.43E-26 |
| C1qb | 7.52E-30 | 0.282315 | 1 | 1 | 1.77E-25 |
| Ecscr | 2.45E-29 | -0.38762 | 0.554 | 0.706 | 5.77E-25 |
| Gm14303 | 3.49E-29 | 0.449009 | 0.833 | 0.753 | 8.24E-25 |
| Gm3511 | 7.41E-29 | 0.517944 | 0.654 | 0.534 | 1.75E-24 |
| Golm1 | 8.66E-29 | -0.39429 | 0.773 | 0.877 | 2.04E-24 |
| Rpl17 | 1.06E-28 | 0.662539 | 0.834 | 0.778 | 2.51E-24 |
| Fscn1 | 1.16E-28 | -0.34581 | 0.742 | 0.859 | 2.73E-24 |
| Ivns1abp | 1.96E-28 | -0.37659 | 0.759 | 0.852 | 4.62E-24 |
| Lsp1 | 2.25E-28 | 1.112383 | 0.407 | 0.284 | 5.3E-24 |
| Ptgs1 | 5.79E-28 | -0.39392 | 0.888 | 0.946 | 1.36E-23 |
| Rpl23a | 6.49E-28 | 0.567711 | 0.808 | 0.745 | 1.53E-23 |
| Gm9843 | 1.78E-27 | 0.577108 | 0.88 | 0.834 | 4.21E-23 |
| Cmtm6 | 2.04E-27 | -0.33906 | 0.738 | 0.854 | 4.81E-23 |
| Ftl2-ps | 5.47E-27 | 0.500127 | 0.948 | 0.914 | 1.29E-22 |
| Ubc | 5.94E-27 | -0.31111 | 0.946 | 0.974 | 1.4E-22 |
| Rps16 | 7.35E-27 | 0.535066 | 0.815 | 0.76 | 1.73E-22 |
| Slc40a1 | 9.01E-27 | -0.49619 | 0.378 | 0.522 | 2.13E-22 |
| Fam102b | 1.78E-26 | -0.51314 | 0.543 | 0.665 | 4.2E-22 |
| Slc7a8 | 2.18E-26 | -0.36368 | 0.589 | 0.723 | 5.15E-22 |
| Ltc4s | 2.32E-26 | -0.35856 | 0.753 | 0.867 | 5.46E-22 |
| Pmepa1 | 1.01E-25 | -0.36783 | 0.745 | 0.845 | 2.38E-21 |
| Pdia3 | 1.62E-25 | -0.32413 | 0.917 | 0.961 | 3.82E-21 |

| | | | | | |
|---|---|---|---|---|---|
| Rps27rt | 3.69E-25 | 0.519782 | 0.783 | 0.687 | 8.71E-21 |
| Cd74 | 4.33E-25 | 2.274675 | 0.245 | 0.153 | 1.02E-20 |
| Rpl10a | 4.86E-25 | 0.739517 | 0.718 | 0.643 | 1.15E-20 |
| Rpl19 | 4.96E-25 | 0.531116 | 0.842 | 0.794 | 1.17E-20 |
| Bank1 | 8.97E-25 | -1.27763 | 0.095 | 0.2 | 2.12E-20 |
| Scoc | 1.01E-24 | -0.36941 | 0.725 | 0.831 | 2.37E-20 |
| Rps25 | 1.27E-24 | 0.505263 | 0.888 | 0.852 | 2.99E-20 |
| Rps19-ps6 | 1.45E-24 | 0.430895 | 0.716 | 0.618 | 3.42E-20 |
| Gpr165 | 4E-24 | -0.94267 | 0.151 | 0.267 | 9.43E-20 |
| Rpl34 | 4.18E-24 | 0.528336 | 0.828 | 0.778 | 9.87E-20 |
| Rpsa | 4.88E-24 | 0.6468 | 0.82 | 0.774 | 1.15E-19 |
| Gm10443 | 9.95E-24 | 0.446595 | 0.479 | 0.37 | 2.35E-19 |
| Gm11361 | 1.04E-23 | 0.499053 | 0.542 | 0.434 | 2.45E-19 |
| Tsc22d4 | 1.21E-23 | -0.38635 | 0.548 | 0.667 | 2.86E-19 |
| Entpd1 | 1.25E-23 | -0.29287 | 0.874 | 0.94 | 2.95E-19 |
| Tmem204 | 1.66E-23 | -1.07401 | 0.117 | 0.225 | 3.92E-19 |
| 1700017B0 | 1.98E-23 | -0.41808 | 0.575 | 0.694 | 4.68E-19 |
| Maf | 2.59E-23 | -0.41605 | 0.832 | 0.897 | 6.11E-19 |
| Fam110a | 3.14E-23 | -0.69123 | 0.237 | 0.364 | 7.41E-19 |
| Slco2b1 | 4.91E-23 | -0.32084 | 0.877 | 0.937 | 1.16E-18 |
| Rps7 | 1.15E-22 | 0.601799 | 0.822 | 0.776 | 2.71E-18 |
| Gm29216 | 1.63E-22 | 0.404362 | 0.806 | 0.759 | 3.84E-18 |
| Cyth4 | 1.82E-22 | -0.35966 | 0.846 | 0.894 | 4.28E-18 |
| Rps26 | 1.85E-22 | 0.613712 | 0.729 | 0.664 | 4.37E-18 |
| AU020206 | 2.78E-22 | 0.793488 | 0.581 | 0.473 | 6.56E-18 |
| Rps27a | 4.3E-22 | 0.56173 | 0.826 | 0.78 | 1.02E-17 |
| Rps17 | 4.74E-22 | 0.504023 | 0.78 | 0.719 | 1.12E-17 |
| Elmo1 | 5.23E-22 | -0.37534 | 0.609 | 0.724 | 1.23E-17 |
| Plod1 | 5.37E-22 | -0.54417 | 0.516 | 0.628 | 1.27E-17 |
| Rps21 | 7.36E-22 | 0.559969 | 0.821 | 0.762 | 1.74E-17 |
| Rpl13a | 7.79E-22 | 0.543598 | 0.866 | 0.824 | 1.84E-17 |
| Uba52 | 9.65E-22 | 0.579228 | 0.802 | 0.721 | 2.28E-17 |
| Rps13 | 1.88E-21 | 0.587589 | 0.807 | 0.766 | 4.42E-17 |
| Rps5 | 2.25E-21 | 0.643484 | 0.767 | 0.718 | 5.31E-17 |
| Ddit4 | 2.49E-21 | -0.8531 | 0.143 | 0.25 | 5.88E-17 |
| Ctsz | 3.51E-21 | 0.395156 | 0.992 | 0.993 | 8.28E-17 |
| Rpl14 | 8.73E-21 | 0.545975 | 0.813 | 0.756 | 2.06E-16 |
| Mertk | 1.62E-20 | -0.34848 | 0.81 | 0.872 | 3.83E-16 |
| Gm10736 | 1.69E-20 | 0.43217 | 0.594 | 0.489 | 3.99E-16 |
| Hpgd | 2.21E-20 | -0.36519 | 0.64 | 0.746 | 5.21E-16 |
| Fermt3 | 3.41E-20 | -0.40926 | 0.68 | 0.773 | 8.03E-16 |
| Gm2000 | 3.54E-20 | 0.552655 | 0.655 | 0.571 | 8.35E-16 |
| Tm9sf2 | 5.02E-20 | -0.29831 | 0.672 | 0.769 | 1.18E-15 |
| Nav3 | 5.06E-20 | -0.37278 | 0.541 | 0.656 | 1.19E-15 |
| Rpl31 | 5.18E-20 | 0.494449 | 0.823 | 0.785 | 1.22E-15 |
| Rplp0 | 5.53E-20 | 0.499567 | 0.912 | 0.89 | 1.3E-15 |
| Gm28437 | 6.29E-20 | 0.543945 | 0.855 | 0.825 | 1.48E-15 |

| | | | | | |
|---|---|---|---|---|---|
| Gm6977 | 6.67E-20 | 0.610189 | 0.579 | 0.479 | 1.57E-15 |
| Tpt1 | 6.69E-20 | 0.505883 | 0.895 | 0.878 | 1.58E-15 |
| Nrros | 6.72E-20 | -0.49649 | 0.544 | 0.64 | 1.58E-15 |
| Gm10132 | 9.13E-20 | 0.663748 | 0.654 | 0.555 | 2.15E-15 |
| Rps3a1 | 9.32E-20 | 0.521747 | 0.866 | 0.837 | 2.2E-15 |
| Rpl12 | 1.05E-19 | 0.610196 | 0.731 | 0.659 | 2.48E-15 |
| Hspa8 | 1.07E-19 | -0.26727 | 0.904 | 0.945 | 2.52E-15 |
| Rps16-ps2 | 1.98E-19 | 0.474666 | 0.65 | 0.559 | 4.66E-15 |
| Rap1b | 2.03E-19 | -0.40235 | 0.697 | 0.773 | 4.79E-15 |
| G530011O( | 2.06E-19 | -0.50715 | 0.27 | 0.385 | 4.85E-15 |
| Pnp | 2.07E-19 | -0.3562 | 0.478 | 0.591 | 4.89E-15 |
| Rpl10a-ps1 | 2.17E-19 | 0.614863 | 0.618 | 0.528 | 5.12E-15 |
| Arhgap5 | 2.71E-19 | -0.34723 | 0.666 | 0.761 | 6.4E-15 |
| St3gal5 | 2.77E-19 | -0.53435 | 0.504 | 0.611 | 6.55E-15 |
| Numb | 2.91E-19 | -0.35182 | 0.473 | 0.592 | 6.86E-15 |
| Rps10 | 3.62E-19 | 0.557312 | 0.836 | 0.786 | 8.55E-15 |
| Mtss1 | 6.57E-19 | -0.71103 | 0.262 | 0.37 | 1.55E-14 |
| Gm7331 | 6.93E-19 | 0.536861 | 0.624 | 0.528 | 1.63E-14 |
| Scamp5 | 7.59E-19 | -0.43997 | 0.462 | 0.578 | 1.79E-14 |
| Gm5905 | 1.09E-18 | 0.429133 | 0.773 | 0.693 | 2.58E-14 |
| Chsy1 | 1.13E-18 | -0.53693 | 0.376 | 0.496 | 2.67E-14 |
| Pla2g15 | 1.2E-18 | -0.3417 | 0.737 | 0.803 | 2.82E-14 |
| Gal3st4 | 1.24E-18 | -0.26567 | 0.461 | 0.58 | 2.92E-14 |
| Qk | 1.65E-18 | -0.2788 | 0.894 | 0.936 | 3.89E-14 |
| Rps4x | 1.82E-18 | 0.562843 | 0.843 | 0.802 | 4.29E-14 |
| 2410006H1 | 1.83E-18 | 1.205602 | 0.282 | 0.198 | 4.32E-14 |
| Gm6204 | 2.15E-18 | 0.336497 | 0.671 | 0.582 | 5.06E-14 |
| Plxdc2 | 2.4E-18 | -0.27254 | 0.87 | 0.931 | 5.65E-14 |
| Rab3il1 | 2.42E-18 | -0.36088 | 0.758 | 0.828 | 5.71E-14 |
| Il16 | 2.57E-18 | -0.50489 | 0.468 | 0.574 | 6.07E-14 |
| Sgk1 | 2.67E-18 | -0.51806 | 0.49 | 0.584 | 6.3E-14 |
| Eef1a1 | 3.19E-18 | 0.338855 | 0.993 | 0.994 | 7.53E-14 |
| Sema4b | 3.65E-18 | -0.90566 | 0.121 | 0.213 | 8.62E-14 |
| mt-Co3 | 6.15E-18 | 0.601713 | 0.925 | 0.902 | 1.45E-13 |
| Rpl38 | 6.62E-18 | 0.496037 | 0.691 | 0.616 | 1.56E-13 |
| Rps18-ps3 | 6.82E-18 | 0.590628 | 0.642 | 0.556 | 1.61E-13 |
| Gas5 | 8.08E-18 | 0.681925 | 0.432 | 0.333 | 1.9E-13 |
| Dusp7 | 1.1E-17 | -0.3554 | 0.482 | 0.588 | 2.59E-13 |
| Gm9794 | 1.39E-17 | 0.436733 | 0.684 | 0.606 | 3.28E-13 |
| Fcrl1 | 1.41E-17 | -0.66808 | 0.187 | 0.287 | 3.33E-13 |
| Mef2a | 3.54E-17 | -0.33669 | 0.743 | 0.81 | 8.36E-13 |
| Gm15710 | 3.56E-17 | 0.445267 | 0.789 | 0.723 | 8.39E-13 |
| Rpl19-ps11 | 4.11E-17 | 0.507202 | 0.67 | 0.589 | 9.69E-13 |
| Rps8 | 4.25E-17 | 0.490175 | 0.848 | 0.813 | 1E-12 |
| Hfe | 4.81E-17 | -0.46367 | 0.342 | 0.452 | 1.13E-12 |
| Gm10180 | 5.08E-17 | 0.303988 | 0.482 | 0.389 | 1.2E-12 |
| Ms4a6b | 5.09E-17 | -0.41509 | 0.262 | 0.372 | 1.2E-12 |

| | | | | | |
|---|---|---|---|---|---|
| Rpl39 | 5.76E-17 | 0.563993 | 0.768 | 0.699 | 1.36E-12 |
| Zfas1 | 5.8E-17 | 0.906953 | 0.203 | 0.127 | 1.37E-12 |
| Csnk1e | 5.82E-17 | -0.41796 | 0.521 | 0.622 | 1.37E-12 |
| Adrb2 | 6.82E-17 | -0.3524 | 0.553 | 0.652 | 1.61E-12 |
| Prpsap2 | 8.78E-17 | -0.52001 | 0.254 | 0.359 | 2.07E-12 |
| Gm10073 | 9.03E-17 | 0.457293 | 0.651 | 0.568 | 2.13E-12 |
| Sh2b3 | 1.04E-16 | -0.38562 | 0.341 | 0.449 | 2.45E-12 |
| Gp9 | 1.25E-16 | -0.86298 | 0.139 | 0.228 | 2.96E-12 |
| Ubl3 | 1.31E-16 | -0.39622 | 0.606 | 0.691 | 3.09E-12 |
| Tmem173 | 1.81E-16 | -0.25509 | 0.838 | 0.905 | 4.27E-12 |
| Emp3 | 1.88E-16 | 1.271731 | 0.202 | 0.128 | 4.44E-12 |
| Il10ra | 2.18E-16 | -0.26741 | 0.637 | 0.741 | 5.15E-12 |
| Gm8730 | 2.27E-16 | 0.475489 | 0.776 | 0.723 | 5.36E-12 |
| Xbp1 | 2.32E-16 | -0.35872 | 0.526 | 0.63 | 5.48E-12 |
| Tram1 | 2.61E-16 | -0.34903 | 0.571 | 0.673 | 6.15E-12 |
| Rpl36-ps2 | 3.09E-16 | 0.40614 | 0.565 | 0.478 | 7.3E-12 |
| Prkab1 | 3.15E-16 | -0.67855 | 0.282 | 0.381 | 7.42E-12 |
| Ly86 | 3.75E-16 | 0.28499 | 0.99 | 0.991 | 8.84E-12 |
| mt-Atp6 | 3.93E-16 | 0.524994 | 0.857 | 0.825 | 9.28E-12 |
| Rpl11 | 4.03E-16 | 0.526609 | 0.76 | 0.713 | 9.5E-12 |
| Mylip | 4.32E-16 | -0.41306 | 0.614 | 0.699 | 1.02E-11 |
| Rpsa-ps10 | 5.17E-16 | 0.549392 | 0.611 | 0.532 | 1.22E-11 |
| Rpl10 | 6.32E-16 | 0.3821 | 0.917 | 0.901 | 1.49E-11 |
| Mir692-2 | 6.84E-16 | 0.304181 | 0.393 | 0.301 | 1.61E-11 |
| 0610040J0 | 7.2E-16 | -0.42147 | 0.448 | 0.543 | 1.7E-11 |
| Tspo | 8.15E-16 | 0.914194 | 0.36 | 0.274 | 1.92E-11 |
| Cd83 | 8.34E-16 | 0.778257 | 0.351 | 0.26 | 1.97E-11 |
| Rps2 | 9.04E-16 | 0.542624 | 0.737 | 0.675 | 2.13E-11 |
| Timp2 | 1.07E-15 | 0.436754 | 0.854 | 0.824 | 2.52E-11 |
| Bin1 | 1.28E-15 | -0.27361 | 0.855 | 0.913 | 3.02E-11 |
| Rps3 | 1.31E-15 | 0.404484 | 0.917 | 0.897 | 3.09E-11 |
| Rpl3 | 1.43E-15 | 0.503035 | 0.836 | 0.821 | 3.36E-11 |
| Fcer1g | 1.44E-15 | 0.261685 | 0.99 | 0.993 | 3.4E-11 |
| Gm5805 | 1.52E-15 | 0.457566 | 0.556 | 0.474 | 3.6E-11 |
| Naalad2 | 1.81E-15 | 1.010952 | 0.22 | 0.143 | 4.27E-11 |
| Rnf13 | 2.09E-15 | -0.28278 | 0.711 | 0.776 | 4.93E-11 |
| Rpl27a | 2.15E-15 | 0.446269 | 0.782 | 0.733 | 5.07E-11 |
| Gm15590 | 2.17E-15 | 0.320694 | 0.542 | 0.453 | 5.12E-11 |
| Mgat1 | 2.42E-15 | -0.43625 | 0.484 | 0.574 | 5.7E-11 |
| Gm11942 | 2.7E-15 | 0.330879 | 0.492 | 0.408 | 6.36E-11 |
| Rab1a | 3.02E-15 | -0.43256 | 0.569 | 0.659 | 7.12E-11 |
| Srgap2 | 3.16E-15 | -0.27639 | 0.757 | 0.831 | 7.46E-11 |
| Rps6ka1 | 3.2E-15 | -0.29848 | 0.74 | 0.813 | 7.54E-11 |
| Eef2k | 3.67E-15 | -0.63675 | 0.355 | 0.448 | 8.65E-11 |
| Chst7 | 4.64E-15 | -0.64103 | 0.193 | 0.286 | 1.1E-10 |
| Cttnbp2nl | 4.65E-15 | -0.25126 | 0.652 | 0.755 | 1.1E-10 |
| Rpl15 | 4.9E-15 | 0.506298 | 0.73 | 0.672 | 1.16E-10 |

| | | | | | |
|---|---|---|---|---|---|
| Snx17 | 7.71E-15 | -0.47396 | 0.41 | 0.501 | 1.82E-10 |
| Rpl26 | 7.79E-15 | 0.519142 | 0.797 | 0.763 | 1.84E-10 |
| Tmsb10 | 8.14E-15 | 0.614288 | 0.492 | 0.412 | 1.92E-10 |
| mt-Cytb | 9.27E-15 | 0.45338 | 0.972 | 0.967 | 2.19E-10 |
| Siglece | 9.69E-15 | -0.32246 | 0.273 | 0.372 | 2.29E-10 |
| Gm11478 | 1.04E-14 | 0.464764 | 0.729 | 0.663 | 2.46E-10 |
| Capza2 | 1.43E-14 | -0.26006 | 0.856 | 0.89 | 3.37E-10 |
| Olfr77 | 1.46E-14 | 0.339375 | 0.38 | 0.294 | 3.45E-10 |
| St3gal6 | 1.46E-14 | -0.25501 | 0.722 | 0.804 | 3.46E-10 |
| Ykt6 | 1.73E-14 | -0.38118 | 0.307 | 0.405 | 4.07E-10 |
| Sox4 | 2.27E-14 | -0.5325 | 0.326 | 0.422 | 5.34E-10 |
| Arhgef40 | 2.33E-14 | -0.28897 | 0.367 | 0.469 | 5.51E-10 |
| Rps15 | 2.63E-14 | 0.481105 | 0.776 | 0.737 | 6.21E-10 |
| Cdk5r1 | 3.44E-14 | -0.46909 | 0.191 | 0.283 | 8.1E-10 |
| Rpl7 | 5E-14 | 0.442682 | 0.832 | 0.792 | 1.18E-09 |
| Tnfsf13b | 5.3E-14 | 0.75921 | 0.239 | 0.162 | 1.25E-09 |
| Pdia6 | 5.78E-14 | -0.30399 | 0.665 | 0.736 | 1.36E-09 |
| Arhgap27 | 6.74E-14 | -0.61786 | 0.253 | 0.342 | 1.59E-09 |
| Vav1 | 7.04E-14 | -0.39816 | 0.588 | 0.661 | 1.66E-09 |
| Rpl22 | 7.79E-14 | 0.515192 | 0.594 | 0.517 | 1.84E-09 |
| Vcp | 8.82E-14 | -0.32738 | 0.481 | 0.576 | 2.08E-09 |
| Sipa1 | 1.07E-13 | -0.31297 | 0.537 | 0.635 | 2.52E-09 |
| Gm11808 | 1.17E-13 | 0.487771 | 0.551 | 0.473 | 2.76E-09 |
| Sesn1 | 1.21E-13 | -0.53467 | 0.346 | 0.439 | 2.85E-09 |
| Asb2 | 1.32E-13 | -0.50247 | 0.281 | 0.375 | 3.11E-09 |
| Rps10-ps2 | 1.42E-13 | 0.500144 | 0.646 | 0.571 | 3.35E-09 |
| Bst2 | 1.44E-13 | 0.721695 | 0.449 | 0.369 | 3.4E-09 |
| Rpl28 | 1.7E-13 | 0.437936 | 0.766 | 0.723 | 4E-09 |
| Tmem63a | 1.7E-13 | -0.32602 | 0.371 | 0.469 | 4.01E-09 |
| Rpl17-ps3 | 1.84E-13 | 0.49096 | 0.449 | 0.375 | 4.35E-09 |
| Ccng2 | 2.03E-13 | -0.41963 | 0.273 | 0.365 | 4.79E-09 |
| Capg | 2.27E-13 | 1.372391 | 0.219 | 0.153 | 5.36E-09 |
| Rilpl1 | 2.43E-13 | -0.45863 | 0.239 | 0.328 | 5.73E-09 |
| Btg2 | 2.75E-13 | -0.39241 | 0.499 | 0.584 | 6.49E-09 |
| Gm8430 | 2.79E-13 | 0.537257 | 0.573 | 0.496 | 6.58E-09 |
| Rps15a | 2.82E-13 | 0.616868 | 0.648 | 0.596 | 6.65E-09 |
| Gm7536 | 2.99E-13 | 0.510142 | 0.641 | 0.562 | 7.06E-09 |
| Tifab | 3.04E-13 | -0.38445 | 0.49 | 0.577 | 7.17E-09 |
| Zfp710 | 3.44E-13 | -0.29338 | 0.645 | 0.727 | 8.11E-09 |
| Cyba | 3.76E-13 | 0.296306 | 0.971 | 0.973 | 8.88E-09 |
| Gtf2h2 | 3.87E-13 | -0.29421 | 0.533 | 0.628 | 9.12E-09 |
| Gm10925 | 4.08E-13 | 0.533799 | 0.854 | 0.823 | 9.63E-09 |
| Tgm2 | 4.82E-13 | -0.55005 | 0.188 | 0.27 | 1.14E-08 |
| Rpl36a | 5.02E-13 | 0.590071 | 0.578 | 0.511 | 1.19E-08 |
| Cxxc5 | 6.12E-13 | -0.26235 | 0.613 | 0.707 | 1.44E-08 |
| Rpl14-ps1 | 6.29E-13 | 0.331613 | 0.639 | 0.563 | 1.48E-08 |
| Ncf1 | 7.06E-13 | -0.27865 | 0.658 | 0.735 | 1.66E-08 |

| | | | | | |
|---|---|---|---|---|---|
| Wdr89 | 8.13E-13 | 0.283529 | 0.657 | 0.59 | 1.92E-08 |
| Stard3 | 9.29E-13 | -0.35903 | 0.376 | 0.472 | 2.19E-08 |
| Rpl6 | 1.18E-12 | 0.472154 | 0.79 | 0.753 | 2.78E-08 |
| Gm6277 | 1.2E-12 | -0.84176 | 0.143 | 0.219 | 2.82E-08 |
| mt-Co1 | 1.29E-12 | 0.353254 | 0.989 | 0.985 | 3.04E-08 |
| Ddx5 | 1.37E-12 | -0.30289 | 0.724 | 0.789 | 3.23E-08 |
| Gm3699 | 1.43E-12 | 0.423668 | 0.408 | 0.326 | 3.37E-08 |
| Rpl31-ps8 | 1.5E-12 | 0.407654 | 0.64 | 0.579 | 3.55E-08 |
| Rtn1 | 1.53E-12 | -0.5763 | 0.305 | 0.391 | 3.61E-08 |
| Fgd2 | 1.71E-12 | -0.35134 | 0.687 | 0.747 | 4.04E-08 |
| Atxn7l3b | 1.94E-12 | -0.35091 | 0.395 | 0.486 | 4.58E-08 |
| Gcn1l1 | 2.1E-12 | -0.48607 | 0.37 | 0.464 | 4.95E-08 |
| Slc44a2 | 2.39E-12 | -0.4384 | 0.354 | 0.443 | 5.65E-08 |
| Fez2 | 2.46E-12 | -0.29239 | 0.41 | 0.501 | 5.8E-08 |
| Itga9 | 2.49E-12 | -0.63178 | 0.207 | 0.291 | 5.88E-08 |
| Rpl37rt | 2.58E-12 | 0.275648 | 0.706 | 0.635 | 6.08E-08 |
| Gm12715 | 2.65E-12 | -0.25901 | 0.641 | 0.724 | 6.26E-08 |
| Snx18 | 2.77E-12 | -0.28147 | 0.563 | 0.648 | 6.54E-08 |
| Gm6394 | 2.88E-12 | 0.280948 | 0.575 | 0.489 | 6.8E-08 |
| Tspan14 | 3.23E-12 | -0.36628 | 0.483 | 0.571 | 7.62E-08 |
| Abl1 | 3.61E-12 | -0.47274 | 0.24 | 0.327 | 8.5E-08 |
| Ggta1 | 3.67E-12 | -1.06457 | 0.134 | 0.206 | 8.66E-08 |
| Itpripl1 | 3.78E-12 | -0.27238 | 0.635 | 0.729 | 8.92E-08 |
| Gna15 | 3.87E-12 | -0.28042 | 0.605 | 0.689 | 9.13E-08 |
| Lcp2 | 3.97E-12 | -0.30365 | 0.508 | 0.595 | 9.36E-08 |
| Gm10275 | 4.4E-12 | 0.465213 | 0.465 | 0.388 | 1.04E-07 |
| Eng | 4.53E-12 | -0.33755 | 0.429 | 0.519 | 1.07E-07 |
| Rpl9 | 5.05E-12 | 0.488519 | 0.789 | 0.75 | 1.19E-07 |
| Tuba1a | 5.13E-12 | -0.37489 | 0.526 | 0.61 | 1.21E-07 |
| Nptn | 5.6E-12 | -0.27055 | 0.547 | 0.619 | 1.32E-07 |
| Mlec | 5.78E-12 | -0.27363 | 0.396 | 0.491 | 1.36E-07 |
| Fchsd2 | 5.89E-12 | -0.53385 | 0.342 | 0.429 | 1.39E-07 |
| Hebp1 | 6.03E-12 | 0.885695 | 0.227 | 0.161 | 1.42E-07 |
| Grap | 6.07E-12 | -0.43337 | 0.228 | 0.313 | 1.43E-07 |
| Rcsd1 | 7.56E-12 | -0.40496 | 0.371 | 0.46 | 1.78E-07 |
| Inka1 | 7.73E-12 | -0.31493 | 0.371 | 0.465 | 1.82E-07 |
| Gfm2 | 1.06E-11 | -0.2986 | 0.97 | 0.986 | 2.51E-07 |
| Gtf2a1 | 1.06E-11 | -0.41746 | 0.308 | 0.389 | 2.51E-07 |
| Rpl3-ps1 | 1.15E-11 | 0.443731 | 0.724 | 0.694 | 2.71E-07 |
| Amfr | 1.33E-11 | -0.39494 | 0.36 | 0.444 | 3.14E-07 |
| Gm10800 | 1.34E-11 | 0.325457 | 0.735 | 0.711 | 3.17E-07 |
| Tcf7l2 | 1.47E-11 | -0.28703 | 0.363 | 0.452 | 3.48E-07 |
| Rps12-ps9 | 1.49E-11 | 0.310044 | 0.497 | 0.423 | 3.51E-07 |
| Gm28661 | 1.54E-11 | 0.430396 | 0.751 | 0.731 | 3.64E-07 |
| Zfp652 | 1.75E-11 | -0.33499 | 0.627 | 0.694 | 4.12E-07 |
| Gm12918 | 1.76E-11 | 0.376843 | 0.561 | 0.496 | 4.16E-07 |
| Sla | 1.93E-11 | -0.36941 | 0.463 | 0.545 | 4.54E-07 |

| | | | | | |
|---|---|---|---|---|---|
| Tnfrsf21 | 1.98E-11 | -0.40279 | 0.428 | 0.513 | 4.67E-07 |
| Ccnd1 | 2.1E-11 | -0.38858 | 0.215 | 0.296 | 4.96E-07 |
| Hnrnpf | 2.14E-11 | -0.26787 | 0.52 | 0.596 | 5.05E-07 |
| Rpl35a | 2.2E-11 | 0.405343 | 0.771 | 0.732 | 5.18E-07 |
| Tjp1 | 2.34E-11 | -0.43002 | 0.258 | 0.342 | 5.53E-07 |
| Csnk1a1 | 2.47E-11 | -0.31623 | 0.439 | 0.519 | 5.82E-07 |
| Tmem206 | 2.57E-11 | -0.56056 | 0.273 | 0.353 | 6.06E-07 |
| Slc35c2 | 2.77E-11 | -0.42267 | 0.382 | 0.464 | 6.52E-07 |
| Clstn1 | 2.8E-11 | -0.42656 | 0.35 | 0.432 | 6.61E-07 |
| Ptp4a3 | 2.9E-11 | -0.34245 | 0.452 | 0.531 | 6.84E-07 |
| Cryl1 | 2.91E-11 | -0.39205 | 0.253 | 0.333 | 6.86E-07 |
| Nr3c1 | 3.27E-11 | -0.27307 | 0.437 | 0.528 | 7.71E-07 |
| Gm3362 | 3.43E-11 | 0.546623 | 0.433 | 0.356 | 8.1E-07 |
| Gm10095 | 4.02E-11 | 0.464038 | 0.487 | 0.413 | 9.49E-07 |
| Gas6 | 4.25E-11 | -0.40944 | 0.407 | 0.482 | 1E-06 |
| Pdia4 | 4.32E-11 | -0.49076 | 0.31 | 0.392 | 1.02E-06 |
| Soga1 | 4.46E-11 | -0.26017 | 0.496 | 0.586 | 1.05E-06 |
| Rpl23a-ps3 | 4.55E-11 | 0.399853 | 0.538 | 0.467 | 1.07E-06 |
| Dtx4 | 4.82E-11 | -0.30354 | 0.372 | 0.458 | 1.14E-06 |
| Surf4 | 4.9E-11 | -0.32069 | 0.543 | 0.618 | 1.16E-06 |
| Rgmb | 5.1E-11 | -0.33375 | 0.375 | 0.461 | 1.2E-06 |
| Rpl18 | 5.57E-11 | 0.398661 | 0.815 | 0.799 | 1.31E-06 |
| Mafg | 5.83E-11 | -0.30744 | 0.444 | 0.525 | 1.38E-06 |
| Rpl34-ps1 | 5.89E-11 | 0.313403 | 0.564 | 0.483 | 1.39E-06 |
| Ddx6 | 6.12E-11 | -0.4566 | 0.308 | 0.381 | 1.44E-06 |
| Atp6v0a2 | 7.32E-11 | -0.31982 | 0.348 | 0.432 | 1.73E-06 |
| Tpt1-ps3 | 7.95E-11 | 0.469419 | 0.667 | 0.62 | 1.87E-06 |
| Orai2 | 1.03E-10 | -0.60189 | 0.163 | 0.233 | 2.43E-06 |
| Rps26-ps1 | 1.06E-10 | 0.252252 | 0.392 | 0.317 | 2.5E-06 |
| Rnf167 | 1.25E-10 | -0.33056 | 0.445 | 0.524 | 2.96E-06 |
| Cables1 | 1.27E-10 | -0.43569 | 0.215 | 0.29 | 2.99E-06 |
| Dnajb9 | 1.31E-10 | -0.43321 | 0.256 | 0.333 | 3.09E-06 |
| Rhoh | 1.31E-10 | -0.26416 | 0.392 | 0.48 | 3.1E-06 |
| Fli1 | 1.38E-10 | -0.25749 | 0.569 | 0.645 | 3.26E-06 |
| mt-Rnr1 | 1.47E-10 | 0.477889 | 0.886 | 0.888 | 3.46E-06 |
| Gosr2 | 1.59E-10 | -0.57182 | 0.139 | 0.206 | 3.75E-06 |
| Man1c1 | 1.59E-10 | -0.27884 | 0.439 | 0.521 | 3.75E-06 |
| Cdk6 | 1.95E-10 | -0.29346 | 0.363 | 0.446 | 4.6E-06 |
| Crlf2 | 2.35E-10 | 0.590216 | 0.452 | 0.372 | 5.54E-06 |
| Csf2ra | 2.69E-10 | 1.034774 | 0.331 | 0.266 | 6.36E-06 |
| Gbp7 | 2.73E-10 | -0.56021 | 0.205 | 0.275 | 6.45E-06 |
| Tm6sf1 | 2.99E-10 | -0.39153 | 0.377 | 0.459 | 7.05E-06 |
| Slc3a2 | 3.17E-10 | -0.28332 | 0.674 | 0.733 | 7.47E-06 |
| Akirin1 | 3.73E-10 | -0.30683 | 0.299 | 0.375 | 8.81E-06 |
| Dapp1 | 3.85E-10 | -0.29872 | 0.329 | 0.41 | 9.07E-06 |
| H2-M3 | 4.07E-10 | 0.753843 | 0.248 | 0.184 | 9.59E-06 |
| Gm4294 | 4.16E-10 | 0.27266 | 0.39 | 0.327 | 9.81E-06 |

| | | | | | |
|---|---|---|---|---|---|
| Cd48 | 4.16E-10 | 0.693578 | 0.444 | 0.379 | 9.82E-06 |
| Vcp-rs | 4.29E-10 | -0.31352 | 0.236 | 0.315 | 1.01E-05 |
| mt-Co2 | 4.48E-10 | 0.46669 | 0.863 | 0.84 | 1.06E-05 |
| Rps15a-ps6 | 4.62E-10 | 0.468429 | 0.296 | 0.226 | 1.09E-05 |
| Gm9625 | 5.08E-10 | 0.343923 | 0.539 | 0.477 | 1.2E-05 |
| Rpl24 | 5.22E-10 | 0.507304 | 0.66 | 0.604 | 1.23E-05 |
| Sqor | 5.26E-10 | -0.45375 | 0.244 | 0.319 | 1.24E-05 |
| Rapgef5 | 5.59E-10 | -0.40649 | 0.293 | 0.374 | 1.32E-05 |
| Neat1 | 6.18E-10 | 0.701211 | 0.246 | 0.183 | 1.46E-05 |
| Dnajb11 | 6.58E-10 | -0.44358 | 0.312 | 0.387 | 1.55E-05 |
| Nckap5l | 6.99E-10 | -0.46103 | 0.17 | 0.24 | 1.65E-05 |
| Cdk2ap1 | 7.25E-10 | -0.25962 | 0.343 | 0.43 | 1.71E-05 |
| Rps3a3 | 7.28E-10 | 0.262159 | 0.418 | 0.35 | 1.72E-05 |
| Gm26917 | 7.41E-10 | 0.477451 | 0.943 | 0.951 | 1.75E-05 |
| Gm5148 | 7.59E-10 | 0.282656 | 0.502 | 0.435 | 1.79E-05 |
| Rpl19-ps1 | 7.96E-10 | 0.333577 | 0.275 | 0.212 | 1.88E-05 |
| Rpl22l1 | 8E-10 | 0.593022 | 0.307 | 0.239 | 1.89E-05 |
| Hs6st1 | 8.04E-10 | -0.32158 | 0.242 | 0.317 | 1.9E-05 |
| I830077J02 | 9.39E-10 | -0.39101 | 0.334 | 0.41 | 2.21E-05 |
| Ifi30 | 1.3E-09 | 0.511272 | 0.531 | 0.468 | 3.06E-05 |
| Kif21b | 1.38E-09 | -0.30877 | 0.313 | 0.392 | 3.26E-05 |
| Rps6 | 1.78E-09 | 0.386024 | 0.777 | 0.749 | 4.2E-05 |
| Tnfrsf13b | 1.8E-09 | -0.46127 | 0.26 | 0.331 | 4.24E-05 |
| Stt3a | 1.81E-09 | -0.25409 | 0.509 | 0.576 | 4.26E-05 |
| Sash3 | 1.85E-09 | -0.30983 | 0.333 | 0.412 | 4.37E-05 |
| Lpcat3 | 1.88E-09 | -0.27689 | 0.47 | 0.547 | 4.44E-05 |
| Cib1 | 1.89E-09 | -0.31007 | 0.219 | 0.292 | 4.45E-05 |
| Klf3 | 1.89E-09 | -0.30598 | 0.52 | 0.588 | 4.46E-05 |
| Rptor | 2.08E-09 | -0.26211 | 0.39 | 0.474 | 4.9E-05 |
| Srm | 2.17E-09 | -0.43969 | 0.166 | 0.232 | 5.11E-05 |
| Slc16a6 | 2.3E-09 | -0.27329 | 0.467 | 0.548 | 5.42E-05 |
| Gm8129 | 2.55E-09 | 0.388294 | 0.329 | 0.265 | 6.03E-05 |
| Npl | 2.89E-09 | -0.43977 | 0.159 | 0.224 | 6.82E-05 |
| Lpcat1 | 3.07E-09 | -0.25753 | 0.442 | 0.519 | 7.24E-05 |
| Rpl30-ps1 | 3.14E-09 | 0.276587 | 0.235 | 0.176 | 7.41E-05 |
| Gm6136 | 3.29E-09 | 0.373834 | 0.581 | 0.525 | 7.76E-05 |
| Tm9sf3 | 3.58E-09 | -0.27046 | 0.414 | 0.493 | 8.44E-05 |
| Bnip2 | 3.62E-09 | -0.50825 | 0.239 | 0.307 | 8.54E-05 |
| Mfng | 3.82E-09 | -0.37273 | 0.348 | 0.422 | 9.02E-05 |
| Slc25a37 | 3.91E-09 | -0.5615 | 0.171 | 0.236 | 9.22E-05 |
| Nars | 4.09E-09 | -0.32676 | 0.436 | 0.507 | 9.64E-05 |
| Gm15500 | 4.27E-09 | 0.415405 | 0.652 | 0.601 | 0.000101 |
| Trim8 | 4.28E-09 | -0.3041 | 0.415 | 0.493 | 0.000101 |
| Tcf4 | 4.47E-09 | -0.35998 | 0.572 | 0.634 | 0.000106 |
| Sypl | 4.47E-09 | -0.32516 | 0.39 | 0.463 | 0.000106 |
| Zmiz1 | 4.48E-09 | -0.41922 | 0.58 | 0.642 | 0.000106 |
| Gmpr | 4.74E-09 | -0.49679 | 0.152 | 0.216 | 0.000112 |

| | | | | | |
|---|---|---|---|---|---|
| Cox4i1 | 5.01E-09 | 0.383511 | 0.799 | 0.775 | 0.000118 |
| Rpl13-ps6 | 5.03E-09 | 0.25898 | 0.577 | 0.509 | 0.000119 |
| Cep68 | 5.08E-09 | -0.41487 | 0.246 | 0.32 | 0.00012 |
| Rpl4 | 5.09E-09 | 0.373966 | 0.848 | 0.829 | 0.00012 |
| Parvg | 5.6E-09 | -0.28368 | 0.383 | 0.457 | 0.000132 |
| Gm8186 | 5.91E-09 | 0.500026 | 0.328 | 0.267 | 0.00014 |
| Gm10163 | 6.04E-09 | 0.340743 | 0.482 | 0.421 | 0.000143 |
| Spred1 | 6.1E-09 | -0.7359 | 0.168 | 0.23 | 0.000144 |
| Rin2 | 6.4E-09 | -0.38178 | 0.488 | 0.55 | 0.000151 |
| Tspan13 | 6.82E-09 | -0.32151 | 0.468 | 0.538 | 0.000161 |
| Slc25a45 | 7.16E-09 | -0.50726 | 0.271 | 0.338 | 0.000169 |
| Crebbp | 7.52E-09 | -0.32962 | 0.508 | 0.569 | 0.000177 |
| Irf2bp2 | 8.57E-09 | -0.28963 | 0.308 | 0.381 | 0.000202 |
| Gm15536 | 8.61E-09 | 0.321612 | 0.276 | 0.216 | 0.000203 |
| Gm9616 | 9.26E-09 | 0.313351 | 0.525 | 0.464 | 0.000218 |
| Tubgcp5 | 9.53E-09 | -0.3468 | 0.384 | 0.465 | 0.000225 |
| Casp8 | 1E-08 | -0.33682 | 0.411 | 0.474 | 0.000237 |
| Tle3 | 1.1E-08 | -0.26057 | 0.246 | 0.317 | 0.000259 |
| Gm13456 | 1.1E-08 | 0.284008 | 0.959 | 0.948 | 0.00026 |
| Cxcl16 | 1.17E-08 | 0.846053 | 0.303 | 0.244 | 0.000277 |
| Lrp10 | 1.26E-08 | -0.39825 | 0.408 | 0.472 | 0.000298 |
| Map7d1 | 1.32E-08 | -0.516 | 0.277 | 0.34 | 0.000312 |
| Kdm6b | 1.51E-08 | -0.42367 | 0.225 | 0.292 | 0.000356 |
| Kat6a | 1.53E-08 | -0.49571 | 0.266 | 0.334 | 0.00036 |
| Ssr1 | 1.78E-08 | -0.37793 | 0.594 | 0.652 | 0.00042 |
| Dnajc3 | 1.83E-08 | -0.29904 | 0.424 | 0.49 | 0.000431 |
| Spsb1 | 1.9E-08 | -0.43151 | 0.256 | 0.325 | 0.000448 |
| Malat1 | 2E-08 | 0.282221 | 0.99 | 0.992 | 0.000472 |
| Sort1 | 2.11E-08 | -0.30687 | 0.449 | 0.52 | 0.000498 |
| Mlph | 2.42E-08 | -0.52618 | 0.346 | 0.411 | 0.000572 |
| Gm10288 | 2.43E-08 | 0.411982 | 0.482 | 0.419 | 0.000572 |
| Cdh23 | 2.48E-08 | -0.52211 | 0.197 | 0.26 | 0.000585 |
| Rap1a | 2.48E-08 | -0.28902 | 0.408 | 0.471 | 0.000586 |
| Kcnk6 | 2.57E-08 | -0.32273 | 0.48 | 0.554 | 0.000606 |
| Ggt5 | 2.79E-08 | -0.44616 | 0.201 | 0.263 | 0.000657 |
| Tmem100 | 2.81E-08 | -0.33569 | 0.295 | 0.362 | 0.000662 |
| Tmem144 | 2.84E-08 | -0.64481 | 0.145 | 0.203 | 0.000671 |
| Gm13680 | 2.95E-08 | 0.277744 | 0.463 | 0.396 | 0.000695 |
| Nmt1 | 3.17E-08 | -0.3188 | 0.516 | 0.587 | 0.000747 |
| Map2k1 | 3.2E-08 | -0.44607 | 0.399 | 0.459 | 0.000754 |
| Cep152 | 3.75E-08 | -0.75566 | 0.165 | 0.221 | 0.000886 |
| Srsf6 | 4.07E-08 | -0.32346 | 0.273 | 0.34 | 0.000961 |
| Uba2 | 4.25E-08 | -0.48332 | 0.22 | 0.284 | 0.001001 |
| Rps3a2 | 4.42E-08 | 0.332597 | 0.423 | 0.358 | 0.001042 |
| Inpp4b | 4.68E-08 | -0.61794 | 0.192 | 0.252 | 0.001104 |
| Bcl9l | 4.84E-08 | -0.32442 | 0.181 | 0.243 | 0.001142 |
| Gm3379 | 4.89E-08 | 0.432092 | 0.614 | 0.562 | 0.001153 |

| | | | | | |
|---|---|---|---|---|---|
| Nup88 | 4.99E-08 | -0.44926 | 0.231 | 0.294 | 0.001177 |
| Rpl5 | 5.79E-08 | 0.37896 | 0.809 | 0.778 | 0.001367 |
| Samsn1 | 5.81E-08 | -0.35048 | 0.39 | 0.464 | 0.001371 |
| Etv5 | 6.07E-08 | -0.37067 | 0.351 | 0.418 | 0.001431 |
| Gm6472 | 6.41E-08 | 0.347358 | 0.5 | 0.445 | 0.001512 |
| Anxa5 | 6.57E-08 | 0.904153 | 0.21 | 0.159 | 0.00155 |
| B4galt4 | 6.7E-08 | -0.37395 | 0.241 | 0.307 | 0.001579 |
| Bcl2l1 | 6.72E-08 | -0.40532 | 0.352 | 0.415 | 0.001584 |
| Rpl21-ps15 | 6.89E-08 | 0.431929 | 0.367 | 0.308 | 0.001624 |
| Prkar1a | 7.18E-08 | -0.30474 | 0.526 | 0.581 | 0.001693 |
| Pacsin2 | 7.64E-08 | -0.28858 | 0.392 | 0.456 | 0.001802 |
| Ppp2ca | 7.79E-08 | -0.32444 | 0.453 | 0.517 | 0.001837 |
| Abcc3 | 8E-08 | -0.36335 | 0.355 | 0.427 | 0.001888 |
| Slamf9 | 8.11E-08 | 0.561131 | 0.38 | 0.321 | 0.001913 |
| Tagap | 8.4E-08 | -0.39492 | 0.31 | 0.376 | 0.00198 |
| Ly6e | 8.4E-08 | 0.306675 | 0.954 | 0.948 | 0.001982 |
| Hhex | 8.95E-08 | -0.25592 | 0.321 | 0.394 | 0.002112 |
| Actr1a | 9.57E-08 | -0.47358 | 0.269 | 0.331 | 0.002257 |
| Ptpn1 | 9.68E-08 | -0.27386 | 0.575 | 0.63 | 0.002284 |
| Camk2n1 | 9.92E-08 | -0.38499 | 0.233 | 0.294 | 0.002341 |
| Rps13-ps2 | 1.01E-07 | 0.30596 | 0.554 | 0.503 | 0.002382 |
| Sat1 | 1.06E-07 | 0.395917 | 0.863 | 0.847 | 0.002498 |
| Rps10-ps1 | 1.08E-07 | 0.480908 | 0.546 | 0.482 | 0.002552 |
| Ppm1g | 1.08E-07 | -0.50548 | 0.243 | 0.306 | 0.002556 |
| Creld2 | 1.13E-07 | -0.32374 | 0.206 | 0.267 | 0.002658 |
| Lasp1 | 1.18E-07 | -0.25774 | 0.378 | 0.446 | 0.002772 |
| Il21r | 1.23E-07 | -0.29425 | 0.404 | 0.473 | 0.002894 |
| Yipf4 | 1.24E-07 | -0.3744 | 0.286 | 0.348 | 0.002927 |
| Lpxn | 1.27E-07 | -0.31455 | 0.325 | 0.39 | 0.002988 |
| Gm24270 | 1.28E-07 | 0.406182 | 0.75 | 0.749 | 0.003029 |
| Pxk | 1.31E-07 | -0.56757 | 0.179 | 0.234 | 0.003085 |
| Uqcrh | 1.32E-07 | 0.596251 | 0.442 | 0.391 | 0.003116 |
| Cd300a | 1.35E-07 | -0.27233 | 0.519 | 0.579 | 0.003187 |
| Gm10790 | 1.44E-07 | -0.2598 | 0.541 | 0.605 | 0.003391 |
| Mgat2 | 1.45E-07 | -0.33462 | 0.304 | 0.368 | 0.00341 |
| Sec14l1 | 1.45E-07 | -0.28357 | 0.373 | 0.438 | 0.00343 |
| Rpl9-ps6 | 1.48E-07 | 0.445519 | 0.628 | 0.585 | 0.00349 |
| Mgat5 | 1.52E-07 | -0.30208 | 0.331 | 0.398 | 0.003586 |
| G3bp1 | 1.57E-07 | -0.34337 | 0.368 | 0.431 | 0.003694 |
| Plod3 | 1.62E-07 | -0.29994 | 0.512 | 0.575 | 0.003831 |
| Edem1 | 1.62E-07 | -0.29789 | 0.529 | 0.591 | 0.003832 |
| Ptbp1 | 1.73E-07 | -0.54467 | 0.246 | 0.303 | 0.004082 |
| Mbnl2 | 1.77E-07 | -0.39114 | 0.375 | 0.44 | 0.004173 |
| Gm15772 | 1.85E-07 | 0.305809 | 0.501 | 0.445 | 0.004364 |
| Bid | 1.89E-07 | -0.34414 | 0.199 | 0.259 | 0.004454 |
| Sh3bgrl3 | 2.04E-07 | 0.345625 | 0.815 | 0.812 | 0.004805 |
| Ski | 2.11E-07 | -0.25777 | 0.47 | 0.53 | 0.004981 |

| | | | | | |
|---|---|---|---|---|---|
| Snta1 | 2.16E-07 | -0.47374 | 0.255 | 0.319 | 0.005105 |
| Ccdc71 | 2.18E-07 | -0.36009 | 0.146 | 0.2 | 0.005154 |
| Pten | 2.22E-07 | -0.31864 | 0.357 | 0.42 | 0.005225 |
| Rab18 | 2.31E-07 | -0.43094 | 0.212 | 0.271 | 0.005458 |
| Mtmr3 | 2.34E-07 | -0.52877 | 0.161 | 0.217 | 0.005516 |
| Sbf2 | 2.43E-07 | -0.26531 | 0.292 | 0.357 | 0.005724 |
| Idh2 | 2.49E-07 | -0.35927 | 0.379 | 0.444 | 0.005877 |
| Ttc7 | 2.65E-07 | -0.26195 | 0.188 | 0.249 | 0.006256 |
| Prrc2a | 2.75E-07 | -0.32889 | 0.307 | 0.371 | 0.006477 |
| Tlr12 | 2.78E-07 | -0.4111 | 0.221 | 0.279 | 0.006563 |
| Cln8 | 3.07E-07 | -0.42291 | 0.278 | 0.337 | 0.007245 |
| Gm9320 | 3.08E-07 | 0.340789 | 0.208 | 0.156 | 0.007276 |
| Slc46a1 | 3.19E-07 | -0.3122 | 0.394 | 0.455 | 0.00753 |
| Ac165079.1 | 3.21E-07 | 0.3202 | 0.274 | 0.221 | 0.007581 |
| Akna | 3.47E-07 | -0.3101 | 0.402 | 0.467 | 0.008194 |
| D1Ertd622e | 4.09E-07 | -0.25244 | 0.198 | 0.255 | 0.009642 |
| Gpr155 | 4.16E-07 | -0.33504 | 0.237 | 0.295 | 0.009805 |
| Fam32a | 4.5E-07 | -0.29892 | 0.197 | 0.254 | 0.010622 |
| Mcfd2 | 4.67E-07 | -0.26268 | 0.391 | 0.453 | 0.011005 |
| Zmynd11 | 4.74E-07 | -0.43726 | 0.219 | 0.274 | 0.01117 |
| Ddx3x | 4.8E-07 | -0.25102 | 0.336 | 0.397 | 0.011324 |
| Kat7 | 4.85E-07 | -0.2944 | 0.181 | 0.236 | 0.01145 |
| Cerk | 4.89E-07 | -0.32174 | 0.352 | 0.417 | 0.011529 |
| Arhgap12 | 5.07E-07 | -0.67041 | 0.208 | 0.265 | 0.011964 |
| Supt20 | 5.62E-07 | -0.27951 | 0.237 | 0.298 | 0.013254 |
| Gm6988 | 5.77E-07 | 0.26463 | 0.276 | 0.224 | 0.013611 |
| Actr3 | 6.04E-07 | -0.28432 | 0.48 | 0.536 | 0.014252 |
| Alkbh5 | 6.05E-07 | -0.49336 | 0.316 | 0.376 | 0.014261 |
| Ndfip2 | 6.16E-07 | -0.42791 | 0.275 | 0.333 | 0.014533 |
| Ubash3b | 6.54E-07 | -0.27584 | 0.559 | 0.608 | 0.015424 |
| Cd47 | 6.98E-07 | -0.28948 | 0.609 | 0.655 | 0.016458 |
| Usf1 | 7.27E-07 | -0.35834 | 0.19 | 0.247 | 0.017154 |
| Ralb | 7.33E-07 | -0.36834 | 0.387 | 0.445 | 0.017284 |
| Kcnk13 | 7.57E-07 | -0.41681 | 0.275 | 0.332 | 0.017845 |
| Rpl29 | 7.67E-07 | 0.356366 | 0.675 | 0.643 | 0.018085 |
| Pld1 | 8.54E-07 | -0.26099 | 0.391 | 0.461 | 0.020154 |
| Prag1 | 8.6E-07 | -0.37247 | 0.196 | 0.25 | 0.020284 |
| Gm43028 | 8.82E-07 | 0.278999 | 0.359 | 0.307 | 0.020796 |
| Capn3 | 8.86E-07 | -0.25958 | 0.359 | 0.422 | 0.020893 |
| Atp2a2 | 8.99E-07 | -0.28976 | 0.254 | 0.316 | 0.021198 |
| Abcd3 | 8.99E-07 | -0.53344 | 0.17 | 0.225 | 0.021211 |
| Mxi1 | 9E-07 | -0.35545 | 0.152 | 0.203 | 0.021223 |
| Rack1 | 9.3E-07 | 0.370541 | 0.729 | 0.694 | 0.021938 |
| Zfp36 | 9.42E-07 | -0.25359 | 0.413 | 0.476 | 0.022215 |
| Ak1 | 9.62E-07 | -0.53175 | 0.2 | 0.253 | 0.022702 |
| Filip1l | 1.11E-06 | -0.4313 | 0.239 | 0.295 | 0.026128 |
| Gm9703 | 1.12E-06 | 0.259784 | 0.307 | 0.256 | 0.026439 |

| | | | | | |
|---|---|---|---|---|---|
| Ncbp3 | 1.14E-06 | -0.44847 | 0.186 | 0.239 | 0.026832 |
| Snrpg | 1.19E-06 | 0.345327 | 0.394 | 0.337 | 0.028022 |
| Rpl36-ps12 | 1.22E-06 | 0.415743 | 0.255 | 0.206 | 0.028795 |
| Dbnl | 1.26E-06 | -0.30965 | 0.373 | 0.434 | 0.029722 |
| Ddah2 | 1.27E-06 | -0.36936 | 0.158 | 0.209 | 0.029949 |
| Cd68 | 1.34E-06 | 0.255486 | 0.949 | 0.956 | 0.031597 |
| Il1a | 1.48E-06 | -0.4365 | 0.159 | 0.21 | 0.034989 |
| Poldip3 | 1.48E-06 | -0.32751 | 0.329 | 0.389 | 0.034994 |
| Jam2 | 1.62E-06 | -0.26863 | 0.263 | 0.324 | 0.038155 |
| Gm8960 | 1.65E-06 | 0.38761 | 0.323 | 0.273 | 0.039005 |
| Jun | 1.68E-06 | -0.44025 | 0.335 | 0.39 | 0.039699 |
| Gm5835 | 1.74E-06 | 0.392147 | 0.405 | 0.352 | 0.041071 |
| Gm10036 | 1.78E-06 | 0.362209 | 0.364 | 0.311 | 0.041965 |
| Wbp1l | 1.86E-06 | -0.70066 | 0.172 | 0.222 | 0.043828 |
| Slc38a10 | 1.9E-06 | -0.49023 | 0.294 | 0.347 | 0.044917 |
| Dcaf12 | 1.95E-06 | -0.43747 | 0.215 | 0.266 | 0.046015 |
| Elk3 | 2E-06 | -0.31963 | 0.348 | 0.401 | 0.047209 |
| Exosc10 | 2.07E-06 | -0.45708 | 0.165 | 0.216 | 0.048924 |
| Rpl8 | 2.1E-06 | 0.383268 | 0.737 | 0.708 | 0.049555 |
| Gm8995 | 2.1E-06 | 0.704404 | 0.23 | 0.182 | 0.0496 |

#### Supplementry Table 2. DAM DEGs compared to HS

| GeneName | p_val | avg_log2FC | pct.1 | pct.2 | p_val_adj |
|---|---|---|---|---|---|
| Clec7a | 0 | 6.077894 | 0.7 | 0.025 | 0 |
| Lpl | 0 | 7.182072 | 0.565 | 0.052 | 0 |
| Cst7 | 4.4E-251 | 5.033391 | 0.473 | 0.046 | 1E-246 |
| Axl | 2E-246 | 5.136287 | 0.408 | 0.032 | 4.7E-242 |
| Itgax | 8.7E-240 | 4.42447 | 0.289 | 0.012 | 2.1E-235 |
| Gas2l3 | 1.2E-202 | 4.744386 | 0.24 | 0.009 | 2.8E-198 |
| Lilrb4a | 3.6E-195 | 3.813066 | 0.283 | 0.016 | 8.5E-191 |
| P2ry12 | 7.6E-185 | -1.97909 | 0.922 | 1 | 1.8E-180 |
| Csf1 | 1.4E-181 | 3.882426 | 0.377 | 0.04 | 3.2E-177 |
| Cd63 | 2.8E-174 | 2.626947 | 0.989 | 0.684 | 6.5E-170 |
| Lgals3 | 1.6E-171 | 8.197114 | 0.424 | 0.062 | 3.8E-167 |
| Selplg | 1.2E-168 | -1.68995 | 0.966 | 1 | 2.8E-164 |
| Cd74 | 7.9E-167 | 5.10919 | 0.614 | 0.153 | 1.9E-162 |
| Fam20c | 7.1E-165 | 4.31389 | 0.316 | 0.029 | 1.7E-160 |
| Cd63-ps | 6.4E-162 | 2.312401 | 0.935 | 0.465 | 1.5E-157 |
| Slc2a5 | 1.7E-157 | -2.80257 | 0.5 | 0.944 | 3.9E-153 |
| Tmem119 | 3.5E-149 | -1.72875 | 0.899 | 1 | 8.2E-145 |
| Fgr | 3.3E-143 | 4.225019 | 0.24 | 0.018 | 7.8E-139 |
| Ccl4 | 1.6E-136 | 7.664959 | 0.251 | 0.022 | 3.9E-132 |
| Ctsb | 1.2E-128 | 1.471438 | 1 | 0.999 | 2.9E-124 |
| Ftl1 | 6.9E-124 | 1.932415 | 1 | 0.991 | 1.6E-119 |
| Capg | 3.6E-121 | 3.537601 | 0.554 | 0.153 | 8.6E-117 |
| Cox6a2 | 4.2E-121 | 3.668783 | 0.209 | 0.016 | 9.8E-117 |
| Chst2 | 1.6E-119 | 3.581734 | 0.251 | 0.026 | 3.7E-115 |
| Apoe | 4.6E-109 | 6.884149 | 0.744 | 0.376 | 1.1E-104 |
| Cd9 | 6.9E-109 | 1.657737 | 1 | 0.965 | 1.6E-104 |
| Ccl3 | 7.1E-108 | 6.152176 | 0.307 | 0.047 | 1.7E-103 |
| Ftl1-ps1 | 5.3E-105 | 1.844764 | 0.991 | 0.874 | 1.2E-100 |
| Ftl2-ps | 7.8E-105 | 1.838599 | 0.998 | 0.914 | 1.8E-100 |
| Cd83 | 1.7E-99 | 2.521748 | 0.684 | 0.26 | 3.99E-95 |
| Olfr111 | 1.05E-98 | 2.372131 | 0.28 | 0.041 | 2.48E-94 |
| Cstb | 3.45E-96 | 2.383318 | 0.686 | 0.263 | 8.13E-92 |
| Fxyd5 | 2.61E-92 | 4.364522 | 0.312 | 0.059 | 6.16E-88 |
| Fth1 | 2.7E-91 | 1.847698 | 1 | 0.993 | 6.38E-87 |
| Ank | 1.91E-90 | 2.640128 | 0.401 | 0.093 | 4.52E-86 |
| Plaur | 7.51E-90 | 3.077057 | 0.33 | 0.065 | 1.77E-85 |
| Il2rg | 1.19E-89 | 3.976835 | 0.253 | 0.038 | 2.8E-85 |
| Cxcl16 | 3.27E-89 | 2.667459 | 0.632 | 0.244 | 7.71E-85 |
| Siglech | 5.29E-86 | -1.12159 | 0.969 | 0.999 | 1.25E-81 |
| Ctsz | 1.26E-85 | 1.238292 | 1 | 0.993 | 2.96E-81 |
| Cd52 | 4.66E-83 | 2.372275 | 0.778 | 0.411 | 1.1E-78 |
| P2ry13 | 7.25E-83 | -1.4448 | 0.809 | 0.977 | 1.71E-78 |
| H2-D1 | 1.08E-82 | 1.824812 | 0.975 | 0.867 | 2.56E-78 |

| | | | | | |
|---|---|---|---|---|---|
| Vsir | 5.14E-79 | -1.225 | 0.888 | 0.985 | 1.21E-74 |
| Lyz2 | 3.24E-78 | 2.823566 | 0.89 | 0.608 | 7.63E-74 |
| Ptgs1 | 3.03E-77 | -1.66412 | 0.704 | 0.946 | 7.15E-73 |
| Pkm | 2.46E-74 | 1.920632 | 0.872 | 0.559 | 5.8E-70 |
| H2-K1 | 6.03E-74 | 1.739643 | 0.948 | 0.766 | 1.42E-69 |
| Mif | 1.7E-73 | 2.305119 | 0.693 | 0.347 | 4.02E-69 |
| Cd164 | 2.79E-73 | -1.42892 | 0.724 | 0.946 | 6.58E-69 |
| Mir692-2 | 4.85E-73 | 1.060639 | 0.657 | 0.301 | 1.15E-68 |
| Gm15590 | 6.3E-72 | 1.209917 | 0.776 | 0.453 | 1.49E-67 |
| Pld3 | 1.18E-71 | 1.779893 | 0.879 | 0.576 | 2.79E-67 |
| Clic4 | 1.5E-71 | 4.119204 | 0.231 | 0.04 | 3.55E-67 |
| Atp1a3 | 5.25E-71 | 2.983836 | 0.211 | 0.032 | 1.24E-66 |
| Mt1 | 6.92E-70 | 1.832159 | 0.664 | 0.297 | 1.63E-65 |
| Cx3cr1 | 3.08E-68 | -0.81453 | 0.991 | 1 | 7.28E-64 |
| Cd72 | 1.24E-63 | 3.757167 | 0.204 | 0.034 | 2.93E-59 |
| Ifi27l2a | 1.28E-63 | 2.399345 | 0.305 | 0.074 | 3.02E-59 |
| Adgrg1 | 2.03E-63 | -1.52297 | 0.619 | 0.905 | 4.79E-59 |
| Crybb1 | 2.3E-62 | -1.56464 | 0.433 | 0.794 | 5.43E-58 |
| Aplp2 | 4.15E-61 | 2.16013 | 0.578 | 0.257 | 9.8E-57 |
| Ftl1-ps2 | 2.55E-60 | 0.970362 | 0.583 | 0.271 | 6.01E-56 |
| Tpi1 | 4.08E-60 | 1.791057 | 0.626 | 0.284 | 9.63E-56 |
| Olfml3 | 5.39E-60 | -1.00954 | 0.953 | 0.996 | 1.27E-55 |
| Gapdh | 1.15E-59 | 1.326678 | 0.969 | 0.904 | 2.72E-55 |
| C3ar1 | 7.77E-59 | 1.683094 | 0.825 | 0.553 | 1.83E-54 |
| Mfsd12 | 2.01E-58 | 2.12024 | 0.433 | 0.149 | 4.74E-54 |
| Ltc4s | 2.74E-58 | -1.48783 | 0.585 | 0.867 | 6.47E-54 |
| Prdx1 | 5.27E-58 | 1.75217 | 0.791 | 0.495 | 1.24E-53 |
| Aldoa | 6.18E-58 | 1.490202 | 0.917 | 0.704 | 1.46E-53 |
| Gm6560 | 2.81E-57 | 1.290054 | 0.628 | 0.306 | 6.62E-53 |
| Zfhx3 | 7.45E-57 | -1.34948 | 0.76 | 0.948 | 1.76E-52 |
| Rpsa | 5.35E-56 | 1.395306 | 0.928 | 0.774 | 1.26E-51 |
| Cd68 | 2.15E-55 | 1.117707 | 0.991 | 0.956 | 5.07E-51 |
| Anxa5 | 2.73E-55 | 2.450052 | 0.43 | 0.159 | 6.43E-51 |
| Ifngr1 | 5.72E-55 | -1.01268 | 0.865 | 0.972 | 1.35E-50 |
| Atp6v0c-ps | 1.01E-54 | 1.199668 | 0.883 | 0.651 | 2.37E-50 |
| Ctsd | 1.43E-54 | 0.878837 | 1 | 1 | 3.36E-50 |
| Pgk1 | 2.61E-54 | 1.6338 | 0.48 | 0.18 | 6.16E-50 |
| H2-Q7 | 9.32E-54 | 3.681242 | 0.247 | 0.059 | 2.2E-49 |
| Susd3 | 1.69E-53 | -1.48899 | 0.574 | 0.846 | 3.98E-49 |
| Gm6977 | 5.57E-52 | 1.411219 | 0.756 | 0.479 | 1.31E-47 |
| Rhoc | 9.76E-52 | 2.259636 | 0.321 | 0.094 | 2.3E-47 |
| Atp6v0c | 1.03E-51 | 1.200321 | 0.948 | 0.828 | 2.42E-47 |
| B2m | 4.34E-51 | 0.957495 | 1 | 0.998 | 1.02E-46 |
| Rhob | 2.36E-50 | -1.05717 | 0.735 | 0.928 | 5.56E-46 |
| Fcrls | 4.78E-50 | -0.96549 | 0.848 | 0.975 | 1.13E-45 |
| Glipr1 | 1.62E-49 | 2.470451 | 0.224 | 0.05 | 3.83E-45 |
| H2-Q6 | 3.7E-49 | 3.334437 | 0.22 | 0.05 | 8.72E-45 |

| | | | | | |
|---|---|---|---|---|---|
| Sparc | 4.4E-49 | -0.74146 | 0.989 | 1 | 1.04E-44 |
| Adrb2 | 9.03E-49 | -1.89572 | 0.314 | 0.652 | 2.13E-44 |
| Gpr34 | 6.99E-48 | -0.84005 | 0.915 | 0.992 | 1.65E-43 |
| Lgals3bp | 8.58E-48 | 1.82203 | 0.771 | 0.503 | 2.02E-43 |
| Adssl1 | 4.71E-47 | 2.264828 | 0.249 | 0.064 | 1.11E-42 |
| C530043K1 | 8.51E-47 | 2.508594 | 0.265 | 0.071 | 2.01E-42 |
| Csf1r | 8.89E-47 | -0.7447 | 0.989 | 1 | 2.1E-42 |
| Rnase4 | 9.96E-46 | -1.02658 | 0.816 | 0.955 | 2.35E-41 |
| Lpcat2 | 1.26E-45 | -0.83423 | 0.897 | 0.984 | 2.98E-41 |
| Ecscr | 4.35E-45 | -1.58337 | 0.383 | 0.706 | 1.03E-40 |
| Nceh1 | 5.5E-45 | 2.391975 | 0.238 | 0.062 | 1.3E-40 |
| Hmox1 | 8.1E-45 | 2.166582 | 0.363 | 0.129 | 1.91E-40 |
| Ldha | 9.41E-45 | 1.653336 | 0.783 | 0.54 | 2.22E-40 |
| Ly9 | 1.86E-44 | 2.046863 | 0.289 | 0.086 | 4.38E-40 |
| Tubb6 | 2.25E-43 | 2.846596 | 0.229 | 0.06 | 5.32E-39 |
| Tmem173 | 2.82E-43 | -1.20547 | 0.673 | 0.905 | 6.65E-39 |
| Apbb2 | 3.14E-43 | 1.490849 | 0.397 | 0.148 | 7.41E-39 |
| Rps2 | 4.23E-43 | 1.25201 | 0.877 | 0.675 | 9.97E-39 |
| Slamf8 | 8.11E-43 | 2.220362 | 0.294 | 0.091 | 1.91E-38 |
| Sulf2 | 9.94E-43 | 2.137752 | 0.224 | 0.057 | 2.35E-38 |
| Maf | 1.14E-41 | -1.02457 | 0.686 | 0.897 | 2.7E-37 |
| Cpd | 1.48E-41 | 2.023695 | 0.274 | 0.082 | 3.49E-37 |
| Lrrc3 | 4.75E-41 | -0.90291 | 0.63 | 0.896 | 1.12E-36 |
| Pea15a | 6.22E-41 | 1.310273 | 0.693 | 0.403 | 1.47E-36 |
| Selenow | 8.5E-41 | 1.584512 | 0.415 | 0.167 | 2.01E-36 |
| Rpl41 | 1.31E-40 | 0.999237 | 0.978 | 0.934 | 3.08E-36 |
| Selenop | 2.19E-40 | -0.74476 | 0.948 | 0.991 | 5.18E-36 |
| Rpsa-ps10 | 4.32E-40 | 1.100842 | 0.796 | 0.532 | 1.02E-35 |
| Sdc3 | 4.82E-40 | 1.84103 | 0.318 | 0.109 | 1.14E-35 |
| Rap2b | 5.11E-40 | 1.703062 | 0.278 | 0.086 | 1.2E-35 |
| Gm2574 | 8.27E-40 | 1.110372 | 0.691 | 0.43 | 1.95E-35 |
| Sall1 | 8.86E-40 | -1.08613 | 0.581 | 0.856 | 2.09E-35 |
| Hif1a | 3.35E-39 | 1.666727 | 0.603 | 0.324 | 7.91E-35 |
| Lrba | 2.95E-38 | -1.94362 | 0.224 | 0.547 | 6.96E-34 |
| Cadm1 | 3.58E-38 | 1.321908 | 0.661 | 0.387 | 8.44E-34 |
| Ifitm3 | 6.77E-38 | 3.007813 | 0.318 | 0.121 | 1.6E-33 |
| Il6ra | 1.21E-37 | -1.17486 | 0.74 | 0.903 | 2.85E-33 |
| Upk1b | 1.27E-37 | -2.36985 | 0.166 | 0.473 | 2.99E-33 |
| Hsp90ab1 | 1.63E-37 | 1.000769 | 0.944 | 0.844 | 3.85E-33 |
| Bin2 | 1.65E-37 | -0.98832 | 0.762 | 0.909 | 3.89E-33 |
| Gm22774 | 1.02E-36 | 0.461128 | 0.285 | 0.095 | 2.41E-32 |
| Bola2 | 1.24E-36 | 0.505566 | 0.41 | 0.163 | 2.92E-32 |
| Bcl2a1d | 1.31E-36 | 2.707899 | 0.352 | 0.143 | 3.1E-32 |
| Id2 | 1.61E-36 | 2.622469 | 0.359 | 0.147 | 3.8E-32 |
| Rplp0 | 1.66E-36 | 1.078411 | 0.962 | 0.89 | 3.92E-32 |
| Gm5559 | 3.18E-36 | 1.054743 | 0.897 | 0.717 | 7.51E-32 |
| Rps6ka1 | 4.44E-36 | -1.20773 | 0.576 | 0.813 | 1.05E-31 |

| | | | | | |
|---|---|---|---|---|---|
| Tlr2 | 7.1E-36 | 2.028891 | 0.608 | 0.35 | 1.68E-31 |
| Otulinl | 7.86E-36 | -0.97761 | 0.791 | 0.93 | 1.85E-31 |
| Bcl2a1b | 8.02E-36 | 1.044835 | 0.738 | 0.458 | 1.89E-31 |
| Cyba | 1.84E-35 | 0.812907 | 0.991 | 0.973 | 4.34E-31 |
| Pmepa1 | 2.32E-35 | -1.05358 | 0.623 | 0.845 | 5.48E-31 |
| Entpd1 | 1.23E-34 | -0.88292 | 0.774 | 0.94 | 2.9E-30 |
| Csf2ra | 1.85E-34 | 1.690845 | 0.502 | 0.266 | 4.37E-30 |
| Emp3 | 3.71E-34 | 1.807811 | 0.33 | 0.128 | 8.76E-30 |
| Abhd15 | 4.1E-34 | -2.14612 | 0.262 | 0.552 | 9.66E-30 |
| Gnas | 4.47E-34 | 1.186862 | 0.859 | 0.708 | 1.05E-29 |
| Myo1e | 6.27E-34 | 1.642644 | 0.262 | 0.086 | 1.48E-29 |
| Serinc3 | 6.8E-34 | -0.78532 | 0.919 | 0.996 | 1.6E-29 |
| Gm7336 | 9.59E-34 | 1.060873 | 0.574 | 0.326 | 2.26E-29 |
| Slco2b1 | 1.5E-33 | -0.95834 | 0.756 | 0.937 | 3.53E-29 |
| Tyrobp | 1.92E-33 | 0.737189 | 0.998 | 0.996 | 4.53E-29 |
| Ccr5 | 2.11E-33 | -1.27833 | 0.426 | 0.717 | 4.97E-29 |
| Dpp7 | 2.7E-33 | 1.45352 | 0.336 | 0.132 | 6.38E-29 |
| Gm10076 | 2.89E-33 | 1.014114 | 0.96 | 0.893 | 6.81E-29 |
| Rps20 | 2.99E-33 | 0.974291 | 0.93 | 0.807 | 7.06E-29 |
| Lgals1 | 3.16E-33 | 3.188592 | 0.211 | 0.064 | 7.45E-29 |
| Cd9-ps | 3.41E-33 | 0.418176 | 0.26 | 0.086 | 8.05E-29 |
| Slc16a3 | 4.08E-33 | 1.672201 | 0.422 | 0.193 | 9.62E-29 |
| Rab3il1 | 7.43E-33 | -1.18018 | 0.643 | 0.828 | 1.75E-28 |
| Cmtm7 | 1.04E-32 | -1.06432 | 0.601 | 0.815 | 2.46E-28 |
| Rpl32 | 1.66E-32 | 0.860711 | 0.948 | 0.863 | 3.91E-28 |
| Col27a1 | 3.58E-32 | -1.4901 | 0.265 | 0.563 | 8.44E-28 |
| Mir692-1 | 4.72E-32 | 0.389711 | 0.249 | 0.083 | 1.11E-27 |
| Cmtm6 | 5.44E-32 | -1.12137 | 0.628 | 0.854 | 1.28E-27 |
| Gm3699 | 5.96E-32 | 0.935061 | 0.572 | 0.326 | 1.41E-27 |
| Gm12164 | 6.46E-32 | 0.429591 | 0.316 | 0.124 | 1.52E-27 |
| Eno1 | 8.19E-32 | 1.006307 | 0.872 | 0.695 | 1.93E-27 |
| Gm10790 | 1.08E-31 | -1.60352 | 0.334 | 0.605 | 2.55E-27 |
| Pld4 | 1.35E-31 | -0.7708 | 0.852 | 0.959 | 3.19E-27 |
| Nme1 | 1.87E-31 | 1.340088 | 0.437 | 0.206 | 4.4E-27 |
| Abi3 | 2.11E-31 | -1.02423 | 0.643 | 0.851 | 4.99E-27 |
| Nrp2 | 2.42E-31 | 1.830474 | 0.33 | 0.134 | 5.7E-27 |
| Il16 | 2.5E-31 | -1.76075 | 0.318 | 0.574 | 5.89E-27 |
| Iqgap1 | 4.1E-31 | 2.508845 | 0.242 | 0.083 | 9.67E-27 |
| Lgmn | 8.08E-31 | -0.50168 | 1 | 1 | 1.91E-26 |
| Zfp710 | 8.22E-31 | -1.37829 | 0.484 | 0.727 | 1.94E-26 |
| Tanc2 | 1.62E-30 | -1.05467 | 0.628 | 0.831 | 3.82E-26 |
| Glul | 1.67E-30 | -0.84559 | 0.796 | 0.938 | 3.93E-26 |
| Ttyh2 | 1.78E-30 | 1.974845 | 0.209 | 0.064 | 4.2E-26 |
| AU020206 | 1.89E-30 | 1.155345 | 0.715 | 0.473 | 4.45E-26 |
| Txnip | 2.21E-30 | -1.38924 | 0.469 | 0.699 | 5.21E-26 |
| Abca9 | 2.58E-30 | -1.26788 | 0.422 | 0.695 | 6.07E-26 |
| Gna15 | 2.77E-30 | -1.3484 | 0.453 | 0.689 | 6.54E-26 |

| | | | | | |
|---|---|---|---|---|---|
| Calm2 | 4.64E-30 | -0.81834 | 0.874 | 0.953 | 1.1E-25 |
| Gm8730 | 5.24E-30 | 0.961228 | 0.87 | 0.723 | 1.24E-25 |
| Rps2-ps10 | 5.72E-30 | 0.822492 | 0.478 | 0.248 | 1.35E-25 |
| Rpl10a | 6.14E-30 | 0.892375 | 0.85 | 0.643 | 1.45E-25 |
| Ssh2 | 8.97E-30 | -0.93582 | 0.608 | 0.843 | 2.12E-25 |
| Rps19 | 1.05E-29 | 0.881874 | 0.937 | 0.863 | 2.48E-25 |
| Tpm4 | 1.08E-29 | 2.092251 | 0.312 | 0.129 | 2.55E-25 |
| Arhgap5 | 1.14E-29 | -1.18814 | 0.54 | 0.761 | 2.69E-25 |
| Bst2 | 1.53E-29 | 1.348252 | 0.612 | 0.369 | 3.62E-25 |
| Rps28 | 2.97E-29 | 0.826747 | 0.978 | 0.892 | 7.01E-25 |
| Uba52 | 3.63E-29 | 0.915126 | 0.886 | 0.721 | 8.56E-25 |
| Fgd2 | 4.11E-29 | -1.22575 | 0.522 | 0.747 | 9.69E-25 |
| Dusp7 | 4.28E-29 | -2.02262 | 0.332 | 0.588 | 1.01E-24 |
| Hnrnpa1 | 4.49E-29 | 1.360111 | 0.507 | 0.28 | 1.06E-24 |
| Rps18 | 7.25E-29 | 0.902259 | 0.942 | 0.833 | 1.71E-24 |
| Pcna | 7.25E-29 | 1.127407 | 0.446 | 0.217 | 1.71E-24 |
| St3gal6 | 1.36E-28 | -1.00297 | 0.587 | 0.804 | 3.2E-24 |
| Alox5ap | 1.38E-28 | -1.03116 | 0.578 | 0.777 | 3.26E-24 |
| Cdkn1a | 1.52E-28 | 2.809829 | 0.262 | 0.102 | 3.58E-24 |
| Npm1 | 2.04E-28 | 1.187436 | 0.744 | 0.523 | 4.81E-24 |
| Nme2 | 2.14E-28 | 0.996983 | 0.59 | 0.336 | 5.06E-24 |
| Rps5 | 2.42E-28 | 1.068638 | 0.865 | 0.718 | 5.71E-24 |
| F11r | 2.63E-28 | -0.69007 | 0.877 | 0.965 | 6.19E-24 |
| Ivns1abp | 4.51E-28 | -0.89545 | 0.661 | 0.852 | 1.06E-23 |
| Scamp2 | 4.67E-28 | -0.81334 | 0.83 | 0.931 | 1.1E-23 |
| Slc11a2 | 4.72E-28 | 1.793702 | 0.24 | 0.083 | 1.11E-23 |
| Rpl35 | 7.02E-28 | 0.823158 | 0.93 | 0.807 | 1.66E-23 |
| C5ar2 | 7.08E-28 | -1.48373 | 0.404 | 0.658 | 1.67E-23 |
| Rpl3 | 7.15E-28 | 0.828248 | 0.942 | 0.821 | 1.69E-23 |
| Npc2 | 8.67E-28 | 0.917283 | 0.944 | 0.82 | 2.04E-23 |
| Rplp1 | 9.39E-28 | 0.697788 | 0.982 | 0.952 | 2.22E-23 |
| Marcks | 1.16E-27 | -0.57055 | 0.978 | 0.996 | 2.75E-23 |
| Stab1 | 1.37E-27 | -0.72151 | 0.583 | 0.822 | 3.23E-23 |
| Atp5g1 | 1.41E-27 | 1.166624 | 0.583 | 0.349 | 3.34E-23 |
| Atf3 | 1.84E-27 | 3.528644 | 0.283 | 0.119 | 4.34E-23 |
| Ncl | 3.61E-27 | 1.144489 | 0.776 | 0.568 | 8.51E-23 |
| Srgap2 | 3.73E-27 | -0.91713 | 0.63 | 0.831 | 8.8E-23 |
| Elmo1 | 4.28E-27 | -1.08775 | 0.482 | 0.724 | 1.01E-22 |
| Myl6 | 4.6E-27 | 0.927988 | 0.803 | 0.602 | 1.09E-22 |
| Rps8 | 4.89E-27 | 0.867548 | 0.922 | 0.813 | 1.15E-22 |
| Plekhm2 | 5.23E-27 | 1.924692 | 0.258 | 0.098 | 1.23E-22 |
| Gm11808 | 5.66E-27 | 0.874499 | 0.691 | 0.473 | 1.33E-22 |
| Gm10359 | 5.88E-27 | 0.709751 | 0.52 | 0.303 | 1.39E-22 |
| Atox1 | 8.18E-27 | 0.968256 | 0.731 | 0.498 | 1.93E-22 |
| Rack1 | 1.01E-26 | 1.028802 | 0.854 | 0.694 | 2.39E-22 |
| Rtn4rl1 | 1.15E-26 | -1.24098 | 0.439 | 0.684 | 2.71E-22 |
| Rpl38 | 1.2E-26 | 0.939508 | 0.809 | 0.616 | 2.83E-22 |

| | | | | | |
|---|---|---|---|---|---|
| Itpripl1 | 2.14E-26 | -1.33879 | 0.509 | 0.729 | 5.06E-22 |
| Eef1a1 | 2.18E-26 | 0.762766 | 0.998 | 0.994 | 5.14E-22 |
| Tpst2 | 2.32E-26 | -1.35047 | 0.507 | 0.701 | 5.47E-22 |
| Rplp2 | 2.38E-26 | 0.690697 | 0.955 | 0.895 | 5.62E-22 |
| Golm1 | 2.82E-26 | -0.88725 | 0.74 | 0.877 | 6.66E-22 |
| Ckb | 4.23E-26 | -0.73403 | 0.798 | 0.895 | 9.99E-22 |
| Crip1 | 4.58E-26 | 2.743031 | 0.233 | 0.089 | 1.08E-21 |
| Gm47428 | 5.34E-26 | 0.415898 | 0.226 | 0.08 | 1.26E-21 |
| Gm2000 | 5.36E-26 | 0.932758 | 0.765 | 0.571 | 1.26E-21 |
| Ctc1 | 6.03E-26 | -1.47158 | 0.401 | 0.63 | 1.42E-21 |
| Nav3 | 6.3E-26 | -1.39343 | 0.428 | 0.656 | 1.49E-21 |
| Qk | 8.19E-26 | -0.81653 | 0.816 | 0.936 | 1.93E-21 |
| Fam110a | 9.36E-26 | -2.35764 | 0.13 | 0.364 | 2.21E-21 |
| Slc15a3 | 9.47E-26 | 1.633184 | 0.592 | 0.378 | 2.23E-21 |
| Sdc4 | 1.1E-25 | 1.951827 | 0.359 | 0.175 | 2.6E-21 |
| Scpep1 | 1.5E-25 | 0.911968 | 0.605 | 0.366 | 3.53E-21 |
| Pde3b | 1.6E-25 | -0.8974 | 0.596 | 0.799 | 3.77E-21 |
| Rps26 | 1.62E-25 | 0.832327 | 0.859 | 0.664 | 3.83E-21 |
| Rpl13 | 1.78E-25 | 0.739341 | 0.975 | 0.938 | 4.2E-21 |
| Ywhah | 2.36E-25 | -0.78698 | 0.85 | 0.919 | 5.58E-21 |
| Kcnj2 | 2.61E-25 | 1.680512 | 0.22 | 0.077 | 6.15E-21 |
| Gpi1 | 4.74E-25 | 1.089276 | 0.561 | 0.336 | 1.12E-20 |
| Rpl23 | 5.66E-25 | 0.880602 | 0.933 | 0.82 | 1.34E-20 |
| Hexb | 5.87E-25 | -0.39941 | 1 | 1 | 1.38E-20 |
| Hsp90aa1 | 7.08E-25 | 1.138832 | 0.724 | 0.533 | 1.67E-20 |
| Arhgap45 | 7.47E-25 | -0.88812 | 0.709 | 0.86 | 1.76E-20 |
| Gm12537 | 8.16E-25 | 0.796173 | 0.5 | 0.291 | 1.92E-20 |
| Itga5 | 8.79E-25 | 2.102758 | 0.222 | 0.081 | 2.07E-20 |
| Pdgfa | 9.26E-25 | 1.246589 | 0.307 | 0.131 | 2.19E-20 |
| Rpl3-ps1 | 1.49E-24 | 0.897756 | 0.857 | 0.694 | 3.51E-20 |
| Gm21399 | 1.53E-24 | 0.477203 | 0.343 | 0.16 | 3.62E-20 |
| Commd8 | 1.62E-24 | -1.10611 | 0.462 | 0.689 | 3.82E-20 |
| Gm9625 | 3.71E-24 | 0.93805 | 0.684 | 0.477 | 8.74E-20 |
| Gm13456 | 3.88E-24 | 0.742554 | 0.978 | 0.948 | 9.14E-20 |
| Rpl12 | 4.29E-24 | 0.957524 | 0.814 | 0.659 | 1.01E-19 |
| Gm12346 | 4.49E-24 | 0.721042 | 0.478 | 0.273 | 1.06E-19 |
| Gpr165 | 5.04E-24 | -2.90121 | 0.052 | 0.267 | 1.19E-19 |
| Uap1l1 | 8.44E-24 | 1.308545 | 0.357 | 0.169 | 1.99E-19 |
| Il10ra | 9.68E-24 | -0.97609 | 0.518 | 0.741 | 2.28E-19 |
| Eng | 1.13E-23 | -1.43858 | 0.28 | 0.519 | 2.68E-19 |
| Sft2d1 | 1.34E-23 | -0.72205 | 0.783 | 0.896 | 3.15E-19 |
| Dtnbp1 | 1.53E-23 | 1.028535 | 0.413 | 0.21 | 3.62E-19 |
| Eno1b | 1.56E-23 | 0.953913 | 0.742 | 0.536 | 3.68E-19 |
| Lpar6 | 1.8E-23 | -1.13923 | 0.401 | 0.632 | 4.24E-19 |
| Rpsa-ps9 | 1.93E-23 | 0.32079 | 0.478 | 0.273 | 4.56E-19 |
| Rpl14 | 1.95E-23 | 0.841892 | 0.895 | 0.756 | 4.59E-19 |
| Fblim1 | 1.99E-23 | 1.634682 | 0.249 | 0.098 | 4.7E-19 |

| | | | | | |
|---|---|---|---|---|---|
| Ctsl | 2.08E-23 | 0.817774 | 0.982 | 0.946 | 4.9E-19 |
| Anp32b | 2.31E-23 | 1.377361 | 0.435 | 0.232 | 5.44E-19 |
| Ccl9 | 2.57E-23 | 1.924587 | 0.556 | 0.374 | 6.06E-19 |
| Rps13 | 2.92E-23 | 0.752971 | 0.899 | 0.766 | 6.89E-19 |
| Wasf2 | 3.08E-23 | -0.83047 | 0.72 | 0.862 | 7.26E-19 |
| Hpgd | 3.22E-23 | -0.9025 | 0.558 | 0.746 | 7.59E-19 |
| Fam102b | 3.44E-23 | -1.44705 | 0.464 | 0.665 | 8.12E-19 |
| Rsrp1 | 6.04E-23 | -0.73536 | 0.776 | 0.911 | 1.42E-18 |
| Gm12166 | 6.17E-23 | -0.74631 | 0.639 | 0.817 | 1.46E-18 |
| Rpl19 | 6.53E-23 | 0.76645 | 0.913 | 0.794 | 1.54E-18 |
| Nrip1 | 7.05E-23 | -1.17508 | 0.419 | 0.654 | 1.66E-18 |
| Plekho2 | 7.26E-23 | 1.636775 | 0.296 | 0.131 | 1.71E-18 |
| Dgkz | 9.62E-23 | 1.375091 | 0.475 | 0.265 | 2.27E-18 |
| Rps27a | 1.09E-22 | 0.864892 | 0.906 | 0.78 | 2.58E-18 |
| Rpl18 | 1.47E-22 | 0.778568 | 0.908 | 0.799 | 3.47E-18 |
| Scamp5 | 1.71E-22 | -1.23478 | 0.348 | 0.578 | 4.05E-18 |
| Rgs2 | 1.86E-22 | -0.88408 | 0.691 | 0.841 | 4.38E-18 |
| 0610040J0 | 2.3E-22 | -1.30748 | 0.325 | 0.543 | 5.43E-18 |
| Ang | 2.99E-22 | -1.07862 | 0.552 | 0.755 | 7.06E-18 |
| Ndufa1 | 3.21E-22 | 0.990842 | 0.48 | 0.275 | 7.57E-18 |
| Got1 | 3.26E-22 | 1.282883 | 0.24 | 0.095 | 7.7E-18 |
| Renbp | 3.62E-22 | 1.028883 | 0.536 | 0.32 | 8.55E-18 |
| Ptma | 4.31E-22 | 0.869738 | 0.791 | 0.629 | 1.02E-17 |
| Ddx39 | 5.04E-22 | 1.504622 | 0.222 | 0.085 | 1.19E-17 |
| Gm10180 | 5.43E-22 | 0.475925 | 0.583 | 0.389 | 1.28E-17 |
| Gm4149 | 6.41E-22 | 0.558869 | 0.917 | 0.813 | 1.51E-17 |
| Dusp6 | 6.67E-22 | -1.04371 | 0.529 | 0.731 | 1.57E-17 |
| Hspe1 | 7.1E-22 | 1.003459 | 0.605 | 0.39 | 1.68E-17 |
| Eif3a | 9.2E-22 | 1.425439 | 0.478 | 0.284 | 2.17E-17 |
| Numb | 9.59E-22 | -0.6409 | 0.357 | 0.592 | 2.26E-17 |
| Gm15710 | 1.58E-21 | 0.647733 | 0.863 | 0.723 | 3.73E-17 |
| Gm9118 | 1.69E-21 | 0.707274 | 0.35 | 0.171 | 3.99E-17 |
| Laptm5 | 2.13E-21 | -0.41104 | 1 | 1 | 5.02E-17 |
| Eif4a1 | 2.32E-21 | 1.130153 | 0.776 | 0.615 | 5.47E-17 |
| Rpl17 | 2.4E-21 | 0.807778 | 0.872 | 0.778 | 5.66E-17 |
| Rpl10a-ps1 | 2.73E-21 | 0.721704 | 0.72 | 0.528 | 6.44E-17 |
| Cebpd | 2.87E-21 | -1.51574 | 0.274 | 0.482 | 6.78E-17 |
| Pgam1 | 3.2E-21 | 0.984668 | 0.525 | 0.314 | 7.54E-17 |
| Dock10 | 3.86E-21 | -1.0813 | 0.372 | 0.591 | 9.11E-17 |
| Gal3st4 | 4.88E-21 | -1.06984 | 0.379 | 0.58 | 1.15E-16 |
| Gm3839 | 5.33E-21 | 0.360893 | 0.325 | 0.157 | 1.26E-16 |
| Rnf167 | 5.4E-21 | -1.45939 | 0.327 | 0.524 | 1.27E-16 |
| Rpl5 | 5.76E-21 | 0.81484 | 0.877 | 0.778 | 1.36E-16 |
| Bcl2a1a | 6.39E-21 | 1.083884 | 0.325 | 0.155 | 1.51E-16 |
| Rps16 | 7.21E-21 | 0.761442 | 0.872 | 0.76 | 1.7E-16 |
| Ctsa | 7.83E-21 | 0.535461 | 0.989 | 0.972 | 1.85E-16 |
| Hfe | 8.76E-21 | -1.66776 | 0.235 | 0.452 | 2.07E-16 |

| | | | | | |
|---|---|---|---|---|---|
| Myo9b | 1E-20 | 0.915272 | 0.392 | 0.204 | 2.36E-16 |
| Pebp1 | 1.02E-20 | 1.053443 | 0.489 | 0.291 | 2.42E-16 |
| Plek | 1.26E-20 | 1.291956 | 0.702 | 0.518 | 2.97E-16 |
| Gm49369 | 1.41E-20 | 0.861164 | 0.85 | 0.795 | 3.33E-16 |
| Bmp2k | 1.65E-20 | -0.64976 | 0.805 | 0.933 | 3.9E-16 |
| Sec61g | 1.79E-20 | 0.675301 | 0.596 | 0.372 | 4.23E-16 |
| St3gal5 | 2.27E-20 | -1.16653 | 0.404 | 0.611 | 5.35E-16 |
| Anxa4 | 2.41E-20 | 1.298397 | 0.267 | 0.116 | 5.68E-16 |
| Cox8a | 2.54E-20 | 0.81676 | 0.776 | 0.605 | 5.98E-16 |
| Rpl37a | 2.55E-20 | 0.565348 | 0.964 | 0.929 | 6.01E-16 |
| Gm4735 | 2.75E-20 | 0.9005 | 0.563 | 0.364 | 6.49E-16 |
| Ppia | 2.82E-20 | 0.584551 | 0.982 | 0.969 | 6.66E-16 |
| Tpd52 | 2.94E-20 | 0.903434 | 0.561 | 0.355 | 6.94E-16 |
| Cotl1 | 3.17E-20 | 0.789993 | 0.895 | 0.789 | 7.48E-16 |
| Mtss1 | 4.03E-20 | -1.43929 | 0.152 | 0.37 | 9.51E-16 |
| Eef2k | 4.29E-20 | -1.73933 | 0.229 | 0.448 | 1.01E-15 |
| Slc16a6 | 4.33E-20 | -1.53185 | 0.345 | 0.548 | 1.02E-15 |
| P2rx4 | 6.14E-20 | 0.939392 | 0.587 | 0.389 | 1.45E-15 |
| Ms4a6b | 6.37E-20 | -1.63603 | 0.155 | 0.372 | 1.5E-15 |
| Rps25 | 6.38E-20 | 0.726672 | 0.955 | 0.852 | 1.51E-15 |
| Rps15 | 6.73E-20 | 0.727696 | 0.877 | 0.737 | 1.59E-15 |
| Gm7331 | 7E-20 | 0.577813 | 0.704 | 0.528 | 1.65E-15 |
| Zfp36l2 | 8.45E-20 | -1.58834 | 0.285 | 0.504 | 1.99E-15 |
| Cfl1 | 9.25E-20 | 0.719787 | 0.973 | 0.968 | 2.18E-15 |
| Gm5138 | 1.11E-19 | 0.977614 | 0.626 | 0.455 | 2.62E-15 |
| Gbp7 | 1.12E-19 | -2.12416 | 0.081 | 0.275 | 2.64E-15 |
| Tagln2 | 1.13E-19 | 1.223101 | 0.399 | 0.222 | 2.66E-15 |
| Rps27l | 1.51E-19 | 0.877728 | 0.543 | 0.341 | 3.57E-15 |
| Siglece | 1.64E-19 | -1.68335 | 0.166 | 0.372 | 3.87E-15 |
| Olfr77 | 1.75E-19 | 0.570516 | 0.489 | 0.294 | 4.12E-15 |
| Sft2d2 | 2.02E-19 | -1.63159 | 0.22 | 0.427 | 4.77E-15 |
| Unc93b1 | 2.18E-19 | -0.46727 | 0.969 | 0.991 | 5.14E-15 |
| Eef1g | 2.31E-19 | 0.980495 | 0.592 | 0.399 | 5.45E-15 |
| 1700017B0 | 2.56E-19 | -1.18367 | 0.513 | 0.694 | 6.05E-15 |
| Chd9 | 3.02E-19 | -0.97457 | 0.397 | 0.604 | 7.11E-15 |
| Ccl6 | 3.1E-19 | 1.599794 | 0.599 | 0.421 | 7.3E-15 |
| AI506816 | 3.18E-19 | 1.31366 | 0.419 | 0.254 | 7.49E-15 |
| Adora3 | 3.37E-19 | -1.0042 | 0.305 | 0.523 | 7.94E-15 |
| Rgs10 | 5.03E-19 | -0.49365 | 0.895 | 0.966 | 1.19E-14 |
| Uqcr11 | 6.04E-19 | 0.890847 | 0.594 | 0.397 | 1.43E-14 |
| Cttnbp2nl | 7.46E-19 | -0.84749 | 0.563 | 0.755 | 1.76E-14 |
| Gm4609 | 7.67E-19 | 0.715276 | 0.504 | 0.314 | 1.81E-14 |
| Gm9844 | 7.71E-19 | 0.369819 | 0.231 | 0.097 | 1.82E-14 |
| Pag1 | 7.97E-19 | -0.88861 | 0.581 | 0.779 | 1.88E-14 |
| Rps10 | 9.32E-19 | 0.697469 | 0.922 | 0.786 | 2.2E-14 |
| Rps16-ps2 | 9.54E-19 | 0.819142 | 0.715 | 0.559 | 2.25E-14 |
| Rps7 | 1.01E-18 | 0.822457 | 0.863 | 0.776 | 2.39E-14 |

| | | | | | |
|---|---|---|---|---|---|
| Rpl15 | 1.3E-18 | 0.854126 | 0.807 | 0.672 | 3.06E-14 |
| Rpl9 | 1.42E-18 | 0.751444 | 0.874 | 0.75 | 3.35E-14 |
| Tmem204 | 1.76E-18 | -3.6097 | 0.052 | 0.225 | 4.16E-14 |
| Fcgr3 | 1.76E-18 | -0.53637 | 0.879 | 0.961 | 4.16E-14 |
| Gng10 | 1.81E-18 | -0.8058 | 0.639 | 0.768 | 4.27E-14 |
| Inka1 | 1.93E-18 | -1.51355 | 0.271 | 0.465 | 4.54E-14 |
| Cd33 | 2.02E-18 | -0.72964 | 0.592 | 0.775 | 4.77E-14 |
| Anxa2 | 2.12E-18 | 4.092816 | 0.222 | 0.1 | 4.99E-14 |
| Tuba1a | 2.17E-18 | -1.24822 | 0.446 | 0.61 | 5.13E-14 |
| Soga1 | 2.29E-18 | -1.0417 | 0.379 | 0.586 | 5.41E-14 |
| Fry | 2.54E-18 | -2.26095 | 0.085 | 0.269 | 5.99E-14 |
| Gm5526 | 2.68E-18 | 0.556712 | 0.513 | 0.321 | 6.33E-14 |
| Rpl14-ps1 | 2.79E-18 | 0.637021 | 0.744 | 0.563 | 6.57E-14 |
| Mertk | 3.25E-18 | -0.62735 | 0.717 | 0.872 | 7.66E-14 |
| Ggt5 | 3.57E-18 | -2.7071 | 0.081 | 0.263 | 8.43E-14 |
| Snn | 3.81E-18 | -1.67416 | 0.175 | 0.382 | 8.99E-14 |
| Clec4a2 | 4.06E-18 | -1.5203 | 0.135 | 0.337 | 9.57E-14 |
| Cd84 | 5.61E-18 | 0.923308 | 0.818 | 0.693 | 1.32E-13 |
| Timp2 | 6.22E-18 | 0.694013 | 0.913 | 0.824 | 1.47E-13 |
| Pnp | 6.54E-18 | -1.00328 | 0.408 | 0.591 | 1.54E-13 |
| Gm10275 | 6.55E-18 | 0.785182 | 0.572 | 0.388 | 1.55E-13 |
| S1pr1 | 7.17E-18 | 0.650854 | 0.496 | 0.309 | 1.69E-13 |
| Git2 | 7.53E-18 | -1.01868 | 0.592 | 0.726 | 1.78E-13 |
| Gm29667 | 8.51E-18 | 0.627838 | 0.489 | 0.309 | 2.01E-13 |
| Ctsf | 8.94E-18 | -1.09615 | 0.516 | 0.666 | 2.11E-13 |
| Tmsb10 | 1.05E-17 | 0.910494 | 0.592 | 0.412 | 2.49E-13 |
| Ybx1 | 1.24E-17 | 0.770289 | 0.76 | 0.585 | 2.92E-13 |
| Prkab1 | 1.49E-17 | -1.5509 | 0.186 | 0.381 | 3.52E-13 |
| Slc2a1 | 1.63E-17 | 1.342255 | 0.381 | 0.217 | 3.84E-13 |
| Flna | 1.7E-17 | 0.64876 | 0.249 | 0.112 | 4.01E-13 |
| Sipa1 | 1.72E-17 | -0.87918 | 0.433 | 0.635 | 4.06E-13 |
| Gm4691 | 1.92E-17 | 0.344885 | 0.222 | 0.096 | 4.52E-13 |
| Mtus1 | 2.06E-17 | -1.20095 | 0.383 | 0.592 | 4.87E-13 |
| Pabpc1 | 2.22E-17 | 0.764586 | 0.816 | 0.658 | 5.25E-13 |
| Gm4366 | 2.37E-17 | 0.730654 | 0.43 | 0.253 | 5.58E-13 |
| Gm10224 | 2.42E-17 | 0.469948 | 0.318 | 0.164 | 5.72E-13 |
| Zfp691 | 2.61E-17 | -1.22615 | 0.451 | 0.642 | 6.16E-13 |
| Rps12 | 2.62E-17 | 0.736205 | 0.901 | 0.812 | 6.17E-13 |
| Rps14 | 2.68E-17 | 0.716528 | 0.964 | 0.914 | 6.32E-13 |
| Chsy1 | 3.47E-17 | -1.63623 | 0.307 | 0.496 | 8.19E-13 |
| Pcna-ps2 | 3.47E-17 | 0.607946 | 0.26 | 0.122 | 8.2E-13 |
| Asb2 | 3.59E-17 | -1.69135 | 0.179 | 0.375 | 8.46E-13 |
| Rps17 | 4.06E-17 | 0.628469 | 0.861 | 0.719 | 9.58E-13 |
| Rpl23a | 4.11E-17 | 0.717149 | 0.886 | 0.745 | 9.7E-13 |
| Bax | 5.01E-17 | 0.972265 | 0.439 | 0.265 | 1.18E-12 |
| Adap2 | 5.13E-17 | -0.57404 | 0.818 | 0.928 | 1.21E-12 |
| Fcrl1 | 5.15E-17 | -1.67866 | 0.101 | 0.287 | 1.21E-12 |

| | | | | | |
|---|---|---|---|---|---|
| Rps15a | 5.15E-17 | 0.773132 | 0.738 | 0.596 | 1.22E-12 |
| Akr1a1 | 5.78E-17 | 0.792006 | 0.868 | 0.73 | 1.36E-12 |
| Scoc | 6.13E-17 | -0.69433 | 0.686 | 0.831 | 1.45E-12 |
| Sh3bgrl3 | 6.31E-17 | 0.724002 | 0.906 | 0.812 | 1.49E-12 |
| Tspo | 6.49E-17 | 1.057933 | 0.446 | 0.274 | 1.53E-12 |
| Rpl11 | 6.74E-17 | 0.628311 | 0.877 | 0.713 | 1.59E-12 |
| Gnl3 | 8.41E-17 | 1.415383 | 0.28 | 0.139 | 1.98E-12 |
| Siglecf | 8.74E-17 | 1.438606 | 0.217 | 0.094 | 2.06E-12 |
| Rpl31 | 8.82E-17 | 0.617915 | 0.886 | 0.785 | 2.08E-12 |
| Commd4 | 9.04E-17 | 0.918614 | 0.361 | 0.197 | 2.13E-12 |
| Hint1 | 9.78E-17 | 0.871714 | 0.574 | 0.387 | 2.31E-12 |
| Atp5k | 1.11E-16 | 0.550771 | 0.466 | 0.277 | 2.63E-12 |
| Gm6477 | 1.16E-16 | 0.719161 | 0.26 | 0.123 | 2.74E-12 |
| Ulk2 | 1.26E-16 | -0.99972 | 0.392 | 0.575 | 2.97E-12 |
| Hmgn2 | 1.5E-16 | 0.728402 | 0.556 | 0.357 | 3.53E-12 |
| Tpt1 | 1.59E-16 | 0.704792 | 0.93 | 0.878 | 3.74E-12 |
| Eprs | 1.72E-16 | 0.904488 | 0.419 | 0.248 | 4.06E-12 |
| Idh2 | 1.73E-16 | -1.4754 | 0.258 | 0.444 | 4.09E-12 |
| Rps19-ps6 | 1.75E-16 | 0.579654 | 0.78 | 0.618 | 4.12E-12 |
| Cndp2 | 1.93E-16 | 0.750832 | 0.652 | 0.459 | 4.55E-12 |
| Rpl28 | 1.97E-16 | 0.684788 | 0.845 | 0.723 | 4.64E-12 |
| Calm1 | 2.23E-16 | 0.759246 | 0.863 | 0.748 | 5.25E-12 |
| Slc40a1 | 2.67E-16 | -1.02859 | 0.318 | 0.522 | 6.31E-12 |
| Gtf2h2 | 2.74E-16 | -0.90204 | 0.442 | 0.628 | 6.47E-12 |
| Aprt | 2.91E-16 | 1.220797 | 0.305 | 0.16 | 6.87E-12 |
| Ntpcr | 2.95E-16 | -0.93532 | 0.462 | 0.634 | 6.97E-12 |
| I830077J02 | 3.09E-16 | -1.7066 | 0.222 | 0.41 | 7.29E-12 |
| Tgfbr1 | 3.14E-16 | -0.57315 | 0.843 | 0.939 | 7.4E-12 |
| Tsc22d4 | 3.69E-16 | -0.98342 | 0.498 | 0.667 | 8.7E-12 |
| Grn | 3.72E-16 | 0.452658 | 1 | 0.995 | 8.77E-12 |
| Amdhd2 | 3.79E-16 | 1.511662 | 0.238 | 0.111 | 8.94E-12 |
| Rps23 | 3.85E-16 | 0.559873 | 0.955 | 0.887 | 9.08E-12 |
| Tmem59 | 3.97E-16 | -0.64481 | 0.816 | 0.884 | 9.37E-12 |
| Lat2 | 3.99E-16 | 0.803908 | 0.536 | 0.358 | 9.4E-12 |
| Rpl18a | 4.04E-16 | 0.513122 | 0.991 | 0.986 | 9.54E-12 |
| Gm9794 | 4.42E-16 | 0.763507 | 0.765 | 0.606 | 1.04E-11 |
| Fscn1 | 5.34E-16 | -0.77435 | 0.706 | 0.859 | 1.26E-11 |
| B4galt1 | 5.52E-16 | -0.64688 | 0.762 | 0.87 | 1.3E-11 |
| Fau | 5.61E-16 | 0.600848 | 0.955 | 0.899 | 1.32E-11 |
| Elob | 5.66E-16 | 0.72274 | 0.652 | 0.47 | 1.33E-11 |
| Fam162a | 5.89E-16 | 1.152366 | 0.242 | 0.113 | 1.39E-11 |
| Adap2os | 6.65E-16 | -0.57799 | 0.821 | 0.93 | 1.57E-11 |
| Rpl39 | 6.88E-16 | 0.599991 | 0.832 | 0.699 | 1.62E-11 |
| Ranbp1 | 6.98E-16 | 0.937143 | 0.446 | 0.273 | 1.65E-11 |
| Gm11539 | 8.03E-16 | 0.532887 | 0.46 | 0.292 | 1.89E-11 |
| Zfp36l1 | 9.26E-16 | -0.49529 | 0.617 | 0.78 | 2.18E-11 |
| Pla2g15 | 9.35E-16 | -0.78954 | 0.695 | 0.803 | 2.21E-11 |

| | | | | | |
|---|---|---|---|---|---|
| Slc25a5 | 1.11E-15 | 0.648309 | 0.89 | 0.811 | 2.61E-11 |
| Rpl6 | 1.11E-15 | 0.689472 | 0.861 | 0.753 | 2.62E-11 |
| mt-Tp | 1.12E-15 | 1.207495 | 0.307 | 0.164 | 2.65E-11 |
| Gm5915 | 1.14E-15 | 0.459784 | 0.354 | 0.199 | 2.69E-11 |
| Cbfa2t3 | 1.17E-15 | -1.57882 | 0.195 | 0.375 | 2.77E-11 |
| Rpsa-ps2 | 1.26E-15 | 0.406863 | 0.327 | 0.181 | 2.98E-11 |
| Cyth4 | 1.32E-15 | -0.702 | 0.814 | 0.894 | 3.11E-11 |
| Atp6v1a | 1.42E-15 | 1.300861 | 0.372 | 0.21 | 3.35E-11 |
| Inpp5d | 1.58E-15 | -0.53513 | 0.785 | 0.919 | 3.74E-11 |
| Gm10039 | 1.94E-15 | 0.552882 | 0.386 | 0.215 | 4.57E-11 |
| Rps15a-ps6 | 2.1E-15 | 0.875778 | 0.386 | 0.226 | 4.96E-11 |
| Sla | 2.26E-15 | -0.96503 | 0.357 | 0.545 | 5.33E-11 |
| Eif4a-ps4 | 2.46E-15 | 0.833529 | 0.574 | 0.411 | 5.8E-11 |
| Mien1 | 2.5E-15 | 0.865947 | 0.332 | 0.179 | 5.89E-11 |
| Lamp1 | 2.52E-15 | 0.500841 | 0.98 | 0.947 | 5.96E-11 |
| Rpl19-ps11 | 2.55E-15 | 0.676117 | 0.751 | 0.589 | 6.01E-11 |
| Mitf | 2.55E-15 | 0.772774 | 0.413 | 0.245 | 6.01E-11 |
| Plcl2 | 2.7E-15 | -1.04194 | 0.321 | 0.52 | 6.36E-11 |
| Rpl4 | 2.79E-15 | 0.594457 | 0.915 | 0.829 | 6.59E-11 |
| Cst3 | 2.9E-15 | -0.31287 | 1 | 1 | 6.83E-11 |
| Rogdi | 3.07E-15 | -0.99769 | 0.404 | 0.585 | 7.24E-11 |
| Slc11a1 | 3.31E-15 | 0.729749 | 0.765 | 0.595 | 7.81E-11 |
| Gpx1 | 3.56E-15 | 0.668065 | 0.85 | 0.708 | 8.4E-11 |
| Epb41l2 | 4.11E-15 | -0.69778 | 0.787 | 0.89 | 9.7E-11 |
| Txn1 | 4.3E-15 | 0.845465 | 0.408 | 0.246 | 1.01E-10 |
| Cox6c | 4.45E-15 | 0.772329 | 0.621 | 0.441 | 1.05E-10 |
| Neat1 | 4.45E-15 | 1.175108 | 0.325 | 0.183 | 1.05E-10 |
| Whrn | 4.8E-15 | -1.18029 | 0.186 | 0.374 | 1.13E-10 |
| Gm15503 | 4.89E-15 | -0.99848 | 0.247 | 0.443 | 1.15E-10 |
| Kcnk6 | 4.97E-15 | -1.18213 | 0.383 | 0.554 | 1.17E-10 |
| Gaa | 5.28E-15 | 0.81678 | 0.534 | 0.365 | 1.24E-10 |
| Ssr4 | 5.4E-15 | 0.697244 | 0.776 | 0.626 | 1.27E-10 |
| Rpl36 | 6.69E-15 | 0.604909 | 0.899 | 0.799 | 1.58E-10 |
| Pld1 | 7.63E-15 | -1.55567 | 0.269 | 0.461 | 1.8E-10 |
| Rpl34 | 7.92E-15 | 0.577609 | 0.899 | 0.778 | 1.87E-10 |
| Atp6v1b2 | 8.03E-15 | 0.806319 | 0.498 | 0.316 | 1.89E-10 |
| Arhgap27 | 8.04E-15 | -1.61888 | 0.166 | 0.342 | 1.9E-10 |
| Arl5c | 8.08E-15 | 2.178234 | 0.211 | 0.097 | 1.91E-10 |
| Rpl30 | 8.98E-15 | 0.603626 | 0.933 | 0.858 | 2.12E-10 |
| Rpl36-ps2 | 9.34E-15 | 0.598946 | 0.637 | 0.478 | 2.2E-10 |
| Gusb | 1.03E-14 | 0.668885 | 0.771 | 0.622 | 2.43E-10 |
| Gm11361 | 1.18E-14 | 0.523127 | 0.594 | 0.434 | 2.78E-10 |
| Ctsc | 1.18E-14 | 0.618712 | 0.91 | 0.842 | 2.78E-10 |
| Rpl9-ps6 | 1.22E-14 | 0.720169 | 0.76 | 0.585 | 2.88E-10 |
| Mgat1 | 1.25E-14 | -1.18866 | 0.41 | 0.574 | 2.94E-10 |
| Gm10443 | 1.31E-14 | 0.422183 | 0.538 | 0.37 | 3.09E-10 |
| Gm15500 | 1.41E-14 | 1.007298 | 0.74 | 0.601 | 3.33E-10 |

| | | | | | |
|---|---|---|---|---|---|
| Rpl36a | 1.43E-14 | 0.744968 | 0.688 | 0.511 | 3.38E-10 |
| Tmem160 | 1.56E-14 | 0.506068 | 0.496 | 0.311 | 3.68E-10 |
| Grina | 1.57E-14 | 0.588562 | 0.722 | 0.557 | 3.71E-10 |
| Rps11 | 1.58E-14 | 0.634669 | 0.944 | 0.911 | 3.73E-10 |
| Eef1b2 | 1.67E-14 | 0.749019 | 0.61 | 0.445 | 3.94E-10 |
| Gm2606 | 1.68E-14 | 0.284983 | 0.28 | 0.145 | 3.96E-10 |
| Eif4ebp1 | 1.69E-14 | 1.135066 | 0.287 | 0.151 | 3.99E-10 |
| Cysltr1 | 1.9E-14 | -1.24124 | 0.157 | 0.336 | 4.48E-10 |
| Rps3 | 1.9E-14 | 0.589385 | 0.951 | 0.897 | 4.48E-10 |
| Vim | 2.49E-14 | 3.562918 | 0.209 | 0.101 | 5.89E-10 |
| Zfp652 | 2.51E-14 | -0.75948 | 0.525 | 0.694 | 5.93E-10 |
| Dok3 | 2.68E-14 | -2.00685 | 0.123 | 0.286 | 6.31E-10 |
| Btg2 | 2.83E-14 | -1.11864 | 0.433 | 0.584 | 6.67E-10 |
| Rps24 | 2.84E-14 | 0.677567 | 0.91 | 0.818 | 6.71E-10 |
| Prpsap2 | 3.35E-14 | -1.31088 | 0.184 | 0.359 | 7.89E-10 |
| Akirin2 | 3.62E-14 | -1.36553 | 0.265 | 0.428 | 8.55E-10 |
| Cox4i1 | 3.87E-14 | 0.61091 | 0.863 | 0.775 | 9.13E-10 |
| Kctd12 | 3.92E-14 | -0.62438 | 0.594 | 0.756 | 9.26E-10 |
| Dleu2 | 4.1E-14 | -0.45662 | 0.691 | 0.85 | 9.67E-10 |
| Dip2b | 4.17E-14 | -0.85874 | 0.437 | 0.603 | 9.84E-10 |
| Rps18-ps3 | 4.22E-14 | 0.716906 | 0.686 | 0.556 | 9.95E-10 |
| Tm6sf1 | 4.4E-14 | -1.26336 | 0.291 | 0.459 | 1.04E-09 |
| Dnase2a | 4.46E-14 | 0.652312 | 0.632 | 0.46 | 1.05E-09 |
| Fli1 | 4.76E-14 | -1.08302 | 0.48 | 0.645 | 1.12E-09 |
| Cd48 | 4.77E-14 | 0.818443 | 0.549 | 0.379 | 1.12E-09 |
| Ptp4a3 | 5.06E-14 | -1.15076 | 0.374 | 0.531 | 1.19E-09 |
| Hsd17b12 | 5.56E-14 | 0.556819 | 0.482 | 0.313 | 1.31E-09 |
| Rpl21 | 6.71E-14 | 0.632738 | 0.942 | 0.917 | 1.58E-09 |
| Cxxc5 | 6.82E-14 | -0.84893 | 0.572 | 0.707 | 1.61E-09 |
| Grk2 | 6.93E-14 | -0.894 | 0.453 | 0.619 | 1.63E-09 |
| Gm5844 | 6.99E-14 | 0.826813 | 0.379 | 0.228 | 1.65E-09 |
| Rpl23a-ps3 | 7.08E-14 | 0.596745 | 0.628 | 0.467 | 1.67E-09 |
| Clcn7 | 7.1E-14 | 0.727343 | 0.502 | 0.327 | 1.67E-09 |
| Tceal9 | 7.29E-14 | 1.337632 | 0.22 | 0.106 | 1.72E-09 |
| Lyl1 | 8.6E-14 | -1.11265 | 0.35 | 0.51 | 2.03E-09 |
| Bank1 | 8.64E-14 | -2.47156 | 0.058 | 0.2 | 2.04E-09 |
| Sars | 9.38E-14 | 0.855642 | 0.417 | 0.261 | 2.21E-09 |
| Psen2 | 9.68E-14 | 1.347228 | 0.238 | 0.118 | 2.28E-09 |
| C5ar1 | 1.03E-13 | 1.186034 | 0.576 | 0.426 | 2.44E-09 |
| Itgam | 1.15E-13 | -0.67565 | 0.679 | 0.813 | 2.72E-09 |
| Dusp3 | 1.16E-13 | 0.94041 | 0.518 | 0.356 | 2.73E-09 |
| Tcn2 | 1.18E-13 | -0.65749 | 0.742 | 0.836 | 2.78E-09 |
| Gm1840 | 1.21E-13 | 0.433156 | 0.287 | 0.156 | 2.86E-09 |
| Csmd3 | 1.25E-13 | -1.26867 | 0.231 | 0.412 | 2.96E-09 |
| Atp5e | 1.28E-13 | 0.737219 | 0.657 | 0.481 | 3.03E-09 |
| Ddit4 | 1.51E-13 | -1.41647 | 0.099 | 0.25 | 3.55E-09 |
| Smpdl3a | 1.53E-13 | 1.280016 | 0.224 | 0.109 | 3.61E-09 |

| | | | | | |
|---|---|---|---|---|---|
| Txndc17 | 1.59E-13 | 0.776801 | 0.397 | 0.241 | 3.74E-09 |
| Frmd4a | 1.78E-13 | -0.65713 | 0.574 | 0.754 | 4.19E-09 |
| Tomm20 | 1.93E-13 | 0.838091 | 0.498 | 0.332 | 4.55E-09 |
| Scd2 | 2.03E-13 | 0.653616 | 0.28 | 0.148 | 4.79E-09 |
| Tifab | 2.21E-13 | -0.95367 | 0.408 | 0.577 | 5.22E-09 |
| Serf2 | 2.27E-13 | 0.353011 | 0.857 | 0.721 | 5.34E-09 |
| Gng5 | 2.58E-13 | 0.592872 | 0.796 | 0.664 | 6.08E-09 |
| Cox6a1 | 2.75E-13 | 0.690417 | 0.697 | 0.554 | 6.49E-09 |
| Ccng2 | 2.93E-13 | -1.27459 | 0.197 | 0.365 | 6.91E-09 |
| Rpl26 | 3.29E-13 | 0.598853 | 0.848 | 0.763 | 7.75E-09 |
| Txn2 | 3.33E-13 | 0.826033 | 0.372 | 0.225 | 7.85E-09 |
| Creg1 | 3.39E-13 | 0.543475 | 0.834 | 0.695 | 7.99E-09 |
| Slc25a3 | 3.4E-13 | 0.694727 | 0.697 | 0.543 | 8.02E-09 |
| Fam221b | 3.45E-13 | 0.649544 | 0.583 | 0.422 | 8.13E-09 |
| Prune2 | 3.47E-13 | 0.637924 | 0.413 | 0.253 | 8.19E-09 |
| Cdk5r1 | 3.65E-13 | -1.56992 | 0.126 | 0.283 | 8.62E-09 |
| Nfam1 | 3.71E-13 | -0.74011 | 0.462 | 0.632 | 8.75E-09 |
| Rps19-ps4 | 3.74E-13 | 0.527133 | 0.217 | 0.106 | 8.82E-09 |
| Ogfrl1 | 3.75E-13 | -1.00783 | 0.417 | 0.571 | 8.85E-09 |
| Med12l | 3.85E-13 | -0.90679 | 0.381 | 0.554 | 9.09E-09 |
| Naa50 | 3.89E-13 | 1.328196 | 0.233 | 0.117 | 9.17E-09 |
| Gm4294 | 4.01E-13 | 0.482179 | 0.487 | 0.327 | 9.45E-09 |
| Pgam1-ps2 | 4.3E-13 | 0.375263 | 0.309 | 0.174 | 1.01E-08 |
| Gm3511 | 4.3E-13 | 0.544867 | 0.682 | 0.534 | 1.01E-08 |
| Sesn1 | 4.38E-13 | -0.96929 | 0.269 | 0.439 | 1.03E-08 |
| Tnfaip8l2 | 4.75E-13 | -0.79073 | 0.473 | 0.62 | 1.12E-08 |
| Gm8129 | 5.6E-13 | 0.609637 | 0.413 | 0.265 | 1.32E-08 |
| Ran | 5.84E-13 | 0.860152 | 0.543 | 0.388 | 1.38E-08 |
| Banf1 | 5.86E-13 | 0.716927 | 0.401 | 0.244 | 1.38E-08 |
| Rtn1 | 5.9E-13 | -1.02037 | 0.222 | 0.391 | 1.39E-08 |
| Gm5879 | 5.97E-13 | 0.563659 | 0.484 | 0.329 | 1.41E-08 |
| Plin2 | 6.05E-13 | 1.145575 | 0.22 | 0.109 | 1.43E-08 |
| Arhgef40 | 6.08E-13 | -0.89406 | 0.289 | 0.469 | 1.43E-08 |
| Snta1 | 6.13E-13 | -1.30255 | 0.155 | 0.319 | 1.45E-08 |
| Cycs | 6.23E-13 | 0.801626 | 0.464 | 0.297 | 1.47E-08 |
| Rpl37 | 6.54E-13 | 0.475339 | 0.951 | 0.886 | 1.54E-08 |
| Slc6a6 | 6.55E-13 | 0.611033 | 0.554 | 0.383 | 1.54E-08 |
| Ypel3 | 6.86E-13 | -0.59203 | 0.509 | 0.671 | 1.62E-08 |
| Tmem256 | 7.1E-13 | 0.549526 | 0.444 | 0.279 | 1.67E-08 |
| Tmem100 | 7.89E-13 | -1.31423 | 0.2 | 0.362 | 1.86E-08 |
| Myl12a | 8.15E-13 | 0.760668 | 0.648 | 0.48 | 1.92E-08 |
| Gm8430 | 8.3E-13 | 0.578596 | 0.648 | 0.496 | 1.96E-08 |
| Dpysl2 | 8.36E-13 | -0.69568 | 0.78 | 0.878 | 1.97E-08 |
| Soat1 | 8.75E-13 | 0.893182 | 0.48 | 0.318 | 2.06E-08 |
| Clec4a3 | 8.81E-13 | -1.1474 | 0.253 | 0.415 | 2.08E-08 |
| Psme1 | 8.92E-13 | 0.720479 | 0.646 | 0.48 | 2.1E-08 |
| Gm10132 | 8.95E-13 | 0.725989 | 0.704 | 0.555 | 2.11E-08 |

| | | | | | |
|---|---|---|---|---|---|
| Gm11836 | 9.06E-13 | 0.278873 | 0.211 | 0.103 | 2.14E-08 |
| Ncoa3 | 9.27E-13 | -0.78245 | 0.621 | 0.759 | 2.19E-08 |
| Mef2a | 9.4E-13 | -0.6641 | 0.682 | 0.81 | 2.22E-08 |
| Alas1 | 9.5E-13 | 0.867162 | 0.386 | 0.239 | 2.24E-08 |
| Foxn3 | 1.04E-12 | -0.60445 | 0.769 | 0.872 | 2.45E-08 |
| Arid1a | 1.1E-12 | -0.76413 | 0.469 | 0.632 | 2.59E-08 |
| Rpl27a | 1.11E-12 | 0.588739 | 0.848 | 0.733 | 2.61E-08 |
| Mybbp1a | 1.14E-12 | 0.695889 | 0.321 | 0.184 | 2.69E-08 |
| Irgm2 | 1.22E-12 | -1.22741 | 0.217 | 0.383 | 2.87E-08 |
| Gm33699 | 1.28E-12 | -1.34763 | 0.105 | 0.258 | 3.01E-08 |
| Nap1l1 | 1.31E-12 | 0.817538 | 0.274 | 0.149 | 3.1E-08 |
| Rpl31-ps8 | 1.43E-12 | 0.635603 | 0.72 | 0.579 | 3.36E-08 |
| Gng12 | 1.44E-12 | 1.263691 | 0.274 | 0.152 | 3.39E-08 |
| Slamf9 | 1.46E-12 | 0.840773 | 0.473 | 0.321 | 3.46E-08 |
| Napa | 1.47E-12 | 0.926025 | 0.453 | 0.303 | 3.48E-08 |
| Atp6v0a2 | 1.53E-12 | -1.41416 | 0.276 | 0.432 | 3.61E-08 |
| Uqcrq | 1.53E-12 | 0.697585 | 0.518 | 0.37 | 3.62E-08 |
| Gp9 | 1.6E-12 | -2.36907 | 0.09 | 0.228 | 3.77E-08 |
| Gm10073 | 1.62E-12 | 0.396632 | 0.724 | 0.568 | 3.82E-08 |
| Fchsd2 | 1.75E-12 | -1.32431 | 0.267 | 0.429 | 4.12E-08 |
| Hspa9 | 1.77E-12 | 1.032402 | 0.388 | 0.242 | 4.18E-08 |
| Lipa | 1.95E-12 | 0.97218 | 0.576 | 0.436 | 4.59E-08 |
| Ifitm10 | 2.03E-12 | -0.81274 | 0.368 | 0.528 | 4.78E-08 |
| Mettl1 | 2.13E-12 | 1.085493 | 0.211 | 0.103 | 5.03E-08 |
| Cox5a | 2.37E-12 | 0.74399 | 0.453 | 0.29 | 5.6E-08 |
| Eif5a | 2.51E-12 | 0.713583 | 0.742 | 0.591 | 5.92E-08 |
| Slfn2 | 2.61E-12 | 1.003437 | 0.238 | 0.122 | 6.16E-08 |
| Gm12918 | 2.68E-12 | 0.547611 | 0.632 | 0.496 | 6.32E-08 |
| Gm10123 | 2.83E-12 | 0.613385 | 0.948 | 0.927 | 6.68E-08 |
| Cops9 | 2.94E-12 | 0.520929 | 0.567 | 0.403 | 6.93E-08 |
| Sh2b3 | 2.94E-12 | -0.8881 | 0.276 | 0.449 | 6.95E-08 |
| Rpl24 | 2.95E-12 | 0.637279 | 0.731 | 0.604 | 6.97E-08 |
| Dapp1 | 3E-12 | -1.18356 | 0.253 | 0.41 | 7.08E-08 |
| Rpl27 | 3.05E-12 | 0.584197 | 0.807 | 0.688 | 7.19E-08 |
| Glmp | 3.18E-12 | 0.543256 | 0.702 | 0.545 | 7.51E-08 |
| H2-Ab1 | 3.22E-12 | 3.659096 | 0.26 | 0.154 | 7.59E-08 |
| Rps10-ps2 | 3.23E-12 | 0.620029 | 0.726 | 0.571 | 7.62E-08 |
| Sema4b | 3.25E-12 | -1.74381 | 0.072 | 0.213 | 7.66E-08 |
| Rpl22 | 3.5E-12 | 0.657044 | 0.684 | 0.517 | 8.26E-08 |
| Bri3bp | 3.67E-12 | 1.349591 | 0.202 | 0.099 | 8.65E-08 |
| Atp5l | 3.74E-12 | 0.579023 | 0.637 | 0.472 | 8.82E-08 |
| Gm15501 | 3.98E-12 | 0.674999 | 0.621 | 0.466 | 9.39E-08 |
| Degs1 | 4.07E-12 | 0.734064 | 0.439 | 0.286 | 9.6E-08 |
| Eif5 | 4.11E-12 | 0.6765 | 0.574 | 0.409 | 9.69E-08 |
| Ikzf1 | 4.26E-12 | -0.86452 | 0.386 | 0.553 | 1E-07 |
| Gm26917 | 4.39E-12 | 1.228374 | 0.89 | 0.951 | 1.04E-07 |
| Crlf2 | 4.42E-12 | 0.868293 | 0.525 | 0.372 | 1.04E-07 |

| | | | | | |
|---|---|---|---|---|---|
| Pip4k2a | 4.71E-12 | -0.82066 | 0.419 | 0.579 | 1.11E-07 |
| Lrpap1 | 4.76E-12 | 0.928833 | 0.581 | 0.436 | 1.12E-07 |
| Atp6v1g1 | 4.78E-12 | 0.570459 | 0.776 | 0.605 | 1.13E-07 |
| Colgalt1 | 4.89E-12 | 0.572599 | 0.451 | 0.295 | 1.15E-07 |
| Atp6v1c1 | 4.96E-12 | 0.680138 | 0.525 | 0.359 | 1.17E-07 |
| Gm14586 | 5.4E-12 | 0.51258 | 0.59 | 0.436 | 1.27E-07 |
| Rpl38-ps2 | 5.88E-12 | 0.429206 | 0.343 | 0.209 | 1.39E-07 |
| Gm10736 | 6.13E-12 | 0.593734 | 0.626 | 0.489 | 1.45E-07 |
| Gm8186 | 6.15E-12 | 0.631377 | 0.419 | 0.267 | 1.45E-07 |
| Capn3 | 6.21E-12 | -1.07974 | 0.262 | 0.422 | 1.46E-07 |
| Zfp69 | 6.76E-12 | -1.43684 | 0.217 | 0.369 | 1.59E-07 |
| Rps13-ps1 | 6.82E-12 | 0.34526 | 0.442 | 0.289 | 1.61E-07 |
| Hspd1 | 7.25E-12 | 0.792912 | 0.37 | 0.229 | 1.71E-07 |
| Dock4 | 7.34E-12 | -0.7898 | 0.442 | 0.6 | 1.73E-07 |
| Gm4617 | 7.45E-12 | 0.432993 | 0.41 | 0.266 | 1.76E-07 |
| Cox5b | 7.52E-12 | 0.640768 | 0.543 | 0.398 | 1.77E-07 |
| Tpt1-ps3 | 7.55E-12 | 0.555496 | 0.735 | 0.62 | 1.78E-07 |
| Tnfrsf13b | 7.56E-12 | -1.44705 | 0.179 | 0.331 | 1.78E-07 |
| Gm28437 | 7.68E-12 | 0.682643 | 0.881 | 0.825 | 1.81E-07 |
| Etv5 | 8.19E-12 | -0.89193 | 0.251 | 0.418 | 1.93E-07 |
| C1qbp | 8.4E-12 | 0.709531 | 0.361 | 0.22 | 1.98E-07 |
| Ac165079.1 | 8.46E-12 | 0.305183 | 0.359 | 0.221 | 2E-07 |
| Tcf7l2 | 9.07E-12 | -1.29077 | 0.3 | 0.452 | 2.14E-07 |
| Gabarap | 9.24E-12 | 0.566229 | 0.841 | 0.727 | 2.18E-07 |
| Nop56 | 9.25E-12 | 0.924024 | 0.345 | 0.21 | 2.18E-07 |
| Itgb5 | 9.39E-12 | -0.33723 | 0.944 | 0.969 | 2.22E-07 |
| Klf3 | 9.61E-12 | -0.85664 | 0.435 | 0.588 | 2.27E-07 |
| Gm11942 | 9.94E-12 | 0.427144 | 0.545 | 0.408 | 2.34E-07 |
| Galnt7 | 1.01E-11 | 1.147603 | 0.217 | 0.109 | 2.39E-07 |
| Fam49b | 1.06E-11 | -0.67954 | 0.639 | 0.749 | 2.5E-07 |
| Ifrd1 | 1.07E-11 | 1.343457 | 0.231 | 0.122 | 2.53E-07 |
| Gm8995 | 1.1E-11 | 0.974081 | 0.312 | 0.182 | 2.59E-07 |
| Ncf1 | 1.12E-11 | -0.63998 | 0.608 | 0.735 | 2.64E-07 |
| Ptms | 1.15E-11 | 0.730597 | 0.83 | 0.693 | 2.72E-07 |
| Atp5mpl | 1.19E-11 | 0.708608 | 0.419 | 0.273 | 2.82E-07 |
| Srgn | 1.2E-11 | 0.708851 | 0.599 | 0.457 | 2.83E-07 |
| Csnk1e | 1.2E-11 | -0.62137 | 0.473 | 0.622 | 2.83E-07 |
| Cd53 | 1.27E-11 | -0.44928 | 0.937 | 0.969 | 3E-07 |
| Anapc15 | 1.3E-11 | 1.278435 | 0.202 | 0.1 | 3.06E-07 |
| Cep68 | 1.34E-11 | -1.31908 | 0.164 | 0.32 | 3.17E-07 |
| Rpl3-ps2 | 1.35E-11 | 0.309635 | 0.303 | 0.18 | 3.17E-07 |
| Gm3788 | 1.37E-11 | 0.419472 | 0.224 | 0.117 | 3.23E-07 |
| Snrpg | 1.38E-11 | 0.555707 | 0.5 | 0.337 | 3.26E-07 |
| Tmbim1 | 1.44E-11 | 0.83647 | 0.318 | 0.188 | 3.4E-07 |
| Ctdnep1 | 1.6E-11 | 1.382858 | 0.296 | 0.179 | 3.78E-07 |
| Psap | 1.65E-11 | 0.421211 | 1 | 0.999 | 3.9E-07 |
| Bri3 | 1.76E-11 | 0.636656 | 0.726 | 0.573 | 4.14E-07 |

| | | | | | |
|---|---|---|---|---|---|
| Ap1b1 | 2.06E-11 | -0.84029 | 0.296 | 0.459 | 4.86E-07 |
| Atp5b | 2.18E-11 | 0.553751 | 0.836 | 0.75 | 5.14E-07 |
| Gba | 2.23E-11 | 0.663883 | 0.572 | 0.428 | 5.27E-07 |
| Tlr1 | 2.27E-11 | 0.64078 | 0.291 | 0.164 | 5.35E-07 |
| Rpl7 | 2.32E-11 | 0.512198 | 0.879 | 0.792 | 5.46E-07 |
| Gm7536 | 2.36E-11 | 0.45885 | 0.704 | 0.562 | 5.56E-07 |
| Creld2 | 2.52E-11 | 0.92719 | 0.41 | 0.267 | 5.95E-07 |
| Tbca | 2.54E-11 | 0.845537 | 0.325 | 0.198 | 6E-07 |
| Usmg5 | 2.59E-11 | 0.632746 | 0.491 | 0.336 | 6.1E-07 |
| Limd2 | 2.6E-11 | -0.51419 | 0.666 | 0.8 | 6.14E-07 |
| Rpl18-ps2 | 2.64E-11 | 0.519031 | 0.484 | 0.344 | 6.23E-07 |
| Snx18 | 2.66E-11 | -0.81152 | 0.498 | 0.648 | 6.28E-07 |
| Rpl10 | 2.68E-11 | 0.529311 | 0.948 | 0.901 | 6.31E-07 |
| Cln5 | 2.74E-11 | 0.560851 | 0.406 | 0.251 | 6.46E-07 |
| Sdf2l1 | 2.77E-11 | 0.799887 | 0.599 | 0.458 | 6.53E-07 |
| Atp1a1 | 2.85E-11 | 0.591969 | 0.623 | 0.47 | 6.73E-07 |
| Syngr1 | 2.86E-11 | 0.625528 | 0.901 | 0.839 | 6.73E-07 |
| Zbtb20 | 2.97E-11 | -0.8778 | 0.298 | 0.459 | 7E-07 |
| Tgif1 | 2.99E-11 | 0.859921 | 0.321 | 0.192 | 7.06E-07 |
| Gm5529 | 3.02E-11 | 0.446403 | 0.471 | 0.341 | 7.13E-07 |
| mt-Nd1 | 3.12E-11 | 0.610749 | 0.955 | 0.935 | 7.35E-07 |
| Rpl35a | 3.34E-11 | 0.580182 | 0.814 | 0.732 | 7.88E-07 |
| Tmem273 | 3.46E-11 | -1.57732 | 0.17 | 0.311 | 8.16E-07 |
| Nupr1 | 3.58E-11 | 1.448045 | 0.235 | 0.129 | 8.45E-07 |
| Atp6v1f | 3.6E-11 | 0.551733 | 0.693 | 0.529 | 8.49E-07 |
| Higd1a | 3.6E-11 | 0.71597 | 0.291 | 0.167 | 8.5E-07 |
| Nhp2 | 3.69E-11 | 1.086692 | 0.231 | 0.123 | 8.7E-07 |
| Vegfb | 3.78E-11 | 0.81221 | 0.404 | 0.266 | 8.92E-07 |
| Ubash3b | 3.79E-11 | -0.64705 | 0.464 | 0.608 | 8.94E-07 |
| Rpl7a | 3.85E-11 | 0.670627 | 0.8 | 0.691 | 9.07E-07 |
| Cox7c | 3.85E-11 | 0.397001 | 0.717 | 0.555 | 9.09E-07 |
| Srm | 3.93E-11 | 0.914204 | 0.365 | 0.232 | 9.27E-07 |
| Fcgr2b | 4.11E-11 | -0.53675 | 0.684 | 0.802 | 9.71E-07 |
| Hpgds | 4.27E-11 | -0.46679 | 0.879 | 0.944 | 1.01E-06 |
| Cox7b | 4.65E-11 | 0.722235 | 0.442 | 0.295 | 1.1E-06 |
| Rbbp7 | 4.87E-11 | 0.63352 | 0.417 | 0.271 | 1.15E-06 |
| Mrpl33 | 5.21E-11 | 0.827016 | 0.305 | 0.184 | 1.23E-06 |
| Mapk14 | 5.23E-11 | -1.36753 | 0.238 | 0.382 | 1.23E-06 |
| Ighm | 5.49E-11 | -0.66854 | 0.525 | 0.666 | 1.3E-06 |
| Rpl28-ps1 | 5.58E-11 | 0.673821 | 0.578 | 0.426 | 1.32E-06 |
| Nfia | 5.92E-11 | -1.0209 | 0.312 | 0.472 | 1.4E-06 |
| Uqcc2 | 6.05E-11 | 0.742721 | 0.368 | 0.23 | 1.43E-06 |
| Ubc | 6.1E-11 | -0.26345 | 0.939 | 0.974 | 1.44E-06 |
| Pan3 | 6.27E-11 | -0.39544 | 0.565 | 0.708 | 1.48E-06 |
| Polr2l | 6.63E-11 | 0.424456 | 0.291 | 0.165 | 1.56E-06 |
| Dnajb14 | 6.71E-11 | 0.55235 | 0.341 | 0.207 | 1.58E-06 |
| Spg21 | 7.13E-11 | 0.892979 | 0.37 | 0.234 | 1.68E-06 |

| | | | | | |
|---|---|---|---|---|---|
| Gm13436 | 7.38E-11 | 0.56462 | 0.617 | 0.47 | 1.74E-06 |
| Gsto1 | 7.67E-11 | 0.88705 | 0.238 | 0.128 | 1.81E-06 |
| Gm32036 | 7.71E-11 | -0.80776 | 0.316 | 0.471 | 1.82E-06 |
| Abl1 | 8.06E-11 | -1.70594 | 0.184 | 0.327 | 1.9E-06 |
| Tle3 | 8.26E-11 | -1.10852 | 0.173 | 0.317 | 1.95E-06 |
| Gm10080 | 8.45E-11 | 0.333282 | 0.334 | 0.206 | 1.99E-06 |
| Abcc3 | 8.46E-11 | -0.85655 | 0.269 | 0.427 | 1.99E-06 |
| Slirp | 9.1E-11 | 0.494133 | 0.271 | 0.152 | 2.15E-06 |
| Tubgcp5 | 1E-10 | -0.99891 | 0.318 | 0.465 | 2.37E-06 |
| Arpc1b | 1.02E-10 | 0.559755 | 0.87 | 0.78 | 2.4E-06 |
| Pfkl | 1.03E-10 | 0.655989 | 0.401 | 0.261 | 2.43E-06 |
| Akna | 1.19E-10 | -0.9245 | 0.316 | 0.467 | 2.81E-06 |
| Ppp1r18 | 1.23E-10 | -0.71975 | 0.478 | 0.626 | 2.91E-06 |
| Agmo | 1.26E-10 | -1.41122 | 0.112 | 0.245 | 2.98E-06 |
| Cdh23 | 1.28E-10 | -1.70769 | 0.128 | 0.26 | 3.01E-06 |
| Elovl1 | 1.39E-10 | 0.567238 | 0.511 | 0.358 | 3.28E-06 |
| Sort1 | 1.4E-10 | -0.91038 | 0.363 | 0.52 | 3.29E-06 |
| Rps6 | 1.41E-10 | 0.658504 | 0.841 | 0.749 | 3.33E-06 |
| Atp5j | 1.41E-10 | 0.472938 | 0.668 | 0.507 | 3.33E-06 |
| Timm50 | 1.44E-10 | 0.713296 | 0.303 | 0.18 | 3.39E-06 |
| Gm33858 | 1.52E-10 | -1.20308 | 0.179 | 0.32 | 3.58E-06 |
| Snrpf | 1.53E-10 | 0.469013 | 0.352 | 0.217 | 3.61E-06 |
| Rps13-ps2 | 1.57E-10 | 0.598096 | 0.641 | 0.503 | 3.69E-06 |
| Tomm70a | 1.68E-10 | 0.581429 | 0.307 | 0.183 | 3.97E-06 |
| Gm11560 | 1.7E-10 | 0.3373 | 0.415 | 0.273 | 4E-06 |
| Sppl3 | 1.75E-10 | -1.11316 | 0.334 | 0.469 | 4.13E-06 |
| Eif1a | 1.83E-10 | 0.955595 | 0.296 | 0.175 | 4.32E-06 |
| Rpl29 | 1.84E-10 | 0.749732 | 0.76 | 0.643 | 4.34E-06 |
| Tmem170b | 1.97E-10 | -0.85928 | 0.244 | 0.398 | 4.64E-06 |
| Hebp1 | 2.01E-10 | 0.721397 | 0.278 | 0.161 | 4.75E-06 |
| Fnip2 | 2.04E-10 | 1.205394 | 0.249 | 0.141 | 4.82E-06 |
| Gm9800 | 2.07E-10 | 0.458836 | 0.348 | 0.219 | 4.89E-06 |
| Tjp1 | 2.25E-10 | -1.17372 | 0.202 | 0.342 | 5.31E-06 |
| 1810022K0 | 2.48E-10 | 0.559022 | 0.379 | 0.236 | 5.84E-06 |
| Bag1 | 2.48E-10 | 0.699078 | 0.446 | 0.311 | 5.85E-06 |
| Plxdc2 | 2.67E-10 | -0.36392 | 0.845 | 0.931 | 6.3E-06 |
| B4galnt1 | 2.77E-10 | 0.947248 | 0.213 | 0.113 | 6.53E-06 |
| Cyb561a3 | 2.78E-10 | -1.01597 | 0.354 | 0.487 | 6.55E-06 |
| Ifi30 | 2.91E-10 | 0.769172 | 0.61 | 0.468 | 6.86E-06 |
| Mgat4a | 3.1E-10 | -0.68366 | 0.583 | 0.699 | 7.3E-06 |
| Pitpna | 3.26E-10 | 0.833153 | 0.54 | 0.412 | 7.69E-06 |
| Echs1 | 3.36E-10 | 0.698098 | 0.35 | 0.221 | 7.92E-06 |
| Ankrd44 | 3.46E-10 | -1.42491 | 0.339 | 0.471 | 8.17E-06 |
| Usp50 | 3.54E-10 | 0.678253 | 0.267 | 0.15 | 8.35E-06 |
| Ufm1 | 3.58E-10 | 0.994437 | 0.229 | 0.127 | 8.46E-06 |
| Psmb6 | 3.71E-10 | 0.696678 | 0.587 | 0.454 | 8.75E-06 |
| mt-Rnr2 | 3.83E-10 | 0.413483 | 0.987 | 0.991 | 9.04E-06 |

| | | | | | |
|---|---|---|---|---|---|
| Nutf2 | 3.87E-10 | 0.576855 | 0.372 | 0.234 | 9.12E-06 |
| Polr2f | 3.89E-10 | 0.587513 | 0.283 | 0.164 | 9.19E-06 |
| Rps6-ps4 | 4.11E-10 | 0.529956 | 0.614 | 0.497 | 9.69E-06 |
| Gm44951 | 4.17E-10 | -0.48981 | 0.419 | 0.568 | 9.83E-06 |
| Eif3i | 4.24E-10 | 0.941662 | 0.363 | 0.238 | 1E-05 |
| Prmt1 | 4.25E-10 | 0.748553 | 0.318 | 0.201 | 1E-05 |
| Eif3e | 4.27E-10 | 0.579499 | 0.397 | 0.262 | 1.01E-05 |
| Tkt | 4.38E-10 | 0.413818 | 0.496 | 0.343 | 1.03E-05 |
| Rps3a1 | 4.68E-10 | 0.504451 | 0.901 | 0.837 | 1.1E-05 |
| Gm9843 | 4.76E-10 | 0.477464 | 0.926 | 0.834 | 1.12E-05 |
| Tspan18 | 5.21E-10 | -2.58227 | 0.092 | 0.207 | 1.23E-05 |
| Pfkfb3 | 5.39E-10 | 0.790879 | 0.357 | 0.227 | 1.27E-05 |
| Cox7a2 | 5.48E-10 | 0.502131 | 0.487 | 0.342 | 1.29E-05 |
| Rcsd1 | 5.49E-10 | -1.12568 | 0.325 | 0.46 | 1.3E-05 |
| Gm10819 | 5.53E-10 | -0.49915 | 0.262 | 0.15 | 1.3E-05 |
| Mapkapk2 | 5.71E-10 | 0.587939 | 0.357 | 0.228 | 1.35E-05 |
| Cd302 | 5.73E-10 | -0.88933 | 0.336 | 0.469 | 1.35E-05 |
| Morf4l2 | 5.82E-10 | 0.701518 | 0.43 | 0.291 | 1.37E-05 |
| Hpf1 | 5.83E-10 | 0.68687 | 0.303 | 0.183 | 1.38E-05 |
| Edf1 | 5.99E-10 | 0.632524 | 0.578 | 0.436 | 1.41E-05 |
| Tent5c | 6.6E-10 | 0.783419 | 0.511 | 0.386 | 1.56E-05 |
| Pik3cg | 6.75E-10 | -1.21001 | 0.296 | 0.432 | 1.59E-05 |
| Fcer1g | 7.1E-10 | 0.388975 | 0.998 | 0.993 | 1.67E-05 |
| Gm6170 | 7.29E-10 | 0.367624 | 0.484 | 0.356 | 1.72E-05 |
| Zfas1 | 7.43E-10 | 1.192628 | 0.226 | 0.127 | 1.75E-05 |
| Dennd1c | 7.83E-10 | -1.68774 | 0.157 | 0.289 | 1.85E-05 |
| Gpr84 | 7.94E-10 | 1.224755 | 0.567 | 0.457 | 1.87E-05 |
| Slc46a1 | 8.91E-10 | -1.00941 | 0.325 | 0.455 | 2.1E-05 |
| Wdr43 | 8.93E-10 | 0.722814 | 0.291 | 0.176 | 2.11E-05 |
| Rpl8 | 9.23E-10 | 0.6504 | 0.8 | 0.708 | 2.18E-05 |
| Coro1c | 9.3E-10 | 0.442529 | 0.363 | 0.227 | 2.19E-05 |
| Sdf4 | 9.33E-10 | 0.576715 | 0.7 | 0.568 | 2.2E-05 |
| Gm14494 | 9.44E-10 | 0.52644 | 0.296 | 0.179 | 2.23E-05 |
| Twf2 | 9.8E-10 | -0.81297 | 0.509 | 0.619 | 2.31E-05 |
| Ndufa4 | 9.85E-10 | 0.563357 | 0.561 | 0.411 | 2.32E-05 |
| Rbm39 | 1.01E-09 | -0.48576 | 0.823 | 0.9 | 2.39E-05 |
| Actr3 | 1.02E-09 | 0.591593 | 0.661 | 0.536 | 2.4E-05 |
| Rps25-ps1 | 1.02E-09 | 0.332147 | 0.406 | 0.277 | 2.41E-05 |
| Frmd4b | 1.04E-09 | -0.82987 | 0.397 | 0.535 | 2.46E-05 |
| Usp2 | 1.08E-09 | -0.43774 | 0.556 | 0.703 | 2.55E-05 |
| Plbd2 | 1.1E-09 | 0.510321 | 0.504 | 0.357 | 2.59E-05 |
| Elp5 | 1.1E-09 | 0.782324 | 0.22 | 0.119 | 2.6E-05 |
| Casp8 | 1.12E-09 | -0.91591 | 0.336 | 0.474 | 2.65E-05 |
| Cox6b1 | 1.19E-09 | 0.522557 | 0.641 | 0.497 | 2.8E-05 |
| Hist3h2a | 1.23E-09 | -1.04448 | 0.157 | 0.292 | 2.89E-05 |
| Rpl22l1 | 1.24E-09 | 0.640559 | 0.361 | 0.239 | 2.92E-05 |
| Ptpn18 | 1.27E-09 | -0.49989 | 0.581 | 0.726 | 3E-05 |

| | | | | | |
|---|---|---|---|---|---|
| Plxnb2 | 1.29E-09 | -0.57289 | 0.614 | 0.735 | 3.04E-05 |
| mt-Co1 | 1.29E-09 | 0.580127 | 0.987 | 0.985 | 3.05E-05 |
| Slc25a45 | 1.34E-09 | -1.06329 | 0.204 | 0.338 | 3.17E-05 |
| Rgmb | 1.36E-09 | -0.77264 | 0.323 | 0.461 | 3.22E-05 |
| Uqcrh | 1.38E-09 | 0.469652 | 0.538 | 0.391 | 3.26E-05 |
| Gm5805 | 1.42E-09 | 0.46974 | 0.608 | 0.474 | 3.36E-05 |
| Hspd1-ps3 | 1.45E-09 | 0.458944 | 0.213 | 0.118 | 3.42E-05 |
| Gm28417 | 1.56E-09 | -1.86861 | 0.096 | 0.213 | 3.69E-05 |
| Nop10 | 1.62E-09 | 0.587866 | 0.339 | 0.212 | 3.82E-05 |
| Ube2l3 | 1.62E-09 | 0.415279 | 0.561 | 0.403 | 3.83E-05 |
| Psmb2 | 1.63E-09 | 0.598015 | 0.489 | 0.347 | 3.85E-05 |
| Arpc2 | 1.63E-09 | 0.549598 | 0.91 | 0.839 | 3.85E-05 |
| Mpi | 1.72E-09 | -0.98283 | 0.166 | 0.298 | 4.05E-05 |
| Rhog | 1.89E-09 | -0.51479 | 0.709 | 0.789 | 4.47E-05 |
| Pip4p1 | 1.91E-09 | -0.66036 | 0.61 | 0.716 | 4.49E-05 |
| Rbm3 | 1.91E-09 | 0.607552 | 0.426 | 0.291 | 4.5E-05 |
| Gtf2f1 | 1.95E-09 | 0.509299 | 0.24 | 0.135 | 4.59E-05 |
| Mfap3 | 1.98E-09 | -0.83688 | 0.489 | 0.615 | 4.66E-05 |
| Picalm | 2.1E-09 | -0.6384 | 0.626 | 0.74 | 4.94E-05 |
| Eif5b | 2.12E-09 | 0.616091 | 0.381 | 0.254 | 5.01E-05 |
| Sult1a1 | 2.13E-09 | -1.22925 | 0.146 | 0.272 | 5.02E-05 |
| Ybx3 | 2.16E-09 | 1.102196 | 0.202 | 0.11 | 5.1E-05 |
| Mydgf | 2.29E-09 | 0.4063 | 0.504 | 0.36 | 5.41E-05 |
| Atp6v0e | 2.37E-09 | 0.599414 | 0.612 | 0.466 | 5.59E-05 |
| Eif2s1 | 2.39E-09 | 0.764076 | 0.368 | 0.243 | 5.63E-05 |
| Sod2 | 2.5E-09 | 0.670143 | 0.3 | 0.185 | 5.9E-05 |
| Gm15427 | 2.57E-09 | 0.462459 | 0.953 | 0.942 | 6.06E-05 |
| Glb1 | 2.74E-09 | 0.828925 | 0.379 | 0.256 | 6.47E-05 |
| Rpl17-ps3 | 2.78E-09 | 0.497607 | 0.504 | 0.375 | 6.55E-05 |
| Rapgef5 | 2.79E-09 | -0.67685 | 0.235 | 0.374 | 6.58E-05 |
| Bin1 | 2.87E-09 | -0.48882 | 0.881 | 0.913 | 6.78E-05 |
| Cd82 | 2.88E-09 | -0.66063 | 0.457 | 0.595 | 6.8E-05 |
| Thrsp | 2.9E-09 | -0.75171 | 0.179 | 0.313 | 6.85E-05 |
| Slc12a9 | 3E-09 | -0.5974 | 0.314 | 0.463 | 7.07E-05 |
| Nav2 | 3.05E-09 | -1.14113 | 0.283 | 0.415 | 7.2E-05 |
| Nsa2 | 3.07E-09 | 0.410521 | 0.395 | 0.264 | 7.25E-05 |
| Rasgef1b | 3.27E-09 | 0.542539 | 0.2 | 0.106 | 7.71E-05 |
| Ddx21 | 3.48E-09 | 1.069169 | 0.269 | 0.161 | 8.2E-05 |
| Snrpa1 | 3.55E-09 | 0.750245 | 0.222 | 0.125 | 8.38E-05 |
| Oxct1 | 3.63E-09 | 0.579406 | 0.63 | 0.502 | 8.57E-05 |
| Tcp1 | 3.78E-09 | 0.394449 | 0.408 | 0.27 | 8.92E-05 |
| Clasp2 | 3.88E-09 | -0.32872 | 0.655 | 0.796 | 9.16E-05 |
| Rrp1 | 3.94E-09 | 0.492733 | 0.493 | 0.363 | 9.29E-05 |
| Cask | 4.04E-09 | -0.81779 | 0.267 | 0.405 | 9.52E-05 |
| Plekha1 | 4.05E-09 | 0.643424 | 0.298 | 0.188 | 9.55E-05 |
| Rpl34-ps1 | 4.13E-09 | 0.391973 | 0.61 | 0.483 | 9.74E-05 |
| Taf10 | 4.15E-09 | 0.442465 | 0.516 | 0.387 | 9.79E-05 |

| | | | | | |
|---|---|---|---|---|---|
| Csf3r | 4.2E-09 | -0.50942 | 0.733 | 0.854 | 9.9E-05 |
| Krtcap2 | 4.89E-09 | 0.517318 | 0.585 | 0.452 | 0.000115 |
| Vps29 | 5.07E-09 | 0.621131 | 0.469 | 0.333 | 0.00012 |
| Garnl3 | 5.08E-09 | -1.14482 | 0.108 | 0.225 | 0.00012 |
| Rps27 | 5.11E-09 | 0.446082 | 0.984 | 0.95 | 0.000121 |
| Hnrnpm | 5.2E-09 | 0.461442 | 0.531 | 0.395 | 0.000123 |
| Eif1ax | 5.56E-09 | 0.860766 | 0.202 | 0.111 | 0.000131 |
| Nr3c1 | 5.8E-09 | -0.67374 | 0.392 | 0.528 | 0.000137 |
| Gpr146 | 5.9E-09 | -0.97978 | 0.28 | 0.415 | 0.000139 |
| Gm6180 | 6E-09 | 0.506064 | 0.507 | 0.397 | 0.000141 |
| Il11ra1 | 6.13E-09 | 0.833586 | 0.296 | 0.186 | 0.000145 |
| Vmp1 | 6.25E-09 | 0.367473 | 0.383 | 0.249 | 0.000148 |
| Rpl13a | 6.67E-09 | 0.626508 | 0.879 | 0.824 | 0.000157 |
| Ndufa7 | 6.72E-09 | 0.617928 | 0.531 | 0.4 | 0.000158 |
| Asna1 | 6.79E-09 | 0.714886 | 0.249 | 0.146 | 0.00016 |
| Mfsd1 | 7.12E-09 | 0.60151 | 0.563 | 0.426 | 0.000168 |
| Hacd2 | 7.28E-09 | 0.714054 | 0.354 | 0.235 | 0.000172 |
| Gm28661 | 7.43E-09 | 0.512608 | 0.796 | 0.731 | 0.000175 |
| Sash3 | 7.64E-09 | -0.74213 | 0.274 | 0.412 | 0.00018 |
| Gm14760 | 7.75E-09 | 0.251425 | 0.233 | 0.136 | 0.000183 |
| Hnrnpab | 7.89E-09 | 0.518107 | 0.679 | 0.55 | 0.000186 |
| Rps9 | 7.92E-09 | 0.360117 | 0.982 | 0.95 | 0.000187 |
| Gm12070 | 7.92E-09 | 0.519121 | 0.217 | 0.126 | 0.000187 |
| Sqor | 8.23E-09 | -1.21774 | 0.197 | 0.319 | 0.000194 |
| Ndufc1 | 8.23E-09 | 0.270942 | 0.318 | 0.196 | 0.000194 |
| Prelid1 | 8.34E-09 | 0.857973 | 0.549 | 0.449 | 0.000197 |
| Tor3a | 8.4E-09 | 0.75747 | 0.352 | 0.237 | 0.000198 |
| March1 | 8.43E-09 | -1.34827 | 0.204 | 0.334 | 0.000199 |
| Sirt2 | 8.58E-09 | 0.737408 | 0.444 | 0.325 | 0.000203 |
| Prkca | 8.96E-09 | -0.76501 | 0.101 | 0.216 | 0.000211 |
| Ptpn1 | 9.19E-09 | -0.88153 | 0.554 | 0.63 | 0.000217 |
| Mfng | 9.78E-09 | -0.74481 | 0.287 | 0.422 | 0.000231 |
| Gm6472 | 1E-08 | 0.495985 | 0.578 | 0.445 | 0.000236 |
| Phgdh | 1.01E-08 | 0.693303 | 0.507 | 0.38 | 0.000237 |
| Ggh | 1.03E-08 | 0.731097 | 0.309 | 0.199 | 0.000242 |
| Psme2 | 1.07E-08 | 0.783942 | 0.585 | 0.491 | 0.000251 |
| Slc29a3 | 1.13E-08 | -0.57359 | 0.666 | 0.767 | 0.000267 |
| Rgs19 | 1.14E-08 | -0.77648 | 0.388 | 0.506 | 0.00027 |
| Zfp217 | 1.18E-08 | -1.06472 | 0.186 | 0.315 | 0.000279 |
| Ggta1 | 1.2E-08 | -1.86455 | 0.094 | 0.206 | 0.000283 |
| Etf1 | 1.21E-08 | 0.875552 | 0.276 | 0.171 | 0.000286 |
| Lifr | 1.24E-08 | -1.4147 | 0.195 | 0.326 | 0.000292 |
| Tcf4 | 1.24E-08 | -0.76758 | 0.52 | 0.634 | 0.000292 |
| Arid3b | 1.24E-08 | -1.63405 | 0.143 | 0.264 | 0.000293 |
| Ndufa13 | 1.31E-08 | 0.461012 | 0.614 | 0.471 | 0.000309 |
| Iah1 | 1.33E-08 | 1.011601 | 0.224 | 0.129 | 0.000314 |
| Srp14 | 1.35E-08 | 0.546293 | 0.457 | 0.326 | 0.000318 |

| | | | | | |
|---|---|---|---|---|---|
| Plod1 | 1.39E-08 | -0.72572 | 0.529 | 0.628 | 0.000328 |
| Notch2 | 1.49E-08 | -0.59629 | 0.574 | 0.697 | 0.000351 |
| Gm10288 | 1.53E-08 | 0.366249 | 0.552 | 0.419 | 0.00036 |
| Tnfsf13b | 1.53E-08 | 0.571707 | 0.267 | 0.162 | 0.000361 |
| mt-Atp6 | 1.54E-08 | 0.685231 | 0.85 | 0.825 | 0.000364 |
| Asph | 1.59E-08 | 0.504616 | 0.767 | 0.642 | 0.000375 |
| Pid1 | 1.64E-08 | -1.0566 | 0.197 | 0.319 | 0.000387 |
| H2afz | 1.64E-08 | 0.583126 | 0.614 | 0.501 | 0.000387 |
| Sgpl1 | 1.65E-08 | 0.620166 | 0.583 | 0.447 | 0.00039 |
| Cd180 | 1.66E-08 | 0.607657 | 0.552 | 0.439 | 0.000391 |
| Gm9493 | 1.68E-08 | 0.291327 | 0.35 | 0.24 | 0.000397 |
| Tomm7 | 1.78E-08 | 0.548824 | 0.43 | 0.3 | 0.00042 |
| Pa2g4 | 1.83E-08 | 0.649406 | 0.327 | 0.212 | 0.000432 |
| Rpl27-ps3 | 1.93E-08 | 0.417063 | 0.478 | 0.354 | 0.000455 |
| Phb | 1.95E-08 | 0.71291 | 0.377 | 0.26 | 0.00046 |
| Wdfy2 | 2.05E-08 | -0.8615 | 0.166 | 0.288 | 0.000484 |
| Rhoh | 2.16E-08 | -0.79201 | 0.359 | 0.48 | 0.00051 |
| Dnm2 | 2.17E-08 | -0.64034 | 0.561 | 0.687 | 0.000511 |
| Galc | 2.18E-08 | 0.640656 | 0.202 | 0.115 | 0.000514 |
| Rnf149 | 2.18E-08 | 0.529657 | 0.312 | 0.197 | 0.000514 |
| Gm10925 | 2.28E-08 | 0.61648 | 0.865 | 0.823 | 0.000537 |
| Rps29 | 2.29E-08 | 0.358528 | 0.987 | 0.972 | 0.00054 |
| Mlph | 2.32E-08 | -1.05163 | 0.283 | 0.411 | 0.000546 |
| Wdr83os | 2.37E-08 | 0.559165 | 0.357 | 0.238 | 0.000559 |
| Filip1l | 2.45E-08 | -1.03522 | 0.175 | 0.295 | 0.000577 |
| Nedd8 | 2.58E-08 | 0.676436 | 0.525 | 0.399 | 0.00061 |
| Gfm2 | 2.77E-08 | -0.52546 | 0.939 | 0.986 | 0.000654 |
| Tspan13 | 2.8E-08 | -0.85786 | 0.428 | 0.538 | 0.00066 |
| Btf3 | 2.82E-08 | 0.628959 | 0.601 | 0.476 | 0.000664 |
| Adam10 | 2.86E-08 | -0.47299 | 0.695 | 0.796 | 0.000676 |
| Syncrip | 2.87E-08 | 0.835191 | 0.296 | 0.19 | 0.000678 |
| Rnaset2b | 2.99E-08 | -0.36371 | 0.913 | 0.951 | 0.000706 |
| Gm11518 | 3E-08 | 0.399468 | 0.623 | 0.478 | 0.000709 |
| Uqcr10 | 3.04E-08 | 0.377105 | 0.442 | 0.311 | 0.000716 |
| Hypk | 3.1E-08 | 0.634741 | 0.265 | 0.161 | 0.000732 |
| Phb2 | 3.12E-08 | 0.623549 | 0.401 | 0.279 | 0.000735 |
| Cib1 | 3.14E-08 | -1.18243 | 0.175 | 0.292 | 0.000741 |
| Cct3 | 3.38E-08 | 0.401632 | 0.43 | 0.302 | 0.000798 |
| Zdhhc14 | 3.49E-08 | -1.33128 | 0.096 | 0.201 | 0.000822 |
| Gm6204 | 3.51E-08 | 0.443094 | 0.693 | 0.582 | 0.000829 |
| Tifa | 3.61E-08 | -1.10656 | 0.28 | 0.396 | 0.000851 |
| H2afj | 3.76E-08 | 0.477994 | 0.426 | 0.294 | 0.000887 |
| Orai1 | 3.78E-08 | -0.59112 | 0.518 | 0.639 | 0.000891 |
| Tlr12 | 3.88E-08 | -1.11978 | 0.164 | 0.279 | 0.000915 |
| St13 | 3.93E-08 | 0.301448 | 0.78 | 0.635 | 0.000928 |
| Igbp1 | 4.04E-08 | 0.737636 | 0.247 | 0.15 | 0.000952 |
| Calm3 | 4.1E-08 | 0.508071 | 0.359 | 0.241 | 0.000967 |

| | | | | | |
|---|---|---|---|---|---|
| Themis2 | 4.25E-08 | -1.4869 | 0.238 | 0.356 | 0.001002 |
| Rps12-ps9 | 4.43E-08 | 0.385804 | 0.54 | 0.423 | 0.001045 |
| Rnf128 | 4.44E-08 | 1.268462 | 0.226 | 0.137 | 0.001047 |
| Eif3c | 4.58E-08 | 0.341975 | 0.572 | 0.439 | 0.001081 |
| Erf | 4.58E-08 | -0.49171 | 0.298 | 0.417 | 0.001081 |
| Kif21b | 4.59E-08 | -0.80905 | 0.26 | 0.392 | 0.001082 |
| Cnot8 | 4.69E-08 | -0.82534 | 0.372 | 0.496 | 0.001106 |
| Minos1 | 4.72E-08 | 0.495439 | 0.516 | 0.392 | 0.001114 |
| Cct5 | 4.73E-08 | 0.549365 | 0.491 | 0.368 | 0.001116 |
| Rps27a-ps2 | 4.76E-08 | 0.601012 | 0.397 | 0.278 | 0.001122 |
| Rpl19-ps1 | 4.81E-08 | 0.304849 | 0.321 | 0.212 | 0.001135 |
| Gm4332 | 4.89E-08 | 0.295981 | 0.318 | 0.212 | 0.001153 |
| Lyst | 4.9E-08 | 1.023911 | 0.217 | 0.128 | 0.001155 |
| Mrpl12 | 4.99E-08 | 0.788852 | 0.28 | 0.175 | 0.001177 |
| Got2 | 5.38E-08 | 0.370225 | 0.262 | 0.161 | 0.00127 |
| P4hb | 5.45E-08 | 0.469942 | 0.901 | 0.829 | 0.001284 |
| Ndufa12 | 5.72E-08 | 0.626147 | 0.341 | 0.227 | 0.00135 |
| Ceacam1 | 5.76E-08 | -1.47287 | 0.126 | 0.234 | 0.001359 |
| Rars | 5.81E-08 | 0.478544 | 0.213 | 0.123 | 0.001371 |
| Slc7a8 | 5.83E-08 | -0.36505 | 0.608 | 0.723 | 0.001374 |
| Gm11478 | 5.98E-08 | 0.805891 | 0.742 | 0.663 | 0.001411 |
| Mrpl39 | 6.01E-08 | 0.778975 | 0.222 | 0.13 | 0.001419 |
| Naca | 6.03E-08 | 0.488485 | 0.731 | 0.599 | 0.001423 |
| Zfp706 | 6.14E-08 | -0.71557 | 0.574 | 0.661 | 0.001448 |
| Gtf2h5 | 6.37E-08 | 0.60191 | 0.217 | 0.125 | 0.001502 |
| Prkcsh | 6.49E-08 | 0.84069 | 0.374 | 0.268 | 0.001532 |
| Tspan14 | 6.53E-08 | -0.81229 | 0.457 | 0.571 | 0.001541 |
| Hmgn1 | 6.66E-08 | 0.503622 | 0.466 | 0.345 | 0.001571 |
| Abracl | 6.97E-08 | 0.414004 | 0.305 | 0.192 | 0.001644 |
| Capns1 | 7.02E-08 | 0.766375 | 0.419 | 0.307 | 0.001655 |
| Rnf13 | 7.04E-08 | -0.46247 | 0.697 | 0.776 | 0.00166 |
| Rpl36a-ps2 | 7.11E-08 | 0.362672 | 0.336 | 0.23 | 0.001677 |
| Bmyc | 7.13E-08 | -0.62466 | 0.374 | 0.502 | 0.001682 |
| Rbx1 | 7.15E-08 | 0.37597 | 0.576 | 0.45 | 0.001686 |
| Ndufb7 | 7.69E-08 | 0.376007 | 0.543 | 0.401 | 0.001813 |
| Sod1 | 7.91E-08 | 0.545368 | 0.731 | 0.612 | 0.001866 |
| mt-Co3 | 7.98E-08 | 0.782468 | 0.91 | 0.902 | 0.001882 |
| Numa1 | 8.41E-08 | -1.06537 | 0.294 | 0.401 | 0.001983 |
| Psmd1 | 8.65E-08 | 0.667268 | 0.428 | 0.307 | 0.002041 |
| Tns1 | 8.75E-08 | 0.82114 | 0.341 | 0.231 | 0.002065 |
| Arglu1 | 8.86E-08 | -0.64477 | 0.756 | 0.862 | 0.002091 |
| Impdh2 | 8.88E-08 | 0.785506 | 0.231 | 0.139 | 0.002095 |
| mt-Rnr1 | 8.96E-08 | 0.506217 | 0.895 | 0.888 | 0.002114 |
| Atp5d | 9.02E-08 | 0.455242 | 0.567 | 0.429 | 0.002127 |
| Arhgap12 | 9.32E-08 | -1.00202 | 0.148 | 0.265 | 0.002197 |
| Mvp | 9.81E-08 | 0.458319 | 0.274 | 0.172 | 0.002315 |
| Fez2 | 9.98E-08 | -0.73009 | 0.379 | 0.501 | 0.002355 |

| | | | | | |
|---|---|---|---|---|---|
| Snx3 | 1E-07 | 0.480976 | 0.578 | 0.442 | 0.002366 |
| Dbi | 1.02E-07 | 0.815745 | 0.341 | 0.234 | 0.002398 |
| Ogt | 1.06E-07 | -0.61733 | 0.439 | 0.567 | 0.002491 |
| Sat1 | 1.07E-07 | 0.436106 | 0.899 | 0.847 | 0.002528 |
| Gng5-ps | 1.11E-07 | 0.570966 | 0.374 | 0.263 | 0.002627 |
| Dock8 | 1.15E-07 | -0.65102 | 0.419 | 0.53 | 0.002724 |
| Nckap5l | 1.18E-07 | -1.18463 | 0.13 | 0.24 | 0.002793 |
| Pfdn5 | 1.2E-07 | 0.515678 | 0.709 | 0.592 | 0.002831 |
| Cryba4 | 1.21E-07 | -1.0246 | 0.161 | 0.277 | 0.002843 |
| Pgd | 1.27E-07 | 0.626484 | 0.46 | 0.338 | 0.002985 |
| Ddx5 | 1.34E-07 | -0.58666 | 0.729 | 0.789 | 0.003172 |
| Pwwp2a | 1.38E-07 | -0.92194 | 0.318 | 0.442 | 0.003264 |
| Txndc5 | 1.38E-07 | 0.834365 | 0.314 | 0.216 | 0.003265 |
| Ubl5 | 1.39E-07 | 0.466645 | 0.54 | 0.418 | 0.003287 |
| Rps21 | 1.44E-07 | 0.460005 | 0.834 | 0.762 | 0.003407 |
| Sec61b | 1.46E-07 | 0.452641 | 0.424 | 0.299 | 0.003435 |
| Nop58 | 1.47E-07 | 0.867053 | 0.267 | 0.172 | 0.003456 |
| Gm14303 | 1.58E-07 | 0.540929 | 0.825 | 0.753 | 0.003739 |
| Hjurp | 1.59E-07 | -1.4472 | 0.121 | 0.222 | 0.003756 |
| Kars | 1.6E-07 | 0.472793 | 0.314 | 0.202 | 0.003775 |
| Zbtb18 | 1.62E-07 | -1.32277 | 0.126 | 0.231 | 0.003815 |
| Fis1 | 1.62E-07 | 0.663551 | 0.484 | 0.361 | 0.003821 |
| Gm3531 | 1.62E-07 | 0.320994 | 0.39 | 0.271 | 0.003822 |
| Slc9a9 | 1.64E-07 | -0.94995 | 0.258 | 0.373 | 0.003865 |
| Gm16118 | 1.65E-07 | -0.93647 | 0.123 | 0.23 | 0.003893 |
| Gltp | 1.66E-07 | 0.490706 | 0.498 | 0.375 | 0.003913 |
| Plxna4 | 1.72E-07 | -1.12783 | 0.121 | 0.227 | 0.004048 |
| Gm3379 | 1.75E-07 | 0.477583 | 0.661 | 0.562 | 0.00413 |
| Rpl30-ps10 | 1.75E-07 | 0.299814 | 0.305 | 0.208 | 0.004135 |
| Ndufb6 | 1.77E-07 | 0.706651 | 0.291 | 0.187 | 0.004179 |
| Gm9320 | 1.79E-07 | 0.511434 | 0.251 | 0.156 | 0.004228 |
| Rhobtb1 | 1.85E-07 | -1.65888 | 0.146 | 0.253 | 0.004354 |
| Itga6 | 1.85E-07 | -0.72168 | 0.48 | 0.591 | 0.004375 |
| Tma7 | 1.87E-07 | 0.400163 | 0.529 | 0.4 | 0.004415 |
| Ppp1r9a | 1.89E-07 | -0.73113 | 0.26 | 0.382 | 0.00445 |
| Gm10053 | 1.91E-07 | 0.328761 | 0.262 | 0.168 | 0.004509 |
| Jam2 | 1.98E-07 | -1.10529 | 0.211 | 0.324 | 0.00467 |
| Nutf2-ps1 | 2.01E-07 | 0.46037 | 0.253 | 0.158 | 0.004738 |
| Rps10-ps1 | 2.02E-07 | 0.340983 | 0.614 | 0.482 | 0.00476 |
| Myo1f | 2.03E-07 | 0.575129 | 0.643 | 0.535 | 0.004785 |
| Retreg2 | 2.03E-07 | 0.515897 | 0.386 | 0.268 | 0.004786 |
| Agpat5 | 2.04E-07 | 1.252642 | 0.226 | 0.14 | 0.004802 |
| Arpc3 | 2.04E-07 | 0.435782 | 0.722 | 0.617 | 0.004806 |
| Mrpl54 | 2.05E-07 | 0.649065 | 0.231 | 0.14 | 0.004834 |
| Gm10036 | 2.07E-07 | 0.253095 | 0.428 | 0.311 | 0.004875 |
| Chd4 | 2.19E-07 | 0.381966 | 0.439 | 0.325 | 0.005168 |
| Hspa8 | 2.2E-07 | 0.409683 | 0.948 | 0.945 | 0.005187 |

| | | | | | |
|---|---|---|---|---|---|
| Mroh1 | 2.31E-07 | 0.901342 | 0.226 | 0.139 | 0.005441 |
| 9930111J2 | 2.39E-07 | -1.27339 | 0.209 | 0.325 | 0.005631 |
| Fermt3 | 2.41E-07 | -0.52217 | 0.691 | 0.773 | 0.005689 |
| Bcl9l | 2.44E-07 | -0.91001 | 0.135 | 0.243 | 0.005753 |
| Gm6136 | 2.45E-07 | 0.500831 | 0.632 | 0.525 | 0.005782 |
| Cmtm3 | 2.51E-07 | 0.477626 | 0.502 | 0.383 | 0.005919 |
| Nop16 | 2.52E-07 | 0.791319 | 0.226 | 0.136 | 0.00595 |
| Dad1 | 2.54E-07 | 0.48199 | 0.695 | 0.592 | 0.005986 |
| Gm7901 | 2.6E-07 | 0.51991 | 0.379 | 0.267 | 0.00614 |
| Rpl13-ps6 | 2.63E-07 | 0.416187 | 0.603 | 0.509 | 0.006209 |
| Oaz1 | 2.66E-07 | 0.334373 | 0.886 | 0.789 | 0.006277 |
| Gcnt1 | 2.73E-07 | -0.641 | 0.392 | 0.524 | 0.006437 |
| Slc1a3 | 2.75E-07 | -0.77267 | 0.437 | 0.561 | 0.006498 |
| Rpl17-ps8 | 2.76E-07 | 0.363471 | 0.251 | 0.158 | 0.006503 |
| Slc46a3 | 2.77E-07 | -1.11524 | 0.211 | 0.324 | 0.006541 |
| Ubxn1 | 2.78E-07 | 0.679027 | 0.372 | 0.267 | 0.00655 |
| Chst7 | 2.87E-07 | -0.94539 | 0.175 | 0.286 | 0.006768 |
| Ccl12 | 2.9E-07 | 2.544734 | 0.267 | 0.184 | 0.00683 |
| Smdt1 | 2.95E-07 | 0.389879 | 0.415 | 0.292 | 0.006954 |
| Pnisr | 3.02E-07 | -0.64697 | 0.516 | 0.612 | 0.007125 |
| Fmnl3 | 3.12E-07 | -0.52189 | 0.413 | 0.531 | 0.007361 |
| BC035044 | 3.21E-07 | -1.06352 | 0.15 | 0.258 | 0.007564 |
| Ypel5 | 3.28E-07 | -0.87503 | 0.233 | 0.345 | 0.007737 |
| Vav2 | 3.28E-07 | -1.52949 | 0.128 | 0.231 | 0.007742 |
| Snrpd2 | 3.37E-07 | 0.809836 | 0.294 | 0.191 | 0.007945 |
| Trem2 | 3.41E-07 | 0.347511 | 0.991 | 0.988 | 0.008044 |
| Tomm40 | 3.45E-07 | 0.719218 | 0.274 | 0.178 | 0.008129 |
| A830008E2 | 3.45E-07 | -1.0621 | 0.184 | 0.292 | 0.008133 |
| Gm5835 | 3.45E-07 | 0.461816 | 0.462 | 0.352 | 0.008138 |
| Eif3l | 3.47E-07 | 0.579139 | 0.401 | 0.289 | 0.008182 |
| Fam167b | 3.72E-07 | -1.17912 | 0.114 | 0.214 | 0.008773 |
| Ndufa11 | 3.79E-07 | 0.360007 | 0.374 | 0.255 | 0.008938 |
| Cep152 | 4.08E-07 | -1.61527 | 0.123 | 0.221 | 0.009629 |
| G530011O( | 4.1E-07 | -0.76756 | 0.269 | 0.385 | 0.009661 |
| Gm12428 | 4.21E-07 | -0.3648 | 0.135 | 0.239 | 0.009936 |
| Cct8 | 4.21E-07 | 0.445926 | 0.507 | 0.388 | 0.009942 |
| D8Ertd738( | 4.22E-07 | 0.422971 | 0.498 | 0.371 | 0.009957 |
| Gas6 | 4.25E-07 | -0.61119 | 0.365 | 0.482 | 0.010017 |
| Notch1 | 4.28E-07 | -1.01299 | 0.226 | 0.334 | 0.010096 |
| Sqstm1 | 4.4E-07 | 0.659961 | 0.711 | 0.606 | 0.01037 |
| E230029C( | 4.54E-07 | -0.37251 | 0.549 | 0.666 | 0.010708 |
| Cd34 | 4.58E-07 | 0.43953 | 0.628 | 0.51 | 0.010803 |
| Erh | 4.59E-07 | 0.422243 | 0.473 | 0.343 | 0.010838 |
| Galnt12 | 4.72E-07 | -1.09831 | 0.105 | 0.204 | 0.01114 |
| Gm9234 | 4.74E-07 | 0.437827 | 0.552 | 0.462 | 0.011174 |
| Pkig | 4.76E-07 | -0.84503 | 0.309 | 0.412 | 0.011219 |
| Map3k3 | 4.81E-07 | -1.18955 | 0.283 | 0.384 | 0.01134 |

| | | | | | |
|---|---|---|---|---|---|
| Gm10163 | 4.88E-07 | -0.33903 | 0.538 | 0.421 | 0.0115 |
| Tsc22d3 | 4.91E-07 | -1.58611 | 0.166 | 0.263 | 0.011579 |
| Ppfia4 | 4.92E-07 | -0.5033 | 0.574 | 0.678 | 0.011601 |
| Usp21 | 5.32E-07 | -1.12727 | 0.256 | 0.362 | 0.012559 |
| Mrps24 | 5.65E-07 | 0.772317 | 0.274 | 0.183 | 0.013318 |
| Pdcd5 | 6E-07 | 0.524506 | 0.37 | 0.26 | 0.014163 |
| Cct2 | 6.04E-07 | 0.327462 | 0.41 | 0.296 | 0.014256 |
| Bid | 6.08E-07 | -1.03649 | 0.155 | 0.259 | 0.014343 |
| mt-Nd3 | 6.09E-07 | 0.462797 | 0.664 | 0.537 | 0.014365 |
| Llph | 6.09E-07 | 0.566377 | 0.296 | 0.196 | 0.014368 |
| Rps27rt | 6.1E-07 | 0.401698 | 0.767 | 0.687 | 0.014397 |
| Stambpl1 | 6.21E-07 | -0.97966 | 0.222 | 0.33 | 0.014645 |
| Atxn10 | 6.27E-07 | 0.537736 | 0.466 | 0.345 | 0.014801 |
| Mib2 | 6.3E-07 | -1.15277 | 0.112 | 0.21 | 0.014861 |
| mt-Cytb | 6.35E-07 | 0.555264 | 0.966 | 0.967 | 0.014968 |
| Fbrsl1 | 6.36E-07 | -0.7205 | 0.3 | 0.414 | 0.014994 |
| Igsf8 | 6.55E-07 | 0.495053 | 0.323 | 0.219 | 0.015452 |
| Arrb2 | 6.64E-07 | -0.34012 | 0.621 | 0.72 | 0.015664 |
| Ramp1 | 6.67E-07 | 0.894961 | 0.271 | 0.182 | 0.015741 |
| Pou2f2 | 6.75E-07 | -0.42306 | 0.639 | 0.751 | 0.015926 |
| Tcof1 | 6.79E-07 | 0.546091 | 0.278 | 0.183 | 0.016017 |
| Rilpl1 | 6.86E-07 | -0.34046 | 0.209 | 0.328 | 0.016192 |
| Itpr2 | 6.98E-07 | -0.8408 | 0.41 | 0.517 | 0.016459 |
| Snu13 | 7.11E-07 | 0.509919 | 0.345 | 0.241 | 0.016768 |
| Emc1 | 7.13E-07 | 0.713407 | 0.204 | 0.123 | 0.016814 |
| Noc2l | 7.15E-07 | 0.627466 | 0.291 | 0.193 | 0.016873 |
| Rere | 7.18E-07 | -0.36825 | 0.271 | 0.391 | 0.016947 |
| Eif3b | 7.25E-07 | 0.383382 | 0.377 | 0.266 | 0.017099 |
| Gm45640 | 7.31E-07 | -0.53183 | 0.323 | 0.451 | 0.017244 |
| Mbnl1 | 7.33E-07 | -0.34351 | 0.778 | 0.842 | 0.017289 |
| Hspa4 | 7.45E-07 | 0.463858 | 0.484 | 0.367 | 0.017582 |
| Zscan26 | 7.72E-07 | -0.51684 | 0.623 | 0.727 | 0.018211 |
| App | 7.78E-07 | 0.470239 | 0.552 | 0.448 | 0.018364 |
| Apobec1 | 7.8E-07 | 0.811344 | 0.343 | 0.245 | 0.018404 |
| Sigmar1 | 7.91E-07 | 0.653513 | 0.298 | 0.201 | 0.018667 |
| Slc35b1 | 8.1E-07 | 0.598338 | 0.374 | 0.27 | 0.019115 |
| Tuba1c | 8.11E-07 | 1.695054 | 0.457 | 0.4 | 0.019127 |
| B4galt4 | 8.32E-07 | -0.75169 | 0.197 | 0.307 | 0.019621 |
| Rapgef1 | 8.37E-07 | -0.90625 | 0.251 | 0.362 | 0.019739 |
| Drap1 | 8.41E-07 | 0.503056 | 0.433 | 0.316 | 0.01983 |
| Gm43028 | 8.44E-07 | 0.382415 | 0.408 | 0.307 | 0.0199 |
| Rpl17-ps10 | 8.77E-07 | 0.362798 | 0.296 | 0.199 | 0.020682 |
| Tln2 | 9.09E-07 | -0.8192 | 0.195 | 0.301 | 0.02144 |
| Camk2n1 | 9.18E-07 | -0.8522 | 0.191 | 0.294 | 0.021656 |
| Gm26825 | 9.96E-07 | 0.400124 | 0.807 | 0.743 | 0.023484 |
| Gm5560 | 9.99E-07 | 0.346089 | 0.453 | 0.345 | 0.023559 |
| 1500011K1 | 1.03E-06 | 0.464693 | 0.249 | 0.159 | 0.024415 |

| | | | | | |
|---|---|---|---|---|---|
| Gm5552 | 1.06E-06 | 0.284615 | 0.253 | 0.163 | 0.025113 |
| Snx13 | 1.08E-06 | -1.00426 | 0.249 | 0.355 | 0.025529 |
| Gm6277 | 1.1E-06 | -0.83777 | 0.123 | 0.219 | 0.02586 |
| Rtcb | 1.1E-06 | 0.377258 | 0.401 | 0.284 | 0.025953 |
| Lpin2 | 1.1E-06 | -0.56016 | 0.316 | 0.438 | 0.025979 |
| Ucp2 | 1.11E-06 | 0.474667 | 0.717 | 0.649 | 0.026183 |
| Ccdc86 | 1.12E-06 | 0.589949 | 0.213 | 0.131 | 0.026484 |
| Actn4 | 1.19E-06 | 0.516902 | 0.354 | 0.252 | 0.028062 |
| Rpl17-ps5 | 1.2E-06 | 0.291825 | 0.332 | 0.235 | 0.028226 |
| Camk2d | 1.25E-06 | -0.68455 | 0.312 | 0.414 | 0.029532 |
| Grap | 1.27E-06 | -0.98174 | 0.209 | 0.313 | 0.029948 |
| Atp5h | 1.28E-06 | 0.520762 | 0.563 | 0.442 | 0.030191 |
| Dohh | 1.29E-06 | 0.74167 | 0.213 | 0.13 | 0.030406 |
| Stard3 | 1.31E-06 | -0.78321 | 0.372 | 0.472 | 0.030972 |
| Frrs1 | 1.33E-06 | 0.531459 | 0.48 | 0.367 | 0.031288 |
| Aif1 | 1.35E-06 | -0.31829 | 0.868 | 0.929 | 0.031817 |
| Zcrb1 | 1.36E-06 | 0.804322 | 0.253 | 0.165 | 0.032034 |
| 2410015M | 1.38E-06 | 0.410295 | 0.39 | 0.282 | 0.032651 |
| Ubb | 1.39E-06 | 0.295653 | 0.984 | 0.976 | 0.032791 |
| Pnrc1 | 1.42E-06 | -0.60998 | 0.428 | 0.524 | 0.033576 |
| Por | 1.44E-06 | 0.650296 | 0.352 | 0.25 | 0.033979 |
| Cyc1 | 1.45E-06 | 0.631936 | 0.404 | 0.3 | 0.034145 |
| Gas5 | 1.45E-06 | 0.577579 | 0.451 | 0.333 | 0.034254 |
| Mgll | 1.46E-06 | -0.77444 | 0.251 | 0.361 | 0.034404 |
| Pald1 | 1.46E-06 | -0.87547 | 0.361 | 0.473 | 0.034421 |
| Hhex | 1.47E-06 | -0.80513 | 0.294 | 0.394 | 0.034564 |
| Gm9888 | 1.49E-06 | -1.04085 | 0.291 | 0.401 | 0.035175 |
| Trim8 | 1.49E-06 | -0.59384 | 0.379 | 0.493 | 0.035188 |
| Sall3 | 1.52E-06 | -0.67932 | 0.442 | 0.56 | 0.035819 |
| Ctso | 1.54E-06 | -0.82183 | 0.287 | 0.393 | 0.036225 |
| Tns3 | 1.54E-06 | 0.769759 | 0.435 | 0.33 | 0.036286 |
| Phyhd1 | 1.59E-06 | -0.56305 | 0.444 | 0.54 | 0.037558 |
| Ndufb9 | 1.6E-06 | 0.570018 | 0.491 | 0.385 | 0.037772 |
| Dnmt3a | 1.61E-06 | 0.445798 | 0.552 | 0.425 | 0.038012 |
| Commd9 | 1.62E-06 | 0.44857 | 0.247 | 0.158 | 0.038135 |
| Ankrd12 | 1.66E-06 | 0.661959 | 0.309 | 0.214 | 0.039103 |
| Atxn7l3b | 1.67E-06 | -0.83619 | 0.388 | 0.486 | 0.039385 |
| Gm10146 | 1.71E-06 | 0.37097 | 0.2 | 0.121 | 0.040275 |
| Zkscan3 | 1.73E-06 | -0.63691 | 0.278 | 0.393 | 0.040709 |
| Irf2 | 1.74E-06 | -0.60315 | 0.229 | 0.335 | 0.041043 |
| Prdx6 | 1.74E-06 | 0.980119 | 0.213 | 0.134 | 0.04107 |
| Clip1 | 1.75E-06 | 0.441878 | 0.475 | 0.362 | 0.04129 |
| 1810058I2 | 1.77E-06 | 0.654586 | 0.296 | 0.197 | 0.041796 |
| Ubl3 | 1.81E-06 | -0.61362 | 0.623 | 0.691 | 0.042804 |
| Maml3 | 1.83E-06 | -1.04172 | 0.177 | 0.277 | 0.043251 |
| Sdcbp | 1.84E-06 | 0.451146 | 0.814 | 0.71 | 0.04352 |
| Ints6l | 1.89E-06 | -0.63853 | 0.238 | 0.347 | 0.044544 |

| | | | | | |
|---|---|---|---|---|---|
| Oaz1-ps | 1.91E-06 | 0.27868 | 0.756 | 0.653 | 0.045122 |
| Gm7931 | 1.92E-06 | 0.387993 | 0.229 | 0.148 | 0.045241 |
| Klf7 | 1.96E-06 | -0.79289 | 0.184 | 0.285 | 0.046206 |

#### Supplementry Table 2. IRM DEGs compared to HS

| GeneName | p_val | avg_log2FC | pct.1 | pct.2 | p_val_adj |
|---|---|---|---|---|---|
| Ifi204 | 0 | 4.990574 | 0.494 | 0.03 | 0 |
| Ifi27l2a | 0 | 4.793118 | 0.658 | 0.074 | 0 |
| Ifit3 | 0 | 5.376836 | 0.487 | 0.034 | 0 |
| Ifit3b | 0 | 5.643706 | 0.349 | 0.014 | 0 |
| Ifitm3 | 0 | 5.874757 | 0.781 | 0.121 | 0 |
| Irf7 | 0 | 5.105054 | 0.422 | 0.027 | 0 |
| Isg15 | 0 | 4.335334 | 0.446 | 0.032 | 0 |
| Oasl2 | 0 | 5.320428 | 0.517 | 0.029 | 0 |
| Rtp4 | 0 | 4.024983 | 0.716 | 0.093 | 0 |
| Zbp1 | 0 | 5.532375 | 0.339 | 0.01 | 0 |
| Usp18 | 8.1E-302 | 4.983595 | 0.54 | 0.063 | 1.9E-297 |
| Oas1a | 9.6E-300 | 4.347674 | 0.457 | 0.038 | 2.3E-295 |
| Lgals3bp | 1.6E-270 | 3.151595 | 0.984 | 0.503 | 3.8E-266 |
| Ifit2 | 2.7E-255 | 5.775395 | 0.321 | 0.018 | 6.4E-251 |
| Oas2 | 6.5E-238 | 4.577842 | 0.233 | 0.006 | 1.5E-233 |
| Sdc3 | 4E-225 | 3.46352 | 0.591 | 0.109 | 9.3E-221 |
| Ifi213 | 1.1E-221 | 5.657882 | 0.247 | 0.01 | 2.5E-217 |
| Ifi206 | 1.6E-220 | 4.968905 | 0.22 | 0.006 | 3.8E-216 |
| Ifi209 | 2.9E-212 | 4.274804 | 0.344 | 0.03 | 6.8E-208 |
| Sp100 | 3.3E-211 | 3.602858 | 0.519 | 0.086 | 7.9E-207 |
| Nlrc5 | 2.1E-207 | 3.506937 | 0.316 | 0.025 | 5E-203 |
| Phf11b | 5.7E-194 | 3.764004 | 0.471 | 0.076 | 1.3E-189 |
| Tgtp2 | 4.2E-193 | 2.873395 | 0.21 | 0.008 | 9.9E-189 |
| Slfn5 | 5.5E-193 | 4.799545 | 0.39 | 0.051 | 1.3E-188 |
| Ifi207 | 2.3E-185 | 5.610109 | 0.247 | 0.015 | 5.5E-181 |
| Slfn2 | 6.7E-174 | 2.597513 | 0.564 | 0.122 | 1.6E-169 |
| Ifit1 | 6E-167 | 5.095461 | 0.215 | 0.012 | 1.4E-162 |
| Bst2 | 1.5E-166 | 2.953554 | 0.817 | 0.369 | 3.5E-162 |
| H2-D1 | 6.7E-164 | 2.237341 | 0.993 | 0.867 | 1.6E-159 |
| Mx1 | 4.7E-161 | 4.557081 | 0.243 | 0.019 | 1.1E-156 |
| H2-Q7 | 1.6E-154 | 4.49404 | 0.372 | 0.059 | 3.8E-150 |
| Xaf1 | 2.6E-154 | 2.852349 | 0.533 | 0.123 | 6.1E-150 |
| Stat1 | 8.9E-154 | 2.910137 | 0.661 | 0.203 | 2.1E-149 |
| Gm4951 | 3.5E-151 | 3.610714 | 0.34 | 0.047 | 8.3E-147 |
| H2-Q6 | 2E-149 | 4.108228 | 0.344 | 0.05 | 4.7E-145 |
| Axl | 1.9E-143 | 4.208273 | 0.279 | 0.032 | 4.5E-139 |
| H2-K1 | 1.9E-139 | 2.097964 | 0.975 | 0.766 | 4.5E-135 |
| Ccl12 | 2.2E-134 | 4.880258 | 0.57 | 0.184 | 5.1E-130 |
| B2m | 1.7E-127 | 1.263367 | 1 | 0.998 | 3.9E-123 |
| Phf11d | 1.1E-123 | 3.598302 | 0.384 | 0.079 | 2.5E-119 |
| Ube2l6 | 5.1E-118 | 3.036011 | 0.363 | 0.071 | 1.2E-113 |
| Fcrls | 3.6E-116 | -1.43784 | 0.772 | 0.975 | 8.6E-112 |
| Rnf213 | 5.9E-111 | 2.48686 | 0.473 | 0.127 | 1.4E-106 |

| | | | | | |
|---|---|---|---|---|---|
| Dhx58 | 1.6E-110 | 2.959724 | 0.342 | 0.066 | 3.8E-106 |
| Isg20 | 3.4E-110 | 4.240651 | 0.215 | 0.025 | 8E-106 |
| Fgl2 | 1.6E-107 | 3.71169 | 0.282 | 0.046 | 3.9E-103 |
| Parp14 | 2.5E-99 | 2.825376 | 0.515 | 0.17 | 5.78E-95 |
| Ly6a | 9.7E-99 | 4.164725 | 0.42 | 0.121 | 2.3E-94 |
| Stat2 | 5.87E-98 | 2.985919 | 0.404 | 0.105 | 1.38E-93 |
| Ly6e | 2.71E-91 | 1.317569 | 0.986 | 0.948 | 6.39E-87 |
| Trim30a | 3.56E-87 | 2.243954 | 0.679 | 0.32 | 8.4E-83 |
| P2ry12 | 5.35E-80 | -0.79647 | 0.995 | 1 | 1.26E-75 |
| Prr5l | 8.48E-79 | 3.709376 | 0.206 | 0.033 | 2E-74 |
| Eif2ak2 | 7.37E-77 | 2.318504 | 0.416 | 0.132 | 1.74E-72 |
| Tor3a | 1.27E-74 | 2.482213 | 0.545 | 0.237 | 3E-70 |
| Gm12250 | 2.76E-73 | 3.114309 | 0.219 | 0.04 | 6.52E-69 |
| Herc6 | 2.38E-72 | 2.738032 | 0.397 | 0.127 | 5.61E-68 |
| Tspo | 6.02E-70 | 2.165861 | 0.577 | 0.274 | 1.42E-65 |
| H2-T23 | 4.61E-64 | 1.576062 | 0.859 | 0.653 | 1.09E-59 |
| Irgm1 | 3.75E-62 | 1.946012 | 0.434 | 0.163 | 8.84E-58 |
| Clec2d | 1.52E-57 | 2.116069 | 0.242 | 0.059 | 3.58E-53 |
| Trim30d | 3.7E-57 | 1.858245 | 0.534 | 0.246 | 8.72E-53 |
| H2-Q4 | 4.74E-57 | 1.045786 | 0.256 | 0.065 | 1.12E-52 |
| Ddx60 | 9.48E-55 | 2.182451 | 0.231 | 0.057 | 2.24E-50 |
| Shisa5 | 1.27E-54 | 1.304469 | 0.843 | 0.613 | 3E-50 |
| Ifi35 | 2.09E-54 | 2.075041 | 0.48 | 0.218 | 4.94E-50 |
| Tap1 | 3.26E-54 | 1.850853 | 0.462 | 0.191 | 7.69E-50 |
| Slc2a5 | 7.36E-53 | -0.90866 | 0.757 | 0.944 | 1.74E-48 |
| Gm8995 | 5.53E-52 | 2.066647 | 0.432 | 0.182 | 1.31E-47 |
| Atrip | 5.85E-50 | 0.818827 | 0.481 | 0.21 | 1.38E-45 |
| Cd52 | 6.06E-50 | 1.7565 | 0.661 | 0.411 | 1.43E-45 |
| Ddx58 | 7.83E-48 | 1.954778 | 0.342 | 0.124 | 1.85E-43 |
| Fcgr1 | 2.15E-46 | 1.313402 | 0.861 | 0.744 | 5.08E-42 |
| Adar | 4.52E-43 | 1.536661 | 0.425 | 0.189 | 1.07E-38 |
| Pml | 8.4E-43 | 2.614942 | 0.257 | 0.083 | 1.98E-38 |
| AU020206 | 5.07E-41 | 1.223541 | 0.723 | 0.473 | 1.2E-36 |
| Cd164 | 5.17E-41 | -0.78482 | 0.841 | 0.946 | 1.22E-36 |
| Fth1 | 1.82E-40 | 0.797705 | 0.996 | 0.993 | 4.3E-36 |
| Igtp | 1.06E-39 | 1.710161 | 0.519 | 0.276 | 2.49E-35 |
| Epsti1 | 3.73E-39 | 1.523525 | 0.386 | 0.167 | 8.79E-35 |
| Gm15753 | 8.09E-39 | 2.798842 | 0.302 | 0.118 | 1.91E-34 |
| Ctss | 1.25E-38 | 0.534021 | 0.996 | 1 | 2.94E-34 |
| Parp12 | 7.45E-38 | 1.555262 | 0.42 | 0.195 | 1.76E-33 |
| Samhd1 | 1.38E-35 | 1.599854 | 0.481 | 0.253 | 3.26E-31 |
| Stab1 | 1.98E-35 | -0.78108 | 0.58 | 0.822 | 4.68E-31 |
| Slfn8 | 2.01E-35 | 1.803264 | 0.21 | 0.066 | 4.75E-31 |
| Fscn1 | 1.66E-34 | -0.86042 | 0.67 | 0.859 | 3.9E-30 |
| Parp9 | 2.65E-33 | 1.394236 | 0.347 | 0.152 | 6.26E-29 |
| Znfx1 | 2.72E-33 | 1.685145 | 0.459 | 0.243 | 6.42E-29 |
| Gpr34 | 2.99E-33 | -0.5669 | 0.951 | 0.992 | 7.06E-29 |

| | | | | | |
|---|---|---|---|---|---|
| Dtx3l | 1.05E-32 | 1.33704 | 0.42 | 0.206 | 2.47E-28 |
| Lgals9 | 1.32E-32 | 1.0822 | 0.824 | 0.694 | 3.1E-28 |
| Ifih1 | 3.38E-32 | 2.25252 | 0.266 | 0.105 | 7.98E-28 |
| Sp110 | 1.47E-31 | 1.666655 | 0.494 | 0.284 | 3.48E-27 |
| Grn | 2.43E-31 | 0.542111 | 0.996 | 0.995 | 5.72E-27 |
| Ogfr | 2.03E-30 | 1.435189 | 0.36 | 0.169 | 4.79E-26 |
| Ecscr | 3.11E-30 | -0.89834 | 0.467 | 0.706 | 7.33E-26 |
| Ctsc | 1.04E-29 | 0.912785 | 0.922 | 0.842 | 2.45E-25 |
| Upk1b | 1.87E-29 | -1.23279 | 0.224 | 0.473 | 4.41E-25 |
| Rhob | 3.14E-29 | -0.6788 | 0.831 | 0.928 | 7.4E-25 |
| H2-T24 | 5.53E-29 | 1.264383 | 0.46 | 0.248 | 1.3E-24 |
| Trim34a | 8.58E-29 | 1.835115 | 0.261 | 0.106 | 2.02E-24 |
| Trim25 | 2.12E-28 | 1.789564 | 0.224 | 0.084 | 4.99E-24 |
| Lyz2 | 9.06E-28 | 1.177991 | 0.744 | 0.608 | 2.14E-23 |
| Zc3hav1 | 2.39E-27 | 1.29921 | 0.526 | 0.32 | 5.64E-23 |
| Cx3cr1 | 5.49E-27 | -0.4146 | 0.996 | 1 | 1.29E-22 |
| Nampt | 1.58E-26 | 1.735865 | 0.245 | 0.1 | 3.73E-22 |
| Psme1 | 5.62E-26 | 1.180046 | 0.674 | 0.48 | 1.33E-21 |
| Selenow | 1.86E-25 | 1.645309 | 0.335 | 0.167 | 4.38E-21 |
| Psmb8 | 4.85E-24 | 0.904516 | 0.741 | 0.565 | 1.14E-19 |
| Trim12c | 5.6E-24 | 1.467151 | 0.337 | 0.172 | 1.32E-19 |
| Slc7a8 | 1.41E-23 | -0.75919 | 0.511 | 0.723 | 3.32E-19 |
| Zup1 | 3.46E-23 | 1.39416 | 0.527 | 0.344 | 8.16E-19 |
| 4930599N2 | 6.71E-23 | 1.646904 | 0.238 | 0.105 | 1.58E-18 |
| Gpr165 | 1.21E-22 | -1.82323 | 0.079 | 0.267 | 2.85E-18 |
| Ltc4s | 1.21E-22 | -0.66871 | 0.691 | 0.867 | 2.87E-18 |
| P2ry13 | 1.52E-22 | -0.54113 | 0.922 | 0.977 | 3.58E-18 |
| Selenop | 1.53E-22 | -0.48064 | 0.979 | 0.991 | 3.62E-18 |
| Trim14 | 2.41E-22 | 1.207065 | 0.309 | 0.152 | 5.69E-18 |
| Glul | 3.08E-22 | -0.5743 | 0.832 | 0.938 | 7.26E-18 |
| Rplp1 | 4.38E-22 | 0.662737 | 0.97 | 0.952 | 1.03E-17 |
| Psme2 | 9.66E-22 | 1.098072 | 0.653 | 0.491 | 2.28E-17 |
| Entpd1 | 2.03E-21 | -0.5602 | 0.862 | 0.94 | 4.79E-17 |
| Scoc | 2.94E-21 | -0.67472 | 0.677 | 0.831 | 6.94E-17 |
| Csf2ra | 1.17E-20 | 1.242961 | 0.436 | 0.266 | 2.76E-16 |
| Calm2 | 1.58E-20 | -0.52646 | 0.871 | 0.953 | 3.73E-16 |
| Rpl41 | 2.97E-20 | 0.65869 | 0.972 | 0.934 | 7.01E-16 |
| Slc40a1 | 4.58E-20 | -1.0467 | 0.325 | 0.522 | 1.08E-15 |
| Cd74 | 4.97E-20 | 3.100736 | 0.282 | 0.153 | 1.17E-15 |
| Gm26917 | 6.78E-20 | 0.980873 | 0.97 | 0.951 | 1.6E-15 |
| Ly9 | 8.34E-20 | 1.389764 | 0.203 | 0.086 | 1.97E-15 |
| Tmem106a | 8.43E-20 | 1.429307 | 0.265 | 0.129 | 1.99E-15 |
| Ly86 | 1.08E-19 | 0.449734 | 0.993 | 0.991 | 2.54E-15 |
| Uba7 | 2.77E-19 | 0.958891 | 0.367 | 0.206 | 6.53E-15 |
| 1600014C1 | 4.97E-19 | 1.138012 | 0.374 | 0.215 | 1.17E-14 |
| Irf1 | 8.99E-19 | 1.538428 | 0.323 | 0.179 | 2.12E-14 |
| Trim12a | 3.83E-18 | 1.29586 | 0.561 | 0.412 | 9.03E-14 |

| | | | | | |
|---|---|---|---|---|---|
| Adgrg1 | 6.39E-18 | -0.45499 | 0.801 | 0.905 | 1.51E-13 |
| Il16 | 7.06E-18 | -0.71513 | 0.386 | 0.574 | 1.67E-13 |
| Ftl1-ps1 | 9.01E-18 | 0.566062 | 0.917 | 0.874 | 2.12E-13 |
| Rpsa | 9.41E-18 | 0.764071 | 0.857 | 0.774 | 2.22E-13 |
| Ctsz | 1.87E-17 | 0.477714 | 0.995 | 0.993 | 4.4E-13 |
| Lrba | 2E-17 | -0.87514 | 0.354 | 0.547 | 4.72E-13 |
| Csf1r | 3.24E-17 | -0.37458 | 0.995 | 1 | 7.65E-13 |
| Cd9 | 5.05E-17 | 0.546424 | 0.979 | 0.965 | 1.19E-12 |
| Calm1 | 5.2E-17 | 0.811533 | 0.808 | 0.748 | 1.23E-12 |
| Rnase4 | 6.39E-17 | -0.45742 | 0.894 | 0.955 | 1.51E-12 |
| Npc2 | 6.73E-17 | 0.666224 | 0.873 | 0.82 | 1.59E-12 |
| Sema4b | 7.85E-17 | -1.73315 | 0.067 | 0.213 | 1.85E-12 |
| Tnfsf13b | 8.71E-17 | 0.977361 | 0.3 | 0.162 | 2.05E-12 |
| Elmo1 | 1.62E-16 | -0.65175 | 0.578 | 0.724 | 3.83E-12 |
| Psmb9 | 1.82E-16 | 0.930992 | 0.473 | 0.316 | 4.3E-12 |
| Sox4 | 3.53E-16 | -1.03605 | 0.25 | 0.422 | 8.32E-12 |
| Casp4 | 4.57E-16 | 1.453653 | 0.212 | 0.102 | 1.08E-11 |
| Ftl1 | 5.5E-16 | 0.483149 | 0.993 | 0.991 | 1.3E-11 |
| Emp3 | 5.53E-16 | 1.53309 | 0.245 | 0.128 | 1.3E-11 |
| Rnase6 | 1.08E-15 | 1.512847 | 0.217 | 0.108 | 2.55E-11 |
| Abcc3 | 1.24E-15 | -0.98023 | 0.259 | 0.427 | 2.92E-11 |
| Hck | 1.37E-15 | 0.767721 | 0.728 | 0.613 | 3.23E-11 |
| Gm46224 | 1.56E-15 | 1.131491 | 0.261 | 0.138 | 3.68E-11 |
| Tent5a | 1.72E-15 | 0.989947 | 0.543 | 0.398 | 4.05E-11 |
| Irf9 | 2.4E-15 | 1.182558 | 0.326 | 0.194 | 5.66E-11 |
| Rps19 | 3.25E-15 | 0.642101 | 0.912 | 0.863 | 7.67E-11 |
| Tor1aip1 | 4.32E-15 | 0.923051 | 0.684 | 0.567 | 1.02E-10 |
| Golm1 | 5.58E-15 | -0.45769 | 0.735 | 0.877 | 1.32E-10 |
| Tuba1a | 5.87E-15 | -0.70602 | 0.453 | 0.61 | 1.38E-10 |
| Naalad2 | 1.12E-14 | 1.307908 | 0.263 | 0.143 | 2.64E-10 |
| S1pr1 | 1.16E-14 | -0.90402 | 0.155 | 0.309 | 2.74E-10 |
| Abca9 | 1.18E-14 | -0.5202 | 0.538 | 0.695 | 2.79E-10 |
| Mlph | 1.25E-14 | -0.78246 | 0.247 | 0.411 | 2.96E-10 |
| Rpl32 | 1.34E-14 | 0.633712 | 0.894 | 0.863 | 3.16E-10 |
| Ddit4 | 1.38E-14 | -1.14289 | 0.104 | 0.25 | 3.27E-10 |
| Ctsh | 1.65E-14 | 0.501416 | 0.958 | 0.947 | 3.89E-10 |
| Cmtm6 | 1.74E-14 | -0.51298 | 0.751 | 0.854 | 4.11E-10 |
| Hsp90ab1 | 1.77E-14 | 0.63978 | 0.901 | 0.844 | 4.18E-10 |
| Rpl13 | 2.14E-14 | 0.544118 | 0.949 | 0.938 | 5.04E-10 |
| H2-M3 | 2.95E-14 | 1.030704 | 0.31 | 0.184 | 6.96E-10 |
| Capg | 3E-14 | 1.516079 | 0.27 | 0.153 | 7.08E-10 |
| Rps18 | 3.5E-14 | 0.660503 | 0.891 | 0.833 | 8.25E-10 |
| Eef1a1 | 3.88E-14 | 0.351002 | 0.998 | 0.994 | 9.15E-10 |
| Zfas1 | 4.86E-14 | 0.868174 | 0.238 | 0.127 | 1.15E-09 |
| Naaa | 5.96E-14 | 1.100453 | 0.367 | 0.234 | 1.41E-09 |
| C1qb | 7.64E-14 | 0.265767 | 0.996 | 1 | 1.8E-09 |
| Vsir | 7.79E-14 | -0.33962 | 0.951 | 0.985 | 1.84E-09 |

| | | | | | |
|---|---|---|---|---|---|
| Rpl18a | 8.89E-14 | 0.449931 | 0.991 | 0.986 | 2.1E-09 |
| Plxdc2 | 9E-14 | -0.3822 | 0.847 | 0.931 | 2.12E-09 |
| Tmem184b | 9.58E-14 | 0.717068 | 0.469 | 0.323 | 2.26E-09 |
| Cebpb | 2.43E-13 | 0.763216 | 0.647 | 0.529 | 5.73E-09 |
| Gm6977 | 2.46E-13 | 0.471594 | 0.608 | 0.479 | 5.81E-09 |
| Ccdc86 | 3.33E-13 | 1.246807 | 0.24 | 0.131 | 7.85E-09 |
| Fam102b | 3.94E-13 | -0.57622 | 0.52 | 0.665 | 9.29E-09 |
| Rpl14 | 8.89E-13 | 0.65316 | 0.822 | 0.756 | 2.1E-08 |
| Dopey2 | 9.04E-13 | 1.344603 | 0.272 | 0.158 | 2.13E-08 |
| Rpl36 | 1.04E-12 | 0.591037 | 0.859 | 0.799 | 2.45E-08 |
| Sirpa | 1.16E-12 | -0.41662 | 0.912 | 0.958 | 2.73E-08 |
| Tmem204 | 1.18E-12 | -1.56937 | 0.101 | 0.225 | 2.79E-08 |
| Rtn4rl1 | 1.79E-12 | -0.53934 | 0.545 | 0.684 | 4.23E-08 |
| Cyba | 1.93E-12 | 0.370174 | 0.975 | 0.973 | 4.55E-08 |
| Gm10076 | 2.57E-12 | 0.502758 | 0.926 | 0.893 | 6.05E-08 |
| Fchsd2 | 2.59E-12 | -0.87982 | 0.28 | 0.429 | 6.12E-08 |
| Fam110a | 2.95E-12 | -0.87007 | 0.213 | 0.364 | 6.95E-08 |
| Sall1 | 3.76E-12 | -0.45431 | 0.771 | 0.856 | 8.86E-08 |
| Fcrl1 | 3.79E-12 | -1.18665 | 0.152 | 0.287 | 8.93E-08 |
| Gm10443 | 3.8E-12 | 0.493483 | 0.499 | 0.37 | 8.95E-08 |
| Rpsa-ps10 | 5.6E-12 | 0.641312 | 0.642 | 0.532 | 1.32E-07 |
| Man1c1 | 5.67E-12 | -0.74665 | 0.376 | 0.521 | 1.34E-07 |
| Csnk1e | 6.81E-12 | -0.53521 | 0.476 | 0.622 | 1.61E-07 |
| Rps28 | 6.83E-12 | 0.668479 | 0.938 | 0.892 | 1.61E-07 |
| Ivns1abp | 9.29E-12 | -0.41328 | 0.767 | 0.852 | 2.19E-07 |
| Tmsb10 | 1.4E-11 | 0.793748 | 0.522 | 0.412 | 3.3E-07 |
| Rps20 | 1.47E-11 | 0.598189 | 0.854 | 0.807 | 3.46E-07 |
| Dusp6 | 1.54E-11 | -0.53143 | 0.616 | 0.731 | 3.63E-07 |
| Luc7l3 | 1.6E-11 | 0.87476 | 0.515 | 0.391 | 3.77E-07 |
| Apobec3 | 1.75E-11 | 0.798859 | 0.624 | 0.49 | 4.13E-07 |
| Maf | 2E-11 | -0.44398 | 0.808 | 0.897 | 4.72E-07 |
| Ifngr1 | 2.01E-11 | -0.3044 | 0.931 | 0.972 | 4.74E-07 |
| Crybb1 | 2.57E-11 | -0.35852 | 0.653 | 0.794 | 6.07E-07 |
| Ckb | 2.59E-11 | -0.36585 | 0.804 | 0.895 | 6.11E-07 |
| Apobec1 | 3.04E-11 | 0.635826 | 0.374 | 0.245 | 7.18E-07 |
| Sat1 | 3.27E-11 | 0.472013 | 0.892 | 0.847 | 7.72E-07 |
| C1qa | 3.31E-11 | 0.25598 | 0.996 | 1 | 7.8E-07 |
| Eef2k | 3.54E-11 | -0.71143 | 0.3 | 0.448 | 8.34E-07 |
| Ctsb | 3.69E-11 | 0.354751 | 0.996 | 0.999 | 8.69E-07 |
| Anxa6 | 3.7E-11 | 0.591998 | 0.3 | 0.183 | 8.72E-07 |
| Rps7 | 4.34E-11 | 0.624178 | 0.836 | 0.776 | 1.02E-06 |
| Rps24 | 4.82E-11 | 0.616517 | 0.887 | 0.818 | 1.14E-06 |
| Rplp2 | 5.16E-11 | 0.544683 | 0.924 | 0.895 | 1.22E-06 |
| Tpp1 | 5.88E-11 | -0.42609 | 0.656 | 0.771 | 1.39E-06 |
| Prpsap2 | 6.29E-11 | -0.82371 | 0.217 | 0.359 | 1.48E-06 |
| Cd47 | 6.35E-11 | 0.710414 | 0.744 | 0.655 | 1.5E-06 |
| Rpl24 | 6.53E-11 | 0.604368 | 0.695 | 0.604 | 1.54E-06 |

| | | | | | |
|---|---|---|---|---|---|
| 9930111J2 | 7.24E-11 | 1.186306 | 0.441 | 0.325 | 1.71E-06 |
| Tmem219 | 7.94E-11 | 0.74738 | 0.499 | 0.376 | 1.87E-06 |
| Lsp1 | 8.02E-11 | 1.013047 | 0.404 | 0.284 | 1.89E-06 |
| Etnk1 | 8.3E-11 | 0.963483 | 0.466 | 0.343 | 1.96E-06 |
| Gm26740 | 8.48E-11 | 1.15614 | 0.217 | 0.123 | 2E-06 |
| Idh2 | 8.99E-11 | -0.82788 | 0.307 | 0.444 | 2.12E-06 |
| Rps11 | 9.18E-11 | 0.487432 | 0.944 | 0.911 | 2.16E-06 |
| Bank1 | 1.15E-10 | -1.1889 | 0.088 | 0.2 | 2.7E-06 |
| Ftl2-ps | 1.15E-10 | 0.429517 | 0.954 | 0.914 | 2.71E-06 |
| Ppia | 1.19E-10 | 0.367856 | 0.974 | 0.969 | 2.82E-06 |
| Rps6ka1 | 1.27E-10 | -0.45271 | 0.714 | 0.813 | 3E-06 |
| Trafd1 | 1.29E-10 | 0.784228 | 0.476 | 0.347 | 3.05E-06 |
| Clstn1 | 1.38E-10 | -0.66703 | 0.298 | 0.432 | 3.26E-06 |
| Wasf2 | 1.52E-10 | -0.41379 | 0.781 | 0.862 | 3.59E-06 |
| Qk | 2.13E-10 | -0.34272 | 0.868 | 0.936 | 5.03E-06 |
| 1700017B0 | 2.19E-10 | -0.36972 | 0.552 | 0.694 | 5.16E-06 |
| Pmepa1 | 2.65E-10 | -0.4085 | 0.739 | 0.845 | 6.25E-06 |
| Adgre1 | 2.76E-10 | -0.47456 | 0.489 | 0.629 | 6.51E-06 |
| Gm13456 | 3.22E-10 | 0.34117 | 0.972 | 0.948 | 7.6E-06 |
| Rpl37a | 3.89E-10 | 0.439006 | 0.947 | 0.929 | 9.18E-06 |
| Dusp7 | 3.92E-10 | -0.46015 | 0.446 | 0.588 | 9.24E-06 |
| Gm15427 | 4.03E-10 | 0.522486 | 0.958 | 0.942 | 9.51E-06 |
| Gm10073 | 4.27E-10 | 0.338565 | 0.681 | 0.568 | 1.01E-05 |
| Slc35c2 | 4.32E-10 | -0.73286 | 0.337 | 0.464 | 1.02E-05 |
| Abcg1 | 4.58E-10 | -0.28259 | 0.506 | 0.657 | 1.08E-05 |
| Ifi30 | 5.27E-10 | 0.594614 | 0.58 | 0.468 | 1.24E-05 |
| Rnf13 | 5.31E-10 | -0.45337 | 0.686 | 0.776 | 1.25E-05 |
| Rps12 | 6.78E-10 | 0.557285 | 0.866 | 0.812 | 1.6E-05 |
| Usp25 | 6.8E-10 | 0.704635 | 0.295 | 0.186 | 1.61E-05 |
| Nmi | 6.81E-10 | 1.183418 | 0.245 | 0.15 | 1.61E-05 |
| Tle3 | 6.84E-10 | -0.81596 | 0.192 | 0.317 | 1.61E-05 |
| Apoe | 6.96E-10 | 2.659278 | 0.455 | 0.376 | 1.64E-05 |
| Rplp0 | 7.48E-10 | 0.472729 | 0.914 | 0.89 | 1.76E-05 |
| Inpp4b | 7.49E-10 | -1.15345 | 0.136 | 0.252 | 1.77E-05 |
| Gp9 | 8.16E-10 | -0.92734 | 0.113 | 0.228 | 1.92E-05 |
| Cbfa2t3 | 8.18E-10 | -0.51234 | 0.238 | 0.375 | 1.93E-05 |
| Sort1 | 8.36E-10 | -0.44514 | 0.379 | 0.52 | 1.97E-05 |
| Cited2 | 9.32E-10 | 0.970689 | 0.295 | 0.193 | 2.2E-05 |
| Rps13 | 1.02E-09 | 0.489505 | 0.822 | 0.766 | 2.4E-05 |
| Sgk1 | 1.04E-09 | -0.57063 | 0.469 | 0.584 | 2.45E-05 |
| Prkab1 | 1.04E-09 | -0.79105 | 0.263 | 0.381 | 2.46E-05 |
| Zfhx3 | 1.06E-09 | -0.38965 | 0.899 | 0.948 | 2.5E-05 |
| Sqor | 1.09E-09 | -0.7149 | 0.194 | 0.319 | 2.56E-05 |
| Lpcat3 | 1.1E-09 | -0.44098 | 0.411 | 0.547 | 2.6E-05 |
| Hpgd | 1.12E-09 | -0.48562 | 0.647 | 0.746 | 2.64E-05 |
| Clec4a3 | 1.29E-09 | -0.40394 | 0.277 | 0.415 | 3.05E-05 |
| Cables1 | 1.31E-09 | -0.76411 | 0.171 | 0.29 | 3.1E-05 |

| | | | | | |
|---|---|---|---|---|---|
| Ulk2 | 1.35E-09 | -0.35179 | 0.434 | 0.575 | 3.19E-05 |
| Rpl19 | 1.6E-09 | 0.516465 | 0.836 | 0.794 | 3.77E-05 |
| Zfp36l1 | 1.88E-09 | -0.39767 | 0.667 | 0.78 | 4.44E-05 |
| Rps2 | 1.98E-09 | 0.678416 | 0.743 | 0.675 | 4.66E-05 |
| Klhl24 | 2.04E-09 | -0.6521 | 0.254 | 0.38 | 4.82E-05 |
| Sesn1 | 2.13E-09 | -0.64029 | 0.314 | 0.439 | 5.01E-05 |
| Arhgef40 | 2.25E-09 | -0.53389 | 0.34 | 0.469 | 5.31E-05 |
| Rpl38 | 2.27E-09 | 0.639295 | 0.686 | 0.616 | 5.35E-05 |
| Rnf114 | 2.4E-09 | 0.740493 | 0.464 | 0.349 | 5.67E-05 |
| Tspan4 | 2.46E-09 | -0.52018 | 0.332 | 0.464 | 5.8E-05 |
| Rpl35 | 2.68E-09 | 0.535651 | 0.857 | 0.807 | 6.33E-05 |
| Cdk5r1 | 2.75E-09 | -0.69105 | 0.168 | 0.283 | 6.49E-05 |
| Plcg1 | 2.85E-09 | -1.05355 | 0.102 | 0.204 | 6.72E-05 |
| Nrip1 | 2.88E-09 | -0.32123 | 0.522 | 0.654 | 6.8E-05 |
| Slco2b1 | 3.14E-09 | -0.3244 | 0.854 | 0.937 | 7.4E-05 |
| Btg2 | 3.21E-09 | -0.37214 | 0.471 | 0.584 | 7.58E-05 |
| Hfe | 3.25E-09 | -0.37665 | 0.317 | 0.452 | 7.66E-05 |
| Gm3511 | 3.31E-09 | 0.449064 | 0.635 | 0.534 | 7.81E-05 |
| Rpl30 | 3.46E-09 | 0.462205 | 0.891 | 0.858 | 8.17E-05 |
| Pik3cg | 3.65E-09 | -0.57541 | 0.3 | 0.432 | 8.62E-05 |
| Tor1aip2 | 3.71E-09 | 0.757803 | 0.432 | 0.319 | 8.75E-05 |
| Lrch3 | 3.75E-09 | 1.000619 | 0.21 | 0.123 | 8.84E-05 |
| Tmem135 | 4.36E-09 | -0.48646 | 0.247 | 0.371 | 0.000103 |
| Rps9 | 4.51E-09 | 0.36576 | 0.958 | 0.95 | 0.000106 |
| Mfng | 4.55E-09 | -0.75588 | 0.303 | 0.422 | 0.000107 |
| Tpt1 | 4.61E-09 | 0.437907 | 0.907 | 0.878 | 0.000109 |
| Mtss1 | 4.63E-09 | -0.53716 | 0.252 | 0.37 | 0.000109 |
| Ptma | 4.88E-09 | 0.684912 | 0.698 | 0.629 | 0.000115 |
| Rps29 | 4.88E-09 | 0.467879 | 0.986 | 0.972 | 0.000115 |
| Rpl23 | 5.31E-09 | 0.582491 | 0.861 | 0.82 | 0.000125 |
| Gas6 | 5.35E-09 | -0.69083 | 0.363 | 0.482 | 0.000126 |
| Rps27a | 5.7E-09 | 0.543753 | 0.827 | 0.78 | 0.000134 |
| Ccr5 | 5.93E-09 | -0.27615 | 0.582 | 0.717 | 0.00014 |
| Trim56 | 5.98E-09 | 1.155628 | 0.236 | 0.147 | 0.000141 |
| Psap | 6.1E-09 | 0.252665 | 0.996 | 0.999 | 0.000144 |
| Susd3 | 6.12E-09 | -0.28855 | 0.734 | 0.846 | 0.000144 |
| Rpl11 | 6.3E-09 | 0.674959 | 0.776 | 0.713 | 0.000149 |
| Tnfrsf21 | 8.38E-09 | -0.65754 | 0.402 | 0.513 | 0.000198 |
| C5ar2 | 8.61E-09 | -0.34807 | 0.538 | 0.658 | 0.000203 |
| Cd81 | 9.15E-09 | 0.28288 | 0.996 | 0.999 | 0.000216 |
| Acer3 | 9.24E-09 | 0.742335 | 0.545 | 0.434 | 0.000218 |
| Zfp706 | 9.76E-09 | -0.3453 | 0.519 | 0.661 | 0.00023 |
| Rps14 | 9.82E-09 | 0.450637 | 0.931 | 0.914 | 0.000232 |
| Gal3st4 | 1.27E-08 | -0.38126 | 0.455 | 0.58 | 0.000299 |
| Dnm2 | 1.28E-08 | -0.38789 | 0.575 | 0.687 | 0.000303 |
| Tnfrsf13b | 1.32E-08 | -0.63148 | 0.217 | 0.331 | 0.000311 |
| Gm7331 | 1.37E-08 | 0.49309 | 0.621 | 0.528 | 0.000322 |

| | | | | | |
|---|---|---|---|---|---|
| Rpl36-ps2 | 1.44E-08 | 0.489574 | 0.568 | 0.478 | 0.00034 |
| Cttnbp2nl | 1.48E-08 | -0.33389 | 0.66 | 0.755 | 0.000349 |
| Serinc1 | 1.56E-08 | -0.43318 | 0.55 | 0.657 | 0.000368 |
| Creb3l2 | 1.65E-08 | -1.09174 | 0.134 | 0.233 | 0.000389 |
| Lat2 | 1.67E-08 | 0.611004 | 0.464 | 0.358 | 0.000394 |
| Scamp5 | 1.75E-08 | -0.37814 | 0.446 | 0.578 | 0.000413 |
| Gpr183 | 1.82E-08 | -0.37086 | 0.55 | 0.673 | 0.000429 |
| Gm11518 | 1.82E-08 | 0.435001 | 0.578 | 0.478 | 0.00043 |
| Rnf130 | 1.83E-08 | -0.3146 | 0.73 | 0.812 | 0.000431 |
| Rpl12 | 1.9E-08 | 0.623789 | 0.73 | 0.659 | 0.000448 |
| Rpl37 | 1.92E-08 | 0.455055 | 0.912 | 0.886 | 0.000454 |
| Nupr1 | 2.08E-08 | 1.094618 | 0.212 | 0.129 | 0.00049 |
| Tmpo | 2.16E-08 | 0.852716 | 0.287 | 0.195 | 0.000509 |
| Ptafr | 2.17E-08 | -0.43543 | 0.654 | 0.783 | 0.000511 |
| Prdx1 | 2.24E-08 | 0.703527 | 0.593 | 0.495 | 0.000528 |
| Mylip | 2.63E-08 | -0.3699 | 0.577 | 0.699 | 0.00062 |
| Gatm | 2.63E-08 | 0.667492 | 0.358 | 0.252 | 0.00062 |
| Rps8 | 2.75E-08 | 0.528555 | 0.857 | 0.813 | 0.000648 |
| Sh2b3 | 3.16E-08 | -0.60615 | 0.326 | 0.449 | 0.000744 |
| Ptgs1 | 3.33E-08 | -0.33322 | 0.864 | 0.946 | 0.000785 |
| Cd302 | 3.36E-08 | -0.50076 | 0.354 | 0.469 | 0.000792 |
| Gm4617 | 3.4E-08 | 0.477058 | 0.365 | 0.266 | 0.000802 |
| Rpl14-ps1 | 4.05E-08 | 0.287388 | 0.661 | 0.563 | 0.000954 |
| Gm2000 | 4.48E-08 | 0.419622 | 0.646 | 0.571 | 0.001056 |
| Gm29216 | 4.5E-08 | 0.360595 | 0.804 | 0.759 | 0.001062 |
| Rpl27 | 4.79E-08 | 0.433885 | 0.757 | 0.688 | 0.001129 |
| Anxa5 | 4.96E-08 | 0.691189 | 0.247 | 0.159 | 0.00117 |
| Gm8430 | 5.04E-08 | 0.406732 | 0.594 | 0.496 | 0.001188 |
| Cxcl16 | 5.18E-08 | 1.222873 | 0.337 | 0.244 | 0.001222 |
| Rps19-ps6 | 5.33E-08 | 0.373192 | 0.698 | 0.618 | 0.001256 |
| Gm11478 | 6.39E-08 | 0.567126 | 0.734 | 0.663 | 0.001507 |
| Malat1 | 6.63E-08 | 0.30367 | 0.996 | 0.992 | 0.001564 |
| Rpl13a | 6.82E-08 | 0.52495 | 0.854 | 0.824 | 0.00161 |
| Ccng2 | 7.35E-08 | -0.64624 | 0.25 | 0.365 | 0.001735 |
| Ms4a6c | 8.64E-08 | 1.269448 | 0.206 | 0.131 | 0.002037 |
| Fermt3 | 9.08E-08 | -0.48557 | 0.681 | 0.773 | 0.002141 |
| Uba52 | 9.19E-08 | 0.524716 | 0.783 | 0.721 | 0.002169 |
| Tm9sf2 | 9.41E-08 | -0.25098 | 0.668 | 0.769 | 0.00222 |
| Itga9 | 9.79E-08 | -0.51263 | 0.185 | 0.291 | 0.002309 |
| Gm6277 | 1.02E-07 | -0.49558 | 0.122 | 0.219 | 0.002406 |
| Eif4ebp1 | 1.02E-07 | 1.003588 | 0.235 | 0.151 | 0.002408 |
| Numb | 1.04E-07 | -0.49376 | 0.469 | 0.592 | 0.002444 |
| Sppl2a | 1.19E-07 | 0.54778 | 0.684 | 0.589 | 0.002813 |
| Nckap1l | 1.24E-07 | -0.31654 | 0.582 | 0.69 | 0.002936 |
| Rpl31 | 1.27E-07 | 0.477549 | 0.817 | 0.785 | 0.002995 |
| Commd8 | 1.27E-07 | -0.49236 | 0.614 | 0.689 | 0.003002 |
| Mertk | 1.31E-07 | -0.38288 | 0.808 | 0.872 | 0.003086 |

| | | | | | |
|---|---|---|---|---|---|
| Rps16 | 1.43E-07 | 0.497209 | 0.799 | 0.76 | 0.003365 |
| Gm14303 | 1.44E-07 | 0.408796 | 0.808 | 0.753 | 0.003394 |
| Sgk3 | 1.54E-07 | 0.903022 | 0.296 | 0.205 | 0.003643 |
| Rpl39 | 1.57E-07 | 0.611371 | 0.767 | 0.699 | 0.003699 |
| Rpl10a | 1.62E-07 | 0.612448 | 0.718 | 0.643 | 0.003818 |
| Gm33699 | 1.77E-07 | -0.59385 | 0.159 | 0.258 | 0.004166 |
| Ap1b1 | 1.9E-07 | -0.853 | 0.354 | 0.459 | 0.004485 |
| Ifitm10 | 1.91E-07 | -0.34429 | 0.402 | 0.528 | 0.0045 |
| March5 | 1.95E-07 | 0.837078 | 0.231 | 0.151 | 0.004603 |
| Tdrd7 | 1.97E-07 | 0.867649 | 0.222 | 0.14 | 0.00464 |
| Slamf9 | 1.98E-07 | 0.661596 | 0.413 | 0.321 | 0.00466 |
| Sh3bgrl3 | 2.16E-07 | 0.473569 | 0.829 | 0.812 | 0.005091 |
| Mgll | 2.24E-07 | -0.69848 | 0.257 | 0.361 | 0.005277 |
| Nfam1 | 2.39E-07 | -0.27866 | 0.519 | 0.632 | 0.005635 |
| Ninj1 | 2.43E-07 | -0.5514 | 0.284 | 0.392 | 0.005742 |
| Gm9385 | 2.52E-07 | 0.344773 | 0.402 | 0.311 | 0.005943 |
| Txnip | 2.54E-07 | -0.42065 | 0.614 | 0.699 | 0.005989 |
| Pfkfb3 | 2.57E-07 | 0.734058 | 0.319 | 0.227 | 0.006062 |
| Gm10790 | 2.61E-07 | -0.39833 | 0.492 | 0.605 | 0.006145 |
| Hmox2 | 2.66E-07 | 0.692901 | 0.531 | 0.45 | 0.006272 |
| mt-Co3 | 2.85E-07 | 0.406479 | 0.912 | 0.902 | 0.006724 |
| Eng | 2.96E-07 | -0.34611 | 0.409 | 0.519 | 0.006987 |
| Sla | 3.2E-07 | -0.46813 | 0.443 | 0.545 | 0.007538 |
| Sqstm1 | 3.22E-07 | -0.46195 | 0.52 | 0.606 | 0.007604 |
| Aif1 | 3.45E-07 | 0.371934 | 0.921 | 0.929 | 0.008149 |
| Gm5453 | 3.57E-07 | 0.255165 | 0.259 | 0.177 | 0.008427 |
| Rpl4 | 3.84E-07 | 0.415442 | 0.873 | 0.829 | 0.009062 |
| Rpl23a | 3.89E-07 | 0.462778 | 0.801 | 0.745 | 0.009173 |
| Pik3ap1 | 4.13E-07 | 0.758749 | 0.354 | 0.262 | 0.00974 |
| Ppp1r18 | 4.18E-07 | -0.41271 | 0.517 | 0.626 | 0.00986 |
| Rpl19-ps11 | 4.26E-07 | 0.394089 | 0.663 | 0.589 | 0.010056 |
| Rps16-ps2 | 4.37E-07 | 0.418628 | 0.64 | 0.559 | 0.010302 |
| Khk | 4.38E-07 | -0.58665 | 0.194 | 0.293 | 0.010332 |
| mt-Co2 | 4.45E-07 | 0.386332 | 0.852 | 0.84 | 0.010501 |
| Neat1 | 4.54E-07 | 1.049127 | 0.265 | 0.183 | 0.010704 |
| Cfh | 4.57E-07 | -0.25212 | 0.716 | 0.779 | 0.010788 |
| Pald1 | 4.59E-07 | -0.6398 | 0.367 | 0.473 | 0.010817 |
| Ypel3 | 4.6E-07 | -0.34906 | 0.568 | 0.671 | 0.010862 |
| Gm4149 | 4.86E-07 | 0.362468 | 0.85 | 0.813 | 0.011457 |
| Slc46a1 | 4.87E-07 | -0.40469 | 0.339 | 0.455 | 0.011493 |
| Fam111a | 5E-07 | 0.96745 | 0.333 | 0.248 | 0.011787 |
| Chsy1 | 5.41E-07 | -0.40518 | 0.372 | 0.496 | 0.012763 |
| Gm10288 | 5.43E-07 | 0.640286 | 0.508 | 0.419 | 0.012809 |
| Gm10123 | 5.48E-07 | 0.303615 | 0.929 | 0.927 | 0.012922 |
| 2410006H1 | 5.54E-07 | 0.761119 | 0.28 | 0.198 | 0.013067 |
| Gcn1l1 | 5.56E-07 | -0.56527 | 0.367 | 0.464 | 0.013111 |
| Pip4k2a | 6E-07 | -0.36644 | 0.471 | 0.579 | 0.014153 |

| | | | | | |
|---|---|---|---|---|---|
| Mb21d1 | 6.31E-07 | 0.562755 | 0.277 | 0.19 | 0.014875 |
| Tmem160 | 6.8E-07 | 0.60154 | 0.404 | 0.311 | 0.016049 |
| Scpep1 | 6.92E-07 | 0.522122 | 0.466 | 0.366 | 0.016322 |
| Sash3 | 6.92E-07 | -0.45862 | 0.307 | 0.412 | 0.016332 |
| Gm15266 | 7.1E-07 | 0.291958 | 0.695 | 0.649 | 0.016739 |
| Chst7 | 7.74E-07 | -0.62431 | 0.189 | 0.286 | 0.018248 |
| Gm9800 | 8.2E-07 | 0.361973 | 0.305 | 0.219 | 0.019345 |
| Rps15 | 8.51E-07 | 0.517721 | 0.781 | 0.737 | 0.020064 |
| Tgfb1 | 9E-07 | -0.26274 | 0.233 | 0.338 | 0.021224 |
| Ets1 | 9.34E-07 | -0.47414 | 0.249 | 0.351 | 0.022023 |
| Gm6548 | 9.41E-07 | 1.535606 | 0.314 | 0.236 | 0.022196 |
| Ankle2 | 9.49E-07 | 0.752012 | 0.464 | 0.371 | 0.022394 |
| Parvg | 9.6E-07 | -0.42815 | 0.353 | 0.457 | 0.022647 |
| Atxn1 | 1.01E-06 | -1.16361 | 0.139 | 0.222 | 0.023888 |
| mt-Co1 | 1.09E-06 | 0.32 | 0.988 | 0.985 | 0.025742 |
| Gm47283 | 1.11E-06 | 0.83538 | 0.381 | 0.293 | 0.026279 |
| Rnf167 | 1.12E-06 | -0.27056 | 0.414 | 0.524 | 0.026475 |
| Fcer1g | 1.12E-06 | 0.255409 | 0.986 | 0.993 | 0.02648 |
| Gm3699 | 1.18E-06 | 0.485496 | 0.416 | 0.326 | 0.027733 |
| Rpl21 | 1.19E-06 | 0.364431 | 0.91 | 0.917 | 0.028019 |
| Glns-ps1 | 1.25E-06 | -0.25209 | 0.152 | 0.241 | 0.029511 |
| Edem2 | 1.29E-06 | -0.3288 | 0.476 | 0.586 | 0.030415 |
| Cxxc5 | 1.29E-06 | -0.31609 | 0.61 | 0.707 | 0.030518 |
| Mxd4 | 1.32E-06 | -0.99298 | 0.238 | 0.33 | 0.031121 |
| Rpl9 | 1.34E-06 | 0.523562 | 0.788 | 0.75 | 0.031708 |
| Chmp4b | 1.36E-06 | 0.620439 | 0.233 | 0.155 | 0.032033 |
| Tap2 | 1.39E-06 | 0.532222 | 0.437 | 0.34 | 0.032762 |
| Pdlim4 | 1.39E-06 | -0.50414 | 0.259 | 0.356 | 0.032834 |
| Cox8a | 1.45E-06 | 0.592016 | 0.661 | 0.605 | 0.03413 |
| Cd33 | 1.49E-06 | -0.34882 | 0.688 | 0.775 | 0.03522 |
| Gm9625 | 1.5E-06 | 0.433728 | 0.559 | 0.477 | 0.035385 |
| Ccdc93 | 1.55E-06 | -0.62969 | 0.187 | 0.284 | 0.0366 |
| Rps5 | 1.62E-06 | 0.460835 | 0.78 | 0.718 | 0.03831 |
| Nfatc1 | 1.69E-06 | -0.31242 | 0.362 | 0.455 | 0.039782 |
| Slc37a2 | 1.71E-06 | -0.6148 | 0.349 | 0.443 | 0.040411 |
| 0610040J0 | 1.73E-06 | -0.43935 | 0.45 | 0.543 | 0.04084 |
| Mef2c | 1.87E-06 | -0.31254 | 0.787 | 0.846 | 0.044103 |
| Dok3 | 1.99E-06 | -0.58266 | 0.196 | 0.286 | 0.046914 |
| Nsd3 | 2.02E-06 | 0.736295 | 0.524 | 0.432 | 0.047596 |
| Ubash3b | 2.12E-06 | -0.49192 | 0.52 | 0.608 | 0.049894 |
| Timp2 | 2.12E-06 | 0.402495 | 0.859 | 0.824 | 0.049945 |

### Supplementry Table 2. TNFRM DEGs compared to HS

| GeneName | p_val | avg_log2FC | pct.1 | pct.2 | p_val_adj |
|---|---|---|---|---|---|
| Marcksl1 | 4.66E-60 | 4.504526 | 0.554 | 0.085 | 1.1E-55 |
| Cd83 | 4.93E-59 | 4.417118 | 0.87 | 0.26 | 1.16E-54 |
| Vcam1 | 9.49E-52 | 4.831763 | 0.326 | 0.032 | 2.24E-47 |
| H2-Q6 | 3.06E-45 | 3.99853 | 0.38 | 0.05 | 7.22E-41 |
| Gpr84 | 7.25E-34 | 3.702977 | 0.859 | 0.457 | 1.71E-29 |
| Slamf8 | 9.64E-33 | 3.53992 | 0.446 | 0.091 | 2.27E-28 |
| Bcl3 | 1.8E-32 | 4.855389 | 0.239 | 0.028 | 4.24E-28 |
| Cd74 | 1.45E-31 | 4.671314 | 0.554 | 0.153 | 3.42E-27 |
| Ccl4 | 2.04E-30 | 7.370941 | 0.207 | 0.022 | 4.8E-26 |
| Ccl12 | 7.19E-30 | 5.57537 | 0.598 | 0.184 | 1.7E-25 |
| Tlr2 | 1.29E-28 | 3.228101 | 0.783 | 0.35 | 3.03E-24 |
| Icam1 | 4.52E-28 | 2.680786 | 0.815 | 0.371 | 1.07E-23 |
| Nfkbia | 1.72E-25 | 2.858498 | 0.848 | 0.465 | 4.07E-21 |
| Relb | 1.74E-25 | 2.279148 | 0.446 | 0.106 | 4.1E-21 |
| P2ry12 | 2.67E-25 | -1.18988 | 0.978 | 1 | 6.29E-21 |
| H2-Q7 | 1.02E-24 | 3.505388 | 0.315 | 0.059 | 2.41E-20 |
| H2-Q4 | 4.32E-22 | 4.226103 | 0.315 | 0.065 | 1.02E-17 |
| Cxcl16 | 1.39E-21 | 3.127727 | 0.63 | 0.244 | 3.27E-17 |
| Nfkbie | 5.54E-21 | 2.48775 | 0.413 | 0.11 | 1.31E-16 |
| Herpud1 | 6.11E-19 | 2.219006 | 0.837 | 0.576 | 1.44E-14 |
| H2-K1 | 9.46E-19 | 2.005177 | 0.957 | 0.766 | 2.23E-14 |
| Nfkb2 | 8.68E-18 | 3.307389 | 0.391 | 0.111 | 2.05E-13 |
| Cd52 | 8.91E-18 | 2.099418 | 0.761 | 0.411 | 2.1E-13 |
| Csf1 | 1.17E-17 | 3.322522 | 0.217 | 0.04 | 2.77E-13 |
| Birc3 | 6.49E-17 | 2.437112 | 0.391 | 0.116 | 1.53E-12 |
| Tnfaip2 | 1.21E-16 | 3.750465 | 0.337 | 0.093 | 2.85E-12 |
| Arl5c | 7.37E-16 | 3.722787 | 0.337 | 0.097 | 1.74E-11 |
| Fcrls | 1.51E-15 | -1.09825 | 0.804 | 0.975 | 3.56E-11 |
| Bcl2a1a | 2.75E-15 | 1.898455 | 0.446 | 0.155 | 6.5E-11 |
| Hpn | 1.25E-14 | 2.535154 | 0.261 | 0.063 | 2.95E-10 |
| Nfkbiz | 1.29E-14 | 3.246476 | 0.315 | 0.088 | 3.03E-10 |
| H2-D1 | 2.48E-14 | 1.517266 | 0.967 | 0.867 | 5.85E-10 |
| Sdc4 | 3.9E-14 | 2.888679 | 0.457 | 0.175 | 9.19E-10 |
| Rab20 | 1.26E-13 | 2.418571 | 0.435 | 0.169 | 2.98E-09 |
| Tnfaip3 | 2.33E-13 | 2.241723 | 0.293 | 0.08 | 5.5E-09 |
| C5ar1 | 5.33E-13 | 2.047568 | 0.663 | 0.426 | 1.26E-08 |
| Ccl3 | 6.16E-13 | 4.171485 | 0.207 | 0.047 | 1.45E-08 |
| Ccrl2 | 1.58E-12 | 2.52569 | 0.217 | 0.052 | 3.72E-08 |
| Hcar2 | 1.77E-12 | 3.681302 | 0.217 | 0.053 | 4.17E-08 |
| Scoc | 7.15E-12 | -1.58579 | 0.598 | 0.831 | 1.69E-07 |
| Rnf19b | 1.02E-11 | 1.510619 | 0.522 | 0.234 | 2.4E-07 |
| Bcl6 | 2.49E-11 | 1.837739 | 0.326 | 0.111 | 5.88E-07 |
| Ier3 | 2.89E-11 | 3.255592 | 0.217 | 0.057 | 6.82E-07 |

| | | | | | |
|---|---|---|---|---|---|
| Slc15a3 | 3.33E-11 | 1.732533 | 0.652 | 0.378 | 7.85E-07 |
| Plekho2 | 3.54E-11 | 1.910236 | 0.359 | 0.131 | 8.36E-07 |
| Rrad | 3.84E-11 | 2.589888 | 0.25 | 0.074 | 9.06E-07 |
| Gm26917 | 2.04E-10 | 1.304377 | 1 | 0.951 | 4.82E-06 |
| Plaur | 2.81E-10 | 2.08319 | 0.228 | 0.065 | 6.63E-06 |
| Slc2a5 | 3.14E-10 | -0.84575 | 0.674 | 0.944 | 7.41E-06 |
| Fth1 | 4.58E-10 | 0.811785 | 1 | 0.993 | 1.08E-05 |
| Arhgap5 | 5.28E-10 | -1.05878 | 0.457 | 0.761 | 1.25E-05 |
| Hivep3 | 9.34E-10 | 1.635007 | 0.696 | 0.419 | 2.2E-05 |
| Cebpb | 1.4E-09 | 1.411919 | 0.772 | 0.529 | 3.31E-05 |
| Itga5 | 1.92E-09 | 2.483611 | 0.25 | 0.081 | 4.54E-05 |
| Ifi30 | 2.01E-09 | 1.552632 | 0.739 | 0.468 | 4.75E-05 |
| Siglech | 2.28E-09 | -0.63907 | 0.978 | 0.999 | 5.38E-05 |
| Rtn4rl1 | 2.9E-09 | -1.05963 | 0.348 | 0.684 | 6.83E-05 |
| Cd164 | 3.03E-09 | -0.88582 | 0.772 | 0.946 | 7.16E-05 |
| Maf | 7.53E-09 | -1.14527 | 0.696 | 0.897 | 0.000178 |
| Nfkbib | 7.58E-09 | 1.824311 | 0.402 | 0.18 | 0.000179 |
| Mat2a | 9.96E-09 | 1.465524 | 0.511 | 0.255 | 0.000235 |
| Selplg | 1.07E-08 | -0.50422 | 1 | 1 | 0.000253 |
| Mthfd2 | 1.31E-08 | 1.937222 | 0.261 | 0.091 | 0.000309 |
| Cd9 | 1.39E-08 | 0.723925 | 0.978 | 0.965 | 0.000328 |
| Cdkn1a | 1.95E-08 | 3.316142 | 0.272 | 0.102 | 0.00046 |
| Lpar6 | 2.05E-08 | -1.2827 | 0.326 | 0.632 | 0.000485 |
| Mertk | 2.3E-08 | -1.17914 | 0.685 | 0.872 | 0.000544 |
| Glul | 2.97E-08 | -0.89728 | 0.826 | 0.938 | 0.0007 |
| Ptgs1 | 3.27E-08 | -0.8576 | 0.826 | 0.946 | 0.00077 |
| Vasp | 4.27E-08 | 1.524561 | 0.63 | 0.394 | 0.001006 |
| Ncoa3 | 4.67E-08 | -1.71597 | 0.587 | 0.759 | 0.001101 |
| Mcl1 | 5.17E-08 | 1.612591 | 0.707 | 0.513 | 0.001221 |
| P2ry13 | 5.56E-08 | -0.85742 | 0.902 | 0.977 | 0.001312 |
| Tagln2 | 6.38E-08 | 1.729472 | 0.435 | 0.222 | 0.001504 |
| Tnfaip8l2 | 7.56E-08 | -1.33097 | 0.348 | 0.62 | 0.001784 |
| Vsir | 9.4E-08 | -0.57732 | 0.967 | 0.985 | 0.002217 |
| Apobec3 | 9.58E-08 | 1.450031 | 0.707 | 0.49 | 0.00226 |
| Slfn2 | 1.84E-07 | 1.565491 | 0.293 | 0.122 | 0.004346 |
| Nupr1 | 2.47E-07 | 1.886546 | 0.304 | 0.129 | 0.005829 |
| Gm43305 | 3.12E-07 | 2.112398 | 0.75 | 0.591 | 0.007368 |
| Ctsz | 3.96E-07 | 0.678964 | 1 | 0.993 | 0.009348 |
| Stk38l | 4.17E-07 | 2.175914 | 0.217 | 0.079 | 0.009842 |
| Cep85 | 4.38E-07 | 1.545327 | 0.717 | 0.51 | 0.010327 |
| C1qa | 4.66E-07 | 0.466745 | 1 | 1 | 0.010999 |
| Cpd | 5.04E-07 | 1.605602 | 0.228 | 0.082 | 0.011896 |
| Rgs2 | 5.54E-07 | -0.95637 | 0.696 | 0.841 | 0.013061 |
| Mapkapk2 | 6.03E-07 | 1.652413 | 0.435 | 0.228 | 0.014218 |
| Mvp | 7.27E-07 | 1.402729 | 0.37 | 0.172 | 0.01715 |
| Rassf4 | 7.51E-07 | 1.48415 | 0.543 | 0.34 | 0.017707 |
| Bin2 | 8.47E-07 | -0.71818 | 0.761 | 0.909 | 0.019975 |

| | | | | | |
|---|---|---|---|---|---|
| Tank | 8.64E-07 | 1.162629 | 0.261 | 0.102 | 0.020387 |
| Tnip1 | 1.12E-06 | 1.651648 | 0.207 | 0.073 | 0.026488 |
| Selenop | 1.13E-06 | -0.67703 | 0.946 | 0.991 | 0.026737 |
| Pea15a | 1.33E-06 | 1.286989 | 0.62 | 0.403 | 0.031446 |
| Calm2 | 1.35E-06 | -0.80803 | 0.902 | 0.953 | 0.031934 |
| AI506816 | 1.51E-06 | 0.93401 | 0.478 | 0.254 | 0.035728 |
| C3ar1 | 1.54E-06 | 1.109106 | 0.75 | 0.553 | 0.036384 |
| Il1a | 1.66E-06 | 1.668128 | 0.402 | 0.21 | 0.039154 |
| B2m | 1.86E-06 | 0.564121 | 1 | 0.998 | 0.043825 |
| Golm1 | 1.96E-06 | -0.94872 | 0.783 | 0.877 | 0.04634 |

#### Supplementry Table 2. MHCIIM DEGs compared to HS

| GeneName | p_val | avg_log2FC | pct.1 | pct.2 | p_val_adj |
|---|---|---|---|---|---|
| Cox6a2 | 3.8E-102 | 4.150708 | 0.212 | 0.016 | 8.97E-98 |
| Chst2 | 1.41E-96 | 3.312582 | 0.253 | 0.026 | 3.33E-92 |
| Capg | 4.88E-75 | 3.233228 | 0.516 | 0.153 | 1.15E-70 |
| Fcrls | 1.79E-71 | -1.50772 | 0.753 | 0.975 | 4.22E-67 |
| Ccl6 | 2.52E-68 | 2.42048 | 0.792 | 0.421 | 5.94E-64 |
| C1qb | 4.62E-67 | 0.93352 | 1 | 1 | 1.09E-62 |
| H2-D1 | 7.92E-64 | 1.582777 | 0.981 | 0.867 | 1.87E-59 |
| Siglech | 1.54E-61 | 1.125863 | 0.994 | 0.999 | 3.64E-57 |
| Haao | 1.23E-60 | 3.081556 | 0.24 | 0.039 | 2.89E-56 |
| C1qa | 2.04E-59 | 0.872235 | 1 | 1 | 4.82E-55 |
| Cd74 | 5.2E-59 | 5.446729 | 0.458 | 0.153 | 1.23E-54 |
| Ptgs1 | 1.16E-56 | -1.59244 | 0.644 | 0.946 | 2.74E-52 |
| Rad51b | 2.38E-52 | 3.098534 | 0.212 | 0.034 | 5.63E-48 |
| Ly6e | 3.24E-49 | 1.374974 | 0.99 | 0.948 | 7.65E-45 |
| Mafb | 3.91E-49 | -1.54339 | 0.606 | 0.913 | 9.23E-45 |
| Lsp1 | 6.61E-49 | 2.006979 | 0.635 | 0.284 | 1.56E-44 |
| H2-Ab1 | 1.07E-48 | 4.379906 | 0.426 | 0.154 | 2.53E-44 |
| Crlf2 | 1.28E-47 | 1.693934 | 0.718 | 0.372 | 3.03E-43 |
| Rnase6 | 1.76E-46 | 2.268126 | 0.369 | 0.108 | 4.15E-42 |
| Sirpa | 1.49E-44 | -1.37615 | 0.808 | 0.958 | 3.51E-40 |
| Ctsd | 2.19E-41 | 0.733719 | 1 | 1 | 5.17E-37 |
| Gapdh | 4.72E-41 | 1.176667 | 0.965 | 0.904 | 1.11E-36 |
| Serinc3 | 8.57E-40 | -1.00808 | 0.946 | 0.996 | 2.02E-35 |
| H2-K1 | 1.07E-35 | 1.454611 | 0.901 | 0.766 | 2.52E-31 |
| Abhd12 | 3.52E-35 | -0.98464 | 0.869 | 0.977 | 8.29E-31 |
| Abca1 | 2.09E-33 | -2.25966 | 0.196 | 0.556 | 4.94E-29 |
| Ctss | 1.1E-32 | 0.637279 | 1 | 1 | 2.61E-28 |
| Cd52 | 9.68E-30 | 1.688261 | 0.66 | 0.411 | 2.28E-25 |
| Rrbp1 | 1.6E-29 | -1.0713 | 0.75 | 0.93 | 3.77E-25 |
| Nupr1 | 4.94E-28 | 2.21413 | 0.337 | 0.129 | 1.17E-23 |
| Frmd4a | 5.87E-28 | -1.3541 | 0.455 | 0.754 | 1.39E-23 |
| Mertk | 1.93E-27 | -1.09575 | 0.609 | 0.872 | 4.56E-23 |
| Slc2a5 | 2.81E-27 | 0.803957 | 0.971 | 0.944 | 6.62E-23 |
| Fcer1g | 4.21E-27 | 0.653079 | 1 | 0.993 | 9.93E-23 |
| Ly86 | 2.03E-26 | 0.643136 | 0.99 | 0.991 | 4.78E-22 |
| Cst3 | 2.62E-26 | 0.478724 | 1 | 1 | 6.19E-22 |
| Ecscr | 3.06E-26 | -1.10556 | 0.391 | 0.706 | 7.23E-22 |
| Surf4 | 3.36E-25 | -1.52086 | 0.311 | 0.618 | 7.93E-21 |
| Pmepa1 | 1.24E-24 | -0.89809 | 0.596 | 0.845 | 2.91E-20 |
| Spint1 | 1.71E-24 | -2.30488 | 0.208 | 0.487 | 4.02E-20 |
| Ctsa | 2.13E-24 | -0.74205 | 0.856 | 0.972 | 5.03E-20 |
| Srgn | 2.34E-24 | 1.293802 | 0.696 | 0.457 | 5.51E-20 |
| Tmsb4x | 3.05E-24 | 0.593583 | 1 | 1 | 7.2E-20 |

| | | | | | |
|---|---|---|---|---|---|
| Stab1 | 6.18E-24 | -1.05552 | 0.558 | 0.822 | 1.46E-19 |
| Bsg | 8.52E-24 | 0.831313 | 0.939 | 0.859 | 2.01E-19 |
| Acp2 | 1.1E-23 | -1.02308 | 0.333 | 0.633 | 2.6E-19 |
| Laptm5 | 7.56E-23 | 0.521748 | 1 | 1 | 1.78E-18 |
| C1qc | 9.4E-23 | 0.520617 | 1 | 1 | 2.22E-18 |
| Itpripl1 | 1.29E-22 | -1.22804 | 0.423 | 0.729 | 3.05E-18 |
| Fth1 | 1.51E-22 | 0.68168 | 0.997 | 0.993 | 3.57E-18 |
| Ppia | 1.8E-22 | 0.625537 | 0.987 | 0.969 | 4.24E-18 |
| Sparc | 1.3E-21 | 0.513378 | 0.997 | 1 | 3.07E-17 |
| Syngr1 | 1.75E-21 | 0.946299 | 0.901 | 0.839 | 4.12E-17 |
| Hck | 5.58E-21 | -1.29663 | 0.333 | 0.613 | 1.32E-16 |
| Tspan7 | 8.33E-21 | 1.156403 | 0.712 | 0.532 | 1.97E-16 |
| Ank | 1.14E-20 | 1.616582 | 0.253 | 0.093 | 2.7E-16 |
| Pnpla7 | 4.3E-20 | -1.2411 | 0.282 | 0.552 | 1.01E-15 |
| Rnaset2a | 7.59E-20 | 0.70606 | 0.968 | 0.948 | 1.79E-15 |
| Scamp5 | 1.39E-19 | -0.96041 | 0.295 | 0.578 | 3.28E-15 |
| Lpar6 | 1.78E-19 | -1.18933 | 0.381 | 0.632 | 4.19E-15 |
| C3ar1 | 1.81E-19 | -1.29376 | 0.279 | 0.553 | 4.26E-15 |
| Lyz2 | 4.06E-19 | -1.35799 | 0.359 | 0.608 | 9.57E-15 |
| Rab14 | 5.78E-19 | -0.75064 | 0.526 | 0.794 | 1.36E-14 |
| Ctsz | 8.85E-19 | -0.58874 | 0.958 | 0.993 | 2.09E-14 |
| Slc40a1 | 1.02E-18 | -1.67323 | 0.276 | 0.522 | 2.41E-14 |
| Cd81 | 1.18E-18 | 0.509911 | 1 | 0.999 | 2.78E-14 |
| Mpc1 | 1.67E-18 | 0.942382 | 0.708 | 0.498 | 3.94E-14 |
| Rps14 | 2.1E-18 | 0.657686 | 0.939 | 0.914 | 4.95E-14 |
| Pkm | 3.78E-18 | 1.168199 | 0.724 | 0.559 | 8.91E-14 |
| Tyrobp | 5.1E-18 | 0.581039 | 0.994 | 0.996 | 1.2E-13 |
| Ighm | 5.25E-18 | 0.885756 | 0.817 | 0.666 | 1.24E-13 |
| Gm10076 | 6.39E-18 | 0.748257 | 0.949 | 0.893 | 1.51E-13 |
| Itm2b | 6.7E-18 | 0.402589 | 1 | 1 | 1.58E-13 |
| Sdf4 | 8.35E-18 | 1.088053 | 0.728 | 0.568 | 1.97E-13 |
| Syngr2 | 9.73E-18 | 0.807393 | 0.901 | 0.819 | 2.29E-13 |
| Ptpra | 1.55E-17 | -0.97982 | 0.369 | 0.621 | 3.66E-13 |
| Rbm39 | 2.46E-17 | -0.77402 | 0.734 | 0.9 | 5.81E-13 |
| Prex1 | 2.87E-17 | -1.00207 | 0.276 | 0.539 | 6.77E-13 |
| Selenop | 3.59E-17 | -0.63581 | 0.981 | 0.991 | 8.46E-13 |
| Rpl41 | 5.73E-17 | 0.639468 | 0.968 | 0.934 | 1.35E-12 |
| Tgfbi | 6.25E-17 | 1.531956 | 0.401 | 0.217 | 1.47E-12 |
| Trem2 | 8.41E-17 | 0.58553 | 0.994 | 0.988 | 1.98E-12 |
| Plxdc2 | 1.26E-16 | -0.73559 | 0.856 | 0.931 | 2.97E-12 |
| Snx5 | 1.3E-16 | -0.86815 | 0.619 | 0.81 | 3.07E-12 |
| Bmyc | 1.42E-16 | -1.26156 | 0.256 | 0.502 | 3.35E-12 |
| Gm5559 | 2.33E-16 | 0.610031 | 0.885 | 0.717 | 5.49E-12 |
| Lat2 | 2.64E-16 | 1.062616 | 0.548 | 0.358 | 6.23E-12 |
| Ptpn1 | 2.93E-16 | -1.10499 | 0.394 | 0.63 | 6.9E-12 |
| H2-Oa | 4.71E-16 | 1.646289 | 0.353 | 0.182 | 1.11E-11 |
| Tmem176b | 5.56E-16 | 0.820284 | 0.862 | 0.794 | 1.31E-11 |

| | | | | | |
|---|---|---|---|---|---|
| Lpcat2 | 6.81E-16 | -0.55951 | 0.92 | 0.984 | 1.61E-11 |
| Ccr5 | 9.57E-16 | -1.25214 | 0.526 | 0.717 | 2.26E-11 |
| Soga1 | 1.04E-15 | -0.90593 | 0.337 | 0.586 | 2.45E-11 |
| Rgs19 | 1.4E-15 | -1.15837 | 0.263 | 0.506 | 3.3E-11 |
| Cd14 | 5.83E-15 | -0.75467 | 0.561 | 0.772 | 1.38E-10 |
| Ccl9 | 8.02E-15 | 1.178863 | 0.545 | 0.374 | 1.89E-10 |
| Crybb1 | 9.67E-15 | -0.65931 | 0.564 | 0.794 | 2.28E-10 |
| Gm10359 | 1.44E-14 | 0.434546 | 0.49 | 0.303 | 3.4E-10 |
| Siglece | 2.18E-14 | -1.58573 | 0.16 | 0.372 | 5.14E-10 |
| Mpc1-ps | 2.4E-14 | 0.673333 | 0.628 | 0.438 | 5.67E-10 |
| Gm7336 | 3.02E-14 | 0.532316 | 0.532 | 0.326 | 7.12E-10 |
| Pafah1b3 | 5.25E-14 | 1.642428 | 0.266 | 0.126 | 1.24E-09 |
| Gm49369 | 7.39E-14 | 0.629187 | 0.84 | 0.795 | 1.74E-09 |
| Rps27rt | 9.02E-14 | 0.60905 | 0.817 | 0.687 | 2.13E-09 |
| Gm2574 | 1.17E-13 | 0.570256 | 0.593 | 0.43 | 2.76E-09 |
| Tspo | 1.19E-13 | 1.079901 | 0.449 | 0.274 | 2.81E-09 |
| Hfe | 1.3E-13 | -1.00934 | 0.234 | 0.452 | 3.07E-09 |
| Stard9 | 1.4E-13 | -1.4461 | 0.112 | 0.311 | 3.3E-09 |
| Glul | 1.49E-13 | -0.53694 | 0.859 | 0.938 | 3.52E-09 |
| Fnbp1 | 1.59E-13 | -1.19696 | 0.279 | 0.501 | 3.76E-09 |
| Rpsa | 1.68E-13 | 0.692679 | 0.869 | 0.774 | 3.97E-09 |
| AW112010 | 1.69E-13 | 1.806284 | 0.228 | 0.1 | 3.98E-09 |
| St6galnac4 | 1.86E-13 | -1.12792 | 0.256 | 0.467 | 4.38E-09 |
| Ptprj | 1.86E-13 | -0.76377 | 0.324 | 0.556 | 4.39E-09 |
| Il10rb | 2.05E-13 | -0.95416 | 0.388 | 0.597 | 4.84E-09 |
| Npc2 | 2.35E-13 | 0.683182 | 0.885 | 0.82 | 5.55E-09 |
| Notch1 | 2.42E-13 | -1.65387 | 0.135 | 0.334 | 5.72E-09 |
| Lrrc25 | 3.21E-13 | -0.95092 | 0.231 | 0.449 | 7.58E-09 |
| Rin2 | 3.3E-13 | -1.20334 | 0.343 | 0.55 | 7.79E-09 |
| Fads1 | 3.44E-13 | -1.74799 | 0.096 | 0.282 | 8.11E-09 |
| Gm12537 | 3.56E-13 | 0.429548 | 0.478 | 0.291 | 8.39E-09 |
| Cyb561a3 | 3.59E-13 | 0.800224 | 0.663 | 0.487 | 8.47E-09 |
| Ly6a | 3.69E-13 | 2.829571 | 0.247 | 0.121 | 8.69E-09 |
| Hspa5 | 3.73E-13 | -0.70758 | 0.728 | 0.866 | 8.79E-09 |
| Sept11 | 4.88E-13 | 1.71304 | 0.221 | 0.098 | 1.15E-08 |
| Pdia3 | 5.58E-13 | -0.49531 | 0.888 | 0.961 | 1.32E-08 |
| Edem2 | 6.18E-13 | -0.87769 | 0.369 | 0.586 | 1.46E-08 |
| Apbb1ip | 6.2E-13 | -0.75307 | 0.554 | 0.749 | 1.46E-08 |
| Rps11 | 1.02E-12 | 0.549245 | 0.949 | 0.911 | 2.4E-08 |
| Rpl21 | 1.5E-12 | 0.531908 | 0.929 | 0.917 | 3.53E-08 |
| Tnfsf13b | 1.83E-12 | 1.195382 | 0.311 | 0.162 | 4.32E-08 |
| Ehd4 | 2.12E-12 | -0.89554 | 0.288 | 0.496 | 5E-08 |
| Tanc2 | 2.29E-12 | -0.71323 | 0.686 | 0.831 | 5.4E-08 |
| Lgals3bp | 2.62E-12 | 1.114446 | 0.638 | 0.503 | 6.18E-08 |
| Tm6sf1 | 3.01E-12 | -1.25773 | 0.263 | 0.459 | 7.1E-08 |
| Rps27 | 3.14E-12 | 0.569035 | 0.958 | 0.95 | 7.4E-08 |
| Selenow | 3.47E-12 | 1.48762 | 0.304 | 0.167 | 8.18E-08 |

| | | | | | |
|---|---|---|---|---|---|
| Clic1 | 3.86E-12 | 0.551269 | 0.891 | 0.854 | 9.11E-08 |
| Hpgd | 4.1E-12 | -0.81569 | 0.606 | 0.746 | 9.67E-08 |
| Rpl3 | 4.77E-12 | 0.630955 | 0.897 | 0.821 | 1.12E-07 |
| Cd82 | 4.94E-12 | -0.8309 | 0.388 | 0.595 | 1.17E-07 |
| Fam110a | 5.26E-12 | -1.33936 | 0.176 | 0.364 | 1.24E-07 |
| Actb | 6.86E-12 | 0.417527 | 1 | 1 | 1.62E-07 |
| Kcnk6 | 7.81E-12 | -0.78671 | 0.333 | 0.554 | 1.84E-07 |
| Igsf6 | 9.84E-12 | -3.25955 | 0.048 | 0.202 | 2.32E-07 |
| Sox4 | 9.87E-12 | -1.25325 | 0.24 | 0.422 | 2.33E-07 |
| Rnase4 | 1.04E-11 | 0.523635 | 0.962 | 0.955 | 2.45E-07 |
| Pkig | 1.12E-11 | -0.90145 | 0.205 | 0.412 | 2.63E-07 |
| Epb41l2 | 1.21E-11 | -0.62829 | 0.779 | 0.89 | 2.86E-07 |
| Abl1 | 1.22E-11 | -0.82552 | 0.131 | 0.327 | 2.88E-07 |
| Ptafr | 1.29E-11 | -0.75062 | 0.609 | 0.783 | 3.05E-07 |
| Lmo2 | 1.77E-11 | -0.783 | 0.433 | 0.632 | 4.18E-07 |
| Ntpcr | 2.32E-11 | -0.59575 | 0.426 | 0.634 | 5.48E-07 |
| Apoe | 2.53E-11 | 3.233313 | 0.481 | 0.376 | 5.97E-07 |
| Nfe2l2 | 2.55E-11 | -0.90117 | 0.458 | 0.643 | 6.02E-07 |
| Cd300a | 2.63E-11 | -0.89933 | 0.381 | 0.579 | 6.21E-07 |
| Rpn2 | 2.7E-11 | -0.75421 | 0.551 | 0.735 | 6.37E-07 |
| Abcc3 | 3.4E-11 | -0.7215 | 0.228 | 0.427 | 8.03E-07 |
| Set | 3.74E-11 | -0.81814 | 0.429 | 0.619 | 8.81E-07 |
| Cep152 | 4.83E-11 | -2.23233 | 0.067 | 0.221 | 1.14E-06 |
| Hexa | 5.77E-11 | 0.452276 | 0.974 | 0.972 | 1.36E-06 |
| Cx3cr1 | 5.85E-11 | -0.30284 | 1 | 1 | 1.38E-06 |
| Tspan3 | 6.06E-11 | 0.871533 | 0.628 | 0.483 | 1.43E-06 |
| Tmem250- | 6.44E-11 | -1.08211 | 0.356 | 0.541 | 1.52E-06 |
| Ubb | 6.48E-11 | 0.483249 | 0.984 | 0.976 | 1.53E-06 |
| Snta1 | 6.88E-11 | -1.51624 | 0.144 | 0.319 | 1.62E-06 |
| Mafg | 7.01E-11 | -1.12328 | 0.34 | 0.525 | 1.65E-06 |
| Fkbp2 | 7.03E-11 | 0.881896 | 0.587 | 0.426 | 1.66E-06 |
| Bcl2l1 | 7.5E-11 | -1.04548 | 0.228 | 0.415 | 1.77E-06 |
| Selplg | 9.45E-11 | 0.34326 | 1 | 1 | 2.23E-06 |
| Rpl13a | 9.91E-11 | 0.664653 | 0.881 | 0.824 | 2.34E-06 |
| Arhgdib | 1.07E-10 | 0.766324 | 0.817 | 0.75 | 2.52E-06 |
| Hnrnpc | 1.09E-10 | 0.877017 | 0.638 | 0.497 | 2.58E-06 |
| Serpine2 | 1.29E-10 | 0.516451 | 0.929 | 0.91 | 3.04E-06 |
| Rpl19 | 1.34E-10 | 0.649712 | 0.878 | 0.794 | 3.15E-06 |
| Tmem104 | 1.36E-10 | -0.43261 | 0.196 | 0.387 | 3.21E-06 |
| Ggta1 | 1.42E-10 | -1.86974 | 0.058 | 0.206 | 3.34E-06 |
| Zmynd8 | 1.43E-10 | -1.04598 | 0.221 | 0.406 | 3.37E-06 |
| Rpl13 | 1.44E-10 | 0.446985 | 0.968 | 0.938 | 3.39E-06 |
| Sppl2a | 1.5E-10 | -0.94149 | 0.417 | 0.589 | 3.55E-06 |
| Sdf2l1 | 1.64E-10 | 1.087342 | 0.567 | 0.458 | 3.87E-06 |
| Tgm2 | 1.64E-10 | -1.56442 | 0.106 | 0.27 | 3.87E-06 |
| 1700017B0 | 1.66E-10 | -0.84507 | 0.519 | 0.694 | 3.91E-06 |
| Ifi30 | 1.84E-10 | -0.92508 | 0.282 | 0.468 | 4.35E-06 |

| | | | | | |
|---|---|---|---|---|---|
| Ctsf | 1.94E-10 | -0.72154 | 0.487 | 0.666 | 4.57E-06 |
| Acss1 | 2.02E-10 | -1.78832 | 0.09 | 0.247 | 4.78E-06 |
| Trim47 | 2.31E-10 | -2.69024 | 0.058 | 0.201 | 5.46E-06 |
| Dido1 | 2.34E-10 | -1.83522 | 0.103 | 0.262 | 5.52E-06 |
| Itga6 | 2.51E-10 | -0.94394 | 0.426 | 0.591 | 5.91E-06 |
| Abi1 | 3.08E-10 | -1.32086 | 0.212 | 0.383 | 7.26E-06 |
| Ube2e3 | 3.12E-10 | -1.38475 | 0.112 | 0.274 | 7.36E-06 |
| Cd34 | 3.43E-10 | -1.07689 | 0.353 | 0.51 | 8.1E-06 |
| Brd3 | 3.57E-10 | -2.02646 | 0.138 | 0.298 | 8.41E-06 |
| Slc11a1 | 4.22E-10 | -0.64565 | 0.391 | 0.595 | 9.95E-06 |
| Slc35c2 | 5E-10 | -0.8582 | 0.282 | 0.464 | 1.18E-05 |
| Asah1 | 5.24E-10 | -0.64542 | 0.753 | 0.845 | 1.24E-05 |
| Bri3 | 5.29E-10 | 0.791826 | 0.673 | 0.573 | 1.25E-05 |
| Eng | 5.47E-10 | -0.91353 | 0.349 | 0.519 | 1.29E-05 |
| Tpd52l2 | 5.62E-10 | -1.29053 | 0.115 | 0.278 | 1.33E-05 |
| Otulinl | 5.79E-10 | -0.55816 | 0.862 | 0.93 | 1.37E-05 |
| Fam102b | 5.92E-10 | -0.69763 | 0.478 | 0.665 | 1.4E-05 |
| Slc12a6 | 5.97E-10 | -0.89218 | 0.176 | 0.351 | 1.41E-05 |
| Gpr137b | 6.37E-10 | -1.65558 | 0.096 | 0.249 | 1.5E-05 |
| Il17ra | 6.97E-10 | -0.97225 | 0.317 | 0.51 | 1.64E-05 |
| Csnk2a1 | 7.13E-10 | -1.20348 | 0.324 | 0.494 | 1.68E-05 |
| Cd63 | 7.47E-10 | 0.6886 | 0.782 | 0.684 | 1.76E-05 |
| Ramp1 | 8.41E-10 | 1.579104 | 0.308 | 0.182 | 1.98E-05 |
| Laptm4b | 8.78E-10 | 1.280694 | 0.327 | 0.201 | 2.07E-05 |
| Pip4k2a | 8.86E-10 | -0.67179 | 0.404 | 0.579 | 2.09E-05 |
| Ssr4 | 9.06E-10 | 0.804667 | 0.715 | 0.626 | 2.14E-05 |
| Gm2606 | 9.55E-10 | 0.44605 | 0.269 | 0.145 | 2.25E-05 |
| Tmem50a | 9.74E-10 | 0.504918 | 0.917 | 0.863 | 2.3E-05 |
| Rplp1 | 1.16E-09 | 0.439709 | 0.958 | 0.952 | 2.73E-05 |
| Eef1a1 | 1.16E-09 | 0.44614 | 0.994 | 0.994 | 2.74E-05 |
| Rpl36 | 1.18E-09 | 0.499436 | 0.869 | 0.799 | 2.79E-05 |
| Rpl37 | 1.19E-09 | 0.452816 | 0.942 | 0.886 | 2.8E-05 |
| Gm5514 | 1.3E-09 | -0.47987 | 0.337 | 0.516 | 3.08E-05 |
| Tpst2 | 1.32E-09 | -0.70265 | 0.535 | 0.701 | 3.11E-05 |
| Rps3a1 | 1.38E-09 | 0.559965 | 0.891 | 0.837 | 3.26E-05 |
| Tmem63a | 1.41E-09 | -0.85554 | 0.292 | 0.469 | 3.34E-05 |
| Rgl2 | 1.65E-09 | -0.56252 | 0.256 | 0.443 | 3.88E-05 |
| Tmed3 | 1.7E-09 | 0.769081 | 0.619 | 0.497 | 4.02E-05 |
| Tpi1 | 1.9E-09 | 1.202609 | 0.413 | 0.284 | 4.47E-05 |
| Rassf2 | 1.99E-09 | -0.71022 | 0.24 | 0.423 | 4.68E-05 |
| Med22 | 2.38E-09 | -1.56153 | 0.083 | 0.229 | 5.62E-05 |
| Rpl7a | 2.72E-09 | -0.6539 | 0.526 | 0.691 | 6.41E-05 |
| Agps | 2.72E-09 | -1.45958 | 0.17 | 0.331 | 6.42E-05 |
| Ndufa1 | 2.86E-09 | 0.995531 | 0.404 | 0.275 | 6.74E-05 |
| Myl6 | 2.95E-09 | 0.72392 | 0.705 | 0.602 | 6.96E-05 |
| Dip2b | 3.12E-09 | -0.973 | 0.452 | 0.603 | 7.35E-05 |
| Pfn1 | 3.13E-09 | 0.383144 | 0.974 | 0.958 | 7.38E-05 |

| | | | | | |
|---|---|---|---|---|---|
| Rplp2 | 3.27E-09 | 0.4985 | 0.92 | 0.895 | 7.72E-05 |
| Plod3 | 3.35E-09 | -0.59759 | 0.391 | 0.575 | 7.89E-05 |
| Map1s | 3.45E-09 | -1.16481 | 0.163 | 0.324 | 8.13E-05 |
| Secisbp2l | 3.64E-09 | -0.80936 | 0.186 | 0.351 | 8.58E-05 |
| Ctsh | 3.67E-09 | 0.509022 | 0.949 | 0.947 | 8.65E-05 |
| Tor1b | 3.79E-09 | -1.20807 | 0.167 | 0.325 | 8.94E-05 |
| Camsap1 | 3.83E-09 | -1.32013 | 0.067 | 0.204 | 9.03E-05 |
| Rps23 | 4.12E-09 | 0.489068 | 0.917 | 0.887 | 9.71E-05 |
| Trpc4ap | 4.24E-09 | -0.80325 | 0.304 | 0.481 | 0.0001 |
| P2ry6 | 4.36E-09 | -0.51741 | 0.644 | 0.812 | 0.000103 |
| Camk2n1 | 4.57E-09 | -0.93769 | 0.138 | 0.294 | 0.000108 |
| Tagap | 4.66E-09 | -0.77845 | 0.208 | 0.376 | 0.00011 |
| Gpx1 | 4.79E-09 | 0.749928 | 0.785 | 0.708 | 0.000113 |
| Itgav | 4.91E-09 | -0.69949 | 0.221 | 0.387 | 0.000116 |
| Gm6560 | 5.11E-09 | 0.463489 | 0.458 | 0.306 | 0.00012 |
| Dennd4a | 5.41E-09 | -0.95682 | 0.404 | 0.558 | 0.000128 |
| Dnajc5 | 6.06E-09 | -0.95773 | 0.202 | 0.364 | 0.000143 |
| Rpl32 | 6.2E-09 | 0.545337 | 0.891 | 0.863 | 0.000146 |
| Csnk1e | 6.51E-09 | -0.74459 | 0.468 | 0.622 | 0.000154 |
| Stx16 | 6.55E-09 | -1.07823 | 0.154 | 0.312 | 0.000155 |
| Camk1d | 6.87E-09 | -1.07051 | 0.131 | 0.282 | 0.000162 |
| Sqor | 6.87E-09 | -0.97612 | 0.16 | 0.319 | 0.000162 |
| Cd302 | 7.05E-09 | -0.75411 | 0.292 | 0.469 | 0.000166 |
| Rpl9 | 7.24E-09 | 0.596677 | 0.817 | 0.75 | 0.000171 |
| Tmem160 | 8.27E-09 | 0.959781 | 0.446 | 0.311 | 0.000195 |
| Atp5c1 | 9.41E-09 | -0.74795 | 0.314 | 0.49 | 0.000222 |
| Rpl18 | 9.64E-09 | 0.521925 | 0.856 | 0.799 | 0.000227 |
| Cox4i1 | 9.9E-09 | 0.58182 | 0.833 | 0.775 | 0.000234 |
| Cebpb | 1.04E-08 | -0.48136 | 0.343 | 0.529 | 0.000245 |
| Rpl18a | 1.11E-08 | 0.499811 | 0.978 | 0.986 | 0.000262 |
| Sh3bgrl3 | 1.16E-08 | 0.569174 | 0.875 | 0.812 | 0.000274 |
| Rgmb | 1.21E-08 | -0.81934 | 0.295 | 0.461 | 0.000284 |
| Gnai2 | 1.27E-08 | 0.44163 | 0.965 | 0.954 | 0.000299 |
| Lrrc8a | 1.3E-08 | -0.85331 | 0.317 | 0.489 | 0.000307 |
| Gm10123 | 1.34E-08 | 0.431498 | 0.946 | 0.927 | 0.000316 |
| Cep68 | 1.4E-08 | -0.77282 | 0.163 | 0.32 | 0.00033 |
| Atp2c1 | 1.46E-08 | -0.74453 | 0.266 | 0.432 | 0.000345 |
| Dad1 | 1.54E-08 | 0.699712 | 0.683 | 0.592 | 0.000364 |
| Selenof | 1.59E-08 | 0.589089 | 0.737 | 0.643 | 0.000376 |
| Tmem204 | 1.61E-08 | -1.46563 | 0.09 | 0.225 | 0.000379 |
| Mvb12b | 1.68E-08 | -0.95091 | 0.157 | 0.308 | 0.000397 |
| Api5 | 1.71E-08 | -1.97935 | 0.08 | 0.21 | 0.000402 |
| Cbfa2t3 | 1.89E-08 | 0.921604 | 0.519 | 0.375 | 0.000447 |
| Slc12a9 | 1.94E-08 | -1.03585 | 0.308 | 0.463 | 0.000458 |
| Mpi | 1.95E-08 | -0.96639 | 0.151 | 0.298 | 0.00046 |
| Dgkz | 2.04E-08 | -1.41825 | 0.128 | 0.265 | 0.000481 |
| Cd164 | 2.52E-08 | 0.5774 | 0.946 | 0.946 | 0.000594 |

| | | | | | |
|---|---|---|---|---|---|
| Vav2 | 2.55E-08 | -1.48055 | 0.093 | 0.231 | 0.000601 |
| Rpl4 | 2.7E-08 | 0.532647 | 0.878 | 0.829 | 0.000636 |
| Litaf | 2.78E-08 | -0.83982 | 0.324 | 0.484 | 0.000656 |
| Usp8 | 2.86E-08 | -0.41652 | 0.16 | 0.318 | 0.000674 |
| Gatm | 3.05E-08 | -1.45271 | 0.115 | 0.252 | 0.00072 |
| Vkorc1 | 3.07E-08 | 0.868412 | 0.513 | 0.388 | 0.000724 |
| Plagl2 | 3.07E-08 | -1.46361 | 0.103 | 0.237 | 0.000724 |
| Mgat1 | 3.22E-08 | 0.853789 | 0.67 | 0.574 | 0.000759 |
| Man1c1 | 3.28E-08 | -0.64634 | 0.349 | 0.521 | 0.000774 |
| Clec4a2 | 3.29E-08 | -0.75615 | 0.186 | 0.337 | 0.000775 |
| Rpl37a | 3.61E-08 | 0.298021 | 0.968 | 0.929 | 0.000851 |
| Zswim6 | 3.63E-08 | -1.40114 | 0.103 | 0.237 | 0.000855 |
| Ncf1 | 3.75E-08 | -0.45333 | 0.567 | 0.735 | 0.000886 |
| Rps9 | 4.06E-08 | 0.420529 | 0.971 | 0.95 | 0.000959 |
| Cds2 | 4.25E-08 | -1.25824 | 0.154 | 0.302 | 0.001003 |
| Slc16a3 | 4.26E-08 | 1.18176 | 0.308 | 0.193 | 0.001005 |
| Gm4654 | 4.33E-08 | -0.2761 | 0.269 | 0.153 | 0.001022 |
| Rpl36-ps2 | 4.69E-08 | 0.694618 | 0.622 | 0.478 | 0.001105 |
| Map3k3 | 5.03E-08 | -0.64257 | 0.228 | 0.384 | 0.001185 |
| Dync1i2 | 5.2E-08 | -0.65493 | 0.266 | 0.432 | 0.001226 |
| H13 | 5.53E-08 | -0.78935 | 0.247 | 0.4 | 0.001305 |
| Csad | 5.85E-08 | -0.69128 | 0.221 | 0.381 | 0.001381 |
| Gnpda1 | 5.93E-08 | -1.70766 | 0.08 | 0.205 | 0.001399 |
| Pdlim4 | 6.35E-08 | -0.57225 | 0.199 | 0.356 | 0.001499 |
| Slc35f6 | 7.09E-08 | -1.24918 | 0.125 | 0.264 | 0.001672 |
| Ssfa2 | 7.18E-08 | -1.00163 | 0.099 | 0.234 | 0.001694 |
| Mapre1 | 7.36E-08 | -0.67653 | 0.301 | 0.467 | 0.001735 |
| Dlgap4 | 7.7E-08 | -0.94587 | 0.224 | 0.372 | 0.001817 |
| Gm5138 | 8.23E-08 | 0.4063 | 0.599 | 0.455 | 0.00194 |
| Mif | 8.24E-08 | 0.803908 | 0.481 | 0.347 | 0.001943 |
| Gm10134 | 8.68E-08 | -1.17455 | 0.24 | 0.388 | 0.002046 |
| Vars | 8.86E-08 | 1.220048 | 0.292 | 0.176 | 0.002089 |
| Themis2 | 9.03E-08 | -0.70596 | 0.202 | 0.356 | 0.00213 |
| Gngt2 | 9.2E-08 | 0.647392 | 0.699 | 0.607 | 0.002169 |
| Ncoa3 | 9.71E-08 | -0.47881 | 0.599 | 0.759 | 0.002289 |
| Msl1 | 9.88E-08 | -0.86501 | 0.167 | 0.31 | 0.002331 |
| Spcs1 | 1.12E-07 | 0.577104 | 0.635 | 0.51 | 0.002643 |
| Vps18 | 1.17E-07 | -1.1607 | 0.135 | 0.271 | 0.002763 |
| Rapgef1 | 1.21E-07 | -0.71005 | 0.215 | 0.362 | 0.002849 |
| Elob | 1.22E-07 | 0.752594 | 0.59 | 0.47 | 0.002866 |
| Swi5 | 1.29E-07 | -0.81444 | 0.189 | 0.336 | 0.003036 |
| Oaz1 | 1.31E-07 | 0.526667 | 0.821 | 0.789 | 0.003092 |
| Eno1 | 1.32E-07 | 0.584395 | 0.772 | 0.695 | 0.003111 |
| Fau | 1.36E-07 | 0.49302 | 0.936 | 0.899 | 0.003219 |
| Scn1b | 1.39E-07 | -1.19652 | 0.179 | 0.316 | 0.003277 |
| Caprin1 | 1.4E-07 | -0.78494 | 0.173 | 0.312 | 0.003308 |
| Rps3 | 1.44E-07 | 0.441228 | 0.926 | 0.897 | 0.003407 |

| | | | | | |
|---|---|---|---|---|---|
| Cyb5r3 | 1.46E-07 | 1.212046 | 0.279 | 0.169 | 0.003454 |
| Rapgef6 | 1.48E-07 | -0.90012 | 0.157 | 0.303 | 0.003489 |
| Gp9 | 1.49E-07 | 1.032802 | 0.346 | 0.228 | 0.003506 |
| Arl4c | 1.54E-07 | -0.77977 | 0.337 | 0.486 | 0.003624 |
| Cox6b1 | 1.56E-07 | 0.621254 | 0.619 | 0.497 | 0.003674 |
| Hnrnpa1 | 1.57E-07 | 0.658752 | 0.413 | 0.28 | 0.003697 |
| Sec16a | 1.59E-07 | -1.34634 | 0.087 | 0.208 | 0.003761 |
| Arid3b | 1.65E-07 | -0.98037 | 0.128 | 0.264 | 0.003898 |
| Rpl34 | 1.66E-07 | 0.508673 | 0.837 | 0.778 | 0.003921 |
| Rps17 | 1.72E-07 | 0.514044 | 0.804 | 0.719 | 0.004061 |
| Ppm1h | 1.75E-07 | -0.6753 | 0.458 | 0.597 | 0.00413 |
| Rps25 | 1.81E-07 | 0.433664 | 0.881 | 0.852 | 0.004276 |
| Ctsb | 1.84E-07 | 0.322331 | 1 | 0.999 | 0.004335 |
| Capn3 | 2.12E-07 | -0.35777 | 0.26 | 0.422 | 0.004999 |
| Zfp120 | 2.16E-07 | -2.08981 | 0.083 | 0.201 | 0.005101 |
| Psmb1 | 2.44E-07 | 0.749729 | 0.587 | 0.464 | 0.005751 |
| Map1lc3a | 2.63E-07 | -0.93828 | 0.167 | 0.301 | 0.006197 |
| Fam217b | 2.67E-07 | -0.90363 | 0.103 | 0.227 | 0.006292 |
| Nfatc2 | 2.7E-07 | -1.05769 | 0.087 | 0.207 | 0.006361 |
| Atp5h | 2.73E-07 | 0.585598 | 0.571 | 0.442 | 0.00643 |
| Twf2 | 2.74E-07 | -0.48671 | 0.452 | 0.619 | 0.006453 |
| Cd37 | 2.85E-07 | 0.463197 | 0.897 | 0.877 | 0.006714 |
| Fgd2 | 2.85E-07 | -0.51892 | 0.606 | 0.747 | 0.006719 |
| Krtcap2 | 2.94E-07 | 0.814908 | 0.554 | 0.452 | 0.006946 |
| Sh2b3 | 3.03E-07 | -0.53812 | 0.298 | 0.449 | 0.007145 |
| Rplp0 | 3.05E-07 | 0.54969 | 0.897 | 0.89 | 0.007187 |
| Nucb1 | 3.22E-07 | 0.633938 | 0.683 | 0.58 | 0.007604 |
| Gpr137b-ps | 3.31E-07 | -1.14378 | 0.212 | 0.352 | 0.007799 |
| Grn | 3.33E-07 | -0.27402 | 0.978 | 0.995 | 0.007866 |
| Ppig | 3.46E-07 | -0.81653 | 0.154 | 0.291 | 0.008154 |
| H2afz | 3.85E-07 | 0.707786 | 0.615 | 0.501 | 0.009089 |
| Arap3 | 4.12E-07 | -1.05778 | 0.115 | 0.237 | 0.00972 |
| Gapvd1 | 4.28E-07 | -0.83459 | 0.228 | 0.365 | 0.010089 |
| Dcaf17 | 4.43E-07 | -1.22509 | 0.151 | 0.277 | 0.010455 |
| Uba52 | 4.55E-07 | 0.530311 | 0.798 | 0.721 | 0.010726 |
| Snrpb | 4.56E-07 | -0.63576 | 0.349 | 0.51 | 0.010764 |
| Tspan13 | 4.9E-07 | 0.689806 | 0.625 | 0.538 | 0.011565 |
| Tnfrsf21 | 5.26E-07 | -0.67133 | 0.381 | 0.513 | 0.012397 |
| Os9 | 5.62E-07 | 0.767292 | 0.574 | 0.467 | 0.013261 |
| Plcg1 | 5.87E-07 | -1.04582 | 0.087 | 0.204 | 0.013849 |
| Smox | 6.42E-07 | -0.75073 | 0.179 | 0.319 | 0.015136 |
| Klhdc8b | 6.46E-07 | 0.992138 | 0.301 | 0.196 | 0.015234 |
| Prkcd | 6.73E-07 | -0.65726 | 0.596 | 0.725 | 0.015886 |
| Mavs | 6.96E-07 | -2.13207 | 0.112 | 0.227 | 0.016427 |
| Rpl14 | 7.02E-07 | 0.546363 | 0.808 | 0.756 | 0.016556 |
| Rpl31 | 7.18E-07 | 0.544719 | 0.84 | 0.785 | 0.016942 |
| Stau1 | 7.22E-07 | -1.22584 | 0.189 | 0.318 | 0.017028 |

| | | | | | |
|---|---|---|---|---|---|
| Tmbim6 | 7.26E-07 | 0.380499 | 0.974 | 0.959 | 0.017123 |
| S100a13 | 7.43E-07 | 0.750275 | 0.253 | 0.152 | 0.017515 |
| Cdk9 | 7.53E-07 | -1.00023 | 0.202 | 0.332 | 0.017757 |
| Slc16a6 | 7.57E-07 | -0.7861 | 0.417 | 0.548 | 0.017855 |
| H2-T23 | 7.98E-07 | 0.554744 | 0.769 | 0.653 | 0.018817 |
| Prpf6 | 8E-07 | -1.56295 | 0.103 | 0.215 | 0.018864 |
| Il10ra | 8.09E-07 | -0.59872 | 0.631 | 0.741 | 0.019092 |
| Entpd1 | 8.57E-07 | -0.44504 | 0.862 | 0.94 | 0.020225 |
| Ggt5 | 8.66E-07 | -1.05502 | 0.138 | 0.263 | 0.020422 |
| Gm6204 | 8.98E-07 | 0.341159 | 0.702 | 0.582 | 0.021173 |
| Vps28 | 9.44E-07 | 0.761962 | 0.506 | 0.396 | 0.022259 |
| Gpr107 | 9.46E-07 | -1.32656 | 0.151 | 0.273 | 0.022314 |
| Fermt3 | 1.02E-06 | -0.60334 | 0.683 | 0.773 | 0.023972 |
| Tor4a | 1.07E-06 | -1.48191 | 0.147 | 0.262 | 0.02529 |
| Ssb | 1.09E-06 | -0.71305 | 0.304 | 0.447 | 0.025596 |
| Rps8 | 1.1E-06 | 0.463388 | 0.853 | 0.813 | 0.025952 |
| Dnase2a | 1.12E-06 | -0.63631 | 0.308 | 0.46 | 0.026322 |
| Ldhb | 1.12E-06 | -0.45959 | 0.667 | 0.809 | 0.026392 |
| Ndufs3 | 1.13E-06 | -0.69074 | 0.125 | 0.246 | 0.026584 |
| Fry | 1.14E-06 | -1.09016 | 0.147 | 0.269 | 0.026922 |
| Arrdc1 | 1.19E-06 | -0.75989 | 0.138 | 0.26 | 0.028122 |
| Wdfy3 | 1.21E-06 | -0.95859 | 0.231 | 0.365 | 0.028474 |
| Gm10736 | 1.21E-06 | 0.258924 | 0.622 | 0.489 | 0.02848 |
| Abhd17c | 1.22E-06 | -0.84391 | 0.343 | 0.474 | 0.028741 |
| Cd63-ps | 1.24E-06 | 0.320043 | 0.596 | 0.465 | 0.029162 |
| Cdan1 | 1.27E-06 | -1.35205 | 0.096 | 0.207 | 0.030064 |
| Irf2bp2 | 1.31E-06 | -0.63968 | 0.244 | 0.381 | 0.030883 |
| Arl14ep | 1.33E-06 | -1.52834 | 0.109 | 0.223 | 0.031404 |
| Rpl27a | 1.39E-06 | 0.65614 | 0.801 | 0.733 | 0.032896 |
| Fam53b | 1.4E-06 | -0.5788 | 0.167 | 0.298 | 0.032948 |
| Gm10269 | 1.43E-06 | -0.49348 | 0.205 | 0.337 | 0.03369 |
| Zfp217 | 1.54E-06 | -0.80439 | 0.192 | 0.315 | 0.036435 |
| Rpl23a | 1.55E-06 | 0.423918 | 0.804 | 0.745 | 0.036574 |
| Nrip1 | 1.56E-06 | -0.33673 | 0.506 | 0.654 | 0.036757 |
| Rps5 | 1.59E-06 | 0.558494 | 0.772 | 0.718 | 0.037414 |
| Upk1b | 1.59E-06 | 0.415101 | 0.609 | 0.473 | 0.037497 |
| Ninj1 | 1.61E-06 | -0.67576 | 0.253 | 0.392 | 0.037974 |
| Grk2 | 1.62E-06 | -0.47733 | 0.478 | 0.619 | 0.038273 |
| Rpl17 | 1.65E-06 | 0.51883 | 0.824 | 0.778 | 0.03902 |
| Ighj1 | 1.67E-06 | 0.426908 | 0.288 | 0.179 | 0.039462 |
| Rnaset2b | 1.69E-06 | 0.365613 | 0.981 | 0.951 | 0.039756 |
| Chst12 | 1.72E-06 | -1.02591 | 0.112 | 0.228 | 0.040569 |
| Rhoa | 1.72E-06 | 0.458792 | 0.897 | 0.842 | 0.04062 |
| Ap1b1 | 1.75E-06 | -0.46229 | 0.317 | 0.459 | 0.04121 |
| Hk2 | 1.78E-06 | -0.48761 | 0.67 | 0.784 | 0.041957 |
| Rhoh | 1.79E-06 | -0.54789 | 0.333 | 0.48 | 0.042133 |
| Eif2s2 | 1.83E-06 | -1.34904 | 0.269 | 0.395 | 0.043086 |

| | | | | | |
|---|---|---|---|---|---|
| Eif3f | 1.88E-06 | 0.887643 | 0.522 | 0.436 | 0.044301 |
| Itch | 1.88E-06 | -0.98636 | 0.144 | 0.272 | 0.044375 |
| Ddx5 | 1.96E-06 | -0.72023 | 0.705 | 0.789 | 0.046258 |
| Tomm34 | 2E-06 | -0.90072 | 0.17 | 0.298 | 0.047274 |
| Dnm2 | 2.02E-06 | -0.63343 | 0.571 | 0.687 | 0.047618 |
| Edem1 | 2.03E-06 | 0.901295 | 0.663 | 0.591 | 0.047837 |
| Ints6l | 2.05E-06 | -0.35307 | 0.212 | 0.347 | 0.048411 |
| Adrb1 | 2.07E-06 | -0.68811 | 0.189 | 0.32 | 0.048899 |
| Clec4a3 | 2.09E-06 | -0.82218 | 0.285 | 0.415 | 0.049246 |
| Rpsa-ps10 | 2.1E-06 | 0.447731 | 0.67 | 0.532 | 0.049482 |
| Fyb | 2.1E-06 | -0.39251 | 0.548 | 0.687 | 0.049627 |

#### Supplementry Table 2. CycM DEGs compared to HS

| GeneName | p_val | avg_log2FC | pct.1 | pct.2 | p_val_adj |
|---|---|---|---|---|---|
| Mki67 | 0 | 7.92966 | 0.625 | 0.007 | 0 |
| Nuf2 | 0 | 7.384774 | 0.475 | 0.002 | 0 |
| Top2a | 1.7E-228 | 7.971913 | 0.575 | 0.009 | 3.9E-224 |
| Cdca8 | 6.6E-216 | 6.55856 | 0.475 | 0.006 | 1.6E-211 |
| Knstrn | 7.1E-215 | 6.270567 | 0.425 | 0.005 | 1.7E-210 |
| Racgap1 | 1.6E-208 | 7.560089 | 0.525 | 0.008 | 3.8E-204 |
| Dlgap5 | 5.2E-198 | 5.988046 | 0.25 | 0.001 | 1.2E-193 |
| Cenpe | 8.1E-190 | 6.424152 | 0.425 | 0.006 | 1.9E-185 |
| Ccna2 | 2.1E-188 | 7.757909 | 0.575 | 0.012 | 5E-184 |
| Melk | 7.2E-180 | 5.581301 | 0.2 | 0.001 | 1.7E-175 |
| Birc5 | 9.4E-179 | 6.814788 | 0.4 | 0.005 | 2.2E-174 |
| Kifc1 | 2.9E-178 | 6.816497 | 0.4 | 0.005 | 6.7E-174 |
| Ncapg | 8.8E-164 | 5.740918 | 0.35 | 0.004 | 2.1E-159 |
| Ckap2l | 1.1E-162 | 6.055529 | 0.575 | 0.014 | 2.7E-158 |
| Cenpa | 9.7E-158 | 6.504528 | 0.575 | 0.014 | 2.3E-153 |
| Lockd | 2.4E-156 | 3.698264 | 0.275 | 0.002 | 5.6E-152 |
| Ndc80 | 9.9E-156 | 6.730219 | 0.4 | 0.006 | 2.3E-151 |
| Foxm1 | 3.5E-155 | 5.146133 | 0.4 | 0.006 | 8.2E-151 |
| Arhgef39 | 1.5E-154 | 5.912901 | 0.45 | 0.008 | 3.5E-150 |
| Prc1 | 2.3E-153 | 6.498407 | 0.425 | 0.007 | 5.4E-149 |
| Kif23 | 2.9E-153 | 6.279027 | 0.325 | 0.004 | 6.8E-149 |
| Sgo2a | 1.3E-152 | 4.785876 | 0.25 | 0.002 | 3.1E-148 |
| Nusap1 | 5.5E-152 | 5.762997 | 0.45 | 0.009 | 1.3E-147 |
| Ccnb2 | 7.3E-151 | 7.822846 | 0.45 | 0.009 | 1.7E-146 |
| Kif20a | 3E-145 | 6.457738 | 0.325 | 0.004 | 7.1E-141 |
| Prr11 | 1.4E-137 | 4.58581 | 0.3 | 0.004 | 3.3E-133 |
| Anln | 1.6E-136 | 4.879797 | 0.375 | 0.006 | 3.7E-132 |
| Bub1b | 1.1E-134 | 5.753393 | 0.325 | 0.005 | 2.6E-130 |
| Fignl1 | 2.3E-130 | 4.743418 | 0.35 | 0.006 | 5.3E-126 |
| Aspm | 5.9E-129 | 4.479042 | 0.25 | 0.003 | 1.4E-124 |
| 4930579G2 | 1.9E-128 | 4.771883 | 0.375 | 0.007 | 4.5E-124 |
| Cdca2 | 9.9E-127 | 5.77882 | 0.3 | 0.004 | 2.3E-122 |
| Cip2a | 3.4E-126 | 3.822212 | 0.275 | 0.003 | 7.9E-122 |
| Knl1 | 9.2E-123 | 5.875952 | 0.275 | 0.004 | 2.2E-118 |
| Ncapd2 | 9.7E-121 | 5.747501 | 0.525 | 0.016 | 2.3E-116 |
| Cdk1 | 1.4E-115 | 6.46051 | 0.475 | 0.014 | 3.2E-111 |
| Kif22 | 4E-115 | 6.321288 | 0.375 | 0.008 | 9.4E-111 |
| Rad51ap1 | 4.1E-114 | 4.794829 | 0.325 | 0.006 | 9.6E-110 |
| Pbk | 2.4E-113 | 6.823007 | 0.2 | 0.002 | 5.7E-109 |
| Cdc20 | 9.9E-111 | 7.052419 | 0.35 | 0.007 | 2.3E-106 |
| Cenpk | 5.4E-108 | 4.426982 | 0.25 | 0.003 | 1.3E-103 |
| Cdca3 | 1.1E-106 | 6.096758 | 0.275 | 0.004 | 2.6E-102 |
| Plk1 | 9.1E-105 | 7.243936 | 0.25 | 0.004 | 2.1E-100 |

| | | | | | |
|---|---|---|---|---|---|
| Cenpf | 1.5E-103 | 6.021732 | 0.55 | 0.021 | 3.5E-99 |
| Gm11223 | 9.8E-102 | 4.930332 | 0.875 | 0.063 | 2.3E-97 |
| Trim59 | 2.6E-101 | 4.618539 | 0.25 | 0.004 | 6.23E-97 |
| Tpx2 | 1E-100 | 5.185233 | 0.225 | 0.003 | 2.42E-96 |
| Hmmr | 6.07E-97 | 5.265615 | 0.225 | 0.003 | 1.43E-92 |
| Mis18bp1 | 1.31E-95 | 5.77085 | 0.25 | 0.004 | 3.08E-91 |
| Kif2c | 5.75E-95 | 4.403097 | 0.25 | 0.004 | 1.36E-90 |
| Tacc3 | 2.17E-92 | 5.01396 | 0.5 | 0.02 | 5.12E-88 |
| Lmnb1 | 1.54E-90 | 5.366583 | 0.525 | 0.023 | 3.63E-86 |
| Aurkb | 4.38E-90 | 5.090023 | 0.25 | 0.004 | 1.03E-85 |
| Esco2 | 1.11E-89 | 5.070272 | 0.2 | 0.003 | 2.62E-85 |
| Gen1 | 8.21E-89 | 4.568677 | 0.35 | 0.01 | 1.94E-84 |
| Clspn | 3.89E-88 | 5.768722 | 0.25 | 0.005 | 9.19E-84 |
| Rad51 | 2.24E-85 | 5.419065 | 0.3 | 0.007 | 5.29E-81 |
| Ccnf | 9.52E-85 | 6.40834 | 0.225 | 0.004 | 2.25E-80 |
| Cit | 3.96E-80 | 5.20088 | 0.375 | 0.013 | 9.35E-76 |
| Spc25 | 2.21E-75 | 4.532944 | 0.3 | 0.008 | 5.21E-71 |
| Kif20b | 2.02E-74 | 4.654278 | 0.275 | 0.007 | 4.78E-70 |
| Tcf19 | 1.99E-73 | 6.50238 | 0.3 | 0.009 | 4.7E-69 |
| Asf1b | 1.04E-72 | 5.435535 | 0.375 | 0.014 | 2.46E-68 |
| Spc24 | 1.33E-72 | 6.446321 | 0.375 | 0.014 | 3.15E-68 |
| Mybl1 | 2.41E-71 | 1.785573 | 0.2 | 0.004 | 5.7E-67 |
| Cks1b | 5.88E-71 | 4.572826 | 0.5 | 0.026 | 1.39E-66 |
| Stmn1 | 6.41E-71 | 5.716221 | 0.9 | 0.102 | 1.51E-66 |
| Aurka | 2.5E-70 | 5.204939 | 0.275 | 0.007 | 5.91E-66 |
| Mad2l1 | 3.94E-70 | 4.574022 | 0.45 | 0.021 | 9.29E-66 |
| Gas2l3 | 7.18E-69 | 5.379139 | 0.3 | 0.009 | 1.69E-64 |
| Kifc5b | 8.38E-65 | 3.822672 | 0.35 | 0.014 | 1.98E-60 |
| Cdkn2c | 1.06E-64 | 5.289239 | 0.5 | 0.03 | 2.5E-60 |
| Gm4739 | 6.17E-63 | 2.292424 | 0.775 | 0.075 | 1.46E-58 |
| Mastl | 2.72E-61 | 3.839749 | 0.275 | 0.009 | 6.43E-57 |
| Kif15 | 1.33E-60 | 5.438764 | 0.375 | 0.017 | 3.14E-56 |
| H2afx | 2.47E-60 | 5.151073 | 0.675 | 0.061 | 5.83E-56 |
| Cdkn1a | 9.55E-59 | 6.194143 | 0.825 | 0.102 | 2.25E-54 |
| Gm4316 | 1.08E-57 | 3.268697 | 0.425 | 0.024 | 2.54E-53 |
| Tk1 | 1.2E-55 | 5.451561 | 0.55 | 0.042 | 2.84E-51 |
| Gm42047 | 8.48E-55 | 4.80397 | 0.45 | 0.028 | 2E-50 |
| Cks2 | 1.06E-54 | 4.278094 | 0.6 | 0.052 | 2.5E-50 |
| Uhrf1 | 1.81E-53 | 4.844673 | 0.2 | 0.005 | 4.27E-49 |
| Dtl | 3E-53 | 2.810314 | 0.2 | 0.005 | 7.07E-49 |
| Ccne2 | 6.45E-50 | 5.131232 | 0.325 | 0.016 | 1.52E-45 |
| Smc2 | 2.17E-49 | 4.524488 | 0.5 | 0.039 | 5.11E-45 |
| Pclaf | 2.25E-49 | 4.71653 | 0.375 | 0.021 | 5.3E-45 |
| Gm15452 | 1.17E-48 | 0.437796 | 0.25 | 0.009 | 2.77E-44 |
| Zgrf1 | 4E-48 | 2.991624 | 0.275 | 0.012 | 9.43E-44 |
| Ncapg2 | 1.69E-44 | 4.036473 | 0.275 | 0.013 | 3.98E-40 |
| Prim1 | 3.34E-42 | 4.356212 | 0.275 | 0.013 | 7.88E-38 |

| | | | | | |
|---|---|---|---|---|---|
| Nsl1 | 9.05E-42 | 3.115745 | 0.375 | 0.025 | 2.14E-37 |
| Cchcr1 | 1.4E-41 | 3.502299 | 0.225 | 0.009 | 3.31E-37 |
| Incenp | 8.21E-41 | 4.84188 | 0.4 | 0.03 | 1.94E-36 |
| Smc4 | 8.9E-41 | 4.885552 | 0.55 | 0.059 | 2.1E-36 |
| Slc43a3 | 9.55E-41 | 5.070786 | 0.45 | 0.038 | 2.25E-36 |
| Bora | 1.26E-40 | 4.264521 | 0.3 | 0.017 | 2.97E-36 |
| Lmnb2 | 3.68E-39 | 4.152611 | 0.375 | 0.027 | 8.67E-35 |
| Atad2 | 6.82E-39 | 4.209799 | 0.275 | 0.015 | 1.61E-34 |
| Gm19945 | 2.88E-38 | 0.439876 | 0.2 | 0.008 | 6.78E-34 |
| Wdhd1 | 3.31E-37 | 3.706997 | 0.3 | 0.018 | 7.82E-33 |
| Hmgb2 | 6.05E-37 | 4.529073 | 0.875 | 0.193 | 1.43E-32 |
| Tubb4b-ps1 | 6.85E-37 | 1.656878 | 0.675 | 0.095 | 1.62E-32 |
| Hist1h2ab | 1.41E-35 | 3.740814 | 0.325 | 0.022 | 3.34E-31 |
| Ezh2 | 1.7E-33 | 4.182954 | 0.375 | 0.032 | 4.01E-29 |
| Fbxo5 | 7.84E-33 | 5.003196 | 0.225 | 0.012 | 1.85E-28 |
| Pmf1 | 1.13E-32 | 3.893659 | 0.425 | 0.042 | 2.67E-28 |
| Hist2h2ac | 1.38E-32 | 3.986557 | 0.375 | 0.033 | 3.26E-28 |
| Gm6750 | 3.77E-32 | 1.238368 | 0.525 | 0.065 | 8.9E-28 |
| Dna2 | 5.88E-31 | 4.136119 | 0.2 | 0.01 | 1.39E-26 |
| Gm42427 | 2.72E-30 | 0.688931 | 0.4 | 0.04 | 6.43E-26 |
| Hspb6 | 2.24E-29 | 3.900726 | 0.25 | 0.016 | 5.29E-25 |
| Nt5dc2 | 3.01E-29 | 3.663329 | 0.5 | 0.065 | 7.11E-25 |
| Cbx5 | 4.31E-29 | 2.689875 | 0.6 | 0.093 | 1.02E-24 |
| Enkd1 | 9.56E-29 | 3.341405 | 0.2 | 0.01 | 2.26E-24 |
| Gm12919 | 1.9E-28 | 0.519585 | 0.275 | 0.02 | 4.48E-24 |
| Ac125028.1 | 2.55E-28 | 0.765881 | 0.225 | 0.013 | 6.02E-24 |
| Ddx39 | 3.66E-28 | 2.881128 | 0.575 | 0.085 | 8.62E-24 |
| Hist1h2ae | 2.42E-27 | 3.873588 | 0.425 | 0.051 | 5.7E-23 |
| Smtn | 2.44E-27 | 5.178914 | 0.225 | 0.014 | 5.75E-23 |
| Kcna3 | 1.33E-26 | 2.800702 | 0.275 | 0.021 | 3.13E-22 |
| Tyms | 2.22E-25 | 3.127336 | 0.525 | 0.079 | 5.23E-21 |
| Prdx4 | 1.44E-24 | 2.614306 | 0.8 | 0.189 | 3.4E-20 |
| Tubb4b | 1.59E-24 | 3.79729 | 0.775 | 0.216 | 3.75E-20 |
| Atad5 | 1.69E-24 | 3.182428 | 0.25 | 0.019 | 4E-20 |
| Lin9 | 2.32E-24 | 2.729886 | 0.425 | 0.055 | 5.48E-20 |
| Hist1h1d | 2.81E-24 | 3.06122 | 0.3 | 0.028 | 6.62E-20 |
| Hmgn2-ps1 | 4.02E-24 | 1.354143 | 0.625 | 0.119 | 9.47E-20 |
| Gm10182 | 1.29E-23 | 0.334196 | 0.25 | 0.02 | 3.03E-19 |
| Gm6682 | 1.8E-23 | 1.683588 | 0.6 | 0.119 | 4.26E-19 |
| Mcm7 | 4.94E-23 | 3.083392 | 0.425 | 0.058 | 1.17E-18 |
| Polh | 6.04E-23 | 1.901695 | 0.425 | 0.057 | 1.42E-18 |
| H2afv | 7.18E-23 | 2.898965 | 0.75 | 0.19 | 1.69E-18 |
| Gm20628 | 7.3E-23 | 1.567644 | 0.2 | 0.013 | 1.72E-18 |
| Ncapd3 | 1.46E-22 | 1.819481 | 0.4 | 0.051 | 3.45E-18 |
| Hmgn2 | 5.26E-22 | 3.159456 | 0.925 | 0.357 | 1.24E-17 |
| Cdkn2d | 5.31E-22 | 3.6214 | 0.375 | 0.048 | 1.25E-17 |
| Dbf4 | 5.8E-22 | 3.607376 | 0.275 | 0.026 | 1.37E-17 |

| | | | | | |
|---|---|---|---|---|---|
| Gm10282 | 6.58E-22 | 2.517966 | 0.65 | 0.146 | 1.55E-17 |
| Fen1 | 8.24E-22 | 2.873818 | 0.375 | 0.047 | 1.94E-17 |
| Mcm2 | 9.02E-22 | 2.733867 | 0.275 | 0.026 | 2.13E-17 |
| Haus5 | 9.06E-22 | 3.264892 | 0.225 | 0.018 | 2.14E-17 |
| Fancd2 | 2.03E-21 | 2.921531 | 0.2 | 0.014 | 4.79E-17 |
| Rnf26 | 3.58E-21 | 1.902974 | 0.55 | 0.098 | 8.45E-17 |
| Rrm1 | 3.82E-21 | 3.455325 | 0.425 | 0.064 | 9.01E-17 |
| Ifi27l2a | 6.05E-21 | 4.49998 | 0.45 | 0.074 | 1.43E-16 |
| Ptgr1 | 6.07E-21 | 2.416665 | 0.425 | 0.061 | 1.43E-16 |
| Tmpo | 7.05E-21 | 3.126369 | 0.725 | 0.195 | 1.66E-16 |
| Rdm1 | 8.32E-21 | 3.082126 | 0.475 | 0.079 | 1.96E-16 |
| Mcm5 | 9.04E-21 | 3.135529 | 0.275 | 0.027 | 2.13E-16 |
| Hist1h2ap | 1.12E-20 | 2.068161 | 0.425 | 0.067 | 2.65E-16 |
| Ramp1 | 1.92E-20 | 3.234966 | 0.725 | 0.182 | 4.52E-16 |
| Gm13007 | 2.17E-20 | 0.271262 | 0.2 | 0.015 | 5.13E-16 |
| Tagln2 | 2.82E-20 | 2.359297 | 0.775 | 0.222 | 6.66E-16 |
| Cenpl | 3.4E-20 | 2.824647 | 0.275 | 0.028 | 8.02E-16 |
| Suv39h1 | 5.06E-20 | 3.065473 | 0.35 | 0.045 | 1.19E-15 |
| Haspin | 6.1E-20 | 2.953226 | 0.4 | 0.058 | 1.44E-15 |
| Ska2 | 9.15E-20 | 2.705609 | 0.25 | 0.024 | 2.16E-15 |
| Rfwd3 | 1.06E-19 | 3.483686 | 0.5 | 0.094 | 2.49E-15 |
| Gm7931 | 1.29E-19 | 1.640123 | 0.625 | 0.148 | 3.04E-15 |
| Tubb4b-ps2 | 2.24E-19 | 0.431115 | 0.2 | 0.016 | 5.29E-15 |
| Kntc1 | 3.56E-19 | 2.043995 | 0.275 | 0.029 | 8.4E-15 |
| Kpna2 | 3.65E-19 | 3.430182 | 0.55 | 0.114 | 8.6E-15 |
| Selenoh | 3.81E-19 | 2.466039 | 0.425 | 0.067 | 8.99E-15 |
| Dnajc9 | 1.76E-18 | 3.847964 | 0.4 | 0.063 | 4.14E-14 |
| Cenpt | 1.9E-18 | 2.361325 | 0.4 | 0.06 | 4.49E-14 |
| Nemp1 | 2.07E-18 | 2.082566 | 0.3 | 0.036 | 4.88E-14 |
| Nptxr | 2.13E-18 | 2.415256 | 0.225 | 0.021 | 5.02E-14 |
| Tex30 | 2.73E-18 | 2.59478 | 0.35 | 0.048 | 6.43E-14 |
| Cntrob | 3.43E-18 | 1.910542 | 0.25 | 0.026 | 8.09E-14 |
| Slc20a1 | 4.23E-18 | 2.596857 | 0.3 | 0.037 | 9.99E-14 |
| Ube2s | 5E-18 | 3.003582 | 0.65 | 0.171 | 1.18E-13 |
| Rgs12 | 5.34E-18 | 2.31412 | 0.25 | 0.026 | 1.26E-13 |
| Cmc2 | 8.69E-18 | 2.99544 | 0.325 | 0.043 | 2.05E-13 |
| Rangap1 | 9.22E-18 | 2.729159 | 0.5 | 0.1 | 2.18E-13 |
| Cep57l1 | 1.27E-17 | 1.997467 | 0.4 | 0.063 | 2.99E-13 |
| Hist1h2ao | 1.5E-17 | 3.524325 | 0.525 | 0.122 | 3.54E-13 |
| Zfp367 | 1.52E-17 | 3.349184 | 0.35 | 0.051 | 3.58E-13 |
| AI506816 | 1.89E-17 | 2.364969 | 0.8 | 0.254 | 4.47E-13 |
| Arhgap19 | 3.94E-17 | 2.497797 | 0.25 | 0.027 | 9.3E-13 |
| Nde1 | 4.15E-17 | 2.204743 | 0.575 | 0.13 | 9.8E-13 |
| Gm10184 | 5.27E-17 | 1.294945 | 0.375 | 0.06 | 1.24E-12 |
| E2f6 | 2.08E-16 | 2.33304 | 0.25 | 0.028 | 4.9E-12 |
| Rbl1 | 2.17E-16 | 2.839319 | 0.35 | 0.055 | 5.12E-12 |
| Haus3 | 2.8E-16 | 2.386973 | 0.35 | 0.054 | 6.6E-12 |

| | | | | | |
|---|---|---|---|---|---|
| Nup210 | 4.45E-16 | 4.092047 | 0.225 | 0.024 | 1.05E-11 |
| Sae1 | 5.87E-16 | 2.309792 | 0.525 | 0.118 | 1.38E-11 |
| Gm10143 | 1.04E-15 | 2.122564 | 0.2 | 0.019 | 2.45E-11 |
| Cdk5rap2 | 1.61E-15 | 2.261178 | 0.35 | 0.056 | 3.8E-11 |
| Gm3226 | 1.74E-15 | 1.132009 | 0.675 | 0.202 | 4.09E-11 |
| H2afz | 1.96E-15 | 2.734596 | 0.9 | 0.501 | 4.63E-11 |
| H2-Q4 | 2.42E-15 | 1.108327 | 0.375 | 0.065 | 5.72E-11 |
| Wdr76 | 2.5E-15 | 1.855199 | 0.35 | 0.058 | 5.89E-11 |
| Lamb2 | 2.77E-15 | 2.639677 | 0.325 | 0.051 | 6.54E-11 |
| Reep4 | 4.26E-15 | 2.556771 | 0.45 | 0.092 | 1.01E-10 |
| Gm10357 | 4.56E-15 | 0.456041 | 0.225 | 0.026 | 1.07E-10 |
| Rfc5 | 4.58E-15 | 2.534287 | 0.275 | 0.037 | 1.08E-10 |
| Cep89 | 5.96E-15 | 3.193856 | 0.225 | 0.026 | 1.41E-10 |
| Glipr1 | 7.76E-15 | 2.878055 | 0.325 | 0.05 | 1.83E-10 |
| Hdgf | 1.29E-14 | 1.697983 | 0.7 | 0.208 | 3.05E-10 |
| Nup62 | 1.71E-14 | 2.246917 | 0.55 | 0.135 | 4.03E-10 |
| Sdk1 | 2.28E-14 | 1.754166 | 0.35 | 0.059 | 5.38E-10 |
| Aaas | 3.08E-14 | 2.27104 | 0.35 | 0.06 | 7.28E-10 |
| Alg8 | 4.82E-14 | 1.464863 | 0.625 | 0.165 | 1.14E-09 |
| Anp32b | 4.83E-14 | 2.20776 | 0.7 | 0.232 | 1.14E-09 |
| Tuba1c | 4.85E-14 | 3.989047 | 0.8 | 0.4 | 1.14E-09 |
| Jpt2 | 6.69E-14 | 1.949794 | 0.425 | 0.088 | 1.58E-09 |
| Rbmx2 | 6.9E-14 | 1.603387 | 0.2 | 0.022 | 1.63E-09 |
| Usp1 | 6.91E-14 | 2.661457 | 0.375 | 0.071 | 1.63E-09 |
| 4930503L1 | 9.14E-14 | 2.795681 | 0.2 | 0.022 | 2.16E-09 |
| Gm14150 | 1.04E-13 | 1.065155 | 0.775 | 0.305 | 2.45E-09 |
| Ctsd | 2.54E-13 | -1.19002 | 1 | 1 | 6E-09 |
| P2ry12 | 3.64E-13 | -1.25396 | 1 | 1 | 8.58E-09 |
| Malat1 | 4.63E-13 | -1.21917 | 1 | 0.992 | 1.09E-08 |
| Ckap5 | 6.26E-13 | 2.356305 | 0.5 | 0.128 | 1.48E-08 |
| Cln6 | 6.67E-13 | 2.265078 | 0.35 | 0.066 | 1.57E-08 |
| Tuba1b | 7.66E-13 | 1.9677 | 0.975 | 0.779 | 1.81E-08 |
| 1700025G( | 7.96E-13 | 1.902292 | 0.225 | 0.029 | 1.88E-08 |
| Brip1os | 1.01E-12 | 2.086826 | 0.6 | 0.181 | 2.37E-08 |
| Pidd1 | 1.4E-12 | 3.248951 | 0.2 | 0.024 | 3.31E-08 |
| Dynlt1a | 1.5E-12 | 3.691899 | 0.25 | 0.038 | 3.54E-08 |
| Tfdp1 | 1.55E-12 | 2.756954 | 0.35 | 0.068 | 3.66E-08 |
| Gm13577 | 1.61E-12 | 1.097526 | 0.575 | 0.162 | 3.79E-08 |
| Gm13461 | 2.01E-12 | 0.88437 | 0.575 | 0.173 | 4.75E-08 |
| Gm26623 | 2.72E-12 | -0.84113 | 0.3 | 0.051 | 6.41E-08 |
| Pomt1 | 2.89E-12 | 1.901443 | 0.425 | 0.096 | 6.81E-08 |
| Hyls1 | 3.07E-12 | 3.227306 | 0.225 | 0.032 | 7.23E-08 |
| Arl6ip1 | 3.48E-12 | 2.203696 | 0.95 | 0.76 | 8.2E-08 |
| Calm3 | 3.49E-12 | 1.696658 | 0.725 | 0.241 | 8.22E-08 |
| Poc1a | 3.81E-12 | 3.511212 | 0.225 | 0.031 | 8.99E-08 |
| Nucks1 | 4.66E-12 | 1.794894 | 0.75 | 0.292 | 1.1E-07 |
| Gm17383 | 6.73E-12 | 2.139714 | 0.925 | 0.572 | 1.59E-07 |

| | | | | | |
|---|---|---|---|---|---|
| Nup107 | 7.53E-12 | 2.10042 | 0.275 | 0.045 | 1.78E-07 |
| Lmf2 | 8.68E-12 | 2.46354 | 0.6 | 0.2 | 2.05E-07 |
| Orc6 | 8.82E-12 | 2.999815 | 0.275 | 0.046 | 2.08E-07 |
| Anp32e | 1.45E-11 | 1.482335 | 0.6 | 0.179 | 3.41E-07 |
| Arhgap11a | 1.55E-11 | 0.594458 | 0.275 | 0.045 | 3.65E-07 |
| Rcc1 | 1.85E-11 | 2.653957 | 0.25 | 0.039 | 4.37E-07 |
| Ncaph2 | 2.6E-11 | 1.905837 | 0.65 | 0.247 | 6.13E-07 |
| Dck | 3.1E-11 | 2.236565 | 0.35 | 0.074 | 7.3E-07 |
| Hirip3 | 3.15E-11 | 3.066325 | 0.2 | 0.027 | 7.43E-07 |
| Rad21 | 3.98E-11 | 1.047645 | 0.5 | 0.14 | 9.38E-07 |
| Ran | 4.14E-11 | 1.78444 | 0.825 | 0.388 | 9.76E-07 |
| Dnmt1 | 4.83E-11 | 2.480126 | 0.325 | 0.067 | 1.14E-06 |
| Pold1 | 6.1E-11 | 3.617789 | 0.225 | 0.035 | 1.44E-06 |
| Lbr | 8.81E-11 | 1.749515 | 0.575 | 0.184 | 2.08E-06 |
| Rad54l | 9.22E-11 | 3.120965 | 0.275 | 0.052 | 2.17E-06 |
| Gm9844 | 9.41E-11 | 0.524743 | 0.4 | 0.097 | 2.22E-06 |
| Jade1 | 9.83E-11 | 2.079774 | 0.375 | 0.085 | 2.32E-06 |
| Fbxo34 | 1.01E-10 | 2.093832 | 0.35 | 0.078 | 2.37E-06 |
| Slc2a5 | 1.06E-10 | -1.84498 | 0.8 | 0.944 | 2.5E-06 |
| Gm5620 | 1.12E-10 | 1.154322 | 0.625 | 0.233 | 2.65E-06 |
| Hmgb1-ps7 | 1.23E-10 | 1.134927 | 0.475 | 0.133 | 2.89E-06 |
| Hjurp | 1.3E-10 | 2.738308 | 0.6 | 0.222 | 3.06E-06 |
| G2e3 | 1.35E-10 | 2.061257 | 0.3 | 0.059 | 3.18E-06 |
| Nudc-ps1 | 1.57E-10 | 0.513885 | 0.425 | 0.105 | 3.69E-06 |
| Kmt5a | 1.57E-10 | 1.751636 | 0.475 | 0.128 | 3.7E-06 |
| Gm15387 | 1.79E-10 | 0.571267 | 0.45 | 0.121 | 4.22E-06 |
| Tpm4 | 1.89E-10 | 2.503234 | 0.45 | 0.129 | 4.45E-06 |
| Ndc1 | 1.96E-10 | 2.440637 | 0.2 | 0.029 | 4.62E-06 |
| Fcgr3 | 2.16E-10 | -1.7273 | 0.775 | 0.961 | 5.1E-06 |
| Snrpd1 | 2.38E-10 | 1.584927 | 0.525 | 0.156 | 5.62E-06 |
| Hmgb1-ps2 | 2.75E-10 | 0.855826 | 0.575 | 0.191 | 6.49E-06 |
| Ccng1 | 2.83E-10 | 1.471018 | 0.45 | 0.121 | 6.68E-06 |
| Dbr1 | 2.84E-10 | 2.027734 | 0.275 | 0.051 | 6.7E-06 |
| Mast2 | 2.94E-10 | 2.317144 | 0.325 | 0.07 | 6.93E-06 |
| Cenpq | 3.11E-10 | 3.017688 | 0.225 | 0.037 | 7.34E-06 |
| Ccdc34 | 3.24E-10 | 1.786113 | 0.3 | 0.06 | 7.64E-06 |
| Htra3 | 3.33E-10 | 2.541357 | 0.35 | 0.083 | 7.86E-06 |
| Jpt1 | 3.44E-10 | 1.597699 | 0.825 | 0.401 | 8.11E-06 |
| Pgp | 3.55E-10 | 2.266463 | 0.55 | 0.192 | 8.37E-06 |
| Pcgf5 | 4.28E-10 | 2.062247 | 0.275 | 0.053 | 1.01E-05 |
| Rbm3 | 4.77E-10 | 1.54059 | 0.725 | 0.291 | 1.13E-05 |
| Cfap43 | 4.88E-10 | 2.655776 | 0.2 | 0.03 | 1.15E-05 |
| Selenop | 6.26E-10 | -1.35191 | 0.975 | 0.991 | 1.48E-05 |
| Emp3 | 6.59E-10 | 1.93944 | 0.45 | 0.128 | 1.55E-05 |
| Xpo1 | 6.97E-10 | 1.353341 | 0.4 | 0.103 | 1.64E-05 |
| Mis12 | 6.99E-10 | 1.264875 | 0.4 | 0.1 | 1.65E-05 |
| Ncaph | 7.16E-10 | 1.342373 | 0.425 | 0.11 | 1.69E-05 |

| | | | | | |
|---|---|---|---|---|---|
| Gm3756 | 8.74E-10 | 0.597356 | 0.575 | 0.206 | 2.06E-05 |
| Hmgb1-ps1 | 9.39E-10 | 0.299403 | 0.275 | 0.053 | 2.21E-05 |
| Gm21596 | 9.59E-10 | 1.168077 | 0.7 | 0.276 | 2.26E-05 |
| Slc24a5 | 9.73E-10 | 0.376582 | 0.225 | 0.037 | 2.3E-05 |
| Sugp2 | 1.08E-09 | 1.814098 | 0.35 | 0.084 | 2.55E-05 |
| Nudc | 1.1E-09 | 1.388762 | 0.725 | 0.304 | 2.59E-05 |
| Chac2 | 1.35E-09 | 1.324225 | 0.2 | 0.03 | 3.18E-05 |
| Nup188 | 1.38E-09 | 3.069536 | 0.275 | 0.055 | 3.24E-05 |
| Cdk2 | 1.44E-09 | 2.47405 | 0.35 | 0.087 | 3.4E-05 |
| Taf9 | 1.5E-09 | 1.304918 | 0.225 | 0.038 | 3.54E-05 |
| Mum1 | 1.51E-09 | 2.206494 | 0.35 | 0.084 | 3.56E-05 |
| Maz | 1.68E-09 | 2.018147 | 0.525 | 0.179 | 3.96E-05 |
| Rbbp8 | 1.95E-09 | 3.214684 | 0.225 | 0.039 | 4.59E-05 |
| Smc1a | 1.96E-09 | 1.51215 | 0.625 | 0.226 | 4.62E-05 |
| Gm12346 | 2.17E-09 | 0.967898 | 0.625 | 0.273 | 5.11E-05 |
| Lig1 | 2.64E-09 | 3.351432 | 0.225 | 0.04 | 6.22E-05 |
| Haus1 | 2.94E-09 | 2.173229 | 0.225 | 0.039 | 6.93E-05 |
| Cnnm4 | 2.96E-09 | 1.583922 | 0.575 | 0.195 | 6.99E-05 |
| Stom | 2.99E-09 | 3.296245 | 0.2 | 0.033 | 7.04E-05 |
| Raly | 3.1E-09 | 1.402191 | 0.825 | 0.406 | 7.3E-05 |
| Ppil1 | 3.16E-09 | 2.077582 | 0.25 | 0.048 | 7.45E-05 |
| Dek | 3.3E-09 | 2.331461 | 0.65 | 0.283 | 7.78E-05 |
| Prpf3 | 3.4E-09 | 2.628726 | 0.3 | 0.066 | 8.02E-05 |
| Sesn2 | 3.48E-09 | 2.176339 | 0.25 | 0.048 | 8.2E-05 |
| Nasp | 4.07E-09 | 0.424583 | 0.3 | 0.065 | 9.6E-05 |
| Rbbp7 | 4.12E-09 | 1.522286 | 0.675 | 0.271 | 9.72E-05 |
| Haus8 | 4.47E-09 | 2.284773 | 0.25 | 0.048 | 0.000105 |
| Milr1 | 4.53E-09 | 1.83027 | 0.35 | 0.087 | 0.000107 |
| Rfc3 | 5.47E-09 | 1.333071 | 0.275 | 0.057 | 0.000129 |
| Gm7870 | 6.47E-09 | 0.431192 | 0.325 | 0.076 | 0.000153 |
| Tmsb10 | 6.47E-09 | 1.844007 | 0.8 | 0.412 | 0.000153 |
| Trim28 | 7.01E-09 | 1.442631 | 0.725 | 0.321 | 0.000165 |
| Tubb5 | 7.17E-09 | 2.492982 | 0.95 | 0.809 | 0.000169 |
| Galns | 8.23E-09 | 1.996849 | 0.25 | 0.049 | 0.000194 |
| Sass6 | 8.86E-09 | 2.514651 | 0.275 | 0.059 | 0.000209 |
| Tipin | 9.25E-09 | 2.628373 | 0.275 | 0.059 | 0.000218 |
| Nemp2 | 9.73E-09 | 0.447765 | 0.225 | 0.04 | 0.00023 |
| Gm5844 | 1.11E-08 | 0.680997 | 0.6 | 0.228 | 0.000262 |
| Nsd2 | 1.13E-08 | 2.677408 | 0.375 | 0.106 | 0.000268 |
| Cbx3 | 1.2E-08 | 1.47841 | 0.8 | 0.418 | 0.000283 |
| Zhx3 | 1.35E-08 | 0.734199 | 0.4 | 0.114 | 0.000319 |
| Lsm2 | 1.38E-08 | 1.850935 | 0.4 | 0.116 | 0.000326 |
| Recql | 1.42E-08 | 2.743941 | 0.275 | 0.062 | 0.000334 |
| Stt3b | 1.67E-08 | 1.848015 | 0.5 | 0.176 | 0.000394 |
| Cpd | 1.78E-08 | 2.167835 | 0.325 | 0.082 | 0.000419 |
| Tubg1 | 1.78E-08 | 1.786098 | 0.325 | 0.081 | 0.00042 |
| Hat1 | 1.79E-08 | 1.991265 | 0.375 | 0.105 | 0.000421 |

| | | | | | |
|---|---|---|---|---|---|
| Psip1 | 1.9E-08 | 1.123708 | 0.5 | 0.167 | 0.000449 |
| Mcm6 | 2.3E-08 | 1.840088 | 0.25 | 0.052 | 0.000542 |
| Eri1 | 2.32E-08 | 2.202089 | 0.35 | 0.094 | 0.000546 |
| Adgrl1 | 2.35E-08 | 2.746899 | 0.225 | 0.042 | 0.000555 |
| Ube2l6 | 2.36E-08 | 2.077804 | 0.3 | 0.071 | 0.000557 |
| Dtymk | 2.56E-08 | 2.069694 | 0.375 | 0.107 | 0.000603 |
| Unc93b1 | 2.76E-08 | -1.1256 | 0.975 | 0.991 | 0.000652 |
| Cnot6 | 3.72E-08 | 2.312691 | 0.35 | 0.095 | 0.000878 |
| Nap1l1 | 3.8E-08 | 1.754085 | 0.45 | 0.149 | 0.000895 |
| Efemp2 | 3.82E-08 | 1.014953 | 0.225 | 0.043 | 0.000901 |
| Ccl12 | 3.85E-08 | 2.055223 | 0.5 | 0.184 | 0.000908 |
| Evi2 | 3.98E-08 | 0.415036 | 0.325 | 0.08 | 0.000938 |
| Psat1 | 4.18E-08 | 1.609839 | 0.35 | 0.092 | 0.000985 |
| Haus4 | 4.38E-08 | 1.569813 | 0.3 | 0.071 | 0.001034 |
| Ssna1 | 4.7E-08 | 1.33571 | 0.525 | 0.184 | 0.00111 |
| Ipo9 | 4.75E-08 | 0.924824 | 0.45 | 0.141 | 0.001121 |
| Gm4617 | 4.91E-08 | 0.72585 | 0.65 | 0.266 | 0.001159 |
| Actn4 | 5E-08 | 1.078151 | 0.625 | 0.252 | 0.001179 |
| Vsir | 5.15E-08 | -1.0664 | 0.95 | 0.985 | 0.001214 |
| Gm9800 | 5.19E-08 | 0.475755 | 0.575 | 0.219 | 0.001224 |
| Hsp90aa1 | 5.36E-08 | 1.854657 | 0.85 | 0.533 | 0.001265 |
| Rhno1 | 5.53E-08 | 1.205445 | 0.525 | 0.184 | 0.001303 |
| Pcyox1l | 6E-08 | 1.542029 | 0.425 | 0.13 | 0.001415 |
| Ankrd13a | 6.02E-08 | 1.182206 | 0.65 | 0.266 | 0.00142 |
| Nsmce1 | 6.47E-08 | 0.890678 | 0.525 | 0.182 | 0.001526 |
| Hmgn1 | 6.81E-08 | 1.581966 | 0.7 | 0.345 | 0.001606 |
| Cep135 | 7.08E-08 | 1.578099 | 0.25 | 0.054 | 0.00167 |
| Calu | 7.67E-08 | 1.332957 | 0.675 | 0.281 | 0.001808 |
| Sdc3 | 8.08E-08 | 1.272034 | 0.375 | 0.109 | 0.001905 |
| Uevld | 8.15E-08 | 1.541524 | 0.25 | 0.054 | 0.001921 |
| Gm21399 | 8.24E-08 | 0.772704 | 0.475 | 0.16 | 0.001943 |
| Anapc5 | 8.24E-08 | 1.704923 | 0.6 | 0.256 | 0.001945 |
| Nrm | 8.33E-08 | 1.756989 | 0.575 | 0.235 | 0.001965 |
| Fbxo45 | 8.35E-08 | 2.441659 | 0.2 | 0.038 | 0.001969 |
| Cdc25a | 8.41E-08 | 1.067847 | 0.2 | 0.037 | 0.001983 |
| Gm17201 | 9.3E-08 | 0.603918 | 0.25 | 0.054 | 0.002193 |
| Myef2 | 9.39E-08 | 1.670236 | 0.425 | 0.139 | 0.002215 |
| Siglech | 1E-07 | -1.00439 | 1 | 0.999 | 0.002367 |
| Cenpb | 1.04E-07 | 1.641452 | 0.4 | 0.125 | 0.002457 |
| Slc25a25 | 1.07E-07 | 2.746985 | 0.25 | 0.056 | 0.002516 |
| Apba1 | 1.1E-07 | 0.76268 | 0.525 | 0.184 | 0.002593 |
| Gm16374 | 1.15E-07 | 0.373517 | 0.3 | 0.076 | 0.00272 |
| Miip | 1.35E-07 | 1.937463 | 0.3 | 0.078 | 0.003192 |
| Hmgn3 | 1.4E-07 | 2.640217 | 0.2 | 0.039 | 0.003311 |
| Ptma | 1.41E-07 | 1.383131 | 0.9 | 0.629 | 0.003319 |
| Ppia | 1.44E-07 | 1.021212 | 1 | 0.969 | 0.00339 |
| Fkbp2 | 1.54E-07 | 1.277789 | 0.775 | 0.426 | 0.003622 |

| | | | | | |
|---|---|---|---|---|---|
| Chek2 | 1.59E-07 | 1.441732 | 0.225 | 0.047 | 0.003752 |
| Pak4 | 1.59E-07 | 1.467514 | 0.25 | 0.056 | 0.003759 |
| Tmem129 | 1.7E-07 | 0.938607 | 0.35 | 0.097 | 0.004007 |
| Bex3 | 1.72E-07 | 1.38905 | 0.225 | 0.047 | 0.004048 |
| Gm3788 | 1.94E-07 | 1.025748 | 0.375 | 0.117 | 0.004581 |
| Cenpc1 | 2.06E-07 | 1.751162 | 0.225 | 0.048 | 0.004866 |
| Rnaseh2b | 2.1E-07 | 1.80066 | 0.275 | 0.067 | 0.004958 |
| Pcna-ps2 | 2.18E-07 | 1.216283 | 0.375 | 0.122 | 0.005154 |
| Iqgap1 | 2.25E-07 | 3.587777 | 0.3 | 0.083 | 0.005302 |
| Ranbp3 | 2.25E-07 | 1.2023 | 0.475 | 0.168 | 0.005311 |
| Stard7 | 2.28E-07 | 1.136805 | 0.4 | 0.123 | 0.005377 |
| Gm5518 | 2.37E-07 | 0.739489 | 0.55 | 0.237 | 0.005589 |
| Tmem134 | 2.42E-07 | 1.220221 | 0.475 | 0.166 | 0.005718 |
| Alyref | 2.49E-07 | 1.673273 | 0.5 | 0.193 | 0.005881 |
| Flna | 2.52E-07 | 1.895303 | 0.375 | 0.112 | 0.00594 |
| Pla2g4a | 2.69E-07 | 0.833521 | 0.55 | 0.205 | 0.006357 |
| Timm50 | 2.73E-07 | 1.283472 | 0.5 | 0.18 | 0.006435 |
| Fam111a | 2.89E-07 | 1.427549 | 0.575 | 0.248 | 0.006821 |
| Gapdh | 2.97E-07 | 1.004657 | 1 | 0.904 | 0.007008 |
| Pcyox1 | 2.98E-07 | 1.083924 | 0.675 | 0.304 | 0.00702 |
| Coq2 | 3.07E-07 | 0.927068 | 0.5 | 0.182 | 0.007239 |
| Hmgb1 | 3.67E-07 | 1.111144 | 0.95 | 0.768 | 0.00866 |
| Hist1h1e | 3.89E-07 | 2.719607 | 0.45 | 0.179 | 0.009185 |
| Gpr34 | 3.91E-07 | -0.96444 | 0.95 | 0.992 | 0.009231 |
| Ppp2r5d | 4.21E-07 | 1.758313 | 0.4 | 0.128 | 0.009935 |
| Cbfb | 4.89E-07 | 0.907034 | 0.5 | 0.188 | 0.011531 |
| Ccdc77 | 5.24E-07 | 2.257623 | 0.2 | 0.041 | 0.012363 |
| Tmem109 | 5.71E-07 | 0.837044 | 0.85 | 0.433 | 0.013473 |
| Nxt2 | 5.91E-07 | 1.251526 | 0.4 | 0.131 | 0.013935 |
| Chuk | 6.28E-07 | 1.159909 | 0.4 | 0.131 | 0.014821 |
| Sfpq | 6.3E-07 | 1.155024 | 0.675 | 0.343 | 0.014868 |
| Pom121 | 7.47E-07 | 0.921291 | 0.6 | 0.257 | 0.017624 |
| Grk6 | 7.48E-07 | 1.597832 | 0.35 | 0.106 | 0.017653 |
| Ac122335.: | 7.68E-07 | 1.548556 | 0.2 | 0.042 | 0.018128 |
| Mafb | 8.27E-07 | -1.72315 | 0.85 | 0.913 | 0.019502 |
| Cd300c2 | 8.71E-07 | -2.04941 | 0.6 | 0.809 | 0.020539 |
| Clpb | 8.99E-07 | 0.703483 | 0.35 | 0.105 | 0.021211 |
| Cdkn2aipnl | 9.08E-07 | 1.390702 | 0.525 | 0.213 | 0.021419 |
| Tmem242 | 9.39E-07 | 1.945181 | 0.35 | 0.109 | 0.022147 |
| Gm10123 | 9.49E-07 | 0.765528 | 1 | 0.927 | 0.022377 |
| Ing2 | 9.63E-07 | 1.73348 | 0.3 | 0.083 | 0.022712 |
| Fam92a | 9.93E-07 | 2.593843 | 0.2 | 0.042 | 0.023415 |
| Rhob | 1.03E-06 | -1.55963 | 0.825 | 0.928 | 0.024202 |
| Zfp131 | 1.07E-06 | 1.021967 | 0.375 | 0.124 | 0.025326 |
| Larp7 | 1.18E-06 | 1.511703 | 0.3 | 0.084 | 0.027739 |
| Arl4c | 1.29E-06 | 0.837133 | 0.875 | 0.486 | 0.030531 |
| tpr | 1.32E-06 | 0.539051 | 0.65 | 0.293 | 0.031198 |

| | | | | | |
|---|---|---|---|---|---|
| Ctsa | 1.44E-06 | -1.2237 | 1 | 0.972 | 0.033954 |
| Tdp2 | 1.46E-06 | 1.422534 | 0.25 | 0.061 | 0.034414 |
| Gm4204 | 1.5E-06 | 0.439067 | 0.25 | 0.062 | 0.035302 |
| Smarca4 | 1.5E-06 | 1.651838 | 0.525 | 0.222 | 0.035327 |
| Ruvbl2 | 1.56E-06 | 1.541731 | 0.4 | 0.138 | 0.036874 |
| Hist1h4d | 1.57E-06 | 2.74843 | 0.55 | 0.302 | 0.036948 |
| Abhd2 | 1.6E-06 | 1.293359 | 0.425 | 0.155 | 0.037741 |
| Hexb | 1.63E-06 | -0.61863 | 1 | 1 | 0.038479 |
| Nup160 | 1.68E-06 | 1.443929 | 0.225 | 0.054 | 0.039561 |
| Srsf7 | 1.72E-06 | 1.168254 | 0.575 | 0.251 | 0.040605 |
| Mrpl28 | 1.74E-06 | 1.509143 | 0.375 | 0.124 | 0.041003 |
| Eif1ad | 1.74E-06 | 1.096175 | 0.4 | 0.132 | 0.041029 |
| H1f0 | 1.74E-06 | 1.432393 | 0.525 | 0.216 | 0.041093 |
| Ctsb | 1.84E-06 | -0.8899 | 1 | 0.999 | 0.043511 |
| Aldoa | 2.05E-06 | -2.41393 | 0.425 | 0.704 | 0.048297 |

Supplementry Table 3. Core microglia activation genes

| Genename | IntM | eDAM | DAM | IRM | TNFRM | MHCIIM |
|---|---|---|---|---|---|---|
| 1700017B( | -0.26849 | -0.41808 | -1.18367 | -0.36972 | 0 | -0.84507 |
| Abcc3 | -0.45771 | -0.36335 | -0.85655 | -0.98023 | 0 | -0.7215 |
| B2m | 0.719262 | 0.733424 | 0.957495 | 1.263367 | 0.564121 | 0 |
| Cd164 | -0.68232 | -0.69844 | -1.42892 | -0.78482 | -0.88582 | 0.5774 |
| Cd52 | 1.003163 | 1.146513 | 2.372275 | 1.7565 | 2.099418 | 1.688261 |
| Cd74 | 1.591875 | 2.274675 | 5.10919 | 3.100736 | 4.671314 | 5.446729 |
| Cd9 | 0.531919 | 0.592123 | 1.657737 | 0.546424 | 0.723925 | 0 |
| Ctsz | 0.34984 | 0.395156 | 1.238292 | 0.477714 | 0.678964 | -0.58874 |
| Cx3cr1 | -0.34961 | -0.30427 | -0.81453 | -0.4146 | 0 | -0.30284 |
| Eng | -0.61909 | -0.33755 | -1.43858 | -0.34611 | 0 | -0.91353 |
| Entpd1 | -0.3086 | -0.29287 | -0.88292 | -0.5602 | 0 | -0.44504 |
| Fam102b | -0.6913 | -0.51314 | -1.44705 | -0.57622 | 0 | -0.69763 |
| Fam110a | -0.69763 | -0.69123 | -2.35764 | -0.87007 | 0 | -1.33936 |
| Fcer1g | 0.331415 | 0.261685 | 0.388975 | 0.255409 | 0 | 0.653079 |
| Fcrls | -1.23854 | -1.0043 | -0.96549 | -1.43784 | -1.09825 | -1.50772 |
| Fermt3 | -0.46637 | -0.40926 | -0.52217 | -0.48557 | 0 | -0.60334 |
| Fth1 | 0.656733 | 0.743374 | 1.847698 | 0.797705 | 0.811785 | 0.68168 |
| Gm10076 | 0.530763 | 0.681361 | 1.014114 | 0.502758 | 0 | 0.748257 |
| Gm26917 | 0.733967 | 0.477451 | 1.228374 | 0.980873 | 1.304377 | 0 |
| Golm1 | -0.28009 | -0.39429 | -0.88725 | -0.45769 | -0.94872 | 0 |
| Gp9 | -0.36454 | -0.86298 | -2.36907 | -0.92734 | 0 | 1.032802 |
| H2-D1 | 1.486782 | 1.536861 | 1.824812 | 2.237341 | 1.517266 | 1.582777 |
| H2-K1 | 1.274552 | 1.277628 | 1.739643 | 2.097964 | 2.005177 | 1.454611 |
| Hfe | -0.49762 | -0.46367 | -1.66776 | -0.37665 | 0 | -1.00934 |
| Hpgd | -0.45599 | -0.36519 | -0.9025 | -0.48562 | 0 | -0.81569 |
| Ifi30 | 0.616825 | 0.511272 | 0.769172 | 0.594614 | 1.552632 | -0.92508 |
| Lgals3bp | 0.908913 | 1.117367 | 1.82203 | 3.151595 | 0 | 1.114446 |
| Lyz2 | 1.340087 | 1.804824 | 2.823566 | 1.177991 | 0 | -1.35799 |
| Maf | -0.52264 | -0.41605 | -1.02457 | -0.44398 | -1.14527 | 0 |
| Mertk | 0 | -0.34848 | -0.62735 | -0.38288 | -1.17914 | -1.09575 |
| Npc2 | 0.507641 | 0.645434 | 0.917283 | 0.666224 | 0 | 0.683182 |
| P2ry12 | -0.69244 | -0.57154 | -1.97909 | -0.79647 | -1.18988 | 0 |
| P2ry13 | -0.75731 | -0.51959 | -1.4448 | -0.54113 | -0.85742 | 0 |
| Plxdc2 | -0.47225 | -0.27254 | -0.36392 | -0.3822 | 0 | -0.73559 |
| Ptgs1 | -0.6365 | -0.39392 | -1.66412 | -0.33322 | -0.8576 | -1.59244 |
| Rpl13 | 0.564876 | 0.553917 | 0.739341 | 0.544118 | 0 | 0.446985 |
| Rpl13a | 0.479053 | 0.543598 | 0.626508 | 0.52495 | 0 | 0.664653 |
| Rpl14 | 0.570669 | 0.545975 | 0.841892 | 0.65316 | 0 | 0.546363 |
| Rpl18a | 0.582444 | 0.618053 | 0.513122 | 0.449931 | 0 | 0.499811 |
| Rpl19 | 0.515159 | 0.531116 | 0.76645 | 0.516465 | 0 | 0.649712 |
| Rpl21 | 0.540041 | 0.545979 | 0.632738 | 0.364431 | 0 | 0.531908 |
| Rpl23a | 0.524462 | 0.567711 | 0.717149 | 0.462778 | 0 | 0.423918 |
| Rpl31 | 0.410098 | 0.494449 | 0.617915 | 0.477549 | 0 | 0.544719 |
| Rpl32 | 0.632672 | 0.714658 | 0.860711 | 0.633712 | 0 | 0.545337 |
| Rpl36 | 0.593094 | 0.616679 | 0.604909 | 0.591037 | 0 | 0.499436 |
| Rpl36-ps2 | 0.455611 | 0.40614 | 0.598946 | 0.489574 | 0 | 0.694618 |

| | | | | | | |
|---|---|---|---|---|---|---|
| Rpl37 | 0.539868 | 0.562669 | 0.475339 | 0.455055 | 0 | 0.452816 |
| Rpl37a | 0.52625 | 0.593614 | 0.565348 | 0.439006 | 0 | 0.298021 |
| Rpl41 | 0.562055 | 0.639709 | 0.999237 | 0.65869 | 0 | 0.639468 |
| Rpl9 | 0.529351 | 0.488519 | 0.751444 | 0.523562 | 0 | 0.596677 |
| Rplp0 | 0.516894 | 0.499567 | 1.078411 | 0.472729 | 0 | 0.54969 |
| Rplp1 | 0.604784 | 0.575806 | 0.697788 | 0.662737 | 0 | 0.439709 |
| Rplp2 | 0.544784 | 0.58314 | 0.690697 | 0.544683 | 0 | 0.4985 |
| Rps11 | 0.579932 | 0.554083 | 0.634669 | 0.487432 | 0 | 0.549245 |
| Rps14 | 0.554939 | 0.588228 | 0.716528 | 0.450637 | 0 | 0.657686 |
| Rps5 | 0.507655 | 0.643484 | 1.068638 | 0.460835 | 0 | 0.558494 |
| Rps8 | 0.520423 | 0.490175 | 0.867548 | 0.528555 | 0 | 0.463388 |
| Rps9 | 0.472374 | 0.593567 | 0.360117 | 0.36576 | 0 | 0.420529 |
| Rpsa | 0.60582 | 0.6468 | 1.395306 | 0.764071 | 0 | 0.692679 |
| Rpsa-ps10 | 0.352871 | 0.549392 | 1.100842 | 0.641312 | 0 | 0.447731 |
| Scamp5 | -0.39338 | -0.43997 | -1.23478 | -0.37814 | 0 | -0.96041 |
| Scoc | -0.40846 | -0.36941 | -0.69433 | -0.67472 | -1.58579 | 0 |
| Selenop | -0.43411 | 0 | -0.74476 | -0.48064 | -0.67703 | -0.63581 |
| Selplg | -0.34153 | -0.32684 | -1.68995 | 0 | -0.50422 | 0.34326 |
| Sh2b3 | -0.49459 | -0.38562 | -0.8881 | -0.60615 | 0 | -0.53812 |
| Sh3bgrl3 | 0.458092 | 0.345625 | 0.724002 | 0.473569 | 0 | 0.569174 |
| Slc2a5 | -0.91933 | -0.8212 | -2.80257 | -0.90866 | -0.84575 | 0.803957 |
| Slc40a1 | -0.50711 | -0.49619 | -1.02859 | -1.0467 | 0 | -1.67323 |
| Sqor | -0.31439 | -0.45375 | -1.21774 | -0.7149 | 0 | -0.97612 |
| Stab1 | -0.37231 | -0.54638 | -0.72151 | -0.78108 | 0 | -1.05552 |
| Tmem204 | -1.18505 | -1.07401 | -3.6097 | -1.56937 | 0 | -1.46563 |
| Tnfsf13b | 0.566494 | 0.75921 | 0.571707 | 0.977361 | 0 | 1.195382 |
| Tspo | 0.858365 | 0.914194 | 1.057933 | 2.165861 | 0 | 1.079901 |
| Uba52 | 0.592881 | 0.579228 | 0.915126 | 0.524716 | 0 | 0.530311 |
| Upk1b | -1.25455 | -1.09015 | -2.36985 | -1.23279 | 0 | 0.415101 |

### Supplementry Table 4. Aging DEGs in Cerebellum

| Gene | p_val | avg_log2FC | pct.1 | pct.2 | p_val_adj |
|---|---|---|---|---|---|
| Gm15564 | 2.5E-148 | -1.41575 | 0.12 | 0.911 | 5.8E-144 |
| C1qa | 2.4E-122 | 1.853858 | 1 | 0.992 | 5.6E-118 |
| C1qc | 2.6E-120 | 1.747765 | 1 | 0.992 | 6.2E-116 |
| Gcg | 1.5E-115 | -2.73344 | 0 | 0.443 | 3.6E-111 |
| H2-D1 | 4.6E-112 | 2.807493 | 0.994 | 0.967 | 1.1E-107 |
| Itm2b | 9.5E-110 | 1.647215 | 1 | 0.996 | 2.3E-105 |
| Ins2 | 1.9E-105 | -1.16339 | 0 | 0.407 | 4.5E-101 |
| Cd81 | 7.4E-104 | 1.826646 | 0.999 | 0.996 | 1.7E-99 |
| Fth1 | 7.7E-100 | 2.327446 | 0.997 | 1 | 1.82E-95 |
| mt-Co1 | 4.22E-98 | 3.060626 | 0.985 | 0.972 | 9.96E-94 |
| B2m | 1.83E-94 | 2.21048 | 0.996 | 1 | 4.32E-90 |
| Cd9 | 4.32E-93 | 2.46679 | 0.989 | 0.939 | 1.02E-88 |
| C1qb | 3.84E-92 | 1.276743 | 1 | 0.992 | 9.06E-88 |
| Gsn | 9.94E-91 | 0.727125 | 0.061 | 0.618 | 2.35E-86 |
| Tmsb4x | 1.99E-90 | 1.623375 | 1 | 1 | 4.7E-86 |
| Ctss | 5.18E-90 | 1.378233 | 1 | 0.992 | 1.22E-85 |
| Ftl1 | 3.06E-88 | 2.124822 | 0.998 | 0.988 | 7.21E-84 |
| Ctsz | 1.33E-87 | 1.932903 | 0.994 | 0.988 | 3.15E-83 |
| mt-Co3 | 3.75E-84 | 2.912468 | 0.955 | 0.809 | 8.86E-80 |
| Cst3 | 6.27E-81 | 1.190546 | 1 | 1 | 1.48E-76 |
| Lgmn | 2.84E-79 | 1.384215 | 0.999 | 0.996 | 6.69E-75 |
| H2-K1 | 3.02E-77 | 2.796553 | 0.951 | 0.907 | 7.12E-73 |
| mt-Cytb | 3.83E-76 | 2.996795 | 0.956 | 0.915 | 9.02E-72 |
| Ins1 | 1.14E-71 | -0.68319 | 0 | 0.28 | 2.68E-67 |
| Ct010467.1 | 3.8E-71 | 1.077502 | 1 | 1 | 8.97E-67 |
| Eef1a1 | 7.04E-71 | 1.741832 | 0.982 | 0.988 | 1.66E-66 |
| Ac154912.3 | 7.06E-70 | -1.37631 | 0.045 | 0.467 | 1.67E-65 |
| Ctsb | 1.03E-66 | 1.714736 | 0.998 | 0.996 | 2.44E-62 |
| Dcn | 3.98E-66 | -0.54632 | 0.006 | 0.297 | 9.38E-62 |
| Tmem119 | 4.64E-66 | 1.795846 | 0.979 | 0.988 | 1.09E-61 |
| Laptm5 | 2.91E-65 | 1.36608 | 0.998 | 0.996 | 6.87E-61 |
| mt-Atp6 | 3.74E-65 | 2.714994 | 0.878 | 0.646 | 8.82E-61 |
| mt-Co2 | 1.82E-63 | 3.137367 | 0.876 | 0.663 | 4.3E-59 |
| Ctsd | 5.48E-63 | 1.242285 | 0.999 | 0.996 | 1.29E-58 |
| Sparc | 1.79E-62 | 1.558502 | 0.987 | 1 | 4.23E-58 |
| Rpl18a | 1.87E-62 | 1.736399 | 0.971 | 0.996 | 4.41E-58 |
| Rpl13 | 6.53E-61 | 1.919899 | 0.952 | 0.935 | 1.54E-56 |
| Cyba | 1.6E-59 | 1.94654 | 0.953 | 0.947 | 3.77E-55 |
| Rn7sk | 3.18E-57 | -0.93987 | 0.16 | 0.695 | 7.51E-53 |
| Cfl1 | 1.33E-56 | 2.885331 | 0.921 | 0.911 | 3.15E-52 |
| Ctsh | 2.69E-56 | 1.748036 | 0.954 | 0.976 | 6.34E-52 |
| Fkbp5 | 3.85E-56 | -0.80071 | 0.159 | 0.667 | 9.08E-52 |
| Hexb | 4.34E-56 | 0.916174 | 1 | 0.992 | 1.02E-51 |

| | | | | | |
|---|---|---|---|---|---|
| Tyrobp | 7.67E-53 | 1.425143 | 0.993 | 0.988 | 1.81E-48 |
| Chga | 9.58E-53 | -0.60052 | 0 | 0.207 | 2.26E-48 |
| Adap2os | 6.43E-52 | 2.680755 | 0.947 | 0.955 | 1.52E-47 |
| Ppy | 1.06E-51 | -0.50043 | 0 | 0.203 | 2.49E-47 |
| Fcer1g | 1.42E-51 | 1.547273 | 0.978 | 0.98 | 3.35E-47 |
| Gm10925 | 1.67E-51 | 2.32933 | 0.88 | 0.675 | 3.95E-47 |
| Gm44860 | 1.26E-50 | -0.98753 | 0.265 | 0.793 | 2.98E-46 |
| Rps9 | 1.34E-50 | 1.627412 | 0.962 | 0.992 | 3.16E-46 |
| Selplg | 5.51E-50 | 1.14387 | 0.996 | 0.992 | 1.3E-45 |
| Hist1h1e | 1.2E-49 | -0.93554 | 0.095 | 0.504 | 2.82E-45 |
| Gm15427 | 2.8E-48 | 2.334009 | 0.936 | 0.955 | 6.6E-44 |
| Trem2 | 6.32E-47 | 1.436874 | 0.963 | 0.976 | 1.49E-42 |
| Gm28437 | 3.92E-46 | 1.927982 | 0.891 | 0.748 | 9.25E-42 |
| Ftl1-ps1 | 4.35E-46 | 2.038969 | 0.918 | 0.874 | 1.03E-41 |
| Serinc3 | 6.22E-46 | -1.05051 | 0.932 | 0.996 | 1.47E-41 |
| Ddit4 | 6.67E-46 | -0.3457 | 0.036 | 0.333 | 1.57E-41 |
| Gm23935 | 2.2E-45 | 0.478417 | 0.864 | 1 | 5.19E-41 |
| mt-Nd1 | 1.34E-44 | 2.531548 | 0.901 | 0.87 | 3.16E-40 |
| Adrb1 | 6.48E-44 | -0.39148 | 0.124 | 0.545 | 1.53E-39 |
| Ctsl | 3.35E-43 | 1.430644 | 0.954 | 0.98 | 7.91E-39 |
| Gm8464 | 5.62E-43 | -0.34471 | 0.011 | 0.224 | 1.33E-38 |
| Lifr | 1.49E-42 | -0.33656 | 0.171 | 0.618 | 3.52E-38 |
| Ac154912.: | 3.7E-42 | -0.38465 | 0.076 | 0.431 | 8.72E-38 |
| Ftl2-ps | 9.13E-42 | 1.862339 | 0.964 | 0.943 | 2.15E-37 |
| Gm24245 | 1.89E-41 | -0.34918 | 0.045 | 0.333 | 4.47E-37 |
| Pfn1 | 3.03E-41 | 2.046108 | 0.916 | 0.919 | 7.15E-37 |
| Gm15710 | 1.01E-40 | 2.251601 | 0.788 | 0.585 | 2.39E-36 |
| Ubb | 5.28E-40 | 1.415949 | 0.964 | 0.988 | 1.25E-35 |
| Pld4 | 8.47E-40 | 1.636016 | 0.94 | 0.927 | 2E-35 |
| Rpl13a | 1.38E-39 | 2.829219 | 0.823 | 0.772 | 3.26E-35 |
| Lyz2 | 1.42E-39 | 2.611937 | 0.866 | 0.858 | 3.35E-35 |
| Gm1821 | 7.55E-39 | 1.750526 | 0.935 | 0.959 | 1.78E-34 |
| Malat1 | 8.18E-39 | 1.174955 | 0.982 | 0.992 | 1.93E-34 |
| Ctsa | 1.22E-38 | 1.495706 | 0.948 | 0.959 | 2.87E-34 |
| Tlr7 | 1.78E-37 | -0.57398 | 0.224 | 0.659 | 4.2E-33 |
| Gm10076 | 2.2E-37 | 1.826505 | 0.93 | 0.939 | 5.2E-33 |
| Grn | 2.48E-37 | 1.141457 | 0.98 | 0.996 | 5.86E-33 |
| Akna | 2.92E-37 | -0.2813 | 0.177 | 0.593 | 6.9E-33 |
| Rnu3a | 3.03E-37 | -1.30516 | 0.012 | 0.203 | 7.16E-33 |
| Rplp1 | 3.37E-37 | 1.566727 | 0.928 | 0.923 | 7.96E-33 |
| Runx1 | 6.98E-37 | -1.042 | 0.208 | 0.63 | 1.65E-32 |
| Rps18 | 1.5E-36 | 1.848362 | 0.88 | 0.837 | 3.53E-32 |
| Jmjd1c | 2.5E-36 | -0.33916 | 0.151 | 0.549 | 5.9E-32 |
| Gm9843 | 1.09E-35 | 2.509165 | 0.839 | 0.679 | 2.56E-31 |
| Fos | 1.7E-35 | -3.00161 | 0.131 | 0.508 | 4.02E-31 |
| Cd63 | 4.02E-35 | 2.409304 | 0.85 | 0.797 | 9.48E-31 |
| Rplp0 | 5.42E-35 | 1.778761 | 0.891 | 0.894 | 1.28E-30 |

| | | | | | |
|---|---|---|---|---|---|
| Aif1 | 6.22E-35 | 2.016538 | 0.874 | 0.829 | 1.47E-30 |
| Rps11 | 8.37E-35 | 1.563963 | 0.916 | 0.907 | 1.97E-30 |
| Olfml3 | 1.32E-34 | 1.206352 | 0.967 | 0.988 | 3.1E-30 |
| Gm43813 | 4.91E-34 | -0.51623 | 0.068 | 0.358 | 1.16E-29 |
| Jun | 5.28E-34 | -1.45783 | 0.234 | 0.61 | 1.25E-29 |
| Lamp1 | 1.79E-33 | 1.50678 | 0.965 | 0.972 | 4.22E-29 |
| Rnaset2a | 3.7E-33 | 1.990393 | 0.895 | 0.874 | 8.73E-29 |
| Sat1 | 5.67E-33 | 2.222304 | 0.888 | 0.878 | 1.34E-28 |
| Sbno2 | 6.59E-33 | -0.48895 | 0.147 | 0.512 | 1.55E-28 |
| Cd68 | 1.8E-32 | 1.543406 | 0.924 | 0.911 | 4.25E-28 |
| Rps3 | 2.99E-32 | 1.959612 | 0.86 | 0.898 | 7.05E-28 |
| Gm13456 | 1.08E-31 | 1.445988 | 0.95 | 0.959 | 2.54E-27 |
| 0610039K1 | 1.25E-31 | -1.04824 | 0.029 | 0.24 | 2.96E-27 |
| March1 | 3.36E-31 | -0.97468 | 0.174 | 0.545 | 7.92E-27 |
| P2ry12 | 8.12E-31 | 1.274377 | 0.985 | 0.984 | 1.92E-26 |
| Klf2 | 1.38E-30 | -0.75046 | 0.066 | 0.329 | 3.25E-26 |
| Abca1 | 1.67E-30 | -0.66105 | 0.252 | 0.646 | 3.95E-26 |
| Rgs10 | 8.02E-30 | 1.62688 | 0.888 | 0.87 | 1.89E-25 |
| Slc8a1 | 3.18E-29 | -1.65971 | 0.092 | 0.374 | 7.51E-25 |
| Rps3a1 | 6.43E-29 | 1.907011 | 0.828 | 0.797 | 1.52E-24 |
| Nisch | 7.5E-29 | 0.260968 | 0.265 | 0.687 | 1.77E-24 |
| Gm8797 | 1.05E-28 | 1.692104 | 0.854 | 0.894 | 2.48E-24 |
| Tagap | 1.82E-28 | -0.73537 | 0.175 | 0.52 | 4.3E-24 |
| Zfp36l2 | 2.17E-28 | 0.549885 | 0.281 | 0.756 | 5.13E-24 |
| Rps14 | 4.18E-28 | 1.385959 | 0.915 | 0.923 | 9.87E-24 |
| Rpl41 | 5.78E-28 | 1.392128 | 0.955 | 0.963 | 1.36E-23 |
| Rps20 | 9.95E-28 | 1.966447 | 0.834 | 0.785 | 2.35E-23 |
| Psap | 1.11E-27 | 1.102581 | 0.994 | 0.996 | 2.61E-23 |
| Abca9 | 1.22E-27 | -0.45248 | 0.409 | 0.825 | 2.89E-23 |
| Rps27rt | 2.75E-27 | 1.907456 | 0.723 | 0.537 | 6.48E-23 |
| Npc2 | 2.98E-27 | 2.237958 | 0.815 | 0.797 | 7.02E-23 |
| Gm44861 | 3.32E-27 | -0.71103 | 0.041 | 0.252 | 7.82E-23 |
| Dhx9 | 3.32E-27 | -0.64012 | 0.076 | 0.333 | 7.83E-23 |
| Rps18-ps3 | 3.61E-27 | 2.298733 | 0.643 | 0.398 | 8.52E-23 |
| Nptn | 3.67E-27 | -0.86079 | 0.382 | 0.736 | 8.66E-23 |
| Dusp1 | 4.45E-27 | -1.81031 | 0.129 | 0.439 | 1.05E-22 |
| Rnaset2b | 4.86E-27 | 2.061368 | 0.904 | 0.854 | 1.15E-22 |
| Rpl3 | 6.17E-27 | 2.125217 | 0.815 | 0.817 | 1.45E-22 |
| Skil | 6.41E-27 | 2.383291 | 0.839 | 0.711 | 1.51E-22 |
| Gm45223 | 1.91E-26 | -0.41 | 0.093 | 0.366 | 4.49E-22 |
| Ptafr | 2.05E-26 | 2.47661 | 0.802 | 0.663 | 4.84E-22 |
| Dpysl2 | 4.47E-26 | 2.304653 | 0.876 | 0.829 | 1.06E-21 |
| Sun2 | 4.81E-26 | -0.67601 | 0.323 | 0.687 | 1.13E-21 |
| Gab2 | 7.81E-26 | -0.27533 | 0.098 | 0.374 | 1.84E-21 |
| Per1 | 1.19E-25 | -0.26862 | 0.04 | 0.24 | 2.82E-21 |
| Gm42908 | 1.56E-25 | 2.122818 | 0.123 | 0.435 | 3.68E-21 |
| Gm28661 | 2.4E-25 | 1.616137 | 0.765 | 0.642 | 5.67E-21 |

| | | | | | |
|---|---|---|---|---|---|
| Ly86 | 3.25E-25 | 1.011205 | 0.969 | 0.98 | 7.67E-21 |
| S100a9 | 3.58E-25 | 1.180533 | 0.053 | 0.28 | 8.44E-21 |
| Tmem50a | 3.81E-25 | 1.88121 | 0.818 | 0.797 | 8.98E-21 |
| Chka | 7.05E-25 | -0.60348 | 0.118 | 0.398 | 1.66E-20 |
| Basp1 | 8.71E-25 | 2.272563 | 0.848 | 0.911 | 2.05E-20 |
| Fau | 1.58E-24 | 1.429503 | 0.893 | 0.854 | 3.73E-20 |
| G530011O( | 3.14E-24 | -0.25962 | 0.178 | 0.508 | 7.41E-20 |
| Tab2 | 4.49E-24 | 0.708326 | 0.183 | 0.528 | 1.06E-19 |
| Hist1h4d | 6.17E-24 | -1.1064 | 0.187 | 0.516 | 1.46E-19 |
| Sfpq | 8.44E-24 | 0.410369 | 0.166 | 0.5 | 1.99E-19 |
| Rpl35 | 1.12E-23 | 1.767814 | 0.857 | 0.789 | 2.63E-19 |
| Rapgef5 | 1.32E-23 | -0.3423 | 0.164 | 0.476 | 3.12E-19 |
| Gm4149 | 1.86E-23 | 1.674577 | 0.81 | 0.756 | 4.39E-19 |
| Gm13340 | 2.01E-23 | 1.261134 | 0.711 | 0.533 | 4.75E-19 |
| Rassf2 | 2.19E-23 | 0.493294 | 0.241 | 0.614 | 5.18E-19 |
| Zfp36 | 5.65E-23 | 0.486619 | 0.304 | 0.732 | 1.33E-18 |
| Rps25 | 7.34E-23 | 1.588249 | 0.867 | 0.87 | 1.73E-18 |
| Zfp91 | 8.21E-23 | -0.68871 | 0.087 | 0.329 | 1.94E-18 |
| Atp6v0c-ps | 1.13E-22 | 2.229001 | 0.638 | 0.476 | 2.66E-18 |
| Rps19 | 1.36E-22 | 1.36797 | 0.89 | 0.874 | 3.21E-18 |
| Atp6v0c | 1.41E-22 | 1.825365 | 0.809 | 0.821 | 3.32E-18 |
| Gm29216 | 1.97E-22 | 1.294318 | 0.74 | 0.589 | 4.64E-18 |
| Apc | 2.39E-22 | -0.77463 | 0.121 | 0.39 | 5.63E-18 |
| Gm47664 | 4.36E-22 | -0.34849 | 0.047 | 0.24 | 1.03E-17 |
| Gm2a | 5.61E-22 | 2.057421 | 0.786 | 0.573 | 1.32E-17 |
| Calm2 | 1.01E-21 | 1.852447 | 0.873 | 0.915 | 2.39E-17 |
| Socs3 | 1.34E-21 | -0.95251 | 0.065 | 0.272 | 3.16E-17 |
| 6820431F2 | 3.07E-21 | -0.2574 | 0.156 | 0.443 | 7.24E-17 |
| Egr1 | 3.29E-21 | -0.38661 | 0.056 | 0.252 | 7.75E-17 |
| Mdm4 | 4.16E-21 | 0.43749 | 0.183 | 0.508 | 9.82E-17 |
| Zfp445 | 5.08E-21 | -0.51441 | 0.142 | 0.411 | 1.2E-16 |
| Ppia | 5.53E-21 | 0.947491 | 0.95 | 0.967 | 1.3E-16 |
| Hexa | 6.57E-21 | 1.187797 | 0.931 | 0.955 | 1.55E-16 |
| Atp8a1 | 6.86E-21 | 0.603368 | 0.263 | 0.63 | 1.62E-16 |
| Numb | 7.41E-21 | -0.268 | 0.309 | 0.667 | 1.75E-16 |
| Nmt1 | 7.55E-21 | 0.565682 | 0.309 | 0.671 | 1.78E-16 |
| Cd63-ps | 7.65E-21 | 1.688035 | 0.638 | 0.455 | 1.8E-16 |
| Dhrs3 | 8.08E-21 | 2.154343 | 0.848 | 0.797 | 1.91E-16 |
| Tbc1d14 | 1.06E-20 | -0.42489 | 0.161 | 0.443 | 2.5E-16 |
| Wdr26 | 1.27E-20 | 0.334631 | 0.214 | 0.549 | 2.99E-16 |
| Csf2rb2 | 1.34E-20 | -0.27188 | 0.092 | 0.325 | 3.17E-16 |
| Bsg | 1.62E-20 | 2.027589 | 0.806 | 0.785 | 3.83E-16 |
| S100a8 | 1.77E-20 | 1.271623 | 0.065 | 0.28 | 4.18E-16 |
| Rpl30 | 1.91E-20 | 1.42613 | 0.869 | 0.837 | 4.51E-16 |
| Parp8 | 2.16E-20 | 0.349163 | 0.099 | 0.341 | 5.09E-16 |
| Zfp691 | 2.19E-20 | 1.52944 | 0.647 | 0.407 | 5.16E-16 |
| Hnrnpm | 2.23E-20 | 0.549183 | 0.269 | 0.638 | 5.27E-16 |

| | | | | | |
|---|---|---|---|---|---|
| Ccr5 | 3.37E-20 | -0.46303 | 0.467 | 0.797 | 7.94E-16 |
| Gm37472 | 3.46E-20 | 0.851315 | 0.101 | 0.35 | 8.17E-16 |
| Zscan26 | 3.62E-20 | 2.336951 | 0.808 | 0.691 | 8.53E-16 |
| Pik3cg | 3.62E-20 | -0.49692 | 0.164 | 0.435 | 8.53E-16 |
| Sik2 | 3.99E-20 | -0.71198 | 0.058 | 0.252 | 9.4E-16 |
| Ssfa2 | 4.97E-20 | 0.844505 | 0.099 | 0.341 | 1.17E-15 |
| Fmnl1 | 7.18E-20 | 1.311434 | 0.228 | 0.581 | 1.69E-15 |
| Syngr1 | 8.82E-20 | 2.171922 | 0.771 | 0.707 | 2.08E-15 |
| Rpl4 | 9.47E-20 | 1.934048 | 0.801 | 0.837 | 2.23E-15 |
| Gm45221 | 1.01E-19 | -1.88948 | 0.058 | 0.244 | 2.38E-15 |
| Kat6a | 1.03E-19 | -0.57802 | 0.135 | 0.39 | 2.42E-15 |
| Otulinl | 1.21E-19 | 1.590533 | 0.883 | 0.89 | 2.86E-15 |
| Srsf2 | 1.73E-19 | 2.088425 | 0.876 | 0.858 | 4.07E-15 |
| Gm11478 | 1.73E-19 | 2.40675 | 0.61 | 0.463 | 4.07E-15 |
| Rpsa | 1.82E-19 | 1.9068 | 0.781 | 0.821 | 4.3E-15 |
| Bhlhe41 | 1.94E-19 | 2.190557 | 0.862 | 0.854 | 4.57E-15 |
| Gm5559 | 1.95E-19 | 2.48943 | 0.591 | 0.423 | 4.59E-15 |
| Hnrnpul2 | 2.04E-19 | 0.262258 | 0.049 | 0.228 | 4.81E-15 |
| Clic1 | 2.08E-19 | 1.795906 | 0.778 | 0.72 | 4.91E-15 |
| Ifngr1 | 2.11E-19 | 1.22863 | 0.943 | 0.972 | 4.99E-15 |
| Il4ra | 2.67E-19 | -0.42639 | 0.249 | 0.557 | 6.3E-15 |
| Rpl23a-ps3 | 2.92E-19 | 1.803137 | 0.507 | 0.268 | 6.9E-15 |
| Plek | 3.17E-19 | 0.576086 | 0.378 | 0.785 | 7.49E-15 |
| Rps7 | 3.43E-19 | 1.952847 | 0.782 | 0.813 | 8.1E-15 |
| Serpine2 | 4.78E-19 | 1.609396 | 0.848 | 0.813 | 1.13E-14 |
| Gm17147 | 5.86E-19 | -0.31113 | 0.084 | 0.301 | 1.38E-14 |
| Gpr155 | 9.44E-19 | -0.46295 | 0.089 | 0.305 | 2.23E-14 |
| Scamp5 | 1.01E-18 | 0.333979 | 0.244 | 0.565 | 2.39E-14 |
| Gm5879 | 1.06E-18 | 1.042292 | 0.41 | 0.13 | 2.5E-14 |
| Abhd12 | 1.23E-18 | 1.215497 | 0.911 | 0.955 | 2.89E-14 |
| Rpl10 | 1.34E-18 | 1.495147 | 0.842 | 0.882 | 3.17E-14 |
| Spi1 | 1.89E-18 | 2.078744 | 0.785 | 0.768 | 4.45E-14 |
| Gramd1a | 2.2E-18 | 0.375438 | 0.112 | 0.354 | 5.18E-14 |
| Cmtm6 | 2.52E-18 | -0.44869 | 0.544 | 0.809 | 5.94E-14 |
| Rpl6 | 2.66E-18 | 2.059673 | 0.737 | 0.663 | 6.28E-14 |
| Ywhah | 3.08E-18 | 1.518435 | 0.862 | 0.894 | 7.27E-14 |
| Elmo1 | 3.44E-18 | 0.301632 | 0.438 | 0.797 | 8.11E-14 |
| Pik3r1 | 3.82E-18 | 0.548371 | 0.336 | 0.703 | 9.02E-14 |
| Mpeg1 | 4.34E-18 | -0.31829 | 0.685 | 0.951 | 1.02E-13 |
| Ly6e | 4.66E-18 | 1.60691 | 0.912 | 0.984 | 1.1E-13 |
| Cerk | 5.04E-18 | 0.289543 | 0.176 | 0.463 | 1.19E-13 |
| Gm15266 | 5.39E-18 | 1.699228 | 0.553 | 0.358 | 1.27E-13 |
| Rpl37a | 5.41E-18 | 1.144451 | 0.931 | 0.923 | 1.28E-13 |
| Gm10123 | 6.44E-18 | 1.63122 | 0.884 | 0.927 | 1.52E-13 |
| Rps6 | 6.52E-18 | 1.596915 | 0.762 | 0.642 | 1.54E-13 |
| Atf6 | 6.82E-18 | 0.266761 | 0.103 | 0.333 | 1.61E-13 |
| Rpl3-ps1 | 7.13E-18 | 2.0499 | 0.692 | 0.598 | 1.68E-13 |

| | | | | | |
|---|---|---|---|---|---|
| Sft2d1 | 8.38E-18 | 1.74384 | 0.782 | 0.809 | 1.98E-13 |
| Tmem245 | 9.86E-18 | -0.45477 | 0.114 | 0.346 | 2.32E-13 |
| Gm37420 | 1.33E-17 | -0.51084 | 0.089 | 0.301 | 3.13E-13 |
| Picalm | 1.39E-17 | 0.42725 | 0.541 | 0.943 | 3.27E-13 |
| Gm6977 | 1.5E-17 | 1.915321 | 0.631 | 0.512 | 3.55E-13 |
| Nek7 | 1.55E-17 | 1.059916 | 0.206 | 0.512 | 3.66E-13 |
| Arsb | 1.9E-17 | -0.29145 | 0.268 | 0.557 | 4.48E-13 |
| Gbp7 | 1.99E-17 | 0.406922 | 0.135 | 0.382 | 4.7E-13 |
| Tsc22d2 | 2.28E-17 | -0.39431 | 0.146 | 0.394 | 5.38E-13 |
| Rpl21 | 2.3E-17 | 1.303495 | 0.88 | 0.931 | 5.43E-13 |
| Gm6900 | 2.9E-17 | 0.945824 | 0.273 | 0.024 | 6.85E-13 |
| Ppp1r10 | 2.9E-17 | 0.798834 | 0.129 | 0.374 | 6.85E-13 |
| F11r | 2.98E-17 | 1.651467 | 0.886 | 0.919 | 7.02E-13 |
| Tbc1d5 | 3.14E-17 | 0.474292 | 0.118 | 0.358 | 7.4E-13 |
| Wipf1 | 3.85E-17 | 1.612904 | 0.169 | 0.451 | 9.08E-13 |
| Laptm4a | 4.34E-17 | 1.686832 | 0.824 | 0.874 | 1.02E-12 |
| Sod1 | 4.4E-17 | 2.422648 | 0.653 | 0.492 | 1.04E-12 |
| Rgs1 | 4.53E-17 | -0.27919 | 0.065 | 0.252 | 1.07E-12 |
| Ppp1r12c | 4.87E-17 | 0.518374 | 0.13 | 0.374 | 1.15E-12 |
| Cltc | 7.24E-17 | 0.728031 | 0.391 | 0.752 | 1.71E-12 |
| Diaph2 | 7.3E-17 | 0.71681 | 0.131 | 0.378 | 1.72E-12 |
| H2-Q7 | 7.44E-17 | 4.811146 | 0.337 | 0.085 | 1.75E-12 |
| Xpr1 | 8.75E-17 | 0.349274 | 0.083 | 0.28 | 2.07E-12 |
| Rpl23a | 9.23E-17 | 1.63444 | 0.784 | 0.764 | 2.18E-12 |
| Kdm3a | 1.03E-16 | -1.01417 | 0.06 | 0.232 | 2.42E-12 |
| Bod1l | 1.03E-16 | 0.714144 | 0.118 | 0.354 | 2.43E-12 |
| Rps27 | 1.03E-16 | 1.119585 | 0.964 | 0.955 | 2.44E-12 |
| Gm16378 | 1.04E-16 | 0.803596 | 0.266 | 0.024 | 2.46E-12 |
| Gm43028 | 1.1E-16 | 1.039914 | 0.362 | 0.106 | 2.59E-12 |
| Gm44705 | 1.11E-16 | 0.919083 | 0.254 | 0.016 | 2.62E-12 |
| Fbxl14 | 1.13E-16 | -0.79024 | 0.07 | 0.248 | 2.66E-12 |
| Rhob | 1.15E-16 | 0.562927 | 0.678 | 0.98 | 2.72E-12 |
| Tm6sf1 | 1.2E-16 | 0.293452 | 0.249 | 0.573 | 2.82E-12 |
| Marcks | 1.27E-16 | 1.12397 | 0.966 | 0.996 | 3E-12 |
| Lgals3bp | 1.28E-16 | 2.567349 | 0.719 | 0.663 | 3.01E-12 |
| Atrx | 1.34E-16 | 1.166192 | 0.203 | 0.504 | 3.16E-12 |
| Atp6v0b | 1.35E-16 | 1.815216 | 0.838 | 0.833 | 3.17E-12 |
| D1Ertd622( | 1.55E-16 | -0.56039 | 0.117 | 0.333 | 3.66E-12 |
| Map4k4 | 1.62E-16 | 0.360584 | 0.268 | 0.602 | 3.82E-12 |
| Map3k3 | 1.99E-16 | 0.684861 | 0.189 | 0.467 | 4.7E-12 |
| Bri3 | 2E-16 | 2.904392 | 0.605 | 0.476 | 4.73E-12 |
| Tle4 | 2.2E-16 | 0.281046 | 0.088 | 0.28 | 5.19E-12 |
| Rpl32 | 2.37E-16 | 1.336516 | 0.864 | 0.911 | 5.59E-12 |
| Rpl18 | 2.67E-16 | 1.549695 | 0.767 | 0.715 | 6.3E-12 |
| Cd2ap | 2.85E-16 | 0.470347 | 0.204 | 0.496 | 6.72E-12 |
| Setd5 | 3.6E-16 | 0.539358 | 0.2 | 0.484 | 8.5E-12 |
| Adap2 | 3.61E-16 | 1.75648 | 0.878 | 0.939 | 8.52E-12 |

| | | | | | |
|---|---|---|---|---|---|
| Rplp2 | 3.94E-16 | 1.365017 | 0.876 | 0.919 | 9.3E-12 |
| Siglech | 4.3E-16 | 0.924998 | 0.971 | 0.992 | 1.01E-11 |
| 9930111J2 | 4.69E-16 | -0.66084 | 0.049 | 0.207 | 1.11E-11 |
| Itsn2 | 5.19E-16 | 0.29636 | 0.185 | 0.451 | 1.22E-11 |
| Zdhhc9 | 5.49E-16 | 0.498303 | 0.132 | 0.374 | 1.29E-11 |
| Cotl1 | 6.29E-16 | 2.45567 | 0.718 | 0.679 | 1.48E-11 |
| Phf14 | 6.54E-16 | 0.554766 | 0.193 | 0.472 | 1.54E-11 |
| Rpl17 | 7.45E-16 | 1.74675 | 0.777 | 0.78 | 1.76E-11 |
| Extl3 | 8.65E-16 | 0.424825 | 0.198 | 0.48 | 2.04E-11 |
| Prpf4b | 9.9E-16 | 0.598616 | 0.205 | 0.48 | 2.33E-11 |
| Spen | 1.04E-15 | 0.426961 | 0.069 | 0.248 | 2.46E-11 |
| Cmklr1 | 1.16E-15 | 0.519033 | 0.134 | 0.366 | 2.75E-11 |
| Rps23 | 1.61E-15 | 1.182986 | 0.878 | 0.874 | 3.81E-11 |
| Aff4 | 1.75E-15 | 0.460803 | 0.124 | 0.354 | 4.13E-11 |
| Unc93b1 | 1.76E-15 | 0.918802 | 0.969 | 0.988 | 4.14E-11 |
| Arhgap12 | 1.96E-15 | -0.32557 | 0.122 | 0.337 | 4.62E-11 |
| Cep170 | 2.18E-15 | 0.261232 | 0.145 | 0.382 | 5.15E-11 |
| Tlr13 | 2.67E-15 | -0.5207 | 0.156 | 0.382 | 6.29E-11 |
| Lnpep | 2.74E-15 | 0.51228 | 0.299 | 0.602 | 6.47E-11 |
| Rpl36 | 2.76E-15 | 1.352362 | 0.851 | 0.813 | 6.5E-11 |
| Prrc2a | 3.26E-15 | 1.010502 | 0.125 | 0.35 | 7.68E-11 |
| Gm12918 | 3.81E-15 | 1.270591 | 0.548 | 0.354 | 8.99E-11 |
| Tjp1 | 3.82E-15 | 1.014627 | 0.144 | 0.39 | 9.02E-11 |
| Hnrnpl | 4.2E-15 | 1.613935 | 0.269 | 0.573 | 9.92E-11 |
| Usp8 | 4.38E-15 | 0.672312 | 0.166 | 0.423 | 1.03E-10 |
| Tubb5 | 4.45E-15 | 1.78053 | 0.78 | 0.679 | 1.05E-10 |
| Phykpl | 4.8E-15 | 0.798941 | 0.094 | 0.285 | 1.13E-10 |
| Nktr | 5.25E-15 | 0.466483 | 0.169 | 0.423 | 1.24E-10 |
| Ubr4 | 5.55E-15 | -0.86566 | 0.097 | 0.289 | 1.31E-10 |
| Hps3 | 5.74E-15 | 0.563054 | 0.276 | 0.581 | 1.35E-10 |
| Rpl7 | 6.09E-15 | 1.588154 | 0.789 | 0.789 | 1.44E-10 |
| Rasa1 | 6.43E-15 | 0.402731 | 0.229 | 0.516 | 1.52E-10 |
| Tmbim6 | 6.97E-15 | 1.266337 | 0.923 | 0.915 | 1.64E-10 |
| Rock2 | 7.23E-15 | 0.438782 | 0.275 | 0.573 | 1.71E-10 |
| Gm14011 | 7.29E-15 | -0.54297 | 0.061 | 0.224 | 1.72E-10 |
| Itm2c | 7.49E-15 | 1.651668 | 0.822 | 0.854 | 1.77E-10 |
| Gm3362 | 7.75E-15 | 1.103473 | 0.401 | 0.163 | 1.83E-10 |
| Dock10 | 9.09E-15 | 0.520146 | 0.398 | 0.744 | 2.14E-10 |
| Setd2 | 9.43E-15 | 0.404143 | 0.158 | 0.394 | 2.22E-10 |
| Lpcat2 | 1.01E-14 | 0.963198 | 0.95 | 0.963 | 2.38E-10 |
| Kmt2a | 1.05E-14 | 0.344776 | 0.156 | 0.394 | 2.48E-10 |
| H3f3b | 1.08E-14 | 1.508169 | 0.838 | 0.841 | 2.55E-10 |
| Gnb1 | 1.11E-14 | 2.264073 | 0.837 | 0.874 | 2.62E-10 |
| Hnrnpu | 1.19E-14 | -0.26391 | 0.253 | 0.524 | 2.82E-10 |
| Pkn2 | 1.35E-14 | 0.292473 | 0.114 | 0.325 | 3.18E-10 |
| Stab1 | 1.39E-14 | 0.542657 | 0.426 | 0.756 | 3.28E-10 |
| Mms19 | 1.42E-14 | -0.30599 | 0.11 | 0.309 | 3.34E-10 |

| | | | | | |
|---|---|---|---|---|---|
| Gm12537 | 1.48E-14 | 1.134109 | 0.3 | 0.073 | 3.49E-10 |
| Sidt2 | 1.52E-14 | -0.2834 | 0.1 | 0.293 | 3.58E-10 |
| Eif4g3 | 1.74E-14 | 0.610444 | 0.213 | 0.492 | 4.1E-10 |
| Polr2a | 1.77E-14 | 0.979139 | 0.178 | 0.439 | 4.17E-10 |
| Chd2 | 1.99E-14 | 0.875277 | 0.147 | 0.39 | 4.7E-10 |
| Mob1a | 2.01E-14 | 0.32295 | 0.332 | 0.622 | 4.74E-10 |
| Zfp871 | 2.01E-14 | -0.84948 | 0.081 | 0.256 | 4.75E-10 |
| Klhl18 | 2.03E-14 | 0.61246 | 0.098 | 0.293 | 4.78E-10 |
| Tbc1d10b | 2.28E-14 | -0.26785 | 0.054 | 0.207 | 5.39E-10 |
| Xiap | 2.35E-14 | 0.348593 | 0.194 | 0.443 | 5.54E-10 |
| Slc25a5 | 2.42E-14 | 1.897555 | 0.741 | 0.703 | 5.72E-10 |
| Gm34280 | 2.5E-14 | 0.350975 | 0.124 | 0.341 | 5.89E-10 |
| Zfp106 | 2.5E-14 | 0.401807 | 0.177 | 0.431 | 5.89E-10 |
| Ppp1r9a | 2.54E-14 | 0.498738 | 0.204 | 0.467 | 5.99E-10 |
| Gm11942 | 2.61E-14 | 1.313789 | 0.46 | 0.248 | 6.15E-10 |
| Ang | 2.62E-14 | 2.494037 | 0.747 | 0.764 | 6.18E-10 |
| Rnf216 | 2.64E-14 | 0.289626 | 0.144 | 0.37 | 6.23E-10 |
| Tet3 | 2.67E-14 | -0.31205 | 0.137 | 0.35 | 6.29E-10 |
| Ftl1-ps2 | 2.75E-14 | 0.783472 | 0.42 | 0.183 | 6.49E-10 |
| Frmd4a | 3.41E-14 | 0.364122 | 0.54 | 0.898 | 8.05E-10 |
| 5031439G( | 3.5E-14 | 0.796612 | 0.103 | 0.301 | 8.25E-10 |
| Gm6204 | 3.69E-14 | 1.802594 | 0.551 | 0.419 | 8.71E-10 |
| Heatr5a | 3.9E-14 | 0.56381 | 0.272 | 0.581 | 9.2E-10 |
| Apba1 | 4.03E-14 | 0.29109 | 0.083 | 0.264 | 9.5E-10 |
| Rpl23 | 4.53E-14 | 1.66073 | 0.799 | 0.785 | 1.07E-09 |
| Rreb1 | 5.41E-14 | 0.439477 | 0.163 | 0.398 | 1.28E-09 |
| Rps24 | 5.49E-14 | 1.746327 | 0.793 | 0.833 | 1.3E-09 |
| Tm9sf3 | 5.54E-14 | 0.319184 | 0.278 | 0.569 | 1.31E-09 |
| Crkl | 6.16E-14 | -0.62887 | 0.062 | 0.215 | 1.45E-09 |
| Rps16-ps2 | 6.39E-14 | 1.773008 | 0.579 | 0.439 | 1.51E-09 |
| Atp6v0a1 | 6.41E-14 | 1.337119 | 0.188 | 0.455 | 1.51E-09 |
| Itga9 | 6.72E-14 | -0.25873 | 0.074 | 0.244 | 1.58E-09 |
| Rps16 | 7.2E-14 | 1.331682 | 0.783 | 0.687 | 1.7E-09 |
| Gm8730 | 8.29E-14 | 1.553391 | 0.719 | 0.72 | 1.96E-09 |
| Sult1a1 | 8.3E-14 | 0.640645 | 0.138 | 0.366 | 1.96E-09 |
| Ranbp2 | 9.05E-14 | -0.46975 | 0.099 | 0.28 | 2.14E-09 |
| Arih1 | 9.56E-14 | -0.28701 | 0.084 | 0.26 | 2.25E-09 |
| Stk4 | 1.09E-13 | 0.542214 | 0.09 | 0.272 | 2.56E-09 |
| Arhgef2 | 1.2E-13 | 0.539203 | 0.231 | 0.5 | 2.83E-09 |
| Cdc37l1 | 1.21E-13 | 1.256632 | 0.084 | 0.264 | 2.85E-09 |
| Fus | 1.24E-13 | 0.98597 | 0.335 | 0.642 | 2.93E-09 |
| Strn3 | 1.3E-13 | 0.406478 | 0.157 | 0.394 | 3.08E-09 |
| Itga6 | 1.4E-13 | 0.336645 | 0.382 | 0.699 | 3.3E-09 |
| Gm2000 | 1.46E-13 | 1.582642 | 0.61 | 0.455 | 3.45E-09 |
| Rcan1 | 1.52E-13 | -1.18373 | 0.06 | 0.207 | 3.58E-09 |
| Usp4 | 1.53E-13 | 0.400477 | 0.192 | 0.439 | 3.6E-09 |
| Abl2 | 1.58E-13 | 0.260668 | 0.075 | 0.24 | 3.73E-09 |

| | | | | | |
|---|---|---|---|---|---|
| Gm26857 | 1.59E-13 | 0.465985 | 0.093 | 0.272 | 3.76E-09 |
| Stau1 | 1.72E-13 | 0.575961 | 0.166 | 0.402 | 4.06E-09 |
| Cop1 | 1.74E-13 | 0.370528 | 0.107 | 0.301 | 4.11E-09 |
| Gapdh | 1.75E-13 | 1.300807 | 0.819 | 0.85 | 4.14E-09 |
| Zfp292 | 1.82E-13 | 0.561944 | 0.126 | 0.333 | 4.29E-09 |
| Mlxipl | 1.84E-13 | 1.409166 | 0.186 | 0.451 | 4.34E-09 |
| Zbtb18 | 1.85E-13 | 0.496692 | 0.096 | 0.28 | 4.37E-09 |
| Nav3 | 2.03E-13 | 0.616858 | 0.443 | 0.801 | 4.8E-09 |
| Mkln1 | 2.12E-13 | 0.369876 | 0.21 | 0.467 | 5E-09 |
| Arhgap39 | 2.14E-13 | 0.304169 | 0.245 | 0.52 | 5.04E-09 |
| Tns3 | 2.27E-13 | 0.839517 | 0.197 | 0.451 | 5.35E-09 |
| Arfgef1 | 2.31E-13 | 0.257862 | 0.073 | 0.236 | 5.44E-09 |
| Ac165079.: | 2.42E-13 | 0.96889 | 0.288 | 0.073 | 5.7E-09 |
| Dpy19l4 | 2.5E-13 | 0.863453 | 0.135 | 0.35 | 5.89E-09 |
| Tmx3 | 2.56E-13 | 0.594527 | 0.093 | 0.276 | 6.03E-09 |
| Bmf | 2.56E-13 | 0.364672 | 0.076 | 0.24 | 6.04E-09 |
| Taok1 | 2.58E-13 | 0.655178 | 0.307 | 0.606 | 6.09E-09 |
| Fam208a | 2.6E-13 | 0.497065 | 0.248 | 0.541 | 6.12E-09 |
| Ttyh3 | 2.84E-13 | 0.643134 | 0.153 | 0.378 | 6.7E-09 |
| Tnrc6a | 2.9E-13 | 0.264826 | 0.181 | 0.407 | 6.84E-09 |
| Rpl19 | 2.95E-13 | 1.422397 | 0.789 | 0.805 | 6.97E-09 |
| Il17ra | 2.96E-13 | 0.27342 | 0.303 | 0.573 | 6.99E-09 |
| Tmem87b | 3.01E-13 | 0.479411 | 0.112 | 0.305 | 7.1E-09 |
| Chd7 | 3.08E-13 | 0.778202 | 0.285 | 0.585 | 7.26E-09 |
| Prdm2 | 3.13E-13 | -0.3949 | 0.093 | 0.268 | 7.38E-09 |
| Hsd17b11 | 3.49E-13 | 0.602354 | 0.112 | 0.305 | 8.23E-09 |
| Pcnx3 | 3.78E-13 | 0.25975 | 0.114 | 0.305 | 8.92E-09 |
| Numa1 | 4.11E-13 | 0.569623 | 0.15 | 0.37 | 9.7E-09 |
| Rlf | 4.53E-13 | 0.436837 | 0.076 | 0.24 | 1.07E-08 |
| Rptor | 4.65E-13 | 0.801388 | 0.195 | 0.439 | 1.1E-08 |
| Mbnl2 | 4.84E-13 | 0.803153 | 0.219 | 0.48 | 1.14E-08 |
| Ash1l | 4.93E-13 | 0.43023 | 0.206 | 0.459 | 1.16E-08 |
| Rpl34 | 5.24E-13 | 1.329807 | 0.794 | 0.732 | 1.23E-08 |
| Ldhb | 5.36E-13 | 2.18332 | 0.684 | 0.581 | 1.26E-08 |
| Usp24 | 5.42E-13 | 0.318119 | 0.121 | 0.321 | 1.28E-08 |
| Lfng | 5.85E-13 | 1.81382 | 0.137 | 0.362 | 1.38E-08 |
| Kidins220 | 5.94E-13 | 0.471462 | 0.093 | 0.272 | 1.4E-08 |
| Rab35 | 6.11E-13 | 1.23652 | 0.117 | 0.317 | 1.44E-08 |
| Scaf8 | 6.32E-13 | 0.343748 | 0.107 | 0.297 | 1.49E-08 |
| Dtx4 | 6.61E-13 | 0.560278 | 0.203 | 0.439 | 1.56E-08 |
| Foxj2 | 6.76E-13 | 0.361976 | 0.084 | 0.252 | 1.6E-08 |
| Rps5 | 7E-13 | 1.852775 | 0.728 | 0.74 | 1.65E-08 |
| Eif1 | 7.33E-13 | 1.550277 | 0.837 | 0.85 | 1.73E-08 |
| Rbbp6 | 7.86E-13 | 0.685781 | 0.209 | 0.463 | 1.85E-08 |
| Dennd4a | 8.33E-13 | 0.543068 | 0.418 | 0.752 | 1.97E-08 |
| Gnai2 | 8.39E-13 | 1.331738 | 0.873 | 0.894 | 1.98E-08 |
| Gm8960 | 8.65E-13 | 1.245616 | 0.334 | 0.122 | 2.04E-08 |

| | | | | | |
|---|---|---|---|---|---|
| Ngp | 9.34E-13 | 3.544685 | 0.054 | 0.203 | 2.2E-08 |
| Caprin1 | 9.43E-13 | 0.399119 | 0.18 | 0.411 | 2.22E-08 |
| Foxk2 | 1E-12 | 0.392808 | 0.076 | 0.236 | 2.36E-08 |
| Atf7 | 1.04E-12 | 0.292012 | 0.112 | 0.301 | 2.46E-08 |
| Dock2 | 1.05E-12 | 0.490534 | 0.269 | 0.545 | 2.48E-08 |
| Tmem59 | 1.2E-12 | 1.438734 | 0.79 | 0.833 | 2.83E-08 |
| Tle3 | 1.2E-12 | 0.356711 | 0.105 | 0.289 | 2.83E-08 |
| Kmt5b | 1.26E-12 | 0.438098 | 0.128 | 0.325 | 2.96E-08 |
| Etv5 | 1.3E-12 | -0.27921 | 0.137 | 0.333 | 3.06E-08 |
| Gm16192 | 1.31E-12 | 0.867348 | 0.08 | 0.248 | 3.08E-08 |
| Ube2h | 1.34E-12 | 0.678847 | 0.226 | 0.488 | 3.17E-08 |
| Dazap1 | 1.37E-12 | 0.941479 | 0.139 | 0.346 | 3.24E-08 |
| Gpr146 | 1.37E-12 | 0.349727 | 0.266 | 0.533 | 3.24E-08 |
| Inpp4b | 1.39E-12 | 0.487032 | 0.09 | 0.264 | 3.27E-08 |
| Gm38042 | 1.42E-12 | 0.666195 | 0.091 | 0.264 | 3.35E-08 |
| Herc4 | 1.48E-12 | 0.419068 | 0.136 | 0.341 | 3.49E-08 |
| H2-Q6 | 1.54E-12 | 3.686993 | 0.3 | 0.093 | 3.64E-08 |
| Slc2a5 | 1.62E-12 | 1.929758 | 0.702 | 0.569 | 3.82E-08 |
| Fubp1 | 1.78E-12 | 0.618529 | 0.193 | 0.435 | 4.2E-08 |
| N4bp2l2 | 1.89E-12 | 0.394492 | 0.159 | 0.378 | 4.46E-08 |
| Itpripl2 | 1.94E-12 | 0.478277 | 0.091 | 0.26 | 4.57E-08 |
| Gm4202 | 2E-12 | 0.51182 | 0.311 | 0.622 | 4.71E-08 |
| Irf2 | 2.13E-12 | 0.599146 | 0.201 | 0.447 | 5.03E-08 |
| Ktn1 | 2.16E-12 | 0.665137 | 0.193 | 0.419 | 5.09E-08 |
| Pmp22 | 2.21E-12 | 1.759766 | 0.766 | 0.756 | 5.2E-08 |
| Ogt | 2.36E-12 | 0.886102 | 0.377 | 0.654 | 5.56E-08 |
| Klf3 | 2.52E-12 | 0.277731 | 0.35 | 0.634 | 5.95E-08 |
| Rsrp1 | 2.67E-12 | 1.564072 | 0.839 | 0.874 | 6.3E-08 |
| Prrc2c | 2.8E-12 | 0.66978 | 0.391 | 0.683 | 6.62E-08 |
| Fam117b | 2.92E-12 | 0.324521 | 0.175 | 0.407 | 6.88E-08 |
| Myo1c | 3.07E-12 | 1.267614 | 0.095 | 0.272 | 7.23E-08 |
| Cd14 | 3.14E-12 | 1.322769 | 0.803 | 0.821 | 7.41E-08 |
| Zc3h4 | 3.31E-12 | 0.348201 | 0.09 | 0.26 | 7.81E-08 |
| Brox | 3.34E-12 | -0.40788 | 0.101 | 0.272 | 7.88E-08 |
| Timp2 | 3.36E-12 | 1.523931 | 0.78 | 0.789 | 7.93E-08 |
| Rps6kb1 | 3.46E-12 | 0.609749 | 0.166 | 0.39 | 8.16E-08 |
| Gtf3c2 | 3.49E-12 | 0.415441 | 0.139 | 0.341 | 8.23E-08 |
| Irf8 | 3.62E-12 | 1.833842 | 0.776 | 0.801 | 8.54E-08 |
| Rbpj | 3.66E-12 | 0.573133 | 0.142 | 0.35 | 8.62E-08 |
| Oaz1 | 3.81E-12 | 2.148566 | 0.696 | 0.703 | 8.99E-08 |
| Kcnq1ot1 | 4.02E-12 | 0.78439 | 0.21 | 0.459 | 9.49E-08 |
| Ascc3 | 4.31E-12 | 1.07264 | 0.088 | 0.256 | 1.02E-07 |
| BC037034 | 4.64E-12 | 0.740428 | 0.161 | 0.382 | 1.09E-07 |
| Rpl11 | 4.7E-12 | 1.718395 | 0.715 | 0.711 | 1.11E-07 |
| Ncoa3 | 4.83E-12 | 1.667356 | 0.776 | 0.74 | 1.14E-07 |
| 1810026B0 | 4.89E-12 | 1.519566 | 0.136 | 0.341 | 1.15E-07 |
| Gm44694 | 5.16E-12 | -0.26793 | 0.07 | 0.22 | 1.22E-07 |

| | | | | | |
|---|---|---|---|---|---|
| Ap3b1 | 5.25E-12 | 0.525233 | 0.127 | 0.321 | 1.24E-07 |
| Fnip1 | 5.4E-12 | 0.597143 | 0.159 | 0.374 | 1.27E-07 |
| Pten | 5.55E-12 | 0.824231 | 0.244 | 0.508 | 1.31E-07 |
| Mapk14 | 5.64E-12 | 0.33769 | 0.193 | 0.415 | 1.33E-07 |
| Rab43 | 5.67E-12 | 0.719313 | 0.12 | 0.313 | 1.34E-07 |
| Mtdh | 5.67E-12 | 1.481203 | 0.853 | 0.878 | 1.34E-07 |
| Ctnna1 | 5.8E-12 | 0.792313 | 0.134 | 0.333 | 1.37E-07 |
| Pcm1 | 5.93E-12 | 0.529961 | 0.1 | 0.268 | 1.4E-07 |
| Cic | 6.36E-12 | 0.735795 | 0.091 | 0.26 | 1.5E-07 |
| Rdx | 6.48E-12 | 0.648936 | 0.186 | 0.419 | 1.53E-07 |
| Rpl14 | 7.5E-12 | 1.814864 | 0.774 | 0.833 | 1.77E-07 |
| Rapgef1 | 7.7E-12 | 0.517573 | 0.155 | 0.366 | 1.82E-07 |
| Rpl27a | 7.79E-12 | 1.696595 | 0.747 | 0.756 | 1.84E-07 |
| Clk4 | 8.97E-12 | 0.699183 | 0.112 | 0.293 | 2.12E-07 |
| Klf6 | 1.03E-11 | 0.524913 | 0.29 | 0.573 | 2.43E-07 |
| Prkd3 | 1.06E-11 | 0.850296 | 0.088 | 0.252 | 2.5E-07 |
| Zeb2 | 1.06E-11 | 0.361147 | 0.414 | 0.732 | 2.5E-07 |
| Hk3 | 1.1E-11 | -0.63183 | 0.105 | 0.272 | 2.61E-07 |
| Taok2 | 1.11E-11 | 0.407927 | 0.153 | 0.358 | 2.62E-07 |
| Vav2 | 1.14E-11 | 1.288208 | 0.099 | 0.272 | 2.7E-07 |
| Gls | 1.27E-11 | 1.156019 | 0.136 | 0.333 | 2.99E-07 |
| Tmem104 | 1.28E-11 | 0.837488 | 0.194 | 0.431 | 3.02E-07 |
| Smchd1 | 1.56E-11 | 0.543794 | 0.123 | 0.313 | 3.68E-07 |
| Cd300c2 | 1.57E-11 | 1.609453 | 0.762 | 0.801 | 3.7E-07 |
| Dst | 1.59E-11 | 0.461697 | 0.387 | 0.675 | 3.74E-07 |
| Hs6st1 | 1.77E-11 | 1.078743 | 0.119 | 0.309 | 4.18E-07 |
| Spast | 1.83E-11 | -0.35353 | 0.065 | 0.207 | 4.31E-07 |
| Arrb1 | 1.87E-11 | 0.379837 | 0.069 | 0.215 | 4.4E-07 |
| Mtus1 | 1.92E-11 | 0.44831 | 0.355 | 0.626 | 4.53E-07 |
| Senp6 | 1.97E-11 | 1.422645 | 0.156 | 0.378 | 4.66E-07 |
| Cdk5r1 | 2.01E-11 | 0.980185 | 0.063 | 0.203 | 4.75E-07 |
| Dpp8 | 2.02E-11 | 0.341916 | 0.13 | 0.321 | 4.76E-07 |
| Git2 | 2.02E-11 | 0.597867 | 0.488 | 0.801 | 4.76E-07 |
| Slc12a6 | 2.18E-11 | 0.948546 | 0.212 | 0.467 | 5.14E-07 |
| Rpl24 | 2.22E-11 | 1.994085 | 0.601 | 0.5 | 5.23E-07 |
| Gigyf2 | 2.36E-11 | 1.062051 | 0.108 | 0.289 | 5.57E-07 |
| Galnt11 | 2.47E-11 | -0.3403 | 0.093 | 0.252 | 5.82E-07 |
| Sec16a | 2.5E-11 | 0.354995 | 0.075 | 0.224 | 5.89E-07 |
| Rpl17-ps3 | 2.51E-11 | 1.637407 | 0.416 | 0.248 | 5.92E-07 |
| Cnot1 | 2.58E-11 | 0.642559 | 0.102 | 0.272 | 6.09E-07 |
| Lag3 | 2.6E-11 | 1.755123 | 0.686 | 0.545 | 6.12E-07 |
| Fcgr2b | 2.6E-11 | 1.65284 | 0.735 | 0.728 | 6.13E-07 |
| Anks1 | 2.92E-11 | 0.403822 | 0.08 | 0.232 | 6.89E-07 |
| Mfsd14b | 3.08E-11 | -0.94983 | 0.097 | 0.248 | 7.26E-07 |
| Gm2574 | 3.12E-11 | 1.24959 | 0.379 | 0.191 | 7.36E-07 |
| Notch2 | 3.12E-11 | 1.485874 | 0.723 | 0.61 | 7.37E-07 |
| Gpr34 | 3.26E-11 | 1.033423 | 0.945 | 0.959 | 7.68E-07 |

| | | | | | |
|---|---|---|---|---|---|
| Rpl18-ps1 | 3.27E-11 | 0.743178 | 0.312 | 0.118 | 7.72E-07 |
| Ids | 3.71E-11 | 0.665611 | 0.079 | 0.232 | 8.75E-07 |
| Arhgap5 | 3.76E-11 | 0.473804 | 0.531 | 0.858 | 8.88E-07 |
| Dock8 | 3.79E-11 | 0.411401 | 0.359 | 0.638 | 8.94E-07 |
| Fbxw11 | 3.91E-11 | -0.28957 | 0.143 | 0.329 | 9.22E-07 |
| Ncbp3 | 4E-11 | 0.695888 | 0.1 | 0.268 | 9.43E-07 |
| Rpl27-ps3 | 4E-11 | 1.482569 | 0.351 | 0.159 | 9.43E-07 |
| Ipo7 | 4.18E-11 | -1.08315 | 0.086 | 0.236 | 9.85E-07 |
| Wdfy2 | 4.46E-11 | 0.823514 | 0.12 | 0.301 | 1.05E-06 |
| Rbm26 | 4.61E-11 | 2.17215 | 0.724 | 0.744 | 1.09E-06 |
| Capns1 | 4.74E-11 | 1.254336 | 0.155 | 0.366 | 1.12E-06 |
| Cntrl | 4.77E-11 | 1.062634 | 0.124 | 0.313 | 1.13E-06 |
| Ddx6 | 4.93E-11 | 0.426965 | 0.193 | 0.411 | 1.16E-06 |
| Stag2 | 4.96E-11 | 0.401143 | 0.156 | 0.346 | 1.17E-06 |
| Rnf44 | 5.03E-11 | 1.20936 | 0.175 | 0.402 | 1.19E-06 |
| Ripk1 | 5.14E-11 | 0.393792 | 0.11 | 0.285 | 1.21E-06 |
| Zfp280d | 5.28E-11 | -0.36937 | 0.088 | 0.24 | 1.25E-06 |
| Ddx21 | 5.3E-11 | 0.743388 | 0.086 | 0.244 | 1.25E-06 |
| Arhgef12 | 5.44E-11 | 0.256127 | 0.088 | 0.244 | 1.28E-06 |
| Rnase4 | 5.46E-11 | 1.373006 | 0.837 | 0.898 | 1.29E-06 |
| Casc3 | 5.49E-11 | 0.825257 | 0.078 | 0.228 | 1.29E-06 |
| Rgmb | 5.55E-11 | 0.615999 | 0.229 | 0.472 | 1.31E-06 |
| Mon2 | 5.62E-11 | 1.54287 | 0.159 | 0.378 | 1.33E-06 |
| Uso1 | 5.85E-11 | 0.628717 | 0.095 | 0.256 | 1.38E-06 |
| Zfp655 | 5.97E-11 | -0.46013 | 0.081 | 0.228 | 1.41E-06 |
| A630001G2 | 5.98E-11 | 0.491222 | 0.253 | 0.504 | 1.41E-06 |
| Uty | 6.1E-11 | -0.47902 | 0.072 | 0.211 | 1.44E-06 |
| Vps39 | 6.11E-11 | 0.486677 | 0.077 | 0.228 | 1.44E-06 |
| Atf4 | 6.24E-11 | 0.770692 | 0.148 | 0.35 | 1.47E-06 |
| Zzz3 | 6.28E-11 | 0.253908 | 0.08 | 0.228 | 1.48E-06 |
| Mtr | 6.75E-11 | 1.035064 | 0.102 | 0.272 | 1.59E-06 |
| Dip2b | 7.04E-11 | 1.003449 | 0.379 | 0.667 | 1.66E-06 |
| Zfp148 | 7.05E-11 | 0.55727 | 0.133 | 0.321 | 1.66E-06 |
| Rps10 | 7.34E-11 | 1.446878 | 0.757 | 0.793 | 1.73E-06 |
| Epb41l2 | 7.58E-11 | 0.313398 | 0.702 | 0.939 | 1.79E-06 |
| Rapgef6 | 7.59E-11 | 0.914938 | 0.174 | 0.394 | 1.79E-06 |
| Rab3gap1 | 8.33E-11 | 0.353998 | 0.134 | 0.317 | 1.97E-06 |
| Gm2225 | 8.36E-11 | 0.809996 | 0.324 | 0.13 | 1.97E-06 |
| Ago1 | 8.39E-11 | 0.756793 | 0.075 | 0.22 | 1.98E-06 |
| Gdi2 | 8.44E-11 | 1.607227 | 0.835 | 0.825 | 1.99E-06 |
| Wdfy4 | 8.74E-11 | 0.829199 | 0.068 | 0.207 | 2.06E-06 |
| Pik3ap1 | 8.77E-11 | 0.287757 | 0.169 | 0.362 | 2.07E-06 |
| Jak1 | 8.77E-11 | 0.372588 | 0.333 | 0.598 | 2.07E-06 |
| Trmt1 | 8.96E-11 | 0.347075 | 0.078 | 0.228 | 2.11E-06 |
| Kif21b | 9.64E-11 | 1.071586 | 0.178 | 0.39 | 2.27E-06 |
| Zmym4 | 9.78E-11 | 0.394311 | 0.091 | 0.248 | 2.31E-06 |
| Cysltr1 | 9.81E-11 | 0.699118 | 0.214 | 0.455 | 2.31E-06 |

| | | | | | |
|---|---|---|---|---|---|
| Hnrnph1 | 1.01E-10 | 0.307429 | 0.207 | 0.423 | 2.39E-06 |
| Gm10073 | 1.01E-10 | 1.322049 | 0.604 | 0.52 | 2.39E-06 |
| Gm6180 | 1.02E-10 | 0.856026 | 0.319 | 0.126 | 2.41E-06 |
| Iars | 1.02E-10 | -0.62984 | 0.069 | 0.203 | 2.41E-06 |
| Tmem209 | 1.09E-10 | 0.436597 | 0.097 | 0.26 | 2.56E-06 |
| mt-Nd4 | 1.11E-10 | 2.116549 | 0.776 | 0.858 | 2.63E-06 |
| Wdr44 | 1.12E-10 | 0.437452 | 0.13 | 0.309 | 2.63E-06 |
| Zfp217 | 1.15E-10 | -0.47072 | 0.135 | 0.305 | 2.72E-06 |
| Akap13 | 1.2E-10 | 0.803734 | 0.464 | 0.768 | 2.83E-06 |
| Setdb1 | 1.28E-10 | 0.265583 | 0.086 | 0.236 | 3.01E-06 |
| Rap1gds1 | 1.28E-10 | 1.070259 | 0.316 | 0.589 | 3.01E-06 |
| Thrap3 | 1.28E-10 | 0.711769 | 0.261 | 0.516 | 3.03E-06 |
| Ubr2 | 1.3E-10 | 0.4053 | 0.149 | 0.341 | 3.06E-06 |
| Sema4d | 1.45E-10 | 0.507315 | 0.371 | 0.646 | 3.41E-06 |
| Map3k2 | 1.51E-10 | 0.799521 | 0.103 | 0.272 | 3.56E-06 |
| Gng10 | 1.57E-10 | 1.504911 | 0.689 | 0.679 | 3.7E-06 |
| Phip | 1.58E-10 | 0.41805 | 0.175 | 0.382 | 3.72E-06 |
| Map4 | 1.62E-10 | 0.812732 | 0.25 | 0.508 | 3.83E-06 |
| Specc1 | 1.64E-10 | 0.757534 | 0.206 | 0.439 | 3.87E-06 |
| Rnf111 | 1.65E-10 | 0.812161 | 0.149 | 0.341 | 3.89E-06 |
| Tln1 | 1.67E-10 | 0.574028 | 0.388 | 0.679 | 3.94E-06 |
| Myo18a | 1.67E-10 | 0.790781 | 0.123 | 0.297 | 3.94E-06 |
| Brd9 | 1.69E-10 | 0.58897 | 0.136 | 0.313 | 4E-06 |
| Pom121 | 1.7E-10 | 0.649962 | 0.099 | 0.26 | 4.01E-06 |
| 4833420G1 | 1.74E-10 | 0.30871 | 0.133 | 0.313 | 4.1E-06 |
| Cd48 | 1.82E-10 | 2.408996 | 0.516 | 0.39 | 4.29E-06 |
| Tcf12 | 1.83E-10 | 0.447775 | 0.083 | 0.232 | 4.33E-06 |
| Creb1 | 1.86E-10 | 0.407982 | 0.117 | 0.289 | 4.4E-06 |
| Zc3h13 | 1.99E-10 | 0.397473 | 0.072 | 0.211 | 4.69E-06 |
| Lasp1 | 2.03E-10 | 0.403722 | 0.181 | 0.386 | 4.79E-06 |
| Iffo1 | 2.11E-10 | 1.130971 | 0.223 | 0.459 | 4.98E-06 |
| Ctnnb1 | 2.11E-10 | -0.36145 | 0.171 | 0.362 | 4.99E-06 |
| Ddx17 | 2.14E-10 | 0.569291 | 0.289 | 0.541 | 5.05E-06 |
| Rere | 2.16E-10 | 0.634266 | 0.2 | 0.419 | 5.1E-06 |
| Dleu2 | 2.2E-10 | 0.612513 | 0.78 | 0.943 | 5.19E-06 |
| Sipa1 | 2.23E-10 | 0.524738 | 0.361 | 0.646 | 5.26E-06 |
| Arpc1b | 2.3E-10 | 1.909754 | 0.711 | 0.711 | 5.43E-06 |
| Rhoh | 2.33E-10 | 0.550346 | 0.288 | 0.549 | 5.5E-06 |
| Gm11518 | 2.39E-10 | 2.001585 | 0.419 | 0.285 | 5.64E-06 |
| Bptf | 2.45E-10 | 0.801548 | 0.219 | 0.443 | 5.77E-06 |
| Zfp36l1 | 2.58E-10 | 0.358339 | 0.587 | 0.911 | 6.08E-06 |
| Plcl2 | 2.61E-10 | 0.316038 | 0.232 | 0.455 | 6.17E-06 |
| Ankrd11 | 2.63E-10 | 0.759591 | 0.365 | 0.65 | 6.19E-06 |
| Gm6170 | 2.65E-10 | 0.851713 | 0.407 | 0.22 | 6.26E-06 |
| Phrf1 | 2.71E-10 | -0.37762 | 0.07 | 0.203 | 6.4E-06 |
| Snn | 2.72E-10 | 0.767675 | 0.102 | 0.268 | 6.42E-06 |
| Dennd5a | 2.74E-10 | 0.465611 | 0.14 | 0.325 | 6.45E-06 |

| | | | | | |
|---|---|---|---|---|---|
| Camsap1 | 2.76E-10 | 0.64516 | 0.082 | 0.228 | 6.51E-06 |
| Insr | 2.81E-10 | 0.322387 | 0.111 | 0.276 | 6.62E-06 |
| Rbm33 | 2.96E-10 | 0.799274 | 0.102 | 0.264 | 6.98E-06 |
| Irf2bp2 | 3.06E-10 | 1.514053 | 0.194 | 0.415 | 7.21E-06 |
| Rpl9 | 3.14E-10 | 1.470667 | 0.735 | 0.772 | 7.41E-06 |
| Gm10180 | 3.18E-10 | 1.02287 | 0.456 | 0.301 | 7.5E-06 |
| Smo | 3.18E-10 | 0.520211 | 0.068 | 0.203 | 7.5E-06 |
| Pla2g4a | 3.54E-10 | -0.99989 | 0.099 | 0.248 | 8.36E-06 |
| Txlna | 3.56E-10 | -0.33171 | 0.079 | 0.22 | 8.39E-06 |
| Acin1 | 3.77E-10 | 0.640597 | 0.213 | 0.439 | 8.89E-06 |
| Rbm28 | 3.77E-10 | 0.283471 | 0.104 | 0.26 | 8.89E-06 |
| Fmnl3 | 3.87E-10 | 0.677178 | 0.289 | 0.528 | 9.13E-06 |
| Trim28 | 4.12E-10 | 0.607439 | 0.163 | 0.354 | 9.71E-06 |
| Prpf38b | 4.18E-10 | 1.138036 | 0.262 | 0.516 | 9.87E-06 |
| Tmem131 | 4.26E-10 | 0.603099 | 0.077 | 0.22 | 1E-05 |
| Rps4x | 4.32E-10 | 1.7499 | 0.755 | 0.789 | 1.02E-05 |
| Mark2 | 4.35E-10 | 0.847178 | 0.194 | 0.415 | 1.03E-05 |
| Ophn1 | 4.47E-10 | 0.416986 | 0.384 | 0.659 | 1.05E-05 |
| Aak1 | 4.59E-10 | 0.306731 | 0.073 | 0.207 | 1.08E-05 |
| Gm12166 | 4.75E-10 | 1.754984 | 0.666 | 0.667 | 1.12E-05 |
| Mtmr3 | 4.98E-10 | 0.264066 | 0.104 | 0.264 | 1.18E-05 |
| Rps13 | 5.54E-10 | 1.475928 | 0.748 | 0.764 | 1.31E-05 |
| Fam149b | 5.69E-10 | 0.53687 | 0.084 | 0.228 | 1.34E-05 |
| Slc43a2 | 5.7E-10 | 0.297788 | 0.124 | 0.293 | 1.34E-05 |
| Whrn | 5.83E-10 | 1.304346 | 0.173 | 0.386 | 1.38E-05 |
| Ppp1r9b | 6.06E-10 | 1.413081 | 0.108 | 0.276 | 1.43E-05 |
| Trrap | 6.11E-10 | 0.380616 | 0.116 | 0.28 | 1.44E-05 |
| Smarca4 | 6.2E-10 | 0.728364 | 0.098 | 0.252 | 1.46E-05 |
| Lrch1 | 6.27E-10 | 0.326202 | 0.246 | 0.48 | 1.48E-05 |
| Rps23-ps1 | 6.43E-10 | 1.100621 | 0.42 | 0.26 | 1.52E-05 |
| Sipa1l1 | 6.5E-10 | 0.309748 | 0.085 | 0.228 | 1.53E-05 |
| Naip5 | 6.5E-10 | 0.42353 | 0.104 | 0.264 | 1.53E-05 |
| Arid5a | 7.18E-10 | 1.02537 | 0.078 | 0.22 | 1.69E-05 |
| Snrnp70 | 7.25E-10 | 1.435689 | 0.271 | 0.528 | 1.71E-05 |
| Ncoa2 | 7.33E-10 | 0.741694 | 0.077 | 0.215 | 1.73E-05 |
| Cbl | 7.46E-10 | 0.387957 | 0.163 | 0.354 | 1.76E-05 |
| Arid3a | 7.56E-10 | 1.394898 | 0.099 | 0.26 | 1.78E-05 |
| Gdi1 | 7.68E-10 | 0.722 | 0.174 | 0.378 | 1.81E-05 |
| Epn1 | 7.74E-10 | 1.373023 | 0.181 | 0.407 | 1.82E-05 |
| Olfr620 | 7.78E-10 | 0.536932 | 0.207 | 0.045 | 1.84E-05 |
| Cyld | 7.81E-10 | 1.079405 | 0.106 | 0.268 | 1.84E-05 |
| Kdm1a | 7.87E-10 | 0.845803 | 0.076 | 0.215 | 1.86E-05 |
| Strada | 8.77E-10 | 0.443633 | 0.136 | 0.317 | 2.07E-05 |
| Zfp672 | 8.92E-10 | 0.288786 | 0.083 | 0.224 | 2.1E-05 |
| Washc4 | 8.96E-10 | 0.799951 | 0.255 | 0.496 | 2.11E-05 |
| Itch | 9.04E-10 | 0.897727 | 0.161 | 0.354 | 2.13E-05 |
| Nfatc1 | 9.06E-10 | 0.965491 | 0.214 | 0.447 | 2.14E-05 |

| | | | | | |
|---|---|---|---|---|---|
| Tlr4 | 9.28E-10 | -0.67801 | 0.113 | 0.268 | 2.19E-05 |
| Gm10359 | 9.42E-10 | 0.638734 | 0.296 | 0.114 | 2.22E-05 |
| Ykt6 | 9.86E-10 | 1.058189 | 0.157 | 0.346 | 2.33E-05 |
| Mbtps1 | 1.02E-09 | -0.64668 | 0.084 | 0.22 | 2.4E-05 |
| Arglu1 | 1.03E-09 | 1.620227 | 0.834 | 0.886 | 2.42E-05 |
| Vps13c | 1.06E-09 | 0.423316 | 0.109 | 0.268 | 2.51E-05 |
| Bmpr2 | 1.07E-09 | 0.343341 | 0.155 | 0.333 | 2.52E-05 |
| Safb2 | 1.09E-09 | 1.086164 | 0.196 | 0.419 | 2.58E-05 |
| Sall1 | 1.1E-09 | 0.715295 | 0.52 | 0.817 | 2.6E-05 |
| Trim8 | 1.11E-09 | 0.279158 | 0.264 | 0.496 | 2.62E-05 |
| H2-DMa | 1.15E-09 | 1.975818 | 0.655 | 0.622 | 2.72E-05 |
| 9930017N2 | 1.17E-09 | 0.968665 | 0.115 | 0.28 | 2.75E-05 |
| 9930021J0 | 1.27E-09 | 1.234251 | 0.085 | 0.228 | 2.99E-05 |
| Usp15 | 1.37E-09 | 1.077322 | 0.173 | 0.374 | 3.23E-05 |
| Crlf2 | 1.39E-09 | 3.625684 | 0.429 | 0.285 | 3.28E-05 |
| Pgs1 | 1.4E-09 | 0.778224 | 0.133 | 0.309 | 3.3E-05 |
| Ddx3y | 1.41E-09 | 0.796201 | 0.15 | 0.337 | 3.33E-05 |
| Arid4a | 1.53E-09 | 0.955202 | 0.172 | 0.37 | 3.61E-05 |
| Hltf | 1.54E-09 | 0.282743 | 0.125 | 0.289 | 3.63E-05 |
| Spag9 | 1.56E-09 | 0.428786 | 0.204 | 0.411 | 3.69E-05 |
| Nipbl | 1.58E-09 | 1.036392 | 0.272 | 0.508 | 3.73E-05 |
| Slc36a1 | 1.6E-09 | 0.696364 | 0.127 | 0.293 | 3.77E-05 |
| Cnot2 | 1.62E-09 | 0.393411 | 0.197 | 0.402 | 3.81E-05 |
| Sec14l1 | 1.75E-09 | 0.859907 | 0.227 | 0.439 | 4.12E-05 |
| Dhx15 | 1.8E-09 | -0.63306 | 0.132 | 0.293 | 4.26E-05 |
| Rassf3 | 1.82E-09 | 0.666072 | 0.1 | 0.252 | 4.28E-05 |
| Zc3h7a | 1.83E-09 | 0.68183 | 0.198 | 0.402 | 4.32E-05 |
| Pld1 | 1.9E-09 | 0.662642 | 0.244 | 0.463 | 4.49E-05 |
| Rhobtb1 | 1.96E-09 | 0.861378 | 0.138 | 0.321 | 4.61E-05 |
| Gm11560 | 1.97E-09 | 2.411663 | 0.266 | 0.106 | 4.65E-05 |
| Csf2rb | 2E-09 | 0.546556 | 0.263 | 0.496 | 4.71E-05 |
| Ccdc93 | 2.04E-09 | 0.598798 | 0.123 | 0.289 | 4.81E-05 |
| Smurf2 | 2.15E-09 | 0.486081 | 0.151 | 0.333 | 5.07E-05 |
| Gtf2ird2 | 2.21E-09 | 1.1626 | 0.144 | 0.325 | 5.21E-05 |
| Pkn1 | 2.22E-09 | 0.826641 | 0.282 | 0.533 | 5.24E-05 |
| Dcaf8 | 2.22E-09 | 0.630386 | 0.137 | 0.313 | 5.25E-05 |
| Pias2 | 2.24E-09 | 0.652745 | 0.073 | 0.203 | 5.28E-05 |
| Dmtf1 | 2.25E-09 | 0.709576 | 0.119 | 0.285 | 5.31E-05 |
| Pik3r5 | 2.38E-09 | 0.44059 | 0.125 | 0.289 | 5.6E-05 |
| Cenpb | 2.39E-09 | 0.333458 | 0.076 | 0.207 | 5.63E-05 |
| Dctn6 | 2.55E-09 | 0.419426 | 0.089 | 0.232 | 6.02E-05 |
| Tdrd7 | 2.58E-09 | 0.299886 | 0.079 | 0.211 | 6.08E-05 |
| Trim30d | 2.7E-09 | 1.096303 | 0.184 | 0.39 | 6.36E-05 |
| Add1 | 2.71E-09 | 0.796929 | 0.265 | 0.488 | 6.39E-05 |
| Tnfrsf11a | 2.71E-09 | 0.778022 | 0.211 | 0.431 | 6.39E-05 |
| Elk4 | 2.74E-09 | -0.43645 | 0.081 | 0.211 | 6.46E-05 |
| Med12l | 2.79E-09 | 0.532253 | 0.389 | 0.687 | 6.57E-05 |

| | | | | | |
|---|---|---|---|---|---|
| Efhd2 | 2.83E-09 | 1.942119 | 0.743 | 0.793 | 6.69E-05 |
| Washc2 | 2.86E-09 | 0.890095 | 0.119 | 0.285 | 6.74E-05 |
| Lrrc58 | 2.87E-09 | 0.386721 | 0.238 | 0.451 | 6.77E-05 |
| Sh3bgrl3 | 2.9E-09 | 1.280194 | 0.776 | 0.813 | 6.84E-05 |
| Ankrd44 | 2.93E-09 | 0.881994 | 0.308 | 0.577 | 6.9E-05 |
| Rps12-ps9 | 2.98E-09 | 1.886103 | 0.419 | 0.276 | 7.04E-05 |
| Rc3h2 | 3.1E-09 | 0.336747 | 0.095 | 0.24 | 7.32E-05 |
| Ppp6r3 | 3.15E-09 | 0.6188 | 0.155 | 0.333 | 7.44E-05 |
| Gm6988 | 3.16E-09 | 0.834172 | 0.28 | 0.114 | 7.46E-05 |
| Smarca2 | 3.27E-09 | 0.885781 | 0.182 | 0.382 | 7.71E-05 |
| Odr4 | 3.28E-09 | 0.394888 | 0.099 | 0.248 | 7.73E-05 |
| Rpl28 | 3.44E-09 | 1.655302 | 0.69 | 0.691 | 8.12E-05 |
| Hdac4 | 3.49E-09 | 1.358456 | 0.083 | 0.22 | 8.24E-05 |
| Irf2bpl | 3.52E-09 | 0.836541 | 0.183 | 0.37 | 8.3E-05 |
| Papd4 | 3.54E-09 | 0.925991 | 0.119 | 0.28 | 8.34E-05 |
| Ythdc1 | 3.69E-09 | 1.405168 | 0.195 | 0.415 | 8.71E-05 |
| Tmem170b | 3.72E-09 | 0.546644 | 0.237 | 0.451 | 8.77E-05 |
| Golga4 | 3.87E-09 | 0.308966 | 0.088 | 0.228 | 9.13E-05 |
| 8030442B0 | 4.04E-09 | 0.435719 | 0.08 | 0.215 | 9.53E-05 |
| Gm21092 | 4.11E-09 | 0.507394 | 0.08 | 0.215 | 9.69E-05 |
| E230029C0 | 4.12E-09 | 0.54291 | 0.558 | 0.882 | 9.71E-05 |
| Prrc1 | 4.27E-09 | 1.258209 | 0.126 | 0.301 | 0.000101 |
| Manba | 4.29E-09 | -0.37097 | 0.113 | 0.26 | 0.000101 |
| Itgam | 4.37E-09 | 0.89599 | 0.538 | 0.817 | 0.000103 |
| Gm15503 | 4.39E-09 | 0.540674 | 0.243 | 0.467 | 0.000103 |
| Mvb12b | 4.44E-09 | 1.615779 | 0.116 | 0.276 | 0.000105 |
| Actr2 | 4.49E-09 | 1.674938 | 0.871 | 0.886 | 0.000106 |
| Cspp1 | 4.51E-09 | 0.698918 | 0.1 | 0.248 | 0.000106 |
| Rnf145 | 4.75E-09 | -0.48417 | 0.1 | 0.24 | 0.000112 |
| U2af2 | 4.81E-09 | 1.437492 | 0.189 | 0.394 | 0.000113 |
| Asph | 4.92E-09 | 0.46312 | 0.476 | 0.744 | 0.000116 |
| Xpo6 | 5.01E-09 | 0.367733 | 0.107 | 0.256 | 0.000118 |
| Senp3 | 5.06E-09 | 0.311409 | 0.097 | 0.232 | 0.000119 |
| Bnip2 | 5.1E-09 | 0.489597 | 0.183 | 0.378 | 0.00012 |
| Reep3 | 5.17E-09 | 0.352909 | 0.301 | 0.537 | 0.000122 |
| Cnot6l | 5.31E-09 | 0.549569 | 0.101 | 0.248 | 0.000125 |
| Myo9b | 5.31E-09 | 0.795487 | 0.152 | 0.333 | 0.000125 |
| Cd52 | 5.55E-09 | 1.94853 | 0.606 | 0.561 | 0.000131 |
| Mroh1 | 5.67E-09 | 0.556233 | 0.091 | 0.232 | 0.000134 |
| Dclre1c | 5.71E-09 | 0.306113 | 0.09 | 0.228 | 0.000135 |
| Ppp3r1 | 5.82E-09 | -0.60069 | 0.104 | 0.244 | 0.000137 |
| Tmcc3 | 5.86E-09 | 0.367444 | 0.389 | 0.663 | 0.000138 |
| Gm44667 | 6.01E-09 | 0.978414 | 0.088 | 0.228 | 0.000142 |
| Tln2 | 6.22E-09 | 0.422621 | 0.141 | 0.309 | 0.000147 |
| Abcd3 | 6.27E-09 | -0.39286 | 0.082 | 0.211 | 0.000148 |
| Syngr2 | 6.35E-09 | 1.613487 | 0.712 | 0.732 | 0.00015 |
| Trf | 6.44E-09 | 1.018683 | 0.914 | 0.931 | 0.000152 |

| | | | | | |
|---|---|---|---|---|---|
| Slain2 | 6.54E-09 | 0.416359 | 0.08 | 0.211 | 0.000154 |
| Rin2 | 6.63E-09 | 0.495195 | 0.32 | 0.557 | 0.000156 |
| Erbin | 6.85E-09 | 0.513127 | 0.164 | 0.35 | 0.000161 |
| Rpl15 | 6.86E-09 | 1.395062 | 0.697 | 0.589 | 0.000162 |
| Kdm2a | 6.91E-09 | 0.940096 | 0.229 | 0.455 | 0.000163 |
| Tpt1 | 7.16E-09 | 1.378707 | 0.796 | 0.943 | 0.000169 |
| Hivep3 | 7.25E-09 | 1.017336 | 0.265 | 0.5 | 0.000171 |
| Ppp4r1l-ps | 7.49E-09 | 0.40586 | 0.082 | 0.215 | 0.000177 |
| Ptms | 7.64E-09 | 3.03432 | 0.602 | 0.573 | 0.00018 |
| Mogs | 7.68E-09 | 0.279308 | 0.135 | 0.293 | 0.000181 |
| Arpc3 | 7.77E-09 | 2.028771 | 0.558 | 0.463 | 0.000183 |
| Baz1b | 7.98E-09 | 0.463032 | 0.113 | 0.268 | 0.000188 |
| Msn | 8.06E-09 | 0.430919 | 0.466 | 0.699 | 0.00019 |
| Sart1 | 8.07E-09 | 0.566554 | 0.076 | 0.203 | 0.00019 |
| Lgals8 | 8.07E-09 | 0.852258 | 0.148 | 0.325 | 0.00019 |
| Prex1 | 8.15E-09 | 0.508431 | 0.344 | 0.585 | 0.000192 |
| Dusp7 | 8.47E-09 | 1.033775 | 0.21 | 0.402 | 0.0002 |
| Pank2 | 8.48E-09 | 0.537189 | 0.121 | 0.28 | 0.0002 |
| Nfam1 | 8.49E-09 | 0.429499 | 0.429 | 0.691 | 0.0002 |
| Gm26825 | 8.6E-09 | 1.65613 | 0.587 | 0.589 | 0.000203 |
| Tbc1d23 | 8.68E-09 | 0.444785 | 0.144 | 0.313 | 0.000205 |
| Kpna4 | 8.92E-09 | 1.126832 | 0.269 | 0.528 | 0.00021 |
| Phkb | 8.97E-09 | 0.655919 | 0.076 | 0.203 | 0.000212 |
| Shoc2 | 9.14E-09 | 0.524725 | 0.076 | 0.203 | 0.000216 |
| Sipa1l2 | 9.16E-09 | 0.504758 | 0.187 | 0.382 | 0.000216 |
| Ep400 | 9.19E-09 | 0.570109 | 0.14 | 0.309 | 0.000217 |
| Zfp638 | 9.46E-09 | 0.468238 | 0.167 | 0.35 | 0.000223 |
| Ankhd1 | 9.59E-09 | 1.361804 | 0.144 | 0.321 | 0.000226 |
| Trpc4ap | 9.6E-09 | 0.339188 | 0.26 | 0.472 | 0.000227 |
| Rps2 | 1E-08 | 1.478526 | 0.712 | 0.724 | 0.000236 |
| Gabarap | 1E-08 | 1.596559 | 0.676 | 0.659 | 0.000236 |
| Kif5b | 1.01E-08 | 0.499752 | 0.213 | 0.415 | 0.000238 |
| Rps2-ps10 | 1.02E-08 | 1.254964 | 0.325 | 0.175 | 0.00024 |
| Ubn2 | 1.03E-08 | 0.67295 | 0.343 | 0.622 | 0.000243 |
| Lhfpl2 | 1.05E-08 | 0.645772 | 0.265 | 0.484 | 0.000247 |
| Cmpk1 | 1.06E-08 | 0.589653 | 0.141 | 0.313 | 0.000249 |
| Zfand5 | 1.07E-08 | 0.8264 | 0.226 | 0.439 | 0.000252 |
| Tnpo3 | 1.1E-08 | 0.668283 | 0.156 | 0.333 | 0.00026 |
| Pxn | 1.11E-08 | 0.738562 | 0.093 | 0.232 | 0.000263 |
| AU040320 | 1.12E-08 | 0.782328 | 0.097 | 0.24 | 0.000265 |
| Elf4 | 1.18E-08 | 0.761515 | 0.09 | 0.224 | 0.000278 |
| Luc7l2 | 1.26E-08 | 0.785947 | 0.406 | 0.679 | 0.000298 |
| Ppp4r3b | 1.29E-08 | 0.413335 | 0.125 | 0.285 | 0.000305 |
| Cept1 | 1.33E-08 | 0.698971 | 0.23 | 0.459 | 0.000313 |
| Tmem135 | 1.34E-08 | 0.824368 | 0.158 | 0.341 | 0.000317 |
| Atxn2 | 1.39E-08 | 0.800861 | 0.135 | 0.297 | 0.000329 |
| Hdlbp | 1.4E-08 | 0.614542 | 0.135 | 0.301 | 0.000331 |

| | | | | | |
|---|---|---|---|---|---|
| Rock1 | 1.41E-08 | 0.896398 | 0.225 | 0.447 | 0.000333 |
| Otud4 | 1.41E-08 | 0.471642 | 0.099 | 0.24 | 0.000333 |
| Rpl10a | 1.47E-08 | 1.75854 | 0.691 | 0.691 | 0.000347 |
| Dnajc5 | 1.5E-08 | 0.676553 | 0.208 | 0.398 | 0.000354 |
| Gmeb1 | 1.52E-08 | 0.49607 | 0.106 | 0.252 | 0.000358 |
| Dcaf7 | 1.61E-08 | 0.531055 | 0.174 | 0.354 | 0.000379 |
| Gtpbp1 | 1.63E-08 | 0.483859 | 0.116 | 0.264 | 0.000385 |
| Snx10 | 1.68E-08 | 0.565895 | 0.17 | 0.354 | 0.000396 |
| Syncrip | 1.68E-08 | 1.159645 | 0.11 | 0.26 | 0.000396 |
| Tagap1 | 1.69E-08 | 0.98061 | 0.133 | 0.297 | 0.000399 |
| Gm10224 | 1.72E-08 | 0.745534 | 0.216 | 0.065 | 0.000406 |
| Psme4 | 1.77E-08 | 1.059187 | 0.14 | 0.305 | 0.000418 |
| Crot | 1.81E-08 | 0.349421 | 0.093 | 0.228 | 0.000427 |
| 4932438A1 | 1.81E-08 | 0.913518 | 0.117 | 0.268 | 0.000428 |
| Susd3 | 1.83E-08 | 1.688339 | 0.719 | 0.744 | 0.000433 |
| Gm10443 | 1.86E-08 | 0.853768 | 0.406 | 0.244 | 0.000438 |
| Rpl7a-ps7 | 1.86E-08 | 1.255994 | 0.216 | 0.069 | 0.000438 |
| Crk | 1.87E-08 | -0.32664 | 0.149 | 0.305 | 0.000442 |
| BC005537 | 1.89E-08 | 0.991336 | 0.206 | 0.398 | 0.000445 |
| Nck1 | 1.94E-08 | 0.567083 | 0.091 | 0.228 | 0.000459 |
| Akr1a1 | 1.97E-08 | 1.873321 | 0.715 | 0.711 | 0.000464 |
| Tnrc6c | 1.98E-08 | 0.692732 | 0.211 | 0.431 | 0.000467 |
| Tmod3 | 2.02E-08 | 0.422486 | 0.173 | 0.358 | 0.000476 |
| Zbtb4 | 2.04E-08 | 1.097132 | 0.105 | 0.252 | 0.00048 |
| Mtpn | 2.09E-08 | 0.565557 | 0.23 | 0.431 | 0.000493 |
| Pdlim4 | 2.16E-08 | 0.548503 | 0.144 | 0.313 | 0.000509 |
| Pnpla8 | 2.2E-08 | 0.346228 | 0.135 | 0.293 | 0.000519 |
| Snta1 | 2.2E-08 | 0.646231 | 0.136 | 0.305 | 0.00052 |
| Mib1 | 2.24E-08 | 0.886909 | 0.107 | 0.252 | 0.000528 |
| Rpl37 | 2.27E-08 | 0.981517 | 0.898 | 0.866 | 0.000534 |
| Lcp2 | 2.39E-08 | 0.619148 | 0.368 | 0.602 | 0.000564 |
| Eif3a | 2.4E-08 | 0.717442 | 0.211 | 0.419 | 0.000566 |
| Arid3b | 2.49E-08 | 1.059772 | 0.146 | 0.305 | 0.000586 |
| Ifitm3 | 2.53E-08 | 2.017061 | 0.373 | 0.224 | 0.000596 |
| Kansl1 | 2.56E-08 | 0.722682 | 0.275 | 0.512 | 0.000604 |
| Sppl3 | 2.58E-08 | 0.493557 | 0.27 | 0.5 | 0.000608 |
| Rbm6 | 2.64E-08 | 0.515382 | 0.209 | 0.402 | 0.000622 |
| Ap3d1 | 2.64E-08 | 0.678541 | 0.164 | 0.337 | 0.000623 |
| Fam193b | 2.65E-08 | 0.874871 | 0.085 | 0.215 | 0.000626 |
| Kmt2e | 2.75E-08 | 1.216326 | 0.336 | 0.593 | 0.00065 |
| Cdk12 | 2.76E-08 | 1.029612 | 0.195 | 0.402 | 0.00065 |
| Abcc3 | 2.81E-08 | 0.337108 | 0.146 | 0.297 | 0.000662 |
| Capzb | 2.82E-08 | 1.678282 | 0.692 | 0.654 | 0.000665 |
| Stt3b | 2.87E-08 | -0.43722 | 0.093 | 0.22 | 0.000676 |
| Mob2 | 2.89E-08 | 1.331395 | 0.096 | 0.232 | 0.000681 |
| Il6st | 3E-08 | 0.459853 | 0.289 | 0.492 | 0.000707 |
| Gm13408 | 3.06E-08 | 0.803509 | 0.269 | 0.11 | 0.000722 |

| | | | | | |
|---|---|---|---|---|---|
| Rpl26 | 3.09E-08 | 1.470994 | 0.703 | 0.728 | 0.000728 |
| Zzef1 | 3.1E-08 | 0.801816 | 0.102 | 0.244 | 0.00073 |
| Tal1 | 3.11E-08 | 0.718504 | 0.085 | 0.215 | 0.000734 |
| Gm11361 | 3.18E-08 | 1.099878 | 0.501 | 0.398 | 0.000749 |
| Mecp2 | 3.26E-08 | 1.502208 | 0.149 | 0.325 | 0.00077 |
| Rpl36-ps2 | 3.28E-08 | 2.373697 | 0.538 | 0.419 | 0.000773 |
| Fam102b | 3.29E-08 | 0.645052 | 0.331 | 0.557 | 0.000776 |
| Crebrf | 3.33E-08 | 0.9229 | 0.124 | 0.28 | 0.000786 |
| Acly | 3.34E-08 | -0.45772 | 0.149 | 0.297 | 0.000787 |
| Rps6-ps4 | 3.45E-08 | 1.282368 | 0.492 | 0.362 | 0.000815 |
| Atp1a1 | 3.49E-08 | 0.452864 | 0.324 | 0.533 | 0.000823 |
| Cfap74 | 3.52E-08 | 0.744447 | 0.135 | 0.293 | 0.00083 |
| Stk10 | 3.58E-08 | 1.241598 | 0.183 | 0.378 | 0.000845 |
| Bst2 | 3.72E-08 | 2.152842 | 0.526 | 0.451 | 0.000876 |
| Ogdh | 3.78E-08 | 0.521312 | 0.19 | 0.37 | 0.000891 |
| Zfp467 | 3.86E-08 | 0.735736 | 0.158 | 0.325 | 0.000911 |
| Tmem273 | 3.88E-08 | 1.115298 | 0.154 | 0.329 | 0.000916 |
| Pdcd6ip | 3.89E-08 | 0.257121 | 0.182 | 0.354 | 0.000919 |
| 1700020I1 | 3.93E-08 | 0.806121 | 0.152 | 0.321 | 0.000927 |
| Rbm5 | 3.95E-08 | 1.090149 | 0.269 | 0.488 | 0.000931 |
| Npc1 | 4.14E-08 | 0.862107 | 0.177 | 0.358 | 0.000978 |
| Sh3bp2 | 4.17E-08 | 0.888266 | 0.124 | 0.276 | 0.000984 |
| Elk3 | 4.23E-08 | 0.444597 | 0.211 | 0.402 | 0.000999 |
| Cap1 | 4.39E-08 | 0.423071 | 0.286 | 0.508 | 0.001035 |
| Bcl6 | 4.39E-08 | 1.405856 | 0.096 | 0.232 | 0.001036 |
| Rsf1 | 4.5E-08 | 1.295437 | 0.112 | 0.26 | 0.001062 |
| Adcy7 | 4.6E-08 | 0.444522 | 0.199 | 0.382 | 0.001086 |
| Ash2l | 4.98E-08 | 0.805677 | 0.082 | 0.211 | 0.001174 |
| Chn2 | 4.98E-08 | 1.132347 | 0.191 | 0.394 | 0.001175 |
| Zfc3h1 | 5.07E-08 | 0.70119 | 0.162 | 0.333 | 0.001197 |
| Gm10020 | 5.14E-08 | 0.575128 | 0.243 | 0.093 | 0.001212 |
| P3h2 | 5.14E-08 | 0.457567 | 0.141 | 0.297 | 0.001213 |
| Rab8b | 5.15E-08 | 0.3147 | 0.194 | 0.382 | 0.001216 |
| Camk2d | 5.15E-08 | 0.914228 | 0.28 | 0.512 | 0.001216 |
| Kras | 5.21E-08 | 0.665623 | 0.159 | 0.333 | 0.00123 |
| Usp40 | 5.27E-08 | 0.492035 | 0.107 | 0.248 | 0.001244 |
| Rpl9-ps6 | 5.55E-08 | 1.647774 | 0.618 | 0.557 | 0.001309 |
| Mybbp1a | 5.67E-08 | -0.40906 | 0.106 | 0.24 | 0.001336 |
| Plxnb2 | 5.67E-08 | 0.478229 | 0.429 | 0.679 | 0.001337 |
| Rps28 | 5.78E-08 | 0.958984 | 0.919 | 0.923 | 0.001363 |
| Pip4k2a | 6.01E-08 | 0.891346 | 0.326 | 0.557 | 0.001417 |
| Dync1h1 | 6.25E-08 | 0.478272 | 0.137 | 0.293 | 0.001474 |
| Gmfb | 6.29E-08 | 0.279609 | 0.106 | 0.244 | 0.001483 |
| A130071D( | 6.34E-08 | 2.079771 | 0.105 | 0.244 | 0.001495 |
| Gm21811 | 6.67E-08 | 0.853721 | 0.091 | 0.224 | 0.001574 |
| Ifi27l2a | 6.67E-08 | 1.837652 | 0.398 | 0.244 | 0.001574 |
| Abi3 | 6.74E-08 | 1.944921 | 0.731 | 0.841 | 0.00159 |

| | | | | | |
|---|---|---|---|---|---|
| Ecpas | 6.75E-08 | 0.547574 | 0.112 | 0.256 | 0.001593 |
| Kctd20 | 6.76E-08 | 0.323884 | 0.091 | 0.215 | 0.001594 |
| Vps4a | 6.79E-08 | 1.00436 | 0.101 | 0.24 | 0.001603 |
| Fcho2 | 6.84E-08 | 0.871814 | 0.251 | 0.472 | 0.001615 |
| Dennd1c | 7.09E-08 | 1.231902 | 0.132 | 0.289 | 0.001672 |
| Abcb1b | 7.17E-08 | 1.198866 | 0.098 | 0.232 | 0.001692 |
| Fam13b | 7.42E-08 | 0.937585 | 0.081 | 0.207 | 0.00175 |
| Stag1 | 7.43E-08 | 0.264094 | 0.152 | 0.313 | 0.001753 |
| Plagl2 | 7.45E-08 | 0.789871 | 0.122 | 0.268 | 0.001757 |
| Dock4 | 7.52E-08 | 0.97945 | 0.464 | 0.764 | 0.001774 |
| Map3k11 | 7.72E-08 | 0.491665 | 0.136 | 0.289 | 0.001821 |
| Ube2r2 | 7.95E-08 | 1.0295 | 0.141 | 0.301 | 0.001874 |
| Pfkfb3 | 8.22E-08 | 0.511248 | 0.199 | 0.378 | 0.00194 |
| Marf1 | 8.37E-08 | 2.319487 | 0.632 | 0.63 | 0.001975 |
| Slc25a37 | 8.41E-08 | 0.628511 | 0.088 | 0.215 | 0.001984 |
| Gm3379 | 8.47E-08 | 1.953851 | 0.474 | 0.415 | 0.001999 |
| Mgat4b | 8.48E-08 | 0.55493 | 0.124 | 0.272 | 0.002 |
| Csnk1d | 8.49E-08 | 0.764735 | 0.167 | 0.341 | 0.002002 |
| Pi4k2a | 8.6E-08 | 0.66652 | 0.124 | 0.276 | 0.00203 |
| Rhoq | 8.67E-08 | 1.309979 | 0.1 | 0.236 | 0.002045 |
| Cd37 | 8.71E-08 | 1.390463 | 0.775 | 0.874 | 0.002054 |
| Ccdc50 | 8.8E-08 | 1.015863 | 0.269 | 0.492 | 0.002076 |
| Dock11 | 9.09E-08 | 0.460377 | 0.144 | 0.297 | 0.002145 |
| Bicd2 | 9.17E-08 | 0.415676 | 0.109 | 0.248 | 0.002163 |
| Mapk1 | 9.5E-08 | 0.524679 | 0.15 | 0.309 | 0.002242 |
| Gm26542 | 9.77E-08 | 1.049094 | 0.09 | 0.224 | 0.002305 |
| Txndc11 | 9.78E-08 | 0.73429 | 0.147 | 0.305 | 0.002307 |
| Mbnl1 | 1.01E-07 | 0.695469 | 0.684 | 0.931 | 0.002387 |
| Snx30 | 1.02E-07 | 0.828307 | 0.083 | 0.207 | 0.002405 |
| Cds1 | 1.04E-07 | 0.625602 | 0.1 | 0.236 | 0.002451 |
| Gm4294 | 1.1E-07 | 0.766988 | 0.393 | 0.248 | 0.002593 |
| Atf1 | 1.11E-07 | 0.714003 | 0.102 | 0.236 | 0.002609 |
| Map3k5 | 1.13E-07 | 0.409199 | 0.117 | 0.26 | 0.002676 |
| Ttc28 | 1.14E-07 | 1.573782 | 0.131 | 0.285 | 0.002694 |
| Kpnb1 | 1.15E-07 | 0.695273 | 0.138 | 0.297 | 0.002702 |
| Psmb8 | 1.17E-07 | 2.195515 | 0.563 | 0.5 | 0.002755 |
| Vps13b | 1.18E-07 | 0.476328 | 0.083 | 0.203 | 0.002793 |
| Gm8129 | 1.19E-07 | 1.10989 | 0.265 | 0.122 | 0.002817 |
| AI837181 | 1.2E-07 | 0.786328 | 0.113 | 0.26 | 0.00284 |
| Pbrm1 | 1.21E-07 | 0.870194 | 0.196 | 0.378 | 0.002861 |
| Fgfr1op2 | 1.24E-07 | 0.763597 | 0.188 | 0.37 | 0.002936 |
| Ski | 1.25E-07 | 1.098127 | 0.243 | 0.447 | 0.002958 |
| Arap3 | 1.25E-07 | 1.19343 | 0.104 | 0.24 | 0.002959 |
| Arcn1 | 1.28E-07 | 0.568711 | 0.164 | 0.329 | 0.003024 |
| Plekha1 | 1.32E-07 | 1.592616 | 0.135 | 0.297 | 0.003103 |
| Agmo | 1.33E-07 | 0.37287 | 0.112 | 0.252 | 0.00314 |
| Rassf4 | 1.35E-07 | 0.633032 | 0.271 | 0.488 | 0.003187 |

| | | | | | |
|---|---|---|---|---|---|
| Arhgap30 | 1.36E-07 | 1.037439 | 0.218 | 0.407 | 0.003197 |
| Mycbp2 | 1.36E-07 | 0.560089 | 0.447 | 0.744 | 0.00321 |
| Qrich1 | 1.39E-07 | -0.29804 | 0.15 | 0.301 | 0.003291 |
| Ndrg3 | 1.4E-07 | 0.947101 | 0.109 | 0.248 | 0.003292 |
| Golph3 | 1.4E-07 | -0.51005 | 0.085 | 0.203 | 0.003295 |
| Edem1 | 1.45E-07 | 0.580657 | 0.347 | 0.589 | 0.003409 |
| D5Ertd579 | 1.47E-07 | 0.485233 | 0.084 | 0.203 | 0.003474 |
| Sall3 | 1.48E-07 | 0.791502 | 0.369 | 0.606 | 0.003495 |
| Csk | 1.5E-07 | 0.98384 | 0.216 | 0.411 | 0.003543 |
| Cnp | 1.52E-07 | 1.391478 | 0.192 | 0.39 | 0.003584 |
| Slc15a3 | 1.52E-07 | 0.970676 | 0.218 | 0.407 | 0.003589 |
| Clk1 | 1.53E-07 | 1.154481 | 0.311 | 0.541 | 0.003604 |
| Trappc8 | 1.55E-07 | 0.693474 | 0.117 | 0.26 | 0.003665 |
| Uvrag | 1.56E-07 | 0.498592 | 0.112 | 0.248 | 0.003691 |
| Npl | 1.59E-07 | 0.437872 | 0.101 | 0.236 | 0.003744 |
| Rgl1 | 1.6E-07 | 0.443152 | 0.09 | 0.215 | 0.003773 |
| Ccdc82 | 1.6E-07 | 0.759948 | 0.146 | 0.301 | 0.003776 |
| Ccdc88a | 1.61E-07 | 0.708705 | 0.122 | 0.264 | 0.003795 |
| Osbpl9 | 1.63E-07 | 0.628713 | 0.183 | 0.358 | 0.003836 |
| Fnbp4 | 1.64E-07 | 0.927071 | 0.139 | 0.293 | 0.003877 |
| Acadm | 1.65E-07 | 0.616052 | 0.111 | 0.248 | 0.003882 |
| Zcchc7 | 1.65E-07 | 0.814991 | 0.171 | 0.341 | 0.003888 |
| Ptprc | 1.66E-07 | 0.647875 | 0.594 | 0.874 | 0.003908 |
| Acss1 | 1.68E-07 | -0.40497 | 0.1 | 0.22 | 0.003969 |
| Sept2 | 1.7E-07 | 1.704993 | 0.765 | 0.707 | 0.004012 |
| Arhgdia | 1.72E-07 | 2.035223 | 0.696 | 0.728 | 0.004059 |
| Smarcc2 | 1.78E-07 | 0.629066 | 0.128 | 0.276 | 0.004191 |
| Acox3 | 1.79E-07 | 0.556698 | 0.192 | 0.362 | 0.004214 |
| Cep350 | 1.82E-07 | 0.357704 | 0.133 | 0.276 | 0.004286 |
| Stx2 | 1.84E-07 | 0.859286 | 0.133 | 0.285 | 0.004335 |
| Wdr37 | 1.84E-07 | 1.594919 | 0.089 | 0.215 | 0.004349 |
| Atp7a | 1.87E-07 | 0.537925 | 0.127 | 0.272 | 0.0044 |
| Skp1a | 1.88E-07 | 0.769902 | 0.198 | 0.382 | 0.004443 |
| Htt | 1.9E-07 | 0.303462 | 0.112 | 0.248 | 0.004489 |
| Rps10-ps1 | 1.95E-07 | 2.84974 | 0.441 | 0.354 | 0.004595 |
| Vrk1 | 1.97E-07 | 0.42501 | 0.121 | 0.264 | 0.004657 |
| Dgkd | 1.99E-07 | 0.491309 | 0.135 | 0.28 | 0.004684 |
| Abl1 | 2.07E-07 | 0.861943 | 0.125 | 0.268 | 0.004893 |
| Dnajc13 | 2.12E-07 | 0.495636 | 0.147 | 0.301 | 0.004994 |
| Zfp740 | 2.17E-07 | 1.250708 | 0.183 | 0.362 | 0.005121 |
| Lemd2 | 2.2E-07 | 0.530211 | 0.125 | 0.268 | 0.005188 |
| Nuak1 | 2.21E-07 | 1.145837 | 0.242 | 0.459 | 0.005205 |
| Larp4 | 2.25E-07 | 0.548214 | 0.112 | 0.248 | 0.005312 |
| Hipk1 | 2.25E-07 | 0.532088 | 0.128 | 0.272 | 0.005314 |
| Gspt1 | 2.37E-07 | 0.728663 | 0.205 | 0.394 | 0.005593 |
| Rps27a | 2.38E-07 | 1.29063 | 0.759 | 0.797 | 0.005611 |
| Cbfa2t3 | 2.4E-07 | 0.909568 | 0.178 | 0.346 | 0.005667 |

| | | | | | |
|---|---|---|---|---|---|
| Emsy | 2.43E-07 | 1.399525 | 0.088 | 0.211 | 0.005728 |
| Gm5138 | 2.53E-07 | 1.925415 | 0.272 | 0.138 | 0.005976 |
| Colgalt1 | 2.67E-07 | 0.561364 | 0.188 | 0.366 | 0.006294 |
| Pds5a | 2.68E-07 | 0.818425 | 0.187 | 0.362 | 0.006324 |
| Shisa5 | 2.7E-07 | 1.914188 | 0.645 | 0.659 | 0.006358 |
| 2610005L0 | 2.74E-07 | 0.823056 | 0.182 | 0.358 | 0.006457 |
| Casp4 | 2.8E-07 | 0.950685 | 0.085 | 0.203 | 0.006612 |
| Rps15a | 2.86E-07 | 1.666145 | 0.613 | 0.577 | 0.006735 |
| Zkscan3 | 2.9E-07 | 0.690785 | 0.278 | 0.496 | 0.00685 |
| Sp3 | 2.91E-07 | 0.486513 | 0.197 | 0.358 | 0.006866 |
| Erp29 | 2.91E-07 | 1.716909 | 0.698 | 0.74 | 0.006873 |
| Arhgef40 | 2.94E-07 | 1.206141 | 0.2 | 0.378 | 0.006927 |
| Ubn1 | 3.01E-07 | 0.853331 | 0.158 | 0.321 | 0.007104 |
| Thrsp | 3.04E-07 | 2.821564 | 0.251 | 0.11 | 0.007169 |
| Serf2 | 3.05E-07 | 1.835308 | 0.705 | 0.638 | 0.007186 |
| Gm7536 | 3.07E-07 | 1.693275 | 0.542 | 0.512 | 0.007249 |
| Sgk3 | 3.16E-07 | 0.819622 | 0.167 | 0.333 | 0.007454 |
| Slc12a2 | 3.19E-07 | 0.352157 | 0.127 | 0.264 | 0.007535 |
| Zmiz1 | 3.24E-07 | 0.755637 | 0.367 | 0.589 | 0.007633 |
| Itgb3 | 3.46E-07 | 0.583792 | 0.09 | 0.207 | 0.008156 |
| Hps4 | 3.49E-07 | 0.667841 | 0.434 | 0.687 | 0.008232 |
| Atf7ip | 3.63E-07 | 0.816697 | 0.299 | 0.508 | 0.008574 |
| Cebpz | 3.77E-07 | 1.273212 | 0.226 | 0.411 | 0.008902 |
| Kif1b | 3.8E-07 | 1.278804 | 0.144 | 0.301 | 0.008975 |
| Camk1d | 3.83E-07 | 0.719786 | 0.181 | 0.354 | 0.009023 |
| Plcg2 | 3.95E-07 | 0.91597 | 0.211 | 0.39 | 0.009309 |
| Dido1 | 3.96E-07 | 1.433078 | 0.146 | 0.301 | 0.009331 |
| Cul1 | 3.99E-07 | 0.57846 | 0.253 | 0.447 | 0.009412 |
| Herpud2 | 4.02E-07 | 0.663063 | 0.192 | 0.37 | 0.009473 |
| Fam217b | 4.02E-07 | 1.095708 | 0.085 | 0.203 | 0.009492 |
| Gpr137b | 4.03E-07 | 0.854537 | 0.15 | 0.305 | 0.009505 |
| Kpna3 | 4.07E-07 | -0.39627 | 0.109 | 0.232 | 0.009589 |
| Zup1 | 4.08E-07 | 0.858764 | 0.281 | 0.504 | 0.009635 |
| Ubtf | 4.12E-07 | 0.619494 | 0.13 | 0.272 | 0.009716 |
| P2rx7 | 4.15E-07 | 0.814059 | 0.359 | 0.593 | 0.009791 |
| Btbd1 | 4.2E-07 | 0.35418 | 0.132 | 0.276 | 0.009915 |
| Gpr107 | 4.25E-07 | 0.560061 | 0.147 | 0.297 | 0.010014 |
| Gm10132 | 4.27E-07 | 2.012176 | 0.594 | 0.589 | 0.010069 |
| Gpat4 | 4.3E-07 | 0.625661 | 0.091 | 0.211 | 0.010151 |
| C130089K0 | 4.33E-07 | 1.212941 | 0.216 | 0.39 | 0.01022 |
| Dcaf12 | 4.44E-07 | 0.625085 | 0.116 | 0.252 | 0.010462 |
| N4bp1 | 4.44E-07 | 0.531593 | 0.122 | 0.256 | 0.010468 |
| Pitpna | 4.48E-07 | 1.222955 | 0.307 | 0.549 | 0.010557 |
| Tnrc18 | 4.5E-07 | 1.414653 | 0.105 | 0.236 | 0.010606 |
| Chd6 | 4.6E-07 | 0.607581 | 0.106 | 0.236 | 0.010857 |
| Snx4 | 4.67E-07 | 0.716899 | 0.099 | 0.224 | 0.011008 |
| Anp32a | 4.69E-07 | 0.594782 | 0.198 | 0.374 | 0.011064 |

| | | | | | |
|---|---|---|---|---|---|
| Sec24b | 4.71E-07 | 0.683066 | 0.124 | 0.256 | 0.011107 |
| Gm12428 | 4.72E-07 | 0.819671 | 0.221 | 0.085 | 0.011144 |
| Tnip2 | 4.75E-07 | 0.761012 | 0.1 | 0.224 | 0.011207 |
| Casd1 | 4.77E-07 | 1.044334 | 0.108 | 0.24 | 0.011252 |
| Tmem127 | 4.83E-07 | 0.697384 | 0.227 | 0.415 | 0.011395 |
| Tgoln1 | 4.89E-07 | 0.697018 | 0.223 | 0.407 | 0.011524 |
| Cpsf6 | 4.97E-07 | 0.535715 | 0.186 | 0.346 | 0.011732 |
| Hsbp1 | 5.02E-07 | 2.507151 | 0.457 | 0.337 | 0.011846 |
| Epb41 | 5.11E-07 | 0.508849 | 0.093 | 0.211 | 0.012053 |
| Gm9888 | 5.17E-07 | 0.522808 | 0.238 | 0.427 | 0.012193 |
| Eed | 5.18E-07 | 0.989692 | 0.159 | 0.321 | 0.012213 |
| Herc2 | 5.25E-07 | 1.1027 | 0.344 | 0.577 | 0.012388 |
| Epc1 | 5.3E-07 | 0.639602 | 0.101 | 0.228 | 0.012507 |
| Retreg1 | 5.31E-07 | 0.92027 | 0.309 | 0.528 | 0.012522 |
| Gm10157 | 5.36E-07 | 0.572542 | 0.211 | 0.077 | 0.012634 |
| Vrk2 | 5.36E-07 | 1.429559 | 0.105 | 0.24 | 0.012655 |
| Dync1i2 | 5.45E-07 | 1.307412 | 0.231 | 0.423 | 0.012865 |
| Arhgap27 | 5.56E-07 | -0.29722 | 0.116 | 0.244 | 0.013115 |
| Rps10-ps2 | 5.72E-07 | 1.865515 | 0.535 | 0.459 | 0.013495 |
| Thumpd3 | 5.8E-07 | 0.849558 | 0.152 | 0.305 | 0.013673 |
| Wsb1 | 5.94E-07 | 0.845494 | 0.281 | 0.492 | 0.014017 |
| Mfap3 | 5.95E-07 | 0.60307 | 0.406 | 0.63 | 0.014035 |
| Gltp | 5.99E-07 | 2.478 | 0.419 | 0.297 | 0.014133 |
| Morc3 | 6.06E-07 | 0.800101 | 0.232 | 0.427 | 0.014291 |
| Tomm70a | 6.07E-07 | 0.410605 | 0.118 | 0.248 | 0.014318 |
| Gm49369 | 6.13E-07 | 1.583773 | 0.665 | 0.947 | 0.014449 |
| Usp12 | 6.14E-07 | 0.537011 | 0.088 | 0.203 | 0.014476 |
| Rabgap1 | 6.21E-07 | 0.693245 | 0.148 | 0.301 | 0.014644 |
| Tlr3 | 6.29E-07 | 0.880959 | 0.128 | 0.268 | 0.014837 |
| Cox4i1 | 6.59E-07 | 1.381987 | 0.716 | 0.756 | 0.015542 |
| Gm37954 | 6.6E-07 | 1.343513 | 0.213 | 0.411 | 0.015558 |
| March7 | 6.68E-07 | 0.660789 | 0.184 | 0.346 | 0.015756 |
| Gng2 | 6.69E-07 | 1.045997 | 0.218 | 0.411 | 0.01579 |
| Ppp1ca | 6.82E-07 | 1.766499 | 0.583 | 0.52 | 0.016092 |
| Tmem126a | 6.95E-07 | 2.870799 | 0.462 | 0.382 | 0.016397 |
| Snx14 | 7.11E-07 | -0.26048 | 0.097 | 0.211 | 0.016762 |
| Cds2 | 7.11E-07 | 0.940641 | 0.146 | 0.289 | 0.016765 |
| Trim2 | 7.12E-07 | 0.619848 | 0.101 | 0.224 | 0.016787 |
| Ino80d | 7.32E-07 | 0.624707 | 0.092 | 0.211 | 0.017266 |
| Ddx50 | 7.36E-07 | 0.960277 | 0.192 | 0.374 | 0.017367 |
| Gcnt1 | 7.61E-07 | 1.004973 | 0.295 | 0.512 | 0.017946 |
| Atxn2l | 7.65E-07 | 1.002 | 0.188 | 0.366 | 0.018046 |
| Ganab | 7.68E-07 | -0.28678 | 0.1 | 0.215 | 0.018105 |
| Cbfb | 7.78E-07 | -0.31672 | 0.112 | 0.236 | 0.018363 |
| Cd33 | 8.03E-07 | 0.596561 | 0.548 | 0.805 | 0.018953 |
| Herc1 | 8.14E-07 | 1.048684 | 0.197 | 0.37 | 0.019204 |
| Pdzd8 | 8.42E-07 | 0.464287 | 0.145 | 0.293 | 0.019854 |

| | | | | | |
|---|---|---|---|---|---|
| Prdx1 | 8.54E-07 | 1.898302 | 0.529 | 0.447 | 0.020146 |
| Baz1a | 8.58E-07 | 0.941956 | 0.112 | 0.244 | 0.020245 |
| Oaz1-ps | 8.61E-07 | 1.343762 | 0.541 | 0.492 | 0.020313 |
| Clint1 | 8.64E-07 | 0.644262 | 0.217 | 0.398 | 0.020377 |
| Sltm | 8.64E-07 | 1.071214 | 0.167 | 0.325 | 0.020386 |
| Cmip | 8.68E-07 | 1.092235 | 0.139 | 0.289 | 0.020473 |
| Mtss1 | 8.87E-07 | 1.336797 | 0.131 | 0.276 | 0.020919 |
| Pcyox1 | 9.07E-07 | 0.257532 | 0.135 | 0.272 | 0.021403 |
| Btg2 | 9.08E-07 | 0.362263 | 0.424 | 0.65 | 0.021419 |
| Tmx4 | 9.18E-07 | 0.450345 | 0.127 | 0.264 | 0.021663 |
| Ap2a2 | 9.44E-07 | 0.553479 | 0.253 | 0.455 | 0.022268 |
| Leng8 | 9.46E-07 | 1.091446 | 0.267 | 0.472 | 0.022314 |
| Rp2 | 9.48E-07 | 1.112616 | 0.116 | 0.248 | 0.022356 |
| Endod1 | 9.53E-07 | -0.29644 | 0.113 | 0.236 | 0.022485 |
| Vps4b | 9.73E-07 | 1.030939 | 0.177 | 0.341 | 0.022954 |
| Fkbp15 | 9.78E-07 | 0.826983 | 0.285 | 0.496 | 0.023076 |
| Tpm3 | 9.89E-07 | 2.001244 | 0.796 | 0.89 | 0.023329 |
| Tk2 | 9.97E-07 | 0.432692 | 0.113 | 0.244 | 0.023509 |
| Lrrc14 | 1E-06 | 1.095344 | 0.094 | 0.215 | 0.023632 |
| Map7d1 | 1.02E-06 | 0.255907 | 0.139 | 0.276 | 0.023982 |
| Gabpb2 | 1.02E-06 | 0.909203 | 0.106 | 0.232 | 0.024094 |
| Ctcf | 1.04E-06 | 0.638462 | 0.135 | 0.276 | 0.024455 |
| Otud5 | 1.04E-06 | 0.640716 | 0.142 | 0.285 | 0.024473 |
| Memo1 | 1.05E-06 | 0.552378 | 0.111 | 0.236 | 0.024865 |
| Fam120b | 1.06E-06 | 0.479123 | 0.174 | 0.333 | 0.024913 |
| Ss18 | 1.06E-06 | 0.549 | 0.122 | 0.256 | 0.024957 |
| Gm17511 | 1.07E-06 | 0.673219 | 0.22 | 0.093 | 0.025347 |
| Noc2l | 1.08E-06 | 0.468 | 0.137 | 0.276 | 0.025412 |
| Arl8b | 1.09E-06 | 0.302599 | 0.289 | 0.472 | 0.025785 |
| Nfkb1 | 1.13E-06 | 0.715437 | 0.308 | 0.52 | 0.026572 |
| Dpagt1 | 1.14E-06 | 1.12331 | 0.13 | 0.276 | 0.026927 |
| Rpl3-ps2 | 1.15E-06 | 0.425063 | 0.241 | 0.106 | 0.027163 |
| Glyr1 | 1.16E-06 | 0.73934 | 0.136 | 0.276 | 0.027313 |
| Xpo7 | 1.16E-06 | 0.394901 | 0.102 | 0.224 | 0.027407 |
| Ap1s2 | 1.18E-06 | 0.984825 | 0.134 | 0.272 | 0.02776 |
| Asxl2 | 1.18E-06 | 0.379028 | 0.215 | 0.386 | 0.027845 |
| Pacs2 | 1.18E-06 | 1.53543 | 0.162 | 0.321 | 0.027884 |
| Nbeal1 | 1.24E-06 | 0.442473 | 0.097 | 0.211 | 0.029282 |
| Itpkb | 1.29E-06 | 0.737181 | 0.088 | 0.203 | 0.030478 |
| Ccni | 1.29E-06 | 0.297808 | 0.418 | 0.622 | 0.030516 |
| Cr1l | 1.3E-06 | 0.872412 | 0.268 | 0.476 | 0.030749 |
| Vps54 | 1.32E-06 | 0.943699 | 0.173 | 0.333 | 0.031058 |
| Rasa4 | 1.34E-06 | 0.787381 | 0.247 | 0.431 | 0.031671 |
| Tmem86a | 1.34E-06 | 2.138761 | 0.601 | 0.561 | 0.031701 |
| Kdm2b | 1.37E-06 | 1.04143 | 0.135 | 0.276 | 0.032406 |
| Gatad2b | 1.39E-06 | 1.052552 | 0.195 | 0.374 | 0.03282 |
| Stambpl1 | 1.4E-06 | 0.726511 | 0.2 | 0.37 | 0.03304 |

| | | | | | |
|---|---|---|---|---|---|
| Usp21 | 1.41E-06 | 1.065672 | 0.203 | 0.374 | 0.033145 |
| Gtf2a2 | 1.41E-06 | 0.713481 | 0.161 | 0.317 | 0.033308 |
| Rasgrp3 | 1.42E-06 | 0.349789 | 0.335 | 0.508 | 0.033482 |
| Lsm14a | 1.43E-06 | 0.539824 | 0.103 | 0.224 | 0.033624 |
| Lpgat1 | 1.44E-06 | 0.417058 | 0.129 | 0.264 | 0.034004 |
| Gm5148 | 1.45E-06 | 0.588971 | 0.238 | 0.427 | 0.03426 |
| Pxk | 1.46E-06 | 0.614113 | 0.106 | 0.228 | 0.034399 |
| Magt1 | 1.49E-06 | 0.384128 | 0.217 | 0.386 | 0.035228 |
| Npepps | 1.54E-06 | 1.12425 | 0.175 | 0.337 | 0.036276 |
| Smc6 | 1.55E-06 | 0.420777 | 0.148 | 0.289 | 0.036531 |
| Rpl17-ps5 | 1.55E-06 | 0.979823 | 0.264 | 0.13 | 0.036566 |
| Ubfd1 | 1.56E-06 | 0.816617 | 0.131 | 0.268 | 0.036863 |
| Pcf11 | 1.57E-06 | 1.022333 | 0.176 | 0.341 | 0.037074 |
| Birc3 | 1.58E-06 | 1.013705 | 0.09 | 0.207 | 0.037253 |
| Orc3 | 1.62E-06 | 1.233909 | 0.132 | 0.276 | 0.038244 |
| Sqor | 1.63E-06 | 1.161367 | 0.14 | 0.289 | 0.038543 |
| Papola | 1.67E-06 | 0.840668 | 0.273 | 0.48 | 0.039301 |
| Ctbs | 1.67E-06 | 0.54747 | 0.111 | 0.236 | 0.039479 |
| Atad2b | 1.68E-06 | 0.645417 | 0.135 | 0.268 | 0.039543 |
| Epb41l3 | 1.69E-06 | 0.904735 | 0.125 | 0.26 | 0.039792 |
| Selenop | 1.71E-06 | 0.776568 | 0.948 | 0.984 | 0.040402 |
| Sesn1 | 1.72E-06 | 0.488938 | 0.238 | 0.419 | 0.040603 |
| Mxd4 | 1.76E-06 | 0.839775 | 0.173 | 0.325 | 0.041618 |
| Rnf2 | 1.77E-06 | -0.58371 | 0.107 | 0.22 | 0.04177 |
| Tulp4 | 1.77E-06 | 0.668792 | 0.189 | 0.337 | 0.041794 |
| Ube3a | 1.81E-06 | 1.095739 | 0.221 | 0.407 | 0.042699 |
| Faf2 | 1.81E-06 | 0.698808 | 0.168 | 0.321 | 0.042756 |
| Brd2 | 1.82E-06 | 1.35453 | 0.245 | 0.447 | 0.042841 |
| Selenot | 1.86E-06 | 0.624432 | 0.167 | 0.309 | 0.043813 |
| Ywhag | 1.86E-06 | 1.74993 | 0.147 | 0.297 | 0.043984 |
| Mgrn1 | 1.87E-06 | 0.838228 | 0.174 | 0.325 | 0.044167 |
| Gtf2a1 | 1.95E-06 | 0.420486 | 0.22 | 0.378 | 0.045971 |
| Gm7336 | 1.96E-06 | 1.308075 | 0.294 | 0.179 | 0.046167 |
| Frg1 | 2.01E-06 | 1.223491 | 0.169 | 0.325 | 0.047428 |
| Synrg | 2.02E-06 | 0.322784 | 0.098 | 0.211 | 0.047735 |
| Zfp398 | 2.03E-06 | 0.616515 | 0.106 | 0.228 | 0.047938 |
| Camk1 | 2.05E-06 | 1.792914 | 0.685 | 0.699 | 0.048437 |
| Ccnl2 | 2.06E-06 | 0.544528 | 0.161 | 0.309 | 0.048583 |
| Med13l | 2.1E-06 | 1.417919 | 0.159 | 0.305 | 0.049447 |
| Gpx1 | 2.1E-06 | 1.517859 | 0.65 | 0.618 | 0.049537 |
| Limd2 | 2.1E-06 | 2.078228 | 0.633 | 0.638 | 0.049567 |
| Sh3kbp1 | 2.11E-06 | 1.169347 | 0.218 | 0.398 | 0.049717 |

### Supplementry Table 4. Aging DEGs in Cortex

| Gene | p_val | avg_log2FC | pct.1 | pct.2 | p_val_adj |
|---|---|---|---|---|---|
| S100a8 | 1.4E-136 | -1.95309 | 0.004 | 0.52 | 3.3E-132 |
| C1qa | 4.3E-120 | 2.006581 | 1 | 1 | 1E-115 |
| S100a9 | 3.9E-114 | -1.79785 | 0.004 | 0.441 | 9.3E-110 |
| Ct010467.1 | 2.1E-111 | 1.219254 | 1 | 1 | 5.1E-107 |
| Lgmn | 3.1E-108 | 1.522331 | 1 | 1 | 7.4E-104 |
| Selplg | 1.48E-98 | 1.495924 | 0.997 | 1 | 3.49E-94 |
| Hexb | 2.11E-97 | 1.093001 | 1 | 1 | 4.98E-93 |
| C1qc | 3.81E-97 | 1.644847 | 1 | 1 | 8.98E-93 |
| Laptm5 | 1.69E-96 | 1.572493 | 1 | 1 | 4E-92 |
| Ngp | 1.26E-92 | -1.20625 | 0.006 | 0.384 | 2.98E-88 |
| Tmem119 | 7.54E-92 | 1.648647 | 0.997 | 1 | 1.78E-87 |
| Cd81 | 3.39E-91 | 1.56412 | 0.998 | 1 | 7.99E-87 |
| Itm2b | 1.17E-88 | 1.299334 | 1 | 1 | 2.77E-84 |
| Cfl1 | 3.08E-88 | 2.831777 | 0.964 | 0.991 | 7.26E-84 |
| Ins2 | 7.01E-83 | -1.99923 | 0.008 | 0.354 | 1.65E-78 |
| mt-Co1 | 1.45E-82 | 2.004782 | 0.99 | 1 | 3.43E-78 |
| Gcg | 1.18E-80 | -2.69023 | 0 | 0.301 | 2.78E-76 |
| Sparc | 1.07E-79 | 1.428147 | 0.997 | 1 | 2.53E-75 |
| Plp1 | 2.01E-79 | 0.543983 | 0.027 | 0.437 | 4.74E-75 |
| P2ry12 | 8.94E-78 | 1.319135 | 0.997 | 1 | 2.11E-73 |
| Eef1a1 | 1.94E-76 | 1.626355 | 0.995 | 1 | 4.57E-72 |
| Ctsb | 1.44E-75 | 1.517297 | 0.998 | 1 | 3.4E-71 |
| Gsn | 6.46E-70 | 0.456469 | 0.064 | 0.533 | 1.52E-65 |
| Ctsd | 1.02E-69 | 1.06006 | 1 | 1 | 2.42E-65 |
| Psap | 5.41E-69 | 1.375392 | 0.999 | 1 | 1.28E-64 |
| Cd9 | 2.57E-68 | 1.959819 | 0.975 | 0.991 | 6.05E-64 |
| Ctsz | 2.31E-65 | 1.517976 | 0.995 | 1 | 5.45E-61 |
| Iapp | 1.14E-61 | -0.79015 | 0 | 0.231 | 2.68E-57 |
| Malat1 | 1.51E-60 | 1.395726 | 0.974 | 1 | 3.56E-56 |
| Ins1 | 1.82E-60 | -1.04135 | 0.004 | 0.249 | 4.29E-56 |
| Ftl1 | 5.05E-60 | 1.736279 | 0.989 | 1 | 1.19E-55 |
| Fth1 | 3.23E-57 | 1.732452 | 0.997 | 1 | 7.62E-53 |
| C1qb | 1.18E-56 | 1.130378 | 1 | 1 | 2.78E-52 |
| mt-Co2 | 1.74E-55 | 2.272692 | 0.93 | 0.642 | 4.1E-51 |
| Slc2a5 | 2.18E-55 | 1.906715 | 0.934 | 0.956 | 5.14E-51 |
| Gm1821 | 2.34E-54 | 1.544619 | 0.971 | 0.983 | 5.51E-50 |
| Cyba | 4.48E-53 | 1.73309 | 0.975 | 1 | 1.06E-48 |
| Fos | 3.4E-52 | -1.21245 | 0.12 | 0.576 | 8.02E-48 |
| Trem2 | 3.79E-51 | 1.613965 | 0.982 | 1 | 8.93E-47 |
| B2m | 3.94E-51 | 1.427054 | 0.996 | 1 | 9.3E-47 |
| Gm15710 | 4.19E-51 | 2.075646 | 0.806 | 0.284 | 9.89E-47 |
| Siglech | 5.46E-51 | 1.25478 | 0.995 | 1 | 1.29E-46 |
| Pld4 | 5.49E-51 | 1.710181 | 0.948 | 1 | 1.3E-46 |

| | | | | | |
|---|---|---|---|---|---|
| mt-Cytb | 9.08E-51 | 1.684504 | 0.975 | 0.996 | 2.14E-46 |
| Olfml3 | 1.04E-50 | 1.237013 | 0.988 | 1 | 2.45E-46 |
| Unc93b1 | 1.05E-50 | 1.250821 | 0.986 | 1 | 2.48E-46 |
| mt-Co3 | 2.35E-49 | 1.6385 | 0.966 | 0.764 | 5.54E-45 |
| Rnaset2a | 7.2E-49 | 1.626602 | 0.962 | 0.974 | 1.7E-44 |
| Tmbim6 | 2.67E-48 | 1.483447 | 0.967 | 1 | 6.29E-44 |
| Ltf | 2.64E-46 | -0.77955 | 0.004 | 0.201 | 6.24E-42 |
| Gm11478 | 2.55E-45 | 2.689593 | 0.722 | 0.201 | 6.01E-41 |
| Gm23935 | 1.87E-44 | -0.8164 | 0.836 | 1 | 4.41E-40 |
| Pfn1 | 2.09E-44 | 1.822958 | 0.954 | 0.996 | 4.93E-40 |
| Krt15 | 4.7E-44 | -0.79249 | 0.035 | 0.314 | 1.11E-39 |
| F11r | 1.91E-43 | 1.763818 | 0.947 | 1 | 4.51E-39 |
| Cx3cr1 | 2.35E-43 | 1.069247 | 1 | 1 | 5.55E-39 |
| Rpl13a | 2.48E-43 | 2.578267 | 0.859 | 0.751 | 5.85E-39 |
| Ubb | 2.04E-42 | 1.547364 | 0.977 | 1 | 4.81E-38 |
| Abi3 | 2.05E-42 | 2.307179 | 0.864 | 0.965 | 4.83E-38 |
| Gm5559 | 2.1E-42 | 2.64645 | 0.761 | 0.31 | 4.94E-38 |
| Ctss | 2.84E-42 | 1.000591 | 1 | 1 | 6.7E-38 |
| Bsg | 4.04E-42 | 2.09094 | 0.869 | 0.961 | 9.53E-38 |
| Lcn2 | 8.32E-42 | -0.77421 | 0.011 | 0.214 | 1.96E-37 |
| Gm24270 | 1.33E-41 | -1.13517 | 0.415 | 0.769 | 3.14E-37 |
| Lpcat2 | 2.29E-41 | 1.142142 | 0.984 | 1 | 5.41E-37 |
| Irf8 | 3.64E-41 | 2.218168 | 0.891 | 0.926 | 8.58E-37 |
| Cst3 | 2.54E-40 | 1.131909 | 1 | 1 | 5.98E-36 |
| Ifngr1 | 3.48E-40 | 1.1135 | 0.975 | 0.991 | 8.21E-36 |
| Otulinl | 3.65E-40 | 1.658309 | 0.931 | 0.969 | 8.61E-36 |
| Ctsh | 4.05E-39 | 1.528448 | 0.94 | 0.996 | 9.56E-35 |
| Selenop | 4.47E-39 | 1.074497 | 0.989 | 1 | 1.05E-34 |
| Itm2c | 4.74E-39 | 1.58214 | 0.911 | 1 | 1.12E-34 |
| Rpl13 | 1.13E-37 | 1.870934 | 0.943 | 0.956 | 2.66E-33 |
| Rgs10 | 1.35E-37 | 1.392476 | 0.956 | 1 | 3.18E-33 |
| Laptm4a | 1.36E-37 | 1.626195 | 0.903 | 0.996 | 3.2E-33 |
| Cd68 | 1.57E-37 | 1.338766 | 0.954 | 1 | 3.7E-33 |
| Rps9 | 1.17E-36 | 1.571397 | 0.951 | 0.987 | 2.76E-32 |
| Ptgs1 | 1.98E-36 | 1.70153 | 0.929 | 0.978 | 4.67E-32 |
| Gm24187 | 4.19E-36 | -0.99036 | 0.47 | 0.843 | 9.89E-32 |
| Rnaset2b | 5.38E-36 | 1.504746 | 0.975 | 0.934 | 1.27E-31 |
| Ctsa | 1.11E-35 | 1.310819 | 0.967 | 0.996 | 2.63E-31 |
| Abhd12 | 1.13E-35 | 1.186248 | 0.964 | 1 | 2.67E-31 |
| Adap2os | 1.14E-35 | 2.220091 | 0.947 | 0.965 | 2.7E-31 |
| Gm8797 | 2E-35 | 1.226642 | 0.933 | 0.803 | 4.72E-31 |
| Dusp1 | 2.13E-35 | -1.20671 | 0.081 | 0.389 | 5.03E-31 |
| Tmsb4x | 2.33E-35 | 1.314515 | 1 | 1 | 5.51E-31 |
| Cotl1 | 9.93E-35 | 2.636815 | 0.809 | 0.869 | 2.34E-30 |
| Ptgds | 2.17E-34 | 0.600692 | 0.016 | 0.205 | 5.13E-30 |
| Gm15564 | 3.68E-34 | -0.34221 | 0.113 | 0.415 | 8.68E-30 |
| Spi1 | 1.45E-33 | 2.14472 | 0.861 | 0.974 | 3.41E-29 |

| | | | | | |
|---|---|---|---|---|---|
| Calm2 | 1.53E-33 | 1.379358 | 0.945 | 0.996 | 3.61E-29 |
| Rps27rt | 6.27E-33 | 1.422843 | 0.714 | 0.249 | 1.48E-28 |
| Lars2 | 9.68E-33 | -0.66968 | 0.755 | 1 | 2.28E-28 |
| Vsir | 1.57E-32 | 1.085818 | 0.967 | 1 | 3.69E-28 |
| Clic1 | 2E-32 | 1.543501 | 0.874 | 0.939 | 4.73E-28 |
| Gm42418 | 2.84E-32 | 3.428877 | 0.687 | 0.406 | 6.69E-28 |
| Ctsl | 3.42E-32 | 1.288013 | 0.946 | 0.996 | 8.06E-28 |
| Gm3379 | 4.41E-32 | 1.869461 | 0.636 | 0.201 | 1.04E-27 |
| Serpine2 | 7.37E-32 | 1.53516 | 0.903 | 0.969 | 1.74E-27 |
| Hexa | 8.24E-32 | 1.101542 | 0.957 | 0.991 | 1.94E-27 |
| Bin1 | 1.03E-31 | 1.690961 | 0.898 | 0.987 | 2.42E-27 |
| Gm15427 | 1.06E-31 | 1.479973 | 0.949 | 0.843 | 2.49E-27 |
| Grn | 1.12E-31 | 0.94005 | 0.994 | 1 | 2.65E-27 |
| Actb | 6.35E-31 | 0.957061 | 1 | 1 | 1.5E-26 |
| Rps3 | 7.09E-31 | 1.68354 | 0.901 | 0.961 | 1.67E-26 |
| H2-D1 | 1.37E-30 | 1.89344 | 0.894 | 0.983 | 3.23E-26 |
| Rplp0 | 1.95E-30 | 1.563331 | 0.898 | 0.974 | 4.6E-26 |
| Ac154912.3 | 2.47E-30 | -1.99699 | 0.05 | 0.288 | 5.84E-26 |
| Fcrls | 5.84E-30 | 1.024678 | 0.975 | 1 | 1.38E-25 |
| Trf | 6.15E-30 | 1.077375 | 0.97 | 0.991 | 1.45E-25 |
| Gpr34 | 1.73E-29 | 1.013255 | 0.989 | 1 | 4.07E-25 |
| Gm6977 | 8.55E-29 | 1.455023 | 0.632 | 0.214 | 2.02E-24 |
| Mptx1 | 2.26E-28 | 0.731705 | 0.024 | 0.21 | 5.33E-24 |
| Rsrp1 | 3.31E-28 | 1.836131 | 0.9 | 0.965 | 7.81E-24 |
| Gm15266 | 4.76E-28 | 1.213174 | 0.623 | 0.175 | 1.12E-23 |
| Rpl23a-ps3 | 5.54E-28 | 1.15386 | 0.553 | 0.127 | 1.31E-23 |
| mt-Atp6 | 5.95E-28 | 1.27201 | 0.902 | 0.616 | 1.4E-23 |
| Atp6v0c-ps | 1.21E-27 | 1.573574 | 0.741 | 0.362 | 2.85E-23 |
| Gm9843 | 1.49E-27 | 1.675078 | 0.867 | 0.616 | 3.51E-23 |
| Rnase4 | 1.73E-27 | 1.154967 | 0.928 | 0.991 | 4.08E-23 |
| H3f3b | 1.75E-27 | 1.343693 | 0.903 | 0.991 | 4.12E-23 |
| Rpl3 | 1.78E-27 | 1.938416 | 0.835 | 0.86 | 4.21E-23 |
| Atp6v0c | 2.82E-27 | 1.618328 | 0.842 | 0.926 | 6.66E-23 |
| Gm12918 | 2.96E-27 | 0.998722 | 0.556 | 0.127 | 6.97E-23 |
| Rps16-ps2 | 3.32E-27 | 1.649893 | 0.623 | 0.214 | 7.82E-23 |
| Zfhx3 | 5.53E-27 | 1.593745 | 0.939 | 0.965 | 1.3E-22 |
| Rps3a1 | 1.35E-26 | 1.629166 | 0.856 | 0.869 | 3.18E-22 |
| Gm6204 | 1.49E-26 | 1.367613 | 0.623 | 0.201 | 3.51E-22 |
| Gm11518 | 1.58E-26 | 2.085473 | 0.513 | 0.127 | 3.73E-22 |
| Gm26825 | 1.65E-26 | 1.676247 | 0.794 | 0.511 | 3.88E-22 |
| Tpm3 | 3.58E-26 | 2.02973 | 0.899 | 0.987 | 8.44E-22 |
| Rps18-ps3 | 4.84E-26 | 1.278715 | 0.648 | 0.253 | 1.14E-21 |
| Scamp2 | 5.18E-26 | 1.383914 | 0.891 | 0.987 | 1.22E-21 |
| Ly86 | 5.84E-26 | 1.01496 | 0.989 | 1 | 1.38E-21 |
| Gm10123 | 7.02E-26 | 1.32289 | 0.935 | 0.917 | 1.66E-21 |
| Rpl10 | 7.53E-26 | 1.568293 | 0.904 | 0.943 | 1.78E-21 |
| Adap2 | 1.38E-25 | 1.546553 | 0.929 | 0.987 | 3.24E-21 |

| | | | | | |
|---|---|---|---|---|---|
| Serinc3 | 2.26E-25 | -0.55233 | 0.996 | 1 | 5.33E-21 |
| Ftl1-ps1 | 2.62E-25 | 1.224576 | 0.906 | 0.585 | 6.17E-21 |
| Gm8960 | 3.36E-25 | 1.06002 | 0.381 | 0.022 | 7.93E-21 |
| Gm6394 | 3.86E-25 | 1.217144 | 0.547 | 0.148 | 9.11E-21 |
| Sft2d1 | 4.24E-25 | 1.33395 | 0.904 | 0.978 | 1E-20 |
| Gm10359 | 6.39E-25 | 0.939029 | 0.427 | 0.052 | 1.51E-20 |
| Tmem59 | 6.86E-25 | 1.211473 | 0.882 | 0.983 | 1.62E-20 |
| Aif1 | 1.28E-24 | 1.411259 | 0.933 | 0.965 | 3.03E-20 |
| Gnai2 | 1.52E-24 | 1.37148 | 0.947 | 1 | 3.59E-20 |
| Gapdh | 4.22E-24 | 1.321668 | 0.891 | 0.987 | 9.95E-20 |
| Jun | 8.31E-24 | -0.43518 | 0.331 | 0.764 | 1.96E-19 |
| Gm11942 | 1.59E-23 | 0.942476 | 0.467 | 0.096 | 3.75E-19 |
| Syngr1 | 1.77E-23 | 1.898464 | 0.828 | 0.856 | 4.18E-19 |
| Rpl27-ps3 | 2.02E-23 | 1.217996 | 0.434 | 0.07 | 4.76E-19 |
| mt-Nd1 | 2.15E-23 | 1.170367 | 0.949 | 0.991 | 5.08E-19 |
| Rpl18a | 2.24E-23 | 1.287411 | 0.982 | 0.996 | 5.29E-19 |
| Gm49369 | 3.59E-23 | 1.212535 | 0.812 | 0.795 | 8.47E-19 |
| Gstm1 | 3.71E-23 | 0.792965 | 0.086 | 0.336 | 8.75E-19 |
| Tmem50a | 1.65E-22 | 1.483469 | 0.84 | 0.948 | 3.9E-18 |
| Rpl10-ps3 | 1.7E-22 | 1.837235 | 0.665 | 0.297 | 4E-18 |
| Gm5879 | 1.95E-22 | 0.873832 | 0.481 | 0.105 | 4.6E-18 |
| Gnb1 | 2.14E-22 | 2.150764 | 0.865 | 0.93 | 5.05E-18 |
| Gm3362 | 2.38E-22 | 0.298038 | 0.401 | 0.048 | 5.6E-18 |
| Srsf2 | 2.83E-22 | 1.681741 | 0.885 | 0.856 | 6.67E-18 |
| Rpl4 | 3.98E-22 | 1.774778 | 0.82 | 0.895 | 9.4E-18 |
| Gm5138 | 4.19E-22 | 1.908926 | 0.477 | 0.122 | 9.87E-18 |
| Gm7536 | 4.38E-22 | 1.448209 | 0.619 | 0.275 | 1.03E-17 |
| Rps23-ps1 | 4.61E-22 | 0.92486 | 0.455 | 0.096 | 1.09E-17 |
| Gm16378 | 5.88E-22 | 0.716014 | 0.346 | 0.022 | 1.39E-17 |
| Syngr2 | 1E-21 | 1.586564 | 0.82 | 0.93 | 2.36E-17 |
| Npc2 | 1.39E-21 | 1.706959 | 0.842 | 0.921 | 3.27E-17 |
| Egr1 | 1.63E-21 | -1.26049 | 0.046 | 0.231 | 3.85E-17 |
| P2ry13 | 1.67E-21 | 1.020691 | 0.97 | 1 | 3.93E-17 |
| Gm2574 | 1.68E-21 | 1.200944 | 0.542 | 0.192 | 3.96E-17 |
| Gm43028 | 1.71E-21 | 0.688334 | 0.377 | 0.044 | 4.02E-17 |
| Anxa2 | 1.94E-21 | 0.355032 | 0.067 | 0.284 | 4.59E-17 |
| Bin2 | 2.11E-21 | 1.371792 | 0.885 | 0.991 | 4.97E-17 |
| Gm7336 | 2.58E-21 | 1.427368 | 0.456 | 0.114 | 6.08E-17 |
| Marcks | 4.04E-21 | 0.864838 | 0.989 | 1 | 9.53E-17 |
| Mtdh | 7.71E-21 | 1.455688 | 0.93 | 0.987 | 1.82E-16 |
| Crlf2 | 9.23E-21 | 3.891894 | 0.469 | 0.175 | 2.18E-16 |
| Ywhah | 1.29E-20 | 1.195356 | 0.891 | 0.991 | 3.05E-16 |
| Rps10-ps1 | 1.83E-20 | 2.160187 | 0.522 | 0.188 | 4.31E-16 |
| Fkbp5 | 2.22E-20 | -0.45005 | 0.237 | 0.568 | 5.23E-16 |
| Capzb | 2.27E-20 | 1.656293 | 0.801 | 0.878 | 5.36E-16 |
| Gm6472 | 2.29E-20 | 0.99951 | 0.49 | 0.135 | 5.4E-16 |
| Arhgdia | 5.88E-20 | 1.989124 | 0.797 | 0.908 | 1.39E-15 |

| | | | | | |
|---|---|---|---|---|---|
| Gm8730 | 9E-20 | 1.253955 | 0.758 | 0.467 | 2.12E-15 |
| Rps6-ps4 | 9.02E-20 | 0.985597 | 0.586 | 0.223 | 2.13E-15 |
| Gm12537 | 1.34E-19 | 0.991624 | 0.412 | 0.087 | 3.15E-15 |
| Cd37 | 1.58E-19 | 1.428584 | 0.846 | 0.969 | 3.74E-15 |
| Ptafr | 2.03E-19 | 2.108529 | 0.818 | 0.808 | 4.78E-15 |
| Rps6 | 2.47E-19 | 1.575736 | 0.811 | 0.747 | 5.82E-15 |
| Slc25a5 | 2.76E-19 | 1.553226 | 0.836 | 0.934 | 6.5E-15 |
| Rpl7a-ps7 | 2.94E-19 | 1.013026 | 0.323 | 0.026 | 6.94E-15 |
| Gm6900 | 3.29E-19 | 0.739543 | 0.292 | 0.009 | 7.76E-15 |
| Gm6180 | 4.09E-19 | 0.660992 | 0.435 | 0.096 | 9.65E-15 |
| Gm6170 | 4.24E-19 | 0.482797 | 0.485 | 0.131 | 1E-14 |
| Gm10073 | 5.64E-19 | 0.845773 | 0.588 | 0.21 | 1.33E-14 |
| Camk1 | 8.15E-19 | 1.596635 | 0.787 | 0.869 | 1.92E-14 |
| Sat1 | 9.56E-19 | 1.39279 | 0.884 | 0.921 | 2.26E-14 |
| Arpc1b | 9.84E-19 | 1.899704 | 0.778 | 0.908 | 2.32E-14 |
| Atp6v0b | 9.86E-19 | 1.353712 | 0.883 | 0.956 | 2.33E-14 |
| Gm10288 | 1.25E-18 | 0.968505 | 0.491 | 0.153 | 2.95E-14 |
| Lair1 | 1.69E-18 | 1.435284 | 0.877 | 0.974 | 3.98E-14 |
| Ftl2-ps | 2.03E-18 | 1.346914 | 0.94 | 0.908 | 4.78E-14 |
| Tmem86a | 2.29E-18 | 2.035722 | 0.759 | 0.843 | 5.41E-14 |
| Rpl18-ps1 | 2.41E-18 | 0.412844 | 0.353 | 0.048 | 5.69E-14 |
| Rps12-ps9 | 3.07E-18 | 1.568245 | 0.412 | 0.105 | 7.25E-14 |
| Cd53 | 5.07E-18 | 0.854545 | 0.961 | 1 | 1.2E-13 |
| Zfp691 | 5.51E-18 | 2.101083 | 0.695 | 0.48 | 1.3E-13 |
| Rps4x | 5.97E-18 | 1.798546 | 0.808 | 0.812 | 1.41E-13 |
| Rps11 | 6.18E-18 | 1.34259 | 0.935 | 0.952 | 1.46E-13 |
| Rpl3-ps1 | 7.87E-18 | 1.440059 | 0.735 | 0.524 | 1.86E-13 |
| Rps7 | 9.59E-18 | 1.704275 | 0.793 | 0.782 | 2.26E-13 |
| Wdr89 | 9.66E-18 | 1.306159 | 0.567 | 0.218 | 2.28E-13 |
| H2-K1 | 1.05E-17 | 1.931831 | 0.788 | 0.9 | 2.47E-13 |
| Canx | 1.14E-17 | 1.260602 | 0.944 | 0.991 | 2.7E-13 |
| Ppia | 1.16E-17 | 1.089255 | 0.977 | 1 | 2.74E-13 |
| Rpl17-ps3 | 1.23E-17 | 0.915492 | 0.424 | 0.109 | 2.91E-13 |
| Gm9625 | 1.98E-17 | 1.010708 | 0.549 | 0.21 | 4.66E-13 |
| Gm44705 | 2.17E-17 | 0.661504 | 0.261 | 0.004 | 5.11E-13 |
| Per1 | 2.86E-17 | -1.08923 | 0.055 | 0.227 | 6.74E-13 |
| Gm5835 | 4.27E-17 | 1.070747 | 0.425 | 0.118 | 1.01E-12 |
| Comt | 4.35E-17 | 1.502834 | 0.819 | 0.843 | 1.03E-12 |
| Rps10-ps2 | 5.03E-17 | 1.684445 | 0.582 | 0.301 | 1.19E-12 |
| Gm13456 | 6.07E-17 | 0.882259 | 0.968 | 0.9 | 1.43E-12 |
| Hspa8 | 6.13E-17 | 1.039547 | 0.929 | 1 | 1.45E-12 |
| Susd3 | 6.58E-17 | 1.623932 | 0.799 | 0.908 | 1.55E-12 |
| Gm9616 | 7.66E-17 | 1.088363 | 0.528 | 0.205 | 1.81E-12 |
| Rpl19-ps11 | 8.46E-17 | 1.217237 | 0.65 | 0.336 | 2E-12 |
| Ckb | 8.64E-17 | 1.171959 | 0.859 | 0.991 | 2.04E-12 |
| Skil | 9.06E-17 | 1.903509 | 0.828 | 0.769 | 2.14E-12 |
| Gm5905 | 9.68E-17 | 1.279518 | 0.696 | 0.397 | 2.28E-12 |

| | | | | | |
|---|---|---|---|---|---|
| Ywhae | 1.27E-16 | 1.838178 | 0.806 | 0.838 | 2.99E-12 |
| Gm4149 | 1.54E-16 | 1.085511 | 0.796 | 0.502 | 3.64E-12 |
| Dpysl2 | 1.55E-16 | 1.343547 | 0.893 | 0.956 | 3.65E-12 |
| Arglu1 | 1.81E-16 | 1.770944 | 0.886 | 0.821 | 4.27E-12 |
| Oaz1 | 1.84E-16 | 1.574276 | 0.813 | 0.882 | 4.35E-12 |
| Il6ra | 1.92E-16 | 1.808097 | 0.863 | 0.969 | 4.53E-12 |
| Upk1b | 2.01E-16 | 2.344642 | 0.563 | 0.336 | 4.75E-12 |
| Gm10020 | 2.04E-16 | 0.475072 | 0.287 | 0.026 | 4.8E-12 |
| Gm2225 | 2.2E-16 | 0.586224 | 0.336 | 0.057 | 5.18E-12 |
| Rn7sk | 2.57E-16 | -1.13173 | 0.184 | 0.397 | 6.05E-12 |
| Hspa5 | 2.84E-16 | 1.220762 | 0.867 | 0.978 | 6.69E-12 |
| Actr2 | 2.87E-16 | 1.706347 | 0.906 | 0.948 | 6.77E-12 |
| Rps2-ps10 | 2.98E-16 | 0.858469 | 0.333 | 0.057 | 7.03E-12 |
| Gm11263 | 3.1E-16 | 0.53906 | 0.323 | 0.048 | 7.32E-12 |
| Ftl1-ps2 | 5.35E-16 | 0.423571 | 0.344 | 0.061 | 1.26E-11 |
| Gm43305 | 5.57E-16 | 1.774764 | 0.586 | 0.314 | 1.31E-11 |
| Ldhb | 7.62E-16 | 1.587289 | 0.794 | 0.948 | 1.8E-11 |
| Arpc4 | 8.37E-16 | 1.624277 | 0.817 | 0.93 | 1.97E-11 |
| Anxa3 | 9.4E-16 | 1.48428 | 0.806 | 0.878 | 2.22E-11 |
| Fcer1g | 9.42E-16 | 1.020688 | 0.99 | 1 | 2.22E-11 |
| Dhrs3 | 1.23E-15 | 1.658054 | 0.853 | 0.956 | 2.91E-11 |
| Rpl6 | 1.28E-15 | 1.802549 | 0.752 | 0.747 | 3.03E-11 |
| Lamp1 | 1.43E-15 | 1.024698 | 0.874 | 0.996 | 3.36E-11 |
| Crybb1 | 1.62E-15 | 1.881458 | 0.759 | 0.847 | 3.82E-11 |
| Gm8430 | 2.2E-15 | 1.207378 | 0.532 | 0.223 | 5.19E-11 |
| Rplp1 | 2.32E-15 | 1.297632 | 0.957 | 0.969 | 5.47E-11 |
| Rpl9-ps6 | 3.13E-15 | 1.60435 | 0.67 | 0.476 | 7.38E-11 |
| Fcgr3 | 3.7E-15 | 0.864799 | 0.959 | 0.996 | 8.72E-11 |
| Gm10180 | 4.86E-15 | 0.647466 | 0.415 | 0.122 | 1.15E-10 |
| Lag3 | 5.41E-15 | 1.349598 | 0.834 | 0.948 | 1.28E-10 |
| Gm6988 | 5.53E-15 | 0.444681 | 0.327 | 0.057 | 1.3E-10 |
| Coro1a | 9.27E-15 | 1.217524 | 0.838 | 0.952 | 2.19E-10 |
| Basp1 | 9.85E-15 | 2.053306 | 0.836 | 0.965 | 2.32E-10 |
| Akr1a1 | 9.89E-15 | 1.734675 | 0.748 | 0.769 | 2.33E-10 |
| Rps13-ps2 | 9.97E-15 | 1.101884 | 0.546 | 0.245 | 2.35E-10 |
| Rpl31-ps8 | 1.08E-14 | 1.24587 | 0.573 | 0.275 | 2.55E-10 |
| Gm29667 | 1.63E-14 | 1.31223 | 0.312 | 0.066 | 3.84E-10 |
| Golm1 | 2.21E-14 | 1.305725 | 0.87 | 0.943 | 5.21E-10 |
| Fcgr2b | 2.77E-14 | 1.418118 | 0.83 | 0.926 | 6.53E-10 |
| Ac165079.: | 2.79E-14 | 0.687332 | 0.313 | 0.061 | 6.59E-10 |
| Rpl10a-ps1 | 3.42E-14 | 1.350115 | 0.573 | 0.31 | 8.07E-10 |
| Rps20 | 3.43E-14 | 1.573061 | 0.802 | 0.834 | 8.08E-10 |
| Gm4617 | 4.84E-14 | 0.925782 | 0.328 | 0.074 | 1.14E-09 |
| Gm12166 | 5.75E-14 | 1.144574 | 0.849 | 0.729 | 1.36E-09 |
| Gm11539 | 5.85E-14 | 0.664386 | 0.326 | 0.07 | 1.38E-09 |
| Gm12286 | 6.86E-14 | 0.467851 | 0.253 | 0.026 | 1.62E-09 |
| Sh2b3 | 8.98E-14 | 0.423049 | 0.329 | 0.624 | 2.12E-09 |

| | | | | | |
|---|---|---|---|---|---|
| Gm15772 | 9.51E-14 | 0.978391 | 0.461 | 0.179 | 2.24E-09 |
| Rps27a-ps2 | 9.81E-14 | 1.14454 | 0.32 | 0.074 | 2.31E-09 |
| Gdi2 | 1.08E-13 | 1.122095 | 0.911 | 0.956 | 2.56E-09 |
| Qk | 1.4E-13 | 1.217155 | 0.928 | 0.969 | 3.3E-09 |
| Cd63-ps | 1.53E-13 | 1.150022 | 0.562 | 0.293 | 3.61E-09 |
| Rbm39 | 1.56E-13 | 1.299511 | 0.892 | 0.939 | 3.68E-09 |
| Ppib | 1.83E-13 | 1.43322 | 0.729 | 0.838 | 4.32E-09 |
| Gm13408 | 2.26E-13 | 1.095651 | 0.282 | 0.048 | 5.33E-09 |
| Rpl37rt | 2.94E-13 | 0.441914 | 0.61 | 0.297 | 6.94E-09 |
| Gm5453 | 3.02E-13 | 1.624273 | 0.235 | 0.022 | 7.11E-09 |
| Cyth4 | 3.07E-13 | 1.24153 | 0.874 | 0.978 | 7.25E-09 |
| Gm11560 | 3.62E-13 | 2.070257 | 0.299 | 0.074 | 8.53E-09 |
| Clta | 4.06E-13 | 1.294058 | 0.821 | 0.917 | 9.57E-09 |
| Actg1 | 4.2E-13 | 1.112642 | 0.814 | 0.978 | 9.9E-09 |
| Gm10231 | 5.18E-13 | 1.139774 | 0.281 | 0.057 | 1.22E-08 |
| Gm6210 | 5.45E-13 | 0.470788 | 0.471 | 0.179 | 1.29E-08 |
| mt-Rnr2 | 5.67E-13 | 0.827046 | 0.989 | 0.996 | 1.34E-08 |
| Atp2a2 | 5.81E-13 | 0.494761 | 0.203 | 0.445 | 1.37E-08 |
| Rpl28-ps1 | 6.98E-13 | 1.387646 | 0.415 | 0.157 | 1.65E-08 |
| Erp29 | 9.17E-13 | 1.55181 | 0.757 | 0.869 | 2.16E-08 |
| H2-DMa | 1.11E-12 | 1.73845 | 0.68 | 0.707 | 2.63E-08 |
| Gm4294 | 1.19E-12 | 0.484222 | 0.399 | 0.131 | 2.81E-08 |
| Ywhaz | 1.26E-12 | 1.361361 | 0.887 | 0.934 | 2.97E-08 |
| Arpc2 | 1.31E-12 | 1.277025 | 0.834 | 0.934 | 3.1E-08 |
| Zscan26 | 1.35E-12 | 1.845884 | 0.815 | 0.803 | 3.19E-08 |
| Rpl9 | 1.39E-12 | 1.399218 | 0.766 | 0.751 | 3.28E-08 |
| Rpsa | 1.48E-12 | 1.549843 | 0.759 | 0.873 | 3.48E-08 |
| Mcl1 | 1.98E-12 | 0.323317 | 0.384 | 0.703 | 4.66E-08 |
| Gm17241 | 2.05E-12 | 0.650881 | 0.376 | 0.114 | 4.84E-08 |
| Rpl36-ps2 | 2.08E-12 | 1.788153 | 0.511 | 0.227 | 4.91E-08 |
| Atf3 | 2.17E-12 | 1.152035 | 0.074 | 0.236 | 5.11E-08 |
| Eno1b | 2.18E-12 | 1.90117 | 0.602 | 0.454 | 5.14E-08 |
| Gm13493 | 2.2E-12 | 0.800915 | 0.771 | 0.489 | 5.19E-08 |
| Rps3a2 | 2.37E-12 | 0.743823 | 0.41 | 0.144 | 5.58E-08 |
| Rpl17 | 2.44E-12 | 1.452795 | 0.796 | 0.755 | 5.75E-08 |
| Gm5805 | 2.57E-12 | 0.519711 | 0.423 | 0.144 | 6.07E-08 |
| Gm8226 | 2.89E-12 | 0.629074 | 0.231 | 0.026 | 6.82E-08 |
| Gm14760 | 3.57E-12 | 0.440509 | 0.221 | 0.022 | 8.41E-08 |
| Pmp22 | 3.97E-12 | 1.434978 | 0.772 | 0.821 | 9.36E-08 |
| Gm6063 | 4.31E-12 | 0.549598 | 0.273 | 0.057 | 1.02E-07 |
| Abcg1 | 4.87E-12 | 2.180639 | 0.693 | 0.598 | 1.15E-07 |
| Cd300c2 | 5.53E-12 | 1.277923 | 0.811 | 0.965 | 1.31E-07 |
| Limd2 | 6.12E-12 | 1.777571 | 0.754 | 0.856 | 1.44E-07 |
| Cd14 | 6.45E-12 | 1.437928 | 0.766 | 0.921 | 1.52E-07 |
| Mbp | 7.74E-12 | 1.460632 | 0.097 | 0.275 | 1.82E-07 |
| Ubc | 7.85E-12 | 0.676936 | 0.969 | 0.991 | 1.85E-07 |
| Tmed9 | 8.17E-12 | 2.150288 | 0.639 | 0.686 | 1.93E-07 |

| | | | | | |
|---|---|---|---|---|---|
| Gng10 | 8.74E-12 | 1.293677 | 0.749 | 0.882 | 2.06E-07 |
| Rhoa | 1.02E-11 | 1.305392 | 0.849 | 0.921 | 2.42E-07 |
| Api5 | 1.11E-11 | -0.52189 | 0.172 | 0.362 | 2.62E-07 |
| Gpx4-ps2 | 1.37E-11 | 1.469937 | 0.443 | 0.231 | 3.23E-07 |
| Notch2 | 1.59E-11 | 2.167217 | 0.736 | 0.624 | 3.75E-07 |
| Gm5514 | 1.69E-11 | 0.929238 | 0.535 | 0.275 | 3.98E-07 |
| Rpl21-ps15 | 2.18E-11 | 0.642378 | 0.349 | 0.109 | 5.15E-07 |
| Gm11361 | 2.41E-11 | 0.434891 | 0.454 | 0.179 | 5.69E-07 |
| Mafb | 2.46E-11 | 1.490053 | 0.856 | 0.987 | 5.81E-07 |
| Arrb2 | 2.51E-11 | 2.477426 | 0.676 | 0.729 | 5.92E-07 |
| Gm12396 | 2.55E-11 | 1.051091 | 0.377 | 0.148 | 6.02E-07 |
| Atp6ap1 | 2.64E-11 | 1.142085 | 0.782 | 0.93 | 6.22E-07 |
| Rpl7 | 2.66E-11 | 1.352722 | 0.781 | 0.838 | 6.26E-07 |
| Tagap | 2.68E-11 | -0.7139 | 0.335 | 0.559 | 6.32E-07 |
| Rps3a3 | 3.34E-11 | 0.38854 | 0.423 | 0.157 | 7.88E-07 |
| Rpsa-ps9 | 3.39E-11 | 0.723948 | 0.353 | 0.122 | 7.99E-07 |
| Foxn3 | 3.61E-11 | 1.59864 | 0.867 | 0.904 | 8.52E-07 |
| Gm10443 | 3.62E-11 | 0.310446 | 0.332 | 0.092 | 8.55E-07 |
| Gm10175 | 3.76E-11 | 0.95841 | 0.21 | 0.026 | 8.87E-07 |
| Rps19-ps6 | 4.28E-11 | 0.919439 | 0.644 | 0.376 | 1.01E-06 |
| Cramp1l | 4.38E-11 | -0.53797 | 0.082 | 0.231 | 1.03E-06 |
| Gm9800 | 4.97E-11 | 0.868115 | 0.279 | 0.07 | 1.17E-06 |
| Rpl15-ps3 | 5.12E-11 | 0.659682 | 0.316 | 0.092 | 1.21E-06 |
| Rpl11 | 5.53E-11 | 1.439782 | 0.753 | 0.699 | 1.31E-06 |
| Rpl13a-ps1 | 5.98E-11 | 0.589562 | 0.2 | 0.022 | 1.41E-06 |
| Gm10240 | 7.21E-11 | 0.749566 | 0.411 | 0.162 | 1.7E-06 |
| Eif1 | 7.58E-11 | 1.238599 | 0.881 | 0.904 | 1.79E-06 |
| Gm5637 | 7.97E-11 | 0.866716 | 0.327 | 0.109 | 1.88E-06 |
| Rpl13-ps6 | 8.07E-11 | 0.816658 | 0.492 | 0.231 | 1.9E-06 |
| Rps13-ps1 | 8.9E-11 | 0.381531 | 0.305 | 0.083 | 2.1E-06 |
| Gm13680 | 8.94E-11 | 1.060295 | 0.43 | 0.192 | 2.11E-06 |
| Ecscr | 1.07E-10 | 1.597727 | 0.679 | 0.703 | 2.52E-06 |
| Cnbp | 1.26E-10 | 1.154473 | 0.792 | 0.891 | 2.98E-06 |
| Oaz1-ps | 1.29E-10 | 1.191042 | 0.72 | 0.546 | 3.05E-06 |
| Gm10163 | 1.37E-10 | 2.133043 | 0.448 | 0.214 | 3.22E-06 |
| Gm11808 | 1.4E-10 | 0.964839 | 0.441 | 0.214 | 3.29E-06 |
| Gm5869 | 1.4E-10 | 0.395965 | 0.328 | 0.096 | 3.31E-06 |
| Gm6863 | 1.5E-10 | 1.178179 | 0.396 | 0.179 | 3.54E-06 |
| Efhd2 | 1.58E-10 | 1.641744 | 0.799 | 0.974 | 3.72E-06 |
| Tpt1-ps3 | 1.64E-10 | 0.709838 | 0.605 | 0.332 | 3.87E-06 |
| Rps18 | 2.01E-10 | 1.329345 | 0.847 | 0.882 | 4.75E-06 |
| mt-Nd4 | 2.29E-10 | 0.984228 | 0.896 | 0.987 | 5.39E-06 |
| Gm12191 | 2.54E-10 | 0.781214 | 0.274 | 0.07 | 5.99E-06 |
| Gm10275 | 2.54E-10 | 1.043505 | 0.366 | 0.144 | 6E-06 |
| Rpl14-ps1 | 2.65E-10 | 1.204511 | 0.598 | 0.38 | 6.25E-06 |
| Gm10224 | 2.73E-10 | 0.386335 | 0.209 | 0.031 | 6.45E-06 |
| Rps14 | 2.86E-10 | 1.102696 | 0.915 | 0.913 | 6.75E-06 |

| | | | | | |
|---|---|---|---|---|---|
| Gm12892 | 3.11E-10 | 0.437949 | 0.245 | 0.052 | 7.35E-06 |
| Rpl9-ps7 | 3.34E-10 | 0.930213 | 0.361 | 0.144 | 7.89E-06 |
| Tmsb10 | 3.64E-10 | 2.58056 | 0.441 | 0.279 | 8.59E-06 |
| Gm14303 | 3.74E-10 | 0.644223 | 0.655 | 0.393 | 8.81E-06 |
| Tpt1 | 3.78E-10 | 1.256329 | 0.842 | 0.939 | 8.92E-06 |
| Gm5586 | 3.97E-10 | 0.47989 | 0.211 | 0.035 | 9.38E-06 |
| Commd8 | 4.04E-10 | 1.558957 | 0.695 | 0.651 | 9.52E-06 |
| Calr | 4.35E-10 | 0.89834 | 0.905 | 0.991 | 1.03E-05 |
| Cdk2ap1 | 4.65E-10 | 0.33559 | 0.306 | 0.59 | 1.1E-05 |
| Rpl19 | 5.26E-10 | 1.283208 | 0.813 | 0.838 | 1.24E-05 |
| Gm9493 | 5.81E-10 | 0.454971 | 0.27 | 0.07 | 1.37E-05 |
| Olfr77 | 5.88E-10 | 0.487229 | 0.299 | 0.092 | 1.39E-05 |
| Ang | 6.08E-10 | 1.852652 | 0.747 | 0.834 | 1.43E-05 |
| Rpl34-ps1 | 6.13E-10 | 0.902876 | 0.469 | 0.223 | 1.45E-05 |
| Arf1 | 6.17E-10 | 1.305091 | 0.758 | 0.917 | 1.45E-05 |
| Gm8172 | 6.47E-10 | 1.31163 | 0.456 | 0.262 | 1.53E-05 |
| Bhlhe41 | 6.49E-10 | 1.790085 | 0.833 | 0.852 | 1.53E-05 |
| Rpl3-ps2 | 6.55E-10 | 0.410723 | 0.262 | 0.07 | 1.55E-05 |
| Gm7935 | 6.91E-10 | 0.66577 | 0.211 | 0.039 | 1.63E-05 |
| Gm14494 | 7.6E-10 | 0.743506 | 0.238 | 0.052 | 1.79E-05 |
| Gm5560 | 7.84E-10 | 0.413263 | 0.421 | 0.17 | 1.85E-05 |
| Rbm26 | 7.9E-10 | 1.693803 | 0.732 | 0.716 | 1.86E-05 |
| Cmtm7 | 8.25E-10 | 1.253195 | 0.803 | 0.882 | 1.95E-05 |
| Zfp36 | 9.5E-10 | 0.585185 | 0.456 | 0.777 | 2.24E-05 |
| Cd63 | 1.04E-09 | 1.562877 | 0.736 | 0.812 | 2.46E-05 |
| Rgmb | 1.11E-09 | 0.271591 | 0.338 | 0.581 | 2.62E-05 |
| Rps10 | 1.13E-09 | 1.386701 | 0.755 | 0.799 | 2.65E-05 |
| Hpgds | 1.16E-09 | 0.959118 | 0.927 | 0.996 | 2.73E-05 |
| Gm2a | 1.17E-09 | 1.490488 | 0.774 | 0.755 | 2.76E-05 |
| Capza2 | 1.19E-09 | 0.894058 | 0.893 | 0.965 | 2.81E-05 |
| Ncoa3 | 1.28E-09 | 1.752108 | 0.771 | 0.738 | 3.02E-05 |
| Mylip | 1.3E-09 | 1.737158 | 0.679 | 0.611 | 3.06E-05 |
| Rpl14 | 1.38E-09 | 1.455056 | 0.771 | 0.799 | 3.26E-05 |
| Gm9794 | 1.41E-09 | 0.884081 | 0.602 | 0.358 | 3.33E-05 |
| Ndfip1 | 1.41E-09 | 1.35933 | 0.7 | 0.803 | 3.34E-05 |
| Gm10051 | 1.51E-09 | 0.739943 | 0.24 | 0.057 | 3.55E-05 |
| Rps25-ps1 | 1.61E-09 | 0.36023 | 0.304 | 0.092 | 3.8E-05 |
| Gm16437 | 1.67E-09 | 0.557959 | 0.258 | 0.07 | 3.93E-05 |
| Arhgdib | 1.78E-09 | 1.476896 | 0.743 | 0.852 | 4.2E-05 |
| Gm2000 | 1.81E-09 | 0.808792 | 0.573 | 0.314 | 4.27E-05 |
| Creg1 | 1.92E-09 | 1.444701 | 0.715 | 0.742 | 4.52E-05 |
| Tesk1 | 1.94E-09 | 0.269886 | 0.072 | 0.205 | 4.59E-05 |
| Gna15 | 1.95E-09 | 1.569851 | 0.669 | 0.681 | 4.61E-05 |
| Col6a3 | 1.96E-09 | 4.217716 | 0.229 | 0.061 | 4.62E-05 |
| Bri3 | 2.01E-09 | 2.519494 | 0.587 | 0.581 | 4.74E-05 |
| Pdia3 | 2.23E-09 | 0.599593 | 0.955 | 1 | 5.26E-05 |
| Sec24b | 2.54E-09 | 0.364473 | 0.139 | 0.31 | 5.99E-05 |

| | | | | | |
|---|---|---|---|---|---|
| Gm8129 | 2.59E-09 | 0.561381 | 0.233 | 0.057 | 6.12E-05 |
| Bmp2k | 2.87E-09 | 1.343289 | 0.929 | 0.987 | 6.78E-05 |
| Rpsa-ps10 | 2.91E-09 | 0.910212 | 0.594 | 0.384 | 6.87E-05 |
| Gm10157 | 3.23E-09 | 0.364483 | 0.221 | 0.048 | 7.63E-05 |
| Usp2 | 3.27E-09 | 1.982909 | 0.663 | 0.729 | 7.71E-05 |
| Alox5ap | 3.52E-09 | 1.245026 | 0.761 | 0.9 | 8.3E-05 |
| Ifitm10 | 3.91E-09 | 2.852004 | 0.516 | 0.48 | 9.22E-05 |
| Gm10257 | 4.33E-09 | 0.580376 | 0.238 | 0.061 | 0.000102 |
| Hist1h4d | 4.36E-09 | -0.60349 | 0.264 | 0.459 | 0.000103 |
| Ttyh3 | 4.42E-09 | 1.1752 | 0.217 | 0.428 | 0.000104 |
| H3f3a-ps2 | 4.55E-09 | 0.548011 | 0.221 | 0.048 | 0.000107 |
| Hmgb1 | 5.28E-09 | 1.2475 | 0.778 | 0.799 | 0.000125 |
| Gm9703 | 5.3E-09 | 0.269305 | 0.272 | 0.079 | 0.000125 |
| Gm12428 | 5.44E-09 | 0.392416 | 0.316 | 0.109 | 0.000128 |
| Gm10036 | 5.44E-09 | 0.667549 | 0.343 | 0.135 | 0.000128 |
| Rps15a-ps6 | 5.81E-09 | 1.469312 | 0.263 | 0.074 | 0.000137 |
| Sept2 | 5.95E-09 | 1.329209 | 0.76 | 0.79 | 0.00014 |
| Bst2 | 6.65E-09 | 2.454507 | 0.435 | 0.306 | 0.000157 |
| Gm3511 | 6.71E-09 | 0.471155 | 0.453 | 0.218 | 0.000158 |
| Rpl19-ps1 | 6.74E-09 | 0.422939 | 0.254 | 0.07 | 0.000159 |
| Spcs2 | 7.32E-09 | 1.189718 | 0.718 | 0.721 | 0.000173 |
| Gm6136 | 7.51E-09 | 1.239014 | 0.567 | 0.38 | 0.000177 |
| Jade2 | 7.88E-09 | 0.840248 | 0.179 | 0.367 | 0.000186 |
| Ctsc | 7.97E-09 | 0.87151 | 0.872 | 0.991 | 0.000188 |
| Tpm3-rs7 | 8.59E-09 | 1.131398 | 0.585 | 0.393 | 0.000203 |
| Ddost | 8.6E-09 | 1.561419 | 0.703 | 0.803 | 0.000203 |
| Arhgap5 | 9.64E-09 | 1.648175 | 0.721 | 0.734 | 0.000227 |
| Gm7308 | 1.02E-08 | 0.386312 | 0.258 | 0.074 | 0.000241 |
| Arap1 | 1.03E-08 | 0.69489 | 0.272 | 0.485 | 0.000244 |
| Lrrc3 | 1.08E-08 | 1.458878 | 0.873 | 0.948 | 0.000256 |
| Rpl30-ps10 | 1.26E-08 | 0.451244 | 0.22 | 0.052 | 0.000298 |
| Kcnk6 | 1.29E-08 | 0.863101 | 0.419 | 0.716 | 0.000304 |
| Arl6ip1 | 1.32E-08 | 1.240972 | 0.765 | 0.825 | 0.000313 |
| Gm8566 | 1.5E-08 | 0.338904 | 0.219 | 0.052 | 0.000353 |
| Ltc4s | 1.59E-08 | 1.294829 | 0.87 | 0.934 | 0.000376 |
| Ac114990.3 | 1.59E-08 | 0.994101 | 0.256 | 0.087 | 0.000376 |
| Rpl17-ps5 | 1.64E-08 | 0.461976 | 0.264 | 0.083 | 0.000386 |
| Rpl27a | 1.64E-08 | 1.828062 | 0.76 | 0.721 | 0.000387 |
| Gm29216 | 1.66E-08 | 0.377236 | 0.776 | 0.467 | 0.000391 |
| Gm15501 | 1.8E-08 | 1.121339 | 0.468 | 0.249 | 0.000424 |
| Zfp710 | 1.81E-08 | 2.443777 | 0.68 | 0.694 | 0.000426 |
| Selenof | 2.03E-08 | 1.246997 | 0.685 | 0.742 | 0.000478 |
| Gm28661 | 2.04E-08 | 0.494739 | 0.812 | 0.55 | 0.000481 |
| Rps24-ps3 | 2.05E-08 | 0.472054 | 0.255 | 0.074 | 0.000484 |
| Ptms | 2.05E-08 | 2.555303 | 0.651 | 0.699 | 0.000484 |
| Eno1 | 2.1E-08 | 1.60145 | 0.697 | 0.808 | 0.000495 |
| Tubb5 | 2.17E-08 | 1.478432 | 0.793 | 0.869 | 0.000511 |

| | | | | | |
|---|---|---|---|---|---|
| Gm12350 | 2.26E-08 | 1.420089 | 0.258 | 0.079 | 0.000534 |
| C5ar2 | 2.29E-08 | 2.063334 | 0.694 | 0.655 | 0.000539 |
| Ppp1r3b | 2.39E-08 | 0.332925 | 0.138 | 0.293 | 0.000565 |
| Lat2 | 3.13E-08 | 2.16392 | 0.405 | 0.271 | 0.000738 |
| Gm3839 | 3.7E-08 | 0.297605 | 0.208 | 0.052 | 0.000872 |
| Gm4735 | 3.86E-08 | 0.703694 | 0.454 | 0.245 | 0.000911 |
| Maf | 4.04E-08 | 1.235676 | 0.867 | 0.921 | 0.000954 |
| Itpripl1 | 4.06E-08 | 1.652763 | 0.74 | 0.79 | 0.000958 |
| Akna | 4.34E-08 | 0.785571 | 0.309 | 0.524 | 0.001024 |
| Coa5 | 4.42E-08 | -0.45226 | 0.118 | 0.258 | 0.001042 |
| Gm5915 | 5.02E-08 | 1.395608 | 0.239 | 0.074 | 0.001184 |
| Ptp4a2 | 5.13E-08 | 1.628269 | 0.747 | 0.777 | 0.001211 |
| Id2 | 5.57E-08 | -0.79469 | 0.152 | 0.31 | 0.001313 |
| Hs6st1 | 5.78E-08 | 0.678744 | 0.222 | 0.419 | 0.001363 |
| Tcn2 | 6.82E-08 | 0.948641 | 0.808 | 0.978 | 0.001609 |
| Pycard | 7.15E-08 | 1.18753 | 0.793 | 0.856 | 0.001688 |
| Gm10059 | 7.99E-08 | 0.264259 | 0.252 | 0.079 | 0.001885 |
| Saraf | 8.03E-08 | 1.096812 | 0.73 | 0.908 | 0.001895 |
| Gpx1 | 8.68E-08 | 1.469835 | 0.694 | 0.734 | 0.002048 |
| Snhg20 | 8.82E-08 | 2.152966 | 0.32 | 0.153 | 0.002081 |
| Selenok | 8.91E-08 | 1.494347 | 0.658 | 0.707 | 0.002102 |
| Tmem87b | 9.43E-08 | -0.28152 | 0.118 | 0.258 | 0.002225 |
| Rps2 | 1.2E-07 | 1.52236 | 0.693 | 0.703 | 0.002837 |
| Mfsd14b | 1.28E-07 | -0.56709 | 0.172 | 0.319 | 0.003021 |
| Rpl18 | 1.38E-07 | 1.248089 | 0.789 | 0.865 | 0.003246 |
| Rpl7-ps7 | 1.38E-07 | 0.334044 | 0.285 | 0.109 | 0.003256 |
| Pla2g15 | 1.55E-07 | 1.168274 | 0.787 | 0.895 | 0.003655 |
| Gm13453 | 1.57E-07 | 0.513919 | 0.302 | 0.122 | 0.003709 |
| Evi2a | 1.72E-07 | 1.453922 | 0.62 | 0.633 | 0.004046 |
| Tapbp | 1.8E-07 | 1.304276 | 0.772 | 0.838 | 0.004248 |
| Gm9385 | 1.82E-07 | 0.550738 | 0.319 | 0.131 | 0.004293 |
| Gm3724 | 1.89E-07 | 1.347672 | 0.282 | 0.109 | 0.004458 |
| Hsp90b1 | 1.91E-07 | 0.630139 | 0.909 | 0.996 | 0.004498 |
| Nrros | 1.92E-07 | 1.501948 | 0.634 | 0.651 | 0.004519 |
| Cbx3 | 1.93E-07 | 1.823767 | 0.494 | 0.341 | 0.004551 |
| B4galt1 | 2.23E-07 | 1.257671 | 0.872 | 0.908 | 0.005252 |
| Arhgap45 | 2.37E-07 | 1.119313 | 0.84 | 0.93 | 0.005599 |
| Zfp385a | 2.41E-07 | 2.312739 | 0.273 | 0.114 | 0.005693 |
| Lemd2 | 2.44E-07 | 0.347368 | 0.167 | 0.328 | 0.005755 |
| Atp5b | 2.83E-07 | 1.206902 | 0.725 | 0.93 | 0.006682 |
| Rpl32 | 3E-07 | 1.218032 | 0.849 | 0.869 | 0.007077 |
| Rps11-ps1 | 3.09E-07 | 1.103751 | 0.207 | 0.057 | 0.007283 |
| Gm9347 | 3.27E-07 | 0.482734 | 0.246 | 0.087 | 0.007725 |
| Rps19 | 3.31E-07 | 1.14347 | 0.879 | 0.869 | 0.007808 |
| Rpl21 | 3.39E-07 | 0.919194 | 0.919 | 0.948 | 0.007996 |
| Gm7331 | 3.57E-07 | 0.850808 | 0.47 | 0.284 | 0.008413 |
| Arid5a | 3.63E-07 | 0.563754 | 0.09 | 0.214 | 0.008559 |

| | | | | | |
|---|---|---|---|---|---|
| Lmo2 | 4.2E-07 | 1.914816 | 0.606 | 0.611 | 0.009912 |
| Sidt2 | 4.21E-07 | 0.552249 | 0.212 | 0.384 | 0.009941 |
| Gm12844 | 4.92E-07 | 0.618148 | 0.23 | 0.079 | 0.011605 |
| Snx5 | 5.18E-07 | 1.108441 | 0.81 | 0.93 | 0.012213 |
| Ptpn6 | 5.21E-07 | 1.388934 | 0.704 | 0.843 | 0.012284 |
| Tspan7 | 5.26E-07 | 1.923928 | 0.574 | 0.585 | 0.012402 |
| Cd300a | 6.2E-07 | 1.546928 | 0.616 | 0.616 | 0.014623 |
| Pdia6 | 6.69E-07 | 1.114699 | 0.734 | 0.891 | 0.015776 |
| Gm5529 | 6.86E-07 | 0.302249 | 0.424 | 0.205 | 0.016177 |
| Inpp5d | 6.92E-07 | 1.130797 | 0.898 | 0.974 | 0.016332 |
| Sult1a1 | 7.05E-07 | 0.411021 | 0.236 | 0.432 | 0.016631 |
| Aldoa | 7.08E-07 | 1.444826 | 0.704 | 0.843 | 0.0167 |
| Fuca1 | 7.21E-07 | 1.787906 | 0.546 | 0.511 | 0.017 |
| Nisch | 7.46E-07 | 0.765815 | 0.377 | 0.607 | 0.017598 |
| Ncoa4 | 7.48E-07 | -0.35556 | 0.223 | 0.371 | 0.017651 |
| Gm7901 | 7.7E-07 | 0.996329 | 0.317 | 0.162 | 0.018161 |
| Gm10800 | 7.73E-07 | 1.05445 | 0.645 | 0.472 | 0.018236 |
| Rac1 | 7.8E-07 | 1.327634 | 0.767 | 0.974 | 0.018388 |
| Marf1 | 7.9E-07 | 2.149222 | 0.633 | 0.581 | 0.018625 |
| Mag | 8.33E-07 | 0.938716 | 0.102 | 0.231 | 0.019638 |
| Ssfa2 | 8.41E-07 | 0.518278 | 0.152 | 0.297 | 0.01984 |
| Zfp207 | 8.48E-07 | 1.472291 | 0.626 | 0.511 | 0.020001 |
| Rpl7a | 8.83E-07 | 1.495987 | 0.674 | 0.699 | 0.020835 |
| Bnip3l-ps | 9.27E-07 | 0.703772 | 0.292 | 0.131 | 0.021866 |
| Psmb8 | 9.43E-07 | 1.520252 | 0.602 | 0.585 | 0.022243 |
| Itga9 | 9.97E-07 | 0.861195 | 0.216 | 0.393 | 0.023519 |
| Zfp652 | 1.02E-06 | 1.208055 | 0.701 | 0.498 | 0.024066 |
| Cox4i1 | 1.09E-06 | 1.234884 | 0.764 | 0.843 | 0.025779 |
| Phyhd1 | 1.09E-06 | 1.669115 | 0.548 | 0.524 | 0.025801 |
| Hist1h1e | 1.15E-06 | 0.346082 | 0.203 | 0.384 | 0.027196 |
| Ier2 | 1.22E-06 | 0.362043 | 0.087 | 0.201 | 0.028738 |
| Hnrnpk | 1.22E-06 | 0.92338 | 0.831 | 0.926 | 0.028882 |
| Tpst2 | 1.23E-06 | 1.437645 | 0.667 | 0.755 | 0.028941 |
| Dcaf7 | 1.24E-06 | 0.716542 | 0.238 | 0.437 | 0.029201 |
| Apbb1ip | 1.26E-06 | 2.348592 | 0.663 | 0.76 | 0.029755 |
| Map2k4 | 1.34E-06 | 0.263617 | 0.131 | 0.262 | 0.031722 |
| Cerk | 1.38E-06 | 0.619855 | 0.323 | 0.528 | 0.03247 |
| Cebpb | 1.44E-06 | 0.649282 | 0.434 | 0.782 | 0.033938 |
| Tyrobp | 1.48E-06 | 0.81804 | 0.995 | 1 | 0.034912 |
| Pip4p1 | 1.52E-06 | 1.771739 | 0.687 | 0.769 | 0.035774 |
| Trim28 | 1.52E-06 | 0.497651 | 0.248 | 0.424 | 0.035947 |
| Cdc42 | 1.63E-06 | 0.718584 | 0.871 | 0.996 | 0.038497 |
| Arpc3 | 1.71E-06 | 1.477908 | 0.637 | 0.721 | 0.040425 |
| Ucp2 | 1.77E-06 | 0.388653 | 0.581 | 0.965 | 0.041755 |
| Rgs2 | 1.86E-06 | 1.089901 | 0.806 | 0.913 | 0.043924 |
| Gm38158 | 1.92E-06 | 2.253457 | 0.363 | 0.253 | 0.045388 |
| Iscu | 2.01E-06 | 1.383127 | 0.593 | 0.572 | 0.047404 |

### Supplementry Table 4. Aging DEGs in Hippocampus

| Gene | p_val | avg_log2FC | pct.1 | pct.2 | p_val_adj |
|---|---|---|---|---|---|
| C1qa | 3.2E-166 | 2.20964 | 1 | 1 | 7.5E-162 |
| C1qc | 3.5E-153 | 1.770819 | 1 | 1 | 8.3E-149 |
| Cst3 | 2.5E-144 | 1.566852 | 1 | 1 | 5.9E-140 |
| Plp1 | 4.7E-140 | -0.31776 | 0.023 | 0.589 | 1.1E-135 |
| Itm2b | 4.3E-138 | 1.305902 | 1 | 1 | 1E-133 |
| Cfl1 | 2E-127 | 3.174299 | 0.968 | 0.959 | 4.6E-123 |
| S100a8 | 5E-127 | -0.97569 | 0.007 | 0.482 | 1.2E-122 |
| C1qb | 1.3E-126 | 1.33286 | 1 | 1 | 3E-122 |
| Selplg | 4.5E-126 | 1.387974 | 0.999 | 1 | 1.1E-121 |
| Gsn | 1.2E-125 | -1.15294 | 0.05 | 0.639 | 2.9E-121 |
| Laptm5 | 1.7E-124 | 1.259993 | 1 | 1 | 4E-120 |
| Cd81 | 5.5E-124 | 1.458362 | 1 | 1 | 1.3E-119 |
| Sparc | 9.2E-123 | 1.497359 | 1 | 1 | 2.2E-118 |
| Tmsb4x | 2E-121 | 1.684218 | 1 | 1 | 4.8E-117 |
| B2m | 7.9E-120 | 1.774941 | 0.999 | 1 | 1.9E-115 |
| Gm15564 | 4.9E-119 | -0.37258 | 0.076 | 0.657 | 1.2E-114 |
| S100a9 | 8.6E-119 | -1.05677 | 0.009 | 0.462 | 2E-114 |
| Ac154912.3 | 4E-118 | -3.08926 | 0.032 | 0.544 | 9.4E-114 |
| Lgmn | 3.1E-117 | 1.355586 | 1 | 1 | 7.4E-113 |
| Tmem119 | 4.1E-112 | 1.536644 | 0.997 | 1 | 9.7E-108 |
| mt-Co1 | 2.2E-110 | 1.993499 | 0.994 | 1 | 5.1E-106 |
| Ctss | 2.2E-106 | 1.188542 | 1 | 1 | 5.3E-102 |
| Ngp | 3.7E-102 | -0.89144 | 0 | 0.367 | 8.81E-98 |
| Ins2 | 2.07E-97 | -2.00211 | 0.008 | 0.385 | 4.89E-93 |
| Eef1a1 | 2.46E-97 | 1.72357 | 0.997 | 1 | 5.81E-93 |
| Ctsz | 7.73E-97 | 1.542733 | 0.99 | 0.994 | 1.82E-92 |
| Pfn1 | 4.72E-95 | 2.341094 | 0.953 | 0.979 | 1.11E-90 |
| Fth1 | 6.88E-95 | 1.929984 | 0.99 | 1 | 1.62E-90 |
| mt-Co2 | 1.22E-92 | 2.287388 | 0.943 | 0.754 | 2.87E-88 |
| Trem2 | 6.01E-89 | 1.596797 | 0.992 | 1 | 1.42E-84 |
| mt-Cytb | 6.6E-89 | 1.872819 | 0.981 | 0.982 | 1.56E-84 |
| Fcer1g | 4.04E-86 | 1.401742 | 0.995 | 1 | 9.53E-82 |
| Cd9 | 4.93E-86 | 1.715651 | 0.988 | 0.967 | 1.16E-81 |
| Cyba | 5.1E-86 | 1.967413 | 0.968 | 0.985 | 1.2E-81 |
| mt-Co3 | 6.13E-86 | 1.925971 | 0.965 | 0.861 | 1.45E-81 |
| Ct010467.1 | 4.27E-85 | 0.6433 | 1 | 1 | 1.01E-80 |
| Gcg | 9.3E-85 | -1.25489 | 0.006 | 0.337 | 2.19E-80 |
| Hexb | 7.24E-84 | 0.919576 | 1 | 1 | 1.71E-79 |
| Ftl1 | 1.71E-83 | 1.815528 | 0.987 | 1 | 4.04E-79 |
| mt-Atp6 | 3.95E-82 | 1.914218 | 0.922 | 0.672 | 9.32E-78 |
| Rps9 | 1.54E-81 | 1.971185 | 0.967 | 0.97 | 3.63E-77 |
| Ac154912.1 | 2.58E-79 | -1.45823 | 0.042 | 0.45 | 6.08E-75 |
| Rpl13 | 8.12E-79 | 2.195114 | 0.957 | 0.935 | 1.92E-74 |

| | | | | | |
|---|---|---|---|---|---|
| Ctsb | 2.9E-78 | 1.30712 | 0.999 | 1 | 6.85E-74 |
| Aif1 | 4.29E-75 | 1.989158 | 0.942 | 0.926 | 1.01E-70 |
| Hist1h1e | 1.6E-73 | -1.98004 | 0.105 | 0.574 | 3.77E-69 |
| Mid1 | 4.85E-72 | -1.41348 | 0.068 | 0.488 | 1.14E-67 |
| Psap | 4.87E-71 | 1.208883 | 0.998 | 1 | 1.15E-66 |
| Rgs10 | 6.51E-71 | 1.476395 | 0.971 | 0.988 | 1.54E-66 |
| Iapp | 1.29E-69 | -1.43043 | 0 | 0.254 | 3.04E-65 |
| Fkbp5 | 1.59E-69 | -1.14891 | 0.184 | 0.71 | 3.75E-65 |
| Tubb4a | 9.25E-69 | -1.07605 | 0.002 | 0.26 | 2.18E-64 |
| Rpl18a | 1.16E-68 | 1.684043 | 0.986 | 0.997 | 2.75E-64 |
| Rpl13a | 4.67E-68 | 3.198879 | 0.864 | 0.831 | 1.1E-63 |
| Gm15427 | 2.33E-67 | 2.256625 | 0.942 | 0.896 | 5.5E-63 |
| Olfml3 | 7.27E-67 | 1.110217 | 0.997 | 1 | 1.72E-62 |
| Gm15710 | 3.98E-66 | 2.201802 | 0.789 | 0.382 | 9.39E-62 |
| Pld4 | 4.36E-66 | 1.456462 | 0.964 | 0.973 | 1.03E-61 |
| Jun | 5.06E-64 | -1.39806 | 0.305 | 0.825 | 1.19E-59 |
| Ins1 | 6.57E-63 | -1.39192 | 0.005 | 0.254 | 1.55E-58 |
| Bsg | 1.88E-62 | 2.232939 | 0.879 | 0.92 | 4.44E-58 |
| Siglech | 1.97E-62 | 1.076596 | 0.999 | 1 | 4.65E-58 |
| Rplp1 | 1.42E-61 | 1.816279 | 0.96 | 0.962 | 3.34E-57 |
| Dcn | 1.52E-61 | 1.032114 | 0.036 | 0.37 | 3.58E-57 |
| Ctsd | 2.75E-61 | 0.841997 | 1 | 1 | 6.5E-57 |
| Rnaset2a | 1.14E-59 | 1.725775 | 0.938 | 0.979 | 2.7E-55 |
| Calm2 | 1.83E-58 | 1.597813 | 0.943 | 0.988 | 4.31E-54 |
| Ubb | 9.71E-58 | 1.378349 | 0.984 | 0.988 | 2.29E-53 |
| Clic1 | 1.09E-57 | 1.875196 | 0.889 | 0.888 | 2.57E-53 |
| Laptm4a | 3.29E-57 | 1.634788 | 0.912 | 0.962 | 7.75E-53 |
| Rplp0 | 4.21E-57 | 2.003586 | 0.914 | 0.941 | 9.93E-53 |
| 4933406I1[illegible] | 2.4E-56 | -1.48616 | 0.095 | 0.488 | 5.66E-52 |
| Rps11 | 6.12E-56 | 1.756573 | 0.93 | 0.938 | 1.44E-51 |
| Slc2a5 | 6.47E-56 | 1.585221 | 0.947 | 0.932 | 1.53E-51 |
| Tyrobp | 1.06E-55 | 1.110912 | 0.997 | 1 | 2.51E-51 |
| Chga | 1.38E-55 | -0.79141 | 0 | 0.204 | 3.25E-51 |
| Abhd12 | 1.64E-55 | 1.265977 | 0.974 | 1 | 3.86E-51 |
| Rps3 | 4.13E-55 | 1.941693 | 0.906 | 0.935 | 9.75E-51 |
| mt-Nd1 | 4.5E-55 | 1.542602 | 0.962 | 0.964 | 1.06E-50 |
| Adap2os | 3.46E-53 | 1.914587 | 0.948 | 0.976 | 8.17E-49 |
| Basp1 | 8.31E-53 | 2.432305 | 0.903 | 0.97 | 1.96E-48 |
| F11r | 1.74E-52 | 1.620341 | 0.953 | 0.988 | 4.1E-48 |
| Gm44860 | 6.15E-52 | -1.28051 | 0.234 | 0.669 | 1.45E-47 |
| Ppia | 6.16E-52 | 1.323139 | 0.963 | 0.991 | 1.45E-47 |
| Ctsh | 8.91E-52 | 1.386568 | 0.96 | 0.967 | 2.1E-47 |
| Gm5559 | 9.17E-52 | 2.596438 | 0.774 | 0.5 | 2.16E-47 |
| Irf8 | 1.53E-51 | 2.201351 | 0.878 | 0.876 | 3.62E-47 |
| Gm15726 | 2.12E-51 | -2.06549 | 0.023 | 0.275 | 5.01E-47 |
| Gapdh | 7.3E-51 | 1.539166 | 0.936 | 0.962 | 1.72E-46 |
| Rps3a1 | 1.12E-50 | 1.937884 | 0.88 | 0.885 | 2.65E-46 |

| | | | | | |
|---|---|---|---|---|---|
| Cd68 | 2.29E-50 | 1.365817 | 0.945 | 0.97 | 5.39E-46 |
| Rn7sk | 3.14E-50 | -0.98238 | 0.164 | 0.574 | 7.42E-46 |
| Rps14 | 5.81E-49 | 1.753724 | 0.918 | 0.938 | 1.37E-44 |
| Rps27rt | 8.14E-49 | 1.812534 | 0.763 | 0.459 | 1.92E-44 |
| Gm11478 | 1.04E-48 | 2.894968 | 0.675 | 0.299 | 2.46E-44 |
| mt-Rnr1 | 2.79E-48 | -1.22384 | 0.827 | 0.988 | 6.58E-44 |
| Rnaset2b | 2.87E-48 | 1.651613 | 0.958 | 0.959 | 6.78E-44 |
| P2ry12 | 1.16E-47 | 0.760672 | 0.998 | 1 | 2.73E-43 |
| Gm1821 | 1.82E-47 | 1.494605 | 0.954 | 0.985 | 4.29E-43 |
| Spi1 | 3.9E-47 | 2.159991 | 0.854 | 0.891 | 9.2E-43 |
| Tpt1 | 1.82E-46 | 2.061929 | 0.899 | 0.917 | 4.28E-42 |
| Abi3 | 3.12E-46 | 2.056604 | 0.852 | 0.95 | 7.37E-42 |
| Tmbim6 | 4.73E-46 | 1.29106 | 0.96 | 0.985 | 1.12E-41 |
| Gm4149 | 2.01E-45 | 1.844164 | 0.812 | 0.624 | 4.75E-41 |
| Dhx9 | 5.74E-45 | -1.04374 | 0.09 | 0.423 | 1.35E-40 |
| Jmjd1c | 7.4E-45 | -1.01607 | 0.225 | 0.642 | 1.74E-40 |
| Gm9843 | 2.32E-44 | 2.265571 | 0.86 | 0.725 | 5.48E-40 |
| Cotl1 | 2.34E-44 | 2.803024 | 0.798 | 0.825 | 5.53E-40 |
| Itm2c | 2.53E-44 | 1.530566 | 0.918 | 0.959 | 5.96E-40 |
| Unc93b1 | 2.91E-44 | 1.136095 | 0.991 | 0.997 | 6.86E-40 |
| Tlr7 | 4.37E-44 | -0.67788 | 0.307 | 0.76 | 1.03E-39 |
| Runx1 | 1.03E-43 | -0.80895 | 0.224 | 0.633 | 2.44E-39 |
| Hist1h4d | 1.05E-43 | -1.88335 | 0.219 | 0.645 | 2.47E-39 |
| Gm24270 | 1.9E-43 | -0.38618 | 0.487 | 0.923 | 4.49E-39 |
| Cd37 | 6.33E-43 | 1.706491 | 0.889 | 0.932 | 1.49E-38 |
| H2-D1 | 1.56E-42 | 2.056101 | 0.882 | 0.905 | 3.67E-38 |
| Rpl21 | 2.81E-42 | 1.797746 | 0.921 | 0.956 | 6.62E-38 |
| Gm15266 | 3.02E-42 | 1.672681 | 0.672 | 0.337 | 7.13E-38 |
| Adrb1 | 7.92E-42 | -0.30181 | 0.183 | 0.598 | 1.87E-37 |
| Rpl10 | 8.01E-42 | 1.504062 | 0.918 | 0.935 | 1.89E-37 |
| Rpl3 | 1.36E-41 | 2.038635 | 0.834 | 0.855 | 3.2E-37 |
| Gm10076 | 1.76E-41 | 1.643016 | 0.892 | 0.917 | 4.15E-37 |
| Rgs1 | 3.69E-41 | -1.4392 | 0.08 | 0.388 | 8.72E-37 |
| Npc2 | 4.72E-41 | 2.003904 | 0.848 | 0.879 | 1.11E-36 |
| Mgp | 6.11E-41 | -0.68738 | 0.014 | 0.21 | 1.44E-36 |
| Teddm2 | 9.46E-41 | -0.98577 | 0.068 | 0.364 | 2.23E-36 |
| Per1 | 2.01E-40 | -1.0014 | 0.043 | 0.293 | 4.75E-36 |
| Rps7 | 3.11E-40 | 2.32159 | 0.817 | 0.751 | 7.34E-36 |
| Oaz1 | 4.1E-40 | 2.035757 | 0.816 | 0.802 | 9.68E-36 |
| Ly86 | 5.12E-40 | 0.817101 | 0.998 | 1 | 1.21E-35 |
| Lifr | 5.27E-40 | -0.63461 | 0.209 | 0.607 | 1.24E-35 |
| Fos | 7.07E-40 | -1.59491 | 0.163 | 0.55 | 1.67E-35 |
| Rpl23a-ps3 | 1.42E-39 | 1.433192 | 0.547 | 0.16 | 3.36E-35 |
| Gm12918 | 1.52E-39 | 1.318662 | 0.568 | 0.172 | 3.59E-35 |
| Gm45221 | 5.23E-39 | -1.33089 | 0.07 | 0.349 | 1.23E-34 |
| Gm43028 | 5.6E-39 | 1.019909 | 0.452 | 0.065 | 1.32E-34 |
| Gnb1 | 6.2E-39 | 2.478036 | 0.857 | 0.902 | 1.46E-34 |

| | | | | | |
|---|---|---|---|---|---|
| Ly6e | 5.05E-38 | 1.649376 | 0.934 | 0.985 | 1.19E-33 |
| Plek | 7.82E-38 | -0.35391 | 0.394 | 0.849 | 1.84E-33 |
| Bin1 | 2.13E-37 | 1.533892 | 0.904 | 0.964 | 5.03E-33 |
| Rpl41 | 2.61E-37 | 1.412882 | 0.943 | 0.953 | 6.15E-33 |
| Slc8a1 | 3.64E-37 | -2.03639 | 0.11 | 0.408 | 8.59E-33 |
| Ctsl | 6.41E-37 | 1.080366 | 0.956 | 0.994 | 1.51E-32 |
| Rpl17 | 1.26E-36 | 2.165464 | 0.814 | 0.757 | 2.97E-32 |
| Syngr2 | 2.4E-36 | 1.617815 | 0.854 | 0.899 | 5.67E-32 |
| Rpl4 | 2.66E-36 | 1.866869 | 0.854 | 0.891 | 6.28E-32 |
| Gm47664 | 3.91E-36 | -0.96735 | 0.061 | 0.317 | 9.23E-32 |
| Rplp2 | 5.92E-36 | 1.64675 | 0.889 | 0.87 | 1.4E-31 |
| Atp6v0c-ps | 5.99E-36 | 1.92044 | 0.729 | 0.479 | 1.41E-31 |
| Atp6v0c | 6.02E-36 | 1.778104 | 0.835 | 0.876 | 1.42E-31 |
| Man1a | 6.62E-36 | -0.8605 | 0.293 | 0.672 | 1.56E-31 |
| Gm26669 | 1.66E-35 | -1.29855 | 0.023 | 0.219 | 3.92E-31 |
| Dpysl2 | 2.41E-35 | 2.066976 | 0.892 | 0.914 | 5.69E-31 |
| Tmem50a | 2.41E-35 | 1.510288 | 0.876 | 0.911 | 5.69E-31 |
| Rps18 | 3.56E-35 | 1.732055 | 0.867 | 0.822 | 8.39E-31 |
| Rps18-ps3 | 5.07E-35 | 1.84758 | 0.62 | 0.29 | 1.2E-30 |
| Arhgdia | 5.09E-35 | 2.32131 | 0.808 | 0.864 | 1.2E-30 |
| Bin2 | 9.9E-35 | 1.39158 | 0.912 | 0.964 | 2.34E-30 |
| March1 | 1.14E-34 | -0.52134 | 0.225 | 0.618 | 2.68E-30 |
| Clu | 1.16E-34 | -0.67598 | 0.023 | 0.213 | 2.73E-30 |
| Rps20 | 2.24E-34 | 1.880115 | 0.829 | 0.817 | 5.28E-30 |
| Scamp2 | 3.95E-34 | 1.304563 | 0.912 | 0.967 | 9.32E-30 |
| Syngr1 | 3.96E-34 | 1.719228 | 0.849 | 0.879 | 9.34E-30 |
| Rps24 | 7.35E-34 | 1.923932 | 0.842 | 0.867 | 1.73E-29 |
| Gm14303 | 7.4E-34 | 1.198158 | 0.771 | 0.55 | 1.74E-29 |
| Tmem59 | 1.87E-33 | 1.15928 | 0.892 | 0.959 | 4.41E-29 |
| Rps6 | 2.19E-33 | 1.924985 | 0.787 | 0.692 | 5.16E-29 |
| Sft2d1 | 3.73E-33 | 1.325582 | 0.883 | 0.959 | 8.8E-29 |
| Gm3379 | 7.27E-33 | 1.932863 | 0.615 | 0.343 | 1.72E-28 |
| Gnai2 | 1.02E-32 | 1.254206 | 0.941 | 0.976 | 2.4E-28 |
| Lpcat2 | 1.25E-32 | 0.948255 | 0.974 | 0.997 | 2.95E-28 |
| Marcks | 1.37E-32 | 1.062184 | 0.996 | 1 | 3.24E-28 |
| Rapgef5 | 2.41E-32 | -0.3426 | 0.266 | 0.651 | 5.69E-28 |
| Rpl18 | 2.59E-32 | 1.712672 | 0.846 | 0.84 | 6.11E-28 |
| Grn | 2.96E-32 | 0.818605 | 0.99 | 1 | 6.98E-28 |
| Vsir | 3.72E-32 | 0.870629 | 0.973 | 1 | 8.76E-28 |
| Apc | 4.83E-32 | -0.48618 | 0.165 | 0.491 | 1.14E-27 |
| Gm10925 | 4.99E-32 | 1.009753 | 0.91 | 0.775 | 1.18E-27 |
| Gm6204 | 1E-31 | 1.791211 | 0.587 | 0.278 | 2.36E-27 |
| Rsrp1 | 1.5E-31 | 1.501758 | 0.893 | 0.964 | 3.55E-27 |
| Atp6v0b | 1.72E-31 | 1.423741 | 0.897 | 0.917 | 4.05E-27 |
| Gm26825 | 1.86E-31 | 2.026035 | 0.746 | 0.612 | 4.4E-27 |
| Tsc22d3 | 2.71E-31 | -0.57119 | 0.194 | 0.553 | 6.39E-27 |
| Gm10073 | 5.04E-31 | 1.279818 | 0.625 | 0.296 | 1.19E-26 |

| | | | | | |
|---|---|---|---|---|---|
| Rpl6 | 5.66E-31 | 2.16007 | 0.795 | 0.743 | 1.33E-26 |
| Ctsa | 1.36E-30 | 1.044789 | 0.957 | 0.991 | 3.21E-26 |
| Rps12-ps9 | 1.71E-30 | 1.120209 | 0.495 | 0.154 | 4.04E-26 |
| Coro1a | 1.81E-30 | 1.485882 | 0.848 | 0.893 | 4.26E-26 |
| Gm16378 | 2.25E-30 | 0.815504 | 0.33 | 0.018 | 5.3E-26 |
| Fau | 3.1E-30 | 1.499493 | 0.901 | 0.899 | 7.32E-26 |
| Gm2574 | 3.22E-30 | 1.460028 | 0.52 | 0.204 | 7.6E-26 |
| Mcl1 | 3.33E-30 | -0.25707 | 0.392 | 0.787 | 7.86E-26 |
| Tagap | 3.91E-30 | -0.42757 | 0.295 | 0.669 | 9.22E-26 |
| Rpl32 | 5.42E-30 | 1.597509 | 0.855 | 0.888 | 1.28E-25 |
| Serpine2 | 6.22E-30 | 1.160305 | 0.93 | 0.944 | 1.47E-25 |
| Capzb | 1.01E-29 | 1.881353 | 0.788 | 0.84 | 2.39E-25 |
| Tmed7 | 1.21E-29 | -0.54087 | 0.309 | 0.675 | 2.85E-25 |
| Hexa | 1.26E-29 | 0.880631 | 0.975 | 0.997 | 2.98E-25 |
| Mtdh | 2.24E-29 | 1.497054 | 0.912 | 0.97 | 5.28E-25 |
| Gm45223 | 2.3E-29 | -0.67602 | 0.121 | 0.402 | 5.42E-25 |
| Ptgs1 | 4.55E-29 | 1.338351 | 0.931 | 0.944 | 1.07E-24 |
| Rps25 | 4.64E-29 | 1.732441 | 0.884 | 0.896 | 1.09E-24 |
| Susd3 | 5.07E-29 | 1.587309 | 0.844 | 0.902 | 1.2E-24 |
| Gm3362 | 6.48E-29 | 0.34283 | 0.425 | 0.086 | 1.53E-24 |
| Gm45222 | 1E-28 | -0.89014 | 0.045 | 0.246 | 2.37E-24 |
| Ftl1-ps1 | 1.03E-28 | 1.554706 | 0.866 | 0.793 | 2.42E-24 |
| Gm37420 | 1.27E-28 | -1.00506 | 0.089 | 0.34 | 2.99E-24 |
| Kat6a | 1.46E-28 | -0.60506 | 0.206 | 0.53 | 3.44E-24 |
| Arpc4 | 2.69E-28 | 1.737432 | 0.829 | 0.917 | 6.35E-24 |
| Rpl35 | 2.7E-28 | 1.738985 | 0.817 | 0.772 | 6.37E-24 |
| Gm11942 | 3.03E-28 | 1.117267 | 0.49 | 0.172 | 7.15E-24 |
| Gm2000 | 3.49E-28 | 1.598923 | 0.602 | 0.32 | 8.24E-24 |
| Rpl17-ps3 | 3.85E-28 | 1.384956 | 0.456 | 0.139 | 9.09E-24 |
| G530011O( | 4.79E-28 | -0.49719 | 0.282 | 0.654 | 1.13E-23 |
| Rnase4 | 5.5E-28 | 1.004415 | 0.929 | 0.994 | 1.3E-23 |
| Rpl31-ps8 | 5.56E-28 | 1.923297 | 0.629 | 0.385 | 1.31E-23 |
| Macf1 | 8.51E-28 | -0.47445 | 0.617 | 0.95 | 2.01E-23 |
| Abca1 | 9.23E-28 | -0.48171 | 0.391 | 0.769 | 2.18E-23 |
| Tnks2 | 1.87E-27 | -0.43904 | 0.215 | 0.541 | 4.41E-23 |
| Clta | 1.92E-27 | 1.572263 | 0.833 | 0.899 | 4.52E-23 |
| 8030442B( | 2.25E-27 | -0.68264 | 0.088 | 0.328 | 5.31E-23 |
| H3f3b | 2.31E-27 | 1.179588 | 0.899 | 0.95 | 5.46E-23 |
| Rps4x | 2.99E-27 | 1.912683 | 0.808 | 0.79 | 7.04E-23 |
| Gm10180 | 3.94E-27 | 1.035381 | 0.477 | 0.172 | 9.29E-23 |
| Rps10-ps1 | 4.4E-27 | 2.927266 | 0.499 | 0.219 | 1.04E-22 |
| Gm6900 | 4.99E-27 | 0.845529 | 0.311 | 0.024 | 1.18E-22 |
| Alox5ap | 5.87E-27 | 1.586621 | 0.811 | 0.876 | 1.39E-22 |
| Gm29216 | 6.71E-27 | 0.751441 | 0.815 | 0.609 | 1.58E-22 |
| Ssh2 | 7.48E-27 | -0.58811 | 0.735 | 0.976 | 1.76E-22 |
| Crlf2 | 7.51E-27 | 3.977683 | 0.505 | 0.254 | 1.77E-22 |
| Gm10123 | 1.04E-26 | 1.480841 | 0.909 | 0.964 | 2.46E-22 |

| | | | | | |
|---|---|---|---|---|---|
| Cx3cr1 | 1.19E-26 | 0.581232 | 0.999 | 1 | 2.8E-22 |
| Gm7536 | 1.21E-26 | 1.79378 | 0.588 | 0.337 | 2.85E-22 |
| Rps16-ps2 | 1.43E-26 | 1.692623 | 0.602 | 0.346 | 3.38E-22 |
| H2-K1 | 1.59E-26 | 2.043999 | 0.805 | 0.834 | 3.75E-22 |
| Wdr89 | 1.67E-26 | 1.431016 | 0.591 | 0.317 | 3.94E-22 |
| Zfp691 | 2.06E-26 | 2.338423 | 0.69 | 0.538 | 4.87E-22 |
| Sik2 | 3.15E-26 | -0.81492 | 0.079 | 0.305 | 7.43E-22 |
| Ldhb | 6.63E-26 | 1.666361 | 0.809 | 0.896 | 1.56E-21 |
| Gm8730 | 7.03E-26 | 1.560018 | 0.737 | 0.565 | 1.66E-21 |
| Gm5879 | 7.42E-26 | 1.013309 | 0.401 | 0.101 | 1.75E-21 |
| Rps19 | 9.55E-26 | 1.441349 | 0.861 | 0.882 | 2.25E-21 |
| Ptafr | 1.2E-25 | 1.95841 | 0.802 | 0.781 | 2.84E-21 |
| Gm10359 | 1.26E-25 | 1.014088 | 0.419 | 0.124 | 2.97E-21 |
| Ccnd3 | 1.54E-25 | 0.418562 | 0.22 | 0.574 | 3.64E-21 |
| Abca9 | 1.81E-25 | -0.45682 | 0.551 | 0.896 | 4.26E-21 |
| Rpl9 | 2.3E-25 | 1.859819 | 0.786 | 0.766 | 5.43E-21 |
| Rpl30 | 2.73E-25 | 1.437638 | 0.88 | 0.873 | 6.43E-21 |
| Rps23-ps1 | 3.24E-25 | 0.865092 | 0.437 | 0.136 | 7.64E-21 |
| Rps23 | 3.77E-25 | 1.204458 | 0.887 | 0.879 | 8.9E-21 |
| Arpc1b | 4.35E-25 | 1.897948 | 0.775 | 0.84 | 1.03E-20 |
| Rps10 | 5E-25 | 1.845002 | 0.79 | 0.82 | 1.18E-20 |
| Kmt2c | 1.05E-24 | -0.46052 | 0.218 | 0.524 | 2.47E-20 |
| Usp8 | 1.41E-24 | -0.27319 | 0.238 | 0.559 | 3.33E-20 |
| Tpm3 | 1.73E-24 | 1.840945 | 0.885 | 0.959 | 4.09E-20 |
| Aftph | 2.25E-24 | -0.25171 | 0.276 | 0.607 | 5.31E-20 |
| Pros1 | 2.37E-24 | -0.59675 | 0.653 | 0.917 | 5.6E-20 |
| Trf | 2.49E-24 | 0.869949 | 0.965 | 0.994 | 5.89E-20 |
| Slc25a5 | 2.63E-24 | 1.622144 | 0.809 | 0.852 | 6.21E-20 |
| Inpp4b | 3.35E-24 | -0.31138 | 0.183 | 0.479 | 7.91E-20 |
| Ywhah | 3.67E-24 | 1.120939 | 0.914 | 0.982 | 8.65E-20 |
| Rpl37a | 5.85E-24 | 1.300962 | 0.912 | 0.935 | 1.38E-19 |
| Gm28661 | 6.78E-24 | 0.858483 | 0.854 | 0.692 | 1.6E-19 |
| Rpsa | 7.01E-24 | 1.879884 | 0.789 | 0.837 | 1.65E-19 |
| Kcnq1ot1 | 7.78E-24 | -0.68791 | 0.189 | 0.467 | 1.84E-19 |
| Sat1 | 9.89E-24 | 1.399859 | 0.85 | 0.908 | 2.33E-19 |
| Gm2225 | 1.06E-23 | 0.778937 | 0.341 | 0.065 | 2.5E-19 |
| Rcan1 | 1.1E-23 | -1.04951 | 0.081 | 0.296 | 2.59E-19 |
| Fbxl3 | 1.11E-23 | -0.71689 | 0.157 | 0.426 | 2.61E-19 |
| Ywhae | 1.2E-23 | 2.143139 | 0.788 | 0.84 | 2.82E-19 |
| Rpl3-ps1 | 1.29E-23 | 1.651672 | 0.748 | 0.612 | 3.04E-19 |
| Zfp638 | 1.31E-23 | -0.30588 | 0.18 | 0.462 | 3.09E-19 |
| Serinc3 | 2.03E-23 | -0.33419 | 0.99 | 1 | 4.79E-19 |
| Tmem86a | 2.4E-23 | 2.125881 | 0.733 | 0.749 | 5.67E-19 |
| Gstm1 | 2.7E-23 | 1.747264 | 0.093 | 0.331 | 6.36E-19 |
| Rps13-ps2 | 3.79E-23 | 1.594639 | 0.552 | 0.296 | 8.93E-19 |
| Gm10443 | 4.13E-23 | 0.657536 | 0.424 | 0.13 | 9.74E-19 |
| Arl6ip1 | 4.48E-23 | 1.575046 | 0.803 | 0.802 | 1.06E-18 |

| | | | | | |
|---|---|---|---|---|---|
| Cfh | 4.99E-23 | -0.58207 | 0.686 | 0.959 | 1.18E-18 |
| Gm37494 | 5.35E-23 | -0.5191 | 0.101 | 0.328 | 1.26E-18 |
| Rps10-ps2 | 6.98E-23 | 2.359751 | 0.598 | 0.408 | 1.65E-18 |
| Rpl11 | 8.04E-23 | 1.814278 | 0.739 | 0.692 | 1.9E-18 |
| Ppp1r21 | 8.84E-23 | -0.26513 | 0.202 | 0.494 | 2.09E-18 |
| Mbp | 1.17E-22 | 0.681173 | 0.096 | 0.331 | 2.77E-18 |
| Rpl19 | 1.22E-22 | 1.684111 | 0.797 | 0.787 | 2.88E-18 |
| Gm44861 | 1.35E-22 | -1.09015 | 0.054 | 0.234 | 3.18E-18 |
| Tubb5 | 1.62E-22 | 1.798756 | 0.81 | 0.817 | 3.81E-18 |
| Caprin1 | 1.7E-22 | -0.58761 | 0.237 | 0.541 | 4.01E-18 |
| Rps16 | 1.87E-22 | 1.484945 | 0.796 | 0.76 | 4.41E-18 |
| Gm12537 | 1.89E-22 | 0.966718 | 0.379 | 0.109 | 4.45E-18 |
| Ac165079.: | 2E-22 | 0.850443 | 0.308 | 0.05 | 4.73E-18 |
| Egr1 | 2.23E-22 | -2.65285 | 0.091 | 0.302 | 5.26E-18 |
| Efhd2 | 2.57E-22 | 1.9236 | 0.838 | 0.902 | 6.06E-18 |
| Otulinl | 3.15E-22 | 1.125709 | 0.907 | 0.979 | 7.42E-18 |
| Tlr13 | 4.52E-22 | -0.31777 | 0.231 | 0.524 | 1.07E-17 |
| Gm13456 | 5.33E-22 | 1.178235 | 0.951 | 0.967 | 1.26E-17 |
| Gm44705 | 5.64E-22 | 0.735068 | 0.275 | 0.03 | 1.33E-17 |
| Cd300c2 | 5.99E-22 | 1.468513 | 0.817 | 0.908 | 1.41E-17 |
| Pycard | 6.61E-22 | 1.411097 | 0.833 | 0.846 | 1.56E-17 |
| Rps5 | 7.43E-22 | 1.814556 | 0.767 | 0.79 | 1.75E-17 |
| Rpl28 | 7.67E-22 | 1.995416 | 0.747 | 0.725 | 1.81E-17 |
| Adap2 | 7.68E-22 | 1.402501 | 0.914 | 0.994 | 1.81E-17 |
| mt-Nd4 | 8.19E-22 | 1.203822 | 0.918 | 0.953 | 1.93E-17 |
| Gm5905 | 1.04E-21 | 1.909683 | 0.651 | 0.444 | 2.45E-17 |
| Gm46411 | 1.17E-21 | -1.12038 | 0.046 | 0.21 | 2.77E-17 |
| Rps13 | 1.68E-21 | 1.49884 | 0.79 | 0.746 | 3.97E-17 |
| Rpl10-ps3 | 1.73E-21 | 1.769331 | 0.665 | 0.435 | 4.07E-17 |
| Zfp36 | 2.03E-21 | 0.719487 | 0.355 | 0.766 | 4.79E-17 |
| Rpl7 | 2.24E-21 | 1.556623 | 0.8 | 0.82 | 5.28E-17 |
| Ckb | 2.7E-21 | 1.054995 | 0.881 | 0.967 | 6.37E-17 |
| 0610039K1 | 3.1E-21 | -0.37869 | 0.054 | 0.231 | 7.32E-17 |
| Gm6472 | 3.25E-21 | 1.380737 | 0.451 | 0.198 | 7.66E-17 |
| Ddx3x | 3.32E-21 | -0.36022 | 0.319 | 0.612 | 7.82E-17 |
| Setd5 | 3.34E-21 | -0.41731 | 0.278 | 0.574 | 7.87E-17 |
| Acly | 3.75E-21 | -0.81466 | 0.167 | 0.417 | 8.84E-17 |
| Limd2 | 4.1E-21 | 1.958029 | 0.777 | 0.831 | 9.68E-17 |
| Ubqln1 | 5.66E-21 | -0.38461 | 0.197 | 0.462 | 1.34E-16 |
| Jade2 | 5.91E-21 | 0.456367 | 0.153 | 0.411 | 1.39E-16 |
| Trip12 | 7.17E-21 | -0.7107 | 0.176 | 0.432 | 1.69E-16 |
| Upk1b | 7.18E-21 | 2.240917 | 0.574 | 0.367 | 1.69E-16 |
| Crk | 7.93E-21 | -0.66869 | 0.214 | 0.482 | 1.87E-16 |
| Gab2 | 8.99E-21 | -0.40049 | 0.114 | 0.34 | 2.12E-16 |
| Camk1 | 9.13E-21 | 1.430564 | 0.782 | 0.867 | 2.15E-16 |
| Malat1 | 1.12E-20 | 0.53155 | 0.981 | 1 | 2.64E-16 |
| Ppib | 1.15E-20 | 1.419518 | 0.771 | 0.84 | 2.71E-16 |

| | | | | | |
|---|---|---|---|---|---|
| Top2b | 1.4E-20 | -0.29253 | 0.12 | 0.349 | 3.29E-16 |
| Rpl18-ps1 | 1.54E-20 | 0.507263 | 0.294 | 0.047 | 3.62E-16 |
| Usp24 | 1.55E-20 | -0.36531 | 0.185 | 0.447 | 3.65E-16 |
| Rps19-ps6 | 1.69E-20 | 1.754181 | 0.598 | 0.385 | 3.98E-16 |
| Gm7336 | 1.81E-20 | 1.466289 | 0.434 | 0.186 | 4.27E-16 |
| Pik3cb | 2.41E-20 | -0.50721 | 0.046 | 0.207 | 5.69E-16 |
| Fcgr2b | 2.5E-20 | 1.432728 | 0.806 | 0.864 | 5.89E-16 |
| Actb | 2.58E-20 | 0.683778 | 1 | 1 | 6.09E-16 |
| Dpp8 | 3.52E-20 | -0.33049 | 0.119 | 0.346 | 8.3E-16 |
| Rnf169 | 3.75E-20 | 0.311948 | 0.195 | 0.488 | 8.84E-16 |
| Gm3511 | 4.05E-20 | 1.055675 | 0.544 | 0.293 | 9.56E-16 |
| Zfp871 | 4.39E-20 | -0.5609 | 0.139 | 0.37 | 1.04E-15 |
| Gm8797 | 5.02E-20 | 1.113804 | 0.862 | 0.92 | 1.18E-15 |
| Ftl2-ps | 6.67E-20 | 1.524958 | 0.88 | 0.959 | 1.57E-15 |
| Dhrs3 | 7.37E-20 | 1.825642 | 0.828 | 0.926 | 1.74E-15 |
| Ifngr1 | 7.51E-20 | 0.790295 | 0.959 | 0.994 | 1.77E-15 |
| Gm6170 | 7.93E-20 | 0.676858 | 0.415 | 0.157 | 1.87E-15 |
| Rpl23a | 8.02E-20 | 1.58335 | 0.755 | 0.775 | 1.89E-15 |
| Eif1 | 8.87E-20 | 1.346621 | 0.883 | 0.938 | 2.09E-15 |
| Gm10288 | 9.78E-20 | 1.479361 | 0.438 | 0.189 | 2.31E-15 |
| Rbm39 | 1.46E-19 | 1.365798 | 0.882 | 0.95 | 3.44E-15 |
| Aff4 | 1.48E-19 | -1.03061 | 0.164 | 0.396 | 3.5E-15 |
| Selenop | 2.02E-19 | 0.806563 | 0.985 | 1 | 4.76E-15 |
| Sh3bgrl3 | 2.07E-19 | 1.293199 | 0.806 | 0.888 | 4.88E-15 |
| Gm8960 | 2.19E-19 | 0.906989 | 0.308 | 0.068 | 5.17E-15 |
| Rpl17-ps8 | 3.07E-19 | 0.908983 | 0.228 | 0.015 | 7.23E-15 |
| Sp3 | 3.55E-19 | -0.50434 | 0.178 | 0.42 | 8.38E-15 |
| Gm10240 | 5.32E-19 | 1.426953 | 0.445 | 0.216 | 1.26E-14 |
| Gm8430 | 5.78E-19 | 1.605817 | 0.536 | 0.314 | 1.36E-14 |
| Capza2 | 5.99E-19 | 1.392487 | 0.866 | 0.95 | 1.41E-14 |
| Xpo1 | 6.14E-19 | -0.42547 | 0.072 | 0.251 | 1.45E-14 |
| Gm46367 | 1.14E-18 | -0.47286 | 0.095 | 0.296 | 2.7E-14 |
| Spen | 1.27E-18 | -0.29436 | 0.114 | 0.32 | 3E-14 |
| Abl2 | 1.42E-18 | -0.66409 | 0.124 | 0.331 | 3.36E-14 |
| Chka | 1.44E-18 | -0.35011 | 0.143 | 0.367 | 3.4E-14 |
| Rgs2 | 1.63E-18 | 1.467402 | 0.825 | 0.885 | 3.85E-14 |
| Gtf2i | 1.64E-18 | 0.663091 | 0.317 | 0.663 | 3.86E-14 |
| Arpc3 | 1.68E-18 | 1.944483 | 0.68 | 0.737 | 3.96E-14 |
| Mtmr6 | 1.75E-18 | -0.43557 | 0.184 | 0.42 | 4.12E-14 |
| Rb1cc1 | 1.77E-18 | -0.43267 | 0.183 | 0.429 | 4.18E-14 |
| Akr1a1 | 1.81E-18 | 1.677997 | 0.75 | 0.814 | 4.28E-14 |
| Usp34 | 1.94E-18 | -0.40906 | 0.104 | 0.299 | 4.57E-14 |
| Ecscr | 2.23E-18 | 1.757877 | 0.714 | 0.725 | 5.25E-14 |
| Atrx | 2.37E-18 | 0.561992 | 0.231 | 0.509 | 5.58E-14 |
| Scaf8 | 2.52E-18 | -0.36912 | 0.118 | 0.328 | 5.93E-14 |
| Rpl13a-ps1 | 2.73E-18 | 0.817531 | 0.241 | 0.03 | 6.43E-14 |
| Gm6988 | 2.86E-18 | 0.751339 | 0.307 | 0.077 | 6.76E-14 |

| | | | | | |
|---|---|---|---|---|---|
| Rps27a | 2.88E-18 | 1.451361 | 0.811 | 0.825 | 6.79E-14 |
| Erbin | 3.54E-18 | -0.33345 | 0.186 | 0.432 | 8.34E-14 |
| Erp29 | 3.7E-18 | 1.653904 | 0.781 | 0.893 | 8.72E-14 |
| Crybb1 | 4.12E-18 | 2.025167 | 0.737 | 0.87 | 9.71E-14 |
| Rpl19-ps11 | 4.71E-18 | 1.547906 | 0.575 | 0.385 | 1.11E-13 |
| Arpc2 | 4.82E-18 | 1.297734 | 0.81 | 0.905 | 1.14E-13 |
| Zdhhc5 | 5.71E-18 | -0.64196 | 0.088 | 0.275 | 1.35E-13 |
| Gm6180 | 6.09E-18 | 0.796812 | 0.374 | 0.133 | 1.44E-13 |
| Gm10020 | 7.6E-18 | 0.568302 | 0.25 | 0.038 | 1.79E-13 |
| Ltc4s | 8.02E-18 | 1.39288 | 0.839 | 0.941 | 1.89E-13 |
| Rreb1 | 9.95E-18 | 0.489076 | 0.228 | 0.494 | 2.35E-13 |
| Rpl27-ps3 | 1.04E-17 | 0.999472 | 0.393 | 0.16 | 2.45E-13 |
| Dock4 | 1.05E-17 | 0.362135 | 0.482 | 0.84 | 2.48E-13 |
| Lag3 | 1.07E-17 | 1.321729 | 0.81 | 0.84 | 2.53E-13 |
| Ints6l | 1.34E-17 | -0.40806 | 0.244 | 0.494 | 3.16E-13 |
| Actg1 | 1.47E-17 | 1.184817 | 0.842 | 0.905 | 3.47E-13 |
| Aff1 | 1.65E-17 | -0.43215 | 0.12 | 0.322 | 3.9E-13 |
| Ppp6r1 | 1.68E-17 | -0.5281 | 0.153 | 0.367 | 3.95E-13 |
| Akna | 1.75E-17 | 0.715554 | 0.297 | 0.612 | 4.12E-13 |
| Myo9a | 2.03E-17 | -0.4059 | 0.093 | 0.278 | 4.79E-13 |
| Tpt1-ps3 | 2.05E-17 | 1.245613 | 0.655 | 0.479 | 4.83E-13 |
| Rpl34 | 2.34E-17 | 1.327315 | 0.819 | 0.799 | 5.52E-13 |
| Cox4i1 | 2.4E-17 | 1.406072 | 0.798 | 0.846 | 5.67E-13 |
| Arfgef1 | 2.49E-17 | -0.52897 | 0.118 | 0.32 | 5.87E-13 |
| Anxa3 | 2.67E-17 | 1.206018 | 0.818 | 0.873 | 6.3E-13 |
| Dusp1 | 3.16E-17 | -1.58405 | 0.126 | 0.331 | 7.46E-13 |
| Smarca5 | 3.23E-17 | 0.357597 | 0.227 | 0.503 | 7.63E-13 |
| Nptn | 3.4E-17 | -0.38987 | 0.57 | 0.814 | 8.02E-13 |
| Zfp62 | 3.73E-17 | -0.33914 | 0.14 | 0.352 | 8.81E-13 |
| Ttyh3 | 4.24E-17 | 0.632184 | 0.202 | 0.467 | 1E-12 |
| Hist1h1c | 4.39E-17 | -0.29143 | 0.187 | 0.435 | 1.04E-12 |
| Gm28437 | 4.51E-17 | 0.746273 | 0.905 | 0.831 | 1.06E-12 |
| Rps6-ps4 | 4.94E-17 | 1.172278 | 0.518 | 0.284 | 1.16E-12 |
| Cyth4 | 5.68E-17 | 1.282793 | 0.88 | 0.964 | 1.34E-12 |
| Comt | 6.62E-17 | 1.366905 | 0.816 | 0.831 | 1.56E-12 |
| Rpl23 | 7.24E-17 | 1.511259 | 0.821 | 0.849 | 1.71E-12 |
| Usp9x | 7.77E-17 | -0.48491 | 0.16 | 0.382 | 1.83E-12 |
| Rps13-ps1 | 7.9E-17 | 0.560858 | 0.352 | 0.118 | 1.86E-12 |
| Tle4 | 8.15E-17 | 0.408172 | 0.116 | 0.317 | 1.92E-12 |
| Peak1 | 9.61E-17 | -0.30153 | 0.123 | 0.325 | 2.27E-12 |
| Gm4294 | 1.06E-16 | 0.809558 | 0.426 | 0.195 | 2.5E-12 |
| Chd2 | 1.12E-16 | 0.305489 | 0.171 | 0.414 | 2.63E-12 |
| Rpl14 | 1.24E-16 | 1.87959 | 0.763 | 0.849 | 2.92E-12 |
| Gm11518 | 1.26E-16 | 2.002746 | 0.446 | 0.243 | 2.98E-12 |
| Zfp445 | 1.4E-16 | -0.29894 | 0.181 | 0.408 | 3.29E-12 |
| Gm6394 | 1.44E-16 | 1.091227 | 0.474 | 0.246 | 3.41E-12 |
| A630001G2 | 1.48E-16 | 0.267742 | 0.267 | 0.547 | 3.49E-12 |

| | | | | | |
|---|---|---|---|---|---|
| Rgmb | 1.71E-16 | 0.265483 | 0.331 | 0.615 | 4.03E-12 |
| Vcpip1 | 1.75E-16 | -0.72735 | 0.141 | 0.346 | 4.13E-12 |
| Gm10051 | 1.86E-16 | 1.329301 | 0.276 | 0.071 | 4.38E-12 |
| P3h2 | 1.87E-16 | -0.82392 | 0.176 | 0.391 | 4.42E-12 |
| Rps12 | 2.01E-16 | 1.434388 | 0.802 | 0.834 | 4.75E-12 |
| Pde3b | 2.01E-16 | -0.37155 | 0.712 | 0.95 | 4.75E-12 |
| Rpl27a | 2.07E-16 | 1.707928 | 0.736 | 0.763 | 4.89E-12 |
| Casd1 | 2.26E-16 | 0.474911 | 0.133 | 0.346 | 5.33E-12 |
| Gabarap | 2.47E-16 | 1.620817 | 0.734 | 0.799 | 5.82E-12 |
| 9930021J0 | 2.66E-16 | -0.66158 | 0.106 | 0.29 | 6.28E-12 |
| Gm2245 | 2.97E-16 | -0.44332 | 0.072 | 0.231 | 7.01E-12 |
| Spred1 | 3.24E-16 | -0.31793 | 0.159 | 0.376 | 7.63E-12 |
| Kif3b | 3.35E-16 | -0.65961 | 0.063 | 0.216 | 7.89E-12 |
| Nrp2 | 3.66E-16 | -0.37777 | 0.09 | 0.266 | 8.64E-12 |
| Cmtm7 | 5.15E-16 | 1.265788 | 0.799 | 0.879 | 1.21E-11 |
| Specc1 | 5.32E-16 | 0.275303 | 0.296 | 0.577 | 1.25E-11 |
| Fcrls | 5.58E-16 | 0.66655 | 0.971 | 1 | 1.32E-11 |
| Arhgap12 | 7.78E-16 | -0.4716 | 0.188 | 0.405 | 1.84E-11 |
| Rpl36-ps2 | 7.95E-16 | 1.404077 | 0.508 | 0.293 | 1.87E-11 |
| Snx13 | 8.61E-16 | -0.34584 | 0.276 | 0.538 | 2.03E-11 |
| Lgals9 | 9.55E-16 | 1.823786 | 0.71 | 0.772 | 2.25E-11 |
| Cd63 | 9.61E-16 | 1.960149 | 0.692 | 0.701 | 2.27E-11 |
| Tab2 | 1.01E-15 | 0.405097 | 0.352 | 0.639 | 2.39E-11 |
| Pik3cd | 1.03E-15 | 0.385911 | 0.279 | 0.562 | 2.43E-11 |
| Rpl26 | 1.17E-15 | 1.696402 | 0.748 | 0.751 | 2.77E-11 |
| Oaz1-ps | 1.36E-15 | 1.354875 | 0.698 | 0.586 | 3.2E-11 |
| Ftl1-ps2 | 1.56E-15 | 0.464605 | 0.308 | 0.098 | 3.68E-11 |
| Pten | 1.66E-15 | 0.25174 | 0.315 | 0.651 | 3.92E-11 |
| Rab3gap1 | 1.78E-15 | -0.28818 | 0.16 | 0.373 | 4.2E-11 |
| Arglu1 | 1.94E-15 | 1.254678 | 0.894 | 0.911 | 4.59E-11 |
| Ap1b1 | 2.18E-15 | 0.830168 | 0.334 | 0.666 | 5.13E-11 |
| Cggbp1 | 2.2E-15 | -0.29488 | 0.199 | 0.429 | 5.19E-11 |
| Spcs3 | 2.43E-15 | -0.34491 | 0.197 | 0.411 | 5.72E-11 |
| Cdc27 | 2.62E-15 | -0.27841 | 0.075 | 0.234 | 6.18E-11 |
| Matr3 | 3.15E-15 | -0.49225 | 0.265 | 0.5 | 7.42E-11 |
| Diaph2 | 3.18E-15 | -0.46135 | 0.173 | 0.382 | 7.49E-11 |
| Gm11539 | 3.29E-15 | 0.88314 | 0.325 | 0.112 | 7.77E-11 |
| Nipbl | 3.31E-15 | 0.374537 | 0.338 | 0.636 | 7.8E-11 |
| Rpl15 | 3.35E-15 | 1.345511 | 0.735 | 0.692 | 7.9E-11 |
| Rlf | 3.59E-15 | -0.39689 | 0.094 | 0.266 | 8.46E-11 |
| Bhlhe41 | 3.87E-15 | 1.842975 | 0.826 | 0.885 | 9.12E-11 |
| Hspa8 | 3.94E-15 | 0.976269 | 0.919 | 0.994 | 9.3E-11 |
| Fbxl14 | 3.96E-15 | 0.414251 | 0.114 | 0.296 | 9.35E-11 |
| Ppp1r15b | 4.12E-15 | -0.40015 | 0.11 | 0.29 | 9.73E-11 |
| Tbc1d14 | 4.13E-15 | 0.449816 | 0.206 | 0.444 | 9.74E-11 |
| Mctp1 | 4.85E-15 | -0.33627 | 0.076 | 0.234 | 1.14E-10 |
| Ash1l | 4.91E-15 | 0.327984 | 0.272 | 0.541 | 1.16E-10 |

| | | | | | |
|---|---|---|---|---|---|
| Srsf2 | 5E-15 | 1.585842 | 0.801 | 0.849 | 1.18E-10 |
| Sh2b3 | 5.16E-15 | 0.640348 | 0.298 | 0.571 | 1.22E-10 |
| Stt3b | 5.2E-15 | -0.27948 | 0.128 | 0.317 | 1.23E-10 |
| Epc1 | 5.87E-15 | 0.360569 | 0.165 | 0.379 | 1.38E-10 |
| H2-DMa | 6.15E-15 | 1.815009 | 0.674 | 0.683 | 1.45E-10 |
| Mbnl2 | 6.19E-15 | 0.328297 | 0.361 | 0.663 | 1.46E-10 |
| Tgfb1 | 6.36E-15 | 0.442602 | 0.208 | 0.453 | 1.5E-10 |
| Trpm2 | 7.61E-15 | 0.407773 | 0.13 | 0.322 | 1.79E-10 |
| Rpl7a-ps7 | 8.24E-15 | 0.876599 | 0.269 | 0.074 | 1.94E-10 |
| Rbm6 | 8.4E-15 | -0.29517 | 0.257 | 0.503 | 1.98E-10 |
| Atrn | 8.87E-15 | -0.86882 | 0.093 | 0.257 | 2.09E-10 |
| Gm15772 | 9.37E-15 | 1.376249 | 0.422 | 0.216 | 2.21E-10 |
| Gm43813 | 1.03E-14 | -0.26295 | 0.099 | 0.269 | 2.43E-10 |
| Mia2 | 1.16E-14 | 0.388719 | 0.27 | 0.553 | 2.73E-10 |
| Nufip2 | 1.2E-14 | -0.29496 | 0.176 | 0.376 | 2.83E-10 |
| Gm4202 | 1.29E-14 | 0.27198 | 0.377 | 0.651 | 3.04E-10 |
| C2cd3 | 1.31E-14 | -0.72052 | 0.094 | 0.257 | 3.1E-10 |
| Rpl37rt | 1.39E-14 | 1.174275 | 0.619 | 0.429 | 3.29E-10 |
| Gab1 | 1.4E-14 | 0.308685 | 0.073 | 0.225 | 3.31E-10 |
| Rps27 | 1.48E-14 | 1.001795 | 0.937 | 0.967 | 3.49E-10 |
| Fcgr3 | 1.56E-14 | 0.706402 | 0.933 | 1 | 3.67E-10 |
| Sipa1l1 | 1.65E-14 | 0.847091 | 0.126 | 0.317 | 3.89E-10 |
| Tmem131l | 1.82E-14 | -0.52566 | 0.086 | 0.246 | 4.3E-10 |
| Prrc2b | 1.89E-14 | 0.612542 | 0.139 | 0.337 | 4.46E-10 |
| Gng10 | 2.16E-14 | 1.233132 | 0.763 | 0.861 | 5.1E-10 |
| Rnf216 | 2.17E-14 | 0.375307 | 0.247 | 0.503 | 5.12E-10 |
| Parp8 | 2.25E-14 | -0.38754 | 0.101 | 0.269 | 5.3E-10 |
| Ubn1 | 2.26E-14 | 0.384349 | 0.201 | 0.435 | 5.34E-10 |
| Rabgap1 | 2.48E-14 | -0.26489 | 0.169 | 0.379 | 5.85E-10 |
| Gm38042 | 2.55E-14 | -0.4019 | 0.137 | 0.322 | 6.01E-10 |
| Rpl24 | 2.81E-14 | 1.955906 | 0.627 | 0.565 | 6.63E-10 |
| Gm5805 | 2.91E-14 | 0.885323 | 0.411 | 0.216 | 6.87E-10 |
| Huwe1 | 3.1E-14 | 0.365071 | 0.154 | 0.364 | 7.32E-10 |
| Gm40309 | 3.23E-14 | 0.499293 | 0.127 | 0.322 | 7.63E-10 |
| Rps17 | 3.46E-14 | 1.364982 | 0.759 | 0.769 | 8.16E-10 |
| Rpl37 | 3.46E-14 | 0.97006 | 0.878 | 0.899 | 8.16E-10 |
| Sgk1 | 3.46E-14 | -0.31263 | 0.518 | 0.828 | 8.17E-10 |
| Selenof | 3.69E-14 | 1.352497 | 0.685 | 0.713 | 8.72E-10 |
| Arhgef2 | 3.7E-14 | 0.304788 | 0.307 | 0.556 | 8.74E-10 |
| Pcm1 | 3.77E-14 | -0.83823 | 0.118 | 0.293 | 8.9E-10 |
| Rpl21-ps15 | 3.82E-14 | 1.190289 | 0.309 | 0.109 | 9E-10 |
| B4galt1 | 3.86E-14 | 1.214324 | 0.874 | 0.92 | 9.11E-10 |
| Gm9703 | 4.03E-14 | 0.627955 | 0.289 | 0.095 | 9.51E-10 |
| Gna15 | 5.04E-14 | 1.725454 | 0.7 | 0.76 | 1.19E-09 |
| Dmtf1 | 5.14E-14 | -0.68542 | 0.161 | 0.355 | 1.21E-09 |
| Nf1 | 5.41E-14 | 0.357184 | 0.162 | 0.367 | 1.28E-09 |
| Shoc2 | 6.04E-14 | -0.61123 | 0.101 | 0.266 | 1.42E-09 |

| | | | | | |
|---|---|---|---|---|---|
| Mboat1 | 6.33E-14 | 0.371339 | 0.131 | 0.32 | 1.49E-09 |
| Zscan26 | 6.53E-14 | 1.794479 | 0.773 | 0.837 | 1.54E-09 |
| Stxbp5 | 7.23E-14 | -0.60447 | 0.113 | 0.284 | 1.7E-09 |
| Gm7392 | 7.77E-14 | 0.487306 | 0.205 | 0.033 | 1.83E-09 |
| BC005561 | 8.63E-14 | -0.26899 | 0.099 | 0.263 | 2.04E-09 |
| Zfhx3 | 9.02E-14 | 1.200583 | 0.911 | 0.991 | 2.13E-09 |
| Tnpo1 | 9.47E-14 | -0.39998 | 0.256 | 0.491 | 2.23E-09 |
| Neurl1a | 1.03E-13 | -0.34773 | 0.102 | 0.266 | 2.43E-09 |
| Rpl29 | 1.04E-13 | 1.833773 | 0.672 | 0.66 | 2.45E-09 |
| Gm10163 | 1.1E-13 | 0.921257 | 0.441 | 0.251 | 2.61E-09 |
| Gm10059 | 1.15E-13 | 0.548888 | 0.3 | 0.101 | 2.7E-09 |
| Rpl31 | 1.21E-13 | 1.451308 | 0.78 | 0.864 | 2.86E-09 |
| Cpeb4 | 1.25E-13 | -0.94182 | 0.076 | 0.222 | 2.94E-09 |
| Hnrnpul2 | 1.26E-13 | 0.436115 | 0.082 | 0.234 | 2.97E-09 |
| Shisa5 | 1.26E-13 | 2.02474 | 0.653 | 0.669 | 2.98E-09 |
| Tlr4 | 1.27E-13 | -0.26533 | 0.153 | 0.349 | 2.99E-09 |
| Mon2 | 1.32E-13 | 0.568311 | 0.209 | 0.447 | 3.12E-09 |
| Rpl36 | 1.34E-13 | 1.35004 | 0.805 | 0.834 | 3.17E-09 |
| Ptms | 1.36E-13 | 3.204507 | 0.642 | 0.737 | 3.2E-09 |
| Gm9616 | 1.38E-13 | 1.238281 | 0.469 | 0.284 | 3.26E-09 |
| Esco1 | 1.49E-13 | -0.36854 | 0.109 | 0.281 | 3.51E-09 |
| Man1c1 | 1.55E-13 | 0.409613 | 0.398 | 0.686 | 3.65E-09 |
| Il17ra | 1.61E-13 | 0.593126 | 0.389 | 0.666 | 3.81E-09 |
| Dock8 | 1.73E-13 | 0.38882 | 0.422 | 0.746 | 4.08E-09 |
| Wdr26 | 1.82E-13 | 0.722432 | 0.334 | 0.636 | 4.29E-09 |
| Tmem123 | 1.87E-13 | -0.9131 | 0.128 | 0.302 | 4.41E-09 |
| Kmt5b | 1.98E-13 | 0.469984 | 0.161 | 0.361 | 4.66E-09 |
| Emsy | 2E-13 | 0.289234 | 0.128 | 0.311 | 4.71E-09 |
| Ang | 2.05E-13 | 1.906409 | 0.769 | 0.87 | 4.84E-09 |
| Clint1 | 2.25E-13 | 0.499506 | 0.295 | 0.571 | 5.31E-09 |
| Gm9794 | 2.25E-13 | 1.26777 | 0.592 | 0.45 | 5.31E-09 |
| March6 | 2.35E-13 | -0.37421 | 0.107 | 0.272 | 5.54E-09 |
| Rnf19b | 2.37E-13 | 0.503961 | 0.167 | 0.382 | 5.58E-09 |
| Far1 | 2.53E-13 | -0.46461 | 0.178 | 0.376 | 5.96E-09 |
| Insr | 2.67E-13 | -0.52058 | 0.132 | 0.308 | 6.29E-09 |
| Akap13 | 2.71E-13 | 0.604171 | 0.472 | 0.787 | 6.39E-09 |
| Vps13b | 2.87E-13 | -0.63992 | 0.094 | 0.249 | 6.77E-09 |
| Hnrnpl | 2.98E-13 | 0.734001 | 0.37 | 0.607 | 7.04E-09 |
| Phkb | 3.14E-13 | 0.474616 | 0.093 | 0.251 | 7.42E-09 |
| Mtus1 | 3.39E-13 | 0.377262 | 0.468 | 0.775 | 8E-09 |
| Cysltr1 | 3.49E-13 | -0.28817 | 0.296 | 0.538 | 8.23E-09 |
| Tmem135 | 4.29E-13 | -0.49047 | 0.29 | 0.512 | 1.01E-08 |
| Atf7 | 4.37E-13 | 0.393778 | 0.142 | 0.328 | 1.03E-08 |
| Mark3 | 4.81E-13 | 0.283606 | 0.223 | 0.441 | 1.14E-08 |
| Tm9sf2 | 4.83E-13 | -0.26059 | 0.682 | 0.914 | 1.14E-08 |
| Gm11361 | 4.94E-13 | 0.71593 | 0.445 | 0.254 | 1.17E-08 |
| Ccr5 | 5.34E-13 | -0.36226 | 0.666 | 0.855 | 1.26E-08 |

| | | | | | |
|---|---|---|---|---|---|
| Gm12166 | 5.48E-13 | 1.276885 | 0.808 | 0.828 | 1.29E-08 |
| Zzz3 | 6.43E-13 | -0.51553 | 0.094 | 0.246 | 1.52E-08 |
| Gm14586 | 6.57E-13 | 1.100311 | 0.429 | 0.246 | 1.55E-08 |
| Nipa2 | 6.91E-13 | 0.2729 | 0.179 | 0.382 | 1.63E-08 |
| Dcaf8 | 7.01E-13 | 0.319183 | 0.185 | 0.396 | 1.65E-08 |
| Chmp4b | 7.28E-13 | 0.261661 | 0.089 | 0.243 | 1.72E-08 |
| Arrb2 | 7.57E-13 | 2.52759 | 0.671 | 0.757 | 1.79E-08 |
| Rpl7a | 8.28E-13 | 1.831657 | 0.698 | 0.719 | 1.95E-08 |
| Il6ra | 8.36E-13 | 1.463548 | 0.847 | 0.97 | 1.97E-08 |
| Kdm6a | 8.57E-13 | -0.9576 | 0.072 | 0.21 | 2.02E-08 |
| Msl2 | 9.76E-13 | -0.77907 | 0.129 | 0.299 | 2.3E-08 |
| Slc12a6 | 1.03E-12 | 0.268567 | 0.265 | 0.503 | 2.42E-08 |
| Rpl8 | 1.03E-12 | 1.695293 | 0.717 | 0.754 | 2.44E-08 |
| Fmnl1 | 1.08E-12 | 0.957135 | 0.289 | 0.559 | 2.55E-08 |
| Gm10045 | 1.1E-12 | 1.067957 | 0.202 | 0.041 | 2.59E-08 |
| Tmem65 | 1.15E-12 | -0.46049 | 0.088 | 0.234 | 2.72E-08 |
| Gm16192 | 1.15E-12 | 0.498219 | 0.108 | 0.272 | 2.72E-08 |
| Arhgap45 | 1.19E-12 | 1.363178 | 0.847 | 0.935 | 2.8E-08 |
| Lair1 | 1.21E-12 | 0.932361 | 0.875 | 0.962 | 2.85E-08 |
| Sbno2 | 1.22E-12 | 0.498521 | 0.223 | 0.435 | 2.87E-08 |
| Asxl1 | 1.23E-12 | 0.345645 | 0.113 | 0.275 | 2.91E-08 |
| Galnt11 | 1.27E-12 | 0.376475 | 0.13 | 0.308 | 3.01E-08 |
| Gm13680 | 1.3E-12 | 1.302016 | 0.382 | 0.195 | 3.07E-08 |
| Prkacb | 1.55E-12 | -0.31227 | 0.113 | 0.272 | 3.65E-08 |
| Strn3 | 1.72E-12 | 0.311608 | 0.258 | 0.497 | 4.05E-08 |
| Psmb8 | 1.78E-12 | 2.028744 | 0.594 | 0.574 | 4.21E-08 |
| Hsp90b1 | 1.83E-12 | 0.786031 | 0.932 | 0.985 | 4.32E-08 |
| Rpl34-ps1 | 1.85E-12 | 1.337303 | 0.454 | 0.293 | 4.36E-08 |
| Dennd6a | 1.88E-12 | -0.37857 | 0.097 | 0.251 | 4.43E-08 |
| Fam208a | 1.9E-12 | 0.253313 | 0.286 | 0.53 | 4.49E-08 |
| Mllt10 | 2.01E-12 | 0.744446 | 0.145 | 0.334 | 4.73E-08 |
| Sod1 | 2.09E-12 | 2.206669 | 0.651 | 0.609 | 4.92E-08 |
| Atp2a2 | 2.25E-12 | 0.692322 | 0.25 | 0.488 | 5.32E-08 |
| Ep300 | 2.27E-12 | 0.359233 | 0.347 | 0.601 | 5.36E-08 |
| Rabac1 | 2.29E-12 | 1.481738 | 0.723 | 0.822 | 5.41E-08 |
| Pmepa1 | 2.44E-12 | 1.935033 | 0.806 | 0.84 | 5.76E-08 |
| Twf2 | 2.53E-12 | 1.697309 | 0.627 | 0.648 | 5.98E-08 |
| Atp9b | 2.57E-12 | 0.866022 | 0.117 | 0.287 | 6.07E-08 |
| Atp8a1 | 2.89E-12 | 0.442407 | 0.353 | 0.621 | 6.82E-08 |
| Pdia4 | 2.94E-12 | 0.257369 | 0.33 | 0.621 | 6.94E-08 |
| Arhgdib | 3.11E-12 | 1.457229 | 0.73 | 0.843 | 7.34E-08 |
| Gprasp1 | 3.27E-12 | -0.44733 | 0.116 | 0.275 | 7.71E-08 |
| Gapvd1 | 3.36E-12 | -0.43125 | 0.27 | 0.491 | 7.92E-08 |
| Ncoa3 | 3.48E-12 | 1.672209 | 0.764 | 0.79 | 8.2E-08 |
| Atmin | 3.54E-12 | 0.402614 | 0.136 | 0.317 | 8.36E-08 |
| Lyst | 3.56E-12 | 0.70878 | 0.093 | 0.249 | 8.39E-08 |
| Smo | 3.75E-12 | 0.261705 | 0.098 | 0.251 | 8.84E-08 |

| | | | | | |
|---|---|---|---|---|---|
| Gpr34 | 3.8E-12 | 0.635866 | 0.981 | 0.997 | 8.97E-08 |
| Map4k4 | 3.9E-12 | 0.421658 | 0.366 | 0.627 | 9.2E-08 |
| Tcn2 | 3.96E-12 | 1.014849 | 0.82 | 0.929 | 9.33E-08 |
| Ankrd44 | 4E-12 | 0.497033 | 0.373 | 0.672 | 9.43E-08 |
| Rhoa | 4.17E-12 | 1.144893 | 0.841 | 0.899 | 9.83E-08 |
| Gm10231 | 4.24E-12 | 1.138176 | 0.246 | 0.08 | 1E-07 |
| Rtn4 | 4.26E-12 | 0.409593 | 0.44 | 0.716 | 1E-07 |
| Sgk3 | 4.67E-12 | -0.40978 | 0.142 | 0.317 | 1.1E-07 |
| Shprh | 4.69E-12 | -0.62838 | 0.088 | 0.231 | 1.11E-07 |
| Ripk1 | 4.8E-12 | -0.31792 | 0.155 | 0.328 | 1.13E-07 |
| Nktr | 4.84E-12 | 0.33451 | 0.244 | 0.476 | 1.14E-07 |
| Rps27a-ps2 | 5.6E-12 | 1.168289 | 0.334 | 0.154 | 1.32E-07 |
| Gm34280 | 5.95E-12 | 0.286235 | 0.152 | 0.334 | 1.4E-07 |
| Golgb1 | 5.97E-12 | -0.26425 | 0.113 | 0.269 | 1.41E-07 |
| Atp6v1g1 | 5.99E-12 | 1.525943 | 0.653 | 0.642 | 1.41E-07 |
| Prex1 | 6.07E-12 | 0.572765 | 0.401 | 0.686 | 1.43E-07 |
| R3hdm1 | 6.2E-12 | 0.299243 | 0.232 | 0.447 | 1.46E-07 |
| Fus | 6.32E-12 | 0.753178 | 0.417 | 0.722 | 1.49E-07 |
| Ubtf | 6.5E-12 | 0.420169 | 0.186 | 0.399 | 1.53E-07 |
| Usp2 | 6.92E-12 | 1.917687 | 0.68 | 0.74 | 1.63E-07 |
| Itpripl1 | 6.98E-12 | 1.922356 | 0.725 | 0.814 | 1.65E-07 |
| Chpt1 | 7.44E-12 | -0.60883 | 0.096 | 0.24 | 1.76E-07 |
| B430010I2 | 7.54E-12 | 0.52205 | 0.135 | 0.314 | 1.78E-07 |
| Sort1 | 7.95E-12 | 0.324851 | 0.387 | 0.66 | 1.88E-07 |
| Bri3 | 8.8E-12 | 2.601869 | 0.599 | 0.63 | 2.08E-07 |
| Numa1 | 8.99E-12 | 0.293202 | 0.287 | 0.524 | 2.12E-07 |
| Rps15a-ps6 | 9.28E-12 | 0.717335 | 0.265 | 0.095 | 2.19E-07 |
| Xpo7 | 9.49E-12 | 0.324546 | 0.128 | 0.293 | 2.24E-07 |
| Canx | 9.5E-12 | 0.87891 | 0.928 | 0.982 | 2.24E-07 |
| Tnks | 9.59E-12 | 0.546036 | 0.094 | 0.243 | 2.26E-07 |
| Xpo4 | 1.16E-11 | -0.34124 | 0.072 | 0.201 | 2.74E-07 |
| Ccdc93 | 1.21E-11 | 0.413689 | 0.224 | 0.441 | 2.85E-07 |
| Gm17241 | 1.21E-11 | 1.077016 | 0.336 | 0.16 | 2.87E-07 |
| Zfp106 | 1.28E-11 | 0.47373 | 0.237 | 0.459 | 3.01E-07 |
| Rbm3 | 1.29E-11 | 2.414062 | 0.358 | 0.198 | 3.03E-07 |
| Zfand5 | 1.37E-11 | 0.4734 | 0.317 | 0.601 | 3.23E-07 |
| Chd7 | 1.39E-11 | 0.747288 | 0.313 | 0.556 | 3.27E-07 |
| Pald1 | 1.4E-11 | 0.457963 | 0.396 | 0.654 | 3.29E-07 |
| Rack1 | 1.42E-11 | 2.01591 | 0.711 | 0.796 | 3.36E-07 |
| Secisbp2 | 1.66E-11 | 0.477324 | 0.097 | 0.246 | 3.91E-07 |
| Rpl9-ps6 | 1.83E-11 | 1.819286 | 0.636 | 0.574 | 4.33E-07 |
| Acap2 | 2.1E-11 | 0.531065 | 0.377 | 0.669 | 4.96E-07 |
| Clock | 2.13E-11 | -0.27151 | 0.117 | 0.266 | 5.03E-07 |
| Rbm26 | 2.23E-11 | 1.645229 | 0.723 | 0.802 | 5.26E-07 |
| Serinc1 | 2.32E-11 | -0.27127 | 0.603 | 0.814 | 5.48E-07 |
| Ppfia1 | 2.44E-11 | 0.282245 | 0.072 | 0.201 | 5.76E-07 |
| Rpl28-ps1 | 2.53E-11 | 1.409577 | 0.414 | 0.269 | 5.96E-07 |

| | | | | | |
|---|---|---|---|---|---|
| Tmem245 | 2.65E-11 | 0.577127 | 0.211 | 0.417 | 6.26E-07 |
| Atp2c1 | 2.73E-11 | 0.34084 | 0.334 | 0.595 | 6.45E-07 |
| Pld1 | 2.89E-11 | 0.290571 | 0.348 | 0.583 | 6.82E-07 |
| Btbd1 | 3E-11 | 0.337368 | 0.171 | 0.358 | 7.08E-07 |
| Gm15928 | 3.07E-11 | 0.492149 | 0.231 | 0.464 | 7.24E-07 |
| Asap1 | 3.13E-11 | 0.360905 | 0.334 | 0.609 | 7.38E-07 |
| Nfat5 | 3.19E-11 | 0.268473 | 0.202 | 0.393 | 7.52E-07 |
| Skil | 3.29E-11 | 1.266292 | 0.787 | 0.814 | 7.76E-07 |
| Cd63-ps | 3.29E-11 | 1.523205 | 0.492 | 0.349 | 7.77E-07 |
| Stat3 | 3.32E-11 | 0.321829 | 0.298 | 0.541 | 7.84E-07 |
| Tgfa | 3.47E-11 | 0.647094 | 0.199 | 0.402 | 8.19E-07 |
| Stau1 | 3.54E-11 | 0.543933 | 0.239 | 0.47 | 8.35E-07 |
| 4932438A1 | 3.75E-11 | -0.39182 | 0.13 | 0.29 | 8.83E-07 |
| Ube2h | 3.81E-11 | 0.450965 | 0.312 | 0.553 | 9E-07 |
| Pik3r5 | 3.97E-11 | 0.439322 | 0.22 | 0.411 | 9.36E-07 |
| Rps2-ps10 | 4.33E-11 | 0.917326 | 0.283 | 0.115 | 1.02E-06 |
| Ttc17 | 4.49E-11 | -0.2568 | 0.075 | 0.204 | 1.06E-06 |
| Zfp36l2 | 4.49E-11 | 0.823013 | 0.351 | 0.645 | 1.06E-06 |
| Trim44 | 4.66E-11 | 0.509504 | 0.094 | 0.234 | 1.1E-06 |
| Ccdc186 | 5.13E-11 | 0.422659 | 0.09 | 0.234 | 1.21E-06 |
| Slc35a3 | 5.23E-11 | -0.36294 | 0.138 | 0.299 | 1.23E-06 |
| Senp6 | 5.37E-11 | 0.883215 | 0.213 | 0.435 | 1.27E-06 |
| Hspa13 | 5.42E-11 | -0.37584 | 0.115 | 0.266 | 1.28E-06 |
| Pcmtd2 | 5.64E-11 | -0.63816 | 0.098 | 0.24 | 1.33E-06 |
| Dhx34 | 5.85E-11 | 0.317508 | 0.081 | 0.216 | 1.38E-06 |
| Rpl17-ps5 | 6.1E-11 | 0.748534 | 0.262 | 0.101 | 1.44E-06 |
| Foxj3 | 6.92E-11 | -0.48666 | 0.077 | 0.204 | 1.63E-06 |
| Plcg1 | 7.05E-11 | 0.329162 | 0.126 | 0.284 | 1.66E-06 |
| Ankrd11 | 7.16E-11 | 0.741367 | 0.42 | 0.728 | 1.69E-06 |
| Vwa5a | 7.52E-11 | -0.31008 | 0.151 | 0.311 | 1.77E-06 |
| Sp1 | 7.54E-11 | 0.478172 | 0.135 | 0.299 | 1.78E-06 |
| Mfsd14b | 7.57E-11 | -0.33762 | 0.154 | 0.317 | 1.79E-06 |
| Rpl15-ps3 | 7.76E-11 | 1.29542 | 0.273 | 0.112 | 1.83E-06 |
| Baz2a | 7.77E-11 | -0.45026 | 0.075 | 0.201 | 1.83E-06 |
| Rock2 | 7.98E-11 | 0.315171 | 0.359 | 0.627 | 1.88E-06 |
| Zfp518a | 8.01E-11 | -0.38154 | 0.104 | 0.249 | 1.89E-06 |
| Smarca2 | 8.16E-11 | 0.820423 | 0.252 | 0.464 | 1.93E-06 |
| Cds2 | 8.4E-11 | 0.626747 | 0.211 | 0.411 | 1.98E-06 |
| Spag9 | 8.5E-11 | 0.332081 | 0.261 | 0.485 | 2.01E-06 |
| Thrap3 | 8.86E-11 | 0.350549 | 0.334 | 0.574 | 2.09E-06 |
| Snhg20 | 9.57E-11 | 2.27785 | 0.32 | 0.16 | 2.26E-06 |
| Dnm1l | 9.79E-11 | -0.2581 | 0.081 | 0.21 | 2.31E-06 |
| Dapp1 | 1.08E-10 | 0.272656 | 0.362 | 0.63 | 2.54E-06 |
| Fam117b | 1.09E-10 | 0.461579 | 0.2 | 0.399 | 2.57E-06 |
| Gm5138 | 1.1E-10 | 1.889956 | 0.399 | 0.251 | 2.6E-06 |
| Mag | 1.11E-10 | 0.992054 | 0.111 | 0.272 | 2.61E-06 |
| Ankib1 | 1.25E-10 | -0.25706 | 0.09 | 0.225 | 2.96E-06 |

| | | | | | |
|---|---|---|---|---|---|
| Tnrc18 | 1.33E-10 | 0.531102 | 0.115 | 0.269 | 3.13E-06 |
| Hspa5 | 1.33E-10 | 0.862325 | 0.862 | 0.97 | 3.15E-06 |
| Commd8 | 1.34E-10 | 1.559775 | 0.691 | 0.749 | 3.17E-06 |
| Tmem104 | 1.37E-10 | 0.664191 | 0.248 | 0.473 | 3.23E-06 |
| Kmt2a | 1.37E-10 | 0.439032 | 0.24 | 0.447 | 3.24E-06 |
| Scd2 | 1.41E-10 | 0.85286 | 0.089 | 0.231 | 3.33E-06 |
| Dock2 | 1.45E-10 | 0.539196 | 0.321 | 0.571 | 3.43E-06 |
| Gm13408 | 1.55E-10 | 1.754951 | 0.244 | 0.089 | 3.65E-06 |
| Adipor2 | 1.56E-10 | 0.314777 | 0.127 | 0.284 | 3.68E-06 |
| Ncoa4 | 1.57E-10 | 0.308188 | 0.184 | 0.358 | 3.69E-06 |
| Reep5 | 1.59E-10 | 1.65534 | 0.69 | 0.754 | 3.76E-06 |
| Gm6136 | 1.6E-10 | 1.595165 | 0.525 | 0.393 | 3.78E-06 |
| Hipk2 | 1.66E-10 | 0.425635 | 0.11 | 0.257 | 3.93E-06 |
| Gm9625 | 1.76E-10 | 0.950754 | 0.457 | 0.293 | 4.14E-06 |
| Arf1 | 1.77E-10 | 1.209018 | 0.764 | 0.882 | 4.17E-06 |
| Sec14l1 | 1.78E-10 | 0.73416 | 0.347 | 0.586 | 4.19E-06 |
| Gm8226 | 1.8E-10 | 0.744765 | 0.206 | 0.059 | 4.24E-06 |
| Pomk | 1.82E-10 | -0.40488 | 0.081 | 0.21 | 4.29E-06 |
| Anp32a | 1.84E-10 | 0.622653 | 0.326 | 0.601 | 4.35E-06 |
| Dag1 | 1.89E-10 | -0.34859 | 0.104 | 0.24 | 4.45E-06 |
| Endod1 | 1.95E-10 | 0.302353 | 0.166 | 0.346 | 4.6E-06 |
| Tbc1d5 | 2E-10 | 0.683406 | 0.198 | 0.399 | 4.72E-06 |
| Ssr4 | 2.09E-10 | 1.492976 | 0.652 | 0.666 | 4.92E-06 |
| Adam17 | 2.13E-10 | 0.580761 | 0.345 | 0.592 | 5.03E-06 |
| Sall3 | 2.23E-10 | 0.459649 | 0.394 | 0.639 | 5.25E-06 |
| Cd14 | 2.33E-10 | 1.08462 | 0.764 | 0.873 | 5.5E-06 |
| Rhog | 2.39E-10 | 1.269703 | 0.744 | 0.864 | 5.63E-06 |
| Rab8b | 2.4E-10 | 0.516638 | 0.282 | 0.521 | 5.67E-06 |
| Mapk9 | 2.41E-10 | -0.32645 | 0.088 | 0.219 | 5.69E-06 |
| 6820431F2 | 2.5E-10 | 0.39816 | 0.167 | 0.346 | 5.91E-06 |
| Slc16a7 | 2.51E-10 | -0.34703 | 0.09 | 0.222 | 5.93E-06 |
| Pptc7 | 2.61E-10 | 0.790188 | 0.143 | 0.311 | 6.16E-06 |
| Zranb1 | 2.69E-10 | 0.479687 | 0.189 | 0.373 | 6.35E-06 |
| Gxylt1 | 2.82E-10 | 0.251733 | 0.14 | 0.302 | 6.66E-06 |
| Rsrc1 | 3.26E-10 | 0.252485 | 0.089 | 0.222 | 7.68E-06 |
| Olfr620 | 3.39E-10 | 0.414024 | 0.248 | 0.095 | 7.99E-06 |
| Snn | 3.46E-10 | 0.824477 | 0.241 | 0.453 | 8.16E-06 |
| Ube2q1 | 3.61E-10 | 0.797393 | 0.158 | 0.325 | 8.51E-06 |
| H2-T23 | 3.69E-10 | 1.627306 | 0.69 | 0.707 | 8.72E-06 |
| Ttc7 | 3.96E-10 | 0.994664 | 0.17 | 0.349 | 9.34E-06 |
| Arhgap39 | 3.96E-10 | 0.682047 | 0.335 | 0.583 | 9.35E-06 |
| Fam13b | 3.98E-10 | 0.417866 | 0.099 | 0.243 | 9.4E-06 |
| Nfx1 | 4.02E-10 | 0.542632 | 0.149 | 0.317 | 9.49E-06 |
| Herc1 | 4.06E-10 | 0.397399 | 0.244 | 0.441 | 9.58E-06 |
| Baz1b | 4.21E-10 | 0.261682 | 0.159 | 0.328 | 9.92E-06 |
| Gm11263 | 4.23E-10 | 0.47439 | 0.254 | 0.098 | 9.97E-06 |
| Vps39 | 4.26E-10 | 0.284486 | 0.131 | 0.281 | 1E-05 |

| | | | | | |
|---|---|---|---|---|---|
| Tmem64 | 4.35E-10 | -0.39988 | 0.152 | 0.311 | 1.03E-05 |
| Sept2 | 4.54E-10 | 1.507671 | 0.753 | 0.772 | 1.07E-05 |
| Rpsa-ps10 | 4.65E-10 | 1.470257 | 0.544 | 0.429 | 1.1E-05 |
| Abl1 | 4.65E-10 | 0.895244 | 0.227 | 0.426 | 1.1E-05 |
| Gtpbp2 | 4.72E-10 | 0.915736 | 0.169 | 0.343 | 1.11E-05 |
| Usp25 | 5.11E-10 | -0.34248 | 0.146 | 0.299 | 1.21E-05 |
| Saraf | 5.12E-10 | 0.921542 | 0.773 | 0.873 | 1.21E-05 |
| Lnpep | 5.17E-10 | 0.306868 | 0.433 | 0.686 | 1.22E-05 |
| Cox8a | 5.25E-10 | 1.422456 | 0.633 | 0.607 | 1.24E-05 |
| Pdpk1 | 5.3E-10 | 0.559271 | 0.097 | 0.237 | 1.25E-05 |
| Rpl30-ps10 | 5.32E-10 | 0.7389 | 0.25 | 0.095 | 1.26E-05 |
| Arsb | 5.4E-10 | 0.297898 | 0.381 | 0.615 | 1.27E-05 |
| Abca3 | 5.67E-10 | -0.29906 | 0.093 | 0.222 | 1.34E-05 |
| Mtr | 5.87E-10 | 0.269172 | 0.128 | 0.281 | 1.39E-05 |
| Skp1a | 5.93E-10 | 0.470678 | 0.285 | 0.518 | 1.4E-05 |
| Gmfb | 6E-10 | -0.32286 | 0.153 | 0.308 | 1.41E-05 |
| Rbm28 | 6.11E-10 | 1.050996 | 0.141 | 0.305 | 1.44E-05 |
| Ppp1cb | 6.3E-10 | 0.384488 | 0.124 | 0.266 | 1.49E-05 |
| 2510039O1 | 6.66E-10 | 0.464738 | 0.227 | 0.435 | 1.57E-05 |
| Lsm14a | 6.7E-10 | -0.36484 | 0.128 | 0.275 | 1.58E-05 |
| Calm1 | 6.71E-10 | 1.336215 | 0.742 | 0.873 | 1.58E-05 |
| Pcnt | 6.73E-10 | 0.434139 | 0.082 | 0.207 | 1.59E-05 |
| Hsp90ab1 | 7E-10 | 1.081522 | 0.813 | 0.962 | 1.65E-05 |
| Dicer1 | 7.26E-10 | 0.358193 | 0.108 | 0.249 | 1.71E-05 |
| Sla | 7.42E-10 | 0.345489 | 0.43 | 0.707 | 1.75E-05 |
| Fip1l1 | 7.45E-10 | 0.743217 | 0.14 | 0.305 | 1.76E-05 |
| Gm8129 | 7.51E-10 | 0.573913 | 0.237 | 0.092 | 1.77E-05 |
| Ero1lb | 7.52E-10 | -0.2985 | 0.094 | 0.225 | 1.77E-05 |
| Khdrbs1 | 7.73E-10 | 0.385982 | 0.249 | 0.459 | 1.82E-05 |
| Rps8 | 7.81E-10 | 1.385053 | 0.811 | 0.87 | 1.84E-05 |
| Ythdf3 | 8.58E-10 | -0.78436 | 0.101 | 0.231 | 2.02E-05 |
| Gm5835 | 8.73E-10 | 1.294243 | 0.357 | 0.201 | 2.06E-05 |
| Phykpl | 9.47E-10 | 0.570232 | 0.111 | 0.246 | 2.23E-05 |
| Gna12 | 9.84E-10 | 0.296386 | 0.458 | 0.749 | 2.32E-05 |
| Gm9385 | 1.02E-09 | 1.047399 | 0.313 | 0.16 | 2.41E-05 |
| Zfp451 | 1.05E-09 | 0.287299 | 0.119 | 0.26 | 2.47E-05 |
| Rps28 | 1.05E-09 | 0.778343 | 0.893 | 0.908 | 2.48E-05 |
| Gpx1 | 1.05E-09 | 1.564926 | 0.717 | 0.775 | 2.48E-05 |
| Rpl14-ps1 | 1.07E-09 | 1.232659 | 0.568 | 0.473 | 2.53E-05 |
| Actr2 | 1.14E-09 | 1.282983 | 0.89 | 0.964 | 2.68E-05 |
| Cse1l | 1.18E-09 | -0.55243 | 0.12 | 0.257 | 2.78E-05 |
| Riok3 | 1.19E-09 | 0.394317 | 0.104 | 0.243 | 2.82E-05 |
| Arih1 | 1.26E-09 | 0.416134 | 0.107 | 0.246 | 2.98E-05 |
| Efr3a | 1.28E-09 | 0.333912 | 0.142 | 0.299 | 3.02E-05 |
| Tmpo | 1.29E-09 | 0.560624 | 0.152 | 0.317 | 3.04E-05 |
| Chfr | 1.44E-09 | 0.471122 | 0.114 | 0.257 | 3.39E-05 |
| Tns3 | 1.59E-09 | 0.833893 | 0.229 | 0.432 | 3.76E-05 |

| | | | | | |
|---|---|---|---|---|---|
| Gcn1l1 | 1.6E-09 | 0.407569 | 0.34 | 0.574 | 3.79E-05 |
| Mkln1 | 1.64E-09 | 0.312638 | 0.338 | 0.574 | 3.86E-05 |
| Mtmr4 | 1.67E-09 | 0.425121 | 0.09 | 0.216 | 3.94E-05 |
| Rpl13-ps6 | 1.69E-09 | 1.451753 | 0.434 | 0.284 | 4E-05 |
| mt-Nd2 | 1.71E-09 | 1.232961 | 0.868 | 0.923 | 4.02E-05 |
| Cop1 | 1.71E-09 | -0.33968 | 0.121 | 0.257 | 4.03E-05 |
| Ccnt2 | 1.73E-09 | -0.30258 | 0.176 | 0.334 | 4.09E-05 |
| Mbtps1 | 1.82E-09 | 0.516718 | 0.126 | 0.269 | 4.29E-05 |
| Tut4 | 1.84E-09 | 0.695049 | 0.182 | 0.355 | 4.34E-05 |
| Ociad1 | 1.84E-09 | 1.539396 | 0.669 | 0.772 | 4.35E-05 |
| Mvb12b | 1.87E-09 | 0.745748 | 0.205 | 0.391 | 4.41E-05 |
| Pank2 | 1.89E-09 | -0.62183 | 0.127 | 0.263 | 4.46E-05 |
| Adgrg1 | 1.9E-09 | 1.089933 | 0.872 | 0.959 | 4.49E-05 |
| Slc12a2 | 2.04E-09 | -0.33624 | 0.195 | 0.358 | 4.8E-05 |
| Gm11560 | 2.14E-09 | 2.086433 | 0.231 | 0.098 | 5.06E-05 |
| Lamp1 | 2.15E-09 | 0.94984 | 0.874 | 0.991 | 5.06E-05 |
| Prkd3 | 2.27E-09 | 0.288636 | 0.139 | 0.29 | 5.36E-05 |
| Setdb1 | 2.32E-09 | 0.435676 | 0.126 | 0.269 | 5.46E-05 |
| Gm12191 | 2.53E-09 | 0.73778 | 0.295 | 0.139 | 5.96E-05 |
| Pkd2 | 2.53E-09 | 0.621993 | 0.081 | 0.201 | 5.96E-05 |
| Hmgb1 | 2.55E-09 | 1.366431 | 0.755 | 0.864 | 6.01E-05 |
| Wnk1 | 2.89E-09 | 0.302377 | 0.69 | 0.956 | 6.81E-05 |
| Ubxn7 | 2.98E-09 | -0.51368 | 0.094 | 0.216 | 7.02E-05 |
| E230001N( | 3E-09 | 0.897702 | 0.099 | 0.234 | 7.07E-05 |
| Taok2 | 3.04E-09 | 0.66226 | 0.189 | 0.355 | 7.16E-05 |
| Mafb | 3.08E-09 | 1.222805 | 0.844 | 0.97 | 7.26E-05 |
| Arid5b | 3.12E-09 | 0.719593 | 0.192 | 0.367 | 7.37E-05 |
| Vps4b | 3.19E-09 | 0.360677 | 0.289 | 0.509 | 7.53E-05 |
| Nr2c2 | 3.24E-09 | 0.568429 | 0.287 | 0.494 | 7.64E-05 |
| Elmsan1 | 3.27E-09 | 0.453491 | 0.101 | 0.231 | 7.72E-05 |
| Soga1 | 3.42E-09 | 0.619233 | 0.434 | 0.719 | 8.06E-05 |
| Etv6 | 3.47E-09 | 0.50245 | 0.136 | 0.287 | 8.2E-05 |
| Nisch | 3.52E-09 | 0.913364 | 0.399 | 0.657 | 8.3E-05 |
| Atp7a | 3.59E-09 | 0.433282 | 0.128 | 0.272 | 8.46E-05 |
| Fam193a | 3.6E-09 | 0.57646 | 0.124 | 0.266 | 8.49E-05 |
| Slc36a1 | 3.63E-09 | 0.343929 | 0.15 | 0.302 | 8.57E-05 |
| Nek7 | 3.65E-09 | 0.258612 | 0.299 | 0.503 | 8.62E-05 |
| Ckap5 | 3.72E-09 | 0.68376 | 0.103 | 0.24 | 8.78E-05 |
| Secisbp2l | 3.73E-09 | 0.319857 | 0.285 | 0.497 | 8.79E-05 |
| Notch2 | 3.82E-09 | 1.353879 | 0.716 | 0.719 | 9.01E-05 |
| Heatr5a | 3.87E-09 | 0.335735 | 0.31 | 0.527 | 9.13E-05 |
| Kansl1 | 3.92E-09 | 0.369392 | 0.367 | 0.604 | 9.24E-05 |
| Fndc3a | 4.12E-09 | 0.406853 | 0.192 | 0.373 | 9.72E-05 |
| Wdr81 | 4.26E-09 | 0.64243 | 0.258 | 0.459 | 0.000101 |
| Prpf8 | 4.31E-09 | 0.350547 | 0.203 | 0.376 | 0.000102 |
| Rps24-ps3 | 4.44E-09 | 0.606043 | 0.281 | 0.127 | 0.000105 |
| Itpripl2 | 4.53E-09 | -0.52801 | 0.108 | 0.237 | 0.000107 |

| | | | | | |
|---|---|---|---|---|---|
| Ptk2b | 4.55E-09 | -0.29508 | 0.226 | 0.388 | 0.000107 |
| Cmklr1 | 4.6E-09 | 0.760388 | 0.178 | 0.343 | 0.000108 |
| Ddx3y | 4.7E-09 | 0.342366 | 0.231 | 0.423 | 0.000111 |
| Gabbr1 | 4.77E-09 | 1.103471 | 0.106 | 0.243 | 0.000113 |
| Atp8a2 | 4.86E-09 | 0.524959 | 0.295 | 0.518 | 0.000115 |
| Cenpb | 5.09E-09 | 0.655749 | 0.101 | 0.231 | 0.00012 |
| Rps15a | 5.18E-09 | 1.539449 | 0.627 | 0.595 | 0.000122 |
| Cables1 | 5.37E-09 | 0.325376 | 0.235 | 0.432 | 0.000127 |
| Sipa1l2 | 5.39E-09 | 0.535026 | 0.218 | 0.411 | 0.000127 |
| Hcfc1 | 5.66E-09 | 0.707825 | 0.161 | 0.325 | 0.000134 |
| Gm13340 | 5.7E-09 | 0.629575 | 0.675 | 0.568 | 0.000135 |
| Gm37760 | 5.72E-09 | 0.412641 | 0.107 | 0.24 | 0.000135 |
| Nfatc1 | 5.87E-09 | 0.409609 | 0.326 | 0.538 | 0.000138 |
| Uba52-ps | 5.9E-09 | 0.371335 | 0.206 | 0.074 | 0.000139 |
| Helz | 5.91E-09 | 0.292354 | 0.202 | 0.376 | 0.000139 |
| Tnpo3 | 6.02E-09 | 0.332863 | 0.195 | 0.364 | 0.000142 |
| Frmd4a | 6.2E-09 | 0.277384 | 0.652 | 0.92 | 0.000146 |
| Ythdc1 | 6.58E-09 | 1.009328 | 0.245 | 0.464 | 0.000155 |
| Myo18a | 6.66E-09 | 0.669688 | 0.146 | 0.293 | 0.000157 |
| Eprs | 6.69E-09 | 0.668971 | 0.199 | 0.388 | 0.000158 |
| Cd53 | 7E-09 | 0.514531 | 0.964 | 0.997 | 0.000165 |
| Eif5a | 7.06E-09 | 1.781949 | 0.62 | 0.68 | 0.000167 |
| Gm6063 | 7.09E-09 | 0.380648 | 0.216 | 0.08 | 0.000167 |
| Ccdc50 | 7.1E-09 | 0.647059 | 0.36 | 0.595 | 0.000167 |
| Anapc1 | 7.17E-09 | -0.29247 | 0.096 | 0.219 | 0.000169 |
| Ubr4 | 7.21E-09 | 0.606127 | 0.151 | 0.308 | 0.00017 |
| Tbc1d23 | 7.25E-09 | -0.28224 | 0.193 | 0.352 | 0.000171 |
| Atf7ip | 7.84E-09 | 0.709388 | 0.38 | 0.607 | 0.000185 |
| Pom121 | 7.85E-09 | 1.061288 | 0.176 | 0.355 | 0.000185 |
| Usp15 | 7.94E-09 | 0.2581 | 0.189 | 0.355 | 0.000187 |
| Acin1 | 8.04E-09 | 0.684768 | 0.282 | 0.5 | 0.00019 |
| Rap2a | 8.28E-09 | 0.638855 | 0.232 | 0.444 | 0.000195 |
| Zcchc8 | 8.39E-09 | 0.683739 | 0.16 | 0.317 | 0.000198 |
| Gspt1 | 8.54E-09 | 0.631831 | 0.306 | 0.544 | 0.000201 |
| Coro1b | 8.66E-09 | 1.594752 | 0.568 | 0.562 | 0.000204 |
| Creb1 | 9.13E-09 | 0.404904 | 0.157 | 0.308 | 0.000215 |
| Tmed9 | 9.17E-09 | 1.904838 | 0.625 | 0.737 | 0.000216 |
| Sgpp1 | 9.42E-09 | -0.77072 | 0.106 | 0.225 | 0.000222 |
| Nrf1 | 9.67E-09 | 0.335558 | 0.113 | 0.246 | 0.000228 |
| Serf2 | 9.93E-09 | 1.657832 | 0.716 | 0.746 | 0.000234 |
| Plekho1 | 9.93E-09 | 2.162372 | 0.666 | 0.772 | 0.000234 |
| Gm4617 | 9.99E-09 | 0.862058 | 0.299 | 0.16 | 0.000236 |
| Irf2bp2 | 1.11E-08 | 1.202205 | 0.295 | 0.506 | 0.000263 |
| Rassf4 | 1.13E-08 | 0.593439 | 0.253 | 0.467 | 0.000267 |
| Gm8172 | 1.18E-08 | 2.009907 | 0.349 | 0.228 | 0.000278 |
| Pias2 | 1.26E-08 | -0.32925 | 0.096 | 0.216 | 0.000297 |
| Lbr | 1.29E-08 | 0.439329 | 0.137 | 0.281 | 0.000303 |

| | | | | | |
|---|---|---|---|---|---|
| Zfc3h1 | 1.33E-08 | 0.910945 | 0.147 | 0.302 | 0.000313 |
| Ptpra | 1.35E-08 | 0.298129 | 0.531 | 0.799 | 0.000317 |
| Pacsin2 | 1.39E-08 | 0.266037 | 0.386 | 0.604 | 0.000329 |
| Mapkap1 | 1.41E-08 | 1.067734 | 0.13 | 0.278 | 0.000332 |
| Pfkfb3 | 1.44E-08 | 0.723956 | 0.157 | 0.314 | 0.00034 |
| Ddb1 | 1.45E-08 | 1.025849 | 0.171 | 0.328 | 0.000342 |
| Ppp2r2a | 1.46E-08 | 0.683924 | 0.243 | 0.45 | 0.000345 |
| Rhoq | 1.48E-08 | 0.608196 | 0.185 | 0.358 | 0.000349 |
| Dennd1a | 1.49E-08 | 0.601359 | 0.088 | 0.204 | 0.000351 |
| Ppp1ca | 1.51E-08 | 1.475229 | 0.626 | 0.716 | 0.000355 |
| Pnpla7 | 1.52E-08 | 0.488978 | 0.464 | 0.719 | 0.000359 |
| Klk8 | 1.59E-08 | 1.856572 | 0.442 | 0.349 | 0.000376 |
| Mapk14 | 1.61E-08 | 0.326482 | 0.312 | 0.509 | 0.00038 |
| Nmt1 | 1.66E-08 | 1.030431 | 0.462 | 0.698 | 0.000391 |
| Gm8995 | 1.7E-08 | 0.416569 | 0.154 | 0.305 | 0.000402 |
| Bbx | 1.71E-08 | 0.540019 | 0.099 | 0.225 | 0.000404 |
| Csnk1g2 | 1.73E-08 | 0.57232 | 0.148 | 0.296 | 0.000408 |
| Orai1 | 1.73E-08 | 2.058518 | 0.612 | 0.713 | 0.000408 |
| Pcnx3 | 1.79E-08 | 1.103716 | 0.192 | 0.364 | 0.000422 |
| Prrc2c | 1.92E-08 | 0.596925 | 0.524 | 0.766 | 0.000454 |
| Hp1bp3 | 1.94E-08 | 0.576448 | 0.412 | 0.686 | 0.000457 |
| Gm5560 | 1.95E-08 | 0.498582 | 0.322 | 0.172 | 0.00046 |
| Gbp7 | 1.95E-08 | 0.631821 | 0.218 | 0.402 | 0.000461 |
| Large1 | 1.97E-08 | 0.648588 | 0.096 | 0.219 | 0.000466 |
| Rps3a3 | 2.04E-08 | 0.561459 | 0.376 | 0.234 | 0.000481 |
| Vps36 | 2.07E-08 | 0.437676 | 0.129 | 0.272 | 0.000488 |
| Evi2a | 2.11E-08 | 1.26978 | 0.661 | 0.71 | 0.000497 |
| Rpl10a-ps1 | 2.13E-08 | 1.498721 | 0.533 | 0.447 | 0.000503 |
| Map3k4 | 2.14E-08 | 0.952259 | 0.088 | 0.204 | 0.000504 |
| Rhobtb1 | 2.17E-08 | 0.673069 | 0.211 | 0.393 | 0.000511 |
| Gm29667 | 2.18E-08 | 1.140904 | 0.241 | 0.112 | 0.000515 |
| Ncoa6 | 2.19E-08 | -0.40272 | 0.103 | 0.222 | 0.000517 |
| Syncrip | 2.21E-08 | 0.368342 | 0.151 | 0.299 | 0.000521 |
| Gm10132 | 2.25E-08 | 1.857206 | 0.536 | 0.494 | 0.000531 |
| Camk1d | 2.25E-08 | 0.508701 | 0.218 | 0.391 | 0.000531 |
| Gm9493 | 2.3E-08 | 0.596369 | 0.25 | 0.112 | 0.000542 |
| Otud5 | 2.34E-08 | 0.857541 | 0.19 | 0.358 | 0.000551 |
| Ptp4a1 | 2.35E-08 | -0.33303 | 0.101 | 0.216 | 0.000554 |
| Sec24c | 2.35E-08 | 0.585737 | 0.199 | 0.373 | 0.000554 |
| Nr6a1os | 2.38E-08 | 0.26055 | 0.1 | 0.222 | 0.000561 |
| Manba | 2.45E-08 | -0.41156 | 0.117 | 0.24 | 0.000578 |
| Slc3a2 | 2.45E-08 | 1.149051 | 0.718 | 0.811 | 0.000578 |
| Mrpl37 | 2.46E-08 | 0.326464 | 0.094 | 0.213 | 0.00058 |
| Ino80d | 2.53E-08 | 0.719678 | 0.128 | 0.263 | 0.000598 |
| Ago3 | 2.54E-08 | 0.298709 | 0.179 | 0.334 | 0.000599 |
| Mxi1 | 2.57E-08 | 0.2503 | 0.156 | 0.308 | 0.000605 |
| Tmem260 | 2.57E-08 | -0.39985 | 0.133 | 0.263 | 0.000606 |

| | | | | | |
|---|---|---|---|---|---|
| Sipa1 | 2.66E-08 | 0.295838 | 0.533 | 0.799 | 0.000628 |
| Ube4a | 2.82E-08 | 0.5028 | 0.145 | 0.29 | 0.000664 |
| Gmeb1 | 2.82E-08 | 0.286238 | 0.134 | 0.272 | 0.000665 |
| Ank2 | 2.89E-08 | 0.84611 | 0.178 | 0.334 | 0.000682 |
| Scamp5 | 3.07E-08 | 0.724894 | 0.438 | 0.695 | 0.000724 |
| Jarid2 | 3.12E-08 | 0.748528 | 0.098 | 0.222 | 0.000735 |
| Rnf10 | 3.17E-08 | 0.530678 | 0.199 | 0.373 | 0.000748 |
| Ppp4r1 | 3.2E-08 | 0.520586 | 0.152 | 0.296 | 0.000754 |
| Wdfy4 | 3.26E-08 | 0.637966 | 0.131 | 0.269 | 0.00077 |
| Akap10 | 3.34E-08 | 0.884025 | 0.12 | 0.251 | 0.000787 |
| Cdk8 | 3.37E-08 | 0.75292 | 0.13 | 0.266 | 0.000795 |
| Abcd1 | 3.38E-08 | 0.595798 | 0.205 | 0.376 | 0.000798 |
| Smad3 | 3.39E-08 | 0.669957 | 0.258 | 0.453 | 0.000799 |
| Gm15501 | 3.41E-08 | 1.011586 | 0.43 | 0.281 | 0.000804 |
| Ctcf | 3.51E-08 | 0.267411 | 0.195 | 0.361 | 0.000828 |
| Tapbp | 3.53E-08 | 1.145339 | 0.773 | 0.82 | 0.000833 |
| Dnajb12 | 3.54E-08 | -0.2995 | 0.135 | 0.266 | 0.000835 |
| Pmp22 | 3.56E-08 | 1.146376 | 0.754 | 0.828 | 0.000841 |
| Rbm25 | 3.65E-08 | 0.705067 | 0.46 | 0.743 | 0.000862 |
| Adcy7 | 3.67E-08 | 0.625758 | 0.201 | 0.367 | 0.000867 |
| H2-DMb1 | 3.68E-08 | 1.937364 | 0.484 | 0.453 | 0.000868 |
| Tnrc6a | 3.7E-08 | 0.284901 | 0.266 | 0.435 | 0.000872 |
| Ahcyl1 | 3.76E-08 | -0.37881 | 0.093 | 0.204 | 0.000886 |
| Stmn1 | 3.76E-08 | 1.093019 | 0.088 | 0.207 | 0.000888 |
| Abhd15 | 3.78E-08 | 0.521147 | 0.432 | 0.71 | 0.000893 |
| Peli2 | 3.79E-08 | 0.608232 | 0.12 | 0.251 | 0.000894 |
| Tbc1d9b | 3.88E-08 | 0.279406 | 0.197 | 0.358 | 0.000915 |
| Slc39a10 | 4.02E-08 | 0.269714 | 0.147 | 0.29 | 0.000949 |
| Gm3791 | 4.06E-08 | -0.2795 | 0.173 | 0.322 | 0.000959 |
| Tmem106b | 4.08E-08 | 0.472312 | 0.182 | 0.337 | 0.000963 |
| Hook3 | 4.2E-08 | 0.66781 | 0.332 | 0.553 | 0.000992 |
| Dpp9 | 4.32E-08 | 0.461358 | 0.089 | 0.204 | 0.00102 |
| Kras | 4.36E-08 | 0.834169 | 0.187 | 0.358 | 0.001028 |
| Kpna1 | 4.48E-08 | 0.566191 | 0.145 | 0.287 | 0.001057 |
| Pgs1 | 4.5E-08 | 0.654964 | 0.201 | 0.367 | 0.001061 |
| Fam217b | 4.52E-08 | 0.525886 | 0.166 | 0.317 | 0.001066 |
| Hnrnpm | 4.77E-08 | 1.150578 | 0.354 | 0.604 | 0.001126 |
| Creg1 | 4.87E-08 | 1.244524 | 0.709 | 0.822 | 0.00115 |
| Zfp655 | 4.88E-08 | -0.41756 | 0.101 | 0.219 | 0.001152 |
| Bst2 | 4.9E-08 | 2.893208 | 0.435 | 0.388 | 0.001157 |
| Naip2 | 4.95E-08 | 0.344094 | 0.2 | 0.364 | 0.001168 |
| Senp2 | 5.02E-08 | 0.353016 | 0.277 | 0.462 | 0.001183 |
| Ice1 | 5.05E-08 | 0.456414 | 0.104 | 0.225 | 0.001191 |
| Prdx5 | 5.21E-08 | 2.210099 | 0.519 | 0.536 | 0.001228 |
| Rcsd1 | 5.27E-08 | 0.657759 | 0.41 | 0.683 | 0.001243 |
| Pfdn5 | 5.28E-08 | 1.468832 | 0.607 | 0.624 | 0.001246 |
| Ago1 | 5.33E-08 | 0.314282 | 0.18 | 0.328 | 0.001257 |

| | | | | | |
|---|---|---|---|---|---|
| Nsmaf | 5.41E-08 | 0.585255 | 0.172 | 0.328 | 0.001276 |
| Fcgr1 | 5.45E-08 | 1.331725 | 0.728 | 0.855 | 0.001286 |
| Fam221b | 5.48E-08 | 1.154734 | 0.398 | 0.275 | 0.001293 |
| Tmsb10 | 5.58E-08 | 2.783085 | 0.377 | 0.302 | 0.001315 |
| Csde1 | 5.62E-08 | 0.291066 | 0.338 | 0.541 | 0.001326 |
| Cdc73 | 5.63E-08 | -0.37956 | 0.093 | 0.207 | 0.001329 |
| Pak1 | 5.66E-08 | 0.497908 | 0.23 | 0.411 | 0.001334 |
| Susd6 | 5.85E-08 | 1.443457 | 0.129 | 0.263 | 0.001381 |
| Lgals3bp | 5.91E-08 | 2.45435 | 0.565 | 0.547 | 0.001393 |
| Rapgef1 | 5.95E-08 | 0.825583 | 0.241 | 0.426 | 0.001402 |
| Rptor | 6.23E-08 | 0.738942 | 0.368 | 0.609 | 0.001469 |
| Hivep3 | 6.31E-08 | 0.650911 | 0.328 | 0.55 | 0.001487 |
| Gm13493 | 6.45E-08 | 0.780471 | 0.695 | 0.618 | 0.00152 |
| Gm47283 | 6.46E-08 | 0.865373 | 0.16 | 0.314 | 0.001525 |
| Dirc2 | 6.47E-08 | 0.738601 | 0.218 | 0.405 | 0.001526 |
| Wac | 6.5E-08 | 0.536389 | 0.317 | 0.527 | 0.001534 |
| Gigyf2 | 6.56E-08 | 0.418504 | 0.166 | 0.317 | 0.001548 |
| Stard8 | 6.87E-08 | -0.64886 | 0.109 | 0.225 | 0.00162 |
| Rps26 | 6.95E-08 | 1.420467 | 0.675 | 0.675 | 0.00164 |
| Pds5a | 6.96E-08 | 0.610394 | 0.215 | 0.382 | 0.001641 |
| Abcg1 | 7.31E-08 | 1.846803 | 0.643 | 0.663 | 0.001725 |
| Pmm2 | 7.45E-08 | 1.047613 | 0.138 | 0.281 | 0.001757 |
| Zyg11b | 7.87E-08 | -0.36977 | 0.14 | 0.272 | 0.001856 |
| Rps15 | 7.91E-08 | 1.494802 | 0.711 | 0.772 | 0.001866 |
| Bicd2 | 7.98E-08 | 0.470294 | 0.205 | 0.373 | 0.001883 |
| Yes1 | 8.04E-08 | -0.25911 | 0.116 | 0.237 | 0.001897 |
| Ptprm | 8.1E-08 | 0.385601 | 0.101 | 0.219 | 0.00191 |
| Rai1 | 8.42E-08 | 0.847065 | 0.094 | 0.21 | 0.001986 |
| Samd8 | 8.75E-08 | 0.354572 | 0.294 | 0.482 | 0.002064 |
| Lmo2 | 8.78E-08 | 1.809351 | 0.607 | 0.633 | 0.002071 |
| Ccdc88a | 8.8E-08 | 0.502471 | 0.133 | 0.269 | 0.002075 |
| 5031439G( | 9.03E-08 | 0.600528 | 0.14 | 0.275 | 0.00213 |
| Spg20 | 9.18E-08 | 0.261868 | 0.142 | 0.275 | 0.002165 |
| Ehbp1l1 | 9.48E-08 | 0.668471 | 0.158 | 0.299 | 0.002237 |
| Dad1 | 1.01E-07 | 1.598033 | 0.615 | 0.701 | 0.002388 |
| Rps25-ps1 | 1.02E-07 | 0.492636 | 0.296 | 0.16 | 0.002416 |
| Baz2b | 1.05E-07 | 0.289371 | 0.156 | 0.296 | 0.002485 |
| Rps21 | 1.07E-07 | 1.071754 | 0.742 | 0.766 | 0.002515 |
| Grcc10 | 1.07E-07 | 1.802124 | 0.597 | 0.627 | 0.002526 |
| Vgll4 | 1.09E-07 | 0.851777 | 0.314 | 0.556 | 0.002576 |
| Pkn2 | 1.09E-07 | 0.473023 | 0.152 | 0.296 | 0.002581 |
| Uggt1 | 1.11E-07 | 0.928096 | 0.162 | 0.311 | 0.00262 |
| Elmo2 | 1.15E-07 | 0.308081 | 0.143 | 0.275 | 0.002721 |
| Ncoa2 | 1.17E-07 | 0.517497 | 0.112 | 0.231 | 0.002753 |
| Gm6210 | 1.18E-07 | 0.713148 | 0.412 | 0.29 | 0.002789 |
| Card6 | 1.19E-07 | 0.889196 | 0.14 | 0.278 | 0.002817 |
| Cd180 | 1.2E-07 | 0.591843 | 0.321 | 0.515 | 0.002837 |

| | | | | | |
|---|---|---|---|---|---|
| Stam2 | 1.21E-07 | -0.26923 | 0.098 | 0.21 | 0.002856 |
| Mga | 1.27E-07 | 0.297474 | 0.155 | 0.29 | 0.002992 |
| Gak | 1.28E-07 | 0.73343 | 0.298 | 0.506 | 0.003013 |
| Agpat5 | 1.29E-07 | 0.631074 | 0.107 | 0.228 | 0.003044 |
| Rbbp6 | 1.29E-07 | 0.325764 | 0.302 | 0.497 | 0.003049 |
| Gm11808 | 1.3E-07 | 1.30636 | 0.413 | 0.305 | 0.003057 |
| Irf5 | 1.31E-07 | 1.389571 | 0.68 | 0.746 | 0.003079 |
| Chn2 | 1.32E-07 | 0.991086 | 0.275 | 0.488 | 0.003111 |
| Sec11c | 1.36E-07 | 1.437473 | 0.628 | 0.689 | 0.003205 |
| Trim28 | 1.39E-07 | 0.70555 | 0.273 | 0.462 | 0.003289 |
| Foxj2 | 1.48E-07 | 0.607792 | 0.144 | 0.281 | 0.003482 |
| Slc23a2 | 1.48E-07 | 0.311112 | 0.146 | 0.284 | 0.00349 |
| Nup88 | 1.51E-07 | 0.675272 | 0.239 | 0.429 | 0.003553 |
| Dnajc5 | 1.52E-07 | 0.480399 | 0.297 | 0.488 | 0.003587 |
| Rnf141 | 1.56E-07 | -0.50586 | 0.101 | 0.21 | 0.003669 |
| Ehmt1 | 1.56E-07 | 0.952105 | 0.177 | 0.328 | 0.003684 |
| Gm7331 | 1.61E-07 | 1.268769 | 0.491 | 0.426 | 0.003799 |
| Maml2 | 1.63E-07 | 0.697419 | 0.268 | 0.459 | 0.003844 |
| Plxna4 | 1.64E-07 | 0.474603 | 0.154 | 0.284 | 0.003875 |
| Pkn1 | 1.68E-07 | 0.680429 | 0.341 | 0.574 | 0.003972 |
| Eno1 | 1.69E-07 | 1.442814 | 0.682 | 0.822 | 0.003983 |
| Snta1 | 1.71E-07 | 1.0039 | 0.251 | 0.459 | 0.004041 |
| Slc26a2 | 1.72E-07 | -0.31357 | 0.119 | 0.234 | 0.004058 |
| Klf3 | 1.74E-07 | 0.746716 | 0.49 | 0.766 | 0.004098 |
| Psip1 | 1.75E-07 | 0.733339 | 0.136 | 0.269 | 0.004117 |
| Rpl35a-ps2 | 1.75E-07 | 0.668127 | 0.206 | 0.089 | 0.004118 |
| Usp12 | 1.77E-07 | 0.620072 | 0.118 | 0.237 | 0.004173 |
| Usp4 | 1.82E-07 | 0.339194 | 0.273 | 0.441 | 0.004284 |
| Ptp4a2 | 1.85E-07 | 1.537307 | 0.74 | 0.855 | 0.004352 |
| Slc7a8 | 1.91E-07 | 0.477766 | 0.579 | 0.852 | 0.004506 |
| P2ry13 | 1.92E-07 | 0.570776 | 0.961 | 0.994 | 0.004533 |
| Gm40841 | 1.99E-07 | 0.538906 | 0.107 | 0.228 | 0.004683 |
| Nrp1 | 2.03E-07 | 0.379229 | 0.393 | 0.601 | 0.004789 |
| Usp40 | 2.19E-07 | 0.450778 | 0.146 | 0.278 | 0.005165 |
| Klhl5 | 2.21E-07 | 0.421004 | 0.107 | 0.222 | 0.005216 |
| Parp1 | 2.26E-07 | 1.076285 | 0.195 | 0.361 | 0.00533 |
| Ap3d1 | 2.26E-07 | 0.673341 | 0.203 | 0.367 | 0.005342 |
| Arhgap30 | 2.27E-07 | 0.696404 | 0.318 | 0.518 | 0.005353 |
| Cxxc5 | 2.28E-07 | 1.66487 | 0.674 | 0.793 | 0.005377 |
| Gm37465 | 2.29E-07 | -0.53799 | 0.114 | 0.234 | 0.005394 |
| Rnpepl1 | 2.35E-07 | 0.717697 | 0.219 | 0.385 | 0.005551 |
| Fndc3b | 2.36E-07 | 0.468452 | 0.101 | 0.213 | 0.005565 |
| Zfp710 | 2.36E-07 | 2.271731 | 0.679 | 0.825 | 0.005568 |
| Spcs2 | 2.37E-07 | 1.160132 | 0.682 | 0.743 | 0.005598 |
| Cic | 2.4E-07 | 1.21664 | 0.15 | 0.293 | 0.005655 |
| Ipo9 | 2.48E-07 | 0.424135 | 0.108 | 0.222 | 0.00584 |
| Dido1 | 2.48E-07 | 1.177149 | 0.198 | 0.361 | 0.005857 |

| | | | | | |
|---|---|---|---|---|---|
| Mfhas1 | 2.51E-07 | 0.665786 | 0.188 | 0.334 | 0.005931 |
| Sfswap | 2.52E-07 | 0.842694 | 0.14 | 0.272 | 0.005941 |
| Myl6 | 2.53E-07 | 1.602607 | 0.621 | 0.648 | 0.00598 |
| Gpr137b | 2.54E-07 | 0.647889 | 0.192 | 0.349 | 0.005984 |
| Ankhd1 | 2.55E-07 | 0.364975 | 0.177 | 0.32 | 0.006013 |
| Smarcd2 | 2.56E-07 | 0.505147 | 0.166 | 0.311 | 0.006042 |
| Ncor1 | 2.56E-07 | 0.728134 | 0.365 | 0.607 | 0.006044 |
| Jup | 2.6E-07 | 0.657501 | 0.245 | 0.42 | 0.006127 |
| Phc3 | 2.62E-07 | 0.573192 | 0.172 | 0.32 | 0.006188 |
| Slc15a3 | 2.67E-07 | 0.821581 | 0.295 | 0.491 | 0.006307 |
| Nfkbia | 2.69E-07 | 0.745819 | 0.436 | 0.716 | 0.006338 |
| Gdi2 | 2.73E-07 | 0.880733 | 0.865 | 0.947 | 0.006437 |
| Kcmf1 | 2.77E-07 | 0.594805 | 0.186 | 0.34 | 0.006538 |
| Sh2b1 | 2.79E-07 | 0.454911 | 0.107 | 0.222 | 0.006577 |
| Ighm | 2.87E-07 | 1.371987 | 0.666 | 0.74 | 0.006759 |
| Sgpl1 | 2.94E-07 | 0.574375 | 0.393 | 0.586 | 0.006928 |
| Dip2b | 2.94E-07 | 0.935506 | 0.518 | 0.808 | 0.006939 |
| Tspan4 | 2.97E-07 | 1.74345 | 0.487 | 0.453 | 0.006995 |
| Arid3b | 3.01E-07 | 0.594139 | 0.181 | 0.322 | 0.007096 |
| Tmem181b | 3.02E-07 | -0.32579 | 0.212 | 0.352 | 0.007118 |
| Atp6v1f | 3.03E-07 | 1.528241 | 0.577 | 0.586 | 0.007145 |
| Tubgcp5 | 3.04E-07 | 0.333688 | 0.423 | 0.642 | 0.007173 |
| Cdc42 | 3.05E-07 | 0.736419 | 0.863 | 0.97 | 0.007195 |
| Fbxw2 | 3.17E-07 | 0.856281 | 0.217 | 0.391 | 0.007475 |
| Trpc4ap | 3.33E-07 | 0.445213 | 0.38 | 0.598 | 0.007848 |
| Lrrc41 | 3.39E-07 | 0.343321 | 0.113 | 0.234 | 0.007992 |
| Zbtb18 | 3.48E-07 | 0.417334 | 0.191 | 0.343 | 0.008212 |
| Gm17147 | 3.58E-07 | 0.29269 | 0.108 | 0.222 | 0.008455 |
| Nuak1 | 3.59E-07 | 1.111182 | 0.319 | 0.538 | 0.008476 |
| Sec24b | 3.62E-07 | 0.260124 | 0.157 | 0.284 | 0.00853 |
| Smad4 | 3.65E-07 | 0.471377 | 0.126 | 0.251 | 0.008616 |
| Krt15 | 3.67E-07 | 2.07699 | 0.111 | 0.24 | 0.008658 |
| Il7r | 3.7E-07 | 0.527688 | 0.151 | 0.287 | 0.008721 |
| Hnrnpk | 3.7E-07 | 1.135838 | 0.795 | 0.899 | 0.008738 |
| Ddx23 | 3.82E-07 | 0.963606 | 0.123 | 0.251 | 0.009019 |
| Dstyk | 3.88E-07 | 0.411572 | 0.094 | 0.201 | 0.009164 |
| Appbp2 | 3.92E-07 | 0.491098 | 0.193 | 0.343 | 0.009255 |
| Ppp2r5c | 3.93E-07 | 0.771581 | 0.312 | 0.512 | 0.009271 |
| Ptpn6 | 3.95E-07 | 1.147074 | 0.699 | 0.802 | 0.009307 |
| Rad21 | 3.98E-07 | 0.398323 | 0.104 | 0.216 | 0.009382 |
| Zbtb1 | 4.07E-07 | 0.395284 | 0.174 | 0.317 | 0.009603 |
| Rab11fip5 | 4.1E-07 | 0.451217 | 0.143 | 0.269 | 0.009676 |
| Fbxl20 | 4.17E-07 | 0.642556 | 0.138 | 0.266 | 0.009832 |
| Ckap4 | 4.24E-07 | 0.987897 | 0.202 | 0.367 | 0.010001 |
| Foxn3 | 4.28E-07 | 1.297081 | 0.838 | 0.929 | 0.010103 |
| Ubfd1 | 4.32E-07 | 0.841204 | 0.214 | 0.37 | 0.01019 |
| Psenen | 4.33E-07 | 1.76027 | 0.558 | 0.58 | 0.010209 |

| | | | | | |
|---|---|---|---|---|---|
| Gpbp1 | 4.36E-07 | -0.46051 | 0.145 | 0.263 | 0.010276 |
| Dnajb4 | 4.36E-07 | 0.393429 | 0.113 | 0.231 | 0.010291 |
| Gm46224 | 4.49E-07 | 0.432704 | 0.12 | 0.243 | 0.0106 |
| Ip6k1 | 4.68E-07 | 0.398366 | 0.271 | 0.438 | 0.01103 |
| Fkbp15 | 4.72E-07 | 0.423574 | 0.375 | 0.592 | 0.011131 |
| Kdm2a | 4.74E-07 | 0.365911 | 0.252 | 0.417 | 0.011184 |
| Fgfr1op2 | 4.85E-07 | 0.565931 | 0.288 | 0.485 | 0.011444 |
| Oraov1 | 4.91E-07 | 0.930335 | 0.097 | 0.201 | 0.011578 |
| Arhgef40 | 5.01E-07 | 0.9606 | 0.405 | 0.648 | 0.011815 |
| Tal1 | 5.05E-07 | 0.521078 | 0.125 | 0.246 | 0.011911 |
| Gm44949 | 5.07E-07 | 0.366243 | 0.095 | 0.201 | 0.011964 |
| Tlr6 | 5.1E-07 | -0.34077 | 0.14 | 0.257 | 0.01204 |
| Fubp1 | 5.14E-07 | 0.596601 | 0.294 | 0.479 | 0.012118 |
| Abhd17c | 5.18E-07 | 0.568478 | 0.364 | 0.592 | 0.012225 |
| Kpna4 | 5.19E-07 | 0.934236 | 0.287 | 0.503 | 0.012253 |
| Dmxl1 | 5.26E-07 | 0.710859 | 0.164 | 0.308 | 0.012414 |
| Tln2 | 5.35E-07 | 0.47534 | 0.231 | 0.393 | 0.012627 |
| Map3k2 | 5.36E-07 | 0.860103 | 0.148 | 0.281 | 0.012641 |
| Gm6977 | 5.53E-07 | 0.870938 | 0.465 | 0.367 | 0.013034 |
| Arap3 | 5.58E-07 | 1.232045 | 0.137 | 0.269 | 0.01317 |
| Pwwp2a | 5.64E-07 | 0.553804 | 0.375 | 0.592 | 0.013293 |
| Plcb2 | 5.72E-07 | 0.279323 | 0.111 | 0.225 | 0.013492 |
| Zswim6 | 5.84E-07 | 0.38804 | 0.188 | 0.328 | 0.013785 |
| Ppm1b | 5.88E-07 | 0.416264 | 0.103 | 0.213 | 0.013859 |
| Vps18 | 5.92E-07 | 0.418551 | 0.211 | 0.364 | 0.01397 |
| Hmbox1 | 5.93E-07 | 1.290237 | 0.096 | 0.201 | 0.013997 |
| Nup153 | 5.94E-07 | 0.439768 | 0.105 | 0.216 | 0.014013 |
| Il6st | 6.16E-07 | 0.516177 | 0.395 | 0.609 | 0.014524 |
| P2ry6 | 6.55E-07 | 1.054958 | 0.776 | 0.893 | 0.015443 |
| Atg14 | 6.57E-07 | 0.639164 | 0.107 | 0.219 | 0.015493 |
| Med12l | 6.64E-07 | 0.340486 | 0.497 | 0.751 | 0.015669 |
| Mitf | 6.78E-07 | 0.798367 | 0.218 | 0.399 | 0.015986 |
| Gm42418 | 6.83E-07 | 2.777821 | 0.529 | 0.612 | 0.016102 |
| Plcg2 | 6.95E-07 | 0.66841 | 0.321 | 0.503 | 0.016393 |
| Tnrc6c | 7.02E-07 | 1.010168 | 0.267 | 0.45 | 0.016559 |
| Senp3 | 7.18E-07 | 0.507103 | 0.137 | 0.257 | 0.016935 |
| Paip1 | 7.33E-07 | 0.647268 | 0.236 | 0.396 | 0.017294 |
| H2afz | 7.67E-07 | 1.42525 | 0.548 | 0.53 | 0.018085 |
| Capns1 | 7.74E-07 | 0.855136 | 0.24 | 0.399 | 0.018267 |
| Tmem63a | 7.83E-07 | 0.394304 | 0.398 | 0.609 | 0.018475 |
| Mcur1 | 7.99E-07 | 0.416942 | 0.238 | 0.402 | 0.018848 |
| Arnt | 8.16E-07 | 0.348614 | 0.164 | 0.296 | 0.019237 |
| Gpr146 | 8.24E-07 | 0.514819 | 0.344 | 0.536 | 0.019435 |
| Camk2d | 8.36E-07 | 0.555576 | 0.33 | 0.524 | 0.019717 |
| Nck1 | 8.43E-07 | 0.579795 | 0.17 | 0.317 | 0.019889 |
| Golga4 | 8.5E-07 | 0.885959 | 0.121 | 0.237 | 0.020058 |
| Dync1h1 | 8.74E-07 | 0.917861 | 0.218 | 0.373 | 0.020607 |

| | | | | | |
|---|---|---|---|---|---|
| Gtf3c2 | 8.93E-07 | 0.273693 | 0.202 | 0.337 | 0.021073 |
| Smox | 8.94E-07 | 0.898437 | 0.285 | 0.491 | 0.021099 |
| Pou2f2 | 8.95E-07 | 2.294184 | 0.694 | 0.876 | 0.021122 |
| Zfp658 | 8.97E-07 | 0.254555 | 0.12 | 0.234 | 0.02117 |
| Atad2b | 9.05E-07 | 0.380837 | 0.132 | 0.251 | 0.021349 |
| Fam168b | 9.14E-07 | 0.787182 | 0.154 | 0.281 | 0.021571 |
| Wipf1 | 9.2E-07 | 0.282203 | 0.199 | 0.343 | 0.021693 |
| Nap1l1 | 9.22E-07 | 0.741016 | 0.112 | 0.228 | 0.021749 |
| Gnai3 | 9.26E-07 | -0.26403 | 0.166 | 0.287 | 0.021846 |
| Ncbp3 | 9.37E-07 | 0.710571 | 0.211 | 0.373 | 0.022113 |
| BC005537 | 9.43E-07 | 0.881008 | 0.248 | 0.411 | 0.022248 |
| Atp5b | 9.46E-07 | 1.003296 | 0.737 | 0.893 | 0.022322 |
| Frmd4b | 9.54E-07 | 0.814161 | 0.442 | 0.683 | 0.022497 |
| Zfp277 | 9.75E-07 | 0.436249 | 0.146 | 0.278 | 0.023011 |
| Gm5514 | 1.01E-06 | 1.085282 | 0.529 | 0.423 | 0.023802 |
| Usp19 | 1.06E-06 | 1.179918 | 0.095 | 0.201 | 0.024928 |
| Mef2d | 1.06E-06 | 0.794101 | 0.127 | 0.249 | 0.024973 |
| Elk3 | 1.07E-06 | 0.325502 | 0.317 | 0.488 | 0.025329 |
| Pxk | 1.08E-06 | 0.590949 | 0.205 | 0.358 | 0.025417 |
| Crebrf | 1.09E-06 | 0.747766 | 0.15 | 0.278 | 0.025614 |
| Gm10275 | 1.09E-06 | 1.24469 | 0.358 | 0.251 | 0.025642 |
| Zmiz1 | 1.1E-06 | 0.684677 | 0.497 | 0.74 | 0.025943 |
| Tgoln1 | 1.12E-06 | 0.563342 | 0.325 | 0.506 | 0.026513 |
| Ddx17 | 1.14E-06 | 0.759309 | 0.409 | 0.633 | 0.026858 |
| Slc35e1 | 1.15E-06 | 0.606002 | 0.107 | 0.216 | 0.02718 |
| Eya4 | 1.16E-06 | -0.49236 | 0.107 | 0.21 | 0.027312 |
| Selenok | 1.18E-06 | 1.328614 | 0.629 | 0.713 | 0.027799 |
| Gm26542 | 1.19E-06 | 0.889619 | 0.119 | 0.237 | 0.028165 |
| Gm45606 | 1.22E-06 | 1.033291 | 0.146 | 0.284 | 0.028824 |
| Ddost | 1.22E-06 | 1.218513 | 0.676 | 0.793 | 0.02886 |
| R3hdm2 | 1.24E-06 | 0.39815 | 0.114 | 0.225 | 0.02931 |
| Csf2rb2 | 1.27E-06 | 0.746651 | 0.143 | 0.266 | 0.029985 |
| Dgkd | 1.28E-06 | 0.796247 | 0.192 | 0.337 | 0.030123 |
| Gm7072 | 1.29E-06 | 0.283314 | 0.116 | 0.228 | 0.030345 |
| Il4ra | 1.31E-06 | 0.361121 | 0.375 | 0.571 | 0.030821 |
| Mfng | 1.31E-06 | 2.062185 | 0.426 | 0.349 | 0.030832 |
| Arf6 | 1.33E-06 | 0.688198 | 0.316 | 0.503 | 0.03148 |
| Ssr3 | 1.34E-06 | 0.478024 | 0.292 | 0.456 | 0.031607 |
| Etnk1 | 1.43E-06 | 0.425441 | 0.268 | 0.432 | 0.033651 |
| Lrrc14 | 1.47E-06 | 0.47732 | 0.114 | 0.225 | 0.034744 |
| Git2 | 1.48E-06 | 0.481489 | 0.638 | 0.905 | 0.034911 |
| Fads1 | 1.53E-06 | 0.443792 | 0.213 | 0.364 | 0.036043 |
| Atxn2 | 1.53E-06 | 0.78655 | 0.183 | 0.32 | 0.036129 |
| Arrb1 | 1.53E-06 | 1.278485 | 0.122 | 0.243 | 0.03617 |
| Supt16 | 1.55E-06 | 0.875728 | 0.11 | 0.225 | 0.036494 |
| Hectd4 | 1.56E-06 | 1.343402 | 0.12 | 0.237 | 0.036823 |
| Rbm5 | 1.64E-06 | 0.474875 | 0.347 | 0.544 | 0.038579 |

| | | | | | |
|---|---|---|---|---|---|
| Zc3h13 | 1.64E-06 | 0.6949 | 0.106 | 0.213 | 0.038668 |
| Gramd1a | 1.64E-06 | 0.580135 | 0.224 | 0.379 | 0.038713 |
| Kctd20 | 1.68E-06 | 0.597558 | 0.118 | 0.231 | 0.039621 |
| Naip5 | 1.69E-06 | 0.677995 | 0.119 | 0.231 | 0.039814 |
| Hsd17b11 | 1.72E-06 | 0.696084 | 0.195 | 0.331 | 0.040653 |
| Thoc2 | 1.73E-06 | 0.476066 | 0.211 | 0.355 | 0.040906 |
| Gatad2b | 1.76E-06 | 0.794538 | 0.268 | 0.441 | 0.04142 |
| Zswim8 | 1.78E-06 | 0.735601 | 0.138 | 0.26 | 0.042058 |
| Fam76b | 1.86E-06 | 0.374305 | 0.137 | 0.254 | 0.043867 |
| Slc30a5 | 1.88E-06 | -0.55968 | 0.141 | 0.251 | 0.044337 |
| Ptma | 1.92E-06 | 2.033252 | 0.647 | 0.784 | 0.045199 |
| Bag6 | 1.96E-06 | 1.678119 | 0.126 | 0.246 | 0.046333 |
| Hnrnpdl | 1.99E-06 | 0.30185 | 0.167 | 0.293 | 0.046961 |
| Aldoa | 2E-06 | 1.348043 | 0.702 | 0.825 | 0.047283 |
| Cul1 | 2.07E-06 | 0.333797 | 0.358 | 0.565 | 0.048748 |
| Cep63 | 2.07E-06 | 0.928367 | 0.108 | 0.219 | 0.048829 |
| Rps3a2 | 2.08E-06 | 1.00418 | 0.338 | 0.225 | 0.048952 |
| Tug1 | 2.08E-06 | 0.945149 | 0.227 | 0.388 | 0.048955 |
| Ndrg3 | 2.09E-06 | 0.665251 | 0.151 | 0.278 | 0.049248 |
| Rspry1 | 2.11E-06 | 0.801591 | 0.131 | 0.246 | 0.049723 |

### Supplementry Table 4. Aging DEGs in Striatum

| Gene | p_val | avg_log2FC | pct.1 | pct.2 | p_val_adj |
|---|---|---|---|---|---|
| Gm15564 | 6E-202 | -3.23836 | 0.027 | 0.811 | 1.4E-197 |
| C1qa | 2.2E-155 | 2.343129 | 1 | 1 | 5.2E-151 |
| C1qc | 4.5E-152 | 2.056197 | 1 | 1 | 1.1E-147 |
| Itm2b | 2.6E-145 | 1.645928 | 1 | 1 | 6.2E-141 |
| C1qb | 3.8E-143 | 1.649717 | 1 | 1 | 9E-139 |
| Cst3 | 2.4E-140 | 1.613518 | 1 | 1 | 5.6E-136 |
| Cd81 | 2.2E-139 | 1.960568 | 1 | 0.997 | 5.2E-135 |
| Tmsb4x | 1.9E-138 | 2.01925 | 1 | 1 | 4.4E-134 |
| B2m | 1.2E-134 | 2.22761 | 0.999 | 1 | 2.9E-130 |
| Fth1 | 1.1E-130 | 2.736824 | 0.995 | 0.993 | 2.5E-126 |
| Cd9 | 9.7E-121 | 2.494392 | 0.989 | 0.993 | 2.3E-116 |
| Plp1 | 1.7E-120 | -0.70059 | 0.014 | 0.518 | 3.9E-116 |
| Fcer1g | 1.6E-118 | 1.860022 | 0.998 | 0.997 | 3.8E-114 |
| Ctss | 2.9E-118 | 1.464294 | 1 | 1 | 6.7E-114 |
| Eef1a1 | 1.3E-116 | 2.013449 | 0.997 | 0.997 | 3E-112 |
| Gm24270 | 8.6E-116 | -2.61309 | 0.277 | 0.902 | 2E-111 |
| Cfl1 | 3.8E-113 | 3.14759 | 0.962 | 0.987 | 9E-109 |
| mt-Co1 | 4.5E-113 | 2.397621 | 0.992 | 0.997 | 1.1E-108 |
| Gcg | 4.5E-112 | -2.84839 | 0 | 0.427 | 1.1E-107 |
| Ctsz | 1.7E-110 | 1.974315 | 0.993 | 0.997 | 3.9E-106 |
| Ftl1 | 1.3E-109 | 2.338335 | 0.992 | 1 | 3E-105 |
| Cyba | 2.1E-107 | 2.310095 | 0.978 | 0.984 | 5E-103 |
| Sparc | 1.6E-104 | 1.64283 | 0.998 | 1 | 3.7E-100 |
| Ins2 | 8.1E-104 | -1.21769 | 0 | 0.397 | 1.9E-99 |
| Pfn1 | 1.2E-103 | 2.597359 | 0.964 | 0.977 | 2.9E-99 |
| Rpl18a | 1.5E-100 | 2.193783 | 0.987 | 0.987 | 3.57E-96 |
| Lars2 | 2.5E-100 | -3.10882 | 0.763 | 1 | 6E-96 |
| Lgmn | 2.9E-100 | 1.39708 | 1 | 1 | 6.92E-96 |
| Ubb | 2.8E-99 | 1.938131 | 0.983 | 0.997 | 6.66E-95 |
| Selplg | 8.6E-99 | 1.488028 | 0.997 | 1 | 2.02E-94 |
| Trem2 | 8.72E-98 | 1.838354 | 0.991 | 0.997 | 2.06E-93 |
| Ctsb | 1.98E-94 | 1.587931 | 1 | 1 | 4.67E-90 |
| Tyrobp | 3.87E-94 | 1.529583 | 0.992 | 0.997 | 9.13E-90 |
| Csf1r | 2.59E-93 | -1.22236 | 0.995 | 1 | 6.1E-89 |
| Rpl13 | 2.95E-93 | 2.581873 | 0.959 | 0.938 | 6.97E-89 |
| mt-Cytb | 2.09E-92 | 2.515032 | 0.981 | 0.99 | 4.93E-88 |
| Rps9 | 6.02E-91 | 2.193515 | 0.968 | 0.967 | 1.42E-86 |
| Hexb | 6.26E-87 | 0.935643 | 1 | 1 | 1.48E-82 |
| Gm15427 | 1.03E-84 | 3.396543 | 0.968 | 0.87 | 2.43E-80 |
| Laptm5 | 1.38E-84 | 1.239102 | 1 | 1 | 3.25E-80 |
| Rplp1 | 2.31E-84 | 2.083708 | 0.966 | 0.961 | 5.45E-80 |
| H2-D1 | 3.41E-84 | 2.602176 | 0.962 | 0.961 | 8.04E-80 |
| Tmem119 | 5.31E-83 | 1.611145 | 0.987 | 0.997 | 1.25E-78 |

| | | | | | |
|---|---|---|---|---|---|
| mt-Co3 | 2.03E-82 | 2.582627 | 0.947 | 0.798 | 4.78E-78 |
| Ct010467.1 | 1.2E-80 | 1.351058 | 1 | 1 | 2.83E-76 |
| Gm1821 | 2.5E-80 | 1.846036 | 0.975 | 0.987 | 5.9E-76 |
| Gsn | 5.59E-80 | 0.704185 | 0.091 | 0.619 | 1.32E-75 |
| Ppia | 8.63E-80 | 1.758926 | 0.979 | 0.993 | 2.04E-75 |
| Rps3 | 2.37E-79 | 2.621602 | 0.924 | 0.899 | 5.6E-75 |
| Ctsh | 2.97E-79 | 2.002157 | 0.958 | 0.977 | 7.01E-75 |
| Aif1 | 3.16E-78 | 2.173675 | 0.928 | 0.971 | 7.46E-74 |
| Serinc3 | 3.97E-78 | -1.59243 | 0.945 | 1 | 9.37E-74 |
| mt-Atp6 | 8.81E-77 | 2.492778 | 0.911 | 0.674 | 2.08E-72 |
| Ctsd | 1.04E-76 | 1.12686 | 1 | 1 | 2.44E-72 |
| Rps11 | 3.3E-76 | 2.23684 | 0.94 | 0.919 | 7.79E-72 |
| mt-Co2 | 1.12E-74 | 2.892311 | 0.918 | 0.736 | 2.64E-70 |
| Gm23935 | 5.52E-74 | -2.26792 | 0.926 | 1 | 1.3E-69 |
| Cd68 | 8.15E-74 | 1.694935 | 0.967 | 0.98 | 1.92E-69 |
| Ly86 | 1.13E-73 | 1.337087 | 0.996 | 0.997 | 2.66E-69 |
| Adap2os | 3.5E-73 | 2.714242 | 0.925 | 0.879 | 8.25E-69 |
| Rpl13a | 1.02E-72 | 3.510987 | 0.865 | 0.749 | 2.42E-68 |
| Gm24245 | 1.06E-71 | -0.74555 | 0.015 | 0.342 | 2.49E-67 |
| Hist1h1e | 1.71E-71 | -1.42385 | 0.083 | 0.55 | 4.03E-67 |
| Rps27rt | 7.88E-71 | 2.602018 | 0.812 | 0.394 | 1.86E-66 |
| Rps14 | 6.53E-68 | 2.050916 | 0.933 | 0.928 | 1.54E-63 |
| Gm10076 | 6.62E-68 | 2.307388 | 0.923 | 0.912 | 1.56E-63 |
| Ctsl | 9.28E-68 | 1.555186 | 0.974 | 0.987 | 2.19E-63 |
| Bsg | 6.61E-67 | 2.500758 | 0.9 | 0.909 | 1.56E-62 |
| Iapp | 7.84E-66 | -0.55489 | 0 | 0.257 | 1.85E-61 |
| Ddit4 | 9.01E-66 | -1.19417 | 0.102 | 0.58 | 2.13E-61 |
| Gm15710 | 2.77E-65 | 2.488268 | 0.818 | 0.44 | 6.54E-61 |
| Rnaset2a | 1.27E-64 | 2.015996 | 0.937 | 0.951 | 2.99E-60 |
| Rnaset2b | 1.39E-64 | 2.200094 | 0.967 | 0.951 | 3.27E-60 |
| Basp1 | 7.04E-64 | 2.487242 | 0.914 | 0.967 | 1.66E-59 |
| Adrb1 | 4.18E-63 | -1.06879 | 0.229 | 0.775 | 9.87E-59 |
| Rplp0 | 5.01E-63 | 2.396967 | 0.899 | 0.906 | 1.18E-58 |
| Rps3a1 | 1.15E-62 | 2.460547 | 0.885 | 0.834 | 2.7E-58 |
| Gm14303 | 1.92E-62 | 1.926674 | 0.82 | 0.472 | 4.54E-58 |
| Ins1 | 2.09E-62 | -0.68627 | 0 | 0.244 | 4.92E-58 |
| Calm2 | 3.43E-62 | 1.88239 | 0.938 | 0.971 | 8.09E-58 |
| Pld4 | 3.78E-62 | 1.7436 | 0.957 | 0.974 | 8.93E-58 |
| Gm4149 | 5.43E-62 | 2.627926 | 0.847 | 0.638 | 1.28E-57 |
| Rpl41 | 4.24E-61 | 1.826804 | 0.952 | 0.951 | 1E-56 |
| Laptm4a | 5.98E-61 | 1.983895 | 0.91 | 0.932 | 1.41E-56 |
| Gm9843 | 1.03E-60 | 2.897254 | 0.884 | 0.704 | 2.42E-56 |
| Rgs10 | 4.21E-60 | 1.572671 | 0.947 | 0.98 | 9.93E-56 |
| Rps18 | 1.05E-59 | 2.403856 | 0.88 | 0.811 | 2.48E-55 |
| S100a8 | 4.87E-59 | -3.00102 | 0.042 | 0.378 | 1.15E-54 |
| Chga | 5.07E-59 | -0.85425 | 0 | 0.231 | 1.2E-54 |
| Ppy | 3.69E-58 | -0.87233 | 0 | 0.228 | 8.7E-54 |

| | | | | | |
|---|---|---|---|---|---|
| Ftl2-ps | 3.85E-58 | 2.336141 | 0.932 | 0.945 | 9.08E-54 |
| Rpl10 | 8.94E-58 | 1.992964 | 0.917 | 0.896 | 2.11E-53 |
| Mid1 | 1.46E-57 | -1.06258 | 0.077 | 0.476 | 3.43E-53 |
| Rpl32 | 1.77E-56 | 2.21577 | 0.887 | 0.902 | 4.16E-52 |
| Rplp2 | 2.76E-56 | 2.027181 | 0.916 | 0.938 | 6.51E-52 |
| Olfml3 | 5.56E-56 | 1.247553 | 0.994 | 1 | 1.31E-51 |
| Ftl1-ps1 | 9.17E-56 | 2.081555 | 0.913 | 0.798 | 2.16E-51 |
| Dpysl2 | 6.53E-55 | 2.101103 | 0.921 | 0.896 | 1.54E-50 |
| Ptgds | 8.65E-55 | -0.53232 | 0 | 0.215 | 2.04E-50 |
| Tubb4a | 1.37E-54 | -0.34533 | 0.003 | 0.228 | 3.24E-50 |
| Fkbp5 | 1.4E-54 | -1.29934 | 0.154 | 0.612 | 3.3E-50 |
| S100a9 | 1.52E-54 | -2.54057 | 0.032 | 0.329 | 3.59E-50 |
| Rpl21 | 1.84E-54 | 1.950161 | 0.919 | 0.915 | 4.35E-50 |
| Tm9sf2 | 1.2E-53 | -1.73503 | 0.571 | 0.909 | 2.83E-49 |
| Rps7 | 4.23E-51 | 2.857544 | 0.814 | 0.71 | 9.98E-47 |
| Ac154912.3 | 5.15E-51 | -1.61895 | 0.03 | 0.316 | 1.21E-46 |
| Rps27 | 5.83E-51 | 1.629192 | 0.959 | 0.954 | 1.37E-46 |
| mt-Nd1 | 8.23E-51 | 1.918003 | 0.932 | 0.971 | 1.94E-46 |
| H2-K1 | 1.48E-50 | 2.570755 | 0.87 | 0.84 | 3.5E-46 |
| Rpl37a | 1.99E-50 | 1.843336 | 0.949 | 0.941 | 4.7E-46 |
| Rps20 | 5.09E-49 | 2.5059 | 0.839 | 0.795 | 1.2E-44 |
| Rps24 | 5.54E-49 | 2.384062 | 0.859 | 0.84 | 1.31E-44 |
| Atp6v0c | 1.66E-48 | 2.18455 | 0.84 | 0.866 | 3.93E-44 |
| Jun | 4.97E-48 | -1.04278 | 0.33 | 0.808 | 1.17E-43 |
| Ywhah | 5.93E-48 | 1.610326 | 0.917 | 0.958 | 1.4E-43 |
| Gapdh | 7.04E-48 | 1.750101 | 0.903 | 0.954 | 1.66E-43 |
| Ctsa | 1.49E-47 | 1.388636 | 0.966 | 0.997 | 3.52E-43 |
| Cd63 | 2.15E-47 | 2.772639 | 0.821 | 0.782 | 5.07E-43 |
| Gm44860 | 2.45E-47 | -1.10554 | 0.203 | 0.632 | 5.79E-43 |
| Gm11478 | 6.4E-47 | 3.717679 | 0.663 | 0.261 | 1.51E-42 |
| Fau | 8.12E-47 | 1.937212 | 0.915 | 0.876 | 1.92E-42 |
| Rpl17 | 9.89E-47 | 2.789302 | 0.809 | 0.772 | 2.33E-42 |
| Cmtm6 | 1.19E-46 | -1.59045 | 0.626 | 0.909 | 2.8E-42 |
| Gm25911 | 1.21E-46 | -0.39739 | 0.031 | 0.3 | 2.86E-42 |
| Rps19 | 5.06E-46 | 2.004701 | 0.896 | 0.847 | 1.19E-41 |
| Hexa | 1.75E-45 | 1.191102 | 0.975 | 0.987 | 4.12E-41 |
| Spi1 | 1.76E-45 | 2.284926 | 0.833 | 0.893 | 4.14E-41 |
| Rps18-ps3 | 1.8E-45 | 2.593384 | 0.671 | 0.283 | 4.23E-41 |
| Oaz1 | 3.48E-45 | 2.354388 | 0.814 | 0.814 | 8.21E-41 |
| Npc2 | 1.46E-44 | 2.000043 | 0.858 | 0.879 | 3.43E-40 |
| Clic1 | 2.14E-44 | 1.883906 | 0.86 | 0.912 | 5.05E-40 |
| Gm8797 | 3.89E-44 | 1.465702 | 0.893 | 0.909 | 9.18E-40 |
| Tmed7 | 1.35E-43 | -1.36362 | 0.25 | 0.658 | 3.19E-39 |
| Eif4g2 | 1.6E-43 | -0.83203 | 0.322 | 0.749 | 3.77E-39 |
| Rpl6 | 1.63E-43 | 2.731482 | 0.804 | 0.71 | 3.84E-39 |
| Rps23 | 1.72E-43 | 1.755436 | 0.901 | 0.925 | 4.05E-39 |
| Cd164 | 1.76E-43 | -1.06734 | 0.786 | 0.984 | 4.15E-39 |

| | | | | | |
|---|---|---|---|---|---|
| Rps25 | 2.36E-43 | 2.086762 | 0.891 | 0.827 | 5.57E-39 |
| Rpl30 | 7.98E-43 | 2.004846 | 0.873 | 0.863 | 1.88E-38 |
| Sft2d1 | 8.19E-43 | 1.848792 | 0.865 | 0.919 | 1.93E-38 |
| Gm3379 | 1.14E-42 | 2.527763 | 0.624 | 0.264 | 2.69E-38 |
| Rpl35 | 1.84E-42 | 2.112555 | 0.857 | 0.818 | 4.34E-38 |
| Gm3511 | 2.13E-42 | 1.924167 | 0.65 | 0.264 | 5.02E-38 |
| Gm15266 | 2.38E-42 | 2.214386 | 0.63 | 0.228 | 5.61E-38 |
| Rpl37 | 2.62E-42 | 1.633286 | 0.914 | 0.883 | 6.18E-38 |
| Rpl3 | 3.16E-42 | 2.183243 | 0.835 | 0.831 | 7.46E-38 |
| Rpsa | 4.48E-42 | 2.470055 | 0.814 | 0.788 | 1.06E-37 |
| Tmem59 | 7.51E-42 | 1.610429 | 0.873 | 0.932 | 1.77E-37 |
| Tmem50a | 9.7E-42 | 1.706543 | 0.867 | 0.919 | 2.29E-37 |
| Abhd12 | 9.99E-42 | 1.250793 | 0.952 | 0.99 | 2.36E-37 |
| Malat1 | 1.59E-41 | 0.988488 | 0.981 | 1 | 3.75E-37 |
| Rps28 | 1.68E-41 | 1.56566 | 0.917 | 0.889 | 3.95E-37 |
| Rpl23a-ps3 | 2.9E-41 | 1.950229 | 0.587 | 0.199 | 6.83E-37 |
| Rps6 | 1.18E-40 | 2.120291 | 0.816 | 0.707 | 2.79E-36 |
| Sat1 | 1.51E-40 | 2.102711 | 0.853 | 0.844 | 3.55E-36 |
| Gnb1 | 2.24E-40 | 2.394404 | 0.877 | 0.893 | 5.28E-36 |
| Hist1h4d | 4.81E-40 | -1.16081 | 0.16 | 0.58 | 1.13E-35 |
| Syngr1 | 5.66E-40 | 2.224036 | 0.827 | 0.853 | 1.34E-35 |
| Gm43028 | 6.06E-40 | 1.329749 | 0.464 | 0.046 | 1.43E-35 |
| Rpl4 | 9.86E-40 | 2.063898 | 0.85 | 0.889 | 2.33E-35 |
| Atp6v0c-ps | 1.45E-39 | 2.303954 | 0.713 | 0.436 | 3.42E-35 |
| Per1 | 1.74E-39 | -1.77514 | 0.038 | 0.287 | 4.1E-35 |
| F11r | 4.46E-39 | 1.582579 | 0.937 | 0.971 | 1.05E-34 |
| Srsf2 | 6.87E-39 | 2.39054 | 0.825 | 0.694 | 1.62E-34 |
| Rps16-ps2 | 8.88E-39 | 2.382308 | 0.649 | 0.313 | 2.1E-34 |
| Atp2a2 | 9.41E-39 | -0.83021 | 0.146 | 0.515 | 2.22E-34 |
| Tlr7 | 1.31E-38 | -0.86098 | 0.248 | 0.661 | 3.09E-34 |
| Ngp | 1.35E-38 | -3.53554 | 0.035 | 0.277 | 3.19E-34 |
| Gm10123 | 1.59E-38 | 1.848879 | 0.935 | 0.958 | 3.75E-34 |
| Grn | 1.9E-38 | 0.908926 | 0.991 | 1 | 4.47E-34 |
| Arglu1 | 2.77E-38 | 2.130826 | 0.89 | 0.821 | 6.53E-34 |
| Sh3bgrl3 | 3.59E-38 | 1.838584 | 0.838 | 0.879 | 8.47E-34 |
| Slc8a1 | 1.15E-37 | -1.95521 | 0.082 | 0.378 | 2.71E-33 |
| Cotl1 | 2.19E-37 | 2.734412 | 0.784 | 0.782 | 5.17E-33 |
| Pros1 | 2.43E-37 | -0.9707 | 0.572 | 0.886 | 5.74E-33 |
| Tpt1 | 7.23E-37 | 1.973993 | 0.861 | 0.915 | 1.71E-32 |
| Dhrs3 | 8.81E-37 | 2.11813 | 0.861 | 0.896 | 2.08E-32 |
| Gm8464 | 9.3E-37 | -0.43125 | 0.041 | 0.28 | 2.19E-32 |
| Jmjd1c | 1.46E-36 | -0.84239 | 0.172 | 0.544 | 3.45E-32 |
| Gm3362 | 2.69E-36 | 1.050869 | 0.482 | 0.085 | 6.34E-32 |
| Ikzf1 | 7.95E-36 | -0.43584 | 0.278 | 0.707 | 1.88E-31 |
| Rpl19 | 1.15E-35 | 1.992811 | 0.822 | 0.847 | 2.71E-31 |
| Lamp1 | 1.37E-35 | 1.114112 | 0.972 | 0.997 | 3.22E-31 |
| Nptn | 1.62E-35 | -1.37191 | 0.463 | 0.779 | 3.83E-31 |

| | | | | | |
|---|---|---|---|---|---|
| Rps10 | 3.97E-35 | 2.418265 | 0.78 | 0.779 | 9.35E-31 |
| Gm5559 | 4.54E-35 | 2.987467 | 0.71 | 0.482 | 1.07E-30 |
| Mcl1 | 8.56E-35 | -0.47069 | 0.343 | 0.775 | 2.02E-30 |
| Rps16 | 1.04E-34 | 2.07003 | 0.805 | 0.736 | 2.44E-30 |
| Gm12918 | 1.04E-34 | 1.327218 | 0.584 | 0.215 | 2.46E-30 |
| Nrip1 | 1.06E-34 | -0.57102 | 0.37 | 0.798 | 2.49E-30 |
| Rpl18 | 2.49E-34 | 1.960462 | 0.819 | 0.847 | 5.87E-30 |
| Rgs1 | 3.55E-34 | -0.65572 | 0.066 | 0.342 | 8.38E-30 |
| Kcnk6 | 4.11E-34 | -0.3292 | 0.295 | 0.73 | 9.7E-30 |
| Itm2c | 6.36E-34 | 1.395848 | 0.906 | 0.967 | 1.5E-29 |
| Rpl36 | 6.62E-34 | 1.907968 | 0.845 | 0.788 | 1.56E-29 |
| Cd37 | 6.85E-34 | 1.79208 | 0.839 | 0.935 | 1.62E-29 |
| Atp6v0b | 6.97E-34 | 1.686966 | 0.88 | 0.948 | 1.64E-29 |
| 4933406I1 | 7.19E-34 | -0.62729 | 0.082 | 0.371 | 1.7E-29 |
| Abi3 | 7.52E-34 | 2.065246 | 0.805 | 0.941 | 1.77E-29 |
| Tnfrsf21 | 8.52E-34 | -1.43661 | 0.308 | 0.645 | 2.01E-29 |
| Gm2000 | 9.63E-34 | 2.120327 | 0.649 | 0.345 | 2.27E-29 |
| Teddm2 | 1.18E-33 | -1.10612 | 0.055 | 0.309 | 2.79E-29 |
| Ac154912. | 1.39E-33 | -1.19294 | 0.041 | 0.27 | 3.29E-29 |
| Rps4x | 1.46E-33 | 2.27485 | 0.808 | 0.792 | 3.45E-29 |
| Akna | 2.18E-33 | -0.57486 | 0.203 | 0.57 | 5.13E-29 |
| Fcgr3 | 2.76E-33 | 1.113953 | 0.94 | 0.997 | 6.52E-29 |
| Actb | 3.37E-33 | 0.862956 | 1 | 1 | 7.94E-29 |
| Mpeg1 | 4.87E-33 | -0.87115 | 0.66 | 0.958 | 1.15E-28 |
| Capzb | 4.89E-33 | 1.943434 | 0.805 | 0.85 | 1.15E-28 |
| Rnase4 | 5.05E-33 | 1.306803 | 0.917 | 0.984 | 1.19E-28 |
| Rpl27-ps3 | 8.09E-33 | 1.982781 | 0.45 | 0.088 | 1.91E-28 |
| Gstm1 | 9.6E-33 | 0.628548 | 0.072 | 0.352 | 2.27E-28 |
| Zscan26 | 1.04E-32 | 2.309437 | 0.806 | 0.651 | 2.46E-28 |
| Tsc22d3 | 1.22E-32 | -0.5709 | 0.173 | 0.534 | 2.89E-28 |
| Cd63-ps | 1.56E-32 | 2.512112 | 0.662 | 0.404 | 3.67E-28 |
| Gm10180 | 1.59E-32 | 1.886676 | 0.54 | 0.202 | 3.76E-28 |
| Arpc1b | 1.77E-32 | 2.218756 | 0.767 | 0.798 | 4.17E-28 |
| Siglech | 1.88E-32 | 0.960868 | 0.992 | 1 | 4.43E-28 |
| Mtmr6 | 1.96E-32 | -1.76847 | 0.122 | 0.414 | 4.63E-28 |
| Rpl17-ps3 | 2.27E-32 | 1.862409 | 0.472 | 0.111 | 5.35E-28 |
| Gm10073 | 6.16E-32 | 1.638759 | 0.659 | 0.371 | 1.45E-27 |
| Ncoa4 | 6.32E-32 | -1.54266 | 0.106 | 0.378 | 1.49E-27 |
| Copa | 6.72E-32 | -1.23007 | 0.176 | 0.511 | 1.59E-27 |
| Lyz2 | 6.99E-32 | 2.511126 | 0.786 | 0.853 | 1.65E-27 |
| Ptafr | 1.55E-31 | 2.293913 | 0.773 | 0.668 | 3.66E-27 |
| Gm10443 | 1.67E-31 | 1.03836 | 0.458 | 0.098 | 3.94E-27 |
| Gm10925 | 2.18E-31 | 1.410913 | 0.89 | 0.795 | 5.13E-27 |
| Rps12-ps9 | 2.22E-31 | 1.719802 | 0.487 | 0.13 | 5.24E-27 |
| Gm11942 | 2.81E-31 | 1.310587 | 0.522 | 0.173 | 6.62E-27 |
| Bhlhe41 | 3.1E-31 | 2.162238 | 0.851 | 0.733 | 7.32E-27 |
| Rpl11 | 3.29E-31 | 2.296309 | 0.757 | 0.7 | 7.77E-27 |

| | | | | | |
|---|---|---|---|---|---|
| Rn7sk | 3.38E-31 | -1.01271 | 0.102 | 0.391 | 7.98E-27 |
| Slc25a5 | 5.64E-31 | 1.869814 | 0.817 | 0.876 | 1.33E-26 |
| Rps19-ps6 | 5.88E-31 | 2.21126 | 0.65 | 0.349 | 1.39E-26 |
| Irf8 | 7.58E-31 | 2.042709 | 0.826 | 0.873 | 1.79E-26 |
| Gm6204 | 7.72E-31 | 2.069594 | 0.625 | 0.329 | 1.82E-26 |
| Otulinl | 7.93E-31 | 1.450356 | 0.902 | 0.932 | 1.87E-26 |
| Tubb5 | 1.16E-30 | 2.001377 | 0.83 | 0.743 | 2.73E-26 |
| Rpl23a | 1.43E-30 | 2.000152 | 0.779 | 0.759 | 3.37E-26 |
| Nckap1l | 1.47E-30 | -0.76069 | 0.429 | 0.811 | 3.46E-26 |
| Rps27a | 1.71E-30 | 1.904917 | 0.812 | 0.801 | 4.04E-26 |
| Gm13456 | 2.36E-30 | 1.452079 | 0.965 | 0.928 | 5.57E-26 |
| Gnai2 | 2.48E-30 | 1.484671 | 0.926 | 0.977 | 5.86E-26 |
| G530011O( | 2.54E-30 | -0.25244 | 0.24 | 0.642 | 6E-26 |
| Rapgef5 | 2.83E-30 | -0.8706 | 0.17 | 0.502 | 6.69E-26 |
| H3f3b | 3.24E-30 | 1.443282 | 0.888 | 0.938 | 7.63E-26 |
| Tagap | 3.46E-30 | -1.18341 | 0.244 | 0.59 | 8.16E-26 |
| Rps23-ps1 | 4.17E-30 | 1.305478 | 0.461 | 0.121 | 9.83E-26 |
| Rpl34 | 4.19E-30 | 1.888341 | 0.822 | 0.782 | 9.88E-26 |
| Gm7536 | 4.44E-30 | 1.973767 | 0.633 | 0.384 | 1.05E-25 |
| Fam91a1 | 6.64E-30 | -2.1044 | 0.026 | 0.208 | 1.57E-25 |
| Eif1 | 7.17E-30 | 1.572697 | 0.898 | 0.948 | 1.69E-25 |
| Rps5 | 9.5E-30 | 2.479619 | 0.757 | 0.723 | 2.24E-25 |
| Abcc3 | 9.73E-30 | -0.52351 | 0.21 | 0.567 | 2.3E-25 |
| Numb | 1.29E-29 | -0.61816 | 0.381 | 0.785 | 3.05E-25 |
| Bin1 | 2.05E-29 | 1.577806 | 0.86 | 0.954 | 4.83E-25 |
| Unc93b1 | 2.08E-29 | 0.936437 | 0.988 | 0.997 | 4.9E-25 |
| Slc2a5 | 2.27E-29 | 1.461304 | 0.883 | 0.919 | 5.35E-25 |
| Rap1gds1 | 3.76E-29 | -0.43106 | 0.291 | 0.664 | 8.86E-25 |
| Zfp691 | 4.03E-29 | 2.646309 | 0.699 | 0.528 | 9.51E-25 |
| Rps13 | 4.57E-29 | 2.086985 | 0.764 | 0.756 | 1.08E-24 |
| Anxa3 | 5.05E-29 | 1.825881 | 0.799 | 0.808 | 1.19E-24 |
| Gm16378 | 7.18E-29 | 0.880299 | 0.366 | 0.036 | 1.69E-24 |
| Rpl26 | 9.22E-29 | 2.036539 | 0.781 | 0.772 | 2.18E-24 |
| Ppp1r21 | 1.41E-28 | -0.25476 | 0.169 | 0.505 | 3.32E-24 |
| Abca1 | 1.5E-28 | -0.71941 | 0.351 | 0.743 | 3.55E-24 |
| Wdr89 | 1.6E-28 | 1.518013 | 0.644 | 0.381 | 3.79E-24 |
| Rpl31-ps8 | 2.04E-28 | 2.318117 | 0.61 | 0.352 | 4.82E-24 |
| Rpl23 | 3.9E-28 | 1.882585 | 0.83 | 0.87 | 9.21E-24 |
| Gm8730 | 4.25E-28 | 1.982919 | 0.738 | 0.609 | 1E-23 |
| Rpl7 | 4.54E-28 | 1.717806 | 0.821 | 0.857 | 1.07E-23 |
| Runx1 | 5.09E-28 | -0.28304 | 0.216 | 0.567 | 1.2E-23 |
| Ldhb | 7.59E-28 | 1.99279 | 0.773 | 0.866 | 1.79E-23 |
| Mtdh | 8.11E-28 | 1.572907 | 0.913 | 0.941 | 1.91E-23 |
| Rpl9 | 8.12E-28 | 2.129023 | 0.766 | 0.759 | 1.92E-23 |
| Gm6900 | 1.11E-27 | 0.832336 | 0.355 | 0.036 | 2.63E-23 |
| Bin2 | 1.65E-27 | 1.396161 | 0.865 | 0.941 | 3.88E-23 |
| Tmem170b | 1.93E-27 | -0.26019 | 0.218 | 0.564 | 4.56E-23 |

| | | | | | |
|---|---|---|---|---|---|
| Fbxl14 | 2.02E-27 | -1.513 | 0.058 | 0.27 | 4.76E-23 |
| Serpine2 | 2.42E-27 | 1.200699 | 0.916 | 0.971 | 5.71E-23 |
| Gm2574 | 3.18E-27 | 1.646162 | 0.532 | 0.235 | 7.5E-23 |
| Akr1a1 | 3.77E-27 | 2.050587 | 0.773 | 0.756 | 8.9E-23 |
| Rpl14 | 3.89E-27 | 2.222984 | 0.788 | 0.821 | 9.18E-23 |
| Gm2225 | 6.8E-27 | 0.867864 | 0.393 | 0.068 | 1.6E-22 |
| Ccr5 | 1.08E-26 | -1.0053 | 0.495 | 0.798 | 2.56E-22 |
| Tmem86a | 1.18E-26 | 2.304243 | 0.733 | 0.739 | 2.78E-22 |
| Gm10359 | 1.22E-26 | 1.207221 | 0.404 | 0.088 | 2.87E-22 |
| Rpl36-ps2 | 1.42E-26 | 2.162687 | 0.553 | 0.244 | 3.36E-22 |
| Elmo1 | 2.09E-26 | -0.78862 | 0.476 | 0.798 | 4.92E-22 |
| Rpl31 | 2.31E-26 | 1.900357 | 0.777 | 0.765 | 5.45E-22 |
| Fos | 2.87E-26 | -1.23692 | 0.186 | 0.524 | 6.78E-22 |
| Tpm3 | 2.97E-26 | 2.140432 | 0.84 | 0.925 | 7.01E-22 |
| Mbp | 3.27E-26 | 1.152075 | 0.069 | 0.303 | 7.72E-22 |
| Gm28661 | 3.34E-26 | 1.177186 | 0.819 | 0.664 | 7.88E-22 |
| Spire1 | 3.46E-26 | -0.50407 | 0.182 | 0.489 | 8.16E-22 |
| Gm28437 | 3.99E-26 | 1.105018 | 0.874 | 0.756 | 9.4E-22 |
| Kctd12 | 4.79E-26 | -0.33924 | 0.59 | 0.941 | 1.13E-21 |
| Gm11518 | 5.32E-26 | 2.219933 | 0.482 | 0.186 | 1.25E-21 |
| Gm6394 | 5.89E-26 | 1.718078 | 0.494 | 0.186 | 1.39E-21 |
| Gm8430 | 7.21E-26 | 2.142925 | 0.555 | 0.277 | 1.7E-21 |
| Gpx1 | 8.09E-26 | 2.163054 | 0.764 | 0.752 | 1.91E-21 |
| Arhgdia | 1.67E-25 | 2.33696 | 0.758 | 0.847 | 3.94E-21 |
| Ddb1 | 1.9E-25 | -0.76728 | 0.1 | 0.349 | 4.48E-21 |
| Gm6472 | 1.98E-25 | 1.698654 | 0.477 | 0.176 | 4.67E-21 |
| Gm24187 | 1.99E-25 | -0.40518 | 0.599 | 0.919 | 4.7E-21 |
| Rpl27a | 2.22E-25 | 2.17885 | 0.762 | 0.73 | 5.23E-21 |
| Gm6977 | 2.73E-25 | 1.975018 | 0.596 | 0.329 | 6.45E-21 |
| Myadm | 4.82E-25 | -0.55116 | 0.084 | 0.322 | 1.14E-20 |
| Rpl37rt | 6.31E-25 | 1.741759 | 0.656 | 0.41 | 1.49E-20 |
| Gm4202 | 7.23E-25 | -0.26399 | 0.385 | 0.697 | 1.71E-20 |
| Abcg1 | 8.7E-25 | 2.554575 | 0.74 | 0.707 | 2.05E-20 |
| Fchsd2 | 1.32E-24 | -0.27904 | 0.212 | 0.537 | 3.11E-20 |
| Man1a | 1.42E-24 | -0.81193 | 0.219 | 0.518 | 3.35E-20 |
| Pik3ap1 | 1.49E-24 | -1.13534 | 0.13 | 0.381 | 3.52E-20 |
| Gm12537 | 1.77E-24 | 1.417731 | 0.374 | 0.078 | 4.18E-20 |
| Gm29216 | 1.97E-24 | 1.180711 | 0.766 | 0.58 | 4.64E-20 |
| Rps13-ps2 | 3.19E-24 | 2.480455 | 0.545 | 0.283 | 7.52E-20 |
| Rps10-ps1 | 3.2E-24 | 3.559574 | 0.497 | 0.215 | 7.55E-20 |
| Sun2 | 3.27E-24 | -1.3985 | 0.408 | 0.723 | 7.72E-20 |
| Rhob | 3.74E-24 | -0.68148 | 0.782 | 0.967 | 8.81E-20 |
| Rpl19-ps11 | 4.31E-24 | 2.016105 | 0.636 | 0.436 | 1.02E-19 |
| Msl2 | 4.37E-24 | -1.29269 | 0.08 | 0.303 | 1.03E-19 |
| mt-Nd4 | 4.86E-24 | 1.585129 | 0.878 | 0.935 | 1.15E-19 |
| Gm44705 | 5.14E-24 | 0.833289 | 0.288 | 0.013 | 1.21E-19 |
| Ly6e | 6.2E-24 | 1.459109 | 0.926 | 0.98 | 1.46E-19 |

| | | | | | |
|---|---|---|---|---|---|
| Rps26 | 6.21E-24 | 1.999143 | 0.734 | 0.655 | 1.47E-19 |
| Rpl28 | 7.02E-24 | 2.093321 | 0.733 | 0.726 | 1.66E-19 |
| Ppib | 7.61E-24 | 1.909689 | 0.75 | 0.86 | 1.8E-19 |
| 1700017BC | 8.2E-24 | -0.59637 | 0.441 | 0.765 | 1.94E-19 |
| Fcgr2b | 1.44E-23 | 1.710657 | 0.782 | 0.805 | 3.41E-19 |
| Rps10-ps2 | 1.79E-23 | 2.995389 | 0.587 | 0.384 | 4.22E-19 |
| Slc40a1 | 1.92E-23 | -1.08851 | 0.274 | 0.56 | 4.53E-19 |
| Cd300c2 | 1.99E-23 | 1.681072 | 0.793 | 0.879 | 4.7E-19 |
| Gm45221 | 2.08E-23 | -2.31175 | 0.038 | 0.208 | 4.9E-19 |
| Efhd2 | 2.35E-23 | 1.832385 | 0.813 | 0.922 | 5.54E-19 |
| Tm6sf1 | 2.37E-23 | -0.48043 | 0.271 | 0.612 | 5.59E-19 |
| Rps12 | 2.41E-23 | 1.616356 | 0.807 | 0.811 | 5.69E-19 |
| Gm15726 | 2.48E-23 | -1.46222 | 0.041 | 0.215 | 5.86E-19 |
| Gng10 | 2.75E-23 | 1.795568 | 0.765 | 0.906 | 6.49E-19 |
| Rps17 | 3.13E-23 | 1.80709 | 0.756 | 0.749 | 7.38E-19 |
| Gpr165 | 3.35E-23 | -1.338 | 0.073 | 0.287 | 7.89E-19 |
| Txnip | 3.96E-23 | -0.55597 | 0.518 | 0.831 | 9.34E-19 |
| Cyfip1 | 4.42E-23 | -0.48828 | 0.678 | 0.938 | 1.04E-18 |
| Crlf2 | 4.95E-23 | 3.763698 | 0.462 | 0.202 | 1.17E-18 |
| Syngr2 | 5.44E-23 | 1.773762 | 0.772 | 0.866 | 1.28E-18 |
| Sgk1 | 5.56E-23 | -0.74897 | 0.481 | 0.795 | 1.31E-18 |
| Dhx9 | 5.57E-23 | -0.27939 | 0.088 | 0.322 | 1.31E-18 |
| Vsir | 5.89E-23 | 0.872972 | 0.958 | 0.997 | 1.39E-18 |
| Tnks2 | 5.9E-23 | -0.36494 | 0.194 | 0.489 | 1.39E-18 |
| Clta | 7.75E-23 | 1.786629 | 0.788 | 0.883 | 1.83E-18 |
| Ckb | 8.53E-23 | 1.131637 | 0.858 | 0.958 | 2.01E-18 |
| Man1c1 | 9.47E-23 | -0.55018 | 0.303 | 0.619 | 2.23E-18 |
| Sh2b3 | 1.06E-22 | -0.35794 | 0.239 | 0.541 | 2.5E-18 |
| Cox4i1 | 1.06E-22 | 1.844352 | 0.773 | 0.837 | 2.51E-18 |
| Gm12166 | 1.07E-22 | 1.766531 | 0.785 | 0.801 | 2.52E-18 |
| Gm10240 | 1.37E-22 | 1.877571 | 0.478 | 0.205 | 3.24E-18 |
| Skil | 1.55E-22 | 1.790424 | 0.793 | 0.723 | 3.66E-18 |
| Ifngr1 | 1.63E-22 | 0.971766 | 0.96 | 0.99 | 3.85E-18 |
| Acly | 1.9E-22 | -1.11385 | 0.14 | 0.388 | 4.48E-18 |
| Snx18 | 2.09E-22 | -0.5358 | 0.438 | 0.779 | 4.92E-18 |
| Itgb1 | 2.36E-22 | -0.47246 | 0.393 | 0.746 | 5.56E-18 |
| Rpl18-ps1 | 2.38E-22 | 0.797044 | 0.356 | 0.075 | 5.62E-18 |
| Bmf | 2.57E-22 | -0.45775 | 0.081 | 0.296 | 6.07E-18 |
| Tlr13 | 3.01E-22 | -0.43174 | 0.187 | 0.472 | 7.09E-18 |
| Tlr4 | 3.13E-22 | -1.35988 | 0.099 | 0.322 | 7.39E-18 |
| Vwa5a | 4.12E-22 | -1.69425 | 0.112 | 0.339 | 9.71E-18 |
| Prps2 | 4.61E-22 | -0.46863 | 0.069 | 0.27 | 1.09E-17 |
| Actg1 | 5.7E-22 | 1.429845 | 0.827 | 0.893 | 1.35E-17 |
| Cfh | 6.53E-22 | -0.3654 | 0.635 | 0.945 | 1.54E-17 |
| Ncoa3 | 8.38E-22 | 1.939489 | 0.768 | 0.687 | 1.98E-17 |
| Rnf169 | 9.04E-22 | 0.408073 | 0.138 | 0.41 | 2.13E-17 |
| Apc | 1.2E-21 | -0.60851 | 0.123 | 0.375 | 2.83E-17 |

| | | | | | |
|---|---|---|---|---|---|
| Tmx3 | 1.23E-21 | -0.99186 | 0.117 | 0.349 | 2.91E-17 |
| Ddx3x | 1.37E-21 | -0.76197 | 0.252 | 0.544 | 3.24E-17 |
| Gm26825 | 1.67E-21 | 1.866195 | 0.704 | 0.616 | 3.93E-17 |
| Gm7336 | 2.73E-21 | 1.89074 | 0.431 | 0.173 | 6.45E-17 |
| Gm6988 | 2.92E-21 | 0.865372 | 0.354 | 0.081 | 6.9E-17 |
| Rpl3-ps1 | 3.03E-21 | 2.075493 | 0.75 | 0.645 | 7.14E-17 |
| Mob1a | 3.13E-21 | -0.71361 | 0.428 | 0.73 | 7.38E-17 |
| Itga9 | 3.16E-21 | -0.71259 | 0.15 | 0.41 | 7.44E-17 |
| Sacm1l | 3.49E-21 | 0.355804 | 0.054 | 0.238 | 8.23E-17 |
| Camk1 | 3.81E-21 | 1.759344 | 0.739 | 0.795 | 8.99E-17 |
| Comt | 4E-21 | 1.792988 | 0.769 | 0.798 | 9.43E-17 |
| Rps6-ps4 | 4.31E-21 | 1.51473 | 0.558 | 0.309 | 1.02E-16 |
| Erp29 | 5.38E-21 | 1.861217 | 0.744 | 0.84 | 1.27E-16 |
| Alox5ap | 5.45E-21 | 1.935283 | 0.725 | 0.779 | 1.29E-16 |
| Aftph | 7.75E-21 | -0.94959 | 0.186 | 0.45 | 1.83E-16 |
| Pycard | 7.93E-21 | 1.818515 | 0.764 | 0.785 | 1.87E-16 |
| Rtn4rl1 | 8.29E-21 | -0.6919 | 0.434 | 0.73 | 1.96E-16 |
| Nup153 | 9.46E-21 | -1.01769 | 0.058 | 0.241 | 2.23E-16 |
| Gm45223 | 1.09E-20 | -0.97563 | 0.095 | 0.306 | 2.57E-16 |
| Fmnl1 | 1.4E-20 | 0.538242 | 0.212 | 0.511 | 3.31E-16 |
| H2-DMa | 1.79E-20 | 1.953903 | 0.682 | 0.651 | 4.22E-16 |
| Rasa3 | 2.11E-20 | -0.45014 | 0.125 | 0.362 | 4.98E-16 |
| Gm10163 | 2.18E-20 | 1.712738 | 0.472 | 0.195 | 5.13E-16 |
| Actr2 | 2.2E-20 | 1.515675 | 0.878 | 0.889 | 5.2E-16 |
| Ang | 2.47E-20 | 2.277801 | 0.74 | 0.798 | 5.83E-16 |
| Rpsa-ps10 | 3.05E-20 | 2.240851 | 0.589 | 0.378 | 7.2E-16 |
| Ptms | 3.99E-20 | 3.320143 | 0.688 | 0.694 | 9.41E-16 |
| Zfp62 | 4.12E-20 | -0.72157 | 0.082 | 0.287 | 9.71E-16 |
| Gm10288 | 4.51E-20 | 1.700698 | 0.498 | 0.257 | 1.06E-15 |
| Etnk1 | 4.73E-20 | -0.83495 | 0.167 | 0.427 | 1.12E-15 |
| Gm6136 | 5.04E-20 | 1.916098 | 0.588 | 0.365 | 1.19E-15 |
| Rpl17-ps5 | 5.78E-20 | 1.264193 | 0.318 | 0.062 | 1.36E-15 |
| P3h2 | 5.85E-20 | -1.20855 | 0.147 | 0.375 | 1.38E-15 |
| Gpr155 | 7.55E-20 | -0.90426 | 0.152 | 0.404 | 1.78E-15 |
| Tmem168 | 7.9E-20 | -1.13113 | 0.087 | 0.287 | 1.86E-15 |
| Gm11787 | 8.23E-20 | -0.34936 | 0.043 | 0.202 | 1.94E-15 |
| Gabarap | 1.05E-19 | 1.755401 | 0.728 | 0.795 | 2.48E-15 |
| Oaz1-ps | 1.21E-19 | 1.654507 | 0.707 | 0.635 | 2.85E-15 |
| Socs3 | 1.3E-19 | -0.82227 | 0.058 | 0.231 | 3.06E-15 |
| Slc43a2 | 1.73E-19 | -0.33962 | 0.126 | 0.362 | 4.07E-15 |
| Tgfb1 | 1.82E-19 | 1.0255 | 0.16 | 0.433 | 4.3E-15 |
| Gm5805 | 1.83E-19 | 1.137748 | 0.472 | 0.208 | 4.31E-15 |
| Tm9sf3 | 1.94E-19 | -0.8882 | 0.301 | 0.583 | 4.59E-15 |
| Gm2a | 1.95E-19 | 1.592138 | 0.822 | 0.762 | 4.59E-15 |
| Adgre1 | 2.05E-19 | -1.17276 | 0.461 | 0.687 | 4.84E-15 |
| Tlr3 | 2.07E-19 | -0.3418 | 0.148 | 0.401 | 4.87E-15 |
| Api5 | 2.14E-19 | -1.53758 | 0.094 | 0.29 | 5.05E-15 |

| | | | | | |
|---|---|---|---|---|---|
| Rpl13-ps6 | 2.47E-19 | 1.877748 | 0.483 | 0.235 | 5.82E-15 |
| Cerk | 2.5E-19 | -0.31518 | 0.233 | 0.515 | 5.9E-15 |
| Rpl34-ps1 | 2.61E-19 | 1.627178 | 0.487 | 0.244 | 6.15E-15 |
| Tbc1d14 | 2.63E-19 | -0.89242 | 0.144 | 0.378 | 6.21E-15 |
| Tjp1 | 3.19E-19 | 0.257699 | 0.17 | 0.446 | 7.52E-15 |
| Ssh2 | 3.45E-19 | -0.38538 | 0.674 | 0.951 | 8.14E-15 |
| Mertk | 3.56E-19 | -0.3437 | 0.68 | 0.958 | 8.39E-15 |
| Ac165079.: | 3.58E-19 | 1.38277 | 0.301 | 0.055 | 8.43E-15 |
| Extl3 | 4.48E-19 | -0.48511 | 0.238 | 0.531 | 1.06E-14 |
| Adap2 | 4.72E-19 | 1.553739 | 0.845 | 0.951 | 1.11E-14 |
| Rpl17-ps8 | 4.77E-19 | 1.362958 | 0.249 | 0.02 | 1.13E-14 |
| Tsc22d2 | 5.28E-19 | -1.02861 | 0.146 | 0.375 | 1.25E-14 |
| Uba52 | 7.93E-19 | 1.962358 | 0.763 | 0.772 | 1.87E-14 |
| Zfp871 | 8.17E-19 | -0.80469 | 0.108 | 0.319 | 1.93E-14 |
| Cab39 | 8.76E-19 | -0.5037 | 0.17 | 0.427 | 2.07E-14 |
| Rpl7a-ps7 | 8.98E-19 | 1.249908 | 0.305 | 0.062 | 2.12E-14 |
| Ptbp3 | 9.44E-19 | -0.57415 | 0.303 | 0.596 | 2.23E-14 |
| Rpl9-ps6 | 1.08E-18 | 2.395144 | 0.642 | 0.55 | 2.55E-14 |
| Washc4 | 1.13E-18 | -0.33459 | 0.244 | 0.524 | 2.68E-14 |
| Rps21 | 1.19E-18 | 1.54044 | 0.752 | 0.726 | 2.81E-14 |
| Gm9794 | 1.26E-18 | 1.827655 | 0.617 | 0.472 | 2.98E-14 |
| Trpc4ap | 1.35E-18 | -0.59721 | 0.312 | 0.596 | 3.18E-14 |
| Dpp8 | 1.47E-18 | -0.50466 | 0.1 | 0.309 | 3.46E-14 |
| Sirpa | 1.5E-18 | -0.50386 | 0.862 | 0.99 | 3.54E-14 |
| Cd14 | 1.52E-18 | 1.536216 | 0.782 | 0.866 | 3.6E-14 |
| Galnt1 | 1.62E-18 | -0.85393 | 0.233 | 0.495 | 3.83E-14 |
| Tmem135 | 1.64E-18 | -0.66918 | 0.235 | 0.489 | 3.87E-14 |
| Rsrp1 | 1.79E-18 | 1.76544 | 0.852 | 0.945 | 4.23E-14 |
| Snx13 | 1.95E-18 | -0.44274 | 0.233 | 0.502 | 4.59E-14 |
| Usp24 | 2.28E-18 | -0.60698 | 0.134 | 0.365 | 5.37E-14 |
| Rin2 | 2.33E-18 | -0.58905 | 0.429 | 0.707 | 5.49E-14 |
| Gm8960 | 2.74E-18 | 1.192178 | 0.326 | 0.085 | 6.46E-14 |
| Slc7a8 | 3.08E-18 | -0.37862 | 0.457 | 0.762 | 7.27E-14 |
| Selenot | 3.93E-18 | -0.53343 | 0.183 | 0.423 | 9.28E-14 |
| Caprin1 | 3.95E-18 | -0.73699 | 0.175 | 0.417 | 9.33E-14 |
| Psap | 3.98E-18 | 0.75181 | 0.999 | 1 | 9.4E-14 |
| Lag3 | 4.13E-18 | 1.432556 | 0.81 | 0.834 | 9.75E-14 |
| Cggbp1 | 4.2E-18 | -0.44834 | 0.14 | 0.371 | 9.9E-14 |
| Asph | 4.2E-18 | -0.96531 | 0.519 | 0.752 | 9.91E-14 |
| Vps35 | 5.05E-18 | -1.5652 | 0.246 | 0.476 | 1.19E-13 |
| Plxdc2 | 5.63E-18 | -0.43024 | 0.848 | 0.971 | 1.33E-13 |
| Sipa1 | 5.65E-18 | -0.46096 | 0.46 | 0.759 | 1.33E-13 |
| Arpc4 | 5.71E-18 | 1.817071 | 0.769 | 0.863 | 1.35E-13 |
| Fam234b | 8.68E-18 | -0.77168 | 0.074 | 0.254 | 2.05E-13 |
| Rpl7a | 8.74E-18 | 2.334773 | 0.691 | 0.726 | 2.06E-13 |
| Gprasp1 | 8.79E-18 | -0.86056 | 0.079 | 0.264 | 2.07E-13 |
| Arhgdib | 8.83E-18 | 1.847444 | 0.736 | 0.818 | 2.08E-13 |

| | | | | | |
|---|---|---|---|---|---|
| Hdlbp | 9.24E-18 | -0.42192 | 0.171 | 0.42 | 2.18E-13 |
| Arpc2 | 9.54E-18 | 1.355499 | 0.838 | 0.909 | 2.25E-13 |
| Rps8 | 1.06E-17 | 2.09792 | 0.802 | 0.831 | 2.5E-13 |
| Susd3 | 1.12E-17 | 1.684072 | 0.743 | 0.847 | 2.63E-13 |
| Usp9x | 1.15E-17 | -1.18528 | 0.153 | 0.381 | 2.71E-13 |
| Tmsb10 | 1.2E-17 | 3.392263 | 0.48 | 0.326 | 2.82E-13 |
| Cep68 | 1.23E-17 | -0.25772 | 0.154 | 0.397 | 2.89E-13 |
| Gm5905 | 1.33E-17 | 2.168356 | 0.661 | 0.485 | 3.14E-13 |
| Man2b2 | 1.33E-17 | -1.1491 | 0.355 | 0.586 | 3.15E-13 |
| Hspa8 | 1.52E-17 | 1.111462 | 0.918 | 0.967 | 3.59E-13 |
| Ogdh | 1.78E-17 | -0.39753 | 0.194 | 0.436 | 4.19E-13 |
| Jade2 | 1.88E-17 | 0.418072 | 0.135 | 0.365 | 4.44E-13 |
| Limd2 | 1.93E-17 | 2.0548 | 0.742 | 0.818 | 4.56E-13 |
| Aldoa | 1.95E-17 | 1.918654 | 0.736 | 0.824 | 4.6E-13 |
| Gm11361 | 2.07E-17 | 1.120361 | 0.508 | 0.283 | 4.87E-13 |
| Tmed9 | 2.57E-17 | 2.457202 | 0.65 | 0.678 | 6.07E-13 |
| Tubgcp5 | 2.82E-17 | -0.26796 | 0.283 | 0.573 | 6.65E-13 |
| Pdia4 | 3E-17 | -0.51342 | 0.317 | 0.606 | 7.08E-13 |
| Bri3 | 3.17E-17 | 2.730527 | 0.615 | 0.596 | 7.49E-13 |
| Slc30a9 | 4.86E-17 | -0.86404 | 0.115 | 0.319 | 1.15E-12 |
| Gm6180 | 5.35E-17 | 1.31076 | 0.404 | 0.173 | 1.26E-12 |
| Creg1 | 5.38E-17 | 1.514531 | 0.747 | 0.805 | 1.27E-12 |
| Lrch1 | 5.44E-17 | -0.35917 | 0.206 | 0.472 | 1.28E-12 |
| Scamp2 | 5.67E-17 | 1.125847 | 0.878 | 0.954 | 1.34E-12 |
| Gm5879 | 6.31E-17 | 1.107086 | 0.403 | 0.176 | 1.49E-12 |
| Gm9703 | 7.7E-17 | 0.995892 | 0.324 | 0.091 | 1.82E-12 |
| Gm14586 | 7.71E-17 | 1.418496 | 0.468 | 0.251 | 1.82E-12 |
| Rps26-ps1 | 7.89E-17 | 2.071058 | 0.35 | 0.107 | 1.86E-12 |
| Pip4k2a | 8.09E-17 | 0.347451 | 0.395 | 0.765 | 1.91E-12 |
| Ipo13 | 8.1E-17 | -0.86264 | 0.057 | 0.215 | 1.91E-12 |
| Il17ra | 9.34E-17 | -0.29462 | 0.334 | 0.606 | 2.2E-12 |
| Elmsan1 | 1.06E-16 | -0.50906 | 0.051 | 0.202 | 2.5E-12 |
| Ptpra | 1.14E-16 | -0.33151 | 0.461 | 0.775 | 2.68E-12 |
| Gm9625 | 1.26E-16 | 1.571793 | 0.492 | 0.277 | 2.97E-12 |
| Brox | 1.3E-16 | -1.71312 | 0.107 | 0.29 | 3.07E-12 |
| Gm10059 | 1.32E-16 | 0.789036 | 0.359 | 0.121 | 3.12E-12 |
| Rpl27 | 1.33E-16 | 1.969967 | 0.681 | 0.71 | 3.15E-12 |
| Sla | 1.39E-16 | -0.53259 | 0.394 | 0.681 | 3.28E-12 |
| Il6st | 1.44E-16 | -0.26398 | 0.366 | 0.664 | 3.41E-12 |
| Gm3608 | 1.61E-16 | -0.36965 | 0.14 | 0.352 | 3.81E-12 |
| Serinc1 | 1.68E-16 | -0.49687 | 0.494 | 0.772 | 3.97E-12 |
| Sort1 | 1.91E-16 | -0.40033 | 0.301 | 0.577 | 4.5E-12 |
| Fam210a | 1.93E-16 | -0.25868 | 0.112 | 0.313 | 4.56E-12 |
| Tmem87b | 2.31E-16 | -0.42409 | 0.086 | 0.267 | 5.44E-12 |
| Rbm26 | 2.34E-16 | 1.853336 | 0.696 | 0.651 | 5.52E-12 |
| Kansl1l | 2.39E-16 | -0.75602 | 0.072 | 0.241 | 5.65E-12 |
| Zfp715 | 2.5E-16 | -0.66483 | 0.077 | 0.251 | 5.89E-12 |

| | | | | | |
|---|---|---|---|---|---|
| Gcn1l1 | 2.76E-16 | -0.5156 | 0.222 | 0.476 | 6.51E-12 |
| Gm9616 | 2.91E-16 | 1.700651 | 0.493 | 0.287 | 6.86E-12 |
| Rps2 | 3.51E-16 | 2.08979 | 0.697 | 0.73 | 8.27E-12 |
| Rps29 | 3.78E-16 | 1.038154 | 0.968 | 0.99 | 8.91E-12 |
| Selenof | 4.1E-16 | 1.806307 | 0.68 | 0.736 | 9.67E-12 |
| Gm3699 | 4.37E-16 | 1.337253 | 0.436 | 0.208 | 1.03E-11 |
| Gm5138 | 4.41E-16 | 2.806632 | 0.34 | 0.124 | 1.04E-11 |
| Gm17241 | 5.96E-16 | 1.097693 | 0.378 | 0.15 | 1.41E-11 |
| Dcaf12 | 6.32E-16 | -0.80298 | 0.168 | 0.378 | 1.49E-11 |
| Gm15772 | 6.83E-16 | 1.605647 | 0.454 | 0.248 | 1.61E-11 |
| Gm12191 | 6.99E-16 | 0.832259 | 0.347 | 0.117 | 1.65E-11 |
| Tmem245 | 7.44E-16 | -0.49103 | 0.133 | 0.342 | 1.76E-11 |
| Ppp1ca | 7.79E-16 | 1.851801 | 0.669 | 0.694 | 1.84E-11 |
| Atmin | 7.96E-16 | -0.37186 | 0.094 | 0.277 | 1.88E-11 |
| Gm10231 | 8.23E-16 | 1.326499 | 0.274 | 0.059 | 1.94E-11 |
| Gm6170 | 8.56E-16 | 0.841993 | 0.408 | 0.173 | 2.02E-11 |
| Mfsd14b | 9.67E-16 | -1.75247 | 0.105 | 0.28 | 2.28E-11 |
| Egr1 | 1.12E-15 | -2.31788 | 0.089 | 0.264 | 2.64E-11 |
| Arfgef1 | 1.18E-15 | -0.70292 | 0.068 | 0.228 | 2.77E-11 |
| Amfr | 1.22E-15 | -1.19969 | 0.285 | 0.495 | 2.88E-11 |
| AU040320 | 1.25E-15 | -0.52397 | 0.096 | 0.277 | 2.95E-11 |
| Vcp-rs | 1.33E-15 | -0.50695 | 0.18 | 0.391 | 3.13E-11 |
| Rpl10a-ps1 | 1.43E-15 | 2.181632 | 0.548 | 0.404 | 3.37E-11 |
| Rpl21-ps15 | 1.51E-15 | 1.642318 | 0.344 | 0.124 | 3.57E-11 |
| Gm42908 | 1.57E-15 | 1.108909 | 0.1 | 0.293 | 3.71E-11 |
| Arsb | 1.57E-15 | -1.10596 | 0.262 | 0.479 | 3.71E-11 |
| Itgb5 | 1.65E-15 | -0.33784 | 0.964 | 1 | 3.89E-11 |
| Gm44667 | 1.66E-15 | -0.98329 | 0.063 | 0.218 | 3.91E-11 |
| Itpripl1 | 1.87E-15 | 2.095094 | 0.7 | 0.746 | 4.42E-11 |
| Gm13493 | 2E-15 | 1.271722 | 0.706 | 0.606 | 4.71E-11 |
| Rpsa-ps9 | 2E-15 | 1.118879 | 0.364 | 0.14 | 4.72E-11 |
| Fbxl3 | 2.41E-15 | -0.80407 | 0.133 | 0.329 | 5.69E-11 |
| Slmap | 2.52E-15 | -0.32402 | 0.148 | 0.358 | 5.94E-11 |
| Sgpl1 | 2.67E-15 | -0.37836 | 0.274 | 0.508 | 6.3E-11 |
| Dusp1 | 3.09E-15 | -1.04948 | 0.148 | 0.371 | 7.3E-11 |
| Marcks | 3.22E-15 | 0.747169 | 0.987 | 1 | 7.6E-11 |
| Cmklr1 | 3.3E-15 | 0.561803 | 0.121 | 0.322 | 7.78E-11 |
| Spcs3 | 3.32E-15 | -1.02411 | 0.181 | 0.388 | 7.83E-11 |
| Gm16192 | 3.38E-15 | 0.350482 | 0.09 | 0.27 | 7.98E-11 |
| Calm1 | 3.43E-15 | 1.883446 | 0.732 | 0.814 | 8.09E-11 |
| Gm10020 | 3.75E-15 | 0.659041 | 0.251 | 0.049 | 8.84E-11 |
| Lyn | 3.86E-15 | -0.50722 | 0.759 | 0.958 | 9.11E-11 |
| Rps15a | 4.56E-15 | 2.062938 | 0.635 | 0.586 | 1.08E-10 |
| Ppp1r3b | 4.65E-15 | 0.333063 | 0.082 | 0.251 | 1.1E-10 |
| H2-T23 | 4.66E-15 | 2.015264 | 0.695 | 0.678 | 1.1E-10 |
| Rps13-ps1 | 5.04E-15 | 0.99359 | 0.357 | 0.15 | 1.19E-10 |
| Ywhae | 6.47E-15 | 1.771441 | 0.77 | 0.837 | 1.53E-10 |

| | | | | | |
|---|---|---|---|---|---|
| Gm10045 | 6.52E-15 | 1.896981 | 0.239 | 0.042 | 1.54E-10 |
| Stat6 | 6.77E-15 | -1.01539 | 0.212 | 0.427 | 1.6E-10 |
| Dpy19l4 | 7.95E-15 | -0.39461 | 0.147 | 0.358 | 1.88E-10 |
| Il10ra | 8.41E-15 | -0.38048 | 0.528 | 0.818 | 1.98E-10 |
| Wdfy3 | 8.44E-15 | -0.2694 | 0.209 | 0.45 | 1.99E-10 |
| Sept2 | 8.62E-15 | 1.72816 | 0.741 | 0.726 | 2.03E-10 |
| Wdr81 | 9.08E-15 | -0.37154 | 0.179 | 0.404 | 2.14E-10 |
| Ltc4s | 9.53E-15 | 1.533685 | 0.771 | 0.902 | 2.25E-10 |
| Pik3r5 | 9.96E-15 | -0.58761 | 0.143 | 0.339 | 2.35E-10 |
| Zfp672 | 1.03E-14 | -1.0319 | 0.083 | 0.248 | 2.43E-10 |
| Ptp4a2 | 1.18E-14 | 1.95986 | 0.73 | 0.749 | 2.78E-10 |
| Erbin | 1.25E-14 | -0.36774 | 0.165 | 0.381 | 2.95E-10 |
| Dock2 | 1.28E-14 | 0.315503 | 0.278 | 0.564 | 3.02E-10 |
| Tmbim6 | 1.47E-14 | 0.926721 | 0.949 | 0.974 | 3.48E-10 |
| Fam193a | 1.6E-14 | 0.402198 | 0.083 | 0.254 | 3.77E-10 |
| Rpl24 | 1.68E-14 | 2.088852 | 0.629 | 0.593 | 3.95E-10 |
| Etv5 | 1.74E-14 | -0.64803 | 0.259 | 0.476 | 4.1E-10 |
| Pik3cd | 1.76E-14 | 0.281234 | 0.234 | 0.505 | 4.15E-10 |
| Coro1a | 2.24E-14 | 1.283372 | 0.779 | 0.889 | 5.29E-10 |
| Gm45472 | 2.31E-14 | -0.5479 | 0.161 | 0.362 | 5.46E-10 |
| Setdb1 | 2.34E-14 | -0.6358 | 0.098 | 0.274 | 5.52E-10 |
| Mtpn | 2.4E-14 | -0.92474 | 0.271 | 0.489 | 5.67E-10 |
| Arap1 | 2.53E-14 | -0.30951 | 0.179 | 0.391 | 5.97E-10 |
| Kbtbd2 | 2.58E-14 | -0.64964 | 0.152 | 0.349 | 6.09E-10 |
| Tmem64 | 2.68E-14 | -1.73251 | 0.105 | 0.27 | 6.32E-10 |
| Ttyh3 | 2.77E-14 | -0.25149 | 0.159 | 0.381 | 6.53E-10 |
| Ube2q1 | 2.97E-14 | 0.632676 | 0.123 | 0.313 | 7E-10 |
| Ttpal | 3.05E-14 | -0.59315 | 0.062 | 0.212 | 7.2E-10 |
| Pfkfb3 | 3.09E-14 | -0.34607 | 0.131 | 0.326 | 7.3E-10 |
| Sp3 | 3.14E-14 | -0.87191 | 0.17 | 0.378 | 7.4E-10 |
| Rpl10-ps3 | 3.15E-14 | 1.823615 | 0.667 | 0.485 | 7.44E-10 |
| Gpr84 | 3.36E-14 | 2.780356 | 0.518 | 0.397 | 7.93E-10 |
| Sppl3 | 3.65E-14 | -0.4105 | 0.295 | 0.55 | 8.61E-10 |
| Rpl13a-ps1 | 3.84E-14 | 0.903667 | 0.203 | 0.023 | 9.06E-10 |
| Cd52 | 3.99E-14 | 2.582571 | 0.561 | 0.495 | 9.41E-10 |
| Atp1a1 | 4.05E-14 | -0.93356 | 0.365 | 0.57 | 9.55E-10 |
| Pde3b | 4.05E-14 | -0.33377 | 0.595 | 0.866 | 9.56E-10 |
| Arhgef6 | 4.14E-14 | -0.47861 | 0.158 | 0.362 | 9.77E-10 |
| Wdr26 | 4.15E-14 | 0.413853 | 0.266 | 0.541 | 9.79E-10 |
| Arl6ip1 | 4.39E-14 | 1.521399 | 0.719 | 0.736 | 1.04E-09 |
| Atp6v1g1 | 4.66E-14 | 1.853316 | 0.644 | 0.635 | 1.1E-09 |
| Rpl10a | 5.95E-14 | 2.247949 | 0.673 | 0.661 | 1.4E-09 |
| Gm10689 | 6.31E-14 | 0.512551 | 0.223 | 0.036 | 1.49E-09 |
| Gm8129 | 6.83E-14 | 0.844911 | 0.279 | 0.081 | 1.61E-09 |
| Sidt2 | 7.12E-14 | -0.73742 | 0.142 | 0.329 | 1.68E-09 |
| Zfp24 | 7.95E-14 | -0.30907 | 0.096 | 0.264 | 1.88E-09 |
| Prpf4b | 8.03E-14 | 0.383645 | 0.237 | 0.479 | 1.89E-09 |

| | | | | | |
|---|---|---|---|---|---|
| Pacsin2 | 8.11E-14 | -0.61926 | 0.317 | 0.56 | 1.91E-09 |
| Gm7392 | 8.3E-14 | 0.732146 | 0.227 | 0.042 | 1.96E-09 |
| Rpl15 | 8.39E-14 | 1.547667 | 0.708 | 0.694 | 1.98E-09 |
| Gm10132 | 8.74E-14 | 2.046042 | 0.609 | 0.547 | 2.06E-09 |
| Crk | 9.39E-14 | -0.96217 | 0.183 | 0.388 | 2.21E-09 |
| Rpl38 | 9.49E-14 | 1.445784 | 0.685 | 0.658 | 2.24E-09 |
| Zdhhc9 | 1.02E-13 | -0.86871 | 0.129 | 0.309 | 2.4E-09 |
| Inpp4b | 1.04E-13 | -0.50644 | 0.137 | 0.329 | 2.45E-09 |
| Gm8822 | 1.05E-13 | -0.3759 | 0.16 | 0.358 | 2.48E-09 |
| Myo18a | 1.1E-13 | 0.877121 | 0.1 | 0.28 | 2.6E-09 |
| Ssr4 | 1.11E-13 | 1.742821 | 0.662 | 0.697 | 2.63E-09 |
| Gm10275 | 1.15E-13 | 1.990369 | 0.384 | 0.192 | 2.71E-09 |
| Rere | 1.16E-13 | 0.426377 | 0.208 | 0.459 | 2.74E-09 |
| Gm17511 | 1.17E-13 | 0.567843 | 0.226 | 0.042 | 2.77E-09 |
| Rpl8 | 1.19E-13 | 2.01402 | 0.686 | 0.746 | 2.8E-09 |
| Abcd3 | 1.35E-13 | -0.60122 | 0.127 | 0.309 | 3.19E-09 |
| Sec31a | 1.37E-13 | -0.32471 | 0.158 | 0.358 | 3.24E-09 |
| Abcd2 | 1.38E-13 | -0.28719 | 0.234 | 0.463 | 3.25E-09 |
| Sod1 | 1.44E-13 | 2.361647 | 0.662 | 0.658 | 3.39E-09 |
| Chd6 | 1.46E-13 | 0.292396 | 0.11 | 0.293 | 3.45E-09 |
| Lpcat2 | 1.48E-13 | 0.753177 | 0.959 | 0.99 | 3.49E-09 |
| Eif5a | 1.55E-13 | 2.103134 | 0.638 | 0.651 | 3.67E-09 |
| Ivns1abp | 1.61E-13 | -0.6563 | 0.658 | 0.86 | 3.81E-09 |
| Gm11539 | 1.76E-13 | 1.14326 | 0.347 | 0.14 | 4.15E-09 |
| Timp2 | 1.81E-13 | 1.234367 | 0.805 | 0.896 | 4.27E-09 |
| Ash1l | 1.87E-13 | 0.431722 | 0.191 | 0.417 | 4.4E-09 |
| Aff1 | 2.17E-13 | -0.95462 | 0.081 | 0.235 | 5.12E-09 |
| Gtf2i | 2.55E-13 | 0.398451 | 0.273 | 0.537 | 6.02E-09 |
| Chtf8 | 2.56E-13 | -1.47785 | 0.127 | 0.296 | 6.03E-09 |
| Eno1 | 2.59E-13 | 1.633579 | 0.729 | 0.834 | 6.11E-09 |
| Capza1 | 2.82E-13 | -0.60365 | 0.337 | 0.557 | 6.64E-09 |
| Irf2bp2 | 3.07E-13 | 0.620135 | 0.259 | 0.508 | 7.25E-09 |
| Numa1 | 3.37E-13 | -0.32657 | 0.224 | 0.44 | 7.95E-09 |
| Arpc3 | 3.38E-13 | 2.167378 | 0.626 | 0.668 | 7.96E-09 |
| Crot | 3.57E-13 | -0.7673 | 0.13 | 0.313 | 8.41E-09 |
| Ube2h | 3.68E-13 | 0.279515 | 0.252 | 0.508 | 8.67E-09 |
| Ppp1r15b | 4.1E-13 | -0.95544 | 0.091 | 0.248 | 9.66E-09 |
| Rbm39 | 4.13E-13 | 1.34843 | 0.83 | 0.906 | 9.74E-09 |
| Olfr620 | 4.14E-13 | 0.708653 | 0.24 | 0.059 | 9.76E-09 |
| Gm11263 | 4.21E-13 | 0.645148 | 0.308 | 0.111 | 9.92E-09 |
| Gpd1l | 4.3E-13 | -0.44241 | 0.065 | 0.205 | 1.01E-08 |
| Rpl35a | 4.54E-13 | 1.48328 | 0.709 | 0.779 | 1.07E-08 |
| Gm34280 | 4.56E-13 | 1.171283 | 0.104 | 0.283 | 1.08E-08 |
| Glg1 | 5.32E-13 | -0.28715 | 0.233 | 0.456 | 1.25E-08 |
| Rabac1 | 5.39E-13 | 1.637989 | 0.712 | 0.831 | 1.27E-08 |
| Strn3 | 5.85E-13 | -0.56744 | 0.179 | 0.388 | 1.38E-08 |
| Hnrnpl | 6.06E-13 | 1.907324 | 0.331 | 0.567 | 1.43E-08 |

| | | | | | |
|---|---|---|---|---|---|
| Hnrnpu | 6.77E-13 | -0.27384 | 0.311 | 0.547 | 1.6E-08 |
| Rps2-ps10 | 6.85E-13 | 1.229096 | 0.295 | 0.107 | 1.62E-08 |
| Tmem131l | 7.07E-13 | 0.254504 | 0.08 | 0.235 | 1.67E-08 |
| Cmtm7 | 7.35E-13 | 1.523878 | 0.728 | 0.831 | 1.73E-08 |
| Nbr1 | 8.82E-13 | -0.54472 | 0.132 | 0.313 | 2.08E-08 |
| Trf | 9.03E-13 | 0.781274 | 0.968 | 0.987 | 2.13E-08 |
| Uba1 | 1.01E-12 | -0.5036 | 0.165 | 0.362 | 2.37E-08 |
| Larp4 | 1.01E-12 | -0.54965 | 0.1 | 0.261 | 2.37E-08 |
| Dad1 | 1.09E-12 | 1.886354 | 0.641 | 0.697 | 2.56E-08 |
| Pald1 | 1.13E-12 | -0.39899 | 0.277 | 0.495 | 2.67E-08 |
| Pfdn5 | 1.19E-12 | 2.053913 | 0.599 | 0.593 | 2.81E-08 |
| Saraf | 1.2E-12 | 1.315997 | 0.723 | 0.795 | 2.82E-08 |
| Bank1 | 1.25E-12 | -0.4621 | 0.071 | 0.215 | 2.95E-08 |
| Ankfy1 | 1.26E-12 | -0.35369 | 0.135 | 0.316 | 2.98E-08 |
| Rack1 | 1.29E-12 | 2.188649 | 0.674 | 0.697 | 3.03E-08 |
| Rps3a3 | 1.32E-12 | 0.738326 | 0.405 | 0.208 | 3.11E-08 |
| Gm15501 | 1.36E-12 | 1.603664 | 0.469 | 0.287 | 3.22E-08 |
| Ubqln1 | 1.39E-12 | -0.57074 | 0.196 | 0.397 | 3.28E-08 |
| Cep170 | 1.51E-12 | -0.35313 | 0.193 | 0.401 | 3.57E-08 |
| Rps15a-ps6 | 1.55E-12 | 1.148833 | 0.301 | 0.107 | 3.66E-08 |
| Gm12286 | 1.6E-12 | 0.768484 | 0.211 | 0.039 | 3.78E-08 |
| Rps24-ps3 | 1.61E-12 | 1.19999 | 0.296 | 0.104 | 3.79E-08 |
| Gm4294 | 1.7E-12 | 0.710767 | 0.407 | 0.205 | 4E-08 |
| Man2b1 | 1.9E-12 | -0.34991 | 0.885 | 0.961 | 4.49E-08 |
| Large1 | 1.93E-12 | 0.695659 | 0.075 | 0.225 | 4.55E-08 |
| Fam120b | 1.93E-12 | -0.40282 | 0.19 | 0.397 | 4.55E-08 |
| Stxbp5 | 1.93E-12 | -0.59424 | 0.093 | 0.251 | 4.55E-08 |
| Gm8172 | 1.99E-12 | 1.945374 | 0.455 | 0.303 | 4.7E-08 |
| Csde1 | 2.14E-12 | -0.37238 | 0.28 | 0.518 | 5.05E-08 |
| Gm8226 | 2.17E-12 | 0.853652 | 0.227 | 0.052 | 5.12E-08 |
| Nisch | 2.26E-12 | 0.466756 | 0.327 | 0.59 | 5.32E-08 |
| Slc12a2 | 2.31E-12 | -0.77937 | 0.176 | 0.365 | 5.44E-08 |
| Sgpp1 | 2.31E-12 | -1.68997 | 0.091 | 0.235 | 5.45E-08 |
| Wipf1 | 2.34E-12 | 0.439317 | 0.171 | 0.371 | 5.53E-08 |
| Mbtps1 | 2.35E-12 | -1.17271 | 0.078 | 0.218 | 5.53E-08 |
| Rtn4 | 2.63E-12 | 0.67656 | 0.456 | 0.723 | 6.21E-08 |
| Rpl14-ps1 | 2.66E-12 | 1.809091 | 0.58 | 0.466 | 6.26E-08 |
| Sult1a1 | 2.67E-12 | 0.273033 | 0.204 | 0.443 | 6.3E-08 |
| Gm4617 | 2.76E-12 | 1.243265 | 0.306 | 0.124 | 6.52E-08 |
| Scaf8 | 3.02E-12 | 0.68747 | 0.096 | 0.261 | 7.12E-08 |
| Usp47 | 3.04E-12 | -0.85443 | 0.066 | 0.202 | 7.17E-08 |
| Gm7331 | 3.49E-12 | 1.760405 | 0.535 | 0.43 | 8.22E-08 |
| Rnf139 | 3.75E-12 | -1.32073 | 0.153 | 0.322 | 8.85E-08 |
| B4galt1 | 3.78E-12 | 1.17545 | 0.865 | 0.866 | 8.92E-08 |
| Arhgap39 | 3.81E-12 | 0.291263 | 0.263 | 0.505 | 8.98E-08 |
| Ttc13 | 4.12E-12 | -1.30061 | 0.113 | 0.27 | 9.72E-08 |
| H2-DMb1 | 4.34E-12 | 2.257557 | 0.508 | 0.433 | 1.02E-07 |

| | | | | | |
|---|---|---|---|---|---|
| Arid3b | 4.56E-12 | 0.563798 | 0.128 | 0.306 | 1.08E-07 |
| Dock8 | 4.61E-12 | 0.389005 | 0.353 | 0.632 | 1.09E-07 |
| Gas5 | 4.64E-12 | 2.928467 | 0.416 | 0.251 | 1.09E-07 |
| Far1 | 4.84E-12 | -0.39181 | 0.141 | 0.319 | 1.14E-07 |
| Arf1 | 5.08E-12 | 1.548021 | 0.723 | 0.85 | 1.2E-07 |
| Rps27a-ps2 | 5.44E-12 | 1.687593 | 0.333 | 0.153 | 1.28E-07 |
| mt-Nd2 | 5.51E-12 | 1.850589 | 0.84 | 0.909 | 1.3E-07 |
| Mctp1 | 6.07E-12 | -0.65761 | 0.068 | 0.202 | 1.43E-07 |
| Ddost | 6.94E-12 | 1.695986 | 0.675 | 0.726 | 1.64E-07 |
| Hmgb1 | 6.95E-12 | 1.65919 | 0.729 | 0.814 | 1.64E-07 |
| Ep300 | 7.01E-12 | -0.25677 | 0.252 | 0.479 | 1.65E-07 |
| Gm10224 | 7.4E-12 | 0.689668 | 0.211 | 0.046 | 1.75E-07 |
| Gm37954 | 8.73E-12 | 0.714244 | 0.138 | 0.329 | 2.06E-07 |
| Txndc11 | 8.89E-12 | -0.63625 | 0.179 | 0.371 | 2.1E-07 |
| Rpl28-ps1 | 9.93E-12 | 2.218997 | 0.422 | 0.27 | 2.34E-07 |
| Prdm2 | 1.01E-11 | 0.629032 | 0.096 | 0.257 | 2.38E-07 |
| Chka | 1.04E-11 | -0.52903 | 0.124 | 0.296 | 2.44E-07 |
| Arhgef40 | 1.14E-11 | 0.50949 | 0.251 | 0.495 | 2.69E-07 |
| Tpt1-ps3 | 1.26E-11 | 1.356177 | 0.616 | 0.511 | 2.98E-07 |
| Notch2 | 1.3E-11 | 1.018852 | 0.699 | 0.625 | 3.06E-07 |
| Emc1 | 1.31E-11 | -0.25332 | 0.071 | 0.205 | 3.09E-07 |
| Rsf1 | 1.31E-11 | 0.576168 | 0.119 | 0.29 | 3.1E-07 |
| Arih1 | 1.32E-11 | -0.42711 | 0.077 | 0.215 | 3.12E-07 |
| Nipbl | 1.33E-11 | 0.670987 | 0.272 | 0.534 | 3.13E-07 |
| Cpsf1 | 1.38E-11 | -0.79104 | 0.09 | 0.235 | 3.26E-07 |
| Snhg20 | 1.41E-11 | 2.663734 | 0.34 | 0.169 | 3.32E-07 |
| Ythdf3 | 1.45E-11 | -1.53985 | 0.084 | 0.221 | 3.42E-07 |
| Rlim | 1.54E-11 | -0.27146 | 0.222 | 0.423 | 3.64E-07 |
| Ifnar1 | 1.6E-11 | -0.67812 | 0.152 | 0.326 | 3.78E-07 |
| Lair1 | 1.61E-11 | 1.062429 | 0.858 | 0.958 | 3.79E-07 |
| Gm5453 | 1.69E-11 | 0.92278 | 0.232 | 0.062 | 3.97E-07 |
| C2cd3 | 1.74E-11 | -0.43873 | 0.072 | 0.208 | 4.1E-07 |
| Rps3a2 | 2.03E-11 | 1.213674 | 0.383 | 0.199 | 4.8E-07 |
| Rpl30-ps10 | 2.07E-11 | 0.79962 | 0.252 | 0.081 | 4.89E-07 |
| Rpl35a-ps2 | 2.09E-11 | 0.714603 | 0.222 | 0.055 | 4.93E-07 |
| Cse1l | 2.1E-11 | -0.64605 | 0.086 | 0.228 | 4.94E-07 |
| Bag6 | 2.22E-11 | 1.075799 | 0.113 | 0.283 | 5.23E-07 |
| Lgals3bp | 2.29E-11 | 2.419248 | 0.582 | 0.541 | 5.41E-07 |
| Eif4g3 | 2.46E-11 | 0.495241 | 0.254 | 0.502 | 5.8E-07 |
| Marf1 | 2.46E-11 | 2.119076 | 0.625 | 0.554 | 5.8E-07 |
| Dgkd | 2.55E-11 | 0.49416 | 0.149 | 0.339 | 6.01E-07 |
| Gna12 | 2.56E-11 | -0.32992 | 0.404 | 0.645 | 6.03E-07 |
| 6820431F2 | 2.59E-11 | 0.712684 | 0.142 | 0.329 | 6.11E-07 |
| Upk1b | 2.72E-11 | 2.598007 | 0.425 | 0.277 | 6.41E-07 |
| Shoc2 | 2.76E-11 | -0.71389 | 0.079 | 0.215 | 6.51E-07 |
| Zmiz1 | 2.87E-11 | 0.421452 | 0.426 | 0.71 | 6.77E-07 |
| Fbxl20 | 2.92E-11 | 0.437408 | 0.1 | 0.257 | 6.89E-07 |

| | | | | | |
|---|---|---|---|---|---|
| Cd53 | 2.92E-11 | 0.673961 | 0.958 | 0.997 | 6.9E-07 |
| Myo7a | 3.06E-11 | 0.390885 | 0.079 | 0.218 | 7.22E-07 |
| Spcs2 | 3.1E-11 | 1.569723 | 0.679 | 0.769 | 7.3E-07 |
| Cnbp | 3.14E-11 | 1.415855 | 0.732 | 0.844 | 7.42E-07 |
| Tns3 | 3.27E-11 | -0.34252 | 0.178 | 0.375 | 7.72E-07 |
| Gng5 | 3.28E-11 | 1.470138 | 0.713 | 0.782 | 7.73E-07 |
| Gna15 | 3.29E-11 | 2.006491 | 0.635 | 0.651 | 7.76E-07 |
| Gpbp1 | 3.32E-11 | -0.60557 | 0.106 | 0.254 | 7.82E-07 |
| Mrfap1 | 3.59E-11 | 1.300017 | 0.719 | 0.769 | 8.47E-07 |
| Plxna4 | 3.61E-11 | -0.72799 | 0.094 | 0.235 | 8.51E-07 |
| Mapkapk2 | 3.65E-11 | -0.46317 | 0.179 | 0.365 | 8.6E-07 |
| Kmt2a | 3.99E-11 | 0.951613 | 0.16 | 0.352 | 9.41E-07 |
| Rpl22 | 4E-11 | 2.319478 | 0.563 | 0.511 | 9.44E-07 |
| Ppp6r1 | 4.1E-11 | -0.39308 | 0.125 | 0.287 | 9.67E-07 |
| Gaa | 4.25E-11 | -0.57852 | 0.288 | 0.489 | 1E-06 |
| Gm10157 | 4.33E-11 | 0.732396 | 0.236 | 0.068 | 1.02E-06 |
| Apba1 | 4.76E-11 | 0.510994 | 0.091 | 0.241 | 1.12E-06 |
| Synrg | 5.49E-11 | 0.486695 | 0.089 | 0.235 | 1.3E-06 |
| Ints6l | 5.91E-11 | -0.3386 | 0.199 | 0.384 | 1.39E-06 |
| Gm3150 | 6.13E-11 | 0.639361 | 0.206 | 0.049 | 1.45E-06 |
| Ptma | 6.28E-11 | 2.592464 | 0.667 | 0.746 | 1.48E-06 |
| Slamf9 | 6.71E-11 | 2.618743 | 0.427 | 0.3 | 1.58E-06 |
| Cnot2 | 6.92E-11 | -0.34349 | 0.218 | 0.417 | 1.63E-06 |
| Gmfb | 7.17E-11 | -0.44733 | 0.109 | 0.261 | 1.69E-06 |
| Cap1 | 7.17E-11 | 0.356024 | 0.359 | 0.606 | 1.69E-06 |
| Slc16a7 | 7.22E-11 | -1.13353 | 0.084 | 0.218 | 1.7E-06 |
| Tmpo | 7.48E-11 | -0.35735 | 0.12 | 0.277 | 1.77E-06 |
| Sec63 | 7.66E-11 | 0.29293 | 0.144 | 0.319 | 1.81E-06 |
| Canx | 8.15E-11 | 1.046515 | 0.906 | 0.977 | 1.92E-06 |
| Rffl | 9.37E-11 | -0.56706 | 0.106 | 0.254 | 2.21E-06 |
| Copb2 | 9.49E-11 | -0.61102 | 0.155 | 0.316 | 2.24E-06 |
| Tnfsf13b | 9.54E-11 | 3.35651 | 0.259 | 0.098 | 2.25E-06 |
| Selenok | 1.04E-10 | 1.893628 | 0.621 | 0.661 | 2.46E-06 |
| Rab5a | 1.08E-10 | -0.46548 | 0.196 | 0.381 | 2.54E-06 |
| Pdcd6ip | 1.12E-10 | -0.77702 | 0.2 | 0.375 | 2.63E-06 |
| Abl2 | 1.16E-10 | -0.36037 | 0.092 | 0.228 | 2.73E-06 |
| Mybbp1a | 1.17E-10 | -0.8682 | 0.105 | 0.251 | 2.75E-06 |
| Slc39a1 | 1.18E-10 | 2.147467 | 0.619 | 0.573 | 2.79E-06 |
| Hs6st1 | 1.26E-10 | 0.73217 | 0.169 | 0.358 | 2.97E-06 |
| Gm9385 | 1.27E-10 | 1.126553 | 0.33 | 0.16 | 3E-06 |
| Rnf103 | 1.35E-10 | -1.08127 | 0.14 | 0.287 | 3.18E-06 |
| Tnpo1 | 1.42E-10 | -0.8317 | 0.203 | 0.397 | 3.34E-06 |
| Grcc10 | 1.64E-10 | 2.26208 | 0.604 | 0.648 | 3.86E-06 |
| Zfand3 | 1.65E-10 | 1.956803 | 0.093 | 0.241 | 3.89E-06 |
| Gm40841 | 1.66E-10 | 0.413019 | 0.078 | 0.215 | 3.91E-06 |
| Dync1h1 | 1.69E-10 | 0.44552 | 0.16 | 0.345 | 4E-06 |
| Gm17066 | 1.81E-10 | 1.88181 | 0.569 | 0.511 | 4.28E-06 |

| | | | | | |
|---|---|---|---|---|---|
| Olfr77 | 1.84E-10 | 1.29238 | 0.32 | 0.153 | 4.35E-06 |
| 9930021J0: | 2.03E-10 | 0.366216 | 0.093 | 0.238 | 4.79E-06 |
| Bach2 | 2.06E-10 | 0.459016 | 0.087 | 0.228 | 4.86E-06 |
| Kdm1a | 2.38E-10 | -0.38513 | 0.112 | 0.261 | 5.61E-06 |
| Tnks | 2.39E-10 | -0.58994 | 0.074 | 0.202 | 5.63E-06 |
| Tmem123 | 2.41E-10 | -1.28662 | 0.129 | 0.274 | 5.68E-06 |
| Atp6ap1 | 2.47E-10 | 1.141051 | 0.771 | 0.821 | 5.82E-06 |
| Serf2 | 2.48E-10 | 1.767436 | 0.731 | 0.795 | 5.86E-06 |
| Vamp8 | 2.56E-10 | 1.96774 | 0.593 | 0.625 | 6.03E-06 |
| Fbxl5 | 2.64E-10 | -1.02383 | 0.11 | 0.257 | 6.22E-06 |
| Tor4a | 2.86E-10 | -0.72924 | 0.176 | 0.339 | 6.75E-06 |
| Elmo2 | 2.95E-10 | -0.62973 | 0.107 | 0.251 | 6.97E-06 |
| Srsf9 | 3.02E-10 | 2.112676 | 0.592 | 0.645 | 7.13E-06 |
| Chd2 | 3.04E-10 | 0.958516 | 0.158 | 0.345 | 7.16E-06 |
| Anapc1 | 3.1E-10 | -1.35276 | 0.083 | 0.212 | 7.32E-06 |
| Ppp3r1 | 3.22E-10 | -0.82424 | 0.142 | 0.29 | 7.59E-06 |
| Ctnnb1 | 3.57E-10 | -0.66087 | 0.223 | 0.388 | 8.43E-06 |
| Pacs2 | 3.7E-10 | 0.301417 | 0.134 | 0.3 | 8.72E-06 |
| Prpf8 | 3.72E-10 | 0.348505 | 0.147 | 0.322 | 8.78E-06 |
| Gm11560 | 3.96E-10 | 2.083439 | 0.294 | 0.14 | 9.33E-06 |
| Rhot1 | 4.15E-10 | -1.02144 | 0.136 | 0.287 | 9.78E-06 |
| Snn | 4.28E-10 | 1.673404 | 0.18 | 0.375 | 1.01E-05 |
| Psmb8 | 4.31E-10 | 1.877468 | 0.591 | 0.599 | 1.02E-05 |
| Polr2a | 4.41E-10 | 0.976763 | 0.237 | 0.479 | 1.04E-05 |
| Hk3 | 4.43E-10 | -0.31103 | 0.181 | 0.355 | 1.04E-05 |
| Ap1g1 | 4.5E-10 | 0.604532 | 0.096 | 0.238 | 1.06E-05 |
| Gm9800 | 4.74E-10 | 1.152524 | 0.271 | 0.107 | 1.12E-05 |
| Gm32036 | 4.95E-10 | 2.586234 | 0.489 | 0.414 | 1.17E-05 |
| Rin3 | 5.09E-10 | 0.674449 | 0.192 | 0.375 | 1.2E-05 |
| Plagl2 | 5.17E-10 | 0.505882 | 0.128 | 0.29 | 1.22E-05 |
| Mvb12b | 5.2E-10 | 0.969147 | 0.162 | 0.345 | 1.23E-05 |
| Hspa13 | 5.21E-10 | 0.627298 | 0.104 | 0.254 | 1.23E-05 |
| Ptk2b | 5.27E-10 | -0.368 | 0.178 | 0.349 | 1.24E-05 |
| Anp32a | 5.38E-10 | 0.883751 | 0.283 | 0.554 | 1.27E-05 |
| Zyg11b | 5.42E-10 | -0.53761 | 0.105 | 0.248 | 1.28E-05 |
| C130089K0 | 5.5E-10 | 0.340668 | 0.193 | 0.381 | 1.3E-05 |
| Gm13408 | 5.6E-10 | 0.467101 | 0.255 | 0.091 | 1.32E-05 |
| Tlr6 | 5.69E-10 | -0.65664 | 0.116 | 0.261 | 1.34E-05 |
| Bcor | 5.71E-10 | -0.43608 | 0.083 | 0.215 | 1.35E-05 |
| Rcc2 | 6.08E-10 | 0.456361 | 0.174 | 0.362 | 1.43E-05 |
| Wdr37 | 6.22E-10 | 0.314799 | 0.093 | 0.231 | 1.47E-05 |
| Gm10036 | 6.22E-10 | 1.361113 | 0.35 | 0.195 | 1.47E-05 |
| Dock4 | 6.24E-10 | 0.662501 | 0.442 | 0.765 | 1.47E-05 |
| Reep5 | 6.4E-10 | 1.764249 | 0.656 | 0.717 | 1.51E-05 |
| Lrrc58 | 6.54E-10 | -0.56245 | 0.287 | 0.463 | 1.54E-05 |
| Ndst2 | 6.79E-10 | -0.65114 | 0.125 | 0.274 | 1.6E-05 |
| N4bp1 | 7.08E-10 | 0.386507 | 0.15 | 0.316 | 1.67E-05 |

| | | | | | |
|---|---|---|---|---|---|
| Stard9 | 7.43E-10 | 0.370241 | 0.163 | 0.339 | 1.75E-05 |
| Rps15 | 7.8E-10 | 1.753375 | 0.697 | 0.779 | 1.84E-05 |
| Gm5835 | 8.29E-10 | 1.939848 | 0.375 | 0.221 | 1.96E-05 |
| Rapgef6 | 8.39E-10 | 0.444077 | 0.184 | 0.375 | 1.98E-05 |
| Tgoln1 | 8.43E-10 | 0.416609 | 0.281 | 0.502 | 1.99E-05 |
| Ppm1b | 8.79E-10 | -1.03425 | 0.083 | 0.208 | 2.07E-05 |
| Ppp3cb | 9.09E-10 | 0.585085 | 0.093 | 0.225 | 2.15E-05 |
| Tmed10 | 9.29E-10 | 1.515308 | 0.719 | 0.883 | 2.19E-05 |
| Map3k11 | 9.4E-10 | 0.306809 | 0.137 | 0.296 | 2.22E-05 |
| Bach1 | 9.45E-10 | -0.50388 | 0.19 | 0.358 | 2.23E-05 |
| Fam117b | 9.65E-10 | 0.293957 | 0.168 | 0.352 | 2.28E-05 |
| Atp9b | 9.8E-10 | -0.5739 | 0.093 | 0.225 | 2.31E-05 |
| Ptgs1 | 9.84E-10 | 1.008121 | 0.891 | 0.958 | 2.32E-05 |
| Gm5637 | 9.92E-10 | 1.590103 | 0.325 | 0.163 | 2.34E-05 |
| Nxt2 | 1.01E-09 | -2.10288 | 0.084 | 0.205 | 2.39E-05 |
| Chmp4b | 1.05E-09 | 1.277926 | 0.083 | 0.215 | 2.48E-05 |
| Tnrc6b | 1.05E-09 | 0.79457 | 0.214 | 0.417 | 2.49E-05 |
| Slc33a1 | 1.08E-09 | -1.33216 | 0.086 | 0.208 | 2.55E-05 |
| Prdx1 | 1.09E-09 | 1.833045 | 0.573 | 0.57 | 2.58E-05 |
| C5ar2 | 1.1E-09 | 1.96426 | 0.614 | 0.564 | 2.59E-05 |
| Hnrnpa3 | 1.13E-09 | 0.352732 | 0.173 | 0.349 | 2.67E-05 |
| Mtss1 | 1.14E-09 | 0.277947 | 0.178 | 0.358 | 2.68E-05 |
| Glyr1 | 1.15E-09 | -0.70779 | 0.175 | 0.339 | 2.72E-05 |
| Rasal3 | 1.17E-09 | -0.33836 | 0.375 | 0.58 | 2.77E-05 |
| Mdm2 | 1.18E-09 | -0.75188 | 0.115 | 0.257 | 2.78E-05 |
| Jak1 | 1.19E-09 | 0.466324 | 0.371 | 0.655 | 2.8E-05 |
| Smarca5 | 1.24E-09 | 0.519021 | 0.208 | 0.404 | 2.93E-05 |
| Abcg2 | 1.38E-09 | -0.99394 | 0.115 | 0.251 | 3.26E-05 |
| Rbbp6 | 1.39E-09 | 0.318185 | 0.224 | 0.43 | 3.27E-05 |
| Zc3h14 | 1.39E-09 | -0.34899 | 0.118 | 0.261 | 3.27E-05 |
| Rpl36a | 1.46E-09 | 1.961616 | 0.539 | 0.495 | 3.44E-05 |
| Itfg1 | 1.51E-09 | -0.58817 | 0.252 | 0.427 | 3.56E-05 |
| Gm12396 | 1.54E-09 | 1.404271 | 0.351 | 0.208 | 3.63E-05 |
| Eno1b | 1.58E-09 | 2.328372 | 0.608 | 0.593 | 3.72E-05 |
| Pag1 | 1.64E-09 | 0.285132 | 0.553 | 0.853 | 3.87E-05 |
| Cx3cr1 | 1.65E-09 | -0.36573 | 0.999 | 1 | 3.88E-05 |
| Myl12a | 1.65E-09 | 2.191588 | 0.518 | 0.469 | 3.89E-05 |
| Gm10847 | 1.74E-09 | 3.31064 | 0.306 | 0.163 | 4.1E-05 |
| Pnpla8 | 1.78E-09 | 0.250243 | 0.151 | 0.322 | 4.19E-05 |
| Hipk2 | 1.79E-09 | 0.358353 | 0.096 | 0.235 | 4.22E-05 |
| Gpr146 | 1.9E-09 | 0.528338 | 0.286 | 0.505 | 4.48E-05 |
| Add1 | 1.93E-09 | 0.325746 | 0.288 | 0.495 | 4.56E-05 |
| Nup88 | 1.97E-09 | 0.353084 | 0.205 | 0.401 | 4.65E-05 |
| Atp5b | 2.09E-09 | 1.254857 | 0.734 | 0.85 | 4.94E-05 |
| Hsp90ab1 | 2.18E-09 | 1.07992 | 0.82 | 0.948 | 5.13E-05 |
| Ftl1-ps2 | 2.3E-09 | 0.616741 | 0.331 | 0.179 | 5.43E-05 |
| Ac114990.3 | 2.49E-09 | 1.404297 | 0.245 | 0.104 | 5.87E-05 |

| | | | | | |
|---|---|---|---|---|---|
| Zbtb1 | 2.51E-09 | 0.276902 | 0.149 | 0.313 | 5.92E-05 |
| Rb1cc1 | 2.52E-09 | -0.76016 | 0.147 | 0.3 | 5.95E-05 |
| Rpl12 | 2.55E-09 | 2.077915 | 0.609 | 0.609 | 6.02E-05 |
| P2ry12 | 2.56E-09 | 0.522623 | 0.99 | 1 | 6.03E-05 |
| Herpud2 | 2.57E-09 | -0.34161 | 0.232 | 0.41 | 6.06E-05 |
| Acap2 | 2.57E-09 | 0.928757 | 0.295 | 0.547 | 6.07E-05 |
| Dhx15 | 2.63E-09 | -1.13677 | 0.165 | 0.322 | 6.2E-05 |
| Tbk1 | 2.66E-09 | -0.90833 | 0.132 | 0.27 | 6.26E-05 |
| Zfp579 | 2.72E-09 | 1.6045 | 0.116 | 0.27 | 6.41E-05 |
| Cog4 | 2.78E-09 | -0.35291 | 0.089 | 0.218 | 6.57E-05 |
| Cdk19 | 2.82E-09 | 0.519532 | 0.101 | 0.235 | 6.64E-05 |
| Gm10051 | 2.84E-09 | 1.373908 | 0.224 | 0.078 | 6.7E-05 |
| Naip2 | 2.86E-09 | 0.625629 | 0.166 | 0.342 | 6.75E-05 |
| Gm34589 | 2.94E-09 | 0.629836 | 0.168 | 0.349 | 6.94E-05 |
| Wdfy4 | 2.96E-09 | 0.342835 | 0.089 | 0.218 | 6.99E-05 |
| Fus | 3.02E-09 | 1.268613 | 0.375 | 0.651 | 7.12E-05 |
| Prex1 | 3.18E-09 | 0.450154 | 0.322 | 0.57 | 7.51E-05 |
| Dync1i2 | 3.56E-09 | 0.372846 | 0.295 | 0.502 | 8.39E-05 |
| Rnf130 | 3.56E-09 | 1.143063 | 0.768 | 0.847 | 8.39E-05 |
| Rps6ka4 | 3.64E-09 | -0.50512 | 0.085 | 0.208 | 8.58E-05 |
| Rps25-ps1 | 3.67E-09 | 0.846573 | 0.338 | 0.179 | 8.67E-05 |
| Hspa9 | 3.74E-09 | -0.543 | 0.173 | 0.329 | 8.82E-05 |
| Cnp | 3.76E-09 | 1.380229 | 0.203 | 0.436 | 8.88E-05 |
| Atf7 | 3.78E-09 | 0.297625 | 0.123 | 0.27 | 8.91E-05 |
| Gm26542 | 3.89E-09 | 1.493367 | 0.112 | 0.257 | 9.18E-05 |
| Cebpa | 3.91E-09 | 0.303716 | 0.452 | 0.72 | 9.23E-05 |
| Setd5 | 3.93E-09 | 0.314513 | 0.251 | 0.44 | 9.26E-05 |
| Rab35 | 3.96E-09 | 0.939643 | 0.156 | 0.316 | 9.35E-05 |
| Mprip | 3.96E-09 | -0.76964 | 0.084 | 0.205 | 9.35E-05 |
| Ilkap | 3.99E-09 | 1.010086 | 0.094 | 0.228 | 9.41E-05 |
| St13 | 4.09E-09 | 1.788693 | 0.663 | 0.694 | 9.65E-05 |
| Shisa5 | 4.1E-09 | 1.908442 | 0.627 | 0.697 | 9.67E-05 |
| Dusp3 | 4.14E-09 | 3.149353 | 0.405 | 0.277 | 9.76E-05 |
| Arid5b | 4.16E-09 | 0.578614 | 0.168 | 0.342 | 9.82E-05 |
| Ptgr2 | 4.25E-09 | 0.551016 | 0.09 | 0.221 | 0.0001 |
| Stt3b | 4.37E-09 | -0.69996 | 0.102 | 0.231 | 0.000103 |
| Mir692-2 | 4.73E-09 | 0.562538 | 0.322 | 0.163 | 0.000112 |
| Btbd1 | 4.77E-09 | 0.284625 | 0.163 | 0.329 | 0.000113 |
| Dpy19l1 | 5E-09 | -0.64353 | 0.144 | 0.296 | 0.000118 |
| Atp8a2 | 5.18E-09 | 0.26334 | 0.271 | 0.489 | 0.000122 |
| Erap1 | 5.32E-09 | -0.91285 | 0.094 | 0.215 | 0.000126 |
| Clk2 | 5.36E-09 | 0.365015 | 0.092 | 0.221 | 0.000126 |
| Nmt1 | 5.36E-09 | 0.615393 | 0.443 | 0.651 | 0.000127 |
| Stt3a | 5.43E-09 | -0.50513 | 0.479 | 0.661 | 0.000128 |
| Atp13a2 | 5.52E-09 | -0.64682 | 0.21 | 0.375 | 0.00013 |
| Rpl39 | 5.54E-09 | 1.362065 | 0.695 | 0.7 | 0.000131 |
| Pmp22 | 5.68E-09 | 1.251715 | 0.746 | 0.834 | 0.000134 |

| | | | | | |
|---|---|---|---|---|---|
| Vapa | 5.99E-09 | 1.677435 | 0.644 | 0.648 | 0.000141 |
| Gm9493 | 6.13E-09 | 0.538052 | 0.239 | 0.091 | 0.000145 |
| Gm43305 | 6.19E-09 | 2.467186 | 0.468 | 0.388 | 0.000146 |
| Pomp | 6.28E-09 | 1.91633 | 0.578 | 0.586 | 0.000148 |
| Gpx4-ps2 | 6.34E-09 | 1.921372 | 0.457 | 0.378 | 0.00015 |
| Golph3 | 6.37E-09 | -1.0337 | 0.1 | 0.225 | 0.00015 |
| Cyth4 | 6.72E-09 | 1.220337 | 0.823 | 0.902 | 0.000159 |
| Wdr1 | 6.96E-09 | -0.60891 | 0.419 | 0.596 | 0.000164 |
| Gab2 | 7.19E-09 | 0.580075 | 0.087 | 0.215 | 0.00017 |
| Bst2 | 7.21E-09 | 2.707033 | 0.472 | 0.41 | 0.00017 |
| Vav1 | 7.23E-09 | -0.25949 | 0.524 | 0.72 | 0.00017 |
| Cox8a | 7.36E-09 | 1.617819 | 0.647 | 0.72 | 0.000174 |
| Ccdc50 | 7.46E-09 | 0.441404 | 0.312 | 0.544 | 0.000176 |
| Abcb4 | 7.71E-09 | -0.28637 | 0.097 | 0.225 | 0.000182 |
| Papd4 | 7.94E-09 | -0.31376 | 0.123 | 0.264 | 0.000187 |
| Ppp2r2d | 8.01E-09 | 0.766632 | 0.128 | 0.283 | 0.000189 |
| Pdk1 | 8.01E-09 | -0.47 | 0.204 | 0.378 | 0.000189 |
| Setd3 | 8.02E-09 | -0.26233 | 0.175 | 0.342 | 0.000189 |
| Bzw1 | 8.26E-09 | -0.30394 | 0.215 | 0.384 | 0.000195 |
| Gm15500 | 8.58E-09 | 1.757052 | 0.593 | 0.505 | 0.000202 |
| Cmip | 8.67E-09 | 1.336093 | 0.184 | 0.371 | 0.000205 |
| A630001G2 | 8.76E-09 | 0.637996 | 0.254 | 0.453 | 0.000207 |
| Mfsd14a | 9.13E-09 | -1.15034 | 0.145 | 0.274 | 0.000215 |
| Ociad1 | 9.25E-09 | 1.791253 | 0.623 | 0.7 | 0.000218 |
| Susd6 | 9.4E-09 | -0.77983 | 0.115 | 0.248 | 0.000222 |
| Uba52-ps | 9.47E-09 | 0.525685 | 0.223 | 0.081 | 0.000223 |
| Rps6ka1 | 9.87E-09 | -0.28418 | 0.669 | 0.821 | 0.000233 |
| Usp2 | 9.94E-09 | 2.044362 | 0.647 | 0.736 | 0.000234 |
| Spred1 | 1E-08 | -0.42852 | 0.112 | 0.244 | 0.000237 |
| Mapk1 | 1.01E-08 | -0.41041 | 0.201 | 0.371 | 0.000239 |
| Snx29 | 1.02E-08 | 0.381474 | 0.16 | 0.322 | 0.00024 |
| Ids | 1.06E-08 | -0.31109 | 0.126 | 0.261 | 0.000251 |
| Dip2b | 1.07E-08 | 0.627288 | 0.44 | 0.717 | 0.000252 |
| Qdpr | 1.08E-08 | 1.028471 | 0.178 | 0.371 | 0.000255 |
| Rnf216 | 1.11E-08 | 0.707183 | 0.198 | 0.378 | 0.000262 |
| Flii | 1.11E-08 | -0.55846 | 0.135 | 0.274 | 0.000262 |
| Slc26a2 | 1.16E-08 | -0.30107 | 0.089 | 0.208 | 0.000274 |
| Gm12435 | 1.2E-08 | -0.85871 | 0.143 | 0.293 | 0.000284 |
| Mllt10 | 1.21E-08 | 0.396077 | 0.118 | 0.257 | 0.000285 |
| Agps | 1.26E-08 | 0.324807 | 0.212 | 0.401 | 0.000297 |
| Luc7l2 | 1.27E-08 | 0.617178 | 0.378 | 0.625 | 0.000299 |
| Cdk5r1 | 1.31E-08 | 0.811698 | 0.112 | 0.251 | 0.000309 |
| Lamtor4 | 1.35E-08 | 2.613203 | 0.513 | 0.534 | 0.000319 |
| Atp6v0a1 | 1.35E-08 | 0.33158 | 0.197 | 0.368 | 0.000319 |
| Gm6063 | 1.36E-08 | 0.68057 | 0.212 | 0.075 | 0.000321 |
| Dcaf8 | 1.36E-08 | 0.380334 | 0.149 | 0.306 | 0.000322 |
| Manba | 1.43E-08 | -0.70633 | 0.086 | 0.202 | 0.000337 |

| | | | | | |
|---|---|---|---|---|---|
| Sfi1 | 1.43E-08 | 0.466261 | 0.134 | 0.28 | 0.000337 |
| Hsp90b1 | 1.51E-08 | 0.800607 | 0.881 | 0.971 | 0.000356 |
| Kcmf1 | 1.55E-08 | 0.697653 | 0.135 | 0.29 | 0.000366 |
| Zcchc7 | 1.56E-08 | 0.303807 | 0.182 | 0.349 | 0.000368 |
| Eps15 | 1.57E-08 | -0.28875 | 0.106 | 0.235 | 0.00037 |
| Npc1 | 1.57E-08 | 0.515018 | 0.245 | 0.453 | 0.000371 |
| P4hb | 1.6E-08 | -0.38146 | 0.762 | 0.928 | 0.000377 |
| Gatad2a | 1.6E-08 | -0.39699 | 0.136 | 0.28 | 0.000378 |
| Gm6863 | 1.62E-08 | 1.540702 | 0.407 | 0.283 | 0.000382 |
| Ganab | 1.63E-08 | -0.29832 | 0.13 | 0.267 | 0.000385 |
| Manf | 1.63E-08 | 1.776501 | 0.591 | 0.678 | 0.000385 |
| Phf12 | 1.68E-08 | 0.542105 | 0.104 | 0.238 | 0.000397 |
| P2ry6 | 1.69E-08 | 1.228673 | 0.742 | 0.86 | 0.000398 |
| Slc23a2 | 1.7E-08 | -0.60015 | 0.105 | 0.235 | 0.000401 |
| Gm42659 | 1.7E-08 | 2.571735 | 0.314 | 0.176 | 0.000402 |
| Cd48 | 1.72E-08 | 2.07445 | 0.501 | 0.482 | 0.000407 |
| Tcn2 | 1.81E-08 | 1.13278 | 0.748 | 0.893 | 0.000426 |
| Smad7 | 1.91E-08 | 1.451411 | 0.25 | 0.453 | 0.000451 |
| Maml3 | 1.92E-08 | 0.654737 | 0.12 | 0.261 | 0.000453 |
| Map3k1 | 1.92E-08 | -0.68302 | 0.293 | 0.446 | 0.000453 |
| Kif21b | 1.95E-08 | 0.393237 | 0.21 | 0.394 | 0.000459 |
| Rabl6 | 1.98E-08 | 0.918368 | 0.135 | 0.287 | 0.000467 |
| Csf2rb2 | 2.02E-08 | -0.2841 | 0.088 | 0.205 | 0.000476 |
| Dido1 | 2.05E-08 | 1.197682 | 0.157 | 0.322 | 0.000483 |
| Ppp4r1 | 2.07E-08 | 0.334501 | 0.134 | 0.28 | 0.000489 |
| Pdia6 | 2.1E-08 | 1.167476 | 0.733 | 0.857 | 0.000495 |
| Myl6 | 2.16E-08 | 1.918821 | 0.623 | 0.681 | 0.000511 |
| Lnpep | 2.19E-08 | 0.270636 | 0.358 | 0.567 | 0.000518 |
| Ctcf | 2.26E-08 | 1.043319 | 0.165 | 0.339 | 0.000532 |
| Gng2 | 2.27E-08 | 0.477386 | 0.241 | 0.453 | 0.000535 |
| Ykt6 | 2.28E-08 | 0.270314 | 0.25 | 0.433 | 0.000539 |
| Rpl15-ps3 | 2.52E-08 | 1.145835 | 0.294 | 0.153 | 0.000595 |
| Lbr | 2.61E-08 | -0.40173 | 0.129 | 0.267 | 0.000615 |
| Slc36a1 | 2.63E-08 | 0.987219 | 0.118 | 0.257 | 0.000621 |
| Atxn2l | 2.7E-08 | -0.35944 | 0.207 | 0.378 | 0.000638 |
| Phka2 | 2.88E-08 | 0.473575 | 0.092 | 0.215 | 0.000679 |
| Gm15590 | 2.95E-08 | 0.736304 | 0.515 | 0.388 | 0.000696 |
| Ube2k | 2.99E-08 | 1.215714 | 0.187 | 0.368 | 0.000705 |
| Pcbp1 | 3.08E-08 | 1.220324 | 0.772 | 0.893 | 0.000727 |
| Riok3 | 3.09E-08 | -0.40525 | 0.089 | 0.205 | 0.000729 |
| Sgk3 | 3.16E-08 | -0.34706 | 0.132 | 0.267 | 0.000745 |
| Camsap1 | 3.21E-08 | 0.4358 | 0.108 | 0.238 | 0.000757 |
| Mafb | 3.4E-08 | 1.102831 | 0.822 | 0.964 | 0.000802 |
| Gls | 3.41E-08 | 0.257329 | 0.089 | 0.208 | 0.000805 |
| Gm5915 | 3.47E-08 | 1.564025 | 0.248 | 0.114 | 0.000819 |
| Senp2 | 3.49E-08 | 0.621965 | 0.247 | 0.433 | 0.000822 |
| Abi1 | 3.61E-08 | 0.267658 | 0.281 | 0.469 | 0.000853 |

| | | | | | |
|---|---|---|---|---|---|
| Spop | 3.63E-08 | 0.505843 | 0.159 | 0.313 | 0.000855 |
| Git2 | 3.66E-08 | 0.428745 | 0.565 | 0.84 | 0.000863 |
| Agpat3 | 3.67E-08 | 0.647964 | 0.201 | 0.381 | 0.000865 |
| Gnai3 | 3.67E-08 | -0.9093 | 0.141 | 0.267 | 0.000866 |
| Arfgap1 | 3.72E-08 | 0.30834 | 0.139 | 0.287 | 0.000877 |
| Trip11 | 3.85E-08 | 0.492395 | 0.096 | 0.221 | 0.000907 |
| Sesn1 | 3.86E-08 | -0.42767 | 0.292 | 0.472 | 0.000911 |
| Tbc1d16 | 4.03E-08 | 0.355235 | 0.142 | 0.29 | 0.00095 |
| Hook3 | 4.07E-08 | 0.572818 | 0.275 | 0.489 | 0.000961 |
| Rapgef1 | 4.08E-08 | 0.507854 | 0.176 | 0.345 | 0.000963 |
| Golgb1 | 4.21E-08 | 0.552186 | 0.09 | 0.212 | 0.000992 |
| Uggt1 | 4.24E-08 | 0.411026 | 0.125 | 0.267 | 0.000999 |
| Btbd7 | 4.52E-08 | -0.28017 | 0.095 | 0.215 | 0.001067 |
| Ssr3 | 4.7E-08 | 0.260191 | 0.249 | 0.427 | 0.001109 |
| Med12l | 4.75E-08 | 0.307905 | 0.409 | 0.668 | 0.001119 |
| Spg20 | 4.93E-08 | -0.53451 | 0.117 | 0.241 | 0.001162 |
| Zfp110 | 5.07E-08 | -1.25432 | 0.1 | 0.215 | 0.001195 |
| Disc1 | 5.09E-08 | 1.112382 | 0.095 | 0.225 | 0.0012 |
| Ehmt2 | 5.1E-08 | 0.863243 | 0.096 | 0.221 | 0.001202 |
| Qrich1 | 5.22E-08 | -0.51796 | 0.186 | 0.332 | 0.001231 |
| Kat7 | 5.31E-08 | -0.69963 | 0.152 | 0.287 | 0.001253 |
| Prpsap2 | 5.33E-08 | 0.561761 | 0.247 | 0.463 | 0.001257 |
| B430010I2 | 5.48E-08 | 0.805417 | 0.102 | 0.231 | 0.001294 |
| Mms19 | 5.63E-08 | -0.72193 | 0.123 | 0.254 | 0.001328 |
| Rpl5 | 5.69E-08 | 2.030021 | 0.732 | 0.876 | 0.001342 |
| Gm42418 | 5.79E-08 | 2.619675 | 0.552 | 0.58 | 0.001366 |
| Accs | 6.49E-08 | 0.540902 | 0.119 | 0.254 | 0.001532 |
| Gm8566 | 6.52E-08 | 0.455498 | 0.214 | 0.081 | 0.001538 |
| Atrx | 6.6E-08 | 0.585962 | 0.212 | 0.391 | 0.001557 |
| Lonp2 | 6.64E-08 | -0.2744 | 0.149 | 0.29 | 0.001566 |
| Arhgap4 | 6.72E-08 | -0.54558 | 0.147 | 0.287 | 0.001585 |
| Rnf44 | 6.73E-08 | 0.910461 | 0.226 | 0.42 | 0.001587 |
| Tex261 | 6.95E-08 | 2.661768 | 0.457 | 0.397 | 0.001639 |
| mt-Nd3 | 7.14E-08 | 1.620382 | 0.559 | 0.544 | 0.001685 |
| Nufip2 | 7.25E-08 | -0.26832 | 0.132 | 0.264 | 0.00171 |
| Otud5 | 7.27E-08 | 0.63179 | 0.175 | 0.336 | 0.001714 |
| Ttc7 | 7.51E-08 | 1.121308 | 0.131 | 0.274 | 0.00177 |
| Ndfip1 | 7.53E-08 | 1.39394 | 0.673 | 0.759 | 0.001777 |
| Ppm1h | 7.8E-08 | 1.682662 | 0.641 | 0.616 | 0.00184 |
| Fip1l1 | 8.02E-08 | 1.000548 | 0.115 | 0.251 | 0.001892 |
| Ccng2 | 8.47E-08 | -0.48453 | 0.25 | 0.41 | 0.001999 |
| Nipa2 | 8.93E-08 | -0.2874 | 0.145 | 0.277 | 0.002106 |
| Asap1 | 8.96E-08 | 0.498544 | 0.274 | 0.489 | 0.002114 |
| Zfp740 | 9.65E-08 | 0.339024 | 0.205 | 0.368 | 0.002276 |
| Trim2 | 9.95E-08 | 0.685912 | 0.119 | 0.251 | 0.002347 |
| Lrrc8a | 9.99E-08 | 2.929755 | 0.515 | 0.515 | 0.002357 |
| Kpna3 | 1.01E-07 | 0.483302 | 0.151 | 0.3 | 0.002387 |

| | | | | | |
|---|---|---|---|---|---|
| Wdr33 | 1.03E-07 | 0.424331 | 0.139 | 0.287 | 0.002422 |
| Dusp7 | 1.07E-07 | 0.738424 | 0.336 | 0.55 | 0.002523 |
| Rabgap1 | 1.09E-07 | 0.348812 | 0.138 | 0.28 | 0.002569 |
| Naca | 1.11E-07 | 1.839651 | 0.608 | 0.684 | 0.002628 |
| Gm13680 | 1.14E-07 | 1.384691 | 0.378 | 0.238 | 0.002689 |
| Hectd1 | 1.24E-07 | 0.261224 | 0.18 | 0.345 | 0.002931 |
| Gak | 1.25E-07 | -0.46307 | 0.279 | 0.459 | 0.002937 |
| Mga | 1.35E-07 | 0.253879 | 0.131 | 0.267 | 0.003186 |
| Smndc1 | 1.36E-07 | 0.295221 | 0.128 | 0.267 | 0.003215 |
| Ywhag | 1.37E-07 | 0.66301 | 0.171 | 0.313 | 0.003223 |
| Rnf4 | 1.38E-07 | -0.69474 | 0.151 | 0.287 | 0.003265 |
| Fam221b | 1.42E-07 | 1.410244 | 0.429 | 0.329 | 0.003346 |
| Cd83 | 1.42E-07 | 2.39704 | 0.358 | 0.235 | 0.003359 |
| Bcap31 | 1.45E-07 | 1.488445 | 0.612 | 0.678 | 0.003431 |
| Braf | 1.47E-07 | 0.466158 | 0.096 | 0.215 | 0.003467 |
| Gm11808 | 1.49E-07 | 1.725118 | 0.441 | 0.336 | 0.003517 |
| Sel1l | 1.5E-07 | 0.469428 | 0.225 | 0.397 | 0.003545 |
| Tmem260 | 1.5E-07 | -0.32708 | 0.106 | 0.225 | 0.003548 |
| Glmp | 1.54E-07 | 1.523034 | 0.582 | 0.603 | 0.003638 |
| Slc30a5 | 1.54E-07 | -0.82474 | 0.105 | 0.218 | 0.003638 |
| Bnip2 | 1.55E-07 | 0.392176 | 0.199 | 0.368 | 0.003647 |
| Uba7 | 1.55E-07 | -0.27604 | 0.139 | 0.27 | 0.003649 |
| R3hdm2 | 1.55E-07 | 0.861808 | 0.087 | 0.202 | 0.003654 |
| Atraid | 1.56E-07 | 1.634412 | 0.599 | 0.655 | 0.003681 |
| Tapbp | 1.58E-07 | 1.139881 | 0.766 | 0.749 | 0.003715 |
| Pnpla7 | 1.59E-07 | 0.484947 | 0.416 | 0.645 | 0.003739 |
| Atrn | 1.59E-07 | -0.63505 | 0.092 | 0.205 | 0.003744 |
| Syt11 | 1.6E-07 | 1.414397 | 0.099 | 0.228 | 0.003784 |
| Rnf145 | 1.61E-07 | -0.55037 | 0.096 | 0.208 | 0.003793 |
| Tagap1 | 1.63E-07 | 1.353709 | 0.136 | 0.274 | 0.003847 |
| Stag1 | 1.7E-07 | 0.297523 | 0.137 | 0.277 | 0.004012 |
| Ckap4 | 1.8E-07 | 0.711225 | 0.168 | 0.322 | 0.00425 |
| Rpl29 | 1.8E-07 | 1.594885 | 0.61 | 0.678 | 0.004254 |
| Csf2rb | 1.81E-07 | 0.858262 | 0.247 | 0.443 | 0.004269 |
| Rnf115 | 1.84E-07 | -0.3883 | 0.147 | 0.28 | 0.004332 |
| Ube2j1 | 1.92E-07 | 0.320624 | 0.202 | 0.365 | 0.004532 |
| Gpx4 | 1.93E-07 | 2.136589 | 0.54 | 0.583 | 0.004555 |
| Kpna4 | 1.98E-07 | 1.131912 | 0.237 | 0.433 | 0.004664 |
| Selenos | 2.03E-07 | 2.14288 | 0.584 | 0.58 | 0.00479 |
| Vcpip1 | 2.03E-07 | -0.65217 | 0.132 | 0.254 | 0.004797 |
| Slc35a3 | 2.04E-07 | -0.76762 | 0.103 | 0.215 | 0.004804 |
| Nrd1 | 2.15E-07 | 0.779383 | 0.141 | 0.283 | 0.005063 |
| Prr14l | 2.15E-07 | 0.643134 | 0.201 | 0.378 | 0.005072 |
| Pds5a | 2.17E-07 | 0.660573 | 0.199 | 0.365 | 0.005125 |
| Akap9 | 2.24E-07 | 0.720281 | 0.199 | 0.375 | 0.005281 |
| Kpnb1 | 2.29E-07 | 0.792623 | 0.158 | 0.303 | 0.00539 |
| Znfx1 | 2.29E-07 | 0.510127 | 0.143 | 0.283 | 0.005395 |

| | | | | | |
|---|---|---|---|---|---|
| Mob2 | 2.33E-07 | 1.304396 | 0.1 | 0.225 | 0.005488 |
| Atp5g2 | 2.35E-07 | 2.429465 | 0.502 | 0.528 | 0.005552 |
| Ypel3 | 2.44E-07 | 1.93648 | 0.623 | 0.704 | 0.005762 |
| Supt6 | 2.45E-07 | 0.978383 | 0.095 | 0.212 | 0.005783 |
| Rpl3-ps2 | 2.54E-07 | 0.415759 | 0.207 | 0.081 | 0.006001 |
| Klhl7 | 2.56E-07 | -1.37531 | 0.103 | 0.208 | 0.006034 |
| Stk10 | 2.57E-07 | 0.821382 | 0.225 | 0.41 | 0.006052 |
| Irf2 | 2.58E-07 | 0.581926 | 0.252 | 0.443 | 0.006074 |
| Kdelr2 | 2.58E-07 | 1.305991 | 0.663 | 0.56 | 0.00608 |
| Trmt112-ps | 2.6E-07 | 0.861334 | 0.25 | 0.121 | 0.006131 |
| Ankrd11 | 2.61E-07 | 0.77737 | 0.363 | 0.596 | 0.006162 |
| Usp4 | 2.69E-07 | -0.46904 | 0.199 | 0.339 | 0.006349 |
| Gngt2 | 2.71E-07 | 1.970965 | 0.592 | 0.694 | 0.006396 |
| Ergic1 | 2.74E-07 | 0.431744 | 0.229 | 0.401 | 0.006472 |
| Mavs | 2.79E-07 | -0.30735 | 0.149 | 0.28 | 0.006577 |
| Ccdc88b | 2.87E-07 | 0.746398 | 0.133 | 0.264 | 0.006763 |
| Ccdc47 | 2.89E-07 | 1.006134 | 0.263 | 0.472 | 0.00681 |
| Plekhf2 | 2.91E-07 | -0.80884 | 0.202 | 0.326 | 0.006857 |
| Chmp3 | 2.97E-07 | 0.9516 | 0.247 | 0.453 | 0.006999 |
| Phf3 | 3.2E-07 | 0.426458 | 0.11 | 0.231 | 0.00755 |
| Rnf111 | 3.22E-07 | 0.254525 | 0.159 | 0.3 | 0.007602 |
| Foxn3 | 3.24E-07 | 1.550478 | 0.81 | 0.922 | 0.007634 |
| Srgn | 3.32E-07 | 1.944358 | 0.5 | 0.472 | 0.007838 |
| Bicd2 | 3.33E-07 | 0.330251 | 0.171 | 0.319 | 0.00786 |
| Epn1 | 3.37E-07 | 0.903422 | 0.158 | 0.303 | 0.007961 |
| Tle3 | 3.42E-07 | 0.995423 | 0.161 | 0.309 | 0.008062 |
| Herpud1 | 3.5E-07 | 1.773997 | 0.604 | 0.629 | 0.008252 |
| Ptp4a1 | 3.53E-07 | -0.62003 | 0.108 | 0.218 | 0.008316 |
| Whrn | 3.58E-07 | 1.150485 | 0.193 | 0.355 | 0.008442 |
| Rela | 3.59E-07 | 0.515604 | 0.287 | 0.492 | 0.008463 |
| Eps15l1 | 3.64E-07 | 0.457212 | 0.103 | 0.221 | 0.008589 |
| Skp1a | 3.65E-07 | 0.754938 | 0.305 | 0.515 | 0.0086 |
| Gabpb2 | 3.68E-07 | 0.988148 | 0.122 | 0.251 | 0.008671 |
| Tmx1 | 3.68E-07 | -0.78878 | 0.23 | 0.365 | 0.00869 |
| Fcgr1 | 3.69E-07 | 1.52525 | 0.665 | 0.805 | 0.008714 |
| Mia3 | 3.71E-07 | 0.586678 | 0.22 | 0.394 | 0.008752 |
| Fem1a | 3.84E-07 | -0.38031 | 0.096 | 0.205 | 0.009059 |
| Edem1 | 3.86E-07 | 0.326696 | 0.445 | 0.7 | 0.009114 |
| Prune2 | 3.89E-07 | 2.568504 | 0.301 | 0.182 | 0.009173 |
| Exoc4 | 3.94E-07 | 0.342655 | 0.098 | 0.212 | 0.009283 |
| Ptpa | 3.97E-07 | 1.172449 | 0.223 | 0.414 | 0.009366 |
| Tmem126a | 3.99E-07 | 2.337485 | 0.449 | 0.401 | 0.009408 |
| Atad1 | 4.07E-07 | -1.30306 | 0.129 | 0.241 | 0.009606 |
| Ripk1 | 4.15E-07 | -0.89121 | 0.125 | 0.244 | 0.009781 |
| Abr | 4.24E-07 | 0.700016 | 0.094 | 0.205 | 0.009991 |
| Brk1 | 4.28E-07 | 1.491111 | 0.662 | 0.798 | 0.010103 |
| Usf2 | 4.34E-07 | 1.373793 | 0.158 | 0.309 | 0.010231 |

| | | | | | |
|---|---|---|---|---|---|
| Apoe | 4.35E-07 | 4.010424 | 0.575 | 0.638 | 0.010258 |
| Mboat1 | 4.36E-07 | 0.506338 | 0.098 | 0.212 | 0.010277 |
| Galnt11 | 4.38E-07 | -0.64962 | 0.118 | 0.231 | 0.01033 |
| Mag | 4.43E-07 | 1.886029 | 0.091 | 0.212 | 0.010455 |
| Rpl17-ps10 | 4.49E-07 | 0.944256 | 0.232 | 0.104 | 0.010594 |
| Zswim8 | 4.58E-07 | 1.338571 | 0.106 | 0.228 | 0.010803 |
| Rptor | 4.59E-07 | 0.380143 | 0.255 | 0.436 | 0.010829 |
| Lyst | 4.61E-07 | 0.917263 | 0.095 | 0.212 | 0.01087 |
| Calu | 4.65E-07 | -0.72141 | 0.211 | 0.339 | 0.010959 |
| Kdm2b | 4.76E-07 | 0.270409 | 0.157 | 0.296 | 0.011231 |
| Rcsd1 | 5.22E-07 | 0.457595 | 0.318 | 0.531 | 0.012304 |
| Pik3ca | 5.22E-07 | 0.452679 | 0.116 | 0.235 | 0.012322 |
| Psme1 | 5.24E-07 | 2.403271 | 0.518 | 0.495 | 0.012367 |
| Fbxw11 | 5.37E-07 | -0.71649 | 0.183 | 0.313 | 0.012659 |
| Gm3839 | 5.5E-07 | 0.417709 | 0.227 | 0.104 | 0.012963 |
| Slc12a6 | 5.65E-07 | 0.733308 | 0.245 | 0.423 | 0.013327 |
| Ncbp3 | 5.83E-07 | 0.259917 | 0.135 | 0.264 | 0.013744 |
| Selenow | 5.89E-07 | 2.896283 | 0.264 | 0.147 | 0.013885 |
| Btbd9 | 6.1E-07 | 0.486997 | 0.122 | 0.251 | 0.014383 |
| Mlec | 6.26E-07 | 0.930054 | 0.35 | 0.616 | 0.014771 |
| Gm42602 | 6.51E-07 | 0.312492 | 0.109 | 0.231 | 0.015365 |
| Ldha | 6.52E-07 | 1.676058 | 0.591 | 0.645 | 0.015384 |
| Rgs2 | 6.59E-07 | 1.257548 | 0.747 | 0.902 | 0.015551 |
| Pld3 | 6.6E-07 | 1.651439 | 0.609 | 0.658 | 0.015577 |
| AI837181 | 6.63E-07 | 0.326903 | 0.173 | 0.316 | 0.01564 |
| Slc44a2 | 6.64E-07 | -0.70979 | 0.308 | 0.436 | 0.015655 |
| St6gal1 | 6.7E-07 | 0.28293 | 0.101 | 0.212 | 0.015813 |
| Ccpg1 | 6.76E-07 | 0.730658 | 0.174 | 0.313 | 0.015939 |
| Rsbn1 | 7.33E-07 | 0.347658 | 0.11 | 0.235 | 0.017302 |
| Rbm3 | 7.37E-07 | 2.968741 | 0.301 | 0.199 | 0.017382 |
| Gm12338 | 7.38E-07 | 1.182326 | 0.276 | 0.127 | 0.017403 |
| Snrk | 7.55E-07 | 0.727702 | 0.103 | 0.218 | 0.017801 |
| Rhoa | 7.58E-07 | 1.027397 | 0.841 | 0.847 | 0.017885 |
| Ankrd13a | 7.64E-07 | -0.72947 | 0.186 | 0.313 | 0.018021 |
| Smarcad1 | 7.64E-07 | 0.68174 | 0.107 | 0.225 | 0.018027 |
| Lmo2 | 7.89E-07 | 1.812544 | 0.615 | 0.681 | 0.018617 |
| Cask | 7.96E-07 | 0.575814 | 0.306 | 0.502 | 0.018787 |
| Lhfpl2 | 8E-07 | 0.41753 | 0.28 | 0.456 | 0.018872 |
| Otud1 | 8.33E-07 | -0.76723 | 0.133 | 0.251 | 0.019644 |
| Orai1 | 8.36E-07 | 1.991534 | 0.592 | 0.694 | 0.019725 |
| Csad | 8.61E-07 | 0.746234 | 0.26 | 0.427 | 0.020302 |
| Gm12428 | 8.7E-07 | 0.99416 | 0.246 | 0.127 | 0.020515 |
| Dleu2 | 8.95E-07 | 0.486133 | 0.748 | 0.906 | 0.021105 |
| Akap13 | 8.96E-07 | 0.780585 | 0.417 | 0.655 | 0.021131 |
| Mras | 9.26E-07 | 1.112336 | 0.093 | 0.208 | 0.021832 |
| Chn2 | 9.37E-07 | 0.653838 | 0.228 | 0.404 | 0.022098 |
| Phactr2 | 9.44E-07 | 0.356706 | 0.219 | 0.378 | 0.022261 |

| | | | | | |
|---|---|---|---|---|---|
| Sike1 | 9.72E-07 | 0.266541 | 0.123 | 0.248 | 0.02293 |
| Wnk1 | 9.75E-07 | 0.442415 | 0.618 | 0.902 | 0.023011 |
| Brf1 | 9.81E-07 | 0.766517 | 0.123 | 0.248 | 0.02314 |
| Casc3 | 9.89E-07 | 1.215691 | 0.11 | 0.228 | 0.023319 |
| Sertad2 | 1E-06 | 0.824195 | 0.13 | 0.251 | 0.023706 |
| Eif3c | 1.01E-06 | 0.66488 | 0.35 | 0.57 | 0.023859 |
| Usp12 | 1.04E-06 | 0.37829 | 0.117 | 0.238 | 0.024582 |
| Gm10095 | 1.04E-06 | 1.344625 | 0.429 | 0.326 | 0.024622 |
| Arhgap30 | 1.1E-06 | 0.401266 | 0.224 | 0.381 | 0.026007 |
| Mecp2 | 1.12E-06 | 1.122703 | 0.17 | 0.319 | 0.026409 |
| Lrrfip1 | 1.14E-06 | 0.807911 | 0.217 | 0.391 | 0.026949 |
| Pgrmc2 | 1.16E-06 | 0.937994 | 0.097 | 0.208 | 0.02729 |
| Ptges3 | 1.17E-06 | 1.783011 | 0.534 | 0.489 | 0.02767 |
| Rnf20 | 1.18E-06 | 0.471548 | 0.096 | 0.205 | 0.02776 |
| Efr3a | 1.2E-06 | -0.45045 | 0.133 | 0.251 | 0.028387 |
| Gm5514 | 1.21E-06 | 1.52826 | 0.505 | 0.443 | 0.028444 |
| Kdm5a | 1.21E-06 | 0.345543 | 0.164 | 0.296 | 0.028476 |
| Trrap | 1.21E-06 | 0.674805 | 0.136 | 0.261 | 0.02858 |
| Set | 1.22E-06 | 2.200815 | 0.603 | 0.632 | 0.028769 |
| Ubtf | 1.22E-06 | 0.42768 | 0.144 | 0.274 | 0.028889 |
| Pxk | 1.25E-06 | 0.636246 | 0.148 | 0.283 | 0.029445 |
| Rhoh | 1.26E-06 | 0.794015 | 0.345 | 0.625 | 0.029752 |
| Pkm | 1.26E-06 | 1.623272 | 0.639 | 0.704 | 0.029813 |
| Rgl1 | 1.3E-06 | -0.31358 | 0.124 | 0.238 | 0.030684 |
| Gm8451 | 1.32E-06 | 0.811648 | 0.229 | 0.107 | 0.031214 |
| Suds3 | 1.34E-06 | 0.867497 | 0.101 | 0.212 | 0.031575 |
| Gnl2 | 1.39E-06 | 0.425403 | 0.122 | 0.244 | 0.032762 |
| 2610005L0 | 1.43E-06 | 0.55106 | 0.172 | 0.316 | 0.033822 |
| Pcmtd1 | 1.45E-06 | 0.314481 | 0.123 | 0.241 | 0.034271 |
| Pkn2 | 1.45E-06 | 0.355873 | 0.139 | 0.27 | 0.034303 |
| Cdc42se2 | 1.47E-06 | -0.78509 | 0.122 | 0.231 | 0.034574 |
| Fcrls | 1.47E-06 | 0.698477 | 0.902 | 0.997 | 0.034742 |
| Tmem131 | 1.5E-06 | 0.301731 | 0.116 | 0.231 | 0.035288 |
| Npepps | 1.5E-06 | 0.758912 | 0.181 | 0.326 | 0.03549 |
| Tifab | 1.59E-06 | 0.654655 | 0.427 | 0.674 | 0.0376 |
| Gm10134 | 1.61E-06 | 2.5534 | 0.397 | 0.313 | 0.038071 |
| Atp7a | 1.62E-06 | 0.627072 | 0.115 | 0.231 | 0.038251 |
| Zc3h7a | 1.63E-06 | 0.50445 | 0.197 | 0.336 | 0.038355 |
| Gm8186 | 1.63E-06 | 1.382506 | 0.331 | 0.231 | 0.038465 |
| Cic | 1.65E-06 | 1.988948 | 0.11 | 0.228 | 0.038925 |
| Acin1 | 1.68E-06 | 1.138877 | 0.241 | 0.42 | 0.039514 |
| Foxk2 | 1.7E-06 | 0.660632 | 0.099 | 0.208 | 0.04007 |
| Rpl9-ps7 | 1.73E-06 | 1.41697 | 0.285 | 0.176 | 0.040798 |
| Gm4332 | 1.73E-06 | 0.445132 | 0.204 | 0.088 | 0.04091 |
| Commd8 | 1.75E-06 | 1.608344 | 0.616 | 0.671 | 0.041312 |
| Pak1 | 1.79E-06 | 0.398262 | 0.173 | 0.316 | 0.042328 |
| Itpr2 | 1.81E-06 | 0.581448 | 0.389 | 0.606 | 0.042665 |

| | | | | | |
|---|---|---|---|---|---|
| Tet3 | 1.89E-06 | 1.331368 | 0.16 | 0.303 | 0.044575 |
| Mtr | 1.92E-06 | 0.50884 | 0.11 | 0.225 | 0.0454 |
| Gm5526 | 1.95E-06 | 1.532742 | 0.359 | 0.261 | 0.045958 |
| Gng5-ps | 1.95E-06 | 1.618181 | 0.314 | 0.186 | 0.046081 |
| Rpl39-ps | 1.96E-06 | 0.777533 | 0.212 | 0.098 | 0.04616 |
| Taok2 | 2.02E-06 | 0.457139 | 0.152 | 0.28 | 0.047746 |
| Cntrl | 2.08E-06 | 1.372174 | 0.11 | 0.228 | 0.04897 |
| Uba5 | 2.11E-06 | 1.335608 | 0.159 | 0.306 | 0.049746 |
| Aplp2 | 2.11E-06 | 0.40259 | 0.241 | 0.401 | 0.049838 |

Supplementry Table 5.  Upregulated aging DEGs shared by brain regions

| Gene | Cb | Ctx | Hipc | Str |
|---|---|---|---|---|
| C1qa | 1.853858 | 2.006581 | 2.20964 | 2.343129 |
| C1qc | 1.747765 | 1.644847 | 1.770819 | 2.056197 |
| H2-D1 | 2.807493 | 1.89344 | 2.056101 | 2.602176 |
| Itm2b | 1.647215 | 1.299334 | 1.305902 | 1.645928 |
| Cd81 | 1.826646 | 1.56412 | 1.458362 | 1.960568 |
| Fth1 | 2.327446 | 1.732452 | 1.929984 | 2.736824 |
| mt-Co1 | 3.060626 | 2.004782 | 1.993499 | 2.397621 |
| B2m | 2.21048 | 1.427054 | 1.774941 | 2.22761 |
| Cd9 | 2.46679 | 1.959819 | 1.715651 | 2.494392 |
| C1qb | 1.276743 | 1.130378 | 1.33286 | 1.649717 |
| Tmsb4x | 1.623375 | 1.314515 | 1.684218 | 2.01925 |
| Ctss | 1.378233 | 1.000591 | 1.188542 | 1.464294 |
| Ftl1 | 2.124822 | 1.736279 | 1.815528 | 2.338335 |
| Ctsz | 1.932903 | 1.517976 | 1.542733 | 1.974315 |
| mt-Co3 | 2.912468 | 1.6385 | 1.925971 | 2.582627 |
| Cst3 | 1.190546 | 1.131909 | 1.566852 | 1.613518 |
| Lgmn | 1.384215 | 1.522331 | 1.355586 | 1.39708 |
| H2-K1 | 2.796553 | 1.931831 | 2.043999 | 2.570755 |
| mt-Cytb | 2.996795 | 1.684504 | 1.872819 | 2.515032 |
| Ct010467.: | 1.077502 | 1.219254 | 0.6433 | 1.351058 |
| Eef1a1 | 1.741832 | 1.626355 | 1.72357 | 2.013449 |
| Ctsb | 1.714736 | 1.517297 | 1.30712 | 1.587931 |
| Tmem119 | 1.795846 | 1.648647 | 1.536644 | 1.611145 |
| Laptm5 | 1.36608 | 1.572493 | 1.259993 | 1.239102 |
| mt-Atp6 | 2.714994 | 1.27201 | 1.914218 | 2.492778 |
| mt-Co2 | 3.137367 | 2.272692 | 2.287388 | 2.892311 |
| Ctsd | 1.242285 | 1.06006 | 0.841997 | 1.12686 |
| Sparc | 1.558502 | 1.428147 | 1.497359 | 1.64283 |
| Rpl18a | 1.736399 | 1.287411 | 1.684043 | 2.193783 |
| Rpl13 | 1.919899 | 1.870934 | 2.195114 | 2.581873 |
| Cyba | 1.94654 | 1.733091 | 1.967413 | 2.310095 |
| Cfl1 | 2.885331 | 2.831777 | 3.174299 | 3.14759 |
| Ctsh | 1.748036 | 1.528448 | 1.386568 | 2.002157 |
| Hexb | 0.916174 | 1.093001 | 0.919576 | 0.935643 |
| Tyrobp | 1.425143 | 0.81804 | 1.110912 | 1.529583 |
| Adap2os | 2.680755 | 2.220091 | 1.914587 | 2.714242 |
| Fcer1g | 1.547273 | 1.020688 | 1.401742 | 1.860022 |
| Rps9 | 1.627412 | 1.571397 | 1.971185 | 2.193515 |
| Selplg | 1.14387 | 1.495924 | 1.387974 | 1.488028 |
| Gm15427 | 2.334009 | 1.479973 | 2.256625 | 3.396543 |
| Trem2 | 1.436874 | 1.613965 | 1.596797 | 1.838354 |
| Ftl1-ps1 | 2.038969 | 1.224576 | 1.554706 | 2.081555 |
| mt-Nd1 | 2.531548 | 1.170367 | 1.542602 | 1.918003 |
| Ctsl | 1.430644 | 1.288013 | 1.080366 | 1.555186 |
| Ftl2-ps | 1.862339 | 1.346914 | 1.524958 | 2.336141 |
| Pfn1 | 2.046108 | 1.822958 | 2.341094 | 2.597359 |

| | | | | |
|---|---|---|---|---|
| Gm15710 | 2.251601 | 2.075646 | 2.201802 | 2.488268 |
| Ubb | 1.415949 | 1.547364 | 1.378349 | 1.938131 |
| Pld4 | 1.636016 | 1.710181 | 1.456462 | 1.7436 |
| Rpl13a | 2.829219 | 2.578267 | 3.198879 | 3.510987 |
| Gm1821 | 1.750526 | 1.544619 | 1.494605 | 1.846036 |
| Malat1 | 1.174955 | 1.395726 | 0.53155 | 0.988488 |
| Ctsa | 1.495706 | 1.310819 | 1.044789 | 1.388636 |
| Grn | 1.141457 | 0.94005 | 0.818605 | 0.908926 |
| Rplp1 | 1.566727 | 1.297632 | 1.816279 | 2.083708 |
| Rps18 | 1.848362 | 1.329345 | 1.732055 | 2.403856 |
| Gm9843 | 2.509165 | 1.675078 | 2.265571 | 2.897254 |
| Cd63 | 2.409304 | 1.562877 | 1.960149 | 2.772639 |
| Rplp0 | 1.778761 | 1.563331 | 2.003586 | 2.396967 |
| Aif1 | 2.016538 | 1.411259 | 1.989158 | 2.173675 |
| Rps11 | 1.563963 | 1.34259 | 1.756573 | 2.23684 |
| Olfml3 | 1.206352 | 1.237013 | 1.110217 | 1.247553 |
| Lamp1 | 1.50678 | 1.024698 | 0.94984 | 1.114112 |
| Rnaset2a | 1.990393 | 1.626602 | 1.725775 | 2.015996 |
| Sat1 | 2.222304 | 1.39279 | 1.399859 | 2.102711 |
| Cd68 | 1.543406 | 1.338766 | 1.365817 | 1.694935 |
| Rps3 | 1.959612 | 1.68354 | 1.941693 | 2.621602 |
| Gm13456 | 1.445988 | 0.882259 | 1.178235 | 1.452079 |
| P2ry12 | 1.274377 | 1.319135 | 0.760672 | 0.522623 |
| Rgs10 | 1.62688 | 1.392476 | 1.476395 | 1.572671 |
| Rps3a1 | 1.907011 | 1.629166 | 1.937884 | 2.460547 |
| Nisch | 0.260968 | 0.765815 | 0.913364 | 0.466756 |
| Gm8797 | 1.692104 | 1.226642 | 1.113804 | 1.465702 |
| Rps14 | 1.385959 | 1.102696 | 1.753724 | 2.050916 |
| Rps20 | 1.966447 | 1.573061 | 1.880115 | 2.5059 |
| Psap | 1.102581 | 1.375392 | 1.208883 | 0.75181 |
| Rps27rt | 1.907456 | 1.422843 | 1.812534 | 2.602018 |
| Npc2 | 2.237958 | 1.706959 | 2.003904 | 2.000043 |
| Rps18-ps3 | 2.298733 | 1.278715 | 1.84758 | 2.593384 |
| Rnaset2b | 2.061368 | 1.504746 | 1.651613 | 2.200094 |
| Rpl3 | 2.125217 | 1.938416 | 2.038635 | 2.183243 |
| Skil | 2.383291 | 1.903509 | 1.266292 | 1.790424 |
| Ptafr | 2.47661 | 2.108529 | 1.95841 | 2.293913 |
| Dpysl2 | 2.304653 | 1.343547 | 2.066976 | 2.101103 |
| Gm28661 | 1.616137 | 0.494739 | 0.858483 | 1.177186 |
| Ly86 | 1.011205 | 1.01496 | 0.817101 | 1.337087 |
| Tmem50a | 1.88121 | 1.483469 | 1.510288 | 1.706543 |
| Basp1 | 2.272563 | 2.053306 | 2.432305 | 2.487242 |
| Gm4149 | 1.674577 | 1.085511 | 1.844164 | 2.627926 |
| Atp6v0c-ps | 2.229001 | 1.573574 | 1.92044 | 2.303954 |
| Rps19 | 1.36797 | 1.14347 | 1.441349 | 2.004701 |
| Atp6v0c | 1.825365 | 1.618328 | 1.778104 | 2.18455 |
| Gm29216 | 1.294318 | 0.377236 | 0.751441 | 1.180711 |
| Calm2 | 1.852447 | 1.379358 | 1.597813 | 1.88239 |
| Ppia | 0.947491 | 1.089255 | 1.323139 | 1.758926 |

| | | | | |
|---|---|---|---|---|
| Hexa | 1.187797 | 1.101542 | 0.880631 | 1.191102 |
| Cd63-ps | 1.688035 | 1.150022 | 1.523205 | 2.512112 |
| Dhrs3 | 2.154343 | 1.658054 | 1.825642 | 2.11813 |
| Bsg | 2.027589 | 2.09094 | 2.232939 | 2.500758 |
| Zfp691 | 1.52944 | 2.101083 | 2.338423 | 2.646309 |
| Zscan26 | 2.336951 | 1.845884 | 1.794479 | 2.309437 |
| Syngr1 | 2.171922 | 1.898464 | 1.719228 | 2.224036 |
| Rpl4 | 1.934048 | 1.774778 | 1.866869 | 2.063898 |
| Otulinl | 1.590533 | 1.658309 | 1.125709 | 1.450356 |
| Srsf2 | 2.088425 | 1.681741 | 1.585842 | 2.39054 |
| Gm11478 | 2.40675 | 2.689593 | 2.894968 | 3.717679 |
| Rpsa | 1.9068 | 1.549843 | 1.879884 | 2.470055 |
| Bhlhe41 | 2.190557 | 1.790085 | 1.842975 | 2.162238 |
| Gm5559 | 2.48943 | 2.64645 | 2.596438 | 2.987467 |
| Clic1 | 1.795906 | 1.543501 | 1.875196 | 1.883906 |
| Ifngr1 | 1.22863 | 1.1135 | 0.790295 | 0.971766 |
| Rpl23a-ps3 | 1.803137 | 1.15386 | 1.433192 | 1.950229 |
| Rps7 | 1.952847 | 1.704275 | 2.32159 | 2.857544 |
| Serpine2 | 1.609396 | 1.53516 | 1.160305 | 1.200699 |
| Gm5879 | 1.042292 | 0.873832 | 1.013309 | 1.107086 |
| Abhd12 | 1.215497 | 1.186248 | 1.265977 | 1.250793 |
| Rpl10 | 1.495147 | 1.568293 | 1.504062 | 1.992964 |
| Spi1 | 2.078744 | 2.14472 | 2.159991 | 2.284926 |
| Rpl6 | 2.059673 | 1.802549 | 2.16007 | 2.731482 |
| Ywhah | 1.518435 | 1.195356 | 1.120939 | 1.610326 |
| Gm15266 | 1.699228 | 1.213174 | 1.672681 | 2.214386 |
| Gm10123 | 1.63122 | 1.32289 | 1.480841 | 1.848879 |
| Rps6 | 1.596915 | 1.575736 | 1.924985 | 2.120291 |
| Rpl3-ps1 | 2.0499 | 1.440059 | 1.651672 | 2.075493 |
| Sft2d1 | 1.74384 | 1.33395 | 1.325582 | 1.848792 |
| Gm6977 | 1.915321 | 1.455023 | 0.870938 | 1.975018 |
| Rpl21 | 1.303495 | 0.919194 | 1.797746 | 1.950161 |
| Gm6900 | 0.945824 | 0.739543 | 0.845529 | 0.832336 |
| F11r | 1.651467 | 1.763818 | 1.620341 | 1.582579 |
| Laptm4a | 1.686832 | 1.626195 | 1.634788 | 1.983895 |
| Gm16378 | 0.803596 | 0.716014 | 0.815504 | 0.880299 |
| Gm43028 | 1.039914 | 0.688334 | 1.019909 | 1.329749 |
| Gm44705 | 0.919083 | 0.661504 | 0.735068 | 0.833289 |
| Marcks | 1.12397 | 0.864838 | 1.062184 | 0.747169 |
| Atp6v0b | 1.815216 | 1.353712 | 1.423741 | 1.686966 |
| Bri3 | 2.904392 | 2.519494 | 2.601869 | 2.730527 |
| Rpl32 | 1.336516 | 1.218032 | 1.597509 | 2.21577 |
| Rpl18 | 1.549695 | 1.248089 | 1.712672 | 1.960462 |
| Adap2 | 1.75648 | 1.546553 | 1.402501 | 1.553739 |
| Siglech | 0.924998 | 1.25478 | 1.076596 | 0.960868 |
| Cotl1 | 2.45567 | 2.636815 | 2.803024 | 2.734412 |
| Rpl17 | 1.74675 | 1.452795 | 2.165464 | 2.789302 |
| Unc93b1 | 0.918802 | 1.250821 | 1.136095 | 0.936437 |
| Gm12918 | 1.270591 | 0.998722 | 1.318662 | 1.327218 |

| | | | | |
|---|---|---|---|---|
| Tubb5 | 1.78053 | 1.478432 | 1.798756 | 2.001377 |
| Rpl7 | 1.588154 | 1.352722 | 1.556623 | 1.717806 |
| Tmbim6 | 1.266337 | 1.483447 | 1.29106 | 0.926721 |
| Itm2c | 1.651668 | 1.58214 | 1.530566 | 1.395848 |
| Gm3362 | 1.103473 | 0.298038 | 0.34283 | 1.050869 |
| Lpcat2 | 0.963198 | 1.142142 | 0.948255 | 0.753177 |
| H3f3b | 1.508169 | 1.343693 | 1.179588 | 1.443282 |
| Gnb1 | 2.264073 | 2.150764 | 2.478036 | 2.394404 |
| Gm12537 | 1.134109 | 0.991624 | 0.966718 | 1.417731 |
| Slc25a5 | 1.897555 | 1.553226 | 1.622144 | 1.869814 |
| Gm11942 | 1.313789 | 0.942476 | 1.117267 | 1.310587 |
| Ang | 2.494037 | 1.852652 | 1.906409 | 2.277801 |
| Ftl1-ps2 | 0.783472 | 0.423571 | 0.464605 | 0.616741 |
| Gm6204 | 1.802594 | 1.367613 | 1.791211 | 2.069594 |
| Rps16-ps2 | 1.773008 | 1.649893 | 1.692623 | 2.382308 |
| Gm8730 | 1.553391 | 1.253955 | 1.560018 | 1.982919 |
| Gm2000 | 1.582642 | 0.808792 | 1.598923 | 2.120327 |
| Gapdh | 1.300807 | 1.321668 | 1.539166 | 1.750101 |
| Ac165079. | 0.96889 | 0.687332 | 0.850443 | 1.38277 |
| Rpl19 | 1.422397 | 1.283208 | 1.684111 | 1.992811 |
| Ldhb | 2.18332 | 1.587289 | 1.666361 | 1.99279 |
| Eif1 | 1.550277 | 1.238599 | 1.346621 | 1.572697 |
| Gnai2 | 1.331738 | 1.37148 | 1.254206 | 1.484671 |
| Gm8960 | 1.245616 | 1.06002 | 0.906989 | 1.192178 |
| Tmem59 | 1.438734 | 1.211473 | 1.15928 | 1.610429 |
| Slc2a5 | 1.929758 | 1.906715 | 1.585221 | 1.461304 |
| Pmp22 | 1.759766 | 1.434978 | 1.146376 | 1.251715 |
| Rsrp1 | 1.564072 | 1.836131 | 1.501758 | 1.76544 |
| Cd14 | 1.322769 | 1.437928 | 1.08462 | 1.536216 |
| Irf8 | 1.833842 | 2.218168 | 2.201351 | 2.042709 |
| Oaz1 | 2.148566 | 1.574276 | 2.035757 | 2.354388 |
| Rpl11 | 1.718395 | 1.439782 | 1.814278 | 2.296309 |
| Ncoa3 | 1.667356 | 1.752108 | 1.672209 | 1.939489 |
| Mtdh | 1.481203 | 1.455688 | 1.497054 | 1.572907 |
| Rpl14 | 1.814864 | 1.455056 | 1.87959 | 2.222984 |
| Rpl27a | 1.696595 | 1.828062 | 1.707928 | 2.17885 |
| Cd300c2 | 1.609453 | 1.277923 | 1.468513 | 1.681072 |
| Rpl17-ps3 | 1.637407 | 0.915492 | 1.384956 | 1.862409 |
| Lag3 | 1.755123 | 1.349598 | 1.321729 | 1.432556 |
| Fcgr2b | 1.65284 | 1.418118 | 1.432728 | 1.710657 |
| Gm2574 | 1.24959 | 1.200944 | 1.460028 | 1.646162 |
| Notch2 | 1.485874 | 2.167217 | 1.353879 | 1.018852 |
| Rpl18-ps1 | 0.743178 | 0.412844 | 0.507263 | 0.797044 |
| Rpl27-ps3 | 1.482569 | 1.217996 | 0.999472 | 1.982781 |
| Rbm26 | 2.17215 | 1.693803 | 1.645229 | 1.853336 |
| Rnase4 | 1.373006 | 1.154967 | 1.004415 | 1.306803 |
| Rps10 | 1.446878 | 1.386701 | 1.845002 | 2.418265 |
| Gm2225 | 0.809996 | 0.586224 | 0.778937 | 0.867864 |
| Gm10073 | 1.322049 | 0.845773 | 1.279818 | 1.638759 |

| | | | | |
|---|---|---|---|---|
| Gm6180 | 0.856026 | 0.660992 | 0.796812 | 1.31076 |
| mt-Nd4 | 2.116549 | 0.984228 | 1.203822 | 1.585129 |
| Gng10 | 1.504911 | 1.293677 | 1.233132 | 1.795568 |
| Arpc1b | 1.909754 | 1.899704 | 1.897948 | 2.218756 |
| Gm11518 | 2.001585 | 2.085473 | 2.002746 | 2.219933 |
| Gm6170 | 0.851713 | 0.482797 | 0.676858 | 0.841993 |
| Rpl9 | 1.470667 | 1.399218 | 1.859819 | 2.129023 |
| Gm10180 | 1.02287 | 0.647466 | 1.035381 | 1.886676 |
| Rps4x | 1.7499 | 1.798546 | 1.912683 | 2.27485 |
| Gm12166 | 1.754984 | 1.144574 | 1.276885 | 1.766531 |
| Rps23-ps1 | 1.100621 | 0.92486 | 0.865092 | 1.305478 |
| Gm10359 | 0.638734 | 0.939029 | 1.014088 | 1.207221 |
| Arglu1 | 1.620227 | 1.770944 | 1.254678 | 2.130826 |
| H2-DMa | 1.975818 | 1.73845 | 1.815009 | 1.953903 |
| Crlf2 | 3.625684 | 3.891894 | 3.977683 | 3.763698 |
| Gm11560 | 2.411663 | 2.070257 | 2.086433 | 2.083439 |
| Efhd2 | 1.942119 | 1.641744 | 1.9236 | 1.832385 |
| Rps12-ps9 | 1.886103 | 1.568245 | 1.120209 | 1.719802 |
| Gm6988 | 0.834172 | 0.444681 | 0.751339 | 0.865372 |
| Actr2 | 1.674938 | 1.706347 | 1.282983 | 1.515675 |
| Syngr2 | 1.613487 | 1.586564 | 1.617815 | 1.773762 |
| Trf | 1.018683 | 1.077375 | 0.869949 | 0.781274 |
| Tpt1 | 1.378707 | 1.256329 | 2.061929 | 1.973993 |
| Ptms | 3.03432 | 2.555303 | 3.204507 | 3.320143 |
| Arpc3 | 2.028771 | 1.477908 | 1.944483 | 2.167378 |
| Gm26825 | 1.65613 | 1.676247 | 2.026035 | 1.866195 |
| Rps2-ps10 | 1.254964 | 0.858469 | 0.917326 | 1.229096 |
| Susd3 | 1.688339 | 1.623932 | 1.587309 | 1.684072 |
| Gm10443 | 0.853768 | 0.310446 | 0.657536 | 1.03836 |
| Rpl7a-ps7 | 1.255994 | 1.013026 | 0.876599 | 1.249908 |
| Akr1a1 | 1.873321 | 1.734675 | 1.677997 | 2.050587 |
| Capzb | 1.678282 | 1.656293 | 1.881353 | 1.943434 |
| Gm13408 | 0.803509 | 1.095651 | 1.754951 | 0.467101 |
| Gm11361 | 1.099878 | 0.434891 | 0.71593 | 1.120361 |
| Rpl36-ps2 | 2.373697 | 1.788153 | 1.404077 | 2.162687 |
| Rps6-ps4 | 1.282368 | 0.985597 | 1.172278 | 1.51473 |
| Bst2 | 2.152842 | 2.454507 | 2.893208 | 2.707033 |
| Gm10020 | 0.575128 | 0.475072 | 0.568302 | 0.659041 |
| Rpl9-ps6 | 1.647774 | 1.60435 | 1.819286 | 2.395144 |
| Abi3 | 1.944921 | 2.307179 | 2.056604 | 2.065246 |
| Gm3379 | 1.953851 | 1.869461 | 1.932863 | 2.527763 |
| Cd37 | 1.390463 | 1.428584 | 1.706491 | 1.79208 |
| Gm4294 | 0.766988 | 0.484222 | 0.809558 | 0.710767 |
| Psmb8 | 2.195515 | 1.520252 | 2.028744 | 1.877468 |
| Gm8129 | 1.10989 | 0.561381 | 0.573913 | 0.844911 |
| 2-Sep | 1.704993 | 1.329209 | 1.507671 | 1.72816 |
| Arhgdia | 2.035223 | 1.989124 | 2.32131 | 2.33696 |
| Rps10-ps1 | 2.84974 | 2.160187 | 2.927266 | 3.559574 |
| Gm5138 | 1.925415 | 1.908926 | 1.889956 | 2.806632 |

| | | | | |
|---|---|---|---|---|
| Erp29 | 1.716909 | 1.55181 | 1.653904 | 1.861217 |
| Gm7536 | 1.693275 | 1.448209 | 1.79378 | 1.973767 |
| Rps10-ps2 | 1.865515 | 1.684445 | 2.359751 | 2.995389 |
| Cox4i1 | 1.381987 | 1.234884 | 1.406072 | 1.844352 |
| Oaz1-ps | 1.343762 | 1.191042 | 1.354875 | 1.654507 |
| Tpm3 | 2.001244 | 2.02973 | 1.840945 | 2.140432 |
| Tmem86a | 2.138761 | 2.035722 | 2.125881 | 2.304243 |
| Rpl17-ps5 | 0.979823 | 0.461976 | 0.748534 | 1.264193 |
| Gm7336 | 1.308075 | 1.427368 | 1.466289 | 1.89074 |
| Camk1 | 1.792914 | 1.596635 | 1.430564 | 1.759344 |
| Gpx1 | 1.517859 | 1.469835 | 1.564926 | 2.163054 |
| Limd2 | 2.078228 | 1.777571 | 1.958029 | 2.0548 |
| Ptgs1 | 0 | 1.70153 | 1.338351 | 1.008121 |
| Vsir | 0 | 1.085818 | 0.870629 | 0.872972 |
| Gm42418 | 0 | 3.428877 | 2.777821 | 2.619675 |
| Bin1 | 0 | 1.690961 | 1.533892 | 1.577806 |
| Actb | 0 | 0.957061 | 0.683778 | 0.862956 |
| Fcrls | 0 | 1.024678 | 0.66655 | 0.698477 |
| Scamp2 | 0 | 1.383914 | 1.304563 | 1.125847 |
| Gm6394 | 0 | 1.217144 | 1.091227 | 1.718078 |
| Gstm1 | 0 | 0.792965 | 1.747264 | 0.628548 |
| Rpl10-ps3 | 0 | 1.837235 | 1.769331 | 1.823615 |
| Bin2 | 0 | 1.371792 | 1.39158 | 1.396161 |
| Gm6472 | 0 | 0.99951 | 1.380737 | 1.698654 |
| Gm10288 | 0 | 0.968505 | 1.479361 | 1.700698 |
| Lair1 | 0 | 1.435284 | 0.932361 | 1.062429 |
| Cd53 | 0 | 0.854545 | 0.514531 | 0.673961 |
| Wdr89 | 0 | 1.306159 | 1.431016 | 1.518013 |
| Canx | 0 | 1.260602 | 0.87891 | 1.046515 |
| Gm9625 | 0 | 1.010708 | 0.950754 | 1.571793 |
| Gm5835 | 0 | 1.070747 | 1.294243 | 1.939848 |
| Comt | 0 | 1.502834 | 1.366905 | 1.792988 |
| Hspa8 | 0 | 1.039547 | 0.976269 | 1.111462 |
| Gm9616 | 0 | 1.088363 | 1.238281 | 1.700651 |
| Rpl19-ps1 | 0 | 1.217237 | 1.547906 | 2.016105 |
| Ckb | 0 | 1.171959 | 1.054995 | 1.131637 |
| Gm5905 | 0 | 1.279518 | 1.909683 | 2.168356 |
| Ywhae | 0 | 1.838178 | 2.143139 | 1.771441 |
| Upk1b | 0 | 2.344642 | 2.240917 | 2.598007 |
| Gm11263 | 0 | 0.53906 | 0.47439 | 0.645148 |
| Arpc4 | 0 | 1.624277 | 1.737432 | 1.817071 |
| Anxa3 | 0 | 1.48428 | 1.206018 | 1.825881 |
| Gm8430 | 0 | 1.207378 | 1.605817 | 2.142925 |
| Fcgr3 | 0 | 0.864799 | 0.706402 | 1.113953 |
| Coro1a | 0 | 1.217524 | 1.485882 | 1.283372 |
| Rps13-ps2 | 0 | 1.101884 | 1.594639 | 2.480455 |
| Rpl31-ps8 | 0 | 1.24587 | 1.923297 | 2.318117 |
| Rpl10a-ps1 | 0 | 1.350115 | 1.498721 | 2.181632 |
| Gm4617 | 0 | 0.925782 | 0.862058 | 1.243265 |

| | | | | |
|---|---|---|---|---|
| Gm11539 | 0 | 0.664386 | 0.88314 | 1.14326 |
| Gm15772 | 0 | 0.978391 | 1.376249 | 1.605647 |
| Rps27a-ps | 0 | 1.14454 | 1.168289 | 1.687593 |
| Rbm39 | 0 | 1.299511 | 1.365798 | 1.34843 |
| Ppib | 0 | 1.43322 | 1.419518 | 1.909689 |
| Rpl37rt | 0 | 0.441914 | 1.174275 | 1.741759 |
| Cyth4 | 0 | 1.24153 | 1.282793 | 1.220337 |
| Clta | 0 | 1.294058 | 1.572263 | 1.786629 |
| Actg1 | 0 | 1.112642 | 1.184817 | 1.429845 |
| Gm10231 | 0 | 1.139774 | 1.138176 | 1.326499 |
| Rpl28-ps1 | 0 | 1.387646 | 1.409577 | 2.218997 |
| Arpc2 | 0 | 1.277025 | 1.297734 | 1.355499 |
| Gm17241 | 0 | 0.650881 | 1.077016 | 1.097693 |
| Gm13493 | 0 | 0.800915 | 0.780471 | 1.271722 |
| Rps3a2 | 0 | 0.743823 | 1.00418 | 1.213674 |
| Gm5805 | 0 | 0.519711 | 0.885323 | 1.137748 |
| Gm8226 | 0 | 0.629074 | 0.744765 | 0.853652 |
| Gm6063 | 0 | 0.549598 | 0.380648 | 0.68057 |
| Abcg1 | 0 | 2.180639 | 1.846803 | 2.554575 |
| Mbp | 0 | 1.460632 | 0.681173 | 1.152075 |
| Tmed9 | 0 | 2.150288 | 1.904838 | 2.457202 |
| Rhoa | 0 | 1.305392 | 1.144893 | 1.027397 |
| Gm5514 | 0 | 0.929238 | 1.085282 | 1.52826 |
| Rpl21-ps15 | 0 | 0.642378 | 1.190289 | 1.642318 |
| Mafb | 0 | 1.490053 | 1.222805 | 1.102831 |
| Rps3a3 | 0 | 0.38854 | 0.561459 | 0.738326 |
| Foxn3 | 0 | 1.59864 | 1.297081 | 1.550478 |
| Rps19-ps6 | 0 | 0.919439 | 1.754181 | 2.21126 |
| Rpl15-ps3 | 0 | 0.659682 | 1.29542 | 1.145835 |
| Rpl13a-ps1 | 0 | 0.589562 | 0.817531 | 0.903667 |
| Gm10240 | 0 | 0.749566 | 1.426953 | 1.877571 |
| Rpl13-ps6 | 0 | 0.816658 | 1.451753 | 1.877748 |
| Rps13-ps1 | 0 | 0.381531 | 0.560858 | 0.99359 |
| Gm13680 | 0 | 1.060295 | 1.302016 | 1.384691 |
| Gm10163 | 0 | 2.133043 | 0.921257 | 1.712738 |
| Gm11808 | 0 | 0.964839 | 1.30636 | 1.725118 |
| Tpt1-ps3 | 0 | 0.709838 | 1.245613 | 1.356177 |
| Gm12191 | 0 | 0.781214 | 0.73778 | 0.832259 |
| Gm10275 | 0 | 1.043505 | 1.24469 | 1.990369 |
| Rpl14-ps1 | 0 | 1.204511 | 1.232659 | 1.809091 |
| Tmsb10 | 0 | 2.58056 | 2.783085 | 3.392263 |
| Gm14303 | 0 | 0.644223 | 1.198158 | 1.926674 |
| Commd8 | 0 | 1.558957 | 1.559775 | 1.608344 |
| Gm9493 | 0 | 0.454971 | 0.596369 | 0.538052 |
| Rpl34-ps1 | 0 | 0.902876 | 1.337303 | 1.627178 |
| Arf1 | 0 | 1.305091 | 1.209018 | 1.548021 |
| Gm8172 | 0 | 1.31163 | 2.009907 | 1.945374 |
| Cmtm7 | 0 | 1.253195 | 1.265788 | 1.523878 |
| Gm9794 | 0 | 0.884081 | 1.26777 | 1.827655 |

| | | | | |
|---|---|---|---|---|
| Gm10051 | 0 | 0.739943 | 1.329301 | 1.373908 |
| Rps25-ps1 | 0 | 0.36023 | 0.492636 | 0.846573 |
| Arhgdib | 0 | 1.476896 | 1.457229 | 1.847444 |
| Creg1 | 0 | 1.444701 | 1.244524 | 1.514531 |
| Gna15 | 0 | 1.569851 | 1.725454 | 2.006491 |
| Rpsa-ps10 | 0 | 0.910212 | 1.470257 | 2.240851 |
| Usp2 | 0 | 1.982909 | 1.917687 | 2.044362 |
| Alox5ap | 0 | 1.245026 | 1.586621 | 1.935283 |
| Hmgb1 | 0 | 1.2475 | 1.366431 | 1.65919 |
| Gm9703 | 0 | 0.269305 | 0.627955 | 0.995892 |
| Rps15a-ps | 0 | 1.469312 | 0.717335 | 1.148833 |
| Gm3511 | 0 | 0.471155 | 1.055675 | 1.924167 |
| Spcs2 | 0 | 1.189718 | 1.160132 | 1.569723 |
| Gm6136 | 0 | 1.239014 | 1.595165 | 1.916098 |
| Jade2 | 0 | 0.840248 | 0.456367 | 0.418072 |
| Ddost | 0 | 1.561419 | 1.218513 | 1.695986 |
| Rpl30-ps1( | 0 | 0.451244 | 0.7389 | 0.79962 |
| Arl6ip1 | 0 | 1.240972 | 1.575046 | 1.521399 |
| Ltc4s | 0 | 1.294829 | 1.39288 | 1.533685 |
| Gm15501 | 0 | 1.121339 | 1.011586 | 1.603664 |
| Selenof | 0 | 1.246997 | 1.352497 | 1.806307 |
| Rps24-ps3 | 0 | 0.472054 | 0.606043 | 1.19999 |
| Eno1 | 0 | 1.60145 | 1.442814 | 1.633579 |
| Itpripl1 | 0 | 1.652763 | 1.922356 | 2.095094 |
| Ptp4a2 | 0 | 1.628269 | 1.537307 | 1.95986 |
| Tcn2 | 0 | 0.948641 | 1.014849 | 1.13278 |
| Pycard | 0 | 1.18753 | 1.411097 | 1.818515 |
| Gm10059 | 0 | 0.264259 | 0.548888 | 0.789036 |
| Saraf | 0 | 1.096812 | 0.921542 | 1.315997 |
| Snhg20 | 0 | 2.152966 | 2.27785 | 2.663734 |
| Selenok | 0 | 1.494347 | 1.328614 | 1.893628 |
| Tapbp | 0 | 1.304276 | 1.145339 | 1.139881 |
| Gm9385 | 0 | 0.550738 | 1.047399 | 1.126553 |
| Hsp90b1 | 0 | 0.630139 | 0.786031 | 0.800607 |
| B4galt1 | 0 | 1.257671 | 1.214324 | 1.17545 |
| Atp5b | 0 | 1.206902 | 1.003296 | 1.254857 |
| Gm7331 | 0 | 0.850808 | 1.268769 | 1.760405 |
| Lmo2 | 0 | 1.914816 | 1.809351 | 1.812544 |
| Aldoa | 0 | 1.444826 | 1.348043 | 1.918654 |
| Mag | 0 | 0.938716 | 0.992054 | 1.886029 |
| Rpl7a | 0 | 1.495987 | 1.831657 | 2.334773 |
| Rgs2 | 0 | 1.089901 | 1.467402 | 1.257548 |
| Gm10925 | 2.32933 | 0 | 1.009753 | 1.410913 |
| Gm28437 | 1.927982 | 0 | 0.746273 | 1.105018 |
| Gm10076 | 1.826505 | 0 | 1.643016 | 2.307388 |
| Rpl41 | 1.392128 | 0 | 1.412882 | 1.826804 |
| Fau | 1.429503 | 0 | 1.499493 | 1.937212 |
| Rpl35 | 1.767814 | 0 | 1.738985 | 2.112555 |
| Rps25 | 1.588249 | 0 | 1.732441 | 2.086762 |

| | | | | |
|---|---|---|---|---|
| Nmt1 | 0.565682 | 0 | 1.030431 | 0.615393 |
| Wdr26 | 0.334631 | 0 | 0.722432 | 0.413853 |
| Rpl30 | 1.42613 | 0 | 1.437638 | 2.004846 |
| Fmnl1 | 1.311434 | 0 | 0.957135 | 0.538242 |
| Ly6e | 1.60691 | 0 | 1.649376 | 1.459109 |
| Rpl37a | 1.144451 | 0 | 1.300962 | 1.843336 |
| Wipf1 | 1.612904 | 0 | 0.282203 | 0.439317 |
| Sod1 | 2.422648 | 0 | 2.206669 | 2.361647 |
| Rpl23a | 1.63444 | 0 | 1.58335 | 2.000152 |
| Rps27 | 1.119585 | 0 | 1.001795 | 1.629192 |
| Lgals3bp | 2.567349 | 0 | 2.45435 | 2.419248 |
| Atrx | 1.166192 | 0 | 0.561992 | 0.585962 |
| Rplp2 | 1.365017 | 0 | 1.64675 | 2.027181 |
| Cmklr1 | 0.519033 | 0 | 0.760388 | 0.561803 |
| Rps23 | 1.182986 | 0 | 1.204458 | 1.755436 |
| Lnpep | 0.51228 | 0 | 0.306868 | 0.270636 |
| Rpl36 | 1.352362 | 0 | 1.35004 | 1.907968 |
| Hnrnpl | 1.613935 | 0 | 0.734001 | 1.907324 |
| Kmt2a | 0.344776 | 0 | 0.439032 | 0.951613 |
| Pkn2 | 0.292473 | 0 | 0.473023 | 0.355873 |
| Chd2 | 0.875277 | 0 | 0.305489 | 0.958516 |
| Gm34280 | 0.350975 | 0 | 0.286235 | 1.171283 |
| Rnf216 | 0.289626 | 0 | 0.375307 | 0.707183 |
| Rpl23 | 1.66073 | 0 | 1.511259 | 1.882585 |
| Rps24 | 1.746327 | 0 | 1.923932 | 2.384062 |
| Rps16 | 1.331682 | 0 | 1.484945 | 2.07003 |
| Fus | 0.98597 | 0 | 0.753178 | 1.268613 |
| Arhgap39 | 0.304169 | 0 | 0.682047 | 0.291263 |
| Rptor | 0.801388 | 0 | 0.738942 | 0.380143 |
| Ash1l | 0.43023 | 0 | 0.327984 | 0.431722 |
| Rpl34 | 1.329807 | 0 | 1.327315 | 1.888341 |
| Rps5 | 1.852775 | 0 | 1.814556 | 2.479619 |
| Rbbp6 | 0.685781 | 0 | 0.325764 | 0.318185 |
| Atf7 | 0.292012 | 0 | 0.393778 | 0.297625 |
| Dock2 | 0.490534 | 0 | 0.539196 | 0.315503 |
| Gm16192 | 0.867348 | 0 | 0.498219 | 0.350482 |
| Ube2h | 0.678847 | 0 | 0.450965 | 0.279515 |
| Gpr146 | 0.349727 | 0 | 0.514819 | 0.528338 |
| Fam117b | 0.324521 | 0 | 0.461579 | 0.293957 |
| Cic | 0.735795 | 0 | 1.21664 | 1.988948 |
| Rapgef1 | 0.517573 | 0 | 0.825583 | 0.507854 |
| Taok2 | 0.407927 | 0 | 0.66226 | 0.457139 |
| Git2 | 0.597867 | 0 | 0.481489 | 0.428745 |
| Slc12a6 | 0.948546 | 0 | 0.268567 | 0.733308 |
| Rpl24 | 1.994085 | 0 | 1.955906 | 2.088852 |
| Dock8 | 0.411401 | 0 | 0.38882 | 0.389005 |
| Ncbp3 | 0.695888 | 0 | 0.710571 | 0.259917 |
| A630001G | 0.491222 | 0 | 0.267742 | 0.637996 |
| Mtr | 1.035064 | 0 | 0.269172 | 0.50884 |

| | | | | |
|---|---|---|---|---|
| Dip2b | 1.003449 | 0 | 0.935506 | 0.627288 |
| Wdfy4 | 0.829199 | 0 | 0.637966 | 0.342835 |
| Akap13 | 0.803734 | 0 | 0.604171 | 0.780585 |
| Myo18a | 0.790781 | 0 | 0.669688 | 0.877121 |
| Ankrd11 | 0.759591 | 0 | 0.741367 | 0.77737 |
| Snn | 0.767675 | 0 | 0.824477 | 1.673404 |
| Irf2bp2 | 1.514053 | 0 | 1.202205 | 0.620135 |
| Acin1 | 0.640597 | 0 | 0.684768 | 1.138877 |
| Rps13 | 1.475928 | 0 | 1.49884 | 2.086985 |
| Olfr620 | 0.536932 | 0 | 0.414024 | 0.708653 |
| Nipbl | 1.036392 | 0 | 0.374537 | 0.670987 |
| Slc36a1 | 0.696364 | 0 | 0.343929 | 0.987219 |
| Dcaf8 | 0.630386 | 0 | 0.319183 | 0.380334 |
| Med12l | 0.532253 | 0 | 0.340486 | 0.307905 |
| Sh3bgrl3 | 1.280194 | 0 | 1.293199 | 1.838584 |
| Rpl28 | 1.655302 | 0 | 1.995416 | 2.093321 |
| Mvb12b | 1.615779 | 0 | 0.745748 | 0.969147 |
| Rpl15 | 1.395062 | 0 | 1.345511 | 1.547667 |
| Prex1 | 0.508431 | 0 | 0.572765 | 0.450154 |
| Kpna4 | 1.126832 | 0 | 0.934236 | 1.131912 |
| Gabarap | 1.596559 | 0 | 1.620817 | 1.755401 |
| Rpl37 | 0.981517 | 0 | 0.97006 | 1.633286 |
| Arid3b | 1.059772 | 0 | 0.594139 | 0.563798 |
| Rpl26 | 1.470994 | 0 | 1.696402 | 2.036539 |
| Chn2 | 1.132347 | 0 | 0.991086 | 0.653838 |
| Rps28 | 0.958984 | 0 | 0.778343 | 1.56566 |
| Dync1h1 | 0.478272 | 0 | 0.917861 | 0.44552 |
| Dock4 | 0.97945 | 0 | 0.362135 | 0.662501 |
| Ccdc50 | 1.015863 | 0 | 0.647059 | 0.441404 |
| Bicd2 | 0.415676 | 0 | 0.470294 | 0.330251 |
| Gm26542 | 1.049094 | 0 | 0.889619 | 1.493367 |
| Arhgap30 | 1.037439 | 0 | 0.696404 | 0.401266 |
| Atp7a | 0.537925 | 0 | 0.433282 | 0.627072 |
| Skp1a | 0.769902 | 0 | 0.470678 | 0.754938 |
| Dgkd | 0.491309 | 0 | 0.796247 | 0.49416 |
| Rps27a | 1.29063 | 0 | 1.451361 | 1.904917 |
| Pds5a | 0.818425 | 0 | 0.610394 | 0.660573 |
| Shisa5 | 1.914188 | 0 | 2.02474 | 1.908442 |
| Rps15a | 1.666145 | 0 | 1.539449 | 2.062938 |
| Arhgef40 | 1.206141 | 0 | 0.9606 | 0.50949 |
| Serf2 | 1.835308 | 0 | 1.657832 | 1.767436 |
| Zmiz1 | 0.755637 | 0 | 0.684677 | 0.421452 |
| Dido1 | 1.433078 | 0 | 1.177149 | 1.197682 |
| Ubtf | 0.619494 | 0 | 0.420169 | 0.42768 |
| Btbd1 | 0.35418 | 0 | 0.337368 | 0.284625 |
| Gm10132 | 2.012176 | 0 | 1.857206 | 2.046042 |
| Anp32a | 0.594782 | 0 | 0.622653 | 0.883751 |
| Tgoln1 | 0.697018 | 0 | 0.563342 | 0.416609 |
| Usp12 | 0.537011 | 0 | 0.620072 | 0.37829 |

| | | | | |
|---|---|---|---|---|
| Ppp1ca | 1.766499 | 0 | 1.475229 | 1.851801 |
| Ctcf | 0.638462 | 0 | 0.267411 | 1.043319 |
| Otud5 | 0.640716 | 0 | 0.857541 | 0.63179 |
| Pxk | 0.614113 | 0 | 0.590949 | 0.636246 |

Supplementry Table 6.  Downregulated aging DEGs shared by brain regions

| Gene | Cb | Ctx | Hipc | Str |
|---|---|---|---|---|
| Gm15564 | -1.41575 | -0.34221 | -0.37258 | -3.23836 |
| Gcg | -2.73344 | -2.69023 | -1.25489 | -2.84839 |
| Ins2 | -1.16339 | -1.99923 | -2.00211 | -1.21769 |
| Ins1 | -0.68319 | -1.04135 | -1.39192 | -0.68627 |
| Ac154912. | -1.37631 | -1.99699 | -3.08926 | -1.61895 |
| Rn7sk | -0.93987 | -1.13173 | -0.98238 | -1.01271 |
| Fkbp5 | -0.80071 | -0.45005 | -1.14891 | -1.29934 |
| Serinc3 | -1.05051 | -0.55233 | -0.33419 | -1.59243 |
| Fos | -3.00161 | -1.21245 | -1.59491 | -1.23692 |
| Jun | -1.45783 | -0.43518 | -1.39806 | -1.04278 |
| Tagap | -0.73537 | -0.7139 | -0.42757 | -1.18341 |
| Dusp1 | -1.81031 | -1.20671 | -1.58405 | -1.04948 |
| Per1 | -0.26862 | -1.08923 | -1.0014 | -1.77514 |
| Hist1h4d | -1.1064 | -0.60349 | -1.88335 | -1.16081 |
| Egr1 | -0.38661 | -1.26049 | -2.65285 | -2.31788 |
| Mfsd14b | -0.94983 | -0.56709 | -0.33762 | -1.75247 |
| Acly | -0.45772 | 0 | -0.81466 | -1.11385 |
| Stt3b | -0.43722 | 0 | -0.27948 | -0.69996 |
| Crk | -0.32664 | 0 | -0.66869 | -0.96217 |
| Manba | -0.37097 | 0 | -0.41156 | -0.70633 |
| Tlr4 | -0.67801 | 0 | -0.26533 | -1.35988 |
| Zfp871 | -0.84948 | 0 | -0.5609 | -0.80469 |
| Tlr13 | -0.5207 | 0 | -0.31777 | -0.43174 |
| Rgs1 | -0.27919 | 0 | -1.4392 | -0.65572 |
| Gm45221 | -1.88948 | 0 | -1.33089 | -2.31175 |
| Ccr5 | -0.46303 | 0 | -0.36226 | -1.0053 |
| Apc | -0.77463 | 0 | -0.48618 | -0.60851 |
| Rapgef5 | -0.3423 | 0 | -0.3426 | -0.8706 |
| G530011O | -0.25962 | 0 | -0.49719 | -0.25244 |
| Chka | -0.60348 | 0 | -0.35011 | -0.52903 |
| Gm45223 | -0.41 | 0 | -0.67602 | -0.97563 |
| Nptn | -0.86079 | 0 | -0.38987 | -1.37191 |
| Dhx9 | -0.64012 | 0 | -1.04374 | -0.27939 |
| Slc8a1 | -1.65971 | 0 | -2.03639 | -1.95521 |
| Abca1 | -0.66105 | 0 | -0.48171 | -0.71941 |
| Jmjd1c | -0.33916 | 0 | -1.01607 | -0.84239 |
| Runx1 | -1.042 | 0 | -0.80895 | -0.28304 |
| Tlr7 | -0.57398 | 0 | -0.67788 | -0.86098 |
| Ac154912. | -0.38465 | 0 | -1.45823 | -1.19294 |
| Adrb1 | -0.39148 | 0 | -0.30181 | -1.06879 |
| Gm44860 | -0.98753 | 0 | -1.28051 | -1.10554 |
| Chga | -0.60052 | 0 | -0.79141 | -0.85425 |

Supplementry Table 7.  Upregulated aging DEGs unique to barin regions

| Gene | Cb | Ctx | Hipc | Str |
|---|---|---|---|---|
| Sfpq | 0.410369 | 0 | 0 | 0 |
| Rassf2 | 0.493294 | 0 | 0 | 0 |
| Mdm4 | 0.43749 | 0 | 0 | 0 |
| Gm37472 | 0.851315 | 0 | 0 | 0 |
| Pik3r1 | 0.548371 | 0 | 0 | 0 |
| Atf6 | 0.266761 | 0 | 0 | 0 |
| Picalm | 0.42725 | 0 | 0 | 0 |
| Ppp1r10 | 0.798834 | 0 | 0 | 0 |
| Ppp1r12c | 0.518374 | 0 | 0 | 0 |
| Cltc | 0.728031 | 0 | 0 | 0 |
| H2-Q7 | 4.811146 | 0 | 0 | 0 |
| Xpr1 | 0.349274 | 0 | 0 | 0 |
| Bod1l | 0.714144 | 0 | 0 | 0 |
| Map3k3 | 0.684861 | 0 | 0 | 0 |
| Cd2ap | 0.470347 | 0 | 0 | 0 |
| Itsn2 | 0.29636 | 0 | 0 | 0 |
| Phf14 | 0.554766 | 0 | 0 | 0 |
| Prrc2a | 1.010502 | 0 | 0 | 0 |
| Hps3 | 0.563054 | 0 | 0 | 0 |
| Rasa1 | 0.402731 | 0 | 0 | 0 |
| Dock10 | 0.520146 | 0 | 0 | 0 |
| Setd2 | 0.404143 | 0 | 0 | 0 |
| Stab1 | 0.542657 | 0 | 0 | 0 |
| Klhl18 | 0.61246 | 0 | 0 | 0 |
| Xiap | 0.348593 | 0 | 0 | 0 |
| Ppp1r9a | 0.498738 | 0 | 0 | 0 |
| Stk4 | 0.542214 | 0 | 0 | 0 |
| Cdc37l1 | 1.256632 | 0 | 0 | 0 |
| Itga6 | 0.336645 | 0 | 0 | 0 |
| Gm26857 | 0.465985 | 0 | 0 | 0 |
| Zfp292 | 0.561944 | 0 | 0 | 0 |
| Mlxipl | 1.409166 | 0 | 0 | 0 |
| Nav3 | 0.616858 | 0 | 0 | 0 |
| Taok1 | 0.655178 | 0 | 0 | 0 |
| Lfng | 1.81382 | 0 | 0 | 0 |
| Kidins220 | 0.471462 | 0 | 0 | 0 |
| Dtx4 | 0.560278 | 0 | 0 | 0 |
| Dennd4a | 0.543068 | 0 | 0 | 0 |
| Dazap1 | 0.941479 | 0 | 0 | 0 |
| Herc4 | 0.419068 | 0 | 0 | 0 |
| H2-Q6 | 3.686993 | 0 | 0 | 0 |
| N4bp2l2 | 0.394492 | 0 | 0 | 0 |
| Ktn1 | 0.665137 | 0 | 0 | 0 |
| Ogt | 0.886102 | 0 | 0 | 0 |
| Myo1c | 1.267614 | 0 | 0 | 0 |
| Zc3h4 | 0.348201 | 0 | 0 | 0 |

| | | | | |
|---|---|---|---|---|
| Rps6kb1 | 0.609749 | 0 | 0 | 0 |
| Rbpj | 0.573133 | 0 | 0 | 0 |
| Ascc3 | 1.07264 | 0 | 0 | 0 |
| BC037034 | 0.740428 | 0 | 0 | 0 |
| 1810026B( | 1.519566 | 0 | 0 | 0 |
| Ap3b1 | 0.525233 | 0 | 0 | 0 |
| Fnip1 | 0.597143 | 0 | 0 | 0 |
| Rab43 | 0.719313 | 0 | 0 | 0 |
| Ctnna1 | 0.792313 | 0 | 0 | 0 |
| Rdx | 0.648936 | 0 | 0 | 0 |
| Clk4 | 0.699183 | 0 | 0 | 0 |
| Klf6 | 0.524913 | 0 | 0 | 0 |
| Zeb2 | 0.361147 | 0 | 0 | 0 |
| Vav2 | 1.288208 | 0 | 0 | 0 |
| Smchd1 | 0.543794 | 0 | 0 | 0 |
| Dst | 0.461697 | 0 | 0 | 0 |
| Sec16a | 0.354995 | 0 | 0 | 0 |
| Cnot1 | 0.642559 | 0 | 0 | 0 |
| Anks1 | 0.403822 | 0 | 0 | 0 |
| Wdfy2 | 0.823514 | 0 | 0 | 0 |
| Ddx6 | 0.426965 | 0 | 0 | 0 |
| Stag2 | 0.401143 | 0 | 0 | 0 |
| Ddx21 | 0.743388 | 0 | 0 | 0 |
| Arhgef12 | 0.256127 | 0 | 0 | 0 |
| Uso1 | 0.628717 | 0 | 0 | 0 |
| Atf4 | 0.770692 | 0 | 0 | 0 |
| Zfp148 | 0.55727 | 0 | 0 | 0 |
| Epb41l2 | 0.313398 | 0 | 0 | 0 |
| Trmt1 | 0.347075 | 0 | 0 | 0 |
| Zmym4 | 0.394311 | 0 | 0 | 0 |
| Hnrnph1 | 0.307429 | 0 | 0 | 0 |
| Tmem209 | 0.436597 | 0 | 0 | 0 |
| Wdr44 | 0.437452 | 0 | 0 | 0 |
| Ubr2 | 0.4053 | 0 | 0 | 0 |
| Sema4d | 0.507315 | 0 | 0 | 0 |
| Phip | 0.41805 | 0 | 0 | 0 |
| Map4 | 0.812732 | 0 | 0 | 0 |
| Tln1 | 0.574028 | 0 | 0 | 0 |
| Brd9 | 0.58897 | 0 | 0 | 0 |
| 4833420G: | 0.30871 | 0 | 0 | 0 |
| Tcf12 | 0.447775 | 0 | 0 | 0 |
| Lasp1 | 0.403722 | 0 | 0 | 0 |
| Iffo1 | 1.130971 | 0 | 0 | 0 |
| Bptf | 0.801548 | 0 | 0 | 0 |
| Zfp36l1 | 0.358339 | 0 | 0 | 0 |
| Plcl2 | 0.316038 | 0 | 0 | 0 |
| Dennd5a | 0.465611 | 0 | 0 | 0 |
| Rbm33 | 0.799274 | 0 | 0 | 0 |
| Fmnl3 | 0.677178 | 0 | 0 | 0 |

| | | | | |
|---|---|---|---|---|
| Prpf38b | 1.138036 | 0 | 0 | 0 |
| Mark2 | 0.847178 | 0 | 0 | 0 |
| Ophn1 | 0.416986 | 0 | 0 | 0 |
| Aak1 | 0.306731 | 0 | 0 | 0 |
| Mtmr3 | 0.264066 | 0 | 0 | 0 |
| Fam149b | 0.53687 | 0 | 0 | 0 |
| Ppp1r9b | 1.413081 | 0 | 0 | 0 |
| Smarca4 | 0.728364 | 0 | 0 | 0 |
| Snrnp70 | 1.435689 | 0 | 0 | 0 |
| Cbl | 0.387957 | 0 | 0 | 0 |
| Arid3a | 1.394898 | 0 | 0 | 0 |
| Gdi1 | 0.722 | 0 | 0 | 0 |
| Cyld | 1.079405 | 0 | 0 | 0 |
| Strada | 0.443633 | 0 | 0 | 0 |
| Itch | 0.897727 | 0 | 0 | 0 |
| Vps13c | 0.423316 | 0 | 0 | 0 |
| Bmpr2 | 0.343341 | 0 | 0 | 0 |
| Safb2 | 1.086164 | 0 | 0 | 0 |
| Sall1 | 0.715295 | 0 | 0 | 0 |
| Trim8 | 0.279158 | 0 | 0 | 0 |
| 9930017N2 | 0.968665 | 0 | 0 | 0 |
| Arid4a | 0.955202 | 0 | 0 | 0 |
| Hltf | 0.282743 | 0 | 0 | 0 |
| Rassf3 | 0.666072 | 0 | 0 | 0 |
| Smurf2 | 0.486081 | 0 | 0 | 0 |
| Gtf2ird2 | 1.1626 | 0 | 0 | 0 |
| Dctn6 | 0.419426 | 0 | 0 | 0 |
| Tdrd7 | 0.299886 | 0 | 0 | 0 |
| Trim30d | 1.096303 | 0 | 0 | 0 |
| Tnfrsf11a | 0.778022 | 0 | 0 | 0 |
| Washc2 | 0.890095 | 0 | 0 | 0 |
| Rc3h2 | 0.336747 | 0 | 0 | 0 |
| Ppp6r3 | 0.6188 | 0 | 0 | 0 |
| Odr4 | 0.394888 | 0 | 0 | 0 |
| Hdac4 | 1.358456 | 0 | 0 | 0 |
| Irf2bpl | 0.836541 | 0 | 0 | 0 |
| Gm21092 | 0.507394 | 0 | 0 | 0 |
| E230029C | 0.54291 | 0 | 0 | 0 |
| Prrc1 | 1.258209 | 0 | 0 | 0 |
| Itgam | 0.89599 | 0 | 0 | 0 |
| Gm15503 | 0.540674 | 0 | 0 | 0 |
| Cspp1 | 0.698918 | 0 | 0 | 0 |
| U2af2 | 1.437492 | 0 | 0 | 0 |
| Xpo6 | 0.367733 | 0 | 0 | 0 |
| Reep3 | 0.352909 | 0 | 0 | 0 |
| Cnot6l | 0.549569 | 0 | 0 | 0 |
| Myo9b | 0.795487 | 0 | 0 | 0 |
| Mroh1 | 0.556233 | 0 | 0 | 0 |
| Dclre1c | 0.306113 | 0 | 0 | 0 |

| | | | | |
|---|---|---|---|---|
| Tmcc3 | 0.367444 | 0 | 0 | 0 |
| Slain2 | 0.416359 | 0 | 0 | 0 |
| Ppp4r1l-ps | 0.40586 | 0 | 0 | 0 |
| Mogs | 0.279308 | 0 | 0 | 0 |
| Msn | 0.430919 | 0 | 0 | 0 |
| Sart1 | 0.566554 | 0 | 0 | 0 |
| Lgals8 | 0.852258 | 0 | 0 | 0 |
| Nfam1 | 0.429499 | 0 | 0 | 0 |
| Ep400 | 0.570109 | 0 | 0 | 0 |
| Kif5b | 0.499752 | 0 | 0 | 0 |
| Ubn2 | 0.67295 | 0 | 0 | 0 |
| Cmpk1 | 0.589653 | 0 | 0 | 0 |
| Pxn | 0.738562 | 0 | 0 | 0 |
| Elf4 | 0.761515 | 0 | 0 | 0 |
| Ppp4r3b | 0.413335 | 0 | 0 | 0 |
| Cept1 | 0.698971 | 0 | 0 | 0 |
| Rock1 | 0.896398 | 0 | 0 | 0 |
| Otud4 | 0.471642 | 0 | 0 | 0 |
| Gtpbp1 | 0.483859 | 0 | 0 | 0 |
| Snx10 | 0.565895 | 0 | 0 | 0 |
| Psme4 | 1.059187 | 0 | 0 | 0 |
| Tmod3 | 0.422486 | 0 | 0 | 0 |
| Zbtb4 | 1.097132 | 0 | 0 | 0 |
| Pdlim4 | 0.548503 | 0 | 0 | 0 |
| Mib1 | 0.886909 | 0 | 0 | 0 |
| Lcp2 | 0.619148 | 0 | 0 | 0 |
| Eif3a | 0.717442 | 0 | 0 | 0 |
| Ifitm3 | 2.017061 | 0 | 0 | 0 |
| Fam193b | 0.874871 | 0 | 0 | 0 |
| Kmt2e | 1.216326 | 0 | 0 | 0 |
| Cdk12 | 1.029612 | 0 | 0 | 0 |
| Zzef1 | 0.801816 | 0 | 0 | 0 |
| Fam102b | 0.645052 | 0 | 0 | 0 |
| Cfap74 | 0.744447 | 0 | 0 | 0 |
| Zfp467 | 0.735736 | 0 | 0 | 0 |
| Tmem273 | 1.115298 | 0 | 0 | 0 |
| 1700020I1 | 0.806121 | 0 | 0 | 0 |
| Sh3bp2 | 0.888266 | 0 | 0 | 0 |
| Bcl6 | 1.405856 | 0 | 0 | 0 |
| Ash2l | 0.805677 | 0 | 0 | 0 |
| Plxnb2 | 0.478229 | 0 | 0 | 0 |
| A130071D | 2.079771 | 0 | 0 | 0 |
| Gm21811 | 0.853721 | 0 | 0 | 0 |
| Ifi27l2a | 1.837652 | 0 | 0 | 0 |
| Ecpas | 0.547574 | 0 | 0 | 0 |
| Vps4a | 1.00436 | 0 | 0 | 0 |
| Fcho2 | 0.871814 | 0 | 0 | 0 |
| Dennd1c | 1.231902 | 0 | 0 | 0 |
| Abcb1b | 1.198866 | 0 | 0 | 0 |

| | | | | |
|---|---|---|---|---|
| Ube2r2 | 1.0295 | 0 | 0 | 0 |
| Slc25a37 | 0.628511 | 0 | 0 | 0 |
| Mgat4b | 0.55493 | 0 | 0 | 0 |
| Csnk1d | 0.764735 | 0 | 0 | 0 |
| Pi4k2a | 0.66652 | 0 | 0 | 0 |
| Dock11 | 0.460377 | 0 | 0 | 0 |
| Mbnl1 | 0.695469 | 0 | 0 | 0 |
| Snx30 | 0.828307 | 0 | 0 | 0 |
| Cds1 | 0.625602 | 0 | 0 | 0 |
| Atf1 | 0.714003 | 0 | 0 | 0 |
| Map3k5 | 0.409199 | 0 | 0 | 0 |
| Ttc28 | 1.573782 | 0 | 0 | 0 |
| Pbrm1 | 0.870194 | 0 | 0 | 0 |
| Ski | 1.098127 | 0 | 0 | 0 |
| Arcn1 | 0.568711 | 0 | 0 | 0 |
| Plekha1 | 1.592616 | 0 | 0 | 0 |
| Agmo | 0.37287 | 0 | 0 | 0 |
| Mycbp2 | 0.560089 | 0 | 0 | 0 |
| D5Ertd579 | 0.485233 | 0 | 0 | 0 |
| Csk | 0.98384 | 0 | 0 | 0 |
| Clk1 | 1.154481 | 0 | 0 | 0 |
| Trappc8 | 0.693474 | 0 | 0 | 0 |
| Uvrag | 0.498592 | 0 | 0 | 0 |
| Npl | 0.437872 | 0 | 0 | 0 |
| Ccdc82 | 0.759948 | 0 | 0 | 0 |
| Osbpl9 | 0.628713 | 0 | 0 | 0 |
| Fnbp4 | 0.927071 | 0 | 0 | 0 |
| Acadm | 0.616052 | 0 | 0 | 0 |
| Ptprc | 0.647875 | 0 | 0 | 0 |
| Smarcc2 | 0.629066 | 0 | 0 | 0 |
| Acox3 | 0.556698 | 0 | 0 | 0 |
| Cep350 | 0.357704 | 0 | 0 | 0 |
| Stx2 | 0.859286 | 0 | 0 | 0 |
| Htt | 0.303462 | 0 | 0 | 0 |
| Vrk1 | 0.42501 | 0 | 0 | 0 |
| Dnajc13 | 0.495636 | 0 | 0 | 0 |
| Hipk1 | 0.532088 | 0 | 0 | 0 |
| Cbfa2t3 | 0.909568 | 0 | 0 | 0 |
| Colgalt1 | 0.561364 | 0 | 0 | 0 |
| Casp4 | 0.950685 | 0 | 0 | 0 |
| Zkscan3 | 0.690785 | 0 | 0 | 0 |
| Thrsp | 2.821564 | 0 | 0 | 0 |
| Itgb3 | 0.583792 | 0 | 0 | 0 |
| Hps4 | 0.667841 | 0 | 0 | 0 |
| Cebpz | 1.273212 | 0 | 0 | 0 |
| Kif1b | 1.278804 | 0 | 0 | 0 |
| Zup1 | 0.858764 | 0 | 0 | 0 |
| P2rx7 | 0.814059 | 0 | 0 | 0 |
| Gpr107 | 0.560061 | 0 | 0 | 0 |

| | | | | |
|---|---|---|---|---|
| Gpat4 | 0.625661 | 0 | 0 | 0 |
| Pitpna | 1.222955 | 0 | 0 | 0 |
| Snx4 | 0.716899 | 0 | 0 | 0 |
| Tnip2 | 0.761012 | 0 | 0 | 0 |
| Tmem127 | 0.697384 | 0 | 0 | 0 |
| Cpsf6 | 0.535715 | 0 | 0 | 0 |
| Hsbp1 | 2.507151 | 0 | 0 | 0 |
| Epb41 | 0.508849 | 0 | 0 | 0 |
| Gm9888 | 0.522808 | 0 | 0 | 0 |
| Eed | 0.989692 | 0 | 0 | 0 |
| Herc2 | 1.1027 | 0 | 0 | 0 |
| Retreg1 | 0.92027 | 0 | 0 | 0 |
| Vrk2 | 1.429559 | 0 | 0 | 0 |
| Thumpd3 | 0.849558 | 0 | 0 | 0 |
| Wsb1 | 0.845494 | 0 | 0 | 0 |
| Mfap3 | 0.60307 | 0 | 0 | 0 |
| Gltp | 2.478 | 0 | 0 | 0 |
| Morc3 | 0.800101 | 0 | 0 | 0 |
| Tomm70a | 0.410605 | 0 | 0 | 0 |
| 7-Mar | 0.660789 | 0 | 0 | 0 |
| Ddx50 | 0.960277 | 0 | 0 | 0 |
| Gcnt1 | 1.004973 | 0 | 0 | 0 |
| Cd33 | 0.596561 | 0 | 0 | 0 |
| Pdzd8 | 0.464287 | 0 | 0 | 0 |
| Baz1a | 0.941956 | 0 | 0 | 0 |
| Sltm | 1.071214 | 0 | 0 | 0 |
| Pcyox1 | 0.257532 | 0 | 0 | 0 |
| Btg2 | 0.362263 | 0 | 0 | 0 |
| Tmx4 | 0.450345 | 0 | 0 | 0 |
| Ap2a2 | 0.553479 | 0 | 0 | 0 |
| Leng8 | 1.091446 | 0 | 0 | 0 |
| Rp2 | 1.112616 | 0 | 0 | 0 |
| Tk2 | 0.432692 | 0 | 0 | 0 |
| Map7d1 | 0.255907 | 0 | 0 | 0 |
| Memo1 | 0.552378 | 0 | 0 | 0 |
| Ss18 | 0.549 | 0 | 0 | 0 |
| Noc2l | 0.468 | 0 | 0 | 0 |
| Arl8b | 0.302599 | 0 | 0 | 0 |
| Nfkb1 | 0.715437 | 0 | 0 | 0 |
| Dpagt1 | 1.12331 | 0 | 0 | 0 |
| Ap1s2 | 0.984825 | 0 | 0 | 0 |
| Asxl2 | 0.379028 | 0 | 0 | 0 |
| Nbeal1 | 0.442473 | 0 | 0 | 0 |
| Itpkb | 0.737181 | 0 | 0 | 0 |
| Ccni | 0.297808 | 0 | 0 | 0 |
| Cr1l | 0.872412 | 0 | 0 | 0 |
| Vps54 | 0.943699 | 0 | 0 | 0 |
| Rasa4 | 0.787381 | 0 | 0 | 0 |
| Stambpl1 | 0.726511 | 0 | 0 | 0 |

| | | | | |
|---|---|---|---|---|
| Usp21 | 1.065672 | 0 | 0 | 0 |
| Gtf2a2 | 0.713481 | 0 | 0 | 0 |
| Rasgrp3 | 0.349789 | 0 | 0 | 0 |
| Lpgat1 | 0.417058 | 0 | 0 | 0 |
| Gm5148 | 0.588971 | 0 | 0 | 0 |
| Magt1 | 0.384128 | 0 | 0 | 0 |
| Smc6 | 0.420777 | 0 | 0 | 0 |
| Pcf11 | 1.022333 | 0 | 0 | 0 |
| Birc3 | 1.013705 | 0 | 0 | 0 |
| Orc3 | 1.233909 | 0 | 0 | 0 |
| Sqor | 1.161367 | 0 | 0 | 0 |
| Papola | 0.840668 | 0 | 0 | 0 |
| Ctbs | 0.54747 | 0 | 0 | 0 |
| Epb41l3 | 0.904735 | 0 | 0 | 0 |
| Mxd4 | 0.839775 | 0 | 0 | 0 |
| Tulp4 | 0.668792 | 0 | 0 | 0 |
| Ube3a | 1.095739 | 0 | 0 | 0 |
| Faf2 | 0.698808 | 0 | 0 | 0 |
| Brd2 | 1.35453 | 0 | 0 | 0 |
| Mgrn1 | 0.838228 | 0 | 0 | 0 |
| Gtf2a1 | 0.420486 | 0 | 0 | 0 |
| Frg1 | 1.223491 | 0 | 0 | 0 |
| Zfp398 | 0.616515 | 0 | 0 | 0 |
| Ccnl2 | 0.544528 | 0 | 0 | 0 |
| Med13l | 1.417919 | 0 | 0 | 0 |
| Sh3kbp1 | 1.169347 | 0 | 0 | 0 |
| Mptx1 | 0 | 0.731705 | 0 | 0 |
| Anxa2 | 0 | 0.355032 | 0 | 0 |
| Golm1 | 0 | 1.305725 | 0 | 0 |
| Qk | 0 | 1.217155 | 0 | 0 |
| mt-Rnr2 | 0 | 0.827046 | 0 | 0 |
| Ywhaz | 0 | 1.361361 | 0 | 0 |
| Atf3 | 0 | 1.152035 | 0 | 0 |
| Gm14760 | 0 | 0.440509 | 0 | 0 |
| Ubc | 0 | 0.676936 | 0 | 0 |
| Gm10175 | 0 | 0.95841 | 0 | 0 |
| Gm5869 | 0 | 0.395965 | 0 | 0 |
| Gm12892 | 0 | 0.437949 | 0 | 0 |
| Gm5586 | 0 | 0.47989 | 0 | 0 |
| Calr | 0 | 0.89834 | 0 | 0 |
| Cdk2ap1 | 0 | 0.33559 | 0 | 0 |
| Gm7935 | 0 | 0.66577 | 0 | 0 |
| Gm14494 | 0 | 0.743506 | 0 | 0 |
| Hpgds | 0 | 0.959118 | 0 | 0 |
| Mylip | 0 | 1.737158 | 0 | 0 |
| Gm16437 | 0 | 0.557959 | 0 | 0 |
| Tesk1 | 0 | 0.269886 | 0 | 0 |
| Col6a3 | 0 | 4.217716 | 0 | 0 |
| Pdia3 | 0 | 0.599593 | 0 | 0 |

| | | | | |
|---|---|---|---|---|
| Bmp2k | 0 | 1.343289 | 0 | 0 |
| Ifitm10 | 0 | 2.852004 | 0 | 0 |
| Gm10257 | 0 | 0.580376 | 0 | 0 |
| H3f3a-ps2 | 0 | 0.548011 | 0 | 0 |
| Rpl19-ps1 | 0 | 0.422939 | 0 | 0 |
| Ctsc | 0 | 0.87151 | 0 | 0 |
| Tpm3-rs7 | 0 | 1.131398 | 0 | 0 |
| Gm7308 | 0 | 0.386312 | 0 | 0 |
| Lrrc3 | 0 | 1.458878 | 0 | 0 |
| Gm12350 | 0 | 1.420089 | 0 | 0 |
| Lat2 | 0 | 2.16392 | 0 | 0 |
| Gm4735 | 0 | 0.703694 | 0 | 0 |
| Maf | 0 | 1.235676 | 0 | 0 |
| Rpl7-ps7 | 0 | 0.334044 | 0 | 0 |
| Pla2g15 | 0 | 1.168274 | 0 | 0 |
| Gm13453 | 0 | 0.513919 | 0 | 0 |
| Gm3724 | 0 | 1.347672 | 0 | 0 |
| Nrros | 0 | 1.501948 | 0 | 0 |
| Cbx3 | 0 | 1.823767 | 0 | 0 |
| Zfp385a | 0 | 2.312739 | 0 | 0 |
| Rps11-ps1 | 0 | 1.103751 | 0 | 0 |
| Gm9347 | 0 | 0.482734 | 0 | 0 |
| Gm12844 | 0 | 0.618148 | 0 | 0 |
| Snx5 | 0 | 1.108441 | 0 | 0 |
| Tspan7 | 0 | 1.923928 | 0 | 0 |
| Cd300a | 0 | 1.546928 | 0 | 0 |
| Gm5529 | 0 | 0.302249 | 0 | 0 |
| Inpp5d | 0 | 1.130797 | 0 | 0 |
| Fuca1 | 0 | 1.787906 | 0 | 0 |
| Gm7901 | 0 | 0.996329 | 0 | 0 |
| Gm10800 | 0 | 1.05445 | 0 | 0 |
| Rac1 | 0 | 1.327634 | 0 | 0 |
| Zfp207 | 0 | 1.472291 | 0 | 0 |
| Bnip3l-ps | 0 | 0.703772 | 0 | 0 |
| Zfp652 | 0 | 1.208055 | 0 | 0 |
| Phyhd1 | 0 | 1.669115 | 0 | 0 |
| Ier2 | 0 | 0.362043 | 0 | 0 |
| Tpst2 | 0 | 1.437645 | 0 | 0 |
| Apbb1ip | 0 | 2.348592 | 0 | 0 |
| Map2k4 | 0 | 0.263617 | 0 | 0 |
| Cebpb | 0 | 0.649282 | 0 | 0 |
| Pip4p1 | 0 | 1.771739 | 0 | 0 |
| Ucp2 | 0 | 0.388653 | 0 | 0 |
| Gm38158 | 0 | 2.253457 | 0 | 0 |
| Iscu | 0 | 1.383127 | 0 | 0 |
| Rspry1 | 0 | 0 | 0.801591 | 0 |
| Tug1 | 0 | 0 | 0.945149 | 0 |
| Cep63 | 0 | 0 | 0.928367 | 0 |
| Hnrnpdl | 0 | 0 | 0.30185 | 0 |

| | | | | |
|---|---|---|---|---|
| Fam76b | 0 | 0 | 0.374305 | 0 |
| Thoc2 | 0 | 0 | 0.476066 | 0 |
| Hectd4 | 0 | 0 | 1.343402 | 0 |
| Supt16 | 0 | 0 | 0.875728 | 0 |
| Fads1 | 0 | 0 | 0.443792 | 0 |
| Arf6 | 0 | 0 | 0.688198 | 0 |
| Mfng | 0 | 0 | 2.062185 | 0 |
| Gm7072 | 0 | 0 | 0.283314 | 0 |
| Gm45606 | 0 | 0 | 1.033291 | 0 |
| Slc35e1 | 0 | 0 | 0.606002 | 0 |
| Mef2d | 0 | 0 | 0.794101 | 0 |
| Usp19 | 0 | 0 | 1.179918 | 0 |
| Zfp277 | 0 | 0 | 0.436249 | 0 |
| Frmd4b | 0 | 0 | 0.814161 | 0 |
| Nap1l1 | 0 | 0 | 0.741016 | 0 |
| Fam168b | 0 | 0 | 0.787182 | 0 |
| Zfp658 | 0 | 0 | 0.254555 | 0 |
| Pou2f2 | 0 | 0 | 2.294184 | 0 |
| Smox | 0 | 0 | 0.898437 | 0 |
| Arnt | 0 | 0 | 0.348614 | 0 |
| Mcur1 | 0 | 0 | 0.416942 | 0 |
| Tmem63a | 0 | 0 | 0.394304 | 0 |
| H2afz | 0 | 0 | 1.42525 | 0 |
| Paip1 | 0 | 0 | 0.647268 | 0 |
| Mitf | 0 | 0 | 0.798367 | 0 |
| Atg14 | 0 | 0 | 0.639164 | 0 |
| Hmbox1 | 0 | 0 | 1.290237 | 0 |
| Vps18 | 0 | 0 | 0.418551 | 0 |
| Zswim6 | 0 | 0 | 0.38804 | 0 |
| Plcb2 | 0 | 0 | 0.279323 | 0 |
| Pwwp2a | 0 | 0 | 0.553804 | 0 |
| Dmxl1 | 0 | 0 | 0.710859 | 0 |
| Abhd17c | 0 | 0 | 0.568478 | 0 |
| Gm44949 | 0 | 0 | 0.366243 | 0 |
| Oraov1 | 0 | 0 | 0.930335 | 0 |
| Ip6k1 | 0 | 0 | 0.398366 | 0 |
| Gm46224 | 0 | 0 | 0.432704 | 0 |
| Dnajb4 | 0 | 0 | 0.393429 | 0 |
| Psenen | 0 | 0 | 1.76027 | 0 |
| Rab11fip5 | 0 | 0 | 0.451217 | 0 |
| Rad21 | 0 | 0 | 0.398323 | 0 |
| Ppp2r5c | 0 | 0 | 0.771581 | 0 |
| Appbp2 | 0 | 0 | 0.491098 | 0 |
| Dstyk | 0 | 0 | 0.411572 | 0 |
| Ddx23 | 0 | 0 | 0.963606 | 0 |
| Il7r | 0 | 0 | 0.527688 | 0 |
| Smad4 | 0 | 0 | 0.471377 | 0 |
| Lrrc41 | 0 | 0 | 0.343321 | 0 |
| Fbxw2 | 0 | 0 | 0.856281 | 0 |

| | | | | |
|---|---|---|---|---|
| Atp6v1f | 0 | 0 | 1.528241 | 0 |
| Tspan4 | 0 | 0 | 1.74345 | 0 |
| Ighm | 0 | 0 | 1.371987 | 0 |
| Sh2b1 | 0 | 0 | 0.454911 | 0 |
| Nfkbia | 0 | 0 | 0.745819 | 0 |
| Phc3 | 0 | 0 | 0.573192 | 0 |
| Jup | 0 | 0 | 0.657501 | 0 |
| Ncor1 | 0 | 0 | 0.728134 | 0 |
| Smarcd2 | 0 | 0 | 0.505147 | 0 |
| Sfswap | 0 | 0 | 0.842694 | 0 |
| Mfhas1 | 0 | 0 | 0.665786 | 0 |
| Ipo9 | 0 | 0 | 0.424135 | 0 |
| Fndc3b | 0 | 0 | 0.468452 | 0 |
| Rnpepl1 | 0 | 0 | 0.717697 | 0 |
| Cxxc5 | 0 | 0 | 1.66487 | 0 |
| Parp1 | 0 | 0 | 1.076285 | 0 |
| Klhl5 | 0 | 0 | 0.421004 | 0 |
| Nrp1 | 0 | 0 | 0.379229 | 0 |
| Psip1 | 0 | 0 | 0.733339 | 0 |
| Maml2 | 0 | 0 | 0.697419 | 0 |
| Ehmt1 | 0 | 0 | 0.952105 | 0 |
| Sec11c | 0 | 0 | 1.437473 | 0 |
| Irf5 | 0 | 0 | 1.389571 | 0 |
| Agpat5 | 0 | 0 | 0.631074 | 0 |
| Cd180 | 0 | 0 | 0.591843 | 0 |
| Card6 | 0 | 0 | 0.889196 | 0 |
| Vgll4 | 0 | 0 | 0.851777 | 0 |
| Baz2b | 0 | 0 | 0.289371 | 0 |
| Ehbp1l1 | 0 | 0 | 0.668471 | 0 |
| Samd8 | 0 | 0 | 0.354572 | 0 |
| Rai1 | 0 | 0 | 0.847065 | 0 |
| Ptprm | 0 | 0 | 0.385601 | 0 |
| Pmm2 | 0 | 0 | 1.047613 | 0 |
| Wac | 0 | 0 | 0.536389 | 0 |
| Dirc2 | 0 | 0 | 0.738601 | 0 |
| Gm47283 | 0 | 0 | 0.865373 | 0 |
| Nsmaf | 0 | 0 | 0.585255 | 0 |
| Prdx5 | 0 | 0 | 2.210099 | 0 |
| Ice1 | 0 | 0 | 0.456414 | 0 |
| Kpna1 | 0 | 0 | 0.566191 | 0 |
| Dpp9 | 0 | 0 | 0.461358 | 0 |
| Tmem106b | 0 | 0 | 0.472312 | 0 |
| Slc39a10 | 0 | 0 | 0.269714 | 0 |
| Tbc1d9b | 0 | 0 | 0.279406 | 0 |
| Peli2 | 0 | 0 | 0.608232 | 0 |
| Abhd15 | 0 | 0 | 0.521147 | 0 |
| Stmn1 | 0 | 0 | 1.093019 | 0 |
| Rbm25 | 0 | 0 | 0.705067 | 0 |
| Smad3 | 0 | 0 | 0.669957 | 0 |

| | | | | |
|---|---|---|---|---|
| Abcd1 | 0 | 0 | 0.595798 | 0 |
| Cdk8 | 0 | 0 | 0.75292 | 0 |
| Akap10 | 0 | 0 | 0.884025 | 0 |
| Rnf10 | 0 | 0 | 0.530678 | 0 |
| Jarid2 | 0 | 0 | 0.748528 | 0 |
| Ank2 | 0 | 0 | 0.84611 | 0 |
| Ube4a | 0 | 0 | 0.5028 | 0 |
| Mxi1 | 0 | 0 | 0.2503 | 0 |
| Ago3 | 0 | 0 | 0.298709 | 0 |
| Mrpl37 | 0 | 0 | 0.326464 | 0 |
| Slc3a2 | 0 | 0 | 1.149051 | 0 |
| Nr6a1os | 0 | 0 | 0.26055 | 0 |
| Sec24c | 0 | 0 | 0.585737 | 0 |
| Map3k4 | 0 | 0 | 0.952259 | 0 |
| Vps36 | 0 | 0 | 0.437676 | 0 |
| Hp1bp3 | 0 | 0 | 0.576448 | 0 |
| Csnk1g2 | 0 | 0 | 0.57232 | 0 |
| Bbx | 0 | 0 | 0.540019 | 0 |
| Gm8995 | 0 | 0 | 0.416569 | 0 |
| Klk8 | 0 | 0 | 1.856572 | 0 |
| Dennd1a | 0 | 0 | 0.601359 | 0 |
| Ppp2r2a | 0 | 0 | 0.683924 | 0 |
| Mapkap1 | 0 | 0 | 1.067734 | 0 |
| Plekho1 | 0 | 0 | 2.162372 | 0 |
| Nrf1 | 0 | 0 | 0.335558 | 0 |
| Coro1b | 0 | 0 | 1.594752 | 0 |
| Zcchc8 | 0 | 0 | 0.683739 | 0 |
| Rap2a | 0 | 0 | 0.638855 | 0 |
| Eprs | 0 | 0 | 0.668971 | 0 |
| Helz | 0 | 0 | 0.292354 | 0 |
| Gm37760 | 0 | 0 | 0.412641 | 0 |
| Hcfc1 | 0 | 0 | 0.707825 | 0 |
| Cables1 | 0 | 0 | 0.325376 | 0 |
| Gabbr1 | 0 | 0 | 1.103471 | 0 |
| Fndc3a | 0 | 0 | 0.406853 | 0 |
| Secisbp2l | 0 | 0 | 0.319857 | 0 |
| Ckap5 | 0 | 0 | 0.68376 | 0 |
| Etv6 | 0 | 0 | 0.50245 | 0 |
| Soga1 | 0 | 0 | 0.619233 | 0 |
| Nr2c2 | 0 | 0 | 0.568429 | 0 |
| E230001N | 0 | 0 | 0.897702 | 0 |
| Pkd2 | 0 | 0 | 0.621993 | 0 |
| Adgrg1 | 0 | 0 | 1.089933 | 0 |
| Tut4 | 0 | 0 | 0.695049 | 0 |
| Mtmr4 | 0 | 0 | 0.425121 | 0 |
| Chfr | 0 | 0 | 0.471122 | 0 |
| Zfp451 | 0 | 0 | 0.287299 | 0 |
| Khdrbs1 | 0 | 0 | 0.385982 | 0 |
| Dicer1 | 0 | 0 | 0.358193 | 0 |

| | | | | |
|---|---|---|---|---|
| Pcnt | 0 | 0 | 0.434139 | 0 |
| 2510039O[illegible] | 0 | 0 | 0.464738 | 0 |
| Ppp1cb | 0 | 0 | 0.384488 | 0 |
| Pdpk1 | 0 | 0 | 0.559271 | 0 |
| Gtpbp2 | 0 | 0 | 0.915736 | 0 |
| Nfx1 | 0 | 0 | 0.542632 | 0 |
| Rsrc1 | 0 | 0 | 0.252485 | 0 |
| Gxylt1 | 0 | 0 | 0.251733 | 0 |
| Zranb1 | 0 | 0 | 0.479687 | 0 |
| Pptc7 | 0 | 0 | 0.790188 | 0 |
| Rhog | 0 | 0 | 1.269703 | 0 |
| Adam17 | 0 | 0 | 0.580761 | 0 |
| Adipor2 | 0 | 0 | 0.314777 | 0 |
| Scd2 | 0 | 0 | 0.85286 | 0 |
| Dapp1 | 0 | 0 | 0.272656 | 0 |
| Sp1 | 0 | 0 | 0.478172 | 0 |
| Plcg1 | 0 | 0 | 0.329162 | 0 |
| Dhx34 | 0 | 0 | 0.317508 | 0 |
| Ccdc186 | 0 | 0 | 0.422659 | 0 |
| Trim44 | 0 | 0 | 0.509504 | 0 |
| Tgfa | 0 | 0 | 0.647094 | 0 |
| Stat3 | 0 | 0 | 0.321829 | 0 |
| Nfat5 | 0 | 0 | 0.268473 | 0 |
| Gm15928 | 0 | 0 | 0.492149 | 0 |
| Atp2c1 | 0 | 0 | 0.34084 | 0 |
| Ppfia1 | 0 | 0 | 0.282245 | 0 |
| Secisbp2 | 0 | 0 | 0.477324 | 0 |
| R3hdm1 | 0 | 0 | 0.299243 | 0 |
| Twf2 | 0 | 0 | 1.697309 | 0 |
| Pmepa1 | 0 | 0 | 1.935033 | 0 |
| Asxl1 | 0 | 0 | 0.345645 | 0 |
| Mark3 | 0 | 0 | 0.283606 | 0 |
| Rnf19b | 0 | 0 | 0.503961 | 0 |
| Nf1 | 0 | 0 | 0.357184 | 0 |
| Gm40309 | 0 | 0 | 0.499293 | 0 |
| Huwe1 | 0 | 0 | 0.365071 | 0 |
| Prrc2b | 0 | 0 | 0.612542 | 0 |
| Gab1 | 0 | 0 | 0.308685 | 0 |
| Mia2 | 0 | 0 | 0.388719 | 0 |
| Trpm2 | 0 | 0 | 0.407773 | 0 |
| Ap1b1 | 0 | 0 | 0.830168 | 0 |
| Lgals9 | 0 | 0 | 1.823786 | 0 |
| Ccnd3 | 0 | 0 | 0.418562 | 0 |
| Sacm1l | 0 | 0 | 0 | 0.355804 |
| Uba52 | 0 | 0 | 0 | 1.962358 |
| Rps26-ps1 | 0 | 0 | 0 | 2.071058 |
| Rpl27 | 0 | 0 | 0 | 1.969967 |
| Rps29 | 0 | 0 | 0 | 1.038154 |
| Gm3699 | 0 | 0 | 0 | 1.337253 |

| | | | | |
|---|---|---|---|---|
| Gpr84 | 0 | 0 | 0 | 2.780356 |
| Gm10689 | 0 | 0 | 0 | 0.512551 |
| Rpl38 | 0 | 0 | 0 | 1.445784 |
| Rpl35a | 0 | 0 | 0 | 1.48328 |
| Gas5 | 0 | 0 | 0 | 2.928467 |
| Myo7a | 0 | 0 | 0 | 0.390885 |
| Gng5 | 0 | 0 | 0 | 1.470138 |
| Mrfap1 | 0 | 0 | 0 | 1.300017 |
| Rpl22 | 0 | 0 | 0 | 2.319478 |
| Gm3150 | 0 | 0 | 0 | 0.639361 |
| Slamf9 | 0 | 0 | 0 | 2.618743 |
| Sec63 | 0 | 0 | 0 | 0.29293 |
| Tnfsf13b | 0 | 0 | 0 | 3.35651 |
| Slc39a1 | 0 | 0 | 0 | 2.147467 |
| Zfand3 | 0 | 0 | 0 | 1.956803 |
| Gm17066 | 0 | 0 | 0 | 1.88181 |
| Bach2 | 0 | 0 | 0 | 0.459016 |
| Vamp8 | 0 | 0 | 0 | 1.96774 |
| Srsf9 | 0 | 0 | 0 | 2.112676 |
| Ap1g1 | 0 | 0 | 0 | 0.604532 |
| Gm32036 | 0 | 0 | 0 | 2.586234 |
| Rin3 | 0 | 0 | 0 | 0.674449 |
| Rcc2 | 0 | 0 | 0 | 0.456361 |
| Stard9 | 0 | 0 | 0 | 0.370241 |
| Ppp3cb | 0 | 0 | 0 | 0.585085 |
| Tmed10 | 0 | 0 | 0 | 1.515308 |
| Tnrc6b | 0 | 0 | 0 | 0.79457 |
| Hnrnpa3 | 0 | 0 | 0 | 0.352732 |
| Rpl36a | 0 | 0 | 0 | 1.961616 |
| Pag1 | 0 | 0 | 0 | 0.285132 |
| Myl12a | 0 | 0 | 0 | 2.191588 |
| Gm10847 | 0 | 0 | 0 | 3.31064 |
| Rpl12 | 0 | 0 | 0 | 2.077915 |
| Zfp579 | 0 | 0 | 0 | 1.6045 |
| Cdk19 | 0 | 0 | 0 | 0.519532 |
| Gm34589 | 0 | 0 | 0 | 0.629836 |
| Rnf130 | 0 | 0 | 0 | 1.143063 |
| Cebpa | 0 | 0 | 0 | 0.303716 |
| Ilkap | 0 | 0 | 0 | 1.010086 |
| St13 | 0 | 0 | 0 | 1.788693 |
| Dusp3 | 0 | 0 | 0 | 3.149353 |
| Ptgr2 | 0 | 0 | 0 | 0.551016 |
| Mir692-2 | 0 | 0 | 0 | 0.562538 |
| Clk2 | 0 | 0 | 0 | 0.365015 |
| Rpl39 | 0 | 0 | 0 | 1.362065 |
| Vapa | 0 | 0 | 0 | 1.677435 |
| Pomp | 0 | 0 | 0 | 1.91633 |
| Ppp2r2d | 0 | 0 | 0 | 0.766632 |
| Gm15500 | 0 | 0 | 0 | 1.757052 |

| | | | | |
|---|---|---|---|---|
| Snx29 | 0 | 0 | 0 | 0.381474 |
| Qdpr | 0 | 0 | 0 | 1.028471 |
| Agps | 0 | 0 | 0 | 0.324807 |
| Lamtor4 | 0 | 0 | 0 | 2.613203 |
| Sfi1 | 0 | 0 | 0 | 0.466261 |
| Manf | 0 | 0 | 0 | 1.776501 |
| Phf12 | 0 | 0 | 0 | 0.542105 |
| Gm42659 | 0 | 0 | 0 | 2.571735 |
| Smad7 | 0 | 0 | 0 | 1.451411 |
| Maml3 | 0 | 0 | 0 | 0.654737 |
| Rabl6 | 0 | 0 | 0 | 0.918368 |
| Phka2 | 0 | 0 | 0 | 0.473575 |
| Gm15590 | 0 | 0 | 0 | 0.736304 |
| Ube2k | 0 | 0 | 0 | 1.215714 |
| Pcbp1 | 0 | 0 | 0 | 1.220324 |
| Abi1 | 0 | 0 | 0 | 0.267658 |
| Spop | 0 | 0 | 0 | 0.505843 |
| Agpat3 | 0 | 0 | 0 | 0.647964 |
| Arfgap1 | 0 | 0 | 0 | 0.30834 |
| Trip11 | 0 | 0 | 0 | 0.492395 |
| Tbc1d16 | 0 | 0 | 0 | 0.355235 |
| Disc1 | 0 | 0 | 0 | 1.112382 |
| Ehmt2 | 0 | 0 | 0 | 0.863243 |
| Prpsap2 | 0 | 0 | 0 | 0.561761 |
| Rpl5 | 0 | 0 | 0 | 2.030021 |
| Accs | 0 | 0 | 0 | 0.540902 |
| Tex261 | 0 | 0 | 0 | 2.661768 |
| mt-Nd3 | 0 | 0 | 0 | 1.620382 |
| Ppm1h | 0 | 0 | 0 | 1.682662 |
| Lrrc8a | 0 | 0 | 0 | 2.929755 |
| Wdr33 | 0 | 0 | 0 | 0.424331 |
| Naca | 0 | 0 | 0 | 1.839651 |
| Hectd1 | 0 | 0 | 0 | 0.261224 |
| Smndc1 | 0 | 0 | 0 | 0.295221 |
| Cd83 | 0 | 0 | 0 | 2.39704 |
| Bcap31 | 0 | 0 | 0 | 1.488445 |
| Braf | 0 | 0 | 0 | 0.466158 |
| Sel1l | 0 | 0 | 0 | 0.469428 |
| Glmp | 0 | 0 | 0 | 1.523034 |
| Atraid | 0 | 0 | 0 | 1.634412 |
| Syt11 | 0 | 0 | 0 | 1.414397 |
| Ube2j1 | 0 | 0 | 0 | 0.320624 |
| Gpx4 | 0 | 0 | 0 | 2.136589 |
| Selenos | 0 | 0 | 0 | 2.14288 |
| Nrd1 | 0 | 0 | 0 | 0.779383 |
| Prr14l | 0 | 0 | 0 | 0.643134 |
| Akap9 | 0 | 0 | 0 | 0.720281 |
| Znfx1 | 0 | 0 | 0 | 0.510127 |
| Atp5g2 | 0 | 0 | 0 | 2.429465 |

| | | | | |
|---|---|---|---|---|
| Ypel3 | 0 | 0 | 0 | 1.93648 |
| Supt6 | 0 | 0 | 0 | 0.978383 |
| Kdelr2 | 0 | 0 | 0 | 1.305991 |
| Trmt112-ps | 0 | 0 | 0 | 0.861334 |
| Gngt2 | 0 | 0 | 0 | 1.970965 |
| Ergic1 | 0 | 0 | 0 | 0.431744 |
| Ccdc88b | 0 | 0 | 0 | 0.746398 |
| Ccdc47 | 0 | 0 | 0 | 1.006134 |
| Chmp3 | 0 | 0 | 0 | 0.9516 |
| Phf3 | 0 | 0 | 0 | 0.426458 |
| Srgn | 0 | 0 | 0 | 1.944358 |
| Herpud1 | 0 | 0 | 0 | 1.773997 |
| Rela | 0 | 0 | 0 | 0.515604 |
| Eps15l1 | 0 | 0 | 0 | 0.457212 |
| Mia3 | 0 | 0 | 0 | 0.586678 |
| Prune2 | 0 | 0 | 0 | 2.568504 |
| Exoc4 | 0 | 0 | 0 | 0.342655 |
| Ptpa | 0 | 0 | 0 | 1.172449 |
| Abr | 0 | 0 | 0 | 0.700016 |
| Brk1 | 0 | 0 | 0 | 1.491111 |
| Usf2 | 0 | 0 | 0 | 1.373793 |
| Apoe | 0 | 0 | 0 | 4.010424 |
| Rpl17-ps1( | 0 | 0 | 0 | 0.944256 |
| Pik3ca | 0 | 0 | 0 | 0.452679 |
| Psme1 | 0 | 0 | 0 | 2.403271 |
| Selenow | 0 | 0 | 0 | 2.896283 |
| Btbd9 | 0 | 0 | 0 | 0.486997 |
| Mlec | 0 | 0 | 0 | 0.930054 |
| Gm42602 | 0 | 0 | 0 | 0.312492 |
| Ldha | 0 | 0 | 0 | 1.676058 |
| Pld3 | 0 | 0 | 0 | 1.651439 |
| St6gal1 | 0 | 0 | 0 | 0.28293 |
| Ccpg1 | 0 | 0 | 0 | 0.730658 |
| Rsbn1 | 0 | 0 | 0 | 0.347658 |
| Gm12338 | 0 | 0 | 0 | 1.182326 |
| Snrk | 0 | 0 | 0 | 0.727702 |
| Smarcad1 | 0 | 0 | 0 | 0.68174 |
| Cask | 0 | 0 | 0 | 0.575814 |
| Csad | 0 | 0 | 0 | 0.746234 |
| Mras | 0 | 0 | 0 | 1.112336 |
| Phactr2 | 0 | 0 | 0 | 0.356706 |
| Sike1 | 0 | 0 | 0 | 0.266541 |
| Brf1 | 0 | 0 | 0 | 0.766517 |
| Sertad2 | 0 | 0 | 0 | 0.824195 |
| Eif3c | 0 | 0 | 0 | 0.66488 |
| Gm10095 | 0 | 0 | 0 | 1.344625 |
| Lrrfip1 | 0 | 0 | 0 | 0.807911 |
| Pgrmc2 | 0 | 0 | 0 | 0.937994 |
| Ptges3 | 0 | 0 | 0 | 1.783011 |

| | | | | |
|---|---|---|---|---|
| Rnf20 | 0 | 0 | 0 | 0.471548 |
| Kdm5a | 0 | 0 | 0 | 0.345543 |
| Set | 0 | 0 | 0 | 2.200815 |
| Pkm | 0 | 0 | 0 | 1.623272 |
| Gm8451 | 0 | 0 | 0 | 0.811648 |
| Suds3 | 0 | 0 | 0 | 0.867497 |
| Gnl2 | 0 | 0 | 0 | 0.425403 |
| Pcmtd1 | 0 | 0 | 0 | 0.314481 |
| Tifab | 0 | 0 | 0 | 0.654655 |
| Gm10134 | 0 | 0 | 0 | 2.5534 |
| Gm8186 | 0 | 0 | 0 | 1.382506 |
| Gm4332 | 0 | 0 | 0 | 0.445132 |
| Itpr2 | 0 | 0 | 0 | 0.581448 |
| Gm5526 | 0 | 0 | 0 | 1.532742 |
| Gng5-ps | 0 | 0 | 0 | 1.618181 |
| Rpl39-ps | 0 | 0 | 0 | 0.777533 |
| Uba5 | 0 | 0 | 0 | 1.335608 |
| Aplp2 | 0 | 0 | 0 | 0.40259 |

Supplementry Table 8.  PB-Old DEGs in Cerebellum

| Gene | p_val | avg_log2FC | pct.1 | pct.2 | p_val_adj |
|---|---|---|---|---|---|
| Lars2 | 2E-106 | -2.54006 | 1 | 1 | 4.7E-102 |
| Gm23935 | 1.7E-105 | -2.3308 | 0.996 | 1 | 3.9E-101 |
| Gm24187 | 3.5E-105 | -2.37828 | 0.988 | 1 | 8.2E-101 |
| Tmsb4x | 1.2E-99 | 1.738904 | 1 | 1 | 2.78E-95 |
| Gm24270 | 4.43E-88 | -1.826 | 0.965 | 0.996 | 1.05E-83 |
| C1qc | 1.17E-87 | 1.278777 | 1 | 1 | 2.76E-83 |
| Rps29 | 7.04E-74 | 1.7668 | 0.988 | 0.996 | 1.66E-69 |
| C1qb | 1.33E-73 | 1.039568 | 1 | 1 | 3.13E-69 |
| Cst3 | 4.6E-69 | 1.013218 | 1 | 1 | 1.08E-64 |
| Rn7sk | 2.8E-66 | -1.83279 | 0.621 | 0.98 | 6.62E-62 |
| Cd81 | 1.32E-60 | 1.227584 | 1 | 1 | 3.12E-56 |
| Gm15564 | 2.39E-60 | -1.34561 | 0.789 | 0.98 | 5.63E-56 |
| C1qa | 9.46E-59 | 0.883386 | 1 | 1 | 2.23E-54 |
| Eef1a1 | 8.36E-54 | 1.244237 | 1 | 0.993 | 1.97E-49 |
| Rps27 | 7.96E-53 | 1.621248 | 0.969 | 0.944 | 1.88E-48 |
| Itm2b | 1.47E-52 | 0.924103 | 1 | 1 | 3.46E-48 |
| Ubb | 6.01E-52 | 1.790176 | 0.945 | 0.916 | 1.42E-47 |
| Gm14303 | 4.57E-50 | 1.349315 | 0.965 | 0.842 | 1.08E-45 |
| mt-Co3 | 3.13E-48 | 1.631635 | 0.922 | 0.814 | 7.39E-44 |
| Rps11 | 5.26E-47 | 1.707563 | 0.934 | 0.903 | 1.24E-42 |
| Gm24245 | 1.1E-46 | -1.23358 | 0.367 | 0.835 | 2.61E-42 |
| Adap2os | 2.63E-46 | 1.855779 | 0.953 | 0.894 | 6.21E-42 |
| Csf1r | 3.9E-46 | -0.91174 | 1 | 1 | 9.2E-42 |
| Selplg | 6.12E-46 | 0.911343 | 1 | 1 | 1.44E-41 |
| Ppia | 7.72E-45 | 1.403557 | 0.961 | 0.942 | 1.82E-40 |
| Rpl37a | 1.09E-42 | 1.633081 | 0.945 | 0.926 | 2.57E-38 |
| Rps28 | 3.31E-42 | 1.634578 | 0.965 | 0.937 | 7.8E-38 |
| mt-Co2 | 1.5E-41 | 1.629688 | 0.898 | 0.758 | 3.53E-37 |
| Rplp1 | 3.87E-40 | 1.382569 | 0.957 | 0.961 | 9.14E-36 |
| Rpl37 | 5.89E-40 | 1.543539 | 0.938 | 0.896 | 1.39E-35 |
| Ctsh | 8.76E-40 | 1.360967 | 0.965 | 0.957 | 2.07E-35 |
| Ctss | 1.15E-39 | 0.792712 | 1 | 1 | 2.72E-35 |
| Rps9 | 2.31E-39 | 1.369782 | 0.965 | 0.959 | 5.44E-35 |
| Cd9 | 2.67E-39 | 1.235419 | 0.973 | 0.933 | 6.29E-35 |
| mt-Co1 | 3.01E-39 | 1.197052 | 0.992 | 0.985 | 7.09E-35 |
| Fcer1g | 9.75E-39 | 1.065283 | 0.98 | 0.983 | 2.3E-34 |
| Rpl13 | 1.31E-38 | 1.508989 | 0.953 | 0.937 | 3.1E-34 |
| Fth1 | 1.44E-38 | 1.094234 | 0.996 | 0.993 | 3.41E-34 |
| Gm25911 | 5.43E-38 | -1.05877 | 0.273 | 0.755 | 1.28E-33 |
| Pfn1 | 6.15E-38 | 1.465489 | 0.941 | 0.926 | 1.45E-33 |
| Ftl1 | 3.26E-37 | 1.108003 | 0.992 | 0.996 | 7.69E-33 |
| Rps27rt | 5.19E-37 | 1.396379 | 0.902 | 0.682 | 1.23E-32 |
| Sparc | 1.24E-36 | 0.928659 | 1 | 1 | 2.91E-32 |
| Malat1 | 1.5E-36 | 0.776146 | 1 | 0.994 | 3.54E-32 |

| | | | | | |
|---|---|---|---|---|---|
| mt-Atp6 | 2.12E-36 | 1.432646 | 0.914 | 0.755 | 5E-32 |
| Gm15536 | 3.41E-36 | 1.116462 | 0.676 | 0.258 | 8.05E-32 |
| Rps14 | 1.6E-34 | 1.492302 | 0.922 | 0.933 | 3.77E-30 |
| Rpl18a | 2.15E-34 | 1.148728 | 0.996 | 0.994 | 5.07E-30 |
| mt-Cytb | 3.12E-34 | 1.412418 | 0.965 | 0.937 | 7.35E-30 |
| Rpl41 | 5.45E-34 | 1.445234 | 0.965 | 0.941 | 1.28E-29 |
| Ctsz | 2.4E-33 | 0.901717 | 0.992 | 0.998 | 5.65E-29 |
| Tyrobp | 3.05E-33 | 0.973812 | 0.992 | 0.998 | 7.2E-29 |
| Srsf2 | 6.21E-33 | 1.840298 | 0.941 | 0.883 | 1.46E-28 |
| Fau | 1.25E-32 | 1.37125 | 0.938 | 0.909 | 2.95E-28 |
| Cfl1 | 2.37E-32 | 1.244019 | 0.965 | 0.933 | 5.59E-28 |
| Lgmn | 4.63E-32 | 0.595741 | 1 | 1 | 1.09E-27 |
| Ly86 | 7.38E-32 | 0.927943 | 0.988 | 0.993 | 1.74E-27 |
| Gm28661 | 4.22E-31 | 1.246369 | 0.852 | 0.639 | 9.95E-27 |
| Gm47283 | 4.88E-31 | 2.237016 | 0.754 | 0.435 | 1.15E-26 |
| Gm10443 | 5.12E-31 | 1.202522 | 0.738 | 0.411 | 1.21E-26 |
| Rnaset2a | 5.45E-31 | 1.437945 | 0.926 | 0.87 | 1.29E-26 |
| Gm1821 | 9.07E-31 | 1.336905 | 0.938 | 0.861 | 2.14E-26 |
| Gm7618 | 5.75E-30 | 0.854258 | 0.574 | 0.204 | 1.36E-25 |
| Tmem119 | 1.42E-29 | 0.819325 | 1 | 0.998 | 3.36E-25 |
| Hexb | 1.86E-29 | 0.467064 | 1 | 1 | 4.4E-25 |
| Rplp0 | 2.84E-29 | 1.538313 | 0.898 | 0.885 | 6.7E-25 |
| Slc39a1 | 6.83E-29 | 1.212785 | 0.719 | 0.37 | 1.61E-24 |
| Arglu1 | 2.39E-28 | 1.154175 | 0.961 | 0.825 | 5.63E-24 |
| Gm28437 | 6.05E-28 | 1.076699 | 0.879 | 0.753 | 1.43E-23 |
| Rps3 | 1.74E-27 | 1.456765 | 0.902 | 0.888 | 4.1E-23 |
| B2m | 1.76E-27 | 0.774834 | 1 | 1 | 4.16E-23 |
| Rgs10 | 2.5E-27 | 1.282575 | 0.914 | 0.928 | 5.9E-23 |
| Olfml3 | 1.71E-26 | 0.88933 | 0.996 | 0.996 | 4.03E-22 |
| Cyba | 1.81E-26 | 1.129355 | 0.953 | 0.963 | 4.26E-22 |
| Rps3a1 | 9.31E-26 | 1.406176 | 0.875 | 0.805 | 2.2E-21 |
| Rplp2 | 1.01E-25 | 1.242943 | 0.902 | 0.9 | 2.37E-21 |
| Gm10059 | 1.47E-25 | 0.821334 | 0.633 | 0.294 | 3.46E-21 |
| Actb | 6.2E-25 | 0.582831 | 1 | 1 | 1.46E-20 |
| Zscan26 | 1.65E-24 | 1.916024 | 0.801 | 0.675 | 3.89E-20 |
| Gm13340 | 2.06E-24 | 0.989059 | 0.816 | 0.619 | 4.86E-20 |
| Itgb5 | 2.55E-24 | -0.91802 | 0.969 | 0.991 | 6.02E-20 |
| Dpysl2 | 2.89E-24 | 1.726517 | 0.883 | 0.768 | 6.81E-20 |
| Ftl2-ps | 9.55E-24 | 1.033743 | 0.957 | 0.913 | 2.25E-19 |
| Trem2 | 9.81E-24 | 0.912825 | 0.961 | 0.987 | 2.31E-19 |
| Aif1 | 1.62E-23 | 1.510168 | 0.863 | 0.823 | 3.83E-19 |
| Gm3511 | 3.89E-23 | 1.040714 | 0.828 | 0.628 | 9.18E-19 |
| Man2b1 | 5.32E-23 | -1.1075 | 0.875 | 0.965 | 1.25E-18 |
| Gm8797 | 5.59E-23 | 0.917979 | 0.824 | 0.597 | 1.32E-18 |
| Basp1 | 5.9E-23 | 0.979286 | 0.961 | 0.965 | 1.39E-18 |
| Rpl35 | 8.4E-23 | 1.450591 | 0.848 | 0.818 | 1.98E-18 |
| Gm10800 | 8.49E-23 | -0.59239 | 0.824 | 0.954 | 2E-18 |

| | | | | | |
|---|---|---|---|---|---|
| Clic1 | 9.88E-23 | 1.648088 | 0.801 | 0.723 | 2.33E-18 |
| Gm5805 | 3.8E-22 | 1.195493 | 0.777 | 0.636 | 8.97E-18 |
| Rps18 | 6.69E-22 | 1.510051 | 0.848 | 0.829 | 1.58E-17 |
| Zfp691 | 1.48E-21 | 1.276568 | 0.781 | 0.515 | 3.49E-17 |
| Ftl1-ps1 | 1.67E-21 | 0.953072 | 0.965 | 0.903 | 3.94E-17 |
| Rps6 | 2.06E-21 | 1.450795 | 0.785 | 0.675 | 4.87E-17 |
| Sall1 | 3.25E-21 | -0.79939 | 0.656 | 0.937 | 7.67E-17 |
| Rpl21 | 4.47E-21 | 1.164367 | 0.953 | 0.937 | 1.05E-16 |
| Ncoa3 | 3.53E-20 | 1.392066 | 0.812 | 0.771 | 8.32E-16 |
| Rps19 | 3.69E-20 | 1.339741 | 0.883 | 0.875 | 8.71E-16 |
| Gm10736 | 5.95E-20 | 1.234231 | 0.727 | 0.528 | 1.4E-15 |
| Ftl1-ps2 | 6.24E-20 | 0.78706 | 0.531 | 0.24 | 1.47E-15 |
| Gm13341 | 8.94E-20 | 0.743136 | 0.457 | 0.177 | 2.11E-15 |
| Gm11942 | 9.23E-20 | 0.817921 | 0.684 | 0.385 | 2.18E-15 |
| Rpl13a | 1.06E-19 | 1.213293 | 0.855 | 0.799 | 2.51E-15 |
| Rpl36 | 1.11E-19 | 1.347517 | 0.875 | 0.814 | 2.61E-15 |
| Rpl4 | 1.85E-19 | 1.403769 | 0.824 | 0.823 | 4.36E-15 |
| mt-Atp8 | 2.56E-19 | 0.562076 | 0.328 | 0.084 | 6.04E-15 |
| Gm29216 | 4.41E-19 | 0.801156 | 0.883 | 0.768 | 1.04E-14 |
| Gm6977 | 7.15E-19 | 0.951613 | 0.711 | 0.487 | 1.69E-14 |
| Ctsl | 7.27E-19 | 0.878608 | 0.938 | 0.944 | 1.71E-14 |
| Gng10 | 8.25E-19 | 1.557056 | 0.723 | 0.597 | 1.95E-14 |
| Gm3362 | 9.31E-19 | 1.103243 | 0.633 | 0.37 | 2.19E-14 |
| Cd14 | 9.64E-19 | 1.275704 | 0.828 | 0.799 | 2.27E-14 |
| Cmtm6 | 1.12E-18 | -1.34793 | 0.641 | 0.81 | 2.65E-14 |
| Rpl17 | 1.54E-18 | 1.518295 | 0.82 | 0.82 | 3.63E-14 |
| Itpripl1 | 1.62E-18 | 2.300939 | 0.695 | 0.589 | 3.83E-14 |
| Plxnb2 | 2.44E-18 | -0.76992 | 0.531 | 0.849 | 5.76E-14 |
| Gm17066 | 2.48E-18 | 1.275132 | 0.684 | 0.496 | 5.84E-14 |
| Gm10073 | 2.55E-18 | 0.759423 | 0.797 | 0.578 | 6.01E-14 |
| Ptafr | 2.63E-18 | 1.637303 | 0.789 | 0.686 | 6.2E-14 |
| Rnaset2b | 4.05E-18 | 1.32473 | 0.93 | 0.87 | 9.54E-14 |
| Rpl30 | 5.14E-18 | 1.379499 | 0.871 | 0.862 | 1.21E-13 |
| Plxdc2 | 7.1E-18 | -0.98196 | 0.855 | 0.955 | 1.67E-13 |
| Bst2 | 7.88E-18 | 2.502859 | 0.574 | 0.372 | 1.86E-13 |
| Gm10123 | 9.25E-18 | 0.897002 | 0.949 | 0.901 | 2.18E-13 |
| Sirpa | 1.47E-17 | -0.76748 | 0.895 | 0.976 | 3.47E-13 |
| Gm9843 | 1.48E-17 | 1.133857 | 0.926 | 0.87 | 3.5E-13 |
| Gm11478 | 1.62E-17 | 0.850202 | 0.875 | 0.749 | 3.82E-13 |
| Sat1 | 1.63E-17 | 1.348091 | 0.836 | 0.846 | 3.84E-13 |
| Rsrp1 | 1.71E-17 | 1.110736 | 0.969 | 0.92 | 4.02E-13 |
| Rps25 | 1.9E-17 | 1.444652 | 0.879 | 0.809 | 4.48E-13 |
| Rpl27a | 1.92E-17 | 1.635977 | 0.789 | 0.719 | 4.53E-13 |
| Rpl10 | 2.28E-17 | 1.06388 | 0.887 | 0.905 | 5.37E-13 |
| Sun2 | 2.31E-17 | -1.85165 | 0.453 | 0.704 | 5.44E-13 |
| Rps7 | 3.87E-17 | 1.467226 | 0.797 | 0.794 | 9.13E-13 |
| Rpl23a-ps3 | 4.64E-17 | 0.963715 | 0.695 | 0.5 | 1.09E-12 |

| | | | | | |
|---|---|---|---|---|---|
| Rpl23 | 4.65E-17 | 1.370475 | 0.863 | 0.855 | 1.1E-12 |
| Gm15590 | 8.07E-17 | 0.602753 | 0.707 | 0.468 | 1.9E-12 |
| Rhob | 1.57E-16 | -0.84254 | 0.887 | 0.939 | 3.71E-12 |
| Tmem50a | 3.31E-16 | 1.274311 | 0.832 | 0.79 | 7.81E-12 |
| Rps24 | 4.48E-16 | 1.366812 | 0.871 | 0.896 | 1.06E-11 |
| Tm9sf2 | 5.73E-16 | -1.40484 | 0.633 | 0.783 | 1.35E-11 |
| Rps27a | 5.74E-16 | 1.437633 | 0.809 | 0.784 | 1.36E-11 |
| Oaz1 | 7.02E-16 | 1.549218 | 0.746 | 0.643 | 1.66E-11 |
| Rpl15 | 7.24E-16 | 1.45836 | 0.707 | 0.641 | 1.71E-11 |
| Wdr81 | 7.81E-16 | -0.83025 | 0.219 | 0.52 | 1.84E-11 |
| Akna | 8.73E-16 | -0.57867 | 0.27 | 0.613 | 2.06E-11 |
| Rpl32 | 9.73E-16 | 1.204792 | 0.891 | 0.924 | 2.3E-11 |
| Calm1 | 1E-15 | 1.599025 | 0.758 | 0.686 | 2.37E-11 |
| Rpl13-ps3 | 1.1E-15 | 0.539054 | 0.293 | 0.082 | 2.59E-11 |
| Gm10076 | 1.16E-15 | 0.959605 | 0.922 | 0.907 | 2.74E-11 |
| Gm9625 | 1.5E-15 | 0.886993 | 0.734 | 0.552 | 3.54E-11 |
| Laptm5 | 1.56E-15 | 0.463463 | 1 | 1 | 3.69E-11 |
| Rps16 | 2.16E-15 | 1.45552 | 0.777 | 0.77 | 5.09E-11 |
| Gm5735 | 2.17E-15 | 0.365062 | 0.254 | 0.059 | 5.12E-11 |
| Gm26917 | 2.21E-15 | -0.81869 | 0.984 | 0.994 | 5.2E-11 |
| Gm15427 | 2.31E-15 | 0.703069 | 0.977 | 0.97 | 5.46E-11 |
| Rpl17-ps8 | 3.24E-15 | 1.248981 | 0.324 | 0.104 | 7.65E-11 |
| Tmem176b | 4.64E-15 | 1.045682 | 0.879 | 0.881 | 1.09E-10 |
| Tmem59 | 5.01E-15 | 1.213703 | 0.805 | 0.771 | 1.18E-10 |
| Soga1 | 5.11E-15 | -0.57011 | 0.414 | 0.725 | 1.2E-10 |
| Npc2 | 5.27E-15 | 1.189323 | 0.832 | 0.829 | 1.24E-10 |
| Skil | 6.05E-15 | 1.477123 | 0.766 | 0.68 | 1.43E-10 |
| Serinc3 | 7.12E-15 | -0.44276 | 0.992 | 1 | 1.68E-10 |
| Serf2 | 7.33E-15 | 1.224878 | 0.77 | 0.656 | 1.73E-10 |
| Rbm26 | 1.05E-14 | 1.508933 | 0.723 | 0.654 | 2.47E-10 |
| Slc25a5 | 1.57E-14 | 1.749236 | 0.73 | 0.66 | 3.71E-10 |
| Rps12 | 1.59E-14 | 1.056709 | 0.879 | 0.903 | 3.75E-10 |
| mt-Nd1 | 1.7E-14 | 1.089186 | 0.91 | 0.872 | 4.01E-10 |
| Mir692-2 | 1.89E-14 | 0.691969 | 0.598 | 0.359 | 4.45E-10 |
| Gm5560 | 2.11E-14 | 0.744749 | 0.434 | 0.197 | 4.98E-10 |
| Cd164 | 2.37E-14 | -0.78122 | 0.75 | 0.888 | 5.58E-10 |
| Rps4x | 2.95E-14 | 1.137807 | 0.836 | 0.825 | 6.95E-10 |
| Gm10263 | 3.04E-14 | 0.446971 | 0.273 | 0.078 | 7.17E-10 |
| Sft2d1 | 3.95E-14 | 1.304914 | 0.766 | 0.773 | 9.31E-10 |
| Rpl27-ps3 | 4E-14 | 1.350313 | 0.543 | 0.314 | 9.44E-10 |
| Bhlhe41 | 4.63E-14 | 1.28557 | 0.836 | 0.753 | 1.09E-09 |
| Rreb1 | 5.19E-14 | -1.21471 | 0.184 | 0.468 | 1.23E-09 |
| Rpl27 | 5.43E-14 | 1.457262 | 0.723 | 0.658 | 1.28E-09 |
| Gm8960 | 5.68E-14 | 0.661505 | 0.477 | 0.23 | 1.34E-09 |
| Gm2000 | 6.62E-14 | 0.921962 | 0.758 | 0.612 | 1.56E-09 |
| 1700017B0 | 9.81E-14 | -0.56435 | 0.426 | 0.703 | 2.31E-09 |
| H3f3b | 9.98E-14 | 1.178117 | 0.82 | 0.786 | 2.35E-09 |

| | | | | | |
|---|---|---|---|---|---|
| Nckap1l | 1.14E-13 | -0.73545 | 0.5 | 0.757 | 2.68E-09 |
| Oasl2 | 1.38E-13 | 2.742759 | 0.23 | 0.056 | 3.25E-09 |
| Tgfbr2 | 1.63E-13 | -0.40936 | 0.77 | 0.918 | 3.85E-09 |
| Gm21738 | 1.64E-13 | -0.6884 | 0.41 | 0.678 | 3.88E-09 |
| Cx3cr1 | 1.71E-13 | -0.45622 | 1 | 0.998 | 4.03E-09 |
| Rps3a3 | 1.73E-13 | 0.770783 | 0.559 | 0.342 | 4.09E-09 |
| Sod1 | 1.76E-13 | 1.975502 | 0.629 | 0.498 | 4.15E-09 |
| H2-D1 | 1.79E-13 | 0.955794 | 0.914 | 0.955 | 4.23E-09 |
| Hsp90ab1 | 1.96E-13 | 1.255169 | 0.844 | 0.781 | 4.63E-09 |
| Gm11361 | 2.49E-13 | 0.864441 | 0.648 | 0.459 | 5.87E-09 |
| Gm15710 | 2.59E-13 | 0.836084 | 0.824 | 0.736 | 6.1E-09 |
| Gm7536 | 2.65E-13 | 1.046541 | 0.727 | 0.578 | 6.26E-09 |
| Hspa8 | 2.66E-13 | 0.874413 | 0.918 | 0.857 | 6.28E-09 |
| P2ry13 | 2.77E-13 | -0.81767 | 0.895 | 0.959 | 6.53E-09 |
| Rpsa | 3.55E-13 | 1.262357 | 0.828 | 0.816 | 8.37E-09 |
| Man1c1 | 3.6E-13 | -1.72949 | 0.324 | 0.561 | 8.49E-09 |
| Rtp4 | 3.65E-13 | 3.097013 | 0.332 | 0.138 | 8.61E-09 |
| Dhrs3 | 3.65E-13 | 1.343781 | 0.828 | 0.81 | 8.61E-09 |
| Pld4 | 4.26E-13 | 0.729705 | 0.91 | 0.942 | 1.01E-08 |
| Gm4613 | 5.25E-13 | 0.420745 | 0.285 | 0.093 | 1.24E-08 |
| Gcn1l1 | 5.7E-13 | -0.74331 | 0.211 | 0.498 | 1.34E-08 |
| Gm10925 | 6.43E-13 | 0.775669 | 0.859 | 0.747 | 1.52E-08 |
| Rpl14 | 7.27E-13 | 1.481326 | 0.77 | 0.775 | 1.71E-08 |
| A630001G2 | 8.05E-13 | -0.89127 | 0.285 | 0.571 | 1.9E-08 |
| Ac154187.2 | 8.55E-13 | 0.516095 | 0.211 | 0.048 | 2.02E-08 |
| Oaz1-ps | 9.06E-13 | 1.269942 | 0.656 | 0.494 | 2.14E-08 |
| Gm5529 | 9.39E-13 | 0.701143 | 0.422 | 0.206 | 2.21E-08 |
| Rpl39 | 1.33E-12 | 1.562482 | 0.789 | 0.796 | 3.14E-08 |
| Ac123956.1 | 1.4E-12 | 0.43859 | 0.266 | 0.084 | 3.3E-08 |
| Rpl36-ps2 | 2.5E-12 | 0.933601 | 0.703 | 0.498 | 5.9E-08 |
| Tubb5 | 2.85E-12 | 1.193845 | 0.766 | 0.697 | 6.72E-08 |
| Gm4617 | 2.86E-12 | 1.015864 | 0.402 | 0.186 | 6.75E-08 |
| Gm26857 | 3.33E-12 | -1.24251 | 0.117 | 0.353 | 7.86E-08 |
| Rpl26 | 3.37E-12 | 1.559824 | 0.762 | 0.77 | 7.95E-08 |
| Sall3 | 3.59E-12 | -0.7982 | 0.488 | 0.747 | 8.47E-08 |
| Il17ra | 3.79E-12 | -1.09858 | 0.379 | 0.623 | 8.94E-08 |
| Slc7a8 | 4.01E-12 | -0.53701 | 0.43 | 0.714 | 9.45E-08 |
| Gm12918 | 4.94E-12 | 0.820654 | 0.672 | 0.504 | 1.17E-07 |
| Rps20 | 5.11E-12 | 1.055379 | 0.863 | 0.885 | 1.2E-07 |
| Gnb1 | 7.69E-12 | 1.495441 | 0.836 | 0.827 | 1.81E-07 |
| Rpl28 | 7.81E-12 | 1.410819 | 0.742 | 0.734 | 1.84E-07 |
| Ac165079.1 | 8.32E-12 | 0.631903 | 0.387 | 0.18 | 1.96E-07 |
| Gm10180 | 1.01E-11 | 0.494615 | 0.527 | 0.325 | 2.38E-07 |
| Rps16-ps2 | 1.01E-11 | 1.163798 | 0.715 | 0.604 | 2.38E-07 |
| Pros1 | 1.1E-11 | -0.70207 | 0.617 | 0.771 | 2.6E-07 |
| Gm34280 | 1.21E-11 | -0.67686 | 0.176 | 0.431 | 2.86E-07 |
| Rpl36-ps12 | 1.25E-11 | 0.282714 | 0.457 | 0.236 | 2.94E-07 |

| | | | | | |
|---|---|---|---|---|---|
| Gm13192 | 1.48E-11 | 0.354562 | 0.254 | 0.082 | 3.5E-07 |
| Gm12166 | 1.49E-11 | 1.141936 | 0.707 | 0.634 | 3.51E-07 |
| Gm8464 | 1.82E-11 | 0.500746 | 0.367 | 0.173 | 4.29E-07 |
| Rpl17-ps5 | 1.98E-11 | 0.584163 | 0.441 | 0.227 | 4.68E-07 |
| Gdi2 | 2.6E-11 | 0.935131 | 0.906 | 0.835 | 6.13E-07 |
| Gm11518 | 2.68E-11 | 1.215997 | 0.672 | 0.571 | 6.33E-07 |
| Tpm3 | 3.33E-11 | 1.154571 | 0.879 | 0.874 | 7.85E-07 |
| Ptma | 3.8E-11 | 1.720791 | 0.637 | 0.526 | 8.98E-07 |
| Gm13408 | 4.14E-11 | 1.501713 | 0.395 | 0.195 | 9.76E-07 |
| Gm22774 | 4.29E-11 | 0.457541 | 0.301 | 0.119 | 1.01E-06 |
| Sbno2 | 4.32E-11 | -0.94291 | 0.223 | 0.468 | 1.02E-06 |
| Gm4609 | 4.62E-11 | -0.69852 | 0.152 | 0.372 | 1.09E-06 |
| Gm14173 | 4.71E-11 | 0.353387 | 0.281 | 0.102 | 1.11E-06 |
| Abca1 | 5.03E-11 | -1.01154 | 0.324 | 0.576 | 1.19E-06 |
| Rpl6 | 5.3E-11 | 1.420576 | 0.777 | 0.749 | 1.25E-06 |
| Mob1a | 5.59E-11 | -0.47456 | 0.438 | 0.708 | 1.32E-06 |
| Rpl7 | 6.06E-11 | 1.203458 | 0.797 | 0.792 | 1.43E-06 |
| Gm44860 | 6.07E-11 | -0.77469 | 0.344 | 0.606 | 1.43E-06 |
| Tmed7 | 6.39E-11 | -0.72023 | 0.18 | 0.416 | 1.51E-06 |
| Cd33 | 6.51E-11 | -0.47808 | 0.617 | 0.851 | 1.54E-06 |
| Elmo1 | 6.86E-11 | -0.78046 | 0.578 | 0.771 | 1.62E-06 |
| Adgre1 | 7.16E-11 | -1.25166 | 0.426 | 0.63 | 1.69E-06 |
| Calm2 | 7.33E-11 | 1.119158 | 0.863 | 0.883 | 1.73E-06 |
| B4galt1 | 7.4E-11 | 1.024093 | 0.832 | 0.788 | 1.75E-06 |
| Rpl3 | 7.54E-11 | 1.064094 | 0.816 | 0.853 | 1.78E-06 |
| Gm9512 | 8.37E-11 | 0.353862 | 0.242 | 0.078 | 1.97E-06 |
| Rpl11 | 8.42E-11 | 1.678385 | 0.723 | 0.684 | 1.99E-06 |
| Rnu3a | 8.46E-11 | -0.78443 | 0.055 | 0.251 | 2E-06 |
| Gm6988 | 8.84E-11 | 0.477539 | 0.316 | 0.128 | 2.09E-06 |
| Gm13456 | 1.02E-10 | 0.509203 | 0.98 | 0.922 | 2.4E-06 |
| Rpl17-ps3 | 1.21E-10 | 0.845984 | 0.566 | 0.381 | 2.84E-06 |
| Slfn2 | 1.23E-10 | 2.250531 | 0.375 | 0.203 | 2.91E-06 |
| Alox5ap | 1.48E-10 | 1.404658 | 0.723 | 0.691 | 3.49E-06 |
| Lrrc8d | 1.83E-10 | -1.83091 | 0.141 | 0.355 | 4.31E-06 |
| Anxa3 | 1.83E-10 | 1.424061 | 0.75 | 0.73 | 4.32E-06 |
| Nuak1 | 2.32E-10 | -0.32074 | 0.328 | 0.608 | 5.48E-06 |
| Ccnd3 | 2.41E-10 | -1.73244 | 0.266 | 0.474 | 5.69E-06 |
| Rps25-ps1 | 2.59E-10 | 0.598055 | 0.426 | 0.221 | 6.11E-06 |
| Ac154540.: | 2.63E-10 | 0.463563 | 0.305 | 0.126 | 6.21E-06 |
| Gm10221 | 2.73E-10 | 0.49531 | 0.262 | 0.095 | 6.44E-06 |
| Gm10020 | 2.95E-10 | 0.541426 | 0.34 | 0.158 | 6.95E-06 |
| Scaf1 | 3.93E-10 | -0.2614 | 0.273 | 0.524 | 9.28E-06 |
| Gapdh | 4.84E-10 | 0.95561 | 0.789 | 0.833 | 1.14E-05 |
| Gm8129 | 5.12E-10 | 1.059114 | 0.527 | 0.366 | 1.21E-05 |
| Gm8566 | 5.24E-10 | 0.395057 | 0.262 | 0.099 | 1.24E-05 |
| Gm6180 | 5.75E-10 | 0.643049 | 0.527 | 0.342 | 1.36E-05 |
| Gm8430 | 6.69E-10 | 0.913144 | 0.719 | 0.565 | 1.58E-05 |

| | | | | | |
|---|---|---|---|---|---|
| Rpsa-ps9 | 6.72E-10 | 0.609427 | 0.43 | 0.253 | 1.58E-05 |
| Csf3r | 7.26E-10 | -0.55121 | 0.746 | 0.948 | 1.71E-05 |
| Gm14165 | 7.34E-10 | 0.602632 | 0.488 | 0.305 | 1.73E-05 |
| Gm5822 | 7.59E-10 | 0.477008 | 0.246 | 0.089 | 1.79E-05 |
| Gm9800 | 7.93E-10 | 0.947904 | 0.344 | 0.16 | 1.87E-05 |
| Hk2 | 8.03E-10 | 1.215995 | 0.801 | 0.797 | 1.89E-05 |
| Slfn5 | 8.08E-10 | 2.712555 | 0.246 | 0.089 | 1.91E-05 |
| Bsg | 8.19E-10 | 1.100988 | 0.742 | 0.779 | 1.93E-05 |
| Kif21b | 8.7E-10 | -0.37401 | 0.297 | 0.548 | 2.05E-05 |
| Mir703 | 9.64E-10 | 0.52515 | 0.418 | 0.227 | 2.27E-05 |
| Nptn | 1.02E-09 | -1.11458 | 0.523 | 0.662 | 2.41E-05 |
| Rps11-ps2 | 1.04E-09 | -0.85542 | 0.352 | 0.164 | 2.45E-05 |
| Gm7589 | 1.1E-09 | 0.431638 | 0.242 | 0.087 | 2.59E-05 |
| Gm8730 | 1.11E-09 | 0.766543 | 0.812 | 0.764 | 2.62E-05 |
| Dusp7 | 1.28E-09 | -0.53457 | 0.367 | 0.597 | 3.02E-05 |
| Rpl10a | 1.28E-09 | 1.503773 | 0.73 | 0.693 | 3.03E-05 |
| Rpl38 | 1.3E-09 | 1.174501 | 0.672 | 0.599 | 3.07E-05 |
| Rpl15-ps3 | 1.31E-09 | 0.645426 | 0.383 | 0.201 | 3.09E-05 |
| Spcs2 | 1.43E-09 | 1.599538 | 0.578 | 0.472 | 3.37E-05 |
| Ly6e | 1.47E-09 | 0.878069 | 0.969 | 0.985 | 3.47E-05 |
| Notch1 | 1.61E-09 | -0.912 | 0.199 | 0.429 | 3.79E-05 |
| Rps10 | 1.67E-09 | 1.016894 | 0.82 | 0.838 | 3.93E-05 |
| Gm2387 | 1.73E-09 | 0.43908 | 0.305 | 0.136 | 4.07E-05 |
| Ctcf | 1.75E-09 | -1.14881 | 0.082 | 0.27 | 4.13E-05 |
| Lrp5 | 1.78E-09 | -1.62621 | 0.102 | 0.29 | 4.2E-05 |
| Rpl23a | 1.83E-09 | 1.014115 | 0.75 | 0.784 | 4.32E-05 |
| E230029C0 | 1.86E-09 | -0.34267 | 0.598 | 0.874 | 4.39E-05 |
| Rin3 | 1.94E-09 | -0.9003 | 0.191 | 0.403 | 4.57E-05 |
| Rpl7a-ps7 | 1.94E-09 | 0.832711 | 0.418 | 0.242 | 4.58E-05 |
| Kdelr2 | 2E-09 | 1.106408 | 0.59 | 0.398 | 4.71E-05 |
| Gm26825 | 2.01E-09 | 0.974518 | 0.695 | 0.602 | 4.74E-05 |
| Gm17147 | 2.09E-09 | -1.2307 | 0.148 | 0.359 | 4.93E-05 |
| Dtx3l | 2.82E-09 | 1.288958 | 0.344 | 0.162 | 6.66E-05 |
| Rpl34 | 3.01E-09 | 1.261783 | 0.754 | 0.797 | 7.1E-05 |
| Cbx3 | 3.06E-09 | 0.287785 | 0.477 | 0.288 | 7.22E-05 |
| Gm8451 | 3.15E-09 | 0.513154 | 0.367 | 0.182 | 7.44E-05 |
| Lpgat1 | 3.17E-09 | -0.34545 | 0.133 | 0.342 | 7.48E-05 |
| Rbm39 | 3.52E-09 | 0.963805 | 0.887 | 0.935 | 8.3E-05 |
| Rps23 | 3.65E-09 | 0.810105 | 0.863 | 0.898 | 8.61E-05 |
| Slc37a2 | 4.05E-09 | -1.39262 | 0.316 | 0.53 | 9.55E-05 |
| Rpl3-ps2 | 4.07E-09 | 0.462545 | 0.324 | 0.156 | 9.6E-05 |
| Rps13 | 4.57E-09 | 1.187206 | 0.793 | 0.818 | 0.000108 |
| Gm10288 | 4.82E-09 | 0.913614 | 0.555 | 0.392 | 0.000114 |
| Gm2225 | 4.82E-09 | 0.494471 | 0.359 | 0.184 | 0.000114 |
| Nfatc1 | 4.87E-09 | -0.6699 | 0.352 | 0.599 | 0.000115 |
| Tlr7 | 5.16E-09 | -0.3644 | 0.309 | 0.528 | 0.000122 |
| Gm6548 | 5.79E-09 | 1.371925 | 0.402 | 0.21 | 0.000137 |

| | | | | | |
|---|---|---|---|---|---|
| Fam221b | 6.13E-09 | 0.894023 | 0.582 | 0.396 | 0.000145 |
| Man2b2 | 6.48E-09 | -1.78044 | 0.277 | 0.452 | 0.000153 |
| Klhl9 | 6.51E-09 | 1.734693 | 0.492 | 0.357 | 0.000154 |
| Ifi204 | 7.74E-09 | 2.524267 | 0.234 | 0.089 | 0.000183 |
| Gm5148 | 7.94E-09 | -0.84643 | 0.473 | 0.593 | 0.000187 |
| Slc8a1 | 8.15E-09 | -2.67396 | 0.074 | 0.245 | 0.000192 |
| H2-K1 | 8.23E-09 | 1.000359 | 0.828 | 0.857 | 0.000194 |
| Adam10 | 8.25E-09 | 1.019205 | 0.777 | 0.719 | 0.000195 |
| Wtap | 8.51E-09 | 0.715 | 0.605 | 0.375 | 0.000201 |
| Gm9616 | 8.9E-09 | 0.555136 | 0.641 | 0.467 | 0.00021 |
| Wdr89 | 1.02E-08 | 0.66992 | 0.777 | 0.671 | 0.000241 |
| Fchsd2 | 1.1E-08 | -0.37236 | 0.27 | 0.504 | 0.00026 |
| Rpl35a | 1.17E-08 | 1.371364 | 0.762 | 0.758 | 0.000277 |
| Laptm4a | 1.19E-08 | 0.831814 | 0.844 | 0.849 | 0.000281 |
| Abcc3 | 1.2E-08 | -0.73175 | 0.164 | 0.372 | 0.000283 |
| Gm4202 | 1.24E-08 | -0.69419 | 0.375 | 0.572 | 0.000292 |
| Gm6394 | 1.24E-08 | 0.548743 | 0.707 | 0.539 | 0.000293 |
| Plcg2 | 1.3E-08 | -0.81722 | 0.223 | 0.442 | 0.000307 |
| Amfr | 1.57E-08 | -1.60134 | 0.301 | 0.481 | 0.00037 |
| Zmiz1 | 1.66E-08 | -0.47083 | 0.516 | 0.736 | 0.000392 |
| Fcgr3 | 1.72E-08 | 0.64766 | 0.906 | 0.946 | 0.000405 |
| Gm3379 | 1.77E-08 | 0.947247 | 0.668 | 0.565 | 0.000417 |
| Rpl18-ps1 | 1.82E-08 | 0.465831 | 0.379 | 0.21 | 0.00043 |
| Spcs2-ps | 1.82E-08 | 0.373703 | 0.23 | 0.089 | 0.00043 |
| Tnfrsf21 | 1.88E-08 | -0.92454 | 0.309 | 0.489 | 0.000445 |
| Aftph | 1.96E-08 | -0.45692 | 0.227 | 0.439 | 0.000462 |
| Fcgr1 | 2.1E-08 | 1.312187 | 0.758 | 0.743 | 0.000494 |
| Sfswap | 2.22E-08 | -1.30489 | 0.121 | 0.301 | 0.000524 |
| Rapgef1 | 2.22E-08 | -0.99462 | 0.234 | 0.428 | 0.000524 |
| Mib2 | 2.27E-08 | -1.06308 | 0.117 | 0.299 | 0.000537 |
| Olfr620 | 2.41E-08 | -0.38589 | 0.148 | 0.333 | 0.000569 |
| Gm6170 | 2.56E-08 | 0.427606 | 0.52 | 0.34 | 0.000603 |
| Gm6210 | 2.6E-08 | 0.476421 | 0.441 | 0.273 | 0.000613 |
| H2-Ab1 | 2.74E-08 | 0.690512 | 0.082 | 0.253 | 0.000646 |
| Gm14760 | 2.79E-08 | 0.366737 | 0.203 | 0.072 | 0.000659 |
| Usp18 | 3.06E-08 | 2.724457 | 0.266 | 0.123 | 0.000722 |
| Rpl5 | 3.06E-08 | 1.406941 | 0.789 | 0.783 | 0.000722 |
| Pdgfb | 3.16E-08 | -0.39626 | 0.355 | 0.58 | 0.000745 |
| Tlr13 | 3.23E-08 | -0.28946 | 0.18 | 0.387 | 0.000761 |
| F11r | 3.24E-08 | 0.599282 | 0.961 | 0.981 | 0.000764 |
| Hltf | 3.29E-08 | -0.49307 | 0.086 | 0.258 | 0.000775 |
| Gm4149 | 3.31E-08 | 0.705946 | 0.906 | 0.861 | 0.00078 |
| Ubc | 3.59E-08 | 0.66106 | 0.941 | 0.952 | 0.000847 |
| Gm32849 | 3.59E-08 | -2.1246 | 0.051 | 0.201 | 0.000848 |
| Prkcd | 3.7E-08 | -0.77957 | 0.676 | 0.82 | 0.000872 |
| Gm42418 | 4E-08 | -0.40682 | 0.848 | 0.978 | 0.000945 |
| Set | 4.06E-08 | 1.475689 | 0.582 | 0.487 | 0.000957 |

| | | | | | |
|---|---|---|---|---|---|
| Rpl19 | 4.08E-08 | 0.973843 | 0.773 | 0.848 | 0.000963 |
| Rps21 | 4.16E-08 | 1.123406 | 0.801 | 0.851 | 0.00098 |
| Arpc1b | 4.31E-08 | 1.261327 | 0.75 | 0.784 | 0.001017 |
| Eef2 | 4.36E-08 | -0.77081 | 0.699 | 0.812 | 0.00103 |
| Arsb | 4.47E-08 | -0.76532 | 0.516 | 0.688 | 0.001054 |
| Arap1 | 4.47E-08 | -0.36718 | 0.242 | 0.463 | 0.001054 |
| Bag6 | 4.56E-08 | -0.59674 | 0.078 | 0.243 | 0.001075 |
| Lhfpl2 | 4.81E-08 | -0.28063 | 0.336 | 0.546 | 0.001135 |
| Rpl30-ps10 | 4.83E-08 | 0.390027 | 0.391 | 0.21 | 0.001139 |
| Rps6-ps4 | 4.98E-08 | 0.674485 | 0.602 | 0.455 | 0.001175 |
| Camta2 | 5.45E-08 | -1.07543 | 0.191 | 0.385 | 0.001285 |
| Gm4824 | 5.51E-08 | -0.41689 | 0.215 | 0.08 | 0.0013 |
| Ac140300.: | 5.75E-08 | 0.392534 | 0.336 | 0.175 | 0.001357 |
| Ctsa | 5.79E-08 | 0.617737 | 0.914 | 0.978 | 0.001366 |
| Gm9385 | 6E-08 | 0.61734 | 0.508 | 0.336 | 0.001417 |
| Actr2 | 6.86E-08 | 0.981503 | 0.883 | 0.842 | 0.001619 |
| Syngr2 | 7.21E-08 | 1.044268 | 0.738 | 0.717 | 0.001701 |
| Abca9 | 7.78E-08 | -0.45809 | 0.57 | 0.775 | 0.001835 |
| Rps5 | 7.96E-08 | 1.339647 | 0.723 | 0.734 | 0.001878 |
| Daglb | 8.21E-08 | -1.17798 | 0.656 | 0.703 | 0.001938 |
| Ifngr1 | 8.45E-08 | 0.527512 | 0.973 | 0.95 | 0.001993 |
| Gm10240 | 9.59E-08 | 0.913826 | 0.547 | 0.416 | 0.002262 |
| Gm5637 | 9.6E-08 | 0.849697 | 0.496 | 0.355 | 0.002264 |
| Slc23a2 | 9.61E-08 | -1.48829 | 0.113 | 0.283 | 0.002266 |
| Pald1 | 9.68E-08 | -0.40092 | 0.262 | 0.457 | 0.002282 |
| Tnrc18 | 1.01E-07 | -0.41538 | 0.137 | 0.316 | 0.002382 |
| Slc43a2 | 1.04E-07 | -1.25012 | 0.172 | 0.355 | 0.002454 |
| Mlxip | 1.04E-07 | -0.6147 | 0.227 | 0.424 | 0.002459 |
| Kcnk6 | 1.05E-07 | -0.41367 | 0.438 | 0.69 | 0.002487 |
| Neat1 | 1.06E-07 | -0.86636 | 0.262 | 0.467 | 0.002498 |
| Ogt | 1.13E-07 | -0.4529 | 0.5 | 0.725 | 0.002659 |
| Gm4294 | 1.2E-07 | 0.663735 | 0.438 | 0.294 | 0.002836 |
| St13 | 1.22E-07 | 1.64994 | 0.605 | 0.511 | 0.002879 |
| Cd68 | 1.24E-07 | 0.798966 | 0.887 | 0.926 | 0.002923 |
| Gm7600 | 1.25E-07 | 0.342295 | 0.242 | 0.102 | 0.002939 |
| Myadm | 1.27E-07 | -0.8497 | 0.09 | 0.247 | 0.002985 |
| Epb41l2 | 1.29E-07 | -0.3151 | 0.812 | 0.948 | 0.003045 |
| Rpl18 | 1.32E-07 | 1.20831 | 0.734 | 0.829 | 0.003118 |
| Snx22 | 1.38E-07 | -0.71398 | 0.355 | 0.191 | 0.003256 |
| Rasa1 | 1.44E-07 | -1.31839 | 0.297 | 0.465 | 0.003399 |
| Usf2 | 1.54E-07 | -0.25794 | 0.156 | 0.351 | 0.003627 |
| Fbxw11 | 1.6E-07 | -1.13724 | 0.184 | 0.359 | 0.003765 |
| Cdk13 | 1.6E-07 | -0.87686 | 0.281 | 0.481 | 0.00377 |
| Lrch1 | 1.68E-07 | -0.47269 | 0.328 | 0.546 | 0.003962 |
| Cox4i1 | 1.68E-07 | 1.230821 | 0.699 | 0.73 | 0.003969 |
| Siglech | 1.74E-07 | 0.389189 | 1 | 0.998 | 0.004108 |
| Abhd12 | 1.77E-07 | 0.453949 | 0.957 | 0.98 | 0.004166 |

| | | | | | |
|---|---|---|---|---|---|
| Rps27a-ps2 | 1.83E-07 | 0.986729 | 0.453 | 0.294 | 0.004306 |
| Atp6v0b | 1.89E-07 | 1.027214 | 0.805 | 0.874 | 0.004449 |
| Cic | 1.9E-07 | 0.300803 | 0.148 | 0.329 | 0.004494 |
| Fnbp1 | 1.98E-07 | -0.25225 | 0.402 | 0.613 | 0.004673 |
| Ccr5 | 1.98E-07 | -1.15943 | 0.711 | 0.773 | 0.004682 |
| Dnm2 | 1.99E-07 | -0.28589 | 0.504 | 0.712 | 0.0047 |
| Lrp1 | 2.03E-07 | -0.35012 | 0.734 | 0.9 | 0.004777 |
| Kansl1 | 2.09E-07 | -0.33356 | 0.387 | 0.589 | 0.004941 |
| Sec24b | 2.12E-07 | -1.00549 | 0.156 | 0.331 | 0.004994 |
| Rin2 | 2.15E-07 | -1.08887 | 0.398 | 0.587 | 0.005081 |
| Tpt1 | 2.16E-07 | 0.812274 | 0.875 | 0.924 | 0.0051 |
| Tmf1 | 2.27E-07 | 0.385487 | 0.59 | 0.396 | 0.005348 |
| Lrrc8a | 2.36E-07 | 2.009235 | 0.492 | 0.392 | 0.005579 |
| Irf2bpl | 2.37E-07 | -0.98563 | 0.207 | 0.379 | 0.005581 |
| Ppib | 2.44E-07 | 1.076554 | 0.656 | 0.626 | 0.005759 |
| Spi1 | 2.46E-07 | 1.087281 | 0.762 | 0.816 | 0.005796 |
| Arhgdib | 2.61E-07 | 1.462122 | 0.684 | 0.69 | 0.006158 |
| Gm10224 | 2.68E-07 | 0.474334 | 0.297 | 0.16 | 0.006326 |
| Ifitm3 | 2.71E-07 | 3.077387 | 0.344 | 0.214 | 0.006396 |
| Scamp5 | 2.79E-07 | -0.48306 | 0.355 | 0.55 | 0.006571 |
| Sowahc | 2.9E-07 | -0.85423 | 0.266 | 0.463 | 0.006845 |
| Gm6136 | 3.04E-07 | 0.714769 | 0.652 | 0.52 | 0.007161 |
| Ikzf1 | 3.06E-07 | -0.3904 | 0.457 | 0.662 | 0.00722 |
| Mpeg1 | 3.11E-07 | -0.31377 | 0.836 | 0.942 | 0.007345 |
| Sipa1 | 3.14E-07 | -0.54909 | 0.473 | 0.651 | 0.007399 |
| Evi2a | 3.44E-07 | 1.215216 | 0.684 | 0.621 | 0.008118 |
| Pik3cg | 3.49E-07 | -0.37585 | 0.273 | 0.465 | 0.008241 |
| Rpl10a-ps1 | 3.52E-07 | 0.785324 | 0.719 | 0.602 | 0.0083 |
| Rps17 | 3.6E-07 | 1.152619 | 0.715 | 0.719 | 0.008504 |
| Csde1 | 3.67E-07 | -1.33834 | 0.297 | 0.472 | 0.008655 |
| Lyn | 3.7E-07 | -0.61078 | 0.797 | 0.872 | 0.008727 |
| Epn1 | 3.83E-07 | -0.98617 | 0.27 | 0.448 | 0.009024 |
| Slc40a1 | 3.85E-07 | -0.6031 | 0.27 | 0.442 | 0.00909 |
| Sec62 | 3.96E-07 | 1.196223 | 0.59 | 0.517 | 0.009335 |
| Asph | 3.97E-07 | -0.79278 | 0.605 | 0.732 | 0.009373 |
| Prrc2b | 4.17E-07 | -1.04633 | 0.141 | 0.307 | 0.00984 |
| Elf4 | 4.37E-07 | -0.57662 | 0.148 | 0.323 | 0.010303 |
| Ifi27l2a | 4.48E-07 | 2.02259 | 0.367 | 0.234 | 0.01056 |
| Sh3bgrl3 | 4.52E-07 | 0.984973 | 0.734 | 0.799 | 0.010662 |
| Syk | 4.55E-07 | -0.35708 | 0.312 | 0.522 | 0.010727 |
| Gm5835 | 4.65E-07 | 0.612868 | 0.535 | 0.398 | 0.010963 |
| Psma3 | 4.75E-07 | 1.820404 | 0.5 | 0.4 | 0.011216 |
| Vav1 | 4.76E-07 | -0.9054 | 0.492 | 0.649 | 0.01123 |
| Map3k1 | 4.81E-07 | -1.51088 | 0.328 | 0.465 | 0.011336 |
| Cox7c | 4.88E-07 | 1.417947 | 0.555 | 0.474 | 0.011513 |
| Ppp1r12c | 4.89E-07 | -1.5504 | 0.184 | 0.355 | 0.011531 |
| Ang4 | 4.97E-07 | 0.477581 | 0.254 | 0.123 | 0.011731 |

| | | | | | |
|---|---|---|---|---|---|
| Canx | 5.07E-07 | 0.776379 | 0.914 | 0.875 | 0.01197 |
| Rnase4 | 5.18E-07 | 0.780404 | 0.883 | 0.944 | 0.012218 |
| Gm5559 | 5.5E-07 | 0.704895 | 0.723 | 0.708 | 0.012985 |
| Prdx1 | 5.61E-07 | 1.467831 | 0.496 | 0.381 | 0.01324 |
| Gm6807 | 5.7E-07 | 2.131051 | 0.289 | 0.149 | 0.013453 |
| Cebpa | 5.8E-07 | -0.86433 | 0.508 | 0.621 | 0.013678 |
| Mafg | 6.26E-07 | -0.40222 | 0.32 | 0.522 | 0.014772 |
| Pkn1 | 6.31E-07 | -0.40922 | 0.332 | 0.559 | 0.014887 |
| Rps3a2 | 6.33E-07 | 0.589433 | 0.492 | 0.359 | 0.014934 |
| Ap1b1 | 6.65E-07 | -0.58325 | 0.258 | 0.444 | 0.015683 |
| Rpl24 | 7.65E-07 | 1.071717 | 0.656 | 0.619 | 0.018044 |
| Pum2 | 8E-07 | -0.40508 | 0.223 | 0.409 | 0.018879 |
| Capzb | 8.09E-07 | 1.354845 | 0.629 | 0.643 | 0.019092 |
| Gm10051 | 8.12E-07 | 0.821484 | 0.359 | 0.216 | 0.019154 |
| Hjurp | 8.26E-07 | -1.37693 | 0.113 | 0.27 | 0.019493 |
| Cd53 | 8.55E-07 | 0.574228 | 0.961 | 0.922 | 0.020158 |
| Kdm2b | 8.68E-07 | -0.61416 | 0.176 | 0.346 | 0.020486 |
| Tns3 | 8.83E-07 | -0.62914 | 0.297 | 0.494 | 0.020841 |
| Rps6ka1 | 8.92E-07 | -0.85078 | 0.688 | 0.799 | 0.021033 |
| Fbxl20 | 9.09E-07 | -0.70879 | 0.105 | 0.262 | 0.021432 |
| Gm9493 | 9.34E-07 | 0.40131 | 0.461 | 0.303 | 0.022035 |
| Gm8172 | 9.86E-07 | 0.923979 | 0.566 | 0.459 | 0.023269 |
| Plod3 | 1.01E-06 | -0.75913 | 0.371 | 0.561 | 0.023737 |
| Cdkn1b | 1.02E-06 | 0.269304 | 0.375 | 0.21 | 0.023993 |
| Cacna1d | 1.23E-06 | -1.08328 | 0.133 | 0.294 | 0.029119 |
| Tet3 | 1.27E-06 | -0.32757 | 0.242 | 0.424 | 0.029972 |
| Pacs2 | 1.33E-06 | -0.9636 | 0.195 | 0.372 | 0.031318 |
| Anks3 | 1.35E-06 | -0.69928 | 0.086 | 0.232 | 0.031797 |
| Gm5453 | 1.38E-06 | 0.443991 | 0.34 | 0.188 | 0.032532 |
| Mef2d | 1.45E-06 | -0.36025 | 0.102 | 0.258 | 0.03421 |
| Grn | 1.47E-06 | 0.425193 | 0.996 | 1 | 0.034611 |
| Sfpq | 1.49E-06 | -0.60955 | 0.246 | 0.414 | 0.035132 |
| Gm44861 | 1.51E-06 | -2.26532 | 0.105 | 0.245 | 0.035661 |
| Prdm2 | 1.51E-06 | -0.67119 | 0.137 | 0.301 | 0.035681 |
| P3h2 | 1.52E-06 | -2.73684 | 0.129 | 0.264 | 0.035914 |
| Map1s | 1.53E-06 | -0.81738 | 0.227 | 0.398 | 0.036086 |
| Itgb2 | 1.58E-06 | -0.82639 | 0.695 | 0.771 | 0.037239 |
| Bmp2k | 1.59E-06 | 0.403861 | 0.965 | 0.922 | 0.037547 |
| Fgd3 | 1.63E-06 | -1.19512 | 0.113 | 0.264 | 0.038354 |
| Ncor2 | 1.66E-06 | 0.284567 | 0.293 | 0.504 | 0.039223 |
| Pxn | 1.78E-06 | -0.87319 | 0.141 | 0.294 | 0.041872 |
| Hsp90b1 | 2.04E-06 | 0.727176 | 0.883 | 0.913 | 0.04814 |
| Fam193a | 2.08E-06 | -0.54488 | 0.133 | 0.292 | 0.049025 |
| Gm16192 | 2.1E-06 | -0.66567 | 0.109 | 0.26 | 0.049568 |
| Gm15266 | 2.11E-06 | 0.450153 | 0.93 | 0.92 | 0.049692 |

Supplementry Table 8.  PB-Old DEGs in Cortex

| Gene | p_val | avg_log2FC | pct.1 | pct.2 | p_val_adj |
|---|---|---|---|---|---|
| Tmsb4x | 4.61E-50 | 0.963258 | 1 | 1 | 1.09E-45 |
| Csf1r | 6.82E-49 | -0.91751 | 1 | 1 | 1.61E-44 |
| Ct010467.1 | 5.24E-43 | -0.41587 | 1 | 1 | 1.24E-38 |
| Malat1 | 7.03E-39 | 0.641393 | 0.99 | 1 | 1.66E-34 |
| Rps27 | 1.36E-38 | 1.098681 | 0.979 | 0.96 | 3.21E-34 |
| Cst3 | 7.12E-37 | 0.602975 | 1 | 1 | 1.68E-32 |
| Gm10925 | 8.43E-37 | 1.487322 | 0.864 | 0.76 | 1.99E-32 |
| Gm28661 | 1.17E-33 | 1.167965 | 0.807 | 0.623 | 2.76E-29 |
| Rps29 | 7.96E-33 | 0.858268 | 0.996 | 0.993 | 1.88E-28 |
| mt-Co2 | 2.42E-32 | 1.440806 | 0.87 | 0.785 | 5.71E-28 |
| Arglu1 | 1.61E-29 | 1.547104 | 0.918 | 0.791 | 3.81E-25 |
| mt-Atp6 | 1.42E-28 | 1.083309 | 0.872 | 0.807 | 3.36E-24 |
| Gm28437 | 1.37E-27 | 1.245446 | 0.841 | 0.76 | 3.22E-23 |
| Lars2 | 5.49E-26 | -0.73172 | 1 | 1 | 1.3E-21 |
| mt-Co3 | 4.73E-25 | 1.318383 | 0.91 | 0.908 | 1.11E-20 |
| Gm15564 | 3.65E-24 | -0.87223 | 0.678 | 0.949 | 8.61E-20 |
| Zfp691 | 1.05E-23 | 0.895268 | 0.774 | 0.554 | 2.47E-19 |
| Gm15266 | 2.71E-23 | 0.669287 | 0.875 | 0.718 | 6.39E-19 |
| Tgfbr2 | 8.18E-23 | -0.84452 | 0.839 | 0.96 | 1.93E-18 |
| Zscan26 | 8.6E-23 | 2.283122 | 0.741 | 0.601 | 2.03E-18 |
| Adap2os | 1.74E-22 | 1.669568 | 0.948 | 0.908 | 4.11E-18 |
| Serinc3 | 2.11E-22 | -0.67025 | 1 | 0.996 | 4.97E-18 |
| Rps28 | 3.07E-22 | 0.943665 | 0.95 | 0.922 | 7.24E-18 |
| Cmtm6 | 3.81E-22 | -1.28447 | 0.858 | 0.913 | 9E-18 |
| Sirpa | 9.4E-22 | -0.73032 | 0.971 | 0.989 | 2.22E-17 |
| C1qb | 9.51E-21 | 0.464025 | 1 | 1 | 2.24E-16 |
| Dpysl2 | 1.33E-19 | 1.629542 | 0.889 | 0.854 | 3.15E-15 |
| Gm4609 | 2.15E-19 | -0.63614 | 0.324 | 0.576 | 5.07E-15 |
| Sparc | 6.74E-19 | 0.598321 | 1 | 1 | 1.59E-14 |
| mt-Co1 | 8.27E-19 | 1.04605 | 0.994 | 0.989 | 1.95E-14 |
| Cd164 | 1.64E-18 | -0.6898 | 0.973 | 0.984 | 3.87E-14 |
| Ly86 | 1.86E-18 | 0.741059 | 0.996 | 0.987 | 4.38E-14 |
| Skil | 3.39E-18 | 1.480595 | 0.793 | 0.673 | 8.01E-14 |
| Tm9sf2 | 6.12E-18 | -1.15165 | 0.759 | 0.83 | 1.44E-13 |
| Cx3cr1 | 6.58E-18 | -0.50805 | 1 | 1 | 1.55E-13 |
| Calm2 | 7.12E-18 | 0.923725 | 0.983 | 0.958 | 1.68E-13 |
| Rps11 | 7.34E-18 | 0.85521 | 0.941 | 0.912 | 1.73E-13 |
| Rpl13 | 1.08E-17 | 0.753338 | 0.967 | 0.946 | 2.54E-13 |
| Ppia | 1.49E-17 | 0.736521 | 0.979 | 0.973 | 3.51E-13 |
| Slc7a8 | 2.68E-17 | -0.94816 | 0.738 | 0.87 | 6.32E-13 |
| Fau | 4.99E-17 | 0.773764 | 0.952 | 0.933 | 1.18E-12 |
| Rps3a1 | 5.51E-17 | 1.038028 | 0.883 | 0.839 | 1.3E-12 |
| mt-Cytb | 6.99E-17 | 1.047845 | 0.983 | 0.966 | 1.65E-12 |
| Adgrg1 | 2.31E-16 | -0.75818 | 0.912 | 0.949 | 5.45E-12 |

| | | | | | |
|---|---|---|---|---|---|
| Sun2 | 2.76E-16 | -1.59971 | 0.623 | 0.755 | 6.52E-12 |
| 1700017B0 | 4.23E-16 | -0.96927 | 0.722 | 0.856 | 9.99E-12 |
| Rnaset2a | 5.05E-16 | 0.665033 | 0.966 | 0.939 | 1.19E-11 |
| Slc39a1 | 5.28E-16 | 0.921241 | 0.617 | 0.442 | 1.25E-11 |
| Man2b1 | 1.58E-15 | -0.77554 | 0.952 | 0.957 | 3.74E-11 |
| B2m | 1.8E-15 | 0.59978 | 1 | 1 | 4.24E-11 |
| Itm2b | 3.23E-15 | 0.521775 | 1 | 1 | 7.62E-11 |
| Soga1 | 3.66E-15 | -1.01507 | 0.602 | 0.765 | 8.63E-11 |
| Sall1 | 4.74E-15 | -0.6872 | 0.879 | 0.966 | 1.12E-10 |
| Actr2 | 5.29E-14 | 1.061076 | 0.952 | 0.901 | 1.25E-09 |
| Aif1 | 5.93E-14 | 0.806483 | 0.925 | 0.942 | 1.4E-09 |
| Gns | 1.03E-13 | -0.68555 | 0.944 | 0.975 | 2.44E-09 |
| Bhlhe41 | 1.19E-13 | 1.377255 | 0.787 | 0.711 | 2.81E-09 |
| Rps25 | 1.23E-13 | 1.152679 | 0.86 | 0.843 | 2.9E-09 |
| Itgb5 | 2.04E-13 | -0.47183 | 0.99 | 1 | 4.8E-09 |
| Rpl37a | 2.15E-13 | 0.660468 | 0.962 | 0.953 | 5.07E-09 |
| Rpl18a | 2.21E-13 | 0.602754 | 0.996 | 0.995 | 5.22E-09 |
| Rnaset2b | 2.27E-13 | 0.616326 | 0.967 | 0.964 | 5.35E-09 |
| Fcer1g | 2.86E-13 | 0.494923 | 0.994 | 0.995 | 6.75E-09 |
| Rhob | 2.96E-13 | -0.57144 | 0.948 | 0.964 | 6.99E-09 |
| mt-Nd1 | 3.09E-13 | 0.893654 | 0.962 | 0.948 | 7.3E-09 |
| Rgs10 | 4.31E-13 | 0.632297 | 0.979 | 0.984 | 1.02E-08 |
| Ncoa3 | 4.7E-13 | 1.176552 | 0.81 | 0.722 | 1.11E-08 |
| Gm5148 | 7.19E-13 | -0.82423 | 0.588 | 0.715 | 1.69E-08 |
| Ubb | 9.78E-13 | 0.753764 | 0.989 | 0.978 | 2.31E-08 |
| Rplp1 | 1E-12 | 0.675077 | 0.985 | 0.978 | 2.37E-08 |
| Tyrobp | 1.05E-12 | 0.444063 | 0.996 | 0.996 | 2.49E-08 |
| Srsf2 | 1.52E-12 | 1.374816 | 0.862 | 0.821 | 3.58E-08 |
| Bmp2k | 1.62E-12 | 0.681839 | 0.96 | 0.917 | 3.83E-08 |
| Rpl41 | 2.18E-12 | 0.629267 | 0.962 | 0.949 | 5.14E-08 |
| Rps9 | 2.21E-12 | 0.717288 | 0.964 | 0.964 | 5.21E-08 |
| Rpl37 | 2.63E-12 | 0.855851 | 0.927 | 0.913 | 6.2E-08 |
| Rpl17 | 3.22E-12 | 1.031731 | 0.826 | 0.8 | 7.59E-08 |
| Gm25911 | 3.99E-12 | -0.2838 | 0.515 | 0.736 | 9.41E-08 |
| Rpl30 | 4.53E-12 | 0.932731 | 0.887 | 0.89 | 1.07E-07 |
| Gm10076 | 5.01E-12 | 0.641985 | 0.902 | 0.901 | 1.18E-07 |
| Notch2 | 5.05E-12 | 0.264726 | 0.757 | 0.617 | 1.19E-07 |
| Rps19 | 6.71E-12 | 0.790362 | 0.897 | 0.875 | 1.58E-07 |
| Gm8129 | 7.28E-12 | 0.711729 | 0.441 | 0.28 | 1.72E-07 |
| Gm26917 | 1.16E-11 | -0.39942 | 0.966 | 0.998 | 2.74E-07 |
| Ptafr | 1.44E-11 | 1.42956 | 0.818 | 0.765 | 3.39E-07 |
| Gm29216 | 1.52E-11 | 0.449282 | 0.864 | 0.821 | 3.58E-07 |
| Gm17494 | 1.53E-11 | 0.738367 | 0.343 | 0.181 | 3.6E-07 |
| Rbm26 | 1.66E-11 | 1.595343 | 0.728 | 0.675 | 3.92E-07 |
| Wtap | 2.28E-11 | 0.537373 | 0.584 | 0.379 | 5.38E-07 |
| Oaz1 | 2.56E-11 | 0.881935 | 0.803 | 0.767 | 6.04E-07 |
| Kdelr2 | 2.58E-11 | 0.524019 | 0.67 | 0.486 | 6.08E-07 |

| | | | | | |
|---|---|---|---|---|---|
| Rplp2 | 2.68E-11 | 0.693092 | 0.944 | 0.928 | 6.32E-07 |
| Rpl35 | 2.82E-11 | 0.902376 | 0.874 | 0.843 | 6.66E-07 |
| Rpl38 | 2.98E-11 | 0.97409 | 0.709 | 0.635 | 7.02E-07 |
| Atp2a2 | 4.22E-11 | -0.55846 | 0.276 | 0.466 | 9.96E-07 |
| Cd9 | 5.8E-11 | 0.674338 | 0.962 | 0.93 | 1.37E-06 |
| Pfn1 | 5.93E-11 | 0.599741 | 0.973 | 0.964 | 1.4E-06 |
| Rps18 | 6.51E-11 | 0.758879 | 0.914 | 0.879 | 1.54E-06 |
| Entpd1 | 6.87E-11 | -0.39097 | 0.985 | 0.98 | 1.62E-06 |
| Cbx3 | 7.78E-11 | 0.745838 | 0.515 | 0.352 | 1.84E-06 |
| Alox5ap | 8.82E-11 | 0.90203 | 0.789 | 0.745 | 2.08E-06 |
| Itpripl1 | 8.96E-11 | 1.841716 | 0.761 | 0.736 | 2.11E-06 |
| Gm19810 | 9.18E-11 | -0.42852 | 0.178 | 0.339 | 2.16E-06 |
| Gm24276 | 9.91E-11 | 0.352704 | 0.322 | 0.157 | 2.34E-06 |
| Sowahc | 1.04E-10 | -1.28159 | 0.372 | 0.527 | 2.45E-06 |
| Gm17066 | 1.08E-10 | 1.4933 | 0.588 | 0.518 | 2.55E-06 |
| Tgfa | 1.17E-10 | -0.31545 | 0.255 | 0.457 | 2.76E-06 |
| Myadm | 1.28E-10 | -0.99597 | 0.148 | 0.301 | 3.02E-06 |
| Rsrp1 | 1.38E-10 | 0.714943 | 0.941 | 0.912 | 3.25E-06 |
| Gm10123 | 1.67E-10 | 0.487158 | 0.944 | 0.949 | 3.93E-06 |
| Cap1 | 1.82E-10 | -0.78938 | 0.439 | 0.59 | 4.29E-06 |
| Cdk5r1 | 1.91E-10 | -0.91789 | 0.282 | 0.469 | 4.51E-06 |
| Tubb5 | 1.95E-10 | 0.830264 | 0.891 | 0.816 | 4.59E-06 |
| Gm13340 | 2.23E-10 | 0.466122 | 0.736 | 0.625 | 5.26E-06 |
| Rps24 | 2.25E-10 | 0.864166 | 0.868 | 0.866 | 5.31E-06 |
| Ftl1-ps1 | 2.43E-10 | 0.560339 | 0.9 | 0.917 | 5.74E-06 |
| Rpl13a | 2.8E-10 | 0.695778 | 0.862 | 0.847 | 6.61E-06 |
| Gm4202 | 2.84E-10 | -0.60731 | 0.596 | 0.727 | 6.71E-06 |
| Rps3a3 | 4.19E-10 | 0.50984 | 0.462 | 0.303 | 9.88E-06 |
| Gna12 | 8.34E-10 | -0.8497 | 0.594 | 0.729 | 1.97E-05 |
| Pdcd4 | 9.79E-10 | 0.776536 | 0.485 | 0.347 | 2.31E-05 |
| Cebpa | 1.17E-09 | -1.05601 | 0.6 | 0.682 | 2.76E-05 |
| Sept2 | 1.18E-09 | 1.425084 | 0.774 | 0.691 | 2.79E-05 |
| Man1c1 | 1.2E-09 | -1.14331 | 0.536 | 0.661 | 2.83E-05 |
| Cd81 | 1.37E-09 | 0.507524 | 0.998 | 1 | 3.24E-05 |
| C5ar2 | 1.38E-09 | 1.268844 | 0.699 | 0.605 | 3.25E-05 |
| Sec61a1 | 1.45E-09 | -0.88388 | 0.67 | 0.751 | 3.41E-05 |
| Ftl1 | 1.52E-09 | 0.485796 | 0.996 | 0.998 | 3.58E-05 |
| Rpl11 | 1.59E-09 | 0.896316 | 0.761 | 0.74 | 3.75E-05 |
| Rps20 | 1.66E-09 | 0.81501 | 0.858 | 0.872 | 3.92E-05 |
| Eef2 | 1.78E-09 | -0.76682 | 0.824 | 0.883 | 4.2E-05 |
| Abcg1 | 2.3E-09 | 1.599892 | 0.715 | 0.655 | 5.42E-05 |
| Rps14 | 2.42E-09 | 0.611133 | 0.941 | 0.939 | 5.71E-05 |
| Fmnl1 | 2.64E-09 | -0.37254 | 0.366 | 0.551 | 6.23E-05 |
| C1qc | 3.17E-09 | 0.414332 | 1 | 1 | 7.48E-05 |
| Gm2225 | 3.41E-09 | 0.356069 | 0.362 | 0.209 | 8.04E-05 |
| Comt | 3.54E-09 | 0.839971 | 0.776 | 0.699 | 8.34E-05 |
| Fth1 | 4.34E-09 | 0.474817 | 0.998 | 1 | 0.000102 |

| | | | | | |
|---|---|---|---|---|---|
| Eef1a1 | 5.05E-09 | 0.665915 | 0.996 | 0.996 | 0.000119 |
| Rps27a | 8.53E-09 | 0.82988 | 0.824 | 0.83 | 0.000201 |
| Dusp7 | 8.53E-09 | -0.50762 | 0.678 | 0.805 | 0.000201 |
| Rapgef1 | 8.9E-09 | -0.77383 | 0.397 | 0.563 | 0.00021 |
| Rpl10 | 1E-08 | 0.701848 | 0.939 | 0.937 | 0.000236 |
| Gm5905 | 1.18E-08 | -0.4571 | 0.801 | 0.845 | 0.000278 |
| Ivns1abp | 1.2E-08 | -0.51729 | 0.902 | 0.939 | 0.000283 |
| Gm14303 | 1.39E-08 | 0.4955 | 0.885 | 0.879 | 0.000327 |
| Rpl34 | 1.42E-08 | 0.915085 | 0.82 | 0.819 | 0.000335 |
| Ptgs1 | 1.63E-08 | 0.623033 | 0.979 | 0.958 | 0.000383 |
| Slc40a1 | 1.85E-08 | -1.03801 | 0.531 | 0.644 | 0.000437 |
| Gm10269 | 2.07E-08 | -0.64292 | 0.46 | 0.576 | 0.000488 |
| Rpl36 | 2.36E-08 | 0.832808 | 0.818 | 0.829 | 0.000558 |
| Ywhah | 2.44E-08 | 0.604367 | 0.937 | 0.921 | 0.000575 |
| Rps4x | 2.65E-08 | 0.937603 | 0.862 | 0.859 | 0.000624 |
| B4galt1 | 3.11E-08 | 0.702278 | 0.889 | 0.865 | 0.000734 |
| Ctss | 3.24E-08 | 0.357702 | 1 | 1 | 0.000765 |
| P4hb | 3.5E-08 | -0.45669 | 0.828 | 0.886 | 0.000826 |
| Gm44705 | 3.79E-08 | 0.348204 | 0.379 | 0.235 | 0.000894 |
| Tpm3 | 4.08E-08 | 0.769711 | 0.92 | 0.93 | 0.000961 |
| St13 | 4.11E-08 | 1.326751 | 0.688 | 0.605 | 0.000969 |
| Sec62 | 4.12E-08 | 1.079255 | 0.68 | 0.644 | 0.000972 |
| Rpl27a | 4.17E-08 | 0.995948 | 0.766 | 0.751 | 0.000983 |
| Ssr1 | 4.41E-08 | 1.364154 | 0.669 | 0.59 | 0.001041 |
| Abcc3 | 4.84E-08 | -0.7915 | 0.443 | 0.587 | 0.001142 |
| Rpl35a | 5.25E-08 | 0.881149 | 0.803 | 0.783 | 0.001238 |
| Rpl21 | 6.02E-08 | 0.61384 | 0.939 | 0.946 | 0.001419 |
| Set | 6.24E-08 | 1.195707 | 0.676 | 0.61 | 0.001473 |
| Rps3 | 6.56E-08 | 0.66732 | 0.923 | 0.933 | 0.001547 |
| Man2b2 | 6.57E-08 | -1.35243 | 0.441 | 0.547 | 0.001549 |
| Nlrp3 | 6.86E-08 | -1.10793 | 0.107 | 0.229 | 0.001618 |
| Lrrc8a | 6.99E-08 | 2.185013 | 0.51 | 0.43 | 0.001649 |
| Slc25a5 | 7.37E-08 | 0.856328 | 0.854 | 0.839 | 0.001739 |
| Nfia | 8.02E-08 | 1.142219 | 0.554 | 0.43 | 0.001891 |
| Ncoa4 | 9.22E-08 | -1.42276 | 0.213 | 0.334 | 0.002175 |
| Rpl7 | 9.88E-08 | 0.792475 | 0.826 | 0.827 | 0.002331 |
| Atp1a1 | 9.91E-08 | -1.11097 | 0.494 | 0.592 | 0.002339 |
| Tifab | 1.05E-07 | -0.60433 | 0.592 | 0.697 | 0.00247 |
| Pdgfb | 1.11E-07 | -0.69706 | 0.473 | 0.623 | 0.002609 |
| Gnb1 | 1.25E-07 | 1.025513 | 0.885 | 0.879 | 0.002944 |
| Rpl26 | 1.38E-07 | 0.851086 | 0.808 | 0.816 | 0.003266 |
| Gm1821 | 1.45E-07 | 0.420362 | 0.962 | 0.962 | 0.00342 |
| Fscn1 | 1.54E-07 | -0.71565 | 0.923 | 0.915 | 0.003636 |
| Actb | 1.63E-07 | 0.294571 | 1 | 1 | 0.003841 |
| Klhl9 | 1.77E-07 | 1.68773 | 0.525 | 0.439 | 0.004175 |
| Lrp10 | 1.79E-07 | -1.10477 | 0.443 | 0.56 | 0.004214 |
| Serf2 | 1.83E-07 | 0.671023 | 0.785 | 0.735 | 0.004307 |

| | | | | | |
|---|---|---|---|---|---|
| Rpl3 | 1.84E-07 | 0.609203 | 0.864 | 0.856 | 0.00435 |
| Sema4d | 1.93E-07 | -0.47653 | 0.546 | 0.708 | 0.004548 |
| Gm11518 | 2.14E-07 | 0.99585 | 0.623 | 0.596 | 0.00504 |
| Rps7 | 2.27E-07 | 0.791799 | 0.801 | 0.796 | 0.005363 |
| Akna | 2.29E-07 | -0.80119 | 0.529 | 0.679 | 0.005407 |
| Rpl23a | 2.72E-07 | 0.624803 | 0.816 | 0.78 | 0.006413 |
| Copa | 2.79E-07 | -0.85919 | 0.343 | 0.487 | 0.006579 |
| Gm9843 | 2.88E-07 | 0.538098 | 0.912 | 0.903 | 0.006793 |
| Hs6st1 | 2.89E-07 | -0.60551 | 0.293 | 0.435 | 0.00681 |
| Rps6ka1 | 2.93E-07 | -0.50341 | 0.852 | 0.91 | 0.006906 |
| Selplg | 3.15E-07 | 0.425804 | 1 | 0.998 | 0.007433 |
| Rps3a2 | 3.24E-07 | 0.36566 | 0.492 | 0.354 | 0.007652 |
| Erf | 3.4E-07 | -1.08575 | 0.469 | 0.561 | 0.008025 |
| Plxnb2 | 3.64E-07 | -0.54476 | 0.776 | 0.888 | 0.008583 |
| Kat6a | 3.75E-07 | -0.85099 | 0.312 | 0.455 | 0.008854 |
| Il10ra | 3.78E-07 | -0.8831 | 0.801 | 0.834 | 0.008918 |
| Ptp4a2 | 4.14E-07 | 1.231031 | 0.785 | 0.744 | 0.009774 |
| mt-Nd2 | 4.53E-07 | 0.946486 | 0.916 | 0.922 | 0.010691 |
| Sft2d1 | 4.58E-07 | 0.607839 | 0.908 | 0.901 | 0.010795 |
| Kctd12 | 4.64E-07 | -0.45007 | 0.795 | 0.879 | 0.01094 |
| Anxa3 | 4.78E-07 | 0.721997 | 0.824 | 0.758 | 0.011281 |
| Lman2 | 5.28E-07 | -0.41224 | 0.412 | 0.567 | 0.012466 |
| Marf1 | 5.85E-07 | 1.641495 | 0.54 | 0.478 | 0.013803 |
| Lrp5 | 6.14E-07 | -0.84458 | 0.184 | 0.316 | 0.014481 |
| Gng10 | 6.15E-07 | 0.614542 | 0.805 | 0.765 | 0.014513 |
| Gm12918 | 6.66E-07 | 0.404316 | 0.609 | 0.505 | 0.015704 |
| Ctsh | 7.25E-07 | 0.610709 | 0.952 | 0.966 | 0.017091 |
| Adam15 | 7.42E-07 | -0.70426 | 0.659 | 0.76 | 0.017494 |
| Sidt2 | 7.64E-07 | -1.13265 | 0.333 | 0.453 | 0.018018 |
| Rgmb | 8.02E-07 | -0.71613 | 0.477 | 0.601 | 0.018929 |
| Gpr165 | 8.17E-07 | -0.77241 | 0.331 | 0.466 | 0.019283 |
| Gm6900 | 8.28E-07 | 0.355023 | 0.437 | 0.319 | 0.019526 |
| Maz | 8.66E-07 | -0.68825 | 0.157 | 0.285 | 0.020436 |
| Abca1 | 8.69E-07 | -0.51841 | 0.594 | 0.733 | 0.02051 |
| Tmem59 | 8.73E-07 | 0.562952 | 0.893 | 0.87 | 0.020601 |
| Sod1 | 1.05E-06 | 1.185317 | 0.649 | 0.581 | 0.024731 |
| Tnfrsf21 | 1.05E-06 | -1.31797 | 0.527 | 0.608 | 0.024763 |
| Rpl6 | 1.06E-06 | 0.866428 | 0.81 | 0.805 | 0.024966 |
| Tmed7 | 1.15E-06 | -0.79871 | 0.446 | 0.552 | 0.027017 |
| Gab1 | 1.16E-06 | -0.95544 | 0.121 | 0.235 | 0.027453 |
| Mtpn | 1.33E-06 | -1.03113 | 0.414 | 0.523 | 0.031435 |
| Sat1 | 1.42E-06 | 0.863203 | 0.881 | 0.874 | 0.033582 |
| Rps27rt | 1.45E-06 | 0.584548 | 0.772 | 0.749 | 0.034233 |
| Adam10 | 1.69E-06 | 1.041617 | 0.814 | 0.764 | 0.039874 |
| Tns1 | 1.7E-06 | -0.71254 | 0.205 | 0.339 | 0.040065 |
| Gm10134 | 1.71E-06 | 1.913979 | 0.391 | 0.294 | 0.040241 |
| Ftl2-ps | 1.85E-06 | 0.426764 | 0.931 | 0.931 | 0.04361 |

| | | | | | |
|---|---|---|---|---|---|
| Msn | 1.86E-06 | -0.2735 | 0.617 | 0.733 | 0.043948 |

Supplementry Table 8.  PB-Old DEGs in Hippocampus

| Gene | p_val | avg_log2FC | pct.1 | pct.2 | p_val_adj |
|---|---|---|---|---|---|
| Gm23935 | 1.1E-132 | -1.72078 | 0.996 | 1 | 2.6E-128 |
| Gm15564 | 4.3E-118 | -2.34287 | 0.696 | 0.985 | 1E-113 |
| Lars2 | 1.6E-116 | -1.80574 | 0.996 | 1 | 3.7E-112 |
| Gm24270 | 6E-111 | -1.60833 | 0.875 | 0.998 | 1.4E-106 |
| Gm24187 | 6E-100 | -1.38678 | 0.899 | 1 | 1.41E-95 |
| Gm24245 | 1.13E-74 | -1.5339 | 0.485 | 0.903 | 2.67E-70 |
| Gm26917 | 5.09E-69 | -1.75323 | 0.924 | 1 | 1.2E-64 |
| Ct010467.1 | 3.1E-64 | -0.48337 | 1 | 1 | 7.32E-60 |
| Rn7sk | 5.57E-55 | -1.5903 | 0.691 | 0.972 | 1.31E-50 |
| Gm25911 | 1.2E-39 | -1.02171 | 0.452 | 0.796 | 2.82E-35 |
| Gm1821 | 8.3E-24 | 0.847146 | 0.953 | 0.931 | 1.96E-19 |
| Soga1 | 8.52E-23 | -0.81993 | 0.548 | 0.843 | 2.01E-18 |
| Gm8797 | 5.14E-19 | 0.866605 | 0.826 | 0.697 | 1.21E-14 |
| Sun2 | 2.19E-18 | -1.09648 | 0.579 | 0.813 | 5.18E-14 |
| Cx3cr1 | 4.95E-17 | -0.44623 | 1 | 1 | 1.17E-12 |
| Gm8464 | 5.21E-17 | -0.38781 | 0.094 | 0.298 | 1.23E-12 |
| Atxn2l | 2.39E-16 | -0.88028 | 0.387 | 0.642 | 5.63E-12 |
| Lrp1 | 2.49E-16 | -0.54048 | 0.797 | 0.97 | 5.86E-12 |
| mt-Co3 | 2.86E-16 | 0.876345 | 0.884 | 0.779 | 6.75E-12 |
| Pou2f2 | 2.91E-16 | -0.5384 | 0.711 | 0.91 | 6.87E-12 |
| Dusp7 | 7.65E-16 | -0.718 | 0.564 | 0.809 | 1.8E-11 |
| Fscn1 | 1.14E-15 | -0.73733 | 0.83 | 0.925 | 2.7E-11 |
| Rreb1 | 7.82E-15 | -1.29387 | 0.307 | 0.536 | 1.84E-10 |
| Nfatc2 | 8.84E-15 | -1.3403 | 0.168 | 0.384 | 2.09E-10 |
| Tgfbr2 | 9.11E-15 | -0.4213 | 0.821 | 0.968 | 2.15E-10 |
| Wdr81 | 9.66E-15 | -1.01016 | 0.418 | 0.622 | 2.28E-10 |
| Rasa4 | 2.66E-14 | -1.21959 | 0.322 | 0.543 | 6.28E-10 |
| Cd81 | 2.98E-14 | 0.66403 | 0.998 | 1 | 7.03E-10 |
| Tgfb1 | 3.51E-14 | -0.80446 | 0.325 | 0.556 | 8.29E-10 |
| mt-Rnr1 | 3.8E-14 | -0.57469 | 0.857 | 0.976 | 8.97E-10 |
| mt-Co2 | 5.03E-14 | 1.055437 | 0.787 | 0.708 | 1.19E-09 |
| Inpp5d | 5.56E-14 | -0.45415 | 0.877 | 0.989 | 1.31E-09 |
| Rnaset2b | 8.05E-14 | 0.831669 | 0.949 | 0.927 | 1.9E-09 |
| Lrp5 | 2.98E-13 | -1.76026 | 0.177 | 0.367 | 7.04E-09 |
| Cdk5r1 | 3.58E-13 | -0.62048 | 0.255 | 0.489 | 8.45E-09 |
| Gm10123 | 4.89E-13 | 0.620616 | 0.929 | 0.888 | 1.15E-08 |
| Gm5148 | 1.89E-12 | -0.72995 | 0.463 | 0.635 | 4.46E-08 |
| mt-Nd6 | 2.2E-12 | -0.57863 | 0.696 | 0.839 | 5.19E-08 |
| Frmd4a | 3.33E-12 | -0.35211 | 0.694 | 0.912 | 7.86E-08 |
| Sowahc | 4.33E-12 | -1.16937 | 0.356 | 0.552 | 1.02E-07 |
| Fchsd2 | 4.65E-12 | -0.42652 | 0.394 | 0.629 | 1.1E-07 |
| Trappc10 | 5.34E-12 | -1.18496 | 0.127 | 0.303 | 1.26E-07 |
| Hps4 | 6.63E-12 | -0.41124 | 0.552 | 0.773 | 1.56E-07 |
| Slc37a2 | 6.78E-12 | -0.80996 | 0.396 | 0.599 | 1.6E-07 |

| | | | | | |
|---|---|---|---|---|---|
| Rere | 1.36E-11 | -0.63402 | 0.351 | 0.564 | 3.21E-07 |
| Pmepa1 | 1.94E-11 | -0.59004 | 0.864 | 0.957 | 4.58E-07 |
| Kcnk6 | 2.88E-11 | -0.40631 | 0.542 | 0.764 | 6.8E-07 |
| Nfatc1 | 2.93E-11 | -0.48134 | 0.438 | 0.644 | 6.91E-07 |
| Ptpa | 3.97E-11 | -0.39674 | 0.297 | 0.511 | 9.37E-07 |
| Zmiz1 | 5.74E-11 | -0.34475 | 0.626 | 0.815 | 1.36E-06 |
| Adap2os | 6.37E-11 | 1.501892 | 0.928 | 0.929 | 1.5E-06 |
| Gcn1l1 | 6.58E-11 | -0.26556 | 0.445 | 0.674 | 1.55E-06 |
| Arhgap39 | 7.19E-11 | -0.73317 | 0.409 | 0.599 | 1.7E-06 |
| Fus | 1.02E-10 | -0.35649 | 0.454 | 0.672 | 2.42E-06 |
| Ogt | 1.12E-10 | -0.43256 | 0.562 | 0.749 | 2.65E-06 |
| Gm26857 | 1.19E-10 | -1.27808 | 0.123 | 0.285 | 2.81E-06 |
| Snrnp70 | 1.41E-10 | -0.3632 | 0.506 | 0.702 | 3.33E-06 |
| Pdgfb | 1.99E-10 | -0.36061 | 0.423 | 0.65 | 4.68E-06 |
| Bmp2k | 2.62E-10 | 1.164558 | 0.966 | 0.946 | 6.19E-06 |
| mt-Rnr2 | 2.98E-10 | -0.39233 | 0.984 | 0.998 | 7.04E-06 |
| Dpp9 | 2.99E-10 | -1.26783 | 0.081 | 0.219 | 7.06E-06 |
| Zscan26 | 3.58E-10 | 2.171788 | 0.714 | 0.655 | 8.45E-06 |
| Tab2 | 4.04E-10 | -0.37404 | 0.465 | 0.693 | 9.52E-06 |
| Ppp1r9b | 4.11E-10 | -0.76369 | 0.277 | 0.466 | 9.69E-06 |
| Mertk | 5.64E-10 | -0.35626 | 0.857 | 0.968 | 1.33E-05 |
| Kansl1 | 6.69E-10 | -0.73943 | 0.358 | 0.558 | 1.58E-05 |
| Tfe3 | 6.71E-10 | -0.51145 | 0.461 | 0.665 | 1.58E-05 |
| Abca1 | 6.98E-10 | -0.51487 | 0.537 | 0.727 | 1.65E-05 |
| mt-Cytb | 7.49E-10 | 0.883332 | 0.946 | 0.959 | 1.77E-05 |
| Ivns1abp | 8.39E-10 | -0.29095 | 0.799 | 0.961 | 1.98E-05 |
| Arglu1 | 8.83E-10 | 1.413991 | 0.857 | 0.848 | 2.08E-05 |
| Zfp691 | 9.28E-10 | 0.789742 | 0.705 | 0.556 | 2.19E-05 |
| Gm42418 | 1.15E-09 | -0.48326 | 0.861 | 0.957 | 2.71E-05 |
| Gm19810 | 1.17E-09 | -0.30379 | 0.114 | 0.262 | 2.76E-05 |
| Sall3 | 1.34E-09 | -0.46951 | 0.577 | 0.766 | 3.15E-05 |
| Map1s | 1.63E-09 | -0.64205 | 0.273 | 0.448 | 3.84E-05 |
| Prex1 | 1.63E-09 | -0.41734 | 0.528 | 0.719 | 3.86E-05 |
| Ints6l | 1.91E-09 | -0.5318 | 0.371 | 0.571 | 4.51E-05 |
| Calm2 | 2.4E-09 | 0.876983 | 0.967 | 0.97 | 5.66E-05 |
| Abca3 | 2.58E-09 | -1.03778 | 0.112 | 0.253 | 6.08E-05 |
| Rps21 | 2.65E-09 | -0.33049 | 0.682 | 0.85 | 6.24E-05 |
| Lrch1 | 2.68E-09 | -0.60356 | 0.316 | 0.498 | 6.32E-05 |
| Fmnl3 | 3.26E-09 | -0.30176 | 0.535 | 0.715 | 7.7E-05 |
| Fkbp15 | 3.77E-09 | -0.46801 | 0.416 | 0.609 | 8.89E-05 |
| Sall1 | 3.85E-09 | -0.26064 | 0.834 | 0.964 | 9.08E-05 |
| Hist1h4d | 4.3E-09 | -1.16578 | 0.259 | 0.408 | 0.000102 |
| Stab1 | 5.31E-09 | -0.35152 | 0.801 | 0.923 | 0.000125 |
| D5Ertd579 | 5.54E-09 | -0.64108 | 0.09 | 0.221 | 0.000131 |
| Srgap2 | 5.72E-09 | -0.27027 | 0.81 | 0.94 | 0.000135 |
| Tsc22d2 | 5.84E-09 | -1.11365 | 0.248 | 0.416 | 0.000138 |
| Dusp6 | 6.96E-09 | -0.4938 | 0.669 | 0.811 | 0.000164 |

| | | | | | |
|---|---|---|---|---|---|
| Rapgef1 | 8.21E-09 | -1.00569 | 0.344 | 0.517 | 0.000194 |
| Phrf1 | 8.51E-09 | -1.1941 | 0.119 | 0.258 | 0.000201 |
| Chd7 | 8.8E-09 | -0.35076 | 0.407 | 0.601 | 0.000208 |
| Zeb2 | 9.03E-09 | -0.47788 | 0.425 | 0.616 | 0.000213 |
| Dennd4a | 9.32E-09 | -0.30879 | 0.528 | 0.723 | 0.00022 |
| Kif21b | 1.09E-08 | -0.33538 | 0.407 | 0.59 | 0.000257 |
| Prpf4b | 1.18E-08 | -0.60217 | 0.251 | 0.41 | 0.000278 |
| Lpin2 | 1.35E-08 | -0.44034 | 0.401 | 0.594 | 0.000318 |
| Ttyh3 | 1.5E-08 | -0.34444 | 0.345 | 0.528 | 0.000353 |
| Itpripl1 | 1.68E-08 | 2.128573 | 0.718 | 0.704 | 0.000396 |
| Selplg | 1.75E-08 | 0.461192 | 1 | 1 | 0.000412 |
| C1qc | 1.97E-08 | 0.552466 | 1 | 1 | 0.000464 |
| Skil | 2.06E-08 | 1.665031 | 0.754 | 0.734 | 0.000485 |
| Il17ra | 2.21E-08 | -0.50251 | 0.441 | 0.603 | 0.000521 |
| Pgd | 2.57E-08 | -0.35207 | 0.257 | 0.429 | 0.000606 |
| Epn1 | 2.62E-08 | -0.49848 | 0.356 | 0.536 | 0.000619 |
| Gm12977 | 2.66E-08 | -1.04028 | 0.141 | 0.277 | 0.000628 |
| mt-Co1 | 2.8E-08 | 0.717031 | 0.966 | 0.985 | 0.00066 |
| Gm15635 | 2.82E-08 | -1.22819 | 0.112 | 0.242 | 0.000665 |
| Akna | 3.11E-08 | -0.42267 | 0.506 | 0.678 | 0.000734 |
| Runx1 | 3.18E-08 | -0.29034 | 0.295 | 0.455 | 0.00075 |
| Gm27201 | 3.22E-08 | -0.91535 | 0.08 | 0.2 | 0.00076 |
| Actr2 | 3.3E-08 | 1.284824 | 0.901 | 0.865 | 0.000778 |
| Washc4 | 3.33E-08 | -0.28842 | 0.389 | 0.543 | 0.000786 |
| Arsb | 3.9E-08 | -0.83348 | 0.544 | 0.674 | 0.00092 |
| Abcd1 | 4.72E-08 | -0.75946 | 0.271 | 0.425 | 0.001113 |
| 4731419I0! | 4.83E-08 | -0.2642 | 0.09 | 0.21 | 0.00114 |
| Ophn1 | 4.86E-08 | -0.44134 | 0.465 | 0.646 | 0.001147 |
| Cdc37l1 | 5.4E-08 | -0.46839 | 0.152 | 0.294 | 0.001274 |
| Rin2 | 5.62E-08 | -0.42122 | 0.508 | 0.693 | 0.001326 |
| Sh2b3 | 5.73E-08 | -0.48308 | 0.45 | 0.646 | 0.001352 |
| Gm49369 | 5.75E-08 | 0.347232 | 0.83 | 0.715 | 0.001357 |
| Arhgef1 | 5.88E-08 | -0.43395 | 0.467 | 0.635 | 0.001387 |
| Wdfy2 | 6.02E-08 | -0.34137 | 0.306 | 0.474 | 0.00142 |
| Atxn2 | 6.02E-08 | -0.25947 | 0.244 | 0.41 | 0.001421 |
| Cd164 | 6.11E-08 | -0.38079 | 0.935 | 0.983 | 0.001442 |
| Cbl | 7.01E-08 | -0.30919 | 0.329 | 0.519 | 0.001653 |
| Gm38042 | 7.47E-08 | -0.6297 | 0.206 | 0.354 | 0.001763 |
| Trrap | 7.52E-08 | -0.91592 | 0.25 | 0.397 | 0.001774 |
| Gm37954 | 8.3E-08 | -0.62423 | 0.154 | 0.292 | 0.001958 |
| Slc39a1 | 8.36E-08 | 1.405293 | 0.586 | 0.509 | 0.001971 |
| Arhgap31 | 8.51E-08 | -0.47773 | 0.277 | 0.44 | 0.002006 |
| Scamp5 | 8.84E-08 | -0.35682 | 0.586 | 0.73 | 0.002085 |
| Gm34589 | 9.49E-08 | -0.95298 | 0.233 | 0.388 | 0.002239 |
| Nrip1 | 1.06E-07 | -0.28586 | 0.627 | 0.779 | 0.002493 |
| Apba1 | 1.09E-07 | -1.13735 | 0.159 | 0.296 | 0.00257 |
| Lrrk1 | 1.13E-07 | -0.43713 | 0.277 | 0.442 | 0.002667 |

| | | | | | |
|---|---|---|---|---|---|
| Dock8 | 1.24E-07 | -0.30119 | 0.501 | 0.682 | 0.002926 |
| Clstn1 | 1.35E-07 | -0.28154 | 0.441 | 0.624 | 0.003185 |
| Mdm4 | 1.48E-07 | -0.40094 | 0.363 | 0.528 | 0.003502 |
| Hexb | 1.6E-07 | 0.360195 | 1 | 1 | 0.003774 |
| Aph1a | 1.68E-07 | -0.51711 | 0.398 | 0.558 | 0.003958 |
| Rnf44 | 1.7E-07 | -0.51446 | 0.365 | 0.536 | 0.004018 |
| Npepps | 1.75E-07 | -0.56355 | 0.244 | 0.408 | 0.004135 |
| Klf3 | 1.88E-07 | -0.37167 | 0.566 | 0.73 | 0.004438 |
| Senp7 | 1.92E-07 | -0.66776 | 0.119 | 0.249 | 0.004537 |
| Plxna4 | 2.33E-07 | -0.38946 | 0.213 | 0.363 | 0.005505 |
| Wnk1 | 2.38E-07 | -0.32041 | 0.756 | 0.91 | 0.005618 |
| Themis2 | 2.52E-07 | -0.35525 | 0.351 | 0.511 | 0.005946 |
| Med12l | 2.56E-07 | -0.56244 | 0.479 | 0.639 | 0.006042 |
| Rps15-ps2 | 2.62E-07 | -0.41336 | 0.242 | 0.382 | 0.006174 |
| Ppp6r1 | 2.72E-07 | -0.51989 | 0.255 | 0.414 | 0.006416 |
| Abcg1 | 2.76E-07 | 1.649241 | 0.694 | 0.644 | 0.006508 |
| Hk3 | 2.83E-07 | -0.81835 | 0.213 | 0.363 | 0.00668 |
| Cant1 | 2.85E-07 | -0.86089 | 0.32 | 0.461 | 0.006724 |
| Git2 | 2.91E-07 | -0.26273 | 0.676 | 0.863 | 0.006857 |
| Gab1 | 2.99E-07 | -0.69698 | 0.101 | 0.219 | 0.007057 |
| Pkn1 | 3E-07 | -0.52301 | 0.427 | 0.588 | 0.007075 |
| Tcirg1 | 3.12E-07 | -0.38465 | 0.456 | 0.622 | 0.007362 |
| Gm37033 | 3.2E-07 | -0.75363 | 0.09 | 0.204 | 0.007538 |
| Gramd1a | 3.7E-07 | -0.61608 | 0.333 | 0.472 | 0.008738 |
| Fmnl1 | 3.97E-07 | -0.51738 | 0.389 | 0.562 | 0.009356 |
| Stk10 | 4.1E-07 | -0.26734 | 0.277 | 0.438 | 0.009663 |
| Hcfc1 | 4.39E-07 | -0.67906 | 0.19 | 0.33 | 0.010364 |
| Trpc4ap | 4.54E-07 | -0.60938 | 0.42 | 0.579 | 0.010709 |
| Nisch | 4.79E-07 | -0.26961 | 0.55 | 0.719 | 0.011295 |
| Myh9 | 5.03E-07 | -0.487 | 0.429 | 0.562 | 0.011875 |
| Cyfip1 | 5.19E-07 | -0.30626 | 0.81 | 0.951 | 0.01224 |
| Slc8a1 | 5.7E-07 | -1.15438 | 0.145 | 0.264 | 0.013455 |
| Nfic | 5.72E-07 | -0.28715 | 0.467 | 0.624 | 0.013499 |
| Vav2 | 5.76E-07 | -0.56138 | 0.217 | 0.358 | 0.013577 |
| Rlim | 5.86E-07 | -0.80076 | 0.309 | 0.448 | 0.013826 |
| Mitf | 5.87E-07 | -0.4137 | 0.168 | 0.298 | 0.013858 |
| Ctsz | 5.92E-07 | 0.511128 | 0.987 | 0.991 | 0.013973 |
| Tm9sf2 | 6E-07 | -0.3733 | 0.75 | 0.876 | 0.014147 |
| Rps6ka1 | 6.23E-07 | -0.31916 | 0.756 | 0.916 | 0.014695 |
| Sp1 | 6.41E-07 | -0.50191 | 0.157 | 0.296 | 0.015122 |
| Rnaset2a | 7.09E-07 | 0.515866 | 0.955 | 0.961 | 0.016735 |
| Rapgef5 | 7.22E-07 | -0.48921 | 0.356 | 0.526 | 0.017029 |
| C130089K0 | 7.65E-07 | -0.49488 | 0.266 | 0.412 | 0.018054 |
| Srsf2 | 8.14E-07 | 1.454829 | 0.835 | 0.843 | 0.019201 |
| Pnpla7 | 8.42E-07 | -0.2744 | 0.521 | 0.691 | 0.019851 |
| Trip11 | 8.5E-07 | -0.45001 | 0.132 | 0.253 | 0.020057 |
| Zc3h4 | 8.85E-07 | -0.86899 | 0.157 | 0.281 | 0.02087 |

| | | | | | |
|---|---|---|---|---|---|
| Eef2 | 8.9E-07 | -0.46164 | 0.749 | 0.867 | 0.020996 |
| Bach2 | 9.03E-07 | -0.96274 | 0.161 | 0.283 | 0.021309 |
| Mdm2 | 9.43E-07 | -0.50404 | 0.168 | 0.298 | 0.022243 |
| Rnf216 | 9.5E-07 | -0.30157 | 0.291 | 0.444 | 0.02242 |
| Tnks2 | 9.62E-07 | -0.43915 | 0.306 | 0.446 | 0.022688 |
| Sympk | 9.8E-07 | -1.73312 | 0.107 | 0.215 | 0.023127 |
| Sipa1l2 | 9.99E-07 | -0.47622 | 0.282 | 0.438 | 0.023561 |
| Klhl9 | 1E-06 | 1.617749 | 0.501 | 0.395 | 0.023685 |
| Arhgef2 | 1.14E-06 | -0.43857 | 0.356 | 0.511 | 0.026987 |
| Mroh1 | 1.18E-06 | -1.25121 | 0.123 | 0.238 | 0.027724 |
| Nhlrc3 | 1.2E-06 | -0.47362 | 0.141 | 0.264 | 0.028205 |
| Evl | 1.21E-06 | -0.48441 | 0.535 | 0.676 | 0.02844 |
| Myadm | 1.24E-06 | -0.52429 | 0.134 | 0.251 | 0.029153 |
| Erf | 1.27E-06 | -0.88597 | 0.438 | 0.575 | 0.030043 |
| Gm15501 | 1.32E-06 | -0.41623 | 0.443 | 0.584 | 0.031085 |
| Mgrn1 | 1.34E-06 | -0.42575 | 0.325 | 0.483 | 0.031705 |
| Bhlhe41 | 1.39E-06 | 1.545427 | 0.807 | 0.783 | 0.032696 |
| Rpl34-ps1 | 1.44E-06 | -0.4239 | 0.445 | 0.592 | 0.033988 |
| Notch1 | 1.48E-06 | -0.72788 | 0.329 | 0.476 | 0.034796 |
| Abl2 | 1.49E-06 | -0.90895 | 0.146 | 0.27 | 0.035165 |
| Ube3c | 1.55E-06 | -0.71398 | 0.163 | 0.29 | 0.036535 |
| Tle3 | 1.6E-06 | -0.52988 | 0.32 | 0.474 | 0.037803 |
| Kmt2a | 1.61E-06 | -0.72426 | 0.32 | 0.466 | 0.037964 |
| Arhgap17 | 1.72E-06 | -0.44292 | 0.456 | 0.603 | 0.040576 |
| Etnk1 | 1.74E-06 | -0.80106 | 0.367 | 0.5 | 0.040965 |
| Yy1 | 1.92E-06 | -0.32725 | 0.222 | 0.365 | 0.045249 |
| Cpsf7 | 1.92E-06 | 0.30497 | 0.116 | 0.232 | 0.045341 |
| Pcnx3 | 1.95E-06 | -0.41873 | 0.3 | 0.446 | 0.046108 |
| Gtf2i | 1.96E-06 | -0.33 | 0.479 | 0.642 | 0.046316 |
| Arap1 | 1.98E-06 | -0.27993 | 0.335 | 0.498 | 0.046777 |
| Ptafr | 1.99E-06 | 1.553808 | 0.769 | 0.773 | 0.046842 |
| Map4 | 1.99E-06 | -0.47549 | 0.28 | 0.418 | 0.046995 |
| Map4k4 | 2.01E-06 | -0.66046 | 0.483 | 0.62 | 0.047299 |
| Gm37494 | 2.01E-06 | -0.73743 | 0.132 | 0.247 | 0.047413 |
| Polr2a | 2.1E-06 | -0.59404 | 0.396 | 0.545 | 0.049577 |
| Cep68 | 2.1E-06 | -0.41603 | 0.306 | 0.455 | 0.049604 |
| Atf7 | 2.12E-06 | -0.58896 | 0.228 | 0.361 | 0.049965 |

Supplementry Table 8.  PB-Old DEGs in Striatum

| Gene | p_val | avg_log2FC | pct.1 | pct.2 | p_val_adj |
|---|---|---|---|---|---|
| Gm24270 | 1.15E-80 | 1.248768 | 0.997 | 1 | 2.72E-76 |
| Gm24245 | 2.85E-74 | 1.340474 | 0.978 | 0.824 | 6.72E-70 |
| Gm23935 | 5.43E-64 | 1.029763 | 1 | 1 | 1.28E-59 |
| Gm15564 | 3.28E-49 | 1.125014 | 0.994 | 0.953 | 7.74E-45 |
| Ctsd | 9.54E-45 | -0.71617 | 1 | 1 | 2.25E-40 |
| Lars2 | 2.43E-43 | 1.007585 | 1 | 1 | 5.73E-39 |
| Gm24187 | 4.18E-42 | 0.852871 | 1 | 1 | 9.85E-38 |
| Krt15 | 7.02E-42 | 0.70106 | 0.349 | 0 | 1.66E-37 |
| Csf1r | 3.57E-32 | -0.74186 | 1 | 1 | 8.41E-28 |
| Hexb | 2.17E-30 | -0.5041 | 1 | 1 | 5.11E-26 |
| Gm49369 | 1.17E-26 | -0.8594 | 0.553 | 0.802 | 2.77E-22 |
| H2-D1 | 4.88E-24 | -1.37581 | 0.911 | 0.931 | 1.15E-19 |
| Krt5 | 6.02E-24 | 0.364807 | 0.207 | 0 | 1.42E-19 |
| Itgb5 | 4.52E-22 | -0.83695 | 0.994 | 0.98 | 1.07E-17 |
| Ctsb | 9.73E-22 | -0.63949 | 1 | 1 | 2.3E-17 |
| Gm25911 | 8.5E-20 | 0.896812 | 0.905 | 0.759 | 2.01E-15 |
| Selplg | 3.2E-19 | -0.50707 | 1 | 0.998 | 7.55E-15 |
| Rps29 | 5.99E-17 | 0.50075 | 0.997 | 0.987 | 1.41E-12 |
| C1qa | 6.16E-17 | -0.34695 | 1 | 1 | 1.45E-12 |
| Psap | 2.27E-16 | -0.47511 | 1 | 0.998 | 5.35E-12 |
| Hexa | 7.09E-16 | -0.72396 | 0.992 | 0.976 | 1.67E-11 |
| Cd81 | 8.8E-16 | -0.4521 | 1 | 0.998 | 2.08E-11 |
| B2m | 1.49E-15 | -0.50088 | 1 | 0.998 | 3.51E-11 |
| Ddit4 | 1.59E-15 | 1.606149 | 0.313 | 0.089 | 3.76E-11 |
| Rn7sk | 3.5E-15 | 0.579014 | 0.989 | 0.933 | 8.27E-11 |
| Fkbp5 | 1.69E-12 | 0.814501 | 0.483 | 0.243 | 4E-08 |
| Rpl31 | 5.56E-12 | 0.742244 | 0.913 | 0.771 | 1.31E-07 |
| Rps21 | 6.91E-12 | 0.794655 | 0.897 | 0.782 | 1.63E-07 |
| Serinc3 | 1.09E-11 | -1.15648 | 0.997 | 0.998 | 2.57E-07 |
| Lgmn | 1.36E-11 | -0.38724 | 1 | 1 | 3.2E-07 |
| C1qb | 1.45E-11 | -0.27237 | 1 | 1 | 3.42E-07 |
| Rps28 | 2.26E-11 | 0.53268 | 0.966 | 0.891 | 5.32E-07 |
| Ctsz | 2.9E-11 | -0.49074 | 1 | 0.996 | 6.85E-07 |
| Itm2b | 2.95E-11 | -0.33987 | 1 | 1 | 6.95E-07 |
| Lamp1 | 6.32E-11 | -0.55612 | 0.989 | 0.987 | 1.49E-06 |
| Adrb1 | 7.14E-11 | 0.445739 | 0.659 | 0.394 | 1.68E-06 |
| Laptm5 | 7.31E-11 | -0.35239 | 1 | 1 | 1.72E-06 |
| Gm5805 | 8.86E-11 | 0.526693 | 0.743 | 0.541 | 2.09E-06 |
| Fau | 1.01E-10 | 0.686763 | 0.964 | 0.909 | 2.38E-06 |
| Il7r | 1.16E-10 | 1.10774 | 0.307 | 0.12 | 2.75E-06 |
| Per1 | 1.66E-10 | 0.914032 | 0.282 | 0.105 | 3.91E-06 |
| Rplp2 | 1.75E-10 | 0.608982 | 0.964 | 0.918 | 4.13E-06 |
| Tmbim6 | 3.39E-10 | -0.6738 | 0.961 | 0.955 | 7.99E-06 |
| Fos | 4.29E-10 | 0.326756 | 0.486 | 0.243 | 1.01E-05 |

| | | | | | |
|---|---|---|---|---|---|
| Gm4609 | 6.34E-10 | -0.65483 | 0.293 | 0.463 | 1.5E-05 |
| H2-K1 | 6.6E-10 | -1.10375 | 0.835 | 0.835 | 1.56E-05 |
| Trf | 7.06E-10 | -0.45319 | 0.994 | 0.978 | 1.67E-05 |
| Lpcat2 | 7.32E-10 | -0.57388 | 0.978 | 0.971 | 1.73E-05 |
| Man2b1 | 7.34E-10 | -0.76664 | 0.972 | 0.967 | 1.73E-05 |
| Tsc22d3 | 8.29E-10 | 0.842212 | 0.338 | 0.151 | 1.96E-05 |
| Tmsb4x | 9.32E-10 | 0.294224 | 1 | 1 | 2.2E-05 |
| Sirpa | 1.76E-09 | -0.79571 | 0.986 | 0.955 | 4.14E-05 |
| Selenop | 2.79E-09 | -0.46947 | 0.997 | 0.998 | 6.57E-05 |
| Pld4 | 6.38E-09 | -0.62776 | 0.983 | 0.949 | 0.00015 |
| P2ry12 | 6.79E-09 | -0.38375 | 1 | 1 | 0.00016 |
| Olfml3 | 7.59E-09 | -0.49862 | 0.997 | 0.989 | 0.000179 |
| Mxd4 | 7.67E-09 | 0.287467 | 0.486 | 0.283 | 0.000181 |
| mt-Co3 | 1.23E-08 | 0.415106 | 0.888 | 0.842 | 0.00029 |
| Grn | 1.47E-08 | -0.45297 | 1 | 0.991 | 0.000346 |
| Gm13456 | 1.88E-08 | -0.60392 | 0.93 | 0.92 | 0.000443 |
| Actb | 2.29E-08 | -0.32281 | 1 | 1 | 0.000539 |
| Camk2n1 | 2.73E-08 | 0.340494 | 0.399 | 0.209 | 0.000644 |
| Pdia3 | 3.31E-08 | -0.61284 | 0.958 | 0.958 | 0.000782 |
| Lag3 | 3.84E-08 | -0.9017 | 0.804 | 0.833 | 0.000905 |
| Ifngr1 | 4.9E-08 | -0.53477 | 0.98 | 0.967 | 0.001155 |
| Rps12 | 5.32E-08 | 0.625995 | 0.93 | 0.855 | 0.001256 |
| Ly86 | 6.69E-08 | -0.39656 | 1 | 0.987 | 0.001578 |
| Anxa2 | 7.72E-08 | -1.38302 | 0.209 | 0.076 | 0.001822 |
| P2ry13 | 7.92E-08 | -0.58471 | 0.98 | 0.949 | 0.001869 |
| Rpl32 | 9.21E-08 | 0.560232 | 0.925 | 0.893 | 0.002173 |
| Cmtm6 | 9.35E-08 | -1.14505 | 0.872 | 0.808 | 0.002206 |
| Rps26 | 1.17E-07 | 0.66311 | 0.757 | 0.644 | 0.002756 |
| Rps27 | 1.22E-07 | 0.398613 | 0.989 | 0.947 | 0.002871 |
| Srpk1 | 1.3E-07 | -0.30334 | 0.358 | 0.174 | 0.003074 |
| Rpl21 | 1.32E-07 | 0.61001 | 0.978 | 0.918 | 0.003121 |
| Rps8 | 1.65E-07 | 0.439421 | 0.947 | 0.871 | 0.003883 |
| Gm10800 | 2.19E-07 | 0.494612 | 0.802 | 0.784 | 0.005166 |
| Rps7 | 2.36E-07 | 0.760208 | 0.883 | 0.788 | 0.005576 |
| Rpl37a | 2.66E-07 | 0.442495 | 0.986 | 0.949 | 0.006277 |
| Tmem119 | 2.74E-07 | -0.36257 | 1 | 0.998 | 0.00647 |
| Rgs19 | 2.79E-07 | 0.287791 | 0.595 | 0.374 | 0.006587 |
| Gm15459 | 2.99E-07 | -0.30994 | 0.131 | 0.276 | 0.007051 |
| B3galnt1 | 3.73E-07 | -0.38335 | 0.299 | 0.131 | 0.008805 |
| Rpl30 | 3.82E-07 | 0.581977 | 0.919 | 0.844 | 0.009008 |
| Gp9 | 4.41E-07 | 0.402437 | 0.31 | 0.147 | 0.010397 |
| Rpl17 | 4.67E-07 | 0.624375 | 0.899 | 0.793 | 0.011024 |
| Plxdc2 | 5.83E-07 | -0.71565 | 0.969 | 0.955 | 0.013743 |
| H2-Aa | 5.92E-07 | -4.19583 | 0.081 | 0.205 | 0.013959 |
| Gm9843 | 6.05E-07 | 0.444749 | 0.939 | 0.842 | 0.01426 |
| Zfp329 | 6.77E-07 | 0.910603 | 0.291 | 0.147 | 0.015968 |
| Unc93b1 | 7.12E-07 | -0.47736 | 1 | 0.989 | 0.016793 |

| | | | | | |
|---|---|---|---|---|---|
| Gm15536 | 7.43E-07 | 0.319724 | 0.344 | 0.187 | 0.017519 |
| mt-Co2 | 8.9E-07 | 0.360069 | 0.846 | 0.737 | 0.020986 |
| Gm28437 | 9.12E-07 | 0.329553 | 0.804 | 0.655 | 0.02151 |
| Gpr34 | 1.01E-06 | -0.46767 | 1 | 0.984 | 0.023878 |
| Rps24 | 1.28E-06 | 0.560944 | 0.925 | 0.84 | 0.030129 |
| Ctss | 1.33E-06 | -0.2754 | 1 | 1 | 0.031434 |
| Rps3a1 | 1.4E-06 | 0.589674 | 0.885 | 0.795 | 0.033045 |
| Rpl17-ps5 | 1.46E-06 | 0.420502 | 0.31 | 0.169 | 0.034466 |
| Bsg | 1.48E-06 | -0.59124 | 0.874 | 0.873 | 0.03481 |
| Tpt1-ps3 | 1.79E-06 | 0.380263 | 0.74 | 0.612 | 0.042207 |

Supplementry Table 9. PB-Young DEGs in Cerebellum

| Gene | p_val | avg_log2FC | pct.1 | pct.2 | p_val_adj |
|---|---|---|---|---|---|
| Lars2 | 2.18E-94 | -3.85205 | 1 | 1 | 5.15E-90 |
| Gm15564 | 1.72E-92 | -3.80549 | 0.872 | 1 | 4.06E-88 |
| Gm23935 | 7.81E-92 | -2.63596 | 1 | 1 | 1.84E-87 |
| Gm24270 | 7.98E-92 | -2.32341 | 0.996 | 1 | 1.88E-87 |
| Gm24245 | 2.16E-91 | -2.75552 | 0.487 | 0.985 | 5.08E-87 |
| Ct010467.1 | 1.29E-85 | -0.8906 | 1 | 1 | 3.05E-81 |
| Gm25911 | 9.86E-82 | -2.17504 | 0.538 | 0.995 | 2.33E-77 |
| Gm26917 | 4.12E-72 | -2.66779 | 0.973 | 0.975 | 9.71E-68 |
| Rn7sk | 4.69E-52 | -1.83751 | 0.815 | 0.975 | 1.11E-47 |
| Tmsb4x | 5.62E-46 | 1.373552 | 1 | 1 | 1.32E-41 |
| Gm42418 | 7.15E-46 | -2.03925 | 0.893 | 0.955 | 1.69E-41 |
| Rps28 | 1.1E-40 | 2.492807 | 0.968 | 0.926 | 2.59E-36 |
| Fos | 4.4E-40 | 7.053368 | 0.59 | 0.025 | 1.04E-35 |
| Rps29 | 7.76E-32 | 2.083488 | 0.996 | 0.995 | 1.83E-27 |
| Rps27 | 9.08E-32 | 1.959199 | 0.989 | 0.965 | 2.14E-27 |
| Gm24187 | 8.54E-31 | -0.73408 | 0.998 | 1 | 2.01E-26 |
| Gm10076 | 2.27E-30 | 1.56309 | 0.964 | 0.871 | 5.35E-26 |
| Gm14303 | 5.83E-30 | 1.68903 | 0.884 | 0.653 | 1.38E-25 |
| Dusp1 | 1.54E-29 | 5.827557 | 0.468 | 0.015 | 3.63E-25 |
| Dpysl2 | 1.29E-26 | 3.36377 | 0.826 | 0.619 | 3.04E-22 |
| Tyrobp | 3.14E-26 | 1.060728 | 0.998 | 0.99 | 7.41E-22 |
| Gm10443 | 4.86E-26 | 1.3369 | 0.674 | 0.272 | 1.15E-21 |
| Egr1 | 7.83E-26 | 4.413516 | 0.422 | 0.01 | 1.85E-21 |
| Cst3 | 1.6E-25 | 0.779518 | 1 | 1 | 3.77E-21 |
| Rpl41 | 3.6E-25 | 1.479009 | 0.977 | 0.906 | 8.49E-21 |
| Gm15266 | 5.01E-25 | 1.779648 | 0.706 | 0.421 | 1.18E-20 |
| Rpl37 | 5.05E-25 | 1.702902 | 0.958 | 0.842 | 1.19E-20 |
| Gsn | 6.26E-25 | 3.327532 | 0.504 | 0.079 | 1.48E-20 |
| mt-Co2 | 1E-24 | 1.949615 | 0.87 | 0.693 | 2.36E-20 |
| Dcn | 3.6E-24 | 7.111447 | 0.431 | 0.04 | 8.5E-20 |
| Nr4a1 | 3.98E-24 | 4.350859 | 0.391 | 0.005 | 9.39E-20 |
| C1qc | 4.16E-24 | 0.609136 | 1 | 1 | 9.82E-20 |
| Fcer1g | 4.06E-23 | 0.911993 | 0.994 | 0.985 | 9.58E-19 |
| Vim | 5.42E-23 | 2.435819 | 0.441 | 0.045 | 1.28E-18 |
| Anxa2 | 6.67E-23 | 1.843465 | 0.447 | 0.045 | 1.57E-18 |
| S100a6 | 9.22E-23 | 4.639732 | 0.384 | 0.015 | 2.18E-18 |
| mt-Cytb | 1.19E-22 | 1.369871 | 0.968 | 0.871 | 2.82E-18 |
| Gm3511 | 1.28E-22 | 1.409949 | 0.803 | 0.554 | 3.02E-18 |
| Ahnak | 1.48E-22 | 3.950309 | 0.37 | 0.005 | 3.48E-18 |
| Zscan26 | 3.03E-22 | 1.443414 | 0.685 | 0.287 | 7.14E-18 |
| C1qb | 9.68E-22 | 0.639585 | 1 | 1 | 2.28E-17 |
| Crip1 | 1.58E-21 | 2.284947 | 0.42 | 0.04 | 3.72E-17 |
| Rpl37a | 3.7E-21 | 1.314878 | 0.989 | 0.965 | 8.72E-17 |
| S100a11 | 5.11E-21 | 3.353124 | 0.357 | 0.01 | 1.2E-16 |

| | | | | | |
|---|---|---|---|---|---|
| Rps25 | 6.86E-21 | 1.534212 | 0.943 | 0.708 | 1.62E-16 |
| Sod1 | 8.42E-21 | 1.826679 | 0.674 | 0.287 | 1.99E-16 |
| Ftl1 | 1.22E-20 | 0.843406 | 1 | 0.995 | 2.87E-16 |
| Gm34280 | 1.92E-20 | -1.70924 | 0.164 | 0.5 | 4.54E-16 |
| Cd9 | 3.4E-20 | 0.96564 | 0.989 | 0.941 | 8.03E-16 |
| Fth1 | 3.64E-20 | 0.820058 | 1 | 0.995 | 8.58E-16 |
| Ifitm2 | 6.11E-20 | 2.480535 | 0.437 | 0.069 | 1.44E-15 |
| Rps27rt | 8.87E-20 | 1.323389 | 0.851 | 0.668 | 2.09E-15 |
| Anxa1 | 1.67E-19 | 4.131772 | 0.328 | 0.005 | 3.94E-15 |
| Rpl38 | 3.8E-19 | 1.78758 | 0.845 | 0.574 | 8.97E-15 |
| Ftl2-ps | 5.53E-19 | 1.057396 | 0.985 | 0.916 | 1.3E-14 |
| S100a10 | 7.42E-19 | 1.87593 | 0.391 | 0.04 | 1.75E-14 |
| Rps19 | 9.44E-19 | 1.458491 | 0.952 | 0.842 | 2.23E-14 |
| Ecm1 | 1.04E-18 | 5.484358 | 0.309 | 0 | 2.46E-14 |
| Gm8129 | 1.24E-18 | 1.766818 | 0.511 | 0.178 | 2.93E-14 |
| Iqgap1 | 2.63E-18 | 1.369123 | 0.37 | 0.04 | 6.2E-14 |
| Klf4 | 2.8E-18 | 2.580409 | 0.303 | 0 | 6.61E-14 |
| Cx3cr1 | 3.13E-18 | -0.64215 | 1 | 1 | 7.39E-14 |
| Gm26857 | 3.75E-18 | -2.22582 | 0.113 | 0.411 | 8.84E-14 |
| Gpx3 | 3.89E-18 | 5.187913 | 0.3 | 0 | 9.16E-14 |
| mt-Co3 | 4.37E-18 | 1.310554 | 0.922 | 0.812 | 1.03E-13 |
| Ier3 | 5.01E-18 | 2.997896 | 0.317 | 0.01 | 1.18E-13 |
| Fosb | 5.4E-18 | 2.693897 | 0.298 | 0 | 1.27E-13 |
| Rpl39 | 8.2E-18 | 2.126216 | 0.857 | 0.673 | 1.93E-13 |
| Gm6977 | 8.59E-18 | 1.068702 | 0.79 | 0.545 | 2.03E-13 |
| Gm11518 | 9.12E-18 | 2.18011 | 0.645 | 0.376 | 2.15E-13 |
| Pltp | 1.45E-17 | 1.789331 | 0.315 | 0.01 | 3.43E-13 |
| Gm4977 | 1.6E-17 | 0.61453 | 0.307 | 0.01 | 3.76E-13 |
| Igfbp7 | 1.64E-17 | 5.385049 | 0.298 | 0.005 | 3.86E-13 |
| Mir6236 | 2.11E-17 | -0.74273 | 0.055 | 0.287 | 4.97E-13 |
| Ptma | 6.77E-17 | 1.494927 | 0.742 | 0.361 | 1.6E-12 |
| Srsf2 | 8.27E-17 | 1.37812 | 0.866 | 0.639 | 1.95E-12 |
| Wnk1 | 1.14E-16 | -1.25322 | 0.765 | 0.906 | 2.69E-12 |
| Ctss | 1.41E-16 | 0.506807 | 1 | 1 | 3.33E-12 |
| Rps18 | 1.82E-16 | 1.346508 | 0.924 | 0.817 | 4.28E-12 |
| Lmna | 1.9E-16 | 3.856954 | 0.317 | 0.025 | 4.48E-12 |
| B2m | 2.06E-16 | 0.66416 | 1 | 1 | 4.86E-12 |
| Serping1 | 2.23E-16 | 5.682668 | 0.282 | 0.005 | 5.26E-12 |
| Sipa1l2 | 4.05E-16 | -1.79053 | 0.256 | 0.55 | 9.55E-12 |
| Rpl13 | 6.12E-16 | 1.049034 | 0.964 | 0.946 | 1.44E-11 |
| Sdc4 | 7.43E-16 | 3.919144 | 0.294 | 0.015 | 1.75E-11 |
| Lgals3 | 7.46E-16 | 0.687445 | 0.3 | 0.015 | 1.76E-11 |
| Col3a1 | 1.21E-15 | 7.0525 | 0.263 | 0 | 2.86E-11 |
| Gm8464 | 1.24E-15 | -0.71783 | 0.263 | 0.545 | 2.92E-11 |
| Tppp3 | 1.49E-15 | 4.199213 | 0.269 | 0.005 | 3.5E-11 |
| Rps16 | 1.7E-15 | 1.46121 | 0.897 | 0.733 | 4.01E-11 |
| Rpl23 | 1.74E-15 | 1.305504 | 0.922 | 0.797 | 4.11E-11 |

| | | | | | |
|---|---|---|---|---|---|
| mt-Atp6 | 1.84E-15 | 1.324233 | 0.876 | 0.708 | 4.35E-11 |
| Col1a2 | 2E-15 | 6.660299 | 0.275 | 0.01 | 4.72E-11 |
| Gm5805 | 2.59E-15 | 1.260225 | 0.744 | 0.51 | 6.1E-11 |
| H2-D1 | 3.24E-15 | 0.76617 | 0.992 | 0.995 | 7.65E-11 |
| Nav2 | 3.44E-15 | -0.85411 | 0.275 | 0.609 | 8.11E-11 |
| Ly6c1 | 3.82E-15 | 4.891872 | 0.277 | 0.015 | 9.01E-11 |
| Rpl34 | 3.91E-15 | 1.621054 | 0.901 | 0.733 | 9.23E-11 |
| Selenow | 4.48E-15 | 1.83854 | 0.431 | 0.109 | 1.06E-10 |
| Gm29667 | 4.55E-15 | 0.955261 | 0.399 | 0.104 | 1.07E-10 |
| Rpl30 | 4.89E-15 | 1.454669 | 0.956 | 0.842 | 1.15E-10 |
| Cd63 | 4.95E-15 | 1.208354 | 0.874 | 0.599 | 1.17E-10 |
| Ppp1r15a | 9.44E-15 | 0.805202 | 0.315 | 0.03 | 2.23E-10 |
| Kdm6b | 9.59E-15 | 1.624545 | 0.353 | 0.054 | 2.26E-10 |
| Id3 | 1.05E-14 | 2.7037 | 0.248 | 0 | 2.47E-10 |
| Mir692-2 | 1.55E-14 | 0.783858 | 0.634 | 0.371 | 3.65E-10 |
| Itm2b | 1.55E-14 | 0.500538 | 1 | 1 | 3.66E-10 |
| Ftl1-ps1 | 1.73E-14 | 0.763772 | 0.981 | 0.926 | 4.08E-10 |
| Fn1 | 1.77E-14 | 5.234328 | 0.252 | 0.005 | 4.18E-10 |
| Rpl36 | 2.04E-14 | 1.391648 | 0.895 | 0.802 | 4.82E-10 |
| Gm10045 | 3.98E-14 | 0.809756 | 0.359 | 0.084 | 9.38E-10 |
| Gas7 | 3.99E-14 | 2.824985 | 0.279 | 0.025 | 9.41E-10 |
| Heg1 | 4.76E-14 | 2.690614 | 0.237 | 0 | 1.12E-09 |
| Rnaset2a | 4.79E-14 | 1.242123 | 0.91 | 0.866 | 1.13E-09 |
| Rps24 | 4.8E-14 | 1.439295 | 0.935 | 0.832 | 1.13E-09 |
| Cavin1 | 4.82E-14 | 3.40921 | 0.246 | 0.005 | 1.14E-09 |
| Gm38042 | 5.16E-14 | -1.58035 | 0.126 | 0.381 | 1.22E-09 |
| Rpl35 | 5.75E-14 | 1.212406 | 0.91 | 0.762 | 1.36E-09 |
| C3 | 6.73E-14 | 3.897711 | 0.277 | 0.025 | 1.59E-09 |
| Pag1 | 7.2E-14 | -0.8939 | 0.683 | 0.896 | 1.7E-09 |
| Plpp3 | 8.45E-14 | 3.660977 | 0.261 | 0.015 | 1.99E-09 |
| Lgals1 | 8.66E-14 | 1.716139 | 0.374 | 0.079 | 2.04E-09 |
| Gadd45b | 8.67E-14 | 2.548126 | 0.233 | 0 | 2.05E-09 |
| Rps3 | 9.42E-14 | 1.195522 | 0.958 | 0.886 | 2.22E-09 |
| mt-Nd1 | 1.05E-13 | 1.194612 | 0.941 | 0.817 | 2.49E-09 |
| Lpl | 1.11E-13 | 1.008051 | 0.315 | 0.045 | 2.61E-09 |
| Rpl18a | 1.13E-13 | 0.883287 | 0.998 | 1 | 2.65E-09 |
| Mgst1 | 1.58E-13 | 2.885312 | 0.229 | 0 | 3.72E-09 |
| Dock2 | 1.61E-13 | -0.53026 | 0.366 | 0.649 | 3.8E-09 |
| Sall1 | 1.68E-13 | -1.30953 | 0.761 | 0.871 | 3.97E-09 |
| Cyba | 1.7E-13 | 0.869338 | 0.977 | 0.921 | 4.01E-09 |
| Gm9843 | 1.8E-13 | 1.523953 | 0.929 | 0.802 | 4.23E-09 |
| Apoe | 1.82E-13 | 0.921738 | 0.67 | 0.317 | 4.3E-09 |
| Fbln2 | 1.9E-13 | 3.88944 | 0.244 | 0.01 | 4.48E-09 |
| Nfatc1 | 1.9E-13 | -0.89965 | 0.374 | 0.649 | 4.48E-09 |
| Akna | 2E-13 | -1.29396 | 0.389 | 0.678 | 4.72E-09 |
| Gda | 2.12E-13 | 2.128243 | 0.227 | 0 | 5E-09 |
| Rps11 | 2.25E-13 | 1.120188 | 0.95 | 0.926 | 5.31E-09 |

| | | | | | |
|---|---|---|---|---|---|
| Rps14 | 2.63E-13 | 1.119607 | 0.96 | 0.921 | 6.2E-09 |
| Gm15427 | 2.74E-13 | 1.058203 | 0.983 | 0.96 | 6.46E-09 |
| Eef1a1 | 2.93E-13 | 0.680117 | 0.996 | 0.99 | 6.9E-09 |
| Prex1 | 3.04E-13 | -1.01974 | 0.414 | 0.688 | 7.18E-09 |
| Gm32849 | 3.14E-13 | -2.32341 | 0.038 | 0.213 | 7.42E-09 |
| Kif21b | 3.45E-13 | -0.97498 | 0.246 | 0.53 | 8.13E-09 |
| C130089K0 | 4.46E-13 | -0.95358 | 0.183 | 0.46 | 1.05E-08 |
| Nedd4 | 4.97E-13 | 3.065284 | 0.229 | 0.005 | 1.17E-08 |
| Sap18 | 5.47E-13 | 1.306285 | 0.5 | 0.193 | 1.29E-08 |
| mt-Co1 | 6.52E-13 | 0.843792 | 0.992 | 0.965 | 1.54E-08 |
| Lum | 6.9E-13 | 5.51848 | 0.218 | 0 | 1.63E-08 |
| Nfil3 | 6.93E-13 | 3.610113 | 0.227 | 0.005 | 1.63E-08 |
| Gstm1 | 7.13E-13 | 0.359368 | 0.3 | 0.04 | 1.68E-08 |
| Gm5559 | 7.27E-13 | 1.610366 | 0.729 | 0.564 | 1.71E-08 |
| Fxyd5 | 7.76E-13 | 0.607989 | 0.349 | 0.069 | 1.83E-08 |
| Rpl11 | 9.02E-13 | 1.239765 | 0.866 | 0.728 | 2.13E-08 |
| Igfbp6 | 9.03E-13 | 4.579605 | 0.225 | 0.005 | 2.13E-08 |
| Rps9 | 9.05E-13 | 0.922958 | 0.981 | 0.921 | 2.14E-08 |
| Nfib | 9.24E-13 | 1.873367 | 0.216 | 0 | 2.18E-08 |
| Dpt | 9.25E-13 | 4.653041 | 0.216 | 0 | 2.18E-08 |
| Cxcl12 | 9.25E-13 | 4.163174 | 0.216 | 0 | 2.18E-08 |
| Gm3362 | 9.7E-13 | 1.379257 | 0.557 | 0.307 | 2.29E-08 |
| Rpl9-ps6 | 9.96E-13 | 1.599647 | 0.731 | 0.48 | 2.35E-08 |
| Fau | 1.03E-12 | 1.19545 | 0.975 | 0.856 | 2.44E-08 |
| Rpl35a | 1.1E-12 | 1.665599 | 0.845 | 0.693 | 2.6E-08 |
| Gm10925 | 1.2E-12 | 1.138477 | 0.849 | 0.777 | 2.83E-08 |
| S100a16 | 1.24E-12 | 2.469864 | 0.214 | 0 | 2.92E-08 |
| Htra1 | 1.24E-12 | 4.012448 | 0.214 | 0 | 2.92E-08 |
| Tagln2 | 1.28E-12 | 0.528826 | 0.46 | 0.144 | 3.01E-08 |
| Zfp691 | 1.48E-12 | 0.904395 | 0.534 | 0.228 | 3.49E-08 |
| Gm5138 | 1.56E-12 | 1.214165 | 0.536 | 0.287 | 3.68E-08 |
| Cald1 | 1.66E-12 | 2.661209 | 0.212 | 0 | 3.91E-08 |
| Emp1 | 1.66E-12 | 2.653523 | 0.212 | 0 | 3.91E-08 |
| Ccdc80 | 1.94E-12 | 2.818034 | 0.223 | 0.005 | 4.59E-08 |
| Gm1821 | 1.98E-12 | 0.925485 | 0.947 | 0.886 | 4.68E-08 |
| Col1a1 | 2.03E-12 | 6.583354 | 0.227 | 0.01 | 4.8E-08 |
| Rplp2 | 2.16E-12 | 1.139843 | 0.956 | 0.896 | 5.1E-08 |
| Sash1 | 2.28E-12 | 1.542003 | 0.221 | 0.005 | 5.38E-08 |
| Gm10736 | 2.35E-12 | 1.100957 | 0.758 | 0.589 | 5.55E-08 |
| Bgn | 2.4E-12 | 5.53985 | 0.227 | 0.01 | 5.65E-08 |
| Ebf1 | 2.48E-12 | 1.689021 | 0.218 | 0.005 | 5.85E-08 |
| Cd63-ps | 2.69E-12 | 1.118851 | 0.723 | 0.495 | 6.35E-08 |
| Mgp | 2.96E-12 | 3.860278 | 0.208 | 0 | 6.97E-08 |
| Cyr61 | 2.96E-12 | 4.065727 | 0.208 | 0 | 6.99E-08 |
| Skil | 3.13E-12 | 0.907055 | 0.716 | 0.465 | 7.37E-08 |
| Rpl32 | 3.26E-12 | 1.029913 | 0.966 | 0.916 | 7.7E-08 |
| Rps10-ps1 | 3.92E-12 | 1.267986 | 0.679 | 0.5 | 9.26E-08 |

| | | | | | |
|---|---|---|---|---|---|
| Ltbp4 | 3.95E-12 | 2.195295 | 0.206 | 0 | 9.32E-08 |
| Fbn1 | 3.98E-12 | 4.069961 | 0.214 | 0.005 | 9.39E-08 |
| Jun | 4.24E-12 | 0.517414 | 0.498 | 0.183 | 1E-07 |
| Cavin3 | 4.92E-12 | 3.630612 | 0.221 | 0.01 | 1.16E-07 |
| Sh3bgrl | 5.02E-12 | 2.224655 | 0.25 | 0.025 | 1.18E-07 |
| Sat1 | 6.51E-12 | 1.120583 | 0.903 | 0.837 | 1.54E-07 |
| Mfap5 | 7.03E-12 | 3.863614 | 0.202 | 0 | 1.66E-07 |
| Rhoj | 7.03E-12 | 2.281355 | 0.202 | 0 | 1.66E-07 |
| Alkbh5 | 7.17E-12 | 0.562679 | 0.363 | 0.094 | 1.69E-07 |
| Pcolce | 7.33E-12 | 4.608396 | 0.218 | 0.01 | 1.73E-07 |
| Snx29 | 7.45E-12 | -0.80891 | 0.172 | 0.436 | 1.76E-07 |
| S100a4 | 7.53E-12 | 2.40725 | 0.212 | 0.005 | 1.78E-07 |
| Socs3 | 8.26E-12 | 2.434878 | 0.244 | 0.02 | 1.95E-07 |
| Cav1 | 8.31E-12 | 1.56941 | 0.212 | 0.005 | 1.96E-07 |
| Plec | 8.61E-12 | 2.102489 | 0.288 | 0.05 | 2.03E-07 |
| Arglu1 | 8.82E-12 | 0.679963 | 0.824 | 0.535 | 2.08E-07 |
| Mt1 | 9.14E-12 | 1.822055 | 0.452 | 0.168 | 2.16E-07 |
| Slit3 | 9.35E-12 | 1.836501 | 0.2 | 0 | 2.21E-07 |
| Sod3 | 9.36E-12 | 3.463559 | 0.2 | 0 | 2.21E-07 |
| Tnxb | 9.36E-12 | 3.217948 | 0.2 | 0 | 2.21E-07 |
| Lamc1 | 1.11E-11 | 1.748737 | 0.21 | 0.005 | 2.61E-07 |
| Hspg2 | 1.17E-11 | 2.177556 | 0.223 | 0.015 | 2.75E-07 |
| Xdh | 1.25E-11 | 2.16176 | 0.214 | 0.01 | 2.94E-07 |
| Nid1 | 1.26E-11 | 3.292507 | 0.206 | 0.005 | 2.97E-07 |
| Plxnb2 | 1.38E-11 | -1.25389 | 0.691 | 0.842 | 3.25E-07 |
| Rpl15 | 1.39E-11 | 1.214253 | 0.805 | 0.5 | 3.29E-07 |
| Tpm4 | 1.56E-11 | 0.793286 | 0.376 | 0.099 | 3.67E-07 |
| Fstl1 | 1.61E-11 | 5.00498 | 0.221 | 0.015 | 3.8E-07 |
| Cd44 | 1.66E-11 | 0.836701 | 0.225 | 0.015 | 3.91E-07 |
| Sptbn1 | 1.71E-11 | 1.200516 | 0.296 | 0.064 | 4.04E-07 |
| Rpl23a-ps3 | 1.73E-11 | 1.137555 | 0.62 | 0.441 | 4.07E-07 |
| Sertad1 | 1.73E-11 | 1.396863 | 0.267 | 0.035 | 4.07E-07 |
| Gm17066 | 1.8E-11 | 0.658312 | 0.529 | 0.257 | 4.25E-07 |
| Uqcrh | 1.82E-11 | 1.830192 | 0.601 | 0.297 | 4.29E-07 |
| Canx | 2.02E-11 | 1.197868 | 0.868 | 0.792 | 4.77E-07 |
| Cd81 | 2.07E-11 | 0.486889 | 0.996 | 1 | 4.89E-07 |
| Emb | 2.2E-11 | 2.836643 | 0.3 | 0.074 | 5.19E-07 |
| C1s1 | 2.24E-11 | 3.928771 | 0.202 | 0.005 | 5.29E-07 |
| mt-Nd2 | 2.29E-11 | 1.55899 | 0.908 | 0.752 | 5.41E-07 |
| Rps16-ps2 | 2.54E-11 | 1.108357 | 0.754 | 0.55 | 5.98E-07 |
| Sik1 | 2.6E-11 | -0.64557 | 0.204 | 0.005 | 6.13E-07 |
| Slfn5 | 2.6E-11 | 1.90505 | 0.387 | 0.114 | 6.14E-07 |
| B430010I2 | 2.63E-11 | -2.28813 | 0.116 | 0.322 | 6.2E-07 |
| Psap | 2.78E-11 | -0.46302 | 1 | 1 | 6.56E-07 |
| Oat | 3.03E-11 | 1.581995 | 0.309 | 0.059 | 7.15E-07 |
| Aebp1 | 3.16E-11 | 4.413556 | 0.208 | 0.01 | 7.46E-07 |
| Actr2 | 3.2E-11 | 1.038996 | 0.853 | 0.743 | 7.54E-07 |

| | | | | | |
|---|---|---|---|---|---|
| Sall3 | 3.46E-11 | -0.87494 | 0.405 | 0.683 | 8.17E-07 |
| Dazap1 | 4.8E-11 | -0.99006 | 0.229 | 0.47 | 1.13E-06 |
| Csf3r | 4.81E-11 | -0.79933 | 0.792 | 0.916 | 1.14E-06 |
| Gm17087 | 5.26E-11 | -0.39128 | 0.111 | 0.312 | 1.24E-06 |
| Rpl17 | 5.34E-11 | 1.45747 | 0.884 | 0.772 | 1.26E-06 |
| Rps17 | 5.45E-11 | 1.24794 | 0.832 | 0.678 | 1.29E-06 |
| Rps10 | 5.82E-11 | 1.106761 | 0.908 | 0.812 | 1.37E-06 |
| Eef1d | 7.22E-11 | 1.483995 | 0.555 | 0.272 | 1.7E-06 |
| Nuak1 | 8.35E-11 | -1.0701 | 0.328 | 0.574 | 1.97E-06 |
| Junb | 8.44E-11 | 1.004682 | 0.637 | 0.312 | 1.99E-06 |
| mt-Nd3 | 9.71E-11 | 1.507763 | 0.618 | 0.381 | 2.29E-06 |
| Btf3 | 1.02E-10 | 1.315948 | 0.592 | 0.292 | 2.41E-06 |
| Snrpf | 1.12E-10 | 2.318254 | 0.382 | 0.134 | 2.64E-06 |
| Clic4 | 1.19E-10 | 0.756873 | 0.286 | 0.054 | 2.81E-06 |
| Rpl9 | 1.25E-10 | 1.050981 | 0.891 | 0.663 | 2.94E-06 |
| Gpx1 | 1.41E-10 | 1.067547 | 0.775 | 0.5 | 3.33E-06 |
| Slc43a3 | 1.5E-10 | -0.45984 | 0.21 | 0.015 | 3.53E-06 |
| S100a13 | 1.52E-10 | 1.217003 | 0.332 | 0.079 | 3.58E-06 |
| Plac8 | 1.59E-10 | 1.698738 | 0.208 | 0.015 | 3.74E-06 |
| Prdx1 | 1.59E-10 | 1.077875 | 0.624 | 0.322 | 3.74E-06 |
| Rplp1 | 1.79E-10 | 0.935927 | 0.989 | 0.95 | 4.23E-06 |
| Lrrc3 | 1.89E-10 | -1.27178 | 0.811 | 0.837 | 4.46E-06 |
| Prnp | 2E-10 | 0.379717 | 0.218 | 0.02 | 4.71E-06 |
| Pfn1 | 2.07E-10 | 0.861314 | 0.947 | 0.911 | 4.87E-06 |
| Bicra | 2.14E-10 | -2.60516 | 0.074 | 0.248 | 5.04E-06 |
| Tsc22d3 | 2.23E-10 | 0.33141 | 0.462 | 0.158 | 5.27E-06 |
| Rps20 | 2.25E-10 | 1.127292 | 0.926 | 0.827 | 5.31E-06 |
| Inpp5d | 2.6E-10 | -0.75186 | 0.859 | 0.96 | 6.14E-06 |
| Gm10123 | 2.85E-10 | 0.872207 | 0.941 | 0.926 | 6.72E-06 |
| Igf1 | 2.89E-10 | 1.119879 | 0.206 | 0.015 | 6.83E-06 |
| Rnaset2b | 2.92E-10 | 1.01573 | 0.918 | 0.827 | 6.89E-06 |
| Dock10 | 2.93E-10 | -1.02889 | 0.45 | 0.683 | 6.91E-06 |
| Ubb | 3E-10 | 0.75962 | 0.979 | 0.911 | 7.08E-06 |
| Prdx2 | 3.11E-10 | 1.524851 | 0.426 | 0.163 | 7.34E-06 |
| Marf1 | 3.26E-10 | 1.336704 | 0.544 | 0.292 | 7.69E-06 |
| Rcn3 | 3.35E-10 | 3.55459 | 0.233 | 0.035 | 7.9E-06 |
| Gm28661 | 3.37E-10 | 1.135127 | 0.739 | 0.683 | 7.96E-06 |
| Plin2 | 3.51E-10 | 0.261136 | 0.284 | 0.054 | 8.29E-06 |
| Vav2 | 3.73E-10 | -0.27493 | 0.179 | 0.406 | 8.8E-06 |
| Gm45221 | 3.93E-10 | -1.92736 | 0.092 | 0.272 | 9.27E-06 |
| Idh1 | 4.24E-10 | 1.130632 | 0.284 | 0.059 | 9.99E-06 |
| Map1lc3b | 4.54E-10 | 0.870657 | 0.529 | 0.238 | 1.07E-05 |
| Fchsd2 | 4.65E-10 | -1.11139 | 0.233 | 0.47 | 1.1E-05 |
| Rpl24 | 4.91E-10 | 1.203491 | 0.782 | 0.584 | 1.16E-05 |
| Gm24276 | 5.64E-10 | 0.582931 | 0.277 | 0.069 | 1.33E-05 |
| Selenom | 5.77E-10 | 1.120416 | 0.229 | 0.03 | 1.36E-05 |
| Rtraf | 6.48E-10 | 1.29572 | 0.445 | 0.183 | 1.53E-05 |

| | | | | | |
|---|---|---|---|---|---|
| Tubb5 | 6.53E-10 | 0.569987 | 0.798 | 0.49 | 1.54E-05 |
| Fundc2 | 6.7E-10 | 0.685444 | 0.25 | 0.04 | 1.58E-05 |
| Ifi207 | 7.03E-10 | 1.006153 | 0.303 | 0.074 | 1.66E-05 |
| Exosc2 | 7.31E-10 | 0.487246 | 0.372 | 0.139 | 1.72E-05 |
| Klf9 | 7.37E-10 | 0.669994 | 0.208 | 0.02 | 1.74E-05 |
| Gm26542 | 7.41E-10 | -0.77207 | 0.107 | 0.292 | 1.75E-05 |
| Lyz2 | 7.68E-10 | 0.915712 | 0.842 | 0.752 | 1.81E-05 |
| Ptafr | 7.9E-10 | 1.436938 | 0.66 | 0.525 | 1.86E-05 |
| Nfkbia | 8.16E-10 | 0.69928 | 0.63 | 0.317 | 1.93E-05 |
| Calm2 | 8.37E-10 | 1.069605 | 0.914 | 0.782 | 1.97E-05 |
| Ly6a | 8.39E-10 | 0.398916 | 0.447 | 0.178 | 1.98E-05 |
| Pim1 | 8.87E-10 | 1.042591 | 0.214 | 0.025 | 2.09E-05 |
| Prr13 | 9.48E-10 | 1.388245 | 0.489 | 0.228 | 2.24E-05 |
| Nav1 | 9.76E-10 | 1.75068 | 0.336 | 0.109 | 2.3E-05 |
| Lrrc8a | 1.05E-09 | 1.74837 | 0.429 | 0.193 | 2.48E-05 |
| Gm17511 | 1.07E-09 | 0.482842 | 0.307 | 0.089 | 2.53E-05 |
| Gm2225 | 1.2E-09 | 0.533013 | 0.382 | 0.149 | 2.84E-05 |
| Chd7 | 1.25E-09 | -1.10518 | 0.441 | 0.658 | 2.95E-05 |
| Rpl23a | 1.26E-09 | 1.02278 | 0.866 | 0.752 | 2.97E-05 |
| Cdkn1a | 1.27E-09 | 0.772629 | 0.338 | 0.099 | 2.99E-05 |
| Rap2b | 1.45E-09 | 0.861036 | 0.244 | 0.045 | 3.42E-05 |
| Ndufa1 | 1.56E-09 | 1.731548 | 0.443 | 0.188 | 3.68E-05 |
| Rpl27 | 1.71E-09 | 1.18165 | 0.815 | 0.644 | 4.03E-05 |
| Heatr5a | 1.75E-09 | -1.29459 | 0.363 | 0.579 | 4.14E-05 |
| Rpl18-ps1 | 1.96E-09 | 0.522902 | 0.439 | 0.218 | 4.62E-05 |
| Gm12918 | 1.96E-09 | 0.700517 | 0.689 | 0.505 | 4.62E-05 |
| Rpl31 | 1.97E-09 | 1.127925 | 0.889 | 0.797 | 4.65E-05 |
| Gm11478 | 2.01E-09 | 1.167163 | 0.807 | 0.738 | 4.73E-05 |
| Nbl1 | 2.16E-09 | 1.273826 | 0.216 | 0.03 | 5.1E-05 |
| Gm37306 | 2.2E-09 | -1.54205 | 0.111 | 0.292 | 5.19E-05 |
| Dynlrb1 | 2.26E-09 | 1.591786 | 0.462 | 0.203 | 5.33E-05 |
| Rpl36a | 2.62E-09 | 1.550856 | 0.683 | 0.495 | 6.17E-05 |
| Gm44860 | 2.62E-09 | -0.71893 | 0.368 | 0.604 | 6.17E-05 |
| Sun2 | 2.64E-09 | -1.85673 | 0.557 | 0.693 | 6.23E-05 |
| Gm43305 | 2.78E-09 | 1.607455 | 0.647 | 0.495 | 6.57E-05 |
| Igfbp4 | 2.82E-09 | 0.818981 | 0.313 | 0.089 | 6.66E-05 |
| Elmo1 | 2.86E-09 | -0.27819 | 0.473 | 0.718 | 6.74E-05 |
| H2-T-ps | 2.93E-09 | 2.500186 | 0.202 | 0.025 | 6.92E-05 |
| Rps13 | 2.96E-09 | 1.212956 | 0.889 | 0.777 | 6.99E-05 |
| Serf2 | 2.98E-09 | 1.076935 | 0.782 | 0.614 | 7.03E-05 |
| Rps21 | 2.99E-09 | 1.381269 | 0.926 | 0.837 | 7.06E-05 |
| Srgap2 | 3.13E-09 | -0.51849 | 0.708 | 0.896 | 7.39E-05 |
| Ost4 | 3.24E-09 | 1.755257 | 0.559 | 0.347 | 7.63E-05 |
| Qsox1 | 3.38E-09 | 1.720252 | 0.206 | 0.025 | 7.98E-05 |
| Sept2 | 3.39E-09 | 1.110674 | 0.727 | 0.5 | 8E-05 |
| Rpl13a-ps1 | 3.53E-09 | 0.846934 | 0.426 | 0.228 | 8.33E-05 |
| Ece1 | 3.64E-09 | 3.154873 | 0.202 | 0.025 | 8.58E-05 |

| | | | | | |
|---|---|---|---|---|---|
| Cox6c | 3.75E-09 | 1.875351 | 0.559 | 0.322 | 8.84E-05 |
| Rras | 3.87E-09 | 1.78615 | 0.242 | 0.045 | 9.12E-05 |
| Hmgb1 | 4.17E-09 | 1.209383 | 0.763 | 0.525 | 9.84E-05 |
| Rpl15-ps3 | 4.2E-09 | 0.932122 | 0.389 | 0.168 | 9.91E-05 |
| Gm6421 | 4.21E-09 | 0.735065 | 0.309 | 0.094 | 9.93E-05 |
| Dynll1 | 4.31E-09 | 1.177426 | 0.618 | 0.342 | 0.000102 |
| Oaz1 | 4.32E-09 | 1.030389 | 0.819 | 0.599 | 0.000102 |
| Hnrnpc | 4.5E-09 | 0.949408 | 0.536 | 0.262 | 0.000106 |
| Dhrs3 | 4.73E-09 | 1.373422 | 0.849 | 0.782 | 0.000112 |
| Itgb5 | 4.75E-09 | -0.66394 | 0.964 | 0.98 | 0.000112 |
| Dnajb1 | 4.77E-09 | 0.992224 | 0.412 | 0.158 | 0.000113 |
| Cct2 | 4.87E-09 | 1.432518 | 0.37 | 0.139 | 0.000115 |
| Rpl36al | 4.92E-09 | 1.530102 | 0.603 | 0.356 | 0.000116 |
| Ncl | 4.95E-09 | 0.827591 | 0.664 | 0.356 | 0.000117 |
| Rpl17-ps5 | 5.21E-09 | 0.724782 | 0.435 | 0.208 | 0.000123 |
| Ankrd44 | 5.33E-09 | -0.49895 | 0.416 | 0.673 | 0.000126 |
| Gm28437 | 5.45E-09 | 0.801868 | 0.847 | 0.733 | 0.000128 |
| Man2b1 | 5.49E-09 | -0.70377 | 0.914 | 0.955 | 0.000129 |
| Gm6988 | 5.72E-09 | 0.515651 | 0.34 | 0.124 | 0.000135 |
| Gm8430 | 5.95E-09 | 1.121078 | 0.651 | 0.485 | 0.00014 |
| Gm13456 | 5.98E-09 | 0.716843 | 0.971 | 0.95 | 0.000141 |
| Grk2 | 6E-09 | -0.63107 | 0.54 | 0.757 | 0.000142 |
| Cenpb | 6.06E-09 | 0.487314 | 0.307 | 0.089 | 0.000143 |
| Rpl13a | 6.16E-09 | 0.909556 | 0.903 | 0.797 | 0.000145 |
| Rpl17-ps3 | 6.39E-09 | 0.914159 | 0.58 | 0.361 | 0.000151 |
| Gm3379 | 6.41E-09 | 1.093604 | 0.725 | 0.559 | 0.000151 |
| Rps27l | 6.87E-09 | 1.503544 | 0.492 | 0.243 | 0.000162 |
| Zc3hav1 | 7.33E-09 | 1.739815 | 0.46 | 0.228 | 0.000173 |
| Serpinb6a | 7.7E-09 | 1.290704 | 0.239 | 0.05 | 0.000182 |
| Gm10288 | 7.72E-09 | 0.591283 | 0.601 | 0.366 | 0.000182 |
| Ppia | 7.97E-09 | 0.695359 | 0.958 | 0.95 | 0.000188 |
| Gm44951 | 8.45E-09 | -0.77235 | 0.399 | 0.668 | 0.000199 |
| Got1 | 8.68E-09 | 0.950157 | 0.25 | 0.054 | 0.000205 |
| Arsg | 8.88E-09 | -0.92878 | 0.158 | 0.347 | 0.000209 |
| Mrps24 | 9.75E-09 | 1.185774 | 0.315 | 0.094 | 0.00023 |
| Rpl10-ps3 | 1E-08 | 1.012556 | 0.75 | 0.525 | 0.000237 |
| Atox1 | 1.01E-08 | 1.057577 | 0.582 | 0.312 | 0.000239 |
| Aif1 | 1.03E-08 | 1.019245 | 0.863 | 0.847 | 0.000243 |
| Gm36985 | 1.07E-08 | 0.391619 | 0.261 | 0.069 | 0.000253 |
| Uqcrb | 1.08E-08 | 2.099825 | 0.435 | 0.208 | 0.000254 |
| Numb | 1.12E-08 | -0.90126 | 0.353 | 0.569 | 0.000264 |
| Rps10-ps2 | 1.12E-08 | 1.161095 | 0.777 | 0.614 | 0.000264 |
| Gm14586 | 1.12E-08 | 0.69683 | 0.616 | 0.411 | 0.000264 |
| Specc1 | 1.16E-08 | -0.46937 | 0.282 | 0.52 | 0.000273 |
| Rps12-ps9 | 1.25E-08 | 1.220403 | 0.658 | 0.5 | 0.000294 |
| Frmd4a | 1.28E-08 | -0.66062 | 0.693 | 0.851 | 0.000301 |
| Kdelr1 | 1.28E-08 | 0.681658 | 0.464 | 0.208 | 0.000301 |

| | | | | | |
|---|---|---|---|---|---|
| Spire1 | 1.29E-08 | -0.55827 | 0.233 | 0.455 | 0.000303 |
| Myl12a | 1.29E-08 | 0.986771 | 0.611 | 0.351 | 0.000305 |
| Rps5 | 1.3E-08 | 1.056267 | 0.868 | 0.738 | 0.000307 |
| Ccnd3 | 1.33E-08 | -1.7997 | 0.391 | 0.535 | 0.000313 |
| Tubb6 | 1.33E-08 | 0.747708 | 0.233 | 0.045 | 0.000314 |
| Gm15710 | 1.4E-08 | 0.88517 | 0.88 | 0.802 | 0.000329 |
| Akr1a1 | 1.4E-08 | 0.967999 | 0.805 | 0.609 | 0.00033 |
| Hcfc1r1 | 1.57E-08 | 1.263752 | 0.345 | 0.119 | 0.00037 |
| Ybx3 | 1.6E-08 | 0.627088 | 0.294 | 0.079 | 0.000377 |
| 1810037I1 | 1.73E-08 | 1.908223 | 0.357 | 0.129 | 0.000407 |
| Ctsz | 1.74E-08 | 0.546515 | 0.996 | 0.975 | 0.00041 |
| Twf1 | 1.96E-08 | 1.567694 | 0.37 | 0.153 | 0.000463 |
| Prdx6 | 2.03E-08 | 0.989686 | 0.286 | 0.084 | 0.000478 |
| Eif4g1 | 2.05E-08 | 0.575598 | 0.597 | 0.381 | 0.000484 |
| Soga1 | 2.16E-08 | -1 | 0.487 | 0.693 | 0.000509 |
| H3f3b | 2.18E-08 | 0.780197 | 0.872 | 0.688 | 0.000513 |
| Diaph1 | 2.29E-08 | 0.772574 | 0.25 | 0.059 | 0.000541 |
| Cggbp1 | 2.31E-08 | 0.357121 | 0.254 | 0.059 | 0.000545 |
| Impdh2 | 2.43E-08 | 1.226731 | 0.25 | 0.059 | 0.000574 |
| Atp6v0e | 2.56E-08 | 1.273737 | 0.58 | 0.337 | 0.000605 |
| Irf7 | 2.6E-08 | 1.470095 | 0.29 | 0.089 | 0.000613 |
| Tmsb10 | 2.65E-08 | 1.092403 | 0.643 | 0.441 | 0.000626 |
| Elob | 2.66E-08 | 1.310008 | 0.607 | 0.376 | 0.000627 |
| Trem2 | 2.69E-08 | 0.5738 | 0.973 | 0.975 | 0.000634 |
| Capn2 | 2.72E-08 | -0.92728 | 0.229 | 0.045 | 0.000642 |
| Rbm26 | 2.84E-08 | 0.99269 | 0.649 | 0.465 | 0.000669 |
| Uqcr10 | 2.84E-08 | 1.386466 | 0.437 | 0.203 | 0.000671 |
| Otub1 | 2.9E-08 | 1.048755 | 0.429 | 0.203 | 0.000685 |
| Gm3699 | 2.91E-08 | 1.134995 | 0.555 | 0.342 | 0.000688 |
| Ndufa8 | 2.96E-08 | 1.391971 | 0.38 | 0.149 | 0.000699 |
| Epn1 | 3.07E-08 | -1.34345 | 0.317 | 0.5 | 0.000725 |
| Nenf | 3.28E-08 | 1.541522 | 0.34 | 0.124 | 0.000773 |
| Rpl34-ps1 | 3.28E-08 | 1.044545 | 0.689 | 0.554 | 0.000774 |
| Taldo1 | 3.3E-08 | 1.040873 | 0.567 | 0.307 | 0.000778 |
| Ier3ip1 | 3.36E-08 | 1.416195 | 0.546 | 0.332 | 0.000791 |
| Phb2 | 3.38E-08 | 1.251843 | 0.399 | 0.173 | 0.000798 |
| Rhoc | 3.39E-08 | 0.741918 | 0.254 | 0.064 | 0.000801 |
| Acta1 | 3.54E-08 | -0.25316 | 0.074 | 0.223 | 0.000836 |
| Atp5j | 3.64E-08 | 1.261344 | 0.613 | 0.351 | 0.000858 |
| Gm10263 | 3.75E-08 | 0.330922 | 0.231 | 0.054 | 0.000884 |
| Tns3 | 3.8E-08 | -1.08684 | 0.324 | 0.515 | 0.000897 |
| Prdx5 | 3.9E-08 | 1.158216 | 0.586 | 0.361 | 0.000921 |
| Dusp3 | 3.91E-08 | 0.82513 | 0.498 | 0.248 | 0.000922 |
| Ly86 | 3.93E-08 | 0.613914 | 0.987 | 0.98 | 0.000928 |
| Nedd8 | 4.05E-08 | 1.243896 | 0.506 | 0.257 | 0.000956 |
| Capg | 4.12E-08 | 0.590066 | 0.452 | 0.208 | 0.000971 |
| Rps3a1 | 4.26E-08 | 1.069186 | 0.901 | 0.827 | 0.001004 |

| | | | | | |
|---|---|---|---|---|---|
| Tomm7 | 4.34E-08 | 1.573832 | 0.437 | 0.203 | 0.001023 |
| Errfi1 | 4.39E-08 | 0.41341 | 0.275 | 0.074 | 0.001036 |
| Gm9794 | 4.51E-08 | 0.786467 | 0.828 | 0.644 | 0.001064 |
| Snrpb | 4.51E-08 | 0.748397 | 0.597 | 0.317 | 0.001065 |
| Esd | 4.68E-08 | 0.850412 | 0.397 | 0.163 | 0.001103 |
| Gm15523 | 4.68E-08 | -1.37106 | 0.082 | 0.233 | 0.001105 |
| Ifrd1 | 4.77E-08 | 0.492561 | 0.279 | 0.079 | 0.001125 |
| Tceal9 | 5.23E-08 | 0.27469 | 0.25 | 0.064 | 0.001234 |
| Tmf1 | 5.92E-08 | 0.659421 | 0.494 | 0.257 | 0.001396 |
| Uap1 | 5.93E-08 | 1.230979 | 0.265 | 0.074 | 0.001398 |
| Cox6b1 | 6.02E-08 | 1.154784 | 0.616 | 0.386 | 0.001419 |
| Brk1 | 6.11E-08 | 1.131887 | 0.712 | 0.47 | 0.00144 |
| Rpl30-ps10 | 6.22E-08 | 0.691501 | 0.399 | 0.193 | 0.001468 |
| Rps12 | 6.4E-08 | 1.108351 | 0.939 | 0.866 | 0.001509 |
| Psmb8 | 6.58E-08 | 1.2191 | 0.702 | 0.475 | 0.001553 |
| Srgn | 7.08E-08 | 1.244352 | 0.544 | 0.317 | 0.001669 |
| Rpl19 | 7.49E-08 | 0.935657 | 0.889 | 0.797 | 0.001766 |
| Nckap1l | 7.83E-08 | -0.51139 | 0.517 | 0.748 | 0.001848 |
| Dip2b | 8.01E-08 | -0.62173 | 0.443 | 0.658 | 0.001889 |
| Basp1 | 8.11E-08 | 0.901573 | 0.92 | 0.911 | 0.001913 |
| Copb2 | 8.16E-08 | 1.439974 | 0.298 | 0.099 | 0.001924 |
| Pcnp | 8.85E-08 | 1.316397 | 0.433 | 0.218 | 0.002088 |
| Tspo | 8.93E-08 | 0.945614 | 0.595 | 0.337 | 0.002106 |
| C4b | 8.98E-08 | 1.444014 | 0.231 | 0.054 | 0.002118 |
| Mertk | 9.12E-08 | -0.74657 | 0.729 | 0.896 | 0.002152 |
| Rpl21 | 9.13E-08 | 0.823055 | 0.964 | 0.916 | 0.002153 |
| Cox8a | 9.24E-08 | 1.094803 | 0.744 | 0.52 | 0.002181 |
| Rps27a-ps2 | 9.29E-08 | 1.118613 | 0.405 | 0.213 | 0.002191 |
| Rps13-ps2 | 9.3E-08 | 0.803168 | 0.681 | 0.505 | 0.002194 |
| Dbi | 1.02E-07 | 0.828436 | 0.389 | 0.158 | 0.002405 |
| Gm47087 | 1.08E-07 | -0.61381 | 0.074 | 0.218 | 0.002547 |
| Etf1 | 1.16E-07 | 0.74304 | 0.284 | 0.089 | 0.002727 |
| Sec61g | 1.16E-07 | 1.481572 | 0.532 | 0.312 | 0.002746 |
| Lrch1 | 1.17E-07 | -1.34758 | 0.311 | 0.49 | 0.002761 |
| Wtap | 1.18E-07 | -0.4577 | 0.475 | 0.203 | 0.002777 |
| Vamp8 | 1.19E-07 | 1.052963 | 0.601 | 0.386 | 0.002818 |
| Rps26 | 1.2E-07 | 1.079764 | 0.851 | 0.723 | 0.002832 |
| Ac165079.: | 1.2E-07 | 0.569953 | 0.359 | 0.163 | 0.002832 |
| Gm10180 | 1.32E-07 | 0.567508 | 0.603 | 0.416 | 0.003121 |
| Gm10134 | 1.36E-07 | 1.806412 | 0.261 | 0.089 | 0.003197 |
| Selplg | 1.36E-07 | 0.421882 | 0.994 | 0.99 | 0.003202 |
| Cyb5r3 | 1.38E-07 | 0.738835 | 0.345 | 0.129 | 0.003264 |
| Txndc17 | 1.39E-07 | 0.893187 | 0.424 | 0.178 | 0.003267 |
| Gm10059 | 1.44E-07 | 0.469181 | 0.515 | 0.327 | 0.00339 |
| Cstb | 1.44E-07 | 1.241606 | 0.46 | 0.228 | 0.003398 |
| Cd47 | 1.44E-07 | 0.382089 | 0.737 | 0.465 | 0.003407 |
| Sfr1 | 1.45E-07 | 0.620464 | 0.378 | 0.153 | 0.003412 |

| | | | | | |
|---|---|---|---|---|---|
| Rps7 | 1.49E-07 | 0.80915 | 0.914 | 0.762 | 0.003521 |
| Gm10020 | 1.51E-07 | 0.570749 | 0.338 | 0.153 | 0.003568 |
| Olfml2b | 1.53E-07 | 1.333601 | 0.206 | 0.045 | 0.003602 |
| Tceal8 | 1.6E-07 | 0.505457 | 0.223 | 0.05 | 0.003768 |
| Rps27a | 1.63E-07 | 1.177421 | 0.866 | 0.777 | 0.003844 |
| Rpl3-ps2 | 1.64E-07 | 0.434376 | 0.357 | 0.158 | 0.003861 |
| Nme1 | 1.67E-07 | 1.250938 | 0.359 | 0.149 | 0.003934 |
| Rps15a | 1.67E-07 | 1.089892 | 0.773 | 0.594 | 0.003948 |
| mt-Rnr2 | 1.69E-07 | -0.5167 | 0.994 | 1 | 0.00398 |
| Rgs10 | 1.71E-07 | 0.7213 | 0.922 | 0.881 | 0.004041 |
| Degs1 | 1.75E-07 | 0.505428 | 0.435 | 0.198 | 0.004135 |
| Ifi211 | 1.78E-07 | 1.417248 | 0.218 | 0.054 | 0.004203 |
| Adap2os | 1.81E-07 | 1.053516 | 0.784 | 0.718 | 0.004275 |
| Zbp1 | 1.85E-07 | 1.734311 | 0.231 | 0.064 | 0.004369 |
| D8Ertd738 | 1.86E-07 | 1.066121 | 0.521 | 0.277 | 0.004392 |
| Rpl22l1 | 1.89E-07 | 1.259534 | 0.456 | 0.228 | 0.004452 |
| Cebpd | 1.99E-07 | 0.664953 | 0.519 | 0.272 | 0.004686 |
| Atp6v1g1 | 2.02E-07 | 1.414673 | 0.632 | 0.446 | 0.004773 |
| Smdt1 | 2.04E-07 | 1.40397 | 0.424 | 0.218 | 0.004801 |
| Gm3791 | 2.05E-07 | -1.00621 | 0.09 | 0.243 | 0.00484 |
| Ndufb9 | 2.1E-07 | 1.231135 | 0.515 | 0.287 | 0.004965 |
| Ndufs8 | 2.13E-07 | 1.137349 | 0.412 | 0.188 | 0.005033 |
| Itpripl1 | 2.18E-07 | 2.908089 | 0.508 | 0.391 | 0.005142 |
| Sept7 | 2.2E-07 | 0.773869 | 0.553 | 0.302 | 0.005182 |
| Gm37033 | 2.21E-07 | -3.17803 | 0.118 | 0.267 | 0.005202 |
| Sfi1 | 2.21E-07 | -1.46616 | 0.298 | 0.485 | 0.005219 |
| Mlf2 | 2.21E-07 | 0.633727 | 0.426 | 0.198 | 0.005223 |
| Sgk3 | 2.25E-07 | -1.0824 | 0.204 | 0.381 | 0.005303 |
| H3f3c | 2.3E-07 | 0.553081 | 0.204 | 0.045 | 0.005421 |
| Ndufa4 | 2.31E-07 | 1.614974 | 0.462 | 0.248 | 0.005454 |
| Psmd8 | 2.39E-07 | 0.964122 | 0.418 | 0.193 | 0.005636 |
| Rps18-ps3 | 2.4E-07 | 0.601146 | 0.714 | 0.554 | 0.005662 |
| H2afj | 2.4E-07 | 0.711678 | 0.468 | 0.238 | 0.005664 |
| Gm5879 | 2.41E-07 | 0.621541 | 0.458 | 0.262 | 0.005679 |
| Gm15500 | 2.44E-07 | 1.384068 | 0.714 | 0.584 | 0.005751 |
| Sap18b | 2.46E-07 | 0.569072 | 0.282 | 0.094 | 0.005814 |
| Slc8a1 | 2.47E-07 | -2.05002 | 0.101 | 0.248 | 0.005826 |
| 4933406I1 | 2.49E-07 | -1.159 | 0.069 | 0.208 | 0.005867 |
| Gna12 | 2.52E-07 | -1.28205 | 0.517 | 0.653 | 0.005948 |
| Ctsl | 2.52E-07 | 0.616561 | 0.966 | 0.916 | 0.005954 |
| Npc2 | 2.55E-07 | 0.743489 | 0.914 | 0.817 | 0.006023 |
| Tex264 | 2.58E-07 | 0.842637 | 0.303 | 0.109 | 0.006079 |
| Ubl7 | 2.6E-07 | 1.806432 | 0.292 | 0.104 | 0.006124 |
| Laptm4a | 2.68E-07 | 0.710939 | 0.887 | 0.792 | 0.00632 |
| Gm4617 | 2.73E-07 | 0.689393 | 0.38 | 0.178 | 0.006429 |
| Mpc2 | 2.78E-07 | 1.676534 | 0.342 | 0.139 | 0.006563 |
| Clic1 | 2.88E-07 | 0.812827 | 0.838 | 0.668 | 0.006784 |

| | | | | | |
|---|---|---|---|---|---|
| Rpl27-ps3 | 2.89E-07 | 1.023886 | 0.498 | 0.312 | 0.006806 |
| Slc25a5 | 2.96E-07 | 1.098267 | 0.748 | 0.594 | 0.006976 |
| Ndufc1 | 3.03E-07 | 0.974122 | 0.321 | 0.119 | 0.007154 |
| Rexo2 | 3.05E-07 | 0.343828 | 0.332 | 0.129 | 0.007185 |
| Gm10243 | 3.06E-07 | 1.425049 | 0.294 | 0.119 | 0.007209 |
| Serpinf1 | 3.07E-07 | 0.724501 | 0.317 | 0.124 | 0.007244 |
| Epb41l2 | 3.11E-07 | -0.66366 | 0.821 | 0.916 | 0.007328 |
| Txn1 | 3.15E-07 | 0.9792 | 0.458 | 0.218 | 0.007435 |
| Timm13 | 3.27E-07 | 1.014383 | 0.395 | 0.173 | 0.007725 |
| Nme2 | 3.38E-07 | 1.098355 | 0.548 | 0.302 | 0.007978 |
| Ac154667.: | 3.48E-07 | 0.440634 | 0.223 | 0.064 | 0.008206 |
| Klf2 | 3.52E-07 | -0.78652 | 0.342 | 0.119 | 0.008299 |
| Ndufb1-ps | 3.55E-07 | 1.827116 | 0.418 | 0.228 | 0.008364 |
| Morf4l2 | 3.7E-07 | 0.793216 | 0.389 | 0.178 | 0.008718 |
| Eif3e | 3.79E-07 | 0.755054 | 0.466 | 0.238 | 0.00893 |
| Hps3 | 3.99E-07 | -0.50886 | 0.347 | 0.564 | 0.009408 |
| Btg2 | 4.1E-07 | 0.317104 | 0.592 | 0.322 | 0.009674 |
| Ndufa5 | 4.15E-07 | 1.226229 | 0.286 | 0.099 | 0.009788 |
| Gm3788 | 4.17E-07 | 0.385589 | 0.288 | 0.109 | 0.009833 |
| Ikzf1 | 4.2E-07 | -0.66897 | 0.439 | 0.629 | 0.009906 |
| Rpl21-ps15 | 4.4E-07 | 0.718406 | 0.519 | 0.351 | 0.010368 |
| Ndufa13 | 4.42E-07 | 0.908055 | 0.569 | 0.337 | 0.010426 |
| Mmp2 | 4.51E-07 | 1.03447 | 0.324 | 0.129 | 0.010641 |
| Fkbp9 | 4.53E-07 | 0.519696 | 0.225 | 0.059 | 0.010679 |
| Rpl27a | 4.71E-07 | 1.347717 | 0.859 | 0.718 | 0.011113 |
| Cox5a | 4.71E-07 | 1.106153 | 0.376 | 0.158 | 0.011117 |
| Actn4 | 4.74E-07 | -0.31512 | 0.401 | 0.173 | 0.011191 |
| Sem1 | 4.92E-07 | 0.760967 | 0.54 | 0.307 | 0.011607 |
| Eif1a | 4.99E-07 | 1.036914 | 0.3 | 0.114 | 0.011766 |
| Entpd1 | 5.08E-07 | -0.64822 | 0.84 | 0.931 | 0.011993 |
| Odc1 | 5.09E-07 | 0.632748 | 0.204 | 0.045 | 0.011997 |
| Mlph | 5.13E-07 | -1.17746 | 0.21 | 0.391 | 0.012113 |
| Pttg1ip | 5.14E-07 | 0.771069 | 0.483 | 0.257 | 0.012134 |
| Fam114a1 | 5.27E-07 | 0.388859 | 0.208 | 0.05 | 0.012425 |
| Plcb2 | 5.36E-07 | -1.72615 | 0.139 | 0.297 | 0.01265 |
| Sdhd | 5.37E-07 | 1.44422 | 0.345 | 0.144 | 0.012672 |
| Trpm2 | 5.38E-07 | -0.61317 | 0.137 | 0.302 | 0.012692 |
| Sde2 | 5.42E-07 | 0.866592 | 0.223 | 0.054 | 0.012788 |
| Rps24-ps3 | 5.45E-07 | 0.672308 | 0.525 | 0.371 | 0.012847 |
| Pdcl3 | 5.49E-07 | 1.853486 | 0.214 | 0.054 | 0.012948 |
| Vat1 | 5.97E-07 | 0.817842 | 0.231 | 0.064 | 0.01408 |
| Dagla | 6.03E-07 | -1.61979 | 0.195 | 0.371 | 0.014218 |
| Atp5g3 | 6.03E-07 | 0.934184 | 0.498 | 0.272 | 0.014227 |
| Gm17494 | 6.06E-07 | 1.644539 | 0.334 | 0.173 | 0.0143 |
| Dap | 6.07E-07 | 0.671217 | 0.309 | 0.114 | 0.014316 |
| Acadl | 6.17E-07 | 0.749673 | 0.273 | 0.094 | 0.01455 |
| Impact | 6.18E-07 | 1.680953 | 0.361 | 0.168 | 0.014579 |

| | | | | | |
|---|---|---|---|---|---|
| Gm15635 | 6.47E-07 | -2.04506 | 0.095 | 0.238 | 0.015263 |
| Eef1b2 | 6.59E-07 | 0.950101 | 0.653 | 0.426 | 0.015543 |
| Tfg | 6.64E-07 | 0.723239 | 0.324 | 0.129 | 0.015674 |
| Mroh1 | 6.67E-07 | -2.06373 | 0.162 | 0.312 | 0.015726 |
| Gm11560 | 6.78E-07 | 1.072903 | 0.41 | 0.213 | 0.015982 |
| Sparc | 7.01E-07 | 0.42932 | 0.996 | 0.985 | 0.016546 |
| Gm11361 | 7.31E-07 | 0.680032 | 0.647 | 0.47 | 0.017234 |
| D17H6S53E | 7.36E-07 | 0.708277 | 0.206 | 0.054 | 0.017355 |
| H2afz | 7.38E-07 | 0.8634 | 0.58 | 0.337 | 0.01741 |
| Cd14 | 7.49E-07 | 0.858941 | 0.754 | 0.624 | 0.017662 |
| Rpl28 | 7.66E-07 | 1.041051 | 0.819 | 0.693 | 0.018077 |
| Scaf1 | 7.76E-07 | -1.02599 | 0.405 | 0.559 | 0.018299 |
| Gm8960 | 7.84E-07 | 0.53233 | 0.458 | 0.282 | 0.018496 |
| Metrnl | 7.86E-07 | -0.45362 | 0.202 | 0.045 | 0.018545 |
| Arl1 | 7.89E-07 | 0.846836 | 0.378 | 0.173 | 0.018622 |
| Eif3i | 7.93E-07 | 1.265611 | 0.397 | 0.183 | 0.0187 |
| 2410015M | 7.93E-07 | 1.432389 | 0.366 | 0.163 | 0.018714 |
| Pcna | 7.98E-07 | 0.70542 | 0.321 | 0.124 | 0.018834 |
| Ccdc124 | 8.09E-07 | 1.632821 | 0.284 | 0.104 | 0.01908 |
| Sumo3 | 8.21E-07 | 0.731936 | 0.382 | 0.168 | 0.019361 |
| Tmem258 | 8.25E-07 | 1.037388 | 0.468 | 0.262 | 0.019454 |
| Top1 | 8.45E-07 | 0.61692 | 0.408 | 0.188 | 0.019935 |
| Chchd1 | 8.52E-07 | 1.51103 | 0.246 | 0.079 | 0.020096 |
| Ddah2 | 8.53E-07 | 1.325013 | 0.244 | 0.074 | 0.020129 |
| Ifitm3 | 8.82E-07 | 1.356132 | 0.63 | 0.416 | 0.020816 |
| Fam221b | 8.84E-07 | 0.679398 | 0.525 | 0.327 | 0.020842 |
| Psmc4 | 8.86E-07 | 0.591629 | 0.313 | 0.124 | 0.020904 |
| Cycs | 8.88E-07 | 1.157977 | 0.376 | 0.168 | 0.020942 |
| C1qa | 8.89E-07 | 0.272501 | 1 | 1 | 0.020979 |
| Snx3 | 8.97E-07 | 0.481294 | 0.515 | 0.262 | 0.021156 |
| Sub1 | 8.97E-07 | 1.110697 | 0.487 | 0.272 | 0.021161 |
| Psma6 | 9.27E-07 | 1.084573 | 0.462 | 0.257 | 0.021863 |
| Arpp19 | 9.46E-07 | 1.358306 | 0.422 | 0.218 | 0.022323 |
| Psmd6 | 9.54E-07 | 1.518463 | 0.296 | 0.114 | 0.022507 |
| Cdk13 | 9.73E-07 | -1.60501 | 0.374 | 0.505 | 0.022955 |
| Rsrp1 | 9.94E-07 | 0.703823 | 0.937 | 0.827 | 0.023446 |
| Taok2 | 9.97E-07 | -1.37301 | 0.246 | 0.411 | 0.023525 |
| Tomm22 | 1.02E-06 | 1.389977 | 0.351 | 0.153 | 0.024029 |
| Eif3h | 1.02E-06 | 0.949021 | 0.494 | 0.272 | 0.024048 |
| Bloc1s1 | 1.05E-06 | 1.192306 | 0.342 | 0.139 | 0.024856 |
| Ostc | 1.06E-06 | 0.610931 | 0.447 | 0.218 | 0.024895 |
| Gas5 | 1.07E-06 | 1.750377 | 0.565 | 0.371 | 0.025234 |
| Rpl31-ps8 | 1.07E-06 | 0.698562 | 0.737 | 0.584 | 0.025277 |
| Gm11450 | 1.08E-06 | 0.341901 | 0.206 | 0.054 | 0.025525 |
| 1810058I24 | 1.12E-06 | 0.905978 | 0.368 | 0.153 | 0.026388 |
| Med28 | 1.13E-06 | 0.708045 | 0.431 | 0.208 | 0.026558 |
| Atp5d | 1.13E-06 | 0.792816 | 0.536 | 0.327 | 0.026678 |

| | | | | | |
|---|---|---|---|---|---|
| Mmp14 | 1.13E-06 | 0.555617 | 0.342 | 0.158 | 0.026717 |
| Rpl29 | 1.15E-06 | 0.703774 | 0.756 | 0.559 | 0.027122 |
| Tax1bp1 | 1.16E-06 | 0.876127 | 0.523 | 0.292 | 0.027462 |
| Gm21811 | 1.18E-06 | -0.91968 | 0.128 | 0.282 | 0.027749 |
| Gng10 | 1.2E-06 | 1.022709 | 0.718 | 0.609 | 0.028228 |
| Lrpap1 | 1.21E-06 | 0.578634 | 0.525 | 0.287 | 0.028498 |
| Mrpl33 | 1.21E-06 | 0.823532 | 0.351 | 0.149 | 0.028544 |
| Zeb2 | 1.22E-06 | -1.35848 | 0.561 | 0.688 | 0.028709 |
| Gm44667 | 1.22E-06 | -0.97687 | 0.086 | 0.223 | 0.028726 |
| Rpl10 | 1.23E-06 | 0.742399 | 0.964 | 0.911 | 0.029051 |
| Kdelr2 | 1.23E-06 | 0.337434 | 0.55 | 0.297 | 0.029086 |
| Arhgap39 | 1.27E-06 | -0.74631 | 0.359 | 0.559 | 0.02995 |
| Litaf | 1.28E-06 | 0.505696 | 0.538 | 0.287 | 0.030275 |
| Gm10275 | 1.31E-06 | 1.221432 | 0.582 | 0.436 | 0.030836 |
| Rpl30-ps8 | 1.31E-06 | 0.321762 | 0.221 | 0.069 | 0.031006 |
| Glrx2 | 1.34E-06 | 3.055286 | 0.21 | 0.059 | 0.031507 |
| Tanc2 | 1.36E-06 | -0.60823 | 0.702 | 0.866 | 0.032043 |
| Slc48a1 | 1.42E-06 | 1.300623 | 0.252 | 0.084 | 0.033442 |
| Cnn2 | 1.46E-06 | 0.717185 | 0.21 | 0.054 | 0.034341 |
| Csnk2b | 1.47E-06 | 0.607533 | 0.397 | 0.183 | 0.034693 |
| Ube2l6 | 1.48E-06 | 0.944353 | 0.258 | 0.084 | 0.034853 |
| Eif4ebp1 | 1.49E-06 | 1.18574 | 0.313 | 0.124 | 0.035162 |
| Gm3531 | 1.52E-06 | 1.288078 | 0.357 | 0.178 | 0.035917 |
| Ndufs6 | 1.53E-06 | 0.655844 | 0.359 | 0.158 | 0.036137 |
| Zfp706 | 1.56E-06 | 0.856052 | 0.588 | 0.356 | 0.036734 |
| Sumo2 | 1.59E-06 | 1.279872 | 0.458 | 0.248 | 0.037503 |
| Chchd2 | 1.6E-06 | 0.629122 | 0.597 | 0.361 | 0.037646 |
| Znhit1 | 1.64E-06 | 2.145343 | 0.254 | 0.089 | 0.03879 |
| Sptssa | 1.64E-06 | 0.805235 | 0.395 | 0.183 | 0.038802 |
| Arf1 | 1.68E-06 | 0.809575 | 0.725 | 0.545 | 0.03952 |
| Elof1 | 1.7E-06 | 0.737732 | 0.229 | 0.069 | 0.04011 |
| Gm38056 | 1.73E-06 | -1.93088 | 0.237 | 0.391 | 0.040696 |
| Usmg5 | 1.78E-06 | 1.500137 | 0.487 | 0.287 | 0.041923 |
| E230029C0 | 1.81E-06 | -0.75372 | 0.67 | 0.812 | 0.042725 |
| Gm27201 | 1.81E-06 | -0.96553 | 0.084 | 0.213 | 0.04278 |
| Snrpd3 | 1.82E-06 | 0.886764 | 0.313 | 0.129 | 0.042909 |
| Gnb1 | 1.85E-06 | 0.919987 | 0.828 | 0.733 | 0.043591 |
| Eif3f | 1.86E-06 | 1.022994 | 0.658 | 0.47 | 0.043915 |
| Ccdc152 | 1.89E-06 | 0.777969 | 0.3 | 0.129 | 0.04463 |
| Rpl4 | 1.89E-06 | 0.550435 | 0.899 | 0.738 | 0.044683 |
| Fgl2 | 1.91E-06 | 0.672549 | 0.244 | 0.079 | 0.045101 |
| Vmp1 | 1.91E-06 | 0.913995 | 0.38 | 0.183 | 0.045141 |
| Cfap74 | 1.95E-06 | -0.80721 | 0.147 | 0.307 | 0.045951 |
| Hsbp1 | 1.96E-06 | 1.47336 | 0.544 | 0.342 | 0.046213 |
| Creb3 | 1.98E-06 | 0.99081 | 0.361 | 0.163 | 0.046809 |
| Nop53 | 2E-06 | 0.984523 | 0.55 | 0.342 | 0.047241 |
| Hint1 | 2E-06 | 1.043597 | 0.544 | 0.312 | 0.047265 |

| | | | | | |
|---|---|---|---|---|---|
| Rap1gds1 | 2E-06 | -0.47388 | 0.456 | 0.639 | 0.047287 |
| Cerk | 2.02E-06 | -0.82323 | 0.328 | 0.52 | 0.047567 |
| Gm6055 | 2.02E-06 | 1.038188 | 0.233 | 0.074 | 0.047714 |
| Commd1 | 2.04E-06 | 1.060925 | 0.303 | 0.119 | 0.048205 |
| Gm7536 | 2.04E-06 | 1.058503 | 0.706 | 0.569 | 0.048237 |
| Git2 | 2.07E-06 | -0.63515 | 0.624 | 0.787 | 0.048801 |
| Ufc1 | 2.07E-06 | 0.566979 | 0.248 | 0.079 | 0.048867 |
| Rreb1 | 2.12E-06 | -1.87806 | 0.263 | 0.421 | 0.049997 |

Supplementry Table 9. PB-Young DEGs in Cortex

| Gene | p_val | avg_log2FC | pct.1 | pct.2 | p_val_adj |
|---|---|---|---|---|---|
| Csf1r | 6.94E-64 | -1.37834 | 1 | 1 | 1.64E-59 |
| P2ry12 | 2.88E-50 | -0.91943 | 1 | 1 | 6.79E-46 |
| Cx3cr1 | 3.46E-47 | -0.92047 | 1 | 1 | 8.17E-43 |
| Itgb5 | 9.95E-44 | -1.26155 | 0.986 | 0.98 | 2.35E-39 |
| C1qc | 2.72E-43 | 0.626205 | 1 | 1 | 6.42E-39 |
| Gpr34 | 1.76E-41 | -1.1437 | 0.984 | 0.983 | 4.14E-37 |
| Psap | 1.39E-37 | -0.7962 | 1 | 1 | 3.28E-33 |
| Man2b1 | 6.17E-37 | -1.56268 | 0.895 | 0.931 | 1.45E-32 |
| C1qb | 3.48E-34 | 0.611707 | 1 | 1 | 8.2E-30 |
| Sirpa | 4.46E-33 | -1.57304 | 0.888 | 0.965 | 1.05E-28 |
| Ct010467.1 | 7.78E-33 | -0.38141 | 1 | 1 | 1.83E-28 |
| Serinc3 | 2.1E-29 | -2.03488 | 0.988 | 0.986 | 4.95E-25 |
| Cmtm6 | 2.69E-29 | -2.01966 | 0.758 | 0.856 | 6.34E-25 |
| P2ry13 | 1.14E-28 | -1.2203 | 0.953 | 0.954 | 2.7E-24 |
| Hexb | 8.15E-28 | -0.43276 | 1 | 1 | 1.92E-23 |
| Selenop | 3.1E-24 | -0.74098 | 0.984 | 0.986 | 7.31E-20 |
| Tmsb4x | 9.69E-24 | 0.810816 | 0.998 | 0.997 | 2.28E-19 |
| Fcer1g | 3.93E-23 | 0.695907 | 0.995 | 0.986 | 9.27E-19 |
| Plxdc2 | 1.53E-22 | -1.6171 | 0.83 | 0.879 | 3.62E-18 |
| Rhob | 3.25E-22 | -1.15249 | 0.902 | 0.919 | 7.68E-18 |
| Rnaset2a | 1.06E-20 | 1.008659 | 0.965 | 0.928 | 2.49E-16 |
| S100a8 | 2E-20 | -1.28087 | 0.007 | 0.202 | 4.72E-16 |
| Lamp1 | 2.54E-20 | -0.78639 | 0.967 | 0.963 | 5.99E-16 |
| Gm1821 | 5.06E-20 | 0.792653 | 0.953 | 0.948 | 1.19E-15 |
| Lars2 | 1.99E-18 | -1.6954 | 1 | 1 | 4.7E-14 |
| Marcks | 2.48E-18 | -0.59487 | 0.998 | 0.994 | 5.86E-14 |
| Cst3 | 3.25E-18 | 0.562647 | 1 | 1 | 7.67E-14 |
| Olfr620 | 1.78E-16 | -0.56539 | 0.102 | 0.331 | 4.19E-12 |
| Asah1 | 1.97E-16 | -1.41889 | 0.793 | 0.824 | 4.65E-12 |
| Rps27 | 2.4E-16 | 1.244812 | 0.942 | 0.925 | 5.66E-12 |
| Gm13493 | 4.05E-16 | -0.73163 | 0.774 | 0.865 | 9.56E-12 |
| Tm9sf2 | 4.09E-16 | -1.6229 | 0.684 | 0.761 | 9.64E-12 |
| Pfn1 | 4.52E-16 | 0.726277 | 0.958 | 0.948 | 1.07E-11 |
| C1qa | 6.74E-16 | 0.263216 | 1 | 1 | 1.59E-11 |
| Gm8797 | 1.12E-15 | 0.632752 | 0.923 | 0.862 | 2.63E-11 |
| Fos | 2.89E-15 | -0.41256 | 0.205 | 0.017 | 6.83E-11 |
| Gm14303 | 4.02E-15 | 1.06973 | 0.863 | 0.778 | 9.47E-11 |
| Capza2 | 4.83E-15 | -1.25453 | 0.863 | 0.862 | 1.14E-10 |
| Lyn | 4.83E-15 | -1.47254 | 0.835 | 0.839 | 1.14E-10 |
| Cd164 | 9.48E-15 | -0.80717 | 0.963 | 0.963 | 2.24E-10 |
| Tyrobp | 1E-14 | 0.609338 | 0.993 | 0.983 | 2.37E-10 |
| Sall1 | 1.55E-14 | -0.94313 | 0.853 | 0.948 | 3.65E-10 |
| Pros1 | 2.21E-14 | -1.46534 | 0.549 | 0.709 | 5.22E-10 |
| Actb | 2.42E-14 | 0.29991 | 1 | 1 | 5.72E-10 |

| | | | | | |
|---|---|---|---|---|---|
| Rpl18a | 2.91E-14 | 0.892505 | 0.991 | 0.963 | 6.86E-10 |
| Rnaset2b | 3.45E-14 | 0.931313 | 0.958 | 0.954 | 8.14E-10 |
| Tmbim6 | 5.84E-14 | -0.71951 | 0.958 | 0.939 | 1.38E-09 |
| Rps29 | 9.17E-14 | 1.248966 | 0.981 | 0.986 | 2.16E-09 |
| Rpl37 | 2.23E-13 | 1.315972 | 0.865 | 0.87 | 5.25E-09 |
| Rpl13 | 4.6E-13 | 0.982423 | 0.914 | 0.899 | 1.09E-08 |
| Lpcat2 | 5.99E-13 | -0.62013 | 0.974 | 0.954 | 1.41E-08 |
| Zfp36l2 | 8.17E-13 | -1.02213 | 0.333 | 0.594 | 1.93E-08 |
| Adap2os | 8.42E-13 | 0.883969 | 0.891 | 0.769 | 1.99E-08 |
| Gsn | 8.53E-13 | 0.353042 | 0.2 | 0.029 | 2.01E-08 |
| Gm10076 | 1.2E-12 | 0.995618 | 0.891 | 0.844 | 2.83E-08 |
| Lgmn | 1.21E-12 | -0.32164 | 1 | 1 | 2.84E-08 |
| Adgrg1 | 1.4E-12 | -0.82916 | 0.865 | 0.89 | 3.3E-08 |
| Ftl1 | 3.04E-12 | 0.463174 | 0.993 | 0.988 | 7.16E-08 |
| Rplp1 | 3.11E-12 | 1.000996 | 0.937 | 0.934 | 7.33E-08 |
| Rplp2 | 6.68E-12 | 1.011662 | 0.902 | 0.85 | 1.58E-07 |
| Rpl41 | 7.93E-12 | 1.069424 | 0.916 | 0.885 | 1.87E-07 |
| Fmnl1 | 9.64E-12 | -1.58634 | 0.291 | 0.493 | 2.27E-07 |
| Rps6ka1 | 1.12E-11 | -1.09555 | 0.77 | 0.807 | 2.64E-07 |
| Gm47283 | 3.63E-11 | 1.691372 | 0.356 | 0.15 | 8.56E-07 |
| Gm17494 | 3.98E-11 | -0.47158 | 0.358 | 0.159 | 9.39E-07 |
| Slc7a8 | 4.78E-11 | -1.2227 | 0.651 | 0.752 | 1.13E-06 |
| Ivns1abp | 7.31E-11 | -1.06913 | 0.847 | 0.879 | 1.73E-06 |
| Rpl18-ps1 | 7.42E-11 | 0.426909 | 0.391 | 0.182 | 1.75E-06 |
| Sun2 | 1.05E-10 | -1.52837 | 0.553 | 0.671 | 2.49E-06 |
| Rpl37a | 1.25E-10 | 0.980598 | 0.926 | 0.931 | 2.94E-06 |
| Unc93b1 | 1.48E-10 | -0.43366 | 0.977 | 0.988 | 3.5E-06 |
| Gm3379 | 1.57E-10 | 1.085662 | 0.667 | 0.522 | 3.71E-06 |
| Rpl30 | 2.21E-10 | 1.11117 | 0.856 | 0.804 | 5.21E-06 |
| Rps27rt | 2.52E-10 | 0.868522 | 0.793 | 0.723 | 5.95E-06 |
| Amfr | 2.91E-10 | -2.20734 | 0.349 | 0.49 | 6.86E-06 |
| Tgfbr2 | 3.32E-10 | -0.96379 | 0.828 | 0.867 | 7.83E-06 |
| Serinc1 | 4.03E-10 | -1.45601 | 0.563 | 0.669 | 9.51E-06 |
| Trem2 | 5.52E-10 | 0.484905 | 0.977 | 0.977 | 1.3E-05 |
| Ftl1-ps1 | 6.34E-10 | 0.523855 | 0.951 | 0.951 | 1.49E-05 |
| mt-Rnr2 | 6.76E-10 | -0.53514 | 0.998 | 0.994 | 1.59E-05 |
| Gm10123 | 8.85E-10 | 0.515111 | 0.967 | 0.937 | 2.09E-05 |
| Rps28 | 9.58E-10 | 1.347421 | 0.888 | 0.885 | 2.26E-05 |
| Tnfrsf21 | 9.77E-10 | -1.70481 | 0.386 | 0.524 | 2.3E-05 |
| Gm3362 | 9.89E-10 | 1.04114 | 0.498 | 0.308 | 2.33E-05 |
| Glul | 1.55E-09 | -0.52832 | 0.914 | 0.931 | 3.67E-05 |
| Hnrnpk | 1.55E-09 | -0.98614 | 0.821 | 0.818 | 3.67E-05 |
| Tram1 | 1.67E-09 | -1.54533 | 0.547 | 0.643 | 3.94E-05 |
| Elmo1 | 1.75E-09 | -1.23548 | 0.591 | 0.712 | 4.13E-05 |
| Fth1 | 1.78E-09 | 0.494624 | 0.993 | 0.997 | 4.21E-05 |
| Asph | 2.09E-09 | -2.03196 | 0.484 | 0.579 | 4.94E-05 |
| Rps9 | 2.73E-09 | 0.71714 | 0.937 | 0.911 | 6.44E-05 |

| | | | | | |
|---|---|---|---|---|---|
| Calm1 | 2.78E-09 | 0.959201 | 0.765 | 0.671 | 6.56E-05 |
| Gns | 2.8E-09 | -0.84973 | 0.893 | 0.916 | 6.59E-05 |
| Adgre1 | 2.9E-09 | -1.84189 | 0.57 | 0.648 | 6.84E-05 |
| Daglb | 3.47E-09 | -1.30805 | 0.695 | 0.712 | 8.17E-05 |
| Malat1 | 4.98E-09 | 0.404459 | 0.974 | 0.98 | 0.000117 |
| Hspa8 | 5.25E-09 | -0.60031 | 0.937 | 0.925 | 0.000124 |
| Akr1a1 | 6.08E-09 | 0.812743 | 0.767 | 0.628 | 0.000143 |
| Cdc42 | 6.22E-09 | -0.7867 | 0.886 | 0.882 | 0.000147 |
| Rps18 | 7.36E-09 | 0.866952 | 0.812 | 0.772 | 0.000174 |
| Rps3 | 8.21E-09 | 0.800702 | 0.879 | 0.807 | 0.000194 |
| Rps19 | 8.46E-09 | 0.918145 | 0.844 | 0.787 | 0.0002 |
| Trf | 8.65E-09 | -0.44814 | 0.967 | 0.951 | 0.000204 |
| Slco2b1 | 9.43E-09 | -0.73243 | 0.902 | 0.911 | 0.000222 |
| Rac1 | 1.2E-08 | -0.70519 | 0.884 | 0.882 | 0.000283 |
| Gm28439 | 1.57E-08 | -0.4839 | 0.272 | 0.447 | 0.00037 |
| Gm5805 | 1.58E-08 | 1.151495 | 0.588 | 0.47 | 0.000373 |
| Rpsa | 1.68E-08 | 0.901343 | 0.77 | 0.666 | 0.000397 |
| Gm10443 | 1.77E-08 | 0.749686 | 0.526 | 0.383 | 0.000418 |
| Gm24270 | 1.85E-08 | -0.90454 | 0.956 | 0.977 | 0.000437 |
| Rpl34 | 2.37E-08 | 1.371468 | 0.744 | 0.744 | 0.00056 |
| Rn7sk | 2.39E-08 | -0.38496 | 0.735 | 0.856 | 0.000564 |
| Soga1 | 2.4E-08 | -0.99782 | 0.544 | 0.669 | 0.000565 |
| Ubb | 2.91E-08 | 0.437033 | 0.97 | 0.968 | 0.000687 |
| Nptn | 3.02E-08 | -1.90388 | 0.465 | 0.545 | 0.000712 |
| Mpeg1 | 3.96E-08 | -0.66625 | 0.751 | 0.873 | 0.000935 |
| Skil | 4.33E-08 | 0.462309 | 0.707 | 0.545 | 0.001021 |
| Rpl35 | 4.91E-08 | 1.148765 | 0.802 | 0.772 | 0.001159 |
| Gm12918 | 5.01E-08 | 0.538596 | 0.7 | 0.565 | 0.001183 |
| Rnf13 | 5.78E-08 | -1.35429 | 0.719 | 0.718 | 0.001363 |
| Ccr5 | 6.12E-08 | -1.72571 | 0.609 | 0.663 | 0.001444 |
| Gm15564 | 6.38E-08 | -1.06983 | 0.628 | 0.85 | 0.001504 |
| Tmed7 | 6.45E-08 | -0.72859 | 0.358 | 0.527 | 0.001521 |
| Entpd1 | 7.01E-08 | -0.63514 | 0.947 | 0.951 | 0.001654 |
| Il10ra | 7.26E-08 | -1.35183 | 0.667 | 0.741 | 0.001713 |
| Gm24245 | 1.01E-07 | -0.84643 | 0.367 | 0.522 | 0.002389 |
| Vcp-rs | 1.04E-07 | -0.63965 | 0.214 | 0.369 | 0.002444 |
| Mir692-2 | 1.05E-07 | 0.464718 | 0.519 | 0.357 | 0.002477 |
| Sparc | 1.06E-07 | 0.394658 | 0.998 | 1 | 0.002492 |
| Fcgr3 | 1.17E-07 | 0.430975 | 0.967 | 0.948 | 0.00275 |
| Ly86 | 1.25E-07 | 0.499172 | 0.979 | 0.968 | 0.002953 |
| Rpl23a-ps3 | 1.26E-07 | 0.776137 | 0.598 | 0.455 | 0.002964 |
| Nfkbia | 1.26E-07 | 1.782624 | 0.412 | 0.242 | 0.00298 |
| Ubc | 1.33E-07 | -0.39628 | 0.965 | 0.963 | 0.003134 |
| Gm12435 | 1.33E-07 | 0.786654 | 0.163 | 0.314 | 0.003146 |
| Pik3ap1 | 1.35E-07 | -2.09396 | 0.153 | 0.291 | 0.003176 |
| Msn | 1.36E-07 | -1.02652 | 0.563 | 0.648 | 0.003208 |
| Fau | 1.37E-07 | 1.16295 | 0.867 | 0.865 | 0.003228 |

| | | | | | |
|---|---|---|---|---|---|
| Rplp0 | 1.44E-07 | 0.635042 | 0.872 | 0.813 | 0.003403 |
| Atp5g2 | 1.58E-07 | 0.996389 | 0.558 | 0.409 | 0.003728 |
| Rps14 | 1.61E-07 | 0.895616 | 0.898 | 0.89 | 0.003792 |
| Gm6421 | 1.65E-07 | 0.624529 | 0.347 | 0.19 | 0.003898 |
| Gm10073 | 1.68E-07 | 0.620124 | 0.679 | 0.573 | 0.003952 |
| mt-Cytb | 1.7E-07 | 0.503582 | 0.951 | 0.939 | 0.004015 |
| Gm15427 | 1.82E-07 | 0.902065 | 0.956 | 0.911 | 0.004288 |
| Rpl23 | 1.86E-07 | 1.146422 | 0.788 | 0.767 | 0.004383 |
| Serpine2 | 1.95E-07 | -0.62615 | 0.895 | 0.879 | 0.004588 |
| Itgb2 | 2.09E-07 | -1.11932 | 0.753 | 0.758 | 0.004936 |
| Rps11 | 2.54E-07 | 0.794131 | 0.891 | 0.865 | 0.006 |
| Slc39a1 | 3.03E-07 | -0.32557 | 0.544 | 0.354 | 0.00715 |
| Mbnl1 | 3.26E-07 | -0.76865 | 0.712 | 0.827 | 0.007685 |
| Rpl36 | 3.8E-07 | 1.028213 | 0.802 | 0.769 | 0.00896 |
| Gm3699 | 3.89E-07 | 0.955872 | 0.458 | 0.308 | 0.009188 |
| Basp1 | 4.01E-07 | 0.363139 | 0.94 | 0.965 | 0.00945 |
| Cyba | 4.04E-07 | 0.483708 | 0.967 | 0.96 | 0.009523 |
| Csf3r | 4.16E-07 | -0.78249 | 0.816 | 0.882 | 0.009804 |
| Cd68 | 4.17E-07 | 0.50689 | 0.944 | 0.939 | 0.009839 |
| Arsb | 5.01E-07 | -1.4298 | 0.36 | 0.484 | 0.011827 |
| Orai2 | 5.04E-07 | -1.34151 | 0.147 | 0.285 | 0.011889 |
| Rps3a1 | 6.05E-07 | 0.805232 | 0.821 | 0.752 | 0.014264 |
| Lamp2 | 6.07E-07 | -0.85776 | 0.679 | 0.764 | 0.014328 |
| Rhoa | 6.46E-07 | -0.75974 | 0.826 | 0.807 | 0.015237 |
| Sema4d | 7.28E-07 | -1.2504 | 0.463 | 0.576 | 0.017175 |
| Gm43028 | 7.35E-07 | 0.38549 | 0.447 | 0.294 | 0.01733 |
| Gm3608 | 7.57E-07 | -0.59136 | 0.188 | 0.331 | 0.017862 |
| Rps23 | 8.56E-07 | 0.593641 | 0.907 | 0.867 | 0.020183 |
| Man1c1 | 9.04E-07 | -1.67755 | 0.46 | 0.542 | 0.021315 |
| Smap2 | 9.33E-07 | -0.52907 | 0.814 | 0.853 | 0.022007 |
| Itpripl1 | 9.36E-07 | 0.985547 | 0.677 | 0.55 | 0.022087 |
| Vav1 | 9.57E-07 | -1.43139 | 0.553 | 0.622 | 0.022564 |
| Rps21 | 9.78E-07 | 1.124505 | 0.784 | 0.761 | 0.023078 |
| Rpl10-ps3 | 1.01E-06 | 1.291973 | 0.66 | 0.55 | 0.02385 |
| Gm5138 | 1.09E-06 | 0.54203 | 0.665 | 0.501 | 0.025649 |
| Ctsz | 1.14E-06 | 0.361012 | 0.988 | 0.991 | 0.026844 |
| Oaz1 | 1.18E-06 | 0.810907 | 0.786 | 0.7 | 0.027806 |
| Tmem176b | 1.2E-06 | 0.720304 | 0.793 | 0.72 | 0.028347 |
| Man2b2 | 1.22E-06 | -1.51717 | 0.377 | 0.478 | 0.028745 |
| Gm8566 | 1.37E-06 | 0.361385 | 0.223 | 0.098 | 0.032382 |
| Slc25a5 | 1.54E-06 | 0.670247 | 0.823 | 0.755 | 0.036399 |
| Cd81 | 1.57E-06 | 0.264576 | 1 | 0.994 | 0.037027 |
| Ifngr1 | 1.6E-06 | -0.44924 | 0.942 | 0.928 | 0.037777 |
| Dusp6 | 1.68E-06 | -1.36927 | 0.656 | 0.663 | 0.039668 |
| Gm7964 | 1.75E-06 | -0.73926 | 0.591 | 0.651 | 0.041309 |
| Ftl2-ps | 1.76E-06 | 0.533077 | 0.947 | 0.934 | 0.041488 |
| Gm8960 | 1.92E-06 | 0.388908 | 0.391 | 0.245 | 0.04539 |

| | | | | | |
|---|---|---|---|---|---|
| Ncoa4 | 1.94E-06 | -1.20272 | 0.13 | 0.254 | 0.045817 |
| Hspa5 | 2.08E-06 | -0.87074 | 0.767 | 0.761 | 0.049176 |

Supplementry Table 9.  PB-Young DEGs in Hippocampus

| Gene | p_val | avg_log2FC | pct.1 | pct.2 | p_val_adj |
|---|---|---|---|---|---|
| Ct010467.1 | 5.5E-126 | 1.299782 | 1 | 1 | 1.3E-121 |
| Lars2 | 2.9E-125 | -2.37824 | 1 | 1 | 6.8E-121 |
| Gm24187 | 9.9E-116 | -2.08913 | 0.989 | 1 | 2.3E-111 |
| Gm23935 | 4.8E-115 | -2.06807 | 1 | 1 | 1.1E-110 |
| Gm24270 | 4.8E-110 | -1.86047 | 0.968 | 1 | 1.1E-105 |
| C1qc | 2.15E-92 | 1.162736 | 1 | 1 | 5.08E-88 |
| Tmsb4x | 8.84E-91 | 1.664885 | 1 | 1 | 2.08E-86 |
| Gm24245 | 5.32E-89 | -1.78728 | 0.465 | 0.96 | 1.26E-84 |
| Cst3 | 1.53E-78 | 1.122235 | 1 | 1 | 3.61E-74 |
| C1qa | 4.17E-73 | 0.793758 | 1 | 1 | 9.85E-69 |
| Itm2b | 1.31E-72 | 0.967487 | 1 | 1 | 3.1E-68 |
| Hexb | 6.52E-72 | 0.727319 | 1 | 1 | 1.54E-67 |
| Gm25911 | 1.32E-69 | -1.50371 | 0.511 | 0.933 | 3.12E-65 |
| Actb | 5.46E-67 | 1.046709 | 1 | 1 | 1.29E-62 |
| Selplg | 3.56E-65 | 0.954051 | 1 | 0.997 | 8.4E-61 |
| C1qb | 1.71E-64 | 0.922241 | 1 | 1 | 4.04E-60 |
| Gm15564 | 1.17E-63 | -1.316 | 0.812 | 0.988 | 2.77E-59 |
| Cd81 | 4.79E-62 | 1.068029 | 0.998 | 1 | 1.13E-57 |
| Sparc | 2.4E-59 | 1.158372 | 1 | 0.997 | 5.67E-55 |
| Ubb | 2.22E-57 | 1.617223 | 0.959 | 0.96 | 5.23E-53 |
| Csf1r | 3.68E-54 | -0.7926 | 1 | 1 | 8.69E-50 |
| Olfml3 | 1.42E-49 | 1.220177 | 0.996 | 0.994 | 3.34E-45 |
| Malat1 | 1.9E-49 | 0.971401 | 0.994 | 0.991 | 4.47E-45 |
| Ppia | 1.39E-48 | 1.511473 | 0.97 | 0.945 | 3.28E-44 |
| Fcer1g | 1.67E-48 | 1.225214 | 0.987 | 0.997 | 3.94E-44 |
| Rn7sk | 1.25E-47 | -1.30897 | 0.857 | 0.982 | 2.96E-43 |
| Tmem119 | 3.35E-47 | 0.82877 | 1 | 0.994 | 7.9E-43 |
| P2ry12 | 3.8E-43 | 0.804563 | 0.998 | 0.994 | 8.95E-39 |
| Tyrobp | 4.43E-43 | 1.129483 | 0.994 | 1 | 1.05E-38 |
| Cd9 | 2.17E-41 | 1.493138 | 0.963 | 0.93 | 5.11E-37 |
| Trem2 | 2.6E-41 | 1.038518 | 0.983 | 0.988 | 6.12E-37 |
| Fos | 5.83E-41 | 1.946808 | 0.476 | 0.034 | 1.38E-36 |
| Laptm5 | 1.29E-40 | 0.650048 | 1 | 1 | 3.04E-36 |
| Gm1821 | 4.36E-36 | 1.189085 | 0.946 | 0.936 | 1.03E-31 |
| Siglech | 1.47E-35 | 0.869841 | 1 | 1 | 3.48E-31 |
| Pfn1 | 7.48E-35 | 1.474532 | 0.933 | 0.933 | 1.76E-30 |
| Serinc3 | 1.47E-34 | -0.74 | 1 | 1 | 3.46E-30 |
| Rnase4 | 9.8E-34 | 1.309589 | 0.957 | 0.96 | 2.31E-29 |
| Gm8797 | 6.08E-32 | 1.077132 | 0.853 | 0.652 | 1.43E-27 |
| Rnaset2a | 4.45E-31 | 1.346781 | 0.948 | 0.948 | 1.05E-26 |
| Calm2 | 7.76E-31 | 1.595656 | 0.935 | 0.893 | 1.83E-26 |
| Ly86 | 4.16E-30 | 1.028056 | 0.981 | 0.997 | 9.81E-26 |
| Ctss | 1.81E-29 | 0.594026 | 1 | 1 | 4.26E-25 |
| Cfl1 | 2.18E-29 | 1.046086 | 0.961 | 0.939 | 5.13E-25 |

| | | | | | |
|---|---|---|---|---|---|
| Rgs10 | 2.68E-29 | 1.278716 | 0.944 | 0.97 | 6.31E-25 |
| Gsn | 1.09E-28 | 0.788636 | 0.392 | 0.043 | 2.56E-24 |
| Dusp1 | 1.84E-27 | 2.205597 | 0.335 | 0.018 | 4.34E-23 |
| Sft2d1 | 5.41E-27 | 1.354761 | 0.881 | 0.884 | 1.28E-22 |
| Ftl1 | 6.97E-27 | 1.003197 | 0.987 | 0.997 | 1.64E-22 |
| Slc2a5 | 1.14E-26 | 1.055554 | 0.937 | 0.918 | 2.7E-22 |
| Lgals3bp | 2.53E-26 | -1.10074 | 0.478 | 0.851 | 5.97E-22 |
| Rps29 | 1.18E-25 | 1.321537 | 0.981 | 1 | 2.79E-21 |
| Man2b1 | 2.03E-24 | -0.87216 | 0.916 | 0.988 | 4.8E-20 |
| Aif1 | 4.89E-24 | 1.335937 | 0.885 | 0.924 | 1.15E-19 |
| Mpeg1 | 7.65E-24 | -0.99799 | 0.853 | 0.966 | 1.8E-19 |
| Gm10123 | 1.05E-22 | 0.903437 | 0.939 | 0.902 | 2.47E-18 |
| Basp1 | 2.31E-22 | 1.094328 | 0.974 | 0.985 | 5.44E-18 |
| Dcn | 8.02E-22 | 2.18231 | 0.31 | 0.04 | 1.89E-17 |
| Gapdh | 1.15E-21 | 1.161486 | 0.909 | 0.899 | 2.71E-17 |
| Ctsd | 1.97E-21 | 0.44847 | 1 | 1 | 4.66E-17 |
| Ckb | 2.57E-21 | 1.146702 | 0.866 | 0.802 | 6.06E-17 |
| Abhd12 | 1.12E-19 | 0.810728 | 0.97 | 0.988 | 2.63E-15 |
| Cmtm6 | 2.43E-19 | -1.09192 | 0.781 | 0.887 | 5.72E-15 |
| Lgmn | 2.52E-19 | 0.41118 | 1 | 1 | 5.95E-15 |
| Fcrls | 2.89E-19 | 0.886813 | 0.968 | 0.97 | 6.81E-15 |
| Gm44860 | 3.46E-19 | -1.50687 | 0.288 | 0.576 | 8.16E-15 |
| Gm12166 | 4.61E-19 | 1.258497 | 0.816 | 0.802 | 1.09E-14 |
| Tmem59 | 6.05E-19 | 1.271221 | 0.851 | 0.838 | 1.43E-14 |
| Sirpa | 7.38E-19 | -0.74343 | 0.931 | 0.985 | 1.74E-14 |
| Fcgr3 | 1.26E-18 | 1.024423 | 0.955 | 0.982 | 2.98E-14 |
| Rps27 | 2.23E-18 | 1.255771 | 0.944 | 0.982 | 5.25E-14 |
| AU020206 | 2.74E-18 | -0.45454 | 0.541 | 0.799 | 6.46E-14 |
| Egr1 | 4.49E-18 | -0.84687 | 0.268 | 0.03 | 1.06E-13 |
| Rnaset2b | 6.84E-18 | 1.208822 | 0.935 | 0.933 | 1.61E-13 |
| Eef1a1 | 7.04E-18 | 0.659071 | 0.994 | 1 | 1.66E-13 |
| Pros1 | 1.2E-17 | -1.0621 | 0.671 | 0.826 | 2.84E-13 |
| mt-Cytb | 2.37E-17 | 1.00477 | 0.959 | 0.982 | 5.58E-13 |
| 1700017B0 | 4.94E-17 | -1.09701 | 0.591 | 0.793 | 1.17E-12 |
| Vsir | 7.89E-17 | 0.657964 | 0.994 | 0.994 | 1.86E-12 |
| Rnu3a | 1.16E-16 | -0.67191 | 0.052 | 0.262 | 2.73E-12 |
| Tmem50a | 1.51E-16 | 1.35729 | 0.84 | 0.848 | 3.56E-12 |
| Clic1 | 1.73E-16 | 1.296083 | 0.812 | 0.814 | 4.07E-12 |
| Vim | 3.38E-16 | 0.463848 | 0.238 | 0.027 | 7.98E-12 |
| Ikzf1 | 4.61E-16 | -0.51689 | 0.511 | 0.768 | 1.09E-11 |
| Nr4a1 | 6.06E-16 | 3.148357 | 0.216 | 0.018 | 1.43E-11 |
| Rps28 | 8.46E-16 | 1.263942 | 0.874 | 0.93 | 1.99E-11 |
| H2-Q6 | 1.06E-15 | -2.60631 | 0.052 | 0.244 | 2.49E-11 |
| mt-Atp6 | 1.27E-15 | 1.064866 | 0.844 | 0.787 | 2.99E-11 |
| Ywhah | 2.28E-15 | 0.95146 | 0.905 | 0.905 | 5.37E-11 |
| Ctsz | 3.02E-15 | 0.666954 | 0.983 | 1 | 7.12E-11 |
| Cd68 | 3.08E-15 | 0.876221 | 0.937 | 0.979 | 7.28E-11 |

| | | | | | |
|---|---|---|---|---|---|
| Cyba | 3.59E-15 | 0.8191 | 0.965 | 0.982 | 8.46E-11 |
| Rpl37a | 5.78E-15 | 1.001603 | 0.916 | 0.976 | 1.36E-10 |
| Oaz1 | 5.8E-15 | 1.524997 | 0.758 | 0.72 | 1.37E-10 |
| Syngr2 | 8.4E-15 | 1.164133 | 0.814 | 0.805 | 1.98E-10 |
| Stat1 | 9.44E-15 | -1.28932 | 0.184 | 0.43 | 2.23E-10 |
| Gnai2 | 1.65E-14 | 1.043156 | 0.944 | 0.982 | 3.9E-10 |
| Gm17494 | 1.74E-14 | 1.019478 | 0.385 | 0.146 | 4.11E-10 |
| Gm42418 | 2.02E-14 | -0.48971 | 0.857 | 0.97 | 4.77E-10 |
| Lfng | 3.22E-14 | -0.93575 | 0.225 | 0.445 | 7.61E-10 |
| Itgb5 | 3.25E-14 | -0.49366 | 0.987 | 0.994 | 7.67E-10 |
| Cd53 | 5.32E-14 | 0.774409 | 0.952 | 0.957 | 1.26E-09 |
| Gm13341 | 6.28E-14 | 0.707286 | 0.424 | 0.201 | 1.48E-09 |
| Hexa | 9.77E-14 | 0.675217 | 0.965 | 0.991 | 2.3E-09 |
| Spire1 | 1.18E-13 | -0.76125 | 0.292 | 0.543 | 2.77E-09 |
| Rpl13 | 1.34E-13 | 0.970463 | 0.922 | 0.979 | 3.17E-09 |
| Ltc4s | 2.33E-13 | 1.175715 | 0.842 | 0.857 | 5.49E-09 |
| mt-Nd1 | 2.35E-13 | 0.955147 | 0.939 | 0.933 | 5.54E-09 |
| Actg1 | 4.18E-13 | 1.088022 | 0.844 | 0.823 | 9.86E-09 |
| B2m | 4.42E-13 | 0.503528 | 1 | 1 | 1.04E-08 |
| Gm8464 | 5.28E-13 | 0.543652 | 0.387 | 0.165 | 1.25E-08 |
| Rplp0 | 5.57E-13 | 1.164575 | 0.877 | 0.927 | 1.31E-08 |
| Gm28437 | 5.65E-13 | 0.899505 | 0.807 | 0.759 | 1.33E-08 |
| Gm10925 | 5.74E-13 | 0.839986 | 0.807 | 0.72 | 1.35E-08 |
| Capzb | 5.81E-13 | 1.274887 | 0.781 | 0.735 | 1.37E-08 |
| Bsg | 6.52E-13 | 1.031024 | 0.835 | 0.86 | 1.54E-08 |
| Cnbp | 6.53E-13 | 1.420241 | 0.751 | 0.646 | 1.54E-08 |
| Gm28661 | 6.75E-13 | 0.964518 | 0.732 | 0.665 | 1.59E-08 |
| Alox5ap | 1.08E-12 | 1.407156 | 0.762 | 0.784 | 2.55E-08 |
| Fmnl1 | 1.2E-12 | -0.4998 | 0.335 | 0.588 | 2.84E-08 |
| mt-Co3 | 1.27E-12 | 0.949154 | 0.913 | 0.909 | 2.99E-08 |
| Laptm4a | 1.33E-12 | 0.978208 | 0.881 | 0.884 | 3.14E-08 |
| Fmnl3 | 1.5E-12 | -0.43895 | 0.513 | 0.747 | 3.54E-08 |
| H2-K1 | 1.88E-12 | -0.52457 | 0.738 | 0.924 | 4.42E-08 |
| Hsp90ab1 | 2.61E-12 | 1.052457 | 0.868 | 0.817 | 6.15E-08 |
| Fth1 | 3.47E-12 | 0.579195 | 0.989 | 0.997 | 8.19E-08 |
| Rps9 | 4.23E-12 | 0.859398 | 0.924 | 0.985 | 9.97E-08 |
| Hmgb1 | 5.97E-12 | 1.439347 | 0.76 | 0.68 | 1.41E-07 |
| mt-Co1 | 6.63E-12 | 0.555681 | 0.972 | 0.991 | 1.56E-07 |
| Ppp4r1 | 1.03E-11 | -0.68614 | 0.162 | 0.384 | 2.44E-07 |
| Gm5148 | 1.23E-11 | -0.91702 | 0.545 | 0.71 | 2.9E-07 |
| Ftl2-ps | 1.27E-11 | 0.746097 | 0.905 | 0.96 | 3E-07 |
| Sun2 | 1.37E-11 | -0.71418 | 0.6 | 0.784 | 3.24E-07 |
| Il10ra | 1.59E-11 | -0.76131 | 0.665 | 0.838 | 3.75E-07 |
| Gm7308 | 1.6E-11 | 0.527091 | 0.301 | 0.107 | 3.79E-07 |
| Etnk1 | 1.83E-11 | -0.53621 | 0.355 | 0.588 | 4.31E-07 |
| Sema4d | 2E-11 | -0.84643 | 0.506 | 0.707 | 4.73E-07 |
| H3f3b | 2.07E-11 | 1.02799 | 0.844 | 0.829 | 4.89E-07 |

| | | | | | |
|---|---|---|---|---|---|
| mt-Rnr2 | 2.34E-11 | 0.563125 | 0.991 | 0.994 | 5.53E-07 |
| Rplp1 | 2.42E-11 | 0.84976 | 0.929 | 0.973 | 5.7E-07 |
| Lyn | 2.83E-11 | -0.61807 | 0.807 | 0.921 | 6.67E-07 |
| Sh3bgrl3 | 2.91E-11 | 1.243833 | 0.777 | 0.805 | 6.86E-07 |
| Lrch1 | 2.91E-11 | -0.36375 | 0.312 | 0.567 | 6.87E-07 |
| Pdia3 | 3.55E-11 | 0.663421 | 0.955 | 0.954 | 8.37E-07 |
| Ctsh | 4.01E-11 | 0.747499 | 0.929 | 0.966 | 9.46E-07 |
| mt-Atp8 | 4.25E-11 | 0.436094 | 0.297 | 0.107 | 1E-06 |
| Adgre1 | 4.4E-11 | -0.8626 | 0.595 | 0.759 | 1.04E-06 |
| Rpl41 | 4.9E-11 | 0.998563 | 0.909 | 0.976 | 1.15E-06 |
| Syngr1 | 4.9E-11 | 1.056801 | 0.82 | 0.86 | 1.15E-06 |
| Itgb2 | 5.19E-11 | -0.63936 | 0.751 | 0.86 | 1.22E-06 |
| Ganab | 5.66E-11 | -1.48995 | 0.152 | 0.345 | 1.34E-06 |
| Rpl18a | 5.75E-11 | 0.832597 | 0.985 | 0.997 | 1.36E-06 |
| Bin1 | 6.19E-11 | 0.935194 | 0.898 | 0.954 | 1.46E-06 |
| Ftl1-ps1 | 7.02E-11 | 0.772287 | 0.9 | 0.872 | 1.66E-06 |
| Serpine2 | 7.32E-11 | 0.752725 | 0.903 | 0.89 | 1.73E-06 |
| Comt | 8.28E-11 | 1.376287 | 0.738 | 0.704 | 1.95E-06 |
| Crybb1 | 8.38E-11 | 1.237475 | 0.779 | 0.744 | 1.98E-06 |
| Asah1 | 9.01E-11 | -0.43751 | 0.762 | 0.884 | 2.13E-06 |
| Grk3 | 9.46E-11 | -0.88356 | 0.091 | 0.265 | 2.23E-06 |
| Rpl37 | 1.11E-10 | 1.013624 | 0.864 | 0.936 | 2.63E-06 |
| Gm37306 | 1.2E-10 | -0.9943 | 0.084 | 0.256 | 2.82E-06 |
| Tlr2 | 1.32E-10 | -1.61803 | 0.299 | 0.479 | 3.12E-06 |
| Slc25a5 | 1.46E-10 | 1.427416 | 0.766 | 0.793 | 3.44E-06 |
| Mdm4 | 2.23E-10 | -0.9234 | 0.318 | 0.534 | 5.27E-06 |
| Hsp90b1 | 2.9E-10 | 0.733596 | 0.937 | 0.939 | 6.85E-06 |
| Gm5560 | 4.63E-10 | 0.633868 | 0.452 | 0.265 | 1.09E-05 |
| Ifi27l2a | 5.49E-10 | -2.05704 | 0.08 | 0.232 | 1.3E-05 |
| Gm6421 | 5.89E-10 | 0.958751 | 0.318 | 0.14 | 1.39E-05 |
| Snx18 | 6.96E-10 | -0.57343 | 0.604 | 0.762 | 1.64E-05 |
| Gns | 7.03E-10 | -0.41733 | 0.924 | 0.979 | 1.66E-05 |
| Ttyh3 | 7.91E-10 | -0.5319 | 0.29 | 0.497 | 1.87E-05 |
| Ctsa | 8.95E-10 | 0.7179 | 0.944 | 0.973 | 2.11E-05 |
| Rps11 | 9.01E-10 | 0.943328 | 0.861 | 0.933 | 2.13E-05 |
| Gm13453 | 9.17E-10 | 0.694837 | 0.344 | 0.165 | 2.16E-05 |
| Map3k1 | 1E-09 | -0.86178 | 0.307 | 0.491 | 2.37E-05 |
| Rtp4 | 1.12E-09 | -0.81526 | 0.095 | 0.262 | 2.63E-05 |
| mt-Co2 | 1.19E-09 | 1.089923 | 0.829 | 0.851 | 2.81E-05 |
| Tpt1 | 1.27E-09 | 0.863956 | 0.872 | 0.96 | 2.99E-05 |
| Ldhb | 1.28E-09 | 1.271253 | 0.766 | 0.835 | 3.02E-05 |
| Fam91a1 | 1.28E-09 | -0.76548 | 0.065 | 0.21 | 3.02E-05 |
| Zswim8 | 1.4E-09 | -0.43645 | 0.201 | 0.393 | 3.3E-05 |
| Gng10 | 1.43E-09 | 1.087365 | 0.708 | 0.72 | 3.37E-05 |
| Tnfrsf21 | 1.45E-09 | -1.072 | 0.435 | 0.601 | 3.42E-05 |
| Ccr5 | 1.76E-09 | -0.81566 | 0.684 | 0.802 | 4.16E-05 |
| Adam17 | 1.97E-09 | -0.48106 | 0.405 | 0.616 | 4.64E-05 |

| | | | | | |
|---|---|---|---|---|---|
| Sall3 | 2.04E-09 | -0.4633 | 0.474 | 0.68 | 4.81E-05 |
| Rtn4rl1 | 2.2E-09 | -0.66615 | 0.686 | 0.826 | 5.19E-05 |
| Capn3 | 2.4E-09 | -0.49046 | 0.355 | 0.582 | 5.67E-05 |
| Mir99ahg | 2.55E-09 | -0.67208 | 0.344 | 0.564 | 6.02E-05 |
| Bin2 | 2.83E-09 | 0.850001 | 0.866 | 0.881 | 6.69E-05 |
| Fnbp1 | 3.14E-09 | -0.52032 | 0.426 | 0.631 | 7.4E-05 |
| Ubc | 3.25E-09 | 0.587407 | 0.955 | 0.96 | 7.67E-05 |
| Rpl30 | 3.46E-09 | 1.156609 | 0.835 | 0.918 | 8.16E-05 |
| Leng8 | 3.5E-09 | -0.31766 | 0.377 | 0.604 | 8.26E-05 |
| Slfn2 | 3.64E-09 | -0.56156 | 0.108 | 0.28 | 8.59E-05 |
| Mertk | 4.14E-09 | -0.32384 | 0.823 | 0.97 | 9.76E-05 |
| Wasf2 | 4.3E-09 | 1.171444 | 0.864 | 0.878 | 0.000102 |
| Atf3 | 4.58E-09 | 1.472975 | 0.203 | 0.052 | 0.000108 |
| Spi1 | 4.68E-09 | 1.134173 | 0.82 | 0.887 | 0.00011 |
| Tlr13 | 4.93E-09 | -0.71932 | 0.288 | 0.476 | 0.000116 |
| Psap | 5.2E-09 | -0.26617 | 0.998 | 1 | 0.000123 |
| Rap1gds1 | 5.27E-09 | -0.56609 | 0.489 | 0.686 | 0.000124 |
| Rps18 | 5.77E-09 | 1.08982 | 0.81 | 0.893 | 0.000136 |
| Gm10736 | 6.14E-09 | 0.794821 | 0.695 | 0.625 | 0.000145 |
| Rplp2 | 6.19E-09 | 0.960842 | 0.883 | 0.954 | 0.000146 |
| Rhob | 6.41E-09 | -0.46081 | 0.913 | 0.933 | 0.000151 |
| H2-Q4 | 6.79E-09 | -1.4949 | 0.074 | 0.213 | 0.00016 |
| Pik3ap1 | 7.32E-09 | -1.25535 | 0.188 | 0.348 | 0.000173 |
| Tbc1d9b | 7.42E-09 | -0.71908 | 0.206 | 0.387 | 0.000175 |
| F11r | 7.76E-09 | 0.682359 | 0.961 | 0.973 | 0.000183 |
| E230029C0 | 7.91E-09 | -0.25992 | 0.617 | 0.841 | 0.000187 |
| Asph | 7.97E-09 | -0.65074 | 0.619 | 0.777 | 0.000188 |
| Rabac1 | 8.29E-09 | 1.310115 | 0.712 | 0.723 | 0.000196 |
| Atp6v1g1 | 8.3E-09 | 1.628287 | 0.593 | 0.521 | 0.000196 |
| Selenof | 9.09E-09 | 1.40231 | 0.632 | 0.57 | 0.000214 |
| Rps14 | 9.32E-09 | 0.955881 | 0.89 | 0.954 | 0.00022 |
| Ptma | 1.09E-08 | 1.712359 | 0.632 | 0.534 | 0.000258 |
| Serinc1 | 1.24E-08 | -0.65128 | 0.597 | 0.75 | 0.000293 |
| Gm37033 | 1.36E-08 | -0.71174 | 0.102 | 0.256 | 0.00032 |
| Gsap | 1.37E-08 | -0.40348 | 0.145 | 0.314 | 0.000323 |
| Gm4617 | 1.38E-08 | 0.759456 | 0.357 | 0.192 | 0.000326 |
| Rnf130 | 1.54E-08 | 0.912495 | 0.816 | 0.857 | 0.000363 |
| Spint1 | 1.84E-08 | -0.35345 | 0.392 | 0.622 | 0.000433 |
| Man2b2 | 1.89E-08 | -1.45686 | 0.398 | 0.537 | 0.000446 |
| Plcg2 | 2.02E-08 | -0.61654 | 0.368 | 0.573 | 0.000476 |
| Selenop | 2.04E-08 | 0.510943 | 0.985 | 0.985 | 0.00048 |
| Anxa3 | 2.3E-08 | 1.169244 | 0.788 | 0.817 | 0.000543 |
| Slc38a10 | 2.44E-08 | -1.49563 | 0.288 | 0.451 | 0.000575 |
| Atp6v0b | 2.83E-08 | 0.958376 | 0.851 | 0.912 | 0.000668 |
| Abi3 | 2.89E-08 | 0.893754 | 0.805 | 0.841 | 0.000682 |
| Washc4 | 2.91E-08 | -0.66967 | 0.394 | 0.591 | 0.000687 |
| Gm14303 | 3.82E-08 | 0.716982 | 0.89 | 0.899 | 0.000902 |

| | | | | | |
|---|---|---|---|---|---|
| Ppp1ca | 4.73E-08 | 1.458653 | 0.604 | 0.543 | 0.001116 |
| Camk1 | 5.47E-08 | 1.178385 | 0.766 | 0.796 | 0.001291 |
| Mknk1 | 5.56E-08 | -0.48377 | 0.42 | 0.616 | 0.001312 |
| Hps3 | 6.03E-08 | -0.65269 | 0.372 | 0.549 | 0.001422 |
| Cmtm7 | 6.22E-08 | 1.143592 | 0.79 | 0.841 | 0.001466 |
| Coro1a | 6.36E-08 | 0.893012 | 0.855 | 0.869 | 0.0015 |
| Vav1 | 6.97E-08 | -0.50234 | 0.621 | 0.777 | 0.001643 |
| H3f3c | 7.08E-08 | 0.461778 | 0.223 | 0.082 | 0.00167 |
| Pnpla7 | 7.18E-08 | -0.41289 | 0.506 | 0.701 | 0.001695 |
| Ecscr | 7.5E-08 | 1.407018 | 0.693 | 0.692 | 0.00177 |
| Myo1f | 8E-08 | -0.66435 | 0.47 | 0.652 | 0.001887 |
| Cebpa | 8.3E-08 | -0.97054 | 0.485 | 0.591 | 0.001957 |
| Son | 1.04E-07 | 0.768957 | 0.877 | 0.872 | 0.002448 |
| Rb1 | 1.08E-07 | -0.47346 | 0.084 | 0.22 | 0.002537 |
| Dock2 | 1.1E-07 | 0.607486 | 0.385 | 0.622 | 0.002586 |
| Susd3 | 1.11E-07 | 1.040322 | 0.788 | 0.857 | 0.00261 |
| Cdan1 | 1.14E-07 | -0.66729 | 0.171 | 0.335 | 0.002695 |
| Selenok | 1.15E-07 | 1.533376 | 0.589 | 0.54 | 0.002718 |
| Zdhhc9 | 1.33E-07 | -0.66972 | 0.229 | 0.399 | 0.003145 |
| Camkk2 | 1.35E-07 | -0.44898 | 0.253 | 0.439 | 0.003196 |
| Gm9234 | 1.36E-07 | 0.572967 | 0.578 | 0.424 | 0.003203 |
| Icosl | 1.47E-07 | -0.77997 | 0.31 | 0.473 | 0.003471 |
| Gm44861 | 1.59E-07 | -0.7487 | 0.093 | 0.229 | 0.003748 |
| Calm1 | 1.68E-07 | 1.354521 | 0.716 | 0.671 | 0.003965 |
| Sh2b3 | 1.69E-07 | -0.97248 | 0.431 | 0.585 | 0.003981 |
| Gm5529 | 1.69E-07 | 0.508929 | 0.4 | 0.244 | 0.003991 |
| Rbm14 | 1.7E-07 | -1.73832 | 0.087 | 0.213 | 0.004018 |
| Pycard | 1.73E-07 | 0.944659 | 0.799 | 0.823 | 0.004075 |
| Cdc42 | 1.79E-07 | 0.735007 | 0.881 | 0.86 | 0.004231 |
| Wnk1 | 1.83E-07 | -0.36264 | 0.768 | 0.924 | 0.004317 |
| Gm5905 | 2.26E-07 | -0.43996 | 0.773 | 0.875 | 0.005338 |
| Rps19 | 2.53E-07 | 0.943372 | 0.855 | 0.927 | 0.005965 |
| Arpc3 | 2.54E-07 | 1.543551 | 0.582 | 0.54 | 0.005995 |
| Mms19 | 2.81E-07 | -0.54273 | 0.193 | 0.348 | 0.006625 |
| Slc12a9 | 2.96E-07 | -0.28922 | 0.385 | 0.582 | 0.006982 |
| Arpc4 | 2.99E-07 | 1.31267 | 0.775 | 0.841 | 0.007051 |
| Gm6180 | 3.08E-07 | 0.482223 | 0.548 | 0.396 | 0.007257 |
| Ppp1r9b | 3.08E-07 | -0.60414 | 0.238 | 0.412 | 0.007263 |
| Myo9b | 3.41E-07 | -0.29401 | 0.21 | 0.384 | 0.00805 |
| Kbtbd2 | 3.43E-07 | -0.99759 | 0.203 | 0.36 | 0.008084 |
| Eif5a | 3.49E-07 | 1.506287 | 0.574 | 0.497 | 0.008222 |
| Tmem181a | 3.52E-07 | -0.64488 | 0.117 | 0.259 | 0.008315 |
| Arap1 | 3.57E-07 | -0.81303 | 0.32 | 0.491 | 0.00843 |
| H2-D1 | 3.57E-07 | -0.41347 | 0.87 | 0.951 | 0.00843 |
| Gm6063 | 3.65E-07 | 0.292128 | 0.271 | 0.125 | 0.008622 |
| Arhgef2 | 3.68E-07 | -0.45439 | 0.357 | 0.54 | 0.008684 |
| Gm12164 | 3.89E-07 | 0.258966 | 0.227 | 0.088 | 0.009186 |

| | | | | | |
|---|---|---|---|---|---|
| Adar | 3.97E-07 | -0.27841 | 0.188 | 0.351 | 0.009375 |
| Tcn2 | 4.19E-07 | 0.959106 | 0.842 | 0.854 | 0.009878 |
| Apba1 | 4.19E-07 | -0.55797 | 0.123 | 0.265 | 0.009879 |
| Slc30a9 | 4.27E-07 | -1.10106 | 0.134 | 0.277 | 0.010079 |
| Spcs2 | 5.12E-07 | 1.29246 | 0.623 | 0.573 | 0.012084 |
| Cd9-ps | 5.13E-07 | 0.258001 | 0.203 | 0.076 | 0.012106 |
| Ppib | 5.18E-07 | 0.989278 | 0.706 | 0.704 | 0.012225 |
| Trim34a | 5.24E-07 | -1.04583 | 0.082 | 0.207 | 0.012368 |
| H2-Q7 | 5.41E-07 | -1.80617 | 0.097 | 0.216 | 0.012767 |
| Prkcb | 5.73E-07 | -0.54345 | 0.359 | 0.546 | 0.013519 |
| Jun | 5.81E-07 | 0.343711 | 0.39 | 0.198 | 0.013713 |
| Tuba1a | 6.54E-07 | 1.381657 | 0.578 | 0.454 | 0.01543 |
| Ankrd10 | 6.62E-07 | 0.612276 | 0.374 | 0.607 | 0.015621 |
| Ctsl | 6.76E-07 | 0.574295 | 0.918 | 0.966 | 0.015941 |
| Gm34589 | 6.77E-07 | 0.265577 | 0.229 | 0.412 | 0.015975 |
| Gm14165 | 7.27E-07 | 0.46551 | 0.422 | 0.268 | 0.017153 |
| Rasal3 | 7.75E-07 | -0.66554 | 0.506 | 0.652 | 0.018272 |
| Pmp22 | 7.79E-07 | 0.957251 | 0.784 | 0.732 | 0.018374 |
| Rpl21 | 9.68E-07 | 0.944123 | 0.877 | 0.954 | 0.022843 |
| Eef2 | 9.96E-07 | -0.52742 | 0.747 | 0.854 | 0.023504 |
| Parp14 | 1.05E-06 | -0.58574 | 0.149 | 0.296 | 0.024686 |
| P3h2 | 1.1E-06 | -1.57639 | 0.152 | 0.284 | 0.025943 |
| Parp12 | 1.11E-06 | -1.80644 | 0.141 | 0.274 | 0.026234 |
| Ube3c | 1.16E-06 | -0.37538 | 0.169 | 0.32 | 0.027336 |
| Fyco1 | 1.16E-06 | 0.594493 | 0.182 | 0.354 | 0.027362 |
| Xaf1 | 1.23E-06 | 0.28173 | 0.141 | 0.287 | 0.029021 |
| Gm13408 | 1.26E-06 | 0.255536 | 0.262 | 0.122 | 0.029671 |
| Gpr34 | 1.29E-06 | 0.480862 | 0.987 | 0.985 | 0.030398 |
| Calr | 1.36E-06 | 0.614634 | 0.935 | 0.96 | 0.032128 |
| Arpc1b | 1.4E-06 | 1.346327 | 0.734 | 0.808 | 0.032963 |
| Lpcat2 | 1.41E-06 | 0.452137 | 0.97 | 0.976 | 0.033327 |
| Rassf2 | 1.43E-06 | -0.29853 | 0.348 | 0.524 | 0.033627 |
| Sipa1l2 | 1.5E-06 | -0.43904 | 0.288 | 0.454 | 0.035306 |
| Trim14 | 1.52E-06 | -0.62172 | 0.113 | 0.244 | 0.035766 |
| Abr | 1.6E-06 | -1.38925 | 0.145 | 0.28 | 0.037825 |
| Myo18a | 1.61E-06 | -0.68002 | 0.219 | 0.375 | 0.037892 |
| Lhfpl2 | 1.62E-06 | -0.34393 | 0.418 | 0.564 | 0.038129 |
| Gm5637 | 1.63E-06 | 0.790712 | 0.448 | 0.317 | 0.038462 |
| Gm12715 | 1.75E-06 | 0.716619 | 0.695 | 0.631 | 0.041244 |
| Nfat5 | 1.83E-06 | -0.60975 | 0.238 | 0.39 | 0.043242 |
| Naip2 | 1.84E-06 | -0.31635 | 0.277 | 0.439 | 0.043431 |
| Oaz1-ps | 1.86E-06 | 0.358031 | 0.639 | 0.555 | 0.043788 |
| Rpl18 | 1.92E-06 | 1.178375 | 0.745 | 0.829 | 0.045291 |
| Ac140300.2 | 1.97E-06 | 0.292559 | 0.284 | 0.143 | 0.046375 |

Supplementry Table 9.  PB-Young DEGs in Striatum

| Gene | p_val | avg_log2FC | pct.1 | pct.2 | p_val_adj |
|---|---|---|---|---|---|
| Gm26917 | 3.52E-82 | -2.61022 | 0.883 | 0.988 | 8.3E-78 |
| Ct010467.1 | 4.95E-66 | -0.75531 | 1 | 1 | 1.17E-61 |
| Gm15266 | 5.21E-61 | 1.57157 | 0.88 | 0.459 | 1.23E-56 |
| Dpysl2 | 2.14E-59 | 3.039226 | 0.9 | 0.792 | 5.04E-55 |
| Rps25 | 3.54E-57 | 1.907192 | 0.962 | 0.83 | 8.36E-53 |
| Gm24276 | 2.52E-56 | 1.219605 | 0.526 | 0.05 | 5.95E-52 |
| Arglu1 | 3.33E-53 | 2.581535 | 0.89 | 0.695 | 7.86E-49 |
| Adap2os | 3.58E-53 | 2.121933 | 0.928 | 0.838 | 8.45E-49 |
| Gm23935 | 3.25E-49 | -1.13313 | 1 | 1 | 7.68E-45 |
| Rps29 | 1.57E-47 | 1.073374 | 1 | 0.998 | 3.72E-43 |
| mt-Co1 | 2.54E-46 | 1.137339 | 0.99 | 0.966 | 5.99E-42 |
| Gm14303 | 1.11E-45 | 1.071895 | 0.962 | 0.747 | 2.61E-41 |
| Bmp2k | 1.54E-45 | 1.6445 | 0.966 | 0.78 | 3.63E-41 |
| Slc39a1 | 5.7E-45 | 1.457188 | 0.739 | 0.295 | 1.34E-40 |
| Lars2 | 2.97E-43 | -1.91451 | 0.997 | 1 | 7E-39 |
| Gm24270 | 9.58E-42 | -1.25925 | 0.876 | 0.996 | 2.26E-37 |
| Rn7sk | 1.91E-41 | -1.25009 | 0.491 | 0.918 | 4.5E-37 |
| Tmsb4x | 2.29E-41 | 0.704168 | 1 | 1 | 5.4E-37 |
| Gm2225 | 4.43E-41 | 0.949948 | 0.588 | 0.15 | 1.04E-36 |
| Gm13456 | 4.93E-41 | 1.165503 | 0.983 | 0.932 | 1.16E-36 |
| Actr2 | 2.78E-40 | 1.680933 | 0.938 | 0.822 | 6.55E-36 |
| Marcks | 3.02E-40 | 0.913456 | 1 | 0.996 | 7.13E-36 |
| mt-Co2 | 4.43E-39 | 1.389933 | 0.924 | 0.721 | 1.05E-34 |
| Rps27 | 1.26E-38 | 0.990236 | 0.986 | 0.94 | 2.97E-34 |
| Bhlhe41 | 6.36E-38 | 2.204569 | 0.832 | 0.595 | 1.5E-33 |
| Gm5559 | 6.51E-38 | 1.67121 | 0.852 | 0.647 | 1.54E-33 |
| Sod1 | 2.42E-37 | 2.71569 | 0.732 | 0.425 | 5.7E-33 |
| Gm4977 | 3.26E-37 | 0.79943 | 0.515 | 0.108 | 7.7E-33 |
| Skil | 1.68E-36 | 2.522185 | 0.808 | 0.601 | 3.97E-32 |
| Gm11518 | 2.25E-36 | 2.456642 | 0.687 | 0.377 | 5.3E-32 |
| Zfp691 | 4.06E-36 | 1.745626 | 0.766 | 0.409 | 9.57E-32 |
| Gm6988 | 5.92E-35 | 0.954711 | 0.515 | 0.128 | 1.4E-30 |
| Rps28 | 1.73E-34 | 1.107738 | 0.969 | 0.884 | 4.07E-30 |
| Abcg1 | 4.04E-33 | 2.617919 | 0.787 | 0.619 | 9.53E-29 |
| Zscan26 | 2.55E-32 | 3.140999 | 0.698 | 0.479 | 6.02E-28 |
| Rnaset2b | 6.9E-32 | 1.039085 | 0.979 | 0.928 | 1.63E-27 |
| Gm5138 | 1.46E-31 | 1.164769 | 0.708 | 0.355 | 3.44E-27 |
| Srsf2 | 1.14E-30 | 1.688541 | 0.866 | 0.776 | 2.68E-26 |
| Itpripl1 | 1.3E-30 | 2.996447 | 0.704 | 0.491 | 3.06E-26 |
| Kdelr2 | 2.38E-30 | 1.471029 | 0.718 | 0.343 | 5.61E-26 |
| Cbx3 | 2.5E-30 | 1.160024 | 0.667 | 0.303 | 5.89E-26 |
| Gm29667 | 1.44E-29 | 1.067154 | 0.598 | 0.24 | 3.39E-25 |
| Ncoa3 | 3.28E-29 | 1.928298 | 0.773 | 0.567 | 7.74E-25 |
| Gm13340 | 8.98E-29 | 0.936358 | 0.849 | 0.607 | 2.12E-24 |

| | | | | | |
|---|---|---|---|---|---|
| Gm11478 | 9.94E-29 | 1.189371 | 0.849 | 0.589 | 2.34E-24 |
| Canx | 1.04E-28 | 1.157567 | 0.955 | 0.888 | 2.45E-24 |
| B4galt1 | 2.2E-28 | 1.352555 | 0.876 | 0.687 | 5.19E-24 |
| Gm15564 | 3.6E-28 | -1.60786 | 0.56 | 0.866 | 8.49E-24 |
| Gm10800 | 6.51E-28 | 1.082056 | 0.787 | 0.82 | 1.54E-23 |
| Notch2 | 8.84E-28 | 0.423216 | 0.787 | 0.479 | 2.08E-23 |
| Ptafr | 1.2E-27 | 1.978659 | 0.777 | 0.625 | 2.84E-23 |
| Cdkn1b | 3.85E-27 | 1.524049 | 0.44 | 0.108 | 9.09E-23 |
| Gm8129 | 1.09E-26 | 1.358843 | 0.557 | 0.242 | 2.57E-22 |
| Gnb1 | 1.32E-26 | 1.57838 | 0.859 | 0.796 | 3.11E-22 |
| Gm15427 | 1.88E-26 | 0.742727 | 0.983 | 0.918 | 4.44E-22 |
| Eef1a1 | 3.49E-26 | 0.775317 | 1 | 0.996 | 8.22E-22 |
| Sept2 | 5.22E-26 | 2.045559 | 0.787 | 0.647 | 1.23E-21 |
| Gm6977 | 7.4E-26 | 1.074458 | 0.739 | 0.443 | 1.75E-21 |
| Ssr1 | 8.08E-26 | 2.329968 | 0.711 | 0.503 | 1.91E-21 |
| Gm8566 | 1.36E-25 | 0.753908 | 0.344 | 0.064 | 3.2E-21 |
| Cx3cr1 | 6.8E-25 | -0.70718 | 1 | 1 | 1.6E-20 |
| Gm5805 | 1.06E-24 | 0.982271 | 0.732 | 0.441 | 2.5E-20 |
| Wtap | 1.14E-24 | 0.985525 | 0.66 | 0.271 | 2.68E-20 |
| Dhrs3 | 1.7E-23 | 1.745944 | 0.828 | 0.77 | 4.01E-19 |
| Rpl37a | 2.69E-23 | 0.690973 | 0.979 | 0.938 | 6.34E-19 |
| Gm1821 | 1.06E-22 | 0.648484 | 0.973 | 0.914 | 2.5E-18 |
| Set | 1.06E-22 | 1.746705 | 0.739 | 0.483 | 2.5E-18 |
| Rps27rt | 1.57E-22 | 0.843285 | 0.873 | 0.643 | 3.69E-18 |
| Tubb5 | 1.83E-22 | 1.201204 | 0.869 | 0.651 | 4.31E-18 |
| Alkbh5 | 3.49E-22 | 0.860681 | 0.471 | 0.166 | 8.24E-18 |
| Rnaset2a | 4.06E-22 | 0.915062 | 0.969 | 0.92 | 9.57E-18 |
| Rps25-ps1 | 5.29E-22 | 0.705089 | 0.56 | 0.257 | 1.25E-17 |
| Gm9843 | 9.27E-22 | 1.284822 | 0.914 | 0.756 | 2.19E-17 |
| Ccdc85c | 1.58E-21 | 1.052712 | 0.265 | 0.036 | 3.73E-17 |
| Ftl2-ps | 1.6E-21 | 0.75799 | 0.973 | 0.91 | 3.77E-17 |
| Gm10076 | 2.24E-21 | 0.787799 | 0.955 | 0.878 | 5.28E-17 |
| Rps10-ps1 | 3.92E-21 | 0.809778 | 0.674 | 0.397 | 9.24E-17 |
| Gm10443 | 1.88E-20 | 0.735935 | 0.615 | 0.317 | 4.44E-16 |
| Rpl13a | 2.49E-20 | 0.923393 | 0.904 | 0.776 | 5.86E-16 |
| mt-Cytb | 6.8E-20 | 0.843929 | 0.976 | 0.93 | 1.6E-15 |
| mt-Co3 | 1.08E-19 | 0.82693 | 0.945 | 0.84 | 2.54E-15 |
| Qk | 1.87E-19 | 1.611429 | 0.918 | 0.91 | 4.41E-15 |
| Gm10045 | 2.09E-19 | 0.579854 | 0.333 | 0.086 | 4.93E-15 |
| Cst3 | 5.5E-19 | 0.358973 | 1 | 1 | 1.3E-14 |
| Ptgs1 | 6.11E-19 | 0.802527 | 0.973 | 0.876 | 1.44E-14 |
| Aldoart2 | 6.76E-19 | 0.481434 | 0.247 | 0.04 | 1.6E-14 |
| Tmf1 | 1.07E-18 | 1.257454 | 0.608 | 0.297 | 2.53E-14 |
| C5ar2 | 1.25E-18 | 1.958783 | 0.653 | 0.477 | 2.95E-14 |
| Gm29216 | 1.91E-18 | 0.721024 | 0.876 | 0.721 | 4.5E-14 |
| Sat1 | 2.42E-18 | 1.036275 | 0.9 | 0.828 | 5.71E-14 |
| Rps6 | 3.63E-18 | 0.924695 | 0.869 | 0.705 | 8.57E-14 |

| | | | | | |
|---|---|---|---|---|---|
| Wdr89 | 6.82E-18 | 1.64748 | 0.766 | 0.537 | 1.61E-13 |
| Adam10 | 7.15E-18 | 1.260875 | 0.821 | 0.661 | 1.69E-13 |
| Gm10134 | 8.02E-18 | 3.321819 | 0.471 | 0.244 | 1.89E-13 |
| Gm43305 | 1.2E-17 | 0.325134 | 0.746 | 0.559 | 2.82E-13 |
| Gm36985 | 1.36E-17 | 0.404438 | 0.258 | 0.052 | 3.22E-13 |
| Rpl13a-ps1 | 1.41E-17 | 0.592775 | 0.423 | 0.164 | 3.34E-13 |
| Rps19 | 1.74E-17 | 0.756269 | 0.921 | 0.796 | 4.11E-13 |
| Gm11450 | 2.46E-17 | 0.581568 | 0.265 | 0.056 | 5.81E-13 |
| mt-Atp8 | 2.77E-17 | 0.420548 | 0.33 | 0.092 | 6.54E-13 |
| Gm7536 | 3.15E-17 | 0.893076 | 0.759 | 0.519 | 7.42E-13 |
| Ptges3 | 3.47E-17 | 1.505754 | 0.66 | 0.411 | 8.19E-13 |
| Tmem181b | 3.83E-17 | -1.03668 | 0.392 | 0.12 | 9.05E-13 |
| Rpl17 | 6.23E-17 | 0.849492 | 0.9 | 0.695 | 1.47E-12 |
| Gm17066 | 1.12E-16 | 1.275204 | 0.577 | 0.363 | 2.64E-12 |
| Rps11 | 1.17E-16 | 0.68281 | 0.969 | 0.872 | 2.77E-12 |
| Lrrc8a | 1.22E-16 | 3.226941 | 0.478 | 0.273 | 2.87E-12 |
| Rpl37 | 1.81E-16 | 0.61019 | 0.959 | 0.884 | 4.27E-12 |
| Gm8730 | 2.3E-16 | 0.697396 | 0.856 | 0.661 | 5.43E-12 |
| Gm25911 | 2.4E-16 | -0.68905 | 0.309 | 0.597 | 5.66E-12 |
| Hnrnpk | 2.5E-16 | 0.942159 | 0.863 | 0.768 | 5.89E-12 |
| Gm28661 | 4.35E-16 | 0.719652 | 0.811 | 0.615 | 1.03E-11 |
| Gm3511 | 5.68E-16 | 0.630183 | 0.77 | 0.525 | 1.34E-11 |
| Rplp2 | 5.74E-16 | 0.708752 | 0.952 | 0.864 | 1.35E-11 |
| C1qb | 6.26E-16 | 0.297816 | 1 | 1 | 1.48E-11 |
| Gm8172 | 6.98E-16 | 1.133977 | 0.653 | 0.425 | 1.65E-11 |
| Rps21 | 7.57E-16 | 0.739648 | 0.89 | 0.737 | 1.78E-11 |
| Gm10059 | 7.58E-16 | 0.958834 | 0.553 | 0.279 | 1.79E-11 |
| Gm7964 | 7.86E-16 | 0.94754 | 0.784 | 0.595 | 1.85E-11 |
| Zfhx3 | 8.26E-16 | 1.039251 | 0.935 | 0.924 | 1.95E-11 |
| Marf1 | 1.31E-15 | 2.569502 | 0.546 | 0.375 | 3.08E-11 |
| Rpl7a | 1.32E-15 | 1.03679 | 0.79 | 0.601 | 3.1E-11 |
| Rpl41 | 1.32E-15 | 0.652935 | 0.966 | 0.912 | 3.12E-11 |
| Rps3 | 1.32E-15 | 0.724099 | 0.955 | 0.864 | 3.12E-11 |
| Gm12715 | 1.84E-15 | 0.858111 | 0.808 | 0.655 | 4.33E-11 |
| Gm42418 | 2.16E-15 | -0.8887 | 0.794 | 0.906 | 5.1E-11 |
| Upf1 | 2.42E-15 | -0.57827 | 0.515 | 0.222 | 5.72E-11 |
| Pdcd4 | 2.49E-15 | 1.994977 | 0.491 | 0.277 | 5.86E-11 |
| Gm3379 | 3.27E-15 | 0.835943 | 0.711 | 0.469 | 7.71E-11 |
| Rps16-ps2 | 3.5E-15 | 0.838366 | 0.739 | 0.503 | 8.25E-11 |
| Calm2 | 4.89E-15 | 0.72123 | 0.945 | 0.904 | 1.15E-10 |
| Gm8430 | 5.23E-15 | 0.865212 | 0.68 | 0.435 | 1.23E-10 |
| Gm6136 | 8.24E-15 | 0.926119 | 0.708 | 0.489 | 1.94E-10 |
| Ftl1-ps1 | 8.97E-15 | 0.589709 | 0.955 | 0.868 | 2.12E-10 |
| Rpl38 | 9.54E-15 | 1.003238 | 0.753 | 0.601 | 2.25E-10 |
| Rps16 | 1.14E-14 | 0.802482 | 0.866 | 0.699 | 2.68E-10 |
| mt-Nd2 | 1.18E-14 | 1.129154 | 0.907 | 0.834 | 2.77E-10 |
| Rpl34-ps1 | 1.28E-14 | 0.654987 | 0.698 | 0.425 | 3.03E-10 |

| | | | | | |
|---|---|---|---|---|---|
| Gm7079 | 1.31E-14 | 0.37102 | 0.265 | 0.07 | 3.08E-10 |
| Hsf2 | 1.43E-14 | 1.168389 | 0.247 | 0.058 | 3.37E-10 |
| Aif1 | 1.57E-14 | 0.70185 | 0.948 | 0.916 | 3.7E-10 |
| Rbm26 | 1.63E-14 | 2.03637 | 0.646 | 0.557 | 3.85E-10 |
| Cd164 | 1.81E-14 | -1.09408 | 0.893 | 0.918 | 4.28E-10 |
| Selenop | 1.91E-14 | 0.475257 | 0.997 | 0.992 | 4.52E-10 |
| Rpl27a | 1.96E-14 | 0.779508 | 0.835 | 0.665 | 4.64E-10 |
| Gm46522 | 1.98E-14 | 0.363906 | 0.21 | 0.04 | 4.67E-10 |
| Rpl30 | 2.21E-14 | 0.688062 | 0.931 | 0.814 | 5.21E-10 |
| Rpl36 | 2.49E-14 | 0.734031 | 0.876 | 0.794 | 5.88E-10 |
| Gm44951 | 2.53E-14 | -1.48524 | 0.354 | 0.603 | 5.96E-10 |
| Rpl27-ps3 | 2.99E-14 | 0.858694 | 0.526 | 0.275 | 7.06E-10 |
| Pan3 | 3.03E-14 | 1.458413 | 0.708 | 0.519 | 7.16E-10 |
| Rpl34 | 3.6E-14 | 0.694942 | 0.873 | 0.689 | 8.49E-10 |
| Rpl23a-ps3 | 3.82E-14 | 0.706712 | 0.691 | 0.465 | 9.02E-10 |
| Fau | 4.08E-14 | 0.664545 | 0.938 | 0.856 | 9.62E-10 |
| Rpl6 | 4.87E-14 | 0.931456 | 0.835 | 0.681 | 1.15E-09 |
| Gm5835 | 5.38E-14 | 0.645604 | 0.574 | 0.337 | 1.27E-09 |
| Rpl35 | 5.56E-14 | 0.866387 | 0.869 | 0.766 | 1.31E-09 |
| Gm10231 | 5.73E-14 | 0.589781 | 0.371 | 0.144 | 1.35E-09 |
| Gdi2 | 5.87E-14 | 0.962504 | 0.859 | 0.794 | 1.38E-09 |
| Gm8960 | 6.54E-14 | 0.526058 | 0.443 | 0.202 | 1.54E-09 |
| AI506816 | 8.77E-14 | 0.500514 | 0.478 | 0.214 | 2.07E-09 |
| Gpbp1l1 | 9.75E-14 | 1.206855 | 0.388 | 0.148 | 2.3E-09 |
| Maf | 9.87E-14 | 1.287794 | 0.904 | 0.812 | 2.33E-09 |
| Rpl9-ps6 | 1.69E-13 | 0.987084 | 0.725 | 0.515 | 3.99E-09 |
| Tpm3 | 1.82E-13 | 1.036773 | 0.897 | 0.812 | 4.28E-09 |
| Gm10123 | 1.82E-13 | 0.580568 | 0.952 | 0.924 | 4.29E-09 |
| Zfp710 | 1.83E-13 | 1.605321 | 0.701 | 0.597 | 4.32E-09 |
| Rps3a1 | 2.21E-13 | 0.790394 | 0.883 | 0.796 | 5.21E-09 |
| Abhd2 | 2.28E-13 | -0.4832 | 0.32 | 0.1 | 5.37E-09 |
| Rps27a | 2.43E-13 | 0.762047 | 0.866 | 0.747 | 5.74E-09 |
| Rplp1 | 2.59E-13 | 0.51334 | 0.983 | 0.95 | 6.11E-09 |
| Gng10 | 2.67E-13 | 0.730366 | 0.849 | 0.743 | 6.3E-09 |
| Ftl1 | 2.86E-13 | 0.452672 | 0.993 | 0.988 | 6.74E-09 |
| Cd81 | 2.96E-13 | 0.395278 | 1 | 1 | 6.98E-09 |
| Gm9625 | 3.47E-13 | 0.63518 | 0.656 | 0.427 | 8.18E-09 |
| Gm10263 | 3.87E-13 | 0.37824 | 0.23 | 0.058 | 9.12E-09 |
| Dcn | 4.25E-13 | 1.171935 | 0.265 | 0.08 | 1E-08 |
| Sall1 | 5.41E-13 | -0.84857 | 0.749 | 0.944 | 1.28E-08 |
| Rps9 | 6.63E-13 | 0.458082 | 0.993 | 0.924 | 1.56E-08 |
| Rps24 | 7.55E-13 | 0.714345 | 0.921 | 0.796 | 1.78E-08 |
| Rsrp1 | 7.57E-13 | 0.83528 | 0.935 | 0.878 | 1.78E-08 |
| Cd63-ps | 8.15E-13 | 1.051497 | 0.677 | 0.545 | 1.92E-08 |
| Rpl10-ps1 | 8.5E-13 | 0.444433 | 0.22 | 0.056 | 2E-08 |
| Gm10243 | 1.01E-12 | 2.313578 | 0.302 | 0.106 | 2.38E-08 |
| Klhl9 | 1.03E-12 | 1.527242 | 0.553 | 0.357 | 2.42E-08 |

| | | | | | |
|---|---|---|---|---|---|
| Zfp36l1 | 1.2E-12 | -1.66492 | 0.667 | 0.747 | 2.84E-08 |
| Rps6-ps4 | 1.23E-12 | 0.746644 | 0.694 | 0.465 | 2.9E-08 |
| Ptp4a2 | 1.23E-12 | 1.353561 | 0.763 | 0.663 | 2.9E-08 |
| mt-Atp6 | 1.28E-12 | 0.624291 | 0.88 | 0.768 | 3.03E-08 |
| Gm15500 | 1.36E-12 | 1.465457 | 0.725 | 0.537 | 3.21E-08 |
| Rps12-ps9 | 1.75E-12 | 1.336016 | 0.553 | 0.339 | 4.13E-08 |
| Rps10-ps2 | 2.13E-12 | 0.820639 | 0.746 | 0.571 | 5.03E-08 |
| C130089K0 | 2.17E-12 | -1.67604 | 0.168 | 0.411 | 5.13E-08 |
| Pcnp | 2.45E-12 | 2.738645 | 0.454 | 0.275 | 5.78E-08 |
| Tgfbr2 | 2.46E-12 | -0.99397 | 0.835 | 0.9 | 5.79E-08 |
| Gm6472 | 3.46E-12 | 0.602802 | 0.629 | 0.401 | 8.16E-08 |
| Gm10736 | 4.53E-12 | 0.643804 | 0.77 | 0.603 | 1.07E-07 |
| St13 | 5.01E-12 | 2.077867 | 0.656 | 0.573 | 1.18E-07 |
| H3f3b | 6.12E-12 | 0.737194 | 0.914 | 0.838 | 1.44E-07 |
| Gm24245 | 6.41E-12 | -0.48261 | 0.299 | 0.551 | 1.51E-07 |
| Foxn3 | 6.45E-12 | 1.077783 | 0.869 | 0.83 | 1.52E-07 |
| Lrrc3 | 6.74E-12 | -0.88356 | 0.893 | 0.9 | 1.59E-07 |
| Gm21738 | 9.35E-12 | 0.454503 | 0.454 | 0.244 | 2.21E-07 |
| Slc25a5 | 9.71E-12 | 0.728271 | 0.821 | 0.725 | 2.29E-07 |
| Capza2 | 9.74E-12 | 0.624551 | 0.928 | 0.872 | 2.3E-07 |
| Gm17241 | 1.04E-11 | 0.392463 | 0.68 | 0.469 | 2.46E-07 |
| Rpl26 | 1.13E-11 | 0.850053 | 0.832 | 0.707 | 2.67E-07 |
| Tpm3-rs7 | 1.16E-11 | 0.878248 | 0.656 | 0.451 | 2.74E-07 |
| Gm4149 | 1.21E-11 | 0.508538 | 0.948 | 0.804 | 2.86E-07 |
| Csnk2a1 | 1.25E-11 | 2.351721 | 0.546 | 0.413 | 2.95E-07 |
| Serf2 | 1.41E-11 | 0.627933 | 0.811 | 0.649 | 3.32E-07 |
| Dusp3 | 1.42E-11 | 2.405701 | 0.443 | 0.246 | 3.36E-07 |
| Hmgb1 | 1.44E-11 | 1.023242 | 0.835 | 0.717 | 3.4E-07 |
| Gm15710 | 1.55E-11 | 0.75982 | 0.842 | 0.697 | 3.67E-07 |
| Adap2 | 1.71E-11 | 0.798796 | 0.89 | 0.892 | 4.03E-07 |
| Gm17511 | 1.81E-11 | 0.550182 | 0.309 | 0.128 | 4.28E-07 |
| Rpl19-ps11 | 1.83E-11 | 0.593085 | 0.739 | 0.555 | 4.31E-07 |
| Rpl5 | 1.92E-11 | 1.094683 | 0.845 | 0.776 | 4.53E-07 |
| Oaz1-ps | 1.92E-11 | 0.619078 | 0.746 | 0.539 | 4.54E-07 |
| Rps13-ps4 | 1.94E-11 | 0.472373 | 0.282 | 0.104 | 4.58E-07 |
| Gm5560 | 2.49E-11 | 0.49702 | 0.529 | 0.305 | 5.87E-07 |
| M6pr-ps | 2.67E-11 | 0.629922 | 0.375 | 0.18 | 6.29E-07 |
| Acsl4 | 2.72E-11 | 2.267708 | 0.419 | 0.242 | 6.42E-07 |
| Rpl31 | 2.94E-11 | 0.713441 | 0.859 | 0.703 | 6.94E-07 |
| Rps18-ps3 | 3.07E-11 | 0.574157 | 0.694 | 0.469 | 7.25E-07 |
| Rpl18a | 3.12E-11 | 0.286955 | 0.993 | 0.988 | 7.36E-07 |
| Sema4b | 3.37E-11 | -1.44008 | 0.247 | 0.07 | 7.94E-07 |
| Rps7 | 3.48E-11 | 0.627564 | 0.89 | 0.721 | 8.22E-07 |
| Ly86 | 3.51E-11 | 0.36616 | 0.993 | 0.998 | 8.29E-07 |
| Kdm6b | 4.17E-11 | 2.023007 | 0.313 | 0.132 | 9.85E-07 |
| Rpl19 | 4.51E-11 | 0.619805 | 0.859 | 0.758 | 1.06E-06 |
| Gm9616 | 4.78E-11 | 0.705656 | 0.636 | 0.427 | 1.13E-06 |

| | | | | | |
|---|---|---|---|---|---|
| Gpr34 | 5.92E-11 | 0.540338 | 0.976 | 0.992 | 1.4E-06 |
| Gm47885 | 7.56E-11 | 0.482925 | 0.227 | 0.072 | 1.78E-06 |
| Rpl23 | 7.64E-11 | 0.520781 | 0.921 | 0.812 | 1.8E-06 |
| Akr1a1 | 8.64E-11 | 0.741048 | 0.79 | 0.663 | 2.04E-06 |
| Rps3a2 | 8.95E-11 | 0.617254 | 0.564 | 0.345 | 2.11E-06 |
| Trim12a | 1.01E-10 | 1.010333 | 0.488 | 0.242 | 2.37E-06 |
| Ctsz | 1.38E-10 | 0.368921 | 0.993 | 0.99 | 3.25E-06 |
| Gm10132 | 1.38E-10 | 0.672129 | 0.773 | 0.619 | 3.26E-06 |
| Rps27a-ps2 | 1.47E-10 | 0.759368 | 0.46 | 0.246 | 3.46E-06 |
| Gm10698 | 1.74E-10 | 0.772256 | 0.522 | 0.323 | 4.12E-06 |
| Map3k1 | 2.17E-10 | 1.581927 | 0.553 | 0.403 | 5.13E-06 |
| Rpl7a-ps7 | 2.18E-10 | 0.630267 | 0.378 | 0.192 | 5.15E-06 |
| Pdia3 | 2.26E-10 | 0.526782 | 0.952 | 0.922 | 5.32E-06 |
| Mir703 | 2.31E-10 | 0.462193 | 0.316 | 0.142 | 5.44E-06 |
| Gm26825 | 2.34E-10 | 0.723483 | 0.777 | 0.661 | 5.51E-06 |
| Rpl35a | 2.35E-10 | 0.799799 | 0.845 | 0.699 | 5.55E-06 |
| Rpl31-ps8 | 2.66E-10 | 0.569074 | 0.704 | 0.497 | 6.28E-06 |
| Fam221b | 2.7E-10 | 0.616904 | 0.56 | 0.341 | 6.36E-06 |
| Rps10 | 2.97E-10 | 0.647851 | 0.869 | 0.79 | 7E-06 |
| Soga1 | 2.97E-10 | -1.56574 | 0.399 | 0.605 | 7.02E-06 |
| Rps18 | 2.99E-10 | 0.568746 | 0.904 | 0.794 | 7.05E-06 |
| Rpl17-ps5 | 3.3E-10 | 0.43398 | 0.361 | 0.168 | 7.78E-06 |
| Npm1 | 3.47E-10 | 1.065227 | 0.625 | 0.441 | 8.19E-06 |
| Gm26384 | 3.5E-10 | 0.32861 | 0.21 | 0.064 | 8.25E-06 |
| Ppm1h | 3.7E-10 | 2.195474 | 0.577 | 0.477 | 8.74E-06 |
| Gm15772 | 4.04E-10 | 0.593406 | 0.643 | 0.437 | 9.53E-06 |
| Rbm15 | 4.26E-10 | 0.667443 | 0.347 | 0.162 | 1E-05 |
| Gm8226 | 4.34E-10 | 0.389274 | 0.251 | 0.086 | 1.02E-05 |
| Gm11942 | 4.43E-10 | 0.5153 | 0.595 | 0.393 | 1.04E-05 |
| Rpl17-ps3 | 5.35E-10 | 0.584689 | 0.54 | 0.349 | 1.26E-05 |
| Mylip | 5.5E-10 | 0.92963 | 0.735 | 0.619 | 1.3E-05 |
| Itm2b | 5.64E-10 | 0.278667 | 1 | 1 | 1.33E-05 |
| Gm28437 | 6.31E-10 | 0.600136 | 0.873 | 0.766 | 1.49E-05 |
| Gsk3b | 6.84E-10 | 1.696449 | 0.584 | 0.481 | 1.61E-05 |
| Rpl9 | 8.05E-10 | 0.663547 | 0.856 | 0.685 | 1.9E-05 |
| Gm9794 | 8.72E-10 | 0.609233 | 0.794 | 0.603 | 2.06E-05 |
| Pfn1 | 1.01E-09 | 0.425562 | 0.986 | 0.954 | 2.38E-05 |
| Rpl36-ps2 | 1.05E-09 | 0.956153 | 0.674 | 0.483 | 2.47E-05 |
| Rps26 | 1.08E-09 | 0.884252 | 0.739 | 0.647 | 2.55E-05 |
| Gm2000 | 1.13E-09 | 0.620886 | 0.698 | 0.525 | 2.67E-05 |
| Gm26857 | 1.15E-09 | -2.69126 | 0.096 | 0.273 | 2.71E-05 |
| Gm2a | 1.3E-09 | 0.54653 | 0.814 | 0.611 | 3.06E-05 |
| Rpl4 | 1.4E-09 | 0.62911 | 0.869 | 0.784 | 3.3E-05 |
| Aldoart1 | 1.67E-09 | 0.260052 | 0.206 | 0.064 | 3.93E-05 |
| Inpp5d | 1.92E-09 | -0.78823 | 0.9 | 0.952 | 4.52E-05 |
| P4hb | 2E-09 | -0.8846 | 0.777 | 0.826 | 4.72E-05 |
| Ac165079.: | 2.02E-09 | 0.417139 | 0.323 | 0.148 | 4.77E-05 |

| | | | | | |
|---|---|---|---|---|---|
| Bin2 | 2.14E-09 | 0.606703 | 0.911 | 0.844 | 5.06E-05 |
| Rps3a3 | 2.41E-09 | 0.520774 | 0.515 | 0.331 | 5.68E-05 |
| Fth1 | 2.48E-09 | 0.444248 | 1 | 0.988 | 5.86E-05 |
| Gm12231 | 2.57E-09 | 0.450216 | 0.275 | 0.11 | 6.07E-05 |
| Mrfap1 | 2.6E-09 | 0.880145 | 0.722 | 0.587 | 6.13E-05 |
| Gm12338 | 2.95E-09 | 0.666146 | 0.381 | 0.196 | 6.95E-05 |
| Rps4x | 3.17E-09 | 0.64507 | 0.859 | 0.758 | 7.48E-05 |
| Gm8557 | 3.2E-09 | -1.55846 | 0.247 | 0.096 | 7.55E-05 |
| Gm5905 | 3.2E-09 | 0.393341 | 0.849 | 0.701 | 7.55E-05 |
| Gm3788 | 3.22E-09 | 0.565629 | 0.254 | 0.1 | 7.59E-05 |
| Lrrc58 | 3.22E-09 | 0.937512 | 0.601 | 0.351 | 7.59E-05 |
| Gm11263 | 3.31E-09 | 0.39186 | 0.399 | 0.218 | 7.81E-05 |
| Rpn2 | 3.79E-09 | 0.662171 | 0.777 | 0.633 | 8.94E-05 |
| P2ry12 | 3.85E-09 | 0.346661 | 0.997 | 1 | 9.08E-05 |
| Tmsb10 | 3.91E-09 | 1.149596 | 0.57 | 0.405 | 9.22E-05 |
| Rpl19-ps1 | 3.92E-09 | 0.585082 | 0.368 | 0.196 | 9.26E-05 |
| Rpl39 | 4.24E-09 | 0.632055 | 0.811 | 0.673 | 0.0001 |
| Rpl15 | 4.31E-09 | 0.716183 | 0.777 | 0.611 | 0.000102 |
| Mcmbp | 4.39E-09 | 1.246338 | 0.529 | 0.373 | 0.000104 |
| Gm3362 | 5.05E-09 | -0.54327 | 0.546 | 0.363 | 0.000119 |
| Scoc | 5.08E-09 | 0.757095 | 0.808 | 0.679 | 0.00012 |
| Klf13 | 5.26E-09 | 1.860465 | 0.515 | 0.393 | 0.000124 |
| Rplp0 | 5.35E-09 | 0.421704 | 0.938 | 0.804 | 0.000126 |
| mt-Nd4 | 5.74E-09 | 0.548728 | 0.924 | 0.83 | 0.000135 |
| mt-Nd1 | 5.9E-09 | 0.460953 | 0.945 | 0.9 | 0.000139 |
| Ptma | 5.98E-09 | 0.981802 | 0.711 | 0.551 | 0.000141 |
| Rps2-ps10 | 6.05E-09 | 0.486366 | 0.364 | 0.19 | 0.000143 |
| Oaz1 | 6.29E-09 | 0.493931 | 0.832 | 0.695 | 0.000148 |
| Mir692-2 | 6.43E-09 | 0.388885 | 0.512 | 0.321 | 0.000152 |
| Adar | 6.73E-09 | -2.02342 | 0.134 | 0.309 | 0.000159 |
| Twf1 | 7.44E-09 | 2.355427 | 0.375 | 0.226 | 0.000176 |
| Tial1 | 7.57E-09 | 2.002506 | 0.454 | 0.305 | 0.000179 |
| Ywhae | 9.17E-09 | 0.833885 | 0.808 | 0.741 | 0.000216 |
| Ap2a2 | 9.31E-09 | 0.903118 | 0.447 | 0.273 | 0.00022 |
| Nars | 9.39E-09 | 1.025302 | 0.595 | 0.409 | 0.000221 |
| Kdsr | 9.42E-09 | 0.995172 | 0.309 | 0.138 | 0.000222 |
| Clta | 9.46E-09 | 0.619211 | 0.828 | 0.729 | 0.000223 |
| Gm10288 | 1E-08 | 0.558084 | 0.598 | 0.393 | 0.000236 |
| Nfia | 1.02E-08 | 1.116883 | 0.522 | 0.359 | 0.000241 |
| Rpl11 | 1.15E-08 | 0.542598 | 0.818 | 0.653 | 0.000272 |
| Ctsd | 1.16E-08 | 0.26458 | 1 | 1 | 0.000274 |
| Cfl1 | 1.28E-08 | 0.39405 | 0.993 | 0.968 | 0.000301 |
| Gm14760 | 1.3E-08 | 0.447495 | 0.223 | 0.086 | 0.000308 |
| Nup98 | 1.32E-08 | 1.179272 | 0.371 | 0.206 | 0.000312 |
| Rpl10-ps3 | 1.33E-08 | 0.852501 | 0.691 | 0.505 | 0.000315 |
| Rpsa-ps9 | 1.72E-08 | 0.39562 | 0.423 | 0.242 | 0.000406 |
| Gm7392 | 1.88E-08 | 0.38539 | 0.261 | 0.112 | 0.000445 |

| | | | | | |
|---|---|---|---|---|---|
| Midn | 2.06E-08 | 0.357734 | 0.344 | 0.162 | 0.000486 |
| Rps24-ps3 | 2.12E-08 | 0.479018 | 0.464 | 0.293 | 0.0005 |
| Rpsa | 2.13E-08 | 0.474078 | 0.88 | 0.719 | 0.000502 |
| Rpl13 | 2.25E-08 | 0.466513 | 0.973 | 0.914 | 0.000532 |
| Gm4130 | 2.34E-08 | 0.432605 | 0.491 | 0.307 | 0.000553 |
| D17H6S53E | 2.44E-08 | 2.527045 | 0.244 | 0.108 | 0.000575 |
| Rps15a | 2.47E-08 | 0.717349 | 0.704 | 0.543 | 0.000582 |
| Cd63 | 2.91E-08 | 0.696373 | 0.804 | 0.719 | 0.000687 |
| Entpd1 | 3.2E-08 | -0.65975 | 0.893 | 0.92 | 0.000755 |
| Sipa1l2 | 3.29E-08 | -0.93368 | 0.192 | 0.385 | 0.000776 |
| Ddi2 | 3.43E-08 | 2.235235 | 0.416 | 0.281 | 0.00081 |
| Tram1 | 3.46E-08 | 0.574285 | 0.735 | 0.597 | 0.000817 |
| Atp6ap2 | 3.55E-08 | 0.876405 | 0.667 | 0.553 | 0.000837 |
| Eif5a | 3.67E-08 | 0.695503 | 0.67 | 0.503 | 0.000866 |
| Rpl38-ps2 | 3.81E-08 | 0.407281 | 0.402 | 0.222 | 0.000898 |
| Tmem234 | 3.88E-08 | 1.391144 | 0.526 | 0.377 | 0.000915 |
| Atp5f1 | 3.95E-08 | 0.848108 | 0.567 | 0.391 | 0.000931 |
| Siglech | 4.03E-08 | -0.39524 | 1 | 1 | 0.000951 |
| Fcrls | 4.27E-08 | 0.537576 | 0.924 | 0.85 | 0.001008 |
| Rpsa-ps10 | 4.37E-08 | 0.461277 | 0.722 | 0.523 | 0.001032 |
| Exosc2 | 4.51E-08 | 1.16599 | 0.351 | 0.186 | 0.001064 |
| Sept10 | 4.72E-08 | 1.383094 | 0.309 | 0.152 | 0.001114 |
| Cd9 | 5.12E-08 | 0.442283 | 0.997 | 0.976 | 0.001208 |
| Ccdc152 | 5.18E-08 | 0.258412 | 0.405 | 0.234 | 0.001221 |
| Hnrnpa1 | 5.45E-08 | 0.635959 | 0.409 | 0.222 | 0.001287 |
| Gm5529 | 5.66E-08 | 0.370615 | 0.457 | 0.283 | 0.001334 |
| Sppl2a | 5.73E-08 | 0.788305 | 0.66 | 0.487 | 0.001352 |
| Tmed2 | 6.03E-08 | 0.738027 | 0.725 | 0.575 | 0.001423 |
| Rps14 | 6.21E-08 | 0.492855 | 0.945 | 0.914 | 0.001465 |
| Rpl21-ps15 | 6.24E-08 | 0.41437 | 0.45 | 0.275 | 0.001471 |
| Anp32a | 6.72E-08 | 1.259825 | 0.436 | 0.279 | 0.001586 |
| Gtf2a1 | 6.82E-08 | 1.762636 | 0.478 | 0.333 | 0.001609 |
| Pabpc1 | 7.11E-08 | 0.45914 | 0.777 | 0.611 | 0.001676 |
| Gm9844 | 7.28E-08 | 0.340212 | 0.247 | 0.108 | 0.001717 |
| Rpl28-ps1 | 8.58E-08 | 0.488772 | 0.57 | 0.377 | 0.002023 |
| Arl6ip1 | 8.73E-08 | 0.664531 | 0.804 | 0.695 | 0.002059 |
| Rps8-ps1 | 8.76E-08 | 0.345182 | 0.285 | 0.136 | 0.002067 |
| Btg1 | 9.05E-08 | 0.776962 | 0.711 | 0.565 | 0.002135 |
| Rps13-ps2 | 9.14E-08 | 0.446093 | 0.67 | 0.463 | 0.002157 |
| Gm10275 | 1.02E-07 | 0.85722 | 0.522 | 0.377 | 0.002409 |
| Rps19-ps6 | 1.06E-07 | 0.328656 | 0.78 | 0.593 | 0.002497 |
| Gatad2b | 1.06E-07 | 1.413391 | 0.409 | 0.24 | 0.0025 |
| Gm32036 | 1.11E-07 | 2.628334 | 0.423 | 0.331 | 0.002613 |
| Eno1b | 1.18E-07 | 0.844737 | 0.639 | 0.529 | 0.002779 |
| Cd300c2 | 1.22E-07 | 0.528157 | 0.818 | 0.747 | 0.002871 |
| Rpl10a-ps1 | 1.24E-07 | 0.492803 | 0.667 | 0.471 | 0.002936 |
| Ubl5 | 1.35E-07 | 0.873125 | 0.526 | 0.365 | 0.003189 |

| | | | | | |
|---|---|---|---|---|---|
| Fos | 1.38E-07 | -0.76116 | 0.326 | 0.18 | 0.003262 |
| Hspa5 | 1.42E-07 | 0.584608 | 0.828 | 0.727 | 0.003346 |
| Rps6-ps1 | 1.43E-07 | -0.26998 | 0.216 | 0.088 | 0.003384 |
| Gm43584 | 1.44E-07 | 0.336389 | 0.258 | 0.118 | 0.003406 |
| Hsp90b1 | 1.47E-07 | 0.539318 | 0.935 | 0.886 | 0.003472 |
| Uba52 | 1.54E-07 | 0.533411 | 0.814 | 0.743 | 0.003628 |
| Rpl14-ps1 | 1.58E-07 | 0.517163 | 0.739 | 0.571 | 0.003728 |
| Laptm4a | 1.61E-07 | 0.520016 | 0.918 | 0.886 | 0.003797 |
| Gigyf1 | 1.68E-07 | 0.946985 | 0.416 | 0.259 | 0.003957 |
| Rpl17-ps8 | 1.73E-07 | 0.56797 | 0.251 | 0.112 | 0.004079 |
| Sun2 | 1.75E-07 | -1.79546 | 0.557 | 0.655 | 0.004123 |
| Rps20 | 1.77E-07 | 0.511005 | 0.863 | 0.808 | 0.004182 |
| Rps12 | 1.86E-07 | 0.372307 | 0.876 | 0.784 | 0.004397 |
| Crebbp | 1.89E-07 | 1.95139 | 0.543 | 0.463 | 0.00445 |
| Prpf39 | 2.11E-07 | 1.751523 | 0.278 | 0.132 | 0.004982 |
| Ubb | 2.16E-07 | 0.25104 | 0.986 | 0.94 | 0.005098 |
| Gm6807 | 2.17E-07 | 0.325894 | 0.258 | 0.118 | 0.005126 |
| Gm12918 | 2.24E-07 | 0.511707 | 0.653 | 0.511 | 0.005293 |
| Gm6394 | 2.3E-07 | 0.318434 | 0.67 | 0.463 | 0.005418 |
| Gm14165 | 2.33E-07 | 0.371916 | 0.351 | 0.198 | 0.005504 |
| Lamp1 | 2.36E-07 | 0.382656 | 0.99 | 0.968 | 0.005559 |
| Lrp1 | 2.5E-07 | -0.87769 | 0.739 | 0.842 | 0.005905 |
| Gm38317 | 2.53E-07 | 0.294677 | 0.405 | 0.234 | 0.005965 |
| Camk2n1 | 2.78E-07 | 1.361302 | 0.351 | 0.194 | 0.006554 |
| Ctsb | 3.03E-07 | 0.297445 | 1 | 1 | 0.007144 |
| Gm11361 | 3.06E-07 | 0.385453 | 0.584 | 0.417 | 0.007211 |
| Btg2 | 3.22E-07 | 1.114529 | 0.519 | 0.357 | 0.007594 |
| Ppp1ca | 3.45E-07 | 0.695035 | 0.68 | 0.543 | 0.008148 |
| Rpl3-ps1 | 3.51E-07 | 0.707443 | 0.708 | 0.637 | 0.008286 |
| Ctsh | 3.59E-07 | 0.397599 | 0.966 | 0.908 | 0.00847 |
| Gm10073 | 3.96E-07 | 0.358683 | 0.732 | 0.531 | 0.009346 |
| Fam3c | 4.08E-07 | 0.626413 | 0.45 | 0.263 | 0.009632 |
| Impact | 4.14E-07 | 2.392081 | 0.309 | 0.176 | 0.009761 |
| Ywhah | 4.84E-07 | 0.506298 | 0.9 | 0.856 | 0.011417 |
| Rpl14 | 4.93E-07 | 0.630023 | 0.842 | 0.741 | 0.011629 |
| Gm14586 | 4.94E-07 | 0.442782 | 0.581 | 0.407 | 0.011642 |
| Gsn | 5.2E-07 | 0.559748 | 0.296 | 0.146 | 0.012268 |
| Rpl21 | 5.23E-07 | 0.306453 | 0.966 | 0.878 | 0.012327 |
| Srsf5 | 5.27E-07 | 0.813184 | 0.66 | 0.499 | 0.012439 |
| Gpx1 | 5.42E-07 | 0.588447 | 0.794 | 0.709 | 0.012793 |
| Selenof | 5.6E-07 | 0.626721 | 0.701 | 0.587 | 0.013219 |
| Rpl10 | 5.96E-07 | 0.435757 | 0.935 | 0.854 | 0.01405 |
| Atp5e | 6.21E-07 | 1.017337 | 0.54 | 0.431 | 0.014641 |
| Rpl32 | 6.49E-07 | 0.41379 | 0.928 | 0.876 | 0.015319 |
| Zfp706 | 6.87E-07 | 0.661842 | 0.677 | 0.559 | 0.01621 |
| Capzb | 7.32E-07 | 0.479832 | 0.825 | 0.743 | 0.017273 |
| Gm9774 | 7.54E-07 | 0.890874 | 0.34 | 0.196 | 0.017776 |

| | | | | | |
|---|---|---|---|---|---|
| Rps15 | 7.73E-07 | 0.440322 | 0.832 | 0.691 | 0.01824 |
| Paip1 | 7.83E-07 | 1.591395 | 0.289 | 0.15 | 0.018467 |
| Tmem59 | 8.01E-07 | 0.468062 | 0.883 | 0.79 | 0.0189 |
| Tmem250- | 9.01E-07 | 1.775214 | 0.515 | 0.447 | 0.021261 |
| mt-Nd3 | 9.91E-07 | 0.744676 | 0.636 | 0.519 | 0.023381 |
| Gm5526 | 1E-06 | 0.579894 | 0.436 | 0.297 | 0.023636 |
| Sirt6 | 1.04E-06 | -0.28427 | 0.206 | 0.084 | 0.024553 |
| Ldhb | 1.06E-06 | 0.529193 | 0.814 | 0.729 | 0.024944 |
| Rpl27 | 1.06E-06 | 0.495895 | 0.777 | 0.665 | 0.024991 |
| Gm10020 | 1.07E-06 | 0.372698 | 0.285 | 0.152 | 0.02531 |
| Mfap1b | 1.1E-06 | 0.920268 | 0.275 | 0.13 | 0.025853 |
| Cox6c | 1.21E-06 | 0.85638 | 0.557 | 0.429 | 0.028575 |
| Wapl | 1.23E-06 | 1.496048 | 0.402 | 0.277 | 0.029022 |
| Caml | 1.29E-06 | 0.934381 | 0.309 | 0.162 | 0.030317 |
| B430010I2: | 1.38E-06 | -1.39419 | 0.1 | 0.24 | 0.032532 |
| Gm9493 | 1.41E-06 | 0.340957 | 0.388 | 0.224 | 0.033162 |
| Gm11273 | 1.55E-06 | 0.498906 | 0.34 | 0.19 | 0.0365 |
| Ndufa4 | 1.56E-06 | 0.796551 | 0.519 | 0.375 | 0.036728 |
| Mrpl52 | 1.58E-06 | 0.633443 | 0.481 | 0.317 | 0.037385 |
| Gm6170 | 1.6E-06 | 0.34714 | 0.491 | 0.337 | 0.037845 |
| Otub1 | 1.6E-06 | 0.977972 | 0.436 | 0.293 | 0.037848 |
| Mpc1-ps | 1.65E-06 | 0.756486 | 0.526 | 0.363 | 0.039031 |
| Gm6421 | 1.72E-06 | 0.508937 | 0.306 | 0.166 | 0.040585 |
| Gm10709 | 1.72E-06 | 0.427015 | 0.402 | 0.251 | 0.040628 |
| Aldoa | 1.73E-06 | 0.535925 | 0.749 | 0.643 | 0.040866 |
| Gm34280 | 1.74E-06 | -1.31539 | 0.131 | 0.271 | 0.040928 |
| Rpl12 | 1.79E-06 | 0.466989 | 0.78 | 0.601 | 0.042338 |
| Rpl36a | 1.89E-06 | 0.865774 | 0.625 | 0.525 | 0.044539 |
| Gm7618 | 1.92E-06 | 0.254198 | 0.368 | 0.216 | 0.045202 |
| Cox7a2 | 1.92E-06 | 0.79283 | 0.443 | 0.295 | 0.045398 |
| Gm5822 | 2.01E-06 | 0.318589 | 0.213 | 0.094 | 0.047379 |
| Timmdc1 | 2.03E-06 | 1.005862 | 0.203 | 0.086 | 0.047842 |
| Rps13 | 2.06E-06 | 0.456994 | 0.849 | 0.741 | 0.048706 |
